## Supplementary Figures for "Uncovering the Regional and Cell Specific Bioactivity of Injectable Extracellular Matrix Biomaterials in Myocardial Infarction through Spatial and Single Nucleus Transcriptomics"

**Supplementary Information:**

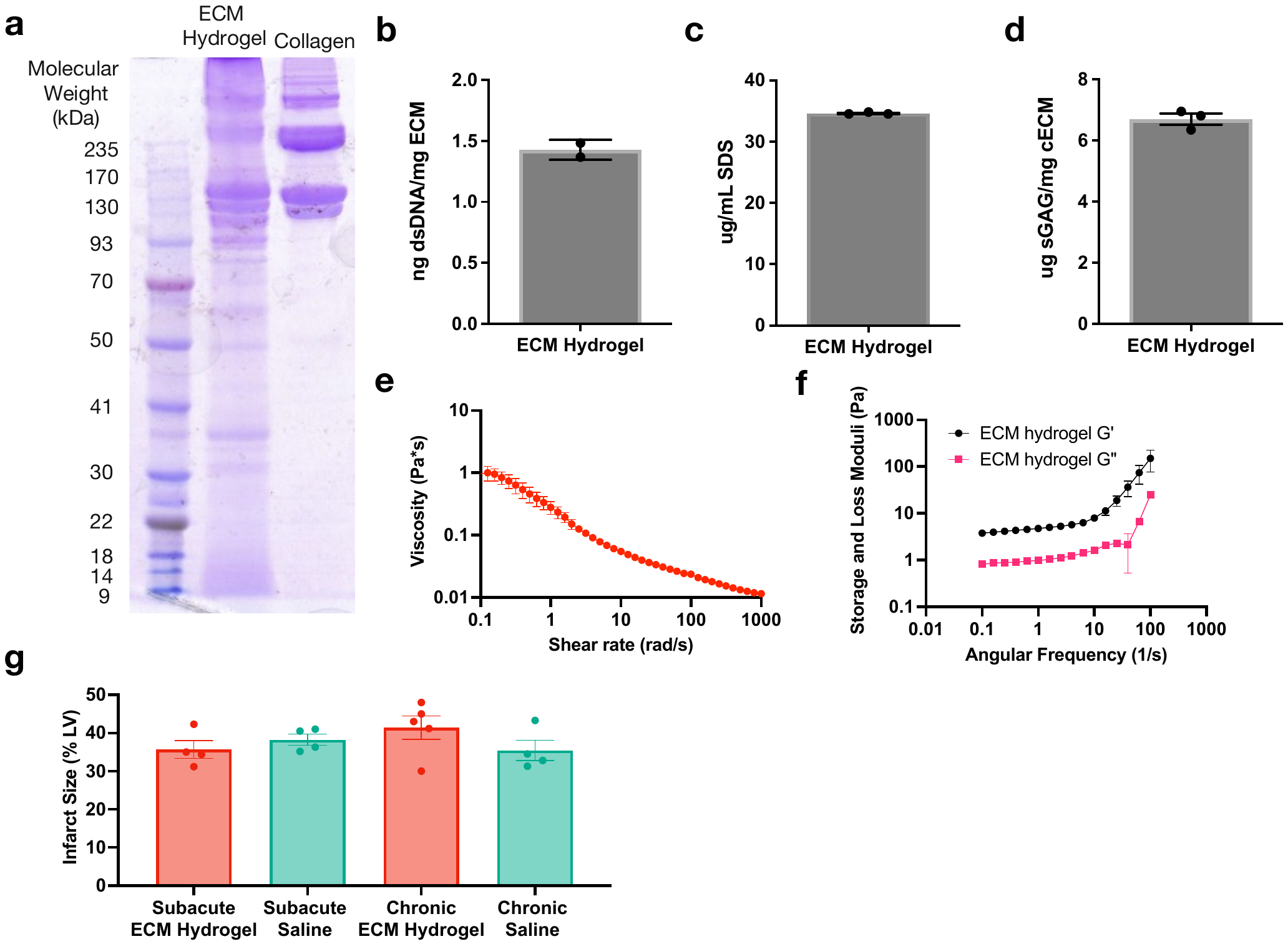

**Supplementary Information Figure 1. Quality control for ECM and biological samples**. **(a-d)** ECM hydrogel quality control metrics are done to ensure that the ECM hydrogel is similar with each batch. **(a)** Gel electrophoresis exhibits a different protein composition compared to collagen. **(b)** Double stranded DNA is quantified to ensure the ECM contains minimal DNA from cellular debris, n = 2 ECM aliquots. **(c)** Sodium dodecyl sulfate (SDS) is quantified to ensure there is no detergent remaining from the decellularization process, n = 3 ECM aliquots. **(d)** Glycosoaminoglycan (GAG) content is quantified to ensure similar GAG concentration across batches, n = 3 ECM aliquots. **(e)** Rheometry is quantified to ensure similar biomechanical properties between ECM hydrogel batches, n = 3 ECM aliquots. **(f)** The storage (G’) and loss (G’’) moduli were also calculated for ECM hydrogel batches, n = 3 ECM aliquots. **(g)** Infarct quantification of each sample, split by condition and MI model, demonstrating consistent infarct sizes, with n = 2-3 per each MI model and its respective treatment condition. Data are presented as mean ± SEM.

**Supplementary Information Figure 2. Strategy for identifying the infarct zone in spatial samples in subacute MI model. (a)** Myocardium (green) was labelled with an anti-alpha-actinin antibody, with a white outline indicating the infarct and the ECM hydrogel fluorescently tagged in light blue. **(b-d)** The adjacent section was used for 10X Visium, with coarse clustering populations identified **(b)**. *Myh6* **(c)** and *Tnnt2* **(d)**, two markers for healthy myocardium, were used to segment and identify clusters that are *Myh6* and *Tnnt2* low, with white outlines overlayed onto the coarse clustering plot, indicating which clusters are infarct specific.

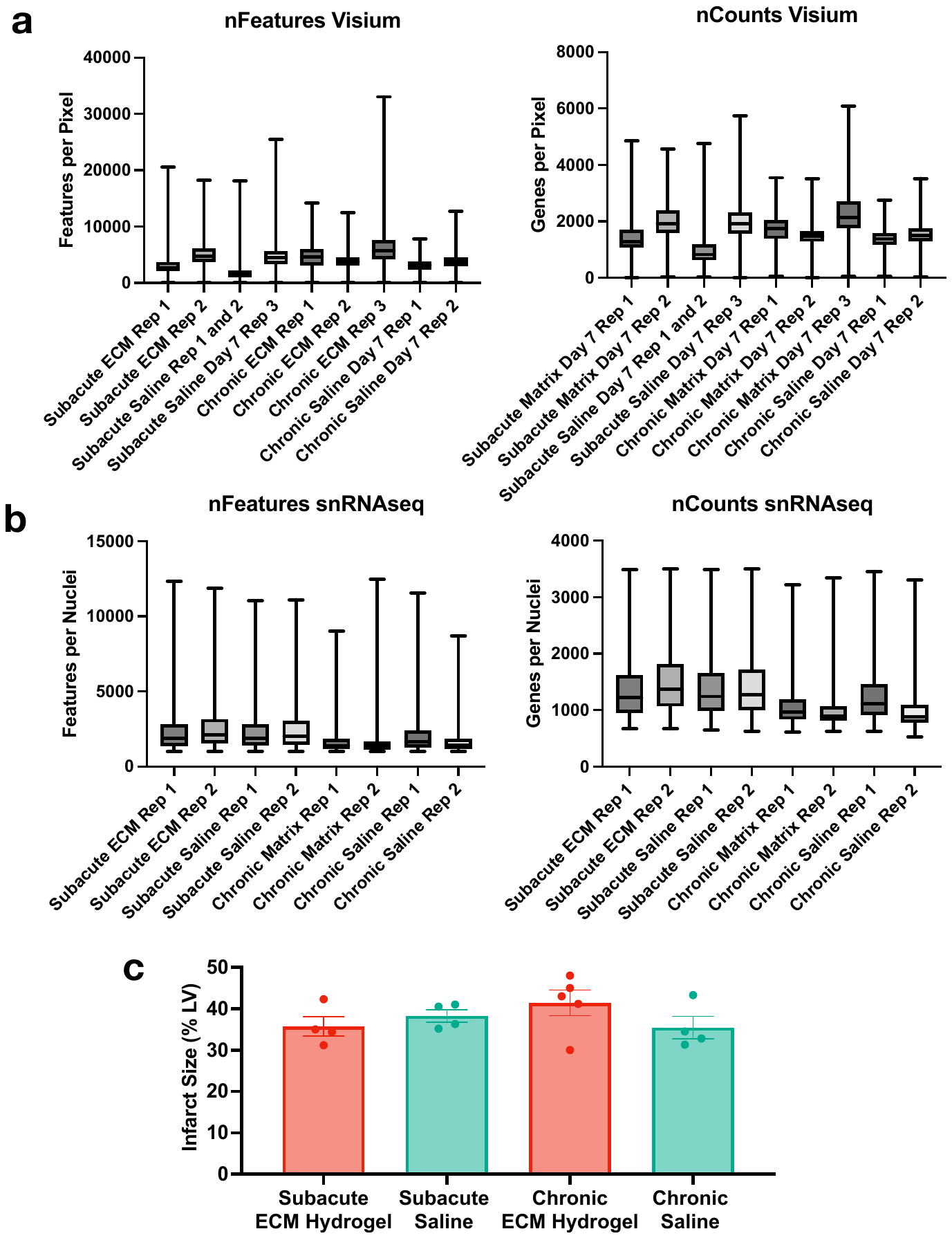

**Supplementary Information Figure 3. Transcriptomic quality control for Visium and snRNAseq samples. (a)** Quality metrics of samples and replicates for Visium samples represented in features per pixel (nFeatures) and genes per sample (nCounts). Data are presented as box and whisker plots. **(b)** Quality metrics of samples and replicates for snRNAseq samples represented in features per nuclei (nFeatures) and genes per nuclei (nCounts). Data are presented as box and whisker plots.

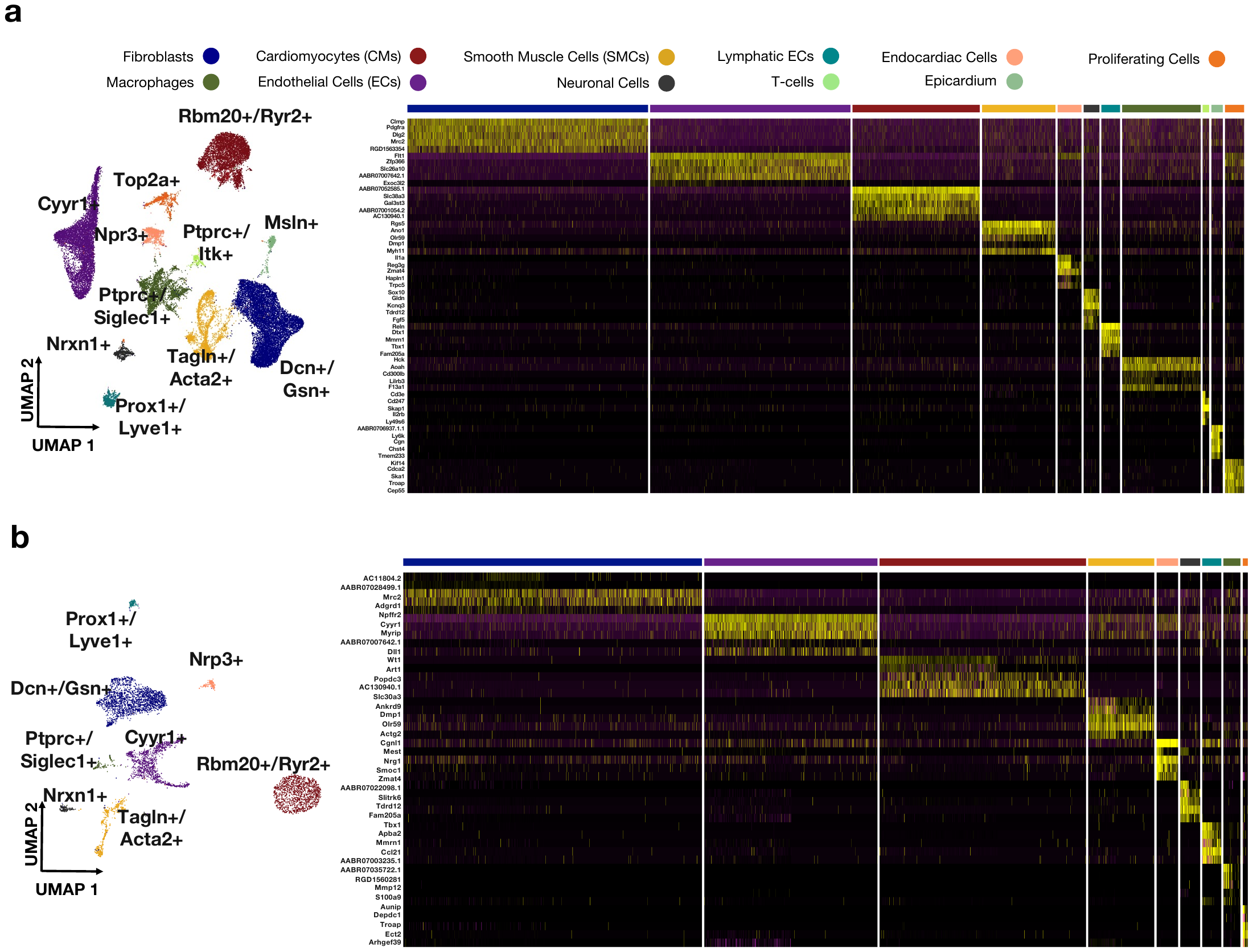

**Supplementary Information Figure 4. Feature subsetting and top genes for canonical cell types. (a)** Gene subsetting strategy overlayed onto UMAP for subacute cell types, alongside heatmap of top 5 genes per each cell type. (**b)** Gene subsetting strategy overlayed onto UMAP for chronic cell types, alongside heatmap of top 5 genes per each cell type.

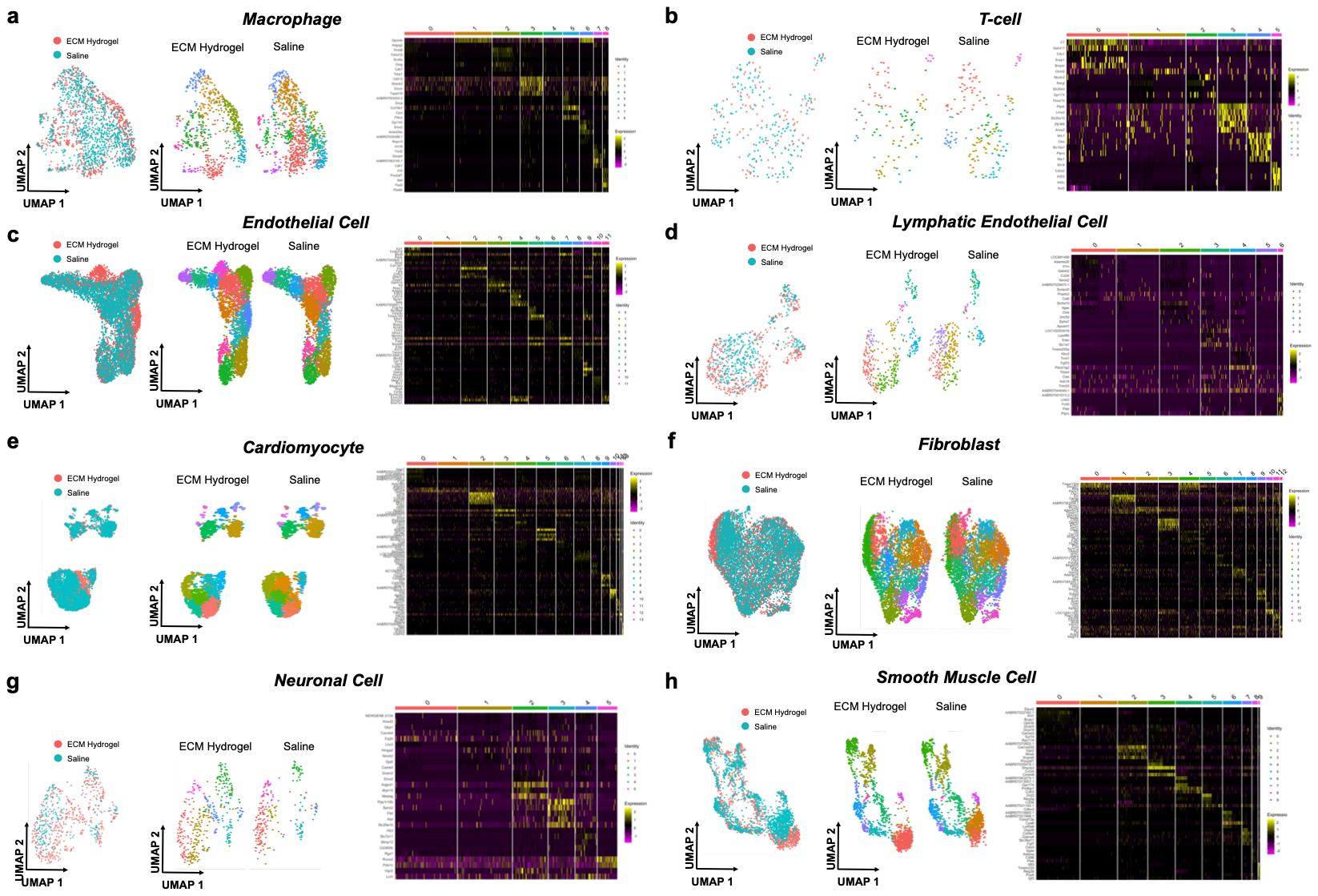

**Supplementary Information Figure 5. Subclusters of subsetted cell types in subacute MI.**

**(a)** Macrophages, **(b)** T-cells**,** **(c)** endothelial cells, **(d)** lymphatic endothelial cells, **(e)** cardiomyocytes, **(f)** fibroblasts, **(g)** neuronal cells, and **(h)** smooth muscle cells were subsetted and reintegrated. They were then reclustered at resolution 1, with the top 5 marker genes for each subcluster.

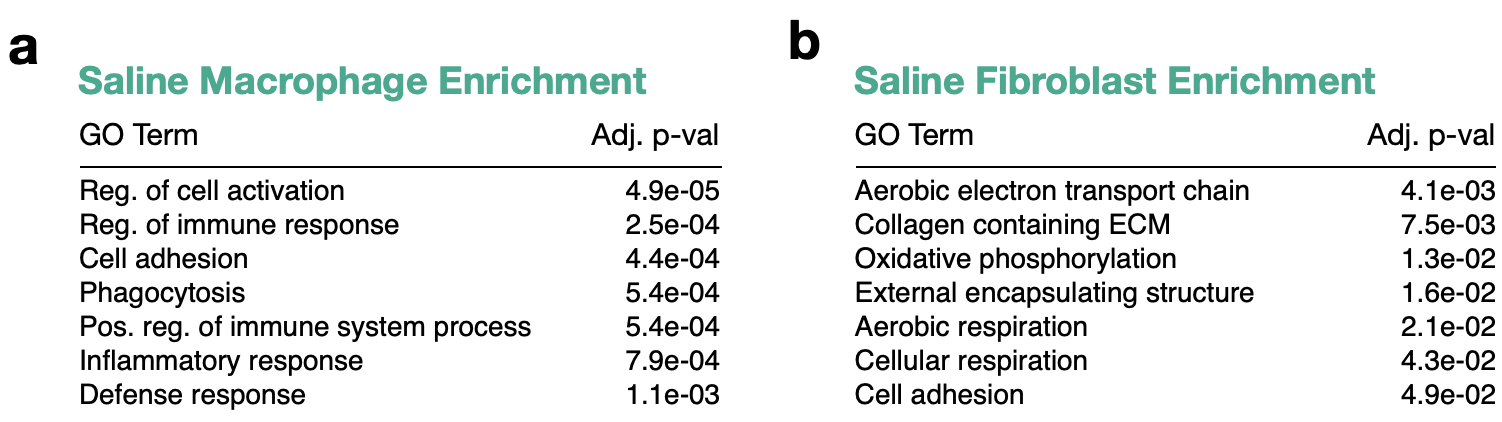

**Supplementary Information Figure 6. Saline GO terms in subacute MI model. (a)** Macrophage enrichment of genes upregulated in saline treatment for the subacute MI model are shown. **(b)** Fibroblast enrichment of genes upregulated in saline treatment for subacute MI model are shown.

**Supplementary Information Figure 7. Strategy for identifying infarct zone in the chronic MI spatial samples**. **(a)** Myocardium (green) was labelled with an anti-alpha-actinin antibody, with a white outline indicating the infarct and the ECM hydrogel fluorescently tagged in light blue. **(b-d)** The adjacent section was used for 10X Visium, with coarse clustering populations identified **(b)**. *Myh6* **(c)** and *Tnnt2* **(d)**, two markers for healthy myocardium, were used to segment and identify clusters that are *Myh6* and *Tnnt2* low, with their white outlines overlayed onto the coarse clustering plot, indicating which clusters are infarct specific.

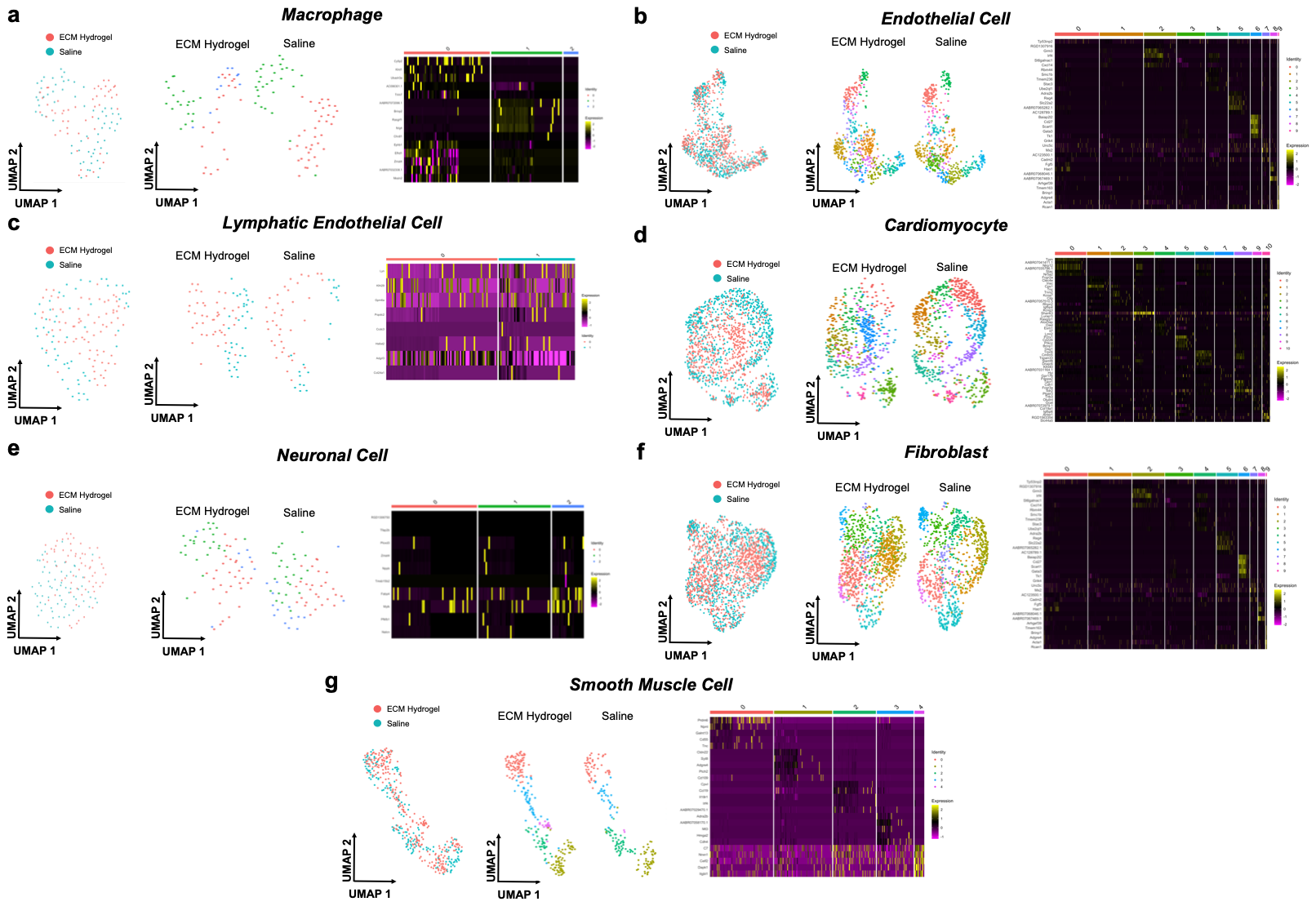

**Supplementary Information Figure 8. Subclusters of subsetted cell types in chronic MI.**

**(a)** Macrophages, **(b)** endothelial cells, **(c)** lymphatic endothelial cells, **(d)** cardiomyocytes, **(e)** neuronal cells, **(f)** fibroblasts, and **(g)** smooth muscle cells were subsetted and reintegrated. They were then reclustered at resolution 1, with the top 5 marker genes for each subcluster.

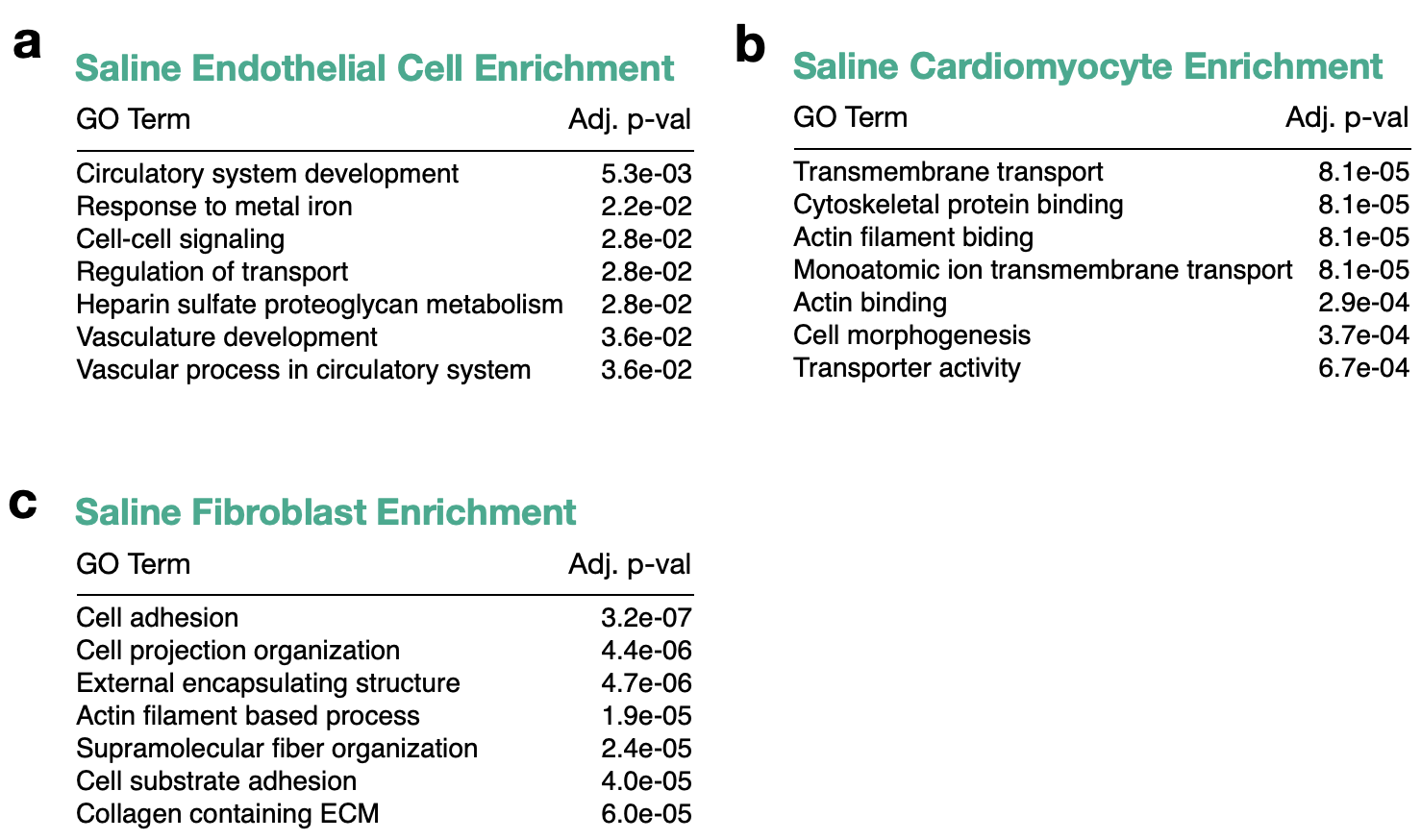

**Supplementary Information Figure 9. Saline GO terms in chronic MI model.** **(a)** Endothelial cell, **(b)** cardiomyocyte, and **(c)** fibroblast enrichment of genes upregulated in saline treatment are shown for the chronic MI model.
