## Supplemental Table 6 for "Uncovering the Regional and Cell Specific Bioactivity of Injectable Extracellular Matrix Biomaterials in Myocardial Infarction through Spatial and Single Nucleus Transcriptomics"

| Supplementary Table 6. Subcluster Markers Genes for Cell Types in Subacute MI | | | | | | | |
|---|---|---|---|---|---|---|---|
| **Cell Type** | **Gene** | **Cluster** | **p_val** | **avg_log2FC** | **pct.1** | **pct.2** | **p_val_adj** |
| Macrophage | Gpnmb | 1 | 8.42E-39 | 1.56795641 | 0.781 | 0.439 | 1.68E-35 |
| Macrophage | Adgrg2 | 1 | 2.94E-22 | 0.5302661 | 0.09 | 0.343 | 5.88E-19 |
| Macrophage | Epha5 | 2 | 2.08E-130 | 1.74592728 | 0.664 | 0.062 | 4.17E-127 |
| Macrophage | Prmt8 | 2 | 1.04E-120 | 4.3349382 | 0.676 | 0.092 | 2.07E-117 |
| Macrophage | Tnfrsf19 | 2 | 2.06E-109 | 2.62720839 | 0.697 | 0.118 | 4.11E-106 |
| Macrophage | Scn9a | 2 | 5.42E-99 | 6.09285109 | 0.502 | 0.05 | 1.08E-95 |
| Macrophage | Neil3 | 2 | 3.68E-97 | 1.51833022 | 0.71 | 0.151 | 7.36E-94 |
| Macrophage | Col9a1 | 2 | 4.96E-93 | 1.48544398 | 0.531 | 0.063 | 9.91E-90 |
| Macrophage | Cacna2d2 | 2 | 4.59E-91 | 1.88835547 | 0.469 | 0.058 | 9.19E-88 |
| Macrophage | Sema3e | 2 | 1.12E-86 | 1.10510661 | 0.726 | 0.181 | 2.24E-83 |
| Macrophage | Caln1 | 2 | 3.72E-82 | 2.39744718 | 0.581 | 0.065 | 7.43E-79 |
| Macrophage | Grm1 | 2 | 1.27E-75 | 0.87760736 | 0.49 | 0.041 | 2.53E-72 |
| Macrophage | Kif11 | 2 | 1.98E-74 | 0.40073116 | 0.349 | 0.088 | 3.97E-71 |
| Macrophage | AABR070473 | 2 | 2.86E-73 | 2.50426524 | 0.556 | 0.102 | 5.72E-70 |
| Macrophage | Fbn2 | 2 | 2.70E-72 | 1.30359928 | 0.714 | 0.187 | 5.40E-69 |
| Macrophage | Adipoq | 2 | 1.05E-71 | 1.85125396 | 0.693 | 0.183 | 2.10E-68 |
| Macrophage | Armc4 | 2 | 2.61E-68 | 2.41193297 | 0.573 | 0.118 | 5.22E-65 |
| Macrophage | Brinp2 | 2 | 1.78E-67 | 0.44472386 | 0.734 | 0.175 | 3.55E-64 |
| Macrophage | Oas3 | 2 | 3.83E-67 | 0.33099909 | 0.349 | 0.067 | 7.66E-64 |
| Macrophage | AABR070651 | 2 | 1.05E-66 | 0.42460622 | 0.386 | 0.035 | 2.10E-63 |
| Macrophage | AABR070179 | 2 | 9.86E-66 | 0.57378942 | 0.631 | 0.128 | 1.97E-62 |
| Macrophage | Alk | 2 | 6.13E-65 | 0.7196904 | 0.415 | 0.063 | 1.23E-61 |
| Macrophage | Chst9 | 2 | 2.88E-64 | 1.25585363 | 0.456 | 0.074 | 5.77E-61 |
| Macrophage | Reg3g | 2 | 2.32E-63 | 2.28008657 | 0.581 | 0.15 | 4.65E-60 |
| Macrophage | Reg3b | 2 | 3.06E-62 | 0.35905156 | 0.651 | 0.171 | 6.12E-59 |
| Macrophage | Lrrc4b | 2 | 9.86E-62 | 1.82508499 | 0.697 | 0.173 | 1.97E-58 |
| Macrophage | Rasgrf1 | 2 | 5.78E-61 | 2.34013495 | 0.556 | 0.122 | 1.16E-57 |
| Macrophage | Ankrd9 | 2 | 1.41E-60 | 1.01691678 | 0.544 | 0.147 | 2.82E-57 |
| Macrophage | Flt4 | 2 | 4.14E-60 | 0.50270866 | 0.697 | 0.185 | 8.28E-57 |

| | | | | | | | |
|---|---|---|---|---|---|---|---|
| Macrophage | Gbp1 | 2 | 9.06E-58 | 1.60392571 | 0.527 | 0.098 | 1.81E-54 |
| Macrophage | Sv2c | 2 | 1.52E-55 | 0.72898576 | 0.481 | 0.102 | 3.03E-52 |
| Macrophage | Fbxw10 | 2 | 2.41E-55 | 2.11183868 | 0.411 | 0.096 | 4.82E-52 |
| Macrophage | Omg | 2 | 4.32E-55 | 3.78169005 | 0.407 | 0.067 | 8.64E-52 |
| Macrophage | Sybu | 2 | 1.21E-54 | 1.64782735 | 0.349 | 0.056 | 2.42E-51 |
| Macrophage | Nuf2 | 2 | 2.24E-54 | 1.22555692 | 0.51 | 0.098 | 4.49E-51 |
| Macrophage | Lrrtm3 | 2 | 2.89E-54 | 1.12257288 | 0.722 | 0.262 | 5.78E-51 |
| Macrophage | Nrg1 | 2 | 6.29E-53 | 1.53885925 | 0.734 | 0.242 | 1.26E-49 |
| Macrophage | Tmem178a | 2 | 3.66E-52 | 1.1749745 | 0.705 | 0.215 | 7.32E-49 |
| Macrophage | Asb15 | 2 | 9.56E-52 | 0.68460965 | 0.531 | 0.095 | 1.91E-48 |
| Macrophage | Gen1 | 2 | 3.53E-50 | 0.55828545 | 0.685 | 0.17 | 7.07E-47 |
| Macrophage | Sypl2 | 2 | 1.07E-49 | 0.3416018 | 0.39 | 0.066 | 2.14E-46 |
| Macrophage | Tnc | 2 | 1.35E-49 | 1.36874707 | 0.556 | 0.12 | 2.70E-46 |
| Macrophage | Fbxo40 | 2 | 4.38E-48 | 0.59805326 | 0.556 | 0.135 | 8.76E-45 |
| Macrophage | Lrrc2 | 2 | 5.63E-48 | 0.8880372 | 0.589 | 0.11 | 1.13E-44 |
| Macrophage | Tesc | 2 | 6.62E-48 | 0.68275767 | 0.631 | 0.147 | 1.32E-44 |
| Macrophage | Rbp1 | 2 | 1.14E-47 | 0.66219256 | 0.432 | 0.116 | 2.28E-44 |
| Macrophage | Sptb | 2 | 1.69E-47 | 0.41083658 | 0.618 | 0.135 | 3.39E-44 |
| Macrophage | Dlgap5 | 2 | 3.00E-47 | 0.50604811 | 0.568 | 0.151 | 6.01E-44 |
| Macrophage | F13a1 | 2 | 3.34E-47 | 1.42171419 | 0.921 | 0.484 | 6.67E-44 |
| Macrophage | Ky | 2 | 4.91E-47 | 0.48974989 | 0.705 | 0.193 | 9.82E-44 |
| Macrophage | Trpc3 | 2 | 1.71E-46 | 0.68108858 | 0.49 | 0.128 | 3.43E-43 |
| Macrophage | Rec114 | 2 | 9.60E-46 | 1.37653852 | 0.643 | 0.212 | 1.92E-42 |
| Macrophage | Iqgap3 | 2 | 5.67E-45 | 1.30958391 | 0.606 | 0.216 | 1.13E-41 |
| Macrophage | Skap1 | 2 | 1.08E-44 | 0.95815971 | 0.622 | 0.185 | 2.16E-41 |
| Macrophage | Bdnf | 2 | 3.21E-44 | 0.86036378 | 0.519 | 0.136 | 6.41E-41 |
| Macrophage | Lmcd1 | 2 | 4.78E-44 | 0.69568985 | 0.718 | 0.221 | 9.56E-41 |
| Macrophage | Cnnm2 | 2 | 4.81E-44 | 1.06095836 | 0.643 | 0.155 | 9.61E-41 |
| Macrophage | Klhl4 | 2 | 7.16E-43 | 0.29134245 | 0.539 | 0.129 | 1.43E-39 |
| Macrophage | Slc1a3 | 2 | 7.22E-43 | 1.18767417 | 0.689 | 0.267 | 1.44E-39 |
| Macrophage | Heyl | 2 | 3.54E-42 | 0.83351838 | 0.564 | 0.121 | 7.07E-39 |

| | | | | | | | |
|---|---|---|---|---|---|---|---|
| Macrophage | Pfkfb1 | 2 | 3.69E-42 | 1.60470221 | 0.415 | 0.125 | 7.38E-39 |
| Macrophage | Cgn | 2 | 6.46E-42 | 1.46869493 | 0.606 | 0.167 | 1.29E-38 |
| Macrophage | Ugdh | 2 | 7.19E-42 | 0.40688514 | 0.705 | 0.215 | 1.44E-38 |
| Macrophage | Stab1 | 2 | 7.58E-42 | 1.33677957 | 0.959 | 0.624 | 1.52E-38 |
| Macrophage | Clu | 2 | 8.78E-42 | 0.94733773 | 0.726 | 0.248 | 1.76E-38 |
| Macrophage | Nrxn3 | 2 | 1.98E-41 | 1.78218733 | 0.477 | 0.095 | 3.96E-38 |
| Macrophage | Samd12 | 2 | 2.04E-41 | 0.80149159 | 0.726 | 0.32 | 4.08E-38 |
| Macrophage | Adamts3 | 2 | 2.07E-41 | 0.87067361 | 0.39 | 0.106 | 4.15E-38 |
| Macrophage | Sntg2 | 2 | 8.76E-41 | 0.77782254 | 0.602 | 0.13 | 1.75E-37 |
| Macrophage | Rims1 | 2 | 1.63E-40 | 2.00459454 | 0.402 | 0.062 | 3.27E-37 |
| Macrophage | Mctp2 | 2 | 1.64E-40 | 1.17139861 | 0.896 | 0.47 | 3.27E-37 |
| Macrophage | Igfbp4 | 2 | 4.65E-40 | 0.78006249 | 0.531 | 0.147 | 9.29E-37 |
| Macrophage | Ccdc148 | 2 | 6.61E-40 | 1.11969835 | 0.61 | 0.121 | 1.32E-36 |
| Macrophage | Lyve1 | 2 | 3.22E-39 | 0.5342913 | 0.805 | 0.325 | 6.45E-36 |
| Macrophage | Art3 | 2 | 3.56E-39 | 0.96129781 | 0.647 | 0.22 | 7.12E-36 |
| Macrophage | Sema3a | 2 | 5.22E-39 | 1.18731922 | 0.544 | 0.163 | 1.04E-35 |
| Macrophage | Ryr3 | 2 | 6.91E-39 | 0.3682457 | 0.556 | 0.143 | 1.38E-35 |
| Macrophage | Cux2 | 2 | 4.86E-38 | 0.27971076 | 0.606 | 0.195 | 9.71E-35 |
| Macrophage | Flnc | 2 | 1.61E-37 | 0.63007447 | 0.535 | 0.185 | 3.23E-34 |
| Macrophage | Abcd2 | 2 | 1.71E-37 | 2.01739848 | 0.382 | 0.129 | 3.42E-34 |
| Macrophage | Nr1h4 | 2 | 2.51E-37 | 1.27068837 | 0.531 | 0.112 | 5.02E-34 |
| Macrophage | Scn7a | 2 | 3.59E-37 | 0.89570811 | 0.461 | 0.089 | 7.18E-34 |
| Macrophage | Nkain2 | 2 | 7.93E-37 | 0.78247276 | 0.502 | 0.118 | 1.59E-33 |
| Macrophage | Pcdha13 | 2 | 8.80E-37 | 1.04015659 | 0.432 | 0.144 | 1.76E-33 |
| Macrophage | Nova1 | 2 | 1.76E-36 | 0.31361985 | 0.423 | 0.153 | 3.51E-33 |
| Macrophage | Fgf10 | 2 | 2.37E-36 | 0.68908176 | 0.448 | 0.086 | 4.74E-33 |
| Macrophage | Plekha7 | 2 | 3.52E-36 | 0.3862775 | 0.651 | 0.222 | 7.04E-33 |
| Macrophage | Ccl2 | 2 | 4.06E-36 | 0.46202069 | 0.598 | 0.193 | 8.13E-33 |
| Macrophage | Nr4a1 | 2 | 4.23E-36 | 0.73596114 | 0.51 | 0.175 | 8.47E-33 |
| Macrophage | Cdh7 | 2 | 5.58E-36 | 2.62238985 | 0.469 | 0.097 | 1.12E-32 |
| Macrophage | Kcnd3 | 2 | 6.46E-36 | 0.32782558 | 0.552 | 0.161 | 1.29E-32 |

| | | | | | | | |
|---|---|---|---|---|---|---|---|
| Macrophage | Colec12 | 2 | 7.14E-36 | 1.52132932 | 0.975 | 0.706 | 1.43E-32 |
| Macrophage | Apobec1 | 2 | 1.51E-35 | 1.24568081 | 0.88 | 0.463 | 3.02E-32 |
| Macrophage | Adhfe1 | 2 | 3.30E-35 | 0.49002322 | 0.535 | 0.183 | 6.60E-32 |
| Macrophage | Cadm3 | 2 | 3.38E-35 | 0.40167843 | 0.461 | 0.155 | 6.76E-32 |
| Macrophage | AC096301.1 | 2 | 3.66E-35 | 0.52224357 | 0.419 | 0.167 | 7.32E-32 |
| Macrophage | Ptn | 2 | 3.67E-35 | 0.88381024 | 0.477 | 0.166 | 7.35E-32 |
| Macrophage | Prodh1 | 2 | 5.53E-35 | 0.92021906 | 0.689 | 0.274 | 1.11E-31 |
| Macrophage | Bace2 | 2 | 8.43E-35 | 0.77479537 | 0.481 | 0.203 | 1.69E-31 |
| Macrophage | Col14a1 | 2 | 1.80E-34 | 0.72751127 | 0.452 | 0.147 | 3.60E-31 |
| Macrophage | Abca17 | 2 | 4.64E-34 | 1.31441096 | 0.905 | 0.51 | 9.28E-31 |
| Macrophage | Anxa3 | 2 | 4.75E-34 | 0.56361791 | 0.502 | 0.163 | 9.51E-31 |
| Macrophage | Ltc4s | 2 | 5.70E-34 | 0.79258173 | 0.664 | 0.291 | 1.14E-30 |
| Macrophage | Dync1i1 | 2 | 7.29E-34 | 0.80220803 | 0.573 | 0.167 | 1.46E-30 |
| Macrophage | AC110709.2 | 2 | 3.73E-33 | 1.57202669 | 0.61 | 0.232 | 7.47E-30 |
| Macrophage | Pld5 | 2 | 4.58E-33 | 0.597503 | 0.647 | 0.248 | 9.15E-30 |
| Macrophage | Frmd5 | 2 | 6.90E-33 | 1.51406329 | 0.452 | 0.199 | 1.38E-29 |
| Macrophage | Mpped2 | 2 | 9.33E-33 | 0.66914957 | 0.519 | 0.177 | 1.87E-29 |
| Macrophage | Prrx1 | 2 | 2.37E-32 | 0.47586379 | 0.485 | 0.171 | 4.74E-29 |
| Macrophage | Prkaa2 | 2 | 7.99E-32 | 0.50401616 | 0.71 | 0.262 | 1.60E-28 |
| Macrophage | Ptpn3 | 2 | 9.21E-32 | 0.61935338 | 0.585 | 0.216 | 1.84E-28 |
| Macrophage | Eva1c | 2 | 1.11E-31 | 0.39966938 | 0.544 | 0.185 | 2.21E-28 |
| Macrophage | Slc16a10 | 2 | 1.13E-31 | 0.53550699 | 0.822 | 0.426 | 2.26E-28 |
| Macrophage | Syt14 | 2 | 1.19E-31 | 0.56817767 | 0.573 | 0.208 | 2.38E-28 |
| Macrophage | Slc22a23 | 2 | 2.28E-31 | 0.38722121 | 0.519 | 0.199 | 4.56E-28 |
| Macrophage | Ppargc1a | 2 | 2.29E-31 | 0.74551498 | 0.477 | 0.191 | 4.58E-28 |
| Macrophage | Mylk | 2 | 2.83E-31 | 0.25688517 | 0.444 | 0.166 | 5.67E-28 |
| Macrophage | Galnt17 | 2 | 3.08E-31 | 0.46766034 | 0.402 | 0.149 | 6.15E-28 |
| Macrophage | Mki67 | 2 | 1.45E-30 | 0.61763114 | 0.548 | 0.249 | 2.89E-27 |
| Macrophage | Lrrc4c | 2 | 3.17E-30 | 0.50825339 | 0.734 | 0.283 | 6.33E-27 |
| Macrophage | Nppb | 2 | 4.41E-30 | 0.58290802 | 0.606 | 0.245 | 8.81E-27 |
| Macrophage | Myoz2 | 2 | 9.91E-30 | 0.75327289 | 0.61 | 0.244 | 1.98E-26 |

| | | | | | | | |
|---|---|---|---|---|---|---|---|
| Macrophage | AC130940.1 | 2 | 1.36E-29 | 0.28119728 | 0.552 | 0.207 | 2.72E-26 |
| Macrophage | Sox6 | 2 | 2.23E-29 | 0.61133977 | 0.539 | 0.208 | 4.46E-26 |
| Macrophage | Pgm5 | 2 | 2.66E-29 | 0.3214564 | 0.515 | 0.205 | 5.32E-26 |
| Macrophage | Gulp1 | 2 | 3.43E-29 | 0.71613548 | 0.502 | 0.187 | 6.86E-26 |
| Macrophage | Sfxn5 | 2 | 4.55E-29 | 0.50532765 | 0.552 | 0.207 | 9.10E-26 |
| Macrophage | Ros1 | 2 | 4.63E-29 | 0.88103158 | 0.568 | 0.211 | 9.26E-26 |
| Macrophage | Tll1 | 2 | 6.83E-29 | 0.27525115 | 0.556 | 0.211 | 1.37E-25 |
| Macrophage | PCOLCE2 | 2 | 7.43E-29 | 0.81194773 | 0.498 | 0.181 | 1.49E-25 |
| Macrophage | Pde1a | 2 | 1.12E-28 | 0.87009571 | 0.452 | 0.189 | 2.23E-25 |
| Macrophage | Fstl5 | 2 | 1.65E-28 | 0.5993059 | 0.51 | 0.104 | 3.30E-25 |
| Macrophage | Thsd7b | 2 | 4.25E-28 | 0.93732745 | 0.407 | 0.133 | 8.51E-25 |
| Macrophage | Col5a3 | 2 | 7.58E-28 | 0.80434508 | 0.502 | 0.204 | 1.52E-24 |
| Macrophage | Ifitm3 | 2 | 9.30E-28 | 0.73426659 | 0.714 | 0.311 | 1.86E-24 |
| Macrophage | Spink8 | 2 | 1.18E-27 | 0.58696161 | 0.643 | 0.241 | 2.35E-24 |
| Macrophage | Aebp1 | 2 | 1.83E-27 | 0.61017374 | 0.502 | 0.227 | 3.66E-24 |
| Macrophage | Tbx20 | 2 | 3.18E-27 | 0.58465095 | 0.548 | 0.252 | 6.36E-24 |
| Macrophage | Nox4 | 2 | 4.55E-27 | 0.26865993 | 0.693 | 0.305 | 9.10E-24 |
| Macrophage | Mrc1 | 2 | 7.35E-27 | 0.90864834 | 0.967 | 0.701 | 1.47E-23 |
| Macrophage | AC134204.1 | 2 | 8.92E-27 | 0.33410439 | 0.581 | 0.284 | 1.78E-23 |
| Macrophage | Smpd3 | 2 | 1.04E-26 | 0.62709867 | 0.635 | 0.293 | 2.07E-23 |
| Macrophage | Col22a1 | 2 | 1.95E-26 | 1.398291 | 0.502 | 0.214 | 3.91E-23 |
| Macrophage | Pcsk5 | 2 | 2.01E-26 | 0.33777871 | 0.49 | 0.212 | 4.01E-23 |
| Macrophage | AABR070515 | 2 | 2.86E-26 | 1.04043101 | 0.589 | 0.284 | 5.72E-23 |
| Macrophage | Gucy1a1 | 2 | 9.34E-26 | 0.61173072 | 0.635 | 0.296 | 1.87E-22 |
| Macrophage | Iqub | 2 | 1.15E-25 | 0.27214329 | 0.527 | 0.276 | 2.31E-22 |
| Macrophage | Olr59 | 2 | 1.23E-25 | 0.31647999 | 0.461 | 0.094 | 2.47E-22 |
| Macrophage | LOC1009114 | 2 | 1.39E-25 | 0.50597649 | 0.407 | 0.102 | 2.77E-22 |
| Macrophage | Rspo1 | 2 | 1.89E-25 | 0.31456805 | 0.585 | 0.283 | 3.77E-22 |
| Macrophage | Mmrn1 | 2 | 2.04E-25 | 0.57930371 | 0.349 | 0.084 | 4.09E-22 |
| Macrophage | Slco2b1 | 2 | 5.93E-25 | 0.80546166 | 0.975 | 0.617 | 1.19E-21 |
| Macrophage | Stk26 | 2 | 7.88E-25 | 0.55382009 | 0.39 | 0.092 | 1.58E-21 |

| | | | | | | | |
|---|---|---|---|---|---|---|---|
| Macrophage | C1qa | 2 | 2.87E-24 | 0.75672222 | 0.701 | 0.302 | 5.74E-21 |
| Macrophage | Mgmt | 2 | 9.35E-24 | 0.76451641 | 0.817 | 0.406 | 1.87E-20 |
| Macrophage | Naca | 2 | 2.10E-23 | 0.30813292 | 0.71 | 0.311 | 4.19E-20 |
| Macrophage | Pf4 | 2 | 3.40E-23 | 0.64687689 | 0.51 | 0.256 | 6.80E-20 |
| Macrophage | Glb1l2 | 2 | 5.70E-23 | 0.3056758 | 0.423 | 0.121 | 1.14E-19 |
| Macrophage | Vash2 | 2 | 1.33E-22 | 1.2571186 | 0.427 | 0.108 | 2.65E-19 |
| Macrophage | Ahr | 2 | 1.64E-22 | 0.51928467 | 0.726 | 0.347 | 3.28E-19 |
| Macrophage | Rgs6 | 2 | 1.78E-22 | 0.94576808 | 0.531 | 0.246 | 3.57E-19 |
| Macrophage | Gramd1b | 2 | 3.25E-22 | 0.91124578 | 0.905 | 0.567 | 6.50E-19 |
| Macrophage | Adap2 | 2 | 8.04E-22 | 0.94618054 | 0.83 | 0.493 | 1.61E-18 |
| Macrophage | Asgr2 | 2 | 1.57E-21 | 0.56589262 | 0.793 | 0.397 | 3.15E-18 |
| Macrophage | Mx1 | 2 | 2.79E-21 | 0.53036506 | 0.647 | 0.318 | 5.58E-18 |
| Macrophage | Alox5 | 2 | 3.32E-21 | 0.76252547 | 0.722 | 0.432 | 6.63E-18 |
| Macrophage | Emilin2 | 2 | 5.30E-21 | 0.49953955 | 0.676 | 0.33 | 1.06E-17 |
| Macrophage | Abcc4 | 2 | 5.58E-21 | 0.41562533 | 0.581 | 0.28 | 1.12E-17 |
| Macrophage | Polr2m | 2 | 8.50E-21 | 0.27369273 | 0.585 | 0.264 | 1.70E-17 |
| Macrophage | Iqgap2 | 2 | 9.45E-21 | 0.99970911 | 0.817 | 0.487 | 1.89E-17 |
| Macrophage | Insr | 2 | 1.73E-20 | 0.2671435 | 0.697 | 0.306 | 3.47E-17 |
| Macrophage | Mertk | 2 | 2.32E-20 | 0.86478981 | 0.913 | 0.608 | 4.63E-17 |
| Macrophage | Nrcam | 2 | 2.56E-20 | 0.35416119 | 0.801 | 0.411 | 5.12E-17 |
| Macrophage | Eda | 2 | 2.95E-20 | 0.40048192 | 0.544 | 0.276 | 5.90E-17 |
| Macrophage | Gpcpd1 | 2 | 7.80E-20 | 0.27643881 | 0.668 | 0.306 | 1.56E-16 |
| Macrophage | Nuak1 | 2 | 9.22E-20 | 0.52108065 | 0.672 | 0.373 | 1.84E-16 |
| Macrophage | Tbc1d4 | 2 | 1.89E-19 | 0.89008742 | 0.934 | 0.651 | 3.78E-16 |
| Macrophage | Nkain3 | 2 | 2.64E-19 | 1.97318734 | 0.365 | 0.066 | 5.28E-16 |
| Macrophage | Mnda | 2 | 6.13E-19 | 0.38996649 | 0.747 | 0.404 | 1.23E-15 |
| Macrophage | Med12l | 2 | 1.03E-18 | 0.25167416 | 0.747 | 0.409 | 2.06E-15 |
| Macrophage | Msr1 | 2 | 1.44E-18 | 0.54134413 | 0.755 | 0.414 | 2.89E-15 |
| Macrophage | AABR070251 | 2 | 3.17E-18 | 0.76988241 | 0.759 | 0.392 | 6.35E-15 |
| Macrophage | Car10 | 2 | 4.59E-18 | 0.64565058 | 0.444 | 0.185 | 9.17E-15 |
| Macrophage | Deptor | 2 | 6.48E-18 | 0.47348311 | 0.759 | 0.38 | 1.30E-14 |

| | | | | | | | |
|---|---|---|---|---|---|---|---|
| Macrophage | Nfatc2 | 2 | 9.62E-18 | 0.64308433 | 0.722 | 0.408 | 1.92E-14 |
| Macrophage | Ubash3b | 2 | 2.56E-17 | 0.78644667 | 0.909 | 0.555 | 5.11E-14 |
| Macrophage | Adgre1 | 2 | 5.13E-17 | 0.64389142 | 0.822 | 0.468 | 1.03E-13 |
| Macrophage | Cdkn2b | 2 | 1.70E-16 | 0.98204914 | 0.427 | 0.126 | 3.40E-13 |
| Macrophage | Ston2 | 2 | 1.02E-15 | 0.46090974 | 0.788 | 0.479 | 2.04E-12 |
| Macrophage | Ivns1abp | 2 | 1.48E-14 | 0.45776125 | 0.751 | 0.44 | 2.96E-11 |
| Macrophage | Actn2 | 2 | 5.35E-14 | 0.3260457 | 0.622 | 0.362 | 1.07E-10 |
| Macrophage | Ctsb | 2 | 6.41E-14 | 0.73974228 | 0.822 | 0.558 | 1.28E-10 |
| Macrophage | Epsti1 | 2 | 1.28E-12 | 0.34508977 | 0.763 | 0.471 | 2.56E-09 |
| Macrophage | Pde7b | 2 | 5.06E-12 | 0.63098392 | 0.759 | 0.509 | 1.01E-08 |
| Macrophage | Slc7a8 | 2 | 1.10E-11 | 0.38694816 | 0.689 | 0.428 | 2.20E-08 |
| Macrophage | Gna15 | 2 | 2.15E-11 | 0.59060742 | 0.838 | 0.564 | 4.29E-08 |
| Macrophage | Adgrg6 | 2 | 8.65E-11 | 0.37602679 | 0.793 | 0.471 | 1.73E-07 |
| Macrophage | Gas7 | 2 | 4.48E-10 | 0.35728564 | 0.863 | 0.562 | 8.95E-07 |
| Macrophage | Stxbp6 | 2 | 6.46E-09 | 0.47133482 | 0.859 | 0.583 | 1.29E-05 |
| Macrophage | Nckap1l | 2 | 1.76E-07 | 0.25808638 | 0.817 | 0.549 | 0.00035159 |
| Macrophage | Ptk2b | 2 | 1.79E-07 | 0.29942796 | 0.805 | 0.541 | 0.00035809 |
| Macrophage | Dgkz | 2 | 2.54E-07 | 0.31034365 | 0.826 | 0.556 | 0.00050812 |
| Macrophage | Thbs1 | 3 | 1.30E-44 | 2.29133975 | 0.277 | 0.015 | 2.59E-41 |
| Macrophage | Cdh13 | 3 | 4.17E-26 | 2.40366346 | 0.549 | 0.214 | 8.33E-23 |
| Macrophage | Shank3 | 3 | 4.39E-26 | 2.84608568 | 0.503 | 0.142 | 8.79E-23 |
| Macrophage | Etl4 | 3 | 1.58E-23 | 2.21682789 | 0.569 | 0.255 | 3.16E-20 |
| Macrophage | Emcn | 3 | 4.58E-22 | 2.33790243 | 0.426 | 0.115 | 9.15E-19 |
| Macrophage | Ptprb | 3 | 5.16E-22 | 1.84155512 | 0.487 | 0.147 | 1.03E-18 |
| Macrophage | Adgrf5 | 3 | 1.33E-20 | 2.26693097 | 0.544 | 0.254 | 2.66E-17 |
| Macrophage | Tspan18 | 3 | 1.74E-20 | 2.61250047 | 0.39 | 0.105 | 3.48E-17 |
| Macrophage | Adgrg3 | 3 | 2.61E-19 | 0.86649221 | 0.462 | 0.205 | 5.22E-16 |
| Macrophage | Cyyr1 | 3 | 3.41E-17 | 2.01485257 | 0.467 | 0.2 | 6.83E-14 |
| Macrophage | Cald1 | 3 | 3.54E-17 | 1.40961009 | 0.456 | 0.182 | 7.07E-14 |
| Macrophage | Dnm3 | 3 | 1.86E-16 | 1.68636508 | 0.533 | 0.256 | 3.72E-13 |
| Macrophage | Clic5 | 3 | 2.45E-16 | 2.00936618 | 0.523 | 0.255 | 4.90E-13 |

| | | | | | | | |
|---|---|---|---|---|---|---|---|
| Macrophage | Mgp | 3 | 9.51E-16 | 1.41770769 | 0.656 | 0.365 | 1.90E-12 |
| Macrophage | Ntn1 | 3 | 2.32E-15 | 2.24419785 | 0.436 | 0.163 | 4.64E-12 |
| Macrophage | Fstl1 | 3 | 3.25E-15 | 1.58761468 | 0.482 | 0.2 | 6.50E-12 |
| Macrophage | Lmo7 | 3 | 1.01E-14 | 1.34727217 | 0.431 | 0.174 | 2.02E-11 |
| Macrophage | Mgll | 3 | 1.14E-14 | 1.39763353 | 0.646 | 0.389 | 2.28E-11 |
| Macrophage | Col1a1 | 3 | 1.38E-14 | 1.52129691 | 0.656 | 0.374 | 2.76E-11 |
| Macrophage | LOC1083526 | 3 | 1.70E-14 | 1.61343456 | 0.446 | 0.174 | 3.40E-11 |
| Macrophage | AABR070070 | 3 | 1.90E-14 | 1.36308091 | 0.585 | 0.311 | 3.80E-11 |
| Macrophage | Atp5f1b | 3 | 2.14E-14 | 1.29714229 | 0.672 | 0.419 | 4.27E-11 |
| Macrophage | Sparc | 3 | 2.24E-14 | 1.2851568 | 0.605 | 0.324 | 4.48E-11 |
| Macrophage | Usp2 | 3 | 2.69E-14 | 1.38060086 | 0.4 | 0.148 | 5.38E-11 |
| Macrophage | Nxn | 3 | 3.61E-14 | 2.01270776 | 0.549 | 0.28 | 7.22E-11 |
| Macrophage | Colgalt2 | 3 | 1.19E-13 | 2.10473811 | 0.405 | 0.106 | 2.37E-10 |
| Macrophage | AABR070275 | 3 | 5.41E-13 | 2.1288461 | 0.328 | 0.077 | 1.08E-09 |
| Macrophage | Scara5 | 3 | 2.40E-12 | 1.59944263 | 0.385 | 0.126 | 4.80E-09 |
| Macrophage | Col8a1 | 3 | 1.75E-11 | 1.0321362 | 0.585 | 0.325 | 3.50E-08 |
| Macrophage | Fbn1 | 3 | 4.66E-11 | 1.05040728 | 0.626 | 0.372 | 9.31E-08 |
| Macrophage | Fbln5 | 3 | 5.64E-11 | 2.19156992 | 0.344 | 0.093 | 1.13E-07 |
| Macrophage | Tmeff2 | 3 | 8.25E-10 | 1.04829691 | 0.431 | 0.18 | 1.65E-06 |
| Macrophage | AABR070033 | 5 | 1.02E-56 | 5.41504671 | 0.373 | 0.037 | 2.04E-53 |
| Macrophage | Snca | 5 | 2.37E-54 | 3.50368284 | 0.373 | 0.038 | 4.73E-51 |
| Macrophage | Cd163 | 5 | 1.05E-50 | 2.88080662 | 0.88 | 0.361 | 2.10E-47 |
| Macrophage | F13a11 | 5 | 3.35E-38 | 2.04084341 | 0.887 | 0.514 | 6.71E-35 |
| Macrophage | Ptger3 | 5 | 5.61E-37 | 2.70458641 | 0.373 | 0.058 | 1.12E-33 |
| Macrophage | Mctp1 | 5 | 3.71E-31 | 2.75188358 | 0.761 | 0.36 | 7.42E-28 |
| Macrophage | Ppp2r2b | 5 | 3.89E-31 | 2.88526875 | 0.683 | 0.19 | 7.78E-28 |
| Macrophage | Kcnt2 | 5 | 1.56E-30 | 1.50509519 | 0.951 | 0.607 | 3.12E-27 |
| Macrophage | Col19a1 | 5 | 5.74E-29 | 3.94777427 | 0.641 | 0.227 | 1.15E-25 |
| Macrophage | Fam71f1 | 5 | 4.02E-25 | 2.32103718 | 0.373 | 0.094 | 8.04E-22 |
| Macrophage | Ccl12 | 5 | 5.80E-25 | 1.03191356 | 0.401 | 0.103 | 1.16E-21 |
| Macrophage | Lamc3 | 5 | 3.35E-23 | 0.6202958 | 0.359 | 0.067 | 6.71E-20 |

| | | | | | | | |
|---|---|---|---|---|---|---|---|
| Macrophage | Lyve11 | 5 | 3.34E-22 | 2.21332458 | 0.697 | 0.364 | 6.67E-19 |
| Macrophage | Cpvl | 5 | 1.25E-21 | 4.62251993 | 0.31 | 0.037 | 2.51E-18 |
| Macrophage | Adam23 | 5 | 1.01E-20 | 3.47392826 | 0.577 | 0.282 | 2.03E-17 |
| Macrophage | Slco2b11 | 5 | 1.57E-20 | 1.12394611 | 0.915 | 0.644 | 3.14E-17 |
| Macrophage | Prc1 | 5 | 8.84E-20 | 0.90998618 | 0.352 | 0.075 | 1.77E-16 |
| Macrophage | Smoc1 | 5 | 1.29E-19 | 1.8528883 | 0.303 | 0.036 | 2.59E-16 |
| Macrophage | Prkcz | 5 | 2.85E-19 | 4.90634072 | 0.43 | 0.138 | 5.70E-16 |
| Macrophage | Bmx | 5 | 4.04E-18 | 1.91449655 | 0.352 | 0.099 | 8.08E-15 |
| Macrophage | Ovol1 | 5 | 2.48E-15 | 1.04019842 | 0.359 | 0.061 | 4.96E-12 |
| Macrophage | Il1rl1 | 5 | 5.97E-13 | 2.04155223 | 0.521 | 0.218 | 1.19E-09 |
| Macrophage | Fbxw101 | 5 | 3.79E-12 | 2.7646719 | 0.401 | 0.116 | 7.59E-09 |
| Macrophage | Fgf101 | 5 | 8.35E-12 | 1.14330448 | 0.373 | 0.115 | 1.67E-08 |
| Macrophage | Olr591 | 5 | 7.49E-10 | 0.92377347 | 0.387 | 0.123 | 1.50E-06 |
| Macrophage | LOC1009114 | 5 | 3.91E-07 | 0.65928501 | 0.394 | 0.121 | 0.00078139 |
| Macrophage | Nkd2 | 5 | 4.02E-07 | 0.56512796 | 0.324 | 0.066 | 0.00080397 |
| Macrophage | Cldn4 | 6 | 1.08E-111 | 5.39498195 | 0.432 | 0.014 | 2.17E-108 |
| Macrophage | Gpr143 | 6 | 7.44E-107 | 6.04800175 | 0.534 | 0.029 | 1.49E-103 |
| Macrophage | Krt17 | 6 | 2.41E-105 | 3.07191175 | 0.458 | 0.016 | 4.82E-102 |
| Macrophage | Emx2 | 6 | 1.50E-99 | 6.192854 | 0.492 | 0.026 | 3.00E-96 |
| Macrophage | Ankrd34c | 6 | 3.00E-95 | 5.95584784 | 0.593 | 0.05 | 6.00E-92 |
| Macrophage | LOC1083484 | 6 | 5.34E-90 | 5.76406515 | 0.441 | 0.028 | 1.07E-86 |
| Macrophage | AABR070354 | 6 | 1.24E-87 | 7.50244785 | 0.297 | 0.006 | 2.48E-84 |
| Macrophage | Hist1h4m | 6 | 2.64E-87 | 5.0418691 | 0.508 | 0.037 | 5.27E-84 |
| Macrophage | AABR070322 | 6 | 3.68E-86 | 3.45066343 | 0.508 | 0.036 | 7.36E-83 |
| Macrophage | Ccl9 | 6 | 1.73E-82 | 2.43891657 | 0.508 | 0.039 | 3.45E-79 |
| Macrophage | Fgf7 | 6 | 2.79E-82 | 2.38503175 | 0.466 | 0.025 | 5.57E-79 |
| Macrophage | Esco2 | 6 | 2.57E-79 | 3.37186036 | 0.5 | 0.039 | 5.15E-76 |
| Macrophage | Susd5 | 6 | 3.67E-78 | 4.48748938 | 0.432 | 0.018 | 7.33E-75 |
| Macrophage | Ropn1l | 6 | 8.97E-78 | 6.11961888 | 0.415 | 0.025 | 1.79E-74 |
| Macrophage | AABR070322 | 6 | 1.41E-76 | 5.51181162 | 0.661 | 0.086 | 2.82E-73 |
| Macrophage | AABR070635 | 6 | 2.37E-71 | 2.82364166 | 0.5 | 0.046 | 4.73E-68 |

| | | | | | | | |
|---|---|---|---|---|---|---|---|
| Macrophage | AABR070339 | 6 | 2.28E-70 | 2.38911502 | 0.475 | 0.036 | 4.56E-67 |
| Macrophage | Rab15 | 6 | 3.20E-68 | 4.03225772 | 0.347 | 0.048 | 6.41E-65 |
| Macrophage | AC128394.2 | 6 | 6.36E-67 | 4.69081731 | 0.483 | 0.014 | 1.27E-63 |
| Macrophage | Slc47a1 | 6 | 2.13E-65 | 5.05691896 | 0.559 | 0.056 | 4.25E-62 |
| Macrophage | Cela1 | 6 | 1.21E-62 | 4.43433494 | 0.627 | 0.111 | 2.42E-59 |
| Macrophage | Tspan33 | 6 | 2.52E-62 | 1.47933113 | 0.466 | 0.057 | 5.04E-59 |
| Macrophage | Dhrs9 | 6 | 5.61E-61 | 3.88904565 | 0.559 | 0.05 | 1.12E-57 |
| Macrophage | Dab1 | 6 | 1.46E-60 | 3.78269926 | 0.297 | 0.013 | 2.93E-57 |
| Macrophage | Ptcd3 | 6 | 5.00E-59 | 3.47140213 | 0.983 | 0.483 | 1.00E-55 |
| Macrophage | Gata3 | 6 | 4.06E-57 | 1.42740881 | 0.483 | 0.042 | 8.12E-54 |
| Macrophage | Atp1b4 | 6 | 8.19E-56 | 2.60295434 | 0.441 | 0.014 | 1.64E-52 |
| Macrophage | Espl1 | 6 | 1.03E-51 | 1.88380167 | 0.508 | 0.076 | 2.07E-48 |
| Macrophage | Atp6v0d2 | 6 | 1.13E-51 | 4.10399199 | 0.822 | 0.139 | 2.27E-48 |
| Macrophage | Ass1 | 6 | 4.85E-47 | 2.59824467 | 0.373 | 0.039 | 9.69E-44 |
| Macrophage | Padi2 | 6 | 7.90E-47 | 3.89473297 | 0.449 | 0.039 | 1.58E-43 |
| Macrophage | NEWGENE-2 | 6 | 1.07E-45 | 1.82155026 | 0.407 | 0.053 | 2.14E-42 |
| Macrophage | Hpgd | 6 | 6.26E-45 | 1.59673107 | 0.415 | 0.039 | 1.25E-41 |
| Macrophage | Gpnmb1 | 6 | 1.43E-44 | 2.45587888 | 0.941 | 0.47 | 2.86E-41 |
| Macrophage | Susd4 | 6 | 3.28E-43 | 2.71612322 | 0.475 | 0.036 | 6.56E-40 |
| Macrophage | Mt3 | 6 | 4.65E-43 | 2.07124069 | 0.424 | 0.031 | 9.29E-40 |
| Macrophage | Spp1 | 6 | 2.22E-42 | 4.06904258 | 0.831 | 0.256 | 4.44E-39 |
| Macrophage | Tm4sf19 | 6 | 1.11E-40 | 4.72238544 | 0.525 | 0.136 | 2.22E-37 |
| Macrophage | Anpep | 6 | 4.01E-40 | 2.473068 | 0.881 | 0.386 | 8.01E-37 |
| Macrophage | Ptchd1 | 6 | 6.73E-39 | 3.16126279 | 0.551 | 0.084 | 1.35E-35 |
| Macrophage | Lilrb4 | 6 | 1.33E-37 | 2.0271382 | 0.975 | 0.616 | 2.66E-34 |
| Macrophage | Mreg | 6 | 4.03E-37 | 3.09266952 | 0.585 | 0.135 | 8.05E-34 |
| Macrophage | Lgals3 | 6 | 7.81E-37 | 2.38005641 | 0.915 | 0.473 | 1.56E-33 |
| Macrophage | Lilrb3 | 6 | 1.43E-33 | 4.07221412 | 0.712 | 0.326 | 2.86E-30 |
| Macrophage | Rab7b | 6 | 1.93E-32 | 2.77191385 | 0.805 | 0.365 | 3.85E-29 |
| Macrophage | Hk3 | 6 | 6.71E-32 | 2.93704772 | 0.763 | 0.276 | 1.34E-28 |
| Macrophage | Mmp12 | 6 | 1.28E-31 | 4.48596447 | 0.669 | 0.108 | 2.57E-28 |

| | | | | | | | |
|---|---|---|---|---|---|---|---|
| Macrophage | Tmem26 | 6 | 7.47E-30 | 3.6271171 | 0.729 | 0.161 | 1.49E-26 |
| Macrophage | Pamr1 | 6 | 9.83E-30 | 4.89517388 | 0.314 | 0.05 | 1.97E-26 |
| Macrophage | AABR070311 | 6 | 1.35E-29 | 3.70692933 | 0.627 | 0.121 | 2.69E-26 |
| Macrophage | Ifitm10 | 6 | 2.19E-29 | 1.08849812 | 0.322 | 0.029 | 4.37E-26 |
| Macrophage | Polq | 6 | 3.63E-29 | 1.12929938 | 0.483 | 0.048 | 7.27E-26 |
| Macrophage | Sorcs2 | 6 | 2.66E-28 | 2.69674966 | 0.39 | 0.132 | 5.32E-25 |
| Macrophage | Prima1 | 6 | 2.88E-28 | 1.45062637 | 0.432 | 0.091 | 5.77E-25 |
| Macrophage | Atp1a3 | 6 | 7.65E-28 | 4.12823503 | 0.373 | 0.096 | 1.53E-24 |
| Macrophage | Dgat2 | 6 | 9.64E-28 | 2.15806468 | 0.542 | 0.172 | 1.93E-24 |
| Macrophage | Alcam | 6 | 5.86E-27 | 1.79439389 | 0.941 | 0.667 | 1.17E-23 |
| Macrophage | AC120486.1( | 6 | 5.57E-26 | 1.04347903 | 0.475 | 0.104 | 1.11E-22 |
| Macrophage | AABR070605 | 6 | 1.33E-25 | 0.88282019 | 0.508 | 0.092 | 2.67E-22 |
| Macrophage | Tec | 6 | 1.37E-25 | 1.71415099 | 0.924 | 0.586 | 2.75E-22 |
| Macrophage | Myo18b | 6 | 9.00E-25 | 2.48323667 | 0.754 | 0.314 | 1.80E-21 |
| Macrophage | Nyap2 | 6 | 1.67E-24 | 4.77608111 | 0.466 | 0.155 | 3.34E-21 |
| Macrophage | Sema3c | 6 | 1.72E-24 | 2.21897459 | 0.737 | 0.211 | 3.44E-21 |
| Macrophage | Smpdl3a | 6 | 1.33E-23 | 1.73548951 | 0.814 | 0.448 | 2.66E-20 |
| Macrophage | Edil3 | 6 | 2.45E-23 | 2.39077642 | 0.72 | 0.274 | 4.90E-20 |
| Macrophage | Pparg | 6 | 7.10E-23 | 2.2267303 | 0.712 | 0.246 | 1.42E-19 |
| Macrophage | Ank3 | 6 | 8.68E-23 | 1.73428997 | 0.797 | 0.331 | 1.74E-19 |
| Macrophage | Matk | 6 | 1.09E-22 | 2.24871286 | 0.729 | 0.36 | 2.18E-19 |
| Macrophage | Slc7a2 | 6 | 4.16E-22 | 2.78635536 | 0.644 | 0.247 | 8.32E-19 |
| Macrophage | Mitf | 6 | 6.39E-22 | 1.41523869 | 0.975 | 0.717 | 1.28E-18 |
| Macrophage | Kcnk2 | 6 | 9.93E-22 | 1.66522309 | 0.339 | 0.027 | 1.99E-18 |
| Macrophage | Metrn | 6 | 1.24E-21 | 2.21632082 | 0.492 | 0.158 | 2.49E-18 |
| Macrophage | Cybb | 6 | 4.27E-21 | 1.60566775 | 0.873 | 0.537 | 8.54E-18 |
| Macrophage | Depdc1 | 6 | 5.04E-21 | 1.22433648 | 0.466 | 0.134 | 1.01E-17 |
| Macrophage | Pld3 | 6 | 2.62E-20 | 2.17859764 | 0.703 | 0.357 | 5.23E-17 |
| Macrophage | Aox3 | 6 | 2.81E-20 | 2.02325608 | 0.39 | 0.054 | 5.62E-17 |
| Macrophage | Mamdc2 | 6 | 3.14E-20 | 2.77011464 | 0.695 | 0.295 | 6.27E-17 |
| Macrophage | Tspan5 | 6 | 5.90E-20 | 2.18815482 | 0.754 | 0.419 | 1.18E-16 |

| | | | | | | | |
|---|---|---|---|---|---|---|---|
| Macrophage | RGD1305807 | 6 | 7.50E-20 | 2.10195555 | 0.678 | 0.207 | 1.50E-16 |
| Macrophage | Rassf4 | 6 | 2.32E-19 | 2.61829157 | 0.686 | 0.293 | 4.65E-16 |
| Macrophage | Cd300lb | 6 | 6.32E-19 | 1.70729076 | 0.703 | 0.406 | 1.26E-15 |
| Macrophage | Clec7a | 6 | 7.51E-19 | 1.80644487 | 0.712 | 0.39 | 1.50E-15 |
| Macrophage | Runx2 | 6 | 8.81E-19 | 1.45516362 | 0.746 | 0.384 | 1.76E-15 |
| Macrophage | Ptgs2 | 6 | 9.52E-19 | 0.37855162 | 0.381 | 0.099 | 1.90E-15 |
| Macrophage | Fabp5 | 6 | 1.80E-18 | 2.23327745 | 0.619 | 0.254 | 3.60E-15 |
| Macrophage | AABR070311 | 6 | 1.88E-18 | 0.70082436 | 0.398 | 0.055 | 3.75E-15 |
| Macrophage | Nr5a2 | 6 | 3.10E-18 | 1.81156199 | 0.398 | 0.039 | 6.20E-15 |
| Macrophage | Lrrtm4 | 6 | 1.04E-17 | 1.64454393 | 0.398 | 0.128 | 2.09E-14 |
| Macrophage | Itga2 | 6 | 1.14E-17 | 1.51902542 | 0.322 | 0.069 | 2.28E-14 |
| Macrophage | Asb9 | 6 | 1.35E-17 | 0.58422419 | 0.373 | 0.028 | 2.70E-14 |
| Macrophage | Comp | 6 | 2.19E-17 | 2.87069403 | 0.339 | 0.017 | 4.38E-14 |
| Macrophage | Slc4a5 | 6 | 3.00E-17 | 1.17574414 | 0.517 | 0.182 | 6.00E-14 |
| Macrophage | Penk | 6 | 3.27E-17 | 2.51896502 | 0.492 | 0.033 | 6.55E-14 |
| Macrophage | Ctsk | 6 | 3.41E-17 | 2.7699798 | 0.466 | 0.064 | 6.82E-14 |
| Macrophage | Col12a1 | 6 | 6.14E-17 | 1.2427595 | 0.517 | 0.064 | 1.23E-13 |
| Macrophage | Cyba | 6 | 3.05E-16 | 1.0936759 | 0.771 | 0.434 | 6.09E-13 |
| Macrophage | Fermt3 | 6 | 3.91E-16 | 1.14287331 | 0.898 | 0.596 | 7.81E-13 |
| Macrophage | Dgki | 6 | 3.95E-16 | 3.28048338 | 0.534 | 0.267 | 7.90E-13 |
| Macrophage | Limch1 | 6 | 4.00E-16 | 1.61375526 | 0.686 | 0.203 | 8.01E-13 |
| Macrophage | Ube2ql1 | 6 | 7.64E-16 | 2.10042705 | 0.525 | 0.166 | 1.53E-12 |
| Macrophage | Adgrg31 | 6 | 2.20E-15 | 0.94340265 | 0.492 | 0.215 | 4.39E-12 |
| Macrophage | Itgam | 6 | 3.23E-15 | 1.25062419 | 0.805 | 0.384 | 6.47E-12 |
| Macrophage | Prph | 6 | 5.43E-15 | 1.92069857 | 0.466 | 0.068 | 1.09E-11 |
| Macrophage | AABR070030 | 6 | 5.54E-15 | 1.68439539 | 0.432 | 0.053 | 1.11E-11 |
| Macrophage | Atf3 | 6 | 6.46E-15 | 2.78069648 | 0.542 | 0.273 | 1.29E-11 |
| Macrophage | Itpkb | 6 | 7.51E-15 | 1.47664401 | 0.873 | 0.549 | 1.50E-11 |
| Macrophage | Robo2 | 6 | 7.67E-15 | 2.67588658 | 0.492 | 0.079 | 1.53E-11 |
| Macrophage | Gpc3 | 6 | 8.16E-15 | 1.15771933 | 0.441 | 0.047 | 1.63E-11 |
| Macrophage | Fn1 | 6 | 1.00E-14 | 1.6700414 | 0.669 | 0.251 | 2.00E-11 |

| | | | | | | | |
|---|---|---|---|---|---|---|---|
| Macrophage | Mgll1 | 6 | 2.36E-14 | 1.97516079 | 0.712 | 0.396 | 4.73E-11 |
| Macrophage | Rgs1 | 6 | 4.13E-14 | 2.86614387 | 0.483 | 0.2 | 8.27E-11 |
| Macrophage | Psap | 6 | 8.21E-14 | 0.9727291 | 0.822 | 0.408 | 1.64E-10 |
| Macrophage | Ap1s3 | 6 | 1.00E-13 | 0.66674874 | 0.517 | 0.085 | 2.01E-10 |
| Macrophage | Apoe | 6 | 1.09E-13 | 1.86090975 | 0.831 | 0.391 | 2.18E-10 |
| Macrophage | Cacna1a | 6 | 2.11E-13 | 1.99198756 | 0.653 | 0.384 | 4.23E-10 |
| Macrophage | Rab11fip1 | 6 | 2.16E-13 | 2.18260784 | 0.5 | 0.15 | 4.33E-10 |
| Macrophage | S100a6 | 6 | 2.68E-13 | 1.14106749 | 0.627 | 0.254 | 5.36E-10 |
| Macrophage | Pi16 | 6 | 7.92E-13 | 1.17466249 | 0.678 | 0.335 | 1.58E-09 |
| Macrophage | Lpl | 6 | 1.01E-12 | 1.65493196 | 0.636 | 0.311 | 2.01E-09 |
| Macrophage | Ccnf | 6 | 1.09E-12 | 1.70354445 | 0.508 | 0.096 | 2.18E-09 |
| Macrophage | Lyz2 | 6 | 1.40E-12 | 1.48492603 | 0.686 | 0.409 | 2.80E-09 |
| Macrophage | Grm8 | 6 | 2.31E-12 | 1.02187126 | 0.305 | 0.031 | 4.63E-09 |
| Macrophage | Gm2a | 6 | 3.15E-12 | 1.79681217 | 0.466 | 0.184 | 6.30E-09 |
| Macrophage | Trim7 | 6 | 3.92E-12 | 1.18570823 | 0.5 | 0.109 | 7.84E-09 |
| Macrophage | Specc1 | 6 | 3.99E-12 | 1.04691849 | 0.864 | 0.583 | 7.97E-09 |
| Macrophage | Angpt2 | 6 | 5.03E-12 | 0.41336387 | 0.424 | 0.063 | 1.01E-08 |
| Macrophage | Syt141 | 6 | 7.24E-12 | 2.1581897 | 0.5 | 0.241 | 1.45E-08 |
| Macrophage | AC111831.1 | 6 | 9.08E-12 | 1.31510444 | 0.398 | 0.074 | 1.82E-08 |
| Macrophage | Lgi4 | 6 | 1.51E-11 | 1.09033165 | 0.475 | 0.087 | 3.01E-08 |
| Macrophage | Tnnt1 | 6 | 1.78E-11 | 1.0092071 | 0.492 | 0.185 | 3.56E-08 |
| Macrophage | Kif22 | 6 | 1.94E-11 | 1.18309422 | 0.5 | 0.141 | 3.88E-08 |
| Macrophage | Kcnn4 | 6 | 2.58E-11 | 1.43700709 | 0.644 | 0.375 | 5.17E-08 |
| Macrophage | Itgb4 | 6 | 2.69E-11 | 0.61862247 | 0.39 | 0.076 | 5.37E-08 |
| Macrophage | Bard1 | 6 | 3.26E-11 | 0.41151352 | 0.492 | 0.193 | 6.51E-08 |
| Macrophage | Gpx1 | 6 | 3.57E-11 | 1.55494037 | 0.593 | 0.22 | 7.15E-08 |
| Macrophage | AABR070010 | 6 | 4.03E-11 | 1.68644575 | 0.322 | 0.047 | 8.07E-08 |
| Macrophage | Kcnj5 | 6 | 6.88E-11 | 1.6887113 | 0.5 | 0.136 | 1.38E-07 |
| Macrophage | Bcar3 | 6 | 7.37E-11 | 1.68284569 | 0.508 | 0.194 | 1.47E-07 |
| Macrophage | Prdx5 | 6 | 1.33E-10 | 2.06556529 | 0.466 | 0.207 | 2.66E-07 |
| Macrophage | Trim55 | 6 | 1.46E-10 | 0.51992406 | 0.627 | 0.317 | 2.92E-07 |

| | | | | | | | |
|---|---|---|---|---|---|---|---|
| Macrophage | Cenpf | 6 | 1.75E-10 | 1.47360587 | 0.483 | 0.123 | 3.50E-07 |
| Macrophage | Shroom2 | 6 | 2.46E-10 | 1.11292666 | 0.534 | 0.2 | 4.91E-07 |
| Macrophage | Fam227b | 6 | 3.04E-10 | 1.37885329 | 0.551 | 0.222 | 6.08E-07 |
| Macrophage | Lmod1 | 6 | 3.12E-10 | 0.69797283 | 0.508 | 0.192 | 6.23E-07 |
| Macrophage | Syk | 6 | 7.84E-10 | 0.79297485 | 0.898 | 0.578 | 1.57E-06 |
| Macrophage | Srl | 6 | 9.55E-10 | 0.46942386 | 0.525 | 0.238 | 1.91E-06 |
| Macrophage | AABR070171 | 6 | 1.17E-09 | 0.56883082 | 0.407 | 0.066 | 2.33E-06 |
| Macrophage | Svep1 | 6 | 1.46E-09 | 0.2918299 | 0.475 | 0.087 | 2.91E-06 |
| Macrophage | Ptprv | 6 | 1.68E-09 | 1.35005715 | 0.373 | 0.12 | 3.36E-06 |
| Macrophage | LOC1003644 | 6 | 1.69E-09 | 0.44222609 | 0.653 | 0.277 | 3.38E-06 |
| Macrophage | Me1 | 6 | 1.86E-09 | 0.44188556 | 0.542 | 0.206 | 3.71E-06 |
| Macrophage | Cdkn1a | 6 | 3.40E-09 | 2.14899408 | 0.424 | 0.152 | 6.80E-06 |
| Macrophage | Nrcam1 | 6 | 4.29E-09 | 0.33349536 | 0.703 | 0.448 | 8.58E-06 |
| Macrophage | Lrp4 | 6 | 4.43E-09 | 0.85848847 | 0.559 | 0.159 | 8.86E-06 |
| Macrophage | Lhfpl2 | 6 | 4.79E-09 | 0.88181391 | 0.678 | 0.398 | 9.57E-06 |
| Macrophage | Galnt16 | 6 | 6.13E-09 | 0.88506129 | 0.458 | 0.131 | 1.23E-05 |
| Macrophage | Tpm2 | 6 | 6.21E-09 | 0.69394147 | 0.424 | 0.085 | 1.24E-05 |
| Macrophage | Rbm38 | 6 | 8.04E-09 | 1.3664179 | 0.483 | 0.159 | 1.61E-05 |
| Macrophage | Dnm31 | 6 | 1.00E-08 | 0.51529137 | 0.61 | 0.263 | 2.01E-05 |
| Macrophage | Oxct1 | 6 | 1.25E-08 | 0.58491206 | 0.686 | 0.386 | 2.51E-05 |
| Macrophage | Errfi1 | 6 | 1.48E-08 | 1.30218366 | 0.559 | 0.27 | 2.97E-05 |
| Macrophage | Lgals1 | 6 | 1.70E-08 | 0.32954286 | 0.686 | 0.385 | 3.40E-05 |
| Macrophage | Trem2 | 6 | 1.93E-08 | 0.69414025 | 0.585 | 0.303 | 3.86E-05 |
| Macrophage | Homer1 | 6 | 1.99E-08 | 0.74155885 | 0.559 | 0.251 | 3.98E-05 |
| Macrophage | Adra1a | 6 | 2.42E-08 | 0.61381923 | 0.492 | 0.176 | 4.84E-05 |
| Macrophage | Npnt | 6 | 2.45E-08 | 0.97466663 | 0.525 | 0.175 | 4.89E-05 |
| Macrophage | Astn2 | 6 | 2.83E-08 | 0.49219448 | 0.466 | 0.145 | 5.67E-05 |
| Macrophage | Pkia | 6 | 5.03E-08 | 0.67181253 | 0.551 | 0.273 | 0.00010053 |
| Macrophage | Abcb4 | 6 | 5.59E-08 | 0.38873304 | 0.703 | 0.3 | 0.0001118 |
| Macrophage | Cadm1 | 6 | 5.84E-08 | 1.01391002 | 0.593 | 0.309 | 0.00011671 |
| Macrophage | Dkk3 | 6 | 6.02E-08 | 0.61962811 | 0.508 | 0.212 | 0.00012048 |

| | | | | | | | |
|---|---|---|---|---|---|---|---|
| Macrophage | Clmp | 6 | 1.18E-07 | 0.3091226 | 0.551 | 0.145 | 0.0002366 |
| Macrophage | Slc37a2 | 6 | 1.25E-07 | 0.68676615 | 0.619 | 0.367 | 0.00024999 |
| Macrophage | Slc20a1 | 6 | 1.96E-07 | 0.58531575 | 0.576 | 0.287 | 0.00039244 |
| Macrophage | Ankrd6 | 6 | 2.38E-07 | 0.96981973 | 0.517 | 0.26 | 0.00047626 |
| Macrophage | Lpcat2 | 6 | 2.81E-07 | 0.48203728 | 0.763 | 0.449 | 0.00056197 |
| Macrophage | Itga4 | 6 | 3.66E-07 | 0.7179896 | 0.754 | 0.478 | 0.00073278 |
| Macrophage | Pfkfb3 | 6 | 4.18E-07 | 0.97660824 | 0.619 | 0.319 | 0.00083694 |
| Macrophage | Cd300a | 6 | 5.03E-07 | 0.56779216 | 0.847 | 0.587 | 0.00100554 |
| Macrophage | LOC1003614 | 6 | 9.39E-07 | 0.26745943 | 0.593 | 0.298 | 0.00187781 |
| Macrophage | Enpp1 | 6 | 1.04E-06 | 0.45775926 | 0.407 | 0.134 | 0.00207956 |
| Macrophage | RGD1559482 | 6 | 2.40E-06 | 0.39286396 | 0.729 | 0.452 | 0.00480566 |
| Macrophage | Nrp2 | 6 | 3.51E-06 | 0.31718783 | 0.822 | 0.543 | 0.00702303 |
| Macrophage | Mis18bp1 | 6 | 1.67E-05 | 1.16328546 | 0.458 | 0.188 | 0.03344996 |
| Macrophage | Krt18 | 7 | 3.79E-74 | 6.62246431 | 0.356 | 0.011 | 7.58E-71 |
| Macrophage | Tnfrsf13b | 7 | 1.08E-61 | 5.35626069 | 0.356 | 0.017 | 2.16E-58 |
| Macrophage | Retn | 7 | 2.47E-61 | 4.6868759 | 0.356 | 0.016 | 4.93E-58 |
| Macrophage | Cd3e | 7 | 1.34E-55 | 5.95548593 | 0.342 | 0.018 | 2.67E-52 |
| Macrophage | Tnni2 | 7 | 4.01E-54 | 6.3794008 | 0.356 | 0.017 | 8.02E-51 |
| Macrophage | Stum | 7 | 7.68E-54 | 4.91400123 | 0.411 | 0.03 | 1.54E-50 |
| Macrophage | Atf7ip2 | 7 | 3.14E-44 | 4.31605119 | 0.356 | 0.012 | 6.28E-41 |
| Macrophage | Khdrbs2 | 7 | 1.06E-43 | 2.10861293 | 0.342 | 0.023 | 2.12E-40 |
| Macrophage | Ajap1 | 7 | 6.70E-36 | 1.58666375 | 0.301 | 0.02 | 1.34E-32 |
| Macrophage | Gcsam | 7 | 5.66E-30 | 7.47017417 | 0.452 | 0.078 | 1.13E-26 |
| Macrophage | Tssk3 | 7 | 3.54E-29 | 6.11953986 | 0.356 | 0.044 | 7.07E-26 |
| Macrophage | Clec9a | 7 | 1.24E-27 | 4.31584496 | 0.411 | 0.066 | 2.48E-24 |
| Macrophage | Fcrl5 | 7 | 6.99E-27 | 5.05756464 | 0.356 | 0.056 | 1.40E-23 |
| Macrophage | Ciita | 7 | 6.82E-25 | 3.5289406 | 0.781 | 0.397 | 1.36E-21 |
| Macrophage | Pde6h | 7 | 9.45E-25 | 2.11424208 | 0.37 | 0.039 | 1.89E-21 |
| Macrophage | Mybpc2 | 7 | 1.18E-24 | 3.18994021 | 0.356 | 0.014 | 2.36E-21 |
| Macrophage | RT1-DOa | 7 | 1.50E-24 | 4.69109325 | 0.342 | 0.082 | 2.99E-21 |
| Macrophage | LOC24906 | 7 | 2.48E-24 | 4.4162892 | 0.397 | 0.039 | 4.96E-21 |

| | | | | | | | |
|---|---|---|---|---|---|---|---|
| Macrophage | Flt3 | 7 | 1.27E-23 | 6.13810468 | 0.685 | 0.153 | 2.54E-20 |
| Macrophage | Cd74 | 7 | 1.06E-22 | 1.97882556 | 0.918 | 0.441 | 2.12E-19 |
| Macrophage | Insc | 7 | 1.76E-22 | 2.21431278 | 0.342 | 0.018 | 3.52E-19 |
| Macrophage | Fam189a2 | 7 | 2.05E-22 | 1.72035722 | 0.356 | 0.028 | 4.11E-19 |
| Macrophage | Exph5 | 7 | 3.89E-21 | 3.11301647 | 0.411 | 0.077 | 7.77E-18 |
| Macrophage | RT1-Db1 | 7 | 1.40E-20 | 2.68076629 | 0.781 | 0.287 | 2.79E-17 |
| Macrophage | Olfm2 | 7 | 6.64E-20 | 3.09355404 | 0.37 | 0.031 | 1.33E-16 |
| Macrophage | Meox1 | 7 | 1.05E-19 | 1.56126036 | 0.356 | 0.057 | 2.11E-16 |
| Macrophage | Hrh2 | 7 | 3.46E-19 | 4.70670278 | 0.479 | 0.124 | 6.91E-16 |
| Macrophage | AABR070410 | 7 | 8.96E-19 | 4.90768761 | 0.671 | 0.192 | 1.79E-15 |
| Macrophage | AABR070637 | 7 | 5.77E-18 | 7.36025784 | 0.384 | 0.068 | 1.15E-14 |
| Macrophage | Kif20a | 7 | 5.51E-17 | 2.26512838 | 0.329 | 0.029 | 1.10E-13 |
| Macrophage | Fcrla | 7 | 1.09E-16 | 3.57224891 | 0.384 | 0.093 | 2.19E-13 |
| Macrophage | Sh3gl3 | 7 | 3.60E-16 | 3.28780958 | 0.356 | 0.051 | 7.19E-13 |
| Macrophage | Cdh1 | 7 | 1.48E-15 | 6.74627031 | 0.452 | 0.02 | 2.96E-12 |
| Macrophage | Slc35f1 | 7 | 1.83E-15 | 2.86184554 | 0.384 | 0.056 | 3.65E-12 |
| Macrophage | Sbspon | 7 | 1.40E-14 | 4.02634639 | 0.37 | 0.104 | 2.80E-11 |
| Macrophage | Crlf1 | 7 | 1.46E-12 | 2.20650008 | 0.37 | 0.053 | 2.91E-09 |
| Macrophage | Treml4 | 7 | 5.25E-12 | 3.41993662 | 0.397 | 0.117 | 1.05E-08 |
| Macrophage | RT1-Da | 7 | 8.31E-12 | 2.26198384 | 0.603 | 0.22 | 1.66E-08 |
| Macrophage | Fcnb | 7 | 8.51E-12 | 6.12948771 | 0.411 | 0.155 | 1.70E-08 |
| Macrophage | Adcy5 | 7 | 3.31E-11 | 4.36988463 | 0.63 | 0.33 | 6.61E-08 |
| Macrophage | Vwc2 | 7 | 1.16E-10 | 4.46840992 | 0.301 | 0.013 | 2.33E-07 |
| Macrophage | Dgkg | 7 | 4.20E-10 | 4.3291921 | 0.548 | 0.243 | 8.41E-07 |
| Macrophage | Slco3a1 | 7 | 8.46E-10 | 2.26659965 | 0.671 | 0.318 | 1.69E-06 |
| Macrophage | Zfp366 | 7 | 1.95E-09 | 3.78902806 | 0.575 | 0.196 | 3.89E-06 |
| Macrophage | Prg4 | 7 | 5.22E-09 | 1.29615695 | 0.315 | 0.025 | 1.04E-05 |
| Macrophage | Bcl11a | 7 | 5.82E-09 | 3.57457364 | 0.562 | 0.256 | 1.16E-05 |
| Macrophage | Galnt3 | 7 | 8.17E-09 | 3.96194574 | 0.37 | 0.038 | 1.63E-05 |
| Macrophage | Tenm4 | 7 | 1.09E-08 | 2.00744899 | 0.37 | 0.029 | 2.18E-05 |
| Macrophage | Fcgr2b | 7 | 1.56E-08 | 2.55925877 | 0.644 | 0.349 | 3.13E-05 |

| | | | | | | | |
|---|---|---|---|---|---|---|---|
| Macrophage | Gpd1 | 7 | 2.07E-08 | 1.61379109 | 0.356 | 0.039 | 4.14E-05 |
| Macrophage | Ccser1 | 7 | 3.00E-08 | 3.56941397 | 0.466 | 0.132 | 6.00E-05 |
| Macrophage | Dscaml1 | 7 | 3.98E-08 | 5.29534025 | 0.521 | 0.208 | 7.96E-05 |
| Macrophage | Gpr141 | 7 | 5.72E-08 | 2.67628703 | 0.425 | 0.148 | 0.0001144 |
| Macrophage | Ckap2 | 7 | 7.55E-08 | 1.96815493 | 0.384 | 0.134 | 0.00015092 |
| Macrophage | RT1-Ba | 7 | 1.49E-07 | 2.01952266 | 0.534 | 0.272 | 0.0002977 |
| Macrophage | Nhs | 7 | 1.88E-07 | 2.05341362 | 0.37 | 0.118 | 0.00037565 |
| Macrophage | Il18r1 | 7 | 1.99E-07 | 5.43141116 | 0.329 | 0.069 | 0.00039878 |
| Macrophage | Lef1 | 7 | 2.80E-07 | 1.11342182 | 0.397 | 0.116 | 0.00056054 |
| Macrophage | Spon1 | 7 | 3.08E-07 | 2.17007247 | 0.534 | 0.176 | 0.00061548 |
| Macrophage | Mob3b | 7 | 8.43E-07 | 2.82573506 | 0.534 | 0.191 | 0.00168536 |
| Macrophage | Parm1 | 7 | 1.27E-06 | 3.69689437 | 0.452 | 0.152 | 0.0025306 |
| Macrophage | Baiap2l1 | 7 | 1.55E-06 | 2.96740344 | 0.493 | 0.124 | 0.00310119 |
| Macrophage | Tmem150c | 7 | 3.74E-06 | 3.20810663 | 0.411 | 0.12 | 0.00747548 |
| Macrophage | Niban1 | 7 | 4.41E-06 | 1.76710001 | 0.685 | 0.421 | 0.00881371 |
| Macrophage | Nr4a3 | 7 | 7.60E-06 | 2.0440287 | 0.397 | 0.1 | 0.0152096 |
| Macrophage | Cspg4 | 7 | 7.85E-06 | 1.10580488 | 0.356 | 0.106 | 0.01569407 |
| Macrophage | Cdca2 | 7 | 8.75E-06 | 0.48108859 | 0.356 | 0.026 | 0.01750908 |
| Macrophage | Il1b | 7 | 1.02E-05 | 3.56741914 | 0.438 | 0.187 | 0.02043863 |
| Macrophage | Cadps2 | 7 | 1.32E-05 | 1.97165605 | 0.521 | 0.096 | 0.02642375 |
| Macrophage | Cr2 | 8 | 4.64E-63 | 13.3361978 | 0.275 | 0.003 | 9.27E-60 |
| Macrophage | Ajap11 | 8 | 3.00E-41 | 5.18909759 | 0.373 | 0.022 | 6.00E-38 |
| Macrophage | Pou2af1 | 8 | 1.07E-40 | 9.92043173 | 0.314 | 0.033 | 2.15E-37 |
| Macrophage | Sell | 8 | 3.20E-34 | 7.08576337 | 0.353 | 0.034 | 6.41E-31 |
| Macrophage | Cd3e1 | 8 | 3.71E-31 | 1.86181378 | 0.314 | 0.023 | 7.43E-28 |
| Macrophage | Pax5 | 8 | 3.21E-28 | 7.09589408 | 0.549 | 0.097 | 6.42E-25 |
| Macrophage | Tnfrsf13b1 | 8 | 6.58E-28 | 3.96528828 | 0.294 | 0.023 | 1.32E-24 |
| Macrophage | Stum1 | 8 | 4.34E-26 | 2.56807232 | 0.353 | 0.037 | 8.69E-23 |
| Macrophage | Pde6h1 | 8 | 6.09E-26 | 7.33161643 | 0.431 | 0.041 | 1.22E-22 |
| Macrophage | Tnni21 | 8 | 8.41E-26 | 2.8789026 | 0.294 | 0.024 | 1.68E-22 |
| Macrophage | Ikzf3 | 8 | 4.84E-25 | 6.4673034 | 0.451 | 0.071 | 9.68E-22 |

| | | | | | | | |
|---|---|---|---|---|---|---|---|
| Macrophage | Retn1 | 8 | 1.18E-24 | 4.46811549 | 0.275 | 0.022 | 2.37E-21 |
| Macrophage | Il34 | 8 | 2.19E-22 | 5.39187432 | 0.549 | 0.131 | 4.38E-19 |
| Macrophage | Tmem163 | 8 | 3.77E-21 | 6.93184579 | 0.49 | 0.107 | 7.54E-18 |
| Macrophage | Fcrl51 | 8 | 8.18E-21 | 3.27010761 | 0.373 | 0.059 | 1.64E-17 |
| Macrophage | Gabrb2 | 8 | 5.04E-19 | 3.6902746 | 0.412 | 0.081 | 1.01E-15 |
| Macrophage | Ighm | 8 | 8.69E-19 | 5.51522602 | 0.725 | 0.202 | 1.74E-15 |
| Macrophage | Aff31 | 8 | 1.69E-18 | 4.02826857 | 0.725 | 0.119 | 3.39E-15 |
| Macrophage | Klrc2 | 8 | 9.60E-18 | 2.73584359 | 0.353 | 0.068 | 1.92E-14 |
| Macrophage | Fcrla1 | 8 | 1.29E-17 | 5.54922734 | 0.471 | 0.094 | 2.58E-14 |
| Macrophage | RT1-DOa1 | 8 | 1.67E-17 | 3.69527169 | 0.353 | 0.085 | 3.34E-14 |
| Macrophage | Kif20a1 | 8 | 5.06E-17 | 1.44250789 | 0.333 | 0.033 | 1.01E-13 |
| Macrophage | Bank1 | 8 | 6.74E-17 | 6.17042207 | 0.706 | 0.292 | 1.35E-13 |
| Macrophage | Olfm21 | 8 | 6.98E-17 | 2.08747464 | 0.333 | 0.037 | 1.40E-13 |
| Macrophage | Itk | 8 | 3.37E-15 | 3.49846891 | 0.294 | 0.043 | 6.75E-12 |
| Macrophage | Meox11 | 8 | 6.79E-15 | 2.1813539 | 0.333 | 0.061 | 1.36E-11 |
| Macrophage | Plac8 | 8 | 1.56E-14 | 4.8197396 | 0.392 | 0.11 | 3.12E-11 |
| Macrophage | Rerg | 8 | 3.21E-14 | 4.12224045 | 0.725 | 0.226 | 6.42E-11 |
| Macrophage | Cspg41 | 8 | 5.21E-13 | 3.48555674 | 0.431 | 0.107 | 1.04E-09 |
| Macrophage | Cmahp | 8 | 2.65E-12 | 5.80642401 | 0.471 | 0.075 | 5.31E-09 |
| Macrophage | Slc35f11 | 8 | 3.93E-12 | 2.51933711 | 0.333 | 0.062 | 7.86E-09 |
| Macrophage | Mob3b1 | 8 | 8.52E-12 | 2.55179437 | 0.647 | 0.192 | 1.70E-08 |
| Macrophage | St8sia1 | 8 | 1.36E-11 | 1.93454902 | 0.392 | 0.1 | 2.72E-08 |
| Macrophage | Gcsam1 | 8 | 2.16E-11 | 0.89206413 | 0.353 | 0.085 | 4.31E-08 |
| Macrophage | Ano1 | 8 | 8.70E-11 | 4.30291512 | 0.588 | 0.205 | 1.74E-07 |
| Macrophage | Abcc9 | 8 | 1.83E-10 | 3.10800498 | 0.647 | 0.303 | 3.67E-07 |
| Macrophage | Pdgfrb | 8 | 6.58E-10 | 2.3841496 | 0.627 | 0.274 | 1.32E-06 |
| Macrophage | Pde3a | 8 | 5.01E-09 | 2.47153844 | 0.608 | 0.329 | 1.00E-05 |
| Macrophage | Bcl11a1 | 8 | 5.52E-09 | 4.26014595 | 0.549 | 0.261 | 1.10E-05 |
| Macrophage | Timp3 | 8 | 6.47E-09 | 2.16308158 | 0.608 | 0.229 | 1.29E-05 |
| Macrophage | Ntng1 | 8 | 7.51E-09 | 3.28178177 | 0.471 | 0.097 | 1.50E-05 |
| Macrophage | Slco4a1 | 8 | 8.50E-09 | 1.10190963 | 0.373 | 0.074 | 1.70E-05 |

| | | | | | | | |
|---|---|---|---|---|---|---|---|
| Macrophage | Gcnt2 | 8 | 9.89E-09 | 2.75911313 | 0.588 | 0.254 | 1.98E-05 |
| Macrophage | Cobll1 | 8 | 1.35E-08 | 2.75590763 | 0.569 | 0.226 | 2.70E-05 |
| Macrophage | Gucy1a2 | 8 | 1.78E-08 | 3.83159494 | 0.412 | 0.116 | 3.55E-05 |
| Macrophage | Adgrl3 | 8 | 2.45E-08 | 3.83422717 | 0.451 | 0.192 | 4.89E-05 |
| Macrophage | Mark1 | 8 | 2.62E-08 | 2.82074936 | 0.51 | 0.175 | 5.23E-05 |
| Macrophage | Fam163a | 8 | 2.69E-08 | 3.74932275 | 0.392 | 0.109 | 5.38E-05 |
| Macrophage | Ncald | 8 | 3.02E-08 | 1.20477934 | 0.784 | 0.398 | 6.04E-05 |
| Macrophage | Sgip1 | 8 | 5.26E-08 | 3.46432853 | 0.529 | 0.146 | 0.00010527 |
| Macrophage | AABR070275 | 8 | 5.79E-08 | 2.91723917 | 0.51 | 0.093 | 0.00011588 |
| Macrophage | Ephb2 | 8 | 6.70E-08 | 2.92957631 | 0.451 | 0.087 | 0.00013395 |
| Macrophage | Slamf6 | 8 | 7.98E-08 | 2.97928313 | 0.314 | 0.063 | 0.0001596 |
| Macrophage | Hs3st3a1 | 8 | 8.01E-08 | 5.10438006 | 0.412 | 0.14 | 0.00016016 |
| Macrophage | Ndst3 | 8 | 8.34E-08 | 2.41410227 | 0.392 | 0.041 | 0.00016678 |
| Macrophage | Cnksr2 | 8 | 9.49E-08 | 3.13198563 | 0.412 | 0.029 | 0.00018973 |
| Macrophage | Has2 | 8 | 1.02E-07 | 1.88159403 | 0.392 | 0.06 | 0.00020412 |
| Macrophage | Gli2 | 8 | 1.03E-07 | 2.44181979 | 0.569 | 0.203 | 0.00020647 |
| Macrophage | Kcnj8 | 8 | 1.30E-07 | 3.8261565 | 0.431 | 0.134 | 0.00026034 |
| Macrophage | Gpr176 | 8 | 1.35E-07 | 3.23064977 | 0.431 | 0.08 | 0.00026989 |
| Macrophage | Prrx11 | 8 | 1.47E-07 | 1.95118053 | 0.529 | 0.205 | 0.00029427 |
| Macrophage | Heyl1 | 8 | 2.47E-07 | 2.02367803 | 0.529 | 0.172 | 0.00049439 |
| Macrophage | Rgs5 | 8 | 2.53E-07 | 3.49148314 | 0.431 | 0.155 | 0.00050505 |
| Macrophage | Adamts16 | 8 | 2.96E-07 | 2.31804849 | 0.353 | 0.044 | 0.00059213 |
| Macrophage | Kynu | 8 | 3.07E-07 | 4.67819452 | 0.529 | 0.203 | 0.0006134 |
| Macrophage | Plcg2 | 8 | 3.15E-07 | 1.58159586 | 0.804 | 0.454 | 0.00062981 |
| Macrophage | Zfp385d | 8 | 3.28E-07 | 1.50855654 | 0.49 | 0.216 | 0.00065518 |
| Macrophage | Cd741 | 8 | 4.60E-07 | 1.08142272 | 0.784 | 0.452 | 0.00092051 |
| Macrophage | Kcnma1 | 8 | 6.53E-07 | 4.27075296 | 0.412 | 0.13 | 0.00130642 |
| Macrophage | Ebf2 | 8 | 8.04E-07 | 2.13413896 | 0.569 | 0.273 | 0.00160844 |
| Macrophage | Cdh4 | 8 | 1.05E-06 | 1.74093785 | 0.294 | 0.012 | 0.00209277 |
| Macrophage | Gpd11 | 8 | 1.23E-06 | 0.83196145 | 0.314 | 0.045 | 0.00246521 |
| Macrophage | Mcam | 8 | 2.72E-06 | 2.67136304 | 0.549 | 0.242 | 0.00544764 |

| | | | | | | | |
|---|---|---|---|---|---|---|---|
| Macrophage | Ccdc80 | 8 | 6.53E-06 | 1.18880281 | 0.647 | 0.256 | 0.01306417 |
| Macrophage | Inpp4b | 8 | 6.59E-06 | 2.07516924 | 0.451 | 0.18 | 0.01317277 |
| Macrophage | Bcar31 | 8 | 7.50E-06 | 2.27133885 | 0.51 | 0.206 | 0.01500843 |
| Macrophage | Cald11 | 8 | 9.09E-06 | 1.46699703 | 0.549 | 0.202 | 0.01817033 |
| Macrophage | Mgmt1 | 8 | 1.01E-05 | 1.16512133 | 0.706 | 0.455 | 0.02015563 |
| Macrophage | Agmo | 8 | 1.14E-05 | 2.76640841 | 0.392 | 0.102 | 0.02276197 |
| Macrophage | Hvcn1 | 8 | 1.21E-05 | 1.38997359 | 0.725 | 0.431 | 0.02421097 |
| Macrophage | Slc12a2 | 8 | 1.32E-05 | 1.7284525 | 0.549 | 0.238 | 0.02644435 |
| Macrophage | Cdca21 | 8 | 1.48E-05 | 1.98813101 | 0.314 | 0.032 | 0.02956065 |
| Endothelial Cell | Xcl1 | 0 | 1.04E-266 | 4.41706115 | 0.301 | 0.027 | 2.09E-263 |
| Endothelial Cell | Tm4sf19 | 0 | 5.32E-127 | 4.5839384 | 0.429 | 0.1 | 1.06E-123 |
| Endothelial Cell | Btnl9 | 0 | 3.06E-67 | 1.49932439 | 0.632 | 0.351 | 6.11E-64 |
| Endothelial Cell | Penk | 0 | 2.59E-35 | 1.99911658 | 0.38 | 0.128 | 5.18E-32 |
| Endothelial Cell | AABR070428 | 1 | 2.03E-80 | 0.45309337 | 0.097 | 0.365 | 4.06E-77 |
| Endothelial Cell | Ninj2 | 1 | 3.53E-71 | 0.38833361 | 0.155 | 0.44 | 7.05E-68 |
| Endothelial Cell | Cst6 | 2 | 0 | 2.5557471 | 0.342 | 0.01 | 0 |
| Endothelial Cell | Fshr | 2 | 0 | 2.56668539 | 0.335 | 0.025 | 0 |
| Endothelial Cell | Col28a1 | 2 | 3.92E-253 | 2.28421175 | 0.274 | 0.012 | 7.84E-250 |
| Endothelial Cell | LOC688282 | 2 | 1.32E-230 | 2.2132227 | 0.382 | 0.063 | 2.63E-227 |
| Endothelial Cell | Col13a1 | 2 | 1.22E-228 | 3.38073921 | 0.488 | 0.093 | 2.45E-225 |
| Endothelial Cell | Insr | 2 | 1.17E-222 | 3.36876368 | 0.801 | 0.427 | 2.34E-219 |
| Endothelial Cell | Cass4 | 2 | 1.01E-196 | 2.19692519 | 0.365 | 0.028 | 2.02E-193 |
| Endothelial Cell | Inhba | 2 | 4.78E-191 | 2.88254934 | 0.423 | 0.134 | 9.57E-188 |
| Endothelial Cell | AABR070015 | 2 | 1.07E-182 | 2.18783318 | 0.326 | 0.029 | 2.13E-179 |
| Endothelial Cell | Tgfa | 2 | 2.11E-182 | 2.98899683 | 0.406 | 0.052 | 4.21E-179 |
| Endothelial Cell | Vwf | 2 | 1.21E-171 | 1.62556716 | 0.845 | 0.504 | 2.41E-168 |
| Endothelial Cell | Pde1c | 2 | 1.60E-131 | 2.06813799 | 0.755 | 0.45 | 3.19E-128 |
| Endothelial Cell | Lmx1a | 2 | 1.49E-129 | 0.96515891 | 0.396 | 0.064 | 2.98E-126 |
| Endothelial Cell | Kif20a | 2 | 1.20E-119 | 1.14793734 | 0.41 | 0.141 | 2.41E-116 |
| Endothelial Cell | Slco2a1 | 2 | 6.23E-118 | 2.03377655 | 0.618 | 0.257 | 1.25E-114 |
| Endothelial Cell | Cntn3 | 2 | 4.07E-113 | 4.43689562 | 0.456 | 0.19 | 8.13E-110 |

| | | | | | | | |
|---|---|---|---|---|---|---|---|
| Endothelial Cell | Samd5 | 2 | 4.67E-99 | 1.79634119 | 0.709 | 0.407 | 9.34E-96 |
| Endothelial Cell | Zbtb7c | 2 | 1.35E-84 | 3.05456146 | 0.53 | 0.221 | 2.69E-81 |
| Endothelial Cell | Lhx6 | 2 | 5.62E-79 | 2.21518212 | 0.608 | 0.317 | 1.12E-75 |
| Endothelial Cell | Unc5c | 2 | 1.56E-76 | 1.7644804 | 0.649 | 0.348 | 3.12E-73 |
| Endothelial Cell | Necab1 | 2 | 1.94E-74 | 0.99890308 | 0.467 | 0.2 | 3.87E-71 |
| Endothelial Cell | Abi3bp | 2 | 5.54E-74 | 1.31660452 | 0.705 | 0.413 | 1.11E-70 |
| Endothelial Cell | Emilin2 | 2 | 3.62E-69 | 2.50213981 | 0.449 | 0.172 | 7.24E-66 |
| Endothelial Cell | Tbx5 | 2 | 2.14E-65 | 1.14730549 | 0.456 | 0.072 | 4.28E-62 |
| Endothelial Cell | Tpx2 | 2 | 2.84E-63 | 0.64647078 | 0.486 | 0.167 | 5.68E-60 |
| Endothelial Cell | Hspb3 | 2 | 2.02E-54 | 0.50674665 | 0.426 | 0.163 | 4.04E-51 |
| Endothelial Cell | Pak3 | 2 | 3.76E-51 | 2.04259622 | 0.362 | 0.098 | 7.51E-48 |
| Endothelial Cell | Olfml2a | 2 | 1.42E-38 | 0.97520204 | 0.51 | 0.26 | 2.84E-35 |
| Endothelial Cell | Ctsz | 2 | 1.88E-34 | 1.0290996 | 0.497 | 0.244 | 3.75E-31 |
| Endothelial Cell | Dupd1 | 3 | 0 | 2.54095184 | 0.333 | 0.014 | 0 |
| Endothelial Cell | Galnt14 | 3 | 4.25E-281 | 5.03019472 | 0.61 | 0.1 | 8.50E-278 |
| Endothelial Cell | Kif14 | 3 | 3.71E-231 | 1.69492698 | 0.327 | 0.046 | 7.42E-228 |
| Endothelial Cell | Cyp4f18 | 3 | 2.52E-180 | 1.53327892 | 0.33 | 0.048 | 5.05E-177 |
| Endothelial Cell | Depdc1 | 3 | 1.04E-170 | 0.95025567 | 0.381 | 0.058 | 2.09E-167 |
| Endothelial Cell | Ska1 | 3 | 3.04E-164 | 1.24404596 | 0.321 | 0.068 | 6.09E-161 |
| Endothelial Cell | Kit | 3 | 1.62E-160 | 2.88929772 | 0.748 | 0.372 | 3.23E-157 |
| Endothelial Cell | Nrp2 | 3 | 1.11E-153 | 1.61231066 | 0.894 | 0.599 | 2.22E-150 |
| Endothelial Cell | Myo16 | 3 | 2.74E-152 | 1.59771669 | 0.503 | 0.138 | 5.48E-149 |
| Endothelial Cell | Ano4 | 3 | 3.90E-149 | 2.3657649 | 0.773 | 0.375 | 7.80E-146 |
| Endothelial Cell | Ltk | 3 | 1.17E-136 | 1.0316684 | 0.371 | 0.08 | 2.34E-133 |
| Endothelial Cell | Rspo1 | 3 | 1.97E-129 | 3.15050791 | 0.517 | 0.09 | 3.93E-126 |
| Endothelial Cell | Hpd | 3 | 1.25E-107 | 0.61754226 | 0.316 | 0.043 | 2.51E-104 |
| Endothelial Cell | Rab15 | 3 | 4.83E-107 | 1.23636601 | 0.329 | 0.036 | 9.67E-104 |
| Endothelial Cell | Fosb | 3 | 5.94E-105 | 1.0419931 | 0.315 | 0.05 | 1.19E-101 |
| Endothelial Cell | Ttc29 | 3 | 2.46E-98 | 0.66015885 | 0.408 | 0.076 | 4.93E-95 |
| Endothelial Cell | Nox4 | 3 | 3.62E-95 | 2.04497475 | 0.698 | 0.417 | 7.24E-92 |
| Endothelial Cell | Ephb1 | 3 | 1.82E-83 | 2.48570373 | 0.652 | 0.377 | 3.63E-80 |

| | | | | | | | |
|---|---|---|---|---|---|---|---|
| Endothelial Cell | Slc4a5 | 3 | 5.29E-83 | 0.81622221 | 0.314 | 0.045 | 1.06E-79 |
| Endothelial Cell | Aox3 | 3 | 4.99E-81 | 0.77112807 | 0.418 | 0.111 | 9.98E-78 |
| Endothelial Cell | Slc1a1 | 3 | 2.21E-79 | 2.1378761 | 0.662 | 0.409 | 4.42E-76 |
| Endothelial Cell | AABR070179 | 3 | 2.19E-77 | 0.48798266 | 0.492 | 0.216 | 4.38E-74 |
| Endothelial Cell | Knl1 | 3 | 2.82E-71 | 1.07887706 | 0.49 | 0.234 | 5.63E-68 |
| Endothelial Cell | Col7a1 | 3 | 3.60E-70 | 1.78016651 | 0.351 | 0.083 | 7.21E-67 |
| Endothelial Cell | Ltc4s | 3 | 6.68E-70 | 0.30661298 | 0.442 | 0.163 | 1.34E-66 |
| Endothelial Cell | Ckap2 | 3 | 9.64E-70 | 1.47659151 | 0.422 | 0.136 | 1.93E-66 |
| Endothelial Cell | Mki67 | 3 | 1.14E-67 | 1.13744703 | 0.56 | 0.24 | 2.28E-64 |
| Endothelial Cell | Ablim2 | 3 | 1.51E-66 | 0.66026497 | 0.556 | 0.253 | 3.01E-63 |
| Endothelial Cell | Mcam | 3 | 2.51E-66 | 1.23130008 | 0.761 | 0.474 | 5.01E-63 |
| Endothelial Cell | Msr1 | 3 | 2.75E-61 | 0.32378785 | 0.317 | 0.047 | 5.50E-58 |
| Endothelial Cell | Osr1 | 3 | 1.72E-58 | 0.31078632 | 0.491 | 0.19 | 3.43E-55 |
| Endothelial Cell | Dlgap1 | 3 | 1.85E-58 | 1.22835681 | 0.597 | 0.27 | 3.69E-55 |
| Endothelial Cell | Pla2r1 | 3 | 2.16E-58 | 1.5909927 | 0.506 | 0.249 | 4.31E-55 |
| Endothelial Cell | Angpt2 | 3 | 2.81E-54 | 2.93498402 | 0.536 | 0.285 | 5.63E-51 |
| Endothelial Cell | Bcat1 | 3 | 2.03E-51 | 0.73065754 | 0.509 | 0.199 | 4.05E-48 |
| Endothelial Cell | Ccl21 | 3 | 3.32E-48 | 0.35896857 | 0.563 | 0.261 | 6.64E-45 |
| Endothelial Cell | Tmem200a | 3 | 1.39E-47 | 0.73750793 | 0.525 | 0.218 | 2.77E-44 |
| Endothelial Cell | Dlgap5 | 3 | 1.08E-42 | 0.39671016 | 0.546 | 0.225 | 2.16E-39 |
| Endothelial Cell | Ptprc | 3 | 3.31E-42 | 0.31755261 | 0.564 | 0.25 | 6.62E-39 |
| Endothelial Cell | Akap12 | 3 | 1.31E-41 | 0.71958058 | 0.572 | 0.274 | 2.61E-38 |
| Endothelial Cell | Prc1 | 3 | 4.36E-41 | 0.33399057 | 0.354 | 0.093 | 8.71E-38 |
| Endothelial Cell | Tbx20 | 3 | 5.35E-41 | 0.41329666 | 0.602 | 0.298 | 1.07E-37 |
| Endothelial Cell | Lyz2 | 3 | 1.29E-38 | 0.36862853 | 0.452 | 0.181 | 2.58E-35 |
| Endothelial Cell | Slc47a1 | 3 | 4.96E-35 | 1.01363889 | 0.4 | 0.115 | 9.92E-32 |
| Endothelial Cell | Fgf12 | 3 | 3.98E-32 | 0.68842428 | 0.533 | 0.275 | 7.95E-29 |
| Endothelial Cell | Trhde | 3 | 1.17E-31 | 1.08204293 | 0.395 | 0.091 | 2.35E-28 |
| Endothelial Cell | Bzw2 | 3 | 2.85E-28 | 0.30273015 | 0.63 | 0.325 | 5.71E-25 |
| Endothelial Cell | Gata4 | 3 | 1.19E-26 | 0.35206991 | 0.496 | 0.245 | 2.38E-23 |
| Endothelial Cell | Fgf23 | 4 | 0 | 5.07957034 | 0.495 | 0.021 | 0 |

| | | | | | | | |
|---|---|---|---|---|---|---|---|
| Endothelial Cell | Stac | 4 | 0 | 4.45375577 | 0.429 | 0.007 | 0 |
| Endothelial Cell | Cxcl14 | 4 | 0 | 5.66090384 | 0.419 | 0.006 | 0 |
| Endothelial Cell | Itprid1 | 4 | 0 | 5.30700728 | 0.386 | 0.012 | 0 |
| Endothelial Cell | Reg3b | 4 | 0 | 3.09836993 | 0.329 | 0.005 | 0 |
| Endothelial Cell | Adarb2 | 4 | 3.93E-300 | 1.18576536 | 0.328 | 0.005 | 7.86E-297 |
| Endothelial Cell | Selp | 4 | 3.43E-270 | 5.80352129 | 0.515 | 0.068 | 6.86E-267 |
| Endothelial Cell | Slc6a19 | 4 | 1.08E-257 | 2.17512756 | 0.439 | 0.038 | 2.16E-254 |
| Endothelial Cell | Vwf1 | 4 | 2.02E-228 | 2.48360023 | 0.947 | 0.51 | 4.03E-225 |
| Endothelial Cell | Kynu | 4 | 3.38E-226 | 1.88749547 | 0.331 | 0.035 | 6.76E-223 |
| Endothelial Cell | AABR070058 | 4 | 4.11E-205 | 4.49636612 | 0.446 | 0.081 | 8.21E-202 |
| Endothelial Cell | Rasa4 | 4 | 2.18E-175 | 2.09743851 | 0.925 | 0.586 | 4.37E-172 |
| Endothelial Cell | Il1r1 | 4 | 4.83E-174 | 1.6898181 | 0.958 | 0.7 | 9.66E-171 |
| Endothelial Cell | Abi3bp1 | 4 | 2.00E-172 | 2.66383041 | 0.862 | 0.412 | 4.01E-169 |
| Endothelial Cell | Met | 4 | 9.85E-170 | 4.10926991 | 0.57 | 0.13 | 1.97E-166 |
| Endothelial Cell | Thsd7a | 4 | 1.18E-164 | 1.73651923 | 0.951 | 0.68 | 2.35E-161 |
| Endothelial Cell | Blk | 4 | 9.01E-159 | 2.37723372 | 0.363 | 0.064 | 1.80E-155 |
| Endothelial Cell | Plvap | 4 | 4.49E-153 | 3.81159183 | 0.373 | 0.095 | 8.98E-150 |
| Endothelial Cell | Tll1 | 4 | 8.67E-149 | 4.04057978 | 0.748 | 0.29 | 1.73E-145 |
| Endothelial Cell | Pgm5 | 4 | 3.38E-143 | 1.97887604 | 0.894 | 0.564 | 6.75E-140 |
| Endothelial Cell | Ropn1l | 4 | 1.75E-142 | 2.89620139 | 0.291 | 0.022 | 3.50E-139 |
| Endothelial Cell | Mctp1 | 4 | 7.79E-136 | 3.97382094 | 0.717 | 0.305 | 1.56E-132 |
| Endothelial Cell | Slco2a11 | 4 | 2.43E-133 | 3.0101468 | 0.688 | 0.268 | 4.86E-130 |
| Endothelial Cell | Mill1 | 4 | 7.03E-132 | 1.70727918 | 0.314 | 0.029 | 1.41E-128 |
| Endothelial Cell | Samd3 | 4 | 3.55E-131 | 3.75295059 | 0.357 | 0.003 | 7.10E-128 |
| Endothelial Cell | Tc2n | 4 | 3.62E-127 | 1.15962363 | 0.304 | 0.041 | 7.25E-124 |
| Endothelial Cell | Nuak1 | 4 | 1.02E-121 | 1.86296449 | 0.84 | 0.502 | 2.03E-118 |
| Endothelial Cell | Ralgapa2 | 4 | 1.84E-121 | 1.67302363 | 0.911 | 0.66 | 3.69E-118 |
| Endothelial Cell | Cnksr2 | 4 | 4.91E-121 | 1.20631401 | 0.466 | 0.061 | 9.83E-118 |
| Endothelial Cell | Ldb2 | 4 | 1.45E-119 | 2.03579749 | 0.778 | 0.368 | 2.90E-116 |
| Endothelial Cell | AABR070720 | 4 | 2.60E-116 | 1.88640984 | 0.34 | 0.038 | 5.19E-113 |
| Endothelial Cell | Susd4 | 4 | 2.10E-107 | 3.77195835 | 0.442 | 0.083 | 4.20E-104 |

| | | | | | | | |
|---|---|---|---|---|---|---|---|
| Endothelial Cell | Sulf1 | 4 | 1.88E-106 | 1.26400993 | 0.842 | 0.476 | 3.77E-103 |
| Endothelial Cell | Atp6v0d2 | 4 | 3.11E-106 | 1.14838016 | 0.291 | 0.021 | 6.22E-103 |
| Endothelial Cell | Gabbr2 | 4 | 5.08E-102 | 1.45147266 | 0.508 | 0.179 | 1.02E-98 |
| Endothelial Cell | Plac8 | 4 | 1.17E-95 | 2.37893664 | 0.391 | 0.122 | 2.33E-92 |
| Endothelial Cell | Asap3 | 4 | 3.09E-94 | 2.83896835 | 0.683 | 0.405 | 6.17E-91 |
| Endothelial Cell | Lhx61 | 4 | 1.61E-89 | 1.73766314 | 0.709 | 0.321 | 3.22E-86 |
| Endothelial Cell | Ptk2b | 4 | 3.09E-87 | 3.03790881 | 0.622 | 0.324 | 6.18E-84 |
| Endothelial Cell | AABR070410 | 4 | 4.00E-87 | 2.15212178 | 0.568 | 0.175 | 8.00E-84 |
| Endothelial Cell | Lgals3 | 4 | 2.49E-86 | 2.53336777 | 0.645 | 0.199 | 4.98E-83 |
| Endothelial Cell | Nlrp3 | 4 | 1.43E-84 | 0.56219973 | 0.439 | 0.07 | 2.86E-81 |
| Endothelial Cell | Ptgs1 | 4 | 4.25E-84 | 2.84456837 | 0.622 | 0.263 | 8.51E-81 |
| Endothelial Cell | Fhad1 | 4 | 3.45E-83 | 1.25181445 | 0.544 | 0.194 | 6.90E-80 |
| Endothelial Cell | Exoc3l2 | 4 | 1.10E-77 | 2.85103922 | 0.599 | 0.282 | 2.20E-74 |
| Endothelial Cell | Adipoq | 4 | 7.16E-77 | 1.44302078 | 0.315 | 0.054 | 1.43E-73 |
| Endothelial Cell | Alox5 | 4 | 8.84E-77 | 0.98932332 | 0.414 | 0.118 | 1.77E-73 |
| Endothelial Cell | Icam1 | 4 | 2.13E-75 | 2.20938639 | 0.571 | 0.256 | 4.26E-72 |
| Endothelial Cell | Plekha7 | 4 | 2.90E-74 | 2.33907384 | 0.62 | 0.256 | 5.80E-71 |
| Endothelial Cell | Chn2 | 4 | 3.38E-74 | 1.55644885 | 0.712 | 0.396 | 6.76E-71 |
| Endothelial Cell | Nsg2 | 4 | 8.96E-73 | 1.13654346 | 0.311 | 0.058 | 1.79E-69 |
| Endothelial Cell | Rgs20 | 4 | 1.00E-71 | 2.09575728 | 0.437 | 0.078 | 2.00E-68 |
| Endothelial Cell | Ptn | 4 | 5.70E-71 | 2.16821517 | 0.66 | 0.4 | 1.14E-67 |
| Endothelial Cell | Samd51 | 4 | 3.63E-68 | 1.36960645 | 0.735 | 0.419 | 7.27E-65 |
| Endothelial Cell | Fbn1 | 4 | 1.65E-61 | 1.1853495 | 0.794 | 0.51 | 3.30E-58 |
| Endothelial Cell | Cttnbp2 | 4 | 2.79E-59 | 1.41781666 | 0.738 | 0.468 | 5.57E-56 |
| Endothelial Cell | Sh3gl3 | 4 | 3.57E-59 | 3.11068889 | 0.322 | 0.067 | 7.14E-56 |
| Endothelial Cell | Ap1s3 | 4 | 7.30E-58 | 1.87822122 | 0.55 | 0.28 | 1.46E-54 |
| Endothelial Cell | Ccn2 | 4 | 2.88E-56 | 1.94775821 | 0.629 | 0.306 | 5.77E-53 |
| Endothelial Cell | Klhl29 | 4 | 1.16E-54 | 3.87978834 | 0.551 | 0.3 | 2.31E-51 |
| Endothelial Cell | Tmcc3 | 4 | 3.29E-53 | 1.23272493 | 0.709 | 0.422 | 6.57E-50 |
| Endothelial Cell | Pde1c1 | 4 | 8.17E-53 | 1.14561695 | 0.75 | 0.465 | 1.63E-49 |
| Endothelial Cell | Apoe | 4 | 9.87E-52 | 1.81512365 | 0.583 | 0.266 | 1.97E-48 |

| | | | | | | | |
|---|---|---|---|---|---|---|---|
| Endothelial Cell | Lhfpl2 | 4 | 4.99E-51 | 1.84716243 | 0.597 | 0.333 | 9.98E-48 |
| Endothelial Cell | Slc44a5 | 4 | 4.83E-50 | 1.53472554 | 0.329 | 0.033 | 9.67E-47 |
| Endothelial Cell | Mid1 | 4 | 1.99E-49 | 1.54407602 | 0.623 | 0.329 | 3.98E-46 |
| Endothelial Cell | Ighm | 4 | 2.96E-47 | 0.57791229 | 0.409 | 0.135 | 5.91E-44 |
| Endothelial Cell | Cers6 | 4 | 2.34E-46 | 1.86631141 | 0.558 | 0.26 | 4.68E-43 |
| Endothelial Cell | Tpx21 | 4 | 7.38E-46 | 0.84489629 | 0.475 | 0.184 | 1.48E-42 |
| Endothelial Cell | Plxdc2 | 4 | 7.39E-46 | 1.27535865 | 0.715 | 0.428 | 1.48E-42 |
| Endothelial Cell | Ntf3 | 4 | 8.19E-46 | 0.92217997 | 0.721 | 0.446 | 1.64E-42 |
| Endothelial Cell | Pdgfc | 4 | 9.14E-45 | 3.20231546 | 0.459 | 0.197 | 1.83E-41 |
| Endothelial Cell | Myo3b | 4 | 3.85E-42 | 1.21255542 | 0.417 | 0.132 | 7.69E-39 |
| Endothelial Cell | Tspan5 | 4 | 8.21E-41 | 1.16026125 | 0.685 | 0.406 | 1.64E-37 |
| Endothelial Cell | Cp | 4 | 2.63E-39 | 3.13244781 | 0.485 | 0.193 | 5.27E-36 |
| Endothelial Cell | Grik2 | 4 | 1.82E-36 | 0.83858291 | 0.665 | 0.389 | 3.63E-33 |
| Endothelial Cell | Inpp4b | 4 | 5.52E-36 | 0.83548201 | 0.652 | 0.357 | 1.10E-32 |
| Endothelial Cell | Adam23 | 4 | 1.53E-34 | 0.62947933 | 0.443 | 0.172 | 3.06E-31 |
| Endothelial Cell | Slc47a11 | 4 | 5.35E-34 | 1.49481891 | 0.413 | 0.122 | 1.07E-30 |
| Endothelial Cell | AABR070544 | 4 | 1.28E-33 | 1.0776542 | 0.601 | 0.326 | 2.56E-30 |
| Endothelial Cell | Ifitm10 | 4 | 1.67E-33 | 0.32489117 | 0.54 | 0.281 | 3.35E-30 |
| Endothelial Cell | Cacnb4 | 4 | 2.37E-33 | 1.43915195 | 0.381 | 0.057 | 4.74E-30 |
| Endothelial Cell | Igfbp4 | 4 | 8.30E-32 | 0.83545121 | 0.567 | 0.278 | 1.66E-28 |
| Endothelial Cell | Ahr | 4 | 6.12E-29 | 0.78213428 | 0.652 | 0.393 | 1.22E-25 |
| Endothelial Cell | Myh11 | 4 | 7.23E-29 | 0.4391951 | 0.479 | 0.169 | 1.45E-25 |
| Endothelial Cell | Ppp1r14c | 4 | 8.49E-29 | 0.29505809 | 0.488 | 0.189 | 1.70E-25 |
| Endothelial Cell | Tnfaip8l3 | 4 | 1.47E-26 | 0.35634549 | 0.422 | 0.119 | 2.93E-23 |
| Endothelial Cell | Ctsz1 | 4 | 3.77E-26 | 0.66791547 | 0.506 | 0.256 | 7.53E-23 |
| Endothelial Cell | Gpcpd1 | 4 | 5.12E-26 | 1.06671812 | 0.565 | 0.306 | 1.02E-22 |
| Endothelial Cell | Heyl | 4 | 7.95E-26 | 0.30394674 | 0.443 | 0.176 | 1.59E-22 |
| Endothelial Cell | Aspa | 4 | 1.07E-25 | 0.26213167 | 0.471 | 0.179 | 2.15E-22 |
| Endothelial Cell | Itga7 | 4 | 1.19E-25 | 0.40451327 | 0.435 | 0.165 | 2.38E-22 |
| Endothelial Cell | Ptpn3 | 4 | 3.39E-24 | 0.27288994 | 0.537 | 0.243 | 6.78E-21 |
| Endothelial Cell | Pappa1 | 4 | 3.05E-23 | 0.51320397 | 0.403 | 0.149 | 6.10E-20 |

| | | | | | | | |
|---|---|---|---|---|---|---|---|
| Endothelial Cell | Rcan1 | 4 | 5.98E-22 | 0.36798099 | 0.489 | 0.228 | 1.20E-18 |
| Endothelial Cell | Tbx18 | 4 | 1.80E-21 | 0.28721952 | 0.43 | 0.176 | 3.59E-18 |
| Endothelial Cell | Fcgr3a | 5 | 0 | 4.39113277 | 0.363 | 0.006 | 0 |
| Endothelial Cell | Slc39a4 | 5 | 0 | 6.11154303 | 0.324 | 0.009 | 0 |
| Endothelial Cell | Chst9 | 5 | 1.45E-298 | 1.77522513 | 0.352 | 0.02 | 2.91E-295 |
| Endothelial Cell | Rab11fip1 | 5 | 1.78E-272 | 2.16733818 | 0.373 | 0.025 | 3.56E-269 |
| Endothelial Cell | Dmp1 | 5 | 8.00E-269 | 1.14128582 | 0.371 | 0.026 | 1.60E-265 |
| Endothelial Cell | Treml4 | 5 | 9.97E-259 | 1.26521934 | 0.292 | 0.016 | 1.99E-255 |
| Endothelial Cell | Nebl | 5 | 8.86E-212 | 2.43170598 | 0.948 | 0.493 | 1.77E-208 |
| Endothelial Cell | Dtx1 | 5 | 1.37E-201 | 1.83054579 | 0.389 | 0.018 | 2.74E-198 |
| Endothelial Cell | Cela3b | 5 | 1.36E-178 | 3.88149351 | 0.295 | 0.032 | 2.72E-175 |
| Endothelial Cell | Sulf11 | 5 | 1.01E-175 | 2.60889296 | 0.899 | 0.476 | 2.03E-172 |
| Endothelial Cell | AABR070635 | 5 | 2.95E-160 | 2.87914924 | 0.262 | 0.006 | 5.89E-157 |
| Endothelial Cell | Esco2 | 5 | 1.68E-149 | 2.05319926 | 0.454 | 0.064 | 3.35E-146 |
| Endothelial Cell | Gja5 | 5 | 1.34E-123 | 3.00417305 | 0.586 | 0.207 | 2.69E-120 |
| Endothelial Cell | St6galnac3 | 5 | 1.50E-121 | 1.78493017 | 0.899 | 0.605 | 3.00E-118 |
| Endothelial Cell | Pclo | 5 | 9.70E-114 | 2.56385215 | 0.436 | 0.026 | 1.94E-110 |
| Endothelial Cell | Tmem100 | 5 | 4.22E-110 | 4.50873366 | 0.666 | 0.199 | 8.44E-107 |
| Endothelial Cell | LOC1003610 | 5 | 1.04E-94 | 1.73905625 | 0.798 | 0.482 | 2.08E-91 |
| Endothelial Cell | Mettl24 | 5 | 7.69E-91 | 2.25610071 | 0.28 | 0.013 | 1.54E-87 |
| Endothelial Cell | Tp63 | 5 | 5.05E-90 | 2.61152469 | 0.402 | 0.102 | 1.01E-86 |
| Endothelial Cell | AC130940.1 | 5 | 1.27E-89 | 2.14699644 | 0.427 | 0.125 | 2.55E-86 |
| Endothelial Cell | Pcsk5 | 5 | 1.34E-83 | 2.85440998 | 0.678 | 0.324 | 2.67E-80 |
| Endothelial Cell | Pdgfd | 5 | 6.73E-82 | 1.91273912 | 0.738 | 0.482 | 1.35E-78 |
| Endothelial Cell | Sox17 | 5 | 1.38E-80 | 2.50644254 | 0.715 | 0.374 | 2.77E-77 |
| Endothelial Cell | AABR070294 | 5 | 1.48E-80 | 0.45937086 | 0.318 | 0.038 | 2.96E-77 |
| Endothelial Cell | Prkcq | 5 | 4.74E-79 | 0.75888892 | 0.44 | 0.077 | 9.49E-76 |
| Endothelial Cell | Smad6 | 5 | 5.32E-77 | 2.16541199 | 0.748 | 0.389 | 1.06E-73 |
| Endothelial Cell | Plcg2 | 5 | 3.16E-74 | 2.89661675 | 0.619 | 0.327 | 6.33E-71 |
| Endothelial Cell | Grm1 | 5 | 3.16E-73 | 1.31996167 | 0.275 | 0.013 | 6.31E-70 |
| Endothelial Cell | Nr4a3 | 5 | 3.66E-72 | 1.59128662 | 0.42 | 0.14 | 7.33E-69 |

| | | | | | | | |
|---|---|---|---|---|---|---|---|
| Endothelial Cell | Htr4 | 5 | 2.05E-70 | 1.33833711 | 0.365 | 0.06 | 4.09E-67 |
| Endothelial Cell | Eva1c | 5 | 2.92E-70 | 1.92145925 | 0.743 | 0.416 | 5.83E-67 |
| Endothelial Cell | Adamts20 | 5 | 6.23E-70 | 2.56420599 | 0.358 | 0.033 | 1.25E-66 |
| Endothelial Cell | Amd1 | 5 | 1.17E-68 | 2.35867982 | 0.656 | 0.365 | 2.33E-65 |
| Endothelial Cell | Pde11a | 5 | 6.35E-60 | 1.15032569 | 0.45 | 0.069 | 1.27E-56 |
| Endothelial Cell | Ifitm101 | 5 | 1.31E-54 | 2.6321914 | 0.565 | 0.281 | 2.61E-51 |
| Endothelial Cell | Chn21 | 5 | 1.20E-52 | 1.71598508 | 0.664 | 0.403 | 2.39E-49 |
| Endothelial Cell | AABR070504 | 5 | 5.33E-50 | 0.78092717 | 0.428 | 0.112 | 1.07E-46 |
| Endothelial Cell | AABR070311 | 5 | 8.10E-50 | 2.6622898 | 0.427 | 0.144 | 1.62E-46 |
| Endothelial Cell | Gabrb2 | 5 | 1.33E-48 | 1.49126663 | 0.436 | 0.152 | 2.66E-45 |
| Endothelial Cell | AABR070252 | 5 | 5.49E-48 | 1.66093044 | 0.704 | 0.386 | 1.10E-44 |
| Endothelial Cell | Efhd1 | 5 | 1.92E-47 | 3.12555694 | 0.544 | 0.287 | 3.85E-44 |
| Endothelial Cell | Ldb21 | 5 | 1.17E-45 | 1.51549455 | 0.671 | 0.382 | 2.34E-42 |
| Endothelial Cell | Sfrp4 | 5 | 2.62E-43 | 1.04363782 | 0.438 | 0.111 | 5.25E-40 |
| Endothelial Cell | Igsf11 | 5 | 2.38E-40 | 0.68269835 | 0.469 | 0.122 | 4.77E-37 |
| Endothelial Cell | Fcgr2b | 5 | 5.15E-40 | 1.3671541 | 0.445 | 0.187 | 1.03E-36 |
| Endothelial Cell | Pla2g7 | 5 | 1.77E-38 | 0.51542371 | 0.35 | 0.049 | 3.53E-35 |
| Endothelial Cell | Cxcl12 | 5 | 4.02E-37 | 1.1007962 | 0.713 | 0.444 | 8.04E-34 |
| Endothelial Cell | Greb1l | 5 | 1.01E-35 | 2.02148593 | 0.5 | 0.194 | 2.01E-32 |
| Endothelial Cell | Pappa2 | 5 | 4.51E-30 | 1.42568885 | 0.396 | 0.129 | 9.01E-27 |
| Endothelial Cell | Marchf1 | 5 | 5.94E-30 | 1.07729866 | 0.436 | 0.157 | 1.19E-26 |
| Endothelial Cell | Rnf144b | 5 | 6.20E-28 | 1.25323047 | 0.656 | 0.399 | 1.24E-24 |
| Endothelial Cell | Musk | 5 | 1.04E-27 | 1.17504639 | 0.415 | 0.15 | 2.08E-24 |
| Endothelial Cell | Slc30a3 | 5 | 4.79E-27 | 0.42282631 | 0.459 | 0.166 | 9.59E-24 |
| Endothelial Cell | Fstl5 | 5 | 2.33E-26 | 0.31381317 | 0.394 | 0.126 | 4.65E-23 |
| Endothelial Cell | Col8a2 | 5 | 2.68E-25 | 1.37527509 | 0.389 | 0.13 | 5.37E-22 |
| Endothelial Cell | Fbln2 | 5 | 5.03E-24 | 1.75980758 | 0.542 | 0.265 | 1.01E-20 |
| Endothelial Cell | Npr3 | 5 | 9.22E-23 | 1.94039948 | 0.469 | 0.206 | 1.84E-19 |
| Endothelial Cell | Anpep | 5 | 2.25E-18 | 0.6386407 | 0.397 | 0.146 | 4.51E-15 |
| Endothelial Cell | Hs6st2 | 5 | 8.49E-16 | 0.5822836 | 0.432 | 0.178 | 1.70E-12 |
| Endothelial Cell | Pip5k1b | 5 | 7.16E-13 | 1.44505777 | 0.441 | 0.17 | 1.43E-09 |

| | | | | | | | |
|---|---|---|---|---|---|---|---|
| Endothelial Cell | Kcnc2 | 5 | 4.21E-12 | 0.4769893 | 0.466 | 0.191 | 8.42E-09 |
| Endothelial Cell | Pnoc | 6 | 3.22E-168 | 2.31201749 | 0.278 | 0.014 | 6.45E-165 |
| Endothelial Cell | Rspo2 | 6 | 3.37E-132 | 2.79249745 | 0.347 | 0.038 | 6.74E-129 |
| Endothelial Cell | Kcnk5 | 6 | 8.15E-46 | 1.14949467 | 0.393 | 0.115 | 1.63E-42 |
| Endothelial Cell | Ccl24 | 6 | 2.27E-43 | 1.82606323 | 0.309 | 0.055 | 4.54E-40 |
| Endothelial Cell | Hmox1 | 6 | 2.39E-36 | 0.28643107 | 0.446 | 0.156 | 4.78E-33 |
| Endothelial Cell | RT1-DOa | 7 | 3.14E-235 | 0.4927365 | 0.327 | 0.021 | 6.29E-232 |
| Endothelial Cell | Myom3 | 7 | 6.96E-100 | 2.25625456 | 0.301 | 0.044 | 1.39E-96 |
| Endothelial Cell | Cxcl121 | 7 | 2.41E-75 | 2.05141095 | 0.784 | 0.444 | 4.81E-72 |
| Endothelial Cell | Unc5b | 7 | 2.92E-72 | 1.97333957 | 0.784 | 0.464 | 5.84E-69 |
| Endothelial Cell | Fstl4 | 7 | 1.41E-57 | 2.08448224 | 0.313 | 0.051 | 2.81E-54 |
| Endothelial Cell | Btnl91 | 7 | 2.41E-56 | 2.0315778 | 0.707 | 0.37 | 4.82E-53 |
| Endothelial Cell | Smad61 | 7 | 3.87E-56 | 2.28679045 | 0.687 | 0.399 | 7.74E-53 |
| Endothelial Cell | Nebl1 | 7 | 1.43E-55 | 1.44055377 | 0.796 | 0.51 | 2.87E-52 |
| Endothelial Cell | Hamp | 7 | 1.48E-53 | 0.43545334 | 0.376 | 0.107 | 2.97E-50 |
| Endothelial Cell | Edar | 7 | 2.21E-51 | 2.30244215 | 0.386 | 0.089 | 4.43E-48 |
| Endothelial Cell | Fgf10 | 7 | 7.14E-17 | 1.54416851 | 0.368 | 0.083 | 1.43E-13 |
| Endothelial Cell | Prdm6 | 7 | 8.33E-17 | 0.31309732 | 0.077 | 0.332 | 1.67E-13 |
| Endothelial Cell | Anpep1 | 7 | 1.28E-08 | 0.72467838 | 0.422 | 0.148 | 2.57E-05 |
| Endothelial Cell | Ikzf3 | 8 | 0 | 0.5948398 | 0.888 | 0.095 | 0 |
| Endothelial Cell | Bcl11b | 8 | 0 | 1.62370811 | 0.746 | 0.075 | 0 |
| Endothelial Cell | Krt17 | 8 | 1.49E-294 | 3.0789858 | 0.639 | 0.071 | 2.99E-291 |
| Endothelial Cell | Dgat2 | 8 | 4.28E-278 | 0.70214318 | 0.821 | 0.145 | 8.56E-275 |
| Endothelial Cell | Hmmr | 8 | 9.15E-273 | 1.52380461 | 0.873 | 0.171 | 1.83E-269 |
| Endothelial Cell | Tenm2 | 8 | 2.26E-252 | 2.75816128 | 0.677 | 0.048 | 4.52E-249 |
| Endothelial Cell | Tlr7 | 8 | 2.06E-247 | 2.02883261 | 0.789 | 0.115 | 4.11E-244 |
| Endothelial Cell | Tspan11 | 8 | 2.13E-246 | 0.50874381 | 0.907 | 0.187 | 4.26E-243 |
| Endothelial Cell | Tenm4 | 8 | 3.03E-246 | 0.44766043 | 0.462 | 0.054 | 6.07E-243 |
| Endothelial Cell | Tbx21 | 8 | 1.60E-231 | 0.49151878 | 0.689 | 0.089 | 3.20E-228 |
| Endothelial Cell | Col25a1 | 8 | 1.55E-219 | 1.90736524 | 0.61 | 0.069 | 3.09E-216 |
| Endothelial Cell | Bank1 | 8 | 7.85E-199 | 1.43995897 | 0.737 | 0.151 | 1.57E-195 |

| | | | | | | | |
|---|---|---|---|---|---|---|---|
| Endothelial Cell | Il1rl1 | 8 | 2.43E-198 | 0.87767386 | 0.73 | 0.134 | 4.85E-195 |
| Endothelial Cell | Atp10b | 8 | 1.09E-195 | 1.75842792 | 0.548 | 0.101 | 2.17E-192 |
| Endothelial Cell | Mark1 | 8 | 3.28E-194 | 1.287227 | 0.928 | 0.196 | 6.55E-191 |
| Endothelial Cell | Hs3st3a1 | 8 | 2.82E-193 | 0.2869596 | 0.4 | 0.053 | 5.64E-190 |
| Endothelial Cell | Stap1 | 8 | 2.93E-193 | 1.03160286 | 0.737 | 0.111 | 5.86E-190 |
| Endothelial Cell | AABR070215 | 8 | 2.55E-185 | 1.37858732 | 0.78 | 0.116 | 5.09E-182 |
| Endothelial Cell | Wwc1 | 8 | 1.39E-183 | 0.82945073 | 0.773 | 0.189 | 2.77E-180 |
| Endothelial Cell | Mapk10 | 8 | 1.97E-178 | 0.88556246 | 0.313 | 0.048 | 3.95E-175 |
| Endothelial Cell | Fhad11 | 8 | 3.31E-177 | 0.36193133 | 0.801 | 0.192 | 6.62E-174 |
| Endothelial Cell | AABR070605 | 8 | 1.28E-176 | 1.72794275 | 0.931 | 0.335 | 2.55E-173 |
| Endothelial Cell | Gpr176 | 8 | 2.14E-171 | 1.34248054 | 0.821 | 0.174 | 4.29E-168 |
| Endothelial Cell | Otulinl | 8 | 7.39E-164 | 1.35264951 | 0.856 | 0.272 | 1.48E-160 |
| Endothelial Cell | Chl1 | 8 | 2.15E-162 | 1.9711789 | 0.775 | 0.189 | 4.30E-159 |
| Endothelial Cell | RGD1305184 | 8 | 1.45E-160 | 0.39612689 | 0.581 | 0.05 | 2.89E-157 |
| Endothelial Cell | Slc37a2 | 8 | 7.44E-159 | 1.12110641 | 0.787 | 0.222 | 1.49E-155 |
| Endothelial Cell | Adgrd1 | 8 | 1.18E-158 | 1.40308688 | 0.849 | 0.218 | 2.35E-155 |
| Endothelial Cell | Baiap2l1 | 8 | 1.89E-157 | 1.39863208 | 0.806 | 0.209 | 3.78E-154 |
| Endothelial Cell | AABR070134 | 8 | 2.40E-157 | 2.14050169 | 0.306 | 0.031 | 4.80E-154 |
| Endothelial Cell | Tpd52l1 | 8 | 5.80E-157 | 1.07730168 | 0.83 | 0.218 | 1.16E-153 |
| Endothelial Cell | Asic2 | 8 | 4.74E-154 | 1.17840909 | 0.706 | 0.175 | 9.48E-151 |
| Endothelial Cell | Cdkn1a | 8 | 1.22E-153 | 1.18828058 | 0.878 | 0.32 | 2.45E-150 |
| Endothelial Cell | Kntc1 | 8 | 3.24E-151 | 1.45071948 | 0.708 | 0.182 | 6.47E-148 |
| Endothelial Cell | Gabrb1 | 8 | 3.52E-151 | 1.47551536 | 0.694 | 0.212 | 7.03E-148 |
| Endothelial Cell | AC118957.1 | 8 | 5.44E-151 | 1.95914983 | 0.455 | 0.054 | 1.09E-147 |
| Endothelial Cell | Atf3 | 8 | 9.83E-151 | 1.00834737 | 0.804 | 0.245 | 1.97E-147 |
| Endothelial Cell | Abcd2 | 8 | 2.21E-148 | 2.33313449 | 0.722 | 0.184 | 4.42E-145 |
| Endothelial Cell | Ccl241 | 8 | 2.30E-146 | 0.34278147 | 0.321 | 0.061 | 4.59E-143 |
| Endothelial Cell | Aldh1a2 | 8 | 3.93E-146 | 0.85013461 | 0.897 | 0.277 | 7.86E-143 |
| Endothelial Cell | Col11a1 | 8 | 1.68E-144 | 0.55091231 | 0.88 | 0.287 | 3.37E-141 |
| Endothelial Cell | Slc10a6 | 8 | 8.17E-141 | 0.5395856 | 0.734 | 0.221 | 1.63E-137 |
| Endothelial Cell | AC121220.1 | 8 | 1.81E-139 | 1.43844141 | 0.502 | 0.087 | 3.62E-136 |

| | | | | | | | |
|---|---|---|---|---|---|---|---|
| Endothelial Cell | Srl | 8 | 4.98E-138 | 0.29766564 | 0.713 | 0.186 | 9.96E-135 |
| Endothelial Cell | Gen1 | 8 | 2.64E-136 | 0.83897632 | 0.854 | 0.244 | 5.28E-133 |
| Endothelial Cell | Plxnc1 | 8 | 4.45E-135 | 0.65204981 | 0.419 | 0.162 | 8.91E-132 |
| Endothelial Cell | AABR070504 | 8 | 2.62E-133 | 0.36410774 | 0.636 | 0.109 | 5.25E-130 |
| Endothelial Cell | Tex22 | 8 | 1.15E-132 | 1.5611974 | 0.423 | 0.171 | 2.30E-129 |
| Endothelial Cell | Loxl2 | 8 | 2.62E-131 | 0.43868939 | 0.873 | 0.312 | 5.23E-128 |
| Endothelial Cell | Tmem17 | 8 | 3.71E-131 | 1.10767642 | 0.789 | 0.235 | 7.41E-128 |
| Endothelial Cell | Mmrn1 | 8 | 1.02E-129 | 1.37278129 | 0.739 | 0.199 | 2.04E-126 |
| Endothelial Cell | LOC1009095 | 8 | 2.34E-127 | 0.89033635 | 0.89 | 0.307 | 4.69E-124 |
| Endothelial Cell | Ndc80 | 8 | 5.78E-127 | 0.58972905 | 0.775 | 0.276 | 1.16E-123 |
| Endothelial Cell | Rassf4 | 8 | 6.73E-127 | 0.85107994 | 0.782 | 0.228 | 1.35E-123 |
| Endothelial Cell | Pfkfb1 | 8 | 6.94E-126 | 1.38542067 | 0.577 | 0.205 | 1.39E-122 |
| Endothelial Cell | Mamdc2 | 8 | 8.27E-126 | 1.25049673 | 0.797 | 0.331 | 1.65E-122 |
| Endothelial Cell | Tnn | 8 | 1.34E-124 | 0.90347139 | 0.529 | 0.197 | 2.68E-121 |
| Endothelial Cell | Gria4 | 8 | 2.80E-123 | 1.37075412 | 0.629 | 0.177 | 5.61E-120 |
| Endothelial Cell | Mtus2 | 8 | 3.89E-123 | 0.50140409 | 0.744 | 0.24 | 7.79E-120 |
| Endothelial Cell | Fgfr2 | 8 | 1.00E-122 | 0.61492316 | 0.828 | 0.241 | 2.01E-119 |
| Endothelial Cell | LOC1009119 | 8 | 4.47E-122 | 0.88351653 | 0.711 | 0.235 | 8.94E-119 |
| Endothelial Cell | Pld5 | 8 | 1.97E-121 | 1.1791475 | 0.77 | 0.25 | 3.94E-118 |
| Endothelial Cell | Knl11 | 8 | 1.29E-120 | 1.20387801 | 0.696 | 0.24 | 2.58E-117 |
| Endothelial Cell | AABR070606 | 8 | 8.05E-120 | 0.97525841 | 0.935 | 0.422 | 1.61E-116 |
| Endothelial Cell | Nmnat2 | 8 | 6.68E-119 | 1.01432406 | 0.679 | 0.228 | 1.34E-115 |
| Endothelial Cell | Kcnk2 | 8 | 1.48E-118 | 1.42048663 | 0.577 | 0.115 | 2.97E-115 |
| Endothelial Cell | Ryr3 | 8 | 1.14E-117 | 0.87954432 | 0.77 | 0.288 | 2.27E-114 |
| Endothelial Cell | Ezr | 8 | 3.75E-117 | 0.67461445 | 0.799 | 0.28 | 7.50E-114 |
| Endothelial Cell | Kcnj8 | 8 | 4.60E-117 | 0.94170597 | 0.703 | 0.166 | 9.21E-114 |
| Endothelial Cell | Nrxn3 | 8 | 6.32E-117 | 1.02790951 | 0.665 | 0.185 | 1.26E-113 |
| Endothelial Cell | Grb14 | 8 | 1.21E-116 | 0.77304484 | 0.856 | 0.298 | 2.42E-113 |
| Endothelial Cell | Gabra4 | 8 | 2.30E-116 | 1.20893072 | 0.864 | 0.347 | 4.59E-113 |
| Endothelial Cell | Alpk2 | 8 | 1.73E-115 | 0.47943799 | 0.792 | 0.244 | 3.45E-112 |
| Endothelial Cell | Tmem26 | 8 | 2.15E-115 | 1.31149202 | 0.842 | 0.331 | 4.30E-112 |

| | | | | | | | |
|---|---|---|---|---|---|---|---|
| Endothelial Cell | Map3k7cl | 8 | 3.66E-115 | 0.70946583 | 0.778 | 0.305 | 7.32E-112 |
| Endothelial Cell | Adgrg6 | 8 | 2.18E-113 | 1.08754637 | 0.651 | 0.184 | 4.36E-110 |
| Endothelial Cell | Prdm61 | 8 | 2.33E-113 | 1.15874431 | 0.823 | 0.289 | 4.66E-110 |
| Endothelial Cell | Palm2 | 8 | 2.63E-113 | 1.12385434 | 0.792 | 0.275 | 5.25E-110 |
| Endothelial Cell | Lurap1l | 8 | 2.64E-113 | 0.7181187 | 0.907 | 0.327 | 5.28E-110 |
| Endothelial Cell | Cyba | 8 | 6.96E-113 | 0.90148573 | 0.821 | 0.296 | 1.39E-109 |
| Endothelial Cell | Prph | 8 | 2.09E-112 | 1.55477933 | 0.514 | 0.121 | 4.18E-109 |
| Endothelial Cell | Dclk1 | 8 | 2.26E-111 | 0.65015837 | 0.708 | 0.215 | 4.52E-108 |
| Endothelial Cell | Cobl | 8 | 6.27E-111 | 0.89818048 | 0.699 | 0.207 | 1.25E-107 |
| Endothelial Cell | Mbp | 8 | 9.51E-111 | 0.74442773 | 0.577 | 0.198 | 1.90E-107 |
| Endothelial Cell | Smoc1 | 8 | 4.07E-110 | 1.53910173 | 0.608 | 0.148 | 8.14E-107 |
| Endothelial Cell | AABR070515 | 8 | 5.29E-110 | 0.30835984 | 0.646 | 0.177 | 1.06E-106 |
| Endothelial Cell | Cd55 | 8 | 8.68E-110 | 0.66623214 | 0.844 | 0.308 | 1.74E-106 |
| Endothelial Cell | Sfrp2 | 8 | 4.25E-109 | 1.42069841 | 0.797 | 0.268 | 8.49E-106 |
| Endothelial Cell | Mmp12 | 8 | 2.35E-108 | 1.24009994 | 0.536 | 0.136 | 4.70E-105 |
| Endothelial Cell | Mnda | 8 | 1.50E-107 | 1.00568603 | 0.778 | 0.295 | 3.01E-104 |
| Endothelial Cell | Camk4 | 8 | 1.71E-107 | 0.8663708 | 0.533 | 0.189 | 3.43E-104 |
| Endothelial Cell | Ror1 | 8 | 2.70E-107 | 0.47818858 | 0.653 | 0.235 | 5.41E-104 |
| Endothelial Cell | Trim50 | 8 | 7.34E-107 | 0.64448432 | 0.742 | 0.231 | 1.47E-103 |
| Endothelial Cell | Acta2 | 8 | 2.04E-105 | 0.8634399 | 0.691 | 0.206 | 4.08E-102 |
| Endothelial Cell | Slc1a11 | 8 | 2.65E-105 | 0.32867665 | 0.931 | 0.412 | 5.30E-102 |
| Endothelial Cell | Aldh1a1 | 8 | 3.71E-105 | 1.2732029 | 0.51 | 0.203 | 7.42E-102 |
| Endothelial Cell | Fras1 | 8 | 1.25E-104 | 1.20293214 | 0.725 | 0.281 | 2.50E-101 |
| Endothelial Cell | Timp1 | 8 | 6.88E-104 | 1.09980305 | 0.749 | 0.245 | 1.38E-100 |
| Endothelial Cell | Kcnma1 | 8 | 8.11E-104 | 1.59225467 | 0.624 | 0.257 | 1.62E-100 |
| Endothelial Cell | Frmd3 | 8 | 1.04E-103 | 0.31910344 | 0.768 | 0.287 | 2.08E-100 |
| Endothelial Cell | Snta1 | 8 | 6.31E-103 | 0.95479259 | 0.722 | 0.252 | 1.26E-99 |
| Endothelial Cell | Trpc3 | 8 | 6.75E-103 | 0.61202928 | 0.663 | 0.182 | 1.35E-99 |
| Endothelial Cell | Sema3c | 8 | 9.93E-102 | 0.81696839 | 0.598 | 0.191 | 1.99E-98 |
| Endothelial Cell | Sgcg | 8 | 3.37E-101 | 0.56417912 | 0.682 | 0.267 | 6.74E-98 |
| Endothelial Cell | Atp8b4 | 8 | 1.19E-99 | 0.44085861 | 0.605 | 0.227 | 2.38E-96 |

| | | | | | | | |
|---|---|---|---|---|---|---|---|
| Endothelial Cell | Miga2 | 8 | 5.20E-99 | 0.78321185 | 0.775 | 0.307 | 1.04E-95 |
| Endothelial Cell | Plcxd3 | 8 | 6.59E-99 | 0.53709632 | 0.687 | 0.24 | 1.32E-95 |
| Endothelial Cell | Cybb | 8 | 2.41E-98 | 1.04429534 | 0.502 | 0.163 | 4.82E-95 |
| Endothelial Cell | Gpm6b | 8 | 8.91E-98 | 1.06540185 | 0.61 | 0.204 | 1.78E-94 |
| Endothelial Cell | Fgf13 | 8 | 3.67E-97 | 1.27688758 | 0.811 | 0.379 | 7.33E-94 |
| Endothelial Cell | Ect2 | 8 | 4.77E-97 | 0.59312108 | 0.488 | 0.167 | 9.53E-94 |
| Endothelial Cell | Spon1 | 8 | 5.13E-97 | 0.9280204 | 0.684 | 0.24 | 1.03E-93 |
| Endothelial Cell | Tagln | 8 | 1.32E-96 | 1.01088247 | 0.639 | 0.255 | 2.63E-93 |
| Endothelial Cell | Klhl4 | 8 | 3.84E-96 | 0.34464613 | 0.895 | 0.359 | 7.68E-93 |
| Endothelial Cell | Ptgfr | 8 | 5.77E-96 | 0.31897247 | 0.438 | 0.175 | 1.15E-92 |
| Endothelial Cell | Cpq | 8 | 1.31E-95 | 0.84349765 | 0.727 | 0.247 | 2.62E-92 |
| Endothelial Cell | Scara5 | 8 | 1.36E-95 | 1.73283304 | 0.787 | 0.264 | 2.73E-92 |
| Endothelial Cell | Cpt1b | 8 | 1.78E-95 | 0.48410403 | 0.55 | 0.178 | 3.56E-92 |
| Endothelial Cell | Ednra | 8 | 1.58E-94 | 0.87391951 | 0.873 | 0.402 | 3.16E-91 |
| Endothelial Cell | Pltp | 8 | 3.37E-94 | 0.7084505 | 0.823 | 0.328 | 6.74E-91 |
| Endothelial Cell | Ctsz2 | 8 | 9.17E-94 | 0.92380131 | 0.653 | 0.257 | 1.83E-90 |
| Endothelial Cell | Myh10 | 8 | 7.89E-93 | 0.60424555 | 0.775 | 0.28 | 1.58E-89 |
| Endothelial Cell | Blnk | 8 | 9.92E-93 | 0.51635749 | 0.749 | 0.37 | 1.98E-89 |
| Endothelial Cell | S100a4 | 8 | 1.14E-92 | 0.65504538 | 0.632 | 0.209 | 2.29E-89 |
| Endothelial Cell | Tmem132d | 8 | 1.34E-92 | 1.03433178 | 0.624 | 0.228 | 2.69E-89 |
| Endothelial Cell | Slc7a2 | 8 | 2.31E-92 | 1.01892606 | 0.667 | 0.241 | 4.61E-89 |
| Endothelial Cell | Ugdh | 8 | 9.98E-92 | 0.59927297 | 0.624 | 0.241 | 2.00E-88 |
| Endothelial Cell | Svep1 | 8 | 1.79E-91 | 0.25567685 | 0.667 | 0.174 | 3.58E-88 |
| Endothelial Cell | Slc1a7 | 8 | 1.82E-91 | 1.72785963 | 0.639 | 0.236 | 3.64E-88 |
| Endothelial Cell | Khdrbs2 | 8 | 2.03E-91 | 1.62699747 | 0.493 | 0.116 | 4.07E-88 |
| Endothelial Cell | Mdga2 | 8 | 2.23E-90 | 1.04860971 | 0.421 | 0.11 | 4.46E-87 |
| Endothelial Cell | Kcnj5 | 8 | 2.92E-90 | 0.6411233 | 0.66 | 0.237 | 5.83E-87 |
| Endothelial Cell | Txlnb | 8 | 3.04E-90 | 0.37071427 | 0.792 | 0.338 | 6.09E-87 |
| Endothelial Cell | Fgf1 | 8 | 5.87E-89 | 0.56170916 | 0.768 | 0.305 | 1.17E-85 |
| Endothelial Cell | Gfra2 | 8 | 8.23E-89 | 1.32977009 | 0.653 | 0.244 | 1.65E-85 |
| Endothelial Cell | Egfl8 | 8 | 2.15E-88 | 0.86871267 | 0.768 | 0.263 | 4.29E-85 |

| | | | | | | | |
|---|---|---|---|---|---|---|---|
| Endothelial Cell | Frmd5 | 8 | 3.07E-88 | 0.64977211 | 0.833 | 0.362 | 6.14E-85 |
| Endothelial Cell | Dgki | 8 | 3.61E-88 | 0.8274035 | 0.474 | 0.161 | 7.22E-85 |
| Endothelial Cell | Tpm2 | 8 | 4.01E-88 | 0.69012033 | 0.754 | 0.299 | 8.02E-85 |
| Endothelial Cell | Ros1 | 8 | 5.38E-88 | 1.17587539 | 0.62 | 0.238 | 1.08E-84 |
| Endothelial Cell | Asb18 | 8 | 5.55E-88 | 0.61751094 | 0.749 | 0.332 | 1.11E-84 |
| Endothelial Cell | Il17rd | 8 | 1.71E-87 | 0.26848247 | 0.5 | 0.181 | 3.42E-84 |
| Endothelial Cell | Abcc8 | 8 | 1.19E-86 | 1.31731022 | 0.4 | 0.144 | 2.38E-83 |
| Endothelial Cell | Slit2 | 8 | 1.37E-86 | 1.00539044 | 0.653 | 0.249 | 2.75E-83 |
| Endothelial Cell | Mreg | 8 | 2.14E-86 | 1.1132419 | 0.4 | 0.063 | 4.28E-83 |
| Endothelial Cell | Pgam2 | 8 | 2.56E-86 | 0.74011153 | 0.646 | 0.242 | 5.12E-83 |
| Endothelial Cell | Col14a1 | 8 | 3.89E-86 | 0.51919 | 0.632 | 0.235 | 7.79E-83 |
| Endothelial Cell | Corin | 8 | 1.27E-85 | 0.57357319 | 0.687 | 0.285 | 2.53E-82 |
| Endothelial Cell | Kif18b | 8 | 3.40E-85 | 0.2565766 | 0.543 | 0.152 | 6.79E-82 |
| Endothelial Cell | Ptprd | 8 | 3.73E-85 | 0.97863641 | 0.679 | 0.259 | 7.46E-82 |
| Endothelial Cell | Col16a1 | 8 | 6.23E-85 | 0.33512778 | 0.596 | 0.208 | 1.25E-81 |
| Endothelial Cell | Ptpn13 | 8 | 3.67E-84 | 1.1028113 | 0.622 | 0.254 | 7.33E-81 |
| Endothelial Cell | LOC1036942 | 8 | 4.63E-84 | 0.41193133 | 0.911 | 0.439 | 9.27E-81 |
| Endothelial Cell | Slc4a3 | 8 | 1.80E-83 | 0.55656093 | 0.644 | 0.276 | 3.60E-80 |
| Endothelial Cell | Enpp1 | 8 | 4.33E-83 | 0.2607837 | 0.835 | 0.342 | 8.65E-80 |
| Endothelial Cell | Uqcr10 | 8 | 2.13E-82 | 0.77039554 | 0.828 | 0.339 | 4.25E-79 |
| Endothelial Cell | Rrad | 8 | 4.97E-82 | 0.39257774 | 0.78 | 0.328 | 9.93E-79 |
| Endothelial Cell | Hvcn1 | 8 | 1.04E-81 | 0.41675723 | 0.737 | 0.263 | 2.08E-78 |
| Endothelial Cell | Alk | 8 | 2.57E-81 | 0.25382739 | 0.596 | 0.25 | 5.14E-78 |
| Endothelial Cell | Pde1a | 8 | 7.29E-81 | 0.64061236 | 0.567 | 0.243 | 1.46E-77 |
| Endothelial Cell | Efna5 | 8 | 1.00E-80 | 0.72693127 | 0.675 | 0.191 | 2.00E-77 |
| Endothelial Cell | Sntb1 | 8 | 7.81E-80 | 0.87467178 | 0.885 | 0.412 | 1.56E-76 |
| Endothelial Cell | Dpp4 | 8 | 7.85E-80 | 1.17041086 | 0.801 | 0.42 | 1.57E-76 |
| Endothelial Cell | Slc38a1 | 8 | 1.78E-79 | 0.28956612 | 0.644 | 0.254 | 3.55E-76 |
| Endothelial Cell | Flvcr2 | 8 | 1.94E-79 | 0.30840046 | 0.768 | 0.309 | 3.88E-76 |
| Endothelial Cell | Nkd2 | 8 | 2.79E-79 | 0.63250008 | 0.739 | 0.369 | 5.59E-76 |
| Endothelial Cell | Acot11 | 8 | 1.56E-78 | 0.86869383 | 0.591 | 0.251 | 3.13E-75 |

| | | | | | | | |
|---|---|---|---|---|---|---|---|
| Endothelial Cell | Bend6 | 8 | 1.59E-78 | 0.288396 | 0.737 | 0.284 | 3.17E-75 |
| Endothelial Cell | Trim63 | 8 | 1.70E-78 | 0.61654745 | 0.644 | 0.265 | 3.40E-75 |
| Endothelial Cell | Meox1 | 8 | 2.71E-78 | 0.33556392 | 0.634 | 0.198 | 5.42E-75 |
| Endothelial Cell | Cdh11 | 8 | 3.22E-78 | 0.33919855 | 0.632 | 0.252 | 6.45E-75 |
| Endothelial Cell | Flnc | 8 | 8.17E-78 | 0.27922305 | 0.555 | 0.21 | 1.63E-74 |
| Endothelial Cell | Aff2 | 8 | 3.62E-77 | 0.73602989 | 0.548 | 0.257 | 7.23E-74 |
| Endothelial Cell | Pi16 | 8 | 5.31E-77 | 0.47211004 | 0.679 | 0.301 | 1.06E-73 |
| Endothelial Cell | Ccn1 | 8 | 6.68E-77 | 0.50162758 | 0.699 | 0.284 | 1.34E-73 |
| Endothelial Cell | Dpt | 8 | 4.38E-75 | 0.61372339 | 0.648 | 0.25 | 8.76E-72 |
| Endothelial Cell | Unc5c1 | 8 | 1.01E-74 | 1.07185495 | 0.844 | 0.362 | 2.02E-71 |
| Endothelial Cell | Tmem82 | 8 | 1.43E-74 | 1.09950394 | 0.639 | 0.292 | 2.85E-71 |
| Endothelial Cell | Slc45a3 | 8 | 1.64E-74 | 0.33295331 | 0.586 | 0.126 | 3.28E-71 |
| Endothelial Cell | Trim55 | 8 | 2.72E-74 | 0.36383144 | 0.711 | 0.304 | 5.43E-71 |
| Endothelial Cell | Podnl1 | 8 | 6.06E-74 | 1.59967666 | 0.328 | 0.045 | 1.21E-70 |
| Endothelial Cell | Ncam1 | 8 | 1.23E-73 | 0.25840334 | 0.536 | 0.246 | 2.46E-70 |
| Endothelial Cell | Mrvi1 | 8 | 1.51E-73 | 0.63342159 | 0.481 | 0.217 | 3.02E-70 |
| Endothelial Cell | Prg4 | 8 | 1.99E-73 | 0.97142036 | 0.536 | 0.239 | 3.98E-70 |
| Endothelial Cell | Fsd2 | 8 | 2.70E-73 | 0.62881518 | 0.658 | 0.304 | 5.41E-70 |
| Endothelial Cell | Efcc1 | 8 | 3.03E-73 | 0.37905422 | 0.653 | 0.292 | 6.05E-70 |
| Endothelial Cell | Jph2 | 8 | 1.39E-72 | 0.61995912 | 0.636 | 0.278 | 2.79E-69 |
| Endothelial Cell | Dmpk | 8 | 3.14E-72 | 0.52847906 | 0.873 | 0.378 | 6.28E-69 |
| Endothelial Cell | Ano5 | 8 | 9.47E-72 | 0.27241572 | 0.665 | 0.231 | 1.89E-68 |
| Endothelial Cell | Srpk3 | 8 | 3.53E-71 | 0.57233857 | 0.56 | 0.268 | 7.06E-68 |
| Endothelial Cell | Smpx | 8 | 6.28E-71 | 1.03637557 | 0.567 | 0.267 | 1.26E-67 |
| Endothelial Cell | Serpine2 | 8 | 1.77E-69 | 0.57354689 | 0.639 | 0.329 | 3.54E-66 |
| Endothelial Cell | Kif20b | 8 | 2.93E-69 | 0.82087293 | 0.56 | 0.206 | 5.86E-66 |
| Endothelial Cell | Csrp3 | 8 | 4.77E-69 | 0.42725157 | 0.67 | 0.289 | 9.54E-66 |
| Endothelial Cell | Col6a6 | 8 | 8.45E-69 | 0.90281735 | 0.653 | 0.296 | 1.69E-65 |
| Endothelial Cell | Mlip | 8 | 9.02E-69 | 0.54360157 | 0.627 | 0.27 | 1.80E-65 |
| Endothelial Cell | Ptpn31 | 8 | 1.57E-68 | 0.41467642 | 0.577 | 0.251 | 3.14E-65 |
| Endothelial Cell | Pkia | 8 | 1.85E-68 | 0.84350408 | 0.761 | 0.357 | 3.70E-65 |

| | | | | | | | |
|---|---|---|---|---|---|---|---|
| Endothelial Cell | Epb41l4b | 8 | 2.34E-68 | 1.30306487 | 0.577 | 0.273 | 4.68E-65 |
| Endothelial Cell | Serping1 | 8 | 2.74E-68 | 1.09830782 | 0.505 | 0.246 | 5.48E-65 |
| Endothelial Cell | Robo2 | 8 | 4.85E-68 | 0.64862581 | 0.531 | 0.226 | 9.70E-65 |
| Endothelial Cell | Tafa2 | 8 | 3.31E-67 | 0.52463422 | 0.715 | 0.328 | 6.61E-64 |
| Endothelial Cell | Gata41 | 8 | 5.14E-67 | 0.41194393 | 0.574 | 0.257 | 1.03E-63 |
| Endothelial Cell | Car10 | 8 | 5.15E-67 | 2.05735913 | 0.462 | 0.12 | 1.03E-63 |
| Endothelial Cell | Nova1 | 8 | 9.04E-67 | 0.76282285 | 0.524 | 0.252 | 1.81E-63 |
| Endothelial Cell | Pde7a | 8 | 2.44E-66 | 0.47500402 | 0.888 | 0.461 | 4.87E-63 |
| Endothelial Cell | Ivns1abp | 8 | 4.26E-66 | 0.40839431 | 0.835 | 0.389 | 8.52E-63 |
| Endothelial Cell | Dtna | 8 | 5.82E-66 | 0.3648502 | 0.624 | 0.297 | 1.16E-62 |
| Endothelial Cell | Runx1 | 8 | 9.26E-66 | 0.62036499 | 0.749 | 0.338 | 1.85E-62 |
| Endothelial Cell | Dapk1 | 8 | 1.11E-65 | 0.48358204 | 0.833 | 0.383 | 2.22E-62 |
| Endothelial Cell | Nmrk2 | 8 | 1.24E-65 | 0.88513419 | 0.397 | 0.118 | 2.47E-62 |
| Endothelial Cell | Gpc3 | 8 | 5.73E-65 | 0.5146978 | 0.333 | 0.066 | 1.15E-61 |
| Endothelial Cell | Ripor2 | 8 | 6.05E-65 | 0.71622781 | 0.766 | 0.324 | 1.21E-61 |
| Endothelial Cell | Cox4i1 | 8 | 7.58E-65 | 0.41546247 | 0.778 | 0.344 | 1.52E-61 |
| Endothelial Cell | Psap | 8 | 1.06E-64 | 0.30403327 | 0.768 | 0.344 | 2.12E-61 |
| Endothelial Cell | Gpam | 8 | 1.66E-64 | 0.44549145 | 0.598 | 0.238 | 3.32E-61 |
| Endothelial Cell | Fat3 | 8 | 2.60E-64 | 1.4566558 | 0.507 | 0.223 | 5.21E-61 |
| Endothelial Cell | Me1 | 8 | 2.70E-64 | 0.91989588 | 0.586 | 0.275 | 5.40E-61 |
| Endothelial Cell | Egfr | 8 | 1.88E-63 | 0.8433258 | 0.5 | 0.25 | 3.75E-60 |
| Endothelial Cell | Slc16a2 | 8 | 4.09E-63 | 0.39069688 | 0.624 | 0.289 | 8.18E-60 |
| Endothelial Cell | AABR070275 | 8 | 9.93E-63 | 1.01196568 | 0.871 | 0.476 | 1.99E-59 |
| Endothelial Cell | Iqgap3 | 8 | 2.53E-62 | 1.65228431 | 0.541 | 0.124 | 5.07E-59 |
| Endothelial Cell | Mis18bp1 | 8 | 4.37E-62 | 0.62826513 | 0.579 | 0.251 | 8.73E-59 |
| Endothelial Cell | Robo1 | 8 | 1.31E-60 | 0.44969058 | 0.603 | 0.313 | 2.61E-57 |
| Endothelial Cell | Kank1 | 8 | 4.04E-60 | 0.3025122 | 0.632 | 0.293 | 8.09E-57 |
| Endothelial Cell | Fam78b | 8 | 4.51E-60 | 0.81463285 | 0.641 | 0.312 | 9.01E-57 |
| Endothelial Cell | Ar | 8 | 9.46E-60 | 0.67621874 | 0.608 | 0.32 | 1.89E-56 |
| Endothelial Cell | Aatk | 8 | 8.08E-59 | 0.98000801 | 0.658 | 0.34 | 1.62E-55 |
| Endothelial Cell | Irs2 | 8 | 8.55E-59 | 0.4321218 | 0.584 | 0.332 | 1.71E-55 |

| | | | | | | | |
|---|---|---|---|---|---|---|---|
| Endothelial Cell | Ngf | 8 | 1.85E-58 | 0.47836902 | 0.603 | 0.303 | 3.69E-55 |
| Endothelial Cell | Rnls | 8 | 4.98E-58 | 0.3561318 | 0.739 | 0.347 | 9.97E-55 |
| Endothelial Cell | Cgnl1 | 8 | 5.43E-58 | 0.37022412 | 0.876 | 0.442 | 1.09E-54 |
| Endothelial Cell | Enah | 8 | 8.67E-58 | 0.33976493 | 0.677 | 0.31 | 1.73E-54 |
| Endothelial Cell | Lrrtm4 | 8 | 1.14E-57 | 1.30040732 | 0.514 | 0.127 | 2.28E-54 |
| Endothelial Cell | Sphkap | 8 | 1.80E-57 | 0.3951992 | 0.701 | 0.295 | 3.59E-54 |
| Endothelial Cell | Neb | 8 | 7.15E-57 | 0.39574621 | 0.565 | 0.178 | 1.43E-53 |
| Endothelial Cell | Mfsd4a | 8 | 1.02E-56 | 0.47232917 | 0.622 | 0.312 | 2.03E-53 |
| Endothelial Cell | Cacna1c | 8 | 3.39E-56 | 0.34497356 | 0.565 | 0.27 | 6.78E-53 |
| Endothelial Cell | Dsp | 8 | 1.77E-55 | 0.27099992 | 0.586 | 0.288 | 3.54E-52 |
| Endothelial Cell | Hcn1 | 8 | 4.83E-55 | 0.73665246 | 0.634 | 0.213 | 9.66E-52 |
| Endothelial Cell | C7 | 8 | 1.33E-54 | 0.8670228 | 0.648 | 0.337 | 2.67E-51 |
| Endothelial Cell | AABR070449 | 8 | 2.48E-54 | 0.66407164 | 0.691 | 0.357 | 4.95E-51 |
| Endothelial Cell | Vegfd | 8 | 5.76E-54 | 0.5717291 | 0.548 | 0.292 | 1.15E-50 |
| Endothelial Cell | Slc12a2 | 8 | 6.81E-54 | 0.94883401 | 0.885 | 0.486 | 1.36E-50 |
| Endothelial Cell | Ntf31 | 8 | 1.12E-53 | 0.55559457 | 0.892 | 0.446 | 2.23E-50 |
| Endothelial Cell | Matn2 | 8 | 1.28E-52 | 0.35833351 | 0.555 | 0.268 | 2.57E-49 |
| Endothelial Cell | Xirp2 | 8 | 2.85E-52 | 0.633971 | 0.567 | 0.304 | 5.70E-49 |
| Endothelial Cell | Sh3bp2 | 8 | 2.52E-51 | 0.52232965 | 0.574 | 0.238 | 5.05E-48 |
| Endothelial Cell | Fap | 8 | 3.37E-51 | 0.46642917 | 0.888 | 0.476 | 6.73E-48 |
| Endothelial Cell | Rgcc | 8 | 7.73E-50 | 0.66660387 | 0.866 | 0.513 | 1.55E-46 |
| Endothelial Cell | Pf4 | 8 | 1.65E-49 | 0.71666074 | 0.467 | 0.173 | 3.30E-46 |
| Endothelial Cell | Fabp5 | 8 | 2.67E-49 | 0.26903087 | 0.797 | 0.37 | 5.34E-46 |
| Endothelial Cell | Itga8 | 8 | 5.49E-49 | 0.76199987 | 0.553 | 0.303 | 1.10E-45 |
| Endothelial Cell | Chsy3 | 8 | 3.35E-48 | 0.32649057 | 0.622 | 0.334 | 6.70E-45 |
| Endothelial Cell | Mfge8 | 8 | 6.81E-48 | 0.29192647 | 0.656 | 0.344 | 1.36E-44 |
| Endothelial Cell | Acsl1 | 8 | 1.39E-47 | 0.32875892 | 0.6 | 0.338 | 2.78E-44 |
| Endothelial Cell | Tmem196 | 8 | 7.41E-47 | 0.30518479 | 0.672 | 0.356 | 1.48E-43 |
| Endothelial Cell | Cyp7b1 | 8 | 1.21E-46 | 0.63422083 | 0.854 | 0.49 | 2.42E-43 |
| Endothelial Cell | Rcan2 | 8 | 1.43E-45 | 0.28113477 | 0.732 | 0.391 | 2.86E-42 |
| Endothelial Cell | Inpp5d | 8 | 1.44E-44 | 0.38923576 | 0.801 | 0.427 | 2.88E-41 |

| | | | | | | | |
|---|---|---|---|---|---|---|---|
| Endothelial Cell | Dcn | 8 | 2.85E-44 | 0.26606735 | 0.579 | 0.302 | 5.70E-41 |
| Endothelial Cell | LOC1003644 | 8 | 2.36E-43 | 0.32869268 | 0.612 | 0.349 | 4.71E-40 |
| Endothelial Cell | Gucy1a2 | 8 | 8.24E-43 | 0.36959429 | 0.612 | 0.353 | 1.65E-39 |
| Endothelial Cell | Acacb | 8 | 1.42E-42 | 0.41548662 | 0.591 | 0.325 | 2.84E-39 |
| Endothelial Cell | AABR070490 | 8 | 7.86E-42 | 0.39539535 | 0.605 | 0.345 | 1.57E-38 |
| Endothelial Cell | Sh3gl2 | 8 | 8.14E-40 | 0.46832224 | 0.833 | 0.473 | 1.63E-36 |
| Endothelial Cell | Wt1 | 8 | 1.02E-39 | 0.45959097 | 0.938 | 0.575 | 2.04E-36 |
| Endothelial Cell | Mki671 | 8 | 4.61E-39 | 0.40580353 | 0.548 | 0.262 | 9.22E-36 |
| Endothelial Cell | Actn2 | 8 | 1.56E-38 | 0.31303705 | 0.624 | 0.357 | 3.13E-35 |
| Endothelial Cell | AABR070251 | 8 | 5.55E-38 | 0.35989274 | 0.703 | 0.424 | 1.11E-34 |
| Endothelial Cell | Sema3d | 8 | 2.73E-37 | 0.77437909 | 0.677 | 0.423 | 5.47E-34 |
| Endothelial Cell | Actc1 | 8 | 2.78E-36 | 0.37641092 | 0.799 | 0.447 | 5.57E-33 |
| Endothelial Cell | Kcng2 | 8 | 4.59E-36 | 0.38538918 | 0.639 | 0.307 | 9.19E-33 |
| Endothelial Cell | Fhod3 | 8 | 1.35E-35 | 0.78732341 | 0.837 | 0.523 | 2.71E-32 |
| Endothelial Cell | Slc26a10 | 8 | 1.67E-35 | 0.86078689 | 0.976 | 0.667 | 3.34E-32 |
| Endothelial Cell | Shb | 8 | 9.92E-35 | 0.34389097 | 0.842 | 0.5 | 1.98E-31 |
| Endothelial Cell | Tmtc2 | 8 | 1.64E-34 | 0.77534271 | 0.911 | 0.593 | 3.28E-31 |
| Endothelial Cell | Herc6 | 8 | 4.88E-33 | 0.50033485 | 0.904 | 0.541 | 9.77E-30 |
| Endothelial Cell | Gpm6a | 8 | 9.22E-32 | 0.60915443 | 0.821 | 0.493 | 1.84E-28 |
| Endothelial Cell | Myl6 | 8 | 3.98E-30 | 0.33047386 | 0.816 | 0.473 | 7.95E-27 |
| Endothelial Cell | Tbc1d4 | 8 | 5.55E-30 | 0.40424329 | 0.78 | 0.448 | 1.11E-26 |
| Endothelial Cell | Trak2 | 8 | 8.61E-30 | 0.30087724 | 0.766 | 0.468 | 1.72E-26 |
| Endothelial Cell | AC110709.2 | 8 | 9.62E-30 | 0.66637778 | 0.447 | 0.152 | 1.92E-26 |
| Endothelial Cell | Aqp1 | 8 | 1.35E-29 | 0.37565921 | 0.785 | 0.496 | 2.70E-26 |
| Endothelial Cell | Dapk2 | 8 | 3.64E-28 | 0.34318603 | 0.842 | 0.524 | 7.28E-25 |
| Endothelial Cell | Slc25a21 | 8 | 3.65E-28 | 0.90218595 | 0.421 | 0.145 | 7.30E-25 |
| Endothelial Cell | Slc6a6 | 8 | 6.17E-27 | 0.5118557 | 0.928 | 0.594 | 1.23E-23 |
| Endothelial Cell | Akr1c15 | 8 | 3.25E-26 | 0.68104599 | 0.931 | 0.642 | 6.51E-23 |
| Endothelial Cell | Polr2m | 8 | 1.05E-25 | 0.25403268 | 0.828 | 0.512 | 2.10E-22 |
| Endothelial Cell | Tspan18 | 8 | 3.33E-21 | 0.44722716 | 0.911 | 0.623 | 6.67E-18 |
| Endothelial Cell | B2m | 8 | 1.41E-20 | 0.27482199 | 0.892 | 0.552 | 2.81E-17 |

| | | | | | | | |
|---|---|---|---|---|---|---|---|
| Endothelial Cell | Cald1 | 8 | 6.48E-16 | 0.46833237 | 0.897 | 0.63 | 1.30E-12 |
| Endothelial Cell | Podxl | 8 | 1.94E-15 | 0.49693585 | 0.919 | 0.66 | 3.87E-12 |
| Endothelial Cell | Zfp366 | 8 | 4.50E-14 | 0.36326235 | 0.957 | 0.702 | 9.00E-11 |
| Endothelial Cell | Timp3 | 8 | 9.64E-13 | 0.26305212 | 0.892 | 0.64 | 1.93E-09 |
| Endothelial Cell | Gpx2 | 9 | 0 | 8.0331954 | 0.367 | 0.001 | 0 |
| Endothelial Cell | Tal2 | 9 | 0 | 6.07804143 | 0.328 | 0.009 | 0 |
| Endothelial Cell | Col9a1 | 9 | 0 | 6.86463527 | 0.261 | 0.007 | 0 |
| Endothelial Cell | AC111804.2 | 9 | 9.77E-264 | 1.91956739 | 0.347 | 0.017 | 1.95E-260 |
| Endothelial Cell | Sbspon | 9 | 2.34E-222 | 6.51125047 | 0.314 | 0.022 | 4.68E-219 |
| Endothelial Cell | AABR070674 | 9 | 5.83E-167 | 4.29551109 | 0.292 | 0.004 | 1.17E-163 |
| Endothelial Cell | Stk26 | 9 | 2.06E-155 | 2.79880413 | 0.375 | 0.009 | 4.12E-152 |
| Endothelial Cell | Sulf12 | 9 | 2.49E-151 | 2.80825312 | 0.961 | 0.487 | 4.98E-148 |
| Endothelial Cell | RGD1564053 | 9 | 6.75E-150 | 3.60243278 | 0.356 | 0.007 | 1.35E-146 |
| Endothelial Cell | Tox2 | 9 | 8.78E-147 | 4.94886303 | 0.556 | 0.125 | 1.76E-143 |
| Endothelial Cell | Dkk2 | 9 | 7.12E-140 | 7.66227355 | 0.8 | 0.11 | 1.42E-136 |
| Endothelial Cell | LOC1003610 | 9 | 8.65E-136 | 3.36196858 | 0.911 | 0.487 | 1.73E-132 |
| Endothelial Cell | Nebl2 | 9 | 9.69E-131 | 2.37132967 | 0.961 | 0.507 | 1.94E-127 |
| Endothelial Cell | Ptprj | 9 | 4.45E-118 | 4.29323059 | 0.85 | 0.313 | 8.89E-115 |
| Endothelial Cell | Ldb22 | 9 | 4.97E-116 | 2.9793544 | 0.864 | 0.382 | 9.93E-113 |
| Endothelial Cell | Tiam1 | 9 | 4.30E-112 | 2.16843875 | 0.367 | 0.021 | 8.59E-109 |
| Endothelial Cell | Gabrp | 9 | 2.50E-108 | 6.83449762 | 0.364 | 0.073 | 4.99E-105 |
| Endothelial Cell | Hoxd3 | 9 | 6.28E-107 | 12.1881762 | 0.289 | 0 | 1.26E-103 |
| Endothelial Cell | Bmx | 9 | 2.00E-101 | 5.26171981 | 0.656 | 0.243 | 4.00E-98 |
| Endothelial Cell | Specc1 | 9 | 2.77E-88 | 2.95609028 | 0.8 | 0.41 | 5.54E-85 |
| Endothelial Cell | S100a3 | 9 | 4.84E-82 | 2.37525463 | 0.336 | 0.052 | 9.67E-79 |
| Endothelial Cell | Gja51 | 9 | 7.09E-80 | 3.74788696 | 0.597 | 0.219 | 1.42E-76 |
| Endothelial Cell | Plcg21 | 9 | 1.30E-78 | 4.34830445 | 0.678 | 0.334 | 2.60E-75 |
| Endothelial Cell | Tmem63c | 9 | 4.50E-78 | 5.24206026 | 0.658 | 0.271 | 9.00E-75 |
| Endothelial Cell | Rab151 | 9 | 1.19E-75 | 1.01237379 | 0.419 | 0.054 | 2.37E-72 |
| Endothelial Cell | Eva1c1 | 9 | 2.60E-74 | 2.45683058 | 0.817 | 0.423 | 5.20E-71 |
| Endothelial Cell | Ifitm102 | 9 | 1.17E-70 | 3.2932183 | 0.717 | 0.284 | 2.34E-67 |

| | | | | | | | |
|---|---|---|---|---|---|---|---|
| Endothelial Cell | Afap1l2 | 9 | 5.31E-70 | 3.14886979 | 0.753 | 0.376 | 1.06E-66 |
| Endothelial Cell | AC118957.11 | 9 | 9.26E-67 | 2.86659507 | 0.328 | 0.063 | 1.85E-63 |
| Endothelial Cell | Olr1 | 9 | 5.31E-66 | 0.52851783 | 0.342 | 0.043 | 1.06E-62 |
| Endothelial Cell | AABR070359 | 9 | 9.58E-66 | 2.04696643 | 0.811 | 0.51 | 1.92E-62 |
| Endothelial Cell | Armc4 | 9 | 4.84E-63 | 2.1180646 | 0.397 | 0.008 | 9.68E-60 |
| Endothelial Cell | Nuak11 | 9 | 8.13E-62 | 1.70607262 | 0.833 | 0.517 | 1.63E-58 |
| Endothelial Cell | Ncam2 | 9 | 1.56E-58 | 2.50778243 | 0.353 | 0.056 | 3.11E-55 |
| Endothelial Cell | Pcsk51 | 9 | 1.51E-57 | 2.77008573 | 0.692 | 0.335 | 3.01E-54 |
| Endothelial Cell | Col8a1 | 9 | 2.49E-57 | 2.22983689 | 0.828 | 0.528 | 4.99E-54 |
| Endothelial Cell | Dgkb | 9 | 1.62E-51 | 3.3909847 | 0.692 | 0.35 | 3.23E-48 |
| Endothelial Cell | Kcnab1 | 9 | 3.71E-50 | 5.22566244 | 0.639 | 0.297 | 7.42E-47 |
| Endothelial Cell | AABR070311 | 9 | 1.43E-48 | 3.83313765 | 0.519 | 0.149 | 2.86E-45 |
| Endothelial Cell | Vsnl1 | 9 | 3.98E-47 | 0.77069934 | 0.392 | 0.102 | 7.96E-44 |
| Endothelial Cell | Sox171 | 9 | 5.31E-46 | 1.94189945 | 0.731 | 0.384 | 1.06E-42 |
| Endothelial Cell | Pdgfd1 | 9 | 3.44E-45 | 1.72734743 | 0.781 | 0.489 | 6.87E-42 |
| Endothelial Cell | Col18a1 | 9 | 1.32E-44 | 2.10032009 | 0.714 | 0.427 | 2.64E-41 |
| Endothelial Cell | Pid1 | 9 | 6.45E-44 | 2.57824973 | 0.692 | 0.31 | 1.29E-40 |
| Endothelial Cell | Rtn4rl1 | 9 | 1.39E-43 | 4.19534461 | 0.625 | 0.3 | 2.78E-40 |
| Endothelial Cell | Kcnn4 | 9 | 4.07E-43 | 3.41817839 | 0.558 | 0.24 | 8.14E-40 |
| Endothelial Cell | Il1rn | 9 | 4.41E-39 | 2.00516493 | 0.303 | 0.037 | 8.82E-36 |
| Endothelial Cell | Atp2b2 | 9 | 9.66E-39 | 4.08908289 | 0.592 | 0.31 | 1.93E-35 |
| Endothelial Cell | Atf31 | 9 | 1.34E-38 | 1.13410907 | 0.033 | 0.286 | 2.68E-35 |
| Endothelial Cell | Fgf7 | 9 | 2.63E-38 | 2.58524648 | 0.358 | 0.058 | 5.25E-35 |
| Endothelial Cell | Ctnnd2 | 9 | 1.39E-35 | 1.48415389 | 0.331 | 0.035 | 2.79E-32 |
| Endothelial Cell | Ptprt | 9 | 2.72E-35 | 0.62955393 | 0.433 | 0.083 | 5.44E-32 |
| Endothelial Cell | Susd41 | 9 | 4.80E-32 | 2.20223274 | 0.433 | 0.099 | 9.60E-29 |
| Endothelial Cell | Chn22 | 9 | 5.11E-29 | 1.47601727 | 0.694 | 0.411 | 1.02E-25 |
| Endothelial Cell | Ephb2 | 9 | 9.95E-28 | 3.28904019 | 0.372 | 0.048 | 1.99E-24 |
| Endothelial Cell | Ttk | 9 | 1.03E-27 | 0.92824299 | 0.317 | 0.061 | 2.06E-24 |
| Endothelial Cell | Slc4a4 | 9 | 2.32E-27 | 2.89452117 | 0.569 | 0.254 | 4.64E-24 |
| Endothelial Cell | Fn1 | 9 | 6.79E-27 | 2.04471342 | 0.614 | 0.315 | 1.36E-23 |

| | | | | | | | |
|---|---|---|---|---|---|---|---|
| Endothelial Cell | Tmem163 | 9 | 1.75E-26 | 1.21352263 | 0.325 | 0.074 | 3.50E-23 |
| Endothelial Cell | Epsti1 | 9 | 2.44E-26 | 0.96426398 | 0.419 | 0.136 | 4.88E-23 |
| Endothelial Cell | Fbln21 | 9 | 2.68E-24 | 2.55335187 | 0.544 | 0.274 | 5.36E-21 |
| Endothelial Cell | Setbp1 | 9 | 5.88E-24 | 3.95201408 | 0.514 | 0.235 | 1.18E-20 |
| Endothelial Cell | AABR070410 | 9 | 1.89E-23 | 2.08211844 | 0.472 | 0.197 | 3.79E-20 |
| Endothelial Cell | Rnd3 | 9 | 1.79E-22 | 2.95229416 | 0.561 | 0.257 | 3.59E-19 |
| Endothelial Cell | Rap1gap2 | 9 | 6.28E-22 | 2.97868576 | 0.561 | 0.267 | 1.26E-18 |
| Endothelial Cell | Arhgap44 | 9 | 2.15E-21 | 3.27751244 | 0.533 | 0.276 | 4.30E-18 |
| Endothelial Cell | Fam189a2 | 9 | 4.68E-21 | 2.18735961 | 0.389 | 0.085 | 9.35E-18 |
| Endothelial Cell | Aldh1a3 | 9 | 1.14E-20 | 0.9161781 | 0.439 | 0.123 | 2.29E-17 |
| Endothelial Cell | Cdkn3 | 9 | 3.39E-19 | 1.1970341 | 0.311 | 0.043 | 6.78E-16 |
| Endothelial Cell | Smad62 | 9 | 6.59E-19 | 1.12402516 | 0.669 | 0.405 | 1.32E-15 |
| Endothelial Cell | Marchf11 | 9 | 1.29E-18 | 1.05230432 | 0.444 | 0.165 | 2.58E-15 |
| Endothelial Cell | Tmem1001 | 9 | 1.98E-18 | 2.27497455 | 0.531 | 0.22 | 3.96E-15 |
| Endothelial Cell | Igfbp41 | 9 | 4.70E-18 | 1.93477572 | 0.581 | 0.29 | 9.41E-15 |
| Endothelial Cell | LOC1009114 | 9 | 1.81E-17 | 1.68277419 | 0.489 | 0.148 | 3.62E-14 |
| Endothelial Cell | Grm8 | 9 | 2.31E-17 | 0.91997577 | 0.417 | 0.15 | 4.62E-14 |
| Endothelial Cell | Fam227b | 9 | 3.71E-17 | 0.64695805 | 0.056 | 0.338 | 7.41E-14 |
| Endothelial Cell | AABR070010 | 9 | 1.16E-16 | 0.3404494 | 0.05 | 0.307 | 2.31E-13 |
| Endothelial Cell | Vcan | 9 | 3.99E-15 | 2.39570913 | 0.531 | 0.25 | 7.98E-12 |
| Endothelial Cell | Has1 | 9 | 2.94E-13 | 0.74574833 | 0.35 | 0.089 | 5.87E-10 |
| Endothelial Cell | S100a6 | 9 | 3.14E-10 | 1.73228947 | 0.508 | 0.215 | 6.27E-07 |
| Endothelial Cell | Mob3b | 9 | 1.60E-08 | 1.69427212 | 0.406 | 0.145 | 3.19E-05 |
| Endothelial Cell | Angpt1 | 9 | 2.40E-08 | 1.32467501 | 0.469 | 0.174 | 4.80E-05 |
| Endothelial Cell | Lgmn | 9 | 2.71E-07 | 1.24620514 | 0.453 | 0.181 | 0.00054105 |
| Endothelial Cell | AABR070015 | 9 | 7.67E-06 | 0.544685 | 0.536 | 0.259 | 0.01534414 |
| Endothelial Cell | Ptchd1 | 10 | 0 | 2.69314273 | 0.971 | 0.039 | 0 |
| Endothelial Cell | Megf10 | 10 | 0 | 3.11797879 | 0.982 | 0.073 | 0 |
| Endothelial Cell | Polq | 10 | 0 | 1.542606 | 0.982 | 0.131 | 0 |
| Endothelial Cell | Klri1 | 10 | 0 | 2.76431055 | 0.968 | 0.129 | 0 |
| Endothelial Cell | Fam135b | 10 | 0 | 1.46554151 | 0.757 | 0.056 | 0 |

| | | | | | | | |
|---|---|---|---|---|---|---|---|
| Endothelial Cell | Hk3 | 10 | 4.95E-296 | 1.83518699 | 0.686 | 0.06 | 9.91E-293 |
| Endothelial Cell | Tmem200a1 | 10 | 1.43E-259 | 1.7932701 | 0.979 | 0.221 | 2.85E-256 |
| Endothelial Cell | Ptgs2 | 10 | 2.19E-252 | 2.04603166 | 0.985 | 0.187 | 4.39E-249 |
| Endothelial Cell | Gria2 | 10 | 8.64E-231 | 0.68428377 | 0.959 | 0.177 | 1.73E-227 |
| Endothelial Cell | Mybphl | 10 | 5.52E-221 | 1.46105353 | 0.639 | 0.042 | 1.10E-217 |
| Endothelial Cell | Ikzf1 | 10 | 2.57E-219 | 1.67961282 | 0.979 | 0.234 | 5.14E-216 |
| Endothelial Cell | Apcdd1l | 10 | 1.17E-216 | 0.92667263 | 0.563 | 0.043 | 2.35E-213 |
| Endothelial Cell | Sv2c | 10 | 1.99E-214 | 1.4493224 | 0.991 | 0.324 | 3.99E-211 |
| Endothelial Cell | Fa2h | 10 | 6.86E-209 | 2.64130324 | 0.651 | 0.057 | 1.37E-205 |
| Endothelial Cell | RGD1559482 | 10 | 4.26E-194 | 1.18411666 | 0.95 | 0.294 | 8.51E-191 |
| Endothelial Cell | B4galnt3 | 10 | 5.86E-193 | 2.66756844 | 0.44 | 0.046 | 1.17E-189 |
| Endothelial Cell | Tafa21 | 10 | 1.53E-190 | 1.52345146 | 0.988 | 0.32 | 3.06E-187 |
| Endothelial Cell | Tm4sf191 | 10 | 3.24E-190 | 1.36420952 | 0.786 | 0.118 | 6.49E-187 |
| Endothelial Cell | Nmrk21 | 10 | 2.66E-187 | 0.27845987 | 0.443 | 0.119 | 5.32E-184 |
| Endothelial Cell | Chrdl1 | 10 | 1.69E-177 | 0.78361078 | 0.977 | 0.349 | 3.38E-174 |
| Endothelial Cell | Sh3bp21 | 10 | 9.33E-177 | 0.66871661 | 0.927 | 0.226 | 1.87E-173 |
| Endothelial Cell | Glb1l2 | 10 | 4.28E-175 | 1.3698317 | 0.941 | 0.262 | 8.56E-172 |
| Endothelial Cell | Trem2 | 10 | 1.13E-174 | 2.14931835 | 0.716 | 0.128 | 2.27E-171 |
| Endothelial Cell | Efna51 | 10 | 3.93E-174 | 1.86753037 | 0.859 | 0.188 | 7.87E-171 |
| Endothelial Cell | Kcnq3 | 10 | 4.31E-173 | 0.30278505 | 0.639 | 0.096 | 8.61E-170 |
| Endothelial Cell | Slc5a1 | 10 | 2.26E-169 | 2.03891661 | 0.628 | 0.1 | 4.51E-166 |
| Endothelial Cell | St8sia1 | 10 | 9.13E-168 | 1.50156829 | 0.569 | 0.08 | 1.83E-164 |
| Endothelial Cell | Pdpn | 10 | 1.49E-166 | 1.65041333 | 0.54 | 0.041 | 2.98E-163 |
| Endothelial Cell | Gen11 | 10 | 1.06E-163 | 1.22288626 | 0.906 | 0.248 | 2.13E-160 |
| Endothelial Cell | PCOLCE2 | 10 | 6.50E-162 | 2.00415345 | 0.804 | 0.195 | 1.30E-158 |
| Endothelial Cell | Mark11 | 10 | 2.41E-157 | 1.77058025 | 0.762 | 0.21 | 4.82E-154 |
| Endothelial Cell | Ndst3 | 10 | 2.28E-153 | 1.40471123 | 0.745 | 0.17 | 4.56E-150 |
| Endothelial Cell | Cdh20 | 10 | 4.88E-153 | 0.75422015 | 0.845 | 0.173 | 9.76E-150 |
| Endothelial Cell | Cdh4 | 10 | 3.24E-149 | 2.03062816 | 0.364 | 0.05 | 6.49E-146 |
| Endothelial Cell | Neil3 | 10 | 5.10E-149 | 0.98851744 | 0.651 | 0.182 | 1.02E-145 |
| Endothelial Cell | Ildr2 | 10 | 2.31E-144 | 1.87509825 | 0.622 | 0.15 | 4.62E-141 |

| | | | | | | | |
|---|---|---|---|---|---|---|---|
| Endothelial Cell | Sntg2 | 10 | 1.69E-143 | 1.04064621 | 0.563 | 0.079 | 3.37E-140 |
| Endothelial Cell | Scube1 | 10 | 7.00E-142 | 0.83068927 | 0.965 | 0.394 | 1.40E-138 |
| Endothelial Cell | Pou2f2 | 10 | 1.10E-141 | 1.2960119 | 0.71 | 0.198 | 2.20E-138 |
| Endothelial Cell | Fabp51 | 10 | 2.62E-141 | 1.108495 | 0.93 | 0.368 | 5.25E-138 |
| Endothelial Cell | Ciita | 10 | 8.08E-140 | 0.37396044 | 0.592 | 0.086 | 1.62E-136 |
| Endothelial Cell | Serpine1 | 10 | 1.27E-139 | 1.81623141 | 0.798 | 0.236 | 2.54E-136 |
| Endothelial Cell | Tnn1 | 10 | 1.42E-139 | 0.62452024 | 0.733 | 0.192 | 2.84E-136 |
| Endothelial Cell | Gpam1 | 10 | 3.25E-137 | 1.11343542 | 0.824 | 0.231 | 6.49E-134 |
| Endothelial Cell | Aspm | 10 | 1.39E-136 | 0.93191128 | 0.516 | 0.097 | 2.79E-133 |
| Endothelial Cell | Iqgap2 | 10 | 1.57E-136 | 0.30006625 | 0.812 | 0.195 | 3.15E-133 |
| Endothelial Cell | AABR070171 | 10 | 2.51E-136 | 0.62984715 | 0.883 | 0.259 | 5.02E-133 |
| Endothelial Cell | Tpm21 | 10 | 9.81E-133 | 0.77233844 | 0.921 | 0.296 | 1.96E-129 |
| Endothelial Cell | Fcrl5 | 10 | 1.54E-131 | 0.48902431 | 0.657 | 0.137 | 3.08E-128 |
| Endothelial Cell | Rimbp2 | 10 | 8.81E-130 | 1.87209869 | 0.994 | 0.442 | 1.76E-126 |
| Endothelial Cell | Mybpc2 | 10 | 3.01E-129 | 1.00943343 | 0.455 | 0.11 | 6.02E-126 |
| Endothelial Cell | Notch3 | 10 | 2.71E-128 | 1.05613238 | 0.751 | 0.223 | 5.42E-125 |
| Endothelial Cell | Fcrla | 10 | 1.56E-127 | 1.86068014 | 0.302 | 0.019 | 3.11E-124 |
| Endothelial Cell | Sgip1 | 10 | 7.38E-125 | 0.71842595 | 0.777 | 0.206 | 1.48E-121 |
| Endothelial Cell | Uqcr101 | 10 | 8.87E-125 | 0.66027827 | 0.959 | 0.338 | 1.77E-121 |
| Endothelial Cell | AABR070336 | 10 | 9.14E-123 | 2.31341087 | 0.273 | 0.017 | 1.83E-119 |
| Endothelial Cell | Cd84 | 10 | 1.05E-121 | 1.22961843 | 0.346 | 0.044 | 2.10E-118 |
| Endothelial Cell | Coro6 | 10 | 6.53E-121 | 0.28967986 | 0.809 | 0.224 | 1.31E-117 |
| Endothelial Cell | Gfra21 | 10 | 6.86E-121 | 1.61124392 | 0.768 | 0.243 | 1.37E-117 |
| Endothelial Cell | Dsg2 | 10 | 1.85E-119 | 0.43800519 | 0.748 | 0.206 | 3.71E-116 |
| Endothelial Cell | Zfhx4 | 10 | 9.61E-119 | 2.06698013 | 0.534 | 0.066 | 1.92E-115 |
| Endothelial Cell | AC111831.1 | 10 | 1.94E-117 | 1.72789479 | 0.411 | 0.143 | 3.87E-114 |
| Endothelial Cell | Smoc11 | 10 | 4.16E-117 | 1.29078078 | 0.61 | 0.153 | 8.32E-114 |
| Endothelial Cell | Tagln1 | 10 | 2.68E-116 | 1.41770363 | 0.777 | 0.253 | 5.37E-113 |
| Endothelial Cell | Scn7a | 10 | 2.69E-116 | 1.35072128 | 0.736 | 0.224 | 5.39E-113 |
| Endothelial Cell | Pcsk6 | 10 | 8.97E-116 | 1.0582201 | 0.877 | 0.259 | 1.79E-112 |
| Endothelial Cell | Srpk31 | 10 | 5.39E-115 | 1.64099467 | 0.848 | 0.258 | 1.08E-111 |

| | | | | | | | |
|---|---|---|---|---|---|---|---|
| Endothelial Cell | Cacna1g | 10 | 2.30E-114 | 0.31732885 | 0.935 | 0.294 | 4.59E-111 |
| Endothelial Cell | Efemp1 | 10 | 2.62E-113 | 0.89552521 | 0.751 | 0.243 | 5.23E-110 |
| Endothelial Cell | Galnt17 | 10 | 3.53E-112 | 1.06562518 | 0.812 | 0.26 | 7.06E-109 |
| Endothelial Cell | Uqcrq | 10 | 4.06E-112 | 0.44482438 | 0.9 | 0.248 | 8.13E-109 |
| Endothelial Cell | Gria41 | 10 | 8.12E-112 | 1.18249642 | 0.686 | 0.179 | 1.62E-108 |
| Endothelial Cell | Tnfrsf19 | 10 | 4.15E-111 | 1.16058569 | 0.51 | 0.121 | 8.31E-108 |
| Endothelial Cell | Trib3 | 10 | 6.62E-111 | 2.00411668 | 0.798 | 0.166 | 1.32E-107 |
| Endothelial Cell | Fbxo40 | 10 | 1.27E-110 | 1.31721661 | 0.622 | 0.115 | 2.53E-107 |
| Endothelial Cell | Cd4 | 10 | 2.66E-110 | 1.68658258 | 0.628 | 0.18 | 5.31E-107 |
| Endothelial Cell | Trpc6 | 10 | 9.65E-110 | 1.28645855 | 0.534 | 0.117 | 1.93E-106 |
| Endothelial Cell | Col24a1 | 10 | 2.12E-109 | 1.04532135 | 0.933 | 0.395 | 4.25E-106 |
| Endothelial Cell | Ctsb | 10 | 5.04E-109 | 0.5777167 | 0.859 | 0.266 | 1.01E-105 |
| Endothelial Cell | Alpl | 10 | 1.43E-108 | 1.22736842 | 0.924 | 0.36 | 2.85E-105 |
| Endothelial Cell | Fhl1 | 10 | 1.84E-108 | 0.7303746 | 0.874 | 0.267 | 3.67E-105 |
| Endothelial Cell | AABR070278 | 10 | 3.40E-108 | 2.10410737 | 0.522 | 0.105 | 6.81E-105 |
| Endothelial Cell | C6 | 10 | 3.75E-105 | 1.95779041 | 0.563 | 0.091 | 7.49E-102 |
| Endothelial Cell | Col12a1 | 10 | 5.08E-105 | 0.57705787 | 0.733 | 0.226 | 1.02E-101 |
| Endothelial Cell | Runx11 | 10 | 5.67E-104 | 0.97182833 | 0.868 | 0.337 | 1.13E-100 |
| Endothelial Cell | Fgf11 | 10 | 1.35E-103 | 0.62613365 | 0.891 | 0.304 | 2.71E-100 |
| Endothelial Cell | Gnao1 | 10 | 1.59E-103 | 1.1238814 | 0.947 | 0.397 | 3.19E-100 |
| Endothelial Cell | Vash2 | 10 | 2.58E-103 | 1.07046216 | 0.522 | 0.115 | 5.15E-100 |
| Endothelial Cell | Col11a11 | 10 | 9.12E-103 | 0.7084033 | 0.877 | 0.293 | 1.82E-99 |
| Endothelial Cell | AABR070172 | 10 | 1.75E-102 | 0.46033711 | 0.692 | 0.176 | 3.51E-99 |
| Endothelial Cell | LOC690045 | 10 | 1.84E-102 | 0.57204795 | 0.85 | 0.274 | 3.68E-99 |
| Endothelial Cell | Cadps | 10 | 6.04E-102 | 0.38146207 | 0.754 | 0.303 | 1.21E-98 |
| Endothelial Cell | Cemip | 10 | 3.25E-101 | 1.22191328 | 0.874 | 0.311 | 6.49E-98 |
| Endothelial Cell | Slco4a1 | 10 | 3.41E-101 | 1.99688546 | 0.367 | 0.037 | 6.81E-98 |
| Endothelial Cell | Slc1a3 | 10 | 4.09E-101 | 1.45274376 | 0.674 | 0.213 | 8.18E-98 |
| Endothelial Cell | Astn2 | 10 | 7.74E-101 | 0.2516787 | 0.727 | 0.233 | 1.55E-97 |
| Endothelial Cell | Lmod1 | 10 | 2.10E-100 | 1.55455964 | 0.818 | 0.255 | 4.20E-97 |
| Endothelial Cell | Acacb1 | 10 | 2.17E-100 | 0.89193602 | 0.93 | 0.313 | 4.33E-97 |

| | | | | | | | |
|---|---|---|---|---|---|---|---|
| Endothelial Cell | Asb2 | 10 | 4.49E-100 | 0.75636489 | 0.748 | 0.231 | 8.98E-97 |
| Endothelial Cell | Nkd21 | 10 | 8.04E-100 | 0.47052138 | 0.953 | 0.363 | 1.61E-96 |
| Endothelial Cell | Aoah | 10 | 1.02E-98 | 1.10239516 | 0.751 | 0.281 | 2.03E-95 |
| Endothelial Cell | Myom2 | 10 | 3.20E-97 | 0.76870072 | 0.727 | 0.23 | 6.40E-94 |
| Endothelial Cell | Ntrk2 | 10 | 7.13E-97 | 1.28212569 | 0.475 | 0.106 | 1.43E-93 |
| Endothelial Cell | Tacr1 | 10 | 2.41E-96 | 1.38099216 | 0.674 | 0.158 | 4.82E-93 |
| Endothelial Cell | Kcnj81 | 10 | 2.09E-95 | 1.02847249 | 0.496 | 0.181 | 4.17E-92 |
| Endothelial Cell | Has2 | 10 | 1.32E-94 | 0.41347844 | 0.616 | 0.23 | 2.64E-91 |
| Endothelial Cell | Dpf3 | 10 | 1.59E-94 | 0.38898174 | 0.883 | 0.316 | 3.18E-91 |
| Endothelial Cell | Pgam21 | 10 | 5.90E-93 | 0.35044267 | 0.736 | 0.242 | 1.18E-89 |
| Endothelial Cell | Matn21 | 10 | 5.23E-92 | 0.7511529 | 0.762 | 0.262 | 1.05E-88 |
| Endothelial Cell | C1qa | 10 | 2.49E-91 | 1.09215587 | 0.88 | 0.317 | 4.97E-88 |
| Endothelial Cell | Arhgap22 | 10 | 4.19E-91 | 0.5196226 | 0.789 | 0.302 | 8.39E-88 |
| Endothelial Cell | Il1rapl1 | 10 | 1.22E-90 | 1.55046275 | 0.672 | 0.253 | 2.44E-87 |
| Endothelial Cell | Apba1 | 10 | 1.76E-90 | 0.83508692 | 0.903 | 0.36 | 3.52E-87 |
| Endothelial Cell | Vwc2 | 10 | 4.17E-90 | 0.79573452 | 0.472 | 0.112 | 8.33E-87 |
| Endothelial Cell | Gpx3 | 10 | 1.00E-89 | 1.13917689 | 0.789 | 0.291 | 2.00E-86 |
| Endothelial Cell | Rcan11 | 10 | 3.00E-89 | 1.4789718 | 0.651 | 0.233 | 6.01E-86 |
| Endothelial Cell | Adamtsl3 | 10 | 6.08E-88 | 1.00776254 | 0.865 | 0.322 | 1.22E-84 |
| Endothelial Cell | Neb1 | 10 | 1.07E-87 | 0.48397441 | 0.607 | 0.18 | 2.14E-84 |
| Endothelial Cell | Kcnh8 | 10 | 2.55E-87 | 0.80732998 | 0.569 | 0.204 | 5.10E-84 |
| Endothelial Cell | Pltp1 | 10 | 4.23E-87 | 0.45791628 | 0.792 | 0.334 | 8.46E-84 |
| Endothelial Cell | Susd1 | 10 | 1.49E-86 | 0.41445387 | 0.669 | 0.234 | 2.99E-83 |
| Endothelial Cell | Vegfd1 | 10 | 1.78E-86 | 1.3486567 | 0.751 | 0.286 | 3.56E-83 |
| Endothelial Cell | Col4a3 | 10 | 2.39E-86 | 0.92696829 | 0.645 | 0.214 | 4.78E-83 |
| Endothelial Cell | Parm1 | 10 | 1.12E-85 | 1.2973615 | 0.663 | 0.237 | 2.24E-82 |
| Endothelial Cell | Lrrc4b | 10 | 2.41E-85 | 1.44071738 | 0.513 | 0.205 | 4.81E-82 |
| Endothelial Cell | Btnl92 | 10 | 6.32E-85 | 1.61895509 | 0.868 | 0.37 | 1.26E-81 |
| Endothelial Cell | Fyb1 | 10 | 1.82E-84 | 0.71169763 | 0.672 | 0.255 | 3.63E-81 |
| Endothelial Cell | Scg2 | 10 | 6.23E-84 | 0.32542124 | 0.463 | 0.177 | 1.25E-80 |
| Endothelial Cell | Dpyd | 10 | 6.52E-83 | 0.76510797 | 0.554 | 0.174 | 1.30E-79 |

| | | | | | | | |
|---|---|---|---|---|---|---|---|
| Endothelial Cell | Cadm2 | 10 | 6.63E-83 | 0.87882407 | 0.378 | 0.082 | 1.33E-79 |
| Endothelial Cell | Csrp31 | 10 | 1.09E-82 | 0.83780196 | 0.73 | 0.29 | 2.18E-79 |
| Endothelial Cell | Col14a11 | 10 | 1.20E-82 | 0.90332281 | 0.639 | 0.239 | 2.40E-79 |
| Endothelial Cell | Abca17 | 10 | 1.49E-82 | 1.24324939 | 0.85 | 0.35 | 2.98E-79 |
| Endothelial Cell | Ckmt2 | 10 | 3.55E-82 | 0.73778899 | 0.886 | 0.358 | 7.10E-79 |
| Endothelial Cell | Musk1 | 10 | 3.93E-82 | 1.55768271 | 0.669 | 0.149 | 7.87E-79 |
| Endothelial Cell | Lrrc4c | 10 | 5.54E-82 | 0.85549227 | 0.622 | 0.243 | 1.11E-78 |
| Endothelial Cell | Gfpt2 | 10 | 8.35E-82 | 0.85138207 | 0.683 | 0.254 | 1.67E-78 |
| Endothelial Cell | Hao1 | 10 | 7.55E-81 | 2.19944128 | 0.323 | 0.021 | 1.51E-77 |
| Endothelial Cell | Corin1 | 10 | 8.77E-81 | 1.04485005 | 0.713 | 0.288 | 1.75E-77 |
| Endothelial Cell | Pygm | 10 | 1.38E-80 | 0.7997986 | 0.674 | 0.275 | 2.77E-77 |
| Endothelial Cell | Gucy1a1 | 10 | 1.44E-80 | 0.48569384 | 0.777 | 0.303 | 2.88E-77 |
| Endothelial Cell | Ckb | 10 | 1.09E-79 | 0.80892701 | 0.463 | 0.181 | 2.19E-76 |
| Endothelial Cell | Ect21 | 10 | 1.31E-79 | 2.02084671 | 0.431 | 0.173 | 2.61E-76 |
| Endothelial Cell | Ctsd | 10 | 1.87E-79 | 0.89906149 | 0.686 | 0.248 | 3.74E-76 |
| Endothelial Cell | Macrod1 | 10 | 2.36E-79 | 0.31875961 | 0.595 | 0.219 | 4.73E-76 |
| Endothelial Cell | Ndc801 | 10 | 4.40E-79 | 0.56037301 | 0.783 | 0.28 | 8.79E-76 |
| Endothelial Cell | Pid11 | 10 | 5.54E-79 | 0.75046962 | 0.862 | 0.304 | 1.11E-75 |
| Endothelial Cell | Atp5mc1 | 10 | 5.81E-79 | 0.8823391 | 0.742 | 0.288 | 1.16E-75 |
| Endothelial Cell | Robo21 | 10 | 9.65E-79 | 0.93556582 | 0.61 | 0.225 | 1.93E-75 |
| Endothelial Cell | Cd44 | 10 | 2.66E-78 | 0.61487407 | 0.891 | 0.383 | 5.32E-75 |
| Endothelial Cell | Adhfe1 | 10 | 5.05E-78 | 0.79925621 | 0.628 | 0.249 | 1.01E-74 |
| Endothelial Cell | Btbd11 | 10 | 1.93E-77 | 0.70214474 | 0.622 | 0.23 | 3.87E-74 |
| Endothelial Cell | Kcnt2 | 10 | 4.01E-77 | 0.40490157 | 0.821 | 0.353 | 8.01E-74 |
| Endothelial Cell | AABR070546 | 10 | 7.78E-77 | 0.59390513 | 0.607 | 0.22 | 1.56E-73 |
| Endothelial Cell | Colec12 | 10 | 1.23E-76 | 0.5290384 | 0.721 | 0.284 | 2.45E-73 |
| Endothelial Cell | Adamts15 | 10 | 1.43E-76 | 0.39177281 | 0.707 | 0.251 | 2.85E-73 |
| Endothelial Cell | Tgfb3 | 10 | 2.08E-76 | 0.48025346 | 0.481 | 0.188 | 4.16E-73 |
| Endothelial Cell | Marchf12 | 10 | 3.12E-76 | 0.90335606 | 0.452 | 0.166 | 6.24E-73 |
| Endothelial Cell | Nusap1 | 10 | 6.16E-76 | 1.10937637 | 0.642 | 0.209 | 1.23E-72 |
| Endothelial Cell | Sctr | 10 | 1.21E-75 | 1.09872474 | 0.487 | 0.073 | 2.41E-72 |

| | | | | | | | |
|---|---|---|---|---|---|---|---|
| Endothelial Cell | Car101 | 10 | 1.48E-75 | 0.6504644 | 0.493 | 0.122 | 2.95E-72 |
| Endothelial Cell | Hspb8 | 10 | 2.68E-75 | 0.92208546 | 0.733 | 0.267 | 5.36E-72 |
| Endothelial Cell | Ptprc1 | 10 | 2.83E-75 | 0.48032496 | 0.689 | 0.268 | 5.67E-72 |
| Endothelial Cell | Bgn | 10 | 7.48E-75 | 1.30078655 | 0.698 | 0.274 | 1.50E-71 |
| Endothelial Cell | Irs1 | 10 | 3.56E-74 | 1.34936197 | 0.628 | 0.249 | 7.12E-71 |
| Endothelial Cell | Iqub | 10 | 5.66E-74 | 1.02685579 | 0.672 | 0.284 | 1.13E-70 |
| Endothelial Cell | Mrc1 | 10 | 6.94E-74 | 0.27631077 | 0.645 | 0.265 | 1.39E-70 |
| Endothelial Cell | AABR070347 | 10 | 7.59E-74 | 0.6075167 | 0.748 | 0.301 | 1.52E-70 |
| Endothelial Cell | AABR070010 | 10 | 2.82E-73 | 0.3991452 | 0.795 | 0.273 | 5.63E-70 |
| Endothelial Cell | LOC1083526 | 10 | 6.41E-73 | 0.68553789 | 0.848 | 0.37 | 1.28E-69 |
| Endothelial Cell | Adra1b | 10 | 1.51E-72 | 0.58578377 | 0.625 | 0.285 | 3.02E-69 |
| Endothelial Cell | Lbp | 10 | 2.29E-72 | 1.23877938 | 0.595 | 0.276 | 4.59E-69 |
| Endothelial Cell | Gbp1 | 10 | 2.83E-72 | 0.53420007 | 0.575 | 0.251 | 5.65E-69 |
| Endothelial Cell | Clybl | 10 | 4.65E-72 | 0.32225015 | 0.686 | 0.268 | 9.30E-69 |
| Endothelial Cell | Ppm1e | 10 | 9.08E-72 | 0.9624894 | 0.499 | 0.219 | 1.82E-68 |
| Endothelial Cell | Enpp2 | 10 | 1.11E-71 | 1.44242972 | 0.449 | 0.12 | 2.23E-68 |
| Endothelial Cell | Dclk11 | 10 | 1.13E-71 | 0.74718117 | 0.551 | 0.227 | 2.26E-68 |
| Endothelial Cell | Srpx | 10 | 1.28E-71 | 1.19191175 | 0.991 | 0.539 | 2.55E-68 |
| Endothelial Cell | Htra3 | 10 | 5.88E-71 | 0.95049421 | 0.745 | 0.312 | 1.18E-67 |
| Endothelial Cell | Nrg4 | 10 | 7.19E-71 | 2.7299645 | 0.413 | 0.04 | 1.44E-67 |
| Endothelial Cell | Fam163a | 10 | 8.45E-71 | 0.58330321 | 0.745 | 0.319 | 1.69E-67 |
| Endothelial Cell | AABR070428 | 10 | 1.19E-70 | 0.52748244 | 0.657 | 0.314 | 2.37E-67 |
| Endothelial Cell | Adgrd11 | 10 | 2.20E-70 | 1.25231816 | 0.63 | 0.234 | 4.40E-67 |
| Endothelial Cell | Plcxd31 | 10 | 2.22E-70 | 1.41513355 | 0.566 | 0.249 | 4.44E-67 |
| Endothelial Cell | Sfxn5 | 10 | 2.97E-70 | 1.12107257 | 0.545 | 0.247 | 5.94E-67 |
| Endothelial Cell | Adamts2 | 10 | 3.33E-70 | 1.21872938 | 0.598 | 0.263 | 6.66E-67 |
| Endothelial Cell | Mmp2 | 10 | 4.20E-70 | 0.70383202 | 0.537 | 0.22 | 8.39E-67 |
| Endothelial Cell | Emid1 | 10 | 4.68E-70 | 1.58005028 | 0.607 | 0.263 | 9.35E-67 |
| Endothelial Cell | Igf1 | 10 | 5.52E-70 | 0.93337962 | 0.686 | 0.284 | 1.10E-66 |
| Endothelial Cell | Hspb6 | 10 | 6.23E-70 | 0.74844336 | 0.528 | 0.221 | 1.25E-66 |
| Endothelial Cell | Pstpip2 | 10 | 1.07E-69 | 1.10817441 | 0.616 | 0.282 | 2.14E-66 |

| | | | | | | | |
|---|---|---|---|---|---|---|---|
| Endothelial Cell | Reg3g | 10 | 1.11E-69 | 0.4472366 | 0.466 | 0.128 | 2.23E-66 |
| Endothelial Cell | Veph1 | 10 | 2.41E-69 | 0.86166952 | 0.68 | 0.291 | 4.81E-66 |
| Endothelial Cell | Slc16a10 | 10 | 2.77E-69 | 1.95220741 | 0.525 | 0.24 | 5.54E-66 |
| Endothelial Cell | S100a41 | 10 | 3.54E-69 | 1.04917256 | 0.516 | 0.219 | 7.09E-66 |
| Endothelial Cell | Mnda1 | 10 | 6.29E-69 | 1.1535042 | 0.704 | 0.303 | 1.26E-65 |
| Endothelial Cell | Myh111 | 10 | 7.22E-69 | 0.60556202 | 0.466 | 0.184 | 1.44E-65 |
| Endothelial Cell | RGD1563354 | 10 | 9.53E-69 | 0.38227178 | 0.478 | 0.156 | 1.91E-65 |
| Endothelial Cell | Tpt1 | 10 | 1.03E-68 | 0.80146489 | 0.771 | 0.304 | 2.06E-65 |
| Endothelial Cell | Adra1a | 10 | 1.26E-68 | 0.43249114 | 0.522 | 0.237 | 2.53E-65 |
| Endothelial Cell | Klhl41 | 10 | 1.75E-68 | 0.38772237 | 0.883 | 0.365 | 3.49E-65 |
| Endothelial Cell | Bcat11 | 10 | 4.10E-68 | 0.35937531 | 0.507 | 0.222 | 8.21E-65 |
| Endothelial Cell | Ripor21 | 10 | 4.45E-68 | 0.93175779 | 0.701 | 0.331 | 8.89E-65 |
| Endothelial Cell | Mertk | 10 | 4.91E-68 | 0.40593649 | 0.718 | 0.301 | 9.81E-65 |
| Endothelial Cell | Aldh1a21 | 10 | 1.05E-67 | 1.27773489 | 0.677 | 0.293 | 2.10E-64 |
| Endothelial Cell | Sgcg1 | 10 | 1.56E-67 | 0.63953524 | 0.666 | 0.272 | 3.12E-64 |
| Endothelial Cell | Atp1b1 | 10 | 2.29E-67 | 1.28564186 | 0.581 | 0.252 | 4.57E-64 |
| Endothelial Cell | Map3k7cl1 | 10 | 2.63E-67 | 0.44412913 | 0.724 | 0.312 | 5.25E-64 |
| Endothelial Cell | Mitf | 10 | 2.88E-67 | 0.52733416 | 0.636 | 0.261 | 5.76E-64 |
| Endothelial Cell | Arhgap15 | 10 | 4.90E-67 | 0.78203168 | 0.818 | 0.413 | 9.81E-64 |
| Endothelial Cell | Il16 | 10 | 1.52E-66 | 0.57046704 | 0.455 | 0.198 | 3.05E-63 |
| Endothelial Cell | Rerg | 10 | 1.29E-65 | 0.39171363 | 0.977 | 0.425 | 2.57E-62 |
| Endothelial Cell | Slit3 | 10 | 1.81E-65 | 0.69634316 | 0.625 | 0.277 | 3.62E-62 |
| Endothelial Cell | Cacna1e | 10 | 1.90E-65 | 1.78361848 | 0.519 | 0.171 | 3.80E-62 |
| Endothelial Cell | AABR070179 | 10 | 5.03E-65 | 0.62866213 | 0.692 | 0.228 | 1.01E-61 |
| Endothelial Cell | Lrrc2 | 10 | 8.70E-65 | 1.08774599 | 0.633 | 0.298 | 1.74E-61 |
| Endothelial Cell | Sdk2 | 10 | 9.43E-65 | 0.88028283 | 0.548 | 0.22 | 1.89E-61 |
| Endothelial Cell | Acsl11 | 10 | 5.43E-64 | 0.63603033 | 0.765 | 0.333 | 1.09E-60 |
| Endothelial Cell | Adcy5 | 10 | 7.16E-64 | 0.34454472 | 0.534 | 0.24 | 1.43E-60 |
| Endothelial Cell | Fhl2 | 10 | 7.34E-64 | 0.83736803 | 0.642 | 0.279 | 1.47E-60 |
| Endothelial Cell | Myom1 | 10 | 1.57E-63 | 0.81130754 | 0.683 | 0.322 | 3.14E-60 |
| Endothelial Cell | Ptpro | 10 | 1.94E-62 | 1.66137198 | 0.633 | 0.256 | 3.88E-59 |

| | | | | | | | |
|---|---|---|---|---|---|---|---|
| Endothelial Cell | AABR070070 | 10 | 3.00E-62 | 0.66104433 | 0.578 | 0.181 | 6.00E-59 |
| Endothelial Cell | Pfkfb2 | 10 | 1.23E-61 | 0.40791498 | 0.636 | 0.295 | 2.46E-58 |
| Endothelial Cell | Sh3kbp1 | 10 | 3.69E-61 | 0.70333638 | 0.9 | 0.445 | 7.39E-58 |
| Endothelial Cell | Kcng21 | 10 | 5.23E-61 | 0.62365146 | 0.768 | 0.305 | 1.05E-57 |
| Endothelial Cell | Slc16a1 | 10 | 7.22E-61 | 0.42544429 | 0.765 | 0.337 | 1.44E-57 |
| Endothelial Cell | Fam227b1 | 10 | 2.65E-60 | 0.97302748 | 0.695 | 0.309 | 5.29E-57 |
| Endothelial Cell | Gpcpd11 | 10 | 2.97E-60 | 0.26963745 | 0.689 | 0.313 | 5.93E-57 |
| Endothelial Cell | Tesc | 10 | 4.94E-60 | 0.49027339 | 0.49 | 0.237 | 9.87E-57 |
| Endothelial Cell | Cst3 | 10 | 5.05E-60 | 0.44745345 | 0.76 | 0.352 | 1.01E-56 |
| Endothelial Cell | Apba2 | 10 | 1.47E-59 | 1.93833982 | 0.472 | 0.11 | 2.94E-56 |
| Endothelial Cell | Scara51 | 10 | 2.61E-59 | 1.37143631 | 0.654 | 0.275 | 5.21E-56 |
| Endothelial Cell | Popdc2 | 10 | 4.47E-59 | 0.90482819 | 0.484 | 0.23 | 8.95E-56 |
| Endothelial Cell | Lpl | 10 | 7.54E-59 | 1.20123354 | 0.959 | 0.522 | 1.51E-55 |
| Endothelial Cell | LOC1025530 | 10 | 1.09E-58 | 0.656839 | 0.44 | 0.167 | 2.18E-55 |
| Endothelial Cell | Myo10 | 10 | 1.75E-58 | 1.31407981 | 0.971 | 0.687 | 3.51E-55 |
| Endothelial Cell | Tgfbi | 10 | 2.66E-58 | 1.49751378 | 0.642 | 0.269 | 5.31E-55 |
| Endothelial Cell | Agbl1 | 10 | 2.78E-58 | 0.89266768 | 0.557 | 0.257 | 5.56E-55 |
| Endothelial Cell | Ldhb | 10 | 4.63E-58 | 0.37484224 | 0.862 | 0.429 | 9.25E-55 |
| Endothelial Cell | Atp5mc3 | 10 | 1.20E-57 | 0.61789923 | 0.786 | 0.367 | 2.40E-54 |
| Endothelial Cell | Kif18b1 | 10 | 1.22E-57 | 1.07963095 | 0.543 | 0.156 | 2.44E-54 |
| Endothelial Cell | Itga4 | 10 | 2.76E-57 | 0.42711612 | 0.628 | 0.249 | 5.52E-54 |
| Endothelial Cell | Gpnmb | 10 | 4.53E-57 | 0.46713091 | 0.689 | 0.33 | 9.06E-54 |
| Endothelial Cell | Rnf207 | 10 | 2.61E-56 | 0.27616084 | 0.516 | 0.243 | 5.22E-53 |
| Endothelial Cell | Slc16a21 | 10 | 3.55E-56 | 0.3719782 | 0.636 | 0.292 | 7.10E-53 |
| Endothelial Cell | Dkk3 | 10 | 1.82E-55 | 0.55045662 | 0.607 | 0.306 | 3.64E-52 |
| Endothelial Cell | Ryr31 | 10 | 1.93E-55 | 0.61740729 | 0.595 | 0.3 | 3.86E-52 |
| Endothelial Cell | Serping11 | 10 | 2.18E-55 | 1.31062253 | 0.522 | 0.248 | 4.35E-52 |
| Endothelial Cell | Hk2 | 10 | 3.80E-55 | 0.5067243 | 0.698 | 0.332 | 7.60E-52 |
| Endothelial Cell | Epb41l4b1 | 10 | 5.14E-55 | 0.77131444 | 0.557 | 0.277 | 1.03E-51 |
| Endothelial Cell | Slc1a71 | 10 | 7.66E-55 | 1.79942764 | 0.531 | 0.244 | 1.53E-51 |
| Endothelial Cell | Mapt | 10 | 6.68E-54 | 0.50099555 | 0.563 | 0.269 | 1.34E-50 |

| | | | | | | | |
|---|---|---|---|---|---|---|---|
| Endothelial Cell | Myoz2 | 10 | 8.80E-54 | 0.92864258 | 0.651 | 0.344 | 1.76E-50 |
| Endothelial Cell | Robo11 | 10 | 8.81E-54 | 0.41144256 | 0.663 | 0.313 | 1.76E-50 |
| Endothelial Cell | Mmrn11 | 10 | 4.54E-53 | 0.90083111 | 0.531 | 0.213 | 9.08E-50 |
| Endothelial Cell | Pappa21 | 10 | 7.32E-53 | 2.48795705 | 0.411 | 0.137 | 1.46E-49 |
| Endothelial Cell | Smyd1 | 10 | 7.06E-52 | 1.0117208 | 0.548 | 0.271 | 1.41E-48 |
| Endothelial Cell | Npr31 | 10 | 7.79E-52 | 0.71453253 | 0.519 | 0.213 | 1.56E-48 |
| Endothelial Cell | Ppp2r3a | 10 | 1.57E-51 | 0.40873591 | 0.862 | 0.402 | 3.15E-48 |
| Endothelial Cell | Ntrk3 | 10 | 5.07E-51 | 1.28346603 | 0.607 | 0.238 | 1.01E-47 |
| Endothelial Cell | Lrrc7 | 10 | 7.13E-51 | 0.96007631 | 0.584 | 0.304 | 1.43E-47 |
| Endothelial Cell | Cnnm2 | 10 | 8.10E-51 | 0.60056559 | 0.584 | 0.318 | 1.62E-47 |
| Endothelial Cell | Timp2 | 10 | 1.40E-50 | 0.66279868 | 0.56 | 0.271 | 2.80E-47 |
| Endothelial Cell | Synpo2 | 10 | 1.95E-50 | 0.60067028 | 0.724 | 0.358 | 3.90E-47 |
| Endothelial Cell | Grb141 | 10 | 2.29E-50 | 0.48611738 | 0.601 | 0.315 | 4.58E-47 |
| Endothelial Cell | Slco2b1 | 10 | 2.71E-50 | 0.90463287 | 0.871 | 0.469 | 5.42E-47 |
| Endothelial Cell | Ca4 | 10 | 7.46E-50 | 0.97197483 | 0.683 | 0.354 | 1.49E-46 |
| Endothelial Cell | Atp5f1b | 10 | 6.06E-49 | 0.60457042 | 0.933 | 0.448 | 1.21E-45 |
| Endothelial Cell | Syt1 | 10 | 6.77E-49 | 0.91924217 | 0.584 | 0.237 | 1.35E-45 |
| Endothelial Cell | Cdon | 10 | 7.53E-49 | 0.31710479 | 0.537 | 0.287 | 1.51E-45 |
| Endothelial Cell | Epha4 | 10 | 2.09E-48 | 0.76450344 | 0.95 | 0.486 | 4.19E-45 |
| Endothelial Cell | Shroom3 | 10 | 3.24E-48 | 0.76511531 | 0.78 | 0.411 | 6.48E-45 |
| Endothelial Cell | Gja1 | 10 | 4.24E-48 | 0.64367679 | 0.812 | 0.391 | 8.48E-45 |
| Endothelial Cell | Frmd51 | 10 | 5.23E-48 | 0.53136867 | 0.636 | 0.375 | 1.05E-44 |
| Endothelial Cell | Flvcr21 | 10 | 7.23E-48 | 0.64025645 | 0.613 | 0.321 | 1.45E-44 |
| Endothelial Cell | Zfp385d | 10 | 1.23E-47 | 0.68159141 | 0.604 | 0.326 | 2.45E-44 |
| Endothelial Cell | Fndc1 | 10 | 2.66E-47 | 0.9411405 | 0.557 | 0.3 | 5.33E-44 |
| Endothelial Cell | Zbtb16 | 10 | 4.02E-47 | 1.16634775 | 0.839 | 0.463 | 8.04E-44 |
| Endothelial Cell | Asb181 | 10 | 9.06E-47 | 0.70194747 | 0.642 | 0.341 | 1.81E-43 |
| Endothelial Cell | Dpp41 | 10 | 1.08E-46 | 0.47049421 | 0.809 | 0.423 | 2.16E-43 |
| Endothelial Cell | Gucy1a21 | 10 | 5.43E-46 | 0.96045721 | 0.651 | 0.353 | 1.09E-42 |
| Endothelial Cell | Adap2 | 10 | 9.45E-46 | 0.69580331 | 0.619 | 0.304 | 1.89E-42 |
| Endothelial Cell | Itgb2 | 10 | 1.07E-45 | 0.89241964 | 0.581 | 0.203 | 2.15E-42 |

| | | | | | | | |
|---|---|---|---|---|---|---|---|
| Endothelial Cell | Pdlim3 | 10 | 3.44E-45 | 0.4849555 | 0.648 | 0.347 | 6.88E-42 |
| Endothelial Cell | Ahr1 | 10 | 5.50E-45 | 0.3383245 | 0.801 | 0.399 | 1.10E-41 |
| Endothelial Cell | Mylk3 | 10 | 1.69E-44 | 0.50117618 | 0.569 | 0.312 | 3.38E-41 |
| Endothelial Cell | Lurap1l1 | 10 | 3.87E-44 | 0.64152558 | 0.677 | 0.343 | 7.73E-41 |
| Endothelial Cell | Esrrg | 10 | 4.54E-44 | 0.28845996 | 0.686 | 0.369 | 9.09E-41 |
| Endothelial Cell | Ninj21 | 10 | 7.70E-44 | 0.52916229 | 0.701 | 0.387 | 1.54E-40 |
| Endothelial Cell | Ncald | 10 | 8.47E-44 | 0.87394935 | 0.921 | 0.489 | 1.69E-40 |
| Endothelial Cell | Tango2 | 10 | 9.76E-44 | 0.38271304 | 0.724 | 0.39 | 1.95E-40 |
| Endothelial Cell | Rxfp1 | 10 | 1.85E-43 | 0.32006939 | 0.566 | 0.269 | 3.70E-40 |
| Endothelial Cell | Nox41 | 10 | 2.89E-43 | 0.38113064 | 0.862 | 0.432 | 5.79E-40 |
| Endothelial Cell | Mfsd4a1 | 10 | 1.99E-42 | 0.87963984 | 0.628 | 0.314 | 3.97E-39 |
| Endothelial Cell | Dusp27 | 10 | 2.40E-42 | 0.60460775 | 0.628 | 0.292 | 4.80E-39 |
| Endothelial Cell | Slc39a8 | 10 | 3.58E-42 | 0.43516964 | 0.587 | 0.272 | 7.15E-39 |
| Endothelial Cell | Trak21 | 10 | 4.09E-42 | 0.63758376 | 0.906 | 0.465 | 8.17E-39 |
| Endothelial Cell | Nlgn1 | 10 | 6.36E-42 | 0.29077778 | 0.443 | 0.069 | 1.27E-38 |
| Endothelial Cell | Sorcs1 | 10 | 8.45E-42 | 0.46217636 | 0.493 | 0.146 | 1.69E-38 |
| Endothelial Cell | Ajap1 | 10 | 2.91E-41 | 1.17113375 | 0.507 | 0.148 | 5.82E-38 |
| Endothelial Cell | Unc13b | 10 | 1.09E-40 | 0.64453242 | 0.994 | 0.601 | 2.19E-37 |
| Endothelial Cell | Ccn3 | 10 | 1.36E-40 | 1.55449704 | 0.566 | 0.312 | 2.72E-37 |
| Endothelial Cell | Psap1 | 10 | 4.82E-40 | 0.48318959 | 0.633 | 0.354 | 9.64E-37 |
| Endothelial Cell | Rilpl1 | 10 | 6.72E-40 | 0.49559314 | 0.674 | 0.363 | 1.34E-36 |
| Endothelial Cell | Efnb2 | 10 | 1.02E-39 | 0.98215836 | 0.982 | 0.646 | 2.04E-36 |
| Endothelial Cell | Slc6a61 | 10 | 6.79E-39 | 0.87496153 | 0.956 | 0.596 | 1.36E-35 |
| Endothelial Cell | Colgalt2 | 10 | 2.57E-38 | 0.5911782 | 0.9 | 0.509 | 5.15E-35 |
| Endothelial Cell | Slc30a31 | 10 | 2.83E-38 | 1.27906994 | 0.537 | 0.173 | 5.65E-35 |
| Endothelial Cell | Prom1 | 10 | 1.22E-37 | 0.98689887 | 0.897 | 0.533 | 2.43E-34 |
| Endothelial Cell | Camk41 | 10 | 1.62E-37 | 1.79805203 | 0.54 | 0.193 | 3.23E-34 |
| Endothelial Cell | Hvcn11 | 10 | 1.66E-37 | 0.76705492 | 0.557 | 0.276 | 3.31E-34 |
| Endothelial Cell | Wipf3 | 10 | 5.28E-37 | 0.35893138 | 0.938 | 0.512 | 1.06E-33 |
| Endothelial Cell | Cxcl122 | 10 | 3.97E-36 | 0.96254813 | 0.707 | 0.454 | 7.94E-33 |
| Endothelial Cell | Crip1 | 10 | 1.04E-35 | 0.53568633 | 0.701 | 0.425 | 2.08E-32 |

| | | | | | | | |
|---|---|---|---|---|---|---|---|
| Endothelial Cell | Tnfaip6 | 10 | 3.49E-35 | 1.62106115 | 0.633 | 0.291 | 6.97E-32 |
| Endothelial Cell | Clic5 | 10 | 4.27E-35 | 0.92086351 | 0.947 | 0.645 | 8.54E-32 |
| Endothelial Cell | Rmdn1 | 10 | 4.60E-35 | 0.26242312 | 0.692 | 0.401 | 9.20E-32 |
| Endothelial Cell | Slc26a101 | 10 | 1.18E-34 | 0.98016582 | 0.947 | 0.672 | 2.36E-31 |
| Endothelial Cell | Kcnn3 | 10 | 1.82E-34 | 0.26362501 | 0.639 | 0.381 | 3.63E-31 |
| Endothelial Cell | Cacnb2 | 10 | 3.17E-34 | 0.60032114 | 0.783 | 0.451 | 6.34E-31 |
| Endothelial Cell | Tmem116 | 10 | 6.69E-34 | 1.02634656 | 0.572 | 0.215 | 1.34E-30 |
| Endothelial Cell | Ano51 | 10 | 1.63E-33 | 0.64929541 | 0.592 | 0.239 | 3.26E-30 |
| Endothelial Cell | Ighm1 | 10 | 1.68E-33 | 0.8986989 | 0.463 | 0.145 | 3.36E-30 |
| Endothelial Cell | Adgre1 | 10 | 1.20E-31 | 1.21112273 | 0.437 | 0.138 | 2.41E-28 |
| Endothelial Cell | Kcnj51 | 10 | 4.69E-30 | 0.60150041 | 0.622 | 0.243 | 9.39E-27 |
| Endothelial Cell | Ptprv | 10 | 6.25E-30 | 1.14188539 | 0.411 | 0.116 | 1.25E-26 |
| Endothelial Cell | Lims2 | 10 | 8.00E-30 | 0.87979699 | 0.979 | 0.68 | 1.60E-26 |
| Endothelial Cell | Unc5b1 | 10 | 3.41E-29 | 0.66067633 | 0.745 | 0.472 | 6.82E-26 |
| Endothelial Cell | AABR070076 | 10 | 1.30E-28 | 0.76758669 | 0.959 | 0.628 | 2.60E-25 |
| Endothelial Cell | Fhod31 | 10 | 2.04E-28 | 0.64809254 | 0.827 | 0.527 | 4.08E-25 |
| Endothelial Cell | Arhgap21 | 10 | 7.42E-28 | 0.30038039 | 0.757 | 0.45 | 1.48E-24 |
| Endothelial Cell | Gcnt2 | 10 | 9.68E-28 | 0.29639766 | 0.806 | 0.461 | 1.94E-24 |
| Endothelial Cell | Lmo7 | 10 | 1.42E-26 | 0.96342908 | 0.968 | 0.713 | 2.85E-23 |
| Endothelial Cell | Cmtm8 | 10 | 3.84E-26 | 0.80894418 | 0.889 | 0.585 | 7.69E-23 |
| Endothelial Cell | Necab11 | 10 | 1.13E-25 | 1.78497326 | 0.484 | 0.223 | 2.26E-22 |
| Endothelial Cell | Pcdh17 | 10 | 9.93E-25 | 0.2969574 | 0.789 | 0.512 | 1.99E-21 |
| Endothelial Cell | Fermt3 | 10 | 4.90E-23 | 0.71744688 | 0.466 | 0.202 | 9.79E-20 |
| Endothelial Cell | Dmd | 10 | 5.72E-23 | 0.69039324 | 0.927 | 0.677 | 1.14E-19 |
| Endothelial Cell | Akr1c151 | 10 | 1.73E-22 | 0.68997926 | 0.953 | 0.644 | 3.45E-19 |
| Endothelial Cell | St6galnac31 | 10 | 4.81E-22 | 0.40235913 | 0.941 | 0.613 | 9.63E-19 |
| Endothelial Cell | Clec7a | 10 | 1.11E-21 | 0.6732723 | 0.499 | 0.233 | 2.22E-18 |
| Endothelial Cell | Dipk2b | 10 | 6.51E-21 | 0.42795144 | 0.941 | 0.594 | 1.30E-17 |
| Endothelial Cell | Prss12 | 10 | 7.92E-19 | 1.25804506 | 0.428 | 0.141 | 1.58E-15 |
| Endothelial Cell | Matk | 10 | 2.96E-16 | 0.56984732 | 0.334 | 0.082 | 5.92E-13 |
| Endothelial Cell | Adamts9 | 10 | 1.77E-15 | 0.3468942 | 0.933 | 0.623 | 3.54E-12 |

| | | | | | | | |
|---|---|---|---|---|---|---|---|
| Endothelial Cell | Palmd | 10 | 2.13E-15 | 0.53208418 | 0.927 | 0.625 | 4.26E-12 |
| Endothelial Cell | Plcb1 | 10 | 6.00E-12 | 0.28581664 | 0.988 | 0.714 | 1.20E-08 |
| Endothelial Cell | Ccnb1 | 11 | 3.63E-41 | 1.23603554 | 0.272 | 0.01 | 7.25E-38 |
| Endothelial Cell | Cdo1 | 11 | 7.20E-40 | 1.34220551 | 0.33 | 0.073 | 1.44E-36 |
| Endothelial Cell | Comp | 11 | 4.33E-34 | 3.26303578 | 0.32 | 0.049 | 8.66E-31 |
| Endothelial Cell | Rundc3a | 11 | 5.59E-34 | 3.02658437 | 0.33 | 0.074 | 1.12E-30 |
| Endothelial Cell | Exoc3l21 | 11 | 1.63E-31 | 2.88635933 | 0.612 | 0.298 | 3.25E-28 |
| Endothelial Cell | Slco2a12 | 11 | 8.68E-29 | 1.81517278 | 0.619 | 0.292 | 1.74E-25 |
| Endothelial Cell | Adamts6 | 11 | 7.33E-26 | 1.66641674 | 0.711 | 0.454 | 1.47E-22 |
| Endothelial Cell | Pak7 | 11 | 4.01E-24 | 1.42612204 | 0.357 | 0.056 | 8.03E-21 |
| Endothelial Cell | Ldb23 | 11 | 7.65E-21 | 1.46819657 | 0.656 | 0.394 | 1.53E-17 |
| Endothelial Cell | Slc47a12 | 11 | 1.07E-14 | 1.75584062 | 0.415 | 0.137 | 2.13E-11 |
| Endothelial Cell | Pdgfc1 | 11 | 1.99E-10 | 1.54293079 | 0.473 | 0.21 | 3.97E-07 |
| Cardiomyocyte | Cd27 | 0 | 4.83E-225 | 1.29569285 | 0.5 | 0.075 | 9.66E-222 |
| Cardiomyocyte | Hao1 | 0 | 1.69E-216 | 1.88950182 | 0.427 | 0.046 | 3.38E-213 |
| Cardiomyocyte | AABR070131 | 0 | 4.87E-143 | 1.91917152 | 0.509 | 0.136 | 9.74E-140 |
| Cardiomyocyte | Coch | 0 | 6.30E-128 | 0.70355691 | 0.343 | 0.061 | 1.26E-124 |
| Cardiomyocyte | LOC499544 | 0 | 4.52E-125 | 1.41370611 | 0.339 | 0.031 | 9.04E-122 |
| Cardiomyocyte | Lsamp | 0 | 3.08E-106 | 1.16803761 | 0.933 | 0.619 | 6.16E-103 |
| Cardiomyocyte | Ptgds | 0 | 3.60E-105 | 0.77353447 | 0.459 | 0.133 | 7.20E-102 |
| Cardiomyocyte | Cd247 | 0 | 4.17E-96 | 0.57650607 | 0.368 | 0.08 | 8.34E-93 |
| Cardiomyocyte | Il13ra2 | 0 | 1.22E-91 | 0.3804388 | 0.397 | 0.087 | 2.44E-88 |
| Cardiomyocyte | Fstl5 | 0 | 2.26E-89 | 1.09000078 | 0.447 | 0.132 | 4.53E-86 |
| Cardiomyocyte | LOC1009098 | 0 | 4.47E-89 | 0.80528121 | 0.449 | 0.163 | 8.94E-86 |
| Cardiomyocyte | E2f7 | 0 | 1.99E-79 | 0.42761745 | 0.468 | 0.107 | 3.98E-76 |
| Cardiomyocyte | AABR070269 | 0 | 5.83E-79 | 2.1812972 | 0.38 | 0.116 | 1.17E-75 |
| Cardiomyocyte | Gpcpd1 | 0 | 9.39E-73 | 0.8584723 | 0.964 | 0.709 | 1.88E-69 |
| Cardiomyocyte | AC130940.1 | 0 | 1.45E-69 | 1.16852909 | 0.76 | 0.404 | 2.90E-66 |
| Cardiomyocyte | Lrrtm3 | 0 | 1.27E-67 | 1.147566 | 0.767 | 0.514 | 2.54E-64 |
| Cardiomyocyte | Scn9a | 0 | 7.84E-67 | 0.75270677 | 0.447 | 0.146 | 1.57E-63 |
| Cardiomyocyte | Hpd | 0 | 8.63E-64 | 1.13402392 | 0.637 | 0.293 | 1.73E-60 |

| | | | | | | | |
|---|---|---|---|---|---|---|---|
| Cardiomyocyte | Corin | 0 | 1.11E-60 | 0.93238574 | 0.901 | 0.635 | 2.21E-57 |
| Cardiomyocyte | Vsnl1 | 0 | 2.38E-55 | 1.1589891 | 0.559 | 0.225 | 4.75E-52 |
| Cardiomyocyte | Fam78b | 0 | 7.78E-52 | 0.87535423 | 0.815 | 0.51 | 1.56E-48 |
| Cardiomyocyte | Cdh8 | 0 | 1.95E-51 | 0.58046764 | 0.421 | 0.146 | 3.91E-48 |
| Cardiomyocyte | AABR070134 | 0 | 3.46E-51 | 1.60917034 | 0.503 | 0.206 | 6.91E-48 |
| Cardiomyocyte | Slc47a1 | 0 | 1.18E-50 | 1.15969917 | 0.513 | 0.241 | 2.36E-47 |
| Cardiomyocyte | Dhrs7c | 0 | 6.46E-50 | 0.90474541 | 0.683 | 0.377 | 1.29E-46 |
| Cardiomyocyte | Ppp1r3a | 0 | 9.33E-49 | 0.9369242 | 0.743 | 0.486 | 1.87E-45 |
| Cardiomyocyte | Hpgd | 0 | 1.00E-48 | 0.64964389 | 0.38 | 0.071 | 2.00E-45 |
| Cardiomyocyte | LOC1036933 | 0 | 6.23E-46 | 0.45024674 | 0.54 | 0.249 | 1.25E-42 |
| Cardiomyocyte | Gpam | 0 | 3.27E-45 | 1.00081879 | 0.636 | 0.362 | 6.54E-42 |
| Cardiomyocyte | Arhgef39 | 0 | 2.12E-44 | 0.3736435 | 0.437 | 0.168 | 4.25E-41 |
| Cardiomyocyte | AABR070030 | 0 | 6.20E-44 | 0.3701839 | 0.596 | 0.335 | 1.24E-40 |
| Cardiomyocyte | Grm1 | 0 | 2.18E-43 | 0.74231005 | 0.468 | 0.183 | 4.36E-40 |
| Cardiomyocyte | Mettl24 | 0 | 1.68E-41 | 0.63637589 | 0.551 | 0.249 | 3.35E-38 |
| Cardiomyocyte | AABR070127 | 0 | 4.14E-40 | 0.61603905 | 0.405 | 0.138 | 8.27E-37 |
| Cardiomyocyte | Sh3rf2 | 0 | 1.43E-39 | 0.48519956 | 0.544 | 0.235 | 2.87E-36 |
| Cardiomyocyte | Tnfrsf19 | 0 | 4.59E-39 | 0.9228716 | 0.497 | 0.205 | 9.18E-36 |
| Cardiomyocyte | Hecw2 | 0 | 1.34E-31 | 0.56907197 | 0.828 | 0.554 | 2.69E-28 |
| Cardiomyocyte | RGD1564053 | 0 | 1.03E-30 | 0.40508905 | 0.54 | 0.269 | 2.06E-27 |
| Cardiomyocyte | AABR070515 | 0 | 6.82E-25 | 0.39670454 | 0.652 | 0.377 | 1.36E-21 |
| Cardiomyocyte | Tmem150c | 0 | 2.12E-23 | 0.94490405 | 0.456 | 0.197 | 4.23E-20 |
| Cardiomyocyte | AABR070349 | 1 | 5.92E-116 | 0.26929143 | 0.025 | 0.304 | 1.18E-112 |
| Cardiomyocyte | Nr4a3 | 1 | 3.54E-114 | 0.26941424 | 0.058 | 0.529 | 7.08E-111 |
| Cardiomyocyte | Nusap1 | 1 | 1.71E-109 | 0.29424828 | 0.047 | 0.375 | 3.43E-106 |
| Cardiomyocyte | Myo5b | 1 | 4.10E-109 | 0.36823975 | 0.046 | 0.297 | 8.21E-106 |
| Cardiomyocyte | Col20a1 | 1 | 6.63E-109 | 0.30211609 | 0.055 | 0.324 | 1.33E-105 |
| Cardiomyocyte | Col13a1 | 1 | 2.86E-106 | 0.26961984 | 0.058 | 0.373 | 5.71E-103 |
| Cardiomyocyte | Myom3 | 1 | 1.22E-105 | 0.60057889 | 0.058 | 0.447 | 2.45E-102 |
| Cardiomyocyte | Fgf7 | 1 | 1.38E-102 | 0.8573973 | 0.056 | 0.418 | 2.76E-99 |
| Cardiomyocyte | Col19a1 | 1 | 5.42E-100 | 0.33684335 | 0.066 | 0.36 | 1.08E-96 |

| | | | | | | | |
|---|---|---|---|---|---|---|---|
| Cardiomyocyte | Kif17 | 1 | 2.72E-99 | 0.49992466 | 0.032 | 0.334 | 5.44E-96 |
| Cardiomyocyte | Fam189a1 | 1 | 3.62E-98 | 0.50976813 | 0.023 | 0.28 | 7.24E-95 |
| Cardiomyocyte | Ankrd55 | 1 | 5.46E-98 | 0.65471662 | 0.044 | 0.341 | 1.09E-94 |
| Cardiomyocyte | Cdo1 | 1 | 2.87E-97 | 0.34947396 | 0.039 | 0.346 | 5.73E-94 |
| Cardiomyocyte | Dok5 | 1 | 2.21E-95 | 0.46603379 | 0.086 | 0.375 | 4.42E-92 |
| Cardiomyocyte | Itgam | 1 | 8.10E-95 | 0.42457482 | 0.047 | 0.319 | 1.62E-91 |
| Cardiomyocyte | Hpd1 | 1 | 3.42E-93 | 0.37143089 | 0.092 | 0.386 | 6.85E-90 |
| Cardiomyocyte | Adgrb3 | 1 | 4.59E-93 | 1.30903671 | 0.032 | 0.3 | 9.17E-90 |
| Cardiomyocyte | Pak7 | 1 | 7.38E-93 | 0.51842167 | 0.047 | 0.313 | 1.48E-89 |
| Cardiomyocyte | Gpd1 | 1 | 5.14E-90 | 0.25972761 | 0.102 | 0.451 | 1.03E-86 |
| Cardiomyocyte | Neurl1 | 1 | 7.48E-88 | 0.37225255 | 0.027 | 0.321 | 1.50E-84 |
| Cardiomyocyte | Slc26a3 | 1 | 2.47E-87 | 0.34306059 | 0.043 | 0.306 | 4.94E-84 |
| Cardiomyocyte | Nkain3 | 1 | 6.34E-87 | 0.30061741 | 0.021 | 0.272 | 1.27E-83 |
| Cardiomyocyte | Myo16 | 1 | 2.02E-84 | 0.40566232 | 0.114 | 0.466 | 4.04E-81 |
| Cardiomyocyte | Gabrb2 | 1 | 9.39E-84 | 0.71387295 | 0.099 | 0.38 | 1.88E-80 |
| Cardiomyocyte | Cdh1 | 1 | 1.73E-83 | 0.65064594 | 0.029 | 0.295 | 3.46E-80 |
| Cardiomyocyte | Themis | 1 | 1.09E-75 | 0.39658739 | 0.011 | 0.291 | 2.18E-72 |
| Cardiomyocyte | Gpam1 | 1 | 3.39E-74 | 0.48855226 | 0.133 | 0.448 | 6.79E-71 |
| Cardiomyocyte | AABR070504 | 1 | 3.50E-67 | 0.45454414 | 0.153 | 0.429 | 7.00E-64 |
| Cardiomyocyte | Fbxw10 | 1 | 2.14E-64 | 0.59491153 | 0.161 | 0.518 | 4.28E-61 |
| Cardiomyocyte | Tec | 1 | 9.77E-61 | 0.25741851 | 0.192 | 0.465 | 1.95E-57 |
| Cardiomyocyte | Myo3b | 1 | 2.27E-60 | 0.44297015 | 0.166 | 0.44 | 4.54E-57 |
| Cardiomyocyte | Cobl | 1 | 6.03E-60 | 0.35908468 | 0.168 | 0.43 | 1.21E-56 |
| Cardiomyocyte | Colq | 1 | 5.87E-59 | 0.43145726 | 0.181 | 0.47 | 1.17E-55 |
| Cardiomyocyte | Kcnj5 | 1 | 1.08E-46 | 0.40506045 | 0.225 | 0.494 | 2.15E-43 |
| Cardiomyocyte | Ano5 | 1 | 5.75E-46 | 0.30711437 | 0.227 | 0.575 | 1.15E-42 |
| Cardiomyocyte | Dsg2 | 1 | 2.78E-44 | 0.25886933 | 0.244 | 0.589 | 5.55E-41 |
| Cardiomyocyte | Pstpip2 | 1 | 1.08E-43 | 0.2759806 | 0.247 | 0.54 | 2.16E-40 |
| Cardiomyocyte | Lrrc10 | 1 | 9.77E-42 | 0.32856989 | 0.247 | 0.558 | 1.95E-38 |
| Cardiomyocyte | Asb9 | 1 | 1.01E-41 | 0.25100348 | 0.256 | 0.6 | 2.01E-38 |
| Cardiomyocyte | Ankrd6 | 1 | 3.13E-39 | 0.28343474 | 0.269 | 0.584 | 6.27E-36 |

| | | | | | | | |
|---|---|---|---|---|---|---|---|
| Cardiomyocyte | Lrrc7 | 1 | 3.94E-30 | 0.55628161 | 0.279 | 0.545 | 7.88E-27 |
| Cardiomyocyte | Dlgap1 | 1 | 3.70E-25 | 0.27815533 | 0.339 | 0.623 | 7.41E-22 |
| Cardiomyocyte | Dcdc5 | 1 | 2.41E-20 | 0.77439974 | 0.307 | 0.588 | 4.81E-17 |
| Cardiomyocyte | Abcb4 | 1 | 2.91E-18 | 0.26793434 | 0.382 | 0.66 | 5.82E-15 |
| Cardiomyocyte | Sfxn5 | 1 | 3.90E-15 | 0.25331276 | 0.386 | 0.662 | 7.81E-12 |
| Cardiomyocyte | Ros1 | 1 | 4.04E-11 | 0.31989864 | 0.399 | 0.665 | 8.08E-08 |
| Cardiomyocyte | Arhgef37 | 1 | 5.14E-09 | 0.45169172 | 0.421 | 0.681 | 1.03E-05 |
| Cardiomyocyte | Ky | 1 | 1.52E-08 | 0.39452128 | 0.428 | 0.687 | 3.04E-05 |
| Cardiomyocyte | Cpe | 1 | 1.71E-06 | 0.26185699 | 0.476 | 0.728 | 0.00341702 |
| Cardiomyocyte | Usp2 | 1 | 2.17E-05 | 0.35672589 | 0.476 | 0.75 | 0.04330131 |
| Cardiomyocyte | Gpc6 | 2 | 1.59E-244 | 2.84449882 | 0.969 | 0.573 | 3.18E-241 |
| Cardiomyocyte | Zfp385d | 2 | 2.58E-206 | 3.1750526 | 0.887 | 0.287 | 5.17E-203 |
| Cardiomyocyte | Clmp | 2 | 5.08E-188 | 4.10958509 | 0.793 | 0.126 | 1.02E-184 |
| Cardiomyocyte | Fbn1 | 2 | 6.02E-179 | 2.98960693 | 0.897 | 0.489 | 1.20E-175 |
| Cardiomyocyte | C1qtnf7 | 2 | 2.88E-170 | 3.53530245 | 0.805 | 0.215 | 5.75E-167 |
| Cardiomyocyte | C7 | 2 | 1.93E-169 | 3.15330664 | 0.851 | 0.374 | 3.85E-166 |
| Cardiomyocyte | Col3a1 | 2 | 9.83E-167 | 1.90476197 | 0.957 | 0.647 | 1.97E-163 |
| Cardiomyocyte | Itgbl1 | 2 | 1.92E-166 | 2.93459221 | 0.834 | 0.438 | 3.84E-163 |
| Cardiomyocyte | AABR070707 | 2 | 7.96E-164 | 2.10460931 | 0.272 | 0.019 | 1.59E-160 |
| Cardiomyocyte | Adamts2 | 2 | 9.31E-159 | 3.26250184 | 0.79 | 0.318 | 1.86E-155 |
| Cardiomyocyte | Cntn2 | 2 | 7.87E-156 | 2.52721538 | 0.273 | 0.012 | 1.57E-152 |
| Cardiomyocyte | Ccdc80 | 2 | 2.21E-149 | 2.82241249 | 0.838 | 0.439 | 4.43E-146 |
| Cardiomyocyte | Fndc1 | 2 | 8.46E-148 | 3.21953594 | 0.778 | 0.251 | 1.69E-144 |
| Cardiomyocyte | Col8a1 | 2 | 4.28E-146 | 3.38396048 | 0.82 | 0.382 | 8.56E-143 |
| Cardiomyocyte | Antxr1 | 2 | 1.94E-142 | 3.87053812 | 0.762 | 0.218 | 3.88E-139 |
| Cardiomyocyte | Ebf2 | 2 | 6.50E-140 | 3.28459868 | 0.752 | 0.216 | 1.30E-136 |
| Cardiomyocyte | Bnc2 | 2 | 1.26E-138 | 3.54320002 | 0.765 | 0.24 | 2.51E-135 |
| Cardiomyocyte | Plxdc2 | 2 | 1.09E-136 | 2.57364488 | 0.833 | 0.352 | 2.18E-133 |
| Cardiomyocyte | Abi3bp | 2 | 4.22E-135 | 3.05951956 | 0.77 | 0.275 | 8.44E-132 |
| Cardiomyocyte | Pdgfra | 2 | 2.58E-133 | 3.4220877 | 0.752 | 0.28 | 5.15E-130 |
| Cardiomyocyte | Fstl1 | 2 | 8.74E-128 | 2.44372268 | 0.805 | 0.278 | 1.75E-124 |

| | | | | | | | |
|---|---|---|---|---|---|---|---|
| Cardiomyocyte | Oprm1 | 2 | 2.68E-126 | 1.11749628 | 0.288 | 0.032 | 5.35E-123 |
| Cardiomyocyte | Prrx1 | 2 | 7.81E-126 | 3.1999766 | 0.752 | 0.215 | 1.56E-122 |
| Cardiomyocyte | Mrc2 | 2 | 2.45E-114 | 3.95234113 | 0.675 | 0.233 | 4.90E-111 |
| Cardiomyocyte | Axl | 2 | 1.01E-108 | 3.35965099 | 0.712 | 0.227 | 2.03E-105 |
| Cardiomyocyte | Plod2 | 2 | 7.04E-105 | 2.8678681 | 0.715 | 0.272 | 1.41E-101 |
| Cardiomyocyte | Col5a2 | 2 | 2.30E-104 | 2.53347487 | 0.727 | 0.262 | 4.61E-101 |
| Cardiomyocyte | Pdgfrb | 2 | 5.16E-104 | 2.48839049 | 0.709 | 0.243 | 1.03E-100 |
| Cardiomyocyte | Dlg2 | 2 | 2.61E-103 | 3.31094788 | 0.709 | 0.281 | 5.21E-100 |
| Cardiomyocyte | Col1a1 | 2 | 6.19E-102 | 1.74693888 | 0.864 | 0.465 | 1.24E-98 |
| Cardiomyocyte | Dcn | 2 | 2.51E-99 | 2.1088528 | 0.79 | 0.374 | 5.03E-96 |
| Cardiomyocyte | Ston2 | 2 | 3.82E-99 | 3.24042792 | 0.677 | 0.217 | 7.64E-96 |
| Cardiomyocyte | Pid1 | 2 | 1.23E-96 | 2.08974549 | 0.753 | 0.375 | 2.46E-93 |
| Cardiomyocyte | Gda | 2 | 2.00E-95 | 3.06775895 | 0.664 | 0.248 | 3.99E-92 |
| Cardiomyocyte | Zeb2 | 2 | 5.23E-93 | 1.29006848 | 0.876 | 0.623 | 1.05E-89 |
| Cardiomyocyte | Gxylt2 | 2 | 8.50E-92 | 3.00274782 | 0.715 | 0.395 | 1.70E-88 |
| Cardiomyocyte | Egfr | 2 | 1.32E-91 | 3.46784607 | 0.657 | 0.331 | 2.63E-88 |
| Cardiomyocyte | Dpysl3 | 2 | 1.90E-90 | 2.78409085 | 0.685 | 0.333 | 3.80E-87 |
| Cardiomyocyte | Timp2 | 2 | 8.14E-90 | 1.78661708 | 0.803 | 0.452 | 1.63E-86 |
| Cardiomyocyte | Map1b | 2 | 1.21E-89 | 1.92772335 | 0.796 | 0.465 | 2.42E-86 |
| Cardiomyocyte | Postn | 2 | 3.61E-88 | 2.55811912 | 0.707 | 0.293 | 7.21E-85 |
| Cardiomyocyte | Tmeff2 | 2 | 1.15E-87 | 3.28200757 | 0.651 | 0.219 | 2.29E-84 |
| Cardiomyocyte | Col6a6 | 2 | 1.85E-86 | 3.90536326 | 0.629 | 0.281 | 3.69E-83 |
| Cardiomyocyte | Fbln1 | 2 | 5.44E-85 | 3.04124307 | 0.657 | 0.202 | 1.09E-81 |
| Cardiomyocyte | Clnk | 2 | 1.04E-81 | 2.15529724 | 0.316 | 0.026 | 2.09E-78 |
| Cardiomyocyte | Cpq | 2 | 1.31E-81 | 2.05122404 | 0.732 | 0.431 | 2.62E-78 |
| Cardiomyocyte | Glis3 | 2 | 2.00E-81 | 3.09077721 | 0.631 | 0.177 | 4.01E-78 |
| Cardiomyocyte | Ptch2 | 2 | 2.57E-81 | 1.26374882 | 0.366 | 0.08 | 5.13E-78 |
| Cardiomyocyte | Hmga2 | 2 | 2.65E-79 | 1.78597052 | 0.376 | 0.11 | 5.29E-76 |
| Cardiomyocyte | Cdkn2b | 2 | 5.77E-79 | 3.1936843 | 0.262 | 0.009 | 1.15E-75 |
| Cardiomyocyte | Col14a1 | 2 | 2.28E-77 | 3.1054044 | 0.624 | 0.228 | 4.57E-74 |
| Cardiomyocyte | Pde1a | 2 | 4.28E-77 | 3.06680134 | 0.651 | 0.357 | 8.55E-74 |

| | | | | | | | |
|---|---|---|---|---|---|---|---|
| Cardiomyocyte | Boc | 2 | 1.57E-76 | 1.71817724 | 0.813 | 0.51 | 3.14E-73 |
| Cardiomyocyte | Col6a2 | 2 | 3.40E-74 | 2.64785455 | 0.636 | 0.263 | 6.81E-71 |
| Cardiomyocyte | Plekha7 | 2 | 1.35E-72 | 3.05693456 | 0.613 | 0.168 | 2.70E-69 |
| Cardiomyocyte | Dclk1 | 2 | 3.12E-71 | 3.05225753 | 0.603 | 0.224 | 6.24E-68 |
| Cardiomyocyte | Chsy3 | 2 | 5.75E-70 | 3.02154802 | 0.649 | 0.353 | 1.15E-66 |
| Cardiomyocyte | Asap3 | 2 | 3.78E-69 | 2.87847835 | 0.631 | 0.304 | 7.57E-66 |
| Cardiomyocyte | Dpt | 2 | 1.87E-66 | 2.1584289 | 0.631 | 0.316 | 3.73E-63 |
| Cardiomyocyte | Adgrd1 | 2 | 1.62E-64 | 3.96618663 | 0.566 | 0.15 | 3.25E-61 |
| Cardiomyocyte | Abca8a | 2 | 1.66E-64 | 3.16046733 | 0.598 | 0.276 | 3.32E-61 |
| Cardiomyocyte | Adamtsl3 | 2 | 1.74E-63 | 3.26621759 | 0.57 | 0.273 | 3.48E-60 |
| Cardiomyocyte | Medag | 2 | 1.95E-63 | 3.42558277 | 0.579 | 0.255 | 3.90E-60 |
| Cardiomyocyte | Mfap5 | 2 | 2.04E-62 | 2.83997977 | 0.608 | 0.306 | 4.08E-59 |
| Cardiomyocyte | Itga8 | 2 | 2.01E-61 | 3.10859871 | 0.608 | 0.23 | 4.03E-58 |
| Cardiomyocyte | Robo1 | 2 | 2.27E-61 | 2.8607521 | 0.589 | 0.271 | 4.54E-58 |
| Cardiomyocyte | Il1r1 | 2 | 1.34E-60 | 1.72519245 | 0.652 | 0.393 | 2.69E-57 |
| Cardiomyocyte | Vcan | 2 | 1.57E-60 | 2.76077043 | 0.589 | 0.207 | 3.14E-57 |
| Cardiomyocyte | Mgp | 2 | 2.53E-60 | 1.36878467 | 0.805 | 0.512 | 5.07E-57 |
| Cardiomyocyte | Cdh11 | 2 | 3.10E-60 | 2.01777077 | 0.596 | 0.223 | 6.20E-57 |
| Cardiomyocyte | Bmper | 2 | 6.38E-59 | 3.89681013 | 0.589 | 0.229 | 1.28E-55 |
| Cardiomyocyte | Aff3 | 2 | 1.35E-58 | 2.55817766 | 0.593 | 0.288 | 2.71E-55 |
| Cardiomyocyte | Adamts17 | 2 | 3.26E-56 | 3.59793577 | 0.571 | 0.217 | 6.52E-53 |
| Cardiomyocyte | Mid1 | 2 | 8.56E-56 | 2.36257348 | 0.618 | 0.148 | 1.71E-52 |
| Cardiomyocyte | Uap1 | 2 | 9.18E-55 | 2.41122256 | 0.671 | 0.381 | 1.84E-51 |
| Cardiomyocyte | Dapk1 | 2 | 1.02E-52 | 2.05084589 | 0.647 | 0.381 | 2.04E-49 |
| Cardiomyocyte | Pde10a | 2 | 1.90E-52 | 3.23477484 | 0.543 | 0.177 | 3.80E-49 |
| Cardiomyocyte | Fap | 2 | 4.24E-51 | 2.59949129 | 0.606 | 0.17 | 8.49E-48 |
| Cardiomyocyte | Gfpt2 | 2 | 2.18E-50 | 3.70875077 | 0.566 | 0.292 | 4.35E-47 |
| Cardiomyocyte | Htra3 | 2 | 2.82E-50 | 2.71287215 | 0.589 | 0.274 | 5.65E-47 |
| Cardiomyocyte | Spon1 | 2 | 4.80E-49 | 3.6175164 | 0.52 | 0.107 | 9.61E-46 |
| Cardiomyocyte | Cacna2d3 | 2 | 1.35E-47 | 3.70374982 | 0.515 | 0.245 | 2.70E-44 |
| Cardiomyocyte | Sparc | 2 | 1.80E-47 | 1.05791101 | 0.776 | 0.481 | 3.59E-44 |

| | | | | | | | |
|---|---|---|---|---|---|---|---|
| Cardiomyocyte | Srgap1 | 2 | 4.22E-47 | 2.25753789 | 0.56 | 0.28 | 8.45E-44 |
| Cardiomyocyte | Kif20a | 2 | 1.49E-46 | 1.24901552 | 0.351 | 0.035 | 2.97E-43 |
| Cardiomyocyte | Sdk2 | 2 | 1.28E-44 | 3.59186719 | 0.503 | 0.164 | 2.56E-41 |
| Cardiomyocyte | RGD1563354 | 2 | 1.93E-44 | 3.7551124 | 0.505 | 0.251 | 3.86E-41 |
| Cardiomyocyte | Fbn2 | 2 | 8.48E-44 | 3.33857425 | 0.417 | 0.091 | 1.70E-40 |
| Cardiomyocyte | Colec12 | 2 | 2.30E-43 | 2.26991953 | 0.55 | 0.291 | 4.60E-40 |
| Cardiomyocyte | Nox4 | 2 | 3.31E-43 | 2.748083 | 0.571 | 0.191 | 6.63E-40 |
| Cardiomyocyte | Ltbp2 | 2 | 1.60E-42 | 2.92928656 | 0.523 | 0.2 | 3.20E-39 |
| Cardiomyocyte | Ar | 2 | 1.50E-41 | 2.97201678 | 0.538 | 0.239 | 3.01E-38 |
| Cardiomyocyte | Tspan5 | 2 | 2.05E-41 | 2.4533279 | 0.538 | 0.218 | 4.11E-38 |
| Cardiomyocyte | Efcc1 | 2 | 2.51E-40 | 3.92788478 | 0.508 | 0.158 | 5.02E-37 |
| Cardiomyocyte | Kif18b | 2 | 8.81E-40 | 1.86104548 | 0.396 | 0.111 | 1.76E-36 |
| Cardiomyocyte | Adamts5 | 2 | 1.66E-39 | 3.35748779 | 0.515 | 0.227 | 3.31E-36 |
| Cardiomyocyte | Loxl2 | 2 | 2.01E-36 | 2.94730174 | 0.528 | 0.27 | 4.03E-33 |
| Cardiomyocyte | Piezo2 | 2 | 1.06E-32 | 2.95507557 | 0.515 | 0.177 | 2.13E-29 |
| Cardiomyocyte | Nrk | 2 | 1.09E-32 | 3.13531251 | 0.447 | 0.179 | 2.17E-29 |
| Cardiomyocyte | Tgfb3 | 2 | 1.42E-31 | 2.85199292 | 0.462 | 0.175 | 2.85E-28 |
| Cardiomyocyte | Abca1 | 2 | 1.30E-29 | 1.11683862 | 0.642 | 0.352 | 2.60E-26 |
| Cardiomyocyte | Igfbp4 | 2 | 3.39E-29 | 2.23310905 | 0.483 | 0.123 | 6.77E-26 |
| Cardiomyocyte | Tnfaip8l3 | 2 | 1.69E-28 | 2.30053044 | 0.351 | 0.092 | 3.37E-25 |
| Cardiomyocyte | Col7a1 | 2 | 1.22E-26 | 2.69790043 | 0.354 | 0.062 | 2.43E-23 |
| Cardiomyocyte | Npas2 | 2 | 1.66E-26 | 1.30523968 | 0.627 | 0.377 | 3.32E-23 |
| Cardiomyocyte | Slc7a2 | 2 | 3.18E-26 | 2.51189113 | 0.475 | 0.187 | 6.37E-23 |
| Cardiomyocyte | Sdc2 | 2 | 1.11E-25 | 1.87987377 | 0.483 | 0.224 | 2.22E-22 |
| Cardiomyocyte | Calcrl | 2 | 1.38E-25 | 1.02056733 | 0.575 | 0.32 | 2.76E-22 |
| Cardiomyocyte | Cacnb4 | 2 | 2.31E-25 | 3.2333842 | 0.402 | 0.145 | 4.61E-22 |
| Cardiomyocyte | Gucy1a1 | 2 | 2.59E-25 | 1.74399382 | 0.475 | 0.2 | 5.18E-22 |
| Cardiomyocyte | Mmp2 | 2 | 2.45E-24 | 1.75528921 | 0.465 | 0.204 | 4.90E-21 |
| Cardiomyocyte | Errfi1 | 2 | 1.73E-23 | 2.24111397 | 0.505 | 0.218 | 3.46E-20 |
| Cardiomyocyte | Pdgfd | 2 | 2.20E-21 | 0.96621287 | 0.626 | 0.373 | 4.41E-18 |
| Cardiomyocyte | Adamts15 | 2 | 5.24E-21 | 2.37813863 | 0.422 | 0.154 | 1.05E-17 |

| | | | | | | | |
|---|---|---|---|---|---|---|---|
| Cardiomyocyte | Sh3gl2 | 2 | 2.16E-19 | 1.0341322 | 0.449 | 0.18 | 4.32E-16 |
| Cardiomyocyte | Opcml | 2 | 3.37E-17 | 2.80951303 | 0.401 | 0.126 | 6.74E-14 |
| Cardiomyocyte | Sema3d | 2 | 3.97E-17 | 1.43179951 | 0.445 | 0.144 | 7.94E-14 |
| Cardiomyocyte | Lurap1l | 2 | 4.79E-17 | 2.19100317 | 0.437 | 0.166 | 9.58E-14 |
| Cardiomyocyte | Col22a1 | 2 | 9.29E-11 | 1.05517185 | 0.45 | 0.189 | 1.86E-07 |
| Cardiomyocyte | Tp63 | 3 | 4.50E-121 | 3.74075526 | 0.352 | 0.047 | 9.00E-118 |
| Cardiomyocyte | Gtsf1 | 3 | 2.46E-109 | 1.93829608 | 0.297 | 0.033 | 4.92E-106 |
| Cardiomyocyte | Ncam1 | 3 | 1.52E-85 | 2.76051668 | 0.78 | 0.477 | 3.05E-82 |
| Cardiomyocyte | LOC689599 | 3 | 6.19E-75 | 3.0774249 | 0.335 | 0.075 | 1.24E-71 |
| Cardiomyocyte | AABR070581 | 3 | 4.10E-42 | 1.80614469 | 0.719 | 0.425 | 8.20E-39 |
| Cardiomyocyte | Nppa | 3 | 9.08E-30 | 1.82622833 | 0.618 | 0.357 | 1.82E-26 |
| Cardiomyocyte | AABR070581 | 3 | 2.00E-17 | 2.44095641 | 0.484 | 0.229 | 4.00E-14 |
| Cardiomyocyte | Mmp16 | 3 | 6.50E-17 | 0.59837403 | 0.065 | 0.324 | 1.30E-13 |
| Cardiomyocyte | Shc4 | 4 | 3.48E-68 | 1.65551057 | 0.471 | 0.112 | 6.96E-65 |
| Cardiomyocyte | Pla2g2a | 4 | 3.02E-60 | 1.08546174 | 0.398 | 0.075 | 6.04E-57 |
| Cardiomyocyte | Slc7a10 | 4 | 5.83E-60 | 2.51507816 | 0.389 | 0.128 | 1.17E-56 |
| Cardiomyocyte | Unc5d | 4 | 1.88E-59 | 0.71896725 | 0.462 | 0.126 | 3.77E-56 |
| Cardiomyocyte | Slit1 | 4 | 1.15E-32 | 2.6484507 | 0.446 | 0.187 | 2.31E-29 |
| Cardiomyocyte | Caln1 | 4 | 2.46E-21 | 0.88856506 | 0.456 | 0.203 | 4.93E-18 |
| Cardiomyocyte | Cpa6 | 4 | 3.22E-20 | 1.01719323 | 0.363 | 0.083 | 6.44E-17 |
| Cardiomyocyte | Shank3 | 5 | 5.35E-216 | 3.63260231 | 0.946 | 0.304 | 1.07E-212 |
| Cardiomyocyte | Knstrn | 5 | 2.32E-197 | 1.4963669 | 0.311 | 0.01 | 4.63E-194 |
| Cardiomyocyte | Fli1 | 5 | 1.28E-195 | 3.45287751 | 0.93 | 0.32 | 2.57E-192 |
| Cardiomyocyte | AABR070584 | 5 | 7.51E-186 | 3.80877102 | 0.35 | 0.006 | 1.50E-182 |
| Cardiomyocyte | Cyyr1 | 5 | 1.42E-182 | 3.68091213 | 0.867 | 0.221 | 2.85E-179 |
| Cardiomyocyte | Blk | 5 | 5.99E-176 | 1.097442 | 0.385 | 0.034 | 1.20E-172 |
| Cardiomyocyte | Itpkb | 5 | 5.40E-172 | 3.64386184 | 0.885 | 0.315 | 1.08E-168 |
| Cardiomyocyte | Dach1 | 5 | 3.24E-171 | 3.54050369 | 0.88 | 0.268 | 6.48E-168 |
| Cardiomyocyte | Flt1 | 5 | 2.71E-169 | 3.49782531 | 0.872 | 0.267 | 5.42E-166 |
| Cardiomyocyte | Prkch | 5 | 2.65E-166 | 3.51970823 | 0.85 | 0.27 | 5.30E-163 |
| Cardiomyocyte | Adgrf5 | 5 | 1.11E-156 | 3.64727444 | 0.835 | 0.28 | 2.21E-153 |

| | | | | | | | |
|---|---|---|---|---|---|---|---|
| Cardiomyocyte | Ptprb | 5 | 1.46E-156 | 3.31066319 | 0.848 | 0.239 | 2.92E-153 |
| Cardiomyocyte | Kitlg | 5 | 2.28E-136 | 3.39401166 | 0.807 | 0.266 | 4.57E-133 |
| Cardiomyocyte | Galnt14 | 5 | 1.46E-129 | 3.25276003 | 0.422 | 0.036 | 2.92E-126 |
| Cardiomyocyte | Ablim3 | 5 | 3.59E-125 | 3.94465461 | 0.748 | 0.342 | 7.17E-122 |
| Cardiomyocyte | Ca3 | 5 | 1.55E-121 | 1.95714552 | 0.317 | 0.028 | 3.10E-118 |
| Cardiomyocyte | Gabra4 | 5 | 5.11E-119 | 1.27424411 | 0.378 | 0.072 | 1.02E-115 |
| Cardiomyocyte | Ankrd34c | 5 | 1.94E-117 | 2.66031796 | 0.385 | 0.068 | 3.89E-114 |
| Cardiomyocyte | Emcn | 5 | 4.19E-110 | 3.35517373 | 0.733 | 0.287 | 8.39E-107 |
| Cardiomyocyte | Adgrl4 | 5 | 6.59E-106 | 3.3961778 | 0.713 | 0.196 | 1.32E-102 |
| Cardiomyocyte | Mfsd4a | 5 | 1.62E-105 | 1.75156121 | 0.389 | 0.027 | 3.23E-102 |
| Cardiomyocyte | Ptch21 | 5 | 1.75E-101 | 1.17168436 | 0.426 | 0.083 | 3.51E-98 |
| Cardiomyocyte | Clec9a | 5 | 5.26E-98 | 2.90797783 | 0.33 | 0.031 | 1.05E-94 |
| Cardiomyocyte | Podnl1 | 5 | 7.10E-94 | 2.42223686 | 0.426 | 0.121 | 1.42E-90 |
| Cardiomyocyte | Nr5a2 | 5 | 2.66E-93 | 4.49985192 | 0.411 | 0.036 | 5.32E-90 |
| Cardiomyocyte | Rasgrf1 | 5 | 1.45E-87 | 1.54249533 | 0.363 | 0.08 | 2.90E-84 |
| Cardiomyocyte | Dnm3 | 5 | 1.56E-85 | 2.98097308 | 0.689 | 0.326 | 3.11E-82 |
| Cardiomyocyte | Prss12 | 5 | 1.63E-85 | 2.486141 | 0.363 | 0.034 | 3.25E-82 |
| Cardiomyocyte | Calcrl1 | 5 | 1.52E-84 | 2.35975901 | 0.754 | 0.31 | 3.04E-81 |
| Cardiomyocyte | Cmtm8 | 5 | 1.60E-84 | 3.67700727 | 0.676 | 0.218 | 3.19E-81 |
| Cardiomyocyte | Fgr | 5 | 9.88E-84 | 0.79040531 | 0.391 | 0.109 | 1.98E-80 |
| Cardiomyocyte | Mx1 | 5 | 2.71E-83 | 1.30848495 | 0.396 | 0.125 | 5.41E-80 |
| Cardiomyocyte | Csmd1 | 5 | 1.62E-80 | 1.15564547 | 0.428 | 0.11 | 3.24E-77 |
| Cardiomyocyte | Hspa1b | 5 | 4.23E-79 | 1.55648033 | 0.359 | 0.087 | 8.47E-76 |
| Cardiomyocyte | Meox2 | 5 | 3.11E-76 | 3.23961656 | 0.657 | 0.256 | 6.22E-73 |
| Cardiomyocyte | Ccdc85a | 5 | 6.53E-75 | 1.46917157 | 0.902 | 0.599 | 1.31E-71 |
| Cardiomyocyte | Efnb2 | 5 | 6.96E-75 | 3.43819779 | 0.67 | 0.207 | 1.39E-71 |
| Cardiomyocyte | Nrp2 | 5 | 3.86E-73 | 2.86572522 | 0.68 | 0.301 | 7.73E-70 |
| Cardiomyocyte | Anxa3 | 5 | 2.63E-71 | 3.66576168 | 0.585 | 0.135 | 5.27E-68 |
| Cardiomyocyte | Neil3 | 5 | 4.59E-71 | 1.65697159 | 0.3 | 0.032 | 9.18E-68 |
| Cardiomyocyte | Dipk2b | 5 | 5.18E-71 | 4.0290428 | 0.648 | 0.242 | 1.04E-67 |
| Cardiomyocyte | Myrip | 5 | 4.24E-69 | 3.4675784 | 0.604 | 0.343 | 8.47E-66 |

| | | | | | | | |
|---|---|---|---|---|---|---|---|
| Cardiomyocyte | Nav3 | 5 | 4.54E-68 | 2.12473073 | 0.785 | 0.53 | 9.07E-65 |
| Cardiomyocyte | Khdrbs2 | 5 | 2.15E-66 | 2.09193281 | 0.372 | 0.088 | 4.30E-63 |
| Cardiomyocyte | Mcf2l | 5 | 1.46E-62 | 3.54520194 | 0.574 | 0.215 | 2.93E-59 |
| Cardiomyocyte | Ednrb | 5 | 1.56E-61 | 2.55888131 | 0.502 | 0.085 | 3.11E-58 |
| Cardiomyocyte | AABR070606 | 5 | 5.00E-60 | 4.0732433 | 0.537 | 0.235 | 9.99E-57 |
| Cardiomyocyte | Podxl | 5 | 6.66E-59 | 3.07011655 | 0.557 | 0.206 | 1.33E-55 |
| Cardiomyocyte | Thsd7a | 5 | 1.32E-58 | 3.04854058 | 0.615 | 0.239 | 2.63E-55 |
| Cardiomyocyte | St6galnac3 | 5 | 1.74E-58 | 3.25707729 | 0.624 | 0.257 | 3.48E-55 |
| Cardiomyocyte | AABR070042 | 5 | 1.65E-57 | 0.30196574 | 0.446 | 0.079 | 3.30E-54 |
| Cardiomyocyte | Mgll | 5 | 5.21E-57 | 1.65527096 | 0.746 | 0.479 | 1.04E-53 |
| Cardiomyocyte | Mx2 | 5 | 1.15E-56 | 2.69285331 | 0.452 | 0.134 | 2.29E-53 |
| Cardiomyocyte | Myo10 | 5 | 1.54E-56 | 2.65972192 | 0.635 | 0.368 | 3.09E-53 |
| Cardiomyocyte | Slc26a10 | 5 | 7.45E-56 | 3.84118563 | 0.567 | 0.274 | 1.49E-52 |
| Cardiomyocyte | Lyn | 5 | 4.14E-53 | 1.724181 | 0.704 | 0.433 | 8.28E-50 |
| Cardiomyocyte | Herc6 | 5 | 4.36E-53 | 3.04267547 | 0.548 | 0.155 | 8.72E-50 |
| Cardiomyocyte | Emid1 | 5 | 1.81E-49 | 2.65770726 | 0.465 | 0.113 | 3.63E-46 |
| Cardiomyocyte | Ikzf3 | 5 | 1.05E-47 | 0.49461771 | 0.357 | 0.075 | 2.11E-44 |
| Cardiomyocyte | Sox13 | 5 | 9.69E-46 | 1.96663419 | 0.661 | 0.397 | 1.94E-42 |
| Cardiomyocyte | Rai14 | 5 | 1.94E-44 | 2.2433391 | 0.522 | 0.195 | 3.88E-41 |
| Cardiomyocyte | Gcnt2 | 5 | 9.87E-44 | 2.88110103 | 0.557 | 0.258 | 1.97E-40 |
| Cardiomyocyte | Ptafr | 5 | 1.71E-43 | 0.89545205 | 0.454 | 0.09 | 3.41E-40 |
| Cardiomyocyte | Fabp4 | 5 | 1.89E-43 | 2.0981568 | 0.591 | 0.307 | 3.78E-40 |
| Cardiomyocyte | Chn1 | 5 | 5.27E-43 | 1.89080559 | 0.626 | 0.276 | 1.05E-39 |
| Cardiomyocyte | Rasa4 | 5 | 1.63E-42 | 3.23417154 | 0.591 | 0.323 | 3.27E-39 |
| Cardiomyocyte | Mgmt | 5 | 4.06E-42 | 1.83373423 | 0.641 | 0.383 | 8.12E-39 |
| Cardiomyocyte | Mcam | 5 | 1.03E-41 | 1.89420768 | 0.598 | 0.235 | 2.07E-38 |
| Cardiomyocyte | Ano2 | 5 | 1.24E-41 | 3.63352439 | 0.426 | 0.149 | 2.47E-38 |
| Cardiomyocyte | Pfkfb3 | 5 | 9.92E-41 | 2.60127646 | 0.591 | 0.275 | 1.98E-37 |
| Cardiomyocyte | Unc5b | 5 | 1.33E-39 | 1.93336251 | 0.624 | 0.31 | 2.65E-36 |
| Cardiomyocyte | Entpd1 | 5 | 1.70E-39 | 1.95240462 | 0.583 | 0.255 | 3.40E-36 |
| Cardiomyocyte | Cadps2 | 5 | 1.35E-38 | 3.05673683 | 0.504 | 0.221 | 2.71E-35 |

| | | | | | | | |
|---|---|---|---|---|---|---|---|
| Cardiomyocyte | Cxcl12 | 5 | 5.05E-37 | 2.62506369 | 0.537 | 0.23 | 1.01E-33 |
| Cardiomyocyte | Wt1 | 5 | 4.19E-35 | 3.339778 | 0.467 | 0.167 | 8.38E-32 |
| Cardiomyocyte | Ccser1 | 5 | 5.13E-32 | 2.23255363 | 0.539 | 0.278 | 1.03E-28 |
| Cardiomyocyte | Epha4 | 5 | 1.62E-31 | 1.89699204 | 0.565 | 0.241 | 3.24E-28 |
| Cardiomyocyte | Car10 | 5 | 3.28E-30 | 0.34039835 | 0.365 | 0.109 | 6.57E-27 |
| Cardiomyocyte | Ndc80 | 5 | 4.26E-28 | 2.346165 | 0.317 | 0.047 | 8.52E-25 |
| Cardiomyocyte | Nxn | 5 | 3.31E-27 | 1.26279393 | 0.663 | 0.401 | 6.63E-24 |
| Cardiomyocyte | Fshr | 5 | 6.59E-27 | 2.22063698 | 0.328 | 0.064 | 1.32E-23 |
| Cardiomyocyte | Lrrtm4 | 5 | 3.25E-25 | 0.99065472 | 0.446 | 0.138 | 6.50E-22 |
| Cardiomyocyte | Nkd2 | 5 | 3.80E-25 | 1.4073823 | 0.433 | 0.181 | 7.61E-22 |
| Cardiomyocyte | Sema3d1 | 5 | 1.08E-24 | 2.11468248 | 0.457 | 0.152 | 2.15E-21 |
| Cardiomyocyte | Mertk | 5 | 1.34E-23 | 1.56596804 | 0.489 | 0.175 | 2.67E-20 |
| Cardiomyocyte | Abcd2 | 5 | 9.88E-23 | 0.46566784 | 0.474 | 0.177 | 1.98E-19 |
| Cardiomyocyte | Ncaph | 5 | 1.41E-22 | 1.10607056 | 0.417 | 0.134 | 2.81E-19 |
| Cardiomyocyte | Cemip2 | 5 | 1.08E-20 | 1.00684877 | 0.502 | 0.217 | 2.15E-17 |
| Cardiomyocyte | Tll1 | 5 | 2.63E-19 | 1.3823574 | 0.533 | 0.271 | 5.26E-16 |
| Cardiomyocyte | Ninj2 | 5 | 1.95E-17 | 0.50621991 | 0.493 | 0.189 | 3.91E-14 |
| Cardiomyocyte | Ccdc148 | 5 | 9.45E-17 | 0.63611269 | 0.45 | 0.175 | 1.89E-13 |
| Cardiomyocyte | Atp5f1e | 5 | 5.95E-11 | 0.32296103 | 0.557 | 0.305 | 1.19E-07 |
| Cardiomyocyte | Tigit | 6 | 1.79E-85 | 3.63635833 | 0.369 | 0.069 | 3.59E-82 |
| Cardiomyocyte | Slc36a2 | 6 | 3.58E-81 | 2.17672385 | 0.44 | 0.107 | 7.16E-78 |
| Cardiomyocyte | AABR070720 | 6 | 7.90E-41 | 1.40723046 | 0.454 | 0.144 | 1.58E-37 |
| Cardiomyocyte | Gata3 | 6 | 5.16E-27 | 0.50972192 | 0.355 | 0.103 | 1.03E-23 |
| Cardiomyocyte | Sema3e | 6 | 1.09E-11 | 0.89848869 | 0.388 | 0.124 | 2.17E-08 |
| Cardiomyocyte | Mcoln2 | 7 | 0 | 2.00712165 | 0.88 | 0.064 | 0 |
| Cardiomyocyte | LOC1083484 | 7 | 0 | 3.72326379 | 0.631 | 0.04 | 0 |
| Cardiomyocyte | RGD1564854 | 7 | 4.56E-275 | 3.03171329 | 0.751 | 0.09 | 9.13E-272 |
| Cardiomyocyte | AABR070637 | 7 | 2.87E-262 | 0.97661714 | 0.758 | 0.088 | 5.74E-259 |
| Cardiomyocyte | Nrg4 | 7 | 1.28E-261 | 2.47760187 | 0.953 | 0.193 | 2.55E-258 |
| Cardiomyocyte | Slc6a15 | 7 | 2.57E-260 | 2.43163979 | 0.81 | 0.09 | 5.14E-257 |
| Cardiomyocyte | Zdbf2 | 7 | 2.12E-249 | 2.13951322 | 0.661 | 0.102 | 4.24E-246 |

| | | | | | | | |
|---|---|---|---|---|---|---|---|
| Cardiomyocyte | Trarg1 | 7 | 1.36E-238 | 2.19009359 | 0.616 | 0.055 | 2.73E-235 |
| Cardiomyocyte | AABR070685 | 7 | 3.53E-236 | 2.79039226 | 0.853 | 0.114 | 7.07E-233 |
| Cardiomyocyte | RGD1564053 | 7 | 1.38E-233 | 1.2759122 | 0.965 | 0.253 | 2.76E-230 |
| Cardiomyocyte | Klrc2 | 7 | 6.89E-233 | 1.75018425 | 0.499 | 0.085 | 1.38E-229 |
| Cardiomyocyte | Tbx21 | 7 | 4.00E-231 | 0.69969886 | 0.791 | 0.113 | 8.01E-228 |
| Cardiomyocyte | Stmn2 | 7 | 1.03E-225 | 3.20908716 | 0.633 | 0.057 | 2.06E-222 |
| Cardiomyocyte | AABR070632 | 7 | 5.56E-225 | 2.71246574 | 0.9 | 0.17 | 1.11E-221 |
| Cardiomyocyte | Susd5 | 7 | 3.23E-215 | 0.35921441 | 0.913 | 0.226 | 6.46E-212 |
| Cardiomyocyte | Olr1 | 7 | 4.93E-215 | 1.50756455 | 0.98 | 0.297 | 9.87E-212 |
| Cardiomyocyte | Slc6a19 | 7 | 1.40E-210 | 2.33280632 | 0.83 | 0.183 | 2.80E-207 |
| Cardiomyocyte | AABR070127 | 7 | 2.54E-202 | 1.51777877 | 0.606 | 0.141 | 5.07E-199 |
| Cardiomyocyte | AC094126.1 | 7 | 4.55E-201 | 1.40783375 | 0.686 | 0.11 | 9.09E-198 |
| Cardiomyocyte | Lyve1 | 7 | 1.17E-200 | 0.4433641 | 0.93 | 0.19 | 2.34E-197 |
| Cardiomyocyte | Depdc1b | 7 | 5.27E-199 | 0.96940639 | 0.738 | 0.08 | 1.05E-195 |
| Cardiomyocyte | Abca4 | 7 | 2.67E-198 | 0.87987464 | 0.963 | 0.24 | 5.33E-195 |
| Cardiomyocyte | Tnnt1 | 7 | 1.13E-195 | 0.98606936 | 0.726 | 0.102 | 2.26E-192 |
| Cardiomyocyte | Grid2 | 7 | 2.35E-195 | 1.33821765 | 0.875 | 0.132 | 4.70E-192 |
| Cardiomyocyte | Slc26a31 | 7 | 4.09E-192 | 1.14363618 | 0.84 | 0.218 | 8.18E-189 |
| Cardiomyocyte | Ly49s6 | 7 | 7.71E-189 | 0.32560285 | 0.938 | 0.217 | 1.54E-185 |
| Cardiomyocyte | Hamp | 7 | 8.38E-189 | 1.09110991 | 0.823 | 0.216 | 1.68E-185 |
| Cardiomyocyte | Rspo2 | 7 | 8.51E-189 | 2.04166625 | 0.928 | 0.216 | 1.70E-185 |
| Cardiomyocyte | Cdo11 | 7 | 1.50E-188 | 1.04244766 | 0.953 | 0.246 | 3.00E-185 |
| Cardiomyocyte | Tpx2 | 7 | 2.78E-188 | 1.42224433 | 0.963 | 0.316 | 5.56E-185 |
| Cardiomyocyte | Slit11 | 7 | 2.10E-187 | 0.76045237 | 0.783 | 0.164 | 4.20E-184 |
| Cardiomyocyte | Col19a11 | 7 | 1.04E-185 | 0.85466597 | 0.96 | 0.262 | 2.08E-182 |
| Cardiomyocyte | Penk | 7 | 6.01E-184 | 1.50751632 | 0.908 | 0.219 | 1.20E-180 |
| Cardiomyocyte | Cdh12 | 7 | 9.53E-180 | 1.33525589 | 0.89 | 0.202 | 1.91E-176 |
| Cardiomyocyte | Ak5 | 7 | 1.08E-178 | 0.57654444 | 0.968 | 0.292 | 2.15E-175 |
| Cardiomyocyte | Lmx1a | 7 | 3.47E-175 | 0.38615992 | 0.843 | 0.17 | 6.94E-172 |
| Cardiomyocyte | Epsti1 | 7 | 4.87E-174 | 0.5044216 | 0.878 | 0.189 | 9.73E-171 |
| Cardiomyocyte | Procr | 7 | 2.59E-173 | 0.78326889 | 0.955 | 0.289 | 5.18E-170 |

| | | | | | | | |
|---|---|---|---|---|---|---|---|
| Cardiomyocyte | AABR070269 | 7 | 2.51E-172 | 3.02529927 | 0.641 | 0.114 | 5.02E-169 |
| Cardiomyocyte | Zp3r | 7 | 1.38E-169 | 0.71091187 | 0.698 | 0.133 | 2.76E-166 |
| Cardiomyocyte | Myo5b1 | 7 | 3.26E-169 | 0.44850913 | 0.865 | 0.209 | 6.52E-166 |
| Cardiomyocyte | Dmp1 | 7 | 5.47E-169 | 0.9066554 | 0.389 | 0.048 | 1.09E-165 |
| Cardiomyocyte | Il22ra1 | 7 | 6.71E-169 | 3.04435408 | 0.681 | 0.138 | 1.34E-165 |
| Cardiomyocyte | Col25a1 | 7 | 1.38E-167 | 1.28175986 | 0.865 | 0.192 | 2.76E-164 |
| Cardiomyocyte | Col13a11 | 7 | 6.81E-167 | 0.58992865 | 0.968 | 0.273 | 1.36E-163 |
| Cardiomyocyte | Brinp2 | 7 | 1.18E-163 | 1.31225145 | 0.913 | 0.222 | 2.35E-160 |
| Cardiomyocyte | Ltk | 7 | 1.71E-163 | 1.15827909 | 0.753 | 0.144 | 3.43E-160 |
| Cardiomyocyte | Fgf9 | 7 | 2.27E-162 | 1.32968869 | 0.938 | 0.288 | 4.53E-159 |
| Cardiomyocyte | Ptprt | 7 | 3.53E-162 | 0.61568858 | 0.905 | 0.271 | 7.07E-159 |
| Cardiomyocyte | F13a1 | 7 | 3.95E-162 | 1.04645413 | 0.773 | 0.229 | 7.90E-159 |
| Cardiomyocyte | Chrdl1 | 7 | 2.22E-159 | 0.53337937 | 0.855 | 0.188 | 4.44E-156 |
| Cardiomyocyte | Armc4 | 7 | 2.71E-159 | 0.58874108 | 0.955 | 0.282 | 5.41E-156 |
| Cardiomyocyte | Kif20b | 7 | 3.58E-159 | 0.70354817 | 0.918 | 0.219 | 7.17E-156 |
| Cardiomyocyte | Il2rb | 7 | 8.56E-159 | 2.21125524 | 0.576 | 0.07 | 1.71E-155 |
| Cardiomyocyte | Gap43 | 7 | 2.23E-157 | 0.41551445 | 0.656 | 0.182 | 4.46E-154 |
| Cardiomyocyte | Kcnq3 | 7 | 8.75E-157 | 1.23286097 | 0.818 | 0.189 | 1.75E-153 |
| Cardiomyocyte | Pfkfb1 | 7 | 2.91E-155 | 0.68201485 | 0.91 | 0.211 | 5.81E-152 |
| Cardiomyocyte | Shroom2 | 7 | 4.97E-155 | 1.41008057 | 0.738 | 0.164 | 9.94E-152 |
| Cardiomyocyte | Rbm44 | 7 | 3.02E-154 | 0.95611971 | 0.753 | 0.196 | 6.05E-151 |
| Cardiomyocyte | Fstl4 | 7 | 7.78E-154 | 2.08948502 | 0.868 | 0.188 | 1.56E-150 |
| Cardiomyocyte | Alk | 7 | 3.00E-153 | 0.54514549 | 0.83 | 0.209 | 5.99E-150 |
| Cardiomyocyte | AABR070131 | 7 | 3.62E-152 | 1.56522971 | 0.693 | 0.148 | 7.25E-149 |
| Cardiomyocyte | Kif171 | 7 | 2.05E-151 | 0.74879649 | 0.895 | 0.238 | 4.11E-148 |
| Cardiomyocyte | Tmem163 | 7 | 1.05E-150 | 0.9503579 | 0.716 | 0.156 | 2.11E-147 |
| Cardiomyocyte | Fmod | 7 | 2.60E-149 | 1.60776855 | 0.741 | 0.155 | 5.19E-146 |
| Cardiomyocyte | Podnl11 | 7 | 5.44E-148 | 0.334424 | 0.631 | 0.107 | 1.09E-144 |
| Cardiomyocyte | RT1-Da | 7 | 3.24E-147 | 0.6608571 | 0.813 | 0.171 | 6.48E-144 |
| Cardiomyocyte | Ttll9 | 7 | 1.14E-146 | 1.00809442 | 0.878 | 0.239 | 2.28E-143 |
| Cardiomyocyte | Neb | 7 | 1.71E-146 | 0.47117541 | 0.835 | 0.236 | 3.41E-143 |

| | | | | | | | |
|---|---|---|---|---|---|---|---|
| Cardiomyocyte | Slc45a3 | 7 | 1.71E-146 | 0.48591778 | 0.815 | 0.192 | 3.43E-143 |
| Cardiomyocyte | Mettl241 | 7 | 2.14E-144 | 0.67997466 | 0.928 | 0.24 | 4.29E-141 |
| Cardiomyocyte | Cadm1 | 7 | 1.08E-143 | 0.26704262 | 0.746 | 0.154 | 2.16E-140 |
| Cardiomyocyte | Gna14 | 7 | 5.27E-143 | 1.73547698 | 0.875 | 0.269 | 1.05E-139 |
| Cardiomyocyte | Adap2 | 7 | 5.26E-142 | 0.93493514 | 0.943 | 0.293 | 1.05E-138 |
| Cardiomyocyte | Megf11 | 7 | 7.67E-142 | 1.06958798 | 0.878 | 0.24 | 1.53E-138 |
| Cardiomyocyte | Edar | 7 | 1.04E-140 | 0.84247765 | 0.651 | 0.112 | 2.08E-137 |
| Cardiomyocyte | Grk3 | 7 | 1.66E-139 | 0.28618955 | 0.815 | 0.182 | 3.32E-136 |
| Cardiomyocyte | Cyp4f18 | 7 | 1.11E-138 | 0.45813458 | 0.648 | 0.225 | 2.22E-135 |
| Cardiomyocyte | Adgrg2 | 7 | 1.67E-138 | 1.1116058 | 0.815 | 0.217 | 3.33E-135 |
| Cardiomyocyte | Pak71 | 7 | 2.35E-136 | 0.37408497 | 0.808 | 0.229 | 4.69E-133 |
| Cardiomyocyte | LOC1036933 | 7 | 5.49E-134 | 0.48421931 | 0.738 | 0.254 | 1.10E-130 |
| Cardiomyocyte | Adamts20 | 7 | 1.47E-133 | 0.30930936 | 0.741 | 0.281 | 2.94E-130 |
| Cardiomyocyte | Lrrc4c | 7 | 2.29E-133 | 1.19127502 | 0.81 | 0.228 | 4.57E-130 |
| Cardiomyocyte | Grip2 | 7 | 1.74E-132 | 0.85790592 | 0.793 | 0.207 | 3.49E-129 |
| Cardiomyocyte | Nyap2 | 7 | 2.63E-132 | 0.94586671 | 0.893 | 0.236 | 5.26E-129 |
| Cardiomyocyte | Prc1 | 7 | 1.63E-130 | 1.00185735 | 0.586 | 0.207 | 3.27E-127 |
| Cardiomyocyte | Cmtm5 | 7 | 1.90E-129 | 1.81618164 | 0.426 | 0.023 | 3.81E-126 |
| Cardiomyocyte | Caln11 | 7 | 5.98E-125 | 1.54657002 | 0.776 | 0.181 | 1.20E-121 |
| Cardiomyocyte | Olfml2b | 7 | 7.61E-125 | 0.4151394 | 0.868 | 0.256 | 1.52E-121 |
| Cardiomyocyte | Ano3 | 7 | 5.12E-123 | 1.38978134 | 0.815 | 0.225 | 1.02E-119 |
| Cardiomyocyte | Sh3rf21 | 7 | 1.10E-121 | 0.52943634 | 0.733 | 0.242 | 2.19E-118 |
| Cardiomyocyte | Igfbp2 | 7 | 1.13E-121 | 1.83876963 | 0.509 | 0.098 | 2.25E-118 |
| Cardiomyocyte | Ttc29 | 7 | 2.24E-120 | 1.53812845 | 0.838 | 0.2 | 4.47E-117 |
| Cardiomyocyte | Vtcn1 | 7 | 5.31E-119 | 1.37867644 | 0.718 | 0.157 | 1.06E-115 |
| Cardiomyocyte | Tshr | 7 | 5.38E-119 | 0.495238 | 0.718 | 0.177 | 1.08E-115 |
| Cardiomyocyte | Fbxw101 | 7 | 2.94E-116 | 0.28080683 | 0.973 | 0.423 | 5.88E-113 |
| Cardiomyocyte | Astn2 | 7 | 1.59E-115 | 0.49397474 | 0.953 | 0.346 | 3.18E-112 |
| Cardiomyocyte | Gpd11 | 7 | 1.21E-114 | 0.6294529 | 0.945 | 0.353 | 2.41E-111 |
| Cardiomyocyte | Adgrb31 | 7 | 3.39E-112 | 0.71482778 | 0.798 | 0.215 | 6.77E-109 |
| Cardiomyocyte | Rnf43 | 7 | 1.44E-109 | 1.0253842 | 0.868 | 0.291 | 2.89E-106 |

| | | | | | | | |
|---|---|---|---|---|---|---|---|
| Cardiomyocyte | AABR070504 | 7 | 2.40E-109 | 0.64458554 | 0.953 | 0.34 | 4.79E-106 |
| Cardiomyocyte | Fermt3 | 7 | 1.18E-108 | 0.57833404 | 0.599 | 0.227 | 2.36E-105 |
| Cardiomyocyte | Ccn5 | 7 | 2.61E-108 | 0.34015311 | 0.843 | 0.27 | 5.21E-105 |
| Cardiomyocyte | Bcl11b | 7 | 2.70E-108 | 0.79460208 | 0.95 | 0.346 | 5.40E-105 |
| Cardiomyocyte | Serpine2 | 7 | 8.43E-107 | 0.77241739 | 0.509 | 0.17 | 1.69E-103 |
| Cardiomyocyte | Hcn4 | 7 | 5.73E-106 | 0.76292434 | 0.648 | 0.242 | 1.15E-102 |
| Cardiomyocyte | Anks1b | 7 | 9.31E-106 | 0.99296209 | 0.983 | 0.373 | 1.86E-102 |
| Cardiomyocyte | Apobec1 | 7 | 1.90E-105 | 0.64504627 | 0.613 | 0.226 | 3.79E-102 |
| Cardiomyocyte | Pcdh19 | 7 | 2.20E-105 | 0.56471633 | 0.676 | 0.208 | 4.40E-102 |
| Cardiomyocyte | Aff2 | 7 | 3.68E-105 | 0.3237859 | 0.943 | 0.324 | 7.37E-102 |
| Cardiomyocyte | Sema3a | 7 | 1.41E-104 | 0.65677459 | 0.83 | 0.303 | 2.82E-101 |
| Cardiomyocyte | Nr4a1 | 7 | 2.13E-103 | 0.38718949 | 0.963 | 0.384 | 4.26E-100 |
| Cardiomyocyte | Nusap11 | 7 | 1.01E-102 | 0.6343051 | 0.828 | 0.284 | 2.02E-99 |
| Cardiomyocyte | Glp2r | 7 | 1.44E-101 | 1.4230645 | 0.509 | 0.165 | 2.88E-98 |
| Cardiomyocyte | Ptgs2 | 7 | 2.79E-101 | 0.35189512 | 0.456 | 0.171 | 5.58E-98 |
| Cardiomyocyte | Ccl21 | 7 | 9.98E-101 | 0.26835718 | 0.484 | 0.188 | 2.00E-97 |
| Cardiomyocyte | AABR070134 | 7 | 2.89E-100 | 1.1103792 | 0.713 | 0.21 | 5.79E-97 |
| Cardiomyocyte | Hmox1 | 7 | 7.04E-100 | 1.23009647 | 0.596 | 0.215 | 1.41E-96 |
| Cardiomyocyte | Sctr | 7 | 1.44E-99 | 0.98837659 | 0.683 | 0.231 | 2.87E-96 |
| Cardiomyocyte | Troap | 7 | 1.92E-98 | 1.44923675 | 0.646 | 0.124 | 3.84E-95 |
| Cardiomyocyte | Etv3l | 7 | 2.97E-98 | 1.03033155 | 0.766 | 0.217 | 5.95E-95 |
| Cardiomyocyte | Adamts3 | 7 | 4.13E-98 | 0.68357007 | 0.858 | 0.316 | 8.26E-95 |
| Cardiomyocyte | Megf10 | 7 | 1.35E-97 | 0.86574079 | 0.798 | 0.297 | 2.70E-94 |
| Cardiomyocyte | Stard10 | 7 | 1.44E-96 | 1.50090363 | 0.569 | 0.161 | 2.89E-93 |
| Cardiomyocyte | Gm2a | 7 | 6.15E-96 | 0.31223005 | 0.88 | 0.338 | 1.23E-92 |
| Cardiomyocyte | Slc7a101 | 7 | 1.02E-95 | 1.68055253 | 0.519 | 0.122 | 2.04E-92 |
| Cardiomyocyte | AABR070635 | 7 | 6.13E-94 | 0.61781042 | 0.758 | 0.308 | 1.23E-90 |
| Cardiomyocyte | Cacna1e | 7 | 1.11E-93 | 0.4682562 | 0.86 | 0.311 | 2.22E-90 |
| Cardiomyocyte | Gpx3 | 7 | 4.99E-92 | 0.46552165 | 0.683 | 0.238 | 9.98E-89 |
| Cardiomyocyte | Sybu | 7 | 7.66E-92 | 0.32647184 | 0.756 | 0.255 | 1.53E-88 |
| Cardiomyocyte | Ptprn2 | 7 | 9.57E-92 | 0.64967659 | 0.771 | 0.263 | 1.91E-88 |

| | | | | | | | |
|---|---|---|---|---|---|---|---|
| Cardiomyocyte | Tuba1b | 7 | 2.18E-91 | 0.65512571 | 0.474 | 0.188 | 4.36E-88 |
| Cardiomyocyte | Sgo2 | 7 | 6.10E-90 | 1.27752491 | 0.681 | 0.186 | 1.22E-86 |
| Cardiomyocyte | Tafa2 | 7 | 5.71E-89 | 0.74257538 | 0.825 | 0.31 | 1.14E-85 |
| Cardiomyocyte | Dgat2 | 7 | 3.27E-88 | 0.72263645 | 0.791 | 0.306 | 6.54E-85 |
| Cardiomyocyte | Sfrp4 | 7 | 1.03E-87 | 1.1135332 | 0.444 | 0.122 | 2.06E-84 |
| Cardiomyocyte | Alpl | 7 | 2.67E-87 | 0.39502594 | 0.878 | 0.327 | 5.33E-84 |
| Cardiomyocyte | Arhgef391 | 7 | 2.69E-87 | 1.4927907 | 0.651 | 0.17 | 5.37E-84 |
| Cardiomyocyte | Sele | 7 | 4.16E-87 | 1.30062738 | 0.691 | 0.2 | 8.32E-84 |
| Cardiomyocyte | Kit | 7 | 1.97E-86 | 0.39627734 | 0.594 | 0.25 | 3.94E-83 |
| Cardiomyocyte | Fam163a | 7 | 2.33E-86 | 0.71316978 | 0.776 | 0.284 | 4.67E-83 |
| Cardiomyocyte | LOC1001348 | 7 | 7.96E-83 | 0.47197796 | 0.818 | 0.333 | 1.59E-79 |
| Cardiomyocyte | Nrg1 | 7 | 4.79E-82 | 0.38002833 | 0.556 | 0.248 | 9.58E-79 |
| Cardiomyocyte | Gpam2 | 7 | 1.86E-79 | 0.30724433 | 0.945 | 0.356 | 3.72E-76 |
| Cardiomyocyte | Pcp4l1 | 7 | 2.52E-78 | 0.69891649 | 0.978 | 0.386 | 5.04E-75 |
| Cardiomyocyte | Pdgfc | 7 | 3.55E-77 | 0.26171525 | 0.516 | 0.237 | 7.11E-74 |
| Cardiomyocyte | Lgals1 | 7 | 4.48E-76 | 0.6566869 | 0.895 | 0.357 | 8.96E-73 |
| Cardiomyocyte | Lmln | 7 | 5.94E-76 | 1.26180084 | 0.678 | 0.28 | 1.19E-72 |
| Cardiomyocyte | Aldh1a1 | 7 | 7.50E-75 | 0.86270562 | 0.521 | 0.24 | 1.50E-71 |
| Cardiomyocyte | Ptgs1 | 7 | 5.62E-73 | 0.46917757 | 0.554 | 0.269 | 1.12E-69 |
| Cardiomyocyte | Rspo1 | 7 | 2.94E-71 | 1.20819364 | 0.671 | 0.174 | 5.88E-68 |
| Cardiomyocyte | Ccl2 | 7 | 2.14E-69 | 1.28607572 | 0.449 | 0.152 | 4.29E-66 |
| Cardiomyocyte | Rcan1 | 7 | 3.60E-68 | 0.43232545 | 0.958 | 0.398 | 7.20E-65 |
| Cardiomyocyte | Kif2c | 7 | 1.31E-67 | 1.43833418 | 0.496 | 0.083 | 2.62E-64 |
| Cardiomyocyte | Neurl11 | 7 | 3.50E-67 | 0.34672553 | 0.731 | 0.239 | 7.01E-64 |
| Cardiomyocyte | Mreg | 7 | 1.38E-66 | 0.54702721 | 0.791 | 0.342 | 2.76E-63 |
| Cardiomyocyte | Fmn1 | 7 | 2.94E-66 | 0.40840402 | 0.865 | 0.368 | 5.87E-63 |
| Cardiomyocyte | Fgfr2 | 7 | 3.00E-66 | 1.76414354 | 0.693 | 0.234 | 6.00E-63 |
| Cardiomyocyte | Myo3b1 | 7 | 5.61E-66 | 0.50238055 | 0.8 | 0.366 | 1.12E-62 |
| Cardiomyocyte | Ppp1r9a | 7 | 1.25E-64 | 0.67609881 | 0.935 | 0.381 | 2.50E-61 |
| Cardiomyocyte | Fa2h | 7 | 1.03E-63 | 1.76373606 | 0.623 | 0.133 | 2.06E-60 |
| Cardiomyocyte | Ano51 | 7 | 2.75E-62 | 0.87560517 | 0.98 | 0.485 | 5.50E-59 |

| | | | | | | | |
|---|---|---|---|---|---|---|---|
| Cardiomyocyte | S100b | 7 | 7.56E-61 | 1.11347558 | 0.374 | 0.084 | 1.51E-57 |
| Cardiomyocyte | Cdca2 | 7 | 3.53E-58 | 1.25334981 | 0.354 | 0.056 | 7.06E-55 |
| Cardiomyocyte | Dupd1 | 7 | 3.83E-58 | 1.2743024 | 0.554 | 0.203 | 7.66E-55 |
| Cardiomyocyte | Slc44a5 | 7 | 4.25E-57 | 0.36469912 | 0.736 | 0.268 | 8.50E-54 |
| Cardiomyocyte | Nrxn3 | 7 | 3.67E-56 | 1.07241923 | 0.648 | 0.238 | 7.35E-53 |
| Cardiomyocyte | Il33 | 7 | 3.72E-56 | 0.32555464 | 0.626 | 0.175 | 7.44E-53 |
| Cardiomyocyte | Hpd2 | 7 | 1.10E-53 | 0.50780337 | 0.623 | 0.319 | 2.20E-50 |
| Cardiomyocyte | Lamc2 | 7 | 2.01E-53 | 0.46150365 | 0.948 | 0.46 | 4.03E-50 |
| Cardiomyocyte | Eef1a2 | 7 | 2.26E-53 | 0.5227374 | 0.808 | 0.359 | 4.51E-50 |
| Cardiomyocyte | Parpbp | 7 | 4.61E-52 | 1.47680096 | 0.434 | 0.1 | 9.22E-49 |
| Cardiomyocyte | Nuf2 | 7 | 4.86E-51 | 0.90343344 | 0.519 | 0.137 | 9.72E-48 |
| Cardiomyocyte | Chst15 | 7 | 5.50E-51 | 0.82230673 | 0.673 | 0.343 | 1.10E-47 |
| Cardiomyocyte | Ppm1e | 7 | 4.29E-49 | 0.75206927 | 0.87 | 0.417 | 8.57E-46 |
| Cardiomyocyte | Ajap1 | 7 | 5.35E-45 | 1.95700783 | 0.304 | 0.053 | 1.07E-41 |
| Cardiomyocyte | Matn3 | 7 | 5.82E-43 | 0.55409412 | 0.484 | 0.176 | 1.16E-39 |
| Cardiomyocyte | Ptgfr | 7 | 7.10E-41 | 0.34640959 | 0.815 | 0.41 | 1.42E-37 |
| Cardiomyocyte | Aspm | 7 | 1.78E-40 | 1.06729589 | 0.608 | 0.203 | 3.57E-37 |
| Cardiomyocyte | Acta1 | 7 | 2.12E-40 | 0.37393743 | 0.978 | 0.512 | 4.24E-37 |
| Cardiomyocyte | Fam189a2 | 7 | 4.52E-40 | 0.27123753 | 0.91 | 0.448 | 9.04E-37 |
| Cardiomyocyte | Npr3 | 7 | 7.84E-40 | 0.34648019 | 0.673 | 0.381 | 1.57E-36 |
| Cardiomyocyte | Kcnj51 | 7 | 1.60E-38 | 0.3494787 | 0.781 | 0.427 | 3.20E-35 |
| Cardiomyocyte | Actn1 | 7 | 3.68E-38 | 0.26691131 | 0.94 | 0.467 | 7.36E-35 |
| Cardiomyocyte | Uqcrq | 7 | 3.75E-36 | 0.26800261 | 0.805 | 0.424 | 7.50E-33 |
| Cardiomyocyte | Bdnf | 7 | 2.65E-34 | 0.4621877 | 0.97 | 0.478 | 5.30E-31 |
| Cardiomyocyte | Thbs4 | 7 | 4.27E-34 | 0.55002195 | 0.671 | 0.285 | 8.55E-31 |
| Cardiomyocyte | Ky1 | 7 | 1.61E-33 | 0.75575201 | 0.993 | 0.62 | 3.22E-30 |
| Cardiomyocyte | Fbxo40 | 7 | 1.94E-33 | 0.50694669 | 0.988 | 0.596 | 3.88E-30 |
| Cardiomyocyte | Adarb2 | 7 | 2.40E-33 | 1.77751052 | 0.479 | 0.142 | 4.80E-30 |
| Cardiomyocyte | Flnc | 7 | 4.64E-33 | 0.33088834 | 0.973 | 0.549 | 9.28E-30 |
| Cardiomyocyte | Cnnm2 | 7 | 5.89E-33 | 0.28306597 | 0.693 | 0.408 | 1.18E-29 |
| Cardiomyocyte | Asb15 | 7 | 4.96E-32 | 0.81872969 | 0.985 | 0.595 | 9.91E-29 |

| | | | | | | | |
|---|---|---|---|---|---|---|---|
| Cardiomyocyte | Mlf1 | 7 | 1.23E-30 | 0.75517537 | 0.98 | 0.566 | 2.46E-27 |
| Cardiomyocyte | Tmem116 | 7 | 2.06E-30 | 0.57710936 | 0.965 | 0.564 | 4.12E-27 |
| Cardiomyocyte | Asb91 | 7 | 1.07E-29 | 0.25708153 | 0.965 | 0.514 | 2.14E-26 |
| Cardiomyocyte | Ros11 | 7 | 2.81E-29 | 0.6012171 | 0.98 | 0.596 | 5.62E-26 |
| Cardiomyocyte | Trim50 | 7 | 2.52E-28 | 0.7432222 | 0.988 | 0.624 | 5.03E-25 |
| Cardiomyocyte | Jph1 | 7 | 2.59E-28 | 0.49881048 | 0.943 | 0.499 | 5.19E-25 |
| Cardiomyocyte | Nlgn1 | 7 | 1.71E-27 | 1.52266125 | 0.589 | 0.286 | 3.43E-24 |
| Cardiomyocyte | Cldn15 | 7 | 2.83E-27 | 1.45089549 | 0.551 | 0.2 | 5.65E-24 |
| Cardiomyocyte | Nfatc2 | 7 | 6.38E-27 | 0.3246595 | 0.958 | 0.509 | 1.28E-23 |
| Cardiomyocyte | Adhfe1 | 7 | 6.76E-27 | 0.79207552 | 0.98 | 0.622 | 1.35E-23 |
| Cardiomyocyte | Prox1 | 7 | 7.51E-27 | 0.43833219 | 0.943 | 0.514 | 1.50E-23 |
| Cardiomyocyte | Glb1l2 | 7 | 1.24E-26 | 0.88882816 | 0.988 | 0.642 | 2.49E-23 |
| Cardiomyocyte | Dsg21 | 7 | 1.36E-26 | 0.43722245 | 0.945 | 0.504 | 2.73E-23 |
| Cardiomyocyte | Gbp1 | 7 | 4.82E-26 | 0.30843443 | 0.938 | 0.49 | 9.64E-23 |
| Cardiomyocyte | Cluh | 7 | 1.23E-25 | 0.84706841 | 0.99 | 0.725 | 2.47E-22 |
| Cardiomyocyte | Rgs6 | 7 | 1.75E-25 | 0.67837574 | 0.993 | 0.617 | 3.51E-22 |
| Cardiomyocyte | Coq8a | 7 | 3.14E-25 | 0.75689089 | 0.97 | 0.56 | 6.29E-22 |
| Cardiomyocyte | Frmd5 | 7 | 5.88E-25 | 0.47272514 | 0.918 | 0.539 | 1.18E-21 |
| Cardiomyocyte | Pank1 | 7 | 9.38E-25 | 0.6498535 | 0.97 | 0.561 | 1.88E-21 |
| Cardiomyocyte | Hspb6 | 7 | 2.22E-24 | 0.48634205 | 0.975 | 0.549 | 4.44E-21 |
| Cardiomyocyte | Atp5mg | 7 | 3.39E-24 | 0.33746433 | 0.908 | 0.498 | 6.78E-21 |
| Cardiomyocyte | Trim7 | 7 | 9.27E-23 | 0.31110009 | 0.978 | 0.591 | 1.85E-19 |
| Cardiomyocyte | Iqgap2 | 7 | 3.11E-22 | 0.26581743 | 0.786 | 0.481 | 6.21E-19 |
| Cardiomyocyte | Abcc4 | 7 | 3.63E-22 | 0.4197701 | 0.97 | 0.572 | 7.27E-19 |
| Cardiomyocyte | Efhd1 | 7 | 9.99E-22 | 0.39720597 | 0.965 | 0.54 | 2.00E-18 |
| Cardiomyocyte | Pip5k1b | 7 | 1.60E-21 | 0.2756269 | 0.983 | 0.581 | 3.21E-18 |
| Cardiomyocyte | Mtus2 | 7 | 1.90E-21 | 0.33077712 | 0.913 | 0.523 | 3.81E-18 |
| Cardiomyocyte | Abcc8 | 7 | 1.26E-20 | 0.4984721 | 0.975 | 0.608 | 2.51E-17 |
| Cardiomyocyte | Ripor2 | 7 | 3.37E-20 | 0.49882451 | 0.958 | 0.573 | 6.74E-17 |
| Cardiomyocyte | Rtn4rl1 | 7 | 1.14E-19 | 0.42302848 | 0.803 | 0.518 | 2.28E-16 |
| Cardiomyocyte | AABR070070 | 7 | 1.20E-19 | 0.49788995 | 0.915 | 0.538 | 2.40E-16 |

| | | | | | | | |
|---|---|---|---|---|---|---|---|
| Cardiomyocyte | Fgf12 | 7 | 1.28E-19 | 0.54559932 | 0.953 | 0.567 | 2.56E-16 |
| Cardiomyocyte | Atp1b1 | 7 | 1.88E-19 | 0.37726532 | 0.965 | 0.561 | 3.76E-16 |
| Cardiomyocyte | Ablim2 | 7 | 2.32E-19 | 0.53212028 | 0.828 | 0.537 | 4.64E-16 |
| Cardiomyocyte | Gramd1b | 7 | 4.73E-19 | 0.45875915 | 0.766 | 0.511 | 9.46E-16 |
| Cardiomyocyte | Cobll1 | 7 | 1.81E-18 | 0.41388228 | 0.88 | 0.549 | 3.62E-15 |
| Cardiomyocyte | Slc25a4 | 7 | 2.70E-18 | 0.62874715 | 0.99 | 0.737 | 5.39E-15 |
| Cardiomyocyte | Ckm | 7 | 2.53E-17 | 0.64724354 | 0.935 | 0.68 | 5.06E-14 |
| Cardiomyocyte | Lrrc4b | 7 | 2.63E-17 | 0.44093149 | 0.923 | 0.574 | 5.26E-14 |
| Cardiomyocyte | Unc45b | 7 | 4.53E-17 | 0.69711084 | 0.985 | 0.706 | 9.06E-14 |
| Cardiomyocyte | Angpt1 | 7 | 7.59E-17 | 0.28575219 | 0.9 | 0.558 | 1.52E-13 |
| Cardiomyocyte | Arhgap44 | 7 | 8.30E-17 | 0.55536608 | 0.983 | 0.635 | 1.66E-13 |
| Cardiomyocyte | Kcnk3 | 7 | 6.81E-16 | 0.27391015 | 0.873 | 0.586 | 1.36E-12 |
| Cardiomyocyte | Srpk3 | 7 | 7.73E-16 | 0.64423871 | 0.88 | 0.627 | 1.55E-12 |
| Cardiomyocyte | Fgf13 | 7 | 9.12E-16 | 0.47352871 | 0.9 | 0.604 | 1.82E-12 |
| Cardiomyocyte | Ankrd9 | 7 | 1.48E-15 | 0.57047016 | 0.878 | 0.6 | 2.96E-12 |
| Cardiomyocyte | Acot11 | 7 | 2.21E-15 | 0.40587196 | 0.99 | 0.637 | 4.42E-12 |
| Cardiomyocyte | AC110709.2 | 7 | 8.65E-15 | 0.29694883 | 0.905 | 0.552 | 1.73E-11 |
| Cardiomyocyte | Pygm | 7 | 1.05E-14 | 0.61494604 | 0.943 | 0.669 | 2.09E-11 |
| Cardiomyocyte | Acad10 | 7 | 1.31E-14 | 0.47317941 | 0.94 | 0.595 | 2.61E-11 |
| Cardiomyocyte | Pfkfb2 | 7 | 1.86E-14 | 0.70427784 | 0.97 | 0.7 | 3.71E-11 |
| Cardiomyocyte | AABR070525 | 7 | 3.63E-13 | 0.51873038 | 0.953 | 0.585 | 7.26E-10 |
| Cardiomyocyte | Sfxn51 | 7 | 3.55E-12 | 0.44282278 | 0.893 | 0.599 | 7.10E-09 |
| Cardiomyocyte | Sptb | 7 | 9.47E-12 | 0.5372015 | 0.995 | 0.717 | 1.89E-08 |
| Cardiomyocyte | Rbm24 | 7 | 1.35E-11 | 0.51353988 | 0.983 | 0.718 | 2.70E-08 |
| Cardiomyocyte | Adra1b | 7 | 1.68E-11 | 0.43157608 | 0.97 | 0.631 | 3.37E-08 |
| Cardiomyocyte | Des | 7 | 1.80E-11 | 0.50005831 | 0.985 | 0.725 | 3.60E-08 |
| Cardiomyocyte | Spink8 | 7 | 5.48E-11 | 0.43040705 | 0.98 | 0.67 | 1.10E-07 |
| Cardiomyocyte | AABR070265 | 7 | 2.07E-10 | 0.37137214 | 0.978 | 0.668 | 4.14E-07 |
| Cardiomyocyte | Usp21 | 7 | 2.96E-10 | 0.45358036 | 0.978 | 0.687 | 5.93E-07 |
| Cardiomyocyte | Kcng2 | 7 | 3.01E-10 | 0.55155274 | 0.983 | 0.7 | 6.01E-07 |
| Cardiomyocyte | Slc25a13 | 7 | 4.40E-10 | 0.46553169 | 0.99 | 0.671 | 8.80E-07 |

| | | | | | | | |
|---|---|---|---|---|---|---|---|
| Cardiomyocyte | Corin1 | 7 | 7.84E-10 | 0.32174596 | 0.968 | 0.649 | 1.57E-06 |
| Cardiomyocyte | Ndrg4 | 7 | 1.58E-09 | 0.52009298 | 0.99 | 0.715 | 3.16E-06 |
| Cardiomyocyte | Cpe1 | 7 | 3.18E-09 | 0.3783383 | 0.97 | 0.667 | 6.37E-06 |
| Cardiomyocyte | Alpk2 | 7 | 3.57E-09 | 0.34116594 | 0.968 | 0.658 | 7.14E-06 |
| Cardiomyocyte | Gal3st3 | 7 | 5.46E-09 | 0.25143975 | 0.98 | 0.631 | 1.09E-05 |
| Cardiomyocyte | Lgr6 | 7 | 2.09E-08 | 0.47966758 | 0.978 | 0.68 | 4.18E-05 |
| Cardiomyocyte | Pkia | 7 | 3.65E-08 | 0.32026 | 0.95 | 0.657 | 7.30E-05 |
| Cardiomyocyte | Myl6 | 7 | 6.32E-08 | 0.43755476 | 0.973 | 0.698 | 0.00012641 |
| Cardiomyocyte | Dpf3 | 7 | 6.69E-08 | 0.31787719 | 0.928 | 0.649 | 0.00013379 |
| Cardiomyocyte | Ntn1 | 7 | 1.96E-07 | 0.28301871 | 0.96 | 0.623 | 0.00039272 |
| Cardiomyocyte | Pcsk6 | 7 | 2.12E-07 | 0.2541417 | 0.968 | 0.677 | 0.00042492 |
| Cardiomyocyte | Art3 | 7 | 2.04E-06 | 0.306902 | 0.978 | 0.7 | 0.00407846 |
| Cardiomyocyte | Disp2 | 8 | 5.50E-158 | 3.86285656 | 0.291 | 0.006 | 1.10E-154 |
| Cardiomyocyte | Slpi | 8 | 1.81E-134 | 3.59133823 | 0.324 | 0.009 | 3.61E-131 |
| Cardiomyocyte | Ccl19 | 8 | 3.48E-121 | 1.64633541 | 0.413 | 0.029 | 6.96E-118 |
| Cardiomyocyte | Clic6 | 8 | 8.47E-99 | 3.98222127 | 0.413 | 0.05 | 1.69E-95 |
| Cardiomyocyte | Hbb | 8 | 2.14E-79 | 2.2479147 | 0.368 | 0.059 | 4.28E-76 |
| Cardiomyocyte | Ska1 | 8 | 1.23E-72 | 1.15206255 | 0.34 | 0.074 | 2.46E-69 |
| Cardiomyocyte | AC128394.2 | 8 | 1.91E-62 | 3.16458161 | 0.421 | 0.053 | 3.83E-59 |
| Cardiomyocyte | AABR070687 | 8 | 1.09E-59 | 1.49591376 | 0.356 | 0.029 | 2.18E-56 |
| Cardiomyocyte | Pak3 | 8 | 2.35E-59 | 2.60027066 | 0.466 | 0.084 | 4.70E-56 |
| Cardiomyocyte | Mgp1 | 8 | 8.68E-56 | 1.95151059 | 0.874 | 0.53 | 1.74E-52 |
| Cardiomyocyte | Col3a11 | 8 | 1.04E-55 | 1.89142273 | 0.927 | 0.671 | 2.09E-52 |
| Cardiomyocyte | Cdh6 | 8 | 7.73E-53 | 0.28260547 | 0.36 | 0.033 | 1.55E-49 |
| Cardiomyocyte | Col1a11 | 8 | 2.51E-52 | 2.07901192 | 0.834 | 0.496 | 5.01E-49 |
| Cardiomyocyte | Cox8b | 8 | 7.36E-51 | 2.13398038 | 0.806 | 0.452 | 1.47E-47 |
| Cardiomyocyte | LOC1083526 | 8 | 1.69E-48 | 2.48930876 | 0.656 | 0.27 | 3.38E-45 |
| Cardiomyocyte | Ndufa4 | 8 | 1.03E-47 | 2.28758899 | 0.781 | 0.355 | 2.05E-44 |
| Cardiomyocyte | Cox6a2 | 8 | 4.16E-43 | 2.22079846 | 0.761 | 0.489 | 8.32E-40 |
| Cardiomyocyte | Tmsb4x | 8 | 7.30E-43 | 2.22521371 | 0.721 | 0.397 | 1.46E-39 |
| Cardiomyocyte | Fth1 | 8 | 3.51E-42 | 2.44194518 | 0.721 | 0.435 | 7.02E-39 |

| | | | | | | | |
|---|---|---|---|---|---|---|---|
| Cardiomyocyte | Tpt1 | 8 | 3.95E-42 | 2.17957852 | 0.709 | 0.399 | 7.90E-39 |
| Cardiomyocyte | Atp5mg1 | 8 | 6.28E-42 | 1.7642995 | 0.81 | 0.516 | 1.26E-38 |
| Cardiomyocyte | Cox4i1 | 8 | 3.80E-40 | 1.57988586 | 0.866 | 0.608 | 7.60E-37 |
| Cardiomyocyte | Atp5mc3 | 8 | 2.90E-39 | 1.98928175 | 0.745 | 0.414 | 5.80E-36 |
| Cardiomyocyte | Atp5me | 8 | 5.08E-39 | 1.74155815 | 0.781 | 0.515 | 1.02E-35 |
| Cardiomyocyte | Sparc1 | 8 | 1.08E-38 | 1.83826072 | 0.769 | 0.503 | 2.17E-35 |
| Cardiomyocyte | Bgn | 8 | 1.97E-38 | 2.44626153 | 0.688 | 0.397 | 3.93E-35 |
| Cardiomyocyte | Atp5f1e1 | 8 | 1.23E-37 | 2.42294821 | 0.623 | 0.313 | 2.47E-34 |
| Cardiomyocyte | Egfl6 | 8 | 4.03E-37 | 2.22743477 | 0.397 | 0.061 | 8.05E-34 |
| Cardiomyocyte | Reg3g | 8 | 3.09E-35 | 1.28579805 | 0.389 | 0.112 | 6.19E-32 |
| Cardiomyocyte | LOC1003644 | 8 | 4.10E-35 | 2.44469997 | 0.599 | 0.211 | 8.20E-32 |
| Cardiomyocyte | Gsn | 8 | 1.12E-34 | 2.11252856 | 0.721 | 0.447 | 2.23E-31 |
| Cardiomyocyte | Igsf11 | 8 | 6.11E-34 | 0.92834768 | 0.478 | 0.16 | 1.22E-30 |
| Cardiomyocyte | LOC687780 | 8 | 5.59E-33 | 2.41797098 | 0.595 | 0.244 | 1.12E-29 |
| Cardiomyocyte | Kank4 | 8 | 1.44E-32 | 0.57517613 | 0.543 | 0.186 | 2.87E-29 |
| Cardiomyocyte | Trem2 | 8 | 7.52E-32 | 1.8204451 | 0.522 | 0.2 | 1.50E-28 |
| Cardiomyocyte | Dcn1 | 8 | 1.04E-31 | 1.41821241 | 0.741 | 0.407 | 2.08E-28 |
| Cardiomyocyte | Cd83 | 8 | 4.58E-31 | 2.14365449 | 0.498 | 0.215 | 9.17E-28 |
| Cardiomyocyte | Uqcr10 | 8 | 1.03E-30 | 2.46980097 | 0.571 | 0.278 | 2.06E-27 |
| Cardiomyocyte | Car101 | 8 | 3.09E-30 | 1.5653235 | 0.413 | 0.118 | 6.18E-27 |
| Cardiomyocyte | Fstl11 | 8 | 2.13E-29 | 1.4528004 | 0.68 | 0.323 | 4.26E-26 |
| Cardiomyocyte | Ppl | 8 | 3.95E-29 | 1.68473556 | 0.453 | 0.152 | 7.89E-26 |
| Cardiomyocyte | Ccl7 | 8 | 5.05E-28 | 0.4889604 | 0.372 | 0.086 | 1.01E-24 |
| Cardiomyocyte | Chl1 | 8 | 5.44E-28 | 0.813282 | 0.409 | 0.103 | 1.09E-24 |
| Cardiomyocyte | Apoe | 8 | 9.11E-28 | 2.41739612 | 0.583 | 0.319 | 1.82E-24 |
| Cardiomyocyte | LOC1001348 | 8 | 1.16E-27 | 2.37934634 | 0.656 | 0.356 | 2.32E-24 |
| Cardiomyocyte | Cox6a1 | 8 | 3.32E-26 | 1.95267815 | 0.656 | 0.343 | 6.64E-23 |
| Cardiomyocyte | Melk | 8 | 4.49E-26 | 1.08285637 | 0.348 | 0.066 | 8.97E-23 |
| Cardiomyocyte | Serpinf1 | 8 | 1.52E-25 | 2.42472778 | 0.595 | 0.249 | 3.04E-22 |
| Cardiomyocyte | Eef1a21 | 8 | 4.72E-24 | 1.76169241 | 0.664 | 0.38 | 9.45E-21 |
| Cardiomyocyte | Mx11 | 8 | 6.02E-24 | 0.28415923 | 0.417 | 0.136 | 1.20E-20 |

| | | | | | | | |
|---|---|---|---|---|---|---|---|
| Cardiomyocyte | Kif22 | 8 | 8.03E-24 | 0.99223831 | 0.389 | 0.109 | 1.61E-20 |
| Cardiomyocyte | RT1-Ba | 8 | 1.06E-23 | 2.5270184 | 0.466 | 0.188 | 2.12E-20 |
| Cardiomyocyte | Npnt | 8 | 5.86E-23 | 0.70302212 | 0.34 | 0.084 | 1.17E-19 |
| Cardiomyocyte | Olfml2a | 8 | 1.80E-22 | 1.80791847 | 0.49 | 0.11 | 3.60E-19 |
| Cardiomyocyte | Stac | 8 | 1.70E-21 | 1.34655338 | 0.441 | 0.153 | 3.39E-18 |
| Cardiomyocyte | Cdh3 | 8 | 8.57E-21 | 1.26734744 | 0.36 | 0.102 | 1.71E-17 |
| Cardiomyocyte | Timp1 | 8 | 9.66E-21 | 2.66924987 | 0.571 | 0.269 | 1.93E-17 |
| Cardiomyocyte | Abca41 | 8 | 1.70E-20 | 0.29978899 | 0.538 | 0.284 | 3.39E-17 |
| Cardiomyocyte | Rasl12 | 8 | 4.20E-20 | 0.68239418 | 0.401 | 0.118 | 8.41E-17 |
| Cardiomyocyte | Ptger3 | 8 | 5.89E-20 | 0.64774366 | 0.502 | 0.221 | 1.18E-16 |
| Cardiomyocyte | Abcd21 | 8 | 1.09E-19 | 1.34385194 | 0.498 | 0.188 | 2.18E-16 |
| Cardiomyocyte | Rspo11 | 8 | 1.49E-19 | 0.99155664 | 0.462 | 0.2 | 2.98E-16 |
| Cardiomyocyte | Pdpn | 8 | 3.04E-18 | 1.51273744 | 0.49 | 0.171 | 6.08E-15 |
| Cardiomyocyte | Gap431 | 8 | 3.24E-16 | 1.36694719 | 0.482 | 0.206 | 6.48E-13 |
| Cardiomyocyte | Cyba | 8 | 4.02E-16 | 1.8372624 | 0.49 | 0.221 | 8.04E-13 |
| Cardiomyocyte | Fras1 | 8 | 4.96E-16 | 0.67706401 | 0.482 | 0.13 | 9.91E-13 |
| Cardiomyocyte | Cotl1 | 8 | 5.71E-15 | 1.49319122 | 0.425 | 0.166 | 1.14E-11 |
| Cardiomyocyte | Nppa1 | 8 | 1.15E-14 | 1.14247674 | 0.628 | 0.37 | 2.29E-11 |
| Cardiomyocyte | AABR070294 | 8 | 2.28E-12 | 1.03799384 | 0.506 | 0.218 | 4.55E-09 |
| Cardiomyocyte | Mnda | 8 | 2.32E-11 | 0.49877391 | 0.518 | 0.225 | 4.63E-08 |
| Cardiomyocyte | Sv2c | 8 | 2.67E-11 | 0.74105536 | 0.518 | 0.219 | 5.35E-08 |
| Cardiomyocyte | Cass4 | 9 | 2.05E-116 | 5.3482919 | 0.356 | 0.045 | 4.11E-113 |
| Cardiomyocyte | Ptprc | 9 | 2.10E-86 | 4.64641279 | 0.871 | 0.327 | 4.20E-83 |
| Cardiomyocyte | Il1rl1 | 9 | 2.90E-85 | 4.05772307 | 0.33 | 0.024 | 5.80E-82 |
| Cardiomyocyte | Itgax | 9 | 8.07E-74 | 4.73388481 | 0.32 | 0.007 | 1.61E-70 |
| Cardiomyocyte | Arhgap15 | 9 | 7.42E-73 | 4.31519731 | 0.835 | 0.325 | 1.48E-69 |
| Cardiomyocyte | Zeb21 | 9 | 9.10E-62 | 2.28462169 | 0.933 | 0.642 | 1.82E-58 |
| Cardiomyocyte | Frmd4b | 9 | 1.33E-55 | 3.40260393 | 0.814 | 0.442 | 2.65E-52 |
| Cardiomyocyte | Dock8 | 9 | 3.98E-55 | 3.44504387 | 0.799 | 0.341 | 7.96E-52 |
| Cardiomyocyte | Tbxas1 | 9 | 7.51E-55 | 5.50457166 | 0.742 | 0.211 | 1.50E-51 |
| Cardiomyocyte | Cd300a | 9 | 1.32E-54 | 5.85076771 | 0.541 | 0.063 | 2.64E-51 |

| | | | | | | | |
|---|---|---|---|---|---|---|---|
| Cardiomyocyte | Fyb1 | 9 | 7.52E-52 | 4.76096329 | 0.722 | 0.223 | 1.50E-48 |
| Cardiomyocyte | Lyn1 | 9 | 4.32E-49 | 2.50004905 | 0.825 | 0.443 | 8.63E-46 |
| Cardiomyocyte | Tlr7 | 9 | 3.72E-48 | 5.40705543 | 0.433 | 0.069 | 7.44E-45 |
| Cardiomyocyte | Cd300lb | 9 | 4.77E-48 | 5.56422015 | 0.304 | 0.037 | 9.54E-45 |
| Cardiomyocyte | Pid11 | 9 | 5.41E-48 | 2.51176206 | 0.83 | 0.403 | 1.08E-44 |
| Cardiomyocyte | Sh3bp2 | 9 | 3.15E-44 | 3.69005459 | 0.742 | 0.409 | 6.30E-41 |
| Cardiomyocyte | Cfh | 9 | 3.15E-44 | 4.0006966 | 0.686 | 0.207 | 6.31E-41 |
| Cardiomyocyte | Gna15 | 9 | 4.06E-42 | 5.57777672 | 0.593 | 0.246 | 8.11E-39 |
| Cardiomyocyte | Aoah | 9 | 6.95E-42 | 5.18029172 | 0.629 | 0.2 | 1.39E-38 |
| Cardiomyocyte | AABR070015 | 9 | 1.23E-41 | 4.59551872 | 0.423 | 0.122 | 2.46E-38 |
| Cardiomyocyte | Inpp5d | 9 | 3.54E-40 | 4.14657949 | 0.706 | 0.347 | 7.07E-37 |
| Cardiomyocyte | Arhgap22 | 9 | 7.24E-40 | 4.54730223 | 0.665 | 0.315 | 1.45E-36 |
| Cardiomyocyte | Ikzf1 | 9 | 1.39E-39 | 5.28984639 | 0.639 | 0.255 | 2.78E-36 |
| Cardiomyocyte | Otulinl | 9 | 2.09E-38 | 5.38051394 | 0.572 | 0.195 | 4.19E-35 |
| Cardiomyocyte | Hk3 | 9 | 2.69E-38 | 4.3230547 | 0.371 | 0.012 | 5.39E-35 |
| Cardiomyocyte | Kcnk13 | 9 | 3.11E-38 | 4.92595855 | 0.577 | 0.191 | 6.23E-35 |
| Cardiomyocyte | Csf1r | 9 | 3.76E-38 | 4.62036814 | 0.675 | 0.324 | 7.52E-35 |
| Cardiomyocyte | Ptprj | 9 | 1.31E-37 | 2.81959369 | 0.732 | 0.264 | 2.62E-34 |
| Cardiomyocyte | Ciita | 9 | 3.86E-37 | 4.06481742 | 0.407 | 0.109 | 7.72E-34 |
| Cardiomyocyte | LOC690045 | 9 | 1.23E-35 | 4.29476649 | 0.665 | 0.318 | 2.46E-32 |
| Cardiomyocyte | Runx1 | 9 | 1.05E-34 | 2.76723492 | 0.711 | 0.395 | 2.11E-31 |
| Cardiomyocyte | Syk | 9 | 2.03E-34 | 4.79999235 | 0.598 | 0.196 | 4.06E-31 |
| Cardiomyocyte | Camk1d | 9 | 2.07E-34 | 3.35130241 | 0.665 | 0.329 | 4.14E-31 |
| Cardiomyocyte | Slc9a9 | 9 | 2.86E-34 | 2.497756 | 0.794 | 0.53 | 5.72E-31 |
| Cardiomyocyte | Nckap1l | 9 | 1.79E-32 | 4.49821729 | 0.479 | 0.165 | 3.58E-29 |
| Cardiomyocyte | Ly86 | 9 | 1.53E-31 | 5.34118722 | 0.577 | 0.289 | 3.06E-28 |
| Cardiomyocyte | Il6r | 9 | 3.36E-30 | 2.23509823 | 0.701 | 0.406 | 6.71E-27 |
| Cardiomyocyte | AABR070311 | 9 | 2.69E-29 | 1.97965858 | 0.289 | 0.023 | 5.38E-26 |
| Cardiomyocyte | Cdh7 | 9 | 2.87E-29 | 2.90884982 | 0.314 | 0.038 | 5.74E-26 |
| Cardiomyocyte | Pou2f2 | 9 | 4.37E-28 | 4.63415553 | 0.454 | 0.196 | 8.75E-25 |
| Cardiomyocyte | Lilrb4 | 9 | 3.56E-26 | 4.55283923 | 0.562 | 0.164 | 7.12E-23 |

| | | | | | | | |
|---|---|---|---|---|---|---|---|
| Cardiomyocyte | Plek | 9 | 1.20E-25 | 4.14082008 | 0.32 | 0.054 | 2.40E-22 |
| Cardiomyocyte | Mrc1 | 9 | 2.15E-25 | 4.20359061 | 0.562 | 0.255 | 4.30E-22 |
| Cardiomyocyte | Ptafr1 | 9 | 3.12E-25 | 4.56398266 | 0.5 | 0.108 | 6.25E-22 |
| Cardiomyocyte | Cd84 | 9 | 3.29E-25 | 5.10759198 | 0.381 | 0.057 | 6.58E-22 |
| Cardiomyocyte | Myo1f | 9 | 3.66E-25 | 3.72761124 | 0.598 | 0.233 | 7.32E-22 |
| Cardiomyocyte | Fli11 | 9 | 2.75E-24 | 1.33072055 | 0.706 | 0.362 | 5.50E-21 |
| Cardiomyocyte | Cd68 | 9 | 1.03E-23 | 2.81647614 | 0.412 | 0.123 | 2.07E-20 |
| Cardiomyocyte | Fermt31 | 9 | 1.35E-23 | 4.19781326 | 0.572 | 0.243 | 2.69E-20 |
| Cardiomyocyte | Abca11 | 9 | 1.47E-23 | 2.12642955 | 0.686 | 0.375 | 2.95E-20 |
| Cardiomyocyte | Itgb2 | 9 | 3.16E-23 | 4.43134317 | 0.412 | 0.096 | 6.32E-20 |
| Cardiomyocyte | Colec121 | 9 | 1.26E-22 | 2.64469551 | 0.634 | 0.309 | 2.52E-19 |
| Cardiomyocyte | Slc4a5 | 9 | 2.90E-22 | 4.60020536 | 0.304 | 0.053 | 5.79E-19 |
| Cardiomyocyte | Pik3r5 | 9 | 9.01E-22 | 5.02911495 | 0.402 | 0.131 | 1.80E-18 |
| Cardiomyocyte | Apba1 | 9 | 1.23E-21 | 3.31119276 | 0.598 | 0.291 | 2.47E-18 |
| Cardiomyocyte | Cd4 | 9 | 1.77E-21 | 4.95791163 | 0.521 | 0.238 | 3.54E-18 |
| Cardiomyocyte | Slco2b1 | 9 | 2.60E-21 | 2.89799418 | 0.603 | 0.274 | 5.20E-18 |
| Cardiomyocyte | Lpcat2 | 9 | 1.23E-20 | 4.0285158 | 0.521 | 0.214 | 2.47E-17 |
| Cardiomyocyte | Chn2 | 9 | 1.58E-20 | 2.24385263 | 0.582 | 0.16 | 3.16E-17 |
| Cardiomyocyte | Hck | 9 | 1.68E-18 | 4.49920614 | 0.464 | 0.209 | 3.36E-15 |
| Cardiomyocyte | Mertk1 | 9 | 4.48E-18 | 3.64953171 | 0.536 | 0.19 | 8.96E-15 |
| Cardiomyocyte | Tmcc3 | 9 | 7.33E-18 | 1.90921622 | 0.675 | 0.404 | 1.47E-14 |
| Cardiomyocyte | Lcp2 | 9 | 8.30E-18 | 3.96613883 | 0.469 | 0.204 | 1.66E-14 |
| Cardiomyocyte | Ptpro | 9 | 6.42E-17 | 5.10886314 | 0.474 | 0.22 | 1.28E-13 |
| Cardiomyocyte | Cybb | 9 | 3.60E-16 | 3.93690216 | 0.495 | 0.233 | 7.19E-13 |
| Cardiomyocyte | Pltp | 9 | 6.12E-15 | 3.31632601 | 0.546 | 0.295 | 1.22E-11 |
| Cardiomyocyte | Entpd11 | 9 | 1.52E-13 | 2.37929859 | 0.541 | 0.274 | 3.03E-10 |
| Cardiomyocyte | Ltc4s | 9 | 1.32E-12 | 5.06717218 | 0.392 | 0.049 | 2.65E-09 |
| Cardiomyocyte | Mbp | 9 | 2.45E-12 | 3.26096544 | 0.402 | 0.142 | 4.90E-09 |
| Cardiomyocyte | Atp8b4 | 9 | 3.61E-12 | 4.59502479 | 0.438 | 0.123 | 7.22E-09 |
| Cardiomyocyte | Mctp2 | 9 | 1.26E-11 | 3.20555704 | 0.485 | 0.228 | 2.53E-08 |
| Cardiomyocyte | Ston21 | 9 | 3.73E-11 | 1.32050226 | 0.562 | 0.26 | 7.45E-08 |

| | | | | | | | |
|---|---|---|---|---|---|---|---|
| Cardiomyocyte | Ptk2b | 9 | 2.23E-10 | 2.48267222 | 0.51 | 0.235 | 4.46E-07 |
| Cardiomyocyte | AABR070280 | 10 | 0 | 7.03502253 | 0.374 | 0.001 | 0 |
| Cardiomyocyte | Ccr6 | 10 | 2.92E-161 | 2.64411115 | 0.259 | 0.005 | 5.85E-158 |
| Cardiomyocyte | Art2b | 10 | 4.33E-159 | 2.84285208 | 0.36 | 0.013 | 8.66E-156 |
| Cardiomyocyte | Cpvl | 10 | 1.45E-77 | 3.99300178 | 0.388 | 0.024 | 2.90E-74 |
| Cardiomyocyte | Gucy1a2 | 10 | 2.40E-67 | 4.31966208 | 0.906 | 0.298 | 4.79E-64 |
| Cardiomyocyte | Pdgfrb1 | 10 | 4.65E-67 | 3.37909163 | 0.942 | 0.28 | 9.30E-64 |
| Cardiomyocyte | Mrvi1 | 10 | 2.55E-63 | 4.55667693 | 0.878 | 0.31 | 5.10E-60 |
| Cardiomyocyte | Cass41 | 10 | 2.84E-63 | 2.08822783 | 0.388 | 0.048 | 5.67E-60 |
| Cardiomyocyte | Anxa8 | 10 | 6.92E-63 | 5.01507381 | 0.288 | 0.027 | 1.38E-59 |
| Cardiomyocyte | Sell | 10 | 5.19E-61 | 1.54050165 | 0.309 | 0.006 | 1.04E-57 |
| Cardiomyocyte | Notch3 | 10 | 6.66E-60 | 5.75199061 | 0.827 | 0.114 | 1.33E-56 |
| Cardiomyocyte | Adgrl3 | 10 | 1.05E-55 | 4.62690445 | 0.827 | 0.257 | 2.10E-52 |
| Cardiomyocyte | Rerg | 10 | 4.50E-53 | 4.47388572 | 0.835 | 0.338 | 8.99E-50 |
| Cardiomyocyte | Rgs5 | 10 | 1.91E-50 | 5.34427406 | 0.784 | 0.345 | 3.82E-47 |
| Cardiomyocyte | Trpc6 | 10 | 1.36E-47 | 4.48576112 | 0.568 | 0.132 | 2.71E-44 |
| Cardiomyocyte | Gucy1a11 | 10 | 4.49E-46 | 3.58103746 | 0.799 | 0.216 | 8.98E-43 |
| Cardiomyocyte | Ano1 | 10 | 7.74E-45 | 5.4603459 | 0.662 | 0.197 | 1.55E-41 |
| Cardiomyocyte | Zeb22 | 10 | 8.46E-44 | 2.30617143 | 0.942 | 0.645 | 1.69E-40 |
| Cardiomyocyte | Cdh61 | 10 | 4.48E-43 | 5.20185632 | 0.295 | 0.042 | 8.97E-40 |
| Cardiomyocyte | Apcdd1l | 10 | 2.75E-42 | 2.4338741 | 0.396 | 0.028 | 5.49E-39 |
| Cardiomyocyte | Zfp385d1 | 10 | 1.51E-41 | 2.09423209 | 0.885 | 0.343 | 3.02E-38 |
| Cardiomyocyte | Egflam | 10 | 7.68E-41 | 3.99916383 | 0.748 | 0.273 | 1.54E-37 |
| Cardiomyocyte | AABR070042 | 10 | 1.23E-38 | 5.07078352 | 0.561 | 0.099 | 2.47E-35 |
| Cardiomyocyte | Mylk | 10 | 3.67E-38 | 2.78545642 | 0.842 | 0.468 | 7.33E-35 |
| Cardiomyocyte | Il34 | 10 | 3.35E-37 | 5.62548199 | 0.59 | 0.123 | 6.70E-34 |
| Cardiomyocyte | Il18r1 | 10 | 9.37E-37 | 2.87599008 | 0.453 | 0.048 | 1.87E-33 |
| Cardiomyocyte | Agap2 | 10 | 1.65E-36 | 5.67565631 | 0.532 | 0.063 | 3.29E-33 |
| Cardiomyocyte | Atp1a3 | 10 | 2.76E-36 | 2.77248291 | 0.388 | 0.011 | 5.51E-33 |
| Cardiomyocyte | Ppfia2 | 10 | 5.13E-36 | 3.597208 | 0.345 | 0.026 | 1.03E-32 |
| Cardiomyocyte | Rasl121 | 10 | 6.35E-36 | 5.59800106 | 0.647 | 0.118 | 1.27E-32 |

| | | | | | | | |
|---|---|---|---|---|---|---|---|
| Cardiomyocyte | Trpc4 | 10 | 3.59E-33 | 6.70556731 | 0.496 | 0.027 | 7.17E-30 |
| Cardiomyocyte | Ebf21 | 10 | 7.42E-32 | 2.1339011 | 0.755 | 0.266 | 1.48E-28 |
| Cardiomyocyte | Neil31 | 10 | 2.56E-31 | 2.79917336 | 0.396 | 0.047 | 5.12E-28 |
| Cardiomyocyte | Cald1 | 10 | 2.78E-31 | 2.25248593 | 0.835 | 0.482 | 5.55E-28 |
| Cardiomyocyte | Msln | 10 | 1.01E-30 | 0.81992285 | 0.417 | 0.039 | 2.01E-27 |
| Cardiomyocyte | Sgip1 | 10 | 1.45E-30 | 4.19422035 | 0.683 | 0.255 | 2.90E-27 |
| Cardiomyocyte | Galnt141 | 10 | 6.02E-30 | 1.48472967 | 0.381 | 0.062 | 1.20E-26 |
| Cardiomyocyte | Plxdc21 | 10 | 6.48E-30 | 1.97316266 | 0.82 | 0.397 | 1.30E-26 |
| Cardiomyocyte | Pde8b | 10 | 2.16E-29 | 4.946938 | 0.662 | 0.284 | 4.31E-26 |
| Cardiomyocyte | Pawr | 10 | 2.26E-29 | 3.34273813 | 0.698 | 0.167 | 4.53E-26 |
| Cardiomyocyte | Hpgd1 | 10 | 1.54E-27 | 4.66377643 | 0.482 | 0.106 | 3.08E-24 |
| Cardiomyocyte | Olr59 | 10 | 5.54E-27 | 5.24208233 | 0.568 | 0.167 | 1.11E-23 |
| Cardiomyocyte | Fgr1 | 10 | 1.67E-26 | 1.06996676 | 0.432 | 0.126 | 3.35E-23 |
| Cardiomyocyte | Ldb2 | 10 | 5.51E-26 | 2.7766034 | 0.64 | 0.221 | 1.10E-22 |
| Cardiomyocyte | Mark1 | 10 | 7.05E-26 | 3.91066516 | 0.655 | 0.263 | 1.41E-22 |
| Cardiomyocyte | Ntng1 | 10 | 1.07E-25 | 5.1123576 | 0.633 | 0.272 | 2.14E-22 |
| Cardiomyocyte | Camk4 | 10 | 9.39E-25 | 0.72516549 | 0.489 | 0.14 | 1.88E-21 |
| Cardiomyocyte | Ngf | 10 | 1.69E-24 | 2.62513422 | 0.676 | 0.259 | 3.39E-21 |
| Cardiomyocyte | Pde5a | 10 | 3.67E-24 | 3.21527169 | 0.64 | 0.262 | 7.34E-21 |
| Cardiomyocyte | Kcnh1 | 10 | 1.01E-22 | 1.37813457 | 0.417 | 0.117 | 2.02E-19 |
| Cardiomyocyte | Med12l | 10 | 7.64E-22 | 3.44572527 | 0.597 | 0.205 | 1.53E-18 |
| Cardiomyocyte | Kcnj8 | 10 | 8.43E-22 | 4.96279431 | 0.561 | 0.138 | 1.69E-18 |
| Cardiomyocyte | Specc1 | 10 | 1.91E-21 | 2.26639944 | 0.698 | 0.293 | 3.82E-18 |
| Cardiomyocyte | P2ry14 | 10 | 2.40E-21 | 4.98360353 | 0.554 | 0.193 | 4.81E-18 |
| Cardiomyocyte | Pid12 | 10 | 5.25E-21 | 2.03118732 | 0.755 | 0.41 | 1.05E-17 |
| Cardiomyocyte | Agmo | 10 | 1.89E-20 | 4.11840106 | 0.597 | 0.326 | 3.77E-17 |
| Cardiomyocyte | Prrx11 | 10 | 4.63E-20 | 2.13952294 | 0.683 | 0.267 | 9.27E-17 |
| Cardiomyocyte | Ptch22 | 10 | 1.70E-19 | 2.61877207 | 0.381 | 0.107 | 3.39E-16 |
| Cardiomyocyte | Igf1 | 10 | 2.20E-19 | 1.6918534 | 0.712 | 0.288 | 4.40E-16 |
| Cardiomyocyte | Myh11 | 10 | 9.61E-19 | 5.06720395 | 0.619 | 0.218 | 1.92E-15 |
| Cardiomyocyte | Cspg4 | 10 | 1.10E-18 | 2.04638587 | 0.669 | 0.329 | 2.20E-15 |

| | | | | | | | |
|---|---|---|---|---|---|---|---|
| Cardiomyocyte | Grin2b | 10 | 2.20E-18 | 1.30760096 | 0.403 | 0.087 | 4.41E-15 |
| Cardiomyocyte | Rassf4 | 10 | 2.64E-18 | 1.39693289 | 0.446 | 0.07 | 5.28E-15 |
| Cardiomyocyte | Cd300a1 | 10 | 2.79E-18 | 0.85105506 | 0.424 | 0.071 | 5.58E-15 |
| Cardiomyocyte | Plek1 | 10 | 3.89E-18 | 1.08303012 | 0.331 | 0.056 | 7.77E-15 |
| Cardiomyocyte | Adcy2 | 10 | 4.33E-18 | 2.32025395 | 0.424 | 0.169 | 8.66E-15 |
| Cardiomyocyte | Trpc3 | 10 | 4.85E-18 | 5.27754769 | 0.46 | 0.125 | 9.70E-15 |
| Cardiomyocyte | Chn11 | 10 | 8.50E-18 | 2.42642953 | 0.626 | 0.298 | 1.70E-14 |
| Cardiomyocyte | Bub1b | 10 | 1.37E-17 | 1.72891757 | 0.417 | 0.082 | 2.73E-14 |
| Cardiomyocyte | Erbb3 | 10 | 2.40E-17 | 1.40698019 | 0.317 | 0.033 | 4.79E-14 |
| Cardiomyocyte | LOC1025530 | 10 | 1.88E-16 | 2.616172 | 0.518 | 0.154 | 3.76E-13 |
| Cardiomyocyte | Lrrc4c1 | 10 | 6.09E-16 | 3.06726351 | 0.525 | 0.266 | 1.22E-12 |
| Cardiomyocyte | Sele1 | 10 | 7.34E-16 | 3.66067425 | 0.482 | 0.231 | 1.47E-12 |
| Cardiomyocyte | Gcnt21 | 10 | 1.06E-15 | 2.579841 | 0.59 | 0.276 | 2.12E-12 |
| Cardiomyocyte | Emid11 | 10 | 1.43E-15 | 2.85007501 | 0.432 | 0.137 | 2.87E-12 |
| Cardiomyocyte | Aatk | 10 | 1.46E-15 | 3.53025511 | 0.547 | 0.189 | 2.91E-12 |
| Cardiomyocyte | Adamts21 | 10 | 1.62E-15 | 1.52212716 | 0.662 | 0.366 | 3.24E-12 |
| Cardiomyocyte | Dach11 | 10 | 3.06E-15 | 1.36355307 | 0.698 | 0.313 | 6.12E-12 |
| Cardiomyocyte | Errfi11 | 10 | 4.04E-15 | 2.32800379 | 0.604 | 0.242 | 8.08E-12 |
| Cardiomyocyte | Ctnnd2 | 10 | 5.90E-15 | 2.41016668 | 0.338 | 0.078 | 1.18E-11 |
| Cardiomyocyte | Kif20a1 | 10 | 7.00E-15 | 3.07276557 | 0.381 | 0.064 | 1.40E-11 |
| Cardiomyocyte | Sparcl1 | 10 | 2.31E-14 | 1.76978479 | 0.511 | 0.23 | 4.61E-11 |
| Cardiomyocyte | Cd841 | 10 | 2.32E-14 | 0.80780859 | 0.417 | 0.06 | 4.64E-11 |
| Cardiomyocyte | Actn11 | 10 | 2.58E-14 | 1.50703063 | 0.763 | 0.497 | 5.16E-11 |
| Cardiomyocyte | Mcam1 | 10 | 2.33E-13 | 2.34474382 | 0.511 | 0.261 | 4.66E-10 |
| Cardiomyocyte | Meox21 | 10 | 6.28E-13 | 1.42335615 | 0.64 | 0.282 | 1.26E-09 |
| Cardiomyocyte | Slc24a3 | 10 | 8.14E-13 | 2.17931683 | 0.561 | 0.226 | 1.63E-09 |
| Cardiomyocyte | AABR070275 | 10 | 9.42E-13 | 2.55407469 | 0.554 | 0.27 | 1.88E-09 |
| Cardiomyocyte | Bcar3 | 10 | 2.49E-12 | 1.36901425 | 0.662 | 0.352 | 4.98E-09 |
| Cardiomyocyte | Map3k7cl | 10 | 3.15E-12 | 3.79826415 | 0.324 | 0.064 | 6.30E-09 |
| Cardiomyocyte | Colgalt2 | 10 | 3.91E-12 | 1.48398455 | 0.633 | 0.38 | 7.82E-09 |
| Cardiomyocyte | Cyp7b1 | 10 | 5.48E-12 | 2.43879819 | 0.532 | 0.212 | 1.10E-08 |

| | | | | | | | |
|---|---|---|---|---|---|---|---|
| Cardiomyocyte | Trpm3 | 10 | 1.25E-11 | 2.42499785 | 0.46 | 0.149 | 2.50E-08 |
| Cardiomyocyte | Egfr1 | 10 | 2.54E-11 | 1.09772061 | 0.64 | 0.362 | 5.08E-08 |
| Cardiomyocyte | Ccdc801 | 10 | 3.29E-11 | 0.91753144 | 0.748 | 0.479 | 6.59E-08 |
| Cardiomyocyte | Map1b1 | 10 | 5.61E-11 | 0.84965673 | 0.755 | 0.497 | 1.12E-07 |
| Cardiomyocyte | Cdh31 | 10 | 1.22E-10 | 2.46265333 | 0.396 | 0.107 | 2.44E-07 |
| Cardiomyocyte | Cnksr2 | 10 | 1.92E-10 | 2.31981264 | 0.46 | 0.154 | 3.84E-07 |
| Cardiomyocyte | Axl1 | 10 | 3.04E-10 | 1.17451913 | 0.576 | 0.276 | 6.08E-07 |
| Cardiomyocyte | Itpkb1 | 10 | 3.19E-10 | 0.59189917 | 0.683 | 0.357 | 6.39E-07 |
| Cardiomyocyte | Nrk1 | 10 | 3.32E-10 | 1.68703319 | 0.468 | 0.204 | 6.64E-07 |
| Cardiomyocyte | AABR070067 | 10 | 3.95E-10 | 1.52534117 | 0.561 | 0.309 | 7.90E-07 |
| Cardiomyocyte | Calcrl2 | 10 | 1.56E-09 | 1.24049585 | 0.626 | 0.343 | 3.12E-06 |
| Cardiomyocyte | Antxr11 | 10 | 1.69E-09 | 1.10833755 | 0.554 | 0.275 | 3.38E-06 |
| Cardiomyocyte | Apln | 10 | 1.93E-09 | 0.93496097 | 0.468 | 0.206 | 3.86E-06 |
| Cardiomyocyte | Ptprj1 | 10 | 2.96E-09 | 1.33091006 | 0.583 | 0.274 | 5.92E-06 |
| Cardiomyocyte | Ugdh | 10 | 8.48E-09 | 1.15306335 | 0.532 | 0.277 | 1.70E-05 |
| Cardiomyocyte | Mis18bp1 | 10 | 1.05E-08 | 0.74022847 | 0.317 | 0.067 | 2.09E-05 |
| Cardiomyocyte | Col5a21 | 10 | 1.60E-08 | 0.83143902 | 0.64 | 0.308 | 3.19E-05 |
| Cardiomyocyte | Crip1 | 10 | 1.83E-08 | 1.28871734 | 0.576 | 0.221 | 3.66E-05 |
| Cardiomyocyte | Acer2 | 10 | 2.25E-08 | 1.74394279 | 0.482 | 0.205 | 4.51E-05 |
| Cardiomyocyte | Mertk2 | 10 | 2.27E-08 | 1.85559732 | 0.475 | 0.196 | 4.54E-05 |
| Cardiomyocyte | Cd63 | 10 | 4.61E-08 | 0.90632916 | 0.583 | 0.277 | 9.22E-05 |
| Cardiomyocyte | Klhl29 | 10 | 5.08E-08 | 1.28799139 | 0.54 | 0.201 | 0.0001017 |
| Cardiomyocyte | Ifitm3 | 10 | 6.55E-08 | 1.55531486 | 0.46 | 0.149 | 0.00013105 |
| Cardiomyocyte | Adam23 | 10 | 1.21E-07 | 1.47491639 | 0.388 | 0.118 | 0.00024189 |
| Cardiomyocyte | Loxl21 | 10 | 1.21E-07 | 0.85391105 | 0.547 | 0.294 | 0.00024275 |
| Cardiomyocyte | Ano4 | 10 | 1.90E-07 | 1.18079514 | 0.388 | 0.096 | 0.00037952 |
| Cardiomyocyte | Heyl | 10 | 1.98E-07 | 1.91811738 | 0.453 | 0.202 | 0.00039509 |
| Cardiomyocyte | Pde10a1 | 10 | 2.16E-07 | 0.93212849 | 0.511 | 0.212 | 0.00043148 |
| Cardiomyocyte | Abca17 | 10 | 2.66E-07 | 1.30118537 | 0.46 | 0.174 | 0.00053161 |
| Cardiomyocyte | Diaph3 | 10 | 7.04E-07 | 0.40581742 | 0.36 | 0.096 | 0.00140856 |
| Cardiomyocyte | Ar1 | 10 | 1.08E-06 | 0.83033057 | 0.518 | 0.268 | 0.00216762 |

| | | | | | | | |
|---|---|---|---|---|---|---|---|
| Cardiomyocyte | Aff31 | 10 | 1.38E-06 | 0.67201655 | 0.59 | 0.317 | 0.00276386 |
| Cardiomyocyte | Fli12 | 10 | 5.91E-06 | 0.35720046 | 0.619 | 0.368 | 0.01182225 |
| Cardiomyocyte | Srgap11 | 10 | 5.92E-06 | 0.41682677 | 0.576 | 0.306 | 0.0118396 |
| Cardiomyocyte | Fhl5 | 10 | 6.93E-06 | 1.46497425 | 0.453 | 0.188 | 0.01386709 |
| Cardiomyocyte | Cemip21 | 10 | 7.85E-06 | 0.66261836 | 0.504 | 0.235 | 0.01570356 |
| Cardiomyocyte | Sema3d2 | 10 | 1.36E-05 | 0.37338767 | 0.453 | 0.172 | 0.0272553 |
| Cardiomyocyte | Tspan51 | 10 | 1.65E-05 | 0.50749521 | 0.525 | 0.248 | 0.03307545 |
| Cardiomyocyte | Wnt9b | 11 | 5.94E-162 | 6.34816752 | 0.486 | 0.013 | 1.19E-158 |
| Cardiomyocyte | Meiob | 11 | 3.28E-104 | 3.79763248 | 0.473 | 0.019 | 6.57E-101 |
| Cardiomyocyte | Stk26 | 11 | 2.73E-43 | 2.49201504 | 0.311 | 0.021 | 5.46E-40 |
| Cardiomyocyte | Gja5 | 11 | 5.46E-36 | 4.39240243 | 0.311 | 0.049 | 1.09E-32 |
| Cardiomyocyte | Sox131 | 11 | 9.85E-32 | 3.27924952 | 0.919 | 0.414 | 1.97E-28 |
| Cardiomyocyte | Cntn6 | 11 | 3.91E-30 | 2.66187213 | 0.284 | 0.018 | 7.83E-27 |
| Cardiomyocyte | Ldb21 | 11 | 4.83E-30 | 3.91856445 | 0.824 | 0.224 | 9.65E-27 |
| Cardiomyocyte | Cep55 | 11 | 2.07E-29 | 3.35540732 | 0.27 | 0.02 | 4.13E-26 |
| Cardiomyocyte | Apba2 | 11 | 5.93E-29 | 5.34474324 | 0.432 | 0.064 | 1.19E-25 |
| Cardiomyocyte | Npnt1 | 11 | 1.13E-27 | 3.83536957 | 0.432 | 0.091 | 2.25E-24 |
| Cardiomyocyte | Ptprb1 | 11 | 6.20E-27 | 2.48542041 | 0.878 | 0.285 | 1.24E-23 |
| Cardiomyocyte | Sulf1 | 11 | 3.20E-22 | 3.77656648 | 0.73 | 0.242 | 6.39E-19 |
| Cardiomyocyte | Mmrn1 | 11 | 3.30E-21 | 5.54892844 | 0.5 | 0.122 | 6.61E-18 |
| Cardiomyocyte | Shank31 | 11 | 9.51E-20 | 2.20290448 | 0.824 | 0.355 | 1.90E-16 |
| Cardiomyocyte | St6galnac31 | 11 | 1.42E-19 | 3.23155644 | 0.689 | 0.284 | 2.84E-16 |
| Cardiomyocyte | Tspan52 | 11 | 4.83E-19 | 2.88553062 | 0.743 | 0.248 | 9.65E-16 |
| Cardiomyocyte | Thsd7a1 | 11 | 1.66E-17 | 3.15259702 | 0.676 | 0.267 | 3.32E-14 |
| Cardiomyocyte | Nxn1 | 11 | 1.90E-17 | 2.32232589 | 0.797 | 0.419 | 3.79E-14 |
| Cardiomyocyte | Itpkb2 | 11 | 2.55E-17 | 1.58619193 | 0.838 | 0.359 | 5.10E-14 |
| Cardiomyocyte | Flt12 | 11 | 1.04E-16 | 2.13468491 | 0.77 | 0.315 | 2.07E-13 |
| Cardiomyocyte | Adgrf51 | 11 | 2.70E-16 | 1.82392661 | 0.77 | 0.323 | 5.41E-13 |
| Cardiomyocyte | Frmd4b1 | 11 | 2.80E-16 | 1.94905228 | 0.77 | 0.451 | 5.59E-13 |
| Cardiomyocyte | Slc26a101 | 11 | 3.17E-16 | 2.72013119 | 0.716 | 0.294 | 6.34E-13 |
| Cardiomyocyte | Srgap12 | 11 | 4.94E-16 | 2.04547416 | 0.797 | 0.306 | 9.88E-13 |

| | | | | | | | |
|---|---|---|---|---|---|---|---|
| Cardiomyocyte | Ppp1r16b | 11 | 9.82E-16 | 2.64377285 | 0.703 | 0.308 | 1.96E-12 |
| Cardiomyocyte | Dmtn | 11 | 1.17E-15 | 5.52033132 | 0.405 | 0.037 | 2.34E-12 |
| Cardiomyocyte | Entpd12 | 11 | 1.83E-15 | 3.00064284 | 0.689 | 0.278 | 3.67E-12 |
| Cardiomyocyte | Podxl1 | 11 | 1.93E-15 | 2.55627848 | 0.689 | 0.231 | 3.85E-12 |
| Cardiomyocyte | Dnm31 | 11 | 2.12E-15 | 1.99055264 | 0.757 | 0.353 | 4.23E-12 |
| Cardiomyocyte | Kif2c1 | 11 | 2.59E-15 | 2.53367979 | 0.459 | 0.111 | 5.17E-12 |
| Cardiomyocyte | Bmx | 11 | 3.90E-15 | 5.06289093 | 0.541 | 0.164 | 7.80E-12 |
| Cardiomyocyte | Tbx1 | 11 | 7.06E-15 | 5.42351697 | 0.257 | 0.005 | 1.41E-11 |
| Cardiomyocyte | Polq | 11 | 1.26E-14 | 2.86290733 | 0.351 | 0.086 | 2.53E-11 |
| Cardiomyocyte | Igf2 | 11 | 1.47E-14 | 3.80930613 | 0.432 | 0.063 | 2.94E-11 |
| Cardiomyocyte | Ptprj2 | 11 | 1.49E-14 | 3.10430628 | 0.649 | 0.277 | 2.97E-11 |
| Cardiomyocyte | Tmem63c | 11 | 2.23E-14 | 6.11364415 | 0.459 | 0.098 | 4.47E-11 |
| Cardiomyocyte | Afap1l2 | 11 | 2.42E-14 | 3.61554432 | 0.568 | 0.267 | 4.84E-11 |
| Cardiomyocyte | Vegfc | 11 | 5.36E-14 | 3.15055195 | 0.662 | 0.253 | 1.07E-10 |
| Cardiomyocyte | Sox17 | 11 | 6.49E-14 | 3.87685584 | 0.608 | 0.234 | 1.30E-10 |
| Cardiomyocyte | Dkk2 | 11 | 7.45E-14 | 4.58907571 | 0.541 | 0.107 | 1.49E-10 |
| Cardiomyocyte | Ccnb1 | 11 | 8.50E-14 | 2.17473966 | 0.459 | 0.173 | 1.70E-10 |
| Cardiomyocyte | Fli13 | 11 | 1.13E-13 | 1.26626879 | 0.797 | 0.369 | 2.26E-10 |
| Cardiomyocyte | LOC1003610 | 11 | 2.28E-13 | 2.09230211 | 0.743 | 0.436 | 4.56E-10 |
| Cardiomyocyte | Pkhd1l1 | 11 | 2.40E-13 | 3.12299405 | 0.622 | 0.154 | 4.80E-10 |
| Cardiomyocyte | Mx12 | 11 | 2.88E-13 | 1.70682517 | 0.405 | 0.146 | 5.77E-10 |
| Cardiomyocyte | Scube1 | 11 | 3.40E-13 | 1.44007095 | 0.459 | 0.078 | 6.80E-10 |
| Cardiomyocyte | Adgrl41 | 11 | 6.04E-13 | 1.93204739 | 0.635 | 0.237 | 1.21E-09 |
| Cardiomyocyte | Flt4 | 11 | 8.61E-13 | 5.02761692 | 0.514 | 0.19 | 1.72E-09 |
| Cardiomyocyte | Myrip1 | 11 | 1.03E-12 | 2.54771937 | 0.649 | 0.363 | 2.07E-09 |
| Cardiomyocyte | P2rx7 | 11 | 1.49E-12 | 3.50418279 | 0.608 | 0.262 | 2.99E-09 |
| Cardiomyocyte | Podnl12 | 11 | 3.81E-12 | 2.80274825 | 0.419 | 0.145 | 7.62E-09 |
| Cardiomyocyte | Ifitm10 | 11 | 7.65E-12 | 4.74516505 | 0.568 | 0.217 | 1.53E-08 |
| Cardiomyocyte | Itgb4 | 11 | 8.91E-12 | 3.338773 | 0.608 | 0.222 | 1.78E-08 |
| Cardiomyocyte | Inpp5d1 | 11 | 1.19E-11 | 2.04122692 | 0.676 | 0.356 | 2.38E-08 |
| Cardiomyocyte | Ablim31 | 11 | 1.34E-11 | 1.75096635 | 0.662 | 0.374 | 2.68E-08 |

| | | | | | | | |
|---|---|---|---|---|---|---|---|
| Cardiomyocyte | Cemip22 | 11 | 1.97E-11 | 1.68694513 | 0.662 | 0.236 | 3.95E-08 |
| Cardiomyocyte | Efnb21 | 11 | 2.46E-11 | 2.56831854 | 0.568 | 0.244 | 4.92E-08 |
| Cardiomyocyte | Meox1 | 11 | 2.52E-11 | 3.85077527 | 0.5 | 0.186 | 5.05E-08 |
| Cardiomyocyte | Bank1 | 11 | 3.22E-11 | 0.67668645 | 0.338 | 0.062 | 6.45E-08 |
| Cardiomyocyte | Tmem200a | 11 | 3.72E-11 | 1.90469734 | 0.311 | 0.058 | 7.43E-08 |
| Cardiomyocyte | Kif4a | 11 | 6.97E-11 | 1.26247162 | 0.486 | 0.152 | 1.39E-07 |
| Cardiomyocyte | Dapk2 | 11 | 8.71E-11 | 1.95631464 | 0.757 | 0.464 | 1.74E-07 |
| Cardiomyocyte | Prkch1 | 11 | 1.07E-10 | 1.41924637 | 0.689 | 0.317 | 2.14E-07 |
| Cardiomyocyte | Kcnip1 | 11 | 2.87E-10 | 1.09802959 | 0.446 | 0.128 | 5.75E-07 |
| Cardiomyocyte | Chn12 | 11 | 4.28E-10 | 1.83045207 | 0.689 | 0.302 | 8.56E-07 |
| Cardiomyocyte | Dipk2b1 | 11 | 4.53E-10 | 2.20812124 | 0.608 | 0.274 | 9.06E-07 |
| Cardiomyocyte | Msr1 | 11 | 4.87E-10 | 1.37284955 | 0.392 | 0.08 | 9.73E-07 |
| Cardiomyocyte | Adgrg3 | 11 | 4.90E-10 | 3.46987537 | 0.595 | 0.278 | 9.81E-07 |
| Cardiomyocyte | Specc11 | 11 | 5.64E-10 | 1.9692148 | 0.581 | 0.3 | 1.13E-06 |
| Cardiomyocyte | Unc5b1 | 11 | 7.32E-10 | 2.87842499 | 0.608 | 0.335 | 1.46E-06 |
| Cardiomyocyte | Dach12 | 11 | 7.78E-10 | 1.01627872 | 0.716 | 0.317 | 1.56E-06 |
| Cardiomyocyte | Nrp21 | 11 | 7.98E-10 | 2.60901102 | 0.676 | 0.33 | 1.60E-06 |
| Cardiomyocyte | Cyyr11 | 11 | 8.01E-10 | 1.79599892 | 0.635 | 0.274 | 1.60E-06 |
| Cardiomyocyte | Mgll1 | 11 | 1.12E-09 | 0.99884503 | 0.797 | 0.499 | 2.24E-06 |
| Cardiomyocyte | Cd842 | 11 | 2.10E-09 | 0.66356103 | 0.378 | 0.065 | 4.21E-06 |
| Cardiomyocyte | AABR070042 | 11 | 2.96E-09 | 0.91111454 | 0.405 | 0.108 | 5.92E-06 |
| Cardiomyocyte | Slc35f1 | 11 | 3.02E-09 | 3.00913318 | 0.568 | 0.277 | 6.04E-06 |
| Cardiomyocyte | Chn21 | 11 | 3.07E-09 | 1.9486436 | 0.514 | 0.171 | 6.15E-06 |
| Cardiomyocyte | Reln | 11 | 3.73E-09 | 6.10030181 | 0.459 | 0.188 | 7.46E-06 |
| Cardiomyocyte | Diaph31 | 11 | 3.75E-09 | 1.41028242 | 0.486 | 0.097 | 7.50E-06 |
| Cardiomyocyte | Capn13 | 11 | 8.00E-09 | 2.48353718 | 0.5 | 0.182 | 1.60E-05 |
| Cardiomyocyte | LOC1009114 | 11 | 1.44E-08 | 4.01818446 | 0.541 | 0.272 | 2.88E-05 |
| Cardiomyocyte | Atp10b | 11 | 2.47E-08 | 4.34533667 | 0.419 | 0.15 | 4.95E-05 |
| Cardiomyocyte | Bcar31 | 11 | 3.66E-08 | 2.31368244 | 0.649 | 0.356 | 7.33E-05 |
| Cardiomyocyte | Ednrb1 | 11 | 4.25E-08 | 2.56169483 | 0.419 | 0.118 | 8.50E-05 |
| Cardiomyocyte | Gcnt22 | 11 | 4.25E-08 | 1.92238712 | 0.595 | 0.28 | 8.51E-05 |

| | | | | | | | |
|---|---|---|---|---|---|---|---|
| Cardiomyocyte | Cmtm81 | 11 | 5.74E-08 | 2.21037813 | 0.554 | 0.255 | 0.00011479 |
| Cardiomyocyte | Ccser11 | 11 | 7.91E-08 | 2.42522244 | 0.554 | 0.297 | 0.00015829 |
| Cardiomyocyte | Meox22 | 11 | 9.73E-08 | 1.26975798 | 0.622 | 0.287 | 0.00019469 |
| Cardiomyocyte | Mbp1 | 11 | 1.11E-07 | 1.24534154 | 0.486 | 0.147 | 0.00022189 |
| Cardiomyocyte | Herc61 | 11 | 2.71E-07 | 1.65441789 | 0.473 | 0.186 | 0.00054285 |
| Cardiomyocyte | Kitlg1 | 11 | 2.72E-07 | 1.91584054 | 0.595 | 0.311 | 0.0005446 |
| Cardiomyocyte | Anxa31 | 11 | 2.88E-07 | 1.6370119 | 0.527 | 0.17 | 0.00057668 |
| Cardiomyocyte | Igfbp41 | 11 | 3.59E-07 | 1.73578915 | 0.486 | 0.161 | 0.00071716 |
| Cardiomyocyte | Aoah1 | 11 | 4.89E-07 | 0.4484686 | 0.514 | 0.212 | 0.00097783 |
| Cardiomyocyte | Sema3d3 | 11 | 6.28E-07 | 3.33202946 | 0.432 | 0.176 | 0.00125569 |
| Cardiomyocyte | Rai141 | 11 | 7.95E-07 | 1.49236085 | 0.554 | 0.219 | 0.0015902 |
| Cardiomyocyte | Ptafr2 | 11 | 8.48E-07 | 0.98732217 | 0.419 | 0.118 | 0.00169502 |
| Cardiomyocyte | Klhl4 | 11 | 8.82E-07 | 3.3722564 | 0.473 | 0.188 | 0.0017639 |
| Cardiomyocyte | Calcrl3 | 11 | 9.09E-07 | 1.15460163 | 0.689 | 0.345 | 0.00181873 |
| Cardiomyocyte | Ston22 | 11 | 1.01E-06 | 1.87904264 | 0.541 | 0.267 | 0.00201418 |
| Cardiomyocyte | Alcam | 11 | 3.22E-06 | 1.04217666 | 0.703 | 0.433 | 0.00644242 |
| Cardiomyocyte | Sorcs1 | 11 | 5.00E-06 | 0.8510073 | 0.514 | 0.235 | 0.00999864 |
| Cardiomyocyte | Ephb1 | 11 | 5.06E-06 | 2.72508616 | 0.514 | 0.243 | 0.01012165 |
| Cardiomyocyte | Pde7a | 11 | 9.34E-06 | 0.57761831 | 0.676 | 0.399 | 0.01868688 |
| Cardiomyocyte | Hmmr | 11 | 1.85E-05 | 1.34142225 | 0.473 | 0.184 | 0.03701454 |
| Cardiomyocyte | Vstm2b | 12 | 2.05E-76 | 6.67775297 | 0.475 | 0.009 | 4.11E-73 |
| Cardiomyocyte | Cntn61 | 12 | 1.41E-50 | 4.00250672 | 0.475 | 0.018 | 2.82E-47 |
| Cardiomyocyte | Wnt9b1 | 12 | 7.45E-43 | 3.77125797 | 0.275 | 0.018 | 1.49E-39 |
| Cardiomyocyte | Cgnl1 | 12 | 8.70E-31 | 5.9924939 | 1 | 0.188 | 1.74E-27 |
| Cardiomyocyte | Pkhd1l11 | 12 | 5.81E-28 | 5.15197007 | 0.975 | 0.154 | 1.16E-24 |
| Cardiomyocyte | Ckap2 | 12 | 5.97E-25 | 4.05373332 | 0.45 | 0.009 | 1.19E-21 |
| Cardiomyocyte | Pgm5 | 12 | 1.52E-24 | 3.54837318 | 0.975 | 0.563 | 3.04E-21 |
| Cardiomyocyte | Hmcn1 | 12 | 1.60E-24 | 4.00929954 | 0.975 | 0.57 | 3.19E-21 |
| Cardiomyocyte | Vwf | 12 | 2.34E-24 | 5.17975103 | 0.925 | 0.176 | 4.68E-21 |
| Cardiomyocyte | Cdh111 | 12 | 3.31E-24 | 5.17708678 | 0.95 | 0.262 | 6.62E-21 |
| Cardiomyocyte | Cemip23 | 12 | 3.63E-24 | 4.46961303 | 0.95 | 0.237 | 7.26E-21 |

| | | | | | | | |
|---|---|---|---|---|---|---|---|
| Cardiomyocyte | Slc9a91 | 12 | 1.42E-21 | 2.96033722 | 1 | 0.536 | 2.84E-18 |
| Cardiomyocyte | Susd4 | 12 | 1.45E-20 | 5.29348721 | 0.5 | 0.022 | 2.90E-17 |
| Cardiomyocyte | Slc5a1 | 12 | 1.98E-20 | 3.57568851 | 0.475 | 0.011 | 3.96E-17 |
| Cardiomyocyte | Fli14 | 12 | 3.99E-20 | 2.60533842 | 0.975 | 0.37 | 7.98E-17 |
| Cardiomyocyte | Chst9 | 12 | 5.60E-20 | 7.25080775 | 0.3 | 0.007 | 1.12E-16 |
| Cardiomyocyte | Chn22 | 12 | 8.85E-20 | 4.68716203 | 0.875 | 0.171 | 1.77E-16 |
| Cardiomyocyte | Exoc3l2 | 12 | 6.95E-19 | 1.95022187 | 0.325 | 0.032 | 1.39E-15 |
| Cardiomyocyte | Shank32 | 12 | 6.38E-18 | 2.52756556 | 0.95 | 0.357 | 1.28E-14 |
| Cardiomyocyte | Zmat4 | 12 | 2.71E-17 | 7.32221546 | 0.625 | 0.204 | 5.43E-14 |
| Cardiomyocyte | Plvap | 12 | 1.10E-16 | 5.42700706 | 0.475 | 0.024 | 2.20E-13 |
| Cardiomyocyte | Ednrb2 | 12 | 2.46E-16 | 4.58509039 | 0.75 | 0.118 | 4.92E-13 |
| Cardiomyocyte | Ltbp1 | 12 | 3.83E-16 | 3.03593549 | 0.9 | 0.457 | 7.66E-13 |
| Cardiomyocyte | Itgb41 | 12 | 9.75E-16 | 4.59416562 | 0.8 | 0.223 | 1.95E-12 |
| Cardiomyocyte | Reg3g1 | 12 | 1.25E-15 | 6.03001396 | 0.525 | 0.122 | 2.50E-12 |
| Cardiomyocyte | Cmahp | 12 | 2.28E-15 | 3.55194141 | 0.5 | 0.031 | 4.55E-12 |
| Cardiomyocyte | Prkch2 | 12 | 2.64E-15 | 2.92624738 | 0.875 | 0.318 | 5.28E-12 |
| Cardiomyocyte | Sncaip | 12 | 3.26E-15 | 5.02699209 | 0.8 | 0.43 | 6.53E-12 |
| Cardiomyocyte | Mfsd4a1 | 12 | 1.10E-14 | 1.91396699 | 0.475 | 0.056 | 2.20E-11 |
| Cardiomyocyte | Flt13 | 12 | 1.44E-14 | 1.95188735 | 0.925 | 0.317 | 2.87E-11 |
| Cardiomyocyte | Klhl291 | 12 | 2.27E-14 | 4.05398294 | 0.8 | 0.206 | 4.55E-11 |
| Cardiomyocyte | Rgs20 | 12 | 2.32E-13 | 4.04415612 | 0.45 | 0.101 | 4.64E-10 |
| Cardiomyocyte | Adgrf52 | 12 | 4.67E-13 | 2.37064749 | 0.8 | 0.326 | 9.35E-10 |
| Cardiomyocyte | Slc12a2 | 12 | 5.64E-13 | 3.01574149 | 0.825 | 0.383 | 1.13E-09 |
| Cardiomyocyte | St6galnac32 | 12 | 3.36E-12 | 3.19441036 | 0.725 | 0.287 | 6.73E-09 |
| Cardiomyocyte | Emcn1 | 12 | 3.58E-12 | 2.9350374 | 0.75 | 0.324 | 7.16E-09 |
| Cardiomyocyte | Ppl1 | 12 | 4.12E-12 | 3.95489627 | 0.575 | 0.163 | 8.24E-09 |
| Cardiomyocyte | Reg3b | 12 | 7.88E-12 | 2.48612022 | 0.55 | 0.191 | 1.58E-08 |
| Cardiomyocyte | Npnt2 | 12 | 1.18E-11 | 2.92586959 | 0.425 | 0.093 | 2.36E-08 |
| Cardiomyocyte | Opcml1 | 12 | 1.35E-11 | 3.7042602 | 0.75 | 0.154 | 2.71E-08 |
| Cardiomyocyte | Nrg11 | 12 | 2.02E-11 | 5.0616251 | 0.7 | 0.269 | 4.04E-08 |
| Cardiomyocyte | Bace2 | 12 | 4.90E-11 | 3.97545758 | 0.65 | 0.271 | 9.79E-08 |

| | | | | | | | |
|---|---|---|---|---|---|---|---|
| Cardiomyocyte | LOC1003610 | 12 | 8.43E-11 | 2.27944895 | 0.9 | 0.437 | 1.69E-07 |
| Cardiomyocyte | P3h2 | 12 | 8.60E-11 | 2.45029711 | 0.775 | 0.321 | 1.72E-07 |
| Cardiomyocyte | Car102 | 12 | 7.87E-10 | 3.72489828 | 0.475 | 0.129 | 1.57E-06 |
| Cardiomyocyte | Acer21 | 12 | 9.40E-10 | 3.92650511 | 0.675 | 0.209 | 1.88E-06 |
| Cardiomyocyte | Ptprb2 | 12 | 1.10E-09 | 2.09966171 | 0.8 | 0.29 | 2.20E-06 |
| Cardiomyocyte | Ifitm101 | 12 | 1.35E-09 | 3.56196827 | 0.65 | 0.219 | 2.71E-06 |
| Cardiomyocyte | Hecw21 | 12 | 1.66E-09 | 1.51577405 | 0.85 | 0.592 | 3.32E-06 |
| Cardiomyocyte | Ntn11 | 12 | 3.05E-09 | 1.38697678 | 0.925 | 0.647 | 6.09E-06 |
| Cardiomyocyte | Frmd4b2 | 12 | 3.20E-09 | 2.00628653 | 0.8 | 0.453 | 6.40E-06 |
| Cardiomyocyte | Lepr | 12 | 4.55E-09 | 4.10068203 | 0.575 | 0.127 | 9.09E-06 |
| Cardiomyocyte | Mid11 | 12 | 5.36E-09 | 2.67895584 | 0.75 | 0.199 | 1.07E-05 |
| Cardiomyocyte | Bmp6 | 12 | 8.40E-09 | 3.47840348 | 0.675 | 0.353 | 1.68E-05 |
| Cardiomyocyte | Entpd13 | 12 | 1.12E-08 | 3.17247949 | 0.675 | 0.281 | 2.23E-05 |
| Cardiomyocyte | Ccnb11 | 12 | 2.19E-08 | 3.1529646 | 0.45 | 0.175 | 4.37E-05 |
| Cardiomyocyte | Slco2a1 | 12 | 2.66E-08 | 3.6203176 | 0.55 | 0.205 | 5.32E-05 |
| Cardiomyocyte | Pawr1 | 12 | 2.72E-08 | 3.70559383 | 0.65 | 0.178 | 5.44E-05 |
| Cardiomyocyte | Spns2 | 12 | 2.75E-08 | 2.64144826 | 0.675 | 0.233 | 5.51E-05 |
| Cardiomyocyte | Cyyr12 | 12 | 4.95E-08 | 1.24425361 | 0.775 | 0.275 | 9.89E-05 |
| Cardiomyocyte | Bmx1 | 12 | 6.05E-08 | 4.68626966 | 0.525 | 0.167 | 0.00012103 |
| Cardiomyocyte | Erc2 | 12 | 8.06E-08 | 3.55982363 | 0.5 | 0.109 | 0.00016111 |
| Cardiomyocyte | Maob | 12 | 8.64E-08 | 0.54557186 | 0.35 | 0.086 | 0.00017276 |
| Cardiomyocyte | Tmem100 | 12 | 1.29E-07 | 3.87399403 | 0.6 | 0.201 | 0.0002575 |
| Cardiomyocyte | Palmd | 12 | 1.69E-07 | 2.26983975 | 0.8 | 0.51 | 0.00033741 |
| Cardiomyocyte | Mgmt1 | 12 | 3.40E-07 | 1.69303324 | 0.75 | 0.404 | 0.00068054 |
| Cardiomyocyte | Kif4a1 | 12 | 4.71E-07 | 1.41265499 | 0.5 | 0.154 | 0.00094136 |
| Cardiomyocyte | Itpkb3 | 12 | 6.24E-07 | 0.95798428 | 0.75 | 0.363 | 0.0012475 |
| Cardiomyocyte | Gucy1a21 | 12 | 6.81E-07 | 2.26581127 | 0.675 | 0.311 | 0.00136289 |
| Cardiomyocyte | Trpc41 | 12 | 7.16E-07 | 0.86486997 | 0.375 | 0.037 | 0.00143199 |
| Cardiomyocyte | Tspan53 | 12 | 8.49E-07 | 1.96980823 | 0.65 | 0.252 | 0.00169717 |
| Cardiomyocyte | Nxn2 | 12 | 8.51E-07 | 1.71880002 | 0.7 | 0.422 | 0.00170189 |
| Cardiomyocyte | Dapk11 | 12 | 1.02E-06 | 1.86648702 | 0.725 | 0.41 | 0.00204803 |

| | | | | | | | |
|---|---|---|---|---|---|---|---|
| Cardiomyocyte | LOC1036942 | 12 | 1.11E-06 | 3.67719097 | 0.45 | 0.045 | 0.00221253 |
| Cardiomyocyte | Cadps21 | 12 | 1.36E-06 | 2.62303023 | 0.575 | 0.244 | 0.00272769 |
| Cardiomyocyte | Ttyh1 | 12 | 1.43E-06 | 3.3943207 | 0.5 | 0.106 | 0.00285312 |
| Cardiomyocyte | Asic2 | 12 | 1.74E-06 | 4.81747657 | 0.5 | 0.125 | 0.00347359 |
| Cardiomyocyte | Efemp1 | 12 | 3.03E-06 | 3.15218454 | 0.6 | 0.268 | 0.00605751 |
| Cardiomyocyte | Smad6 | 12 | 3.61E-06 | 2.06029524 | 0.725 | 0.475 | 0.00722502 |
| Cardiomyocyte | Rgs17 | 12 | 3.90E-06 | 2.93421878 | 0.525 | 0.116 | 0.00779228 |
| Cardiomyocyte | Scube11 | 12 | 4.20E-06 | 1.86626115 | 0.425 | 0.081 | 0.00840218 |
| Cardiomyocyte | Rai142 | 12 | 4.85E-06 | 2.03118999 | 0.675 | 0.221 | 0.00969831 |
| Cardiomyocyte | AABR070275 | 12 | 5.11E-06 | 2.37840188 | 0.625 | 0.275 | 0.01021689 |
| Cardiomyocyte | Nr5a21 | 12 | 5.47E-06 | 1.03971044 | 0.325 | 0.068 | 0.0109489 |
| Cardiomyocyte | Inhba | 12 | 5.97E-06 | 3.98236044 | 0.55 | 0.253 | 0.01194531 |
| Cardiomyocyte | Sulf11 | 12 | 6.35E-06 | 1.32897529 | 0.7 | 0.246 | 0.0127065 |
| Cardiomyocyte | Tnfaip8l31 | 12 | 6.48E-06 | 2.62436522 | 0.475 | 0.12 | 0.012969 |
| Cardiomyocyte | Apoe1 | 12 | 7.42E-06 | 1.57476552 | 0.725 | 0.329 | 0.01484611 |
| Cardiomyocyte | Fth11 | 12 | 7.63E-06 | 1.45759049 | 0.75 | 0.446 | 0.01525303 |
| Cardiomyocyte | Lmcd1 | 12 | 7.90E-06 | 1.18286681 | 0.825 | 0.496 | 0.01579096 |
| Cardiomyocyte | Kcnh11 | 12 | 9.14E-06 | 0.85189003 | 0.45 | 0.123 | 0.01828711 |
| Cardiomyocyte | Ctsz | 12 | 9.18E-06 | 3.28347235 | 0.525 | 0.112 | 0.01835125 |
| Cardiomyocyte | LOC1009114 | 12 | 9.77E-06 | 3.68200579 | 0.55 | 0.274 | 0.01954795 |
| Cardiomyocyte | Eln | 12 | 9.91E-06 | 0.29062039 | 0.925 | 0.662 | 0.01982352 |
| Cardiomyocyte | Gna151 | 12 | 1.08E-05 | 1.71988237 | 0.525 | 0.257 | 0.02161085 |
| Cardiomyocyte | AABR070592 | 12 | 1.55E-05 | 2.47563237 | 0.625 | 0.334 | 0.03095465 |
| Cardiomyocyte | Dpysl31 | 12 | 1.55E-05 | 1.67161687 | 0.7 | 0.372 | 0.03099756 |
| Cardiomyocyte | Camk1d1 | 12 | 1.69E-05 | 1.01048534 | 0.625 | 0.339 | 0.03383159 |
| Cardiomyocyte | Aqp1 | 12 | 1.69E-05 | 2.17190026 | 0.6 | 0.172 | 0.03386366 |
| Cardiomyocyte | Slc16a2 | 12 | 1.80E-05 | 1.73272936 | 0.625 | 0.322 | 0.03602266 |
| Cardiomyocyte | Reln1 | 12 | 1.95E-05 | 2.54028787 | 0.55 | 0.189 | 0.03908076 |
| Cardiomyocyte | Slc16a7 | 12 | 2.49E-05 | 2.67330884 | 0.575 | 0.295 | 0.04988583 |
| Cardiomyocyte | AABR070264 | 13 | 0 | 7.60363429 | 0.36 | 0.004 | 0 |
| Cardiomyocyte | Gfra3 | 13 | 1.87E-111 | 7.48772673 | 0.6 | 0.021 | 3.73E-108 |

| | | | | | | | |
|---|---|---|---|---|---|---|---|
| Cardiomyocyte | Ptprz1 | 13 | 5.00E-78 | 6.4327233 | 0.48 | 0.006 | 1.00E-74 |
| Cardiomyocyte | L1cam | 13 | 2.32E-67 | 6.1413913 | 0.36 | 0.01 | 4.63E-64 |
| Cardiomyocyte | Sox10 | 13 | 3.42E-54 | 6.19044847 | 0.28 | 0.002 | 6.84E-51 |
| Cardiomyocyte | Ngfr | 13 | 2.41E-53 | 9.14916133 | 0.4 | 0.023 | 4.83E-50 |
| Cardiomyocyte | Gldn | 13 | 3.42E-53 | 7.24848845 | 0.72 | 0.005 | 6.83E-50 |
| Cardiomyocyte | Rasgef1c | 13 | 1.86E-40 | 6.84793303 | 0.48 | 0.011 | 3.73E-37 |
| Cardiomyocyte | Cmtm51 | 13 | 5.07E-32 | 7.20739613 | 0.68 | 0.051 | 1.01E-28 |
| Cardiomyocyte | Exoc3l21 | 13 | 1.01E-31 | 3.04080704 | 0.68 | 0.031 | 2.01E-28 |
| Cardiomyocyte | Tdrd12 | 13 | 2.76E-28 | 9.76219207 | 0.56 | 0.02 | 5.52E-25 |
| Cardiomyocyte | Ska11 | 13 | 9.32E-25 | 4.51761795 | 0.72 | 0.083 | 1.86E-21 |
| Cardiomyocyte | Erbb31 | 13 | 3.14E-24 | 7.61813549 | 0.84 | 0.037 | 6.29E-21 |
| Cardiomyocyte | Chl11 | 13 | 4.55E-23 | 7.33127646 | 0.76 | 0.114 | 9.09E-20 |
| Cardiomyocyte | Zfhx4 | 13 | 8.37E-21 | 6.51550623 | 0.52 | 0.004 | 1.67E-17 |
| Cardiomyocyte | Gfra2 | 13 | 9.33E-21 | 7.21668546 | 0.52 | 0.038 | 1.87E-17 |
| Cardiomyocyte | Comp | 13 | 1.17E-18 | 5.15420026 | 0.72 | 0.056 | 2.34E-15 |
| Cardiomyocyte | Scn7a | 13 | 2.75E-18 | 6.72906512 | 0.92 | 0.278 | 5.49E-15 |
| Cardiomyocyte | Lgi4 | 13 | 3.11E-18 | 6.62702811 | 0.92 | 0.148 | 6.22E-15 |
| Cardiomyocyte | Cadm2 | 13 | 4.73E-18 | 7.66635244 | 0.8 | 0.051 | 9.45E-15 |
| Cardiomyocyte | Grik2 | 13 | 1.13E-17 | 6.39412951 | 0.92 | 0.101 | 2.25E-14 |
| Cardiomyocyte | Plac8 | 13 | 7.05E-17 | 3.17904588 | 0.52 | 0.053 | 1.41E-13 |
| Cardiomyocyte | Aatk1 | 13 | 2.30E-16 | 5.27363408 | 0.92 | 0.195 | 4.61E-13 |
| Cardiomyocyte | Adam232 | 13 | 6.52E-16 | 6.5664472 | 0.84 | 0.122 | 1.30E-12 |
| Cardiomyocyte | Igsf112 | 13 | 7.85E-16 | 6.96688663 | 0.64 | 0.173 | 1.57E-12 |
| Cardiomyocyte | Abca8a1 | 13 | 3.17E-15 | 3.46741972 | 0.96 | 0.311 | 6.33E-12 |
| Cardiomyocyte | Zfp536 | 13 | 4.64E-15 | 4.72423379 | 0.92 | 0.38 | 9.28E-12 |
| Cardiomyocyte | Csmd11 | 13 | 4.80E-15 | 6.44627838 | 0.44 | 0.137 | 9.61E-12 |
| Cardiomyocyte | Dab1 | 13 | 1.38E-14 | 5.53349758 | 0.56 | 0.137 | 2.77E-11 |
| Cardiomyocyte | Sorcs11 | 13 | 2.88E-14 | 5.09865538 | 0.88 | 0.236 | 5.76E-11 |
| Cardiomyocyte | Gas7 | 13 | 3.39E-14 | 3.53723646 | 0.88 | 0.227 | 6.78E-11 |
| Cardiomyocyte | Mapk10 | 13 | 5.53E-14 | 4.4764183 | 0.76 | 0.17 | 1.11E-10 |
| Cardiomyocyte | Matn2 | 13 | 6.46E-14 | 5.06141958 | 0.88 | 0.189 | 1.29E-10 |

| | | | | | | | |
|---|---|---|---|---|---|---|---|
| Cardiomyocyte | Tnfrsf9 | 13 | 9.10E-14 | 3.17324693 | 0.68 | 0.099 | 1.82E-10 |
| Cardiomyocyte | Mboat2 | 13 | 9.63E-14 | 4.53886396 | 0.84 | 0.212 | 1.93E-10 |
| Cardiomyocyte | Rassf41 | 13 | 2.79E-13 | 5.81431097 | 0.56 | 0.078 | 5.58E-10 |
| Cardiomyocyte | Il1rapl1 | 13 | 1.13E-12 | 4.83335776 | 0.84 | 0.18 | 2.26E-09 |
| Cardiomyocyte | Afap1l21 | 13 | 1.42E-12 | 4.43669672 | 0.84 | 0.268 | 2.84E-09 |
| Cardiomyocyte | Lrrtm41 | 13 | 2.02E-12 | 6.40225228 | 0.76 | 0.163 | 4.03E-09 |
| Cardiomyocyte | Zeb23 | 13 | 2.60E-12 | 2.22616329 | 0.96 | 0.652 | 5.19E-09 |
| Cardiomyocyte | Gulp1 | 13 | 2.77E-12 | 3.15825965 | 0.92 | 0.392 | 5.53E-09 |
| Cardiomyocyte | AABR070275 | 13 | 2.80E-12 | 3.65678211 | 0.84 | 0.275 | 5.61E-09 |
| Cardiomyocyte | AABR070042 | 13 | 3.55E-12 | 2.77902557 | 0.72 | 0.109 | 7.11E-09 |
| Cardiomyocyte | Alcam1 | 13 | 6.70E-12 | 3.20058741 | 0.88 | 0.435 | 1.34E-08 |
| Cardiomyocyte | Nrxn1 | 13 | 3.26E-11 | 3.73235948 | 0.92 | 0.297 | 6.52E-08 |
| Cardiomyocyte | Itgb42 | 13 | 3.93E-11 | 4.93058683 | 0.76 | 0.225 | 7.85E-08 |
| Cardiomyocyte | Ncam2 | 13 | 5.07E-11 | 7.16663023 | 0.36 | 0.046 | 1.01E-07 |
| Cardiomyocyte | Pde1c | 13 | 5.34E-11 | 3.76804348 | 0.84 | 0.151 | 1.07E-07 |
| Cardiomyocyte | Sv2c1 | 13 | 9.00E-11 | 2.73977595 | 0.84 | 0.23 | 1.80E-07 |
| Cardiomyocyte | Scube12 | 13 | 1.04E-10 | 2.24992345 | 0.56 | 0.082 | 2.08E-07 |
| Cardiomyocyte | Cdh19 | 13 | 1.68E-10 | 6.461342 | 0.64 | 0.148 | 3.36E-07 |
| Cardiomyocyte | Egflam1 | 13 | 2.32E-10 | 2.54704523 | 0.84 | 0.284 | 4.64E-07 |
| Cardiomyocyte | Exph5 | 13 | 3.76E-10 | 2.87268242 | 0.68 | 0.189 | 7.53E-07 |
| Cardiomyocyte | Tenm3 | 13 | 4.66E-10 | 3.92007175 | 0.76 | 0.319 | 9.31E-07 |
| Cardiomyocyte | Bcar32 | 13 | 4.91E-10 | 2.51713227 | 0.84 | 0.358 | 9.82E-07 |
| Cardiomyocyte | Tspan11 | 13 | 7.62E-10 | 4.1020078 | 0.72 | 0.219 | 1.52E-06 |
| Cardiomyocyte | Pappa1 | 13 | 7.76E-10 | 3.60296469 | 0.8 | 0.203 | 1.55E-06 |
| Cardiomyocyte | Iqgap21 | 13 | 8.75E-10 | 2.39849538 | 1 | 0.502 | 1.75E-06 |
| Cardiomyocyte | Bank11 | 13 | 1.23E-09 | 0.57012393 | 0.36 | 0.065 | 2.45E-06 |
| Cardiomyocyte | Fign | 13 | 1.41E-09 | 1.9203967 | 0.92 | 0.643 | 2.81E-06 |
| Cardiomyocyte | Mcoln21 | 13 | 1.46E-09 | 2.48671369 | 0.56 | 0.126 | 2.92E-06 |
| Cardiomyocyte | Col27a1 | 13 | 1.79E-09 | 1.90680664 | 0.84 | 0.357 | 3.58E-06 |
| Cardiomyocyte | Pde8b1 | 13 | 2.13E-09 | 3.12073934 | 0.76 | 0.292 | 4.26E-06 |
| Cardiomyocyte | Gja51 | 13 | 2.26E-09 | 1.15019096 | 0.4 | 0.051 | 4.52E-06 |

| | | | | | | | |
|---|---|---|---|---|---|---|---|
| Cardiomyocyte | Kank41 | 13 | 2.62E-09 | 5.0179697 | 0.68 | 0.2 | 5.24E-06 |
| Cardiomyocyte | Arnt2 | 13 | 3.08E-09 | 2.3434006 | 0.64 | 0.205 | 6.15E-06 |
| Cardiomyocyte | Il331 | 13 | 4.43E-09 | 1.51606771 | 0.72 | 0.208 | 8.87E-06 |
| Cardiomyocyte | Pclo | 13 | 5.81E-09 | 2.68281585 | 0.76 | 0.24 | 1.16E-05 |
| Cardiomyocyte | Col16a1 | 13 | 5.86E-09 | 3.94180552 | 0.8 | 0.323 | 1.17E-05 |
| Cardiomyocyte | Cnksr21 | 13 | 7.58E-09 | 5.49084156 | 0.68 | 0.16 | 1.52E-05 |
| Cardiomyocyte | Nlgn11 | 13 | 7.58E-09 | 4.18240223 | 0.68 | 0.308 | 1.52E-05 |
| Cardiomyocyte | Niban1 | 13 | 8.18E-09 | 3.18569535 | 0.76 | 0.384 | 1.64E-05 |
| Cardiomyocyte | Kcnq31 | 13 | 1.35E-08 | 7.02466652 | 0.6 | 0.236 | 2.70E-05 |
| Cardiomyocyte | Cyp7b13 | 13 | 1.67E-08 | 1.72728672 | 0.8 | 0.218 | 3.35E-05 |
| Cardiomyocyte | Cobl1 | 13 | 1.69E-08 | 2.06887656 | 0.92 | 0.389 | 3.39E-05 |
| Cardiomyocyte | Spon11 | 13 | 2.65E-08 | 2.19580194 | 0.72 | 0.153 | 5.29E-05 |
| Cardiomyocyte | Scg2 | 13 | 3.70E-08 | 2.1653638 | 0.68 | 0.229 | 7.40E-05 |
| Cardiomyocyte | St8sia1 | 13 | 5.24E-08 | 3.58883118 | 0.72 | 0.276 | 0.0001047 |
| Cardiomyocyte | Ntng11 | 13 | 5.29E-08 | 5.2558898 | 0.68 | 0.28 | 0.00010588 |
| Cardiomyocyte | Kcnh8 | 13 | 5.42E-08 | 4.05991253 | 0.76 | 0.251 | 0.00010846 |
| Cardiomyocyte | Icam1 | 13 | 5.77E-08 | 2.45769738 | 0.68 | 0.14 | 0.00011535 |
| Cardiomyocyte | Arhgef26 | 13 | 6.43E-08 | 4.70295607 | 0.6 | 0.11 | 0.00012857 |
| Cardiomyocyte | Aoah2 | 13 | 6.68E-08 | 2.04916366 | 0.72 | 0.214 | 0.00013367 |
| Cardiomyocyte | Tyrobp | 13 | 9.18E-08 | 2.07991468 | 0.68 | 0.19 | 0.0001836 |
| Cardiomyocyte | Reg3b1 | 13 | 1.28E-07 | 0.4295856 | 0.64 | 0.192 | 0.00025664 |
| Cardiomyocyte | Cd831 | 13 | 1.32E-07 | 1.69514249 | 0.68 | 0.227 | 0.00026495 |
| Cardiomyocyte | Lhx6 | 13 | 1.53E-07 | 1.78901745 | 0.68 | 0.3 | 0.00030671 |
| Cardiomyocyte | Kif20a2 | 13 | 1.67E-07 | 3.34294301 | 0.52 | 0.07 | 0.00033346 |
| Cardiomyocyte | Cyfip2 | 13 | 1.71E-07 | 2.94809682 | 0.64 | 0.243 | 0.00034279 |
| Cardiomyocyte | Ptprt1 | 13 | 2.69E-07 | 2.87926262 | 0.72 | 0.319 | 0.00053855 |
| Cardiomyocyte | Gask1b | 13 | 3.01E-07 | 1.27404478 | 0.84 | 0.335 | 0.00060277 |
| Cardiomyocyte | Dclk11 | 13 | 3.09E-07 | 3.25641054 | 0.68 | 0.267 | 0.00061725 |
| Cardiomyocyte | Slc35f11 | 13 | 3.16E-07 | 4.47272584 | 0.6 | 0.28 | 0.00063159 |
| Cardiomyocyte | Bace21 | 13 | 3.81E-07 | 1.60268559 | 0.72 | 0.272 | 0.00076276 |
| Cardiomyocyte | Nrcam | 13 | 4.74E-07 | 2.42420456 | 0.72 | 0.283 | 0.00094737 |

| | | | | | | | |
|---|---|---|---|---|---|---|---|
| Cardiomyocyte | Tnfsf15 | 13 | 5.15E-07 | 3.50350706 | 0.52 | 0.08 | 0.00103066 |
| Cardiomyocyte | Col18a1 | 13 | 5.63E-07 | 2.46323496 | 0.76 | 0.454 | 0.00112501 |
| Cardiomyocyte | Fndc11 | 13 | 6.70E-07 | 1.63565136 | 0.88 | 0.31 | 0.00133941 |
| Cardiomyocyte | Pld3 | 13 | 6.83E-07 | 1.55664122 | 0.8 | 0.334 | 0.00136565 |
| Cardiomyocyte | Mnda1 | 13 | 7.02E-07 | 0.98055106 | 0.76 | 0.237 | 0.00140405 |
| Cardiomyocyte | Gpm6b | 13 | 7.38E-07 | 3.22541353 | 0.64 | 0.222 | 0.00147544 |
| Cardiomyocyte | Aff32 | 13 | 8.36E-07 | 2.66511295 | 0.72 | 0.323 | 0.00167206 |
| Cardiomyocyte | AABR070127 | 13 | 8.61E-07 | 3.63632285 | 0.6 | 0.175 | 0.00172253 |
| Cardiomyocyte | Fam163a1 | 13 | 8.72E-07 | 1.09483607 | 0.64 | 0.321 | 0.0017431 |
| Cardiomyocyte | Rab7b | 13 | 9.75E-07 | 2.2285837 | 0.64 | 0.106 | 0.00194971 |
| Cardiomyocyte | Cybb1 | 13 | 1.26E-06 | 1.26912145 | 0.68 | 0.24 | 0.00251855 |
| Cardiomyocyte | Meox11 | 13 | 1.33E-06 | 0.49566802 | 0.64 | 0.189 | 0.00266479 |
| Cardiomyocyte | Enox1 | 13 | 1.44E-06 | 1.65524705 | 0.76 | 0.312 | 0.00288011 |
| Cardiomyocyte | Nmrk2 | 13 | 1.50E-06 | 1.01649301 | 0.8 | 0.442 | 0.00299269 |
| Cardiomyocyte | Podxl3 | 13 | 1.56E-06 | 2.40250071 | 0.72 | 0.235 | 0.00312931 |
| Cardiomyocyte | Zdbf21 | 13 | 1.68E-06 | 2.4989327 | 0.56 | 0.144 | 0.0033669 |
| Cardiomyocyte | Mbp3 | 13 | 1.72E-06 | 1.54548585 | 0.56 | 0.15 | 0.00344658 |
| Cardiomyocyte | Bmp5 | 13 | 1.85E-06 | 3.19962077 | 0.64 | 0.209 | 0.00369179 |
| Cardiomyocyte | Lgals3 | 13 | 1.87E-06 | 1.59345138 | 0.68 | 0.19 | 0.00374953 |
| Cardiomyocyte | AABR070490 | 13 | 1.90E-06 | 2.66436011 | 0.68 | 0.167 | 0.00380203 |
| Cardiomyocyte | Rgcc | 13 | 1.92E-06 | 2.42583396 | 0.68 | 0.28 | 0.00383686 |
| Cardiomyocyte | Gcnt23 | 13 | 2.00E-06 | 1.58856108 | 0.76 | 0.283 | 0.00399093 |
| Cardiomyocyte | Tmem45b | 13 | 2.01E-06 | 2.64655579 | 0.4 | 0.049 | 0.00401623 |
| Cardiomyocyte | Dnm32 | 13 | 2.09E-06 | 0.96193791 | 0.84 | 0.356 | 0.00418112 |
| Cardiomyocyte | Sybu1 | 13 | 2.29E-06 | 0.81507668 | 0.68 | 0.292 | 0.00458698 |
| Cardiomyocyte | Knl1 | 13 | 2.39E-06 | 2.1590526 | 0.6 | 0.213 | 0.00478682 |
| Cardiomyocyte | Ecm1 | 13 | 2.45E-06 | 1.12092136 | 0.76 | 0.251 | 0.00489905 |
| Cardiomyocyte | Mid12 | 13 | 2.52E-06 | 1.30212329 | 0.8 | 0.201 | 0.00504612 |
| Cardiomyocyte | Ly49s5 | 13 | 2.77E-06 | 2.85792662 | 0.4 | 0.043 | 0.00554885 |
| Cardiomyocyte | Kif18b1 | 13 | 3.59E-06 | 1.61433114 | 0.6 | 0.143 | 0.00717894 |
| Cardiomyocyte | Sorcs2 | 13 | 3.60E-06 | 3.27578648 | 0.6 | 0.308 | 0.00720993 |

| | | | | | | | |
|---|---|---|---|---|---|---|---|
| Cardiomyocyte | Gpd12 | 13 | 3.64E-06 | 1.15455239 | 0.76 | 0.398 | 0.00728207 |
| Cardiomyocyte | Pde10a3 | 13 | 3.92E-06 | 1.36749782 | 0.76 | 0.218 | 0.00784571 |
| Cardiomyocyte | Galnt17 | 13 | 4.16E-06 | 1.75620242 | 0.8 | 0.425 | 0.00832688 |
| Cardiomyocyte | Myo101 | 13 | 4.26E-06 | 1.89503502 | 0.72 | 0.39 | 0.00851091 |
| Cardiomyocyte | Mctp21 | 13 | 4.64E-06 | 1.9258513 | 0.68 | 0.235 | 0.00928211 |
| Cardiomyocyte | Fhl51 | 13 | 4.77E-06 | 1.39154588 | 0.68 | 0.193 | 0.00954135 |
| Cardiomyocyte | Adgrl31 | 13 | 5.10E-06 | 2.26492744 | 0.64 | 0.271 | 0.01019614 |
| Cardiomyocyte | Neurl12 | 13 | 5.11E-06 | 3.19601737 | 0.68 | 0.276 | 0.01021313 |
| Cardiomyocyte | Smpdl3a | 13 | 5.20E-06 | 0.72339579 | 0.76 | 0.265 | 0.01040092 |
| Cardiomyocyte | Cald11 | 13 | 5.75E-06 | 1.4400263 | 0.96 | 0.489 | 0.01149061 |
| Cardiomyocyte | Col7a11 | 13 | 6.62E-06 | 3.2011662 | 0.48 | 0.095 | 0.01324667 |
| Cardiomyocyte | Sh3bp21 | 13 | 6.65E-06 | 0.77271186 | 0.76 | 0.42 | 0.01329621 |
| Cardiomyocyte | Fa2h1 | 13 | 6.66E-06 | 3.47515842 | 0.52 | 0.17 | 0.01332701 |
| Cardiomyocyte | Chn23 | 13 | 6.69E-06 | 1.73026954 | 0.68 | 0.174 | 0.01337413 |
| Cardiomyocyte | Lbp | 13 | 6.92E-06 | 1.10733821 | 0.56 | 0.1 | 0.01383791 |
| Cardiomyocyte | Pip5k1b1 | 13 | 7.10E-06 | 1.35423588 | 1 | 0.611 | 0.01420631 |
| Cardiomyocyte | Itgal | 13 | 7.38E-06 | 1.57777811 | 0.72 | 0.326 | 0.01475986 |
| Cardiomyocyte | Apoe2 | 13 | 9.85E-06 | 1.00114902 | 0.8 | 0.329 | 0.01969755 |
| Cardiomyocyte | AABR070179 | 13 | 1.11E-05 | 2.54290145 | 0.68 | 0.294 | 0.02211267 |
| Cardiomyocyte | Gria1 | 13 | 1.19E-05 | 2.01757895 | 0.32 | 0.03 | 0.02385226 |
| Cardiomyocyte | AC120486.10 | 13 | 1.27E-05 | 3.05989803 | 0.4 | 0.101 | 0.02530109 |
| Cardiomyocyte | Adcy21 | 13 | 1.30E-05 | 1.85594425 | 0.56 | 0.174 | 0.02601593 |
| Cardiomyocyte | Lmln1 | 13 | 1.33E-05 | 1.02194758 | 0.72 | 0.309 | 0.02659944 |
| Cardiomyocyte | Tnfaip6 | 13 | 1.34E-05 | 0.85494574 | 0.44 | 0.146 | 0.02671458 |
| Cardiomyocyte | Tuft11 | 13 | 1.41E-05 | 0.39461928 | 0.68 | 0.238 | 0.0282149 |
| Cardiomyocyte | Camk1d2 | 13 | 1.54E-05 | 0.74038652 | 0.76 | 0.339 | 0.03075975 |
| Cardiomyocyte | Syndig11 | 13 | 1.56E-05 | 0.94895099 | 0.68 | 0.271 | 0.03121788 |
| Cardiomyocyte | Col5a23 | 13 | 1.66E-05 | 1.12976416 | 0.8 | 0.315 | 0.03315994 |
| Cardiomyocyte | Bmx2 | 13 | 1.94E-05 | 0.91099172 | 0.56 | 0.168 | 0.03878403 |
| Cardiomyocyte | Csrp11 | 13 | 2.10E-05 | 1.77515707 | 0.68 | 0.304 | 0.04192198 |
| Cardiomyocyte | Depdc1b1 | 13 | 2.47E-05 | 1.50977842 | 0.4 | 0.131 | 0.04935971 |

| | | | | | | | |
|---|---|---|---|---|---|---|---|
| Cardiomyocyte | AABR070251 | 13 | 2.48E-05 | 1.35326216 | 0.64 | 0.213 | 0.0496328 |
| Fibroblast | Abca8a | 0 | 8.357E-148 | 1.39948175 | 0.802 | 0.53 | 1.671E-144 |
| Fibroblast | Myh6 | 0 | 1.142E-128 | 0.94284586 | 0.887 | 0.761 | 2.284E-125 |
| Fibroblast | Col4a3 | 0 | 7.7653E-93 | 1.52816853 | 0.69 | 0.424 | 1.5531E-89 |
| Fibroblast | Ablim2 | 0 | 1.1375E-75 | 0.33768217 | 0.089 | 0.255 | 2.275E-72 |
| Fibroblast | Tmem132d | 0 | 2.2922E-73 | 2.01423733 | 0.573 | 0.298 | 4.5844E-70 |
| Fibroblast | Kcng2 | 0 | 1.2516E-67 | 0.74376895 | 0.107 | 0.305 | 2.5032E-64 |
| Fibroblast | Aoah | 0 | 2.9399E-67 | 0.51144237 | 0.109 | 0.296 | 5.8798E-64 |
| Fibroblast | Rspo1 | 0 | 2.2697E-66 | 0.26762034 | 0.119 | 0.303 | 4.5394E-63 |
| Fibroblast | Stab1 | 0 | 4.3857E-66 | 0.40456866 | 0.099 | 0.277 | 8.7714E-63 |
| Fibroblast | Nlrp3 | 0 | 5.8226E-66 | 0.33139985 | 0.118 | 0.289 | 1.1645E-62 |
| Fibroblast | Pcdh17 | 0 | 4.3149E-65 | 0.5038527 | 0.106 | 0.261 | 8.6297E-62 |
| Fibroblast | Kcnj8 | 0 | 8.9338E-65 | 0.43722217 | 0.132 | 0.266 | 1.7868E-61 |
| Fibroblast | Grk3 | 0 | 2.5977E-64 | 0.55811744 | 0.109 | 0.262 | 5.1953E-61 |
| Fibroblast | Slco2b1 | 0 | 7.1357E-64 | 1.88399198 | 0.584 | 0.376 | 1.4271E-60 |
| Fibroblast | Cabcoco1 | 0 | 7.7897E-63 | 0.56481941 | 0.123 | 0.299 | 1.5579E-59 |
| Fibroblast | Lrrc4c | 0 | 1.9468E-62 | 0.49660418 | 0.09 | 0.262 | 3.8937E-59 |
| Fibroblast | Nfatc2 | 0 | 2.6718E-59 | 0.37314891 | 0.121 | 0.261 | 5.3436E-56 |
| Fibroblast | Cd68 | 0 | 5.6062E-59 | 0.25341496 | 0.081 | 0.271 | 1.1212E-55 |
| Fibroblast | Aspm | 0 | 1.4799E-58 | 0.69035785 | 0.137 | 0.29 | 2.9599E-55 |
| Fibroblast | Adhfe1 | 0 | 3.0182E-58 | 0.80154345 | 0.115 | 0.266 | 6.0365E-55 |
| Fibroblast | Myo18b | 0 | 3.5646E-58 | 0.34576178 | 0.148 | 0.293 | 7.1291E-55 |
| Fibroblast | Cttnbp2 | 0 | 2.7123E-57 | 0.50504744 | 0.15 | 0.311 | 5.4246E-54 |
| Fibroblast | Kif20a | 0 | 3.1836E-57 | 0.49245421 | 0.128 | 0.271 | 6.3672E-54 |
| Fibroblast | Slc7a8 | 0 | 5.0339E-57 | 1.34381348 | 0.13 | 0.339 | 1.0068E-53 |
| Fibroblast | Pcdh7 | 0 | 8.8705E-57 | 0.36447718 | 0.162 | 0.297 | 1.7741E-53 |
| Fibroblast | Glb1l2 | 0 | 6.0436E-56 | 0.72004153 | 0.127 | 0.282 | 1.2087E-52 |
| Fibroblast | Spink8 | 0 | 1.0556E-55 | 1.06740633 | 0.126 | 0.284 | 2.1112E-52 |
| Fibroblast | Sncaip | 0 | 1.0964E-55 | 0.31614778 | 0.152 | 0.32 | 2.1927E-52 |
| Fibroblast | Nrap | 0 | 1.3348E-55 | 0.79193345 | 0.123 | 0.267 | 2.6695E-52 |
| Fibroblast | Rbp1 | 0 | 3.2694E-55 | 0.36935805 | 0.17 | 0.376 | 6.5387E-52 |

| | | | | | | | |
|---|---|---|---|---|---|---|---|
| Fibroblast | Rcan1 | 0 | 1.1412E-54 | 0.39033907 | 0.163 | 0.315 | 2.2824E-51 |
| Fibroblast | Ddc | 0 | 1.163E-54 | 0.52708408 | 0.096 | 0.262 | 2.3259E-51 |
| Fibroblast | Rgs5 | 0 | 1.2373E-54 | 0.467781 | 0.126 | 0.266 | 2.4746E-51 |
| Fibroblast | Eda2r | 0 | 1.8853E-54 | 0.27852879 | 0.157 | 0.293 | 3.7707E-51 |
| Fibroblast | Sfxn5 | 0 | 6.2114E-54 | 0.42200031 | 0.139 | 0.252 | 1.2423E-50 |
| Fibroblast | Cadps2 | 0 | 7.5107E-54 | 0.37368106 | 0.14 | 0.268 | 1.5021E-50 |
| Fibroblast | Prkaa2 | 0 | 9.4666E-54 | 0.38064346 | 0.126 | 0.253 | 1.8933E-50 |
| Fibroblast | Trim63 | 0 | 1.6262E-53 | 0.47351887 | 0.139 | 0.274 | 3.2523E-50 |
| Fibroblast | Alpk3 | 0 | 1.881E-52 | 1.15163976 | 0.138 | 0.303 | 3.762E-49 |
| Fibroblast | Ptprc | 0 | 4.5486E-52 | 0.46759231 | 0.156 | 0.277 | 9.0972E-49 |
| Fibroblast | LOC1003610 | 0 | 6.0212E-52 | 0.34861594 | 0.155 | 0.298 | 1.2042E-48 |
| Fibroblast | Erbb4 | 0 | 1.0485E-51 | 0.28604836 | 0.161 | 0.278 | 2.0971E-48 |
| Fibroblast | Sacs | 0 | 1.479E-51 | 0.2506729 | 0.14 | 0.272 | 2.958E-48 |
| Fibroblast | AABR070010 | 0 | 2.7448E-51 | 0.58331227 | 0.117 | 0.276 | 5.4895E-48 |
| Fibroblast | Tmem182 | 0 | 3.0312E-51 | 0.41279912 | 0.162 | 0.291 | 6.0624E-48 |
| Fibroblast | Slc16a1 | 0 | 3.3553E-51 | 0.29225342 | 0.152 | 0.275 | 6.7106E-48 |
| Fibroblast | RGD1563354 | 0 | 4.5859E-51 | 1.17605013 | 0.648 | 0.518 | 9.1719E-48 |
| Fibroblast | LOC690045 | 0 | 1.866E-50 | 0.54427693 | 0.155 | 0.278 | 3.7321E-47 |
| Fibroblast | Fyb1 | 0 | 2.8174E-50 | 0.33543508 | 0.15 | 0.26 | 5.6348E-47 |
| Fibroblast | Ppm1e | 0 | 3.4402E-50 | 0.47224709 | 0.157 | 0.297 | 6.8804E-47 |
| Fibroblast | Txlnb | 0 | 4.4652E-50 | 0.4881581 | 0.191 | 0.349 | 8.9303E-47 |
| Fibroblast | AABR070504 | 0 | 5.4239E-50 | 0.91090582 | 0.156 | 0.288 | 1.0848E-46 |
| Fibroblast | Mapt | 0 | 7.4231E-50 | 0.45766763 | 0.144 | 0.261 | 1.4846E-46 |
| Fibroblast | Xkr4 | 0 | 9.574E-50 | 0.65025092 | 0.171 | 0.333 | 1.9148E-46 |
| Fibroblast | Mctp2 | 0 | 2.0819E-49 | 0.30302177 | 0.146 | 0.268 | 4.1637E-46 |
| Fibroblast | Gna15 | 0 | 2.2144E-49 | 0.27637533 | 0.142 | 0.302 | 4.4287E-46 |
| Fibroblast | Art3 | 0 | 3.2352E-49 | 0.65281748 | 0.169 | 0.348 | 6.4703E-46 |
| Fibroblast | Col22a1 | 0 | 5.1319E-49 | 0.38777454 | 0.188 | 0.385 | 1.0264E-45 |
| Fibroblast | Mlf1 | 0 | 8.946E-49 | 0.3744004 | 0.141 | 0.253 | 1.7892E-45 |
| Fibroblast | Acsl1 | 0 | 2.0487E-48 | 0.33902697 | 0.165 | 0.302 | 4.0974E-45 |
| Fibroblast | AABR070171 | 0 | 2.3744E-48 | 0.94090071 | 0.136 | 0.303 | 4.7488E-45 |

| | | | | | | | |
|---|---|---|---|---|---|---|---|
| Fibroblast | Trim54 | 0 | 3.4242E-48 | 0.83796411 | 0.167 | 0.332 | 6.8483E-45 |
| Fibroblast | Trim55 | 0 | 3.6618E-48 | 0.89562258 | 0.145 | 0.261 | 7.3237E-45 |
| Fibroblast | Prkcb | 0 | 4.1273E-48 | 0.2546124 | 0.163 | 0.299 | 8.2545E-45 |
| Fibroblast | Rnf144b | 0 | 4.7633E-48 | 0.33141423 | 0.198 | 0.356 | 9.5267E-45 |
| Fibroblast | Ppara | 0 | 7.3409E-48 | 0.84548848 | 0.16 | 0.306 | 1.4682E-44 |
| Fibroblast | Cox6a1 | 0 | 1.5542E-47 | 0.44416873 | 0.179 | 0.313 | 3.1085E-44 |
| Fibroblast | Lamc2 | 0 | 2.0232E-47 | 0.53549939 | 0.126 | 0.272 | 4.0463E-44 |
| Fibroblast | Ankrd6 | 0 | 2.1301E-47 | 0.51766475 | 0.148 | 0.287 | 4.2602E-44 |
| Fibroblast | Tec | 0 | 3.041E-47 | 0.56955351 | 0.171 | 0.276 | 6.0819E-44 |
| Fibroblast | Esr1 | 0 | 5.4918E-47 | 0.94034809 | 0.182 | 0.325 | 1.0984E-43 |
| Fibroblast | Myom2 | 0 | 1.0468E-46 | 0.57414643 | 0.17 | 0.321 | 2.0936E-43 |
| Fibroblast | AABR070525 | 0 | 1.8734E-46 | 0.46487286 | 0.152 | 0.297 | 3.7467E-43 |
| Fibroblast | Ppp1r14c | 0 | 1.2864E-45 | 0.2507212 | 0.211 | 0.333 | 2.5728E-42 |
| Fibroblast | Fsd2 | 0 | 1.457E-45 | 0.7174485 | 0.184 | 0.314 | 2.914E-42 |
| Fibroblast | Reep1 | 0 | 2.0464E-45 | 0.29532157 | 0.157 | 0.265 | 4.0929E-42 |
| Fibroblast | Efemp1 | 0 | 3.0918E-44 | 0.25366822 | 0.206 | 0.343 | 6.1836E-41 |
| Fibroblast | Pparg | 0 | 3.5129E-44 | 0.43816471 | 0.179 | 0.294 | 7.0259E-41 |
| Fibroblast | Rbm24 | 0 | 4.6647E-44 | 0.87595434 | 0.17 | 0.264 | 9.3295E-41 |
| Fibroblast | Tesc | 0 | 7.6082E-44 | 0.67808746 | 0.196 | 0.302 | 1.5216E-40 |
| Fibroblast | Abca17 | 0 | 1.0539E-43 | 0.48785938 | 0.147 | 0.265 | 2.1078E-40 |
| Fibroblast | Il1rapl2 | 0 | 1.0902E-43 | 0.64626589 | 0.141 | 0.256 | 2.1803E-40 |
| Fibroblast | Asb9 | 0 | 1.2962E-43 | 0.39028467 | 0.131 | 0.258 | 2.5924E-40 |
| Fibroblast | Smpx | 0 | 2.5005E-43 | 0.68233352 | 0.182 | 0.295 | 5.0009E-40 |
| Fibroblast | AABR070265 | 0 | 3.0726E-43 | 0.48154726 | 0.206 | 0.306 | 6.1452E-40 |
| Fibroblast | Eef1a2 | 0 | 1.4806E-42 | 0.87409275 | 0.206 | 0.348 | 2.9612E-39 |
| Fibroblast | Fgf13 | 0 | 3.0645E-42 | 0.69280413 | 0.181 | 0.272 | 6.129E-39 |
| Fibroblast | Unc45b | 0 | 3.4015E-42 | 0.7713597 | 0.186 | 0.309 | 6.803E-39 |
| Fibroblast | Acer2 | 0 | 6.8936E-42 | 0.43741657 | 0.189 | 0.286 | 1.3787E-38 |
| Fibroblast | Sptb | 0 | 1.2368E-41 | 0.449479 | 0.21 | 0.307 | 2.4736E-38 |
| Fibroblast | Myh7b | 0 | 1.3384E-41 | 0.52657885 | 0.217 | 0.364 | 2.6769E-38 |
| Fibroblast | Mypn | 0 | 1.4796E-41 | 0.67292222 | 0.198 | 0.331 | 2.9592E-38 |

| | | | | | | | |
|---|---|---|---|---|---|---|---|
| Fibroblast | Musk | 0 | 1.5173E-41 | 0.28426163 | 0.219 | 0.383 | 3.0347E-38 |
| Fibroblast | Lrrc10 | 0 | 1.9537E-41 | 1.22295688 | 0.15 | 0.263 | 3.9074E-38 |
| Fibroblast | Fam189a1 | 0 | 4.4883E-41 | 0.79257852 | 0.16 | 0.266 | 8.9767E-38 |
| Fibroblast | Popdc2 | 0 | 4.5599E-41 | 0.59876299 | 0.194 | 0.304 | 9.1198E-38 |
| Fibroblast | Pik3ap1 | 0 | 5.5355E-41 | 0.4119417 | 0.194 | 0.299 | 1.1071E-37 |
| Fibroblast | Ckb | 0 | 7.0716E-41 | 0.56970584 | 0.196 | 0.327 | 1.4143E-37 |
| Fibroblast | Pde1c | 0 | 8.4722E-41 | 0.44567001 | 0.181 | 0.264 | 1.6944E-37 |
| Fibroblast | Mtus2 | 0 | 1.164E-40 | 0.53573921 | 0.229 | 0.349 | 2.3279E-37 |
| Fibroblast | Flnc | 0 | 2.1297E-40 | 0.62871788 | 0.225 | 0.356 | 4.2595E-37 |
| Fibroblast | Hvcn1 | 0 | 2.473E-40 | 0.47605958 | 0.21 | 0.315 | 4.9461E-37 |
| Fibroblast | Herc6 | 0 | 3.618E-40 | 0.37320793 | 0.192 | 0.32 | 7.2361E-37 |
| Fibroblast | Tox | 0 | 4.3641E-40 | 0.34314409 | 0.181 | 0.269 | 8.7281E-37 |
| Fibroblast | Tnnt2 | 0 | 5.0978E-40 | 0.96202008 | 0.685 | 0.586 | 1.0196E-36 |
| Fibroblast | Coq8a | 0 | 7.3985E-40 | 0.90707688 | 0.2 | 0.293 | 1.4797E-36 |
| Fibroblast | Dpf3 | 0 | 3.4457E-39 | 0.79281862 | 0.199 | 0.279 | 6.8915E-36 |
| Fibroblast | Itga7 | 0 | 6.9713E-39 | 0.63381601 | 0.168 | 0.26 | 1.3943E-35 |
| Fibroblast | Scn5a | 0 | 9.1849E-39 | 0.71576426 | 0.184 | 0.25 | 1.837E-35 |
| Fibroblast | AABR070515 | 0 | 1.1713E-38 | 0.78426063 | 0.186 | 0.262 | 2.3426E-35 |
| Fibroblast | Abcc8 | 0 | 1.5589E-38 | 0.63284936 | 0.121 | 0.251 | 3.1177E-35 |
| Fibroblast | Pfkfb2 | 0 | 1.8659E-38 | 0.34592458 | 0.224 | 0.296 | 3.7318E-35 |
| Fibroblast | Steap4 | 0 | 3.5031E-38 | 0.90666947 | 0.173 | 0.267 | 7.0061E-35 |
| Fibroblast | AABR070449 | 0 | 3.8676E-38 | 0.57080564 | 0.206 | 0.335 | 7.7353E-35 |
| Fibroblast | Myocd | 0 | 4.8259E-38 | 0.30121795 | 0.205 | 0.305 | 9.6519E-35 |
| Fibroblast | Iqub | 0 | 5.3109E-38 | 0.59944043 | 0.223 | 0.341 | 1.0622E-34 |
| Fibroblast | Arhgap15 | 0 | 6.6497E-38 | 0.58503268 | 0.22 | 0.346 | 1.3299E-34 |
| Fibroblast | Wnk2 | 0 | 6.8964E-38 | 0.57225216 | 0.231 | 0.309 | 1.3793E-34 |
| Fibroblast | Ripor2 | 0 | 7.4289E-38 | 0.26094309 | 0.202 | 0.298 | 1.4858E-34 |
| Fibroblast | Prkcz | 0 | 7.6695E-38 | 0.46726165 | 0.23 | 0.366 | 1.5339E-34 |
| Fibroblast | Dhrs7c | 0 | 8.8126E-38 | 1.40135854 | 0.162 | 0.274 | 1.7625E-34 |
| Fibroblast | AABR070726 | 0 | 1.0109E-37 | 0.66305764 | 0.233 | 0.361 | 2.0218E-34 |
| Fibroblast | AC096301.1 | 0 | 1.065E-37 | 0.40508075 | 0.208 | 0.253 | 2.1301E-34 |

| | | | | | | | |
|---|---|---|---|---|---|---|---|
| Fibroblast | Casq2 | 0 | 2.4647E-37 | 0.46144493 | 0.215 | 0.342 | 4.9293E-34 |
| Fibroblast | Lepr | 0 | 3.5124E-37 | 0.47009315 | 0.223 | 0.343 | 7.0248E-34 |
| Fibroblast | AABR070581 | 0 | 3.6259E-37 | 0.35929676 | 0.13 | 0.258 | 7.2518E-34 |
| Fibroblast | Irf8 | 0 | 9.2079E-37 | 0.61411136 | 0.206 | 0.303 | 1.8416E-33 |
| Fibroblast | Serping1 | 0 | 9.9394E-37 | 0.25708934 | 0.232 | 0.322 | 1.9879E-33 |
| Fibroblast | Maob | 0 | 1.6306E-36 | 0.55629106 | 0.182 | 0.318 | 3.2613E-33 |
| Fibroblast | Igfbp3 | 0 | 2.7947E-36 | 0.41472069 | 0.221 | 0.296 | 5.5894E-33 |
| Fibroblast | Ppp2r2b | 0 | 3.7783E-36 | 0.32940873 | 0.184 | 0.271 | 7.5566E-33 |
| Fibroblast | Cspg4 | 0 | 4.5334E-36 | 0.67103647 | 0.194 | 0.262 | 9.0668E-33 |
| Fibroblast | Efcc1 | 0 | 5.1103E-36 | 1.45983463 | 0.571 | 0.428 | 1.0221E-32 |
| Fibroblast | Tenm4 | 0 | 2.1002E-35 | 1.46596073 | 0.155 | 0.27 | 4.2005E-32 |
| Fibroblast | Naca | 0 | 3.2951E-35 | 0.31645587 | 0.23 | 0.341 | 6.5902E-32 |
| Fibroblast | Rapgef5 | 0 | 3.8216E-35 | 0.35604216 | 0.229 | 0.277 | 7.6432E-32 |
| Fibroblast | Arl15 | 0 | 4.9901E-35 | 0.41722436 | 0.898 | 0.908 | 9.9801E-32 |
| Fibroblast | Fgf12 | 0 | 7.1673E-35 | 0.46043452 | 0.231 | 0.339 | 1.4335E-31 |
| Fibroblast | Shroom3 | 0 | 1.343E-34 | 0.4531513 | 0.212 | 0.333 | 2.686E-31 |
| Fibroblast | Slc22a23 | 0 | 2.2077E-34 | 0.67173664 | 0.208 | 0.271 | 4.4154E-31 |
| Fibroblast | Nmnat2 | 0 | 2.5013E-34 | 1.08710924 | 0.175 | 0.314 | 5.0025E-31 |
| Fibroblast | Fbxo32 | 0 | 5.7669E-34 | 0.4314858 | 0.236 | 0.348 | 1.1534E-30 |
| Fibroblast | Skap1 | 0 | 6.192E-34 | 0.6664245 | 0.133 | 0.26 | 1.2384E-30 |
| Fibroblast | Slc1a1 | 0 | 8.4115E-34 | 0.39586034 | 0.141 | 0.289 | 1.6823E-30 |
| Fibroblast | Bdnf | 0 | 1.4319E-33 | 0.48648078 | 0.219 | 0.3 | 2.8638E-30 |
| Fibroblast | Pla2g5 | 0 | 2.4597E-33 | 0.34190425 | 0.153 | 0.269 | 4.9194E-30 |
| Fibroblast | Slc37a2 | 0 | 8.4271E-33 | 0.70001261 | 0.222 | 0.331 | 1.6854E-29 |
| Fibroblast | Cox5b | 0 | 9.522E-33 | 0.81083858 | 0.213 | 0.299 | 1.9044E-29 |
| Fibroblast | Ntrk3 | 0 | 1.0031E-32 | 0.3305861 | 0.23 | 0.346 | 2.0062E-29 |
| Fibroblast | Ntrk2 | 0 | 1.1923E-32 | 0.60460903 | 0.235 | 0.351 | 2.3845E-29 |
| Fibroblast | Tgfbi | 0 | 1.874E-32 | 0.56199052 | 0.254 | 0.383 | 3.7479E-29 |
| Fibroblast | Gnao1 | 0 | 3.3211E-32 | 0.36115379 | 0.262 | 0.367 | 6.6421E-29 |
| Fibroblast | Aatk | 0 | 3.7854E-32 | 1.13244799 | 0.189 | 0.274 | 7.5708E-29 |
| Fibroblast | Macrod1 | 0 | 3.8965E-32 | 0.54904016 | 0.221 | 0.315 | 7.7929E-29 |

| | | | | | | | |
|---|---|---|---|---|---|---|---|
| Fibroblast | Coro6 | 0 | 3.9757E-32 | 0.69237784 | 0.207 | 0.281 | 7.9515E-29 |
| Fibroblast | Hspb6 | 0 | 4.4394E-32 | 0.93293555 | 0.21 | 0.281 | 8.8788E-29 |
| Fibroblast | RGD1565355 | 0 | 1.9364E-31 | 0.29236932 | 0.194 | 0.262 | 3.8728E-28 |
| Fibroblast | Gsn | 0 | 1.9944E-31 | 0.79585459 | 0.663 | 0.568 | 3.9889E-28 |
| Fibroblast | Myl2 | 0 | 5.577E-31 | 1.0312257 | 0.647 | 0.585 | 1.1154E-27 |
| Fibroblast | Cux2 | 0 | 6.4332E-31 | 1.02540657 | 0.242 | 0.33 | 1.2866E-27 |
| Fibroblast | Mylk3 | 0 | 6.6189E-31 | 0.58949083 | 0.233 | 0.361 | 1.3238E-27 |
| Fibroblast | Pdpn | 0 | 7.6944E-31 | 0.38490016 | 0.251 | 0.294 | 1.5389E-27 |
| Fibroblast | Deptor | 0 | 1.992E-30 | 0.59765386 | 0.26 | 0.395 | 3.984E-27 |
| Fibroblast | Psap | 0 | 6.1144E-30 | 0.30957311 | 0.208 | 0.299 | 1.2229E-26 |
| Fibroblast | Tox3 | 0 | 2.1033E-29 | 0.75128132 | 0.207 | 0.258 | 4.2066E-26 |
| Fibroblast | Mbp | 0 | 3.2486E-29 | 1.06927394 | 0.221 | 0.343 | 6.4973E-26 |
| Fibroblast | Cd74 | 0 | 3.4171E-29 | 0.54686933 | 0.23 | 0.287 | 6.8342E-26 |
| Fibroblast | Ppargc1a | 0 | 5.8061E-29 | 0.87209096 | 0.223 | 0.256 | 1.1612E-25 |
| Fibroblast | Slco5a1 | 0 | 6.1171E-29 | 0.47280732 | 0.252 | 0.298 | 1.2234E-25 |
| Fibroblast | Ebf2 | 0 | 1.1362E-28 | 0.38327313 | 0.874 | 0.861 | 2.2724E-25 |
| Fibroblast | Pcbp3 | 0 | 2.2002E-28 | 0.26474655 | 0.298 | 0.411 | 4.4004E-25 |
| Fibroblast | Asic2 | 0 | 2.3076E-28 | 1.58033095 | 0.195 | 0.282 | 4.6153E-25 |
| Fibroblast | Csrp3 | 0 | 6.1437E-28 | 1.00360516 | 0.231 | 0.298 | 1.2287E-24 |
| Fibroblast | Usp18 | 0 | 8.134E-28 | 0.63855351 | 0.25 | 0.326 | 1.6268E-24 |
| Fibroblast | Lrp4 | 0 | 8.5848E-28 | 0.28493426 | 0.305 | 0.443 | 1.717E-24 |
| Fibroblast | Nppb | 0 | 1.0165E-27 | 0.92745552 | 0.228 | 0.282 | 2.033E-24 |
| Fibroblast | Dsg2 | 0 | 1.1025E-27 | 0.82446641 | 0.18 | 0.261 | 2.205E-24 |
| Fibroblast | Tex22 | 0 | 2.7633E-27 | 1.12716261 | 0.206 | 0.257 | 5.5266E-24 |
| Fibroblast | Hcn1 | 0 | 2.8193E-27 | 0.55811828 | 0.208 | 0.273 | 5.6387E-24 |
| Fibroblast | Enpp2 | 0 | 2.9675E-27 | 0.39854017 | 0.266 | 0.397 | 5.935E-24 |
| Fibroblast | Prom1 | 0 | 3.077E-27 | 0.52448176 | 0.204 | 0.305 | 6.1541E-24 |
| Fibroblast | Apcdd1l | 0 | 5.0018E-27 | 0.99161057 | 0.198 | 0.338 | 1.0004E-23 |
| Fibroblast | Acad10 | 0 | 5.097E-27 | 0.43086845 | 0.272 | 0.353 | 1.0194E-23 |
| Fibroblast | Ptcd3 | 0 | 5.156E-27 | 0.33972877 | 0.26 | 0.322 | 1.0312E-23 |
| Fibroblast | Fgf10 | 0 | 5.6498E-27 | 0.40787366 | 0.252 | 0.347 | 1.13E-23 |

| | | | | | | | |
|---|---|---|---|---|---|---|---|
| Fibroblast | Adamts3 | 0 | 7.6701E-27 | 0.61054579 | 0.248 | 0.312 | 1.534E-23 |
| Fibroblast | Trdn | 0 | 7.9556E-27 | 0.69022922 | 0.225 | 0.292 | 1.5911E-23 |
| Fibroblast | Atp2b2 | 0 | 1.3146E-26 | 0.54330741 | 0.165 | 0.275 | 2.6292E-23 |
| Fibroblast | Slc44a5 | 0 | 1.5039E-26 | 0.57607911 | 0.281 | 0.413 | 3.0079E-23 |
| Fibroblast | Mki67 | 0 | 1.5333E-26 | 0.56310595 | 0.178 | 0.28 | 3.0666E-23 |
| Fibroblast | Ank3 | 0 | 2.6832E-26 | 0.36544829 | 0.238 | 0.31 | 5.3663E-23 |
| Fibroblast | Uqcr10 | 0 | 4.8913E-26 | 0.95741315 | 0.246 | 0.323 | 9.7825E-23 |
| Fibroblast | Trhde | 0 | 5.0256E-26 | 0.59021824 | 0.184 | 0.301 | 1.0051E-22 |
| Fibroblast | Myoz2 | 0 | 5.3919E-26 | 0.79052942 | 0.219 | 0.258 | 1.0784E-22 |
| Fibroblast | Bzw2 | 0 | 6.2155E-26 | 0.42820764 | 0.266 | 0.374 | 1.2431E-22 |
| Fibroblast | Adcy2 | 0 | 1.3159E-25 | 0.57991171 | 0.258 | 0.349 | 2.6318E-22 |
| Fibroblast | LOC687780 | 0 | 1.5319E-25 | 0.43107395 | 0.246 | 0.26 | 3.0638E-22 |
| Fibroblast | Fth1 | 0 | 2.2369E-25 | 0.44342021 | 0.235 | 0.328 | 4.4738E-22 |
| Fibroblast | Cst3 | 0 | 2.7848E-25 | 0.41034178 | 0.269 | 0.317 | 5.5695E-22 |
| Fibroblast | Pgam2 | 0 | 3.0886E-25 | 0.59521103 | 0.261 | 0.333 | 6.1772E-22 |
| Fibroblast | Lyn | 0 | 4.0535E-25 | 0.32319958 | 0.269 | 0.386 | 8.1071E-22 |
| Fibroblast | Mb | 0 | 6.0746E-25 | 0.88159797 | 0.64 | 0.563 | 1.2149E-21 |
| Fibroblast | Ppl | 0 | 1.6525E-24 | 0.53729235 | 0.225 | 0.271 | 3.3049E-21 |
| Fibroblast | Mlip | 0 | 1.835E-24 | 0.58098189 | 0.249 | 0.305 | 3.6701E-21 |
| Fibroblast | Fabp4 | 0 | 1.8513E-24 | 0.58449223 | 0.227 | 0.259 | 3.7025E-21 |
| Fibroblast | Atp5f1e | 0 | 2.5774E-24 | 0.94225612 | 0.237 | 0.338 | 5.1548E-21 |
| Fibroblast | Abcc4 | 0 | 5.7418E-24 | 0.72520476 | 0.275 | 0.38 | 1.1484E-20 |
| Fibroblast | Serf2 | 0 | 5.9672E-24 | 0.34421654 | 0.316 | 0.454 | 1.1934E-20 |
| Fibroblast | Prkar2b | 0 | 6.7956E-24 | 0.7942058 | 0.262 | 0.358 | 1.3591E-20 |
| Fibroblast | Osbp2 | 0 | 1.0926E-23 | 1.12259366 | 0.282 | 0.384 | 2.1853E-20 |
| Fibroblast | Uqcrq | 0 | 1.8688E-23 | 0.59217425 | 0.277 | 0.354 | 3.7375E-20 |
| Fibroblast | Kcnn3 | 0 | 1.9301E-23 | 1.3327474 | 0.57 | 0.534 | 3.8602E-20 |
| Fibroblast | LOC1001348 | 0 | 1.9484E-23 | 0.86416864 | 0.232 | 0.263 | 3.8969E-20 |
| Fibroblast | Lifr | 0 | 4.3013E-23 | 0.54113389 | 0.251 | 0.359 | 8.6026E-20 |
| Fibroblast | Obscn | 0 | 6.1026E-23 | 0.73005163 | 0.261 | 0.314 | 1.2205E-19 |
| Fibroblast | Necab1 | 0 | 9.7031E-23 | 0.99056641 | 0.245 | 0.304 | 1.9406E-19 |

| | | | | | | | |
|---|---|---|---|---|---|---|---|
| Fibroblast | Dlgap1 | 0 | 9.833E-23 | 0.53427065 | 0.31 | 0.412 | 1.9666E-19 |
| Fibroblast | Ank2 | 0 | 1.0259E-22 | 0.39930741 | 0.85 | 0.859 | 2.0519E-19 |
| Fibroblast | Clybl | 0 | 1.5433E-22 | 0.58921641 | 0.294 | 0.439 | 3.0866E-19 |
| Fibroblast | Hspb7 | 0 | 1.8174E-22 | 0.92535497 | 0.273 | 0.368 | 3.6347E-19 |
| Fibroblast | AC110709.2 | 0 | 6.1545E-22 | 0.85832655 | 0.173 | 0.267 | 1.2309E-18 |
| Fibroblast | Xirp2 | 0 | 6.3241E-22 | 0.6784923 | 0.254 | 0.33 | 1.2648E-18 |
| Fibroblast | Frem1 | 0 | 6.7693E-22 | 1.56033971 | 0.503 | 0.379 | 1.3539E-18 |
| Fibroblast | Lmo7 | 0 | 7.9444E-22 | 0.26732455 | 0.31 | 0.401 | 1.5889E-18 |
| Fibroblast | Asb14 | 0 | 1.6675E-21 | 0.8310463 | 0.231 | 0.282 | 3.3349E-18 |
| Fibroblast | Ryr2 | 0 | 2.199E-21 | 0.68355556 | 0.649 | 0.597 | 4.398E-18 |
| Fibroblast | Tmcc3 | 0 | 7.2625E-21 | 0.60116287 | 0.291 | 0.394 | 1.4525E-17 |
| Fibroblast | Hk2 | 0 | 7.6052E-21 | 0.25684545 | 0.364 | 0.498 | 1.521E-17 |
| Fibroblast | Ankrd1 | 0 | 8.8324E-21 | 0.81045971 | 0.625 | 0.584 | 1.7665E-17 |
| Fibroblast | Lpl | 0 | 9.9482E-21 | 0.63616058 | 0.277 | 0.354 | 1.9896E-17 |
| Fibroblast | Nrk | 0 | 1.9137E-20 | 1.04827839 | 0.264 | 0.41 | 3.8274E-17 |
| Fibroblast | Myom1 | 0 | 2.3123E-20 | 0.81543048 | 0.271 | 0.301 | 4.6246E-17 |
| Fibroblast | Pclo | 0 | 2.7408E-20 | 1.13322762 | 0.253 | 0.291 | 5.4816E-17 |
| Fibroblast | Myl3 | 0 | 2.8196E-20 | 1.06289962 | 0.598 | 0.54 | 5.6393E-17 |
| Fibroblast | Il6r | 0 | 4.4641E-20 | 0.34520386 | 0.318 | 0.428 | 8.9281E-17 |
| Fibroblast | Ngf | 0 | 6.5162E-20 | 1.12472549 | 0.599 | 0.575 | 1.3032E-16 |
| Fibroblast | AABR070359 | 0 | 7.4514E-20 | 0.48807232 | 0.767 | 0.764 | 1.4903E-16 |
| Fibroblast | Afap1l2 | 0 | 1.4522E-19 | 0.59956327 | 0.286 | 0.402 | 2.9045E-16 |
| Fibroblast | Acta1 | 0 | 3.2239E-19 | 0.99457799 | 0.244 | 0.267 | 6.4477E-16 |
| Fibroblast | Syndig1 | 0 | 3.6522E-19 | 0.48282537 | 0.321 | 0.468 | 7.3043E-16 |
| Fibroblast | Aldh1a1 | 0 | 4.3032E-19 | 0.82732007 | 0.273 | 0.394 | 8.6064E-16 |
| Fibroblast | AABR070010 | 0 | 7.2539E-19 | 0.75157758 | 0.298 | 0.387 | 1.4508E-15 |
| Fibroblast | Myh7 | 0 | 3.2292E-18 | 1.01673874 | 0.603 | 0.547 | 6.4585E-15 |
| Fibroblast | Htra3 | 0 | 5.6212E-18 | 1.05445736 | 0.546 | 0.515 | 1.1242E-14 |
| Fibroblast | Actn2 | 0 | 6.6887E-18 | 0.5481168 | 0.299 | 0.384 | 1.3377E-14 |
| Fibroblast | AABR070525 | 0 | 1.0111E-17 | 0.7937837 | 0.269 | 0.283 | 2.0222E-14 |
| Fibroblast | Lgals1 | 0 | 1.9299E-17 | 0.35676968 | 0.349 | 0.429 | 3.8597E-14 |

| | | | | | | | |
|---|---|---|---|---|---|---|---|
| Fibroblast | Kcnq1 | 0 | 2.6347E-17 | 0.59841974 | 0.295 | 0.337 | 5.2693E-14 |
| Fibroblast | Tnni3 | 0 | 2.8913E-17 | 1.03681062 | 0.556 | 0.444 | 5.7826E-14 |
| Fibroblast | Adcy5 | 0 | 1.6659E-16 | 0.5132136 | 0.317 | 0.422 | 3.3318E-13 |
| Fibroblast | Btbd11 | 0 | 2.455E-16 | 0.61263061 | 0.301 | 0.384 | 4.91E-13 |
| Fibroblast | Ndufa4 | 0 | 2.5197E-16 | 0.85470902 | 0.3 | 0.363 | 5.0395E-13 |
| Fibroblast | Fabp3 | 0 | 8.797E-16 | 0.90715344 | 0.275 | 0.329 | 1.7594E-12 |
| Fibroblast | Acacb | 0 | 3.0569E-15 | 0.63670554 | 0.281 | 0.294 | 6.1139E-12 |
| Fibroblast | Tnnc1 | 0 | 3.543E-15 | 1.02001993 | 0.231 | 0.276 | 7.0861E-12 |
| Fibroblast | Ntf3 | 0 | 3.5821E-15 | 0.93418191 | 0.272 | 0.348 | 7.1641E-12 |
| Fibroblast | Arhgap22 | 0 | 4.1387E-15 | 0.71913658 | 0.308 | 0.398 | 8.2774E-12 |
| Fibroblast | Atp5mg | 0 | 6.0586E-15 | 0.8024667 | 0.279 | 0.359 | 1.2117E-11 |
| Fibroblast | Dsp | 0 | 1.4733E-14 | 1.06549994 | 0.277 | 0.313 | 2.9467E-11 |
| Fibroblast | Adk | 0 | 2.5109E-14 | 0.31902334 | 0.362 | 0.456 | 5.0219E-11 |
| Fibroblast | Ppargc1b | 0 | 2.5964E-14 | 0.66656747 | 0.312 | 0.366 | 5.1929E-11 |
| Fibroblast | Kcnc2 | 0 | 3.0869E-14 | 1.4626093 | 0.491 | 0.371 | 6.1738E-11 |
| Fibroblast | Pde3a | 0 | 7.8604E-14 | 0.90231436 | 0.568 | 0.503 | 1.5721E-10 |
| Fibroblast | Ccdc141 | 0 | 8.7207E-14 | 0.30578045 | 0.358 | 0.43 | 1.7441E-10 |
| Fibroblast | Oxr1 | 0 | 9.4792E-14 | 0.32576995 | 0.358 | 0.421 | 1.8958E-10 |
| Fibroblast | Agbl1 | 0 | 1.316E-13 | 1.08062381 | 0.288 | 0.394 | 2.632E-10 |
| Fibroblast | Nt5e | 0 | 1.5776E-13 | 0.80428485 | 0.268 | 0.298 | 3.1553E-10 |
| Fibroblast | Tnni3k | 0 | 3.8664E-13 | 0.68556612 | 0.296 | 0.285 | 7.7328E-10 |
| Fibroblast | Sox13 | 0 | 3.9095E-13 | 0.60130961 | 0.309 | 0.414 | 7.819E-10 |
| Fibroblast | Cacna1a | 0 | 4.062E-13 | 0.27255767 | 0.396 | 0.536 | 8.1241E-10 |
| Fibroblast | Insr | 0 | 5.2082E-13 | 0.31020147 | 0.383 | 0.477 | 1.0416E-09 |
| Fibroblast | Ar | 0 | 1.4143E-12 | 0.82585552 | 0.59 | 0.569 | 2.8286E-09 |
| Fibroblast | Col6a2 | 0 | 1.8571E-12 | 0.54009276 | 0.662 | 0.679 | 3.7142E-09 |
| Fibroblast | Mpped2 | 0 | 1.9472E-12 | 0.3481126 | 0.404 | 0.53 | 3.8944E-09 |
| Fibroblast | Cmya5 | 0 | 2.1164E-12 | 0.50398441 | 0.341 | 0.382 | 4.2328E-09 |
| Fibroblast | Magi2 | 0 | 2.2975E-12 | 0.59784928 | 0.322 | 0.336 | 4.595E-09 |
| Fibroblast | Akap12 | 0 | 2.3922E-12 | 0.4373549 | 0.348 | 0.434 | 4.7844E-09 |
| Fibroblast | Arhgap24 | 0 | 2.4698E-12 | 0.60216025 | 0.64 | 0.647 | 4.9396E-09 |

| | | | | | | | |
|---|---|---|---|---|---|---|---|
| Fibroblast | Colec12 | 0 | 2.596E-12 | 0.69650563 | 0.644 | 0.629 | 5.1921E-09 |
| Fibroblast | Atp5mc3 | 0 | 4.3134E-12 | 0.88309269 | 0.32 | 0.358 | 8.6268E-09 |
| Fibroblast | Cacnb2 | 0 | 5.0472E-12 | 0.61660288 | 0.327 | 0.37 | 1.0094E-08 |
| Fibroblast | Dapk1 | 0 | 6.1862E-12 | 0.59627459 | 0.65 | 0.662 | 1.2372E-08 |
| Fibroblast | Serpine2 | 0 | 6.2103E-12 | 0.90749851 | 0.315 | 0.379 | 1.2421E-08 |
| Fibroblast | Atp5mc1 | 0 | 7.1211E-12 | 0.89311354 | 0.313 | 0.327 | 1.4242E-08 |
| Fibroblast | Hsd11b1 | 0 | 7.1994E-12 | 0.96636861 | 0.296 | 0.358 | 1.4399E-08 |
| Fibroblast | Enox1 | 0 | 7.3677E-12 | 0.62530139 | 0.337 | 0.437 | 1.4735E-08 |
| Fibroblast | Des | 0 | 7.5129E-12 | 0.85262418 | 0.271 | 0.331 | 1.5026E-08 |
| Fibroblast | Col6a6 | 0 | 7.6819E-12 | 0.41038323 | 0.715 | 0.741 | 1.5364E-08 |
| Fibroblast | Cyp7b1 | 0 | 1.0343E-11 | 0.36060976 | 0.389 | 0.477 | 2.0686E-08 |
| Fibroblast | Atp2a2 | 0 | 1.0672E-11 | 0.43706971 | 0.72 | 0.755 | 2.1345E-08 |
| Fibroblast | Phyh | 0 | 1.3563E-11 | 0.69916601 | 0.302 | 0.348 | 2.7127E-08 |
| Fibroblast | Dcn | 0 | 1.9222E-11 | 0.37459686 | 0.777 | 0.799 | 3.8444E-08 |
| Fibroblast | Ivns1abp | 0 | 1.9919E-11 | 0.72646535 | 0.306 | 0.36 | 3.9838E-08 |
| Fibroblast | Il1r1 | 0 | 2.0194E-11 | 0.4645796 | 0.702 | 0.723 | 4.0388E-08 |
| Fibroblast | Sgip1 | 0 | 2.3021E-11 | 0.74646005 | 0.327 | 0.412 | 4.6042E-08 |
| Fibroblast | Gpx3 | 0 | 2.5162E-11 | 0.42825326 | 0.345 | 0.389 | 5.0325E-08 |
| Fibroblast | Cryab | 0 | 2.9778E-11 | 0.91161884 | 0.323 | 0.389 | 5.9556E-08 |
| Fibroblast | Atp5me | 0 | 4.3223E-11 | 0.83519847 | 0.308 | 0.382 | 8.6447E-08 |
| Fibroblast | Slc25a4 | 0 | 4.6365E-11 | 1.03114708 | 0.561 | 0.521 | 9.273E-08 |
| Fibroblast | Akap6 | 0 | 1.5233E-10 | 0.70443079 | 0.349 | 0.401 | 3.0466E-07 |
| Fibroblast | AABR070015 | 0 | 2.293E-10 | 0.38909859 | 0.391 | 0.476 | 4.5861E-07 |
| Fibroblast | Dmd | 0 | 6.1235E-10 | 0.33731633 | 0.735 | 0.774 | 1.2247E-06 |
| Fibroblast | Bmper | 0 | 1.2135E-09 | 0.90881505 | 0.547 | 0.545 | 2.427E-06 |
| Fibroblast | Sorbs2 | 0 | 1.7544E-09 | 0.56181317 | 0.31 | 0.325 | 3.5088E-06 |
| Fibroblast | Tll2 | 0 | 1.7894E-09 | 0.98200117 | 0.239 | 0.289 | 3.5787E-06 |
| Fibroblast | Rnls | 0 | 4.0883E-09 | 0.48736326 | 0.366 | 0.428 | 8.1766E-06 |
| Fibroblast | Dgkb | 0 | 4.5112E-09 | 0.59357851 | 0.354 | 0.435 | 9.0225E-06 |
| Fibroblast | AABR070070 | 0 | 6.1885E-09 | 0.5126132 | 0.626 | 0.604 | 1.2377E-05 |
| Fibroblast | Crispld2 | 0 | 9.6607E-09 | 0.99977277 | 0.503 | 0.445 | 1.9321E-05 |

| | | | | | | | |
|---|---|---|---|---|---|---|---|
| Fibroblast | Colgalt2 | 0 | 1.026E-08 | 0.92216972 | 0.527 | 0.515 | 2.052E-05 |
| Fibroblast | Ntn1 | 0 | 2.7725E-08 | 0.42077796 | 0.429 | 0.531 | 5.5451E-05 |
| Fibroblast | Mamdc2 | 0 | 3.2702E-08 | 1.12997795 | 0.302 | 0.344 | 6.5404E-05 |
| Fibroblast | Blnk | 0 | 3.8157E-08 | 1.73761094 | 0.272 | 0.307 | 7.6313E-05 |
| Fibroblast | Ckmt2 | 0 | 6.1249E-08 | 0.98782069 | 0.341 | 0.369 | 0.0001225 |
| Fibroblast | Bmp6 | 0 | 7.5346E-08 | 0.79799702 | 0.531 | 0.479 | 0.00015069 |
| Fibroblast | Oxct1 | 0 | 8.522E-08 | 0.58272184 | 0.406 | 0.505 | 0.00017044 |
| Fibroblast | Zfp385b | 0 | 2.827E-07 | 0.54704522 | 0.391 | 0.47 | 0.00056541 |
| Fibroblast | Gli2 | 0 | 4.1958E-07 | 0.61915855 | 0.592 | 0.596 | 0.00083916 |
| Fibroblast | Ckm | 0 | 4.6349E-07 | 1.16928328 | 0.329 | 0.348 | 0.00092698 |
| Fibroblast | Astn2 | 0 | 7.8021E-07 | 1.65667781 | 0.29 | 0.329 | 0.00156042 |
| Fibroblast | Ldhb | 0 | 7.8457E-07 | 0.72899014 | 0.375 | 0.428 | 0.00156914 |
| Fibroblast | Adra1a | 0 | 8.2098E-07 | 0.7798105 | 0.344 | 0.38 | 0.00164196 |
| Fibroblast | Plxnc1 | 0 | 9.1661E-07 | 1.94799852 | 0.284 | 0.272 | 0.00183322 |
| Fibroblast | Rmdn1 | 0 | 2.2936E-06 | 0.53314079 | 0.398 | 0.499 | 0.00458722 |
| Fibroblast | Tbx20 | 0 | 2.8434E-06 | 0.6913048 | 0.551 | 0.542 | 0.00568681 |
| Fibroblast | Sh3kbp1 | 0 | 1.4417E-05 | 0.62653677 | 0.385 | 0.427 | 0.02883434 |
| Fibroblast | Ror1 | 0 | 1.8536E-05 | 0.4599918 | 0.433 | 0.528 | 0.03707192 |
| Fibroblast | Hs6st2 | 0 | 1.8665E-05 | 0.94588495 | 0.38 | 0.44 | 0.0373304 |
| Fibroblast | Opcml | 0 | 2.363E-05 | 1.38620213 | 0.44 | 0.335 | 0.04725945 |
| Fibroblast | Eln | 1 | 0 | 3.38807974 | 0.987 | 0.829 | 0 |
| Fibroblast | Cemip | 1 | 2.246E-211 | 3.52172138 | 0.749 | 0.3 | 4.492E-208 |
| Fibroblast | Adamtsl2 | 1 | 3.541E-197 | 1.94411193 | 0.813 | 0.381 | 7.082E-194 |
| Fibroblast | Grb14 | 1 | 4.166E-191 | 2.66867943 | 0.756 | 0.375 | 8.332E-188 |
| Fibroblast | Plxdc2 | 1 | 2.937E-190 | 1.20357815 | 0.965 | 0.799 | 5.874E-187 |
| Fibroblast | Nrg1 | 1 | 2.699E-178 | 2.98948138 | 0.708 | 0.301 | 5.398E-175 |
| Fibroblast | Pdlim3 | 1 | 7.394E-178 | 1.97601775 | 0.806 | 0.443 | 1.479E-174 |
| Fibroblast | Hmcn1 | 1 | 2.294E-174 | 1.87178715 | 0.835 | 0.458 | 4.589E-171 |
| Fibroblast | Nox4 | 1 | 1.051E-160 | 2.01247102 | 0.783 | 0.493 | 2.102E-157 |
| Fibroblast | Ccn2 | 1 | 1.522E-139 | 1.64439413 | 0.772 | 0.453 | 3.045E-136 |
| Fibroblast | Gdf6 | 1 | 1.648E-139 | 4.00338059 | 0.592 | 0.269 | 3.297E-136 |

| | | | | | | | |
|---|---|---|---|---|---|---|---|
| Fibroblast | Nrxn1 | 1 | 4.788E-134 | 1.42255324 | 0.83 | 0.532 | 9.576E-131 |
| Fibroblast | Klhl29 | 1 | 3.126E-133 | 1.85792208 | 0.726 | 0.405 | 6.252E-130 |
| Fibroblast | Ltbp2 | 1 | 2.923E-129 | 1.21450149 | 0.8 | 0.433 | 5.847E-126 |
| Fibroblast | AABR070635 | 1 | 3.023E-127 | 3.79626171 | 0.48 | 0.219 | 6.046E-124 |
| Fibroblast | Lbh | 1 | 3.344E-125 | 1.97129151 | 0.727 | 0.466 | 6.687E-122 |
| Fibroblast | Pde5a | 1 | 6.618E-119 | 1.91787462 | 0.715 | 0.41 | 1.324E-115 |
| Fibroblast | Pdgfrb | 1 | 2.247E-118 | 0.98615561 | 0.902 | 0.686 | 4.493E-115 |
| Fibroblast | Myh10 | 1 | 2.26E-111 | 1.34708728 | 0.812 | 0.511 | 4.52E-108 |
| Fibroblast | Boc | 1 | 6.044E-110 | 0.94000137 | 0.916 | 0.738 | 1.209E-106 |
| Fibroblast | Dkk3 | 1 | 2.473E-104 | 1.56445058 | 0.736 | 0.537 | 4.947E-101 |
| Fibroblast | Fshr | 1 | 5.66E-103 | 2.44075509 | 0.332 | 0.117 | 1.132E-99 |
| Fibroblast | Galnt16 | 1 | 7.122E-103 | 1.21780731 | 0.819 | 0.597 | 1.424E-99 |
| Fibroblast | Pcsk5 | 1 | 8.55E-102 | 1.48548007 | 0.785 | 0.575 | 1.71E-98 |
| Fibroblast | Col4a5 | 1 | 3.03E-100 | 1.18491039 | 0.878 | 0.712 | 6.0607E-97 |
| Fibroblast | Sv2c | 1 | 3.1014E-98 | 1.94674844 | 0.696 | 0.48 | 6.2027E-95 |
| Fibroblast | Smoc1 | 1 | 1.332E-93 | 3.41220923 | 0.382 | 0.17 | 2.664E-90 |
| Fibroblast | Dlg2 | 1 | 2.879E-93 | 0.94950148 | 0.897 | 0.732 | 5.758E-90 |
| Fibroblast | Cntn2 | 1 | 6.0933E-89 | 2.50638602 | 0.381 | 0.184 | 1.2187E-85 |
| Fibroblast | Ltbp1 | 1 | 3.0659E-88 | 0.95615542 | 0.865 | 0.644 | 6.1318E-85 |
| Fibroblast | AABR070109 | 1 | 3.2987E-87 | 0.63077955 | 0.286 | 0.061 | 6.5973E-84 |
| Fibroblast | Ccn5 | 1 | 4.6865E-87 | 2.15375502 | 0.622 | 0.4 | 9.373E-84 |
| Fibroblast | Pls3 | 1 | 1.4869E-83 | 0.9690853 | 0.859 | 0.656 | 2.9737E-80 |
| Fibroblast | Agtr1a | 1 | 1.7649E-82 | 2.01842796 | 0.637 | 0.403 | 3.5299E-79 |
| Fibroblast | Ttc29 | 1 | 1.9332E-79 | 0.72065018 | 0.28 | 0.11 | 3.8664E-76 |
| Fibroblast | Pdgfd | 1 | 5.0155E-76 | 1.48445364 | 0.683 | 0.51 | 1.0031E-72 |
| Fibroblast | Rspo3 | 1 | 9.8194E-76 | 0.93722437 | 0.267 | 0.058 | 1.9639E-72 |
| Fibroblast | Cthrc1 | 1 | 1.1691E-75 | 2.00555913 | 0.603 | 0.291 | 2.3382E-72 |
| Fibroblast | LOC689599 | 1 | 8.4968E-75 | 1.09702955 | 0.314 | 0.124 | 1.6994E-71 |
| Fibroblast | Glis3 | 1 | 2.8014E-73 | 0.76000446 | 0.908 | 0.751 | 5.6027E-70 |
| Fibroblast | Slc8a1 | 1 | 3.3697E-72 | 0.64135493 | 0.969 | 0.877 | 6.7394E-69 |
| Fibroblast | Smpd3 | 1 | 5.2335E-72 | 2.35252845 | 0.579 | 0.383 | 1.0467E-68 |

| | | | | | | | |
|---|---|---|---|---|---|---|---|
| Fibroblast | Mtss1 | 1 | 1.2165E-70 | 1.84200375 | 0.619 | 0.351 | 2.433E-67 |
| Fibroblast | Specc1 | 1 | 5.6872E-70 | 1.09780293 | 0.74 | 0.507 | 1.1374E-66 |
| Fibroblast | AABR070252 | 1 | 3.1518E-67 | 0.78027521 | 0.856 | 0.65 | 6.3036E-64 |
| Fibroblast | P4ha3 | 1 | 2.6712E-65 | 2.11866087 | 0.416 | 0.198 | 5.3423E-62 |
| Fibroblast | Gxylt2 | 1 | 5.8908E-65 | 0.87248802 | 0.853 | 0.703 | 1.1782E-61 |
| Fibroblast | Pappa1 | 1 | 5.3488E-62 | 1.2440219 | 0.631 | 0.404 | 1.0698E-58 |
| Fibroblast | Actn1 | 1 | 6.7035E-62 | 0.96993324 | 0.756 | 0.524 | 1.3407E-58 |
| Fibroblast | Itga8 | 1 | 1.3152E-61 | 0.94877332 | 0.762 | 0.578 | 2.6305E-58 |
| Fibroblast | Igf1 | 1 | 1.3745E-61 | 0.94498215 | 0.782 | 0.595 | 2.7489E-58 |
| Fibroblast | Pld5 | 1 | 2.6733E-58 | 2.27642034 | 0.526 | 0.28 | 5.3466E-55 |
| Fibroblast | Osr1 | 1 | 1.4629E-57 | 2.48134883 | 0.544 | 0.384 | 2.9258E-54 |
| Fibroblast | Fcrl1 | 1 | 2.5172E-57 | 1.29485121 | 0.323 | 0.102 | 5.0344E-54 |
| Fibroblast | Prdm6 | 1 | 1.1305E-55 | 1.77045464 | 0.603 | 0.415 | 2.261E-52 |
| Fibroblast | Arnt2 | 1 | 2.0989E-54 | 2.3086998 | 0.521 | 0.361 | 4.1977E-51 |
| Fibroblast | Fgr | 1 | 2.8531E-54 | 0.78142063 | 0.269 | 0.103 | 5.7062E-51 |
| Fibroblast | Bnc2 | 1 | 1.646E-52 | 0.68690621 | 0.879 | 0.729 | 3.2919E-49 |
| Fibroblast | Lmcd1 | 1 | 2.5018E-52 | 1.85625569 | 0.58 | 0.364 | 5.0036E-49 |
| Fibroblast | Fbln5 | 1 | 2.9717E-52 | 1.22979247 | 0.621 | 0.439 | 5.9434E-49 |
| Fibroblast | Pdzrn3 | 1 | 1.8974E-51 | 0.52212309 | 0.976 | 0.91 | 3.7949E-48 |
| Fibroblast | Rcan2 | 1 | 7.5003E-49 | 0.78904707 | 0.787 | 0.619 | 1.5001E-45 |
| Fibroblast | Svil | 1 | 1.4331E-48 | 0.81874024 | 0.781 | 0.627 | 2.8662E-45 |
| Fibroblast | Kif26b | 1 | 1.5535E-48 | 1.99002271 | 0.511 | 0.301 | 3.107E-45 |
| Fibroblast | Epha4 | 1 | 4.5862E-48 | 1.52417362 | 0.571 | 0.387 | 9.1724E-45 |
| Fibroblast | Col24a1 | 1 | 1.8771E-47 | 1.95582993 | 0.53 | 0.353 | 3.7542E-44 |
| Fibroblast | Itgbl1 | 1 | 3.5621E-47 | 0.49235319 | 0.95 | 0.869 | 7.1241E-44 |
| Fibroblast | Mrvi1 | 1 | 2.6175E-46 | 1.36131901 | 0.576 | 0.378 | 5.235E-43 |
| Fibroblast | Npr3 | 1 | 2.6121E-45 | 2.40619057 | 0.489 | 0.27 | 5.2243E-42 |
| Fibroblast | Spock3 | 1 | 4.5815E-45 | 2.08754862 | 0.314 | 0.114 | 9.163E-42 |
| Fibroblast | Ahr | 1 | 1.3202E-44 | 1.30035282 | 0.58 | 0.429 | 2.6405E-41 |
| Fibroblast | Ptprd | 1 | 1.4152E-44 | 0.71762447 | 0.811 | 0.653 | 2.8305E-41 |
| Fibroblast | Mid1 | 1 | 1.6336E-44 | 0.9739077 | 0.723 | 0.577 | 3.2672E-41 |

| | | | | | | | |
|---|---|---|---|---|---|---|---|
| Fibroblast | Tpm1 | 1 | 1.7143E-44 | 0.63786187 | 0.866 | 0.745 | 3.4287E-41 |
| Fibroblast | Ccn3 | 1 | 1.9426E-44 | 2.26428493 | 0.496 | 0.24 | 3.8853E-41 |
| Fibroblast | Bcl11b | 1 | 1.1236E-43 | 0.78136094 | 0.298 | 0.108 | 2.2471E-40 |
| Fibroblast | Srgap3 | 1 | 1.3221E-43 | 0.83362563 | 0.73 | 0.603 | 2.6443E-40 |
| Fibroblast | Abcc9 | 1 | 1.7985E-42 | 0.80664087 | 0.704 | 0.549 | 3.597E-39 |
| Fibroblast | Gramd1b | 1 | 2.2047E-42 | 1.40095412 | 0.58 | 0.452 | 4.4094E-39 |
| Fibroblast | Npnt | 1 | 3.1965E-42 | 1.81468128 | 0.348 | 0.186 | 6.3931E-39 |
| Fibroblast | Fhl2 | 1 | 2.1374E-41 | 1.29727407 | 0.566 | 0.419 | 4.2748E-38 |
| Fibroblast | Col1a1 | 1 | 3.2778E-41 | 0.43442046 | 0.967 | 0.863 | 6.5556E-38 |
| Fibroblast | AABR070311 | 1 | 4.331E-41 | 2.10960365 | 0.398 | 0.188 | 8.6619E-38 |
| Fibroblast | Vegfd | 1 | 1.3375E-39 | 1.39721735 | 0.578 | 0.402 | 2.675E-36 |
| Fibroblast | Ska1 | 1 | 2.8219E-39 | 0.94052719 | 0.264 | 0.09 | 5.6438E-36 |
| Fibroblast | Atp1b1 | 1 | 1.1753E-38 | 1.85694756 | 0.49 | 0.319 | 2.3507E-35 |
| Fibroblast | Ddah1 | 1 | 2.1557E-38 | 1.33684077 | 0.557 | 0.374 | 4.3113E-35 |
| Fibroblast | Tmem17 | 1 | 2.1101E-37 | 1.74162467 | 0.49 | 0.344 | 4.2203E-34 |
| Fibroblast | Tnfsf15 | 1 | 4.4343E-37 | 2.00857983 | 0.316 | 0.147 | 8.8686E-34 |
| Fibroblast | Rgs7bp | 1 | 1.326E-36 | 2.00060523 | 0.273 | 0.198 | 2.6519E-33 |
| Fibroblast | Pmepa1 | 1 | 2.1759E-35 | 0.72591914 | 0.746 | 0.551 | 4.3518E-32 |
| Fibroblast | Aldh1a2 | 1 | 1.1526E-33 | 1.87153776 | 0.483 | 0.341 | 2.3053E-30 |
| Fibroblast | Itgb6 | 1 | 1.4112E-33 | 1.50731486 | 0.268 | 0.165 | 2.8223E-30 |
| Fibroblast | AC118957.1 | 1 | 1.7031E-33 | 1.61088735 | 0.371 | 0.233 | 3.4063E-30 |
| Fibroblast | Setbp1 | 1 | 2.2081E-33 | 0.7439487 | 0.701 | 0.547 | 4.4162E-30 |
| Fibroblast | Hecw2 | 1 | 2.9186E-33 | 1.03253459 | 0.565 | 0.36 | 5.8373E-30 |
| Fibroblast | Zfp385d | 1 | 5.7809E-33 | 0.45210694 | 0.947 | 0.894 | 1.1562E-29 |
| Fibroblast | Tmem82 | 1 | 1.5714E-32 | 2.28483391 | 0.47 | 0.296 | 3.1428E-29 |
| Fibroblast | Insc | 1 | 6.0411E-32 | 0.78689971 | 0.388 | 0.148 | 1.2082E-28 |
| Fibroblast | Sorbs1 | 1 | 1.2745E-31 | 0.5537642 | 0.785 | 0.642 | 2.549E-28 |
| Fibroblast | Niban1 | 1 | 6.9356E-31 | 0.88874221 | 0.684 | 0.524 | 1.3871E-27 |
| Fibroblast | Slc1a7 | 1 | 1.3928E-30 | 1.19090439 | 0.573 | 0.482 | 2.7855E-27 |
| Fibroblast | Timp3 | 1 | 2.7956E-30 | 1.05505633 | 0.64 | 0.474 | 5.5913E-27 |
| Fibroblast | Ptk2b | 1 | 3.4341E-30 | 1.35319098 | 0.571 | 0.435 | 6.8683E-27 |

| | | | | | | | |
|---|---|---|---|---|---|---|---|
| Fibroblast | Col3a1 | 1 | 1.5425E-29 | 0.28160828 | 0.99 | 0.965 | 3.085E-26 |
| Fibroblast | Ptger3 | 1 | 6.5285E-29 | 1.76463818 | 0.359 | 0.248 | 1.3057E-25 |
| Fibroblast | Hpgd | 1 | 1.2249E-28 | 1.02646558 | 0.364 | 0.258 | 2.4498E-25 |
| Fibroblast | Vgll3 | 1 | 1.2332E-28 | 1.37391776 | 0.527 | 0.397 | 2.4664E-25 |
| Fibroblast | Pam | 1 | 1.8377E-28 | 0.52844505 | 0.783 | 0.636 | 3.6753E-25 |
| Fibroblast | Slc1a3 | 1 | 9.8964E-28 | 1.47567146 | 0.47 | 0.253 | 1.9793E-24 |
| Fibroblast | Slc24a3 | 1 | 1.2972E-27 | 1.21419013 | 0.526 | 0.391 | 2.5944E-24 |
| Fibroblast | Fhl1 | 1 | 2.9693E-27 | 0.73814658 | 0.676 | 0.531 | 5.9385E-24 |
| Fibroblast | AABR070707 | 1 | 6.6974E-27 | 2.00984315 | 0.254 | 0.104 | 1.3395E-23 |
| Fibroblast | Arhgap44 | 1 | 7.2222E-27 | 1.33107257 | 0.513 | 0.338 | 1.4444E-23 |
| Fibroblast | Ralgapa2 | 1 | 8.5852E-27 | 0.75771942 | 0.617 | 0.461 | 1.717E-23 |
| Fibroblast | Svep1 | 1 | 9.4435E-27 | 0.97096748 | 0.575 | 0.464 | 1.8887E-23 |
| Fibroblast | C1qtnf3 | 1 | 2.0101E-26 | 1.23699275 | 0.427 | 0.216 | 4.0201E-23 |
| Fibroblast | Vcl | 1 | 4.7071E-26 | 0.53178635 | 0.7 | 0.534 | 9.4143E-23 |
| Fibroblast | Tgfb2 | 1 | 6.2011E-26 | 0.84611669 | 0.617 | 0.527 | 1.2402E-22 |
| Fibroblast | Cpxm2 | 1 | 8.8545E-26 | 2.16389299 | 0.425 | 0.221 | 1.7709E-22 |
| Fibroblast | Gpc6 | 1 | 4.9168E-25 | 0.26579141 | 0.992 | 0.978 | 9.8337E-22 |
| Fibroblast | Kif18b | 1 | 1.0293E-24 | 0.41061767 | 0.251 | 0.139 | 2.0586E-21 |
| Fibroblast | Lmod1 | 1 | 1.0373E-24 | 1.13015983 | 0.516 | 0.332 | 2.0745E-21 |
| Fibroblast | Shc4 | 1 | 2.3835E-24 | 0.51293482 | 0.296 | 0.08 | 4.7669E-21 |
| Fibroblast | Enpp1 | 1 | 3.7544E-24 | 0.59072938 | 0.647 | 0.526 | 7.5087E-21 |
| Fibroblast | Aff3 | 1 | 4.9279E-24 | 0.41259822 | 0.79 | 0.68 | 9.8557E-21 |
| Fibroblast | Tbx18 | 1 | 6.0564E-24 | 0.74534081 | 0.616 | 0.451 | 1.2113E-20 |
| Fibroblast | Smad6 | 1 | 6.1212E-24 | 0.95540724 | 0.567 | 0.41 | 1.2242E-20 |
| Fibroblast | Samd5 | 1 | 6.3677E-24 | 1.76092921 | 0.392 | 0.141 | 1.2735E-20 |
| Fibroblast | Tenm3 | 1 | 6.5051E-24 | 0.59655219 | 0.686 | 0.559 | 1.301E-20 |
| Fibroblast | Scg2 | 1 | 7.8871E-24 | 3.36804077 | 0.347 | 0.183 | 1.5774E-20 |
| Fibroblast | Itgb8 | 1 | 9.1525E-24 | 0.96467062 | 0.561 | 0.434 | 1.8305E-20 |
| Fibroblast | Asap3 | 1 | 1.3245E-23 | 0.5564088 | 0.765 | 0.666 | 2.6489E-20 |
| Fibroblast | Tgfb3 | 1 | 1.4946E-23 | 0.61514523 | 0.649 | 0.496 | 2.9891E-20 |
| Fibroblast | Plxna4 | 1 | 1.6618E-23 | 0.7759731 | 0.548 | 0.377 | 3.3236E-20 |

| | | | | | | | |
|---|---|---|---|---|---|---|---|
| Fibroblast | Vegfc | 1 | 5.0308E-23 | 1.06725329 | 0.51 | 0.376 | 1.0062E-19 |
| Fibroblast | Fn1 | 1 | 1.6251E-22 | 0.25253973 | 0.762 | 0.611 | 3.2501E-19 |
| Fibroblast | Pappa2 | 1 | 2.3608E-22 | 0.6081294 | 0.343 | 0.229 | 4.7215E-19 |
| Fibroblast | Gria3 | 1 | 2.4812E-22 | 0.45078429 | 0.746 | 0.596 | 4.9624E-19 |
| Fibroblast | Tspan5 | 1 | 3.3643E-22 | 0.46544411 | 0.717 | 0.551 | 6.7286E-19 |
| Fibroblast | Adamts6 | 1 | 4.4547E-22 | 0.47832513 | 0.757 | 0.619 | 8.9093E-19 |
| Fibroblast | Map3k7cl | 1 | 7.5078E-22 | 1.47140164 | 0.307 | 0.14 | 1.5016E-18 |
| Fibroblast | Rnd3 | 1 | 4.4102E-21 | 0.71419026 | 0.637 | 0.527 | 8.8203E-18 |
| Fibroblast | Tnfrsf12a | 1 | 6.1637E-21 | 0.28082983 | 0.354 | 0.177 | 1.2327E-17 |
| Fibroblast | Dpt | 1 | 7.2487E-21 | 0.5069014 | 0.794 | 0.712 | 1.4497E-17 |
| Fibroblast | Cit | 1 | 1.191E-20 | 0.99956237 | 0.486 | 0.366 | 2.3819E-17 |
| Fibroblast | Mylk | 1 | 1.5632E-20 | 0.55985874 | 0.679 | 0.557 | 3.1265E-17 |
| Fibroblast | Clmp | 1 | 4.8391E-20 | 0.34477098 | 0.887 | 0.796 | 9.6781E-17 |
| Fibroblast | Map1b | 1 | 5.0207E-20 | 0.44621928 | 0.872 | 0.788 | 1.0041E-16 |
| Fibroblast | Sdk2 | 1 | 8.7579E-20 | 0.60033957 | 0.677 | 0.566 | 1.7516E-16 |
| Fibroblast | Ntf31 | 1 | 1.1387E-19 | 1.03115547 | 0.46 | 0.319 | 2.2774E-16 |
| Fibroblast | Fat3 | 1 | 1.8778E-18 | 0.94906928 | 0.509 | 0.37 | 3.7556E-15 |
| Fibroblast | Cdkl1 | 1 | 3.5713E-18 | 0.63576318 | 0.306 | 0.163 | 7.1425E-15 |
| Fibroblast | Plekha7 | 1 | 8.4811E-18 | 0.43946306 | 0.729 | 0.658 | 1.6962E-14 |
| Fibroblast | Baiap2l1 | 1 | 1.7171E-17 | 1.66910211 | 0.421 | 0.262 | 3.4342E-14 |
| Fibroblast | Fhod3 | 1 | 3.8957E-17 | 0.67045447 | 0.553 | 0.423 | 7.7914E-14 |
| Fibroblast | Col8a2 | 1 | 2.1819E-16 | 1.0704367 | 0.502 | 0.42 | 4.3639E-13 |
| Fibroblast | Tpm2 | 1 | 2.4006E-16 | 0.79911779 | 0.444 | 0.246 | 4.8013E-13 |
| Fibroblast | Prrx1 | 1 | 2.4235E-16 | 0.39090116 | 0.796 | 0.672 | 4.8471E-13 |
| Fibroblast | Fgf9 | 1 | 7.7075E-16 | 2.28963515 | 0.343 | 0.243 | 1.5415E-12 |
| Fibroblast | Ccn4 | 1 | 9.7542E-16 | 1.47157023 | 0.364 | 0.241 | 1.9508E-12 |
| Fibroblast | Nr1h4 | 1 | 1.1103E-15 | 1.58981282 | 0.431 | 0.349 | 2.2206E-12 |
| Fibroblast | Adk1 | 1 | 1.3867E-15 | 0.43211098 | 0.584 | 0.421 | 2.7734E-12 |
| Fibroblast | Rai14 | 1 | 3.325E-15 | 0.32870376 | 0.638 | 0.466 | 6.65E-12 |
| Fibroblast | Sytl2 | 1 | 4.2421E-15 | 1.74011855 | 0.294 | 0.173 | 8.4843E-12 |
| Fibroblast | AABR070605 | 1 | 4.6196E-15 | 1.14528342 | 0.427 | 0.311 | 9.2393E-12 |

| | | | | | | | |
|---|---|---|---|---|---|---|---|
| Fibroblast | Pak3 | 1 | 1.4124E-14 | 2.63118711 | 0.301 | 0.274 | 2.8248E-11 |
| Fibroblast | Kcnma1 | 1 | 4.9528E-14 | 2.06926154 | 0.339 | 0.286 | 9.9055E-11 |
| Fibroblast | Kank1 | 1 | 1.191E-13 | 0.64994406 | 0.494 | 0.403 | 2.3819E-10 |
| Fibroblast | Tyrobp | 1 | 1.4099E-13 | 0.33551617 | 0.303 | 0.201 | 2.8198E-10 |
| Fibroblast | Plbd1 | 1 | 1.5002E-13 | 0.73330985 | 0.324 | 0.201 | 3.0003E-10 |
| Fibroblast | Plod2 | 1 | 1.642E-13 | 0.29445009 | 0.844 | 0.742 | 3.2841E-10 |
| Fibroblast | Slc7a2 | 1 | 1.7101E-13 | 0.61401773 | 0.508 | 0.375 | 3.4201E-10 |
| Fibroblast | Mgmt | 1 | 4.9008E-13 | 0.32559882 | 0.585 | 0.462 | 9.8016E-10 |
| Fibroblast | Greb1l | 1 | 5.04E-13 | 1.18570742 | 0.467 | 0.365 | 1.008E-09 |
| Fibroblast | Ephb2 | 1 | 5.6406E-13 | 0.62123292 | 0.484 | 0.392 | 1.1281E-09 |
| Fibroblast | Snta1 | 1 | 6.375E-13 | 0.62792194 | 0.505 | 0.397 | 1.275E-09 |
| Fibroblast | Chrdl1 | 1 | 7.5135E-13 | 1.73511232 | 0.315 | 0.195 | 1.5027E-09 |
| Fibroblast | Sntg2 | 1 | 1.0153E-12 | 1.25366032 | 0.403 | 0.317 | 2.0305E-09 |
| Fibroblast | Mpped21 | 1 | 1.7404E-12 | 0.40320278 | 0.619 | 0.496 | 3.4808E-09 |
| Fibroblast | Nexn1 | 1 | 4.2211E-12 | 0.42425917 | 0.564 | 0.441 | 8.4422E-09 |
| Fibroblast | Palmd | 1 | 7.1023E-12 | 0.85724648 | 0.444 | 0.297 | 1.4205E-08 |
| Fibroblast | Slco5a11 | 1 | 8.2368E-12 | 0.91825335 | 0.406 | 0.275 | 1.6474E-08 |
| Fibroblast | Gask1b | 1 | 1.6714E-11 | 0.32259567 | 0.672 | 0.544 | 3.3428E-08 |
| Fibroblast | Dmpk | 1 | 1.6722E-11 | 0.41404317 | 0.606 | 0.52 | 3.3444E-08 |
| Fibroblast | Edar | 1 | 2.7016E-11 | 0.71799459 | 0.353 | 0.318 | 5.4032E-08 |
| Fibroblast | Rxfp1 | 1 | 3.3207E-11 | 0.89676855 | 0.281 | 0.133 | 6.6414E-08 |
| Fibroblast | Ednra | 1 | 3.3827E-11 | 0.65598233 | 0.542 | 0.519 | 6.7654E-08 |
| Fibroblast | Angpt2 | 1 | 3.6187E-11 | 1.37495035 | 0.362 | 0.227 | 7.2373E-08 |
| Fibroblast | AABR070215 | 1 | 3.8033E-11 | 1.6790665 | 0.338 | 0.356 | 7.6066E-08 |
| Fibroblast | Piezo2 | 1 | 4.549E-11 | 0.33509281 | 0.535 | 0.427 | 9.0981E-08 |
| Fibroblast | Sgcd | 1 | 6.8806E-11 | 0.49912069 | 0.558 | 0.457 | 1.3761E-07 |
| Fibroblast | Gna14 | 1 | 8.9275E-11 | 1.31357992 | 0.341 | 0.259 | 1.7855E-07 |
| Fibroblast | Cdh2 | 1 | 1.2302E-10 | 0.49089566 | 0.598 | 0.546 | 2.4604E-07 |
| Fibroblast | Tnfrsf11b | 1 | 1.736E-10 | 0.73441156 | 0.48 | 0.396 | 3.472E-07 |
| Fibroblast | Npas2 | 1 | 1.947E-10 | 0.28322813 | 0.687 | 0.596 | 3.894E-07 |
| Fibroblast | Dgki | 1 | 2.0956E-10 | 0.79732591 | 0.421 | 0.28 | 4.1912E-07 |

| | | | | | | | |
|---|---|---|---|---|---|---|---|
| Fibroblast | Pcdha13 | 1 | 2.5859E-10 | 0.30440235 | 0.7 | 0.619 | 5.1717E-07 |
| Fibroblast | Egfl8 | 1 | 4.1579E-10 | 0.87206119 | 0.322 | 0.185 | 8.3158E-07 |
| Fibroblast | Efnb2 | 1 | 5.8466E-10 | 0.60333392 | 0.39 | 0.225 | 1.1693E-06 |
| Fibroblast | Actn21 | 1 | 9.6594E-10 | 0.70010291 | 0.472 | 0.357 | 1.9319E-06 |
| Fibroblast | Alcam | 1 | 1.0354E-09 | 0.46689403 | 0.429 | 0.313 | 2.0707E-06 |
| Fibroblast | Sfrp2 | 1 | 1.764E-09 | 1.12908015 | 0.375 | 0.244 | 3.5279E-06 |
| Fibroblast | Tmem196 | 1 | 1.998E-09 | 0.44591391 | 0.522 | 0.393 | 3.9961E-06 |
| Fibroblast | Lsamp | 1 | 2.342E-09 | 0.38239578 | 0.126 | 0.259 | 4.684E-06 |
| Fibroblast | Slc16a10 | 1 | 3.1966E-09 | 1.90442522 | 0.332 | 0.225 | 6.3931E-06 |
| Fibroblast | Ndst3 | 1 | 3.7854E-09 | 0.52193481 | 0.445 | 0.359 | 7.5707E-06 |
| Fibroblast | Enah | 1 | 4.2388E-09 | 0.25643911 | 0.595 | 0.45 | 8.4777E-06 |
| Fibroblast | Cobll1 | 1 | 5.3483E-09 | 0.522161 | 0.453 | 0.327 | 1.0697E-05 |
| Fibroblast | Olfml2b | 1 | 5.874E-09 | 0.4481635 | 0.485 | 0.402 | 1.1748E-05 |
| Fibroblast | LOC1003610 | 1 | 7.1268E-09 | 0.56537133 | 0.406 | 0.257 | 1.4254E-05 |
| Fibroblast | Slc20a2 | 1 | 1.2836E-08 | 0.40255944 | 0.538 | 0.415 | 2.5673E-05 |
| Fibroblast | Mmp16 | 1 | 2.6092E-08 | 0.69946989 | 0.421 | 0.295 | 5.2183E-05 |
| Fibroblast | Cacna2d3 | 1 | 2.6437E-08 | 0.50313608 | 0.658 | 0.643 | 5.2873E-05 |
| Fibroblast | Slc16a2 | 1 | 2.8415E-08 | 0.61141018 | 0.494 | 0.421 | 5.683E-05 |
| Fibroblast | Loxl1 | 1 | 4.2995E-08 | 0.26236625 | 0.756 | 0.681 | 8.5989E-05 |
| Fibroblast | Epb41l4b | 1 | 4.3816E-08 | 0.97355435 | 0.382 | 0.264 | 8.7632E-05 |
| Fibroblast | Pawr | 1 | 5.0248E-08 | 1.09929906 | 0.338 | 0.246 | 0.0001005 |
| Fibroblast | Dtna | 1 | 6.7461E-08 | 1.160907 | 0.341 | 0.241 | 0.00013492 |
| Fibroblast | AABR070349 | 1 | 8.6855E-08 | 0.80610943 | 0.353 | 0.253 | 0.00017371 |
| Fibroblast | Galnt17 | 1 | 1.0258E-07 | 0.28551823 | 0.717 | 0.662 | 0.00020516 |
| Fibroblast | Crim1 | 1 | 1.0846E-07 | 0.4714586 | 0.465 | 0.368 | 0.00021691 |
| Fibroblast | Cabcoco11 | 1 | 1.6929E-07 | 0.26302648 | 0.416 | 0.252 | 0.00033858 |
| Fibroblast | Alpl | 1 | 2.3755E-07 | 0.61388732 | 0.345 | 0.204 | 0.0004751 |
| Fibroblast | Wdfy4 | 1 | 2.5615E-07 | 1.3190781 | 0.377 | 0.342 | 0.0005123 |
| Fibroblast | Ano2 | 1 | 3.1642E-07 | 0.31607202 | 0.275 | 0.168 | 0.00063285 |
| Fibroblast | Col12a1 | 1 | 3.225E-07 | 0.52636568 | 0.373 | 0.277 | 0.00064501 |
| Fibroblast | Sntb1 | 1 | 3.2541E-07 | 0.80706017 | 0.384 | 0.288 | 0.00065082 |

| | | | | | | | |
|---|---|---|---|---|---|---|---|
| Fibroblast | Pfkfb3 | 1 | 3.2744E-07 | 0.9138182 | 0.412 | 0.335 | 0.00065489 |
| Fibroblast | Cacna1c | 1 | 3.4371E-07 | 0.25966823 | 0.579 | 0.522 | 0.00068742 |
| Fibroblast | Gpr176 | 1 | 3.6369E-07 | 0.73620603 | 0.419 | 0.347 | 0.00072739 |
| Fibroblast | Flrt2 | 1 | 4.3772E-07 | 0.3829369 | 0.563 | 0.507 | 0.00087545 |
| Fibroblast | Acer21 | 1 | 5.5132E-07 | 0.68623656 | 0.38 | 0.256 | 0.00110263 |
| Fibroblast | Ildr2 | 1 | 9.4229E-07 | 0.47133698 | 0.415 | 0.316 | 0.00188458 |
| Fibroblast | Apba1 | 1 | 1.0781E-06 | 0.36001345 | 0.62 | 0.558 | 0.0021561 |
| Fibroblast | Cacna1e | 1 | 1.1029E-06 | 0.42287881 | 0.495 | 0.437 | 0.00220572 |
| Fibroblast | Pde7a | 1 | 1.3428E-06 | 0.33495248 | 0.65 | 0.591 | 0.00268561 |
| Fibroblast | Slit3 | 1 | 1.5623E-06 | 0.39551887 | 0.542 | 0.473 | 0.00312468 |
| Fibroblast | Ncald | 1 | 2.0955E-06 | 0.44312437 | 0.487 | 0.383 | 0.00419107 |
| Fibroblast | Serpine1 | 1 | 2.7735E-06 | 0.41804002 | 0.398 | 0.341 | 0.00554696 |
| Fibroblast | Abca1 | 1 | 9.0148E-06 | 0.26287857 | 0.597 | 0.513 | 0.01802965 |
| Fibroblast | AABR070581 | 1 | 9.6834E-06 | 0.98425799 | 0.298 | 0.23 | 0.01936674 |
| Fibroblast | Rapgef51 | 1 | 1.0154E-05 | 0.58922141 | 0.364 | 0.256 | 0.02030803 |
| Fibroblast | Trabd2b | 1 | 1.298E-05 | 0.33719034 | 0.354 | 0.223 | 0.02596098 |
| Fibroblast | C7 | 2 | 2.901E-108 | 0.85674001 | 0.973 | 0.89 | 5.802E-105 |
| Fibroblast | Fbln1 | 2 | 8.1173E-78 | 1.18926314 | 0.772 | 0.596 | 1.6235E-74 |
| Fibroblast | Boc1 | 2 | 9.8983E-76 | 0.88191392 | 0.877 | 0.744 | 1.9797E-72 |
| Fibroblast | Zfp385d1 | 2 | 1.5671E-60 | 0.64011169 | 0.947 | 0.894 | 3.1342E-57 |
| Fibroblast | Ccdc80 | 2 | 5.9742E-60 | 0.66114209 | 0.953 | 0.887 | 1.1948E-56 |
| Fibroblast | Pid1 | 2 | 2.1372E-39 | 0.67976707 | 0.859 | 0.796 | 4.2745E-36 |
| Fibroblast | Dlg21 | 2 | 2.647E-38 | 0.68628822 | 0.833 | 0.742 | 5.2939E-35 |
| Fibroblast | Srgap31 | 2 | 4.0491E-34 | 0.94068926 | 0.697 | 0.608 | 8.0982E-31 |
| Fibroblast | Plcxd3 | 2 | 5.0647E-32 | 1.88258732 | 0.514 | 0.317 | 1.0129E-28 |
| Fibroblast | Abcc91 | 2 | 1.0913E-26 | 0.89857925 | 0.654 | 0.557 | 2.1825E-23 |
| Fibroblast | Adamts15 | 2 | 2.6759E-25 | 1.32058669 | 0.557 | 0.401 | 5.3518E-22 |
| Fibroblast | Plxdc21 | 2 | 1.0122E-24 | 0.45092138 | 0.88 | 0.811 | 2.0243E-21 |
| Fibroblast | Igf11 | 2 | 1.0643E-24 | 0.61548419 | 0.716 | 0.605 | 2.1286E-21 |
| Fibroblast | Cacna2d31 | 2 | 4.5602E-22 | 0.67861284 | 0.714 | 0.636 | 9.1204E-19 |
| Fibroblast | Fgf7 | 2 | 6.1375E-19 | 0.40870686 | 0.277 | 0.151 | 1.2275E-15 |

| | | | | | | | |
|---|---|---|---|---|---|---|---|
| Fibroblast | Pdgfra | 2 | 2.1278E-18 | 0.42003593 | 0.813 | 0.764 | 4.2556E-15 |
| Fibroblast | Col6a61 | 2 | 3.5322E-18 | 0.42284935 | 0.794 | 0.73 | 7.0644E-15 |
| Fibroblast | Slc8a11 | 2 | 6.9892E-16 | 0.35856602 | 0.923 | 0.883 | 1.3978E-12 |
| Fibroblast | Bnc21 | 2 | 1.5689E-15 | 0.34379821 | 0.797 | 0.741 | 3.1377E-12 |
| Fibroblast | Fat31 | 2 | 2.8349E-15 | 1.20552994 | 0.528 | 0.369 | 5.6698E-12 |
| Fibroblast | Gxylt21 | 2 | 6.0155E-15 | 0.42214195 | 0.786 | 0.713 | 1.2031E-11 |
| Fibroblast | Rcan21 | 2 | 7.3306E-15 | 0.57716002 | 0.696 | 0.632 | 1.4661E-11 |
| Fibroblast | Pcsk51 | 2 | 2.7901E-14 | 0.56960287 | 0.678 | 0.591 | 5.5801E-11 |
| Fibroblast | Frmd3 | 2 | 2.8798E-14 | 0.64252098 | 0.175 | 0.261 | 5.7596E-11 |
| Fibroblast | Lmln | 2 | 3.521E-14 | 0.50871115 | 0.176 | 0.304 | 7.042E-11 |
| Fibroblast | Mgp | 2 | 1.4436E-13 | 0.31774974 | 0.849 | 0.796 | 2.8873E-10 |
| Fibroblast | Aff31 | 2 | 3.2708E-13 | 0.41619658 | 0.719 | 0.69 | 6.5415E-10 |
| Fibroblast | Tgfb31 | 2 | 3.6556E-13 | 0.88498148 | 0.58 | 0.506 | 7.3111E-10 |
| Fibroblast | Postn | 2 | 5.5018E-13 | 0.26688952 | 0.824 | 0.77 | 1.1004E-09 |
| Fibroblast | Wnk21 | 2 | 7.1568E-13 | 0.33485433 | 0.191 | 0.311 | 1.4314E-09 |
| Fibroblast | Mis18bp1 | 2 | 2.2253E-12 | 0.34458261 | 0.191 | 0.294 | 4.4506E-09 |
| Fibroblast | Map1b1 | 2 | 6.4802E-12 | 0.30355357 | 0.839 | 0.793 | 1.296E-08 |
| Fibroblast | Vash2 | 2 | 1.199E-11 | 0.47935099 | 0.107 | 0.258 | 2.398E-08 |
| Fibroblast | Glis31 | 2 | 4.6035E-11 | 0.31932966 | 0.802 | 0.766 | 9.207E-08 |
| Fibroblast | Tc2n | 2 | 1.1522E-10 | 0.32561723 | 0.235 | 0.349 | 2.3044E-07 |
| Fibroblast | Pdgfrb1 | 2 | 2.3218E-10 | 0.35039786 | 0.736 | 0.709 | 4.6435E-07 |
| Fibroblast | Itga81 | 2 | 3.5419E-10 | 0.46926859 | 0.656 | 0.593 | 7.0839E-07 |
| Fibroblast | Mrc2 | 2 | 1.0275E-09 | 0.34609047 | 0.769 | 0.727 | 2.055E-06 |
| Fibroblast | Coq8a1 | 2 | 1.9423E-09 | 0.3019183 | 0.205 | 0.288 | 3.8846E-06 |
| Fibroblast | Trim551 | 2 | 5.7843E-09 | 0.63724411 | 0.182 | 0.252 | 1.1569E-05 |
| Fibroblast | Tmeff2 | 2 | 7.4187E-09 | 0.3177101 | 0.729 | 0.684 | 1.4837E-05 |
| Fibroblast | Hmox1 | 2 | 1.4486E-08 | 0.42502935 | 0.265 | 0.174 | 2.8973E-05 |
| Fibroblast | Slc1a71 | 2 | 2.1267E-08 | 0.92789052 | 0.513 | 0.491 | 4.2534E-05 |
| Fibroblast | Pde1a | 2 | 3.234E-08 | 0.44038718 | 0.704 | 0.654 | 6.468E-05 |
| Fibroblast | Ltbp11 | 2 | 3.3893E-08 | 0.27843701 | 0.703 | 0.666 | 6.7786E-05 |
| Fibroblast | Hcn11 | 2 | 6.1334E-08 | 0.28119468 | 0.183 | 0.273 | 0.00012267 |

| | | | | | | | |
|---|---|---|---|---|---|---|---|
| Fibroblast | Prima1 | 2 | 1.872E-07 | 0.41042656 | 0.216 | 0.279 | 0.00037439 |
| Fibroblast | Lrrtm4 | 2 | 1.9384E-07 | 0.42617353 | 0.338 | 0.279 | 0.00038768 |
| Fibroblast | Tshr | 2 | 2.1414E-07 | 0.44714066 | 0.25 | 0.342 | 0.00042829 |
| Fibroblast | Ncam1 | 2 | 2.3545E-07 | 0.68946833 | 0.557 | 0.497 | 0.00047089 |
| Fibroblast | Dsg21 | 2 | 3.6216E-07 | 0.70578932 | 0.181 | 0.258 | 0.00072432 |
| Fibroblast | Otulinl | 2 | 4.1329E-07 | 0.48808553 | 0.171 | 0.263 | 0.00082658 |
| Fibroblast | Il16 | 2 | 4.6446E-07 | 0.70643815 | 0.558 | 0.53 | 0.00092892 |
| Fibroblast | Fam227b | 2 | 1.1602E-06 | 0.34814507 | 0.277 | 0.319 | 0.00232043 |
| Fibroblast | Tmem821 | 2 | 1.627E-06 | 0.40371338 | 0.255 | 0.326 | 0.00325408 |
| Fibroblast | Unc45b1 | 2 | 2.5884E-06 | 0.42447588 | 0.234 | 0.297 | 0.00517674 |
| Fibroblast | Dpysl3 | 2 | 5.8406E-06 | 0.29310724 | 0.774 | 0.742 | 0.01168128 |
| Fibroblast | Fgfr2 | 2 | 6.6303E-06 | 0.29395781 | 0.273 | 0.32 | 0.01326053 |
| Fibroblast | Smpx1 | 2 | 8.7568E-06 | 0.39658856 | 0.243 | 0.282 | 0.01751363 |
| Fibroblast | Edar1 | 2 | 8.8925E-06 | 0.71335928 | 0.375 | 0.316 | 0.01778509 |
| Fibroblast | Col20a1 | 2 | 8.9695E-06 | 0.46307453 | 0.246 | 0.277 | 0.01793893 |
| Fibroblast | Plxna41 | 2 | 8.9708E-06 | 0.6080505 | 0.484 | 0.387 | 0.01794164 |
| Fibroblast | Slc38a1 | 2 | 1.0784E-05 | 0.39469265 | 0.316 | 0.36 | 0.02156744 |
| Fibroblast | Ndst31 | 2 | 1.0888E-05 | 0.30240669 | 0.298 | 0.379 | 0.02177594 |
| Fibroblast | Eva1c | 2 | 1.1806E-05 | 0.56013994 | 0.256 | 0.317 | 0.02361293 |
| Fibroblast | Adamtsl21 | 2 | 1.4702E-05 | 0.41709575 | 0.498 | 0.425 | 0.02940326 |
| Fibroblast | Tmem26 | 2 | 1.6581E-05 | 0.41352244 | 0.284 | 0.367 | 0.0331625 |
| Fibroblast | Setbp11 | 2 | 1.825E-05 | 0.53803809 | 0.606 | 0.56 | 0.03650002 |
| Fibroblast | Adamts61 | 2 | 2.0585E-05 | 0.29406088 | 0.648 | 0.634 | 0.04116901 |
| Fibroblast | Prrx11 | 2 | 2.4002E-05 | 0.31747977 | 0.707 | 0.684 | 0.04800459 |
| Fibroblast | Trdn1 | 2 | 2.4843E-05 | 0.32225263 | 0.237 | 0.288 | 0.04968554 |
| Fibroblast | Uap1 | 3 | 0 | 2.54227303 | 0.966 | 0.589 | 0 |
| Fibroblast | Fbn1 | 3 | 2.06E-303 | 1.89756252 | 0.998 | 0.932 | 4.12E-300 |
| Fibroblast | Adgrd1 | 3 | 6.465E-252 | 1.92300448 | 0.932 | 0.598 | 1.293E-248 |
| Fibroblast | Fscn2 | 3 | 2.241E-242 | 2.0661477 | 0.308 | 0.028 | 4.483E-239 |
| Fibroblast | Gfpt2 | 3 | 2.256E-237 | 2.77463472 | 0.895 | 0.544 | 4.512E-234 |
| Fibroblast | Cd55 | 3 | 2.704E-217 | 4.05802606 | 0.724 | 0.275 | 5.407E-214 |

| | | | | | | | |
|---|---|---|---|---|---|---|---|
| Fibroblast | Mfap5 | 3 | 3.455E-210 | 1.76086418 | 0.935 | 0.673 | 6.91E-207 |
| Fibroblast | Fstl1 | 3 | 4.253E-209 | 1.3482896 | 0.979 | 0.803 | 8.507E-206 |
| Fibroblast | Creb5 | 3 | 1.944E-198 | 1.46701122 | 0.971 | 0.754 | 3.887E-195 |
| Fibroblast | Cdh13 | 3 | 3.32E-193 | 2.04617444 | 0.873 | 0.55 | 6.64E-190 |
| Fibroblast | Scara5 | 3 | 9.029E-191 | 2.08110045 | 0.843 | 0.471 | 1.806E-187 |
| Fibroblast | Lurap1l | 3 | 6.377E-182 | 2.87559131 | 0.764 | 0.402 | 1.275E-178 |
| Fibroblast | Gap43 | 3 | 2.719E-177 | 4.50649669 | 0.624 | 0.304 | 5.437E-174 |
| Fibroblast | Efhd1 | 3 | 3.224E-172 | 3.59569332 | 0.673 | 0.337 | 6.448E-169 |
| Fibroblast | Pi16 | 3 | 1.394E-168 | 2.22490374 | 0.806 | 0.419 | 2.787E-165 |
| Fibroblast | Axl | 3 | 2.259E-147 | 1.29709215 | 0.928 | 0.656 | 4.519E-144 |
| Fibroblast | Limch1 | 3 | 4.996E-140 | 2.40096367 | 0.739 | 0.367 | 9.991E-137 |
| Fibroblast | Sema3c | 3 | 1.173E-138 | 1.93612646 | 0.774 | 0.412 | 2.345E-135 |
| Fibroblast | Plaur | 3 | 2.166E-137 | 2.25364342 | 0.727 | 0.419 | 4.332E-134 |
| Fibroblast | Tiam1 | 3 | 5.623E-137 | 3.6154693 | 0.595 | 0.224 | 1.125E-133 |
| Fibroblast | Has2 | 3 | 4.48E-133 | 2.49870992 | 0.687 | 0.347 | 8.961E-130 |
| Fibroblast | Cadm3 | 3 | 1.327E-132 | 2.09362477 | 0.718 | 0.375 | 2.653E-129 |
| Fibroblast | Vegfa | 3 | 8.648E-127 | 1.17143279 | 0.915 | 0.637 | 1.73E-123 |
| Fibroblast | Has1 | 3 | 1.576E-126 | 2.94121389 | 0.653 | 0.367 | 3.151E-123 |
| Fibroblast | Ubash3b | 3 | 2.747E-115 | 2.29540937 | 0.688 | 0.409 | 5.494E-112 |
| Fibroblast | Dact2 | 3 | 3.634E-115 | 4.44367646 | 0.537 | 0.221 | 7.268E-112 |
| Fibroblast | Emilin2 | 3 | 1.371E-113 | 2.75254877 | 0.644 | 0.343 | 2.743E-110 |
| Fibroblast | Cers6 | 3 | 3.966E-110 | 1.56936917 | 0.77 | 0.496 | 7.932E-107 |
| Fibroblast | Diaph3 | 3 | 9.532E-109 | 3.10351572 | 0.593 | 0.261 | 1.906E-105 |
| Fibroblast | Kank11 | 3 | 8.61E-107 | 1.91972698 | 0.719 | 0.38 | 1.722E-103 |
| Fibroblast | Vcan | 3 | 5.501E-100 | 1.33353659 | 0.854 | 0.653 | 1.1002E-96 |
| Fibroblast | Adgrg2 | 3 | 1.7171E-97 | 2.5761826 | 0.617 | 0.308 | 3.4342E-94 |
| Fibroblast | Pcsk6 | 3 | 7.5127E-95 | 1.15501888 | 0.884 | 0.669 | 1.5025E-91 |
| Fibroblast | Lmx1a | 3 | 1.6286E-92 | 2.4500571 | 0.609 | 0.356 | 3.2573E-89 |
| Fibroblast | Ston2 | 3 | 1.9812E-91 | 1.03177476 | 0.898 | 0.722 | 3.9624E-88 |
| Fibroblast | Ntn4 | 3 | 5.1841E-91 | 1.58607887 | 0.725 | 0.408 | 1.0368E-87 |
| Fibroblast | Fbln2 | 3 | 2.2003E-90 | 1.38216327 | 0.756 | 0.478 | 4.4005E-87 |

| | | | | | | | |
|---|---|---|---|---|---|---|---|
| Fibroblast | Slc4a4 | 3 | 7.2672E-89 | 1.38931483 | 0.772 | 0.459 | 1.4534E-85 |
| Fibroblast | Npas21 | 3 | 9.1672E-89 | 1.22755983 | 0.829 | 0.581 | 1.8334E-85 |
| Fibroblast | Slco3a1 | 3 | 2.734E-85 | 1.15965331 | 0.793 | 0.521 | 5.4681E-82 |
| Fibroblast | Adamts5 | 3 | 4.4709E-85 | 1.47055586 | 0.773 | 0.509 | 8.9418E-82 |
| Fibroblast | Sulf1 | 3 | 2.9375E-84 | 1.21035748 | 0.829 | 0.6 | 5.875E-81 |
| Fibroblast | Srgap1 | 3 | 6.4606E-82 | 1.02780285 | 0.86 | 0.63 | 1.2921E-78 |
| Fibroblast | Timp2 | 3 | 1.4033E-81 | 0.86255197 | 0.928 | 0.776 | 2.8066E-78 |
| Fibroblast | Tmem100 | 3 | 7.5554E-80 | 2.62397463 | 0.579 | 0.34 | 1.5111E-76 |
| Fibroblast | Dclk1 | 3 | 7.8272E-79 | 0.88203825 | 0.901 | 0.703 | 1.5654E-75 |
| Fibroblast | Medag | 3 | 8.2892E-77 | 1.02581166 | 0.825 | 0.603 | 1.6578E-73 |
| Fibroblast | Rab7b | 3 | 1.3685E-74 | 1.87191088 | 0.632 | 0.372 | 2.737E-71 |
| Fibroblast | Mybpc2 | 3 | 2.8904E-74 | 1.15632553 | 0.324 | 0.093 | 5.7808E-71 |
| Fibroblast | Ackr3 | 3 | 2.7758E-73 | 1.27128116 | 0.677 | 0.442 | 5.5517E-70 |
| Fibroblast | Ptpn22 | 3 | 5.6365E-72 | 0.85631498 | 0.385 | 0.119 | 1.1273E-68 |
| Fibroblast | Tnfrsf11b1 | 3 | 7.0603E-72 | 2.17475844 | 0.63 | 0.381 | 1.4121E-68 |
| Fibroblast | Fam167a | 3 | 2.4404E-71 | 2.24551413 | 0.516 | 0.268 | 4.8808E-68 |
| Fibroblast | Hmga2 | 3 | 1.6861E-70 | 0.64434539 | 0.273 | 0.107 | 3.3723E-67 |
| Fibroblast | Dpp4 | 3 | 7.0785E-70 | 2.44969142 | 0.506 | 0.201 | 1.4157E-66 |
| Fibroblast | Nr4a3 | 3 | 2.2927E-69 | 1.29662282 | 0.511 | 0.271 | 4.5855E-66 |
| Fibroblast | Pde10a | 3 | 3.5451E-69 | 1.06595029 | 0.801 | 0.576 | 7.0902E-66 |
| Fibroblast | Ildr21 | 3 | 3.051E-66 | 2.43978785 | 0.539 | 0.304 | 6.1021E-63 |
| Fibroblast | Anpep | 3 | 2.5298E-62 | 1.82366565 | 0.599 | 0.41 | 5.0597E-59 |
| Fibroblast | Tmem178b | 3 | 2.5136E-59 | 0.88021623 | 0.32 | 0.035 | 5.0272E-56 |
| Fibroblast | Gfra2 | 3 | 3.6983E-58 | 1.82987111 | 0.269 | 0.082 | 7.3966E-55 |
| Fibroblast | Cdon | 3 | 1.3722E-57 | 1.06961095 | 0.726 | 0.506 | 2.7443E-54 |
| Fibroblast | Me1 | 3 | 1.2053E-56 | 1.50659153 | 0.601 | 0.367 | 2.4106E-53 |
| Fibroblast | Dapk11 | 3 | 9.0107E-56 | 0.66613437 | 0.84 | 0.639 | 1.8021E-52 |
| Fibroblast | Esrrg | 3 | 1.9595E-55 | 1.21979419 | 0.732 | 0.521 | 3.919E-52 |
| Fibroblast | Hs3st5 | 3 | 4.3892E-54 | 2.38059561 | 0.519 | 0.26 | 8.7784E-51 |
| Fibroblast | Tmeff21 | 3 | 1.2489E-53 | 0.76339836 | 0.873 | 0.669 | 2.4978E-50 |
| Fibroblast | Negr1 | 3 | 6.6003E-49 | 1.41833807 | 0.658 | 0.465 | 1.3201E-45 |

| | | | | | | | |
|---|---|---|---|---|---|---|---|
| Fibroblast | Akap121 | 3 | 8.3193E-49 | 1.10847576 | 0.638 | 0.397 | 1.6639E-45 |
| Fibroblast | Esco2 | 3 | 3.5024E-48 | 0.94465911 | 0.32 | 0.105 | 7.0048E-45 |
| Fibroblast | Atf3 | 3 | 6.9994E-48 | 1.76275012 | 0.328 | 0.13 | 1.3999E-44 |
| Fibroblast | Fam71f1 | 3 | 4.4566E-47 | 3.24560692 | 0.374 | 0.235 | 8.9132E-44 |
| Fibroblast | Dkk2 | 3 | 3.9916E-46 | 2.30224569 | 0.526 | 0.344 | 7.9832E-43 |
| Fibroblast | Stxbp6 | 3 | 6.6505E-46 | 1.16917328 | 0.6 | 0.349 | 1.3301E-42 |
| Fibroblast | Ptprj | 3 | 1.1321E-44 | 0.78270869 | 0.736 | 0.567 | 2.2642E-41 |
| Fibroblast | Kcnq5 | 3 | 1.773E-43 | 2.58164115 | 0.411 | 0.204 | 3.5461E-40 |
| Fibroblast | Sdc2 | 3 | 1.8996E-42 | 0.90192851 | 0.725 | 0.512 | 3.7993E-39 |
| Fibroblast | Spsb4 | 3 | 2.5682E-42 | 1.85251884 | 0.548 | 0.401 | 5.1365E-39 |
| Fibroblast | Olfml2a | 3 | 2.7208E-42 | 2.87116872 | 0.394 | 0.163 | 5.4416E-39 |
| Fibroblast | Gda | 3 | 2.9323E-42 | 0.67488595 | 0.888 | 0.745 | 5.8647E-39 |
| Fibroblast | Cd44 | 3 | 9.2775E-42 | 1.88251115 | 0.55 | 0.378 | 1.8555E-38 |
| Fibroblast | Tspan18 | 3 | 1.6453E-41 | 0.85720764 | 0.684 | 0.506 | 3.2906E-38 |
| Fibroblast | Myo5b | 3 | 2.3316E-41 | 0.51919505 | 0.279 | 0.119 | 4.6632E-38 |
| Fibroblast | Ugdh | 3 | 1.265E-40 | 2.16039561 | 0.516 | 0.377 | 2.53E-37 |
| Fibroblast | AABR070655 | 3 | 6.3907E-40 | 1.28576417 | 0.331 | 0.187 | 1.2781E-36 |
| Fibroblast | Aldh1a3 | 3 | 1.7215E-39 | 2.95432463 | 0.411 | 0.324 | 3.443E-36 |
| Fibroblast | Ckap2 | 3 | 6.0271E-39 | 1.16510586 | 0.31 | 0.14 | 1.2054E-35 |
| Fibroblast | Penk | 3 | 7.202E-38 | 2.02795492 | 0.362 | 0.207 | 1.4404E-34 |
| Fibroblast | Tnfaip6 | 3 | 1.3764E-37 | 1.76111702 | 0.537 | 0.374 | 2.7527E-34 |
| Fibroblast | Galnt161 | 3 | 3.2514E-37 | 0.64764087 | 0.786 | 0.605 | 6.5029E-34 |
| Fibroblast | Antxr1 | 3 | 7.0721E-37 | 0.66048049 | 0.908 | 0.778 | 1.4144E-33 |
| Fibroblast | Fn11 | 3 | 1.7454E-36 | 0.49248814 | 0.799 | 0.61 | 3.4909E-33 |
| Fibroblast | Gas7 | 3 | 3.7425E-36 | 1.07083845 | 0.604 | 0.447 | 7.4849E-33 |
| Fibroblast | Nckap5 | 3 | 3.7451E-36 | 1.54456849 | 0.498 | 0.306 | 7.4903E-33 |
| Fibroblast | Ccl2 | 3 | 1.4063E-35 | 1.83399447 | 0.32 | 0.128 | 2.8126E-32 |
| Fibroblast | Itga9 | 3 | 1.663E-35 | 0.66775047 | 0.799 | 0.678 | 3.326E-32 |
| Fibroblast | Adarb2 | 3 | 4.6833E-35 | 1.02868885 | 0.299 | 0.132 | 9.3665E-32 |
| Fibroblast | Fndc1 | 3 | 5.72E-35 | 0.51470251 | 0.936 | 0.843 | 1.144E-31 |
| Fibroblast | Nusap1 | 3 | 1.3175E-34 | 0.33723369 | 0.369 | 0.188 | 2.635E-31 |

| | | | | | | | |
|---|---|---|---|---|---|---|---|
| Fibroblast | Syt1 | 3 | 4.6141E-34 | 2.28982169 | 0.397 | 0.187 | 9.2282E-31 |
| Fibroblast | Timp31 | 3 | 9.5379E-34 | 0.72794969 | 0.68 | 0.473 | 1.9076E-30 |
| Fibroblast | Itga2 | 3 | 2.1335E-33 | 1.50604949 | 0.434 | 0.227 | 4.267E-30 |
| Fibroblast | Tyrobp1 | 3 | 3.0438E-33 | 0.69991616 | 0.365 | 0.196 | 6.0876E-30 |
| Fibroblast | Fgf1 | 3 | 8.4708E-33 | 1.17496767 | 0.509 | 0.256 | 1.6942E-29 |
| Fibroblast | Pdlim5 | 3 | 3.6302E-32 | 0.67684251 | 0.771 | 0.663 | 7.2603E-29 |
| Fibroblast | Chn2 | 3 | 5.504E-32 | 0.83828697 | 0.588 | 0.332 | 1.1008E-28 |
| Fibroblast | Slit31 | 3 | 1.155E-31 | 0.7730879 | 0.61 | 0.467 | 2.31E-28 |
| Fibroblast | Abca4 | 3 | 1.912E-31 | 1.80613104 | 0.314 | 0.154 | 3.824E-28 |
| Fibroblast | Ezr | 3 | 3.0393E-31 | 1.43121165 | 0.482 | 0.264 | 6.0785E-28 |
| Fibroblast | Sema3d | 3 | 4.9413E-31 | 1.22402112 | 0.552 | 0.379 | 9.8826E-28 |
| Fibroblast | Adam23 | 3 | 5.0232E-30 | 1.58760049 | 0.428 | 0.197 | 1.0046E-26 |
| Fibroblast | Nrg2 | 3 | 5.6898E-30 | 0.38373142 | 0.43 | 0.187 | 1.138E-26 |
| Fibroblast | Pde7a1 | 3 | 7.9879E-30 | 0.63035954 | 0.741 | 0.582 | 1.5976E-26 |
| Fibroblast | AABR070265 | 3 | 8.7464E-29 | 0.89242489 | 0.448 | 0.273 | 1.7493E-25 |
| Fibroblast | Hecw21 | 3 | 1.0279E-28 | 0.88732169 | 0.578 | 0.363 | 2.0559E-25 |
| Fibroblast | Lpcat2 | 3 | 4.3666E-28 | 0.90547854 | 0.433 | 0.213 | 8.7331E-25 |
| Fibroblast | Loxl11 | 3 | 7.3679E-28 | 0.49130776 | 0.818 | 0.675 | 1.4736E-24 |
| Fibroblast | Trim7 | 3 | 1.5335E-27 | 0.81510882 | 0.447 | 0.231 | 3.0669E-24 |
| Fibroblast | Tspan51 | 3 | 3.664E-26 | 0.59292497 | 0.74 | 0.552 | 7.328E-23 |
| Fibroblast | Ablim21 | 3 | 3.8268E-26 | 0.61820961 | 0.417 | 0.208 | 7.6536E-23 |
| Fibroblast | Col5a3 | 3 | 5.3085E-26 | 0.54732754 | 0.685 | 0.482 | 1.0617E-22 |
| Fibroblast | Rcan22 | 3 | 1.6832E-25 | 0.56201219 | 0.74 | 0.628 | 3.3664E-22 |
| Fibroblast | PCOLCE2 | 3 | 4.0324E-25 | 1.22949523 | 0.475 | 0.344 | 8.0648E-22 |
| Fibroblast | Pi15 | 3 | 4.844E-25 | 1.22492692 | 0.466 | 0.322 | 9.6881E-22 |
| Fibroblast | Kcnk5 | 3 | 5.9393E-25 | 2.16088072 | 0.373 | 0.252 | 1.1879E-21 |
| Fibroblast | Adamts151 | 3 | 8.6411E-25 | 0.81457506 | 0.581 | 0.4 | 1.7282E-21 |
| Fibroblast | C1qtnf7 | 3 | 2.2397E-24 | 0.3863698 | 0.955 | 0.887 | 4.4794E-21 |
| Fibroblast | Mest | 3 | 5.3809E-24 | 1.1891703 | 0.294 | 0.171 | 1.0762E-20 |
| Fibroblast | Stmn4 | 3 | 6.1978E-24 | 1.99119762 | 0.388 | 0.277 | 1.2396E-20 |
| Fibroblast | AABR070215 | 3 | 1.2209E-23 | 0.64491967 | 0.435 | 0.344 | 2.4418E-20 |

| | | | | | | | |
|---|---|---|---|---|---|---|---|
| Fibroblast | Il33 | 3 | 1.4254E-23 | 2.71790988 | 0.429 | 0.324 | 2.8508E-20 |
| Fibroblast | Bmp61 | 3 | 2.0938E-23 | 0.4716154 | 0.65 | 0.469 | 4.1876E-20 |
| Fibroblast | Bcat1 | 3 | 5.6024E-23 | 0.62281013 | 0.663 | 0.512 | 1.1205E-19 |
| Fibroblast | AABR070440 | 3 | 9.1467E-23 | 0.65588225 | 0.567 | 0.37 | 1.8293E-19 |
| Fibroblast | Cdh7 | 3 | 1.0214E-22 | 1.09191998 | 0.398 | 0.241 | 2.0428E-19 |
| Fibroblast | Tbx181 | 3 | 1.0563E-22 | 0.70940887 | 0.602 | 0.456 | 2.1125E-19 |
| Fibroblast | Fras1 | 3 | 1.6884E-22 | 0.89312169 | 0.286 | 0.122 | 3.3769E-19 |
| Fibroblast | Anxa3 | 3 | 3.5604E-22 | 1.67859245 | 0.382 | 0.23 | 7.1207E-19 |
| Fibroblast | Aldh1a21 | 3 | 5.3681E-22 | 0.43602105 | 0.512 | 0.341 | 1.0736E-18 |
| Fibroblast | Nova1 | 3 | 8.7009E-22 | 0.92231295 | 0.499 | 0.327 | 1.7402E-18 |
| Fibroblast | Tpx2 | 3 | 1.0655E-21 | 0.37447071 | 0.41 | 0.185 | 2.1311E-18 |
| Fibroblast | Shb | 3 | 2.2559E-21 | 0.87325458 | 0.517 | 0.374 | 4.5118E-18 |
| Fibroblast | Afap1l21 | 3 | 3.8356E-21 | 0.96987122 | 0.486 | 0.373 | 7.6713E-18 |
| Fibroblast | Zbtb7c | 3 | 8.8755E-20 | 0.99826139 | 0.464 | 0.321 | 1.7751E-16 |
| Fibroblast | Myom3 | 3 | 9.9807E-20 | 1.47524753 | 0.267 | 0.143 | 1.9961E-16 |
| Fibroblast | Cxcl12 | 3 | 1.2807E-19 | 0.40233991 | 0.517 | 0.3 | 2.5614E-16 |
| Fibroblast | Ebf21 | 3 | 1.3273E-19 | 0.33017961 | 0.939 | 0.855 | 2.6546E-16 |
| Fibroblast | Pla2g2a | 3 | 2.5172E-19 | 1.27840546 | 0.328 | 0.164 | 5.0343E-16 |
| Fibroblast | Adamts14 | 3 | 3.0811E-19 | 0.76909874 | 0.552 | 0.432 | 6.1622E-16 |
| Fibroblast | Mark1 | 3 | 3.3078E-19 | 0.46644369 | 0.437 | 0.22 | 6.6157E-16 |
| Fibroblast | Sytl21 | 3 | 5.9051E-19 | 0.44584608 | 0.338 | 0.171 | 1.181E-15 |
| Fibroblast | Tnfaip8l3 | 3 | 1.3242E-18 | 0.60994539 | 0.454 | 0.291 | 2.6485E-15 |
| Fibroblast | Rai141 | 3 | 2.1367E-18 | 0.38533658 | 0.641 | 0.469 | 4.2734E-15 |
| Fibroblast | Flrt21 | 3 | 2.6009E-18 | 0.39839173 | 0.668 | 0.496 | 5.2017E-15 |
| Fibroblast | Crip1 | 3 | 3.9674E-18 | 0.65478906 | 0.474 | 0.335 | 7.9347E-15 |
| Fibroblast | Adcy21 | 3 | 5.3411E-18 | 0.92067806 | 0.446 | 0.323 | 1.0682E-14 |
| Fibroblast | Vcl1 | 3 | 6.8702E-18 | 0.60183975 | 0.657 | 0.542 | 1.374E-14 |
| Fibroblast | Col14a1 | 3 | 8.0888E-18 | 0.41036202 | 0.825 | 0.711 | 1.6178E-14 |
| Fibroblast | Ptgs2 | 3 | 1.2913E-17 | 0.64129035 | 0.429 | 0.315 | 2.5825E-14 |
| Fibroblast | LOC1025530 | 3 | 1.4036E-17 | 0.57009474 | 0.517 | 0.358 | 2.8072E-14 |
| Fibroblast | Pfkfb1 | 3 | 2.0907E-17 | 0.51904966 | 0.326 | 0.212 | 4.1813E-14 |

| | | | | | | | |
|---|---|---|---|---|---|---|---|
| Fibroblast | Ptprn2 | 3 | 2.1556E-17 | 0.93800707 | 0.265 | 0.126 | 4.3112E-14 |
| Fibroblast | Mgmt1 | 3 | 2.245E-17 | 0.45431149 | 0.636 | 0.459 | 4.49E-14 |
| Fibroblast | Filip1l | 3 | 3.5141E-17 | 0.43204681 | 0.742 | 0.619 | 7.0282E-14 |
| Fibroblast | Lims2 | 3 | 3.5694E-17 | 1.50763983 | 0.376 | 0.268 | 7.1388E-14 |
| Fibroblast | Fa2h | 3 | 4.4064E-17 | 1.41228238 | 0.252 | 0.119 | 8.8129E-14 |
| Fibroblast | LOC1003614 | 3 | 7.1907E-17 | 0.35419448 | 0.526 | 0.316 | 1.4381E-13 |
| Fibroblast | Cacna2d32 | 3 | 1.2145E-16 | 0.35644878 | 0.719 | 0.637 | 2.429E-13 |
| Fibroblast | Kif23 | 3 | 1.2353E-16 | 0.3108667 | 0.326 | 0.151 | 2.4705E-13 |
| Fibroblast | Pla2g51 | 3 | 1.2938E-16 | 0.51448076 | 0.417 | 0.232 | 2.5876E-13 |
| Fibroblast | Prph | 3 | 2.8271E-16 | 0.7369364 | 0.355 | 0.2 | 5.6542E-13 |
| Fibroblast | Chst15 | 3 | 2.8845E-16 | 0.65164343 | 0.35 | 0.217 | 5.769E-13 |
| Fibroblast | Efnb21 | 3 | 4.8799E-16 | 1.04228708 | 0.385 | 0.229 | 9.7599E-13 |
| Fibroblast | Ptpn3 | 3 | 5.2594E-16 | 0.42944344 | 0.541 | 0.379 | 1.0519E-12 |
| Fibroblast | Flna | 3 | 6.7712E-16 | 0.45854706 | 0.597 | 0.412 | 1.3542E-12 |
| Fibroblast | Adamts19 | 3 | 7.8204E-16 | 0.33778049 | 0.74 | 0.638 | 1.5641E-12 |
| Fibroblast | Lmln1 | 3 | 2.2659E-15 | 0.31561864 | 0.453 | 0.27 | 4.5318E-12 |
| Fibroblast | Cdkn1a | 3 | 2.5603E-15 | 0.55083693 | 0.47 | 0.33 | 5.1206E-12 |
| Fibroblast | Cacna1e1 | 3 | 2.8268E-15 | 0.45817502 | 0.568 | 0.431 | 5.6536E-12 |
| Fibroblast | Zfp536 | 3 | 7.6013E-15 | 0.57742721 | 0.251 | 0.17 | 1.5203E-11 |
| Fibroblast | Nrxn3 | 3 | 1.0582E-14 | 0.62283279 | 0.434 | 0.29 | 2.1164E-11 |
| Fibroblast | Efemp11 | 3 | 1.2291E-14 | 0.93383475 | 0.431 | 0.31 | 2.4582E-11 |
| Fibroblast | Cyp26b1 | 3 | 1.7155E-14 | 0.63268235 | 0.33 | 0.21 | 3.431E-11 |
| Fibroblast | AABR070070 | 3 | 1.7388E-14 | 0.38214242 | 0.705 | 0.596 | 3.4775E-11 |
| Fibroblast | Loxl2 | 3 | 1.9971E-14 | 0.45290347 | 0.619 | 0.511 | 3.9942E-11 |
| Fibroblast | Tlr7 | 3 | 2.8321E-14 | 0.5374871 | 0.315 | 0.153 | 5.6641E-11 |
| Fibroblast | S100a6 | 3 | 8.585E-14 | 0.68609406 | 0.391 | 0.259 | 1.717E-10 |
| Fibroblast | Tmem1961 | 3 | 1.2821E-13 | 0.50600907 | 0.528 | 0.396 | 2.5641E-10 |
| Fibroblast | Slc7a1 | 3 | 1.3977E-13 | 0.4744484 | 0.508 | 0.362 | 2.7954E-10 |
| Fibroblast | Cpq | 3 | 2.5385E-13 | 0.26999132 | 0.839 | 0.747 | 5.0771E-10 |
| Fibroblast | Hmmr | 3 | 4.3965E-13 | 0.37018638 | 0.254 | 0.08 | 8.793E-10 |
| Fibroblast | Pcdh19 | 3 | 1.0259E-12 | 0.58613194 | 0.466 | 0.365 | 2.0518E-09 |

| | | | | | | | |
|---|---|---|---|---|---|---|---|
| Fibroblast | Ecm1 | 3 | 1.7133E-12 | 1.02897771 | 0.409 | 0.387 | 3.4265E-09 |
| Fibroblast | Fgd2 | 3 | 1.7736E-12 | 0.42155075 | 0.383 | 0.27 | 3.5472E-09 |
| Fibroblast | Igfbp51 | 3 | 2.1403E-12 | 0.51344829 | 0.495 | 0.351 | 4.2806E-09 |
| Fibroblast | Rnls1 | 3 | 2.5654E-12 | 0.58034443 | 0.488 | 0.41 | 5.1309E-09 |
| Fibroblast | Plekha72 | 3 | 2.8287E-12 | 0.31985716 | 0.727 | 0.66 | 5.6575E-09 |
| Fibroblast | Homer1 | 3 | 3.7924E-12 | 0.29737163 | 0.516 | 0.379 | 7.5847E-09 |
| Fibroblast | Kcnab1 | 3 | 3.8693E-12 | 0.73018022 | 0.453 | 0.347 | 7.7386E-09 |
| Fibroblast | Csrp1 | 3 | 5.5533E-12 | 0.2845665 | 0.517 | 0.353 | 1.1107E-08 |
| Fibroblast | Hs3st3a1 | 3 | 6.3345E-12 | 1.34599423 | 0.385 | 0.299 | 1.2669E-08 |
| Fibroblast | Rec114 | 3 | 7.7521E-12 | 0.400196 | 0.282 | 0.163 | 1.5504E-08 |
| Fibroblast | LOC1009119 | 3 | 9.4031E-12 | 1.19162885 | 0.378 | 0.289 | 1.8806E-08 |
| Fibroblast | Frmd5 | 3 | 1.336E-11 | 1.21187641 | 0.31 | 0.228 | 2.6721E-08 |
| Fibroblast | Samd12 | 3 | 1.7991E-11 | 0.78998277 | 0.412 | 0.302 | 3.5982E-08 |
| Fibroblast | Il6r1 | 3 | 2.834E-11 | 0.41757786 | 0.524 | 0.399 | 5.6681E-08 |
| Fibroblast | Hspb8 | 3 | 4.7181E-11 | 1.16219876 | 0.464 | 0.412 | 9.4362E-08 |
| Fibroblast | Slc43a2 | 3 | 8.8157E-11 | 0.39584598 | 0.389 | 0.283 | 1.7631E-07 |
| Fibroblast | Ntrk21 | 3 | 1.7327E-10 | 1.0642606 | 0.388 | 0.327 | 3.4654E-07 |
| Fibroblast | Fhad1 | 3 | 1.7907E-10 | 1.22942746 | 0.302 | 0.122 | 3.5814E-07 |
| Fibroblast | Enpp21 | 3 | 2.1611E-10 | 0.72037483 | 0.437 | 0.37 | 4.3222E-07 |
| Fibroblast | Rtn4rl1 | 3 | 2.2568E-10 | 0.93028331 | 0.295 | 0.163 | 4.5137E-07 |
| Fibroblast | Herc61 | 3 | 2.2698E-10 | 0.3587456 | 0.427 | 0.286 | 4.5396E-07 |
| Fibroblast | Pla2r1 | 3 | 2.7013E-10 | 0.56963513 | 0.38 | 0.252 | 5.4026E-07 |
| Fibroblast | Tmem51 | 3 | 3.6856E-10 | 1.27057622 | 0.33 | 0.247 | 7.3711E-07 |
| Fibroblast | Tbx5 | 3 | 4.3975E-10 | 0.81219656 | 0.373 | 0.279 | 8.795E-07 |
| Fibroblast | LOC1009095 | 3 | 9.1651E-10 | 0.91727702 | 0.354 | 0.252 | 1.833E-06 |
| Fibroblast | Pamr1 | 3 | 9.9928E-10 | 0.58630361 | 0.427 | 0.355 | 1.9986E-06 |
| Fibroblast | Nhs1 | 3 | 1.9257E-09 | 0.25540725 | 0.468 | 0.362 | 3.8513E-06 |
| Fibroblast | Sfrp4 | 3 | 4.4542E-09 | 0.80630023 | 0.315 | 0.332 | 8.9084E-06 |
| Fibroblast | Ptafr1 | 3 | 8.575E-09 | 0.38293307 | 0.401 | 0.289 | 1.715E-05 |
| Fibroblast | Sh3gl2 | 3 | 9.6865E-09 | 0.71100218 | 0.388 | 0.302 | 1.9373E-05 |
| Fibroblast | Enah1 | 3 | 1.0486E-08 | 0.33777009 | 0.534 | 0.46 | 2.0971E-05 |

| | | | | | | | |
|---|---|---|---|---|---|---|---|
| Fibroblast | Kcnk13 | 3 | 1.6673E-08 | 0.7142412 | 0.34 | 0.252 | 3.3345E-05 |
| Fibroblast | Fabp5 | 3 | 2.0178E-08 | 0.61012852 | 0.261 | 0.21 | 4.0357E-05 |
| Fibroblast | Ptk2b1 | 3 | 2.7272E-08 | 0.27410244 | 0.542 | 0.442 | 5.4543E-05 |
| Fibroblast | Galnt172 | 3 | 2.755E-08 | 0.26796169 | 0.76 | 0.658 | 5.51E-05 |
| Fibroblast | AABR070010 | 3 | 2.8414E-08 | 0.57119705 | 0.476 | 0.362 | 5.6828E-05 |
| Fibroblast | Arhgap241 | 3 | 3.3378E-08 | 0.38695402 | 0.709 | 0.639 | 6.6755E-05 |
| Fibroblast | Gnao11 | 3 | 8.0821E-08 | 0.29214046 | 0.433 | 0.341 | 0.00016164 |
| Fibroblast | AABR070067 | 3 | 9.2097E-08 | 0.31682114 | 0.3 | 0.192 | 0.00018419 |
| Fibroblast | Pcp4l1 | 3 | 9.7117E-08 | 0.35754847 | 0.279 | 0.198 | 0.00019423 |
| Fibroblast | Sv2c1 | 3 | 2.5705E-07 | 0.43572734 | 0.537 | 0.504 | 0.0005141 |
| Fibroblast | Polr2m | 3 | 2.6886E-07 | 0.43835044 | 0.481 | 0.426 | 0.00053772 |
| Fibroblast | Rimbp2 | 3 | 5.9485E-07 | 1.21181222 | 0.345 | 0.264 | 0.0011897 |
| Fibroblast | Ltbp12 | 3 | 7.2265E-07 | 0.31512879 | 0.8 | 0.656 | 0.0014453 |
| Fibroblast | Eva1c1 | 3 | 1.1107E-06 | 0.41806655 | 0.367 | 0.303 | 0.0022214 |
| Fibroblast | Nr4a1 | 3 | 1.2773E-06 | 0.95486277 | 0.328 | 0.279 | 0.00255459 |
| Fibroblast | Cotl1 | 3 | 1.3483E-06 | 0.94415006 | 0.397 | 0.324 | 0.00269656 |
| Fibroblast | Tmem63c | 3 | 1.9061E-06 | 1.0819307 | 0.301 | 0.231 | 0.00381212 |
| Fibroblast | Trib3 | 3 | 2.7962E-06 | 0.64522307 | 0.261 | 0.218 | 0.00559239 |
| Fibroblast | Sh3bp2 | 3 | 2.9545E-06 | 1.1534285 | 0.347 | 0.297 | 0.00590902 |
| Fibroblast | Igfbp6 | 3 | 8.322E-06 | 0.84627144 | 0.347 | 0.293 | 0.0166439 |
| Fibroblast | Slc16a11 | 3 | 1.113E-05 | 0.60730319 | 0.311 | 0.25 | 0.02226032 |
| Fibroblast | Plcb1 | 3 | 1.3244E-05 | 0.62106403 | 0.359 | 0.323 | 0.02648703 |
| Fibroblast | Kif17 | 4 | 0 | 2.22614644 | 0.728 | 0.12 | 0 |
| Fibroblast | Zfp5361 | 4 | 0 | 2.08720718 | 0.672 | 0.122 | 0 |
| Fibroblast | Tll21 | 4 | 0 | 2.81117616 | 0.74 | 0.229 | 0 |
| Fibroblast | Knstrn | 4 | 0 | 2.13975979 | 0.565 | 0.055 | 0 |
| Fibroblast | Cxcl10 | 4 | 0 | 3.02952664 | 0.62 | 0.125 | 0 |
| Fibroblast | Adgre1 | 4 | 0 | 1.40772337 | 0.58 | 0.09 | 0 |
| Fibroblast | Capn13 | 4 | 0 | 0.85622123 | 0.569 | 0.083 | 0 |
| Fibroblast | RGD1305807 | 4 | 0 | 0.81909724 | 0.54 | 0.064 | 0 |
| Fibroblast | Acan | 4 | 0 | 0.75649717 | 0.518 | 0.055 | 0 |

| | | | | | | | |
|---|---|---|---|---|---|---|---|
| Fibroblast | Stum | 4 | 0 | 2.12388207 | 0.488 | 0.025 | 0 |
| Fibroblast | Gja5 | 4 | 0 | 1.49940556 | 0.48 | 0.071 | 0 |
| Fibroblast | Col25a1 | 4 | 0 | 1.58786569 | 0.446 | 0.041 | 0 |
| Fibroblast | Zdbf2 | 4 | 0 | 0.808157 | 0.299 | 0.036 | 0 |
| Fibroblast | Abca8a1 | 4 | 1.983E-297 | 2.22749882 | 0.978 | 0.525 | 3.965E-294 |
| Fibroblast | Cdh6 | 4 | 4.257E-284 | 2.02317352 | 0.451 | 0.061 | 8.515E-281 |
| Fibroblast | Cdh4 | 4 | 1.285E-282 | 0.6506466 | 0.607 | 0.138 | 2.57E-279 |
| Fibroblast | Fosb | 4 | 8.805E-276 | 0.26918197 | 0.421 | 0.054 | 1.761E-272 |
| Fibroblast | Slc5a10 | 4 | 1.28E-271 | 2.70659209 | 0.471 | 0.038 | 2.559E-268 |
| Fibroblast | Tmem255a | 4 | 7.601E-263 | 2.10516995 | 0.423 | 0.034 | 1.52E-259 |
| Fibroblast | Cxcl14 | 4 | 3.694E-255 | 1.33864709 | 0.338 | 0.047 | 7.387E-252 |
| Fibroblast | Fbxl2 | 4 | 8.449E-254 | 1.13377274 | 0.495 | 0.109 | 1.69E-250 |
| Fibroblast | Ttll9 | 4 | 1.109E-243 | 1.1321362 | 0.57 | 0.125 | 2.217E-240 |
| Fibroblast | Ptprt | 4 | 1.541E-238 | 2.96931533 | 0.649 | 0.166 | 3.081E-235 |
| Fibroblast | RGD1564053 | 4 | 4.646E-236 | 1.31315721 | 0.429 | 0.056 | 9.292E-233 |
| Fibroblast | Smoc21 | 4 | 3.26E-232 | 1.84465215 | 0.799 | 0.335 | 6.521E-229 |
| Fibroblast | Kcnc21 | 4 | 3.894E-232 | 2.30832965 | 0.846 | 0.337 | 7.787E-229 |
| Fibroblast | Bicdl1 | 4 | 5.928E-226 | 0.75831558 | 0.518 | 0.079 | 1.186E-222 |
| Fibroblast | Cux21 | 4 | 3.712E-225 | 1.76371642 | 0.759 | 0.266 | 7.425E-222 |
| Fibroblast | Sctr | 4 | 5.783E-224 | 2.08355349 | 0.442 | 0.125 | 1.157E-220 |
| Fibroblast | Adap2 | 4 | 8.382E-224 | 0.57618935 | 0.679 | 0.194 | 1.676E-220 |
| Fibroblast | Pde11a | 4 | 3.23E-222 | 0.78710597 | 0.656 | 0.176 | 6.459E-219 |
| Fibroblast | Plxnc11 | 4 | 1.361E-221 | 1.76246473 | 0.709 | 0.225 | 2.723E-218 |
| Fibroblast | Slc22a231 | 4 | 1.464E-221 | 1.78119169 | 0.729 | 0.208 | 2.927E-218 |
| Fibroblast | Asb141 | 4 | 7.933E-219 | 1.50173944 | 0.729 | 0.222 | 1.587E-215 |
| Fibroblast | Lepr1 | 4 | 6.247E-217 | 1.76581587 | 0.732 | 0.278 | 1.249E-213 |
| Fibroblast | AABR070317 | 4 | 4.667E-215 | 0.76628547 | 0.707 | 0.178 | 9.334E-212 |
| Fibroblast | LOC1036933 | 4 | 8.836E-211 | 1.26843416 | 0.665 | 0.23 | 1.767E-207 |
| Fibroblast | Tmem132d1 | 4 | 1.514E-205 | 2.22886015 | 0.755 | 0.292 | 3.029E-202 |
| Fibroblast | Adamts31 | 4 | 3.223E-201 | 1.64096995 | 0.756 | 0.25 | 6.446E-198 |
| Fibroblast | Itgam | 4 | 3.492E-199 | 1.46472667 | 0.536 | 0.153 | 6.985E-196 |

| | | | | | | | |
|---|---|---|---|---|---|---|---|
| Fibroblast | Crispld21 | 4 | 1.422E-198 | 1.95304345 | 0.865 | 0.407 | 2.843E-195 |
| Fibroblast | Smpdl3a | 4 | 9.59E-198 | 0.48140179 | 0.752 | 0.26 | 1.918E-194 |
| Fibroblast | Cadm2 | 4 | 7.769E-193 | 1.31329464 | 0.504 | 0.143 | 1.554E-189 |
| Fibroblast | Pla2g52 | 4 | 4.528E-191 | 1.72124256 | 0.606 | 0.211 | 9.057E-188 |
| Fibroblast | Asic21 | 4 | 6.14E-191 | 2.1515996 | 0.649 | 0.225 | 1.228E-187 |
| Fibroblast | Ppp1r3a | 4 | 3.194E-190 | 0.89961724 | 0.617 | 0.174 | 6.389E-187 |
| Fibroblast | Cd300a | 4 | 8.612E-189 | 1.22814118 | 0.576 | 0.186 | 1.722E-185 |
| Fibroblast | Rgs17 | 4 | 1.301E-187 | 1.75580377 | 0.739 | 0.307 | 2.601E-184 |
| Fibroblast | Hspb3 | 4 | 1.496E-185 | 0.88521407 | 0.497 | 0.104 | 2.992E-182 |
| Fibroblast | Col4a31 | 4 | 4.213E-185 | 1.69033501 | 0.834 | 0.422 | 8.426E-182 |
| Fibroblast | Slco2b11 | 4 | 1.422E-184 | 1.91225185 | 0.773 | 0.366 | 2.843E-181 |
| Fibroblast | Cntn21 | 4 | 1.646E-184 | 0.77310759 | 0.562 | 0.168 | 3.292E-181 |
| Fibroblast | Slc37a21 | 4 | 2.935E-184 | 1.584458 | 0.701 | 0.27 | 5.871E-181 |
| Fibroblast | Polq | 4 | 9.084E-180 | 1.53123969 | 0.493 | 0.127 | 1.817E-176 |
| Fibroblast | Dsg22 | 4 | 1.309E-179 | 0.50269276 | 0.646 | 0.203 | 2.617E-176 |
| Fibroblast | Aldh1a11 | 4 | 1.064E-175 | 1.72067945 | 0.736 | 0.335 | 2.127E-172 |
| Fibroblast | AC111804.2 | 4 | 2.847E-175 | 0.9676715 | 0.343 | 0.09 | 5.695E-172 |
| Fibroblast | Rcan11 | 4 | 1.885E-174 | 0.46002723 | 0.708 | 0.244 | 3.77E-171 |
| Fibroblast | Stk32b | 4 | 3.855E-172 | 1.55092901 | 0.562 | 0.13 | 7.711E-169 |
| Fibroblast | Eya2 | 4 | 4.201E-171 | 1.95467053 | 0.634 | 0.215 | 8.402E-168 |
| Fibroblast | AABR070172 | 4 | 3.528E-170 | 1.13498264 | 0.507 | 0.135 | 7.057E-167 |
| Fibroblast | Prkar2b1 | 4 | 3.863E-170 | 1.29771888 | 0.734 | 0.299 | 7.726E-167 |
| Fibroblast | Fstl5 | 4 | 6.082E-168 | 1.22996483 | 0.57 | 0.175 | 1.216E-164 |
| Fibroblast | Atp2b21 | 4 | 1.264E-167 | 0.70189865 | 0.562 | 0.223 | 2.528E-164 |
| Fibroblast | Cpa6 | 4 | 5.554E-165 | 1.24279852 | 0.463 | 0.134 | 1.111E-161 |
| Fibroblast | Zbtb161 | 4 | 2.682E-164 | 1.55715173 | 0.904 | 0.391 | 5.365E-161 |
| Fibroblast | Mtus21 | 4 | 3.425E-163 | 1.05057297 | 0.719 | 0.286 | 6.849E-160 |
| Fibroblast | Pdcd1lg2 | 4 | 1.445E-162 | 0.74324546 | 0.653 | 0.218 | 2.889E-159 |
| Fibroblast | AABR070015 | 4 | 1.406E-160 | 2.02990201 | 0.366 | 0.043 | 2.812E-157 |
| Fibroblast | Gria41 | 4 | 4.846E-159 | 1.76419523 | 0.701 | 0.315 | 9.691E-156 |
| Fibroblast | Fam189a11 | 4 | 2.995E-157 | 0.36404379 | 0.611 | 0.209 | 5.989E-154 |

| | | | | | | | |
|---|---|---|---|---|---|---|---|
| Fibroblast | Ptgfr | 4 | 4.846E-157 | 0.58186549 | 0.663 | 0.249 | 9.693E-154 |
| Fibroblast | Sgip11 | 4 | 4.852E-157 | 1.64931675 | 0.747 | 0.359 | 9.704E-154 |
| Fibroblast | Glb1l21 | 4 | 7.124E-156 | 1.33484352 | 0.584 | 0.221 | 1.425E-152 |
| Fibroblast | Maob1 | 4 | 2.425E-155 | 1.30079007 | 0.634 | 0.258 | 4.851E-152 |
| Fibroblast | Trim50 | 4 | 1.241E-153 | 0.40272828 | 0.478 | 0.16 | 2.482E-150 |
| Fibroblast | Fgf101 | 4 | 6.639E-151 | 1.55936511 | 0.677 | 0.294 | 1.328E-147 |
| Fibroblast | Cgnl1 | 4 | 3.465E-150 | 0.83917023 | 0.735 | 0.326 | 6.93E-147 |
| Fibroblast | Ccnb1 | 4 | 1.158E-149 | 1.11994454 | 0.286 | 0.051 | 2.316E-146 |
| Fibroblast | Ppl1 | 4 | 3.504E-149 | 1.32870366 | 0.622 | 0.223 | 7.008E-146 |
| Fibroblast | Enpp22 | 4 | 3.253E-146 | 1.43476779 | 0.734 | 0.336 | 6.505E-143 |
| Fibroblast | Shroom2 | 4 | 1.083E-145 | 1.20588657 | 0.33 | 0.103 | 2.165E-142 |
| Fibroblast | Flvcr2 | 4 | 1.679E-145 | 0.61873245 | 0.52 | 0.229 | 3.359E-142 |
| Fibroblast | AABR070687 | 4 | 3.247E-145 | 0.76176314 | 0.538 | 0.157 | 6.494E-142 |
| Fibroblast | Scn1a | 4 | 4.734E-145 | 0.50537553 | 0.584 | 0.188 | 9.468E-142 |
| Fibroblast | Marchf1 | 4 | 8.596E-144 | 0.84849176 | 0.537 | 0.122 | 1.719E-140 |
| Fibroblast | Sgo2 | 4 | 1.359E-143 | 0.86222182 | 0.522 | 0.128 | 2.718E-140 |
| Fibroblast | Gen1 | 4 | 1.977E-143 | 0.78660437 | 0.635 | 0.263 | 3.953E-140 |
| Fibroblast | Efcc11 | 4 | 2.643E-143 | 1.57695635 | 0.749 | 0.415 | 5.287E-140 |
| Fibroblast | P2ry14 | 4 | 4.029E-142 | 0.54840958 | 0.453 | 0.179 | 8.059E-139 |
| Fibroblast | Bdnf1 | 4 | 1.362E-141 | 0.80321434 | 0.626 | 0.249 | 2.724E-138 |
| Fibroblast | Ppargc1a1 | 4 | 1.488E-141 | 0.81258427 | 0.597 | 0.211 | 2.976E-138 |
| Fibroblast | Gna151 | 4 | 9.668E-141 | 1.19206651 | 0.551 | 0.246 | 1.934E-137 |
| Fibroblast | Astn21 | 4 | 1.559E-140 | 2.20753929 | 0.574 | 0.294 | 3.118E-137 |
| Fibroblast | Pdpn1 | 4 | 1.57E-140 | 0.96117464 | 0.66 | 0.245 | 3.14E-137 |
| Fibroblast | Nfatc21 | 4 | 3.772E-140 | 0.31883326 | 0.561 | 0.203 | 7.543E-137 |
| Fibroblast | AABR070010 | 4 | 7.883E-140 | 1.2164007 | 0.715 | 0.335 | 1.577E-136 |
| Fibroblast | Nrcam | 4 | 2.764E-139 | 0.38558347 | 0.596 | 0.237 | 5.529E-136 |
| Fibroblast | Fgf14 | 4 | 3.188E-139 | 0.86917628 | 0.32 | 0.069 | 6.375E-136 |
| Fibroblast | Aspa1 | 4 | 8.192E-139 | 0.38192217 | 0.624 | 0.257 | 1.638E-135 |
| Fibroblast | AABR070414 | 4 | 2.421E-138 | 2.46609299 | 0.303 | 0.068 | 4.842E-135 |
| Fibroblast | Blnk1 | 4 | 2.655E-138 | 2.68235613 | 0.549 | 0.273 | 5.309E-135 |

| | | | | | | | |
|---|---|---|---|---|---|---|---|
| Fibroblast | Heyl | 4 | 3.119E-138 | 1.65420503 | 0.851 | 0.509 | 6.239E-135 |
| Fibroblast | Syt11 | 4 | 7.576E-136 | 0.50399599 | 0.516 | 0.173 | 1.515E-132 |
| Fibroblast | Slc12a2 | 4 | 8.563E-136 | 0.58143919 | 0.612 | 0.233 | 1.713E-132 |
| Fibroblast | Coro61 | 4 | 1.353E-135 | 0.7002894 | 0.634 | 0.228 | 2.707E-132 |
| Fibroblast | Opcml1 | 4 | 1.908E-135 | 1.55987595 | 0.679 | 0.314 | 3.815E-132 |
| Fibroblast | Frem11 | 4 | 1.925E-135 | 1.47900662 | 0.719 | 0.361 | 3.851E-132 |
| Fibroblast | Slc25a21 | 4 | 2.457E-135 | 0.96869636 | 0.354 | 0.133 | 4.913E-132 |
| Fibroblast | Pjvk | 4 | 1.114E-134 | 0.90191142 | 0.253 | 0.055 | 2.228E-131 |
| Fibroblast | Ppp1r9a1 | 4 | 1.161E-133 | 0.44188077 | 0.541 | 0.195 | 2.323E-130 |
| Fibroblast | Itgb4 | 4 | 1.424E-133 | 0.62411207 | 0.482 | 0.152 | 2.848E-130 |
| Fibroblast | RGD1563354 | 4 | 2.801E-133 | 1.45378243 | 0.842 | 0.503 | 5.602E-130 |
| Fibroblast | Frmd4b | 4 | 4.505E-132 | 1.29543534 | 0.788 | 0.401 | 9.009E-129 |
| Fibroblast | Sh3gl3 | 4 | 5.175E-132 | 1.19034692 | 0.579 | 0.243 | 1.035E-128 |
| Fibroblast | Nr4a11 | 4 | 3.221E-131 | 0.59076416 | 0.597 | 0.249 | 6.443E-128 |
| Fibroblast | Epsti1 | 4 | 3.68E-131 | 1.08980195 | 0.384 | 0.121 | 7.359E-128 |
| Fibroblast | Slco5a12 | 4 | 4.832E-131 | 0.78816336 | 0.693 | 0.245 | 9.664E-128 |
| Fibroblast | Ppm1e1 | 4 | 8.626E-131 | 1.13117798 | 0.571 | 0.241 | 1.725E-127 |
| Fibroblast | Serpine21 | 4 | 1.449E-130 | 1.05785267 | 0.709 | 0.33 | 2.898E-127 |
| Fibroblast | Scube1 | 4 | 1.196E-128 | 0.25954896 | 0.661 | 0.329 | 2.392E-125 |
| Fibroblast | Grik2 | 4 | 2.022E-128 | 0.72270289 | 0.441 | 0.155 | 4.044E-125 |
| Fibroblast | Igsf11 | 4 | 7.972E-126 | 1.10795466 | 0.399 | 0.107 | 1.594E-122 |
| Fibroblast | Tspan11 | 4 | 1.492E-125 | 0.71835355 | 0.648 | 0.322 | 2.984E-122 |
| Fibroblast | Mctp1 | 4 | 3.154E-125 | 0.42614545 | 0.663 | 0.297 | 6.309E-122 |
| Fibroblast | Ezh2 | 4 | 4.003E-125 | 0.46344125 | 0.783 | 0.362 | 8.005E-122 |
| Fibroblast | Nppb1 | 4 | 5.85E-125 | 0.54326264 | 0.623 | 0.234 | 1.17E-121 |
| Fibroblast | Gal3st3 | 4 | 3.094E-124 | 0.32111178 | 0.486 | 0.142 | 6.189E-121 |
| Fibroblast | Trhde1 | 4 | 3.871E-124 | 1.60115802 | 0.507 | 0.258 | 7.742E-121 |
| Fibroblast | Cyba | 4 | 4.065E-124 | 0.36460808 | 0.708 | 0.299 | 8.129E-121 |
| Fibroblast | Dpyd | 4 | 9.226E-124 | 1.384093 | 0.721 | 0.35 | 1.845E-120 |
| Fibroblast | Sorcs1 | 4 | 1.469E-123 | 0.98536716 | 0.426 | 0.176 | 2.938E-120 |
| Fibroblast | Tenm41 | 4 | 1.413E-122 | 1.06123421 | 0.515 | 0.222 | 2.825E-119 |

| | | | | | | | |
|---|---|---|---|---|---|---|---|
| Fibroblast | Stap1 | 4 | 3.95E-121 | 0.86509243 | 0.326 | 0.077 | 7.901E-118 |
| Fibroblast | Cd741 | 4 | 2.213E-120 | 0.25460834 | 0.601 | 0.242 | 4.426E-117 |
| Fibroblast | Pik3ap11 | 4 | 9.722E-119 | 0.67404054 | 0.578 | 0.249 | 1.944E-115 |
| Fibroblast | Mmp12 | 4 | 9.505E-117 | 1.54029333 | 0.34 | 0.107 | 1.901E-113 |
| Fibroblast | Crlf1 | 4 | 1.292E-116 | 0.90138945 | 0.535 | 0.226 | 2.584E-113 |
| Fibroblast | Plac8 | 4 | 5.34E-116 | 0.57695561 | 0.303 | 0.108 | 1.068E-112 |
| Fibroblast | Iqub1 | 4 | 8.295E-116 | 0.76801263 | 0.635 | 0.287 | 1.659E-112 |
| Fibroblast | Ccdc148 | 4 | 1.665E-115 | 0.51832768 | 0.379 | 0.126 | 3.33E-112 |
| Fibroblast | Caln1 | 4 | 3.081E-115 | 0.67957478 | 0.394 | 0.105 | 6.161E-112 |
| Fibroblast | Nova11 | 4 | 8.271E-115 | 1.07565838 | 0.641 | 0.311 | 1.654E-111 |
| Fibroblast | Adra1a2 | 4 | 1.297E-113 | 0.71037236 | 0.73 | 0.333 | 2.594E-110 |
| Fibroblast | Anln | 4 | 2.557E-113 | 0.70344954 | 0.56 | 0.217 | 5.113E-110 |
| Fibroblast | Kcnn31 | 4 | 3.868E-113 | 1.54787881 | 0.816 | 0.508 | 7.736E-110 |
| Fibroblast | AABR070265 | 4 | 3.884E-113 | 0.46872772 | 0.68 | 0.246 | 7.767E-110 |
| Fibroblast | Rapgef52 | 4 | 1.378E-112 | 0.62865826 | 0.561 | 0.236 | 2.755E-109 |
| Fibroblast | Dlgap11 | 4 | 5.167E-112 | 0.42508237 | 0.744 | 0.356 | 1.033E-108 |
| Fibroblast | Tbx51 | 4 | 3.655E-111 | 0.75615103 | 0.576 | 0.256 | 7.31E-108 |
| Fibroblast | Mctp21 | 4 | 4.264E-111 | 0.57714857 | 0.545 | 0.216 | 8.527E-108 |
| Fibroblast | Col27a1 | 4 | 5.885E-111 | 0.2964849 | 0.777 | 0.393 | 1.177E-107 |
| Fibroblast | Esr11 | 4 | 1.134E-110 | 0.93879897 | 0.601 | 0.269 | 2.268E-107 |
| Fibroblast | Gria1 | 4 | 2.105E-110 | 1.93912412 | 0.474 | 0.271 | 4.209E-107 |
| Fibroblast | Mypn1 | 4 | 8.434E-110 | 0.49408506 | 0.606 | 0.276 | 1.687E-106 |
| Fibroblast | Insc1 | 4 | 3.27E-109 | 1.52002472 | 0.352 | 0.157 | 6.541E-106 |
| Fibroblast | LOC1001348 | 4 | 2.324E-108 | 0.35512082 | 0.552 | 0.224 | 4.648E-105 |
| Fibroblast | Arhgap221 | 4 | 2.358E-108 | 1.20507253 | 0.67 | 0.352 | 4.715E-105 |
| Fibroblast | Abcd2 | 4 | 2.404E-108 | 0.799912 | 0.531 | 0.173 | 4.809E-105 |
| Fibroblast | Slc30a3 | 4 | 1.319E-107 | 0.92053283 | 0.338 | 0.133 | 2.638E-104 |
| Fibroblast | Asb91 | 4 | 1.089E-106 | 0.56531376 | 0.478 | 0.211 | 2.177E-103 |
| Fibroblast | Sh3gl21 | 4 | 1.435E-106 | 0.72168616 | 0.621 | 0.275 | 2.869E-103 |
| Fibroblast | Slc7a11 | 4 | 7.64E-106 | 0.52860857 | 0.726 | 0.337 | 1.528E-102 |
| Fibroblast | RT1-Ba | 4 | 2.474E-104 | 0.48158766 | 0.304 | 0.109 | 4.949E-101 |

| | | | | | | | |
|---|---|---|---|---|---|---|---|
| Fibroblast | Mbp1 | 4 | 5.658E-104 | 1.12022965 | 0.536 | 0.3 | 1.132E-100 |
| Fibroblast | Grk31 | 4 | 7.54E-104 | 0.72436448 | 0.516 | 0.207 | 1.508E-100 |
| Fibroblast | Egr1 | 4 | 3.416E-103 | 0.80706595 | 0.523 | 0.235 | 6.833E-100 |
| Fibroblast | Ankrd55 | 4 | 1.209E-102 | 0.90544621 | 0.374 | 0.198 | 2.418E-99 |
| Fibroblast | Itga4 | 4 | 1.487E-102 | 0.53495624 | 0.29 | 0.109 | 2.974E-99 |
| Fibroblast | Acad101 | 4 | 1.497E-102 | 0.43284371 | 0.675 | 0.303 | 2.995E-99 |
| Fibroblast | Steap41 | 4 | 1.774E-102 | 0.62724383 | 0.478 | 0.227 | 3.547E-99 |
| Fibroblast | Lrrc2 | 4 | 8.08E-102 | 0.30254988 | 0.455 | 0.191 | 1.6159E-98 |
| Fibroblast | Xkr41 | 4 | 1.391E-101 | 1.09968138 | 0.594 | 0.276 | 2.781E-98 |
| Fibroblast | Igfbp52 | 4 | 2.712E-101 | 1.08942503 | 0.664 | 0.332 | 5.424E-98 |
| Fibroblast | Atp5mc11 | 4 | 6.487E-101 | 0.42542216 | 0.656 | 0.287 | 1.2973E-97 |
| Fibroblast | Fabp31 | 4 | 6.754E-101 | 0.30946425 | 0.639 | 0.284 | 1.3507E-97 |
| Fibroblast | Coq8a2 | 4 | 2.016E-100 | 0.35023999 | 0.546 | 0.248 | 4.0329E-97 |
| Fibroblast | Irf81 | 4 | 4.67E-100 | 0.6782966 | 0.531 | 0.26 | 9.3407E-97 |
| Fibroblast | AABR070070 | 4 | 2.789E-99 | 1.10934478 | 0.297 | 0.155 | 5.5783E-96 |
| Fibroblast | Lmln2 | 4 | 4.629E-99 | 0.54510592 | 0.563 | 0.258 | 9.2577E-96 |
| Fibroblast | Il2ra | 4 | 1.1458E-98 | 2.46401468 | 0.4 | 0.15 | 2.2916E-95 |
| Fibroblast | Osbp21 | 4 | 1.8146E-98 | 0.65698965 | 0.636 | 0.338 | 3.6292E-95 |
| Fibroblast | Ppargc1b1 | 4 | 1.8727E-98 | 1.24448394 | 0.626 | 0.327 | 3.7453E-95 |
| Fibroblast | Ckap2l | 4 | 3.8783E-98 | 0.77360013 | 0.413 | 0.125 | 7.7565E-95 |
| Fibroblast | Lrrc4b | 4 | 4.8887E-98 | 0.91085348 | 0.364 | 0.173 | 9.7773E-95 |
| Fibroblast | Dpf32 | 4 | 8.2121E-98 | 0.42204618 | 0.492 | 0.241 | 1.6424E-94 |
| Fibroblast | Srl | 4 | 2.1316E-97 | 0.47823758 | 0.591 | 0.283 | 4.2631E-94 |
| Fibroblast | Kcnq3 | 4 | 1.1235E-96 | 0.35254892 | 0.421 | 0.127 | 2.247E-93 |
| Fibroblast | Prlr | 4 | 2.2464E-96 | 0.73184327 | 0.369 | 0.204 | 4.4928E-93 |
| Fibroblast | Erbb41 | 4 | 8.8164E-96 | 0.54270243 | 0.487 | 0.234 | 1.7633E-92 |
| Fibroblast | Adgrg6 | 4 | 5.9539E-95 | 0.5085513 | 0.427 | 0.195 | 1.1908E-91 |
| Fibroblast | AABR070525 | 4 | 8.8443E-95 | 0.58257215 | 0.546 | 0.251 | 1.7689E-91 |
| Fibroblast | Prom11 | 4 | 2.1107E-94 | 0.60137997 | 0.521 | 0.264 | 4.2215E-91 |
| Fibroblast | AABR070515 | 4 | 3.1023E-94 | 1.09519713 | 0.454 | 0.228 | 6.2046E-91 |
| Fibroblast | Ntrk22 | 4 | 3.8618E-94 | 0.58110258 | 0.599 | 0.303 | 7.7236E-91 |

| | | | | | | | |
|---|---|---|---|---|---|---|---|
| Fibroblast | Iqgap2 | 4 | 1.1406E-93 | 0.60397985 | 0.576 | 0.297 | 2.2812E-90 |
| Fibroblast | Apobec1 | 4 | 1.5162E-93 | 0.65208325 | 0.539 | 0.212 | 3.0324E-90 |
| Fibroblast | Csf1r | 4 | 2.1057E-93 | 0.33029041 | 0.471 | 0.178 | 4.2114E-90 |
| Fibroblast | Gucy1a21 | 4 | 2.5979E-93 | 1.027619 | 0.852 | 0.459 | 5.1958E-90 |
| Fibroblast | Cnnm1 | 4 | 4.0235E-93 | 1.48846815 | 0.312 | 0.062 | 8.0471E-90 |
| Fibroblast | Ablim3 | 4 | 6.3777E-93 | 0.43866908 | 0.463 | 0.209 | 1.2755E-89 |
| Fibroblast | Abcc41 | 4 | 2.4363E-92 | 0.55585488 | 0.628 | 0.334 | 4.8725E-89 |
| Fibroblast | Synpo2l | 4 | 1.1876E-91 | 1.17079675 | 0.475 | 0.214 | 2.3753E-88 |
| Fibroblast | Fa2h1 | 4 | 3.0592E-91 | 2.78021253 | 0.369 | 0.106 | 6.1185E-88 |
| Fibroblast | Mamdc22 | 4 | 6.1348E-91 | 0.56919647 | 0.637 | 0.304 | 1.227E-87 |
| Fibroblast | Apln | 4 | 1.0801E-90 | 1.5132771 | 0.299 | 0.057 | 2.1601E-87 |
| Fibroblast | Flt1 | 4 | 1.9666E-90 | 0.38530738 | 0.544 | 0.262 | 3.9331E-87 |
| Fibroblast | Ckb1 | 4 | 2.4604E-90 | 0.29180183 | 0.568 | 0.277 | 4.9207E-87 |
| Fibroblast | Ppp1r16b | 4 | 2.5422E-90 | 0.70971552 | 0.465 | 0.246 | 5.0845E-87 |
| Fibroblast | Colec121 | 4 | 1.0034E-89 | 1.21481417 | 0.91 | 0.6 | 2.0069E-86 |
| Fibroblast | Depdc1 | 4 | 1.228E-89 | 0.44273734 | 0.352 | 0.062 | 2.456E-86 |
| Fibroblast | Tesc1 | 4 | 1.2468E-89 | 0.39636896 | 0.524 | 0.259 | 2.4936E-86 |
| Fibroblast | Slco3a11 | 4 | 1.3625E-89 | 1.08800911 | 0.852 | 0.515 | 2.725E-86 |
| Fibroblast | Tox1 | 4 | 2.1706E-89 | 0.31093293 | 0.449 | 0.234 | 4.3411E-86 |
| Fibroblast | Ephb1 | 4 | 3.3153E-89 | 1.60175388 | 0.485 | 0.295 | 6.6306E-86 |
| Fibroblast | Aatk1 | 4 | 4.2038E-89 | 1.38557423 | 0.406 | 0.244 | 8.4076E-86 |
| Fibroblast | Ednrb | 4 | 8.6028E-89 | 0.40056576 | 0.393 | 0.173 | 1.7206E-85 |
| Fibroblast | Cap21 | 4 | 9.2254E-89 | 0.47662988 | 0.633 | 0.309 | 1.8451E-85 |
| Fibroblast | Apcdd1l1 | 4 | 1.6606E-88 | 0.61668012 | 0.505 | 0.295 | 3.3211E-85 |
| Fibroblast | Fabp41 | 4 | 5.8187E-88 | 0.38426939 | 0.505 | 0.225 | 1.1637E-84 |
| Fibroblast | Hs3st3a11 | 4 | 5.0449E-87 | 0.91768044 | 0.472 | 0.289 | 1.009E-83 |
| Fibroblast | Adhfe11 | 4 | 1.1684E-86 | 0.54799543 | 0.418 | 0.223 | 2.3367E-83 |
| Fibroblast | Oprd1 | 4 | 1.6813E-86 | 1.9226775 | 0.32 | 0.195 | 3.3626E-83 |
| Fibroblast | Lamc22 | 4 | 3.0786E-86 | 0.97258247 | 0.419 | 0.23 | 6.1571E-83 |
| Fibroblast | Rimbp21 | 4 | 3.1206E-86 | 0.41160165 | 0.469 | 0.25 | 6.2412E-83 |
| Fibroblast | Dgkb1 | 4 | 1.7176E-85 | 1.03241835 | 0.717 | 0.389 | 3.4352E-82 |

| | | | | | | | |
|---|---|---|---|---|---|---|---|
| Fibroblast | Ano4 | 4 | 2.0345E-85 | 0.85879973 | 0.331 | 0.185 | 4.069E-82 |
| Fibroblast | Sfxn51 | 4 | 2.7748E-85 | 0.74156431 | 0.416 | 0.214 | 5.5496E-82 |
| Fibroblast | AC096301.11 | 4 | 1.0739E-84 | 0.65842611 | 0.418 | 0.227 | 2.1479E-81 |
| Fibroblast | Cacna1d | 4 | 1.7417E-84 | 0.84094843 | 0.255 | 0.167 | 3.4834E-81 |
| Fibroblast | Necab11 | 4 | 1.7866E-84 | 1.53009396 | 0.453 | 0.277 | 3.5732E-81 |
| Fibroblast | Emcn | 4 | 3.4215E-84 | 0.31719095 | 0.374 | 0.196 | 6.8431E-81 |
| Fibroblast | Dync1i1 | 4 | 6.5693E-84 | 0.54967785 | 0.315 | 0.209 | 1.3139E-80 |
| Fibroblast | Sgca | 4 | 1.612E-83 | 0.76370298 | 0.377 | 0.196 | 3.2239E-80 |
| Fibroblast | Zfp385b1 | 4 | 1.7131E-83 | 1.18525187 | 0.713 | 0.429 | 3.4262E-80 |
| Fibroblast | Tbx201 | 4 | 1.7957E-83 | 1.105565 | 0.753 | 0.52 | 3.5915E-80 |
| Fibroblast | Hsd11b11 | 4 | 3.6874E-83 | 1.07479712 | 0.551 | 0.326 | 7.3749E-80 |
| Fibroblast | Pkia1 | 4 | 4.9957E-83 | 0.34079335 | 0.505 | 0.251 | 9.9913E-80 |
| Fibroblast | Aqp1 | 4 | 1.4017E-82 | 0.37874948 | 0.275 | 0.144 | 2.8035E-79 |
| Fibroblast | Myocd1 | 4 | 7.0119E-82 | 0.39937833 | 0.517 | 0.264 | 1.4024E-78 |
| Fibroblast | Rnf43 | 4 | 9.577E-82 | 0.53374364 | 0.428 | 0.153 | 1.9154E-78 |
| Fibroblast | Prox1 | 4 | 1.7858E-81 | 0.56662272 | 0.333 | 0.203 | 3.5716E-78 |
| Fibroblast | Col18a1 | 4 | 8.8981E-81 | 0.50979578 | 0.47 | 0.235 | 1.7796E-77 |
| Fibroblast | Pcdh71 | 4 | 3.4992E-80 | 0.77857199 | 0.472 | 0.254 | 6.9984E-77 |
| Fibroblast | Bmper1 | 4 | 4.4756E-80 | 1.08138336 | 0.741 | 0.523 | 8.9512E-77 |
| Fibroblast | Pparg1 | 4 | 4.087E-79 | 1.01080104 | 0.47 | 0.255 | 8.1739E-76 |
| Fibroblast | Slc26a10 | 4 | 6.2498E-79 | 0.46162407 | 0.454 | 0.241 | 1.25E-75 |
| Fibroblast | Unc13b | 4 | 2.4144E-78 | 0.40208388 | 0.427 | 0.203 | 4.8289E-75 |
| Fibroblast | AABR070251 | 4 | 2.9599E-78 | 0.38594641 | 0.325 | 0.187 | 5.9198E-75 |
| Fibroblast | Ldhb1 | 4 | 4.8176E-78 | 0.36970578 | 0.732 | 0.385 | 9.6351E-75 |
| Fibroblast | Gnao12 | 4 | 6.2682E-78 | 0.51763598 | 0.583 | 0.324 | 1.2536E-74 |
| Fibroblast | Mki671 | 4 | 7.2283E-78 | 0.27241255 | 0.539 | 0.233 | 1.4457E-74 |
| Fibroblast | Dusp27 | 4 | 8.2057E-78 | 0.29378682 | 0.485 | 0.147 | 1.6411E-74 |
| Fibroblast | Colq | 4 | 1.4863E-77 | 1.39361642 | 0.333 | 0.202 | 2.9726E-74 |
| Fibroblast | Fgf122 | 4 | 5.5669E-77 | 0.68617557 | 0.562 | 0.295 | 1.1134E-73 |
| Fibroblast | Bmp62 | 4 | 7.372E-77 | 1.07867865 | 0.661 | 0.467 | 1.4744E-73 |
| Fibroblast | Mlip1 | 4 | 1.2485E-76 | 0.51306723 | 0.536 | 0.269 | 2.497E-73 |

| | | | | | | | |
|---|---|---|---|---|---|---|---|
| Fibroblast | Ar1 | 4 | 5.7725E-76 | 1.15414697 | 0.817 | 0.544 | 1.1545E-72 |
| Fibroblast | Agbl11 | 4 | 7.8512E-76 | 1.12902204 | 0.562 | 0.357 | 1.5702E-72 |
| Fibroblast | Prc1 | 4 | 3.3166E-75 | 0.54207368 | 0.369 | 0.106 | 6.6332E-72 |
| Fibroblast | Tmcc31 | 4 | 1.186E-74 | 0.65074533 | 0.618 | 0.351 | 2.3721E-71 |
| Fibroblast | Scarb1 | 4 | 1.8656E-74 | 0.33297309 | 0.452 | 0.245 | 3.7311E-71 |
| Fibroblast | Hs6st21 | 4 | 4.6025E-74 | 0.71005113 | 0.688 | 0.402 | 9.2049E-71 |
| Fibroblast | Foxp21 | 4 | 7.3929E-74 | 1.01436204 | 0.858 | 0.546 | 1.4786E-70 |
| Fibroblast | Cmya51 | 4 | 2.4385E-73 | 0.3843806 | 0.685 | 0.34 | 4.8771E-70 |
| Fibroblast | Katnal2 | 4 | 3.795E-73 | 1.11094789 | 0.37 | 0.144 | 7.5901E-70 |
| Fibroblast | Grm81 | 4 | 3.6728E-72 | 0.53833346 | 0.47 | 0.221 | 7.3457E-69 |
| Fibroblast | Myh7b1 | 4 | 1.0806E-71 | 0.34234194 | 0.581 | 0.314 | 2.1613E-68 |
| Fibroblast | Gpm6a | 4 | 1.639E-70 | 1.04128854 | 0.705 | 0.469 | 3.2779E-67 |
| Fibroblast | Asb18 | 4 | 2.2152E-70 | 0.52627897 | 0.469 | 0.27 | 4.4304E-67 |
| Fibroblast | Nrk1 | 4 | 3.3183E-70 | 0.65957055 | 0.593 | 0.364 | 6.6366E-67 |
| Fibroblast | Ros1 | 4 | 5.3827E-70 | 0.58058594 | 0.393 | 0.202 | 1.0765E-66 |
| Fibroblast | B2m | 4 | 8.223E-70 | 0.44441317 | 0.784 | 0.396 | 1.6446E-66 |
| Fibroblast | Megf10 | 4 | 1.7308E-69 | 0.55096797 | 0.443 | 0.153 | 3.4616E-66 |
| Fibroblast | Acacb1 | 4 | 3.3733E-69 | 0.55420433 | 0.493 | 0.269 | 6.7466E-66 |
| Fibroblast | Lrrc7 | 4 | 7.5454E-69 | 0.31817287 | 0.292 | 0.193 | 1.5091E-65 |
| Fibroblast | Myh15 | 4 | 1.3205E-68 | 0.58785039 | 0.413 | 0.108 | 2.6409E-65 |
| Fibroblast | Ngf1 | 4 | 1.4543E-68 | 1.07003775 | 0.817 | 0.551 | 2.9086E-65 |
| Fibroblast | Rnf1501 | 4 | 4.8171E-68 | 1.21658444 | 0.485 | 0.315 | 9.6342E-65 |
| Fibroblast | Slc7a81 | 4 | 1.1695E-67 | 0.93282314 | 0.433 | 0.293 | 2.339E-64 |
| Fibroblast | Efemp12 | 4 | 1.1934E-67 | 0.57296922 | 0.523 | 0.3 | 2.3869E-64 |
| Fibroblast | Ank31 | 4 | 2.6873E-67 | 0.45680421 | 0.498 | 0.276 | 5.3745E-64 |
| Fibroblast | Inpp4b | 4 | 6.6533E-67 | 0.27022411 | 0.297 | 0.202 | 1.3307E-63 |
| Fibroblast | Rims1 | 4 | 8.7253E-67 | 0.37529784 | 0.287 | 0.124 | 1.7451E-63 |
| Fibroblast | Ect2 | 4 | 9.8326E-67 | 0.51231108 | 0.453 | 0.166 | 1.9665E-63 |
| Fibroblast | Shank3 | 4 | 4.9227E-66 | 0.25038077 | 0.459 | 0.268 | 9.8454E-63 |
| Fibroblast | Slc43a3 | 4 | 2.9528E-65 | 1.28445928 | 0.439 | 0.309 | 5.9055E-62 |
| Fibroblast | Vgll31 | 4 | 3.3627E-65 | 0.35558082 | 0.66 | 0.385 | 6.7255E-62 |

| | | | | | | | |
|---|---|---|---|---|---|---|---|
| Fibroblast | AABR070606 | 4 | 1.5895E-64 | 0.88426997 | 0.391 | 0.222 | 3.1791E-61 |
| Fibroblast | Slc16a21 | 4 | 3.8972E-64 | 0.55481002 | 0.71 | 0.398 | 7.7944E-61 |
| Fibroblast | Naca1 | 4 | 1.0976E-63 | 0.31497984 | 0.546 | 0.299 | 2.1951E-60 |
| Fibroblast | Negr11 | 4 | 1.5666E-63 | 0.46440308 | 0.77 | 0.453 | 3.1331E-60 |
| Fibroblast | Pcbp31 | 4 | 1.1707E-62 | 0.38321017 | 0.632 | 0.367 | 2.3414E-59 |
| Fibroblast | Txlnb1 | 4 | 1.9218E-61 | 0.30920743 | 0.5 | 0.305 | 3.8436E-58 |
| Fibroblast | Gsn1 | 4 | 3.7891E-61 | 0.82685209 | 0.814 | 0.556 | 7.5783E-58 |
| Fibroblast | Errfi1 | 4 | 6.6917E-61 | 0.4907592 | 0.764 | 0.458 | 1.3383E-57 |
| Fibroblast | Cyfip2 | 4 | 7.8779E-61 | 0.69596413 | 0.347 | 0.112 | 1.5756E-57 |
| Fibroblast | Myom11 | 4 | 4.5508E-60 | 0.36162429 | 0.459 | 0.278 | 9.1016E-57 |
| Fibroblast | Nppa | 4 | 6.2826E-60 | 0.38841265 | 0.257 | 0.205 | 1.2565E-56 |
| Fibroblast | Gpd1 | 4 | 1.0627E-59 | 0.31839157 | 0.42 | 0.128 | 2.1253E-56 |
| Fibroblast | Pde1c1 | 4 | 1.3075E-59 | 0.65517511 | 0.355 | 0.24 | 2.615E-56 |
| Fibroblast | Sh3kbp11 | 4 | 6.0714E-59 | 0.84627717 | 0.634 | 0.396 | 1.2143E-55 |
| Fibroblast | Gpx31 | 4 | 1.9246E-58 | 0.40782902 | 0.622 | 0.355 | 3.8492E-55 |
| Fibroblast | LOC1036899 | 4 | 3.7939E-58 | 0.33782499 | 0.8 | 0.447 | 7.5877E-55 |
| Fibroblast | Tex221 | 4 | 4.0156E-58 | 0.42362151 | 0.339 | 0.239 | 8.0313E-55 |
| Fibroblast | Pygm | 4 | 4.4185E-58 | 0.29061492 | 0.331 | 0.214 | 8.8369E-55 |
| Fibroblast | Ank21 | 4 | 7.1151E-58 | 0.84682404 | 0.967 | 0.845 | 1.423E-54 |
| Fibroblast | Inpp5d | 4 | 7.2679E-58 | 0.27097261 | 0.371 | 0.259 | 1.4536E-54 |
| Fibroblast | Ivns1abp1 | 4 | 1.7791E-57 | 0.98512155 | 0.484 | 0.337 | 3.5583E-54 |
| Fibroblast | Thsd7a | 4 | 2.2058E-57 | 0.29131433 | 0.319 | 0.243 | 4.4116E-54 |
| Fibroblast | Itgal | 4 | 2.2312E-57 | 0.8869893 | 0.396 | 0.181 | 4.4623E-54 |
| Fibroblast | Lyn1 | 4 | 6.343E-57 | 0.4697737 | 0.569 | 0.346 | 1.2686E-53 |
| Fibroblast | Ttyh11 | 4 | 6.4408E-57 | 1.92130792 | 0.317 | 0.139 | 1.2882E-53 |
| Fibroblast | Ntn11 | 4 | 1.5823E-56 | 0.72203035 | 0.763 | 0.487 | 3.1645E-53 |
| Fibroblast | Entpd1 | 4 | 9.7796E-56 | 0.30636717 | 0.505 | 0.354 | 1.9559E-52 |
| Fibroblast | Cit1 | 4 | 1.8286E-54 | 0.58004711 | 0.588 | 0.357 | 3.6572E-51 |
| Fibroblast | Ebf22 | 4 | 2.3336E-54 | 0.62033018 | 0.969 | 0.851 | 4.6671E-51 |
| Fibroblast | Trpc6 | 4 | 1.8926E-53 | 0.6240149 | 0.491 | 0.189 | 3.7853E-50 |
| Fibroblast | AC130940.1 | 4 | 2.5566E-53 | 0.91606956 | 0.402 | 0.158 | 5.1133E-50 |

| | | | | | | | |
|---|---|---|---|---|---|---|---|
| Fibroblast | Rbp11 | 4 | 6.129E-53 | 0.57532527 | 0.477 | 0.33 | 1.2258E-49 |
| Fibroblast | Chn11 | 4 | 1.2067E-52 | 1.37693185 | 0.309 | 0.167 | 2.4133E-49 |
| Fibroblast | Adcy22 | 4 | 1.8172E-52 | 0.91928547 | 0.437 | 0.324 | 3.6345E-49 |
| Fibroblast | Dnm3 | 4 | 2.4506E-52 | 0.34994028 | 0.792 | 0.461 | 4.9013E-49 |
| Fibroblast | Slc44a51 | 4 | 8.4095E-52 | 0.68720025 | 0.549 | 0.375 | 1.6819E-48 |
| Fibroblast | Stk26 | 4 | 6.8876E-51 | 0.74384652 | 0.304 | 0.129 | 1.3775E-47 |
| Fibroblast | Alpk31 | 4 | 1.0867E-50 | 0.42127207 | 0.362 | 0.268 | 2.1735E-47 |
| Fibroblast | Miga2 | 4 | 1.9576E-50 | 0.29911767 | 0.45 | 0.311 | 3.9151E-47 |
| Fibroblast | Lpl1 | 4 | 2.3447E-50 | 0.63206126 | 0.492 | 0.325 | 4.6894E-47 |
| Fibroblast | Cadps21 | 4 | 2.573E-50 | 0.38194591 | 0.336 | 0.238 | 5.146E-47 |
| Fibroblast | Zc3h12d | 4 | 2.8997E-50 | 0.4601551 | 0.35 | 0.197 | 5.7994E-47 |
| Fibroblast | Akap61 | 4 | 8.9844E-50 | 0.36986383 | 0.581 | 0.372 | 1.7969E-46 |
| Fibroblast | Slc4a3 | 4 | 1.9025E-49 | 0.35937301 | 0.471 | 0.308 | 3.805E-46 |
| Fibroblast | Sparcl1 | 4 | 6.7056E-49 | 0.64133129 | 0.681 | 0.456 | 1.3411E-45 |
| Fibroblast | Adcy51 | 4 | 6.8609E-49 | 0.64245847 | 0.56 | 0.389 | 1.3722E-45 |
| Fibroblast | Rbm20 | 4 | 2.0961E-48 | 0.50206915 | 0.807 | 0.499 | 4.1923E-45 |
| Fibroblast | Colgalt21 | 4 | 2.3291E-48 | 0.93819618 | 0.578 | 0.51 | 4.6583E-45 |
| Fibroblast | Cyp7b11 | 4 | 7.21E-48 | 0.31350477 | 0.718 | 0.434 | 1.442E-44 |
| Fibroblast | Inhba | 4 | 1.2773E-47 | 0.58604524 | 0.321 | 0.254 | 2.5547E-44 |
| Fibroblast | Asb2 | 4 | 1.4303E-47 | 0.44712121 | 0.343 | 0.238 | 2.8607E-44 |
| Fibroblast | Megf11 | 4 | 1.521E-47 | 1.3104567 | 0.414 | 0.207 | 3.0419E-44 |
| Fibroblast | Dapk12 | 4 | 2.6432E-47 | 0.7578721 | 0.864 | 0.637 | 5.2863E-44 |
| Fibroblast | Rerg | 4 | 1.9101E-46 | 0.90195224 | 0.421 | 0.328 | 3.8202E-43 |
| Fibroblast | Nmnat21 | 4 | 6.2373E-46 | 1.26757384 | 0.335 | 0.288 | 1.2475E-42 |
| Fibroblast | Htra31 | 4 | 1.1033E-44 | 0.88692976 | 0.588 | 0.511 | 2.2067E-41 |
| Fibroblast | Kcnt2 | 4 | 4.5848E-44 | 0.2897687 | 0.55 | 0.363 | 9.1695E-41 |
| Fibroblast | Eda1 | 4 | 1.0285E-43 | 0.72320517 | 0.864 | 0.646 | 2.057E-40 |
| Fibroblast | Gbp1 | 4 | 1.7853E-43 | 0.34254564 | 0.352 | 0.216 | 3.5706E-40 |
| Fibroblast | Musk1 | 4 | 4.5113E-43 | 0.4738936 | 0.463 | 0.346 | 9.0227E-40 |
| Fibroblast | Adamtsl31 | 4 | 6.8066E-43 | 0.72806479 | 0.893 | 0.717 | 1.3613E-39 |
| Fibroblast | Gpcpd1 | 4 | 7.129E-43 | 0.26347738 | 0.551 | 0.377 | 1.4258E-39 |

| | | | | | | | |
|---|---|---|---|---|---|---|---|
| Fibroblast | Sox6 | 4 | 9.2371E-43 | 0.49615361 | 0.67 | 0.472 | 1.8474E-39 |
| Fibroblast | Wwc1 | 4 | 1.0596E-42 | 0.4530273 | 0.331 | 0.132 | 2.1192E-39 |
| Fibroblast | Nt5e1 | 4 | 1.2088E-42 | 0.41501723 | 0.372 | 0.284 | 2.4176E-39 |
| Fibroblast | Tnik1 | 4 | 2.6812E-42 | 0.98937005 | 0.489 | 0.416 | 5.3624E-39 |
| Fibroblast | Itga71 | 4 | 6.5132E-42 | 0.28814175 | 0.266 | 0.244 | 1.3026E-38 |
| Fibroblast | Themis | 4 | 2.8377E-41 | 0.66883102 | 0.292 | 0.099 | 5.6754E-38 |
| Fibroblast | Calcrl | 4 | 6.2615E-41 | 0.73263435 | 0.811 | 0.584 | 1.2523E-37 |
| Fibroblast | AABR070070 | 4 | 7.1948E-41 | 0.60091283 | 0.863 | 0.578 | 1.439E-37 |
| Fibroblast | Pde3a1 | 4 | 1.1879E-40 | 0.66645639 | 0.639 | 0.499 | 2.3757E-37 |
| Fibroblast | Gpc61 | 4 | 5.3095E-40 | 0.45064255 | 0.999 | 0.978 | 1.0619E-36 |
| Fibroblast | Kifc1 | 4 | 8.3875E-40 | 1.60475059 | 0.325 | 0.127 | 1.6775E-36 |
| Fibroblast | Afap1l22 | 4 | 9.1484E-40 | 0.57213345 | 0.475 | 0.374 | 1.8297E-36 |
| Fibroblast | Fabp51 | 4 | 1.9787E-39 | 0.75022088 | 0.319 | 0.203 | 3.9574E-36 |
| Fibroblast | Cluh | 4 | 3.5712E-39 | 0.39856698 | 0.33 | 0.273 | 7.1425E-36 |
| Fibroblast | Pla2g7 | 4 | 1.4667E-38 | 0.39406274 | 0.411 | 0.148 | 2.9334E-35 |
| Fibroblast | Cspg41 | 4 | 1.2679E-37 | 0.86008407 | 0.265 | 0.25 | 2.5358E-34 |
| Fibroblast | Pcsk61 | 4 | 1.6543E-37 | 0.54339422 | 0.883 | 0.669 | 3.3086E-34 |
| Fibroblast | LOC1003644 | 4 | 6.2958E-37 | 0.29516737 | 0.545 | 0.377 | 1.2592E-33 |
| Fibroblast | Clybl1 | 4 | 2.1661E-36 | 0.86335952 | 0.499 | 0.407 | 4.3322E-33 |
| Fibroblast | Exph5 | 4 | 1.2001E-35 | 0.89529642 | 0.402 | 0.207 | 2.4003E-32 |
| Fibroblast | Vipr21 | 4 | 1.6295E-35 | 0.75320077 | 0.441 | 0.364 | 3.2589E-32 |
| Fibroblast | Btbd111 | 4 | 2.4206E-35 | 0.68498482 | 0.44 | 0.364 | 4.8412E-32 |
| Fibroblast | Arhgap242 | 4 | 5.297E-35 | 0.55059815 | 0.799 | 0.629 | 1.0594E-31 |
| Fibroblast | AC134204.1 | 4 | 1.2952E-34 | 0.36524974 | 0.777 | 0.488 | 2.5904E-31 |
| Fibroblast | Srpx | 4 | 5.387E-34 | 0.3091776 | 0.433 | 0.347 | 1.0774E-30 |
| Fibroblast | Maoa | 4 | 7.0494E-33 | 0.27131403 | 0.432 | 0.334 | 1.4099E-29 |
| Fibroblast | Arl151 | 4 | 1.2776E-32 | 0.47978068 | 0.97 | 0.899 | 2.5551E-29 |
| Fibroblast | Mpped22 | 4 | 4.8872E-32 | 0.55268096 | 0.652 | 0.495 | 9.7744E-29 |
| Fibroblast | Mylk1 | 4 | 5.4388E-32 | 0.38358124 | 0.816 | 0.544 | 1.0878E-28 |
| Fibroblast | Scn7a | 4 | 9.0132E-32 | 0.57336935 | 0.33 | 0.296 | 1.8026E-28 |
| Fibroblast | Rnls2 | 4 | 1.5903E-31 | 0.31062551 | 0.538 | 0.405 | 3.1806E-28 |

| | | | | | | | |
|---|---|---|---|---|---|---|---|
| Fibroblast | Mreg | 4 | 4.12E-31 | 0.58110703 | 0.256 | 0.141 | 8.24E-28 |
| Fibroblast | Insr1 | 4 | 1.8461E-30 | 0.39314069 | 0.606 | 0.446 | 3.6921E-27 |
| Fibroblast | Agap2 | 4 | 2.6794E-30 | 1.00146158 | 0.268 | 0.123 | 5.3588E-27 |
| Fibroblast | Gli21 | 4 | 1.2779E-28 | 0.54936292 | 0.718 | 0.582 | 2.5558E-25 |
| Fibroblast | Col5a31 | 4 | 2.8152E-28 | 0.49676868 | 0.573 | 0.495 | 5.6303E-25 |
| Fibroblast | Rmdn11 | 4 | 8.9778E-28 | 0.39372672 | 0.627 | 0.467 | 1.7956E-24 |
| Fibroblast | Ntrk31 | 4 | 1.3115E-27 | 0.56808608 | 0.359 | 0.325 | 2.623E-24 |
| Fibroblast | Col4a51 | 4 | 1.1882E-26 | 0.52384985 | 0.888 | 0.714 | 2.3763E-23 |
| Fibroblast | Sox131 | 4 | 1.5695E-26 | 0.46954924 | 0.436 | 0.393 | 3.1389E-23 |
| Fibroblast | Med12l | 4 | 1.6801E-26 | 0.42872984 | 0.31 | 0.301 | 3.3602E-23 |
| Fibroblast | Vegfc1 | 4 | 8.7253E-26 | 0.2834255 | 0.465 | 0.384 | 1.7451E-22 |
| Fibroblast | Magi21 | 4 | 1.9207E-25 | 0.36261299 | 0.381 | 0.328 | 3.8414E-22 |
| Fibroblast | Camk4 | 4 | 6.5955E-25 | 0.82211211 | 0.273 | 0.114 | 1.3191E-21 |
| Fibroblast | Fap | 4 | 1.721E-24 | 0.41869017 | 0.869 | 0.61 | 3.442E-21 |
| Fibroblast | Cacna1a1 | 4 | 7.5368E-24 | 0.44471338 | 0.645 | 0.5 | 1.5074E-20 |
| Fibroblast | Vegfa1 | 4 | 1.1893E-23 | 0.33070964 | 0.787 | 0.652 | 2.3786E-20 |
| Fibroblast | AABR070349 | 4 | 1.3233E-23 | 0.29663613 | 0.304 | 0.261 | 2.6466E-20 |
| Fibroblast | Col6a21 | 4 | 1.7258E-23 | 0.49176725 | 0.75 | 0.668 | 3.4516E-20 |
| Fibroblast | AABR070546 | 4 | 1.9045E-23 | 0.41851477 | 0.696 | 0.556 | 3.809E-20 |
| Fibroblast | Itgbl11 | 4 | 3.2678E-23 | 0.38904963 | 0.921 | 0.874 | 6.5356E-20 |
| Fibroblast | Otulinl1 | 4 | 3.5353E-23 | 0.26234735 | 0.39 | 0.237 | 7.0705E-20 |
| Fibroblast | Il1r11 | 4 | 7.7511E-23 | 0.34494305 | 0.872 | 0.702 | 1.5502E-19 |
| Fibroblast | Myl21 | 4 | 3.6824E-22 | 0.3041752 | 0.833 | 0.567 | 7.3649E-19 |
| Fibroblast | Pde4b | 4 | 3.8369E-22 | 0.58557257 | 0.438 | 0.425 | 7.6738E-19 |
| Fibroblast | Ror11 | 4 | 1.9258E-21 | 0.41479947 | 0.614 | 0.502 | 3.8515E-18 |
| Fibroblast | Ryr21 | 4 | 2.2805E-21 | 0.32854454 | 0.73 | 0.59 | 4.5611E-18 |
| Fibroblast | Cldn15 | 4 | 3.7614E-21 | 0.35507838 | 0.392 | 0.238 | 7.5228E-18 |
| Fibroblast | Ackr31 | 4 | 3.0535E-19 | 0.31639242 | 0.521 | 0.46 | 6.107E-16 |
| Fibroblast | Slc9a9 | 4 | 5.8737E-19 | 0.37641757 | 0.728 | 0.572 | 1.1747E-15 |
| Fibroblast | Robo1 | 4 | 1.7348E-18 | 0.41304883 | 0.83 | 0.646 | 3.4696E-15 |
| Fibroblast | Enox11 | 4 | 1.8179E-18 | 0.27061634 | 0.448 | 0.419 | 3.6359E-15 |

| | | | | | | | |
|---|---|---|---|---|---|---|---|
| Fibroblast | Ninj2 | 4 | 6.4221E-18 | 0.27392724 | 0.662 | 0.525 | 1.2844E-14 |
| Fibroblast | Egfr | 4 | 2.9053E-14 | 0.3824275 | 0.885 | 0.695 | 5.8105E-11 |
| Fibroblast | Chsy31 | 4 | 7.2841E-14 | 0.29660004 | 0.572 | 0.54 | 1.4568E-10 |
| Fibroblast | Hs3st2 | 4 | 3.4989E-13 | 0.45013297 | 0.278 | 0.186 | 6.9977E-10 |
| Fibroblast | Arhgap21 | 4 | 3.8826E-12 | 0.34959431 | 0.78 | 0.67 | 7.7652E-09 |
| Fibroblast | Dcn1 | 4 | 6.316E-11 | 0.27522264 | 0.847 | 0.79 | 1.2632E-07 |
| Fibroblast | Kif231 | 4 | 7.2644E-11 | 0.8294177 | 0.331 | 0.151 | 1.4529E-07 |
| Fibroblast | AABR070637 | 4 | 1.7338E-10 | 1.01797355 | 0.269 | 0.125 | 3.4676E-07 |
| Fibroblast | Cdh11 | 4 | 3.9174E-10 | 0.27785729 | 0.795 | 0.678 | 7.8347E-07 |
| Fibroblast | Cpq1 | 4 | 2.2235E-07 | 0.25509422 | 0.813 | 0.75 | 0.0004447 |
| Fibroblast | Dmd1 | 4 | 2.6891E-06 | 0.25417758 | 0.807 | 0.763 | 0.00537818 |
| Fibroblast | Axl1 | 5 | 2.4605E-52 | 1.06733807 | 0.776 | 0.675 | 4.921E-49 |
| Fibroblast | Hhip | 5 | 1.7402E-42 | 0.77530083 | 0.275 | 0.099 | 3.4803E-39 |
| Fibroblast | Uap11 | 5 | 6.7169E-42 | 0.99574121 | 0.75 | 0.616 | 1.3434E-38 |
| Fibroblast | Tfap2b | 5 | 3.6032E-34 | 2.7123861 | 0.265 | 0.105 | 7.2064E-31 |
| Fibroblast | AC110709.21 | 5 | 8.0272E-32 | 0.7242327 | 0.112 | 0.265 | 1.6054E-28 |
| Fibroblast | Slco3a12 | 5 | 1.1675E-30 | 1.1848189 | 0.658 | 0.54 | 2.335E-27 |
| Fibroblast | Cdh131 | 5 | 2.9908E-29 | 1.13448763 | 0.667 | 0.575 | 5.9817E-26 |
| Fibroblast | Vegfa2 | 5 | 1.842E-25 | 0.71657111 | 0.756 | 0.658 | 3.6839E-22 |
| Fibroblast | Mctp22 | 5 | 1.198E-23 | 0.36118364 | 0.084 | 0.264 | 2.3959E-20 |
| Fibroblast | Ebf23 | 5 | 2.5292E-23 | 0.49624687 | 0.877 | 0.862 | 5.0583E-20 |
| Fibroblast | Cxcl101 | 5 | 1.3215E-22 | 0.69990132 | 0.294 | 0.165 | 2.6431E-19 |
| Fibroblast | Mapt1 | 5 | 1.0143E-21 | 0.76962674 | 0.121 | 0.254 | 2.0287E-18 |
| Fibroblast | Pik3cd | 5 | 2.8221E-21 | 0.30869047 | 0.147 | 0.323 | 5.6443E-18 |
| Fibroblast | Trpc3 | 5 | 1.8536E-20 | 0.43497124 | 0.162 | 0.273 | 3.7072E-17 |
| Fibroblast | Cd163 | 5 | 4.3099E-19 | 0.41586627 | 0.147 | 0.27 | 8.6198E-16 |
| Fibroblast | Adgrd11 | 5 | 7.5776E-19 | 0.71845436 | 0.69 | 0.627 | 1.5155E-15 |
| Fibroblast | Adap21 | 5 | 2.5407E-18 | 0.25655077 | 0.151 | 0.252 | 5.0813E-15 |
| Fibroblast | Ptprc1 | 5 | 1.2889E-17 | 0.44623282 | 0.129 | 0.27 | 2.5778E-14 |
| Fibroblast | Tbxas1 | 5 | 1.8883E-17 | 0.57571378 | 0.113 | 0.269 | 3.7765E-14 |
| Fibroblast | Pcsk62 | 5 | 2.1414E-17 | 0.7461386 | 0.729 | 0.687 | 4.2827E-14 |

| | | | | | | | |
|---|---|---|---|---|---|---|---|
| Fibroblast | Scarb11 | 5 | 2.639E-17 | 0.36687228 | 0.142 | 0.278 | 5.2779E-14 |
| Fibroblast | Slfn4 | 5 | 1.0712E-16 | 0.49844109 | 0.191 | 0.314 | 2.1424E-13 |
| Fibroblast | Ptprj1 | 5 | 2.9119E-15 | 0.95021508 | 0.613 | 0.582 | 5.8238E-12 |
| Fibroblast | Arhgef372 | 5 | 1.197E-14 | 0.3730427 | 0.165 | 0.271 | 2.394E-11 |
| Fibroblast | Tmem511 | 5 | 1.2102E-14 | 0.39426393 | 0.129 | 0.267 | 2.4205E-11 |
| Fibroblast | Cspg42 | 5 | 1.7771E-14 | 0.63326834 | 0.149 | 0.26 | 3.5542E-11 |
| Fibroblast | P2rx7 | 5 | 2.0105E-14 | 0.39927069 | 0.209 | 0.31 | 4.021E-11 |
| Fibroblast | AABR070070 | 5 | 4.2885E-14 | 0.76851849 | 0.666 | 0.602 | 8.577E-11 |
| Fibroblast | Camk1d1 | 5 | 5.3655E-14 | 0.43009695 | 0.172 | 0.301 | 1.0731E-10 |
| Fibroblast | Popdc22 | 5 | 1.2644E-13 | 0.38602733 | 0.202 | 0.295 | 2.5288E-10 |
| Fibroblast | Lamc23 | 5 | 2.7561E-13 | 0.39774004 | 0.122 | 0.261 | 5.5122E-10 |
| Fibroblast | Thbs1 | 5 | 8.9985E-13 | 0.34288673 | 0.236 | 0.329 | 1.7997E-09 |
| Fibroblast | RGD1563354 | 5 | 1.179E-12 | 0.8115328 | 0.608 | 0.531 | 2.3579E-09 |
| Fibroblast | Cybb | 5 | 1.5547E-12 | 0.49596007 | 0.181 | 0.269 | 3.1094E-09 |
| Fibroblast | Reep11 | 5 | 1.8784E-12 | 0.54238308 | 0.181 | 0.255 | 3.7567E-09 |
| Fibroblast | Sptb1 | 5 | 1.9373E-12 | 0.28036196 | 0.22 | 0.299 | 3.8747E-09 |
| Fibroblast | Samd121 | 5 | 1.9958E-12 | 0.44692371 | 0.22 | 0.321 | 3.9915E-09 |
| Fibroblast | Stab11 | 5 | 2.6002E-12 | 0.32398935 | 0.187 | 0.255 | 5.2003E-09 |
| Fibroblast | AABR070265 | 5 | 6.4892E-12 | 0.49736198 | 0.232 | 0.296 | 1.2978E-08 |
| Fibroblast | Cabcoco12 | 5 | 2.1716E-11 | 0.2501575 | 0.209 | 0.278 | 4.3432E-08 |
| Fibroblast | Dclk11 | 5 | 4.9057E-11 | 0.5917061 | 0.749 | 0.721 | 9.8113E-08 |
| Fibroblast | Casq21 | 5 | 5.4647E-11 | 0.25984133 | 0.219 | 0.332 | 1.0929E-07 |
| Fibroblast | Gda1 | 5 | 7.6743E-11 | 0.52882524 | 0.764 | 0.759 | 1.5349E-07 |
| Fibroblast | Entpd11 | 5 | 8.3696E-11 | 0.34940672 | 0.303 | 0.375 | 1.6739E-07 |
| Fibroblast | Irs2 | 5 | 8.6472E-11 | 0.35511041 | 0.307 | 0.411 | 1.7294E-07 |
| Fibroblast | Ankrd61 | 5 | 8.9704E-11 | 0.78568015 | 0.187 | 0.273 | 1.7941E-07 |
| Fibroblast | Smpx2 | 5 | 1.1451E-10 | 0.4832577 | 0.202 | 0.284 | 2.2902E-07 |
| Fibroblast | Inhba1 | 5 | 1.1839E-10 | 0.72436966 | 0.195 | 0.267 | 2.3677E-07 |
| Fibroblast | Nrcam1 | 5 | 1.7293E-10 | 1.03551476 | 0.222 | 0.278 | 3.4586E-07 |
| Fibroblast | Egr11 | 5 | 2.0866E-10 | 0.90522372 | 0.192 | 0.271 | 4.1731E-07 |
| Fibroblast | Rgs171 | 5 | 2.7328E-10 | 0.2635275 | 0.269 | 0.359 | 5.4656E-07 |

| | | | | | | | |
|---|---|---|---|---|---|---|---|
| Fibroblast | Sulf11 | 5 | 5.8455E-10 | 0.73623072 | 0.658 | 0.621 | 1.1691E-06 |
| Fibroblast | LOC6900451 | 5 | 6.9184E-10 | 0.34445978 | 0.224 | 0.262 | 1.3837E-06 |
| Fibroblast | Pde10a1 | 5 | 8.002E-10 | 0.69988145 | 0.621 | 0.597 | 1.6004E-06 |
| Fibroblast | Rbm241 | 5 | 8.1979E-10 | 0.38050735 | 0.202 | 0.254 | 1.6396E-06 |
| Fibroblast | Cttnbp21 | 5 | 1.2144E-09 | 0.36434203 | 0.236 | 0.291 | 2.4288E-06 |
| Fibroblast | Pla2r11 | 5 | 1.512E-09 | 0.76884893 | 0.215 | 0.27 | 3.024E-06 |
| Fibroblast | Alpk32 | 5 | 1.5719E-09 | 0.31547942 | 0.214 | 0.283 | 3.1437E-06 |
| Fibroblast | Pclo1 | 5 | 1.573E-09 | 1.30143143 | 0.196 | 0.293 | 3.146E-06 |
| Fibroblast | Zbtb7c1 | 5 | 1.65E-09 | 0.54475853 | 0.266 | 0.342 | 3.3E-06 |
| Fibroblast | Kcng22 | 5 | 1.6535E-09 | 0.48020064 | 0.239 | 0.278 | 3.3071E-06 |
| Fibroblast | Medag1 | 5 | 1.8594E-09 | 0.80083055 | 0.641 | 0.625 | 3.7187E-06 |
| Fibroblast | Usp181 | 5 | 2.1722E-09 | 0.45339098 | 0.283 | 0.317 | 4.3444E-06 |
| Fibroblast | Rnf1502 | 5 | 2.705E-09 | 0.33364447 | 0.25 | 0.34 | 5.4099E-06 |
| Fibroblast | Stmn41 | 5 | 2.8001E-09 | 0.40620174 | 0.196 | 0.297 | 5.6003E-06 |
| Fibroblast | Rspo12 | 5 | 3.8932E-09 | 0.72298932 | 0.211 | 0.28 | 7.7864E-06 |
| Fibroblast | Slc37a22 | 5 | 4.2684E-09 | 0.32163609 | 0.261 | 0.319 | 8.5368E-06 |
| Fibroblast | Adgrg21 | 5 | 4.6427E-09 | 0.71680084 | 0.232 | 0.35 | 9.2854E-06 |
| Fibroblast | Pi161 | 5 | 4.8287E-09 | 0.99302424 | 0.505 | 0.454 | 9.6575E-06 |
| Fibroblast | Prelp | 5 | 5.2056E-09 | 0.26762107 | 0.247 | 0.3 | 1.0411E-05 |
| Fibroblast | Actg2 | 5 | 5.2057E-09 | 0.73157481 | 0.305 | 0.179 | 1.0411E-05 |
| Fibroblast | Pcdh191 | 5 | 5.8396E-09 | 0.30943523 | 0.299 | 0.383 | 1.1679E-05 |
| Fibroblast | Gsn2 | 5 | 6.2436E-09 | 0.56534702 | 0.639 | 0.577 | 1.2487E-05 |
| Fibroblast | Rab7b1 | 5 | 6.4793E-09 | 0.46627908 | 0.281 | 0.409 | 1.2959E-05 |
| Fibroblast | Col27a11 | 5 | 7.9725E-09 | 0.40259057 | 0.33 | 0.441 | 1.5945E-05 |
| Fibroblast | Gli22 | 5 | 8.1407E-09 | 0.66348073 | 0.646 | 0.591 | 1.6281E-05 |
| Fibroblast | Igfbp61 | 5 | 9.4578E-09 | 0.46788797 | 0.234 | 0.304 | 1.8916E-05 |
| Fibroblast | Nr4a12 | 5 | 1.3159E-08 | 0.44306768 | 0.236 | 0.289 | 2.6318E-05 |
| Fibroblast | Rap1gap2 | 5 | 1.3448E-08 | 0.50375171 | 0.217 | 0.265 | 2.6896E-05 |
| Fibroblast | Zbtb162 | 5 | 1.7097E-08 | 0.83263379 | 0.584 | 0.431 | 3.4195E-05 |
| Fibroblast | Vcan1 | 5 | 3.0583E-08 | 0.62227259 | 0.652 | 0.676 | 6.1166E-05 |
| Fibroblast | Ppl2 | 5 | 3.2862E-08 | 0.56403875 | 0.225 | 0.268 | 6.5725E-05 |

| | | | | | | | |
|---|---|---|---|---|---|---|---|
| Fibroblast | Uqcrq1 | 5 | 3.7452E-08 | 0.29653779 | 0.294 | 0.347 | 7.4904E-05 |
| Fibroblast | Hvcn11 | 5 | 3.834E-08 | 0.29402136 | 0.253 | 0.303 | 7.6679E-05 |
| Fibroblast | Lgals11 | 5 | 3.9527E-08 | 0.45199303 | 0.298 | 0.427 | 7.9055E-05 |
| Fibroblast | Lyn2 | 5 | 4.0276E-08 | 0.26869318 | 0.29 | 0.376 | 8.0552E-05 |
| Fibroblast | Colgalt22 | 5 | 4.6678E-08 | 0.86115359 | 0.533 | 0.515 | 9.3356E-05 |
| Fibroblast | Sh3bp21 | 5 | 5.2734E-08 | 0.67952236 | 0.255 | 0.307 | 0.00010547 |
| Fibroblast | Tmem262 | 5 | 6.2424E-08 | 0.30085751 | 0.307 | 0.362 | 0.00012485 |
| Fibroblast | Tex222 | 5 | 6.8342E-08 | 0.38862515 | 0.225 | 0.252 | 0.00013668 |
| Fibroblast | Herc62 | 5 | 8.0615E-08 | 0.43472961 | 0.235 | 0.306 | 0.00016123 |
| Fibroblast | Apobec11 | 5 | 8.1213E-08 | 0.34982892 | 0.178 | 0.251 | 0.00016243 |
| Fibroblast | Creb51 | 5 | 8.5994E-08 | 0.30260675 | 0.794 | 0.775 | 0.00017199 |
| Fibroblast | Ckb2 | 5 | 9.9973E-08 | 0.37258677 | 0.281 | 0.31 | 0.00019995 |
| Fibroblast | Aatk2 | 5 | 1.2447E-07 | 0.47294053 | 0.214 | 0.265 | 0.00024894 |
| Fibroblast | Ppara1 | 5 | 1.9421E-07 | 0.38376102 | 0.269 | 0.285 | 0.00038842 |
| Fibroblast | Sox61 | 5 | 2.006E-07 | 0.70051268 | 0.559 | 0.486 | 0.0004012 |
| Fibroblast | Ttyh12 | 5 | 2.1897E-07 | 0.70291188 | 0.27 | 0.147 | 0.00043795 |
| Fibroblast | Cdh41 | 5 | 2.4878E-07 | 1.06699617 | 0.269 | 0.179 | 0.00049756 |
| Fibroblast | Pkia2 | 5 | 4.2985E-07 | 0.40044756 | 0.271 | 0.278 | 0.00085969 |
| Fibroblast | Spp1 | 5 | 6.4634E-07 | 0.29699428 | 0.251 | 0.258 | 0.00129268 |
| Fibroblast | Irf82 | 5 | 7.3774E-07 | 0.27934644 | 0.273 | 0.29 | 0.00147549 |
| Fibroblast | Mbp2 | 5 | 1.0013E-06 | 0.69522121 | 0.27 | 0.329 | 0.00200253 |
| Fibroblast | Efemp13 | 5 | 1.1856E-06 | 0.36629607 | 0.29 | 0.325 | 0.00237114 |
| Fibroblast | Aldh1a12 | 5 | 1.1968E-06 | 0.44414369 | 0.314 | 0.381 | 0.00239353 |
| Fibroblast | Eef1a21 | 5 | 1.3788E-06 | 0.26595294 | 0.3 | 0.328 | 0.00275766 |
| Fibroblast | Kcnk131 | 5 | 1.6239E-06 | 1.80941031 | 0.221 | 0.264 | 0.00324782 |
| Fibroblast | PCOLCE21 | 5 | 2.0713E-06 | 0.43679394 | 0.304 | 0.362 | 0.0041426 |
| Fibroblast | Il341 | 5 | 2.2375E-06 | 0.52174313 | 0.276 | 0.281 | 0.00447496 |
| Fibroblast | Jph1 | 5 | 2.426E-06 | 0.77575008 | 0.273 | 0.288 | 0.00485205 |
| Fibroblast | Timp21 | 5 | 2.5292E-06 | 0.42044378 | 0.776 | 0.793 | 0.00505834 |
| Fibroblast | Scn5a1 | 5 | 3.1986E-06 | 0.35124656 | 0.256 | 0.238 | 0.00639726 |
| Fibroblast | Ezr1 | 5 | 3.5529E-06 | 0.59338487 | 0.279 | 0.288 | 0.00710585 |

| | | | | | | | |
|---|---|---|---|---|---|---|---|
| Fibroblast | Enox12 | 5 | 4.3899E-06 | 0.29128433 | 0.351 | 0.428 | 0.00877985 |
| Fibroblast | Rnls3 | 5 | 4.7955E-06 | 0.26510161 | 0.328 | 0.426 | 0.00959109 |
| Fibroblast | Tmeff22 | 5 | 1.0003E-05 | 0.47556966 | 0.711 | 0.688 | 0.02000589 |
| Fibroblast | Ivns1abp2 | 5 | 1.1812E-05 | 0.31491254 | 0.302 | 0.356 | 0.02362372 |
| Fibroblast | Lgmn | 5 | 1.4752E-05 | 0.27941225 | 0.323 | 0.327 | 0.02950341 |
| Fibroblast | Col14a11 | 5 | 2.0525E-05 | 0.36069937 | 0.73 | 0.722 | 0.04105085 |
| Fibroblast | Aoah1 | 5 | 2.1341E-05 | 0.91745483 | 0.253 | 0.269 | 0.0426826 |
| Fibroblast | Disp2 | 6 | 1.22E-217 | 2.65884729 | 0.359 | 0.047 | 2.44E-214 |
| Fibroblast | Troap | 6 | 1.612E-123 | 1.65910785 | 0.37 | 0.041 | 3.224E-120 |
| Fibroblast | Ltbp21 | 6 | 1.428E-122 | 1.70543728 | 0.829 | 0.447 | 2.857E-119 |
| Fibroblast | Col8a1 | 6 | 2.202E-117 | 1.30005559 | 0.98 | 0.864 | 4.404E-114 |
| Fibroblast | AABR070121 | 6 | 4.912E-110 | 2.68574612 | 0.259 | 0.023 | 9.825E-107 |
| Fibroblast | Cilp2 | 6 | 7.829E-110 | 2.87339651 | 0.268 | 0.058 | 1.566E-106 |
| Fibroblast | Olr1 | 6 | 1.0709E-82 | 2.08732616 | 0.316 | 0.083 | 2.1418E-79 |
| Fibroblast | Piezo21 | 6 | 5.1685E-80 | 1.96532168 | 0.712 | 0.416 | 1.0337E-76 |
| Fibroblast | Parpbp | 6 | 1.0087E-65 | 2.53420985 | 0.3 | 0.066 | 2.0174E-62 |
| Fibroblast | Podnl1 | 6 | 6.7608E-65 | 1.47932338 | 0.329 | 0.096 | 1.3522E-61 |
| Fibroblast | Runx1 | 6 | 2.3259E-64 | 1.50349338 | 0.727 | 0.506 | 4.6519E-61 |
| Fibroblast | Car10 | 6 | 3.4889E-63 | 1.31165602 | 0.385 | 0.144 | 6.9778E-60 |
| Fibroblast | Ppfia2 | 6 | 7.0383E-63 | 2.14071211 | 0.305 | 0.119 | 1.4077E-59 |
| Fibroblast | Fshr1 | 6 | 7.2057E-63 | 0.60559837 | 0.354 | 0.125 | 1.4411E-59 |
| Fibroblast | Tnfaip8l31 | 6 | 1.8898E-59 | 2.47747002 | 0.562 | 0.285 | 3.7796E-56 |
| Fibroblast | Tgfa | 6 | 4.439E-58 | 2.03520539 | 0.356 | 0.134 | 8.8781E-55 |
| Fibroblast | AABR070252 | 6 | 1.6538E-55 | 0.97991714 | 0.854 | 0.659 | 3.3075E-52 |
| Fibroblast | Ttc291 | 6 | 6.5899E-51 | 1.38540213 | 0.34 | 0.112 | 1.318E-47 |
| Fibroblast | Lancl3 | 6 | 1.4571E-50 | 1.66057451 | 0.424 | 0.207 | 2.9142E-47 |
| Fibroblast | AABR070655 | 6 | 5.2208E-50 | 1.38857943 | 0.346 | 0.189 | 1.0442E-46 |
| Fibroblast | Ddah11 | 6 | 1.4991E-49 | 1.73224477 | 0.623 | 0.376 | 2.9982E-46 |
| Fibroblast | Postn1 | 6 | 7.4115E-49 | 1.06261734 | 0.875 | 0.767 | 1.4823E-45 |
| Fibroblast | B4galnt3 | 6 | 1.0585E-47 | 3.41745319 | 0.409 | 0.192 | 2.1171E-44 |
| Fibroblast | Map1b2 | 6 | 2.9841E-47 | 0.65607998 | 0.917 | 0.788 | 5.9682E-44 |

| | | | | | | | |
|---|---|---|---|---|---|---|---|
| Fibroblast | Lef1 | 6 | 2.5886E-45 | 1.08823398 | 0.3 | 0.11 | 5.1772E-42 |
| Fibroblast | Efcab11 | 6 | 2.8498E-42 | 1.9413489 | 0.271 | 0.063 | 5.6996E-39 |
| Fibroblast | Meox1 | 6 | 4.1673E-39 | 2.26754829 | 0.332 | 0.139 | 8.3347E-36 |
| Fibroblast | Srgap11 | 6 | 6.1494E-39 | 0.7928864 | 0.797 | 0.64 | 1.2299E-35 |
| Fibroblast | Ca3 | 6 | 2.2097E-36 | 0.60607875 | 0.349 | 0.133 | 4.4195E-33 |
| Fibroblast | Slc20a1 | 6 | 5.4375E-36 | 1.90656299 | 0.568 | 0.397 | 1.0875E-32 |
| Fibroblast | Ctnnd2 | 6 | 5.6126E-36 | 0.78564535 | 0.393 | 0.138 | 1.1225E-32 |
| Fibroblast | Ptprv | 6 | 6.4551E-36 | 1.9911134 | 0.515 | 0.316 | 1.291E-32 |
| Fibroblast | Adamts17 | 6 | 1.1794E-35 | 0.90984998 | 0.724 | 0.533 | 2.3588E-32 |
| Fibroblast | Trem2 | 6 | 5.2617E-35 | 0.28581038 | 0.395 | 0.161 | 1.0523E-31 |
| Fibroblast | Plxna42 | 6 | 9.3851E-35 | 1.02680552 | 0.625 | 0.378 | 1.877E-31 |
| Fibroblast | Fn12 | 6 | 1.1971E-34 | 0.76127091 | 0.822 | 0.612 | 2.3942E-31 |
| Fibroblast | Klhl291 | 6 | 2.1353E-33 | 1.17374836 | 0.64 | 0.427 | 4.2706E-30 |
| Fibroblast | Vcan2 | 6 | 1.3033E-31 | 0.78011712 | 0.804 | 0.662 | 2.6065E-28 |
| Fibroblast | Pmepa11 | 6 | 1.8268E-31 | 0.80484076 | 0.733 | 0.56 | 3.6537E-28 |
| Fibroblast | Fstl11 | 6 | 3.6794E-31 | 0.46238271 | 0.929 | 0.811 | 7.3588E-28 |
| Fibroblast | Nlgn1 | 6 | 5.2634E-31 | 2.43567749 | 0.343 | 0.09 | 1.0527E-27 |
| Fibroblast | Unc5b | 6 | 1.8242E-30 | 0.96916958 | 0.673 | 0.454 | 3.6484E-27 |
| Fibroblast | Uchl1 | 6 | 3.1553E-30 | 1.40701956 | 0.286 | 0.127 | 6.3105E-27 |
| Fibroblast | Fam167a1 | 6 | 5.0602E-30 | 1.51879895 | 0.467 | 0.278 | 1.012E-26 |
| Fibroblast | Chst9 | 6 | 3.3078E-29 | 0.35430164 | 0.261 | 0.106 | 6.6156E-26 |
| Fibroblast | AC118957.1 | 6 | 7.389E-29 | 2.31395206 | 0.35 | 0.241 | 1.4778E-25 |
| Fibroblast | Boc2 | 6 | 1.7962E-27 | 0.55068688 | 0.859 | 0.751 | 3.5924E-24 |
| Fibroblast | Itga82 | 6 | 3.664E-27 | 0.76147354 | 0.72 | 0.59 | 7.3281E-24 |
| Fibroblast | Tmem63c1 | 6 | 2.2125E-26 | 1.23616261 | 0.345 | 0.228 | 4.4251E-23 |
| Fibroblast | Il1rapl1 | 6 | 2.6052E-26 | 2.02036607 | 0.499 | 0.244 | 5.2103E-23 |
| Fibroblast | Ccn41 | 6 | 2.9274E-26 | 1.51440651 | 0.458 | 0.238 | 5.8547E-23 |
| Fibroblast | Pdzrn31 | 6 | 1.2439E-25 | 0.4490348 | 0.956 | 0.915 | 2.4877E-22 |
| Fibroblast | Col5a2 | 6 | 1.7263E-25 | 0.55703 | 0.893 | 0.815 | 3.4526E-22 |
| Fibroblast | Dpysl31 | 6 | 2.5074E-23 | 0.59881193 | 0.858 | 0.736 | 5.0147E-20 |
| Fibroblast | Cacna1g | 6 | 3.9253E-23 | 1.18163923 | 0.543 | 0.4 | 7.8506E-20 |

| | | | | | | | |
|---|---|---|---|---|---|---|---|
| Fibroblast | Prrx12 | 6 | 1.4049E-22 | 0.6396553 | 0.817 | 0.675 | 2.8097E-19 |
| Fibroblast | Ncam11 | 6 | 1.8373E-22 | 0.94345812 | 0.645 | 0.492 | 3.6745E-19 |
| Fibroblast | Slc7a21 | 6 | 1.5661E-21 | 0.88593328 | 0.602 | 0.372 | 3.1322E-18 |
| Fibroblast | Cenpe | 6 | 1.0741E-20 | 0.32278044 | 0.34 | 0.148 | 2.1482E-17 |
| Fibroblast | Adamts62 | 6 | 1.4127E-20 | 0.58633315 | 0.751 | 0.626 | 2.8255E-17 |
| Fibroblast | Pde7a2 | 6 | 2.6741E-20 | 0.69793484 | 0.728 | 0.587 | 5.3482E-17 |
| Fibroblast | Pde7b | 6 | 3.1769E-20 | 0.74584379 | 0.681 | 0.583 | 6.3538E-17 |
| Fibroblast | Dkk32 | 6 | 3.2854E-20 | 0.72265405 | 0.657 | 0.553 | 6.5707E-17 |
| Fibroblast | Bend6 | 6 | 5.1563E-20 | 1.30495723 | 0.486 | 0.293 | 1.0313E-16 |
| Fibroblast | Hmcn11 | 6 | 5.5699E-20 | 0.37592737 | 0.696 | 0.487 | 1.114E-16 |
| Fibroblast | Ncald1 | 6 | 7.7635E-20 | 0.97927286 | 0.564 | 0.381 | 1.5527E-16 |
| Fibroblast | AABR070339 | 6 | 1.0332E-19 | 2.02942385 | 0.471 | 0.326 | 2.0665E-16 |
| Fibroblast | Fhod31 | 6 | 1.2722E-19 | 0.94845985 | 0.622 | 0.423 | 2.5443E-16 |
| Fibroblast | Cilp1 | 6 | 2.0707E-19 | 1.97812467 | 0.467 | 0.339 | 4.1413E-16 |
| Fibroblast | Galnt162 | 6 | 2.664E-19 | 0.48115835 | 0.758 | 0.612 | 5.3281E-16 |
| Fibroblast | C1qtnf32 | 6 | 8.3401E-19 | 2.18980042 | 0.406 | 0.228 | 1.668E-15 |
| Fibroblast | Actn11 | 6 | 8.4248E-19 | 0.53615368 | 0.7 | 0.539 | 1.685E-15 |
| Fibroblast | Inhba2 | 6 | 1.1457E-18 | 1.03350265 | 0.451 | 0.244 | 2.2914E-15 |
| Fibroblast | Matn2 | 6 | 1.4669E-18 | 0.8169992 | 0.628 | 0.506 | 2.9338E-15 |
| Fibroblast | Rai142 | 6 | 1.9921E-18 | 0.71302042 | 0.64 | 0.473 | 3.9842E-15 |
| Fibroblast | Chsy32 | 6 | 2.6602E-18 | 0.79171825 | 0.691 | 0.53 | 5.3203E-15 |
| Fibroblast | Thbs4 | 6 | 2.7951E-18 | 2.37146312 | 0.379 | 0.275 | 5.5901E-15 |
| Fibroblast | Cdkn2b | 6 | 3.0444E-18 | 0.85239225 | 0.307 | 0.161 | 6.0887E-15 |
| Fibroblast | Tgfb32 | 6 | 4.5201E-18 | 0.466381 | 0.691 | 0.499 | 9.0402E-15 |
| Fibroblast | Specc11 | 6 | 6.7606E-18 | 0.54918668 | 0.673 | 0.523 | 1.3521E-14 |
| Fibroblast | Efna51 | 6 | 6.8983E-18 | 0.66394333 | 0.503 | 0.368 | 1.3797E-14 |
| Fibroblast | Prima11 | 6 | 8.4539E-18 | 1.36869397 | 0.442 | 0.257 | 1.6908E-14 |
| Fibroblast | Col11a1 | 6 | 2.122E-17 | 1.24223786 | 0.496 | 0.374 | 4.244E-14 |
| Fibroblast | Tnc | 6 | 4.4341E-17 | 2.46227479 | 0.424 | 0.284 | 8.8681E-14 |
| Fibroblast | P4ha31 | 6 | 6.42E-17 | 0.51993751 | 0.35 | 0.214 | 1.284E-13 |
| Fibroblast | Rec1141 | 6 | 6.6E-17 | 0.5158197 | 0.278 | 0.166 | 1.32E-13 |

| | | | | | | | |
|---|---|---|---|---|---|---|---|
| Fibroblast | Prg4 | 6 | 7.0966E-17 | 1.84310475 | 0.39 | 0.214 | 1.4193E-13 |
| Fibroblast | Rspo2 | 6 | 8.1365E-17 | 1.4037501 | 0.296 | 0.1 | 1.6273E-13 |
| Fibroblast | Mmp2 | 6 | 1.7061E-16 | 0.57266495 | 0.647 | 0.451 | 3.4122E-13 |
| Fibroblast | AABR070440 | 6 | 1.7644E-16 | 0.9332829 | 0.537 | 0.377 | 3.5289E-13 |
| Fibroblast | Myl9 | 6 | 2.0676E-16 | 0.34432412 | 0.325 | 0.164 | 4.1353E-13 |
| Fibroblast | AABR070215 | 6 | 2.2156E-16 | 0.89603338 | 0.443 | 0.346 | 4.4313E-13 |
| Fibroblast | Mest1 | 6 | 2.2441E-16 | 0.95595359 | 0.321 | 0.172 | 4.4882E-13 |
| Fibroblast | Nrp2 | 6 | 2.2783E-16 | 0.5793936 | 0.574 | 0.371 | 4.5567E-13 |
| Fibroblast | Gria31 | 6 | 2.698E-16 | 0.53004415 | 0.764 | 0.601 | 5.3961E-13 |
| Fibroblast | Col22a11 | 6 | 2.8451E-16 | 1.218154 | 0.487 | 0.343 | 5.6903E-13 |
| Fibroblast | Bnc22 | 6 | 4.4714E-16 | 0.39798889 | 0.831 | 0.74 | 8.9428E-13 |
| Fibroblast | Fap1 | 6 | 6.3174E-16 | 0.54892642 | 0.736 | 0.628 | 1.2635E-12 |
| Fibroblast | Nr4a31 | 6 | 6.82E-16 | 0.67560096 | 0.387 | 0.287 | 1.364E-12 |
| Fibroblast | Cdkn1a1 | 6 | 1.0181E-15 | 1.70503533 | 0.461 | 0.334 | 2.0362E-12 |
| Fibroblast | Pam2 | 6 | 1.6482E-15 | 0.48530384 | 0.777 | 0.643 | 3.2964E-12 |
| Fibroblast | Col1a11 | 6 | 1.8384E-15 | 0.29023701 | 0.943 | 0.87 | 3.6768E-12 |
| Fibroblast | Bgn | 6 | 2.9504E-15 | 0.49103195 | 0.631 | 0.451 | 5.9008E-12 |
| Fibroblast | Myh101 | 6 | 3.9242E-15 | 0.42850785 | 0.732 | 0.531 | 7.8485E-12 |
| Fibroblast | Agmo1 | 6 | 5.5553E-15 | 1.59559681 | 0.418 | 0.238 | 1.1111E-11 |
| Fibroblast | Ednra1 | 6 | 6.2833E-15 | 0.59454076 | 0.641 | 0.511 | 1.2567E-11 |
| Fibroblast | Nav3 | 6 | 4.8454E-14 | 0.75815195 | 0.479 | 0.341 | 9.6908E-11 |
| Fibroblast | Rab15 | 6 | 6.2433E-14 | 1.3173 | 0.263 | 0.166 | 1.2487E-10 |
| Fibroblast | Robo2 | 6 | 6.9629E-14 | 1.10069698 | 0.469 | 0.335 | 1.3926E-10 |
| Fibroblast | Enah2 | 6 | 8.1312E-14 | 0.73637387 | 0.607 | 0.456 | 1.6262E-10 |
| Fibroblast | Filip1l1 | 6 | 9.7439E-14 | 0.47019021 | 0.749 | 0.622 | 1.9488E-10 |
| Fibroblast | Plod21 | 6 | 1.4324E-13 | 0.44634263 | 0.853 | 0.746 | 2.8647E-10 |
| Fibroblast | Cd441 | 6 | 1.6766E-13 | 0.8090231 | 0.525 | 0.384 | 3.3531E-10 |
| Fibroblast | Ccn21 | 6 | 1.9227E-13 | 0.41638113 | 0.637 | 0.479 | 3.8455E-10 |
| Fibroblast | Creb52 | 6 | 3.55E-13 | 0.40243073 | 0.856 | 0.769 | 7.0999E-10 |
| Fibroblast | Il17rd1 | 6 | 5.2191E-13 | 0.47296449 | 0.569 | 0.409 | 1.0438E-09 |
| Fibroblast | Pcdh192 | 6 | 5.6174E-13 | 0.63813086 | 0.523 | 0.363 | 1.1235E-09 |

| | | | | | | | |
|---|---|---|---|---|---|---|---|
| Fibroblast | Pls31 | 6 | 8.7579E-13 | 0.5215402 | 0.798 | 0.67 | 1.7516E-09 |
| Fibroblast | Dgkg | 6 | 8.8785E-13 | 0.4985735 | 0.287 | 0.135 | 1.7757E-09 |
| Fibroblast | Ptpn13 | 6 | 8.9433E-13 | 0.75014232 | 0.548 | 0.417 | 1.7887E-09 |
| Fibroblast | Cpeb11 | 6 | 1.0288E-12 | 0.59372031 | 0.572 | 0.398 | 2.0576E-09 |
| Fibroblast | Aff32 | 6 | 1.2452E-12 | 0.39867906 | 0.785 | 0.685 | 2.4903E-09 |
| Fibroblast | Cacna1d1 | 6 | 1.7261E-12 | 0.44649166 | 0.365 | 0.159 | 3.4523E-09 |
| Fibroblast | Rnf431 | 6 | 2.8383E-12 | 0.81634147 | 0.296 | 0.17 | 5.6765E-09 |
| Fibroblast | Fgf91 | 6 | 5.2666E-12 | 0.86895682 | 0.372 | 0.245 | 1.0533E-08 |
| Fibroblast | Susd1 | 6 | 5.8523E-12 | 0.5061562 | 0.366 | 0.209 | 1.1705E-08 |
| Fibroblast | Fos | 6 | 1.2733E-11 | 1.26513303 | 0.382 | 0.262 | 2.5466E-08 |
| Fibroblast | Fbln21 | 6 | 1.579E-11 | 0.60415472 | 0.61 | 0.497 | 3.1581E-08 |
| Fibroblast | Antxr11 | 6 | 1.6444E-11 | 0.32599162 | 0.864 | 0.785 | 3.2887E-08 |
| Fibroblast | Pappa22 | 6 | 2.3535E-11 | 0.91650885 | 0.292 | 0.239 | 4.7071E-08 |
| Fibroblast | Tmem116 | 6 | 2.9316E-11 | 0.3516489 | 0.343 | 0.165 | 5.8632E-08 |
| Fibroblast | Trabd2b1 | 6 | 4.386E-11 | 1.24976691 | 0.392 | 0.226 | 8.772E-08 |
| Fibroblast | AC130940.11 | 6 | 5.3111E-11 | 0.41860194 | 0.278 | 0.174 | 1.0622E-07 |
| Fibroblast | Kcnma11 | 6 | 5.8154E-11 | 1.54008265 | 0.384 | 0.285 | 1.1631E-07 |
| Fibroblast | Loxl12 | 6 | 6.233E-11 | 0.40023595 | 0.793 | 0.681 | 1.2466E-07 |
| Fibroblast | Runx2 | 6 | 6.5268E-11 | 0.72057604 | 0.562 | 0.506 | 1.3054E-07 |
| Fibroblast | Enpp11 | 6 | 6.5628E-11 | 0.82056557 | 0.56 | 0.539 | 1.3126E-07 |
| Fibroblast | Dync1i11 | 6 | 1.0203E-10 | 0.94261584 | 0.324 | 0.21 | 2.0407E-07 |
| Fibroblast | Hpgd1 | 6 | 4.8369E-10 | 0.45425534 | 0.343 | 0.265 | 9.6739E-07 |
| Fibroblast | Nrg21 | 6 | 6.6127E-10 | 0.41778386 | 0.366 | 0.199 | 1.3225E-06 |
| Fibroblast | Cthrc11 | 6 | 8.1023E-10 | 0.53547053 | 0.491 | 0.315 | 1.6205E-06 |
| Fibroblast | Spp11 | 6 | 9.1421E-10 | 0.89395329 | 0.389 | 0.246 | 1.8284E-06 |
| Fibroblast | Pdlim31 | 6 | 9.7622E-10 | 0.5334415 | 0.579 | 0.479 | 1.9524E-06 |
| Fibroblast | AABR070546 | 6 | 1.5454E-09 | 0.38188614 | 0.693 | 0.56 | 3.0908E-06 |
| Fibroblast | Homer11 | 6 | 1.6312E-09 | 0.98221345 | 0.46 | 0.387 | 3.2623E-06 |
| Fibroblast | Ror2 | 6 | 1.9079E-09 | 0.45657421 | 0.61 | 0.504 | 3.8157E-06 |
| Fibroblast | Col8a21 | 6 | 3.2656E-09 | 1.15969886 | 0.472 | 0.426 | 6.5313E-06 |
| Fibroblast | Igfbp62 | 6 | 3.3415E-09 | 0.41070521 | 0.445 | 0.285 | 6.6831E-06 |

| | | | | | | | |
|---|---|---|---|---|---|---|---|
| Fibroblast | Cacna1e3 | 6 | 6.3868E-09 | 0.2656808 | 0.531 | 0.437 | 1.2774E-05 |
| Fibroblast | Lgi4 | 6 | 7.153E-09 | 1.16052657 | 0.414 | 0.317 | 1.4306E-05 |
| Fibroblast | Prkcz2 | 6 | 8.4463E-09 | 0.49834147 | 0.455 | 0.336 | 1.6893E-05 |
| Fibroblast | Rbp12 | 6 | 1.067E-08 | 0.5772634 | 0.448 | 0.336 | 2.1339E-05 |
| Fibroblast | Cers61 | 6 | 1.4243E-08 | 0.38242065 | 0.623 | 0.515 | 2.8486E-05 |
| Fibroblast | Srgap32 | 6 | 1.7335E-08 | 0.41963117 | 0.686 | 0.613 | 3.467E-05 |
| Fibroblast | Plaur2 | 6 | 2.2106E-08 | 0.51486875 | 0.55 | 0.442 | 4.4212E-05 |
| Fibroblast | AABR070275 | 6 | 2.8962E-08 | 0.64920079 | 0.431 | 0.295 | 5.7923E-05 |
| Fibroblast | Rab7b2 | 6 | 4.4482E-08 | 0.41391616 | 0.51 | 0.389 | 8.8965E-05 |
| Fibroblast | Col24a11 | 6 | 4.7442E-08 | 0.66652589 | 0.486 | 0.365 | 9.4884E-05 |
| Fibroblast | Pank1 | 6 | 4.7675E-08 | 0.56152962 | 0.427 | 0.311 | 9.535E-05 |
| Fibroblast | Cnnm2 | 6 | 4.7776E-08 | 0.28025017 | 0.484 | 0.355 | 9.5552E-05 |
| Fibroblast | Med12l1 | 6 | 6.4163E-08 | 0.47247931 | 0.437 | 0.29 | 0.00012833 |
| Fibroblast | Ano1 | 6 | 7.5837E-08 | 0.76462742 | 0.275 | 0.146 | 0.00015167 |
| Fibroblast | AABR070525 | 6 | 1.2309E-07 | 0.3210242 | 0.399 | 0.264 | 0.00024618 |
| Fibroblast | Mrc21 | 6 | 1.4678E-07 | 0.25562864 | 0.825 | 0.723 | 0.00029356 |
| Fibroblast | Ptgs21 | 6 | 1.7219E-07 | 1.4207113 | 0.379 | 0.322 | 0.00034438 |
| Fibroblast | Cemip2 | 6 | 1.8413E-07 | 0.4757823 | 0.563 | 0.494 | 0.00036825 |
| Fibroblast | Csrp21 | 6 | 1.872E-07 | 0.49846851 | 0.406 | 0.323 | 0.00037439 |
| Fibroblast | AABR070581 | 6 | 2.2743E-07 | 0.52143592 | 0.63 | 0.537 | 0.00045487 |
| Fibroblast | Sdc21 | 6 | 2.4402E-07 | 0.38435224 | 0.641 | 0.524 | 0.00048804 |
| Fibroblast | Anks1b | 6 | 2.9076E-07 | 0.49920599 | 0.344 | 0.223 | 0.00058152 |
| Fibroblast | Dgki1 | 6 | 3.2297E-07 | 0.66946279 | 0.413 | 0.287 | 0.00064594 |
| Fibroblast | Fmod | 6 | 3.3892E-07 | 0.76209249 | 0.252 | 0.165 | 0.00067785 |
| Fibroblast | Col16a1 | 6 | 3.7479E-07 | 0.73408268 | 0.479 | 0.427 | 0.00074959 |
| Fibroblast | Col27a12 | 6 | 4.6734E-07 | 1.11239796 | 0.462 | 0.429 | 0.00093467 |
| Fibroblast | Sptb2 | 6 | 5.2551E-07 | 0.8895159 | 0.411 | 0.282 | 0.00105103 |
| Fibroblast | Rgcc | 6 | 5.6838E-07 | 0.37155251 | 0.335 | 0.246 | 0.00113676 |
| Fibroblast | Olfml2b1 | 6 | 6.8806E-07 | 0.66202 | 0.474 | 0.406 | 0.00137613 |
| Fibroblast | Hmox11 | 6 | 7.1518E-07 | 0.28653892 | 0.271 | 0.177 | 0.00143035 |
| Fibroblast | Nusap11 | 6 | 1.0544E-06 | 0.49520655 | 0.282 | 0.2 | 0.00210882 |

| | | | | | | | |
|---|---|---|---|---|---|---|---|
| Fibroblast | Sparc | 6 | 1.1161E-06 | 0.25580517 | 0.845 | 0.732 | 0.00223227 |
| Fibroblast | Mgmt2 | 6 | 1.1491E-06 | 0.67843642 | 0.584 | 0.468 | 0.00229821 |
| Fibroblast | Loxl21 | 6 | 1.1695E-06 | 0.3167004 | 0.599 | 0.515 | 0.00233895 |
| Fibroblast | Epha3 | 6 | 1.2306E-06 | 0.26893174 | 0.368 | 0.275 | 0.00246116 |
| Fibroblast | Mpped23 | 6 | 1.3153E-06 | 0.40200731 | 0.587 | 0.504 | 0.00263063 |
| Fibroblast | Palm21 | 6 | 1.7006E-06 | 0.65701987 | 0.469 | 0.42 | 0.00340115 |
| Fibroblast | AC134204.11 | 6 | 3.114E-06 | 0.46585821 | 0.545 | 0.515 | 0.00622795 |
| Fibroblast | Cald1 | 6 | 3.4973E-06 | 0.28832076 | 0.761 | 0.689 | 0.00699462 |
| Fibroblast | Neurl1 | 6 | 3.9495E-06 | 0.96554419 | 0.297 | 0.23 | 0.00789894 |
| Fibroblast | Zbtb7c2 | 6 | 4.1447E-06 | 1.11262349 | 0.382 | 0.332 | 0.00828943 |
| Fibroblast | Klhl4 | 6 | 4.3243E-06 | 0.45954672 | 0.254 | 0.151 | 0.00864856 |
| Fibroblast | Ntng1 | 6 | 5.6319E-06 | 0.43771196 | 0.278 | 0.222 | 0.0112638 |
| Fibroblast | Tc2n2 | 6 | 6.297E-06 | 0.45376614 | 0.428 | 0.328 | 0.0125941 |
| Fibroblast | Aff2 | 6 | 9.2232E-06 | 0.58631331 | 0.463 | 0.414 | 0.01844632 |
| Fibroblast | Serpine11 | 6 | 1.1232E-05 | 1.21443994 | 0.401 | 0.344 | 0.02246425 |
| Fibroblast | Fat32 | 6 | 1.1588E-05 | 0.60578186 | 0.437 | 0.383 | 0.02317658 |
| Fibroblast | Ptpn32 | 6 | 1.2384E-05 | 0.47612649 | 0.45 | 0.391 | 0.0247679 |
| Fibroblast | Rspo13 | 6 | 1.3584E-05 | 0.25433326 | 0.397 | 0.264 | 0.02716714 |
| Fibroblast | Prdx5 | 6 | 1.439E-05 | 0.49707896 | 0.316 | 0.204 | 0.02877952 |
| Fibroblast | Nkd2 | 6 | 1.4782E-05 | 0.76609935 | 0.395 | 0.311 | 0.02956419 |
| Fibroblast | Gxylt22 | 6 | 2.3306E-05 | 0.26164238 | 0.772 | 0.717 | 0.04661131 |
| Fibroblast | Adam12 | 6 | 2.3585E-05 | 0.40185041 | 0.511 | 0.431 | 0.04717066 |
| Fibroblast | Col12a11 | 6 | 2.4161E-05 | 0.74324746 | 0.38 | 0.281 | 0.04832254 |
| Fibroblast | Sphkap | 6 | 2.4937E-05 | 0.60834221 | 0.345 | 0.267 | 0.04987421 |
| Fibroblast | Mlph | 7 | 0 | 2.64050664 | 0.307 | 0.009 | 0 |
| Fibroblast | Fermt1 | 7 | 1.792E-258 | 1.93821055 | 0.27 | 0.009 | 3.583E-255 |
| Fibroblast | Spon1 | 7 | 7.675E-186 | 2.48809134 | 0.898 | 0.514 | 1.535E-182 |
| Fibroblast | Slc4a5 | 7 | 1.046E-170 | 2.0399976 | 0.44 | 0.041 | 2.092E-167 |
| Fibroblast | Cela1 | 7 | 2.051E-155 | 3.76168812 | 0.371 | 0.034 | 4.102E-152 |
| Fibroblast | AABR070213 | 7 | 2.226E-135 | 2.06460271 | 0.279 | 0.043 | 4.451E-132 |
| Fibroblast | C71 | 7 | 1.246E-113 | 1.08234084 | 0.989 | 0.893 | 2.493E-110 |

| | | | | | | | |
|---|---|---|---|---|---|---|---|
| Fibroblast | Igf12 | 7 | 6.098E-111 | 1.80791858 | 0.892 | 0.599 | 1.22E-107 |
| Fibroblast | Shisal1 | 7 | 7.601E-110 | 0.93425713 | 0.295 | 0.049 | 1.52E-106 |
| Fibroblast | Nrg4 | 7 | 9.08E-109 | 1.11261534 | 0.255 | 0.023 | 1.816E-105 |
| Fibroblast | Edar2 | 7 | 4.73E-104 | 3.16872789 | 0.622 | 0.301 | 9.461E-101 |
| Fibroblast | Hspa1b | 7 | 5.512E-102 | 1.4248075 | 0.281 | 0.025 | 1.1024E-98 |
| Fibroblast | Dhrs9 | 7 | 2.357E-100 | 1.81937976 | 0.444 | 0.064 | 4.7143E-97 |
| Fibroblast | Adgre4 | 7 | 9.351E-100 | 1.21524518 | 0.321 | 0.049 | 1.8702E-96 |
| Fibroblast | Tmem178a1 | 7 | 2.089E-88 | 2.60606948 | 0.691 | 0.424 | 4.1779E-85 |
| Fibroblast | Col6a5 | 7 | 1.3683E-87 | 0.61043702 | 0.251 | 0.11 | 2.7367E-84 |
| Fibroblast | Grem2 | 7 | 8.5187E-87 | 3.39187999 | 0.563 | 0.188 | 1.7037E-83 |
| Fibroblast | Neil3 | 7 | 3.4985E-83 | 1.95516499 | 0.382 | 0.095 | 6.997E-80 |
| Fibroblast | Chl1 | 7 | 1.1345E-80 | 2.3854684 | 0.263 | 0.084 | 2.269E-77 |
| Fibroblast | Lifr1 | 7 | 6.827E-78 | 2.14402894 | 0.69 | 0.318 | 1.3654E-74 |
| Fibroblast | Gabrb2 | 7 | 1.3621E-77 | 0.99242386 | 0.381 | 0.054 | 2.7241E-74 |
| Fibroblast | Ccdc801 | 7 | 1.6675E-73 | 0.86295516 | 0.984 | 0.888 | 3.335E-70 |
| Fibroblast | Flrt22 | 7 | 2.9275E-70 | 1.47245397 | 0.779 | 0.495 | 5.8549E-67 |
| Fibroblast | Ano3 | 7 | 2.2582E-68 | 0.40298261 | 0.376 | 0.119 | 4.5164E-65 |
| Fibroblast | Sned11 | 7 | 1.2112E-66 | 2.58802474 | 0.632 | 0.322 | 2.4224E-63 |
| Fibroblast | Svep11 | 7 | 1.4587E-66 | 1.74388534 | 0.754 | 0.458 | 2.9173E-63 |
| Fibroblast | Adamts16 | 7 | 4.463E-65 | 3.27831066 | 0.536 | 0.269 | 8.926E-62 |
| Fibroblast | Postn2 | 7 | 4.8557E-64 | 1.01938483 | 0.94 | 0.764 | 9.7113E-61 |
| Fibroblast | Ptn | 7 | 1.538E-63 | 2.68025325 | 0.583 | 0.338 | 3.0761E-60 |
| Fibroblast | Oprm11 | 7 | 3.9071E-62 | 2.46324835 | 0.502 | 0.242 | 7.8142E-59 |
| Fibroblast | Tacr1 | 7 | 7.1579E-62 | 1.57079498 | 0.354 | 0.109 | 1.4316E-58 |
| Fibroblast | Glis32 | 7 | 2.0227E-59 | 0.85264402 | 0.944 | 0.758 | 4.0454E-56 |
| Fibroblast | Cd682 | 7 | 1.7914E-56 | 0.91346407 | 0.472 | 0.226 | 3.5829E-53 |
| Fibroblast | Cdkn2b1 | 7 | 2.7187E-56 | 0.87299786 | 0.433 | 0.155 | 5.4373E-53 |
| Fibroblast | Dlg22 | 7 | 3.1153E-56 | 0.88554403 | 0.937 | 0.739 | 6.2306E-53 |
| Fibroblast | Nox41 | 7 | 3.9273E-56 | 1.33040211 | 0.781 | 0.511 | 7.8546E-53 |
| Fibroblast | Fgf71 | 7 | 2.0314E-55 | 2.3008493 | 0.257 | 0.159 | 4.0628E-52 |
| Fibroblast | Mnda1 | 7 | 1.1586E-54 | 1.39159898 | 0.702 | 0.484 | 2.3173E-51 |

| | | | | | | | |
|---|---|---|---|---|---|---|---|
| Fibroblast | St8sia1 | 7 | 2.9643E-54 | 0.38180984 | 0.271 | 0.093 | 5.9286E-51 |
| Fibroblast | Rcan23 | 7 | 6.8685E-54 | 0.9791703 | 0.875 | 0.623 | 1.3737E-50 |
| Fibroblast | Ak51 | 7 | 7.7145E-54 | 1.46406322 | 0.553 | 0.277 | 1.5429E-50 |
| Fibroblast | Cd300lb | 7 | 2.5738E-53 | 1.69614227 | 0.266 | 0.04 | 5.1475E-50 |
| Fibroblast | Vwc2 | 7 | 7.263E-53 | 1.48925267 | 0.273 | 0.068 | 1.4526E-49 |
| Fibroblast | Adam121 | 7 | 1.4216E-52 | 1.62365973 | 0.696 | 0.419 | 2.8432E-49 |
| Fibroblast | Ccn51 | 7 | 1.9063E-52 | 1.09942032 | 0.702 | 0.408 | 3.8127E-49 |
| Fibroblast | Adamts152 | 7 | 5.0655E-52 | 1.18756641 | 0.702 | 0.399 | 1.0131E-48 |
| Fibroblast | S100a3 | 7 | 8.2193E-52 | 1.01552326 | 0.281 | 0.058 | 1.6439E-48 |
| Fibroblast | Vcam11 | 7 | 3.3324E-51 | 2.14648638 | 0.578 | 0.298 | 6.6648E-48 |
| Fibroblast | Npnt1 | 7 | 1.5463E-50 | 0.99023904 | 0.42 | 0.191 | 3.0926E-47 |
| Fibroblast | Comp | 7 | 1.61E-50 | 1.35820596 | 0.276 | 0.131 | 3.2201E-47 |
| Fibroblast | Lancl31 | 7 | 3.9094E-47 | 0.33549327 | 0.436 | 0.21 | 7.8188E-44 |
| Fibroblast | Ahr1 | 7 | 1.8812E-46 | 1.45936833 | 0.621 | 0.435 | 3.7625E-43 |
| Fibroblast | Vegfd1 | 7 | 3.3102E-46 | 1.30517895 | 0.668 | 0.407 | 6.6203E-43 |
| Fibroblast | Mettl241 | 7 | 2.9258E-45 | 1.0571739 | 0.44 | 0.21 | 5.8516E-42 |
| Fibroblast | Slit33 | 7 | 4.2195E-44 | 1.36669502 | 0.707 | 0.465 | 8.4389E-41 |
| Fibroblast | Tnn | 7 | 6.074E-44 | 0.98092902 | 0.368 | 0.188 | 1.2148E-40 |
| Fibroblast | Ephb21 | 7 | 2.3677E-43 | 1.51320641 | 0.608 | 0.389 | 4.7354E-40 |
| Fibroblast | Plac81 | 7 | 1.8467E-42 | 1.28348761 | 0.255 | 0.119 | 3.6933E-39 |
| Fibroblast | Met | 7 | 2.2646E-42 | 0.93506588 | 0.274 | 0.115 | 4.5293E-39 |
| Fibroblast | LOC1009114 | 7 | 6.0132E-42 | 2.79160833 | 0.476 | 0.29 | 1.2026E-38 |
| Fibroblast | Tshr1 | 7 | 7.9052E-42 | 3.20894898 | 0.494 | 0.32 | 1.581E-38 |
| Fibroblast | Runx11 | 7 | 9.9869E-41 | 0.9942922 | 0.716 | 0.51 | 1.9974E-37 |
| Fibroblast | Anpep1 | 7 | 1.3626E-40 | 1.00271034 | 0.661 | 0.413 | 2.7253E-37 |
| Fibroblast | Nxn | 7 | 3.011E-40 | 1.11639007 | 0.754 | 0.559 | 6.022E-37 |
| Fibroblast | Chrdl11 | 7 | 3.386E-40 | 0.9903803 | 0.448 | 0.193 | 6.7719E-37 |
| Fibroblast | Sema3e | 7 | 3.8333E-40 | 0.44169296 | 0.329 | 0.117 | 7.6666E-37 |
| Fibroblast | Trpc5 | 7 | 6.5814E-39 | 1.67583506 | 0.266 | 0.162 | 1.3163E-35 |
| Fibroblast | Cass4 | 7 | 1.4276E-38 | 0.64960659 | 0.313 | 0.049 | 2.8553E-35 |
| Fibroblast | Shroom31 | 7 | 2.5688E-38 | 1.62890151 | 0.564 | 0.297 | 5.1377E-35 |

| | | | | | | | |
|---|---|---|---|---|---|---|---|
| Fibroblast | Gpm6b | 7 | 2.7697E-38 | 0.85146239 | 0.76 | 0.515 | 5.5395E-35 |
| Fibroblast | Slc1a72 | 7 | 3.3679E-38 | 1.44395268 | 0.676 | 0.48 | 6.7357E-35 |
| Fibroblast | Angpt1 | 7 | 5.7873E-38 | 1.0735232 | 0.726 | 0.535 | 1.1575E-34 |
| Fibroblast | Svil2 | 7 | 1.6704E-37 | 0.77233975 | 0.848 | 0.631 | 3.3408E-34 |
| Fibroblast | Aox3 | 7 | 2.0269E-37 | 1.86115171 | 0.508 | 0.277 | 4.0539E-34 |
| Fibroblast | Agtr1a1 | 7 | 6.0128E-37 | 1.49751483 | 0.588 | 0.421 | 1.2026E-33 |
| Fibroblast | Igfbp4 | 7 | 7.3204E-37 | 1.54405582 | 0.564 | 0.429 | 1.4641E-33 |
| Fibroblast | Fbln11 | 7 | 3.5875E-36 | 0.81099817 | 0.804 | 0.603 | 7.175E-33 |
| Fibroblast | Ror21 | 7 | 6.5653E-36 | 0.95839665 | 0.715 | 0.499 | 1.3131E-32 |
| Fibroblast | Ednrb1 | 7 | 6.6799E-36 | 0.84171508 | 0.469 | 0.176 | 1.336E-32 |
| Fibroblast | Pappa11 | 7 | 7.2114E-36 | 1.03263497 | 0.677 | 0.414 | 1.4423E-32 |
| Fibroblast | Vipr22 | 7 | 8.974E-36 | 1.05574117 | 0.607 | 0.355 | 1.7948E-32 |
| Fibroblast | Srpx1 | 7 | 9.5699E-36 | 1.28225122 | 0.589 | 0.34 | 1.914E-32 |
| Fibroblast | Prrx13 | 7 | 9.903E-36 | 0.70753713 | 0.878 | 0.674 | 1.9806E-32 |
| Fibroblast | Efemp14 | 7 | 1.5674E-35 | 0.89892137 | 0.577 | 0.305 | 3.1348E-32 |
| Fibroblast | Pdgfd1 | 7 | 2.2195E-35 | 0.95595092 | 0.694 | 0.52 | 4.4389E-32 |
| Fibroblast | Il1r12 | 7 | 2.7041E-35 | 0.79713179 | 0.879 | 0.709 | 5.4081E-32 |
| Fibroblast | Bmper2 | 7 | 3.4125E-35 | 0.83351502 | 0.757 | 0.53 | 6.8251E-32 |
| Fibroblast | Plxdc22 | 7 | 5.4795E-35 | 0.53066954 | 0.955 | 0.81 | 1.0959E-31 |
| Fibroblast | Tmeff23 | 7 | 8.3291E-34 | 0.69315614 | 0.873 | 0.677 | 1.6658E-30 |
| Fibroblast | Gxylt23 | 7 | 9.8927E-34 | 0.72318976 | 0.871 | 0.711 | 1.9785E-30 |
| Fibroblast | Plcxd31 | 7 | 1.4935E-33 | 1.54079651 | 0.511 | 0.327 | 2.987E-30 |
| Fibroblast | Syndig11 | 7 | 4.4182E-33 | 1.36986746 | 0.592 | 0.435 | 8.8364E-30 |
| Fibroblast | Hcn12 | 7 | 4.4633E-33 | 0.9104968 | 0.492 | 0.247 | 8.9267E-30 |
| Fibroblast | Nr1h41 | 7 | 6.3756E-33 | 1.82708344 | 0.531 | 0.347 | 1.2751E-29 |
| Fibroblast | Ptpn221 | 7 | 3.4567E-32 | 1.3897295 | 0.339 | 0.133 | 6.9133E-29 |
| Fibroblast | Boc3 | 7 | 7.5449E-32 | 0.56381041 | 0.925 | 0.748 | 1.509E-28 |
| Fibroblast | Ptk2b2 | 7 | 2.3366E-31 | 0.9687894 | 0.633 | 0.439 | 4.6732E-28 |
| Fibroblast | Runx21 | 7 | 4.3776E-31 | 0.9297214 | 0.682 | 0.499 | 8.7552E-28 |
| Fibroblast | Cp | 7 | 4.4878E-30 | 1.20547253 | 0.539 | 0.291 | 8.9756E-27 |
| Fibroblast | Slit2 | 7 | 4.5646E-30 | 1.06049762 | 0.61 | 0.382 | 9.1293E-27 |

| | | | | | | | |
|---|---|---|---|---|---|---|---|
| Fibroblast | Il331 | 7 | 1.5864E-29 | 1.41933153 | 0.505 | 0.323 | 3.1728E-26 |
| Fibroblast | Nrxn11 | 7 | 2.0523E-29 | 0.9083562 | 0.766 | 0.555 | 4.1045E-26 |
| Fibroblast | Itga83 | 7 | 4.5641E-29 | 0.73978874 | 0.788 | 0.587 | 9.1281E-26 |
| Fibroblast | Abcc92 | 7 | 9.8346E-29 | 0.79829951 | 0.732 | 0.557 | 1.9669E-25 |
| Fibroblast | Tafa2 | 7 | 1.051E-28 | 1.34772985 | 0.343 | 0.126 | 2.102E-25 |
| Fibroblast | Irs1 | 7 | 1.2395E-28 | 0.7394535 | 0.818 | 0.633 | 2.479E-25 |
| Fibroblast | Sorbs11 | 7 | 1.8128E-28 | 0.71403592 | 0.829 | 0.648 | 3.6257E-25 |
| Fibroblast | AABR070252 | 7 | 3.3508E-28 | 0.58211514 | 0.851 | 0.663 | 6.7015E-25 |
| Fibroblast | Nrp21 | 7 | 3.6381E-28 | 1.04103456 | 0.583 | 0.374 | 7.2761E-25 |
| Fibroblast | Mx1 | 7 | 4.1775E-28 | 1.07087783 | 0.415 | 0.198 | 8.355E-25 |
| Fibroblast | Prss12 | 7 | 7.333E-28 | 0.4010764 | 0.348 | 0.139 | 1.4666E-24 |
| Fibroblast | Galnt173 | 7 | 1.9259E-27 | 0.85560419 | 0.809 | 0.659 | 3.8518E-24 |
| Fibroblast | Dpp41 | 7 | 4.3628E-27 | 0.35909225 | 0.439 | 0.218 | 8.7256E-24 |
| Fibroblast | Rspo21 | 7 | 3.7157E-26 | 0.58098096 | 0.295 | 0.103 | 7.4314E-23 |
| Fibroblast | Tm4sf19 | 7 | 4.1094E-26 | 1.2304038 | 0.332 | 0.142 | 8.2188E-23 |
| Fibroblast | Lbp | 7 | 6.8212E-26 | 1.53547895 | 0.508 | 0.305 | 1.3642E-22 |
| Fibroblast | Sctr1 | 7 | 1.4113E-25 | 1.02661891 | 0.368 | 0.143 | 2.8226E-22 |
| Fibroblast | Mest2 | 7 | 1.6507E-25 | 0.97925877 | 0.398 | 0.169 | 3.3014E-22 |
| Fibroblast | Kcnj81 | 7 | 1.7259E-25 | 0.87903581 | 0.442 | 0.231 | 3.4517E-22 |
| Fibroblast | Tmem63c2 | 7 | 1.8574E-25 | 0.49172198 | 0.418 | 0.225 | 3.7149E-22 |
| Fibroblast | Pak31 | 7 | 1.9997E-25 | 0.30052022 | 0.464 | 0.264 | 3.9995E-22 |
| Fibroblast | Cemip21 | 7 | 9.648E-25 | 0.6408851 | 0.701 | 0.485 | 1.9296E-21 |
| Fibroblast | Tnfaip61 | 7 | 1.8061E-24 | 0.78321393 | 0.566 | 0.378 | 3.6122E-21 |
| Fibroblast | Pfkfb11 | 7 | 5.6131E-24 | 0.4588964 | 0.368 | 0.214 | 1.1226E-20 |
| Fibroblast | Sgcg | 7 | 6.8144E-24 | 1.22375893 | 0.466 | 0.305 | 1.3629E-20 |
| Fibroblast | LOC1036899 | 7 | 9.019E-24 | 0.83344172 | 0.671 | 0.47 | 1.8038E-20 |
| Fibroblast | Cemip1 | 7 | 1.1762E-23 | 0.44361135 | 0.571 | 0.34 | 2.3524E-20 |
| Fibroblast | AABR070490 | 7 | 1.258E-23 | 1.10445881 | 0.527 | 0.404 | 2.516E-20 |
| Fibroblast | Fbn2 | 7 | 1.269E-23 | 1.64632333 | 0.509 | 0.379 | 2.538E-20 |
| Fibroblast | Cdh21 | 7 | 2.4828E-23 | 0.75428494 | 0.718 | 0.541 | 4.9655E-20 |
| Fibroblast | Slc8a12 | 7 | 3.0714E-23 | 0.43613381 | 0.951 | 0.883 | 6.1427E-20 |

| | | | | | | | |
|---|---|---|---|---|---|---|---|
| Fibroblast | Slc16a7 | 7 | 1.2212E-22 | 0.53374972 | 0.42 | 0.205 | 2.4423E-19 |
| Fibroblast | Fam227b1 | 7 | 1.24E-22 | 0.78350353 | 0.516 | 0.3 | 2.4799E-19 |
| Fibroblast | Acer22 | 7 | 1.3467E-22 | 0.85172448 | 0.481 | 0.256 | 2.6933E-19 |
| Fibroblast | Alk | 7 | 2.099E-22 | 1.6445662 | 0.277 | 0.18 | 4.1979E-19 |
| Fibroblast | Pamr12 | 7 | 2.5674E-22 | 0.78977356 | 0.513 | 0.352 | 5.1347E-19 |
| Fibroblast | Prdm61 | 7 | 3.5289E-22 | 0.71645069 | 0.621 | 0.425 | 7.0578E-19 |
| Fibroblast | Col6a62 | 7 | 5.6162E-22 | 0.78114395 | 0.848 | 0.729 | 1.1232E-18 |
| Fibroblast | Pcdha132 | 7 | 7.2887E-22 | 0.58594609 | 0.803 | 0.617 | 1.4577E-18 |
| Fibroblast | Slfn41 | 7 | 1.0203E-21 | 0.77538555 | 0.505 | 0.29 | 2.0406E-18 |
| Fibroblast | Ctsz | 7 | 1.1332E-21 | 0.41571818 | 0.251 | 0.129 | 2.2663E-18 |
| Fibroblast | Lrrc17 | 7 | 1.3875E-21 | 0.742291 | 0.641 | 0.426 | 2.7751E-18 |
| Fibroblast | Ncaph | 7 | 1.9938E-21 | 0.46359867 | 0.301 | 0.19 | 3.9875E-18 |
| Fibroblast | Slc10a6 | 7 | 2.8135E-21 | 0.95784826 | 0.274 | 0.19 | 5.6271E-18 |
| Fibroblast | Gda2 | 7 | 3.0346E-21 | 0.54524644 | 0.881 | 0.751 | 6.0691E-18 |
| Fibroblast | Igfbp31 | 7 | 5.1531E-21 | 1.79607897 | 0.428 | 0.275 | 1.0306E-17 |
| Fibroblast | Asap31 | 7 | 8.9609E-21 | 0.60663135 | 0.821 | 0.668 | 1.7922E-17 |
| Fibroblast | Lhfpl21 | 7 | 1.2669E-20 | 0.71368668 | 0.627 | 0.442 | 2.5338E-17 |
| Fibroblast | Antxr12 | 7 | 1.394E-20 | 0.47431489 | 0.912 | 0.783 | 2.7879E-17 |
| Fibroblast | Ano5 | 7 | 1.6421E-20 | 1.26082034 | 0.26 | 0.142 | 3.2841E-17 |
| Fibroblast | Pcsk52 | 7 | 1.6705E-20 | 0.46323475 | 0.757 | 0.59 | 3.3411E-17 |
| Fibroblast | Esr12 | 7 | 2.5725E-20 | 0.33815425 | 0.508 | 0.288 | 5.145E-17 |
| Fibroblast | Pf4 | 7 | 6.5301E-20 | 1.12047428 | 0.266 | 0.074 | 1.306E-16 |
| Fibroblast | Arnt21 | 7 | 9.4071E-20 | 1.15820985 | 0.483 | 0.373 | 1.8814E-16 |
| Fibroblast | Col16a11 | 7 | 1.3612E-19 | 0.61089357 | 0.629 | 0.417 | 2.7224E-16 |
| Fibroblast | Ltbp13 | 7 | 1.4222E-19 | 0.35156915 | 0.856 | 0.658 | 2.8444E-16 |
| Fibroblast | Cdon1 | 7 | 1.5139E-19 | 0.75374635 | 0.643 | 0.52 | 3.0278E-16 |
| Fibroblast | Fn13 | 7 | 2.5706E-19 | 0.33183286 | 0.784 | 0.618 | 5.1412E-16 |
| Fibroblast | Col1a12 | 7 | 8.1599E-19 | 0.32246078 | 0.956 | 0.87 | 1.632E-15 |
| Fibroblast | Filip1l2 | 7 | 8.8954E-19 | 0.52240607 | 0.806 | 0.62 | 1.7791E-15 |
| Fibroblast | Olfml2b2 | 7 | 9.2528E-19 | 0.83851302 | 0.575 | 0.4 | 1.8506E-15 |
| Fibroblast | Ncam12 | 7 | 9.8512E-19 | 0.52532694 | 0.666 | 0.493 | 1.9702E-15 |

| | | | | | | | |
|---|---|---|---|---|---|---|---|
| Fibroblast | Kank4 | 7 | 1.8961E-18 | 1.09414242 | 0.342 | 0.219 | 3.7921E-15 |
| Fibroblast | Fign1 | 7 | 2.394E-18 | 1.02517498 | 0.503 | 0.354 | 4.788E-15 |
| Fibroblast | Slc1a31 | 7 | 2.8851E-18 | 0.84247838 | 0.448 | 0.268 | 5.7702E-15 |
| Fibroblast | Itgb81 | 7 | 6.6968E-18 | 0.52548206 | 0.613 | 0.438 | 1.3394E-14 |
| Fibroblast | Epha7 | 7 | 6.7309E-18 | 1.4976338 | 0.389 | 0.25 | 1.3462E-14 |
| Fibroblast | Grin3a1 | 7 | 7.6987E-18 | 0.48177382 | 0.434 | 0.232 | 1.5397E-14 |
| Fibroblast | Cacna2d2 | 7 | 8.8582E-18 | 0.77206446 | 0.306 | 0.177 | 1.7716E-14 |
| Fibroblast | Hmcn12 | 7 | 9.209E-18 | 0.37445174 | 0.68 | 0.492 | 1.8418E-14 |
| Fibroblast | Mitf | 7 | 1.0715E-17 | 0.74386156 | 0.683 | 0.562 | 2.1429E-14 |
| Fibroblast | Tmem1962 | 7 | 1.4322E-17 | 0.90924997 | 0.55 | 0.399 | 2.8643E-14 |
| Fibroblast | Thsd7b | 7 | 1.7538E-17 | 0.3127165 | 0.27 | 0.187 | 3.5076E-14 |
| Fibroblast | Mylk2 | 7 | 3.1223E-17 | 0.64476024 | 0.658 | 0.566 | 6.2446E-14 |
| Fibroblast | Adipoq | 7 | 3.4751E-17 | 1.37042074 | 0.254 | 0.133 | 6.9502E-14 |
| Fibroblast | Dok5 | 7 | 3.6401E-17 | 1.12573051 | 0.326 | 0.171 | 7.2802E-14 |
| Fibroblast | Cxcl121 | 7 | 4.5521E-17 | 1.25948018 | 0.448 | 0.313 | 9.1042E-14 |
| Fibroblast | Adamts171 | 7 | 7.815E-17 | 0.63925055 | 0.679 | 0.539 | 1.563E-13 |
| Fibroblast | Egfr1 | 7 | 9.7012E-17 | 0.4435605 | 0.843 | 0.705 | 1.9402E-13 |
| Fibroblast | Sntb11 | 7 | 1.7969E-16 | 0.34950641 | 0.489 | 0.286 | 3.5938E-13 |
| Fibroblast | Scube11 | 7 | 2.1668E-16 | 0.85593796 | 0.467 | 0.356 | 4.3336E-13 |
| Fibroblast | Grm82 | 7 | 3.4114E-16 | 0.35655835 | 0.367 | 0.238 | 6.8227E-13 |
| Fibroblast | Cyp7b12 | 7 | 3.5345E-16 | 0.78172386 | 0.614 | 0.453 | 7.0689E-13 |
| Fibroblast | Akr1c15 | 7 | 3.5705E-16 | 1.6458397 | 0.37 | 0.236 | 7.1409E-13 |
| Fibroblast | Baiap2l11 | 7 | 3.8777E-16 | 0.39669322 | 0.439 | 0.27 | 7.7553E-13 |
| Fibroblast | Nrxn31 | 7 | 4.7295E-16 | 1.14063985 | 0.429 | 0.297 | 9.4589E-13 |
| Fibroblast | Mid11 | 7 | 8.1452E-16 | 0.57839283 | 0.701 | 0.588 | 1.629E-12 |
| Fibroblast | Col14a12 | 7 | 1.3072E-15 | 0.42366798 | 0.828 | 0.715 | 2.6145E-12 |
| Fibroblast | Nrk2 | 7 | 1.5975E-15 | 0.40142285 | 0.536 | 0.377 | 3.195E-12 |
| Fibroblast | Trpm3 | 7 | 1.901E-15 | 0.82152138 | 0.451 | 0.277 | 3.802E-12 |
| Fibroblast | Grid2 | 7 | 2.9279E-15 | 0.43086124 | 0.268 | 0.159 | 5.8558E-12 |
| Fibroblast | Cpxm21 | 7 | 3.4235E-15 | 1.34625906 | 0.395 | 0.236 | 6.847E-12 |
| Fibroblast | Cacna2d33 | 7 | 9.0367E-15 | 0.6368342 | 0.715 | 0.64 | 1.8073E-11 |

| | | | | | | | |
|---|---|---|---|---|---|---|---|
| Fibroblast | Sfrp41 | 7 | 1.5447E-14 | 0.60261533 | 0.422 | 0.324 | 3.0894E-11 |
| Fibroblast | Nckap1l | 7 | 1.7295E-14 | 0.50500263 | 0.339 | 0.206 | 3.4589E-11 |
| Fibroblast | Pdzrn32 | 7 | 2.0679E-14 | 0.29007741 | 0.98 | 0.914 | 4.1357E-11 |
| Fibroblast | Abca171 | 7 | 2.5923E-14 | 0.61443504 | 0.37 | 0.238 | 5.1845E-11 |
| Fibroblast | AABR070606 | 7 | 3.5032E-14 | 0.29675943 | 0.392 | 0.228 | 7.0065E-11 |
| Fibroblast | Mrvi11 | 7 | 4.8639E-14 | 0.65007159 | 0.511 | 0.395 | 9.7279E-11 |
| Fibroblast | Adamts2 | 7 | 5.9962E-14 | 0.33452755 | 0.959 | 0.877 | 1.1992E-10 |
| Fibroblast | Epha31 | 7 | 8.2833E-14 | 1.13991152 | 0.39 | 0.275 | 1.6567E-10 |
| Fibroblast | Enah3 | 7 | 8.4592E-14 | 0.31368511 | 0.677 | 0.453 | 1.6918E-10 |
| Fibroblast | Smad61 | 7 | 1.0635E-13 | 0.78049224 | 0.555 | 0.42 | 2.1271E-10 |
| Fibroblast | Gucy1a1 | 7 | 1.2038E-13 | 0.62783235 | 0.665 | 0.522 | 2.4075E-10 |
| Fibroblast | AABR070687 | 7 | 1.2724E-13 | 1.77757582 | 0.382 | 0.183 | 2.5449E-10 |
| Fibroblast | Insr2 | 7 | 1.4653E-13 | 0.387082 | 0.633 | 0.451 | 2.9306E-10 |
| Fibroblast | AABR070410 | 7 | 1.9541E-13 | 0.54313438 | 0.346 | 0.197 | 3.9082E-10 |
| Fibroblast | Setbp12 | 7 | 1.9803E-13 | 0.59951627 | 0.693 | 0.557 | 3.9606E-10 |
| Fibroblast | Fgfr21 | 7 | 3.1376E-13 | 1.23036002 | 0.398 | 0.309 | 6.2752E-10 |
| Fibroblast | Gja1 | 7 | 6.0495E-13 | 0.310017 | 0.511 | 0.347 | 1.2099E-09 |
| Fibroblast | Specc12 | 7 | 6.3682E-13 | 0.31361068 | 0.704 | 0.523 | 1.2736E-09 |
| Fibroblast | RGD1559482 | 7 | 8.9406E-13 | 0.48086644 | 0.265 | 0.184 | 1.7881E-09 |
| Fibroblast | Rxfp11 | 7 | 9.6755E-13 | 0.998582 | 0.282 | 0.142 | 1.9351E-09 |
| Fibroblast | Pltp | 7 | 1.0863E-12 | 0.28674015 | 0.464 | 0.308 | 2.1726E-09 |
| Fibroblast | Osr11 | 7 | 1.2892E-12 | 0.55784728 | 0.524 | 0.395 | 2.5785E-09 |
| Fibroblast | Adra1b | 7 | 1.3166E-12 | 1.1077658 | 0.379 | 0.272 | 2.6332E-09 |
| Fibroblast | AABR070726 | 7 | 1.9686E-12 | 1.5406703 | 0.364 | 0.34 | 3.9373E-09 |
| Fibroblast | Ap1s3 | 7 | 2.5458E-12 | 0.62486165 | 0.455 | 0.328 | 5.0916E-09 |
| Fibroblast | Prkcb1 | 7 | 2.8156E-12 | 1.05204071 | 0.392 | 0.271 | 5.6312E-09 |
| Fibroblast | Arhgef26 | 7 | 2.8799E-12 | 0.45600341 | 0.42 | 0.24 | 5.7597E-09 |
| Fibroblast | Map1b3 | 7 | 4.5606E-12 | 0.62645384 | 0.884 | 0.793 | 9.1213E-09 |
| Fibroblast | Gpam | 7 | 4.6819E-12 | 0.30983168 | 0.397 | 0.259 | 9.3638E-09 |
| Fibroblast | Deptor1 | 7 | 6.952E-12 | 0.61891126 | 0.5 | 0.365 | 1.3904E-08 |
| Fibroblast | Tenm31 | 7 | 8.0989E-12 | 0.39873466 | 0.718 | 0.564 | 1.6198E-08 |

| | | | | | | | |
|---|---|---|---|---|---|---|---|
| Fibroblast | AABR070544 | 7 | 1.0257E-11 | 1.51310436 | 0.395 | 0.3 | 2.0514E-08 |
| Fibroblast | Dkk21 | 7 | 1.5546E-11 | 1.4973594 | 0.434 | 0.357 | 3.1093E-08 |
| Fibroblast | Adamts142 | 7 | 1.6191E-11 | 0.46337606 | 0.563 | 0.436 | 3.2383E-08 |
| Fibroblast | Slc24a32 | 7 | 1.6601E-11 | 0.48412763 | 0.506 | 0.401 | 3.3202E-08 |
| Fibroblast | Fbln51 | 7 | 2.3438E-11 | 0.44051654 | 0.586 | 0.453 | 4.6876E-08 |
| Fibroblast | Ablim22 | 7 | 2.5481E-11 | 0.47916999 | 0.292 | 0.225 | 5.0962E-08 |
| Fibroblast | Rbp13 | 7 | 2.6042E-11 | 0.27695335 | 0.494 | 0.334 | 5.2084E-08 |
| Fibroblast | Efna52 | 7 | 3.8685E-11 | 1.25901308 | 0.379 | 0.379 | 7.7369E-08 |
| Fibroblast | Smpd31 | 7 | 3.8854E-11 | 0.7527409 | 0.502 | 0.4 | 7.7708E-08 |
| Fibroblast | Sfrp21 | 7 | 4.0599E-11 | 1.109019 | 0.35 | 0.254 | 8.1199E-08 |
| Fibroblast | Dapk13 | 7 | 4.2609E-11 | 0.40300687 | 0.763 | 0.653 | 8.5217E-08 |
| Fibroblast | Adamts32 | 7 | 5.7665E-11 | 0.26668573 | 0.453 | 0.291 | 1.1533E-07 |
| Fibroblast | Itgb61 | 7 | 5.9394E-11 | 1.04541646 | 0.274 | 0.17 | 1.1879E-07 |
| Fibroblast | Ptpn131 | 7 | 6.3843E-11 | 0.39358282 | 0.561 | 0.418 | 1.2769E-07 |
| Fibroblast | Ddc1 | 7 | 7.4413E-11 | 0.44224235 | 0.386 | 0.226 | 1.4883E-07 |
| Fibroblast | Ntf32 | 7 | 7.8099E-11 | 0.65978371 | 0.415 | 0.331 | 1.562E-07 |
| Fibroblast | Tmem172 | 7 | 7.9219E-11 | 0.30523534 | 0.461 | 0.355 | 1.5844E-07 |
| Fibroblast | Angpt22 | 7 | 9.031E-11 | 1.47623802 | 0.346 | 0.236 | 1.8062E-07 |
| Fibroblast | Il1rl2 | 7 | 9.418E-11 | 0.25857774 | 0.462 | 0.302 | 1.8836E-07 |
| Fibroblast | Rnf432 | 7 | 1.046E-10 | 0.68313065 | 0.332 | 0.17 | 2.092E-07 |
| Fibroblast | Steap42 | 7 | 1.0509E-10 | 0.77373321 | 0.39 | 0.243 | 2.1018E-07 |
| Fibroblast | Sytl22 | 7 | 1.1889E-10 | 0.96796977 | 0.262 | 0.183 | 2.3779E-07 |
| Fibroblast | Cdh111 | 7 | 1.3097E-10 | 0.35682596 | 0.785 | 0.683 | 2.6194E-07 |
| Fibroblast | Dcn2 | 7 | 1.3268E-10 | 0.2953972 | 0.89 | 0.789 | 2.6536E-07 |
| Fibroblast | Emid1 | 7 | 1.4535E-10 | 0.75110152 | 0.298 | 0.184 | 2.907E-07 |
| Fibroblast | Gpnmb | 7 | 1.7905E-10 | 0.84300666 | 0.418 | 0.301 | 3.581E-07 |
| Fibroblast | Pi151 | 7 | 2.1895E-10 | 0.52941132 | 0.447 | 0.329 | 4.379E-07 |
| Fibroblast | Rnd31 | 7 | 2.2489E-10 | 0.3187913 | 0.665 | 0.531 | 4.4978E-07 |
| Fibroblast | Tmem264 | 7 | 2.2544E-10 | 0.56877653 | 0.447 | 0.351 | 4.5088E-07 |
| Fibroblast | Abca12 | 7 | 2.266E-10 | 0.37886203 | 0.596 | 0.519 | 4.532E-07 |
| Fibroblast | Epha41 | 7 | 3.3129E-10 | 0.93660725 | 0.5 | 0.403 | 6.6258E-07 |

| | | | | | | | |
|---|---|---|---|---|---|---|---|
| Fibroblast | Reep12 | 7 | 4.5834E-10 | 0.74939136 | 0.379 | 0.24 | 9.1668E-07 |
| Fibroblast | Mmp163 | 7 | 5.3171E-10 | 0.41273357 | 0.401 | 0.304 | 1.0634E-06 |
| Fibroblast | Snta11 | 7 | 5.7399E-10 | 0.28926479 | 0.541 | 0.401 | 1.148E-06 |
| Fibroblast | Irf83 | 7 | 8.812E-10 | 0.45630279 | 0.406 | 0.28 | 1.7624E-06 |
| Fibroblast | Aebp1 | 7 | 9.7232E-10 | 0.34627055 | 0.605 | 0.466 | 1.9446E-06 |
| Fibroblast | C61 | 7 | 1.0936E-09 | 0.97730379 | 0.364 | 0.318 | 2.1873E-06 |
| Fibroblast | Cdh71 | 7 | 1.5666E-09 | 0.65801155 | 0.356 | 0.25 | 3.1332E-06 |
| Fibroblast | Scara51 | 7 | 4.1525E-09 | 0.32835654 | 0.589 | 0.503 | 8.3051E-06 |
| Fibroblast | Ldb2 | 7 | 4.4888E-09 | 1.00416284 | 0.331 | 0.238 | 8.9777E-06 |
| Fibroblast | Gulp1 | 7 | 7.5906E-09 | 0.33062904 | 0.619 | 0.485 | 1.5181E-05 |
| Fibroblast | Lcp2 | 7 | 9.8349E-09 | 0.40844942 | 0.281 | 0.198 | 1.967E-05 |
| Fibroblast | Bcar3 | 7 | 1.0132E-08 | 0.91574744 | 0.331 | 0.242 | 2.0263E-05 |
| Fibroblast | Dpysl32 | 7 | 1.163E-08 | 0.2744416 | 0.873 | 0.737 | 2.3261E-05 |
| Fibroblast | Notch3 | 7 | 1.2634E-08 | 0.57467531 | 0.481 | 0.334 | 2.5268E-05 |
| Fibroblast | Plcb11 | 7 | 1.2724E-08 | 0.40377379 | 0.426 | 0.319 | 2.5447E-05 |
| Fibroblast | Mctp11 | 7 | 1.5807E-08 | 0.43531606 | 0.459 | 0.326 | 3.1614E-05 |
| Fibroblast | Lmod11 | 7 | 1.785E-08 | 0.34902048 | 0.461 | 0.347 | 3.5701E-05 |
| Fibroblast | Slc43a21 | 7 | 1.9416E-08 | 0.59045422 | 0.373 | 0.288 | 3.8832E-05 |
| Fibroblast | Sh3gl31 | 7 | 2.058E-08 | 0.62815169 | 0.397 | 0.269 | 4.1159E-05 |
| Fibroblast | Veph11 | 7 | 2.3967E-08 | 0.50859455 | 0.348 | 0.218 | 4.7934E-05 |
| Fibroblast | Rrad | 7 | 2.7401E-08 | 0.4305633 | 0.273 | 0.149 | 5.4803E-05 |
| Fibroblast | Sorcs11 | 7 | 3.3235E-08 | 0.25614587 | 0.26 | 0.198 | 6.647E-05 |
| Fibroblast | Aff33 | 7 | 4.0658E-08 | 0.35229899 | 0.755 | 0.689 | 8.1316E-05 |
| Fibroblast | Ngf2 | 7 | 5.092E-08 | 0.30727748 | 0.671 | 0.572 | 0.00010184 |
| Fibroblast | Parm1 | 7 | 7.8247E-08 | 0.34629792 | 0.292 | 0.186 | 0.00015649 |
| Fibroblast | Tbxas11 | 7 | 7.837E-08 | 0.51762457 | 0.342 | 0.251 | 0.00015674 |
| Fibroblast | Bzw21 | 7 | 1.4172E-07 | 0.27505488 | 0.47 | 0.35 | 0.00028344 |
| Fibroblast | LOC1009095 | 7 | 1.4272E-07 | 0.32996463 | 0.359 | 0.256 | 0.00028543 |
| Fibroblast | Slc7a22 | 7 | 1.6963E-07 | 0.26399175 | 0.475 | 0.385 | 0.00033925 |
| Fibroblast | Tmem1822 | 7 | 2.1209E-07 | 0.25582317 | 0.364 | 0.265 | 0.00042418 |
| Fibroblast | Cacnb41 | 7 | 2.8846E-07 | 0.47252547 | 0.466 | 0.381 | 0.00057691 |

| | | | | | | | |
|---|---|---|---|---|---|---|---|
| Fibroblast | Syn3 | 7 | 3.0559E-07 | 0.58760562 | 0.408 | 0.325 | 0.00061119 |
| Fibroblast | Ndst32 | 7 | 4.0058E-07 | 1.22375444 | 0.361 | 0.37 | 0.00080116 |
| Fibroblast | Slc16a22 | 7 | 5.2559E-07 | 0.47278098 | 0.478 | 0.426 | 0.00105118 |
| Fibroblast | Pcbp32 | 7 | 1.1693E-06 | 0.36332309 | 0.475 | 0.388 | 0.00233862 |
| Fibroblast | Pawr1 | 7 | 1.1868E-06 | 0.79037469 | 0.34 | 0.252 | 0.00237366 |
| Fibroblast | Tango21 | 7 | 1.1893E-06 | 0.25857384 | 0.429 | 0.353 | 0.00237868 |
| Fibroblast | Hs6st22 | 7 | 1.3197E-06 | 0.38677226 | 0.453 | 0.43 | 0.00263931 |
| Fibroblast | Fam180a | 7 | 1.5747E-06 | 0.56467898 | 0.345 | 0.282 | 0.00314949 |
| Fibroblast | Nuak1 | 7 | 2.0496E-06 | 0.28083979 | 0.484 | 0.398 | 0.00409918 |
| Fibroblast | AABR070492 | 7 | 2.6901E-06 | 0.92922364 | 0.266 | 0.216 | 0.00538013 |
| Fibroblast | Myo1f1 | 7 | 2.917E-06 | 0.69570067 | 0.334 | 0.279 | 0.00583407 |
| Fibroblast | Sfxn52 | 7 | 3.4626E-06 | 0.45295933 | 0.312 | 0.229 | 0.00692515 |
| Fibroblast | LOC690097 | 7 | 3.7338E-06 | 0.32340754 | 0.318 | 0.108 | 0.00746758 |
| Fibroblast | AABR070605 | 7 | 3.7744E-06 | 0.77834114 | 0.362 | 0.323 | 0.00754884 |
| Fibroblast | Cldn151 | 7 | 7.3576E-06 | 0.59104838 | 0.234 | 0.256 | 0.01471525 |
| Fibroblast | AABR070275 | 7 | 2.1178E-05 | 0.26442842 | 0.353 | 0.303 | 0.04235543 |
| Fibroblast | Mos | 8 | 2.515E-124 | 3.37871984 | 0.269 | 0.027 | 5.03E-121 |
| Fibroblast | Glp2r | 8 | 1.119E-99 | 1.8454313 | 0.379 | 0.073 | 2.2372E-96 |
| Fibroblast | Rasgrp1 | 8 | 1.8002E-95 | 1.61611337 | 0.345 | 0.057 | 3.6004E-92 |
| Fibroblast | E2f7 | 8 | 2.4336E-88 | 1.59538122 | 0.389 | 0.079 | 4.8672E-85 |
| Fibroblast | Ccdc103 | 8 | 3.4432E-82 | 1.07460394 | 0.337 | 0.084 | 6.8864E-79 |
| Fibroblast | Trpc61 | 8 | 8.2121E-78 | 0.51958783 | 0.554 | 0.201 | 1.6424E-74 |
| Fibroblast | Slc44a52 | 8 | 2.6678E-70 | 1.68918646 | 0.718 | 0.375 | 5.3357E-67 |
| Fibroblast | Susd5 | 8 | 1.1228E-66 | 1.44923073 | 0.257 | 0.043 | 2.2456E-63 |
| Fibroblast | Fos1 | 8 | 3.5777E-65 | 1.84145912 | 0.581 | 0.254 | 7.1554E-62 |
| Fibroblast | Pmfbp1 | 8 | 1.2263E-64 | 1.0829821 | 0.267 | 0.053 | 2.4525E-61 |
| Fibroblast | Kcnh1 | 8 | 9.6022E-58 | 2.04675547 | 0.257 | 0.054 | 1.9204E-54 |
| Fibroblast | L1cam | 8 | 4.116E-57 | 0.58282732 | 0.385 | 0.151 | 8.2319E-54 |
| Fibroblast | Adgrg3 | 8 | 2.4148E-56 | 0.67874975 | 0.39 | 0.111 | 4.8295E-53 |
| Fibroblast | Slit1 | 8 | 6.8877E-53 | 2.04751762 | 0.411 | 0.137 | 1.3775E-49 |
| Fibroblast | AABR070651 | 8 | 5.5797E-50 | 2.06010094 | 0.368 | 0.124 | 1.1159E-46 |

| | | | | | | | |
|---|---|---|---|---|---|---|---|
| Fibroblast | Unc45b2 | 8 | 9.4989E-50 | 0.91950313 | 0.611 | 0.272 | 1.8998E-46 |
| Fibroblast | Scel | 8 | 1.671E-49 | 1.0054074 | 0.284 | 0.071 | 3.342E-46 |
| Fibroblast | Ptch2 | 8 | 3.8368E-49 | 3.34838 | 0.472 | 0.228 | 7.6735E-46 |
| Fibroblast | Mapk101 | 8 | 4.9506E-49 | 0.85446418 | 0.566 | 0.276 | 9.9012E-46 |
| Fibroblast | Col6a63 | 8 | 1.2018E-48 | 1.06082641 | 0.93 | 0.726 | 2.4035E-45 |
| Fibroblast | Rasgef1c | 8 | 3.9573E-48 | 0.56894469 | 0.259 | 0.147 | 7.9147E-45 |
| Fibroblast | Rims11 | 8 | 1.2905E-46 | 1.80609806 | 0.389 | 0.127 | 2.5811E-43 |
| Fibroblast | Adamts192 | 8 | 2.8294E-46 | 1.08344741 | 0.846 | 0.637 | 5.6588E-43 |
| Fibroblast | AABR070525 | 8 | 2.4703E-45 | 0.71468216 | 0.596 | 0.256 | 4.9405E-42 |
| Fibroblast | Mx11 | 8 | 2.8662E-45 | 0.35264619 | 0.463 | 0.198 | 5.7325E-42 |
| Fibroblast | Tex223 | 8 | 4.542E-45 | 0.53592493 | 0.549 | 0.232 | 9.0839E-42 |
| Fibroblast | Lcp21 | 8 | 3.5442E-44 | 0.27326067 | 0.486 | 0.187 | 7.0884E-41 |
| Fibroblast | Col22a12 | 8 | 6.648E-44 | 1.52785718 | 0.617 | 0.34 | 1.3296E-40 |
| Fibroblast | Aff21 | 8 | 2.7981E-43 | 1.13011276 | 0.716 | 0.401 | 5.5962E-40 |
| Fibroblast | Hpd | 8 | 1.2121E-42 | 0.9999607 | 0.35 | 0.095 | 2.4243E-39 |
| Fibroblast | Grm83 | 8 | 1.5865E-41 | 0.70294252 | 0.507 | 0.232 | 3.1729E-38 |
| Fibroblast | Neb1 | 8 | 6.6449E-41 | 1.09580659 | 0.446 | 0.192 | 1.329E-37 |
| Fibroblast | Asb142 | 8 | 1.3284E-40 | 0.84929014 | 0.547 | 0.258 | 2.6568E-37 |
| Fibroblast | Cdh42 | 8 | 1.8201E-40 | 0.73901674 | 0.415 | 0.173 | 3.6402E-37 |
| Fibroblast | Kcnt21 | 8 | 2.572E-40 | 0.6861895 | 0.699 | 0.364 | 5.1441E-37 |
| Fibroblast | Lrrtm41 | 8 | 5.0743E-40 | 1.56619274 | 0.556 | 0.271 | 1.0149E-36 |
| Fibroblast | Afap1l23 | 8 | 6.4157E-40 | 1.03334131 | 0.644 | 0.369 | 1.2831E-36 |
| Fibroblast | Tecrl | 8 | 3.3747E-39 | 1.0159973 | 0.484 | 0.207 | 6.7494E-36 |
| Fibroblast | Hhip1 | 8 | 1.5574E-38 | 3.61220022 | 0.314 | 0.102 | 3.1148E-35 |
| Fibroblast | Lcp1 | 8 | 8.0556E-38 | 0.86450398 | 0.51 | 0.228 | 1.6111E-34 |
| Fibroblast | Nckap1l1 | 8 | 9.094E-38 | 0.28174277 | 0.423 | 0.203 | 1.8188E-34 |
| Fibroblast | Ska11 | 8 | 8.6592E-37 | 0.9509675 | 0.328 | 0.099 | 1.7318E-33 |
| Fibroblast | Thbs41 | 8 | 8.8847E-37 | 0.69225444 | 0.533 | 0.27 | 1.7769E-33 |
| Fibroblast | Sox17 | 8 | 1.2798E-36 | 0.63298121 | 0.413 | 0.137 | 2.5595E-33 |
| Fibroblast | Tmem1823 | 8 | 1.4415E-36 | 0.87165763 | 0.493 | 0.259 | 2.883E-33 |
| Fibroblast | Ebf24 | 8 | 1.6513E-36 | 0.70507998 | 0.962 | 0.858 | 3.3027E-33 |

| | | | | | | | |
|---|---|---|---|---|---|---|---|
| Fibroblast | Itga72 | 8 | 2.7038E-36 | 0.55449406 | 0.505 | 0.231 | 5.4076E-33 |
| Fibroblast | Adamts172 | 8 | 2.5666E-35 | 0.97088335 | 0.789 | 0.535 | 5.1331E-32 |
| Fibroblast | Esco21 | 8 | 3.2965E-35 | 1.83838903 | 0.293 | 0.118 | 6.5929E-32 |
| Fibroblast | Pclo2 | 8 | 1.1193E-34 | 1.16839281 | 0.503 | 0.273 | 2.2386E-31 |
| Fibroblast | Apcdd1l2 | 8 | 2.9447E-34 | 1.10170358 | 0.531 | 0.304 | 5.8893E-31 |
| Fibroblast | Pax5 | 8 | 4.4643E-34 | 0.56102137 | 0.263 | 0.061 | 8.9286E-31 |
| Fibroblast | Kif18b1 | 8 | 5.7501E-34 | 0.42377176 | 0.377 | 0.14 | 1.15E-30 |
| Fibroblast | AABR070515 | 8 | 7.6121E-34 | 0.70016555 | 0.474 | 0.238 | 1.5224E-30 |
| Fibroblast | Tuba1b | 8 | 7.6465E-34 | 0.8408304 | 0.503 | 0.214 | 1.5293E-30 |
| Fibroblast | Kif232 | 8 | 8.5077E-34 | 1.56132108 | 0.364 | 0.158 | 1.7015E-30 |
| Fibroblast | Vash22 | 8 | 6.5689E-33 | 0.47643316 | 0.436 | 0.229 | 1.3138E-29 |
| Fibroblast | Cilp3 | 8 | 8.4941E-33 | 0.74942173 | 0.604 | 0.335 | 1.6988E-29 |
| Fibroblast | Aatk3 | 8 | 1.8614E-32 | 0.57190105 | 0.48 | 0.248 | 3.7229E-29 |
| Fibroblast | Slc1a11 | 8 | 1.2252E-31 | 0.83300503 | 0.472 | 0.254 | 2.4503E-28 |
| Fibroblast | Cotl12 | 8 | 2.1232E-31 | 0.4288954 | 0.564 | 0.318 | 4.2464E-28 |
| Fibroblast | Cacna1g1 | 8 | 2.358E-31 | 1.0529065 | 0.653 | 0.397 | 4.7161E-28 |
| Fibroblast | Ldb21 | 8 | 3.3663E-31 | 0.78089659 | 0.51 | 0.229 | 6.7327E-28 |
| Fibroblast | Hs6st23 | 8 | 7.1537E-31 | 0.80726578 | 0.686 | 0.417 | 1.4307E-27 |
| Fibroblast | Tgfbi2 | 8 | 8.4713E-31 | 1.02751262 | 0.556 | 0.352 | 1.6943E-27 |
| Fibroblast | Hcn13 | 8 | 9.3023E-31 | 0.71763125 | 0.461 | 0.252 | 1.8605E-27 |
| Fibroblast | Bdnf4 | 8 | 1.1869E-30 | 0.3630994 | 0.571 | 0.272 | 2.3739E-27 |
| Fibroblast | Slc38a3 | 8 | 1.2369E-30 | 0.53886765 | 0.461 | 0.219 | 2.4738E-27 |
| Fibroblast | Kcnj3 | 8 | 2.6983E-30 | 0.84873479 | 0.463 | 0.218 | 5.3965E-27 |
| Fibroblast | Cdh72 | 8 | 3.4088E-30 | 1.60202735 | 0.427 | 0.247 | 6.8176E-27 |
| Fibroblast | Plek | 8 | 5.3004E-30 | 1.26161087 | 0.364 | 0.107 | 1.0601E-26 |
| Fibroblast | Grin3a2 | 8 | 7.8132E-30 | 1.33389849 | 0.427 | 0.235 | 1.5626E-26 |
| Fibroblast | AABR070606 | 8 | 8.7687E-30 | 1.97670236 | 0.446 | 0.227 | 1.7537E-26 |
| Fibroblast | Akap122 | 8 | 1.1574E-29 | 0.71104349 | 0.678 | 0.407 | 2.3148E-26 |
| Fibroblast | Sorcs2 | 8 | 1.5424E-29 | 2.06244623 | 0.385 | 0.193 | 3.0849E-26 |
| Fibroblast | AABR070171 | 8 | 9.1891E-29 | 0.66111088 | 0.434 | 0.268 | 1.8378E-25 |
| Fibroblast | Mis18bp11 | 8 | 2.0095E-28 | 0.4995605 | 0.509 | 0.269 | 4.019E-25 |

| | | | | | | | |
|---|---|---|---|---|---|---|---|
| Fibroblast | Synpo2l1 | 8 | 2.703E-28 | 0.40348526 | 0.421 | 0.231 | 5.4059E-25 |
| Fibroblast | Abca172 | 8 | 7.1467E-28 | 0.3996511 | 0.427 | 0.237 | 1.4293E-24 |
| Fibroblast | Ankrd62 | 8 | 1.159E-27 | 0.29868011 | 0.476 | 0.254 | 2.3181E-24 |
| Fibroblast | Ky | 8 | 1.2783E-27 | 0.7442772 | 0.328 | 0.164 | 2.5566E-24 |
| Fibroblast | Adgrg61 | 8 | 1.4291E-27 | 0.41386734 | 0.402 | 0.209 | 2.8582E-24 |
| Fibroblast | Hspb61 | 8 | 1.7865E-27 | 0.51413217 | 0.491 | 0.258 | 3.5729E-24 |
| Fibroblast | Opcml2 | 8 | 2.5445E-27 | 0.74036656 | 0.583 | 0.338 | 5.089E-24 |
| Fibroblast | Popdc23 | 8 | 2.7675E-27 | 0.39488298 | 0.499 | 0.276 | 5.535E-24 |
| Fibroblast | Knl1 | 8 | 4.768E-27 | 0.56423957 | 0.341 | 0.199 | 9.5361E-24 |
| Fibroblast | Susd11 | 8 | 6.022E-27 | 0.28779618 | 0.415 | 0.211 | 1.2044E-23 |
| Fibroblast | Ppargc1b2 | 8 | 7.2808E-27 | 0.42759354 | 0.602 | 0.344 | 1.4562E-23 |
| Fibroblast | Mest3 | 8 | 1.1022E-26 | 1.9164633 | 0.379 | 0.173 | 2.2044E-23 |
| Fibroblast | Rbm242 | 8 | 1.7968E-26 | 0.26516296 | 0.457 | 0.238 | 3.5937E-23 |
| Fibroblast | Shtn1 | 8 | 2.2646E-26 | 0.51323682 | 0.341 | 0.179 | 4.5292E-23 |
| Fibroblast | Hvcn12 | 8 | 2.3332E-26 | 1.00848649 | 0.497 | 0.287 | 4.6664E-23 |
| Fibroblast | Sema5a | 8 | 3.7022E-26 | 0.59973209 | 0.371 | 0.18 | 7.4043E-23 |
| Fibroblast | Dlgap51 | 8 | 9.0556E-26 | 0.35929747 | 0.429 | 0.224 | 1.8111E-22 |
| Fibroblast | Pde1a1 | 8 | 9.523E-26 | 0.76031578 | 0.827 | 0.65 | 1.9046E-22 |
| Fibroblast | Agbl12 | 8 | 1.4826E-25 | 0.55296551 | 0.564 | 0.367 | 2.9652E-22 |
| Fibroblast | Tspan112 | 8 | 2.672E-25 | 1.02193798 | 0.573 | 0.343 | 5.344E-22 |
| Fibroblast | Sparcl11 | 8 | 2.9581E-25 | 0.69478361 | 0.69 | 0.467 | 5.9162E-22 |
| Fibroblast | Adgrl4 | 8 | 4.8885E-25 | 0.36416898 | 0.408 | 0.183 | 9.777E-22 |
| Fibroblast | Cadm1 | 8 | 4.9951E-25 | 0.383919 | 0.27 | 0.082 | 9.9903E-22 |
| Fibroblast | Tfap2b1 | 8 | 5.8323E-25 | 0.58686517 | 0.291 | 0.108 | 1.1665E-21 |
| Fibroblast | Serping11 | 8 | 7.4821E-25 | 0.32270142 | 0.537 | 0.295 | 1.4964E-21 |
| Fibroblast | Pygm1 | 8 | 1.1275E-24 | 0.52891008 | 0.404 | 0.216 | 2.2551E-21 |
| Fibroblast | Tec2 | 8 | 1.2508E-24 | 0.62971941 | 0.467 | 0.248 | 2.5015E-21 |
| Fibroblast | Postn3 | 8 | 2.484E-24 | 0.52515891 | 0.93 | 0.767 | 4.9679E-21 |
| Fibroblast | Acsl11 | 8 | 6.752E-24 | 0.48483856 | 0.459 | 0.271 | 1.3504E-20 |
| Fibroblast | Mboat2 | 8 | 7.9577E-24 | 0.98701741 | 0.589 | 0.37 | 1.5915E-20 |
| Fibroblast | Ptprc2 | 8 | 1.1285E-23 | 0.40398565 | 0.457 | 0.247 | 2.2569E-20 |

| | | | | | | | |
|---|---|---|---|---|---|---|---|
| Fibroblast | Btbd112 | 8 | 1.1312E-23 | 0.55048175 | 0.579 | 0.36 | 2.2623E-20 |
| Fibroblast | Kit2 | 8 | 1.1478E-23 | 0.80017878 | 0.499 | 0.318 | 2.2956E-20 |
| Fibroblast | Slc10a61 | 8 | 1.3373E-23 | 0.61868005 | 0.324 | 0.188 | 2.6746E-20 |
| Fibroblast | Adra1a4 | 8 | 2.5278E-23 | 0.57906668 | 0.615 | 0.36 | 5.0557E-20 |
| Fibroblast | Meox21 | 8 | 2.7215E-23 | 0.92051323 | 0.638 | 0.374 | 5.4431E-20 |
| Fibroblast | Asb15 | 8 | 2.9561E-23 | 1.08843404 | 0.429 | 0.264 | 5.9122E-20 |
| Fibroblast | Cacna2d21 | 8 | 5.2096E-23 | 0.88953175 | 0.432 | 0.171 | 1.0419E-19 |
| Fibroblast | Pde1c2 | 8 | 1.3105E-22 | 1.29304379 | 0.415 | 0.242 | 2.6211E-19 |
| Fibroblast | Megf111 | 8 | 2.2823E-22 | 0.73409461 | 0.43 | 0.216 | 4.5646E-19 |
| Fibroblast | Zfp385b2 | 8 | 2.497E-22 | 0.60779751 | 0.627 | 0.448 | 4.994E-19 |
| Fibroblast | Parm11 | 8 | 3.2922E-22 | 0.56420588 | 0.366 | 0.183 | 6.5844E-19 |
| Fibroblast | RGD1563354 | 8 | 5.8853E-22 | 0.53489595 | 0.728 | 0.527 | 1.1771E-18 |
| Fibroblast | Cyp26b11 | 8 | 7.3912E-22 | 1.16242716 | 0.446 | 0.21 | 1.4782E-18 |
| Fibroblast | Ankrd9 | 8 | 1.1879E-21 | 0.70691247 | 0.421 | 0.217 | 2.3758E-18 |
| Fibroblast | Nrk3 | 8 | 1.2448E-21 | 0.67721169 | 0.543 | 0.379 | 2.4896E-18 |
| Fibroblast | Mlf12 | 8 | 1.2805E-21 | 0.65716647 | 0.381 | 0.228 | 2.5611E-18 |
| Fibroblast | Eef1a22 | 8 | 1.3853E-21 | 0.53434064 | 0.53 | 0.314 | 2.7706E-18 |
| Fibroblast | LOC6877801 | 8 | 2.1312E-21 | 0.26498642 | 0.469 | 0.246 | 4.2624E-18 |
| Fibroblast | Prkaa22 | 8 | 3.1853E-21 | 0.53444035 | 0.375 | 0.226 | 6.3707E-18 |
| Fibroblast | Eda2 | 8 | 3.6496E-21 | 0.71182055 | 0.838 | 0.659 | 7.2993E-18 |
| Fibroblast | Fbxo321 | 8 | 4.7995E-21 | 0.51226728 | 0.52 | 0.32 | 9.599E-18 |
| Fibroblast | Stmn42 | 8 | 5.9971E-21 | 0.39346408 | 0.43 | 0.28 | 1.1994E-17 |
| Fibroblast | Serpine24 | 8 | 6.5885E-21 | 0.35934444 | 0.575 | 0.357 | 1.3177E-17 |
| Fibroblast | Ctsd | 8 | 6.668E-21 | 0.46877246 | 0.37 | 0.216 | 1.3336E-17 |
| Fibroblast | Mrc1 | 8 | 8.9641E-21 | 0.43431781 | 0.387 | 0.215 | 1.7928E-17 |
| Fibroblast | Fgfr22 | 8 | 1.4734E-20 | 0.46305996 | 0.49 | 0.305 | 2.9467E-17 |
| Fibroblast | Lpl2 | 8 | 1.9768E-20 | 0.40498091 | 0.545 | 0.33 | 3.9535E-17 |
| Fibroblast | Adgrb3 | 8 | 2.255E-20 | 0.68679858 | 0.337 | 0.138 | 4.51E-17 |
| Fibroblast | Glb1l22 | 8 | 3.0024E-20 | 0.43054502 | 0.43 | 0.248 | 6.0047E-17 |
| Fibroblast | Ppargc1a2 | 8 | 6.2019E-20 | 0.44313144 | 0.417 | 0.241 | 1.2404E-16 |
| Fibroblast | Abcc82 | 8 | 6.616E-20 | 0.42334443 | 0.307 | 0.227 | 1.3232E-16 |

| | | | | | | | |
|---|---|---|---|---|---|---|---|
| Fibroblast | Abcd21 | 8 | 1.6201E-19 | 0.51916243 | 0.333 | 0.203 | 3.2402E-16 |
| Fibroblast | Ppp1r14c2 | 8 | 1.7272E-19 | 0.38731216 | 0.484 | 0.305 | 3.4544E-16 |
| Fibroblast | Efcc12 | 8 | 1.8142E-19 | 0.31155816 | 0.676 | 0.437 | 3.6284E-16 |
| Fibroblast | Clu | 8 | 2.2015E-19 | 0.43028993 | 0.368 | 0.225 | 4.403E-16 |
| Fibroblast | Rspo14 | 8 | 3.9817E-19 | 0.25266586 | 0.432 | 0.265 | 7.9633E-16 |
| Fibroblast | AABR070172 | 8 | 4.3575E-19 | 0.25216176 | 0.259 | 0.169 | 8.7151E-16 |
| Fibroblast | AABR070179 | 8 | 5.9098E-19 | 1.3377865 | 0.425 | 0.191 | 1.182E-15 |
| Fibroblast | Atp5f1e1 | 8 | 9.6224E-19 | 0.26558625 | 0.501 | 0.313 | 1.9245E-15 |
| Fibroblast | Rgs51 | 8 | 1.5084E-18 | 0.55249997 | 0.408 | 0.235 | 3.0169E-15 |
| Fibroblast | Rimbp22 | 8 | 2.1616E-18 | 0.56904178 | 0.421 | 0.264 | 4.3233E-15 |
| Fibroblast | Nrap1 | 8 | 2.5916E-18 | 0.28309908 | 0.417 | 0.235 | 5.1833E-15 |
| Fibroblast | Pappa23 | 8 | 2.8166E-18 | 0.36179839 | 0.354 | 0.237 | 5.6331E-15 |
| Fibroblast | Colgalt23 | 8 | 4.3639E-18 | 0.51614475 | 0.688 | 0.507 | 8.7279E-15 |
| Fibroblast | Kcnk3 | 8 | 8.1135E-18 | 0.70713235 | 0.364 | 0.166 | 1.6227E-14 |
| Fibroblast | Adcy52 | 8 | 9.0908E-18 | 0.43104585 | 0.594 | 0.396 | 1.8182E-14 |
| Fibroblast | Ptafr2 | 8 | 1.2116E-17 | 0.2980254 | 0.436 | 0.292 | 2.4231E-14 |
| Fibroblast | Pdpn2 | 8 | 3.5003E-17 | 0.46540894 | 0.425 | 0.279 | 7.0005E-14 |
| Fibroblast | Arl152 | 8 | 3.5273E-17 | 0.45723162 | 0.973 | 0.903 | 7.0547E-14 |
| Fibroblast | Iqgap3 | 8 | 3.5534E-17 | 0.80899668 | 0.28 | 0.104 | 7.1067E-14 |
| Fibroblast | Lgmn2 | 8 | 4.6648E-17 | 0.35485752 | 0.488 | 0.318 | 9.3297E-14 |
| Fibroblast | Gpc31 | 8 | 5.606E-17 | 1.43292132 | 0.282 | 0.217 | 1.1212E-13 |
| Fibroblast | Lsamp1 | 8 | 5.7213E-17 | 0.53365706 | 0.366 | 0.236 | 1.1443E-13 |
| Fibroblast | AABR070592 | 8 | 6.5437E-17 | 0.58320958 | 0.737 | 0.543 | 1.3087E-13 |
| Fibroblast | Rtn4rl11 | 8 | 7.5689E-17 | 0.33463548 | 0.259 | 0.172 | 1.5138E-13 |
| Fibroblast | Egr12 | 8 | 1.0155E-16 | 0.63070248 | 0.392 | 0.258 | 2.0309E-13 |
| Fibroblast | Egflam | 8 | 1.0216E-16 | 0.36243157 | 0.274 | 0.186 | 2.0432E-13 |
| Fibroblast | Abi3bp | 8 | 1.0566E-16 | 0.55543904 | 0.939 | 0.815 | 2.1132E-13 |
| Fibroblast | Med12l2 | 8 | 1.2666E-16 | 0.36340845 | 0.461 | 0.293 | 2.5331E-13 |
| Fibroblast | Chst91 | 8 | 1.3408E-16 | 0.30221588 | 0.265 | 0.111 | 2.6817E-13 |
| Fibroblast | Fgd21 | 8 | 1.4873E-16 | 0.68916213 | 0.354 | 0.278 | 2.9745E-13 |
| Fibroblast | AC110709.22 | 8 | 1.5314E-16 | 0.7882004 | 0.379 | 0.246 | 3.0629E-13 |

| | | | | | | | |
|---|---|---|---|---|---|---|---|
| Fibroblast | Pdgfra1 | 8 | 1.8263E-16 | 0.42611589 | 0.914 | 0.761 | 3.6527E-13 |
| Fibroblast | Cemip22 | 8 | 2.0227E-16 | 0.28250537 | 0.722 | 0.487 | 4.0454E-13 |
| Fibroblast | Robo11 | 8 | 2.0485E-16 | 0.52361037 | 0.83 | 0.656 | 4.0969E-13 |
| Fibroblast | Tmem265 | 8 | 2.2203E-16 | 0.47075778 | 0.476 | 0.35 | 4.4406E-13 |
| Fibroblast | Col5a33 | 8 | 2.6264E-16 | 0.58819372 | 0.611 | 0.497 | 5.2527E-13 |
| Fibroblast | Ctnnd21 | 8 | 3.2634E-16 | 0.73937364 | 0.366 | 0.147 | 6.5267E-13 |
| Fibroblast | Enpp12 | 8 | 5.9108E-16 | 0.35885668 | 0.743 | 0.529 | 1.1822E-12 |
| Fibroblast | Islr | 8 | 6.0148E-16 | 0.27828695 | 0.514 | 0.332 | 1.203E-12 |
| Fibroblast | Plbd11 | 8 | 6.339E-16 | 0.56548809 | 0.366 | 0.208 | 1.2678E-12 |
| Fibroblast | Gucy1a11 | 8 | 7.6457E-16 | 0.48197108 | 0.703 | 0.522 | 1.5291E-12 |
| Fibroblast | Uqcrq2 | 8 | 8.9511E-16 | 0.49577485 | 0.48 | 0.334 | 1.7902E-12 |
| Fibroblast | Lrrc171 | 8 | 1.2938E-15 | 0.64512063 | 0.581 | 0.432 | 2.5875E-12 |
| Fibroblast | Gli23 | 8 | 1.4607E-15 | 0.54988698 | 0.731 | 0.588 | 2.9215E-12 |
| Fibroblast | Neurl11 | 8 | 1.656E-15 | 0.47451664 | 0.274 | 0.233 | 3.3121E-12 |
| Fibroblast | Nusap12 | 8 | 1.6818E-15 | 0.80090388 | 0.354 | 0.198 | 3.3636E-12 |
| Fibroblast | Cox6a21 | 8 | 1.7982E-15 | 0.31402103 | 0.522 | 0.341 | 3.5964E-12 |
| Fibroblast | Kif11 | 8 | 1.8975E-15 | 0.95530767 | 0.25 | 0.187 | 3.795E-12 |
| Fibroblast | Pln1 | 8 | 3.0679E-15 | 0.30418156 | 0.651 | 0.446 | 6.1359E-12 |
| Fibroblast | Dlgap12 | 8 | 5.6465E-15 | 0.45737868 | 0.571 | 0.386 | 1.1293E-11 |
| Fibroblast | Ctss | 8 | 7.4724E-15 | 0.72747632 | 0.406 | 0.265 | 1.4945E-11 |
| Fibroblast | Samd122 | 8 | 8.5236E-15 | 0.78436038 | 0.415 | 0.307 | 1.7047E-11 |
| Fibroblast | Matn21 | 8 | 9.4279E-15 | 0.55669011 | 0.65 | 0.508 | 1.8856E-11 |
| Fibroblast | Rbm38 | 8 | 9.761E-15 | 0.44573034 | 0.339 | 0.241 | 1.9522E-11 |
| Fibroblast | Tmem63c3 | 8 | 1.7006E-14 | 0.58080093 | 0.35 | 0.232 | 3.4012E-11 |
| Fibroblast | Il17rd2 | 8 | 3.394E-14 | 0.38784279 | 0.571 | 0.414 | 6.7881E-11 |
| Fibroblast | Tm4sf191 | 8 | 4.6061E-14 | 0.53516653 | 0.272 | 0.148 | 9.2122E-11 |
| Fibroblast | Slc37a23 | 8 | 4.6664E-14 | 0.27569037 | 0.438 | 0.307 | 9.3329E-11 |
| Fibroblast | Pde8b | 8 | 7.2722E-14 | 0.96079025 | 0.309 | 0.121 | 1.4544E-10 |
| Fibroblast | Kcnd32 | 8 | 7.6022E-14 | 0.30445879 | 0.362 | 0.26 | 1.5204E-10 |
| Fibroblast | Fap2 | 8 | 8.4622E-14 | 0.49224883 | 0.815 | 0.626 | 1.6924E-10 |
| Fibroblast | Lrrc4c1 | 8 | 1.1479E-13 | 0.27859852 | 0.307 | 0.232 | 2.2958E-10 |

| | | | | | | | |
|---|---|---|---|---|---|---|---|
| Fibroblast | Bace2 | 8 | 1.2264E-13 | 0.34274736 | 0.478 | 0.344 | 2.4528E-10 |
| Fibroblast | Thsd7b1 | 8 | 1.3862E-13 | 0.84264106 | 0.37 | 0.182 | 2.7723E-10 |
| Fibroblast | Adam122 | 8 | 1.3884E-13 | 0.53694008 | 0.587 | 0.429 | 2.7768E-10 |
| Fibroblast | Sh3kbp13 | 8 | 1.4951E-13 | 0.40903549 | 0.547 | 0.414 | 2.9903E-10 |
| Fibroblast | Prkcb2 | 8 | 1.5155E-13 | 0.29583695 | 0.375 | 0.273 | 3.031E-10 |
| Fibroblast | Myh71 | 8 | 2.3806E-13 | 0.45346087 | 0.737 | 0.545 | 4.7613E-10 |
| Fibroblast | Sdc22 | 8 | 3.0302E-13 | 0.40660711 | 0.707 | 0.524 | 6.0604E-10 |
| Fibroblast | AABR070449 | 8 | 3.0837E-13 | 0.33637361 | 0.434 | 0.308 | 6.1674E-10 |
| Fibroblast | Galnt174 | 8 | 3.2549E-13 | 0.43440689 | 0.811 | 0.66 | 6.5099E-10 |
| Fibroblast | Cyp7b13 | 8 | 3.4039E-13 | 0.34489816 | 0.615 | 0.455 | 6.8078E-10 |
| Fibroblast | Rgs172 | 8 | 3.531E-13 | 0.3931516 | 0.482 | 0.344 | 7.062E-10 |
| Fibroblast | Wipf3 | 8 | 3.8133E-13 | 0.4829942 | 0.423 | 0.296 | 7.6266E-10 |
| Fibroblast | Art31 | 8 | 3.9873E-13 | 0.4669668 | 0.43 | 0.314 | 7.9745E-10 |
| Fibroblast | Ptcd31 | 8 | 4.2239E-13 | 0.36246384 | 0.411 | 0.307 | 8.4478E-10 |
| Fibroblast | AABR070359 | 8 | 5.6216E-13 | 0.45841999 | 0.878 | 0.758 | 1.1243E-09 |
| Fibroblast | Ifitm3 | 8 | 6.8006E-13 | 0.37754268 | 0.488 | 0.329 | 1.3601E-09 |
| Fibroblast | Ror22 | 8 | 1.4302E-12 | 0.35239422 | 0.674 | 0.504 | 2.8605E-09 |
| Fibroblast | Ninj21 | 8 | 1.4313E-12 | 0.40875238 | 0.697 | 0.53 | 2.8627E-09 |
| Fibroblast | Kcnn32 | 8 | 1.5614E-12 | 0.28788245 | 0.674 | 0.532 | 3.1228E-09 |
| Fibroblast | Col27a13 | 8 | 1.5762E-12 | 0.34617343 | 0.585 | 0.423 | 3.1524E-09 |
| Fibroblast | Ccdc802 | 8 | 2.0782E-12 | 0.36468334 | 0.975 | 0.89 | 4.1563E-09 |
| Fibroblast | Myo18b1 | 8 | 2.0996E-12 | 0.72556014 | 0.364 | 0.266 | 4.1993E-09 |
| Fibroblast | Mitf1 | 8 | 2.3339E-12 | 0.31991293 | 0.739 | 0.56 | 4.6677E-09 |
| Fibroblast | Pparg3 | 8 | 2.3939E-12 | 0.27102109 | 0.36 | 0.272 | 4.7878E-09 |
| Fibroblast | AC130940.12 | 8 | 3.1046E-12 | 1.07706852 | 0.291 | 0.176 | 6.2092E-09 |
| Fibroblast | Cnksr2 | 8 | 3.9758E-12 | 0.38244044 | 0.309 | 0.187 | 7.9516E-09 |
| Fibroblast | Ckm1 | 8 | 4.8786E-12 | 0.3197703 | 0.453 | 0.339 | 9.7572E-09 |
| Fibroblast | Cd742 | 8 | 5.7725E-12 | 0.49592715 | 0.368 | 0.273 | 1.1545E-08 |
| Fibroblast | Sulf12 | 8 | 6.844E-12 | 0.27053419 | 0.81 | 0.613 | 1.3688E-08 |
| Fibroblast | Loxl22 | 8 | 1.0973E-11 | 0.26182182 | 0.686 | 0.513 | 2.1946E-08 |
| Fibroblast | Il1rapl21 | 8 | 1.2977E-11 | 0.92927389 | 0.28 | 0.236 | 2.5955E-08 |

| | | | | | | | |
|---|---|---|---|---|---|---|---|
| Fibroblast | Il1rapl11 | 8 | 1.6686E-11 | 0.26802912 | 0.324 | 0.262 | 3.3372E-08 |
| Fibroblast | Gna152 | 8 | 1.9779E-11 | 1.06016627 | 0.366 | 0.272 | 3.9558E-08 |
| Fibroblast | Adamts143 | 8 | 2.3749E-11 | 0.32082011 | 0.545 | 0.439 | 4.7499E-08 |
| Fibroblast | AC134204.12 | 8 | 3.9912E-11 | 0.31949343 | 0.672 | 0.509 | 7.9825E-08 |
| Fibroblast | Slc25a13 | 8 | 5.6517E-11 | 0.41650704 | 0.358 | 0.283 | 1.1303E-07 |
| Fibroblast | C1qtnf71 | 8 | 6.4802E-11 | 0.33675386 | 0.96 | 0.89 | 1.296E-07 |
| Fibroblast | Brca1 | 8 | 6.8106E-11 | 0.46019792 | 0.291 | 0.216 | 1.3621E-07 |
| Fibroblast | Rnf1503 | 8 | 1.1842E-10 | 0.46975943 | 0.396 | 0.329 | 2.3684E-07 |
| Fibroblast | Asb182 | 8 | 1.3548E-10 | 0.4711439 | 0.33 | 0.288 | 2.7095E-07 |
| Fibroblast | Tuft1 | 8 | 4.5388E-10 | 0.85606919 | 0.368 | 0.291 | 9.0777E-07 |
| Fibroblast | Stk261 | 8 | 5.1597E-10 | 0.49162773 | 0.284 | 0.139 | 1.0319E-06 |
| Fibroblast | Pgam21 | 8 | 6.9175E-10 | 0.29060585 | 0.4 | 0.317 | 1.3835E-06 |
| Fibroblast | Gas71 | 8 | 6.9226E-10 | 0.27360305 | 0.583 | 0.456 | 1.3845E-06 |
| Fibroblast | Lhfpl22 | 8 | 7.6274E-10 | 0.39962394 | 0.564 | 0.448 | 1.5255E-06 |
| Fibroblast | Arhgap244 | 8 | 7.6593E-10 | 0.40909161 | 0.794 | 0.638 | 1.5319E-06 |
| Fibroblast | Ptprj2 | 8 | 2.4193E-09 | 0.38299107 | 0.714 | 0.577 | 4.8386E-06 |
| Fibroblast | Il162 | 8 | 2.5849E-09 | 0.31880916 | 0.636 | 0.527 | 5.1699E-06 |
| Fibroblast | Agap21 | 8 | 2.651E-09 | 0.75046474 | 0.276 | 0.13 | 5.3019E-06 |
| Fibroblast | Mgp1 | 8 | 3.6533E-09 | 0.39561943 | 0.905 | 0.796 | 7.3066E-06 |
| Fibroblast | Rnls4 | 8 | 4.3951E-09 | 0.34790718 | 0.539 | 0.411 | 8.7902E-06 |
| Fibroblast | Calcrl1 | 8 | 5.0519E-09 | 0.43946418 | 0.665 | 0.604 | 1.0104E-05 |
| Fibroblast | Eda2r2 | 8 | 5.6229E-09 | 0.51366911 | 0.297 | 0.271 | 1.1246E-05 |
| Fibroblast | AABR070015 | 8 | 9.3239E-09 | 0.28409586 | 0.632 | 0.454 | 1.8648E-05 |
| Fibroblast | Mtus22 | 8 | 2.6306E-08 | 0.42031571 | 0.379 | 0.328 | 5.2612E-05 |
| Fibroblast | Plekha73 | 8 | 3.0184E-08 | 0.33447823 | 0.821 | 0.658 | 6.0368E-05 |
| Fibroblast | Prkar2b2 | 8 | 4.1137E-08 | 0.41800758 | 0.379 | 0.341 | 8.2274E-05 |
| Fibroblast | Grm1 | 8 | 4.3658E-08 | 0.41926654 | 0.257 | 0.103 | 8.7316E-05 |
| Fibroblast | Ston22 | 8 | 8.3349E-08 | 0.28638943 | 0.848 | 0.734 | 0.0001667 |
| Fibroblast | Dpysl33 | 8 | 1.4405E-07 | 0.29830435 | 0.874 | 0.739 | 0.0002881 |
| Fibroblast | Palm22 | 8 | 2.4003E-07 | 0.25598956 | 0.484 | 0.421 | 0.00048006 |
| Fibroblast | Gucy1a23 | 8 | 3.7301E-07 | 0.32858028 | 0.604 | 0.494 | 0.00074601 |

| | | | | | | | |
|---|---|---|---|---|---|---|---|
| Fibroblast | Il1r13 | 8 | 8.8586E-07 | 0.25497234 | 0.811 | 0.715 | 0.00177171 |
| Fibroblast | Egfr2 | 8 | 1.0186E-06 | 0.33900336 | 0.827 | 0.708 | 0.0020371 |
| Fibroblast | Frmd31 | 8 | 1.0914E-06 | 0.37540774 | 0.402 | 0.242 | 0.00218282 |
| Fibroblast | Cmahp | 8 | 1.7315E-06 | 1.58050417 | 0.251 | 0.094 | 0.00346301 |
| Fibroblast | Gask1b1 | 8 | 2.0584E-06 | 0.3528234 | 0.585 | 0.558 | 0.00411676 |
| Fibroblast | Antxr13 | 8 | 4.1537E-06 | 0.25183001 | 0.901 | 0.785 | 0.00830739 |
| Fibroblast | Pcdh193 | 8 | 4.7232E-06 | 0.38348633 | 0.379 | 0.376 | 0.00944635 |
| Fibroblast | Hsd11b13 | 8 | 7.4195E-06 | 0.39711956 | 0.345 | 0.349 | 0.0148391 |
| Fibroblast | Clcnka | 9 | 5.018E-133 | 3.77305835 | 0.341 | 0.051 | 1.004E-129 |
| Fibroblast | Cnn1 | 9 | 6.663E-130 | 3.80222385 | 0.328 | 0.042 | 1.333E-126 |
| Fibroblast | Tnn1 | 9 | 8.137E-125 | 5.20619543 | 0.637 | 0.18 | 1.627E-121 |
| Fibroblast | Specc13 | 9 | 9.625E-121 | 2.2314342 | 0.922 | 0.518 | 1.925E-117 |
| Fibroblast | Fn14 | 9 | 2.094E-113 | 2.36437173 | 0.951 | 0.615 | 4.188E-110 |
| Fibroblast | Podnl11 | 9 | 4.078E-108 | 3.23395903 | 0.477 | 0.099 | 8.157E-105 |
| Fibroblast | Hmcn13 | 9 | 4.08E-106 | 2.13440139 | 0.915 | 0.486 | 8.16E-103 |
| Fibroblast | Ccn22 | 9 | 7.104E-104 | 2.56216222 | 0.869 | 0.475 | 1.421E-100 |
| Fibroblast | Ltbp22 | 9 | 3.023E-102 | 2.04446111 | 0.915 | 0.459 | 6.045E-99 |
| Fibroblast | Gabrp | 9 | 2.339E-101 | 2.4441438 | 0.421 | 0.089 | 4.6788E-98 |
| Fibroblast | Il18r1 | 9 | 4.4013E-97 | 2.94497184 | 0.319 | 0.062 | 8.8027E-94 |
| Fibroblast | Rai143 | 9 | 3.5589E-92 | 2.06841686 | 0.864 | 0.47 | 7.1177E-89 |
| Fibroblast | Adamts20 | 9 | 8.1993E-89 | 1.58888101 | 0.421 | 0.059 | 1.6399E-85 |
| Fibroblast | Runx12 | 9 | 1.7441E-87 | 2.18243144 | 0.835 | 0.51 | 3.4883E-84 |
| Fibroblast | AABR070252 | 9 | 2.4423E-82 | 1.63379703 | 0.951 | 0.663 | 4.8846E-79 |
| Fibroblast | Col11a11 | 9 | 1.9007E-80 | 3.55123969 | 0.701 | 0.37 | 3.8013E-77 |
| Fibroblast | Nsg2 | 9 | 2.3258E-79 | 1.80423816 | 0.277 | 0.024 | 4.6517E-76 |
| Fibroblast | Armc4 | 9 | 2.1918E-75 | 0.70422654 | 0.273 | 0.032 | 4.3836E-72 |
| Fibroblast | Epha32 | 9 | 1.7214E-67 | 3.93596069 | 0.64 | 0.267 | 3.4427E-64 |
| Fibroblast | Kif26b2 | 9 | 6.2259E-64 | 2.583591 | 0.71 | 0.31 | 1.2452E-60 |
| Fibroblast | Col12a12 | 9 | 5.6444E-63 | 3.32698152 | 0.62 | 0.274 | 1.1289E-59 |
| Fibroblast | Vcl2 | 9 | 2.5426E-61 | 1.62444407 | 0.852 | 0.541 | 5.0851E-58 |
| Fibroblast | Robo21 | 9 | 5.713E-61 | 3.18190694 | 0.672 | 0.331 | 1.1426E-57 |

| | | | | | | | |
|---|---|---|---|---|---|---|---|
| Fibroblast | Tnc1 | 9 | 4.2021E-59 | 3.41400279 | 0.606 | 0.282 | 8.4041E-56 |
| Fibroblast | Pawr2 | 9 | 4.8869E-59 | 2.64649423 | 0.65 | 0.24 | 9.7738E-56 |
| Fibroblast | Thbs12 | 9 | 3.6582E-58 | 3.1816816 | 0.633 | 0.307 | 7.3164E-55 |
| Fibroblast | Flt4 | 9 | 8.1052E-57 | 1.6481888 | 0.511 | 0.135 | 1.621E-53 |
| Fibroblast | Hmga21 | 9 | 2.3608E-56 | 0.58632662 | 0.421 | 0.111 | 4.7216E-53 |
| Fibroblast | Comp1 | 9 | 1.1983E-55 | 2.19988753 | 0.411 | 0.128 | 2.3967E-52 |
| Fibroblast | Ano31 | 9 | 6.2897E-54 | 2.66184498 | 0.28 | 0.13 | 1.2579E-50 |
| Fibroblast | Ahr2 | 9 | 1.6637E-53 | 1.76245806 | 0.73 | 0.435 | 3.3273E-50 |
| Fibroblast | Actn12 | 9 | 5.3688E-53 | 1.63851354 | 0.805 | 0.541 | 1.0738E-49 |
| Fibroblast | Col24a12 | 9 | 6.5255E-53 | 1.98470407 | 0.708 | 0.36 | 1.3051E-49 |
| Fibroblast | Rgs7bp1 | 9 | 3.5438E-51 | 2.36216941 | 0.482 | 0.195 | 7.0876E-48 |
| Fibroblast | Tpm21 | 9 | 1.3399E-50 | 1.99094857 | 0.674 | 0.252 | 2.6799E-47 |
| Fibroblast | Gpr1762 | 9 | 7.7891E-50 | 2.57103957 | 0.618 | 0.344 | 1.5578E-46 |
| Fibroblast | Pappa12 | 9 | 1.3272E-49 | 2.20423034 | 0.725 | 0.419 | 2.6544E-46 |
| Fibroblast | Cald11 | 9 | 2.1906E-49 | 1.32593178 | 0.878 | 0.687 | 4.3812E-46 |
| Fibroblast | Col1a13 | 9 | 9.4401E-49 | 0.83511574 | 0.976 | 0.872 | 1.888E-45 |
| Fibroblast | Npnt2 | 9 | 1.8942E-48 | 1.77808914 | 0.455 | 0.195 | 3.7885E-45 |
| Fibroblast | Flna2 | 9 | 2.9674E-48 | 1.50518352 | 0.769 | 0.416 | 5.9347E-45 |
| Fibroblast | Olr11 | 9 | 3.2204E-48 | 4.07626222 | 0.333 | 0.092 | 6.4408E-45 |
| Fibroblast | Csrp12 | 9 | 1.0344E-47 | 1.7670652 | 0.727 | 0.354 | 2.0689E-44 |
| Fibroblast | Mrvi12 | 9 | 1.8921E-47 | 1.90260261 | 0.713 | 0.389 | 3.7842E-44 |
| Fibroblast | Pdgfd2 | 9 | 3.5566E-47 | 1.44760788 | 0.788 | 0.52 | 7.1131E-44 |
| Fibroblast | Nrxn12 | 9 | 5.0119E-47 | 1.36260991 | 0.844 | 0.557 | 1.0024E-43 |
| Fibroblast | Serpine12 | 9 | 2.0267E-46 | 3.03439315 | 0.63 | 0.336 | 4.0534E-43 |
| Fibroblast | Kank41 | 9 | 2.3395E-46 | 2.95857365 | 0.516 | 0.215 | 4.6789E-43 |
| Fibroblast | AABR070414 | 9 | 2.7049E-46 | 1.03000322 | 0.304 | 0.083 | 5.4098E-43 |
| Fibroblast | Lef11 | 9 | 4.5684E-46 | 0.87265355 | 0.338 | 0.117 | 9.1368E-43 |
| Fibroblast | Ccn52 | 9 | 5.5758E-46 | 1.76706894 | 0.703 | 0.415 | 1.1152E-42 |
| Fibroblast | Ptprv1 | 9 | 6.5307E-46 | 2.4773085 | 0.608 | 0.32 | 1.3061E-42 |
| Fibroblast | Kcnk2 | 9 | 2.5594E-45 | 3.27288943 | 0.53 | 0.193 | 5.1188E-42 |
| Fibroblast | Pls32 | 9 | 3.2882E-45 | 1.20402293 | 0.912 | 0.67 | 6.5764E-42 |

| | | | | | | | |
|---|---|---|---|---|---|---|---|
| Fibroblast | Nrp22 | 9 | 2.0309E-44 | 1.52958056 | 0.71 | 0.373 | 4.0618E-41 |
| Fibroblast | Anks1b1 | 9 | 2.9255E-44 | 4.28042314 | 0.557 | 0.219 | 5.851E-41 |
| Fibroblast | LOC1025530 | 9 | 2.9667E-44 | 2.11339797 | 0.655 | 0.362 | 5.9334E-41 |
| Fibroblast | Atp10b | 9 | 4.0359E-44 | 0.86930987 | 0.263 | 0.079 | 8.0718E-41 |
| Fibroblast | Fshr2 | 9 | 6.7855E-44 | 0.50397073 | 0.394 | 0.132 | 1.3571E-40 |
| Fibroblast | Piezo22 | 9 | 4.4449E-43 | 1.65123045 | 0.696 | 0.429 | 8.8899E-40 |
| Fibroblast | Hpgd2 | 9 | 2.1927E-42 | 2.69440142 | 0.53 | 0.26 | 4.3854E-39 |
| Fibroblast | P4ha32 | 9 | 2.6325E-42 | 2.68301072 | 0.457 | 0.215 | 5.265E-39 |
| Fibroblast | Sgcg1 | 9 | 4.0885E-42 | 2.82104031 | 0.582 | 0.304 | 8.1769E-39 |
| Fibroblast | Col8a11 | 9 | 1.1629E-41 | 1.13321416 | 0.961 | 0.87 | 2.3258E-38 |
| Fibroblast | Plod22 | 9 | 1.3391E-41 | 1.00795703 | 0.927 | 0.747 | 2.6782E-38 |
| Fibroblast | Lancl32 | 9 | 1.4397E-41 | 2.46461055 | 0.428 | 0.215 | 2.8795E-38 |
| Fibroblast | Tnfrsf12a1 | 9 | 2.2722E-41 | 1.86516695 | 0.501 | 0.185 | 4.5445E-38 |
| Fibroblast | Nav31 | 9 | 2.7283E-41 | 1.56475284 | 0.71 | 0.337 | 5.4566E-38 |
| Fibroblast | Fgf72 | 9 | 1.6022E-40 | 2.53946285 | 0.377 | 0.156 | 3.2045E-37 |
| Fibroblast | Cdkn2b2 | 9 | 1.7234E-40 | 3.27714555 | 0.358 | 0.165 | 3.4468E-37 |
| Fibroblast | Sntg22 | 9 | 2.0772E-39 | 1.9333015 | 0.633 | 0.314 | 4.1544E-36 |
| Fibroblast | Cemip3 | 9 | 2.765E-39 | 1.80258079 | 0.686 | 0.34 | 5.5299E-36 |
| Fibroblast | Il1rl21 | 9 | 4.2512E-39 | 2.74099133 | 0.535 | 0.303 | 8.5023E-36 |
| Fibroblast | Cdh3 | 9 | 1.0159E-38 | 2.55282876 | 0.273 | 0.084 | 2.0318E-35 |
| Fibroblast | Nrg11 | 9 | 3.0579E-38 | 2.04911811 | 0.628 | 0.339 | 6.1158E-35 |
| Fibroblast | Tnfrsf11b2 | 9 | 4.1695E-38 | 2.01033458 | 0.664 | 0.395 | 8.3391E-35 |
| Fibroblast | Prss121 | 9 | 5.4514E-37 | 2.33357089 | 0.411 | 0.141 | 1.0903E-33 |
| Fibroblast | Itga84 | 9 | 9.8924E-37 | 1.1540941 | 0.808 | 0.591 | 1.9785E-33 |
| Fibroblast | Melk | 9 | 1.5964E-36 | 0.5067542 | 0.285 | 0.022 | 3.1928E-33 |
| Fibroblast | Stac | 9 | 1.6384E-36 | 1.02555455 | 0.353 | 0.094 | 3.2767E-33 |
| Fibroblast | Stk262 | 9 | 6.4207E-36 | 0.26779464 | 0.38 | 0.137 | 1.2841E-32 |
| Fibroblast | Ccn42 | 9 | 1.0189E-35 | 2.3292706 | 0.521 | 0.244 | 2.0377E-32 |
| Fibroblast | AC118957.12 | 9 | 2.1583E-35 | 0.88362925 | 0.533 | 0.238 | 4.3165E-32 |
| Fibroblast | Spon11 | 9 | 2.1263E-34 | 1.51085585 | 0.73 | 0.531 | 4.2527E-31 |
| Fibroblast | AABR070294 | 9 | 2.4571E-34 | 0.91814819 | 0.324 | 0.111 | 4.9142E-31 |

| | | | | | | | |
|---|---|---|---|---|---|---|---|
| Fibroblast | Col5a21 | 9 | 1.4616E-33 | 0.787456 | 0.942 | 0.816 | 2.9232E-30 |
| Fibroblast | Efna53 | 9 | 3.811E-33 | 1.91011776 | 0.569 | 0.371 | 7.622E-30 |
| Fibroblast | Mgmt3 | 9 | 4.6268E-33 | 1.31527349 | 0.735 | 0.466 | 9.2535E-30 |
| Fibroblast | Tpm11 | 9 | 1.3541E-32 | 0.95308471 | 0.895 | 0.754 | 2.7082E-29 |
| Fibroblast | Smoc11 | 9 | 9.1944E-32 | 1.24922434 | 0.389 | 0.188 | 1.8389E-28 |
| Fibroblast | Cntn22 | 9 | 2.218E-31 | 0.7678918 | 0.404 | 0.2 | 4.436E-28 |
| Fibroblast | Csrp22 | 9 | 2.8289E-31 | 2.48763635 | 0.564 | 0.32 | 5.6578E-28 |
| Fibroblast | Itgb82 | 9 | 1.2326E-30 | 1.36519278 | 0.669 | 0.44 | 2.4651E-27 |
| Fibroblast | Il13ra2 | 9 | 1.8543E-30 | 2.60975746 | 0.345 | 0.101 | 3.7087E-27 |
| Fibroblast | Cacna1c1 | 9 | 3.1159E-30 | 0.95913138 | 0.803 | 0.516 | 6.2318E-27 |
| Fibroblast | Ak52 | 9 | 8.8212E-30 | 1.78012435 | 0.528 | 0.284 | 1.7642E-26 |
| Fibroblast | Bnc23 | 9 | 1.7255E-29 | 0.75352493 | 0.944 | 0.738 | 3.451E-26 |
| Fibroblast | Myo10 | 9 | 3.4379E-29 | 1.28846988 | 0.599 | 0.361 | 6.8758E-26 |
| Fibroblast | Plxdc23 | 9 | 3.968E-29 | 0.6794514 | 0.964 | 0.813 | 7.9361E-26 |
| Fibroblast | Angpt4 | 9 | 1.3189E-28 | 2.66770061 | 0.282 | 0.062 | 2.6377E-25 |
| Fibroblast | Pde5a1 | 9 | 1.4882E-28 | 1.26656529 | 0.684 | 0.436 | 2.9765E-25 |
| Fibroblast | AABR070215 | 9 | 1.7038E-28 | 1.80193976 | 0.562 | 0.345 | 3.4076E-25 |
| Fibroblast | Olfml2b3 | 9 | 3.0235E-28 | 1.63789772 | 0.584 | 0.404 | 6.0469E-25 |
| Fibroblast | Myh102 | 9 | 4.8297E-28 | 1.09305423 | 0.752 | 0.539 | 9.6595E-25 |
| Fibroblast | Adamts161 | 9 | 5.3542E-28 | 2.1535575 | 0.501 | 0.277 | 1.0708E-24 |
| Fibroblast | Lbh1 | 9 | 5.4346E-28 | 1.14444882 | 0.72 | 0.488 | 1.0869E-24 |
| Fibroblast | Tagln | 9 | 6.3274E-28 | 2.40180775 | 0.506 | 0.215 | 1.2655E-24 |
| Fibroblast | Filip1l3 | 9 | 6.7441E-28 | 0.84438581 | 0.839 | 0.623 | 1.3488E-24 |
| Fibroblast | Slc24a33 | 9 | 8.7711E-28 | 1.53131488 | 0.628 | 0.398 | 1.7542E-24 |
| Fibroblast | Slit21 | 9 | 8.8026E-28 | 1.15882286 | 0.686 | 0.384 | 1.7605E-24 |
| Fibroblast | Kcnq52 | 9 | 2.6125E-27 | 1.97655539 | 0.513 | 0.212 | 5.225E-24 |
| Fibroblast | Ddah12 | 9 | 4.3355E-27 | 1.91739076 | 0.579 | 0.388 | 8.6709E-24 |
| Fibroblast | Nox42 | 9 | 5.7659E-27 | 1.34653733 | 0.684 | 0.522 | 1.1532E-23 |
| Fibroblast | Rab151 | 9 | 7.9427E-27 | 1.71309262 | 0.355 | 0.166 | 1.5885E-23 |
| Fibroblast | AABR070655 | 9 | 8.7253E-27 | 1.56620218 | 0.265 | 0.199 | 1.7451E-23 |
| Fibroblast | Nxn1 | 9 | 9.8842E-27 | 1.11858366 | 0.776 | 0.563 | 1.9768E-23 |

| | | | | | | | |
|---|---|---|---|---|---|---|---|
| Fibroblast | Zbtb7c3 | 9 | 1.2358E-26 | 1.7216451 | 0.589 | 0.325 | 2.4716E-23 |
| Fibroblast | Matk | 9 | 1.6684E-26 | 0.4236182 | 0.28 | 0.118 | 3.3369E-23 |
| Fibroblast | Pdlim32 | 9 | 2.287E-26 | 1.42980987 | 0.703 | 0.478 | 4.574E-23 |
| Fibroblast | Gcnt2 | 9 | 2.5615E-26 | 1.7100684 | 0.499 | 0.183 | 5.1231E-23 |
| Fibroblast | Jph21 | 9 | 3.0521E-26 | 1.86937388 | 0.564 | 0.281 | 6.1042E-23 |
| Fibroblast | Sorcs21 | 9 | 3.3628E-26 | 1.78028123 | 0.482 | 0.192 | 6.7255E-23 |
| Fibroblast | Lmod12 | 9 | 3.8409E-26 | 1.2124952 | 0.618 | 0.343 | 7.6818E-23 |
| Fibroblast | Flrt23 | 9 | 4.6887E-26 | 1.15674969 | 0.701 | 0.505 | 9.3775E-23 |
| Fibroblast | Dgkg1 | 9 | 5.2919E-26 | 1.60333149 | 0.314 | 0.14 | 1.0584E-22 |
| Fibroblast | Grb141 | 9 | 5.6967E-26 | 1.30073139 | 0.681 | 0.41 | 1.1393E-22 |
| Fibroblast | Ccn31 | 9 | 6.7727E-26 | 1.42863258 | 0.569 | 0.259 | 1.3545E-22 |
| Fibroblast | Svep12 | 9 | 1.1398E-25 | 1.19825073 | 0.672 | 0.469 | 2.2797E-22 |
| Fibroblast | Alk1 | 9 | 1.2332E-25 | 2.15445693 | 0.448 | 0.175 | 2.4664E-22 |
| Fibroblast | Angpt11 | 9 | 1.6727E-25 | 0.98925145 | 0.745 | 0.538 | 3.3453E-22 |
| Fibroblast | Col16a12 | 9 | 2.5902E-25 | 1.10956791 | 0.662 | 0.421 | 5.1805E-22 |
| Fibroblast | AABR070042 | 9 | 5.5038E-25 | 0.66893423 | 0.453 | 0.143 | 1.1008E-21 |
| Fibroblast | Il1rl1 | 9 | 9.3898E-25 | 1.84296778 | 0.36 | 0.094 | 1.878E-21 |
| Fibroblast | Pdgfc1 | 9 | 1.0882E-24 | 3.0585648 | 0.453 | 0.268 | 2.1764E-21 |
| Fibroblast | Kcnip1 | 9 | 1.5657E-24 | 2.72316615 | 0.328 | 0.116 | 3.1313E-21 |
| Fibroblast | Fign2 | 9 | 2.1462E-24 | 1.1069012 | 0.633 | 0.352 | 4.2924E-21 |
| Fibroblast | Kcnn4 | 9 | 2.7002E-24 | 0.85620237 | 0.35 | 0.129 | 5.4004E-21 |
| Fibroblast | Cpxm22 | 9 | 2.9714E-24 | 1.6363943 | 0.533 | 0.233 | 5.9429E-21 |
| Fibroblast | Fmn11 | 9 | 3.0878E-24 | 1.45851077 | 0.599 | 0.367 | 6.1756E-21 |
| Fibroblast | Dnm31 | 9 | 3.5996E-24 | 1.59208377 | 0.669 | 0.487 | 7.1992E-21 |
| Fibroblast | Nuak11 | 9 | 3.9935E-24 | 1.87394985 | 0.535 | 0.397 | 7.9871E-21 |
| Fibroblast | Sgcd1 | 9 | 6.538E-24 | 1.13677324 | 0.684 | 0.46 | 1.3076E-20 |
| Fibroblast | Rspo22 | 9 | 7.1437E-24 | 1.99631977 | 0.326 | 0.106 | 1.4287E-20 |
| Fibroblast | Prrx14 | 9 | 8.6586E-24 | 0.66637275 | 0.888 | 0.678 | 1.7317E-20 |
| Fibroblast | Runx22 | 9 | 9.2415E-24 | 1.10454983 | 0.679 | 0.503 | 1.8483E-20 |
| Fibroblast | Aebp11 | 9 | 1.1449E-23 | 1.0387471 | 0.698 | 0.465 | 2.2897E-20 |
| Fibroblast | Trabd2b2 | 9 | 2.8106E-23 | 2.29218901 | 0.477 | 0.229 | 5.6212E-20 |

| | | | | | | | |
|---|---|---|---|---|---|---|---|
| Fibroblast | Pcdha133 | 9 | 3.1619E-23 | 0.80429334 | 0.827 | 0.621 | 6.3239E-20 |
| Fibroblast | Agtr1a2 | 9 | 4.1519E-23 | 1.21460592 | 0.633 | 0.423 | 8.3037E-20 |
| Fibroblast | Ca31 | 9 | 5.2121E-23 | 0.40862992 | 0.29 | 0.145 | 1.0424E-19 |
| Fibroblast | Cdkn1a2 | 9 | 7.0909E-23 | 1.76772423 | 0.528 | 0.336 | 1.4182E-19 |
| Fibroblast | Slco2a1 | 9 | 1.2134E-22 | 2.09899835 | 0.302 | 0.103 | 2.4268E-19 |
| Fibroblast | Ptafr3 | 9 | 1.4747E-22 | 1.32452981 | 0.499 | 0.291 | 2.9494E-19 |
| Fibroblast | Cenpe1 | 9 | 2.0952E-22 | 0.59372904 | 0.414 | 0.153 | 4.1904E-19 |
| Fibroblast | Map3k7cl1 | 9 | 2.8689E-22 | 1.82954691 | 0.304 | 0.155 | 5.7378E-19 |
| Fibroblast | Adam123 | 9 | 2.966E-22 | 1.23018867 | 0.655 | 0.428 | 5.932E-19 |
| Fibroblast | Postn4 | 9 | 2.9847E-22 | 0.79693007 | 0.883 | 0.771 | 5.9694E-19 |
| Fibroblast | Cdh112 | 9 | 5.8282E-22 | 0.74717042 | 0.856 | 0.683 | 1.1656E-18 |
| Fibroblast | Rnd32 | 9 | 6.5476E-22 | 0.98141525 | 0.737 | 0.531 | 1.3095E-18 |
| Fibroblast | Abcb4 | 9 | 1.254E-21 | 1.77387617 | 0.528 | 0.314 | 2.508E-18 |
| Fibroblast | Bgn1 | 9 | 1.7812E-21 | 0.81724485 | 0.693 | 0.456 | 3.5623E-18 |
| Fibroblast | Inhba3 | 9 | 2.0555E-21 | 1.47584806 | 0.511 | 0.25 | 4.1111E-18 |
| Fibroblast | Gria32 | 9 | 2.6449E-21 | 0.69916542 | 0.813 | 0.606 | 5.2897E-18 |
| Fibroblast | Antxr14 | 9 | 3.4863E-21 | 0.65565455 | 0.888 | 0.787 | 6.9726E-18 |
| Fibroblast | Sdc23 | 9 | 3.8953E-21 | 1.1347205 | 0.727 | 0.525 | 7.7906E-18 |
| Fibroblast | Slc10a62 | 9 | 4.2386E-21 | 1.29211924 | 0.382 | 0.187 | 8.4772E-18 |
| Fibroblast | Col27a14 | 9 | 4.3789E-21 | 0.99328473 | 0.633 | 0.423 | 8.7577E-18 |
| Fibroblast | Cthrc12 | 9 | 5.6058E-21 | 1.30393235 | 0.569 | 0.319 | 1.1212E-17 |
| Fibroblast | Vcam12 | 9 | 8.9023E-21 | 1.61560061 | 0.533 | 0.307 | 1.7805E-17 |
| Fibroblast | Reln | 9 | 3.217E-20 | 2.60231311 | 0.428 | 0.186 | 6.434E-17 |
| Fibroblast | Crim11 | 9 | 4.4645E-20 | 1.15123036 | 0.586 | 0.371 | 8.9291E-17 |
| Fibroblast | Myl91 | 9 | 5.5314E-20 | 1.12007648 | 0.389 | 0.168 | 1.1063E-16 |
| Fibroblast | Mylk4 | 9 | 6.9969E-20 | 0.6588202 | 0.793 | 0.562 | 1.3994E-16 |
| Fibroblast | Pdlim51 | 9 | 7.3579E-20 | 0.7855092 | 0.852 | 0.666 | 1.4716E-16 |
| Fibroblast | Fhl21 | 9 | 1.4255E-19 | 1.12460625 | 0.618 | 0.429 | 2.851E-16 |
| Fibroblast | Sv2c2 | 9 | 2.0824E-19 | 0.91770146 | 0.659 | 0.5 | 4.1648E-16 |
| Fibroblast | Adamts144 | 9 | 2.6114E-19 | 0.65874219 | 0.652 | 0.436 | 5.2228E-16 |
| Fibroblast | Fbxl21 | 9 | 4.3837E-19 | 1.66237892 | 0.341 | 0.14 | 8.7674E-16 |

| | | | | | | | |
|---|---|---|---|---|---|---|---|
| Fibroblast | Acan1 | 9 | 4.8419E-19 | 4.55458051 | 0.253 | 0.096 | 9.6837E-16 |
| Fibroblast | Camk1d2 | 9 | 6.6209E-19 | 1.22591355 | 0.545 | 0.279 | 1.3242E-15 |
| Fibroblast | Adamts63 | 9 | 8.4946E-19 | 0.76302071 | 0.798 | 0.629 | 1.6989E-15 |
| Fibroblast | Mob3b1 | 9 | 2.8243E-18 | 1.27008293 | 0.484 | 0.266 | 5.6486E-15 |
| Fibroblast | C1qtnf33 | 9 | 4.2437E-18 | 1.0092942 | 0.436 | 0.234 | 8.4873E-15 |
| Fibroblast | Katnal21 | 9 | 9.149E-18 | 2.23435534 | 0.328 | 0.161 | 1.8298E-14 |
| Fibroblast | Vegfc2 | 9 | 9.6238E-18 | 0.75968823 | 0.616 | 0.382 | 1.9248E-14 |
| Fibroblast | Gdf61 | 9 | 9.7393E-18 | 1.26922205 | 0.436 | 0.303 | 1.9479E-14 |
| Fibroblast | Gpc62 | 9 | 1.767E-17 | 0.42528401 | 0.99 | 0.979 | 3.5339E-14 |
| Fibroblast | Dpysl34 | 9 | 4.5481E-17 | 0.50473288 | 0.905 | 0.739 | 9.0962E-14 |
| Fibroblast | Enpp13 | 9 | 5.636E-17 | 1.0200819 | 0.703 | 0.533 | 1.1272E-13 |
| Fibroblast | Enah4 | 9 | 5.673E-17 | 0.61031152 | 0.701 | 0.458 | 1.1346E-13 |
| Fibroblast | Rtn4rl12 | 9 | 4.572E-16 | 0.50105251 | 0.423 | 0.166 | 9.1441E-13 |
| Fibroblast | AABR070490 | 9 | 4.8768E-16 | 0.98671333 | 0.611 | 0.403 | 9.7536E-13 |
| Fibroblast | Mid12 | 9 | 4.9317E-16 | 0.62275009 | 0.774 | 0.587 | 9.8633E-13 |
| Fibroblast | Lyz2 | 9 | 6.8973E-16 | 1.09119241 | 0.455 | 0.2 | 1.3795E-12 |
| Fibroblast | Fam180a1 | 9 | 1.7694E-15 | 1.70763181 | 0.443 | 0.279 | 3.5388E-12 |
| Fibroblast | Slc1a32 | 9 | 3.0834E-15 | 0.58808441 | 0.53 | 0.269 | 6.1669E-12 |
| Fibroblast | AABR070067 | 9 | 3.4003E-15 | 2.34846987 | 0.392 | 0.194 | 6.8005E-12 |
| Fibroblast | Palmd1 | 9 | 3.7999E-15 | 1.56841071 | 0.479 | 0.308 | 7.5999E-12 |
| Fibroblast | Nr1h42 | 9 | 5.4447E-15 | 0.36239565 | 0.533 | 0.351 | 1.0889E-11 |
| Fibroblast | Cacna1d2 | 9 | 5.7554E-15 | 0.76433316 | 0.358 | 0.168 | 1.1511E-11 |
| Fibroblast | Asb151 | 9 | 7.6846E-15 | 1.05618408 | 0.443 | 0.265 | 1.5369E-11 |
| Fibroblast | Pdzrn33 | 9 | 9.2593E-15 | 0.46087258 | 0.983 | 0.915 | 1.8519E-11 |
| Fibroblast | Lum | 9 | 1.5152E-14 | 0.87206284 | 0.655 | 0.449 | 3.0304E-11 |
| Fibroblast | AABR070581 | 9 | 1.9661E-14 | 0.78115572 | 0.701 | 0.538 | 3.9322E-11 |
| Fibroblast | Klhl292 | 9 | 2.181E-14 | 1.02963694 | 0.599 | 0.438 | 4.362E-11 |
| Fibroblast | Ralgapa21 | 9 | 3.1229E-14 | 0.73636895 | 0.686 | 0.471 | 6.2459E-11 |
| Fibroblast | Ror23 | 9 | 4.1302E-14 | 0.90141062 | 0.642 | 0.507 | 8.2604E-11 |
| Fibroblast | Ncald2 | 9 | 4.374E-14 | 1.08766228 | 0.56 | 0.389 | 8.748E-11 |
| Fibroblast | Bcar31 | 9 | 5.7313E-14 | 2.02776463 | 0.397 | 0.241 | 1.1463E-10 |

| | | | | | | | |
|---|---|---|---|---|---|---|---|
| Fibroblast | Cyp26b12 | 9 | 6.532E-14 | 1.01612269 | 0.382 | 0.216 | 1.3064E-10 |
| Fibroblast | Ctnnd22 | 9 | 9.5508E-14 | 1.06981237 | 0.358 | 0.15 | 1.9102E-10 |
| Fibroblast | Myl6 | 9 | 9.7474E-14 | 0.54378749 | 0.691 | 0.425 | 1.9495E-10 |
| Fibroblast | Bcat12 | 9 | 1.2595E-13 | 0.69771509 | 0.669 | 0.521 | 2.5191E-10 |
| Fibroblast | Srpx2 | 9 | 1.4681E-13 | 0.5643056 | 0.572 | 0.346 | 2.9362E-10 |
| Fibroblast | Ccdc1411 | 9 | 1.8284E-13 | 0.55057499 | 0.669 | 0.408 | 3.6569E-10 |
| Fibroblast | Rasl12 | 9 | 1.9247E-13 | 1.5031815 | 0.363 | 0.202 | 3.8494E-10 |
| Fibroblast | Brca11 | 9 | 1.9298E-13 | 0.46553624 | 0.46 | 0.209 | 3.8596E-10 |
| Fibroblast | Ncam13 | 9 | 3.8601E-13 | 0.83457172 | 0.647 | 0.498 | 7.7203E-10 |
| Fibroblast | Gpd11 | 9 | 3.8823E-13 | 1.07496695 | 0.251 | 0.154 | 7.7646E-10 |
| Fibroblast | Osr12 | 9 | 4.7296E-13 | 0.42841781 | 0.547 | 0.397 | 9.4592E-10 |
| Fibroblast | Ano11 | 9 | 5.1035E-13 | 2.01774803 | 0.333 | 0.148 | 1.0207E-09 |
| Fibroblast | Tyrobp2 | 9 | 5.6578E-13 | 0.5762716 | 0.375 | 0.206 | 1.1316E-09 |
| Fibroblast | Spp12 | 9 | 5.7578E-13 | 1.51559214 | 0.426 | 0.25 | 1.1516E-09 |
| Fibroblast | Prdm62 | 9 | 7.0702E-13 | 0.27888013 | 0.618 | 0.43 | 1.414E-09 |
| Fibroblast | Cers62 | 9 | 9.6064E-13 | 0.60416192 | 0.659 | 0.518 | 1.9213E-09 |
| Fibroblast | Fhl11 | 9 | 1.2154E-12 | 0.64712873 | 0.676 | 0.543 | 2.4308E-09 |
| Fibroblast | Pak32 | 9 | 1.3553E-12 | 1.29972142 | 0.394 | 0.272 | 2.7107E-09 |
| Fibroblast | Olfm21 | 9 | 1.4158E-12 | 1.88669506 | 0.436 | 0.303 | 2.8316E-09 |
| Fibroblast | Col18a11 | 9 | 1.7173E-12 | 1.05273216 | 0.453 | 0.251 | 3.4345E-09 |
| Fibroblast | Tll1 | 9 | 1.7398E-12 | 0.26770347 | 0.345 | 0.093 | 3.4795E-09 |
| Fibroblast | Lpcat21 | 9 | 2.2352E-12 | 1.05938196 | 0.394 | 0.229 | 4.4705E-09 |
| Fibroblast | Kntc11 | 9 | 3.0807E-12 | 0.81226766 | 0.292 | 0.18 | 6.1613E-09 |
| Fibroblast | Hmox12 | 9 | 3.6131E-12 | 1.18105525 | 0.365 | 0.176 | 7.2262E-09 |
| Fibroblast | Col3a11 | 9 | 3.6924E-12 | 0.33915946 | 0.988 | 0.967 | 7.3848E-09 |
| Fibroblast | Tmem512 | 9 | 3.7023E-12 | 0.29499428 | 0.406 | 0.249 | 7.4045E-09 |
| Fibroblast | AABR070339 | 9 | 4.9057E-12 | 0.74825973 | 0.501 | 0.33 | 9.8115E-09 |
| Fibroblast | Galnt13 | 9 | 5.0773E-12 | 0.93777606 | 0.341 | 0.196 | 1.0155E-08 |
| Fibroblast | Slc8a13 | 9 | 5.7005E-12 | 0.44767676 | 0.942 | 0.886 | 1.1401E-08 |
| Fibroblast | Cit2 | 9 | 8.9141E-12 | 0.52684483 | 0.528 | 0.374 | 1.7828E-08 |
| Fibroblast | Eda2r3 | 9 | 8.9892E-12 | 0.6264205 | 0.47 | 0.264 | 1.7978E-08 |

| | | | | | | | |
|---|---|---|---|---|---|---|---|
| Fibroblast | Vegfd2 | 9 | 9.5608E-12 | 0.84893991 | 0.586 | 0.417 | 1.9122E-08 |
| Fibroblast | Hs6st24 | 9 | 9.6806E-12 | 0.55243691 | 0.591 | 0.424 | 1.9361E-08 |
| Fibroblast | Svil3 | 9 | 9.7061E-12 | 0.50578161 | 0.783 | 0.64 | 1.9412E-08 |
| Fibroblast | Vgll32 | 9 | 9.7083E-12 | 0.83796306 | 0.547 | 0.407 | 1.9417E-08 |
| Fibroblast | Hecw22 | 9 | 1.1642E-11 | 0.43723796 | 0.586 | 0.376 | 2.3285E-08 |
| Fibroblast | Rap1gap21 | 9 | 2.2609E-11 | 0.88848784 | 0.44 | 0.253 | 4.5218E-08 |
| Fibroblast | Chsy33 | 9 | 2.8096E-11 | 0.62407849 | 0.723 | 0.535 | 5.6191E-08 |
| Fibroblast | Map1b4 | 9 | 2.8186E-11 | 0.50040569 | 0.903 | 0.794 | 5.6372E-08 |
| Fibroblast | Polq1 | 9 | 3.3365E-11 | 0.87338911 | 0.258 | 0.161 | 6.6729E-08 |
| Fibroblast | Nkd21 | 9 | 3.8373E-11 | 2.17161384 | 0.401 | 0.314 | 7.6747E-08 |
| Fibroblast | Crlf11 | 9 | 3.8725E-11 | 1.12247291 | 0.46 | 0.249 | 7.745E-08 |
| Fibroblast | Gpx1 | 9 | 4.4308E-11 | 0.46785055 | 0.411 | 0.2 | 8.8615E-08 |
| Fibroblast | Fos2 | 9 | 4.6045E-11 | 0.95863763 | 0.416 | 0.265 | 9.2089E-08 |
| Fibroblast | Kif20a2 | 9 | 8.1368E-11 | 0.75982187 | 0.365 | 0.244 | 1.6274E-07 |
| Fibroblast | Rgcc1 | 9 | 8.401E-11 | 1.46682577 | 0.411 | 0.246 | 1.6802E-07 |
| Fibroblast | Prox11 | 9 | 9.3123E-11 | 0.82910244 | 0.409 | 0.208 | 1.8625E-07 |
| Fibroblast | Adamts173 | 9 | 1.3255E-10 | 0.368406 | 0.713 | 0.541 | 2.6509E-07 |
| Fibroblast | Glis33 | 9 | 1.5315E-10 | 0.33495392 | 0.9 | 0.765 | 3.0631E-07 |
| Fibroblast | Sfrp42 | 9 | 1.6379E-10 | 0.8446147 | 0.443 | 0.326 | 3.2757E-07 |
| Fibroblast | Egr13 | 9 | 1.8446E-10 | 0.4413934 | 0.443 | 0.257 | 3.6893E-07 |
| Fibroblast | Acta2 | 9 | 1.9942E-10 | 2.30075163 | 0.401 | 0.27 | 3.9885E-07 |
| Fibroblast | Setbp13 | 9 | 2.0436E-10 | 0.29853048 | 0.745 | 0.558 | 4.0871E-07 |
| Fibroblast | Pde4b1 | 9 | 3.1579E-10 | 0.30407449 | 0.633 | 0.417 | 6.3158E-07 |
| Fibroblast | Ifitm10 | 9 | 3.2105E-10 | 0.6341358 | 0.326 | 0.183 | 6.4211E-07 |
| Fibroblast | Ndst33 | 9 | 3.7347E-10 | 1.02107629 | 0.479 | 0.365 | 7.4695E-07 |
| Fibroblast | Ifitm31 | 9 | 3.8096E-10 | 0.35501817 | 0.516 | 0.33 | 7.6193E-07 |
| Fibroblast | Oprm12 | 9 | 4.3941E-10 | 0.41960029 | 0.265 | 0.258 | 8.7882E-07 |
| Fibroblast | Cemip23 | 9 | 5.2416E-10 | 0.76043746 | 0.606 | 0.495 | 1.0483E-06 |
| Fibroblast | Cd442 | 9 | 5.8082E-10 | 0.87378056 | 0.509 | 0.391 | 1.1616E-06 |
| Fibroblast | Ccn1 | 9 | 7.5566E-10 | 1.70304665 | 0.382 | 0.275 | 1.5113E-06 |
| Fibroblast | Sema3d2 | 9 | 7.8848E-10 | 0.8293333 | 0.543 | 0.39 | 1.577E-06 |

| | | | | | | | |
|---|---|---|---|---|---|---|---|
| Fibroblast | Slit34 | 9 | 9.2241E-10 | 1.05463432 | 0.533 | 0.479 | 1.8448E-06 |
| Fibroblast | Mtss11 | 9 | 9.629E-10 | 0.61845899 | 0.552 | 0.376 | 1.9258E-06 |
| Fibroblast | Procr | 9 | 1.1666E-09 | 0.7145951 | 0.282 | 0.125 | 2.3331E-06 |
| Fibroblast | Sntb12 | 9 | 1.2163E-09 | 1.39628524 | 0.411 | 0.295 | 2.4327E-06 |
| Fibroblast | Tmem1963 | 9 | 1.3897E-09 | 0.68854042 | 0.579 | 0.402 | 2.7794E-06 |
| Fibroblast | Nexn2 | 9 | 1.4925E-09 | 0.52172786 | 0.623 | 0.449 | 2.985E-06 |
| Fibroblast | Cobl3 | 9 | 1.5166E-09 | 0.70390173 | 0.409 | 0.258 | 3.0331E-06 |
| Fibroblast | Mmp164 | 9 | 1.8347E-09 | 0.81546657 | 0.428 | 0.306 | 3.6695E-06 |
| Fibroblast | Rxfp12 | 9 | 1.94E-09 | 0.28770606 | 0.302 | 0.145 | 3.8801E-06 |
| Fibroblast | Mfge8 | 9 | 2.2251E-09 | 0.64529144 | 0.375 | 0.224 | 4.4503E-06 |
| Fibroblast | Cp1 | 9 | 2.6012E-09 | 0.88585596 | 0.436 | 0.302 | 5.2025E-06 |
| Fibroblast | Cpq3 | 9 | 3.5219E-09 | 0.43997862 | 0.864 | 0.752 | 7.0438E-06 |
| Fibroblast | Ccdc3 | 9 | 3.7645E-09 | 1.18114839 | 0.307 | 0.144 | 7.5291E-06 |
| Fibroblast | Cep55 | 9 | 4.2108E-09 | 0.63528028 | 0.268 | 0.035 | 8.4216E-06 |
| Fibroblast | Pkhd1l1 | 9 | 4.2595E-09 | 0.3272201 | 0.336 | 0.146 | 8.519E-06 |
| Fibroblast | Mitf2 | 9 | 5.3894E-09 | 0.26530841 | 0.725 | 0.563 | 1.0779E-05 |
| Fibroblast | Fbln52 | 9 | 5.9476E-09 | 0.26589934 | 0.657 | 0.453 | 1.1895E-05 |
| Fibroblast | Aqp11 | 9 | 6.0888E-09 | 0.39997012 | 0.307 | 0.151 | 1.2178E-05 |
| Fibroblast | Sphkap1 | 9 | 8.1631E-09 | 1.92144592 | 0.375 | 0.269 | 1.6326E-05 |
| Fibroblast | AABR070651 | 9 | 8.2026E-09 | 1.94093595 | 0.345 | 0.128 | 1.6405E-05 |
| Fibroblast | Slc7a23 | 9 | 8.5319E-09 | 0.61665526 | 0.54 | 0.385 | 1.7064E-05 |
| Fibroblast | Me11 | 9 | 8.5831E-09 | 0.48172483 | 0.506 | 0.386 | 1.7166E-05 |
| Fibroblast | Med12l3 | 9 | 1.0086E-08 | 0.34874643 | 0.46 | 0.295 | 2.0171E-05 |
| Fibroblast | Dgkz | 9 | 3.8312E-08 | 0.34909418 | 0.433 | 0.28 | 7.6625E-05 |
| Fibroblast | Lrrc71 | 9 | 3.8427E-08 | 0.39500111 | 0.372 | 0.196 | 7.6855E-05 |
| Fibroblast | Zeb2 | 9 | 4.1489E-08 | 0.34556057 | 0.934 | 0.883 | 8.2978E-05 |
| Fibroblast | Sdk21 | 9 | 4.3869E-08 | 0.33761128 | 0.689 | 0.575 | 8.7737E-05 |
| Fibroblast | Sparc1 | 9 | 4.6304E-08 | 0.35249825 | 0.856 | 0.736 | 9.2609E-05 |
| Fibroblast | Trpc31 | 9 | 5.8773E-08 | 0.79098877 | 0.406 | 0.258 | 0.00011755 |
| Fibroblast | Myh11 | 9 | 7.4875E-08 | 1.64917932 | 0.367 | 0.281 | 0.00014975 |
| Fibroblast | Arnt22 | 9 | 8.1488E-08 | 0.73573976 | 0.47 | 0.376 | 0.00016298 |

| | | | | | | | |
|---|---|---|---|---|---|---|---|
| Fibroblast | Pde7b1 | 9 | 9.9743E-08 | 0.48013596 | 0.73 | 0.584 | 0.00019949 |
| Fibroblast | Agap22 | 9 | 1.1184E-07 | 0.9794455 | 0.255 | 0.132 | 0.00022369 |
| Fibroblast | Ptpn132 | 9 | 1.2027E-07 | 0.85394397 | 0.484 | 0.425 | 0.00024054 |
| Fibroblast | AC130940.13 | 9 | 1.2642E-07 | 0.9989568 | 0.268 | 0.179 | 0.00025284 |
| Fibroblast | Rcan12 | 9 | 1.2807E-07 | 0.47855919 | 0.472 | 0.283 | 0.00025615 |
| Fibroblast | Atp8b41 | 9 | 1.4306E-07 | 0.9534536 | 0.399 | 0.282 | 0.00028612 |
| Fibroblast | Mcf2l | 9 | 1.453E-07 | 0.27930598 | 0.433 | 0.276 | 0.0002906 |
| Fibroblast | Cnnm21 | 9 | 1.6059E-07 | 0.52079538 | 0.518 | 0.359 | 0.00032119 |
| Fibroblast | Hk21 | 9 | 1.6808E-07 | 0.37225306 | 0.603 | 0.472 | 0.00033615 |
| Fibroblast | AABR070275 | 9 | 1.9471E-07 | 0.37569044 | 0.438 | 0.3 | 0.00038942 |
| Fibroblast | Nr4a13 | 9 | 2.0099E-07 | 0.55810965 | 0.387 | 0.28 | 0.00040199 |
| Fibroblast | AABR070070 | 9 | 2.011E-07 | 0.58104214 | 0.251 | 0.166 | 0.00040221 |
| Fibroblast | Srpk3 | 9 | 2.5249E-07 | 0.4588291 | 0.297 | 0.139 | 0.00050499 |
| Fibroblast | Vash23 | 9 | 2.5517E-07 | 1.67380996 | 0.309 | 0.237 | 0.00051034 |
| Fibroblast | Gpnmb1 | 9 | 2.7063E-07 | 0.55019942 | 0.438 | 0.303 | 0.00054126 |
| Fibroblast | Fam78b1 | 9 | 5.2876E-07 | 0.41118922 | 0.268 | 0.15 | 0.00105753 |
| Fibroblast | Rbfox1 | 9 | 5.5836E-07 | 0.90833638 | 0.433 | 0.318 | 0.00111673 |
| Fibroblast | Gja11 | 9 | 5.6177E-07 | 0.50489215 | 0.504 | 0.352 | 0.00112354 |
| Fibroblast | Arhgef261 | 9 | 5.6592E-07 | 1.38033549 | 0.336 | 0.248 | 0.00113185 |
| Fibroblast | Pdgfrb2 | 9 | 5.9327E-07 | 0.27694763 | 0.825 | 0.707 | 0.00118653 |
| Fibroblast | Slc6a6 | 9 | 6.7606E-07 | 1.39542748 | 0.321 | 0.189 | 0.00135212 |
| Fibroblast | Pcp4l11 | 9 | 8.6049E-07 | 0.4592385 | 0.384 | 0.198 | 0.00172098 |
| Fibroblast | Serf21 | 9 | 8.8756E-07 | 0.30087059 | 0.555 | 0.427 | 0.00177513 |
| Fibroblast | Ankh | 9 | 1.0838E-06 | 0.96655454 | 0.45 | 0.365 | 0.0021676 |
| Fibroblast | Palm23 | 9 | 1.184E-06 | 0.77983265 | 0.482 | 0.422 | 0.00236809 |
| Fibroblast | Gucy1a12 | 9 | 1.2841E-06 | 0.62581176 | 0.579 | 0.53 | 0.00256826 |
| Fibroblast | Arhgap441 | 9 | 1.4996E-06 | 0.58238703 | 0.436 | 0.356 | 0.0029992 |
| Fibroblast | AABR070347 | 9 | 2.6264E-06 | 0.33818262 | 0.37 | 0.222 | 0.00525279 |
| Fibroblast | Mmp21 | 9 | 2.805E-06 | 0.54347765 | 0.589 | 0.462 | 0.00560993 |
| Fibroblast | Slc20a22 | 9 | 3.3312E-06 | 0.31545023 | 0.54 | 0.425 | 0.00666247 |
| Fibroblast | Kcnq11 | 9 | 4.6941E-06 | 0.31443125 | 0.453 | 0.325 | 0.00938827 |

| | | | | | | | |
|---|---|---|---|---|---|---|---|
| Fibroblast | Sorbs12 | 9 | 9.1703E-06 | 0.32175293 | 0.74 | 0.656 | 0.01834063 |
| Fibroblast | Pmepa12 | 9 | 1.1129E-05 | 0.34042476 | 0.684 | 0.57 | 0.02225851 |
| Fibroblast | AABR070544 | 9 | 1.1455E-05 | 0.56013767 | 0.436 | 0.3 | 0.0229096 |
| Fibroblast | Mboat21 | 9 | 1.3735E-05 | 0.37474482 | 0.453 | 0.379 | 0.02747008 |
| Fibroblast | Slc20a11 | 9 | 2.2279E-05 | 0.5263725 | 0.477 | 0.408 | 0.04455888 |
| Fibroblast | Npr32 | 9 | 2.461E-05 | 0.38893175 | 0.414 | 0.291 | 0.04921952 |
| Fibroblast | Chad | 10 | 0 | 6.28760733 | 0.335 | 0.012 | 0 |
| Fibroblast | Atp6v0a4 | 10 | 2.155E-298 | 2.87317836 | 0.335 | 0.012 | 4.311E-295 |
| Fibroblast | Olr292 | 10 | 4.586E-235 | 2.65452808 | 0.314 | 0.004 | 9.173E-232 |
| Fibroblast | Mmp9 | 10 | 7.758E-165 | 1.66617962 | 0.257 | 0.016 | 1.552E-161 |
| Fibroblast | Ccdc1031 | 10 | 2.26E-128 | 2.06158817 | 0.48 | 0.084 | 4.52E-125 |
| Fibroblast | Il1rn | 10 | 1.024E-125 | 3.40128811 | 0.42 | 0.063 | 2.047E-122 |
| Fibroblast | Atp8b3 | 10 | 3.262E-120 | 4.11900106 | 0.344 | 0.009 | 6.523E-117 |
| Fibroblast | Bgn2 | 10 | 7.665E-115 | 2.58348442 | 0.921 | 0.45 | 1.533E-111 |
| Fibroblast | Col3a12 | 10 | 2.071E-107 | 1.42950191 | 0.997 | 0.967 | 4.141E-104 |
| Fibroblast | Col1a14 | 10 | 6.3552E-97 | 1.69242576 | 0.994 | 0.872 | 1.271E-93 |
| Fibroblast | Lum1 | 10 | 3.573E-78 | 2.15348063 | 0.825 | 0.444 | 7.1459E-75 |
| Fibroblast | Gpx33 | 10 | 9.0409E-75 | 2.182879 | 0.804 | 0.368 | 1.8082E-71 |
| Fibroblast | Islr1 | 10 | 4.1973E-74 | 2.98211964 | 0.707 | 0.329 | 8.3947E-71 |
| Fibroblast | RT1-Da | 10 | 5.6124E-74 | 1.553433 | 0.483 | 0.082 | 1.1225E-70 |
| Fibroblast | Sparc2 | 10 | 8.0563E-73 | 1.43789016 | 0.949 | 0.734 | 1.6113E-69 |
| Fibroblast | Fstl12 | 10 | 1.471E-72 | 1.30216604 | 0.976 | 0.816 | 2.9421E-69 |
| Fibroblast | LOC1009119 | 10 | 9.049E-70 | 3.0550251 | 0.716 | 0.283 | 1.8098E-66 |
| Fibroblast | Serpinf1 | 10 | 2.056E-69 | 2.320116 | 0.779 | 0.365 | 4.1119E-66 |
| Fibroblast | Upk1b | 10 | 4.8708E-61 | 1.83227679 | 0.278 | 0.071 | 9.7416E-58 |
| Fibroblast | Myo16 | 10 | 5.0928E-60 | 0.91213362 | 0.369 | 0.081 | 1.0186E-56 |
| Fibroblast | LOC1083526 | 10 | 1.3698E-58 | 1.94576031 | 0.728 | 0.31 | 2.7396E-55 |
| Fibroblast | Plvap | 10 | 3.0791E-57 | 1.70946158 | 0.35 | 0.034 | 6.1583E-54 |
| Fibroblast | S100a61 | 10 | 4.224E-54 | 2.22459991 | 0.662 | 0.258 | 8.448E-51 |
| Fibroblast | Hmox13 | 10 | 1.8839E-53 | 1.62409008 | 0.52 | 0.173 | 3.7677E-50 |
| Fibroblast | Ltc4s | 10 | 4.0027E-52 | 1.40302529 | 0.465 | 0.113 | 8.0054E-49 |

| | | | | | | | |
|---|---|---|---|---|---|---|---|
| Fibroblast | Tmsb4x1 | 10 | 5.9376E-51 | 1.73093424 | 0.74 | 0.39 | 1.1875E-47 |
| Fibroblast | Ano32 | 10 | 1.9367E-50 | 1.37523453 | 0.444 | 0.125 | 3.8734E-47 |
| Fibroblast | Mgp2 | 10 | 8.1631E-49 | 1.33732074 | 0.958 | 0.796 | 1.6326E-45 |
| Fibroblast | Ecm11 | 10 | 3.8257E-48 | 2.2705036 | 0.674 | 0.379 | 7.6513E-45 |
| Fibroblast | Igfbp63 | 10 | 4.0226E-47 | 2.75438619 | 0.601 | 0.288 | 8.0451E-44 |
| Fibroblast | Fbxw101 | 10 | 6.9303E-47 | 1.64602182 | 0.532 | 0.185 | 1.3861E-43 |
| Fibroblast | Ctsz1 | 10 | 4.93E-46 | 0.97018714 | 0.462 | 0.126 | 9.86E-43 |
| Fibroblast | Comp2 | 10 | 6.7677E-45 | 0.66089516 | 0.387 | 0.132 | 1.3535E-41 |
| Fibroblast | Pex5l | 10 | 1.0485E-43 | 1.82505946 | 0.35 | 0.111 | 2.0971E-40 |
| Fibroblast | Pi162 | 10 | 1.1478E-43 | 1.69075736 | 0.737 | 0.449 | 2.2956E-40 |
| Fibroblast | Hmga22 | 10 | 1.3592E-43 | 1.20361929 | 0.353 | 0.116 | 2.7185E-40 |
| Fibroblast | Gsn3 | 10 | 1.8378E-43 | 1.49782173 | 0.855 | 0.573 | 3.6755E-40 |
| Fibroblast | Hhip2 | 10 | 4.9793E-43 | 0.35549474 | 0.369 | 0.105 | 9.9586E-40 |
| Fibroblast | Cd84 | 10 | 8.1164E-43 | 1.35826755 | 0.387 | 0.123 | 1.6233E-39 |
| Fibroblast | LOC1003644 | 10 | 2.5526E-42 | 1.64565503 | 0.683 | 0.384 | 5.1052E-39 |
| Fibroblast | B2m1 | 10 | 1.0846E-40 | 1.47915077 | 0.719 | 0.426 | 2.1691E-37 |
| Fibroblast | Sox171 | 10 | 1.1846E-40 | 0.38347141 | 0.438 | 0.142 | 2.3693E-37 |
| Fibroblast | Crip11 | 10 | 2.025E-40 | 1.65574661 | 0.671 | 0.338 | 4.0501E-37 |
| Fibroblast | Prelp1 | 10 | 4.7638E-40 | 2.72673485 | 0.562 | 0.286 | 9.5275E-37 |
| Fibroblast | Ccl7 | 10 | 1.9815E-39 | 2.29915297 | 0.444 | 0.149 | 3.9629E-36 |
| Fibroblast | Map3k7cl2 | 10 | 3.0707E-39 | 1.08656504 | 0.396 | 0.153 | 6.1415E-36 |
| Fibroblast | Matk1 | 10 | 7.937E-39 | 1.36059081 | 0.417 | 0.114 | 1.5874E-35 |
| Fibroblast | Thsd7b2 | 10 | 1.029E-36 | 0.72180669 | 0.435 | 0.184 | 2.058E-33 |
| Fibroblast | Mmp22 | 10 | 1.5253E-36 | 1.25887047 | 0.785 | 0.456 | 3.0506E-33 |
| Fibroblast | Cd63 | 10 | 2.8449E-36 | 2.00425539 | 0.583 | 0.307 | 5.6899E-33 |
| Fibroblast | Tpt1 | 10 | 1.0721E-35 | 1.77040165 | 0.601 | 0.271 | 2.1442E-32 |
| Fibroblast | Grid21 | 10 | 1.1518E-35 | 1.51880245 | 0.378 | 0.159 | 2.3037E-32 |
| Fibroblast | Mfap51 | 10 | 6.1141E-35 | 0.9152616 | 0.921 | 0.692 | 1.2228E-31 |
| Fibroblast | Ncaph1 | 10 | 1.167E-34 | 1.62011731 | 0.375 | 0.191 | 2.334E-31 |
| Fibroblast | Atf31 | 10 | 3.3666E-34 | 0.83945711 | 0.402 | 0.142 | 6.7332E-31 |
| Fibroblast | Tyrobp3 | 10 | 6.2773E-34 | 1.97046711 | 0.353 | 0.209 | 1.2555E-30 |

| | | | | | | | |
|---|---|---|---|---|---|---|---|
| Fibroblast | Mdga2 | 10 | 1.1972E-33 | 0.97170218 | 0.492 | 0.137 | 2.3944E-30 |
| Fibroblast | RT1-Db1 | 10 | 2.7268E-33 | 0.29767209 | 0.444 | 0.153 | 5.4536E-30 |
| Fibroblast | Lgals12 | 10 | 6.389E-33 | 1.39326008 | 0.668 | 0.408 | 1.2778E-29 |
| Fibroblast | Gap431 | 10 | 4.9243E-32 | 0.38958061 | 0.601 | 0.327 | 9.8485E-29 |
| Fibroblast | Sparcl12 | 10 | 6.7376E-32 | 1.08340455 | 0.776 | 0.469 | 1.3475E-28 |
| Fibroblast | Uchl11 | 10 | 7.1477E-31 | 1.53490897 | 0.396 | 0.131 | 1.4295E-27 |
| Fibroblast | Trpc62 | 10 | 1.084E-30 | 1.19465283 | 0.471 | 0.211 | 2.1679E-27 |
| Fibroblast | Meox11 | 10 | 2.109E-30 | 1.48537742 | 0.381 | 0.147 | 4.2181E-27 |
| Fibroblast | Dnajc6 | 10 | 1.0577E-28 | 1.25431471 | 0.468 | 0.178 | 2.1154E-25 |
| Fibroblast | Stk263 | 10 | 4.9039E-28 | 1.09284876 | 0.444 | 0.136 | 9.8078E-25 |
| Fibroblast | Lgi41 | 10 | 9.987E-28 | 1.30972106 | 0.58 | 0.316 | 1.9974E-24 |
| Fibroblast | LOC1003614 | 10 | 1.0767E-27 | 1.46819866 | 0.595 | 0.328 | 2.1534E-24 |
| Fibroblast | Nusap13 | 10 | 1.1675E-27 | 0.99566553 | 0.483 | 0.197 | 2.335E-24 |
| Fibroblast | Tnfrsf12a2 | 10 | 3.3611E-27 | 0.9367907 | 0.486 | 0.188 | 6.7222E-24 |
| Fibroblast | Penk1 | 10 | 5.2662E-27 | 1.87278279 | 0.423 | 0.216 | 1.0532E-23 |
| Fibroblast | Top2a | 10 | 6.2484E-27 | 0.56527757 | 0.486 | 0.21 | 1.2497E-23 |
| Fibroblast | Psap1 | 10 | 8.5211E-27 | 1.6051461 | 0.523 | 0.277 | 1.7042E-23 |
| Fibroblast | S100a41 | 10 | 1.3508E-26 | 2.72221072 | 0.426 | 0.239 | 2.7017E-23 |
| Fibroblast | Itga22 | 10 | 2.7121E-26 | 0.69309644 | 0.553 | 0.237 | 5.4243E-23 |
| Fibroblast | Gpc32 | 10 | 9.1126E-26 | 0.25533875 | 0.511 | 0.21 | 1.8225E-22 |
| Fibroblast | Igfbp41 | 10 | 1.4243E-25 | 0.88971929 | 0.701 | 0.428 | 2.8486E-22 |
| Fibroblast | Ca32 | 10 | 1.6335E-25 | 0.833965 | 0.323 | 0.145 | 3.2669E-22 |
| Fibroblast | AABR070637 | 10 | 2.5665E-25 | 1.95253677 | 0.369 | 0.132 | 5.133E-22 |
| Fibroblast | LOC1009118 | 10 | 3.4557E-25 | 1.47923188 | 0.526 | 0.285 | 6.9114E-22 |
| Fibroblast | Rims12 | 10 | 5.0675E-25 | 0.49915262 | 0.356 | 0.133 | 1.0135E-21 |
| Fibroblast | Fosb1 | 10 | 6.8223E-25 | 2.20950291 | 0.296 | 0.085 | 1.3645E-21 |
| Fibroblast | Ccl21 | 10 | 7.4834E-25 | 1.35579154 | 0.353 | 0.14 | 1.4967E-21 |
| Fibroblast | Ftl11 | 10 | 8.0999E-25 | 1.8048003 | 0.502 | 0.261 | 1.62E-21 |
| Fibroblast | Ifitm101 | 10 | 8.5326E-25 | 0.95525229 | 0.42 | 0.181 | 1.7065E-21 |
| Fibroblast | Ubash3b1 | 10 | 9.6714E-25 | 0.46834539 | 0.683 | 0.429 | 1.9343E-21 |
| Fibroblast | AABR070492 | 10 | 1.5548E-24 | 0.35589648 | 0.502 | 0.209 | 3.1097E-21 |

| | | | | | | | |
|---|---|---|---|---|---|---|---|
| Fibroblast | Ankrd91 | 10 | 2.2826E-24 | 0.86791845 | 0.511 | 0.218 | 4.5652E-21 |
| Fibroblast | Piezo23 | 10 | 2.3808E-24 | 0.9072863 | 0.731 | 0.43 | 4.7616E-21 |
| Fibroblast | Fn15 | 10 | 2.4242E-24 | 0.79891503 | 0.846 | 0.621 | 4.8485E-21 |
| Fibroblast | Col6a22 | 10 | 2.5039E-24 | 0.75438185 | 0.888 | 0.669 | 5.0079E-21 |
| Fibroblast | Uap12 | 10 | 2.6392E-24 | 0.86208637 | 0.84 | 0.62 | 5.2783E-21 |
| Fibroblast | Actg21 | 10 | 3.9125E-24 | 2.0096695 | 0.372 | 0.183 | 7.825E-21 |
| Fibroblast | Ca4 | 10 | 8.5073E-24 | 0.5370435 | 0.511 | 0.19 | 1.7015E-20 |
| Fibroblast | Nkain3 | 10 | 1.5765E-23 | 1.001009 | 0.423 | 0.17 | 3.1529E-20 |
| Fibroblast | Col13a1 | 10 | 1.7112E-23 | 1.2113116 | 0.263 | 0.085 | 3.4225E-20 |
| Fibroblast | Sfrp43 | 10 | 4.4042E-23 | 1.58758074 | 0.538 | 0.323 | 8.8084E-20 |
| Fibroblast | Ildr22 | 10 | 1.4142E-22 | 0.31535549 | 0.619 | 0.318 | 2.8283E-19 |
| Fibroblast | Hspb31 | 10 | 1.6878E-22 | 0.42848189 | 0.353 | 0.137 | 3.3756E-19 |
| Fibroblast | Tmem200a | 10 | 2.2363E-22 | 1.09783332 | 0.369 | 0.182 | 4.4727E-19 |
| Fibroblast | Inpp5d1 | 10 | 2.4354E-22 | 0.85242325 | 0.523 | 0.261 | 4.8708E-19 |
| Fibroblast | Loxl13 | 10 | 2.6248E-22 | 0.79493001 | 0.876 | 0.683 | 5.2497E-19 |
| Fibroblast | Prph1 | 10 | 3.0842E-22 | 1.45285374 | 0.465 | 0.207 | 6.1684E-19 |
| Fibroblast | Fbln22 | 10 | 3.2766E-22 | 0.74031188 | 0.761 | 0.497 | 6.5532E-19 |
| Fibroblast | Cst31 | 10 | 5.3261E-22 | 1.64521443 | 0.523 | 0.302 | 1.0652E-18 |
| Fibroblast | Tuba1b1 | 10 | 8.6106E-22 | 1.88263992 | 0.474 | 0.221 | 1.7221E-18 |
| Fibroblast | Timp23 | 10 | 1.233E-21 | 0.58992819 | 0.909 | 0.788 | 2.4661E-18 |
| Fibroblast | Prdx51 | 10 | 1.3008E-21 | 1.30905566 | 0.438 | 0.205 | 2.6015E-18 |
| Fibroblast | AABR070581 | 10 | 1.3698E-21 | 1.06103413 | 0.474 | 0.23 | 2.7397E-18 |
| Fibroblast | Grk32 | 10 | 1.5757E-21 | 0.82701809 | 0.477 | 0.23 | 3.1514E-18 |
| Fibroblast | Ptprv2 | 10 | 1.603E-21 | 0.28577179 | 0.598 | 0.322 | 3.206E-18 |
| Fibroblast | Timp33 | 10 | 2.0903E-21 | 0.75710847 | 0.77 | 0.484 | 4.1805E-18 |
| Fibroblast | Cdh132 | 10 | 2.7183E-21 | 0.68804409 | 0.779 | 0.576 | 5.4365E-18 |
| Fibroblast | Msr1 | 10 | 9.8537E-21 | 1.72386485 | 0.323 | 0.161 | 1.9707E-17 |
| Fibroblast | Ifitm32 | 10 | 1.4207E-20 | 1.93044336 | 0.498 | 0.332 | 2.8415E-17 |
| Fibroblast | Vcan3 | 10 | 3.9739E-20 | 0.64998029 | 0.876 | 0.667 | 7.9477E-17 |
| Fibroblast | Frmd32 | 10 | 7.878E-20 | 0.92737283 | 0.444 | 0.244 | 1.5756E-16 |
| Fibroblast | AABR070339 | 10 | 1.1652E-19 | 0.93818095 | 0.547 | 0.33 | 2.3305E-16 |

| | | | | | | | |
|---|---|---|---|---|---|---|---|
| Fibroblast | Cox6a11 | 10 | 1.386E-19 | 1.39613806 | 0.498 | 0.285 | 2.7719E-16 |
| Fibroblast | Slc20a12 | 10 | 1.7154E-19 | 0.51961979 | 0.683 | 0.402 | 3.4307E-16 |
| Fibroblast | Cspg43 | 10 | 1.7223E-19 | 0.56030099 | 0.514 | 0.242 | 3.4446E-16 |
| Fibroblast | Cpeb12 | 10 | 1.7467E-19 | 0.63598719 | 0.692 | 0.402 | 3.4934E-16 |
| Fibroblast | Fth11 | 10 | 2.6712E-19 | 1.27117254 | 0.505 | 0.307 | 5.3425E-16 |
| Fibroblast | Pappa24 | 10 | 2.7091E-19 | 1.0454912 | 0.366 | 0.239 | 5.4183E-16 |
| Fibroblast | P2ry141 | 10 | 2.7203E-19 | 0.61046305 | 0.483 | 0.198 | 5.4407E-16 |
| Fibroblast | Ckap21 | 10 | 3.3615E-19 | 2.14947996 | 0.305 | 0.152 | 6.723E-16 |
| Fibroblast | Uqcr101 | 10 | 4.0357E-19 | 1.33718208 | 0.541 | 0.303 | 8.0714E-16 |
| Fibroblast | Rgcc2 | 10 | 4.2335E-19 | 1.0138538 | 0.483 | 0.245 | 8.467E-16 |
| Fibroblast | Spink81 | 10 | 7.4864E-19 | 0.62129987 | 0.508 | 0.251 | 1.4973E-15 |
| Fibroblast | Hspb71 | 10 | 1.0085E-18 | 0.98674049 | 0.589 | 0.345 | 2.017E-15 |
| Fibroblast | Homer12 | 10 | 1.0147E-18 | 0.46441232 | 0.619 | 0.385 | 2.0294E-15 |
| Fibroblast | Rspo15 | 10 | 1.8255E-18 | 0.41969326 | 0.55 | 0.265 | 3.6509E-15 |
| Fibroblast | Gabrb1 | 10 | 1.9509E-18 | 0.80629342 | 0.278 | 0.139 | 3.9018E-15 |
| Fibroblast | Abcd22 | 10 | 2.3346E-18 | 0.72756588 | 0.414 | 0.203 | 4.6692E-15 |
| Fibroblast | Epsti11 | 10 | 3.1211E-18 | 0.81096508 | 0.269 | 0.143 | 6.2422E-15 |
| Fibroblast | Ptgs1 | 10 | 3.7847E-18 | 0.47758521 | 0.323 | 0.174 | 7.5693E-15 |
| Fibroblast | Rab7b3 | 10 | 4.0877E-18 | 0.66515909 | 0.61 | 0.391 | 8.1755E-15 |
| Fibroblast | Serf22 | 10 | 4.1378E-18 | 0.94547222 | 0.562 | 0.428 | 8.2756E-15 |
| Fibroblast | Fam71f11 | 10 | 4.7207E-18 | 0.45248876 | 0.453 | 0.242 | 9.4414E-15 |
| Fibroblast | Tmem1002 | 10 | 5.5645E-18 | 1.10119973 | 0.556 | 0.358 | 1.1129E-14 |
| Fibroblast | Ltbp23 | 10 | 7.9505E-18 | 0.60310518 | 0.752 | 0.469 | 1.5901E-14 |
| Fibroblast | AC110709.24 | 10 | 1.0147E-17 | 1.04243137 | 0.411 | 0.247 | 2.0295E-14 |
| Fibroblast | Myh111 | 10 | 1.4868E-17 | 1.35330979 | 0.523 | 0.276 | 2.9735E-14 |
| Fibroblast | Fgf11 | 10 | 2.449E-17 | 0.88157555 | 0.508 | 0.274 | 4.898E-14 |
| Fibroblast | Mmp165 | 10 | 2.6697E-17 | 0.27621871 | 0.583 | 0.301 | 5.3395E-14 |
| Fibroblast | Col14a13 | 10 | 2.9487E-17 | 0.55400076 | 0.924 | 0.715 | 5.8973E-14 |
| Fibroblast | Srgap12 | 10 | 3.0873E-17 | 0.45049222 | 0.897 | 0.645 | 6.1746E-14 |
| Fibroblast | LOC6877802 | 10 | 3.161E-17 | 1.57167954 | 0.435 | 0.252 | 6.322E-14 |
| Fibroblast | Uqcrq3 | 10 | 3.2444E-17 | 0.78772036 | 0.58 | 0.334 | 6.4888E-14 |

| | | | | | | | |
|---|---|---|---|---|---|---|---|
| Fibroblast | Col16a13 | 10 | 3.3531E-17 | 0.70522223 | 0.686 | 0.422 | 6.7063E-14 |
| Fibroblast | Ndrg42 | 10 | 3.8303E-17 | 0.61321721 | 0.547 | 0.323 | 7.6606E-14 |
| Fibroblast | Lpcat22 | 10 | 3.8669E-17 | 0.91095818 | 0.45 | 0.228 | 7.7338E-14 |
| Fibroblast | Flna3 | 10 | 4.9803E-17 | 0.59937883 | 0.71 | 0.421 | 9.9606E-14 |
| Fibroblast | Bin2 | 10 | 5.2675E-17 | 0.62120881 | 0.523 | 0.272 | 1.0535E-13 |
| Fibroblast | Gal3st31 | 10 | 5.421E-17 | 0.46352555 | 0.366 | 0.17 | 1.0842E-13 |
| Fibroblast | Polr2m1 | 10 | 5.6117E-17 | 0.60494063 | 0.665 | 0.423 | 1.1223E-13 |
| Fibroblast | Kcnab13 | 10 | 5.9456E-17 | 0.48758518 | 0.589 | 0.35 | 1.1891E-13 |
| Fibroblast | Ccn11 | 10 | 9.6433E-17 | 1.15277764 | 0.48 | 0.272 | 1.9287E-13 |
| Fibroblast | AABR070311 | 10 | 2.3783E-16 | 0.41773662 | 0.441 | 0.205 | 4.7567E-13 |
| Fibroblast | Cd683 | 10 | 2.5669E-16 | 0.27823507 | 0.426 | 0.236 | 5.1338E-13 |
| Fibroblast | Serping12 | 10 | 2.8239E-16 | 1.51205712 | 0.468 | 0.303 | 5.6478E-13 |
| Fibroblast | Slc38a11 | 10 | 3.4653E-16 | 0.60377345 | 0.568 | 0.347 | 6.9306E-13 |
| Fibroblast | Naca2 | 10 | 3.5859E-16 | 0.91987509 | 0.52 | 0.317 | 7.1718E-13 |
| Fibroblast | Trpc32 | 10 | 4.7875E-16 | 0.39410842 | 0.432 | 0.258 | 9.575E-13 |
| Fibroblast | Ezr2 | 10 | 5.3102E-16 | 0.61941432 | 0.511 | 0.279 | 1.062E-12 |
| Fibroblast | Bcl11a | 10 | 5.3323E-16 | 0.94869347 | 0.347 | 0.2 | 1.0665E-12 |
| Fibroblast | Cox6a22 | 10 | 5.8199E-16 | 0.5346957 | 0.619 | 0.342 | 1.164E-12 |
| Fibroblast | Sfrp23 | 10 | 5.9873E-16 | 1.51320927 | 0.438 | 0.254 | 1.1975E-12 |
| Fibroblast | F13a1 | 10 | 6.3825E-16 | 0.29846783 | 0.287 | 0.13 | 1.2765E-12 |
| Fibroblast | Mis18bp12 | 10 | 7.6191E-16 | 0.87494653 | 0.529 | 0.274 | 1.5238E-12 |
| Fibroblast | Grin3a3 | 10 | 9.377E-16 | 1.5703441 | 0.453 | 0.238 | 1.8754E-12 |
| Fibroblast | Ano12 | 10 | 1.1195E-15 | 1.02978233 | 0.287 | 0.152 | 2.2391E-12 |
| Fibroblast | Dpt1 | 10 | 1.14E-15 | 0.52182696 | 0.876 | 0.716 | 2.2799E-12 |
| Fibroblast | Unc5c | 10 | 1.6454E-15 | 0.68607125 | 0.378 | 0.156 | 3.2908E-12 |
| Fibroblast | Stmn43 | 10 | 2.0499E-15 | 0.88339342 | 0.465 | 0.282 | 4.0998E-12 |
| Fibroblast | Cobll11 | 10 | 2.1868E-15 | 0.45252185 | 0.574 | 0.334 | 4.3736E-12 |
| Fibroblast | Tpd52l1 | 10 | 2.4261E-15 | 0.44477123 | 0.426 | 0.224 | 4.8521E-12 |
| Fibroblast | Aff22 | 10 | 4.6506E-15 | 0.58966555 | 0.644 | 0.41 | 9.3012E-12 |
| Fibroblast | Plaur3 | 10 | 5.1422E-15 | 0.55613512 | 0.665 | 0.443 | 1.0284E-11 |
| Fibroblast | Gm2a | 10 | 6.0904E-15 | 2.08010711 | 0.308 | 0.165 | 1.2181E-11 |

| | | | | | | | |
|---|---|---|---|---|---|---|---|
| Fibroblast | Spp13 | 10 | 6.9581E-15 | 1.95037159 | 0.438 | 0.251 | 1.3916E-11 |
| Fibroblast | Robo22 | 10 | 7.3123E-15 | 0.5068419 | 0.556 | 0.338 | 1.4625E-11 |
| Fibroblast | Cdkn1a3 | 10 | 9.3384E-15 | 0.79317366 | 0.529 | 0.338 | 1.8677E-11 |
| Fibroblast | Has22 | 10 | 1.0154E-14 | 0.50262852 | 0.568 | 0.375 | 2.0309E-11 |
| Fibroblast | Pak7 | 10 | 1.0226E-14 | 1.30225398 | 0.302 | 0.096 | 2.0451E-11 |
| Fibroblast | AABR070449 | 10 | 1.2157E-14 | 0.37554331 | 0.538 | 0.307 | 2.4314E-11 |
| Fibroblast | Ctsb | 10 | 1.4726E-14 | 1.33798301 | 0.508 | 0.368 | 2.9452E-11 |
| Fibroblast | Ntn12 | 10 | 1.7576E-14 | 0.42239185 | 0.74 | 0.507 | 3.5153E-11 |
| Fibroblast | Pmepa13 | 10 | 2.5407E-14 | 0.60121395 | 0.764 | 0.568 | 5.0814E-11 |
| Fibroblast | Anxa31 | 10 | 2.8039E-14 | 0.36240986 | 0.447 | 0.239 | 5.6079E-11 |
| Fibroblast | Cd443 | 10 | 2.8116E-14 | 0.88808568 | 0.562 | 0.39 | 5.6231E-11 |
| Fibroblast | Tagln1 | 10 | 3.2473E-14 | 1.87536702 | 0.353 | 0.223 | 6.4945E-11 |
| Fibroblast | Trpm31 | 10 | 3.5424E-14 | 0.58126408 | 0.462 | 0.282 | 7.0847E-11 |
| Fibroblast | AABR070410 | 10 | 3.7204E-14 | 1.22565753 | 0.39 | 0.2 | 7.4408E-11 |
| Fibroblast | B4galnt31 | 10 | 3.8151E-14 | 1.10324497 | 0.287 | 0.207 | 7.6302E-11 |
| Fibroblast | Ackr32 | 10 | 4.4319E-14 | 0.61636023 | 0.671 | 0.459 | 8.8639E-11 |
| Fibroblast | Cd551 | 10 | 5.1126E-14 | 0.48997332 | 0.495 | 0.315 | 1.0225E-10 |
| Fibroblast | Apoe | 10 | 5.4302E-14 | 1.44964884 | 0.447 | 0.309 | 1.086E-10 |
| Fibroblast | Fam167a2 | 10 | 6.13E-14 | 1.07713601 | 0.423 | 0.289 | 1.226E-10 |
| Fibroblast | Col8a12 | 10 | 6.5923E-14 | 0.49588447 | 0.961 | 0.87 | 1.3185E-10 |
| Fibroblast | Tuft11 | 10 | 8.0951E-14 | 0.65197869 | 0.474 | 0.289 | 1.619E-10 |
| Fibroblast | Dcn3 | 10 | 8.5745E-14 | 0.49391485 | 0.9 | 0.792 | 1.7149E-10 |
| Fibroblast | Cdh20 | 10 | 9.0911E-14 | 0.65467028 | 0.462 | 0.205 | 1.8182E-10 |
| Fibroblast | Lamc24 | 10 | 9.7344E-14 | 0.40152405 | 0.429 | 0.243 | 1.9469E-10 |
| Fibroblast | Fgf92 | 10 | 1.1654E-13 | 0.81807542 | 0.402 | 0.25 | 2.3309E-10 |
| Fibroblast | Lgmn3 | 10 | 1.2598E-13 | 0.92325316 | 0.508 | 0.321 | 2.5196E-10 |
| Fibroblast | Fras11 | 10 | 1.7628E-13 | 1.09209645 | 0.272 | 0.134 | 3.5257E-10 |
| Fibroblast | Ptk2b4 | 10 | 1.8292E-13 | 0.50099098 | 0.647 | 0.445 | 3.6584E-10 |
| Fibroblast | Fam227b2 | 10 | 1.9771E-13 | 0.39713164 | 0.535 | 0.307 | 3.9542E-10 |
| Fibroblast | Nrap2 | 10 | 2.1753E-13 | 0.49798267 | 0.453 | 0.238 | 4.3507E-10 |
| Fibroblast | Csrp13 | 10 | 2.3121E-13 | 0.57953777 | 0.613 | 0.361 | 4.6242E-10 |

| | | | | | | | |
|---|---|---|---|---|---|---|---|
| Fibroblast | Pygm2 | 10 | 2.9439E-13 | 0.71495237 | 0.426 | 0.219 | 5.8878E-10 |
| Fibroblast | Ano21 | 10 | 3.3697E-13 | 0.52400717 | 0.269 | 0.178 | 6.7395E-10 |
| Fibroblast | Myl61 | 10 | 4.7907E-13 | 0.79568851 | 0.604 | 0.43 | 9.5813E-10 |
| Fibroblast | Cilp4 | 10 | 5.0285E-13 | 1.36174867 | 0.498 | 0.344 | 1.0057E-09 |
| Fibroblast | Ccn32 | 10 | 5.612E-13 | 0.58274491 | 0.471 | 0.265 | 1.1224E-09 |
| Fibroblast | Thbs42 | 10 | 5.6194E-13 | 2.65340545 | 0.326 | 0.282 | 1.1239E-09 |
| Fibroblast | Adgrd12 | 10 | 5.9736E-13 | 0.3795516 | 0.84 | 0.625 | 1.1947E-09 |
| Fibroblast | Ndufa41 | 10 | 6.7087E-13 | 0.79569702 | 0.535 | 0.347 | 1.3417E-09 |
| Fibroblast | Bend61 | 10 | 6.8215E-13 | 0.54564129 | 0.498 | 0.302 | 1.3643E-09 |
| Fibroblast | Cyba1 | 10 | 7.8596E-13 | 1.57805594 | 0.48 | 0.336 | 1.5719E-09 |
| Fibroblast | L1cam1 | 10 | 8.3899E-13 | 0.62574162 | 0.278 | 0.16 | 1.678E-09 |
| Fibroblast | Me12 | 10 | 1.3307E-12 | 0.31683662 | 0.589 | 0.384 | 2.6615E-09 |
| Fibroblast | Itgb62 | 10 | 1.8348E-12 | 0.35585814 | 0.302 | 0.173 | 3.6695E-09 |
| Fibroblast | Tm4sf192 | 10 | 1.9672E-12 | 0.41471588 | 0.29 | 0.15 | 3.9344E-09 |
| Fibroblast | Rcan13 | 10 | 1.9827E-12 | 1.17473697 | 0.477 | 0.285 | 3.9654E-09 |
| Fibroblast | Nav2 | 10 | 2.078E-12 | 0.28905885 | 0.335 | 0.156 | 4.156E-09 |
| Fibroblast | Atp5f1b1 | 10 | 2.1198E-12 | 0.60042627 | 0.677 | 0.5 | 4.2395E-09 |
| Fibroblast | C1qtnf34 | 10 | 2.9897E-12 | 1.17134859 | 0.329 | 0.239 | 5.9795E-09 |
| Fibroblast | Has12 | 10 | 3.0782E-12 | 0.84509916 | 0.556 | 0.391 | 6.1564E-09 |
| Fibroblast | Il342 | 10 | 4.6668E-12 | 0.71528092 | 0.459 | 0.275 | 9.3335E-09 |
| Fibroblast | Diaph31 | 10 | 5.5101E-12 | 0.49774789 | 0.471 | 0.289 | 1.102E-08 |
| Fibroblast | Des1 | 10 | 5.8117E-12 | 0.97123387 | 0.441 | 0.318 | 1.1623E-08 |
| Fibroblast | Acoxl | 10 | 9.5859E-12 | 1.4568051 | 0.305 | 0.138 | 1.9172E-08 |
| Fibroblast | Treml4 | 10 | 1.0321E-11 | 2.10716454 | 0.278 | 0.075 | 2.0641E-08 |
| Fibroblast | Dusp271 | 10 | 1.1386E-11 | 0.82781075 | 0.278 | 0.178 | 2.2772E-08 |
| Fibroblast | Cadm31 | 10 | 1.2144E-11 | 0.59162334 | 0.526 | 0.406 | 2.4288E-08 |
| Fibroblast | Gpx11 | 10 | 1.255E-11 | 1.73745085 | 0.317 | 0.205 | 2.51E-08 |
| Fibroblast | Tnnc11 | 10 | 1.3525E-11 | 0.7398067 | 0.429 | 0.264 | 2.7051E-08 |
| Fibroblast | Adgrg22 | 10 | 1.5006E-11 | 0.48914029 | 0.492 | 0.335 | 3.0013E-08 |
| Fibroblast | Mfge81 | 10 | 2.4528E-11 | 1.48395198 | 0.35 | 0.226 | 4.9057E-08 |
| Fibroblast | Inhba4 | 10 | 2.7163E-11 | 0.44007993 | 0.462 | 0.254 | 5.4326E-08 |

| | | | | | | | |
|---|---|---|---|---|---|---|---|
| Fibroblast | Susd12 | 10 | 4.1007E-11 | 0.66804931 | 0.369 | 0.217 | 8.2013E-08 |
| Fibroblast | Kitlg | 10 | 4.9335E-11 | 0.35251307 | 0.511 | 0.317 | 9.867E-08 |
| Fibroblast | Hspb81 | 10 | 5.6814E-11 | 1.08061272 | 0.55 | 0.412 | 1.1363E-07 |
| Fibroblast | Nhs3 | 10 | 5.8946E-11 | 0.33525203 | 0.541 | 0.367 | 1.1789E-07 |
| Fibroblast | LOC1001348 | 10 | 6.2015E-11 | 1.11333077 | 0.375 | 0.254 | 1.2403E-07 |
| Fibroblast | Vcam13 | 10 | 1.1952E-10 | 0.60979296 | 0.477 | 0.31 | 2.3905E-07 |
| Fibroblast | Agmo3 | 10 | 1.2613E-10 | 0.86988621 | 0.393 | 0.247 | 2.5226E-07 |
| Fibroblast | Fstl51 | 10 | 1.27E-10 | 0.89304077 | 0.293 | 0.213 | 2.5399E-07 |
| Fibroblast | Ccdc803 | 10 | 1.3779E-10 | 0.34306264 | 0.97 | 0.892 | 2.7558E-07 |
| Fibroblast | Sema3c1 | 10 | 1.5063E-10 | 0.43535081 | 0.625 | 0.443 | 3.0126E-07 |
| Fibroblast | Cox8b1 | 10 | 1.6429E-10 | 0.53828843 | 0.538 | 0.325 | 3.2857E-07 |
| Fibroblast | Aebp12 | 10 | 1.6875E-10 | 0.72890614 | 0.607 | 0.47 | 3.3749E-07 |
| Fibroblast | Lurap1l1 | 10 | 1.7408E-10 | 0.25969598 | 0.628 | 0.433 | 3.4817E-07 |
| Fibroblast | Htra32 | 10 | 1.8519E-10 | 0.37831778 | 0.728 | 0.512 | 3.7037E-07 |
| Fibroblast | Ntrk32 | 10 | 1.8617E-10 | 0.2515846 | 0.529 | 0.321 | 3.7233E-07 |
| Fibroblast | PCOLCE22 | 10 | 1.8656E-10 | 0.43577328 | 0.508 | 0.352 | 3.7313E-07 |
| Fibroblast | Eva1c2 | 10 | 2.3724E-10 | 0.44539587 | 0.471 | 0.304 | 4.7449E-07 |
| Fibroblast | Prg41 | 10 | 3.0276E-10 | 1.82543126 | 0.302 | 0.226 | 6.0552E-07 |
| Fibroblast | Mertk1 | 10 | 3.1106E-10 | 0.64370628 | 0.335 | 0.215 | 6.2212E-07 |
| Fibroblast | Sema3d3 | 10 | 3.2099E-10 | 0.47671305 | 0.571 | 0.391 | 6.4197E-07 |
| Fibroblast | Pik3cd1 | 10 | 3.2801E-10 | 0.47843861 | 0.477 | 0.303 | 6.5603E-07 |
| Fibroblast | Serpine25 | 10 | 4.9781E-10 | 0.83706604 | 0.453 | 0.366 | 9.9561E-07 |
| Fibroblast | AABR070655 | 10 | 5.1988E-10 | 0.96013548 | 0.356 | 0.197 | 1.0398E-06 |
| Fibroblast | P3h2 | 10 | 6.3735E-10 | 0.60432926 | 0.387 | 0.256 | 1.2747E-06 |
| Fibroblast | Pip5k1b | 10 | 6.5008E-10 | 0.88152113 | 0.338 | 0.19 | 1.3002E-06 |
| Fibroblast | AABR070265 | 10 | 7.4851E-10 | 0.47983997 | 0.417 | 0.286 | 1.497E-06 |
| Fibroblast | Adgrl41 | 10 | 7.5087E-10 | 0.31317054 | 0.311 | 0.191 | 1.5017E-06 |
| Fibroblast | Ephb11 | 10 | 9.7866E-10 | 0.69456997 | 0.435 | 0.31 | 1.9573E-06 |
| Fibroblast | Ank32 | 10 | 1.0232E-09 | 0.2689008 | 0.498 | 0.292 | 2.0464E-06 |
| Fibroblast | Cotl13 | 10 | 1.0913E-09 | 0.48334296 | 0.498 | 0.326 | 2.1826E-06 |
| Fibroblast | Reep13 | 10 | 1.2463E-09 | 0.40879531 | 0.356 | 0.245 | 2.4926E-06 |

| | | | | | | | |
|---|---|---|---|---|---|---|---|
| Fibroblast | Postn5 | 10 | 1.3101E-09 | 0.51070723 | 0.858 | 0.773 | 2.6201E-06 |
| Fibroblast | Cryab1 | 10 | 1.468E-09 | 0.68315644 | 0.468 | 0.376 | 2.936E-06 |
| Fibroblast | Rnd33 | 10 | 2.0729E-09 | 0.31984852 | 0.713 | 0.534 | 4.1457E-06 |
| Fibroblast | Olfm22 | 10 | 2.2075E-09 | 0.93696998 | 0.408 | 0.305 | 4.4149E-06 |
| Fibroblast | Fabp52 | 10 | 2.2908E-09 | 0.44810791 | 0.335 | 0.211 | 4.5817E-06 |
| Fibroblast | Lgals3 | 10 | 2.4537E-09 | 1.45659802 | 0.257 | 0.165 | 4.9074E-06 |
| Fibroblast | Tpm22 | 10 | 2.5167E-09 | 0.67237438 | 0.375 | 0.266 | 5.0333E-06 |
| Fibroblast | Prc11 | 10 | 2.7671E-09 | 1.21325939 | 0.293 | 0.128 | 5.5343E-06 |
| Fibroblast | Ctsk1 | 10 | 2.9509E-09 | 1.0405515 | 0.338 | 0.245 | 5.9018E-06 |
| Fibroblast | Myl31 | 10 | 4.0851E-09 | 0.59389417 | 0.713 | 0.543 | 8.1701E-06 |
| Fibroblast | Chn21 | 10 | 4.2575E-09 | 0.35896755 | 0.498 | 0.353 | 8.515E-06 |
| Fibroblast | Galnt131 | 10 | 4.3104E-09 | 0.97537337 | 0.263 | 0.2 | 8.6208E-06 |
| Fibroblast | Egr14 | 10 | 4.4018E-09 | 0.47320364 | 0.39 | 0.26 | 8.8036E-06 |
| Fibroblast | Pdlim52 | 10 | 4.9855E-09 | 0.44696967 | 0.825 | 0.669 | 9.971E-06 |
| Fibroblast | Olfml2b4 | 10 | 5.3264E-09 | 0.47799859 | 0.571 | 0.406 | 1.0653E-05 |
| Fibroblast | Srl1 | 10 | 5.3992E-09 | 0.90700593 | 0.462 | 0.31 | 1.0798E-05 |
| Fibroblast | Jph11 | 10 | 5.8134E-09 | 0.26374118 | 0.429 | 0.281 | 1.1627E-05 |
| Fibroblast | Tnfaip8l33 | 10 | 5.9703E-09 | 0.71106034 | 0.393 | 0.305 | 1.1941E-05 |
| Fibroblast | Asb183 | 10 | 6.2174E-09 | 0.5448628 | 0.42 | 0.286 | 1.2435E-05 |
| Fibroblast | Pdpn3 | 10 | 6.2334E-09 | 0.42808199 | 0.414 | 0.283 | 1.2467E-05 |
| Fibroblast | Axl2 | 10 | 6.8105E-09 | 0.32063032 | 0.867 | 0.677 | 1.3621E-05 |
| Fibroblast | Mt11 | 10 | 6.8187E-09 | 0.74372654 | 0.375 | 0.289 | 1.3637E-05 |
| Fibroblast | Kif20a3 | 10 | 8.333E-09 | 1.14371869 | 0.29 | 0.247 | 1.6666E-05 |
| Fibroblast | Ctsd2 | 10 | 9.0267E-09 | 1.5313864 | 0.326 | 0.221 | 1.8053E-05 |
| Fibroblast | Ddc2 | 10 | 9.0721E-09 | 0.59189537 | 0.378 | 0.232 | 1.8144E-05 |
| Fibroblast | Tbx182 | 10 | 9.5797E-09 | 0.34548074 | 0.634 | 0.465 | 1.9159E-05 |
| Fibroblast | Il163 | 10 | 9.7974E-09 | 0.39046203 | 0.683 | 0.528 | 1.9595E-05 |
| Fibroblast | Nfatc22 | 10 | 1.2199E-08 | 0.50818782 | 0.353 | 0.235 | 2.4397E-05 |
| Fibroblast | Timp11 | 10 | 1.2505E-08 | 2.31079041 | 0.269 | 0.24 | 2.5011E-05 |
| Fibroblast | Tec3 | 10 | 1.3395E-08 | 0.27416151 | 0.42 | 0.254 | 2.679E-05 |
| Fibroblast | Lyz21 | 10 | 1.4712E-08 | 2.11143485 | 0.317 | 0.207 | 2.9425E-05 |

| | | | | | | | |
|---|---|---|---|---|---|---|---|
| Fibroblast | Parm12 | 10 | 1.5328E-08 | 0.29663403 | 0.29 | 0.189 | 3.0655E-05 |
| Fibroblast | Slc25a41 | 10 | 1.608E-08 | 0.39836057 | 0.589 | 0.525 | 3.216E-05 |
| Fibroblast | Dkk33 | 10 | 1.7314E-08 | 0.45550343 | 0.64 | 0.559 | 3.4627E-05 |
| Fibroblast | Ankrd11 | 10 | 1.9562E-08 | 0.5795444 | 0.795 | 0.583 | 3.9124E-05 |
| Fibroblast | Gpcpd11 | 10 | 2.0576E-08 | 0.2943621 | 0.55 | 0.389 | 4.1151E-05 |
| Fibroblast | Sox63 | 10 | 2.2897E-08 | 0.375665 | 0.686 | 0.485 | 4.5793E-05 |
| Fibroblast | Nova12 | 10 | 2.358E-08 | 0.25127448 | 0.477 | 0.34 | 4.716E-05 |
| Fibroblast | Gask1b2 | 10 | 2.4825E-08 | 0.43286024 | 0.731 | 0.553 | 4.965E-05 |
| Fibroblast | Ntng11 | 10 | 2.6238E-08 | 0.7821857 | 0.32 | 0.224 | 5.2476E-05 |
| Fibroblast | Acyp21 | 10 | 2.8172E-08 | 0.29339483 | 0.562 | 0.438 | 5.6343E-05 |
| Fibroblast | Sh3kbp14 | 10 | 3.0413E-08 | 0.25775211 | 0.601 | 0.414 | 6.0826E-05 |
| Fibroblast | Fos3 | 10 | 3.3815E-08 | 0.94844254 | 0.441 | 0.266 | 6.7631E-05 |
| Fibroblast | Cpe | 10 | 5.2827E-08 | 0.48731735 | 0.338 | 0.232 | 0.00010565 |
| Fibroblast | Kif20b | 10 | 5.4521E-08 | 0.64995685 | 0.387 | 0.147 | 0.00010904 |
| Fibroblast | Cfh1 | 10 | 5.5246E-08 | 0.68698484 | 0.414 | 0.308 | 0.00011049 |
| Fibroblast | Rgs6 | 10 | 6.071E-08 | 0.35144037 | 0.257 | 0.227 | 0.00012142 |
| Fibroblast | Slc7a24 | 10 | 6.141E-08 | 0.35849297 | 0.589 | 0.384 | 0.00012282 |
| Fibroblast | Atp5mg1 | 10 | 7.9338E-08 | 0.61827877 | 0.423 | 0.344 | 0.00015868 |
| Fibroblast | Clec7a | 10 | 7.964E-08 | 1.04161518 | 0.251 | 0.145 | 0.00015928 |
| Fibroblast | Nr4a14 | 10 | 9.0322E-08 | 0.2779057 | 0.338 | 0.282 | 0.00018064 |
| Fibroblast | Ppp1r12b | 10 | 9.2706E-08 | 0.39303179 | 0.813 | 0.627 | 0.00018541 |
| Fibroblast | Atp5mc31 | 10 | 9.41E-08 | 0.80373353 | 0.399 | 0.351 | 0.0001882 |
| Fibroblast | Acta21 | 10 | 9.7178E-08 | 2.06198159 | 0.347 | 0.273 | 0.00019436 |
| Fibroblast | Pltp1 | 10 | 1.3242E-07 | 0.91628779 | 0.396 | 0.316 | 0.00026483 |
| Fibroblast | Gja12 | 10 | 1.8066E-07 | 0.58057857 | 0.477 | 0.354 | 0.00036132 |
| Fibroblast | Atp5f1e2 | 10 | 1.9092E-07 | 0.99573535 | 0.372 | 0.321 | 0.00038184 |
| Fibroblast | Atp5me1 | 10 | 2.3484E-07 | 0.50911814 | 0.486 | 0.367 | 0.00046968 |
| Fibroblast | Hs3st3a13 | 10 | 2.4144E-07 | 0.4095764 | 0.426 | 0.304 | 0.00048288 |
| Fibroblast | Cthrc13 | 10 | 2.6605E-07 | 0.73925856 | 0.39 | 0.327 | 0.00053209 |
| Fibroblast | Rbp14 | 10 | 2.992E-07 | 0.4350993 | 0.432 | 0.342 | 0.0005984 |
| Fibroblast | Xkr43 | 10 | 3.169E-07 | 0.44214576 | 0.441 | 0.304 | 0.00063379 |

| | | | | | | | |
|---|---|---|---|---|---|---|---|
| Fibroblast | Pls33 | 10 | 3.2642E-07 | 0.34264231 | 0.834 | 0.675 | 0.00065283 |
| Fibroblast | Acta11 | 10 | 3.656E-07 | 0.95509005 | 0.347 | 0.261 | 0.00073119 |
| Fibroblast | Eda2r4 | 10 | 3.8185E-07 | 0.26558073 | 0.347 | 0.27 | 0.00076371 |
| Fibroblast | Lancl33 | 10 | 3.9524E-07 | 0.34259771 | 0.344 | 0.22 | 0.00079048 |
| Fibroblast | Itga73 | 10 | 3.9767E-07 | 0.89457402 | 0.317 | 0.243 | 0.00079533 |
| Fibroblast | Eef1a23 | 10 | 4.951E-07 | 0.43244571 | 0.417 | 0.323 | 0.00099019 |
| Fibroblast | Fat33 | 10 | 8.4952E-07 | 0.29193583 | 0.538 | 0.382 | 0.00169904 |
| Fibroblast | Mapk102 | 10 | 8.7211E-07 | 1.15832951 | 0.296 | 0.292 | 0.00174421 |
| Fibroblast | Cacna1g2 | 10 | 1.1114E-06 | 0.43689497 | 0.492 | 0.408 | 0.00222287 |
| Fibroblast | Mmp121 | 10 | 1.1633E-06 | 1.26167621 | 0.257 | 0.127 | 0.00232653 |
| Fibroblast | Lbp2 | 10 | 1.3288E-06 | 0.77858238 | 0.405 | 0.315 | 0.00265755 |
| Fibroblast | Cobl4 | 10 | 1.3355E-06 | 0.6271733 | 0.344 | 0.262 | 0.00267102 |
| Fibroblast | Ckb3 | 10 | 1.3938E-06 | 0.99306096 | 0.363 | 0.305 | 0.00278766 |
| Fibroblast | Fabp32 | 10 | 1.4519E-06 | 0.8006441 | 0.393 | 0.318 | 0.00290374 |
| Fibroblast | Tmtc21 | 10 | 1.4564E-06 | 0.7747473 | 0.329 | 0.241 | 0.00291286 |
| Fibroblast | Il1rapl12 | 10 | 1.4681E-06 | 0.77488234 | 0.366 | 0.262 | 0.00293621 |
| Fibroblast | Pde10a2 | 10 | 1.5726E-06 | 0.30012916 | 0.779 | 0.593 | 0.00314521 |
| Fibroblast | Gda3 | 10 | 2.0101E-06 | 0.25053248 | 0.888 | 0.755 | 0.00402012 |
| Fibroblast | Phyh1 | 10 | 2.0927E-06 | 0.34780418 | 0.414 | 0.338 | 0.00418532 |
| Fibroblast | Myh72 | 10 | 2.8033E-06 | 0.37460714 | 0.668 | 0.552 | 0.00560668 |
| Fibroblast | Oprm13 | 10 | 2.9132E-06 | 1.28869512 | 0.151 | 0.262 | 0.00582642 |
| Fibroblast | Cacna1d3 | 10 | 2.9902E-06 | 0.92090595 | 0.323 | 0.171 | 0.00598032 |
| Fibroblast | Nkd22 | 10 | 3.2267E-06 | 0.79952108 | 0.378 | 0.316 | 0.00645344 |
| Fibroblast | Gpnmb2 | 10 | 3.5919E-06 | 1.00142321 | 0.353 | 0.307 | 0.00718376 |
| Fibroblast | Flnc1 | 10 | 3.8156E-06 | 0.62929217 | 0.399 | 0.334 | 0.0076313 |
| Fibroblast | Cox5b1 | 10 | 4.2235E-06 | 1.03033506 | 0.32 | 0.285 | 0.00844704 |
| Fibroblast | Kcnh8 | 10 | 4.8912E-06 | 0.61162981 | 0.302 | 0.205 | 0.00978231 |
| Fibroblast | Mki672 | 10 | 4.9685E-06 | 0.64522958 | 0.218 | 0.266 | 0.00993705 |
| Fibroblast | Nrg22 | 10 | 5.0116E-06 | 0.46168949 | 0.317 | 0.209 | 0.01002312 |
| Fibroblast | Fbln12 | 10 | 5.1799E-06 | 0.32157274 | 0.737 | 0.612 | 0.01035978 |
| Fibroblast | Kcnj82 | 10 | 6.1129E-06 | 0.81203891 | 0.29 | 0.244 | 0.01222582 |

| | | | | | | | |
|---|---|---|---|---|---|---|---|
| Fibroblast | Atp5mc12 | 10 | 7.5917E-06 | 0.58781331 | 0.369 | 0.323 | 0.0151835 |
| Fibroblast | Nppa1 | 10 | 7.5991E-06 | 0.69417761 | 0.332 | 0.206 | 0.01519813 |
| Fibroblast | Fam180a2 | 10 | 7.989E-06 | 0.35967684 | 0.347 | 0.284 | 0.01597806 |
| Fibroblast | Cox4i11 | 10 | 8.0884E-06 | 0.47281599 | 0.438 | 0.417 | 0.0161768 |
| Fibroblast | Adgrl3 | 10 | 9.0807E-06 | 0.34162417 | 0.284 | 0.227 | 0.01816132 |
| Fibroblast | LOC1025530 | 10 | 9.5844E-06 | 0.35967061 | 0.426 | 0.373 | 0.01916883 |
| Fibroblast | Mb1 | 10 | 1.0054E-05 | 0.51761906 | 0.659 | 0.571 | 0.02010871 |
| Fibroblast | Sgcg2 | 10 | 1.5235E-05 | 0.65286123 | 0.372 | 0.314 | 0.0304694 |
| Fibroblast | Col8a23 | 10 | 2.2257E-05 | 0.28569881 | 0.477 | 0.428 | 0.04451381 |
| Fibroblast | Nr4a32 | 10 | 2.2434E-05 | 0.32276591 | 0.432 | 0.29 | 0.04486777 |
| Fibroblast | Chrdl12 | 10 | 2.3344E-05 | 0.25364257 | 0.344 | 0.205 | 0.04668815 |
| Fibroblast | Shank31 | 11 | 1.402E-101 | 3.08218832 | 0.805 | 0.271 | 2.8045E-98 |
| Fibroblast | Etl41 | 11 | 1.4336E-81 | 2.64546865 | 0.763 | 0.279 | 2.8672E-78 |
| Fibroblast | Prkch | 11 | 8.2826E-69 | 2.82973102 | 0.685 | 0.227 | 1.6565E-65 |
| Fibroblast | Adgrf5 | 11 | 3.5374E-67 | 2.901117 | 0.646 | 0.193 | 7.0749E-64 |
| Fibroblast | Cyyr1 | 11 | 7.3569E-67 | 2.77381177 | 0.672 | 0.176 | 1.4714E-63 |
| Fibroblast | Flt11 | 11 | 1.1845E-65 | 2.71459117 | 0.682 | 0.278 | 2.369E-62 |
| Fibroblast | Ptprb | 11 | 4.5548E-64 | 2.62391354 | 0.688 | 0.226 | 9.1096E-61 |
| Fibroblast | Dach1 | 11 | 1.2683E-61 | 2.2170262 | 0.718 | 0.276 | 2.5367E-58 |
| Fibroblast | Fam163a | 11 | 4.5548E-56 | 1.77404132 | 0.344 | 0.156 | 9.1096E-53 |
| Fibroblast | Fli1 | 11 | 6.4267E-52 | 1.51483595 | 0.847 | 0.448 | 1.2853E-48 |
| Fibroblast | Ablim31 | 11 | 1.4902E-51 | 3.02959165 | 0.575 | 0.224 | 2.9804E-48 |
| Fibroblast | Ccdc85a | 11 | 5.5451E-47 | 2.42446107 | 0.575 | 0.202 | 1.109E-43 |
| Fibroblast | Emcn1 | 11 | 1.0472E-45 | 2.58175864 | 0.575 | 0.203 | 2.0944E-42 |
| Fibroblast | Cdh133 | 11 | 3.9608E-45 | 1.15024828 | 0.867 | 0.573 | 7.9216E-42 |
| Fibroblast | Hmcn14 | 11 | 8.5263E-44 | 1.31896576 | 0.834 | 0.493 | 1.7053E-40 |
| Fibroblast | Nrp23 | 11 | 1.164E-41 | 1.25561082 | 0.782 | 0.375 | 2.3281E-38 |
| Fibroblast | Mgll1 | 11 | 1.8326E-41 | 1.7026028 | 0.636 | 0.304 | 3.6653E-38 |
| Fibroblast | Itpkb | 11 | 3.4647E-41 | 1.40033455 | 0.808 | 0.487 | 6.9294E-38 |
| Fibroblast | Lyn3 | 11 | 5.9658E-41 | 1.33275045 | 0.718 | 0.357 | 1.1932E-37 |
| Fibroblast | Ppp1r16b1 | 11 | 2.2501E-40 | 3.30879087 | 0.555 | 0.259 | 4.5003E-37 |

| | | | | | | | |
|---|---|---|---|---|---|---|---|
| Fibroblast | Efnb22 | 11 | 6.9497E-40 | 1.96847329 | 0.571 | 0.234 | 1.3899E-36 |
| Fibroblast | Adgrg31 | 11 | 2.0688E-38 | 2.17532705 | 0.25 | 0.122 | 4.1376E-35 |
| Fibroblast | Sox172 | 11 | 2.8262E-37 | 1.82631041 | 0.315 | 0.147 | 5.6524E-34 |
| Fibroblast | Nav32 | 11 | 8.0757E-37 | 1.31738341 | 0.698 | 0.341 | 1.6151E-33 |
| Fibroblast | Flt41 | 11 | 8.7508E-36 | 2.82802485 | 0.302 | 0.146 | 1.7502E-32 |
| Fibroblast | Kitlg1 | 11 | 1.0878E-35 | 1.49257645 | 0.653 | 0.313 | 2.1756E-32 |
| Fibroblast | Mcf2l1 | 11 | 2.2894E-35 | 2.24898164 | 0.555 | 0.273 | 4.5789E-32 |
| Fibroblast | Adgrl42 | 11 | 9.5908E-35 | 2.45007437 | 0.519 | 0.185 | 1.9182E-31 |
| Fibroblast | Myo101 | 11 | 3.8874E-34 | 1.57965903 | 0.646 | 0.362 | 7.7748E-31 |
| Fibroblast | Col13a11 | 11 | 3.1147E-33 | 1.58075683 | 0.321 | 0.083 | 6.2295E-30 |
| Fibroblast | Rasa4 | 11 | 1.0092E-32 | 2.47413967 | 0.474 | 0.202 | 2.0183E-29 |
| Fibroblast | Anxa32 | 11 | 1.9501E-32 | 2.10406287 | 0.513 | 0.237 | 3.9002E-29 |
| Fibroblast | Tnn2 | 11 | 2.4129E-32 | 1.27814933 | 0.38 | 0.193 | 4.8259E-29 |
| Fibroblast | Podxl | 11 | 2.8885E-32 | 2.54336156 | 0.474 | 0.209 | 5.777E-29 |
| Fibroblast | Ltbp24 | 11 | 4.2535E-32 | 0.8354333 | 0.802 | 0.468 | 8.5069E-29 |
| Fibroblast | Myrip1 | 11 | 5.3566E-32 | 3.0151371 | 0.51 | 0.275 | 1.0713E-28 |
| Fibroblast | Mtss12 | 11 | 1.878E-31 | 1.13384339 | 0.718 | 0.373 | 3.756E-28 |
| Fibroblast | Nuak12 | 11 | 3.5328E-31 | 1.43278151 | 0.669 | 0.395 | 7.0655E-28 |
| Fibroblast | Slc1a12 | 11 | 3.8718E-31 | 2.37684849 | 0.477 | 0.259 | 7.7436E-28 |
| Fibroblast | Rai144 | 11 | 7.1396E-31 | 0.93842709 | 0.805 | 0.476 | 1.4279E-27 |
| Fibroblast | Gcnt21 | 11 | 1.1006E-30 | 2.39546721 | 0.435 | 0.188 | 2.2011E-27 |
| Fibroblast | Mybpc21 | 11 | 1.4761E-30 | 2.55700607 | 0.299 | 0.111 | 2.9522E-27 |
| Fibroblast | Thsd7a1 | 11 | 2.6552E-30 | 1.80875123 | 0.532 | 0.242 | 5.3104E-27 |
| Fibroblast | Hmga23 | 11 | 3.6135E-30 | 1.46359246 | 0.315 | 0.118 | 7.227E-27 |
| Fibroblast | Zfp366 | 11 | 3.7498E-30 | 3.11003641 | 0.429 | 0.173 | 7.4996E-27 |
| Fibroblast | Slc35f11 | 11 | 1.0844E-29 | 2.80663693 | 0.315 | 0.136 | 2.1689E-26 |
| Fibroblast | Slc26a101 | 11 | 2.3614E-29 | 2.54086638 | 0.51 | 0.255 | 4.7228E-26 |
| Fibroblast | Ahr3 | 11 | 6.619E-29 | 0.76265874 | 0.714 | 0.439 | 1.3238E-25 |
| Fibroblast | Fn16 | 11 | 1.1825E-28 | 0.66693706 | 0.919 | 0.62 | 2.365E-25 |
| Fibroblast | Col6a51 | 11 | 2.4589E-28 | 0.27634198 | 0.299 | 0.113 | 4.9179E-25 |
| Fibroblast | Crim12 | 11 | 2.9047E-28 | 1.16669616 | 0.679 | 0.37 | 5.8094E-25 |

| | | | | | | | |
|---|---|---|---|---|---|---|---|
| Fibroblast | Kcnn41 | 11 | 3.7601E-28 | 1.85573772 | 0.276 | 0.134 | 7.5201E-25 |
| Fibroblast | Lef12 | 11 | 1.4302E-27 | 0.98296054 | 0.308 | 0.12 | 2.8604E-24 |
| Fibroblast | LOC1036942 | 11 | 1.5483E-27 | 2.21706105 | 0.429 | 0.176 | 3.0965E-24 |
| Fibroblast | Cmtm8 | 11 | 1.8198E-27 | 3.23736419 | 0.425 | 0.175 | 3.6396E-24 |
| Fibroblast | Chrdl13 | 11 | 5.8992E-27 | 1.06805596 | 0.422 | 0.203 | 1.1798E-23 |
| Fibroblast | Inpp5d2 | 11 | 7.1979E-27 | 1.83832105 | 0.532 | 0.262 | 1.4396E-23 |
| Fibroblast | Clic5 | 11 | 8.5854E-27 | 2.09519669 | 0.487 | 0.162 | 1.7171E-23 |
| Fibroblast | Mgmt4 | 11 | 9.8551E-27 | 0.6953905 | 0.75 | 0.468 | 1.971E-23 |
| Fibroblast | Podnl12 | 11 | 1.148E-26 | 1.62879526 | 0.292 | 0.109 | 2.296E-23 |
| Fibroblast | Meox23 | 11 | 1.7622E-26 | 1.12275913 | 0.63 | 0.38 | 3.5244E-23 |
| Fibroblast | Hecw23 | 11 | 2.0588E-26 | 1.21335751 | 0.636 | 0.377 | 4.1177E-23 |
| Fibroblast | Dnm32 | 11 | 3.5914E-26 | 0.97540999 | 0.731 | 0.487 | 7.1829E-23 |
| Fibroblast | St6galnac3 | 11 | 4.7276E-26 | 3.08423158 | 0.435 | 0.232 | 9.4552E-23 |
| Fibroblast | Cdkn2b3 | 11 | 1.2399E-25 | 0.59760901 | 0.357 | 0.167 | 2.4798E-22 |
| Fibroblast | Pik3r5 | 11 | 2.4296E-25 | 2.38621173 | 0.399 | 0.167 | 4.8592E-22 |
| Fibroblast | Lims22 | 11 | 2.5637E-25 | 1.34825738 | 0.523 | 0.272 | 5.1275E-22 |
| Fibroblast | LOC1003610 | 11 | 4.3017E-25 | 1.35777993 | 0.542 | 0.267 | 8.6034E-22 |
| Fibroblast | Cyth4 | 11 | 4.4962E-25 | 1.44054173 | 0.334 | 0.173 | 8.9924E-22 |
| Fibroblast | Ano22 | 11 | 8.3389E-25 | 2.82173054 | 0.302 | 0.177 | 1.6678E-21 |
| Fibroblast | Pdgfd3 | 11 | 8.3797E-25 | 0.76795626 | 0.76 | 0.524 | 1.6759E-21 |
| Fibroblast | AABR070076 | 11 | 1.2873E-24 | 2.65409207 | 0.464 | 0.222 | 2.5745E-21 |
| Fibroblast | Slc6a61 | 11 | 2.3209E-24 | 1.85823791 | 0.458 | 0.186 | 4.6418E-21 |
| Fibroblast | Cemip4 | 11 | 5.1377E-24 | 1.3161204 | 0.617 | 0.346 | 1.0275E-20 |
| Fibroblast | Cadps22 | 11 | 5.2697E-24 | 2.02355196 | 0.474 | 0.241 | 1.0539E-20 |
| Fibroblast | Specc14 | 11 | 5.5656E-24 | 0.72653639 | 0.799 | 0.526 | 1.1131E-20 |
| Fibroblast | Kntc12 | 11 | 5.7718E-24 | 0.83353794 | 0.354 | 0.179 | 1.1544E-20 |
| Fibroblast | Cd841 | 11 | 8.5379E-24 | 1.9936071 | 0.377 | 0.124 | 1.7076E-20 |
| Fibroblast | Aqp12 | 11 | 2.1617E-23 | 1.55720733 | 0.396 | 0.15 | 4.3234E-20 |
| Fibroblast | Abca41 | 11 | 2.4486E-23 | 0.68073104 | 0.344 | 0.164 | 4.8971E-20 |
| Fibroblast | Piezo24 | 11 | 3.027E-23 | 0.94986136 | 0.701 | 0.432 | 6.054E-20 |
| Fibroblast | Ccn23 | 11 | 3.8804E-23 | 0.82678988 | 0.76 | 0.483 | 7.7608E-20 |

| | | | | | | | |
|---|---|---|---|---|---|---|---|
| Fibroblast | Npnt3 | 11 | 4.4415E-23 | 0.818512 | 0.367 | 0.2 | 8.8831E-20 |
| Fibroblast | Smpd32 | 11 | 4.6243E-23 | 0.76555684 | 0.666 | 0.399 | 9.2486E-20 |
| Fibroblast | Il13ra21 | 11 | 5.7654E-23 | 1.09372877 | 0.266 | 0.106 | 1.1531E-19 |
| Fibroblast | Smoc12 | 11 | 7.0193E-23 | 0.60287185 | 0.321 | 0.192 | 1.4039E-19 |
| Fibroblast | Kif26b3 | 11 | 1.204E-22 | 1.01844132 | 0.581 | 0.318 | 2.408E-19 |
| Fibroblast | Comp3 | 11 | 3.1782E-22 | 0.41256532 | 0.276 | 0.136 | 6.3565E-19 |
| Fibroblast | Met1 | 11 | 4.8561E-22 | 0.89191342 | 0.256 | 0.121 | 9.7122E-19 |
| Fibroblast | Tnfrsf11b3 | 11 | 5.2623E-22 | 0.9210334 | 0.646 | 0.399 | 1.0525E-18 |
| Fibroblast | Dock81 | 11 | 1.5372E-21 | 1.78293065 | 0.526 | 0.332 | 3.0743E-18 |
| Fibroblast | AABR070179 | 11 | 1.6737E-21 | 1.26628174 | 0.38 | 0.198 | 3.3475E-18 |
| Fibroblast | Btnl9 | 11 | 2.0736E-21 | 3.14140215 | 0.302 | 0.121 | 4.1473E-18 |
| Fibroblast | Vwf | 11 | 2.8275E-21 | 2.64903216 | 0.37 | 0.127 | 5.655E-18 |
| Fibroblast | C1qtnf35 | 11 | 2.8699E-21 | 0.90211682 | 0.51 | 0.233 | 5.7398E-18 |
| Fibroblast | Akr1c151 | 11 | 2.9193E-21 | 1.41220161 | 0.471 | 0.237 | 5.8387E-18 |
| Fibroblast | Tyrobp4 | 11 | 3.3783E-21 | 0.68372784 | 0.386 | 0.208 | 6.7566E-18 |
| Fibroblast | Runx13 | 11 | 3.5184E-21 | 0.58525546 | 0.773 | 0.516 | 7.0368E-18 |
| Fibroblast | Erc2 | 11 | 3.6817E-21 | 0.7992259 | 0.321 | 0.123 | 7.3633E-18 |
| Fibroblast | Syk | 11 | 5.043E-21 | 1.85537035 | 0.445 | 0.182 | 1.0086E-17 |
| Fibroblast | Ptgs11 | 11 | 5.2458E-21 | 1.99495853 | 0.321 | 0.174 | 1.0492E-17 |
| Fibroblast | Pgm5 | 11 | 6.5512E-21 | 2.14393049 | 0.435 | 0.236 | 1.3102E-17 |
| Fibroblast | AABR070655 | 11 | 7.3189E-21 | 1.08212593 | 0.383 | 0.196 | 1.4638E-17 |
| Fibroblast | Ifitm102 | 11 | 9.4018E-21 | 1.86598242 | 0.386 | 0.182 | 1.8804E-17 |
| Fibroblast | Sema3a | 11 | 1.0612E-20 | 0.9507266 | 0.442 | 0.171 | 2.1224E-17 |
| Fibroblast | Ccn43 | 11 | 2.804E-20 | 0.96564898 | 0.468 | 0.249 | 5.6081E-17 |
| Fibroblast | AABR070606 | 11 | 2.8821E-20 | 1.92270384 | 0.37 | 0.235 | 5.7643E-17 |
| Fibroblast | Rgcc3 | 11 | 3.061E-20 | 1.85468386 | 0.458 | 0.246 | 6.122E-17 |
| Fibroblast | Col12a13 | 11 | 3.4261E-20 | 1.16601947 | 0.49 | 0.282 | 6.8522E-17 |
| Fibroblast | Adamts91 | 11 | 3.7377E-20 | 1.58857552 | 0.419 | 0.169 | 7.4755E-17 |
| Fibroblast | Prkcq | 11 | 6.5978E-20 | 0.94827809 | 0.26 | 0.076 | 1.3196E-16 |
| Fibroblast | Tnfrsf12a3 | 11 | 6.8546E-20 | 0.27266629 | 0.36 | 0.193 | 1.3709E-16 |
| Fibroblast | Dgkz1 | 11 | 6.9441E-20 | 0.71577946 | 0.523 | 0.278 | 1.3888E-16 |

| | | | | | | | |
|---|---|---|---|---|---|---|---|
| Fibroblast | Samd51 | 11 | 8.2353E-20 | 2.27842408 | 0.36 | 0.166 | 1.6471E-16 |
| Fibroblast | Spns2 | 11 | 9.7982E-20 | 2.97814221 | 0.367 | 0.2 | 1.9596E-16 |
| Fibroblast | Ano33 | 11 | 1.1457E-19 | 0.64355802 | 0.256 | 0.132 | 2.2914E-16 |
| Fibroblast | Tmtc22 | 11 | 2.0321E-19 | 1.30430329 | 0.432 | 0.237 | 4.0641E-16 |
| Fibroblast | Tmem63c4 | 11 | 2.6888E-19 | 1.38519751 | 0.38 | 0.233 | 5.3775E-16 |
| Fibroblast | Plcb12 | 11 | 3.4946E-19 | 0.91939803 | 0.516 | 0.32 | 6.9892E-16 |
| Fibroblast | Ddah13 | 11 | 3.7135E-19 | 0.59708025 | 0.607 | 0.39 | 7.427E-16 |
| Fibroblast | Sncaip2 | 11 | 5.6416E-19 | 2.0452761 | 0.445 | 0.289 | 1.1283E-15 |
| Fibroblast | Epha42 | 11 | 6.0109E-19 | 0.76115219 | 0.62 | 0.403 | 1.2022E-15 |
| Fibroblast | Fermt3 | 11 | 6.6386E-19 | 1.83442751 | 0.383 | 0.22 | 1.3277E-15 |
| Fibroblast | Lancl34 | 11 | 6.7363E-19 | 0.54193087 | 0.377 | 0.219 | 1.3473E-15 |
| Fibroblast | RT1-Db11 | 11 | 8.2916E-19 | 0.90003308 | 0.295 | 0.158 | 1.6583E-15 |
| Fibroblast | Dpp42 | 11 | 1.0505E-18 | 1.50435672 | 0.37 | 0.228 | 2.1009E-15 |
| Fibroblast | Gdf62 | 11 | 1.0565E-18 | 0.74923786 | 0.516 | 0.302 | 2.1129E-15 |
| Fibroblast | Tmem200a1 | 11 | 1.1058E-18 | 0.87415611 | 0.386 | 0.182 | 2.2117E-15 |
| Fibroblast | Lmcd12 | 11 | 1.1166E-18 | 1.04223321 | 0.571 | 0.384 | 2.2332E-15 |
| Fibroblast | Klhl41 | 11 | 1.3768E-18 | 1.84448966 | 0.331 | 0.154 | 2.7536E-15 |
| Fibroblast | Itga23 | 11 | 1.4094E-18 | 1.28493768 | 0.455 | 0.241 | 2.8189E-15 |
| Fibroblast | Dapk2 | 11 | 1.4874E-18 | 1.99292846 | 0.38 | 0.224 | 2.9747E-15 |
| Fibroblast | Palmd2 | 11 | 1.7958E-18 | 1.48464149 | 0.484 | 0.31 | 3.5915E-15 |
| Fibroblast | Filip1l4 | 11 | 2.4398E-18 | 0.41290575 | 0.896 | 0.623 | 4.8796E-15 |
| Fibroblast | Itgam1 | 11 | 2.5398E-18 | 1.11450137 | 0.377 | 0.186 | 5.0796E-15 |
| Fibroblast | Ciita | 11 | 3.3731E-18 | 2.50478099 | 0.263 | 0.108 | 6.7463E-15 |
| Fibroblast | Ncald3 | 11 | 3.4442E-18 | 0.932553 | 0.571 | 0.39 | 6.8884E-15 |
| Fibroblast | Adamts162 | 11 | 3.8196E-18 | 0.55302316 | 0.497 | 0.28 | 7.6391E-15 |
| Fibroblast | Nr4a33 | 11 | 3.9046E-18 | 2.73089806 | 0.487 | 0.289 | 7.8091E-15 |
| Fibroblast | Mest4 | 11 | 3.9607E-18 | 1.55450592 | 0.36 | 0.178 | 7.9215E-15 |
| Fibroblast | Tlr71 | 11 | 5.336E-18 | 1.52847643 | 0.305 | 0.165 | 1.0672E-14 |
| Fibroblast | Prss122 | 11 | 5.7394E-18 | 0.93559154 | 0.26 | 0.149 | 1.1479E-14 |
| Fibroblast | Lpcat23 | 11 | 5.8405E-18 | 1.23595215 | 0.419 | 0.23 | 1.1681E-14 |
| Fibroblast | Ptprv3 | 11 | 7.4227E-18 | 0.78546566 | 0.529 | 0.325 | 1.4845E-14 |

| | | | | | | | |
|---|---|---|---|---|---|---|---|
| Fibroblast | Timp34 | 11 | 7.6372E-18 | 0.63932207 | 0.74 | 0.486 | 1.5274E-14 |
| Fibroblast | Trpc51 | 11 | 1.0416E-17 | 0.9814714 | 0.315 | 0.164 | 2.0831E-14 |
| Fibroblast | Cd444 | 11 | 1.1309E-17 | 0.40650604 | 0.594 | 0.389 | 2.2619E-14 |
| Fibroblast | Nebl | 11 | 1.2929E-17 | 1.37519339 | 0.416 | 0.212 | 2.5857E-14 |
| Fibroblast | Csrp14 | 11 | 1.3843E-17 | 0.55029605 | 0.617 | 0.362 | 2.7686E-14 |
| Fibroblast | Lmo72 | 11 | 1.3988E-17 | 0.81796848 | 0.578 | 0.381 | 2.7976E-14 |
| Fibroblast | Procr1 | 11 | 1.5582E-17 | 0.45845027 | 0.286 | 0.127 | 3.1164E-14 |
| Fibroblast | Tll11 | 11 | 1.6373E-17 | 2.1312734 | 0.263 | 0.098 | 3.2746E-14 |
| Fibroblast | Pcdh171 | 11 | 1.7622E-17 | 2.05839951 | 0.409 | 0.232 | 3.5245E-14 |
| Fibroblast | Grem21 | 11 | 2.1391E-17 | 0.60557267 | 0.399 | 0.207 | 4.2782E-14 |
| Fibroblast | Grb142 | 11 | 2.2504E-17 | 0.8493667 | 0.643 | 0.414 | 4.5009E-14 |
| Fibroblast | Myl92 | 11 | 2.7913E-17 | 0.61698037 | 0.399 | 0.17 | 5.5826E-14 |
| Fibroblast | Flrt24 | 11 | 2.8946E-17 | 0.55284167 | 0.737 | 0.506 | 5.7891E-14 |
| Fibroblast | Vcl3 | 11 | 2.9718E-17 | 0.61709761 | 0.802 | 0.546 | 5.9437E-14 |
| Fibroblast | Ccn33 | 11 | 4.1351E-17 | 1.38198836 | 0.451 | 0.266 | 8.2703E-14 |
| Fibroblast | Ptprc3 | 11 | 6.3262E-17 | 1.20010852 | 0.448 | 0.252 | 1.2652E-13 |
| Fibroblast | Flvcr21 | 11 | 6.374E-17 | 2.15709714 | 0.367 | 0.256 | 1.2748E-13 |
| Fibroblast | Thbs13 | 11 | 6.681E-17 | 0.85483996 | 0.497 | 0.315 | 1.3362E-13 |
| Fibroblast | Hck | 11 | 8.4595E-17 | 1.74284726 | 0.282 | 0.133 | 1.6919E-13 |
| Fibroblast | Agtr1a3 | 11 | 9.7861E-17 | 0.37568554 | 0.643 | 0.425 | 1.9572E-13 |
| Fibroblast | Ptger33 | 11 | 1.1837E-16 | 0.55665946 | 0.438 | 0.256 | 2.3675E-13 |
| Fibroblast | AABR070215 | 11 | 1.2493E-16 | 0.88785197 | 0.516 | 0.348 | 2.4987E-13 |
| Fibroblast | Lurap1l2 | 11 | 1.3712E-16 | 0.52941135 | 0.636 | 0.433 | 2.7425E-13 |
| Fibroblast | Csrp23 | 11 | 1.4078E-16 | 1.06378437 | 0.519 | 0.324 | 2.8156E-13 |
| Fibroblast | Efna54 | 11 | 1.5164E-16 | 0.83736508 | 0.529 | 0.374 | 3.0327E-13 |
| Fibroblast | Tbxas12 | 11 | 1.7956E-16 | 1.75998955 | 0.429 | 0.251 | 3.5912E-13 |
| Fibroblast | Ak53 | 11 | 2.0949E-16 | 1.10866067 | 0.481 | 0.289 | 4.1899E-13 |
| Fibroblast | Ccser1 | 11 | 2.2267E-16 | 2.06678583 | 0.377 | 0.232 | 4.4534E-13 |
| Fibroblast | Nrg12 | 11 | 2.6615E-16 | 0.41302572 | 0.552 | 0.344 | 5.323E-13 |
| Fibroblast | Prom13 | 11 | 2.7552E-16 | 3.22771224 | 0.354 | 0.288 | 5.5104E-13 |
| Fibroblast | Fyb12 | 11 | 3.2659E-16 | 1.4265091 | 0.416 | 0.237 | 6.5318E-13 |

| | | | | | | | |
|---|---|---|---|---|---|---|---|
| Fibroblast | Actn13 | 11 | 4.7516E-16 | 0.42711612 | 0.792 | 0.544 | 9.5032E-13 |
| Fibroblast | Ubash3b2 | 11 | 5.3994E-16 | 0.92141156 | 0.571 | 0.434 | 1.0799E-12 |
| Fibroblast | Lcp22 | 11 | 6.9507E-16 | 1.8013498 | 0.318 | 0.199 | 1.3901E-12 |
| Fibroblast | Fbln23 | 11 | 8.5287E-16 | 0.30599189 | 0.753 | 0.498 | 1.7057E-12 |
| Fibroblast | Pde5a2 | 11 | 1.0839E-15 | 0.57865579 | 0.646 | 0.44 | 2.1678E-12 |
| Fibroblast | Nxn2 | 11 | 1.1771E-15 | 0.43956078 | 0.802 | 0.565 | 2.3541E-12 |
| Fibroblast | Tnc3 | 11 | 1.3343E-15 | 0.87560005 | 0.474 | 0.29 | 2.6687E-12 |
| Fibroblast | Ralgapa22 | 11 | 1.7363E-15 | 0.47112139 | 0.714 | 0.472 | 3.4725E-12 |
| Fibroblast | Ptk2b5 | 11 | 1.7896E-15 | 0.57709064 | 0.653 | 0.445 | 3.5791E-12 |
| Fibroblast | Kcnk21 | 11 | 1.9696E-15 | 1.13380183 | 0.419 | 0.2 | 3.9391E-12 |
| Fibroblast | Mfsd4a | 11 | 1.9868E-15 | 1.61922272 | 0.25 | 0.127 | 3.9736E-12 |
| Fibroblast | Chn22 | 11 | 2.0315E-15 | 0.59486057 | 0.588 | 0.351 | 4.0631E-12 |
| Fibroblast | Sorbs21 | 11 | 2.1201E-15 | 0.51571008 | 0.555 | 0.315 | 4.2403E-12 |
| Fibroblast | Cers63 | 11 | 2.2042E-15 | 0.44559296 | 0.76 | 0.517 | 4.4083E-12 |
| Fibroblast | Plxna43 | 11 | 2.2939E-15 | 0.4833835 | 0.666 | 0.389 | 4.5878E-12 |
| Fibroblast | Kcnq53 | 11 | 2.2954E-15 | 0.63758247 | 0.406 | 0.219 | 4.5908E-12 |
| Fibroblast | Cd300a1 | 11 | 2.894E-15 | 1.35432761 | 0.419 | 0.22 | 5.7879E-12 |
| Fibroblast | Slc10a63 | 11 | 3.066E-15 | 0.9445392 | 0.36 | 0.19 | 6.132E-12 |
| Fibroblast | Ctss2 | 11 | 3.2696E-15 | 1.85693629 | 0.438 | 0.267 | 6.5393E-12 |
| Fibroblast | Dipk2b | 11 | 4.7098E-15 | 2.56384214 | 0.341 | 0.156 | 9.4195E-12 |
| Fibroblast | Cxcl122 | 11 | 5.0286E-15 | 0.4057199 | 0.5 | 0.316 | 1.0057E-11 |
| Fibroblast | Nox43 | 11 | 5.1035E-15 | 0.46179313 | 0.727 | 0.522 | 1.0207E-11 |
| Fibroblast | Camk1d4 | 11 | 5.2088E-15 | 1.10049485 | 0.442 | 0.285 | 1.0418E-11 |
| Fibroblast | Sox132 | 11 | 5.6989E-15 | 0.50965908 | 0.597 | 0.391 | 1.1398E-11 |
| Fibroblast | Msr11 | 11 | 6.3787E-15 | 1.00583124 | 0.282 | 0.163 | 1.2757E-11 |
| Fibroblast | Arnt23 | 11 | 6.4533E-15 | 0.90587409 | 0.536 | 0.375 | 1.2907E-11 |
| Fibroblast | Ank33 | 11 | 7.4035E-15 | 0.81894708 | 0.477 | 0.293 | 1.4807E-11 |
| Fibroblast | Mx12 | 11 | 7.7655E-15 | 1.89536201 | 0.276 | 0.21 | 1.5531E-11 |
| Fibroblast | Chst151 | 11 | 7.7686E-15 | 2.61948432 | 0.36 | 0.226 | 1.5537E-11 |
| Fibroblast | Uchl12 | 11 | 7.9452E-15 | 0.57361658 | 0.276 | 0.136 | 1.589E-11 |
| Fibroblast | Map3k7cl3 | 11 | 8.4667E-15 | 1.63253769 | 0.25 | 0.158 | 1.6933E-11 |

| | | | | | | | |
|---|---|---|---|---|---|---|---|
| Fibroblast | Tagln2 | 11 | 8.6712E-15 | 0.90487408 | 0.432 | 0.221 | 1.7342E-11 |
| Fibroblast | Sybu | 11 | 8.9787E-15 | 1.45243448 | 0.36 | 0.185 | 1.7957E-11 |
| Fibroblast | Tnfaip62 | 11 | 9.0295E-15 | 0.36877406 | 0.591 | 0.384 | 1.8059E-11 |
| Fibroblast | Rapgef53 | 11 | 1.0069E-14 | 1.49848926 | 0.409 | 0.265 | 2.0138E-11 |
| Fibroblast | Sh3bp22 | 11 | 1.234E-14 | 1.15599046 | 0.49 | 0.296 | 2.4679E-11 |
| Fibroblast | Fam71f12 | 11 | 1.2724E-14 | 0.61238306 | 0.39 | 0.244 | 2.5448E-11 |
| Fibroblast | Marchf11 | 11 | 1.2952E-14 | 0.71167374 | 0.282 | 0.161 | 2.5905E-11 |
| Fibroblast | Olfml2b5 | 11 | 1.3793E-14 | 0.40317252 | 0.607 | 0.406 | 2.7586E-11 |
| Fibroblast | Trabd2b3 | 11 | 1.4146E-14 | 0.81802725 | 0.429 | 0.233 | 2.8293E-11 |
| Fibroblast | Ccn53 | 11 | 1.438E-14 | 0.47643012 | 0.607 | 0.422 | 2.8761E-11 |
| Fibroblast | Itga85 | 11 | 1.5236E-14 | 0.35256085 | 0.828 | 0.593 | 3.0472E-11 |
| Fibroblast | Csf1r1 | 11 | 1.595E-14 | 1.04074166 | 0.373 | 0.203 | 3.19E-11 |
| Fibroblast | RGD1565355 | 11 | 1.731E-14 | 0.75746232 | 0.432 | 0.246 | 3.4619E-11 |
| Fibroblast | Vegfc3 | 11 | 1.7313E-14 | 0.2966866 | 0.604 | 0.385 | 3.4626E-11 |
| Fibroblast | Pappa13 | 11 | 1.9829E-14 | 0.60453473 | 0.627 | 0.425 | 3.9657E-11 |
| Fibroblast | Gap432 | 11 | 1.9989E-14 | 0.57289491 | 0.506 | 0.331 | 3.9978E-11 |
| Fibroblast | Bin21 | 11 | 2.635E-14 | 1.3159714 | 0.455 | 0.275 | 5.27E-11 |
| Fibroblast | Npr33 | 11 | 2.698E-14 | 0.82783697 | 0.487 | 0.29 | 5.3961E-11 |
| Fibroblast | Col18a12 | 11 | 2.7024E-14 | 0.49984257 | 0.451 | 0.253 | 5.4048E-11 |
| Fibroblast | Ppp1r9a2 | 11 | 3.0869E-14 | 1.17710098 | 0.36 | 0.227 | 6.1738E-11 |
| Fibroblast | Edar3 | 11 | 3.1682E-14 | 0.30304164 | 0.468 | 0.318 | 6.3363E-11 |
| Fibroblast | Epha33 | 11 | 3.2945E-14 | 1.30989741 | 0.412 | 0.278 | 6.5891E-11 |
| Fibroblast | AABR070251 | 11 | 3.3074E-14 | 1.95912119 | 0.367 | 0.196 | 6.6148E-11 |
| Fibroblast | Ly86 | 11 | 3.3519E-14 | 1.45164086 | 0.39 | 0.203 | 6.7038E-11 |
| Fibroblast | Robo23 | 11 | 3.8562E-14 | 0.59825327 | 0.516 | 0.34 | 7.7125E-11 |
| Fibroblast | Eva1c3 | 11 | 4.6595E-14 | 0.61937426 | 0.461 | 0.305 | 9.3189E-11 |
| Fibroblast | Itgb63 | 11 | 5.0153E-14 | 1.46038027 | 0.292 | 0.173 | 1.0031E-10 |
| Fibroblast | Cdkl11 | 11 | 5.6316E-14 | 1.23179535 | 0.253 | 0.178 | 1.1263E-10 |
| Fibroblast | Herc63 | 11 | 5.7756E-14 | 0.61361707 | 0.455 | 0.295 | 1.1551E-10 |
| Fibroblast | Col24a13 | 11 | 6.0637E-14 | 0.36859756 | 0.532 | 0.37 | 1.2127E-10 |
| Fibroblast | Limch11 | 11 | 6.3708E-14 | 0.54737651 | 0.597 | 0.399 | 1.2742E-10 |

| | | | | | | | |
|---|---|---|---|---|---|---|---|
| Fibroblast | Sntg23 | 11 | 6.4212E-14 | 0.62396048 | 0.5 | 0.322 | 1.2842E-10 |
| Fibroblast | Pawr3 | 11 | 7.3468E-14 | 0.85879145 | 0.403 | 0.253 | 1.4694E-10 |
| Fibroblast | Fabp53 | 11 | 7.6344E-14 | 1.00100481 | 0.386 | 0.21 | 1.5269E-10 |
| Fibroblast | Shroom21 | 11 | 9.24E-14 | 0.88151572 | 0.25 | 0.122 | 1.848E-10 |
| Fibroblast | Tmem513 | 11 | 9.5461E-14 | 0.68319683 | 0.419 | 0.25 | 1.9092E-10 |
| Fibroblast | Ikzf1 | 11 | 1.0699E-13 | 1.86138038 | 0.393 | 0.244 | 2.1399E-10 |
| Fibroblast | Cd684 | 11 | 1.184E-13 | 1.19898166 | 0.39 | 0.238 | 2.368E-10 |
| Fibroblast | Tmcc32 | 11 | 1.3747E-13 | 0.67784726 | 0.536 | 0.373 | 2.7493E-10 |
| Fibroblast | Tpm23 | 11 | 1.5696E-13 | 0.6557813 | 0.435 | 0.265 | 3.1392E-10 |
| Fibroblast | Gpr1763 | 11 | 1.6323E-13 | 0.41778638 | 0.506 | 0.351 | 3.2647E-10 |
| Fibroblast | Kank12 | 11 | 1.7607E-13 | 0.39844485 | 0.627 | 0.408 | 3.5214E-10 |
| Fibroblast | Edil3 | 11 | 1.8277E-13 | 1.9584656 | 0.263 | 0.121 | 3.6554E-10 |
| Fibroblast | Bcl11a1 | 11 | 1.8691E-13 | 0.98890842 | 0.328 | 0.201 | 3.7383E-10 |
| Fibroblast | Lcp11 | 11 | 1.9983E-13 | 1.89267286 | 0.36 | 0.24 | 3.9967E-10 |
| Fibroblast | Stab12 | 11 | 2.3274E-13 | 1.69005452 | 0.37 | 0.246 | 4.6547E-10 |
| Fibroblast | Aldh1a23 | 11 | 2.4127E-13 | 0.64964757 | 0.523 | 0.353 | 4.8254E-10 |
| Fibroblast | Sfrp44 | 11 | 2.4431E-13 | 0.61555752 | 0.464 | 0.326 | 4.8861E-10 |
| Fibroblast | Plbd12 | 11 | 2.6833E-13 | 0.713576 | 0.334 | 0.213 | 5.3665E-10 |
| Fibroblast | Sorcs22 | 11 | 3.0145E-13 | 0.32535901 | 0.364 | 0.199 | 6.029E-10 |
| Fibroblast | Tmem1964 | 11 | 3.03E-13 | 0.31020581 | 0.636 | 0.402 | 6.0601E-10 |
| Fibroblast | Capn131 | 11 | 3.2505E-13 | 1.42704455 | 0.302 | 0.127 | 6.501E-10 |
| Fibroblast | Myh103 | 11 | 3.5786E-13 | 0.51122406 | 0.799 | 0.539 | 7.1572E-10 |
| Fibroblast | Wt1 | 11 | 3.7634E-13 | 1.67800583 | 0.299 | 0.167 | 7.5268E-10 |
| Fibroblast | Sv2c3 | 11 | 3.9265E-13 | 0.39349011 | 0.682 | 0.501 | 7.8529E-10 |
| Fibroblast | Sh3gl23 | 11 | 4.4935E-13 | 0.38089867 | 0.474 | 0.305 | 8.9871E-10 |
| Fibroblast | Tgfa1 | 11 | 4.6093E-13 | 0.40365733 | 0.289 | 0.148 | 9.2187E-10 |
| Fibroblast | Cobl5 | 11 | 4.9527E-13 | 0.42918514 | 0.445 | 0.258 | 9.9054E-10 |
| Fibroblast | Stxbp62 | 11 | 5.0304E-13 | 0.27917324 | 0.562 | 0.368 | 1.0061E-09 |
| Fibroblast | Chn12 | 11 | 5.0905E-13 | 0.34385688 | 0.344 | 0.176 | 1.0181E-09 |
| Fibroblast | Cdkn1a4 | 11 | 5.1812E-13 | 0.66274819 | 0.494 | 0.339 | 1.0362E-09 |
| Fibroblast | Reln1 | 11 | 5.6509E-13 | 1.40423178 | 0.302 | 0.193 | 1.1302E-09 |

| | | | | | | | |
|---|---|---|---|---|---|---|---|
| Fibroblast | Col11a12 | 11 | 5.8644E-13 | 1.22685152 | 0.471 | 0.381 | 1.1729E-09 |
| Fibroblast | Klhl293 | 11 | 6.1443E-13 | 0.43945625 | 0.656 | 0.438 | 1.2289E-09 |
| Fibroblast | Ptafr4 | 11 | 6.2394E-13 | 1.14190724 | 0.442 | 0.296 | 1.2479E-09 |
| Fibroblast | Pkhd1l11 | 11 | 6.3466E-13 | 1.79879734 | 0.299 | 0.15 | 1.2693E-09 |
| Fibroblast | AABR070252 | 11 | 8.6407E-13 | 0.44526301 | 0.893 | 0.668 | 1.7281E-09 |
| Fibroblast | Zbtb7c5 | 11 | 1.0454E-12 | 0.4353293 | 0.49 | 0.331 | 2.0907E-09 |
| Fibroblast | AABR070635 | 11 | 1.0625E-12 | 1.05445934 | 0.39 | 0.246 | 2.125E-09 |
| Fibroblast | Slc24a34 | 11 | 1.2743E-12 | 0.57921078 | 0.588 | 0.402 | 2.5487E-09 |
| Fibroblast | Ano41 | 11 | 1.3393E-12 | 1.82880939 | 0.318 | 0.196 | 2.6785E-09 |
| Fibroblast | Ednrb2 | 11 | 1.4472E-12 | 1.8449692 | 0.269 | 0.193 | 2.8943E-09 |
| Fibroblast | Nckap1l2 | 11 | 1.5494E-12 | 1.52337864 | 0.373 | 0.21 | 3.0989E-09 |
| Fibroblast | Ldb22 | 11 | 1.6619E-12 | 0.99678838 | 0.399 | 0.239 | 3.3238E-09 |
| Fibroblast | Cyba2 | 11 | 1.8464E-12 | 0.61565521 | 0.497 | 0.336 | 3.6929E-09 |
| Fibroblast | Dkk34 | 11 | 1.8491E-12 | 0.27939598 | 0.721 | 0.556 | 3.6982E-09 |
| Fibroblast | Rspo23 | 11 | 1.8913E-12 | 1.06361524 | 0.26 | 0.111 | 3.7826E-09 |
| Fibroblast | Cthrc14 | 11 | 1.9003E-12 | 0.57534394 | 0.5 | 0.324 | 3.8007E-09 |
| Fibroblast | Pstpip2 | 11 | 2.0092E-12 | 0.56909813 | 0.328 | 0.188 | 4.0183E-09 |
| Fibroblast | Sema3d4 | 11 | 2.1721E-12 | 0.86003651 | 0.549 | 0.392 | 4.3442E-09 |
| Fibroblast | Adamts64 | 11 | 2.5389E-12 | 0.43166092 | 0.867 | 0.628 | 5.0777E-09 |
| Fibroblast | Lgr6 | 11 | 2.6543E-12 | 1.33595575 | 0.325 | 0.219 | 5.3085E-09 |
| Fibroblast | Dgki2 | 11 | 3.2803E-12 | 0.27965762 | 0.487 | 0.291 | 6.5606E-09 |
| Fibroblast | Tpx21 | 11 | 3.6764E-12 | 0.48253834 | 0.39 | 0.202 | 7.3527E-09 |
| Fibroblast | Has23 | 11 | 3.9553E-12 | 0.26498464 | 0.555 | 0.376 | 7.9105E-09 |
| Fibroblast | Atp1b12 | 11 | 4.1236E-12 | 0.41040264 | 0.487 | 0.335 | 8.2472E-09 |
| Fibroblast | Entpd12 | 11 | 4.4004E-12 | 1.11981602 | 0.49 | 0.366 | 8.8008E-09 |
| Fibroblast | Slit35 | 11 | 4.4488E-12 | 0.30772793 | 0.623 | 0.477 | 8.8977E-09 |
| Fibroblast | Fam78b2 | 11 | 4.7186E-12 | 1.23568308 | 0.289 | 0.15 | 9.4371E-09 |
| Fibroblast | Srgap13 | 11 | 5.0365E-12 | 0.43436904 | 0.834 | 0.647 | 1.0073E-08 |
| Fibroblast | Fgd22 | 11 | 5.3152E-12 | 0.81474691 | 0.38 | 0.279 | 1.063E-08 |
| Fibroblast | RGD1559482 | 11 | 6.0453E-12 | 1.23743366 | 0.341 | 0.185 | 1.2091E-08 |
| Fibroblast | Smad62 | 11 | 6.1165E-12 | 0.41465904 | 0.643 | 0.422 | 1.2233E-08 |

| | | | | | | | |
|---|---|---|---|---|---|---|---|
| Fibroblast | Cttnbp22 | 11 | 6.1376E-12 | 1.32682207 | 0.396 | 0.283 | 1.2275E-08 |
| Fibroblast | Cpxm23 | 11 | 6.2927E-12 | 0.73718791 | 0.39 | 0.241 | 1.2585E-08 |
| Fibroblast | Pcdh195 | 11 | 7.5983E-12 | 0.71996792 | 0.519 | 0.371 | 1.5197E-08 |
| Fibroblast | Fabp43 | 11 | 7.9351E-12 | 0.46353814 | 0.38 | 0.25 | 1.587E-08 |
| Fibroblast | AABR070440 | 11 | 8.6754E-12 | 0.33435041 | 0.565 | 0.385 | 1.7351E-08 |
| Fibroblast | Alcam2 | 11 | 8.7508E-12 | 0.31923295 | 0.494 | 0.322 | 1.7502E-08 |
| Fibroblast | Ptprn21 | 11 | 9.1301E-12 | 0.32239139 | 0.334 | 0.134 | 1.826E-08 |
| Fibroblast | Atf32 | 11 | 9.9122E-12 | 0.27963025 | 0.302 | 0.146 | 1.9824E-08 |
| Fibroblast | Deptor2 | 11 | 1.047E-11 | 0.69764295 | 0.474 | 0.371 | 2.0941E-08 |
| Fibroblast | Sytl23 | 11 | 1.0531E-11 | 1.00472361 | 0.295 | 0.185 | 2.1063E-08 |
| Fibroblast | Otulinl2 | 11 | 1.1976E-11 | 1.12637805 | 0.409 | 0.247 | 2.3952E-08 |
| Fibroblast | Slc43a22 | 11 | 1.2425E-11 | 0.55157053 | 0.451 | 0.288 | 2.4849E-08 |
| Fibroblast | Jph22 | 11 | 1.2743E-11 | 0.48454414 | 0.422 | 0.289 | 2.5487E-08 |
| Fibroblast | Bend62 | 11 | 1.3071E-11 | 0.67471192 | 0.461 | 0.304 | 2.6141E-08 |
| Fibroblast | Sgcg3 | 11 | 1.3479E-11 | 0.83343742 | 0.445 | 0.311 | 2.6959E-08 |
| Fibroblast | Mmp166 | 11 | 1.3671E-11 | 0.40376285 | 0.477 | 0.305 | 2.7342E-08 |
| Fibroblast | Ca41 | 11 | 1.4256E-11 | 2.93171219 | 0.282 | 0.198 | 2.8512E-08 |
| Fibroblast | Tox31 | 11 | 1.4328E-11 | 1.38405795 | 0.364 | 0.246 | 2.8656E-08 |
| Fibroblast | P4ha33 | 11 | 1.5152E-11 | 1.15066187 | 0.318 | 0.222 | 3.0304E-08 |
| Fibroblast | Gja13 | 11 | 1.5904E-11 | 0.39020513 | 0.526 | 0.353 | 3.1809E-08 |
| Fibroblast | Ap1s31 | 11 | 1.6103E-11 | 0.99019991 | 0.468 | 0.332 | 3.2206E-08 |
| Fibroblast | Pik3cd2 | 11 | 1.7978E-11 | 1.27468447 | 0.429 | 0.305 | 3.5957E-08 |
| Fibroblast | Mctp23 | 11 | 2.0593E-11 | 1.0350769 | 0.38 | 0.245 | 4.1185E-08 |
| Fibroblast | Ccn12 | 11 | 2.0672E-11 | 0.4706863 | 0.432 | 0.274 | 4.1344E-08 |
| Fibroblast | Clec7a1 | 11 | 2.0923E-11 | 2.32186925 | 0.273 | 0.144 | 4.1845E-08 |
| Fibroblast | Hmox14 | 11 | 2.5324E-11 | 0.38535159 | 0.315 | 0.18 | 5.0649E-08 |
| Fibroblast | Tpt11 | 11 | 2.7007E-11 | 0.27896915 | 0.477 | 0.276 | 5.4015E-08 |
| Fibroblast | Anpep2 | 11 | 2.7725E-11 | 0.39148966 | 0.601 | 0.424 | 5.545E-08 |
| Fibroblast | Wipf31 | 11 | 2.8662E-11 | 1.28502576 | 0.38 | 0.301 | 5.7324E-08 |
| Fibroblast | Plaur4 | 11 | 2.9567E-11 | 0.27705184 | 0.617 | 0.446 | 5.9134E-08 |
| Fibroblast | Emid11 | 11 | 3.3168E-11 | 1.34174557 | 0.308 | 0.187 | 6.6336E-08 |

| | | | | | | | |
|---|---|---|---|---|---|---|---|
| Fibroblast | Asgr2 | 11 | 3.7941E-11 | 0.60172937 | 0.315 | 0.167 | 7.5883E-08 |
| Fibroblast | Serpine14 | 11 | 4.2976E-11 | 0.54568416 | 0.497 | 0.343 | 8.5952E-08 |
| Fibroblast | Dmtn | 11 | 4.6951E-11 | 1.09001977 | 0.292 | 0.175 | 9.3902E-08 |
| Fibroblast | Diaph32 | 11 | 4.85E-11 | 0.27075532 | 0.422 | 0.291 | 9.7E-08 |
| Fibroblast | Fcgr2b | 11 | 5.1161E-11 | 1.56045307 | 0.305 | 0.191 | 1.0232E-07 |
| Fibroblast | AABR070544 | 11 | 5.3117E-11 | 1.11661848 | 0.429 | 0.302 | 1.0623E-07 |
| Fibroblast | Slc20a23 | 11 | 5.5505E-11 | 0.4240572 | 0.562 | 0.425 | 1.1101E-07 |
| Fibroblast | Egflam1 | 11 | 6.4319E-11 | 0.84884593 | 0.312 | 0.187 | 1.2864E-07 |
| Fibroblast | Pdgfc2 | 11 | 7.3815E-11 | 0.67882601 | 0.419 | 0.271 | 1.4763E-07 |
| Fibroblast | Scg21 | 11 | 7.7034E-11 | 0.77476831 | 0.331 | 0.198 | 1.5407E-07 |
| Fibroblast | Brca12 | 11 | 9.242E-11 | 0.25234883 | 0.364 | 0.215 | 1.8484E-07 |
| Fibroblast | Arhgap151 | 11 | 9.2436E-11 | 0.86047561 | 0.425 | 0.323 | 1.8487E-07 |
| Fibroblast | Pak33 | 11 | 9.3027E-11 | 1.02452972 | 0.347 | 0.275 | 1.8605E-07 |
| Fibroblast | Mpp7 | 11 | 1.0932E-10 | 0.31422757 | 0.38 | 0.244 | 2.1864E-07 |
| Fibroblast | Icam1 | 11 | 1.1184E-10 | 0.6018481 | 0.354 | 0.226 | 2.2369E-07 |
| Fibroblast | Mrc11 | 11 | 1.1661E-10 | 0.9813314 | 0.338 | 0.221 | 2.3323E-07 |
| Fibroblast | Pamr14 | 11 | 1.272E-10 | 0.81977125 | 0.487 | 0.359 | 2.5441E-07 |
| Fibroblast | Tnfaip8l34 | 11 | 1.3158E-10 | 0.43431168 | 0.445 | 0.304 | 2.6316E-07 |
| Fibroblast | Nr1h43 | 11 | 1.4179E-10 | 0.61791009 | 0.513 | 0.354 | 2.8358E-07 |
| Fibroblast | Fhod32 | 11 | 1.437E-10 | 0.35565041 | 0.614 | 0.434 | 2.874E-07 |
| Fibroblast | Dnah111 | 11 | 1.4855E-10 | 0.30497361 | 0.39 | 0.251 | 2.971E-07 |
| Fibroblast | Ifitm33 | 11 | 1.7577E-10 | 0.30142808 | 0.455 | 0.334 | 3.5155E-07 |
| Fibroblast | Lilrb4 | 11 | 2.0663E-10 | 1.89738074 | 0.37 | 0.25 | 4.1326E-07 |
| Fibroblast | Dpf33 | 11 | 2.242E-10 | 0.89307894 | 0.393 | 0.262 | 4.4841E-07 |
| Fibroblast | Bgn3 | 11 | 2.2452E-10 | 0.4141457 | 0.659 | 0.46 | 4.4905E-07 |
| Fibroblast | Acta22 | 11 | 2.3481E-10 | 0.79220501 | 0.39 | 0.271 | 4.6961E-07 |
| Fibroblast | AABR070428 | 11 | 2.7453E-10 | 0.36466108 | 0.37 | 0.225 | 5.4905E-07 |
| Fibroblast | Igfbp42 | 11 | 3.1227E-10 | 0.43317759 | 0.571 | 0.433 | 6.2455E-07 |
| Fibroblast | Dok51 | 11 | 3.206E-10 | 0.38075475 | 0.312 | 0.177 | 6.412E-07 |
| Fibroblast | Lyz22 | 11 | 3.3993E-10 | 0.41164215 | 0.328 | 0.207 | 6.7986E-07 |
| Fibroblast | Pls34 | 11 | 3.5736E-10 | 0.37161606 | 0.873 | 0.674 | 7.1471E-07 |

| | | | | | | | |
|---|---|---|---|---|---|---|---|
| Fibroblast | LOC1003614 | 11 | 3.7465E-10 | 0.37587301 | 0.523 | 0.331 | 7.493E-07 |
| Fibroblast | Pfkfb32 | 11 | 3.7901E-10 | 0.40611395 | 0.497 | 0.34 | 7.5802E-07 |
| Fibroblast | AABR070275 | 11 | 3.9846E-10 | 0.73163072 | 0.429 | 0.302 | 7.9691E-07 |
| Fibroblast | Baiap2l12 | 11 | 4.2819E-10 | 0.37561878 | 0.432 | 0.276 | 8.5638E-07 |
| Fibroblast | Tec4 | 11 | 4.7514E-10 | 1.18528123 | 0.357 | 0.257 | 9.5028E-07 |
| Fibroblast | Svep13 | 11 | 4.7743E-10 | 0.27897161 | 0.653 | 0.472 | 9.5485E-07 |
| Fibroblast | Pdlim53 | 11 | 5.1586E-10 | 0.37961048 | 0.873 | 0.668 | 1.0317E-06 |
| Fibroblast | Slc16a71 | 11 | 5.6564E-10 | 0.93600842 | 0.347 | 0.215 | 1.1313E-06 |
| Fibroblast | Mark11 | 11 | 5.7118E-10 | 0.4161364 | 0.364 | 0.239 | 1.1424E-06 |
| Fibroblast | Nr4a15 | 11 | 6.0849E-10 | 1.13208064 | 0.383 | 0.281 | 1.217E-06 |
| Fibroblast | Bace22 | 11 | 6.2501E-10 | 0.29963113 | 0.464 | 0.348 | 1.25E-06 |
| Fibroblast | Cntn23 | 11 | 6.2563E-10 | 0.75131323 | 0.325 | 0.205 | 1.2513E-06 |
| Fibroblast | Mid13 | 11 | 6.5325E-10 | 0.39217009 | 0.799 | 0.589 | 1.3065E-06 |
| Fibroblast | Il332 | 11 | 6.5597E-10 | 0.55946055 | 0.451 | 0.331 | 1.3119E-06 |
| Fibroblast | Pdlim33 | 11 | 6.7814E-10 | 0.38981705 | 0.662 | 0.481 | 1.3563E-06 |
| Fibroblast | Tbc1d4 | 11 | 7.6473E-10 | 0.37835775 | 0.519 | 0.377 | 1.5295E-06 |
| Fibroblast | Pdcd1lg21 | 11 | 8.1965E-10 | 2.76197682 | 0.315 | 0.261 | 1.6393E-06 |
| Fibroblast | Ctsd3 | 11 | 9.0249E-10 | 0.25138082 | 0.341 | 0.221 | 1.805E-06 |
| Fibroblast | Ptpro | 11 | 9.9737E-10 | 0.46024399 | 0.334 | 0.203 | 1.9947E-06 |
| Fibroblast | Olfml2a1 | 11 | 1.0572E-09 | 0.26993624 | 0.286 | 0.183 | 2.1143E-06 |
| Fibroblast | Amd1 | 11 | 1.0999E-09 | 0.54656736 | 0.516 | 0.401 | 2.1998E-06 |
| Fibroblast | Apobec12 | 11 | 1.2743E-09 | 1.58339819 | 0.305 | 0.243 | 2.5486E-06 |
| Fibroblast | Vcam14 | 11 | 1.3871E-09 | 0.33759282 | 0.461 | 0.311 | 2.7741E-06 |
| Fibroblast | Col8a24 | 11 | 1.4637E-09 | 0.26087478 | 0.558 | 0.425 | 2.9274E-06 |
| Fibroblast | AABR070410 | 11 | 1.4685E-09 | 1.17926233 | 0.273 | 0.204 | 2.937E-06 |
| Fibroblast | Cd41 | 11 | 1.5144E-09 | 1.05030909 | 0.321 | 0.204 | 3.0287E-06 |
| Fibroblast | Unc5c1 | 11 | 1.9225E-09 | 1.93051798 | 0.286 | 0.159 | 3.845E-06 |
| Fibroblast | Ankh1 | 11 | 2.0445E-09 | 0.47956861 | 0.523 | 0.363 | 4.0891E-06 |
| Fibroblast | Pkia3 | 11 | 2.1016E-09 | 0.4289426 | 0.383 | 0.274 | 4.2032E-06 |
| Fibroblast | Ablim23 | 11 | 2.2453E-09 | 0.58722716 | 0.354 | 0.226 | 4.4906E-06 |
| Fibroblast | AABR070311 | 11 | 2.2543E-09 | 1.05410286 | 0.325 | 0.21 | 4.5086E-06 |

| | | | | | | | |
|---|---|---|---|---|---|---|---|
| Fibroblast | Nckap51 | 11 | 2.2799E-09 | 0.33528924 | 0.464 | 0.321 | 4.5597E-06 |
| Fibroblast | Kcnab14 | 11 | 2.3539E-09 | 0.42036272 | 0.468 | 0.354 | 4.7078E-06 |
| Fibroblast | Unc13b1 | 11 | 2.4156E-09 | 0.7415619 | 0.302 | 0.224 | 4.8313E-06 |
| Fibroblast | Mx2 | 11 | 2.5638E-09 | 0.87455625 | 0.282 | 0.224 | 5.1276E-06 |
| Fibroblast | Naca3 | 11 | 2.7047E-09 | 0.25730309 | 0.468 | 0.319 | 5.4094E-06 |
| Fibroblast | Lbh2 | 11 | 2.762E-09 | 0.33194551 | 0.672 | 0.492 | 5.5239E-06 |
| Fibroblast | Cdo1 | 11 | 2.7643E-09 | 0.76037674 | 0.308 | 0.17 | 5.5286E-06 |
| Fibroblast | Grk33 | 11 | 3.2341E-09 | 1.81649804 | 0.347 | 0.235 | 6.4683E-06 |
| Fibroblast | Pde1c3 | 11 | 3.4086E-09 | 1.22975614 | 0.341 | 0.249 | 6.8173E-06 |
| Fibroblast | Ugdh1 | 11 | 4.0705E-09 | 0.37234386 | 0.519 | 0.387 | 8.1409E-06 |
| Fibroblast | Cldn152 | 11 | 4.1002E-09 | 1.08923332 | 0.315 | 0.252 | 8.2004E-06 |
| Fibroblast | Rtn4rl13 | 11 | 4.2916E-09 | 0.77682727 | 0.286 | 0.173 | 8.5832E-06 |
| Fibroblast | Sntb13 | 11 | 4.3266E-09 | 0.66438133 | 0.393 | 0.297 | 8.6532E-06 |
| Fibroblast | Rnf144b1 | 11 | 4.4802E-09 | 0.54052309 | 0.419 | 0.329 | 8.9605E-06 |
| Fibroblast | Gnao13 | 11 | 5.4438E-09 | 0.36444426 | 0.468 | 0.347 | 1.0888E-05 |
| Fibroblast | Anks1b2 | 11 | 5.9456E-09 | 0.90044014 | 0.357 | 0.229 | 1.1891E-05 |
| Fibroblast | Ptgs23 | 11 | 6.3571E-09 | 0.91282546 | 0.403 | 0.324 | 1.2714E-05 |
| Fibroblast | Kcnk132 | 11 | 6.9524E-09 | 0.42939777 | 0.39 | 0.257 | 1.3905E-05 |
| Fibroblast | Prkcb3 | 11 | 7.5118E-09 | 1.28798784 | 0.357 | 0.276 | 1.5024E-05 |
| Fibroblast | Igsf111 | 11 | 7.7938E-09 | 0.54343458 | 0.328 | 0.131 | 1.5588E-05 |
| Fibroblast | Pparg4 | 11 | 8.0779E-09 | 0.4728552 | 0.36 | 0.274 | 1.6156E-05 |
| Fibroblast | Skap12 | 11 | 8.3788E-09 | 1.16644714 | 0.334 | 0.238 | 1.6758E-05 |
| Fibroblast | Myo1f2 | 11 | 8.6514E-09 | 0.31853976 | 0.377 | 0.279 | 1.7303E-05 |
| Fibroblast | LOC6900453 | 11 | 9.3679E-09 | 1.21465029 | 0.344 | 0.256 | 1.8736E-05 |
| Fibroblast | Cyp26b13 | 11 | 1.0503E-08 | 0.69995343 | 0.315 | 0.22 | 2.1007E-05 |
| Fibroblast | Prg42 | 11 | 1.1657E-08 | 1.26744149 | 0.334 | 0.225 | 2.3313E-05 |
| Fibroblast | AABR070581 | 11 | 1.2192E-08 | 0.68054048 | 0.364 | 0.235 | 2.4384E-05 |
| Fibroblast | Sfrp24 | 11 | 1.263E-08 | 0.48428254 | 0.364 | 0.257 | 2.5259E-05 |
| Fibroblast | Dlgap52 | 11 | 1.3327E-08 | 0.58278758 | 0.347 | 0.231 | 2.6654E-05 |
| Fibroblast | Dkk22 | 11 | 1.3803E-08 | 0.40710598 | 0.474 | 0.359 | 2.7606E-05 |
| Fibroblast | Rap1gap23 | 11 | 1.39E-08 | 0.34420258 | 0.338 | 0.258 | 2.78E-05 |

| | | | | | | | |
|---|---|---|---|---|---|---|---|
| Fibroblast | P3h21 | 11 | 1.9708E-08 | 0.54534088 | 0.331 | 0.258 | 3.9416E-05 |
| Fibroblast | Homer13 | 11 | 2.0589E-08 | 0.28745547 | 0.49 | 0.39 | 4.1178E-05 |
| Fibroblast | Neurl13 | 11 | 2.2864E-08 | 0.65901109 | 0.321 | 0.232 | 4.5727E-05 |
| Fibroblast | Gpnmb3 | 11 | 2.9682E-08 | 1.0282268 | 0.425 | 0.305 | 5.9363E-05 |
| Fibroblast | Mt12 | 11 | 3.2761E-08 | 0.56561531 | 0.406 | 0.289 | 6.5522E-05 |
| Fibroblast | Atp2b22 | 11 | 4.1221E-08 | 0.73326872 | 0.357 | 0.255 | 8.2442E-05 |
| Fibroblast | Nkd23 | 11 | 4.4608E-08 | 0.62136002 | 0.393 | 0.315 | 8.9215E-05 |
| Fibroblast | Oprm14 | 11 | 5.1392E-08 | 0.82388638 | 0.341 | 0.256 | 0.00010278 |
| Fibroblast | Tmem1003 | 11 | 5.4818E-08 | 0.51770708 | 0.432 | 0.363 | 0.00010964 |
| Fibroblast | Fmn12 | 11 | 5.8542E-08 | 0.39721498 | 0.477 | 0.374 | 0.00011708 |
| Fibroblast | LOC6877803 | 11 | 7.8202E-08 | 0.29036034 | 0.377 | 0.254 | 0.0001564 |
| Fibroblast | Igfbp64 | 11 | 7.8878E-08 | 0.43496567 | 0.422 | 0.294 | 0.00015776 |
| Fibroblast | Scarb12 | 11 | 1.108E-07 | 1.33100587 | 0.328 | 0.265 | 0.00022161 |
| Fibroblast | Rcan14 | 11 | 1.1732E-07 | 0.28786793 | 0.377 | 0.289 | 0.00023464 |
| Fibroblast | Pip5k1b1 | 11 | 1.2543E-07 | 0.49535922 | 0.269 | 0.193 | 0.00025085 |
| Fibroblast | B2m2 | 11 | 1.33E-07 | 0.2884654 | 0.575 | 0.431 | 0.00026601 |
| Fibroblast | LOC1009119 | 11 | 1.575E-07 | 0.47600362 | 0.39 | 0.295 | 0.000315 |
| Fibroblast | Gxylt24 | 11 | 1.6269E-07 | 0.26254451 | 0.896 | 0.716 | 0.00032538 |
| Fibroblast | Tspan181 | 11 | 1.6562E-07 | 0.3069735 | 0.662 | 0.52 | 0.00033124 |
| Fibroblast | Reep14 | 11 | 2.2073E-07 | 0.5465857 | 0.338 | 0.246 | 0.00044146 |
| Fibroblast | Slfn42 | 11 | 2.2737E-07 | 0.89707304 | 0.377 | 0.302 | 0.00045474 |
| Fibroblast | Arhgef262 | 11 | 2.3244E-07 | 0.28167531 | 0.302 | 0.25 | 0.00046489 |
| Fibroblast | Ripor21 | 11 | 2.329E-07 | 0.59498482 | 0.367 | 0.281 | 0.00046579 |
| Fibroblast | Fbn21 | 11 | 2.481E-07 | 0.30654627 | 0.487 | 0.384 | 0.0004962 |
| Fibroblast | Slc25a131 | 11 | 2.5136E-07 | 0.3728273 | 0.37 | 0.285 | 0.00050272 |
| Fibroblast | Pik3ap12 | 11 | 2.5811E-07 | 0.82739013 | 0.364 | 0.28 | 0.00051621 |
| Fibroblast | Gna153 | 11 | 2.6186E-07 | 1.57731057 | 0.383 | 0.274 | 0.00052371 |
| Fibroblast | Cotl14 | 11 | 2.6319E-07 | 0.3633428 | 0.416 | 0.329 | 0.00052638 |
| Fibroblast | Cacna2d22 | 11 | 2.6493E-07 | 0.63805725 | 0.318 | 0.181 | 0.00052986 |
| Fibroblast | Trib31 | 11 | 2.9383E-07 | 0.59902468 | 0.269 | 0.22 | 0.00058766 |
| Fibroblast | Ros11 | 11 | 4.4132E-07 | 0.36382724 | 0.331 | 0.218 | 0.00088264 |

| | | | | | | | |
|---|---|---|---|---|---|---|---|
| Fibroblast | Prox12 | 11 | 4.8563E-07 | 0.95995013 | 0.312 | 0.213 | 0.00097125 |
| Fibroblast | Spp14 | 11 | 5.8067E-07 | 0.38913668 | 0.312 | 0.256 | 0.00116135 |
| Fibroblast | Slc12a21 | 11 | 6.6769E-07 | 0.69248024 | 0.344 | 0.269 | 0.00133539 |
| Fibroblast | Wdfy42 | 11 | 7.8451E-07 | 0.76998167 | 0.425 | 0.344 | 0.00156902 |
| Fibroblast | Vash24 | 11 | 8.7292E-07 | 0.40651006 | 0.315 | 0.238 | 0.00174584 |
| Fibroblast | Kcnt22 | 11 | 9.5414E-07 | 0.5368249 | 0.442 | 0.38 | 0.00190829 |
| Fibroblast | Shtn11 | 11 | 9.6763E-07 | 0.5251071 | 0.256 | 0.185 | 0.00193526 |
| Fibroblast | LOC1009095 | 11 | 1.1861E-06 | 0.35874671 | 0.318 | 0.261 | 0.00237218 |
| Fibroblast | Aoah2 | 11 | 1.538E-06 | 1.01016709 | 0.341 | 0.265 | 0.003076 |
| Fibroblast | Rimbp23 | 11 | 1.6494E-06 | 0.57506111 | 0.328 | 0.27 | 0.00329887 |
| Fibroblast | Cybb2 | 11 | 1.8218E-06 | 0.45414582 | 0.302 | 0.261 | 0.00364358 |
| Fibroblast | Abca173 | 11 | 2.0298E-06 | 1.98304623 | 0.253 | 0.247 | 0.00405954 |
| Fibroblast | Cd552 | 11 | 2.7032E-06 | 1.02713181 | 0.412 | 0.318 | 0.00540638 |
| Fibroblast | P2ry142 | 11 | 3.3683E-06 | 0.34576673 | 0.292 | 0.205 | 0.00673669 |
| Fibroblast | Kcng23 | 11 | 4.5061E-06 | 0.61022839 | 0.302 | 0.274 | 0.0090121 |
| Fibroblast | Kif18b2 | 11 | 4.578E-06 | 0.56120824 | 0.253 | 0.149 | 0.00915601 |
| Fibroblast | Cgnl11 | 11 | 4.7052E-06 | 1.24030785 | 0.416 | 0.367 | 0.00941049 |
| Fibroblast | Maob2 | 11 | 5.6562E-06 | 1.30970624 | 0.364 | 0.295 | 0.01131233 |
| Fibroblast | Tafa21 | 11 | 5.9135E-06 | 0.3349468 | 0.26 | 0.137 | 0.01182705 |
| Fibroblast | Slc22a232 | 11 | 6.2815E-06 | 0.58406733 | 0.302 | 0.26 | 0.01256304 |
| Fibroblast | Tox4 | 11 | 6.4517E-06 | 0.46378578 | 0.289 | 0.255 | 0.01290331 |
| Fibroblast | Dact21 | 11 | 7.9662E-06 | 0.6220628 | 0.347 | 0.25 | 0.01593232 |
| Fibroblast | Mertk2 | 11 | 8.0116E-06 | 0.91083705 | 0.25 | 0.218 | 0.01602315 |
| Fibroblast | Nav21 | 11 | 9.2704E-06 | 0.35430751 | 0.273 | 0.158 | 0.01854073 |
| Fibroblast | Bcar32 | 11 | 1.1252E-05 | 0.59119711 | 0.318 | 0.245 | 0.02250423 |
| Fibroblast | Nlrp31 | 11 | 1.4497E-05 | 1.05210462 | 0.351 | 0.26 | 0.02899415 |
| Fibroblast | Col8a13 | 11 | 1.9742E-05 | 0.34042438 | 0.964 | 0.871 | 0.03948338 |
| Fibroblast | Zbtb163 | 12 | 5.2762E-13 | 1.99683654 | 0.703 | 0.44 | 1.0552E-09 |
| Fibroblast | Ctss3 | 12 | 9.9231E-12 | 1.11146104 | 0.051 | 0.276 | 1.9846E-08 |
| Fibroblast | Aox31 | 12 | 3.4689E-09 | 3.42857087 | 0.542 | 0.289 | 6.9379E-06 |
| Fibroblast | Ltbp14 | 12 | 3.7101E-09 | 0.96322179 | 0.814 | 0.669 | 7.4202E-06 |

| | | | | | | | |
|---|---|---|---|---|---|---|---|
| Fibroblast | Art33 | 12 | 4.591E-09 | 0.59512947 | 0.093 | 0.323 | 9.182E-06 |
| Fibroblast | Anln1 | 12 | 1.4441E-08 | 0.49060495 | 0.051 | 0.254 | 2.8882E-05 |
| Fibroblast | Atp2b23 | 12 | 3.6897E-08 | 1.05687389 | 0.034 | 0.261 | 7.3794E-05 |
| Fibroblast | Tec5 | 12 | 6.273E-08 | 0.68634222 | 0.042 | 0.263 | 0.00012546 |
| Fibroblast | Pcdh72 | 12 | 7.6718E-08 | 1.02924657 | 0.076 | 0.279 | 0.00015344 |
| Fibroblast | Tbx203 | 12 | 2.5996E-07 | 1.29973247 | 0.712 | 0.542 | 0.00051991 |
| Fibroblast | Fgfr23 | 12 | 3.1223E-07 | 1.00063766 | 0.085 | 0.317 | 0.00062446 |
| Fibroblast | Ptger34 | 12 | 7.0135E-07 | 0.45572204 | 0.093 | 0.264 | 0.00140269 |
| Fibroblast | Agmo4 | 12 | 8.2842E-07 | 0.9740843 | 0.076 | 0.254 | 0.00165683 |
| Fibroblast | Tox32 | 12 | 1.5622E-06 | 0.31395183 | 0.093 | 0.252 | 0.00312449 |
| Fibroblast | Slc1a33 | 12 | 1.9327E-06 | 1.00363884 | 0.136 | 0.281 | 0.00386541 |
| Fibroblast | Prkcz3 | 12 | 1.9649E-06 | 0.26086016 | 0.136 | 0.348 | 0.0039298 |
| Fibroblast | Lpl3 | 12 | 2.6267E-06 | 0.73723165 | 0.153 | 0.344 | 0.00525347 |
| Fibroblast | Ildr23 | 12 | 2.9293E-06 | 0.26243209 | 0.169 | 0.33 | 0.00585861 |
| Fibroblast | Arhgap152 | 12 | 2.9347E-06 | 0.62198517 | 0.144 | 0.329 | 0.00586931 |
| Fibroblast | Wdfy43 | 12 | 3.437E-06 | 0.85182159 | 0.153 | 0.349 | 0.00687395 |
| Fibroblast | Cit3 | 12 | 5.1905E-06 | 0.54675148 | 0.144 | 0.384 | 0.01038103 |
| Fibroblast | Nlrp32 | 12 | 5.6812E-06 | 1.10561741 | 0.11 | 0.265 | 0.01136239 |
| Fibroblast | Adamts33 | 12 | 7.5943E-06 | 0.43170096 | 0.136 | 0.304 | 0.01518858 |
| Fibroblast | C62 | 12 | 8.7073E-06 | 0.80798465 | 0.102 | 0.324 | 0.01741457 |
| Fibroblast | Itga24 | 12 | 8.7356E-06 | 0.80272846 | 0.076 | 0.25 | 0.01747124 |
| Fibroblast | Stab13 | 12 | 9.214E-06 | 0.73847877 | 0.169 | 0.251 | 0.01842799 |
| Fibroblast | Epb41l4b1 | 12 | 1.2425E-05 | 0.59533998 | 0.144 | 0.28 | 0.02485099 |
| Fibroblast | Baiap2l13 | 12 | 1.2568E-05 | 0.72492202 | 0.161 | 0.283 | 0.02513586 |
| Fibroblast | Lilrb41 | 12 | 1.4427E-05 | 0.6327592 | 0.127 | 0.256 | 0.0288547 |
| Fibroblast | Slc22a233 | 12 | 1.6432E-05 | 0.42718605 | 0.169 | 0.263 | 0.03286383 |
| Fibroblast | Tenm43 | 12 | 1.778E-05 | 0.3847682 | 0.144 | 0.254 | 0.03555911 |
| Fibroblast | Sned12 | 12 | 2.0477E-05 | 1.07059353 | 0.178 | 0.344 | 0.04095436 |
| Fibroblast | Tmem822 | 12 | 2.2544E-05 | 0.28681692 | 0.203 | 0.319 | 0.04508757 |
| Fibroblast | Myom21 | 12 | 2.2837E-05 | 0.65367126 | 0.169 | 0.299 | 0.04567386 |
| Fibroblast | Osr13 | 12 | 2.3345E-05 | 0.3445436 | 0.203 | 0.406 | 0.04668967 |

| | | | | | | | |
|---|---|---|---|---|---|---|---|
| Fibroblast | Macrod11 | 12 | 2.4981E-05 | 0.51438904 | 0.203 | 0.302 | 0.04996155 |
| T-cell | Gpc6 | 0 | 3.46E-24 | 2.2530793 | 0.987 | 0.511 | 6.92E-21 |
| T-cell | C7 | 0 | 1.53E-23 | 3.36093525 | 0.885 | 0.305 | 3.06E-20 |
| T-cell | C1qtnf7 | 0 | 1.40E-21 | 3.03933898 | 0.885 | 0.453 | 2.80E-18 |
| T-cell | Ccdc80 | 0 | 1.55E-21 | 2.63141727 | 0.923 | 0.405 | 3.10E-18 |
| T-cell | Clmp | 0 | 4.29E-21 | 3.03011021 | 0.872 | 0.242 | 8.59E-18 |
| T-cell | Col8a1 | 0 | 1.25E-16 | 2.70168342 | 0.885 | 0.358 | 2.50E-13 |
| T-cell | Adamts2 | 0 | 1.26E-15 | 2.24928678 | 0.821 | 0.311 | 2.52E-12 |
| T-cell | Itgbl1 | 0 | 1.29E-15 | 2.35833575 | 0.821 | 0.442 | 2.58E-12 |
| T-cell | Zfp385d | 0 | 1.68E-15 | 2.08212803 | 0.859 | 0.384 | 3.36E-12 |
| T-cell | Ebf2 | 0 | 3.38E-15 | 2.2031528 | 0.821 | 0.463 | 6.76E-12 |
| T-cell | Bnc2 | 0 | 6.28E-15 | 2.88254653 | 0.756 | 0.2 | 1.26E-11 |
| T-cell | Dcn | 0 | 6.74E-15 | 1.97998569 | 0.808 | 0.274 | 1.35E-11 |
| T-cell | Dlg2 | 0 | 8.80E-15 | 3.01851166 | 0.744 | 0.168 | 1.76E-11 |
| T-cell | Tmeff2 | 0 | 2.72E-14 | 2.88477892 | 0.769 | 0.211 | 5.43E-11 |
| T-cell | Pdzrn3 | 0 | 7.20E-14 | 2.32274934 | 0.795 | 0.342 | 1.44E-10 |
| T-cell | Fbln1 | 0 | 9.16E-14 | 3.22386403 | 0.667 | 0.142 | 1.83E-10 |
| T-cell | Fbn1 | 0 | 9.25E-14 | 2.18417087 | 0.872 | 0.468 | 1.85E-10 |
| T-cell | Col3a1 | 0 | 3.52E-13 | 1.4719285 | 0.949 | 0.658 | 7.04E-10 |
| T-cell | Pdgfra | 0 | 7.62E-13 | 3.19703178 | 0.667 | 0.111 | 1.52E-09 |
| T-cell | Col5a2 | 0 | 8.28E-13 | 1.88120723 | 0.821 | 0.274 | 1.66E-09 |
| T-cell | Col6a6 | 0 | 1.24E-12 | 2.93211479 | 0.744 | 0.416 | 2.48E-09 |
| T-cell | Timp2 | 0 | 1.41E-12 | 1.95693747 | 0.782 | 0.326 | 2.81E-09 |
| T-cell | Cpq | 0 | 2.57E-12 | 2.13104241 | 0.795 | 0.468 | 5.13E-09 |
| T-cell | Postn | 0 | 2.71E-12 | 2.33353497 | 0.705 | 0.268 | 5.42E-09 |
| T-cell | Cacna2d3 | 0 | 3.20E-12 | 3.21100206 | 0.679 | 0.2 | 6.40E-09 |
| T-cell | Abi3bp | 0 | 3.35E-12 | 2.0922145 | 0.756 | 0.321 | 6.70E-09 |
| T-cell | Col4a5 | 0 | 3.69E-12 | 2.62005734 | 0.705 | 0.237 | 7.38E-09 |
| T-cell | Ston2 | 0 | 9.60E-12 | 1.84584152 | 0.756 | 0.237 | 1.92E-08 |
| T-cell | Fndc1 | 0 | 1.59E-11 | 2.29888195 | 0.795 | 0.368 | 3.18E-08 |
| T-cell | Mrc2 | 0 | 2.24E-11 | 2.55125161 | 0.718 | 0.416 | 4.47E-08 |

| | | | | | | | |
|---|---|---|---|---|---|---|---|
| T-cell | Pde1a | 0 | 4.12E-11 | 2.0128357 | 0.731 | 0.332 | 8.23E-08 |
| T-cell | Egfr | 0 | 4.68E-11 | 2.00433222 | 0.731 | 0.416 | 9.36E-08 |
| T-cell | Pcsk6 | 0 | 8.47E-11 | 2.52456138 | 0.654 | 0.253 | 1.69E-07 |
| T-cell | Col14a1 | 0 | 9.11E-11 | 2.71091245 | 0.705 | 0.453 | 1.82E-07 |
| T-cell | Dclk1 | 0 | 3.42E-10 | 1.65524425 | 0.667 | 0.242 | 6.84E-07 |
| T-cell | Mfap5 | 0 | 4.34E-10 | 2.27905881 | 0.731 | 0.426 | 8.69E-07 |
| T-cell | Dapk1 | 0 | 4.66E-10 | 1.6509001 | 0.731 | 0.316 | 9.32E-07 |
| T-cell | Galnt17 | 0 | 2.36E-09 | 3.43223006 | 0.59 | 0.147 | 4.71E-06 |
| T-cell | Adamts5 | 0 | 4.03E-09 | 3.07430643 | 0.615 | 0.147 | 8.05E-06 |
| T-cell | Eda | 0 | 7.50E-09 | 1.93605285 | 0.692 | 0.237 | 1.50E-05 |
| T-cell | Antxr1 | 0 | 1.09E-08 | 2.36436721 | 0.654 | 0.358 | 2.19E-05 |
| T-cell | Mgp | 0 | 1.36E-08 | 1.26770386 | 0.833 | 0.505 | 2.73E-05 |
| T-cell | Plxdc2 | 0 | 1.54E-08 | 1.63952457 | 0.859 | 0.584 | 3.09E-05 |
| T-cell | Map1b | 0 | 1.67E-08 | 1.72246993 | 0.808 | 0.405 | 3.34E-05 |
| T-cell | Ngf | 0 | 1.68E-08 | 2.77561187 | 0.59 | 0.163 | 3.36E-05 |
| T-cell | Glis3 | 0 | 1.88E-08 | 1.78239891 | 0.756 | 0.479 | 3.76E-05 |
| T-cell | Boc | 0 | 2.38E-08 | 2.13080385 | 0.692 | 0.432 | 4.76E-05 |
| T-cell | Cdo1 | 0 | 4.23E-08 | 4.03716049 | 0.321 | 0.058 | 8.45E-05 |
| T-cell | AABR070592 | 0 | 4.99E-08 | 2.39814377 | 0.615 | 0.205 | 9.97E-05 |
| T-cell | Axl | 0 | 6.06E-08 | 1.86255257 | 0.679 | 0.232 | 0.00012117 |
| T-cell | Gria3 | 0 | 6.16E-08 | 2.40650079 | 0.59 | 0.253 | 0.00012319 |
| T-cell | Dpysl3 | 0 | 7.06E-08 | 1.98476307 | 0.667 | 0.3 | 0.00014117 |
| T-cell | Col6a2 | 0 | 7.62E-08 | 1.61984594 | 0.705 | 0.447 | 0.00015247 |
| T-cell | Col1a1 | 0 | 1.13E-07 | 1.24277573 | 0.897 | 0.632 | 0.00022608 |
| T-cell | Asap3 | 0 | 1.28E-07 | 2.0436463 | 0.577 | 0.174 | 0.00025543 |
| T-cell | Creb5 | 0 | 1.56E-07 | 1.47296796 | 0.756 | 0.437 | 0.00031116 |
| T-cell | Foxp2 | 0 | 1.73E-07 | 2.11748272 | 0.615 | 0.179 | 0.00034557 |
| T-cell | Gda | 0 | 2.07E-07 | 1.45014104 | 0.756 | 0.405 | 0.00041357 |
| T-cell | Irs1 | 0 | 2.15E-07 | 3.07878176 | 0.59 | 0.337 | 0.00042961 |
| T-cell | Loxl1 | 0 | 2.19E-07 | 2.05502264 | 0.564 | 0.142 | 0.00043868 |
| T-cell | Adamts19 | 0 | 2.37E-07 | 2.62376274 | 0.577 | 0.153 | 0.00047454 |

| | | | | | | | |
|---|---|---|---|---|---|---|---|
| T-cell | Plod2 | 0 | 3.71E-07 | 1.45865479 | 0.718 | 0.368 | 0.00074291 |
| T-cell | Fstl1 | 0 | 4.14E-07 | 1.66344196 | 0.731 | 0.453 | 0.00082713 |
| T-cell | Abca8a | 0 | 4.46E-07 | 1.85269014 | 0.603 | 0.258 | 0.00089217 |
| T-cell | Pdgfrb | 0 | 5.20E-07 | 1.06258149 | 0.769 | 0.447 | 0.00103932 |
| T-cell | Sulf1 | 0 | 6.53E-07 | 1.31490339 | 0.615 | 0.184 | 0.00130667 |
| T-cell | Adgrd1 | 0 | 7.15E-07 | 2.64434965 | 0.551 | 0.142 | 0.00142946 |
| T-cell | Cdh11 | 0 | 7.57E-07 | 0.95079008 | 0.59 | 0.205 | 0.00151418 |
| T-cell | Adamtsl3 | 0 | 8.14E-07 | 1.87425411 | 0.615 | 0.216 | 0.00162762 |
| T-cell | Gask1b | 0 | 9.43E-07 | 2.44245008 | 0.564 | 0.184 | 0.00188571 |
| T-cell | Htra3 | 0 | 1.30E-06 | 2.01835401 | 0.526 | 0.137 | 0.00260058 |
| T-cell | Plekha7 | 0 | 1.42E-06 | 1.54281844 | 0.603 | 0.2 | 0.00284929 |
| T-cell | Npas2 | 0 | 1.64E-06 | 2.39232357 | 0.538 | 0.137 | 0.00327126 |
| T-cell | Dpt | 0 | 1.80E-06 | 1.94713587 | 0.59 | 0.232 | 0.00360051 |
| T-cell | Vcan | 0 | 2.32E-06 | 2.11428711 | 0.564 | 0.142 | 0.00463324 |
| T-cell | Gpm6b | 0 | 2.86E-06 | 2.55806624 | 0.551 | 0.216 | 0.00572854 |
| T-cell | Tspan5 | 0 | 3.03E-06 | 1.6825911 | 0.654 | 0.274 | 0.00605617 |
| T-cell | Sorcs1 | 0 | 3.31E-06 | 0.69328541 | 0.372 | 0.1 | 0.00662071 |
| T-cell | Pcsk5 | 0 | 3.69E-06 | 2.22916497 | 0.564 | 0.232 | 0.00737932 |
| T-cell | Svep1 | 0 | 4.18E-06 | 3.41363136 | 0.526 | 0.205 | 0.00835846 |
| T-cell | AABR070546 | 0 | 6.68E-06 | 2.95915433 | 0.462 | 0.163 | 0.01335815 |
| T-cell | Mmp2 | 0 | 7.87E-06 | 1.95766044 | 0.551 | 0.132 | 0.01573666 |
| T-cell | Vegfa | 0 | 8.29E-06 | 0.7826983 | 0.718 | 0.395 | 0.0165821 |
| T-cell | Col5a3 | 0 | 1.11E-05 | 2.05035972 | 0.474 | 0.147 | 0.02223842 |
| T-cell | Col4a3 | 0 | 1.34E-05 | 1.53900365 | 0.641 | 0.389 | 0.02673444 |
| T-cell | Igfbp5 | 0 | 1.47E-05 | 2.3012939 | 0.526 | 0.153 | 0.02935319 |
| T-cell | Atp8b4 | 2 | 8.65E-12 | 2.18036176 | 0.949 | 0.493 | 1.73E-08 |
| T-cell | Clnk | 2 | 1.03E-11 | 2.84391435 | 0.897 | 0.528 | 2.06E-08 |
| T-cell | Il2rb | 2 | 1.76E-11 | 2.46038463 | 0.872 | 0.384 | 3.52E-08 |
| T-cell | Mcoln2 | 2 | 1.62E-10 | 3.65710582 | 0.692 | 0.262 | 3.24E-07 |
| T-cell | Rergl | 2 | 7.94E-10 | 3.9293213 | 0.564 | 0.179 | 1.59E-06 |
| T-cell | Ankrd6 | 2 | 2.06E-08 | 2.6151229 | 0.744 | 0.328 | 4.12E-05 |

| | | | | | | | |
|---|---|---|---|---|---|---|---|
| T-cell | Slc26a3 | 2 | 4.41E-08 | 4.5744943 | 0.564 | 0.223 | 8.82E-05 |
| T-cell | Klrc2 | 2 | 1.18E-07 | 2.96800473 | 0.59 | 0.236 | 0.00023626 |
| T-cell | Casr | 2 | 4.09E-07 | 0.81191258 | 0.41 | 0.096 | 0.00081894 |
| T-cell | AC094126.1 | 2 | 4.69E-07 | 0.8568664 | 0.41 | 0.1 | 0.00093823 |
| T-cell | Runx2 | 2 | 4.71E-07 | 1.71318409 | 0.897 | 0.611 | 0.00094248 |
| T-cell | Adamts14 | 2 | 6.47E-07 | 2.20614329 | 0.538 | 0.24 | 0.0012933 |
| T-cell | Mboat2 | 2 | 7.63E-07 | 2.21718579 | 0.744 | 0.345 | 0.00152579 |
| T-cell | Ncr1 | 2 | 9.06E-07 | 2.73098625 | 0.769 | 0.437 | 0.00181157 |
| T-cell | Gpr174 | 2 | 1.19E-06 | 3.47210977 | 0.667 | 0.301 | 0.00238955 |
| T-cell | AABR070014 | 2 | 1.53E-06 | 2.03483918 | 0.538 | 0.197 | 0.00305547 |
| T-cell | Tafa2 | 2 | 1.55E-06 | 2.91763993 | 0.564 | 0.214 | 0.00310576 |
| T-cell | Fbxw10 | 2 | 1.66E-06 | 5.08588163 | 0.564 | 0.284 | 0.0033261 |
| T-cell | Il18r1 | 2 | 1.76E-06 | 1.66281892 | 0.769 | 0.415 | 0.00352748 |
| T-cell | Efcab11 | 2 | 1.96E-06 | 3.44294425 | 0.564 | 0.179 | 0.00392256 |
| T-cell | Klri1 | 2 | 2.23E-06 | 1.62690612 | 0.821 | 0.454 | 0.00446775 |
| T-cell | Ly49s5 | 2 | 7.37E-06 | 1.82561804 | 0.667 | 0.345 | 0.01473104 |
| T-cell | Tbx21 | 2 | 8.35E-06 | 2.00722648 | 0.795 | 0.45 | 0.01669334 |
| T-cell | Rnf43 | 2 | 8.96E-06 | 2.58446408 | 0.769 | 0.445 | 0.01791082 |
| T-cell | Pstpip2 | 2 | 9.08E-06 | 1.73664298 | 0.744 | 0.463 | 0.01815813 |
| T-cell | Ryr2 | 2 | 1.17E-05 | 0.99502251 | 0.923 | 0.672 | 0.02339554 |
| T-cell | Ftl1 | 2 | 1.40E-05 | 2.02642515 | 0.667 | 0.314 | 0.02800877 |
| T-cell | Pik3r5 | 2 | 1.95E-05 | 1.4594663 | 0.718 | 0.454 | 0.0390141 |
| T-cell | Ptprb | 3 | 4.26E-17 | 2.86533787 | 0.972 | 0.466 | 8.52E-14 |
| T-cell | Etl4 | 3 | 7.30E-17 | 2.78284681 | 0.972 | 0.448 | 1.46E-13 |
| T-cell | Shank3 | 3 | 8.66E-15 | 2.54798982 | 0.944 | 0.435 | 1.73E-11 |
| T-cell | Cyyr1 | 3 | 9.32E-15 | 2.76309833 | 0.917 | 0.44 | 1.86E-11 |
| T-cell | Cdh13 | 3 | 1.25E-14 | 2.47687785 | 0.972 | 0.56 | 2.50E-11 |
| T-cell | Dach1 | 3 | 8.26E-12 | 2.75589467 | 0.861 | 0.366 | 1.65E-08 |
| T-cell | Flt1 | 3 | 3.93E-11 | 2.73403999 | 0.833 | 0.297 | 7.85E-08 |
| T-cell | Adgrf5 | 3 | 7.21E-10 | 2.60721782 | 0.806 | 0.453 | 1.44E-06 |
| T-cell | Kitlg | 3 | 3.90E-09 | 2.03906166 | 0.833 | 0.345 | 7.80E-06 |

| | | | | | | | |
|---|---|---|---|---|---|---|---|
| T-cell | Lims2 | 3 | 1.22E-08 | 2.86006573 | 0.75 | 0.409 | 2.44E-05 |
| T-cell | Ccdc85a | 3 | 1.26E-08 | 2.4448586 | 0.806 | 0.5 | 2.52E-05 |
| T-cell | Mgll | 3 | 7.06E-08 | 2.57953177 | 0.778 | 0.28 | 0.00014116 |
| T-cell | AABR070070 | 3 | 4.76E-07 | 1.25074986 | 0.889 | 0.595 | 0.000953 |
| T-cell | Emcn | 3 | 6.11E-07 | 2.17918577 | 0.75 | 0.427 | 0.00122237 |
| T-cell | Ablim3 | 3 | 1.96E-06 | 2.4992316 | 0.694 | 0.246 | 0.00392062 |
| T-cell | Slc26a10 | 3 | 1.96E-06 | 3.58894967 | 0.667 | 0.388 | 0.00392465 |
| T-cell | Crim1 | 3 | 9.40E-06 | 1.84050142 | 0.694 | 0.353 | 0.01879134 |
| T-cell | Frmd4b | 4 | 9.38E-10 | 2.38108345 | 0.9 | 0.403 | 1.88E-06 |
| T-cell | Tbxas1 | 4 | 2.22E-09 | 3.37749156 | 0.8 | 0.437 | 4.45E-06 |
| T-cell | Cfh | 4 | 5.48E-09 | 2.52993774 | 0.767 | 0.197 | 1.10E-05 |
| T-cell | Lgmn | 4 | 6.90E-09 | 3.041091 | 0.8 | 0.277 | 1.38E-05 |
| T-cell | Mrc1 | 4 | 7.72E-09 | 4.07923288 | 0.733 | 0.214 | 1.54E-05 |
| T-cell | Slc9a9 | 4 | 3.50E-08 | 1.47455271 | 0.967 | 0.706 | 7.00E-05 |
| T-cell | LOC690045 | 4 | 9.34E-08 | 2.58705828 | 0.8 | 0.34 | 0.00018673 |
| T-cell | Arhgap22 | 4 | 2.20E-07 | 2.95015156 | 0.767 | 0.445 | 0.00044071 |
| T-cell | Mtss1 | 4 | 4.56E-07 | 1.88187649 | 0.833 | 0.416 | 0.0009121 |
| T-cell | Ctss | 4 | 8.64E-07 | 3.88341744 | 0.7 | 0.218 | 0.00172806 |
| T-cell | Slco2b1 | 4 | 1.28E-06 | 2.18973795 | 0.8 | 0.294 | 0.00255181 |
| T-cell | Cers6 | 4 | 1.29E-06 | 2.0134014 | 0.767 | 0.282 | 0.00257094 |
| T-cell | Pid1 | 4 | 2.57E-06 | 1.483293 | 0.867 | 0.613 | 0.00514635 |
| T-cell | Wdfy4 | 4 | 3.13E-06 | 2.65842996 | 0.633 | 0.223 | 0.00625684 |
| T-cell | Ptprj | 4 | 4.82E-06 | 2.19277856 | 0.8 | 0.525 | 0.00964853 |
| T-cell | Slc16a7 | 4 | 5.29E-06 | 3.98833229 | 0.667 | 0.382 | 0.0105703 |
| T-cell | Alcam | 4 | 1.24E-05 | 2.3283609 | 0.7 | 0.34 | 0.02479546 |
| T-cell | Ect2 | 5 | 2.11E-09 | 5.40294835 | 0.923 | 0.302 | 4.23E-06 |
| T-cell | Cenpf | 5 | 3.42E-09 | 5.42310229 | 0.923 | 0.322 | 6.84E-06 |
| T-cell | Slpi | 5 | 2.47E-08 | 1.05877777 | 0.385 | 0.016 | 4.95E-05 |
| T-cell | Troap | 5 | 3.47E-08 | 6.90152612 | 0.769 | 0.067 | 6.94E-05 |
| T-cell | Ccnf | 5 | 1.54E-07 | 7.17919256 | 0.769 | 0.263 | 0.00030866 |
| T-cell | Kif20b | 5 | 2.02E-07 | 4.5912104 | 0.846 | 0.318 | 0.00040482 |

| | | | | | | | |
|---|---|---|---|---|---|---|---|
| T-cell | Kif11 | 5 | 2.77E-07 | 5.46958081 | 0.846 | 0.196 | 0.00055464 |
| T-cell | Etv3l | 5 | 4.15E-07 | 7.65185156 | 0.538 | 0.11 | 0.00082941 |
| T-cell | Cdca2 | 5 | 7.46E-07 | 7.29900033 | 0.769 | 0.282 | 0.00149209 |
| T-cell | Top2a | 5 | 7.62E-07 | 5.87673841 | 0.769 | 0.071 | 0.00152364 |
| T-cell | Cdkn3 | 5 | 1.02E-06 | 6.25163556 | 0.769 | 0.204 | 0.00204161 |
| T-cell | Casr1 | 5 | 1.35E-06 | 5.73432984 | 0.538 | 0.122 | 0.00270033 |
| T-cell | Cenpe | 5 | 1.53E-06 | 4.4692521 | 0.769 | 0.2 | 0.0030586 |
| T-cell | Kifc1 | 5 | 1.75E-06 | 6.61585218 | 0.769 | 0.298 | 0.00350716 |
| T-cell | AC094126.1 | 5 | 2.69E-06 | 5.86677612 | 0.538 | 0.125 | 0.00538935 |
| T-cell | Knstrn | 5 | 2.79E-06 | 5.47314538 | 0.769 | 0.29 | 0.00558422 |
| T-cell | Sgo2 | 5 | 4.14E-06 | 5.17585121 | 0.769 | 0.325 | 0.00827056 |
| T-cell | AABR070354 | 5 | 4.74E-06 | 4.21075455 | 0.385 | 0.055 | 0.00948654 |
| T-cell | Depdc1 | 5 | 4.89E-06 | 5.14879458 | 0.692 | 0.227 | 0.00977163 |
| T-cell | Kif23 | 5 | 6.21E-06 | 7.53512095 | 0.769 | 0.333 | 0.01241347 |
| T-cell | Ska1 | 5 | 8.34E-06 | 4.83889708 | 0.692 | 0.106 | 0.01668093 |
| T-cell | Melk | 5 | 8.91E-06 | 4.30291121 | 0.615 | 0.153 | 0.01781724 |
| T-cell | Cep55 | 5 | 1.04E-05 | 5.79241815 | 0.538 | 0.098 | 0.02071749 |
| T-cell | Prc1 | 5 | 2.08E-05 | 4.11252859 | 0.769 | 0.255 | 0.04159847 |
| Neuronal Cell | Nrxn3 | 0 | 1.48E-24 | 0.91720187 | 0.123 | 0.517 | 2.96E-21 |
| Neuronal Cell | Fbxw10 | 0 | 3.61E-21 | 0.29684316 | 0.089 | 0.483 | 7.22E-18 |
| Neuronal Cell | Pi15 | 0 | 1.26E-20 | 0.51159551 | 0.151 | 0.561 | 2.51E-17 |
| Neuronal Cell | NEWGENE-2 | 0 | 7.82E-20 | 1.42251684 | 0.073 | 0.453 | 1.56E-16 |
| Neuronal Cell | Pamr1 | 0 | 8.57E-20 | 0.77685934 | 0.201 | 0.53 | 1.71E-16 |
| Neuronal Cell | Nsg2 | 0 | 4.52E-18 | 0.28007613 | 0.017 | 0.342 | 9.04E-15 |
| Neuronal Cell | Hoxd3 | 0 | 5.02E-18 | 2.16078992 | 0.14 | 0.506 | 1.00E-14 |
| Neuronal Cell | Gbp1 | 0 | 4.24E-17 | 1.10793773 | 0.006 | 0.263 | 8.49E-14 |
| Neuronal Cell | Coq8a | 0 | 2.84E-15 | 0.3129431 | 0.223 | 0.57 | 5.68E-12 |
| Neuronal Cell | Asb14 | 0 | 4.69E-15 | 0.47386097 | 0.095 | 0.366 | 9.37E-12 |
| Neuronal Cell | Cacnb4 | 0 | 2.52E-13 | 1.15610319 | 0.207 | 0.578 | 5.04E-10 |
| Neuronal Cell | Wnk2 | 0 | 3.34E-13 | 0.87639818 | 0.251 | 0.576 | 6.68E-10 |
| Neuronal Cell | Bace2 | 0 | 8.89E-13 | 0.32921224 | 0.179 | 0.475 | 1.78E-09 |

| | | | | | | | |
|---|---|---|---|---|---|---|---|
| Neuronal Cell | Iqgap3 | 0 | 1.44E-12 | 0.27098207 | 0.246 | 0.532 | 2.89E-09 |
| Neuronal Cell | AABR070171 | 0 | 1.99E-12 | 0.6496604 | 0.117 | 0.404 | 3.98E-09 |
| Neuronal Cell | Il17rd | 0 | 2.96E-10 | 0.64661985 | 0.218 | 0.51 | 5.92E-07 |
| Neuronal Cell | Gpam | 0 | 4.85E-10 | 0.62626281 | 0.229 | 0.534 | 9.70E-07 |
| Neuronal Cell | Pfkfb2 | 0 | 1.60E-09 | 0.2535003 | 0.285 | 0.545 | 3.20E-06 |
| Neuronal Cell | Pla2r1 | 0 | 4.92E-09 | 0.56612548 | 0.363 | 0.636 | 9.84E-06 |
| Neuronal Cell | Slc6a15 | 0 | 1.01E-08 | 0.32589725 | 0.145 | 0.501 | 2.01E-05 |
| Neuronal Cell | Stum | 0 | 1.59E-08 | 0.4872747 | 0.263 | 0.561 | 3.17E-05 |
| Neuronal Cell | Fa2h | 0 | 1.24E-07 | 1.2530015 | 0.251 | 0.517 | 0.00024824 |
| Neuronal Cell | Fam135b | 1 | 5.85E-23 | 1.03665199 | 0.656 | 0.272 | 1.17E-19 |
| Neuronal Cell | Prodh1 | 1 | 6.87E-23 | 1.65376812 | 0.841 | 0.451 | 1.37E-19 |
| Neuronal Cell | Il1a | 1 | 3.34E-22 | 0.47750383 | 0.732 | 0.328 | 6.68E-19 |
| Neuronal Cell | Lhx3 | 1 | 1.00E-21 | 3.05749109 | 0.809 | 0.36 | 2.00E-18 |
| Neuronal Cell | Krt18 | 1 | 3.05E-21 | 1.96054204 | 0.758 | 0.377 | 6.11E-18 |
| Neuronal Cell | Hmga2 | 1 | 4.42E-20 | 2.73825964 | 0.701 | 0.299 | 8.85E-17 |
| Neuronal Cell | Dok5 | 1 | 1.70E-19 | 1.28345488 | 0.847 | 0.495 | 3.40E-16 |
| Neuronal Cell | Apod | 1 | 1.87E-19 | 0.49988548 | 0.726 | 0.349 | 3.75E-16 |
| Neuronal Cell | AABR070504 | 1 | 4.30E-19 | 1.02995035 | 0.752 | 0.438 | 8.60E-16 |
| Neuronal Cell | Sctr | 1 | 9.66E-19 | 1.95895928 | 0.758 | 0.404 | 1.93E-15 |
| Neuronal Cell | RGD1564053 | 1 | 1.40E-18 | 0.97683163 | 0.809 | 0.387 | 2.80E-15 |
| Neuronal Cell | Cpa6 | 1 | 4.21E-18 | 2.17802887 | 0.828 | 0.467 | 8.42E-15 |
| Neuronal Cell | Mlph | 1 | 4.81E-18 | 0.91540941 | 0.898 | 0.518 | 9.61E-15 |
| Neuronal Cell | Pak3 | 1 | 7.58E-18 | 1.09798752 | 0.713 | 0.364 | 1.52E-14 |
| Neuronal Cell | Fhl5 | 1 | 1.32E-17 | 1.02258597 | 0.809 | 0.425 | 2.63E-14 |
| Neuronal Cell | Tox2 | 1 | 3.68E-17 | 1.86018005 | 0.656 | 0.246 | 7.36E-14 |
| Neuronal Cell | Snca | 1 | 5.36E-17 | 2.43353806 | 0.669 | 0.343 | 1.07E-13 |
| Neuronal Cell | Neurl1 | 1 | 8.89E-17 | 1.42565799 | 0.815 | 0.448 | 1.78E-13 |
| Neuronal Cell | Cacna1e | 1 | 9.79E-17 | 0.50438435 | 0.828 | 0.476 | 1.96E-13 |
| Neuronal Cell | Esr1 | 1 | 1.77E-16 | 1.25198129 | 0.79 | 0.514 | 3.54E-13 |
| Neuronal Cell | S100a3 | 1 | 2.60E-16 | 2.5028183 | 0.809 | 0.438 | 5.19E-13 |
| Neuronal Cell | Slc18a2 | 1 | 3.10E-16 | 1.03060085 | 0.643 | 0.284 | 6.21E-13 |

| | | | | | | | |
|---|---|---|---|---|---|---|---|
| Neuronal Cell | Sfrp4 | 1 | 3.20E-16 | 0.9318274 | 0.822 | 0.537 | 6.40E-13 |
| Neuronal Cell | Dgkg | 1 | 4.08E-16 | 1.3263795 | 0.783 | 0.364 | 8.16E-13 |
| Neuronal Cell | Pax5 | 1 | 5.05E-16 | 1.55109871 | 0.815 | 0.451 | 1.01E-12 |
| Neuronal Cell | Hs3st3a1 | 1 | 6.70E-16 | 1.71296104 | 0.822 | 0.484 | 1.34E-12 |
| Neuronal Cell | Ppp1r14c | 1 | 9.35E-16 | 0.61712601 | 0.86 | 0.537 | 1.87E-12 |
| Neuronal Cell | Pld5 | 1 | 9.72E-16 | 0.26182874 | 0.745 | 0.402 | 1.94E-12 |
| Neuronal Cell | Dusp27 | 1 | 1.06E-15 | 0.31772943 | 0.79 | 0.402 | 2.12E-12 |
| Neuronal Cell | Slc6a151 | 1 | 2.04E-15 | 1.25431139 | 0.688 | 0.305 | 4.08E-12 |
| Neuronal Cell | Mlf1 | 1 | 2.18E-15 | 1.27846999 | 0.732 | 0.425 | 4.36E-12 |
| Neuronal Cell | Ephb1 | 1 | 2.23E-15 | 1.65533129 | 0.701 | 0.274 | 4.47E-12 |
| Neuronal Cell | Nmrk2 | 1 | 2.43E-15 | 3.3910524 | 0.624 | 0.364 | 4.87E-12 |
| Neuronal Cell | Cacna1a | 1 | 3.56E-15 | 0.99732094 | 0.873 | 0.573 | 7.12E-12 |
| Neuronal Cell | Metrn | 1 | 6.12E-15 | 0.58502929 | 0.758 | 0.354 | 1.22E-11 |
| Neuronal Cell | Lef1 | 1 | 8.19E-15 | 1.72146176 | 0.866 | 0.566 | 1.64E-11 |
| Neuronal Cell | Ildr2 | 1 | 1.03E-14 | 1.57443071 | 0.822 | 0.522 | 2.06E-11 |
| Neuronal Cell | Tfap2b | 1 | 1.08E-14 | 1.20169961 | 0.815 | 0.461 | 2.16E-11 |
| Neuronal Cell | Scml4 | 1 | 1.44E-14 | 0.80119719 | 0.586 | 0.305 | 2.88E-11 |
| Neuronal Cell | Ddc | 1 | 2.34E-14 | 1.78217279 | 0.688 | 0.411 | 4.68E-11 |
| Neuronal Cell | S100b | 1 | 5.50E-14 | 1.73916299 | 0.465 | 0.202 | 1.10E-10 |
| Neuronal Cell | Tmem26 | 1 | 5.56E-14 | 1.6224451 | 0.815 | 0.503 | 1.11E-10 |
| Neuronal Cell | Megf10 | 1 | 5.65E-14 | 1.14619178 | 0.847 | 0.516 | 1.13E-10 |
| Neuronal Cell | Gja5 | 1 | 5.93E-14 | 2.7885513 | 0.752 | 0.339 | 1.19E-10 |
| Neuronal Cell | Brca1 | 1 | 7.30E-14 | 1.24282425 | 0.72 | 0.36 | 1.46E-10 |
| Neuronal Cell | Prkcz | 1 | 7.58E-14 | 0.45723226 | 0.834 | 0.501 | 1.52E-10 |
| Neuronal Cell | Rtn4rl1 | 1 | 8.22E-14 | 0.50318428 | 0.726 | 0.335 | 1.64E-10 |
| Neuronal Cell | Nrg4 | 1 | 1.17E-13 | 1.87089416 | 0.688 | 0.32 | 2.34E-10 |
| Neuronal Cell | Anks1b | 1 | 1.54E-13 | 0.57388806 | 0.752 | 0.398 | 3.08E-10 |
| Neuronal Cell | Slc39a8 | 1 | 1.59E-13 | 1.24886682 | 0.732 | 0.394 | 3.19E-10 |
| Neuronal Cell | Rhpn2 | 1 | 1.65E-13 | 0.32594466 | 0.395 | 0.124 | 3.29E-10 |
| Neuronal Cell | Scg2 | 1 | 1.78E-13 | 1.99399091 | 0.809 | 0.421 | 3.57E-10 |
| Neuronal Cell | Corin | 1 | 2.27E-13 | 0.95874726 | 0.592 | 0.253 | 4.53E-10 |

| | | | | | | | |
|---|---|---|---|---|---|---|---|
| Neuronal Cell | Top2a | 1 | 2.98E-13 | 0.31786396 | 0.803 | 0.404 | 5.96E-10 |
| Neuronal Cell | AABR070172 | 1 | 6.04E-13 | 0.33010999 | 0.72 | 0.349 | 1.21E-09 |
| Neuronal Cell | Ctnnd2 | 1 | 1.47E-12 | 2.01294448 | 0.79 | 0.423 | 2.93E-09 |
| Neuronal Cell | Tmem116 | 1 | 1.50E-12 | 1.09059074 | 0.631 | 0.288 | 3.01E-09 |
| Neuronal Cell | Megf11 | 1 | 1.63E-12 | 1.15588997 | 0.783 | 0.491 | 3.26E-09 |
| Neuronal Cell | Vcan | 1 | 1.73E-12 | 1.50033838 | 0.752 | 0.499 | 3.46E-09 |
| Neuronal Cell | Med12l | 1 | 1.94E-12 | 0.35454764 | 0.822 | 0.503 | 3.87E-09 |
| Neuronal Cell | Upk1b | 1 | 2.12E-12 | 0.79880479 | 0.732 | 0.398 | 4.24E-09 |
| Neuronal Cell | Tnc | 1 | 2.60E-12 | 0.84250914 | 0.866 | 0.566 | 5.20E-09 |
| Neuronal Cell | Prkaa2 | 1 | 5.71E-12 | 0.26532038 | 0.834 | 0.453 | 1.14E-08 |
| Neuronal Cell | Dok6 | 1 | 5.91E-12 | 1.96480663 | 0.771 | 0.411 | 1.18E-08 |
| Neuronal Cell | Col20a1 | 1 | 6.66E-12 | 1.15110694 | 0.688 | 0.392 | 1.33E-08 |
| Neuronal Cell | Kank4 | 1 | 1.06E-11 | 0.74206359 | 0.694 | 0.349 | 2.13E-08 |
| Neuronal Cell | Ikzf1 | 1 | 1.23E-11 | 0.34879402 | 0.815 | 0.52 | 2.46E-08 |
| Neuronal Cell | Arpp21 | 1 | 1.42E-11 | 1.40149859 | 0.752 | 0.343 | 2.83E-08 |
| Neuronal Cell | Ptchd1 | 1 | 1.44E-11 | 0.81845738 | 0.694 | 0.379 | 2.88E-08 |
| Neuronal Cell | Spink8 | 1 | 2.46E-11 | 0.31652106 | 0.822 | 0.526 | 4.91E-08 |
| Neuronal Cell | Shc4 | 1 | 3.07E-11 | 1.01301835 | 0.745 | 0.446 | 6.13E-08 |
| Neuronal Cell | Ptprt | 1 | 3.74E-11 | 0.60107869 | 0.847 | 0.512 | 7.48E-08 |
| Neuronal Cell | Hs3st5 | 1 | 4.85E-11 | 0.4253482 | 0.828 | 0.558 | 9.71E-08 |
| Neuronal Cell | Pcdh7 | 1 | 5.39E-11 | 0.54863108 | 0.834 | 0.566 | 1.08E-07 |
| Neuronal Cell | Insc | 1 | 6.31E-11 | 0.92442848 | 0.828 | 0.531 | 1.26E-07 |
| Neuronal Cell | Adcy5 | 1 | 7.60E-11 | 0.700646 | 0.822 | 0.528 | 1.52E-07 |
| Neuronal Cell | Dtna | 1 | 8.92E-11 | 1.23399114 | 0.783 | 0.484 | 1.78E-07 |
| Neuronal Cell | Cyfip2 | 1 | 1.10E-10 | 0.62806075 | 0.745 | 0.413 | 2.20E-07 |
| Neuronal Cell | Cyp26b1 | 1 | 1.12E-10 | 0.97207988 | 0.758 | 0.472 | 2.25E-07 |
| Neuronal Cell | Pfkfb1 | 1 | 1.24E-10 | 0.53485324 | 0.815 | 0.535 | 2.48E-07 |
| Neuronal Cell | Ephb2 | 1 | 1.33E-10 | 0.99210118 | 0.924 | 0.632 | 2.67E-07 |
| Neuronal Cell | Dhrs7c | 1 | 1.38E-10 | 2.37103866 | 0.688 | 0.429 | 2.77E-07 |
| Neuronal Cell | Lrrc2 | 1 | 1.47E-10 | 0.67738357 | 0.739 | 0.434 | 2.93E-07 |
| Neuronal Cell | Stum1 | 1 | 1.60E-10 | 0.63327864 | 0.688 | 0.406 | 3.19E-07 |

| | | | | | | | |
|---|---|---|---|---|---|---|---|
| Neuronal Cell | Col12a1 | 1 | 2.44E-10 | 0.79182075 | 0.707 | 0.415 | 4.89E-07 |
| Neuronal Cell | Spon1 | 1 | 2.88E-10 | 1.00212515 | 0.828 | 0.512 | 5.75E-07 |
| Neuronal Cell | Rnf43 | 1 | 3.83E-10 | 0.84933412 | 0.497 | 0.232 | 7.65E-07 |
| Neuronal Cell | Gpx1 | 1 | 4.02E-10 | 0.34824854 | 0.809 | 0.533 | 8.05E-07 |
| Neuronal Cell | Ndufa4 | 1 | 4.11E-10 | 0.71859903 | 0.713 | 0.425 | 8.21E-07 |
| Neuronal Cell | Cldn15 | 1 | 7.61E-10 | 0.40341986 | 0.662 | 0.387 | 1.52E-06 |
| Neuronal Cell | Tyrobp | 1 | 7.80E-10 | 0.80508045 | 0.809 | 0.497 | 1.56E-06 |
| Neuronal Cell | AABR070655 | 1 | 8.78E-10 | 1.08863104 | 0.624 | 0.291 | 1.76E-06 |
| Neuronal Cell | Tiam1 | 1 | 9.29E-10 | 0.55565888 | 0.739 | 0.406 | 1.86E-06 |
| Neuronal Cell | Ablim2 | 1 | 1.03E-09 | 0.34417406 | 0.669 | 0.381 | 2.06E-06 |
| Neuronal Cell | Prss12 | 1 | 1.11E-09 | 0.90985881 | 0.669 | 0.379 | 2.23E-06 |
| Neuronal Cell | Grm8 | 1 | 1.13E-09 | 0.50315745 | 0.688 | 0.432 | 2.25E-06 |
| Neuronal Cell | Fstl5 | 1 | 1.46E-09 | 1.36039865 | 0.662 | 0.309 | 2.92E-06 |
| Neuronal Cell | Espl1 | 1 | 1.74E-09 | 0.89072016 | 0.732 | 0.406 | 3.48E-06 |
| Neuronal Cell | Crlf1 | 1 | 1.75E-09 | 0.53175814 | 0.732 | 0.381 | 3.51E-06 |
| Neuronal Cell | Necab1 | 1 | 1.80E-09 | 1.0098993 | 0.809 | 0.491 | 3.59E-06 |
| Neuronal Cell | Nmnat2 | 1 | 1.85E-09 | 0.46062289 | 0.688 | 0.354 | 3.71E-06 |
| Neuronal Cell | Itga7 | 1 | 3.04E-09 | 0.52763147 | 0.713 | 0.396 | 6.07E-06 |
| Neuronal Cell | Gm2a | 1 | 3.04E-09 | 0.96043782 | 0.752 | 0.392 | 6.08E-06 |
| Neuronal Cell | Pde8b | 1 | 3.83E-09 | 0.88651714 | 0.924 | 0.672 | 7.66E-06 |
| Neuronal Cell | Myom2 | 1 | 4.16E-09 | 0.43783054 | 0.771 | 0.495 | 8.33E-06 |
| Neuronal Cell | Cpt1b | 1 | 4.84E-09 | 0.29487348 | 0.752 | 0.484 | 9.68E-06 |
| Neuronal Cell | Reln | 1 | 7.82E-09 | 0.89744297 | 0.873 | 0.602 | 1.56E-05 |
| Neuronal Cell | Jph1 | 1 | 8.20E-09 | 0.65470269 | 0.834 | 0.556 | 1.64E-05 |
| Neuronal Cell | P3h2 | 1 | 1.03E-08 | 0.87659699 | 0.86 | 0.545 | 2.06E-05 |
| Neuronal Cell | Myocd | 1 | 1.23E-08 | 0.53125739 | 0.815 | 0.503 | 2.46E-05 |
| Neuronal Cell | Rnf150 | 1 | 1.44E-08 | 0.37517302 | 0.713 | 0.413 | 2.88E-05 |
| Neuronal Cell | Igfbp6 | 1 | 1.89E-08 | 0.41832326 | 0.643 | 0.328 | 3.78E-05 |
| Neuronal Cell | Ckap2 | 1 | 1.97E-08 | 0.95838942 | 0.739 | 0.438 | 3.95E-05 |
| Neuronal Cell | Bzw2 | 1 | 2.02E-08 | 0.52578291 | 0.783 | 0.453 | 4.03E-05 |
| Neuronal Cell | Abcd2 | 1 | 2.05E-08 | 1.66798634 | 0.669 | 0.331 | 4.09E-05 |

| | | | | | | | |
|---|---|---|---|---|---|---|---|
| Neuronal Cell | Tmem196 | 1 | 2.41E-08 | 0.92007018 | 0.854 | 0.562 | 4.82E-05 |
| Neuronal Cell | Mmp16 | 1 | 2.44E-08 | 0.50752018 | 0.777 | 0.463 | 4.87E-05 |
| Neuronal Cell | Ngfr | 1 | 2.46E-08 | 0.62765562 | 0.758 | 0.459 | 4.91E-05 |
| Neuronal Cell | Rnd3 | 1 | 3.30E-08 | 0.59575597 | 0.732 | 0.467 | 6.61E-05 |
| Neuronal Cell | Cdon | 1 | 3.44E-08 | 0.28666159 | 0.828 | 0.547 | 6.88E-05 |
| Neuronal Cell | Lrrtm4 | 1 | 3.62E-08 | 0.6848387 | 0.822 | 0.564 | 7.23E-05 |
| Neuronal Cell | Magi2 | 1 | 4.64E-08 | 1.38964155 | 0.675 | 0.396 | 9.27E-05 |
| Neuronal Cell | Shtn1 | 1 | 4.91E-08 | 1.45257047 | 0.688 | 0.404 | 9.82E-05 |
| Neuronal Cell | Acad10 | 1 | 8.58E-08 | 0.36626483 | 0.79 | 0.497 | 0.00017154 |
| Neuronal Cell | L1cam | 1 | 8.68E-08 | 0.65817966 | 0.803 | 0.533 | 0.00017354 |
| Neuronal Cell | Alpk3 | 1 | 9.34E-08 | 0.86973276 | 0.675 | 0.398 | 0.00018674 |
| Neuronal Cell | Tspan11 | 1 | 9.88E-08 | 0.44955472 | 0.904 | 0.596 | 0.00019765 |
| Neuronal Cell | Cdh8 | 1 | 1.06E-07 | 1.55329544 | 0.573 | 0.303 | 0.00021257 |
| Neuronal Cell | Pex5l | 1 | 1.10E-07 | 0.67931736 | 0.777 | 0.493 | 0.00021909 |
| Neuronal Cell | Acta2 | 1 | 1.42E-07 | 0.52828517 | 0.732 | 0.472 | 0.0002832 |
| Neuronal Cell | Dab1 | 1 | 1.42E-07 | 0.75794166 | 0.809 | 0.539 | 0.00028492 |
| Neuronal Cell | Slc16a1 | 1 | 1.47E-07 | 0.39748288 | 0.611 | 0.28 | 0.00029488 |
| Neuronal Cell | AC128394.2 | 1 | 1.48E-07 | 1.50933462 | 0.573 | 0.303 | 0.00029522 |
| Neuronal Cell | Gask1b | 1 | 1.51E-07 | 0.87076571 | 0.79 | 0.488 | 0.00030111 |
| Neuronal Cell | Cpe | 1 | 1.54E-07 | 0.52365499 | 0.809 | 0.539 | 0.00030776 |
| Neuronal Cell | Cilp | 1 | 1.61E-07 | 0.27051311 | 0.611 | 0.339 | 0.000321 |
| Neuronal Cell | Ezh2 | 1 | 1.85E-07 | 0.38492699 | 0.783 | 0.486 | 0.00036979 |
| Neuronal Cell | Fsd2 | 1 | 2.22E-07 | 0.80634619 | 0.828 | 0.566 | 0.0004447 |
| Neuronal Cell | AABR070347 | 1 | 4.00E-07 | 0.27140517 | 0.809 | 0.539 | 0.00080088 |
| Neuronal Cell | Adamts20 | 1 | 4.27E-07 | 0.60150036 | 0.79 | 0.514 | 0.00085329 |
| Neuronal Cell | Slc44a5 | 1 | 7.12E-07 | 0.97859401 | 0.815 | 0.518 | 0.00142448 |
| Neuronal Cell | Ros1 | 1 | 7.86E-07 | 1.44176026 | 0.777 | 0.472 | 0.00157186 |
| Neuronal Cell | Ttyh1 | 1 | 9.49E-07 | 0.52014445 | 0.892 | 0.623 | 0.00189858 |
| Neuronal Cell | Popdc2 | 1 | 9.72E-07 | 0.33543064 | 0.796 | 0.486 | 0.00194469 |
| Neuronal Cell | Camk4 | 1 | 1.04E-06 | 2.63472919 | 0.79 | 0.488 | 0.00207122 |
| Neuronal Cell | Ckmt2 | 1 | 1.15E-06 | 0.48259411 | 0.815 | 0.52 | 0.00230342 |

| | | | | | | | |
|---|---|---|---|---|---|---|---|
| Neuronal Cell | Nckap5 | 1 | 1.34E-06 | 0.45221486 | 0.675 | 0.404 | 0.00268158 |
| Neuronal Cell | Tesc | 1 | 1.38E-06 | 1.05595806 | 0.777 | 0.516 | 0.0027568 |
| Neuronal Cell | Cep55 | 1 | 1.42E-06 | 0.77714716 | 0.573 | 0.309 | 0.00283907 |
| Neuronal Cell | Kank1 | 1 | 1.71E-06 | 0.26739653 | 0.822 | 0.535 | 0.00341034 |
| Neuronal Cell | Bend6 | 1 | 1.82E-06 | 0.73911243 | 0.771 | 0.438 | 0.0036334 |
| Neuronal Cell | Rilpl1 | 1 | 2.66E-06 | 0.41575325 | 0.809 | 0.526 | 0.00532356 |
| Neuronal Cell | Fgf12 | 1 | 2.92E-06 | 0.47533283 | 0.834 | 0.543 | 0.00583615 |
| Neuronal Cell | Nrxn31 | 1 | 3.34E-06 | 0.5747655 | 0.624 | 0.333 | 0.00667477 |
| Neuronal Cell | Mx1 | 1 | 3.53E-06 | 1.57164111 | 0.79 | 0.522 | 0.00706028 |
| Neuronal Cell | Neb | 1 | 3.93E-06 | 1.53326004 | 0.758 | 0.503 | 0.00786418 |
| Neuronal Cell | Nppa | 1 | 1.38E-05 | 0.47714955 | 0.675 | 0.417 | 0.02761204 |
| Neuronal Cell | Myh7b | 1 | 1.52E-05 | 0.41431183 | 0.828 | 0.566 | 0.03040826 |
| Neuronal Cell | Grem2 | 2 | 3.16E-39 | 3.81741631 | 0.554 | 0.051 | 6.32E-36 |
| Neuronal Cell | Reg3g | 2 | 2.96E-35 | 3.76434141 | 0.624 | 0.121 | 5.93E-32 |
| Neuronal Cell | Gpc6 | 2 | 2.86E-33 | 2.33426047 | 0.931 | 0.588 | 5.71E-30 |
| Neuronal Cell | Cdh3 | 2 | 8.37E-31 | 2.19689348 | 0.475 | 0.041 | 1.67E-27 |
| Neuronal Cell | Adamts16 | 2 | 1.08E-30 | 3.48191424 | 0.416 | 0.041 | 2.16E-27 |
| Neuronal Cell | Npnt | 2 | 3.96E-30 | 2.15984155 | 0.614 | 0.162 | 7.92E-27 |
| Neuronal Cell | Fam180a | 2 | 6.10E-30 | 2.72191513 | 0.535 | 0.104 | 1.22E-26 |
| Neuronal Cell | Zfp385d | 2 | 3.18E-29 | 2.61217234 | 0.891 | 0.388 | 6.36E-26 |
| Neuronal Cell | C1qtnf7 | 2 | 1.33E-28 | 3.01206261 | 0.802 | 0.209 | 2.66E-25 |
| Neuronal Cell | Tbx21 | 2 | 2.38E-28 | 0.93895829 | 0.564 | 0.087 | 4.76E-25 |
| Neuronal Cell | Tbx5 | 2 | 2.65E-28 | 1.01539542 | 0.564 | 0.113 | 5.30E-25 |
| Neuronal Cell | Ebf2 | 2 | 1.54E-27 | 3.02622112 | 0.871 | 0.478 | 3.09E-24 |
| Neuronal Cell | Adamts2 | 2 | 5.11E-27 | 2.25580441 | 0.881 | 0.294 | 1.02E-23 |
| Neuronal Cell | Fmod | 2 | 3.06E-26 | 0.44461071 | 0.614 | 0.205 | 6.11E-23 |
| Neuronal Cell | AABR070586 | 2 | 6.08E-26 | 1.47649485 | 0.505 | 0.1 | 1.22E-22 |
| Neuronal Cell | Unc5d | 2 | 6.79E-26 | 1.10247285 | 0.515 | 0.102 | 1.36E-22 |
| Neuronal Cell | Col8a2 | 2 | 6.65E-25 | 2.66470981 | 0.574 | 0.147 | 1.33E-21 |
| Neuronal Cell | C7 | 2 | 9.32E-25 | 2.65032293 | 0.762 | 0.29 | 1.86E-21 |
| Neuronal Cell | Emx2 | 2 | 9.72E-25 | 3.95614868 | 0.594 | 0.188 | 1.94E-21 |

| | | | | | | | |
|---|---|---|---|---|---|---|---|
| Neuronal Cell | Col6a6 | 2 | 1.07E-24 | 3.30874638 | 0.851 | 0.522 | 2.13E-21 |
| Neuronal Cell | Fbn1 | 2 | 1.08E-24 | 2.09030901 | 0.931 | 0.48 | 2.16E-21 |
| Neuronal Cell | Pdgfrb | 2 | 1.91E-24 | 2.40018271 | 0.901 | 0.638 | 3.81E-21 |
| Neuronal Cell | Pid1 | 2 | 6.32E-24 | 2.32979266 | 0.822 | 0.282 | 1.26E-20 |
| Neuronal Cell | Ttc29 | 2 | 1.46E-23 | 2.54797959 | 0.386 | 0.047 | 2.91E-20 |
| Neuronal Cell | Cpq | 2 | 2.52E-23 | 2.1945138 | 0.812 | 0.245 | 5.05E-20 |
| Neuronal Cell | Cd68 | 2 | 4.36E-23 | 0.30225505 | 0.634 | 0.217 | 8.72E-20 |
| Neuronal Cell | Adgrd1 | 2 | 6.39E-23 | 3.88591181 | 0.634 | 0.068 | 1.28E-19 |
| Neuronal Cell | P2ry14 | 2 | 1.15E-22 | 2.82801066 | 0.644 | 0.22 | 2.29E-19 |
| Neuronal Cell | Ccdc80 | 2 | 1.65E-22 | 2.12105299 | 0.842 | 0.328 | 3.31E-19 |
| Neuronal Cell | Tmem63c | 2 | 2.04E-22 | 0.62559362 | 0.515 | 0.119 | 4.09E-19 |
| Neuronal Cell | AABR070015 | 2 | 2.12E-22 | 1.35076469 | 0.614 | 0.207 | 4.24E-19 |
| Neuronal Cell | Myh15 | 2 | 3.46E-22 | 3.83719198 | 0.495 | 0.109 | 6.91E-19 |
| Neuronal Cell | Itgbl1 | 2 | 5.75E-21 | 2.43239332 | 0.792 | 0.318 | 1.15E-17 |
| Neuronal Cell | Gabrp | 2 | 2.53E-20 | 1.19971815 | 0.554 | 0.139 | 5.07E-17 |
| Neuronal Cell | Boc | 2 | 2.89E-20 | 2.98049101 | 0.733 | 0.169 | 5.79E-17 |
| Neuronal Cell | Pde1a | 2 | 3.57E-20 | 2.37266272 | 0.802 | 0.424 | 7.13E-17 |
| Neuronal Cell | Col8a1 | 2 | 3.61E-20 | 2.74385121 | 0.832 | 0.345 | 7.21E-17 |
| Neuronal Cell | Ighm | 2 | 4.68E-20 | 1.27246297 | 0.545 | 0.205 | 9.36E-17 |
| Neuronal Cell | Slco1a4 | 2 | 4.93E-20 | 0.84251184 | 0.584 | 0.194 | 9.86E-17 |
| Neuronal Cell | Tacr1 | 2 | 5.42E-20 | 0.55021482 | 0.475 | 0.109 | 1.08E-16 |
| Neuronal Cell | Irs1 | 2 | 3.38E-19 | 2.14655343 | 0.861 | 0.478 | 6.77E-16 |
| Neuronal Cell | Eda2r | 2 | 7.89E-19 | 0.71463773 | 0.644 | 0.168 | 1.58E-15 |
| Neuronal Cell | Mfap5 | 2 | 1.50E-18 | 1.98817682 | 0.782 | 0.348 | 3.00E-15 |
| Neuronal Cell | Thbs1 | 2 | 1.86E-18 | 2.26176386 | 0.545 | 0.136 | 3.72E-15 |
| Neuronal Cell | Pdgfra | 2 | 4.56E-18 | 2.32457267 | 0.762 | 0.326 | 9.13E-15 |
| Neuronal Cell | Mrvi1 | 2 | 5.46E-18 | 3.00370865 | 0.525 | 0.113 | 1.09E-14 |
| Neuronal Cell | Npas2 | 2 | 2.07E-17 | 3.50844858 | 0.564 | 0.164 | 4.13E-14 |
| Neuronal Cell | Abi3bp | 2 | 2.56E-17 | 1.72106947 | 0.842 | 0.492 | 5.12E-14 |
| Neuronal Cell | Slc7a8 | 2 | 2.59E-17 | 1.85760279 | 0.436 | 0.113 | 5.17E-14 |
| Neuronal Cell | Prrx1 | 2 | 3.00E-17 | 2.61791341 | 0.673 | 0.211 | 6.00E-14 |

| | | | | | | | |
|---|---|---|---|---|---|---|---|
| Neuronal Cell | Mrc2 | 2 | 3.08E-17 | 2.88051605 | 0.713 | 0.379 | 6.17E-14 |
| Neuronal Cell | Robo1 | 2 | 5.88E-17 | 2.64302522 | 0.762 | 0.418 | 1.18E-13 |
| Neuronal Cell | Dscaml1 | 2 | 7.06E-17 | 0.40405913 | 0.446 | 0.07 | 1.41E-13 |
| Neuronal Cell | Clmp | 2 | 1.55E-16 | 2.98765382 | 0.673 | 0.363 | 3.10E-13 |
| Neuronal Cell | Il1r1 | 2 | 1.61E-16 | 1.53261991 | 0.861 | 0.512 | 3.21E-13 |
| Neuronal Cell | Dlg2 | 2 | 1.93E-16 | 2.95251572 | 0.574 | 0.141 | 3.86E-13 |
| Neuronal Cell | Rbp1 | 2 | 4.89E-16 | 1.82851646 | 0.495 | 0.111 | 9.78E-13 |
| Neuronal Cell | Kcnj8 | 2 | 8.74E-16 | 2.0805886 | 0.495 | 0.151 | 1.75E-12 |
| Neuronal Cell | Egfr | 2 | 3.64E-15 | 2.56443062 | 0.594 | 0.143 | 7.29E-12 |
| Neuronal Cell | Cacna2d3 | 2 | 3.77E-15 | 2.82902923 | 0.713 | 0.315 | 7.55E-12 |
| Neuronal Cell | Ar | 2 | 3.83E-15 | 2.22660553 | 0.723 | 0.303 | 7.65E-12 |
| Neuronal Cell | Pcdha13 | 2 | 4.40E-15 | 1.80571013 | 0.752 | 0.331 | 8.81E-12 |
| Neuronal Cell | AABR070635 | 2 | 1.01E-14 | 3.66234161 | 0.564 | 0.205 | 2.01E-11 |
| Neuronal Cell | Bin2 | 2 | 2.25E-14 | 0.97525098 | 0.455 | 0.124 | 4.49E-11 |
| Neuronal Cell | Piezo2 | 2 | 4.35E-14 | 1.83828901 | 0.713 | 0.307 | 8.70E-11 |
| Neuronal Cell | Trib3 | 2 | 5.20E-14 | 1.67108569 | 0.455 | 0.094 | 1.04E-10 |
| Neuronal Cell | Mylk | 2 | 5.36E-14 | 1.75912484 | 0.713 | 0.249 | 1.07E-10 |
| Neuronal Cell | Slc45a3 | 2 | 6.30E-14 | 0.98888834 | 0.436 | 0.107 | 1.26E-10 |
| Neuronal Cell | Spry3 | 2 | 8.10E-14 | 0.39238498 | 0.495 | 0.149 | 1.62E-10 |
| Neuronal Cell | Fstl1 | 2 | 2.42E-13 | 1.12061816 | 0.97 | 0.646 | 4.85E-10 |
| Neuronal Cell | AABR070546 | 2 | 5.48E-13 | 2.03632058 | 0.762 | 0.386 | 1.10E-09 |
| Neuronal Cell | St8sia1 | 2 | 6.90E-13 | 3.3526767 | 0.485 | 0.124 | 1.38E-09 |
| Neuronal Cell | Il16 | 2 | 8.34E-13 | 1.3916932 | 0.604 | 0.166 | 1.67E-09 |
| Neuronal Cell | Palm2 | 2 | 9.03E-13 | 2.61766056 | 0.634 | 0.186 | 1.81E-09 |
| Neuronal Cell | Myl9 | 2 | 9.75E-13 | 2.92062744 | 0.396 | 0.036 | 1.95E-09 |
| Neuronal Cell | Syk | 2 | 1.17E-12 | 0.89181941 | 0.683 | 0.382 | 2.34E-09 |
| Neuronal Cell | Uap1 | 2 | 1.45E-12 | 2.35575243 | 0.475 | 0.202 | 2.89E-09 |
| Neuronal Cell | Chsy3 | 2 | 1.51E-12 | 2.37368261 | 0.535 | 0.239 | 3.02E-09 |
| Neuronal Cell | Adgrg2 | 2 | 2.39E-12 | 1.69114981 | 0.455 | 0.087 | 4.78E-09 |
| Neuronal Cell | AABR070252 | 2 | 4.95E-12 | 1.496535 | 0.792 | 0.518 | 9.90E-09 |
| Neuronal Cell | Cacna1g | 2 | 5.81E-12 | 2.40673674 | 0.554 | 0.126 | 1.16E-08 |

| | | | | | | | |
|---|---|---|---|---|---|---|---|
| Neuronal Cell | Ptafr | 2 | 9.23E-12 | 0.6389019 | 0.446 | 0.089 | 1.85E-08 |
| Neuronal Cell | Itgal | 2 | 1.15E-11 | 1.30198828 | 0.505 | 0.16 | 2.31E-08 |
| Neuronal Cell | Dpysl3 | 2 | 2.10E-11 | 1.27522362 | 0.703 | 0.362 | 4.21E-08 |
| Neuronal Cell | Cdkn1a | 2 | 2.43E-11 | 0.86906571 | 0.386 | 0.085 | 4.85E-08 |
| Neuronal Cell | Plod2 | 2 | 2.47E-11 | 1.61388367 | 0.683 | 0.296 | 4.93E-08 |
| Neuronal Cell | Gria1 | 2 | 3.96E-11 | 1.53992432 | 0.426 | 0.164 | 7.92E-08 |
| Neuronal Cell | AC111804.2 | 2 | 4.03E-11 | 1.68178109 | 0.505 | 0.077 | 8.06E-08 |
| Neuronal Cell | Tbx20 | 2 | 4.08E-11 | 2.06419285 | 0.525 | 0.192 | 8.15E-08 |
| Neuronal Cell | Antxr1 | 2 | 4.23E-11 | 1.23803892 | 0.782 | 0.444 | 8.47E-08 |
| Neuronal Cell | Cdh11 | 2 | 6.51E-11 | 2.1589666 | 0.545 | 0.213 | 1.30E-07 |
| Neuronal Cell | Il2ra | 2 | 9.33E-11 | 2.29657646 | 0.485 | 0.213 | 1.87E-07 |
| Neuronal Cell | Fndc1 | 2 | 1.90E-10 | 0.83036324 | 0.911 | 0.505 | 3.80E-07 |
| Neuronal Cell | Enpp1 | 2 | 2.24E-10 | 2.05915862 | 0.455 | 0.098 | 4.49E-07 |
| Neuronal Cell | Col25a1 | 2 | 2.25E-10 | 3.31500684 | 0.455 | 0.185 | 4.50E-07 |
| Neuronal Cell | Col1a1 | 2 | 2.30E-10 | 0.8090284 | 0.911 | 0.616 | 4.61E-07 |
| Neuronal Cell | Gxylt2 | 2 | 2.54E-10 | 3.071081 | 0.525 | 0.134 | 5.08E-07 |
| Neuronal Cell | Ninj2 | 2 | 3.02E-10 | 2.57335586 | 0.485 | 0.198 | 6.03E-07 |
| Neuronal Cell | Slc4a4 | 2 | 3.40E-10 | 1.81800996 | 0.594 | 0.294 | 6.80E-07 |
| Neuronal Cell | Timp2 | 2 | 5.14E-10 | 0.95926408 | 0.832 | 0.463 | 1.03E-06 |
| Neuronal Cell | Adamts6 | 2 | 5.18E-10 | 1.6200883 | 0.535 | 0.254 | 1.04E-06 |
| Neuronal Cell | Pde7a | 2 | 5.58E-10 | 1.36156561 | 0.644 | 0.264 | 1.12E-06 |
| Neuronal Cell | Errfi1 | 2 | 5.85E-10 | 2.35727976 | 0.356 | 0.096 | 1.17E-06 |
| Neuronal Cell | Fn1 | 2 | 6.93E-10 | 1.23560576 | 0.634 | 0.269 | 1.39E-06 |
| Neuronal Cell | Cemip | 2 | 8.77E-10 | 2.99751386 | 0.624 | 0.348 | 1.75E-06 |
| Neuronal Cell | Pdlim3 | 2 | 1.31E-09 | 0.98148782 | 0.446 | 0.162 | 2.62E-06 |
| Neuronal Cell | Col6a2 | 2 | 1.68E-09 | 1.37795309 | 0.614 | 0.252 | 3.36E-06 |
| Neuronal Cell | Actn1 | 2 | 2.13E-09 | 1.46737891 | 0.465 | 0.147 | 4.26E-06 |
| Neuronal Cell | Aldh1a2 | 2 | 4.43E-09 | 2.73934902 | 0.515 | 0.23 | 8.86E-06 |
| Neuronal Cell | Dpt | 2 | 4.79E-09 | 1.26581386 | 0.644 | 0.277 | 9.59E-06 |
| Neuronal Cell | Agtr1a | 2 | 4.91E-09 | 1.89912417 | 0.554 | 0.162 | 9.82E-06 |
| Neuronal Cell | Rims1 | 2 | 5.26E-09 | 3.51172374 | 0.347 | 0.085 | 1.05E-05 |

| | | | | | | | |
|---|---|---|---|---|---|---|---|
| Neuronal Cell | Fbln5 | 2 | 5.75E-09 | 0.53697629 | 0.792 | 0.501 | 1.15E-05 |
| Neuronal Cell | Grip2 | 2 | 6.67E-09 | 2.71431852 | 0.515 | 0.122 | 1.33E-05 |
| Neuronal Cell | Nsg21 | 2 | 7.67E-09 | 0.86976769 | 0.475 | 0.207 | 1.53E-05 |
| Neuronal Cell | Colgalt2 | 2 | 8.10E-09 | 1.73292282 | 0.644 | 0.299 | 1.62E-05 |
| Neuronal Cell | Timp1 | 2 | 9.00E-09 | 1.01507108 | 0.535 | 0.137 | 1.80E-05 |
| Neuronal Cell | Gucy1a1 | 2 | 1.22E-08 | 2.18136141 | 0.406 | 0.094 | 2.44E-05 |
| Neuronal Cell | Arnt2 | 2 | 1.30E-08 | 0.86734935 | 0.634 | 0.315 | 2.59E-05 |
| Neuronal Cell | Efcc1 | 2 | 1.43E-08 | 2.53504606 | 0.604 | 0.33 | 2.86E-05 |
| Neuronal Cell | Slc9a9 | 2 | 1.48E-08 | 0.61203218 | 0.832 | 0.569 | 2.96E-05 |
| Neuronal Cell | Ccn3 | 2 | 1.51E-08 | 0.54667952 | 0.634 | 0.307 | 3.02E-05 |
| Neuronal Cell | Gria4 | 2 | 2.56E-08 | 3.3648875 | 0.525 | 0.186 | 5.13E-05 |
| Neuronal Cell | Smad6 | 2 | 2.76E-08 | 0.80545479 | 0.743 | 0.46 | 5.52E-05 |
| Neuronal Cell | Mpped2 | 2 | 2.85E-08 | 0.47761969 | 0.752 | 0.395 | 5.70E-05 |
| Neuronal Cell | Kif20b | 2 | 3.64E-08 | 0.46467647 | 0.515 | 0.213 | 7.27E-05 |
| Neuronal Cell | Pde5a | 2 | 4.73E-08 | 0.84340224 | 0.594 | 0.25 | 9.45E-05 |
| Neuronal Cell | Syn3 | 2 | 6.39E-08 | 0.54969782 | 0.634 | 0.273 | 0.00012775 |
| Neuronal Cell | Runx1 | 2 | 7.20E-08 | 0.91988725 | 0.723 | 0.365 | 0.00014409 |
| Neuronal Cell | Pde3a | 2 | 9.54E-08 | 1.67785461 | 0.673 | 0.373 | 0.00019081 |
| Neuronal Cell | Col24a1 | 2 | 1.02E-07 | 1.60873977 | 0.564 | 0.186 | 0.00020383 |
| Neuronal Cell | Mgp | 2 | 1.14E-07 | 0.59718716 | 0.812 | 0.497 | 0.00022799 |
| Neuronal Cell | Ltbp1 | 2 | 1.24E-07 | 1.32195934 | 0.644 | 0.296 | 0.00024891 |
| Neuronal Cell | Slc16a2 | 2 | 1.33E-07 | 0.55605178 | 0.475 | 0.124 | 0.00026596 |
| Neuronal Cell | Slc20a1 | 2 | 1.34E-07 | 0.99281478 | 0.733 | 0.478 | 0.00026715 |
| Neuronal Cell | Steap4 | 2 | 1.35E-07 | 3.08394449 | 0.535 | 0.181 | 0.00026958 |
| Neuronal Cell | Medag | 2 | 1.49E-07 | 3.96606991 | 0.475 | 0.104 | 0.00029799 |
| Neuronal Cell | Sned1 | 2 | 1.70E-07 | 1.08017995 | 0.465 | 0.175 | 0.00033981 |
| Neuronal Cell | Ppm1e | 2 | 2.21E-07 | 1.70502363 | 0.515 | 0.134 | 0.00044268 |
| Neuronal Cell | Tnn | 2 | 2.82E-07 | 1.63300315 | 0.584 | 0.322 | 0.00056457 |
| Neuronal Cell | Bgn | 2 | 2.95E-07 | 0.59220854 | 0.554 | 0.226 | 0.00059032 |
| Neuronal Cell | Lcp2 | 2 | 3.15E-07 | 0.75368266 | 0.634 | 0.384 | 0.00062909 |
| Neuronal Cell | Asap3 | 2 | 3.57E-07 | 2.14565531 | 0.475 | 0.151 | 0.00071391 |

| | | | | | | | |
|---|---|---|---|---|---|---|---|
| Neuronal Cell | Filip1l | 2 | 3.89E-07 | 0.73656364 | 0.723 | 0.42 | 0.00077785 |
| Neuronal Cell | Ednra | 2 | 4.32E-07 | 1.88282563 | 0.396 | 0.132 | 0.00086495 |
| Neuronal Cell | Bcat1 | 2 | 4.61E-07 | 1.70780916 | 0.455 | 0.143 | 0.00092204 |
| Neuronal Cell | Spp1 | 2 | 5.62E-07 | 0.44394611 | 0.604 | 0.29 | 0.00112408 |
| Neuronal Cell | Slc7a1 | 2 | 6.20E-07 | 0.52005325 | 0.446 | 0.186 | 0.00123983 |
| Neuronal Cell | Osbp2 | 2 | 1.02E-06 | 0.73901188 | 0.624 | 0.269 | 0.00203575 |
| Neuronal Cell | Asgr2 | 2 | 1.21E-06 | 0.69006221 | 0.525 | 0.235 | 0.00241712 |
| Neuronal Cell | Foxp2 | 2 | 1.68E-06 | 1.41011854 | 0.584 | 0.273 | 0.00336119 |
| Neuronal Cell | AABR070179 | 2 | 2.02E-06 | 1.02952764 | 0.515 | 0.218 | 0.00403786 |
| Neuronal Cell | Adamts19 | 2 | 2.13E-06 | 1.9660812 | 0.485 | 0.169 | 0.00426246 |
| Neuronal Cell | Bmp6 | 2 | 3.41E-06 | 1.95829869 | 0.475 | 0.113 | 0.00681691 |
| Neuronal Cell | Igf1 | 2 | 3.75E-06 | 1.93373946 | 0.525 | 0.173 | 0.00749164 |
| Neuronal Cell | Calcrl | 2 | 5.59E-06 | 0.53932347 | 0.584 | 0.286 | 0.0111813 |
| Neuronal Cell | Hs3st2 | 2 | 6.11E-06 | 1.2815153 | 0.396 | 0.137 | 0.01222843 |
| Neuronal Cell | Slit2 | 2 | 6.47E-06 | 1.14179019 | 0.446 | 0.181 | 0.01293674 |
| Neuronal Cell | Loxl2 | 2 | 7.91E-06 | 2.22017952 | 0.406 | 0.094 | 0.01581443 |
| Neuronal Cell | Myo1f | 2 | 8.44E-06 | 0.48030947 | 0.653 | 0.38 | 0.01687456 |
| Neuronal Cell | Ntn1 | 2 | 1.08E-05 | 1.20911492 | 0.446 | 0.162 | 0.02163065 |
| Neuronal Cell | Crim1 | 2 | 2.02E-05 | 0.93250654 | 0.653 | 0.373 | 0.04039585 |
| Neuronal Cell | Icam1 | 2 | 2.24E-05 | 0.79356587 | 0.465 | 0.134 | 0.04480538 |
| Neuronal Cell | Shank3 | 3 | 1.04E-33 | 3.4467036 | 0.961 | 0.665 | 2.08E-30 |
| Neuronal Cell | Cdh13 | 3 | 1.69E-32 | 2.99351618 | 0.987 | 0.525 | 3.38E-29 |
| Neuronal Cell | Ptprb | 3 | 1.99E-30 | 3.57615492 | 0.921 | 0.315 | 3.98E-27 |
| Neuronal Cell | Flt1 | 3 | 1.56E-27 | 3.1889547 | 0.908 | 0.509 | 3.11E-24 |
| Neuronal Cell | Cyyr1 | 3 | 4.87E-26 | 3.34828547 | 0.855 | 0.295 | 9.74E-23 |
| Neuronal Cell | Prkch | 3 | 9.05E-26 | 3.33605511 | 0.855 | 0.201 | 1.81E-22 |
| Neuronal Cell | Fli1 | 3 | 5.79E-24 | 2.88528629 | 0.855 | 0.282 | 1.16E-20 |
| Neuronal Cell | Ppp1r16b | 3 | 8.82E-23 | 4.35719821 | 0.658 | 0.225 | 1.76E-19 |
| Neuronal Cell | Adgrf5 | 3 | 9.94E-22 | 3.16590859 | 0.803 | 0.286 | 1.99E-18 |
| Neuronal Cell | Myrip | 3 | 4.73E-21 | 3.39598246 | 0.842 | 0.511 | 9.47E-18 |
| Neuronal Cell | Ccdc85a | 3 | 4.55E-20 | 2.96207552 | 0.829 | 0.327 | 9.10E-17 |

| | | | | | | | |
|---|---|---|---|---|---|---|---|
| Neuronal Cell | Emcn | 3 | 9.31E-19 | 3.00338052 | 0.803 | 0.29 | 1.86E-15 |
| Neuronal Cell | Itpkb | 3 | 3.95E-18 | 2.14580736 | 0.908 | 0.608 | 7.89E-15 |
| Neuronal Cell | Dach1 | 3 | 3.57E-17 | 3.11225979 | 0.776 | 0.281 | 7.14E-14 |
| Neuronal Cell | Lyn | 3 | 5.38E-16 | 2.02616776 | 0.711 | 0.21 | 1.08E-12 |
| Neuronal Cell | Meox2 | 3 | 7.99E-16 | 3.23022562 | 0.776 | 0.401 | 1.60E-12 |
| Neuronal Cell | Ano2 | 3 | 1.10E-15 | 3.45043967 | 0.513 | 0.174 | 2.21E-12 |
| Neuronal Cell | Calcrl1 | 3 | 1.33E-15 | 2.55188347 | 0.789 | 0.272 | 2.65E-12 |
| Neuronal Cell | Spns2 | 3 | 2.63E-15 | 4.2681894 | 0.526 | 0.149 | 5.25E-12 |
| Neuronal Cell | Clic5 | 3 | 2.79E-15 | 3.29729571 | 0.75 | 0.291 | 5.58E-12 |
| Neuronal Cell | AABR070070 | 3 | 8.60E-15 | 1.6447077 | 0.947 | 0.558 | 1.72E-11 |
| Neuronal Cell | Hmcn1 | 3 | 9.89E-15 | 2.27139006 | 0.75 | 0.234 | 1.98E-11 |
| Neuronal Cell | Crim11 | 3 | 1.65E-14 | 2.09254647 | 0.789 | 0.367 | 3.30E-11 |
| Neuronal Cell | Ablim3 | 3 | 2.96E-14 | 2.6819286 | 0.789 | 0.484 | 5.91E-11 |
| Neuronal Cell | Lmcd1 | 3 | 6.02E-13 | 3.3250267 | 0.618 | 0.147 | 1.20E-09 |
| Neuronal Cell | Fam163a | 3 | 1.05E-12 | 3.8791322 | 0.553 | 0.259 | 2.10E-09 |
| Neuronal Cell | Efnb2 | 3 | 1.68E-12 | 2.48102449 | 0.75 | 0.363 | 3.35E-09 |
| Neuronal Cell | Zfp366 | 3 | 6.88E-12 | 3.77916834 | 0.697 | 0.396 | 1.38E-08 |
| Neuronal Cell | Mcf2l | 3 | 9.44E-12 | 2.62187314 | 0.684 | 0.367 | 1.89E-08 |
| Neuronal Cell | Akr1c15 | 3 | 1.00E-11 | 3.10040469 | 0.684 | 0.254 | 2.01E-08 |
| Neuronal Cell | Plcb1 | 3 | 1.42E-11 | 2.30320711 | 0.803 | 0.514 | 2.84E-08 |
| Neuronal Cell | Dnm3 | 3 | 1.32E-10 | 1.48775852 | 0.842 | 0.415 | 2.64E-07 |
| Neuronal Cell | Thsd7a | 3 | 1.38E-10 | 2.36881252 | 0.618 | 0.187 | 2.76E-07 |
| Neuronal Cell | Nxn | 3 | 1.73E-10 | 1.44546886 | 0.763 | 0.374 | 3.46E-07 |
| Neuronal Cell | RGD1565355 | 3 | 1.78E-10 | 2.12577828 | 0.658 | 0.227 | 3.56E-07 |
| Neuronal Cell | Flt4 | 3 | 2.42E-10 | 4.70197917 | 0.329 | 0.025 | 4.85E-07 |
| Neuronal Cell | Alpl | 3 | 3.33E-10 | 4.74504948 | 0.303 | 0.049 | 6.66E-07 |
| Neuronal Cell | Tspan18 | 3 | 3.51E-10 | 1.94955469 | 0.724 | 0.297 | 7.01E-07 |
| Neuronal Cell | Il1r11 | 3 | 4.19E-10 | 1.91269366 | 0.816 | 0.534 | 8.38E-07 |
| Neuronal Cell | Kitlg | 3 | 4.74E-10 | 2.13424136 | 0.697 | 0.32 | 9.47E-07 |
| Neuronal Cell | Palmd | 3 | 1.32E-09 | 2.94016964 | 0.632 | 0.147 | 2.63E-06 |
| Neuronal Cell | Vwf | 3 | 1.64E-09 | 2.41280779 | 0.671 | 0.223 | 3.27E-06 |

| | | | | | | | |
|---|---|---|---|---|---|---|---|
| Neuronal Cell | Lgr6 | 3 | 7.03E-09 | 4.05519096 | 0.579 | 0.266 | 1.41E-05 |
| Neuronal Cell | Plcg2 | 3 | 7.68E-09 | 0.60229291 | 0.513 | 0.246 | 1.54E-05 |
| Neuronal Cell | Sncaip | 3 | 2.40E-08 | 2.18173917 | 0.434 | 0.108 | 4.80E-05 |
| Neuronal Cell | Wipf3 | 3 | 6.12E-08 | 2.64065645 | 0.618 | 0.27 | 0.00012234 |
| Neuronal Cell | Slc26a10 | 3 | 9.34E-08 | 4.16989601 | 0.566 | 0.174 | 0.00018689 |
| Neuronal Cell | Samd5 | 3 | 1.05E-06 | 3.48943302 | 0.553 | 0.182 | 0.00209863 |
| Neuronal Cell | Lmo7 | 3 | 2.98E-06 | 1.85858709 | 0.605 | 0.342 | 0.00596195 |
| Neuronal Cell | AABR070076 | 3 | 3.15E-06 | 3.93727872 | 0.474 | 0.185 | 0.00630646 |
| Neuronal Cell | Ntn11 | 3 | 3.81E-06 | 2.00335756 | 0.513 | 0.165 | 0.00762742 |
| Neuronal Cell | Mgmt | 3 | 1.34E-05 | 1.17908679 | 0.711 | 0.46 | 0.02672611 |
| Neuronal Cell | Hk3 | 4 | 1.69E-36 | 6.82342392 | 0.435 | 0.026 | 3.39E-33 |
| Neuronal Cell | Slc7a11 | 4 | 3.14E-29 | 4.83864042 | 0.694 | 0.139 | 6.28E-26 |
| Neuronal Cell | Atp1a3 | 4 | 1.04E-28 | 3.53988511 | 0.339 | 0.014 | 2.09E-25 |
| Neuronal Cell | Cd4 | 4 | 1.40E-28 | 3.12908323 | 0.871 | 0.309 | 2.79E-25 |
| Neuronal Cell | Mmp12 | 4 | 6.23E-27 | 7.17399571 | 0.387 | 0.028 | 1.25E-23 |
| Neuronal Cell | RGD1559482 | 4 | 1.26E-23 | 2.91864999 | 0.355 | 0.091 | 2.52E-20 |
| Neuronal Cell | Lmx1a | 4 | 1.71E-22 | 0.7637857 | 0.645 | 0.14 | 3.43E-19 |
| Neuronal Cell | Smoc2 | 4 | 1.77E-21 | 1.02831558 | 0.581 | 0.139 | 3.53E-18 |
| Neuronal Cell | Ky | 4 | 2.89E-21 | 2.24556183 | 0.758 | 0.272 | 5.78E-18 |
| Neuronal Cell | Asgr21 | 4 | 1.11E-20 | 3.14806639 | 0.806 | 0.225 | 2.22E-17 |
| Neuronal Cell | Abca17 | 4 | 1.29E-20 | 3.80859108 | 0.597 | 0.104 | 2.58E-17 |
| Neuronal Cell | Tbxas1 | 4 | 1.54E-20 | 4.5165927 | 0.645 | 0.274 | 3.07E-17 |
| Neuronal Cell | Mreg | 4 | 2.47E-20 | 3.77215388 | 0.339 | 0.047 | 4.94E-17 |
| Neuronal Cell | Itk | 4 | 4.43E-20 | 0.29536493 | 0.677 | 0.188 | 8.85E-17 |
| Neuronal Cell | Uqcrq | 4 | 5.26E-20 | 3.29278223 | 0.871 | 0.289 | 1.05E-16 |
| Neuronal Cell | Plcg21 | 4 | 5.35E-20 | 2.6683207 | 0.613 | 0.242 | 1.07E-16 |
| Neuronal Cell | Blnk | 4 | 6.62E-20 | 2.28823501 | 0.871 | 0.309 | 1.32E-16 |
| Neuronal Cell | Epsti1 | 4 | 1.09E-19 | 4.31762597 | 0.468 | 0.067 | 2.17E-16 |
| Neuronal Cell | Bin21 | 4 | 5.12E-19 | 3.40634534 | 0.565 | 0.135 | 1.02E-15 |
| Neuronal Cell | Cd300lb | 4 | 7.72E-19 | 5.15505436 | 0.387 | 0.046 | 1.54E-15 |
| Neuronal Cell | Slc7a81 | 4 | 1.02E-18 | 2.37456919 | 0.452 | 0.133 | 2.05E-15 |

| | | | | | | | |
|---|---|---|---|---|---|---|---|
| Neuronal Cell | Ly86 | 4 | 8.58E-18 | 4.54125036 | 0.419 | 0.047 | 1.72E-14 |
| Neuronal Cell | P2ry141 | 4 | 1.38E-17 | 1.53219319 | 0.726 | 0.24 | 2.76E-14 |
| Neuronal Cell | Pltp | 4 | 1.73E-17 | 3.29934673 | 0.548 | 0.105 | 3.47E-14 |
| Neuronal Cell | Mcoln2 | 4 | 2.28E-17 | 1.51006254 | 0.306 | 0.005 | 4.55E-14 |
| Neuronal Cell | Prc1 | 4 | 2.35E-17 | 1.86954386 | 0.629 | 0.196 | 4.69E-14 |
| Neuronal Cell | Myo1f1 | 4 | 4.05E-17 | 2.3211907 | 0.871 | 0.375 | 8.10E-14 |
| Neuronal Cell | Rap1gap2 | 4 | 4.89E-17 | 2.64241227 | 0.903 | 0.581 | 9.77E-14 |
| Neuronal Cell | Sptb | 4 | 5.65E-17 | 2.28103166 | 0.855 | 0.389 | 1.13E-13 |
| Neuronal Cell | NEWGENE-2 | 4 | 6.32E-17 | 1.04326733 | 0.758 | 0.3 | 1.26E-13 |
| Neuronal Cell | LOC1009119 | 4 | 8.99E-17 | 2.47317277 | 0.581 | 0.093 | 1.80E-13 |
| Neuronal Cell | Stk26 | 4 | 1.58E-16 | 2.88112237 | 0.306 | 0.032 | 3.15E-13 |
| Neuronal Cell | LOC690097 | 4 | 2.12E-16 | 4.63117754 | 0.565 | 0.16 | 4.24E-13 |
| Neuronal Cell | Cd681 | 4 | 2.30E-16 | 1.36047874 | 0.677 | 0.24 | 4.61E-13 |
| Neuronal Cell | Trem2 | 4 | 2.97E-16 | 3.81209563 | 0.694 | 0.151 | 5.93E-13 |
| Neuronal Cell | Bmx | 4 | 3.28E-16 | 1.64185676 | 0.371 | 0.068 | 6.55E-13 |
| Neuronal Cell | Nkain3 | 4 | 3.55E-16 | 1.20823684 | 0.597 | 0.204 | 7.09E-13 |
| Neuronal Cell | Itgb2 | 4 | 5.32E-16 | 3.0893212 | 0.613 | 0.104 | 1.06E-12 |
| Neuronal Cell | Tagln | 4 | 6.86E-16 | 2.45885708 | 0.435 | 0.082 | 1.37E-12 |
| Neuronal Cell | Edar | 4 | 7.39E-16 | 0.50566091 | 0.484 | 0.047 | 1.48E-12 |
| Neuronal Cell | Pstpip2 | 4 | 8.28E-16 | 3.06048906 | 0.774 | 0.467 | 1.66E-12 |
| Neuronal Cell | LOC690045 | 4 | 9.31E-16 | 3.45439804 | 0.71 | 0.272 | 1.86E-12 |
| Neuronal Cell | Fgf13 | 4 | 1.12E-15 | 1.7821123 | 0.839 | 0.512 | 2.24E-12 |
| Neuronal Cell | Mnda | 4 | 2.66E-15 | 1.46225236 | 0.645 | 0.123 | 5.32E-12 |
| Neuronal Cell | Cfh | 4 | 2.84E-15 | 2.89126974 | 0.903 | 0.598 | 5.68E-12 |
| Neuronal Cell | Rec114 | 4 | 2.91E-15 | 1.73878657 | 0.581 | 0.189 | 5.82E-12 |
| Neuronal Cell | Slco5a1 | 4 | 3.42E-15 | 2.68578391 | 0.79 | 0.27 | 6.84E-12 |
| Neuronal Cell | Marchf1 | 4 | 3.49E-15 | 3.06393058 | 0.758 | 0.386 | 6.97E-12 |
| Neuronal Cell | Des | 4 | 5.49E-15 | 1.96625106 | 0.758 | 0.277 | 1.10E-11 |
| Neuronal Cell | Ncapg | 4 | 5.76E-15 | 1.72649292 | 0.645 | 0.198 | 1.15E-11 |
| Neuronal Cell | Hck | 4 | 9.26E-15 | 3.19413291 | 0.823 | 0.425 | 1.85E-11 |
| Neuronal Cell | Slco2b1 | 4 | 1.07E-14 | 2.52504524 | 0.516 | 0.193 | 2.15E-11 |

| | | | | | | | |
|---|---|---|---|---|---|---|---|
| Neuronal Cell | Camk41 | 4 | 1.15E-14 | 0.32494938 | 0.823 | 0.535 | 2.29E-11 |
| Neuronal Cell | Stard10 | 4 | 1.18E-14 | 2.36727579 | 0.645 | 0.116 | 2.36E-11 |
| Neuronal Cell | Cd96 | 4 | 1.44E-14 | 0.44483084 | 0.661 | 0.202 | 2.88E-11 |
| Neuronal Cell | Ppargc1a | 4 | 1.48E-14 | 1.2978721 | 0.871 | 0.516 | 2.97E-11 |
| Neuronal Cell | Atp2b2 | 4 | 2.03E-14 | 0.88366459 | 0.806 | 0.456 | 4.06E-11 |
| Neuronal Cell | Fosb | 4 | 2.22E-14 | 1.89754801 | 0.597 | 0.172 | 4.43E-11 |
| Neuronal Cell | Cyp26b11 | 4 | 2.27E-14 | 1.05293802 | 0.806 | 0.514 | 4.54E-11 |
| Neuronal Cell | Lsamp | 4 | 2.48E-14 | 0.75628154 | 0.774 | 0.305 | 4.96E-11 |
| Neuronal Cell | Pf4 | 4 | 3.90E-14 | 4.6218756 | 0.339 | 0.056 | 7.79E-11 |
| Neuronal Cell | Nckap1l | 4 | 3.97E-14 | 4.17234333 | 0.516 | 0.167 | 7.95E-11 |
| Neuronal Cell | Slc30a3 | 4 | 5.40E-14 | 1.23932379 | 0.661 | 0.27 | 1.08E-10 |
| Neuronal Cell | Rbp11 | 4 | 6.94E-14 | 1.96057761 | 0.516 | 0.135 | 1.39E-10 |
| Neuronal Cell | Hspb7 | 4 | 8.79E-14 | 2.56619195 | 0.79 | 0.258 | 1.76E-10 |
| Neuronal Cell | Fmod1 | 4 | 9.41E-14 | 2.40603727 | 0.661 | 0.228 | 1.88E-10 |
| Neuronal Cell | Rspo1 | 4 | 9.74E-14 | 1.75205899 | 0.806 | 0.514 | 1.95E-10 |
| Neuronal Cell | Tmsb4x | 4 | 1.02E-13 | 2.2537748 | 0.871 | 0.453 | 2.04E-10 |
| Neuronal Cell | Clec7a | 4 | 1.04E-13 | 2.81321585 | 0.419 | 0.123 | 2.07E-10 |
| Neuronal Cell | Rgs1 | 4 | 1.28E-13 | 6.08401651 | 0.306 | 0.014 | 2.55E-10 |
| Neuronal Cell | Lgmn | 4 | 1.29E-13 | 2.3695945 | 0.823 | 0.475 | 2.57E-10 |
| Neuronal Cell | Lilrb4 | 4 | 1.51E-13 | 4.67087084 | 0.597 | 0.214 | 3.03E-10 |
| Neuronal Cell | Cep551 | 4 | 1.53E-13 | 3.63921564 | 0.726 | 0.337 | 3.05E-10 |
| Neuronal Cell | Cabcoco1 | 4 | 1.97E-13 | 1.38061324 | 0.726 | 0.233 | 3.94E-10 |
| Neuronal Cell | Ighm1 | 4 | 3.01E-13 | 1.47835468 | 0.597 | 0.223 | 6.03E-10 |
| Neuronal Cell | Slc38a3 | 4 | 4.37E-13 | 2.15416286 | 0.516 | 0.118 | 8.74E-10 |
| Neuronal Cell | AC128394.21 | 4 | 4.58E-13 | 1.09488925 | 0.694 | 0.335 | 9.15E-10 |
| Neuronal Cell | Mmrn1 | 4 | 7.08E-13 | 0.82748695 | 0.484 | 0.132 | 1.42E-09 |
| Neuronal Cell | Prkar2b | 4 | 7.10E-13 | 0.47128916 | 0.726 | 0.316 | 1.42E-09 |
| Neuronal Cell | Cdh81 | 4 | 7.15E-13 | 0.35095295 | 0.726 | 0.332 | 1.43E-09 |
| Neuronal Cell | Acta1 | 4 | 8.00E-13 | 2.42913868 | 0.565 | 0.17 | 1.60E-09 |
| Neuronal Cell | Thbs11 | 4 | 8.81E-13 | 0.48974391 | 0.548 | 0.163 | 1.76E-09 |
| Neuronal Cell | Bicdl1 | 4 | 9.20E-13 | 2.14397133 | 0.613 | 0.198 | 1.84E-09 |

| | | | | | | | |
|---|---|---|---|---|---|---|---|
| Neuronal Cell | Lcp21 | 4 | 9.39E-13 | 2.8188796 | 0.774 | 0.386 | 1.88E-09 |
| Neuronal Cell | AABR070015 | 4 | 9.45E-13 | 2.66202921 | 0.629 | 0.233 | 1.89E-09 |
| Neuronal Cell | Kcng2 | 4 | 1.02E-12 | 2.09415606 | 0.71 | 0.263 | 2.05E-09 |
| Neuronal Cell | Fth1 | 4 | 1.18E-12 | 2.25771433 | 0.79 | 0.319 | 2.37E-09 |
| Neuronal Cell | Sphkap | 4 | 1.20E-12 | 1.34688556 | 0.823 | 0.568 | 2.41E-09 |
| Neuronal Cell | AABR070251 | 4 | 1.24E-12 | 0.82893737 | 0.823 | 0.496 | 2.49E-09 |
| Neuronal Cell | Syk1 | 4 | 1.68E-12 | 3.20932524 | 0.823 | 0.388 | 3.35E-09 |
| Neuronal Cell | Ppp1r3a | 4 | 1.85E-12 | 1.73696709 | 0.581 | 0.153 | 3.70E-09 |
| Neuronal Cell | Pik3cd | 4 | 2.07E-12 | 1.98708434 | 0.806 | 0.465 | 4.15E-09 |
| Neuronal Cell | Zc3h12d | 4 | 2.07E-12 | 2.10942646 | 0.742 | 0.319 | 4.15E-09 |
| Neuronal Cell | Hk2 | 4 | 2.12E-12 | 2.12495468 | 0.742 | 0.375 | 4.23E-09 |
| Neuronal Cell | AABR070635 | 4 | 2.15E-12 | 0.89693696 | 0.71 | 0.214 | 4.30E-09 |
| Neuronal Cell | Reg3g1 | 4 | 2.38E-12 | 1.54559019 | 0.548 | 0.163 | 4.77E-09 |
| Neuronal Cell | Knl1 | 4 | 2.48E-12 | 0.98851705 | 0.71 | 0.204 | 4.95E-09 |
| Neuronal Cell | Tbx51 | 4 | 3.06E-12 | 1.73292811 | 0.419 | 0.16 | 6.12E-09 |
| Neuronal Cell | Emid1 | 4 | 3.07E-12 | 1.55770596 | 0.597 | 0.204 | 6.13E-09 |
| Neuronal Cell | Scube1 | 4 | 3.33E-12 | 0.77527096 | 0.516 | 0.102 | 6.66E-09 |
| Neuronal Cell | Lgr61 | 4 | 4.10E-12 | 0.39884733 | 0.742 | 0.256 | 8.21E-09 |
| Neuronal Cell | Wdfy4 | 4 | 4.76E-12 | 3.84564098 | 0.355 | 0.082 | 9.53E-09 |
| Neuronal Cell | Trim54 | 4 | 5.18E-12 | 1.13119978 | 0.71 | 0.282 | 1.04E-08 |
| Neuronal Cell | AABR070179 | 4 | 6.29E-12 | 0.36614798 | 0.661 | 0.223 | 1.26E-08 |
| Neuronal Cell | St8sia11 | 4 | 6.66E-12 | 0.37797308 | 0.548 | 0.142 | 1.33E-08 |
| Neuronal Cell | Rnf207 | 4 | 7.48E-12 | 0.73782159 | 0.774 | 0.325 | 1.50E-08 |
| Neuronal Cell | Pla2g5 | 4 | 9.70E-12 | 0.55560676 | 0.758 | 0.419 | 1.94E-08 |
| Neuronal Cell | Gna15 | 4 | 1.04E-11 | 2.90633026 | 0.71 | 0.426 | 2.07E-08 |
| Neuronal Cell | Frmd4b1 | 4 | 1.04E-11 | 2.07891082 | 0.661 | 0.274 | 2.08E-08 |
| Neuronal Cell | Trpc3 | 4 | 1.17E-11 | 0.62875836 | 0.774 | 0.418 | 2.34E-08 |
| Neuronal Cell | Lum | 4 | 1.26E-11 | 1.98094394 | 0.823 | 0.516 | 2.51E-08 |
| Neuronal Cell | Eef1a2 | 4 | 1.30E-11 | 2.28734696 | 0.645 | 0.204 | 2.60E-08 |
| Neuronal Cell | Scn5a | 4 | 1.32E-11 | 0.81517111 | 0.661 | 0.2 | 2.65E-08 |
| Neuronal Cell | Gria2 | 4 | 1.35E-11 | 3.66511845 | 0.403 | 0.095 | 2.70E-08 |

| | | | | | | | |
|---|---|---|---|---|---|---|---|
| Neuronal Cell | Pamr11 | 4 | 1.79E-11 | 0.60368022 | 0.774 | 0.4 | 3.59E-08 |
| Neuronal Cell | Cd163 | 4 | 1.93E-11 | 3.4019946 | 0.823 | 0.5 | 3.85E-08 |
| Neuronal Cell | Rbm38 | 4 | 2.00E-11 | 0.88672317 | 0.758 | 0.356 | 4.00E-08 |
| Neuronal Cell | Slc4a41 | 4 | 2.55E-11 | 0.86694743 | 0.806 | 0.291 | 5.10E-08 |
| Neuronal Cell | Alox5 | 4 | 2.82E-11 | 3.05559046 | 0.806 | 0.358 | 5.64E-08 |
| Neuronal Cell | Cyth4 | 4 | 2.99E-11 | 1.67226265 | 0.806 | 0.553 | 5.98E-08 |
| Neuronal Cell | Aoah | 4 | 3.05E-11 | 2.80743568 | 0.79 | 0.496 | 6.09E-08 |
| Neuronal Cell | Armc4 | 4 | 3.20E-11 | 0.90483447 | 0.532 | 0.158 | 6.41E-08 |
| Neuronal Cell | Myoz2 | 4 | 3.45E-11 | 1.65613284 | 0.597 | 0.184 | 6.90E-08 |
| Neuronal Cell | Cd55 | 4 | 4.17E-11 | 1.1688548 | 0.355 | 0.095 | 8.34E-08 |
| Neuronal Cell | LOC1003614 | 4 | 4.35E-11 | 1.7007083 | 0.613 | 0.24 | 8.69E-08 |
| Neuronal Cell | Myo18b | 4 | 5.37E-11 | 1.752353 | 0.871 | 0.607 | 1.07E-07 |
| Neuronal Cell | Slc25a21 | 4 | 5.47E-11 | 0.79412137 | 0.726 | 0.396 | 1.09E-07 |
| Neuronal Cell | Adap2 | 4 | 6.50E-11 | 3.54861643 | 0.581 | 0.319 | 1.30E-07 |
| Neuronal Cell | Cd63 | 4 | 7.42E-11 | 2.43419728 | 0.823 | 0.514 | 1.48E-07 |
| Neuronal Cell | Colq | 4 | 8.44E-11 | 0.49659201 | 0.758 | 0.43 | 1.69E-07 |
| Neuronal Cell | Bub1b | 4 | 8.76E-11 | 0.66769814 | 0.629 | 0.228 | 1.75E-07 |
| Neuronal Cell | Gria41 | 4 | 1.02E-10 | 1.01706291 | 0.532 | 0.209 | 2.05E-07 |
| Neuronal Cell | Ftl1 | 4 | 1.05E-10 | 2.09724329 | 0.629 | 0.258 | 2.09E-07 |
| Neuronal Cell | Mb | 4 | 1.23E-10 | 1.86643267 | 0.855 | 0.602 | 2.45E-07 |
| Neuronal Cell | Itgal1 | 4 | 1.28E-10 | 2.80324452 | 0.565 | 0.177 | 2.56E-07 |
| Neuronal Cell | LOC1003644 | 4 | 1.29E-10 | 1.93810113 | 0.613 | 0.281 | 2.58E-07 |
| Neuronal Cell | Aldh1a1 | 4 | 1.41E-10 | 1.67283254 | 0.403 | 0.081 | 2.82E-07 |
| Neuronal Cell | Mylk3 | 4 | 1.51E-10 | 1.55583985 | 0.823 | 0.405 | 3.03E-07 |
| Neuronal Cell | Mtus2 | 4 | 1.72E-10 | 0.36922768 | 0.403 | 0.111 | 3.44E-07 |
| Neuronal Cell | Stab1 | 4 | 1.85E-10 | 2.99020301 | 0.532 | 0.228 | 3.69E-07 |
| Neuronal Cell | Slfn41 | 4 | 1.94E-10 | 1.53623535 | 0.629 | 0.24 | 3.87E-07 |
| Neuronal Cell | Nrap | 4 | 2.12E-10 | 1.53578519 | 0.435 | 0.158 | 4.25E-07 |
| Neuronal Cell | Npnt1 | 4 | 2.49E-10 | 2.24823217 | 0.565 | 0.198 | 4.98E-07 |
| Neuronal Cell | Slco1a41 | 4 | 2.68E-10 | 2.87111417 | 0.597 | 0.219 | 5.37E-07 |
| Neuronal Cell | Rapgef5 | 4 | 2.75E-10 | 0.62994282 | 0.855 | 0.546 | 5.51E-07 |

| | | | | | | | |
|---|---|---|---|---|---|---|---|
| Neuronal Cell | AABR070525 | 4 | 2.77E-10 | 1.62399394 | 0.79 | 0.537 | 5.55E-07 |
| Neuronal Cell | S100a6 | 4 | 2.82E-10 | 1.96363278 | 0.839 | 0.418 | 5.64E-07 |
| Neuronal Cell | Cluh | 4 | 2.87E-10 | 0.43284151 | 0.823 | 0.507 | 5.73E-07 |
| Neuronal Cell | AABR070070 | 4 | 3.07E-10 | 1.15758895 | 0.694 | 0.377 | 6.13E-07 |
| Neuronal Cell | Mfap51 | 4 | 3.28E-10 | 1.23947329 | 0.871 | 0.368 | 6.57E-07 |
| Neuronal Cell | Lbp | 4 | 3.37E-10 | 1.28619793 | 0.823 | 0.563 | 6.73E-07 |
| Neuronal Cell | Rrad | 4 | 3.60E-10 | 0.68668392 | 0.613 | 0.198 | 7.21E-07 |
| Neuronal Cell | Smpx | 4 | 4.02E-10 | 0.26739836 | 0.823 | 0.572 | 8.03E-07 |
| Neuronal Cell | Abcc4 | 4 | 4.05E-10 | 1.20718379 | 0.839 | 0.535 | 8.11E-07 |
| Neuronal Cell | Btbd11 | 4 | 4.33E-10 | 1.684402 | 0.726 | 0.404 | 8.67E-07 |
| Neuronal Cell | Kif20b1 | 4 | 4.36E-10 | 0.615635 | 0.661 | 0.218 | 8.73E-07 |
| Neuronal Cell | Tango2 | 4 | 4.38E-10 | 1.12946987 | 0.774 | 0.33 | 8.75E-07 |
| Neuronal Cell | Rbfox1 | 4 | 4.95E-10 | 1.80828394 | 0.71 | 0.405 | 9.91E-07 |
| Neuronal Cell | Sned11 | 4 | 5.87E-10 | 0.49040114 | 0.516 | 0.189 | 1.17E-06 |
| Neuronal Cell | Fam163a1 | 4 | 6.46E-10 | 0.70041683 | 0.629 | 0.258 | 1.29E-06 |
| Neuronal Cell | Agtr1a1 | 4 | 9.82E-10 | 1.40792417 | 0.548 | 0.189 | 1.96E-06 |
| Neuronal Cell | Arhgef37 | 4 | 1.03E-09 | 0.52185535 | 0.855 | 0.405 | 2.06E-06 |
| Neuronal Cell | Ptprd | 4 | 1.13E-09 | 0.904985 | 0.855 | 0.546 | 2.25E-06 |
| Neuronal Cell | Ccl21 | 4 | 1.16E-09 | 0.96162819 | 0.452 | 0.17 | 2.32E-06 |
| Neuronal Cell | Uchl1 | 4 | 1.22E-09 | 1.71067248 | 0.694 | 0.186 | 2.44E-06 |
| Neuronal Cell | Dapk1 | 4 | 1.36E-09 | 1.39014758 | 0.839 | 0.584 | 2.73E-06 |
| Neuronal Cell | Emx21 | 4 | 1.81E-09 | 1.00784416 | 0.581 | 0.218 | 3.61E-06 |
| Neuronal Cell | Tpt1 | 4 | 1.83E-09 | 1.96496713 | 0.548 | 0.289 | 3.66E-06 |
| Neuronal Cell | Fmn1 | 4 | 1.91E-09 | 1.20228419 | 0.871 | 0.554 | 3.82E-06 |
| Neuronal Cell | Psap | 4 | 2.07E-09 | 1.87723788 | 0.726 | 0.344 | 4.13E-06 |
| Neuronal Cell | Timp11 | 4 | 2.43E-09 | 2.94458731 | 0.5 | 0.168 | 4.86E-06 |
| Neuronal Cell | Fabp3 | 4 | 2.44E-09 | 2.69570045 | 0.742 | 0.479 | 4.89E-06 |
| Neuronal Cell | Atp5mg | 4 | 2.84E-09 | 2.04918209 | 0.758 | 0.414 | 5.69E-06 |
| Neuronal Cell | Eda2r1 | 4 | 2.89E-09 | 1.22544435 | 0.5 | 0.216 | 5.77E-06 |
| Neuronal Cell | Camk1d1 | 4 | 2.91E-09 | 1.44768851 | 0.677 | 0.311 | 5.83E-06 |
| Neuronal Cell | Ndufa41 | 4 | 3.17E-09 | 1.5206081 | 0.806 | 0.463 | 6.34E-06 |

| | | | | | | | |
|---|---|---|---|---|---|---|---|
| Neuronal Cell | Fermt3 | 4 | 4.69E-09 | 1.90970899 | 0.597 | 0.305 | 9.38E-06 |
| Neuronal Cell | Mrc1 | 4 | 5.81E-09 | 2.90930392 | 0.742 | 0.475 | 1.16E-05 |
| Neuronal Cell | Lrrc21 | 4 | 6.84E-09 | 0.33180924 | 0.758 | 0.482 | 1.37E-05 |
| Neuronal Cell | Igfbp5 | 4 | 6.87E-09 | 1.15008577 | 0.71 | 0.323 | 1.37E-05 |
| Neuronal Cell | Kcnj5 | 4 | 7.04E-09 | 0.60565796 | 0.403 | 0.132 | 1.41E-05 |
| Neuronal Cell | Mamdc2 | 4 | 8.01E-09 | 1.36744481 | 0.677 | 0.242 | 1.60E-05 |
| Neuronal Cell | S100a4 | 4 | 8.54E-09 | 1.64154119 | 0.694 | 0.347 | 1.71E-05 |
| Neuronal Cell | Nuak1 | 4 | 1.07E-08 | 0.67226673 | 0.516 | 0.221 | 2.14E-05 |
| Neuronal Cell | Crip1 | 4 | 1.12E-08 | 2.09023515 | 0.823 | 0.514 | 2.24E-05 |
| Neuronal Cell | Lrrc7 | 4 | 1.15E-08 | 2.06767664 | 0.516 | 0.082 | 2.30E-05 |
| Neuronal Cell | Fgfr2 | 4 | 1.17E-08 | 1.86893108 | 0.323 | 0.063 | 2.35E-05 |
| Neuronal Cell | Ctsd | 4 | 1.20E-08 | 1.78443644 | 0.548 | 0.158 | 2.40E-05 |
| Neuronal Cell | Dgat2 | 4 | 1.22E-08 | 0.41605449 | 0.661 | 0.391 | 2.44E-05 |
| Neuronal Cell | Cmya5 | 4 | 1.27E-08 | 1.03638453 | 0.726 | 0.318 | 2.53E-05 |
| Neuronal Cell | AABR070546 | 4 | 1.41E-08 | 0.49403056 | 0.774 | 0.411 | 2.82E-05 |
| Neuronal Cell | Slc43a2 | 4 | 1.52E-08 | 1.36279611 | 0.806 | 0.514 | 3.04E-05 |
| Neuronal Cell | Apoe | 4 | 1.66E-08 | 2.37117662 | 0.629 | 0.196 | 3.32E-05 |
| Neuronal Cell | Dgkz | 4 | 1.67E-08 | 0.94296968 | 0.71 | 0.326 | 3.35E-05 |
| Neuronal Cell | Pdgfra1 | 4 | 1.81E-08 | 0.98102101 | 0.758 | 0.356 | 3.62E-05 |
| Neuronal Cell | Mitf | 4 | 1.88E-08 | 1.14015969 | 0.919 | 0.639 | 3.75E-05 |
| Neuronal Cell | Bcas1 | 4 | 3.13E-08 | 0.73293383 | 0.565 | 0.223 | 6.26E-05 |
| Neuronal Cell | Abca1 | 4 | 3.15E-08 | 1.34116274 | 0.887 | 0.502 | 6.30E-05 |
| Neuronal Cell | Matk | 4 | 3.15E-08 | 2.03307524 | 0.419 | 0.123 | 6.31E-05 |
| Neuronal Cell | Mgp1 | 4 | 3.57E-08 | 2.03691541 | 0.79 | 0.521 | 7.15E-05 |
| Neuronal Cell | Casq2 | 4 | 3.81E-08 | 0.61216758 | 0.839 | 0.568 | 7.62E-05 |
| Neuronal Cell | Gsn | 4 | 3.93E-08 | 1.91635038 | 0.758 | 0.495 | 7.86E-05 |
| Neuronal Cell | Kif22 | 4 | 4.37E-08 | 0.49862332 | 0.726 | 0.365 | 8.73E-05 |
| Neuronal Cell | Med12l1 | 4 | 4.55E-08 | 2.33869949 | 0.823 | 0.556 | 9.09E-05 |
| Neuronal Cell | Tnnc1 | 4 | 4.55E-08 | 1.81135034 | 0.677 | 0.419 | 9.11E-05 |
| Neuronal Cell | Mfge8 | 4 | 5.16E-08 | 2.30170986 | 0.645 | 0.381 | 0.00010312 |
| Neuronal Cell | Aqp1 | 4 | 5.23E-08 | 1.26361504 | 0.694 | 0.384 | 0.00010456 |

| | | | | | | | |
|---|---|---|---|---|---|---|---|
| Neuronal Cell | Il6r | 4 | 5.88E-08 | 2.95483 | 0.484 | 0.188 | 0.00011761 |
| Neuronal Cell | Ano5 | 4 | 6.24E-08 | 1.10265102 | 0.79 | 0.504 | 0.00012483 |
| Neuronal Cell | Atp5me | 4 | 7.22E-08 | 1.78828772 | 0.823 | 0.493 | 0.00014435 |
| Neuronal Cell | Wipf31 | 4 | 1.02E-07 | 1.13927132 | 0.661 | 0.274 | 0.00020391 |
| Neuronal Cell | Asb141 | 4 | 1.08E-07 | 1.3266217 | 0.565 | 0.26 | 0.00021529 |
| Neuronal Cell | Lpl | 4 | 1.18E-07 | 1.05761813 | 0.645 | 0.296 | 0.00023628 |
| Neuronal Cell | Myom21 | 4 | 1.18E-07 | 0.95345334 | 0.839 | 0.533 | 0.00023634 |
| Neuronal Cell | Fabp5 | 4 | 1.19E-07 | 2.55605414 | 0.403 | 0.093 | 0.00023768 |
| Neuronal Cell | Xirp2 | 4 | 1.27E-07 | 1.12955742 | 0.516 | 0.196 | 0.00025409 |
| Neuronal Cell | Ppargc1b | 4 | 1.40E-07 | 0.70115303 | 0.726 | 0.411 | 0.00027931 |
| Neuronal Cell | Polr2m | 4 | 1.61E-07 | 1.35899141 | 0.565 | 0.258 | 0.00032127 |
| Neuronal Cell | Cdkn2b | 4 | 1.61E-07 | 1.15297756 | 0.629 | 0.363 | 0.00032159 |
| Neuronal Cell | Uqcr10 | 4 | 1.69E-07 | 1.57136616 | 0.677 | 0.368 | 0.00033803 |
| Neuronal Cell | Lgals1 | 4 | 1.91E-07 | 1.5239313 | 0.613 | 0.339 | 0.00038243 |
| Neuronal Cell | Cpq1 | 4 | 1.94E-07 | 1.23727487 | 0.661 | 0.3 | 0.00038705 |
| Neuronal Cell | Arhgap22 | 4 | 2.12E-07 | 1.24112731 | 0.742 | 0.409 | 0.00042429 |
| Neuronal Cell | Cox6a1 | 4 | 2.76E-07 | 2.19573003 | 0.403 | 0.096 | 0.00055207 |
| Neuronal Cell | Serf2 | 4 | 2.97E-07 | 2.19637117 | 0.71 | 0.416 | 0.00059433 |
| Neuronal Cell | Cyba | 4 | 3.14E-07 | 1.46144445 | 0.419 | 0.158 | 0.00062837 |
| Neuronal Cell | Htra3 | 4 | 3.69E-07 | 1.16948774 | 0.581 | 0.242 | 0.00073731 |
| Neuronal Cell | Nppa1 | 4 | 4.37E-07 | 1.20512264 | 0.71 | 0.456 | 0.00087352 |
| Neuronal Cell | Ripor2 | 4 | 4.69E-07 | 1.67014318 | 0.677 | 0.396 | 0.00093792 |
| Neuronal Cell | Ptgs1 | 4 | 5.42E-07 | 0.69731806 | 0.371 | 0.102 | 0.00108441 |
| Neuronal Cell | Ppara | 4 | 5.44E-07 | 0.36618087 | 0.597 | 0.289 | 0.00108891 |
| Neuronal Cell | Coq8a1 | 4 | 6.81E-07 | 1.09148196 | 0.71 | 0.446 | 0.00136109 |
| Neuronal Cell | Cd300a | 4 | 7.11E-07 | 2.16870929 | 0.79 | 0.505 | 0.00142101 |
| Neuronal Cell | Cd83 | 4 | 7.68E-07 | 1.84611528 | 0.323 | 0.072 | 0.00153506 |
| Neuronal Cell | Slc25a13 | 4 | 8.17E-07 | 1.73618092 | 0.403 | 0.151 | 0.00163312 |
| Neuronal Cell | Snca1 | 4 | 8.41E-07 | 0.84346451 | 0.71 | 0.393 | 0.00168236 |
| Neuronal Cell | Slc6a6 | 4 | 8.54E-07 | 0.77849207 | 0.661 | 0.391 | 0.00170738 |
| Neuronal Cell | Fhl2 | 4 | 9.36E-07 | 0.51209502 | 0.581 | 0.27 | 0.00187131 |

| | | | | | | | |
|---|---|---|---|---|---|---|---|
| Neuronal Cell | Lcp1 | 4 | 9.98E-07 | 2.19482273 | 0.806 | 0.504 | 0.00199569 |
| Neuronal Cell | Adcy51 | 4 | 1.01E-06 | 0.53397125 | 0.855 | 0.574 | 0.00202699 |
| Neuronal Cell | Scarb1 | 4 | 1.16E-06 | 1.22968829 | 0.516 | 0.212 | 0.00232148 |
| Neuronal Cell | Fcgr2b | 4 | 1.17E-06 | 2.7116219 | 0.355 | 0.053 | 0.00234489 |
| Neuronal Cell | Slc9a91 | 4 | 1.69E-06 | 2.09391238 | 0.903 | 0.579 | 0.00338491 |
| Neuronal Cell | Tnni3 | 4 | 1.79E-06 | 1.30876828 | 0.742 | 0.442 | 0.00357946 |
| Neuronal Cell | Runx11 | 4 | 2.02E-06 | 1.29411588 | 0.677 | 0.395 | 0.00404408 |
| Neuronal Cell | Obscn | 4 | 2.32E-06 | 1.15619324 | 0.629 | 0.333 | 0.00463799 |
| Neuronal Cell | Gpcpd1 | 4 | 2.37E-06 | 0.45496104 | 0.774 | 0.495 | 0.00474496 |
| Neuronal Cell | Fbln2 | 4 | 3.07E-06 | 0.6653638 | 0.661 | 0.319 | 0.00614181 |
| Neuronal Cell | Depdc1b | 4 | 3.18E-06 | 1.62888985 | 0.468 | 0.196 | 0.00636068 |
| Neuronal Cell | Atp5f1b | 4 | 4.03E-06 | 1.32905727 | 0.887 | 0.591 | 0.00806346 |
| Neuronal Cell | Mpz | 4 | 4.50E-06 | 3.6933973 | 0.532 | 0.214 | 0.00900684 |
| Neuronal Cell | Slc37a2 | 4 | 6.38E-06 | 3.31912049 | 0.323 | 0.054 | 0.01275364 |
| Neuronal Cell | Pi16 | 4 | 9.47E-06 | 1.07335432 | 0.468 | 0.191 | 0.01894004 |
| Neuronal Cell | Nebl | 4 | 1.84E-05 | 0.61899008 | 0.758 | 0.468 | 0.03689189 |
| Neuronal Cell | Lgals3 | 4 | 2.45E-05 | 1.00881692 | 0.452 | 0.193 | 0.04904531 |
| Neuronal Cell | Runx2 | 5 | 2.79E-11 | 3.31843785 | 0.789 | 0.504 | 5.58E-08 |
| Neuronal Cell | Pde1c | 5 | 7.70E-10 | 1.92037782 | 0.825 | 0.522 | 1.54E-06 |
| Neuronal Cell | Vipr2 | 5 | 3.99E-06 | 0.47275856 | 0.193 | 0.456 | 0.00797449 |
| Lymphatic Endothelial Cell | LOC681458 | 0 | 6.57E-26 | 5.04382289 | 0.447 | 0.085 | 1.31E-22 |
| Lymphatic Endothelial Cell | Adamts20 | 0 | 1.58E-22 | 3.08391874 | 0.59 | 0.239 | 3.17E-19 |
| Lymphatic Endothelial Cell | Il1rn | 0 | 2.11E-21 | 3.17010995 | 0.547 | 0.21 | 4.22E-18 |
| Lymphatic Endothelial Cell | Scn1a | 0 | 2.28E-20 | 0.47871001 | 0.621 | 0.291 | 4.56E-17 |
| Lymphatic Endothelial Cell | Rapgef5 | 0 | 3.23E-20 | 1.20346979 | 0.925 | 0.606 | 6.46E-17 |
| Lymphatic Endothelial Cell | Adam12 | 0 | 8.10E-20 | 1.89728637 | 0.826 | 0.509 | 1.62E-16 |
| Lymphatic Endothelial Cell | AABR070058 | 0 | 3.33E-18 | 2.2569054 | 0.634 | 0.319 | 6.66E-15 |
| Lymphatic Endothelial Cell | Tmem163 | 0 | 4.66E-18 | 0.625491 | 0.391 | 0.104 | 9.32E-15 |
| Lymphatic Endothelial Cell | Bcas1 | 0 | 9.35E-18 | 0.88372784 | 0.391 | 0.111 | 1.87E-14 |
| Lymphatic Endothelial Cell | Myo3b | 0 | 1.93E-17 | 1.81102443 | 0.621 | 0.366 | 3.86E-14 |
| Lymphatic Endothelial Cell | Gabrb2 | 0 | 4.75E-16 | 2.3810794 | 0.615 | 0.358 | 9.49E-13 |

| | | | | | | | |
|---|---|---|---|---|---|---|---|
| Lymphatic Endothelial Cell | AABR070339 | 0 | 5.15E-16 | 2.34674664 | 0.553 | 0.216 | 1.03E-12 |
| Lymphatic Endothelial Cell | Serping1 | 0 | 5.83E-15 | 1.43580116 | 0.64 | 0.274 | 1.17E-11 |
| Lymphatic Endothelial Cell | Ccl24 | 0 | 5.11E-14 | 2.41134731 | 0.547 | 0.189 | 1.02E-10 |
| Lymphatic Endothelial Cell | Fam205a | 0 | 5.55E-14 | 1.4631813 | 0.733 | 0.385 | 1.11E-10 |
| Lymphatic Endothelial Cell | Sptb | 0 | 6.23E-14 | 1.53811926 | 0.559 | 0.215 | 1.25E-10 |
| Lymphatic Endothelial Cell | AABR070014 | 0 | 4.08E-13 | 2.2326597 | 0.441 | 0.191 | 8.15E-10 |
| Lymphatic Endothelial Cell | C6 | 0 | 4.12E-13 | 0.79565413 | 0.404 | 0.145 | 8.24E-10 |
| Lymphatic Endothelial Cell | Kif11 | 0 | 9.94E-13 | 0.39349711 | 0.441 | 0.114 | 1.99E-09 |
| Lymphatic Endothelial Cell | Usp2 | 0 | 1.52E-12 | 1.32364507 | 0.553 | 0.193 | 3.04E-09 |
| Lymphatic Endothelial Cell | Rrad | 0 | 6.59E-12 | 1.43448548 | 0.615 | 0.293 | 1.32E-08 |
| Lymphatic Endothelial Cell | Abca4 | 0 | 2.86E-11 | 1.66376029 | 0.596 | 0.32 | 5.73E-08 |
| Lymphatic Endothelial Cell | Emid1 | 0 | 4.30E-11 | 1.11213556 | 0.739 | 0.451 | 8.60E-08 |
| Lymphatic Endothelial Cell | RGD1562392 | 0 | 4.94E-11 | 0.34973503 | 0.484 | 0.228 | 9.87E-08 |
| Lymphatic Endothelial Cell | Brca1 | 0 | 9.49E-11 | 0.90840074 | 0.634 | 0.346 | 1.90E-07 |
| Lymphatic Endothelial Cell | Mis18bp1 | 0 | 3.98E-10 | 1.01687201 | 0.609 | 0.346 | 7.97E-07 |
| Lymphatic Endothelial Cell | AABR070449 | 0 | 4.66E-10 | 0.6358933 | 0.634 | 0.303 | 9.31E-07 |
| Lymphatic Endothelial Cell | Col8a2 | 0 | 5.20E-10 | 1.10177458 | 0.677 | 0.419 | 1.04E-06 |
| Lymphatic Endothelial Cell | Trib3 | 0 | 1.89E-09 | 0.87094989 | 0.59 | 0.325 | 3.77E-06 |
| Lymphatic Endothelial Cell | AC096301.1 | 0 | 2.27E-09 | 0.86574296 | 0.59 | 0.337 | 4.55E-06 |
| Lymphatic Endothelial Cell | Cdh4 | 0 | 2.67E-09 | 1.83023346 | 0.41 | 0.153 | 5.33E-06 |
| Lymphatic Endothelial Cell | Fermt3 | 0 | 1.57E-08 | 0.31681534 | 0.41 | 0.138 | 3.15E-05 |
| Lymphatic Endothelial Cell | Cers6 | 0 | 8.28E-08 | 0.602068 | 0.671 | 0.363 | 0.00016554 |
| Lymphatic Endothelial Cell | Lrrc17 | 0 | 8.67E-08 | 0.47361561 | 0.54 | 0.254 | 0.00017347 |
| Lymphatic Endothelial Cell | Flt3 | 0 | 3.02E-07 | 0.42628113 | 0.373 | 0.119 | 0.00060328 |
| Lymphatic Endothelial Cell | Trabd2b | 0 | 4.86E-07 | 0.57259188 | 0.683 | 0.419 | 0.00097278 |
| Lymphatic Endothelial Cell | Asb18 | 1 | 6.36E-13 | 0.69285022 | 0.15 | 0.444 | 1.27E-09 |
| Lymphatic Endothelial Cell | Pygm | 1 | 1.29E-12 | 0.60506352 | 0.17 | 0.508 | 2.58E-09 |
| Lymphatic Endothelial Cell | Pld3 | 1 | 1.83E-12 | 0.28075097 | 0.137 | 0.42 | 3.67E-09 |
| Lymphatic Endothelial Cell | Me3 | 1 | 3.67E-12 | 0.50323488 | 0.209 | 0.533 | 7.33E-09 |
| Lymphatic Endothelial Cell | Slco5a1 | 1 | 5.58E-12 | 1.19999632 | 0.124 | 0.462 | 1.12E-08 |
| Lymphatic Endothelial Cell | Esr1 | 1 | 7.12E-12 | 0.45092366 | 0.163 | 0.466 | 1.42E-08 |

| | | | | | | | |
|---|---|---|---|---|---|---|---|
| Lymphatic Endothelial Cell | Nmrk2 | 1 | 7.82E-12 | 1.78906978 | 0.131 | 0.482 | 1.56E-08 |
| Lymphatic Endothelial Cell | Rspo3 | 1 | 1.37E-11 | 0.31679852 | 0.216 | 0.546 | 2.74E-08 |
| Lymphatic Endothelial Cell | Lrrc2 | 1 | 5.67E-11 | 0.52931976 | 0.229 | 0.561 | 1.13E-07 |
| Lymphatic Endothelial Cell | Obscn | 1 | 1.00E-10 | 0.58592103 | 0.183 | 0.482 | 2.00E-07 |
| Lymphatic Endothelial Cell | S100a4 | 1 | 1.58E-10 | 0.61195298 | 0.15 | 0.412 | 3.15E-07 |
| Lymphatic Endothelial Cell | Mapk10 | 1 | 1.81E-10 | 1.01984995 | 0.222 | 0.519 | 3.61E-07 |
| Lymphatic Endothelial Cell | Lhfpl2 | 1 | 2.00E-10 | 0.39047236 | 0.196 | 0.474 | 4.00E-07 |
| Lymphatic Endothelial Cell | Dtna | 1 | 2.44E-10 | 0.54467241 | 0.19 | 0.474 | 4.88E-07 |
| Lymphatic Endothelial Cell | Pfkfb3 | 1 | 3.19E-10 | 0.32842478 | 0.196 | 0.457 | 6.38E-07 |
| Lymphatic Endothelial Cell | Txlnb | 1 | 5.51E-10 | 0.83565151 | 0.242 | 0.494 | 1.10E-06 |
| Lymphatic Endothelial Cell | Lgmn | 1 | 1.05E-09 | 0.37583316 | 0.203 | 0.455 | 2.10E-06 |
| Lymphatic Endothelial Cell | Pi15 | 1 | 1.35E-09 | 0.90820123 | 0.242 | 0.539 | 2.70E-06 |
| Lymphatic Endothelial Cell | Edil3 | 1 | 3.16E-09 | 0.98838217 | 0.235 | 0.509 | 6.32E-06 |
| Lymphatic Endothelial Cell | Scarb1 | 1 | 3.55E-09 | 0.43270487 | 0.261 | 0.513 | 7.10E-06 |
| Lymphatic Endothelial Cell | Fabp5 | 1 | 3.66E-09 | 0.61425046 | 0.235 | 0.531 | 7.31E-06 |
| Lymphatic Endothelial Cell | AABR070294 | 1 | 2.70E-08 | 1.60095405 | 0.131 | 0.382 | 5.40E-05 |
| Lymphatic Endothelial Cell | Synpo2l | 1 | 2.87E-08 | 1.8791869 | 0.203 | 0.455 | 5.73E-05 |
| Lymphatic Endothelial Cell | Pip5k1b | 1 | 3.93E-08 | 0.43045066 | 0.268 | 0.548 | 7.86E-05 |
| Lymphatic Endothelial Cell | Syt14 | 1 | 9.37E-08 | 0.42435422 | 0.235 | 0.487 | 0.0001873 |
| Lymphatic Endothelial Cell | Kcnj3 | 1 | 1.69E-07 | 1.01682747 | 0.255 | 0.506 | 0.00033708 |
| Lymphatic Endothelial Cell | Col8a21 | 1 | 2.42E-07 | 0.38874242 | 0.268 | 0.528 | 0.00048471 |
| Lymphatic Endothelial Cell | Mylk | 1 | 2.53E-07 | 0.32463944 | 0.327 | 0.582 | 0.00050516 |
| Lymphatic Endothelial Cell | Popdc2 | 1 | 7.53E-07 | 1.57198496 | 0.288 | 0.548 | 0.00150662 |
| Lymphatic Endothelial Cell | Cst6 | 1 | 7.91E-07 | 1.40069006 | 0.261 | 0.529 | 0.00158232 |
| Lymphatic Endothelial Cell | Cntn6 | 2 | 4.65E-30 | 1.79814465 | 0.593 | 0.153 | 9.29E-27 |
| Lymphatic Endothelial Cell | Kif4a | 2 | 2.87E-24 | 1.57688016 | 0.6 | 0.186 | 5.74E-21 |
| Lymphatic Endothelial Cell | AABR070584 | 2 | 8.72E-24 | 1.51059106 | 0.621 | 0.247 | 1.74E-20 |
| Lymphatic Endothelial Cell | Sh3rf2 | 2 | 5.59E-23 | 1.98193419 | 0.621 | 0.249 | 1.12E-19 |
| Lymphatic Endothelial Cell | AABR070294 | 2 | 7.47E-23 | 1.51408877 | 0.641 | 0.255 | 1.49E-19 |
| Lymphatic Endothelial Cell | Kcnj8 | 2 | 5.36E-19 | 0.60920873 | 0.69 | 0.353 | 1.07E-15 |
| Lymphatic Endothelial Cell | Adra1b | 2 | 1.68E-18 | 2.2105483 | 0.71 | 0.371 | 3.35E-15 |

| | | | | | | | |
|---|---|---|---|---|---|---|---|
| Lymphatic Endothelial Cell | Wnk2 | 2 | 4.52E-18 | 1.11643052 | 0.703 | 0.388 | 9.05E-15 |
| Lymphatic Endothelial Cell | Epha3 | 2 | 4.90E-18 | 2.31447492 | 0.634 | 0.28 | 9.81E-15 |
| Lymphatic Endothelial Cell | Nmrk21 | 2 | 5.76E-18 | 1.01179837 | 0.676 | 0.347 | 1.15E-14 |
| Lymphatic Endothelial Cell | Ccdc148 | 2 | 8.25E-18 | 1.39807171 | 0.697 | 0.376 | 1.65E-14 |
| Lymphatic Endothelial Cell | Pfkfb1 | 2 | 1.05E-17 | 1.38700819 | 0.655 | 0.3 | 2.09E-14 |
| Lymphatic Endothelial Cell | Pi151 | 2 | 1.65E-17 | 1.28235945 | 0.697 | 0.426 | 3.29E-14 |
| Lymphatic Endothelial Cell | Slc5a10 | 2 | 6.97E-17 | 2.64273969 | 0.483 | 0.194 | 1.39E-13 |
| Lymphatic Endothelial Cell | Kcng2 | 2 | 1.13E-16 | 1.32589744 | 0.703 | 0.353 | 2.26E-13 |
| Lymphatic Endothelial Cell | Itgax | 2 | 8.07E-16 | 2.98125211 | 0.628 | 0.302 | 1.61E-12 |
| Lymphatic Endothelial Cell | Aspa | 2 | 1.09E-15 | 1.05360863 | 0.697 | 0.34 | 2.18E-12 |
| Lymphatic Endothelial Cell | AABR070581 | 2 | 1.32E-15 | 2.20312133 | 0.483 | 0.171 | 2.64E-12 |
| Lymphatic Endothelial Cell | Susd4 | 2 | 2.31E-15 | 1.84033134 | 0.572 | 0.249 | 4.62E-12 |
| Lymphatic Endothelial Cell | Col19a1 | 2 | 2.80E-15 | 1.29882627 | 0.51 | 0.156 | 5.61E-12 |
| Lymphatic Endothelial Cell | S100a41 | 2 | 7.54E-15 | 0.77702612 | 0.641 | 0.29 | 1.51E-11 |
| Lymphatic Endothelial Cell | Thbs1 | 2 | 1.10E-14 | 1.69552648 | 0.662 | 0.385 | 2.21E-11 |
| Lymphatic Endothelial Cell | Rgs20 | 2 | 1.35E-14 | 1.91499609 | 0.669 | 0.4 | 2.70E-11 |
| Lymphatic Endothelial Cell | Frmd5 | 2 | 2.00E-14 | 0.50134279 | 0.607 | 0.262 | 4.00E-11 |
| Lymphatic Endothelial Cell | Sgo2 | 2 | 2.57E-14 | 1.20757253 | 0.662 | 0.294 | 5.14E-11 |
| Lymphatic Endothelial Cell | Mettl24 | 2 | 2.67E-14 | 0.92541847 | 0.469 | 0.136 | 5.33E-11 |
| Lymphatic Endothelial Cell | Ak5 | 2 | 6.55E-14 | 1.76332755 | 0.69 | 0.413 | 1.31E-10 |
| Lymphatic Endothelial Cell | Slco5a11 | 2 | 7.14E-14 | 1.74094653 | 0.669 | 0.327 | 1.43E-10 |
| Lymphatic Endothelial Cell | Veph1 | 2 | 1.08E-13 | 2.18532414 | 0.517 | 0.216 | 2.16E-10 |
| Lymphatic Endothelial Cell | Gfpt2 | 2 | 2.50E-13 | 0.47319233 | 0.572 | 0.219 | 4.99E-10 |
| Lymphatic Endothelial Cell | Clnk | 2 | 4.75E-13 | 2.97582725 | 0.434 | 0.123 | 9.51E-10 |
| Lymphatic Endothelial Cell | Cit | 2 | 4.81E-13 | 0.74699911 | 0.572 | 0.254 | 9.62E-10 |
| Lymphatic Endothelial Cell | Shroom2 | 2 | 5.03E-13 | 0.61694145 | 0.731 | 0.333 | 1.01E-09 |
| Lymphatic Endothelial Cell | Pygm1 | 2 | 6.84E-13 | 1.01624452 | 0.669 | 0.383 | 1.37E-09 |
| Lymphatic Endothelial Cell | Unc5d | 2 | 1.19E-12 | 3.34454256 | 0.407 | 0.148 | 2.38E-09 |
| Lymphatic Endothelial Cell | Gabrb1 | 2 | 1.51E-12 | 1.39408414 | 0.393 | 0.133 | 3.02E-09 |
| Lymphatic Endothelial Cell | Itgam | 2 | 3.10E-12 | 0.94426948 | 0.676 | 0.347 | 6.20E-09 |
| Lymphatic Endothelial Cell | AABR070265 | 2 | 3.62E-12 | 0.64577767 | 0.51 | 0.211 | 7.24E-09 |

| | | | | | | | |
|---|---|---|---|---|---|---|---|
| Lymphatic Endothelial Cell | Fgf13 | 2 | 4.53E-12 | 1.04401987 | 0.676 | 0.398 | 9.06E-09 |
| Lymphatic Endothelial Cell | Prss12 | 2 | 9.20E-12 | 0.7006389 | 0.745 | 0.415 | 1.84E-08 |
| Lymphatic Endothelial Cell | Podnl1 | 2 | 9.81E-12 | 2.18350839 | 0.6 | 0.31 | 1.96E-08 |
| Lymphatic Endothelial Cell | Me31 | 2 | 1.07E-11 | 0.53658403 | 0.717 | 0.406 | 2.15E-08 |
| Lymphatic Endothelial Cell | Nckap1l | 2 | 1.14E-11 | 0.62968305 | 0.628 | 0.352 | 2.28E-08 |
| Lymphatic Endothelial Cell | Ankrd9 | 2 | 1.87E-11 | 0.69580097 | 0.634 | 0.343 | 3.74E-08 |
| Lymphatic Endothelial Cell | Acer2 | 2 | 5.67E-11 | 0.67415828 | 0.676 | 0.332 | 1.13E-07 |
| Lymphatic Endothelial Cell | Col22a1 | 2 | 6.14E-11 | 0.35219989 | 0.669 | 0.406 | 1.23E-07 |
| Lymphatic Endothelial Cell | Cdca2 | 2 | 6.14E-11 | 1.08114455 | 0.586 | 0.269 | 1.23E-07 |
| Lymphatic Endothelial Cell | Obscn1 | 2 | 8.35E-11 | 0.82095038 | 0.69 | 0.357 | 1.67E-07 |
| Lymphatic Endothelial Cell | Esr11 | 2 | 1.10E-10 | 0.31362039 | 0.634 | 0.348 | 2.20E-07 |
| Lymphatic Endothelial Cell | Adk | 2 | 1.26E-10 | 0.90204456 | 0.759 | 0.443 | 2.52E-07 |
| Lymphatic Endothelial Cell | Tagln | 2 | 1.30E-10 | 1.21303708 | 0.497 | 0.226 | 2.60E-07 |
| Lymphatic Endothelial Cell | Nhs | 2 | 1.61E-10 | 0.77110412 | 0.683 | 0.353 | 3.21E-07 |
| Lymphatic Endothelial Cell | Trib31 | 2 | 1.97E-10 | 0.52178839 | 0.628 | 0.323 | 3.93E-07 |
| Lymphatic Endothelial Cell | Khdrbs2 | 2 | 2.11E-10 | 2.20940169 | 0.676 | 0.358 | 4.22E-07 |
| Lymphatic Endothelial Cell | Slc47a1 | 2 | 3.47E-10 | 0.62330214 | 0.545 | 0.294 | 6.94E-07 |
| Lymphatic Endothelial Cell | Myo3b1 | 2 | 3.80E-10 | 0.73177396 | 0.676 | 0.36 | 7.60E-07 |
| Lymphatic Endothelial Cell | Epha7 | 2 | 5.46E-10 | 2.43274874 | 0.462 | 0.171 | 1.09E-06 |
| Lymphatic Endothelial Cell | Dpp4 | 2 | 7.04E-10 | 0.40448329 | 0.655 | 0.325 | 1.41E-06 |
| Lymphatic Endothelial Cell | Rnls | 2 | 9.06E-10 | 0.45068754 | 0.703 | 0.375 | 1.81E-06 |
| Lymphatic Endothelial Cell | Aatk | 2 | 1.05E-09 | 0.96740187 | 0.531 | 0.254 | 2.11E-06 |
| Lymphatic Endothelial Cell | Mfsd4a | 2 | 1.15E-09 | 0.74556015 | 0.779 | 0.469 | 2.30E-06 |
| Lymphatic Endothelial Cell | Rspo31 | 2 | 1.22E-09 | 1.49942349 | 0.703 | 0.425 | 2.44E-06 |
| Lymphatic Endothelial Cell | Lhfpl21 | 2 | 1.76E-09 | 0.3939489 | 0.628 | 0.367 | 3.52E-06 |
| Lymphatic Endothelial Cell | Asb15 | 2 | 2.07E-09 | 0.72410354 | 0.641 | 0.347 | 4.14E-06 |
| Lymphatic Endothelial Cell | AABR070492 | 2 | 3.06E-09 | 1.83693338 | 0.572 | 0.285 | 6.11E-06 |
| Lymphatic Endothelial Cell | Myom1 | 2 | 6.66E-09 | 0.35969659 | 0.655 | 0.37 | 1.33E-05 |
| Lymphatic Endothelial Cell | Stard10 | 2 | 8.75E-09 | 0.40632115 | 0.683 | 0.378 | 1.75E-05 |
| Lymphatic Endothelial Cell | Tmem196 | 2 | 1.53E-08 | 0.44263546 | 0.71 | 0.451 | 3.06E-05 |
| Lymphatic Endothelial Cell | Tbx1 | 2 | 3.39E-08 | 0.81864514 | 0.814 | 0.549 | 6.79E-05 |

| | | | | | | | |
|---|---|---|---|---|---|---|---|
| Lymphatic Endothelial Cell | Trpc6 | 2 | 4.44E-08 | 0.39520966 | 0.71 | 0.434 | 8.88E-05 |
| Lymphatic Endothelial Cell | Srl | 2 | 7.39E-08 | 0.27881023 | 0.538 | 0.264 | 0.00014783 |
| Lymphatic Endothelial Cell | Cd84 | 2 | 1.46E-07 | 0.57367528 | 0.421 | 0.153 | 0.00029226 |
| Lymphatic Endothelial Cell | Icam1 | 2 | 1.82E-07 | 0.76331726 | 0.724 | 0.431 | 0.00036466 |
| Lymphatic Endothelial Cell | Syt141 | 2 | 2.30E-07 | 0.63314797 | 0.655 | 0.383 | 0.00045918 |
| Lymphatic Endothelial Cell | Ripor2 | 2 | 3.62E-07 | 0.3350349 | 0.634 | 0.363 | 0.0007241 |
| Lymphatic Endothelial Cell | Ddah1 | 2 | 7.47E-07 | 0.42648893 | 0.586 | 0.335 | 0.00149452 |
| Lymphatic Endothelial Cell | Fgf12 | 2 | 8.07E-07 | 0.46168147 | 0.6 | 0.35 | 0.001614 |
| Lymphatic Endothelial Cell | Txlnb1 | 2 | 9.31E-07 | 0.46232542 | 0.648 | 0.393 | 0.001861 |
| Lymphatic Endothelial Cell | Ctsb | 2 | 2.03E-06 | 0.42582042 | 0.724 | 0.441 | 0.00406611 |
| Lymphatic Endothelial Cell | Tmcc3 | 2 | 3.77E-06 | 0.32584856 | 0.731 | 0.443 | 0.00753645 |
| Lymphatic Endothelial Cell | Acot11 | 2 | 7.59E-06 | 1.51769632 | 0.628 | 0.378 | 0.01518854 |
| Lymphatic Endothelial Cell | Apcdd1l | 3 | 7.60E-54 | 6.25568655 | 0.359 | 0.016 | 1.52E-50 |
| Lymphatic Endothelial Cell | LOC1025530 | 3 | 1.12E-45 | 5.59673999 | 0.485 | 0.026 | 2.23E-42 |
| Lymphatic Endothelial Cell | Gpc6 | 3 | 3.53E-40 | 3.19722197 | 0.951 | 0.4 | 7.07E-37 |
| Lymphatic Endothelial Cell | Zfp385d | 3 | 6.47E-40 | 3.45281503 | 0.913 | 0.247 | 1.29E-36 |
| Lymphatic Endothelial Cell | Adarb2 | 3 | 1.49E-38 | 2.89852023 | 0.437 | 0.028 | 2.98E-35 |
| Lymphatic Endothelial Cell | Alox5 | 3 | 9.98E-38 | 0.72310968 | 0.437 | 0.025 | 2.00E-34 |
| Lymphatic Endothelial Cell | Fscn2 | 3 | 2.79E-36 | 2.86345205 | 0.34 | 0.019 | 5.57E-33 |
| Lymphatic Endothelial Cell | Ddc | 3 | 7.35E-36 | 0.95827539 | 0.408 | 0.017 | 1.47E-32 |
| Lymphatic Endothelial Cell | Lypd6b | 3 | 1.19E-31 | 6.20992231 | 0.35 | 0.017 | 2.38E-28 |
| Lymphatic Endothelial Cell | Prg4 | 3 | 3.21E-31 | 5.16067605 | 0.379 | 0.048 | 6.42E-28 |
| Lymphatic Endothelial Cell | Nrg2 | 3 | 5.22E-31 | 0.50727215 | 0.34 | 0.02 | 1.04E-27 |
| Lymphatic Endothelial Cell | Bub1b | 3 | 1.80E-30 | 2.37174439 | 0.388 | 0.042 | 3.60E-27 |
| Lymphatic Endothelial Cell | Slc1a3 | 3 | 2.47E-30 | 2.41442792 | 0.282 | 0.023 | 4.93E-27 |
| Lymphatic Endothelial Cell | Ly49s6 | 3 | 1.05E-29 | 1.60847063 | 0.272 | 0.016 | 2.09E-26 |
| Lymphatic Endothelial Cell | Fgf10 | 3 | 3.29E-29 | 3.72454181 | 0.447 | 0.033 | 6.58E-26 |
| Lymphatic Endothelial Cell | Pdzrn3 | 3 | 8.53E-28 | 3.19038915 | 0.777 | 0.265 | 1.71E-24 |
| Lymphatic Endothelial Cell | Edar | 3 | 6.72E-27 | 5.49210939 | 0.272 | 0.019 | 1.34E-23 |
| Lymphatic Endothelial Cell | Cd247 | 3 | 8.78E-27 | 4.1214322 | 0.282 | 0.023 | 1.76E-23 |
| Lymphatic Endothelial Cell | Antxr1 | 3 | 1.20E-26 | 3.49252612 | 0.825 | 0.426 | 2.40E-23 |

| | | | | | | | |
|---|---|---|---|---|---|---|---|
| Lymphatic Endothelial Cell | Itgbl1 | 3 | 1.94E-26 | 2.78656832 | 0.845 | 0.412 | 3.89E-23 |
| Lymphatic Endothelial Cell | Ccdc80 | 3 | 1.14E-25 | 2.34807522 | 0.922 | 0.416 | 2.28E-22 |
| Lymphatic Endothelial Cell | Cpq | 3 | 4.91E-25 | 2.84776427 | 0.767 | 0.175 | 9.83E-22 |
| Lymphatic Endothelial Cell | Clmp | 3 | 8.71E-25 | 3.56144541 | 0.738 | 0.222 | 1.74E-21 |
| Lymphatic Endothelial Cell | Zeb2 | 3 | 1.66E-24 | 2.34386346 | 0.854 | 0.388 | 3.32E-21 |
| Lymphatic Endothelial Cell | Ank2 | 3 | 1.71E-24 | 2.30159615 | 0.854 | 0.429 | 3.43E-21 |
| Lymphatic Endothelial Cell | Col3a1 | 3 | 2.12E-24 | 2.05027232 | 0.922 | 0.6 | 4.23E-21 |
| Lymphatic Endothelial Cell | C7 | 3 | 2.14E-24 | 2.7288361 | 0.825 | 0.285 | 4.27E-21 |
| Lymphatic Endothelial Cell | C1qtnf7 | 3 | 3.17E-24 | 2.71692133 | 0.767 | 0.267 | 6.35E-21 |
| Lymphatic Endothelial Cell | Adamts2 | 3 | 1.21E-23 | 3.02755687 | 0.806 | 0.498 | 2.41E-20 |
| Lymphatic Endothelial Cell | Rgs17 | 3 | 1.40E-23 | 3.82165483 | 0.35 | 0.04 | 2.80E-20 |
| Lymphatic Endothelial Cell | Col6a2 | 3 | 3.47E-23 | 2.88045285 | 0.709 | 0.119 | 6.95E-20 |
| Lymphatic Endothelial Cell | Col8a1 | 3 | 7.55E-21 | 2.81940918 | 0.757 | 0.296 | 1.51E-17 |
| Lymphatic Endothelial Cell | Pdgfrb | 3 | 2.38E-20 | 2.96844757 | 0.689 | 0.152 | 4.75E-17 |
| Lymphatic Endothelial Cell | Egfr | 3 | 2.51E-20 | 2.93144187 | 0.709 | 0.274 | 5.02E-17 |
| Lymphatic Endothelial Cell | Grin3a | 3 | 4.69E-20 | 1.36636695 | 0.291 | 0.022 | 9.37E-17 |
| Lymphatic Endothelial Cell | Pid1 | 3 | 9.21E-20 | 2.59153665 | 0.68 | 0.177 | 1.84E-16 |
| Lymphatic Endothelial Cell | Dclk1 | 3 | 2.07E-19 | 3.20330239 | 0.748 | 0.42 | 4.15E-16 |
| Lymphatic Endothelial Cell | Col6a6 | 3 | 5.86E-19 | 3.56913074 | 0.68 | 0.223 | 1.17E-15 |
| Lymphatic Endothelial Cell | Ebf2 | 3 | 2.52E-18 | 3.27814505 | 0.757 | 0.487 | 5.05E-15 |
| Lymphatic Endothelial Cell | Arhgap24 | 3 | 3.95E-18 | 2.77919672 | 0.689 | 0.135 | 7.90E-15 |
| Lymphatic Endothelial Cell | Dlg2 | 3 | 6.20E-18 | 2.54356875 | 0.748 | 0.327 | 1.24E-14 |
| Lymphatic Endothelial Cell | Pde5a | 3 | 2.72E-17 | 2.88092491 | 0.631 | 0.302 | 5.43E-14 |
| Lymphatic Endothelial Cell | Mfap5 | 3 | 5.80E-17 | 2.48592429 | 0.602 | 0.141 | 1.16E-13 |
| Lymphatic Endothelial Cell | Galnt17 | 3 | 8.43E-17 | 3.27249605 | 0.631 | 0.119 | 1.69E-13 |
| Lymphatic Endothelial Cell | Vgll3 | 3 | 1.97E-16 | 3.50800004 | 0.437 | 0.025 | 3.93E-13 |
| Lymphatic Endothelial Cell | Col14a1 | 3 | 2.14E-16 | 3.05527059 | 0.641 | 0.251 | 4.28E-13 |
| Lymphatic Endothelial Cell | Adamts14 | 3 | 2.27E-16 | 2.44358784 | 0.417 | 0.073 | 4.55E-13 |
| Lymphatic Endothelial Cell | Pls3 | 3 | 9.33E-16 | 2.02043701 | 0.689 | 0.279 | 1.87E-12 |
| Lymphatic Endothelial Cell | Ltbp2 | 3 | 9.94E-16 | 3.08311996 | 0.65 | 0.209 | 1.99E-12 |
| Lymphatic Endothelial Cell | Cdh11 | 3 | 1.07E-15 | 1.80044153 | 0.67 | 0.256 | 2.13E-12 |

| | | | | | | | |
|---|---|---|---|---|---|---|---|
| Lymphatic Endothelial Cell | Pcdha131 | 3 | 1.23E-15 | 1.92333403 | 0.757 | 0.451 | 2.45E-12 |
| Lymphatic Endothelial Cell | Col1a1 | 3 | 5.17E-15 | 1.68905718 | 0.816 | 0.46 | 1.03E-11 |
| Lymphatic Endothelial Cell | Prrx1 | 3 | 7.41E-15 | 2.95697445 | 0.699 | 0.434 | 1.48E-11 |
| Lymphatic Endothelial Cell | Blnk | 3 | 8.56E-15 | 0.93808865 | 0.388 | 0.05 | 1.71E-11 |
| Lymphatic Endothelial Cell | Axl | 3 | 1.07E-14 | 2.92871156 | 0.699 | 0.417 | 2.14E-11 |
| Lymphatic Endothelial Cell | Slc1a7 | 3 | 1.14E-14 | 5.18766998 | 0.34 | 0.026 | 2.27E-11 |
| Lymphatic Endothelial Cell | Postn | 3 | 1.44E-13 | 2.62462935 | 0.621 | 0.211 | 2.87E-10 |
| Lymphatic Endothelial Cell | Col4a5 | 3 | 2.58E-13 | 2.53478149 | 0.67 | 0.364 | 5.15E-10 |
| Lymphatic Endothelial Cell | Cacna2d2 | 3 | 3.62E-13 | 1.97679704 | 0.437 | 0.14 | 7.25E-10 |
| Lymphatic Endothelial Cell | Fndc1 | 3 | 3.86E-13 | 1.14912381 | 0.922 | 0.662 | 7.73E-10 |
| Lymphatic Endothelial Cell | Sdk2 | 3 | 4.54E-13 | 3.91182472 | 0.505 | 0.144 | 9.08E-10 |
| Lymphatic Endothelial Cell | Bnc2 | 3 | 4.64E-13 | 3.08886556 | 0.534 | 0.13 | 9.28E-10 |
| Lymphatic Endothelial Cell | Pdgfra | 3 | 6.35E-13 | 2.58274024 | 0.592 | 0.163 | 1.27E-09 |
| Lymphatic Endothelial Cell | Cspg4 | 3 | 6.98E-13 | 2.82299502 | 0.495 | 0.127 | 1.40E-09 |
| Lymphatic Endothelial Cell | Ptprj | 3 | 1.11E-12 | 1.95356137 | 0.573 | 0.18 | 2.22E-09 |
| Lymphatic Endothelial Cell | Abca8a | 3 | 1.57E-12 | 2.98399432 | 0.621 | 0.299 | 3.13E-09 |
| Lymphatic Endothelial Cell | Dcn | 3 | 2.07E-12 | 2.01338313 | 0.66 | 0.222 | 4.13E-09 |
| Lymphatic Endothelial Cell | AABR070252 | 3 | 2.18E-12 | 1.50547962 | 0.806 | 0.502 | 4.35E-09 |
| Lymphatic Endothelial Cell | Svil | 3 | 2.37E-12 | 1.34356805 | 0.767 | 0.433 | 4.75E-09 |
| Lymphatic Endothelial Cell | Gria1 | 3 | 2.59E-12 | 2.64384395 | 0.408 | 0.028 | 5.18E-09 |
| Lymphatic Endothelial Cell | Mgp | 3 | 5.31E-12 | 1.35491243 | 0.816 | 0.488 | 1.06E-08 |
| Lymphatic Endothelial Cell | Gpm6b | 3 | 1.24E-11 | 2.89673045 | 0.612 | 0.316 | 2.47E-08 |
| Lymphatic Endothelial Cell | Adgrg2 | 3 | 1.51E-11 | 2.65366345 | 0.447 | 0.043 | 3.02E-08 |
| Lymphatic Endothelial Cell | Tmeff2 | 3 | 1.77E-11 | 3.09813477 | 0.544 | 0.141 | 3.55E-08 |
| Lymphatic Endothelial Cell | Dpt | 3 | 2.02E-11 | 1.67677077 | 0.592 | 0.158 | 4.04E-08 |
| Lymphatic Endothelial Cell | Galnt16 | 3 | 3.63E-11 | 3.02761201 | 0.515 | 0.209 | 7.26E-08 |
| Lymphatic Endothelial Cell | Irs1 | 3 | 4.83E-11 | 2.38040282 | 0.563 | 0.274 | 9.66E-08 |
| Lymphatic Endothelial Cell | Plod2 | 3 | 4.94E-11 | 1.37000557 | 0.835 | 0.561 | 9.87E-08 |
| Lymphatic Endothelial Cell | Gda | 3 | 1.09E-10 | 1.0155454 | 0.728 | 0.398 | 2.19E-07 |
| Lymphatic Endothelial Cell | Boc | 3 | 4.74E-10 | 1.87669177 | 0.592 | 0.322 | 9.48E-07 |
| Lymphatic Endothelial Cell | Il34 | 3 | 4.90E-10 | 2.14900533 | 0.427 | 0.085 | 9.80E-07 |

| | | | | | | | |
|---|---|---|---|---|---|---|---|
| Lymphatic Endothelial Cell | Fstl1 | 3 | 4.94E-10 | 1.26297856 | 0.825 | 0.51 | 9.87E-07 |
| Lymphatic Endothelial Cell | Niban1 | 3 | 8.21E-10 | 2.36478152 | 0.573 | 0.315 | 1.64E-06 |
| Lymphatic Endothelial Cell | Eda | 3 | 1.39E-09 | 2.03718319 | 0.573 | 0.271 | 2.77E-06 |
| Lymphatic Endothelial Cell | Matn2 | 3 | 1.39E-09 | 2.57393985 | 0.437 | 0.065 | 2.79E-06 |
| Lymphatic Endothelial Cell | Medag | 3 | 1.51E-09 | 1.91458856 | 0.524 | 0.183 | 3.03E-06 |
| Lymphatic Endothelial Cell | Nrxn1 | 3 | 1.97E-09 | 2.3495044 | 0.68 | 0.374 | 3.93E-06 |
| Lymphatic Endothelial Cell | Me1 | 3 | 1.99E-09 | 2.39956086 | 0.544 | 0.184 | 3.97E-06 |
| Lymphatic Endothelial Cell | Gucy1a1 | 3 | 3.66E-09 | 1.60396789 | 0.466 | 0.143 | 7.33E-06 |
| Lymphatic Endothelial Cell | Col5a3 | 3 | 6.47E-09 | 2.79480468 | 0.476 | 0.118 | 1.29E-05 |
| Lymphatic Endothelial Cell | AABR070546 | 3 | 6.92E-09 | 2.72778554 | 0.592 | 0.312 | 1.38E-05 |
| Lymphatic Endothelial Cell | Adamts17 | 3 | 1.18E-08 | 2.26740628 | 0.476 | 0.183 | 2.35E-05 |
| Lymphatic Endothelial Cell | Fat3 | 3 | 1.80E-08 | 2.10303968 | 0.505 | 0.124 | 3.60E-05 |
| Lymphatic Endothelial Cell | Adamts19 | 3 | 2.83E-08 | 2.55499946 | 0.515 | 0.122 | 5.66E-05 |
| Lymphatic Endothelial Cell | Ltbp1 | 3 | 2.99E-08 | 0.90403524 | 0.631 | 0.257 | 5.98E-05 |
| Lymphatic Endothelial Cell | Pde1a | 3 | 3.27E-08 | 2.77198815 | 0.485 | 0.095 | 6.54E-05 |
| Lymphatic Endothelial Cell | Pdlim3 | 3 | 5.84E-08 | 1.87305491 | 0.515 | 0.25 | 0.00011681 |
| Lymphatic Endothelial Cell | Sparc | 3 | 7.69E-08 | 0.71646034 | 0.835 | 0.512 | 0.00015383 |
| Lymphatic Endothelial Cell | Mob3b | 3 | 1.15E-07 | 2.66666427 | 0.466 | 0.126 | 0.00023068 |
| Lymphatic Endothelial Cell | Sdc2 | 3 | 1.19E-07 | 2.83914024 | 0.534 | 0.271 | 0.00023813 |
| Lymphatic Endothelial Cell | Gpr176 | 3 | 1.26E-07 | 3.86113488 | 0.447 | 0.195 | 0.00025224 |
| Lymphatic Endothelial Cell | AABR070581 | 3 | 1.44E-07 | 0.74129399 | 0.709 | 0.42 | 0.00028865 |
| Lymphatic Endothelial Cell | Plekha7 | 3 | 1.70E-07 | 3.64848004 | 0.437 | 0.11 | 0.00034076 |
| Lymphatic Endothelial Cell | Atp1b1 | 3 | 1.84E-07 | 1.1422186 | 0.427 | 0.073 | 0.00036779 |
| Lymphatic Endothelial Cell | Npas2 | 3 | 2.09E-07 | 1.48746867 | 0.505 | 0.253 | 0.00041704 |
| Lymphatic Endothelial Cell | Spon1 | 3 | 3.00E-07 | 3.26781643 | 0.427 | 0.093 | 0.00060028 |
| Lymphatic Endothelial Cell | Sox6 | 3 | 3.18E-07 | 1.6880036 | 0.563 | 0.191 | 0.00063617 |
| Lymphatic Endothelial Cell | Ror1 | 3 | 3.60E-07 | 3.05851246 | 0.379 | 0.088 | 0.00072031 |
| Lymphatic Endothelial Cell | Itga8 | 3 | 4.17E-07 | 0.94142748 | 0.573 | 0.259 | 0.00083403 |
| Lymphatic Endothelial Cell | Chsy3 | 3 | 4.27E-07 | 0.8839415 | 0.592 | 0.336 | 0.00085447 |
| Lymphatic Endothelial Cell | Enpp1 | 3 | 4.30E-07 | 2.23176336 | 0.495 | 0.245 | 0.00086036 |
| Lymphatic Endothelial Cell | Loxl1 | 3 | 4.50E-07 | 2.23628939 | 0.437 | 0.14 | 0.00089974 |

| | | | | | | | |
|---|---|---|---|---|---|---|---|
| Lymphatic Endothelial Cell | Il1r1 | 3 | 6.39E-07 | 0.78702133 | 0.67 | 0.364 | 0.0012788 |
| Lymphatic Endothelial Cell | Htra3 | 3 | 6.59E-07 | 2.97029193 | 0.369 | 0.056 | 0.00131829 |
| Lymphatic Endothelial Cell | Vcan | 3 | 9.15E-07 | 2.39795035 | 0.495 | 0.121 | 0.00182918 |
| Lymphatic Endothelial Cell | Ninj2 | 3 | 1.01E-06 | 2.40907221 | 0.456 | 0.192 | 0.00201692 |
| Lymphatic Endothelial Cell | Ptcd3 | 3 | 1.37E-06 | 0.42797426 | 0.65 | 0.33 | 0.00274603 |
| Lymphatic Endothelial Cell | Adamtsl2 | 3 | 1.86E-06 | 1.88008948 | 0.466 | 0.188 | 0.00371154 |
| Lymphatic Endothelial Cell | Cacna1g | 3 | 2.14E-06 | 1.80099654 | 0.485 | 0.116 | 0.00427326 |
| Lymphatic Endothelial Cell | AABR070010 | 3 | 5.16E-06 | 3.47686972 | 0.35 | 0.028 | 0.01031507 |
| Lymphatic Endothelial Cell | Foxp2 | 3 | 1.46E-05 | 1.07924306 | 0.485 | 0.183 | 0.02926931 |
| Lymphatic Endothelial Cell | Greb1l | 3 | 1.60E-05 | 3.75952354 | 0.34 | 0.064 | 0.03207028 |
| Lymphatic Endothelial Cell | Il17rd | 3 | 1.62E-05 | 2.64642549 | 0.408 | 0.14 | 0.03232348 |
| Lymphatic Endothelial Cell | Esrrg | 3 | 1.66E-05 | 2.38826541 | 0.388 | 0.109 | 0.03318587 |
| Lymphatic Endothelial Cell | Pamr1 | 3 | 1.97E-05 | 3.08045062 | 0.282 | 0.019 | 0.03936947 |
| Lymphatic Endothelial Cell | Pdlim5 | 3 | 2.42E-05 | 0.56817405 | 0.757 | 0.491 | 0.04847183 |
| Lymphatic Endothelial Cell | Scml4 | 4 | 6.99E-54 | 0.56759106 | 0.56 | 0.038 | 1.40E-50 |
| Lymphatic Endothelial Cell | Tmem255a | 4 | 2.04E-46 | 4.65042179 | 0.429 | 0.038 | 4.08E-43 |
| Lymphatic Endothelial Cell | Klrc2 | 4 | 1.77E-44 | 4.77793772 | 0.484 | 0.049 | 3.54E-41 |
| Lymphatic Endothelial Cell | Cyp4f18 | 4 | 2.80E-40 | 2.14366793 | 0.538 | 0.094 | 5.61E-37 |
| Lymphatic Endothelial Cell | Flt1 | 4 | 6.46E-40 | 3.81073792 | 0.901 | 0.186 | 1.29E-36 |
| Lymphatic Endothelial Cell | Tnnt1 | 4 | 7.59E-40 | 4.4252373 | 0.33 | 0.015 | 1.52E-36 |
| Lymphatic Endothelial Cell | AC111831.1 | 4 | 1.03E-37 | 0.72756127 | 0.505 | 0.056 | 2.06E-34 |
| Lymphatic Endothelial Cell | Cyyr1 | 4 | 5.27E-34 | 3.02862844 | 0.901 | 0.414 | 1.05E-30 |
| Lymphatic Endothelial Cell | Fgf23 | 4 | 5.62E-34 | 4.5127444 | 0.319 | 0.009 | 1.12E-30 |
| Lymphatic Endothelial Cell | Cyp26b1 | 4 | 2.36E-31 | 2.04590382 | 0.549 | 0.113 | 4.72E-28 |
| Lymphatic Endothelial Cell | Dync1i1 | 4 | 1.56E-30 | 2.61043793 | 0.549 | 0.084 | 3.12E-27 |
| Lymphatic Endothelial Cell | AABR070515 | 4 | 2.03E-29 | 1.4375676 | 0.451 | 0.035 | 4.06E-26 |
| Lymphatic Endothelial Cell | Exoc3l2 | 4 | 4.00E-29 | 4.06980767 | 0.44 | 0.058 | 8.00E-26 |
| Lymphatic Endothelial Cell | Anxa3 | 4 | 9.94E-29 | 3.73180788 | 0.791 | 0.16 | 1.99E-25 |
| Lymphatic Endothelial Cell | Cdh13 | 4 | 9.70E-28 | 2.38842161 | 0.901 | 0.417 | 1.94E-24 |
| Lymphatic Endothelial Cell | Ccdc85a | 4 | 3.93E-27 | 3.19823913 | 0.791 | 0.215 | 7.86E-24 |
| Lymphatic Endothelial Cell | Pdgfc | 4 | 6.31E-27 | 1.10716045 | 0.505 | 0.081 | 1.26E-23 |

| | | | | | | | |
|---|---|---|---|---|---|---|---|
| Lymphatic Endothelial Cell | Cyfip2 | 4 | 7.91E-26 | 1.00656012 | 0.571 | 0.129 | 1.58E-22 |
| Lymphatic Endothelial Cell | Stum | 4 | 1.30E-25 | 1.87690226 | 0.319 | 0.012 | 2.60E-22 |
| Lymphatic Endothelial Cell | Slc18a2 | 4 | 4.15E-25 | 1.91969414 | 0.308 | 0.027 | 8.31E-22 |
| Lymphatic Endothelial Cell | Kcnma1 | 4 | 4.97E-25 | 0.83671391 | 0.549 | 0.119 | 9.93E-22 |
| Lymphatic Endothelial Cell | Dach1 | 4 | 4.80E-24 | 2.41517479 | 0.879 | 0.521 | 9.59E-21 |
| Lymphatic Endothelial Cell | AABR070070 | 4 | 1.24E-23 | 1.87039025 | 0.901 | 0.384 | 2.49E-20 |
| Lymphatic Endothelial Cell | Tafa2 | 4 | 5.31E-23 | 3.91636234 | 0.374 | 0.062 | 1.06E-19 |
| Lymphatic Endothelial Cell | Meox2 | 4 | 6.63E-23 | 2.21695894 | 0.89 | 0.58 | 1.33E-19 |
| Lymphatic Endothelial Cell | Unc5c | 4 | 6.90E-23 | 4.25089826 | 0.725 | 0.212 | 1.38E-19 |
| Lymphatic Endothelial Cell | Kitlg | 4 | 9.05E-23 | 3.11900868 | 0.714 | 0.158 | 1.81E-19 |
| Lymphatic Endothelial Cell | Plac8 | 4 | 6.46E-22 | 1.33316165 | 0.286 | 0.03 | 1.29E-18 |
| Lymphatic Endothelial Cell | Myrip | 4 | 2.04E-21 | 3.2077253 | 0.659 | 0.102 | 4.08E-18 |
| Lymphatic Endothelial Cell | Emilin2 | 4 | 2.08E-21 | 1.12750732 | 0.571 | 0.149 | 4.17E-18 |
| Lymphatic Endothelial Cell | Slc35f1 | 4 | 2.88E-21 | 3.40475589 | 0.571 | 0.116 | 5.77E-18 |
| Lymphatic Endothelial Cell | Gabrb11 | 4 | 3.33E-21 | 1.69987101 | 0.549 | 0.132 | 6.66E-18 |
| Lymphatic Endothelial Cell | AABR070076 | 4 | 1.46E-20 | 3.10838813 | 0.648 | 0.102 | 2.93E-17 |
| Lymphatic Endothelial Cell | Rasa4 | 4 | 7.27E-20 | 3.59492383 | 0.758 | 0.25 | 1.45E-16 |
| Lymphatic Endothelial Cell | Ntf3 | 4 | 7.45E-20 | 3.08158391 | 0.67 | 0.105 | 1.49E-16 |
| Lymphatic Endothelial Cell | Adgrf5 | 4 | 3.49E-19 | 1.33273188 | 0.967 | 0.642 | 6.99E-16 |
| Lymphatic Endothelial Cell | Ccser1 | 4 | 6.27E-19 | 3.14173332 | 0.67 | 0.053 | 1.25E-15 |
| Lymphatic Endothelial Cell | Slc26a10 | 4 | 8.63E-19 | 3.65874539 | 0.758 | 0.303 | 1.73E-15 |
| Lymphatic Endothelial Cell | Plcb1 | 4 | 1.16E-18 | 3.50359007 | 0.582 | 0.091 | 2.32E-15 |
| Lymphatic Endothelial Cell | Tmtc2 | 4 | 1.35E-17 | 2.52968343 | 0.615 | 0.164 | 2.71E-14 |
| Lymphatic Endothelial Cell | Lims2 | 4 | 3.35E-17 | 2.09442647 | 0.692 | 0.248 | 6.70E-14 |
| Lymphatic Endothelial Cell | Akr1c15 | 4 | 5.99E-17 | 3.06111134 | 0.648 | 0.128 | 1.20E-13 |
| Lymphatic Endothelial Cell | Wt1 | 4 | 6.30E-17 | 3.49568555 | 0.692 | 0.371 | 1.26E-13 |
| Lymphatic Endothelial Cell | Myo10 | 4 | 1.36E-16 | 2.99995519 | 0.769 | 0.367 | 2.73E-13 |
| Lymphatic Endothelial Cell | Knl1 | 4 | 2.15E-16 | 0.41133966 | 0.286 | 0.03 | 4.29E-13 |
| Lymphatic Endothelial Cell | Vegfc | 4 | 4.93E-15 | 2.65499187 | 0.659 | 0.166 | 9.85E-12 |
| Lymphatic Endothelial Cell | Palmd | 4 | 5.87E-15 | 2.20861123 | 0.736 | 0.388 | 1.17E-11 |
| Lymphatic Endothelial Cell | Abcb4 | 4 | 2.94E-14 | 1.67098273 | 0.659 | 0.184 | 5.89E-11 |

| | | | | | | | |
|---|---|---|---|---|---|---|---|
| Lymphatic Endothelial Cell | Zfp366 | 4 | 3.12E-14 | 3.51267432 | 0.67 | 0.377 | 6.23E-11 |
| Lymphatic Endothelial Cell | Grik2 | 4 | 5.89E-14 | 1.57724388 | 0.681 | 0.285 | 1.18E-10 |
| Lymphatic Endothelial Cell | Pde1c | 4 | 1.18E-13 | 2.49820367 | 0.692 | 0.414 | 2.37E-10 |
| Lymphatic Endothelial Cell | Cadps2 | 4 | 1.41E-13 | 1.93091485 | 0.648 | 0.225 | 2.83E-10 |
| Lymphatic Endothelial Cell | Cmtm8 | 4 | 1.65E-13 | 3.20547019 | 0.582 | 0.157 | 3.30E-10 |
| Lymphatic Endothelial Cell | Chn1 | 4 | 2.06E-13 | 1.10923292 | 0.835 | 0.473 | 4.11E-10 |
| Lymphatic Endothelial Cell | Clic5 | 4 | 3.02E-13 | 1.54939214 | 0.626 | 0.248 | 6.05E-10 |
| Lymphatic Endothelial Cell | C1qtnf3 | 4 | 4.71E-13 | 2.01945483 | 0.473 | 0.12 | 9.43E-10 |
| Lymphatic Endothelial Cell | Il1r11 | 4 | 2.67E-12 | 1.50521007 | 0.791 | 0.353 | 5.34E-09 |
| Lymphatic Endothelial Cell | Sorbs2 | 4 | 2.68E-12 | 1.51922912 | 0.67 | 0.295 | 5.37E-09 |
| Lymphatic Endothelial Cell | Wipf3 | 4 | 3.38E-12 | 1.52833805 | 0.692 | 0.262 | 6.76E-09 |
| Lymphatic Endothelial Cell | Prom1 | 4 | 4.36E-12 | 3.02590038 | 0.538 | 0.148 | 8.72E-09 |
| Lymphatic Endothelial Cell | Vcam1 | 4 | 8.52E-12 | 2.15849661 | 0.505 | 0.073 | 1.70E-08 |
| Lymphatic Endothelial Cell | Scn9a | 4 | 1.03E-11 | 1.61015125 | 0.505 | 0.213 | 2.07E-08 |
| Lymphatic Endothelial Cell | Lifr | 4 | 1.20E-11 | 2.44012308 | 0.495 | 0.202 | 2.39E-08 |
| Lymphatic Endothelial Cell | Mgmt | 4 | 1.26E-11 | 1.36768529 | 0.791 | 0.463 | 2.53E-08 |
| Lymphatic Endothelial Cell | Pdcd1lg2 | 4 | 1.35E-11 | 4.35135703 | 0.451 | 0.091 | 2.69E-08 |
| Lymphatic Endothelial Cell | Pde5a1 | 4 | 1.68E-11 | 1.07733835 | 0.626 | 0.309 | 3.36E-08 |
| Lymphatic Endothelial Cell | Prima1 | 4 | 1.90E-11 | 1.73412153 | 0.516 | 0.058 | 3.81E-08 |
| Lymphatic Endothelial Cell | Pdgfd | 4 | 2.10E-11 | 1.82066194 | 0.615 | 0.233 | 4.19E-08 |
| Lymphatic Endothelial Cell | Syk | 4 | 2.32E-11 | 1.11574218 | 0.582 | 0.164 | 4.64E-08 |
| Lymphatic Endothelial Cell | Pcdh19 | 4 | 4.03E-11 | 1.97191626 | 0.571 | 0.17 | 8.07E-08 |
| Lymphatic Endothelial Cell | Nav3 | 4 | 5.02E-11 | 1.27553854 | 0.857 | 0.604 | 1.00E-07 |
| Lymphatic Endothelial Cell | Chn2 | 4 | 6.10E-11 | 2.37819333 | 0.637 | 0.324 | 1.22E-07 |
| Lymphatic Endothelial Cell | Ildr2 | 4 | 8.40E-11 | 0.56745641 | 0.538 | 0.043 | 1.68E-07 |
| Lymphatic Endothelial Cell | Ltbp11 | 4 | 1.09E-10 | 2.00203096 | 0.648 | 0.262 | 2.18E-07 |
| Lymphatic Endothelial Cell | Fam167a | 4 | 1.10E-10 | 1.85632646 | 0.495 | 0.154 | 2.20E-07 |
| Lymphatic Endothelial Cell | Gcnt2 | 4 | 1.96E-10 | 1.00255278 | 0.692 | 0.277 | 3.91E-07 |
| Lymphatic Endothelial Cell | Sgo21 | 4 | 2.23E-10 | 1.60212341 | 0.593 | 0.333 | 4.46E-07 |
| Lymphatic Endothelial Cell | Deptor | 4 | 2.37E-10 | 2.62928049 | 0.516 | 0.183 | 4.74E-07 |
| Lymphatic Endothelial Cell | Slc12a2 | 4 | 2.43E-10 | 1.43472818 | 0.681 | 0.341 | 4.86E-07 |

| | | | | | | | |
|---|---|---|---|---|---|---|---|
| Lymphatic Endothelial Cell | RGD1565355 | 4 | 3.32E-10 | 1.93686558 | 0.582 | 0.228 | 6.64E-07 |
| Lymphatic Endothelial Cell | Srpx | 4 | 3.36E-10 | 1.81217697 | 0.626 | 0.374 | 6.72E-07 |
| Lymphatic Endothelial Cell | Pcdh17 | 4 | 3.74E-10 | 2.9959087 | 0.538 | 0.215 | 7.48E-07 |
| Lymphatic Endothelial Cell | Cadm3 | 4 | 8.01E-10 | 0.72278678 | 0.626 | 0.291 | 1.60E-06 |
| Lymphatic Endothelial Cell | Ano4 | 4 | 9.03E-10 | 3.20233657 | 0.297 | 0.024 | 1.81E-06 |
| Lymphatic Endothelial Cell | Pld31 | 4 | 1.12E-09 | 0.59103242 | 0.681 | 0.318 | 2.25E-06 |
| Lymphatic Endothelial Cell | Necab1 | 4 | 1.29E-09 | 1.16602766 | 0.396 | 0.03 | 2.59E-06 |
| Lymphatic Endothelial Cell | Mcf2l | 4 | 1.33E-09 | 2.22407695 | 0.593 | 0.309 | 2.66E-06 |
| Lymphatic Endothelial Cell | Cdkn1a | 4 | 2.36E-09 | 0.78322006 | 0.44 | 0.067 | 4.73E-06 |
| Lymphatic Endothelial Cell | Rimbp2 | 4 | 2.36E-09 | 1.66518939 | 0.549 | 0.242 | 4.73E-06 |
| Lymphatic Endothelial Cell | Sox17 | 4 | 3.77E-09 | 2.95238909 | 0.352 | 0.062 | 7.54E-06 |
| Lymphatic Endothelial Cell | LOC1036942 | 4 | 4.01E-09 | 0.77311474 | 0.407 | 0.09 | 8.02E-06 |
| Lymphatic Endothelial Cell | Samd12 | 4 | 4.41E-09 | 2.29355633 | 0.549 | 0.292 | 8.82E-06 |
| Lymphatic Endothelial Cell | Ca4 | 4 | 9.09E-09 | 3.20574553 | 0.44 | 0.172 | 1.82E-05 |
| Lymphatic Endothelial Cell | Itgb2 | 4 | 1.06E-08 | 0.69653695 | 0.505 | 0.23 | 2.12E-05 |
| Lymphatic Endothelial Cell | Susd1 | 4 | 1.17E-08 | 1.689675 | 0.505 | 0.151 | 2.35E-05 |
| Lymphatic Endothelial Cell | Cenpe | 4 | 3.12E-08 | 0.65814632 | 0.451 | 0.18 | 6.24E-05 |
| Lymphatic Endothelial Cell | AABR070010 | 4 | 3.67E-08 | 0.25238773 | 0.396 | 0.143 | 7.34E-05 |
| Lymphatic Endothelial Cell | Nav2 | 4 | 3.78E-08 | 1.18082688 | 0.527 | 0.274 | 7.56E-05 |
| Lymphatic Endothelial Cell | Slc37a2 | 4 | 3.92E-08 | 0.82563884 | 0.385 | 0.053 | 7.85E-05 |
| Lymphatic Endothelial Cell | Stxbp6 | 4 | 4.62E-08 | 0.64732046 | 0.813 | 0.457 | 9.24E-05 |
| Lymphatic Endothelial Cell | Tmem26 | 4 | 4.78E-08 | 1.3173503 | 0.484 | 0.212 | 9.56E-05 |
| Lymphatic Endothelial Cell | Spp1 | 4 | 5.10E-08 | 0.35601285 | 0.56 | 0.172 | 0.0001019 |
| Lymphatic Endothelial Cell | Sorcs2 | 4 | 5.64E-08 | 0.78218854 | 0.626 | 0.283 | 0.0001128 |
| Lymphatic Endothelial Cell | Cxcl12 | 4 | 6.32E-08 | 2.49107413 | 0.473 | 0.131 | 0.00012639 |
| Lymphatic Endothelial Cell | Kcnab1 | 4 | 9.94E-08 | 1.25521609 | 0.593 | 0.304 | 0.00019871 |
| Lymphatic Endothelial Cell | Efnb2 | 4 | 1.07E-07 | 0.78821745 | 0.769 | 0.438 | 0.00021422 |
| Lymphatic Endothelial Cell | LOC687780 | 4 | 1.16E-07 | 0.41949759 | 0.67 | 0.303 | 0.00023113 |
| Lymphatic Endothelial Cell | AABR070042 | 4 | 1.28E-07 | 1.07880958 | 0.56 | 0.224 | 0.00025594 |
| Lymphatic Endothelial Cell | Herc6 | 4 | 1.31E-07 | 0.87627337 | 0.626 | 0.323 | 0.00026202 |
| Lymphatic Endothelial Cell | Slc4a3 | 4 | 1.45E-07 | 1.02246367 | 0.418 | 0.076 | 0.00028937 |

| | | | | | | | |
|---|---|---|---|---|---|---|---|
| Lymphatic Endothelial Cell | Mcam | 4 | 3.36E-07 | 1.15627178 | 0.396 | 0.139 | 0.00067206 |
| Lymphatic Endothelial Cell | Sybu | 4 | 3.67E-07 | 4.03159847 | 0.308 | 0.026 | 0.00073485 |
| Lymphatic Endothelial Cell | Arhgap44 | 4 | 3.99E-07 | 1.34843602 | 0.462 | 0.202 | 0.00079805 |
| Lymphatic Endothelial Cell | Ptprn2 | 4 | 5.04E-07 | 0.84419572 | 0.571 | 0.283 | 0.00100757 |
| Lymphatic Endothelial Cell | Ptprj1 | 4 | 6.08E-07 | 0.86951807 | 0.516 | 0.195 | 0.00121573 |
| Lymphatic Endothelial Cell | Tmem182 | 4 | 6.79E-07 | 1.71187044 | 0.396 | 0.078 | 0.00135737 |
| Lymphatic Endothelial Cell | Ccnb1 | 4 | 1.09E-06 | 0.77914381 | 0.516 | 0.257 | 0.00218491 |
| Lymphatic Endothelial Cell | Fam78b | 4 | 1.25E-06 | 1.3796341 | 0.681 | 0.417 | 0.00250586 |
| Lymphatic Endothelial Cell | Il18r1 | 4 | 1.43E-06 | 1.31531988 | 0.527 | 0.26 | 0.00286327 |
| Lymphatic Endothelial Cell | Epha4 | 4 | 1.59E-06 | 2.76273563 | 0.385 | 0.09 | 0.00317915 |
| Lymphatic Endothelial Cell | Dipk2b | 4 | 1.79E-06 | 0.88939479 | 0.802 | 0.549 | 0.00358722 |
| Lymphatic Endothelial Cell | Ubash3b | 4 | 1.97E-06 | 0.53288172 | 0.78 | 0.514 | 0.00394615 |
| Lymphatic Endothelial Cell | Ppp1r16b | 4 | 2.43E-06 | 0.50593379 | 0.813 | 0.554 | 0.00485045 |
| Lymphatic Endothelial Cell | Cox6a1 | 4 | 3.39E-06 | 0.71538214 | 0.516 | 0.196 | 0.00677723 |
| Lymphatic Endothelial Cell | Sorbs1 | 4 | 3.47E-06 | 0.48174171 | 0.912 | 0.662 | 0.0069485 |
| Lymphatic Endothelial Cell | Sulf11 | 4 | 1.37E-05 | 0.25766384 | 0.659 | 0.399 | 0.0274381 |
| Lymphatic Endothelial Cell | Hjurp | 4 | 2.33E-05 | 1.45924605 | 0.297 | 0.032 | 0.04664441 |
| Lymphatic Endothelial Cell | Thbs4 | 5 | 2.24E-08 | 0.58498667 | 0.079 | 0.344 | 4.48E-05 |
| Lymphatic Endothelial Cell | Syt142 | 5 | 1.29E-06 | 0.2655366 | 0.158 | 0.467 | 0.00258819 |
| Lymphatic Endothelial Cell | Ctsk | 5 | 5.41E-06 | 0.86380527 | 0.105 | 0.366 | 0.0108188 |
| Lymphatic Endothelial Cell | Asb181 | 5 | 6.18E-06 | 0.71385973 | 0.158 | 0.409 | 0.01235816 |
| Lymphatic Endothelial Cell | Trim55 | 5 | 9.70E-06 | 0.57349151 | 0.132 | 0.406 | 0.01940659 |
| Lymphatic Endothelial Cell | AABR070015 | 6 | 3.01E-46 | 6.92011916 | 0.632 | 0.014 | 6.02E-43 |
| Lymphatic Endothelial Cell | Lilrb3 | 6 | 6.49E-42 | 6.26039986 | 0.632 | 0.025 | 1.30E-38 |
| Lymphatic Endothelial Cell | Fcrl5 | 6 | 1.89E-29 | 5.95942761 | 0.526 | 0.025 | 3.79E-26 |
| Lymphatic Endothelial Cell | Brinp2 | 6 | 5.02E-28 | 5.599655 | 0.526 | 0.016 | 1.00E-24 |
| Lymphatic Endothelial Cell | Stap1 | 6 | 7.52E-28 | 5.5995314 | 0.316 | 0.023 | 1.50E-24 |
| Lymphatic Endothelial Cell | RGD1305807 | 6 | 2.22E-25 | 4.00555037 | 0.474 | 0.023 | 4.44E-22 |
| Lymphatic Endothelial Cell | Matk | 6 | 5.05E-25 | 5.4276887 | 0.474 | 0.018 | 1.01E-21 |
| Lymphatic Endothelial Cell | Cd163 | 6 | 1.42E-18 | 5.25288262 | 0.632 | 0.06 | 2.84E-15 |
| Lymphatic Endothelial Cell | Tm4sf19 | 6 | 1.43E-17 | 3.94884507 | 0.579 | 0.04 | 2.87E-14 |

| | | | | | | | |
|---|---|---|---|---|---|---|---|
| Lymphatic Endothelial Cell | Kcnip1 | 6 | 2.65E-17 | 2.96614656 | 0.316 | 0.037 | 5.29E-14 |
| Lymphatic Endothelial Cell | Txk | 6 | 3.06E-16 | 2.57229049 | 0.474 | 0.037 | 6.12E-13 |
| Lymphatic Endothelial Cell | Ighm | 6 | 8.70E-16 | 0.35956552 | 0.474 | 0.011 | 1.74E-12 |
| Lymphatic Endothelial Cell | Anpep | 6 | 1.72E-15 | 3.4063973 | 0.632 | 0.044 | 3.44E-12 |
| Lymphatic Endothelial Cell | P2ry14 | 6 | 2.86E-15 | 4.50581241 | 0.526 | 0.051 | 5.72E-12 |
| Lymphatic Endothelial Cell | Ltc4s | 6 | 3.13E-15 | 2.873187 | 0.474 | 0.043 | 6.25E-12 |
| Lymphatic Endothelial Cell | Bank1 | 6 | 2.75E-14 | 5.40361679 | 0.526 | 0.022 | 5.50E-11 |
| Lymphatic Endothelial Cell | Cacna1d | 6 | 1.07E-12 | 2.63876375 | 0.368 | 0.036 | 2.13E-09 |
| Lymphatic Endothelial Cell | Plxnc1 | 6 | 1.20E-11 | 5.0517889 | 0.368 | 0.026 | 2.40E-08 |
| Lymphatic Endothelial Cell | Slc1a1 | 6 | 5.19E-11 | 2.85765623 | 0.526 | 0.036 | 1.04E-07 |
| Lymphatic Endothelial Cell | Blnk1 | 6 | 1.33E-10 | 3.95111031 | 0.737 | 0.08 | 2.67E-07 |
| Lymphatic Endothelial Cell | Arhgap15 | 6 | 1.34E-10 | 3.28069393 | 0.947 | 0.414 | 2.68E-07 |
| Lymphatic Endothelial Cell | Fyb1 | 6 | 2.97E-10 | 4.09238247 | 0.895 | 0.481 | 5.94E-07 |
| Lymphatic Endothelial Cell | Ptprc | 6 | 5.27E-10 | 3.87918483 | 0.842 | 0.17 | 1.05E-06 |
| Lymphatic Endothelial Cell | Ctss | 6 | 8.86E-10 | 5.1761078 | 0.842 | 0.202 | 1.77E-06 |
| Lymphatic Endothelial Cell | Depdc1 | 6 | 1.44E-09 | 2.03009667 | 0.316 | 0.034 | 2.87E-06 |
| Lymphatic Endothelial Cell | Zeb21 | 6 | 4.88E-09 | 2.65540148 | 0.947 | 0.439 | 9.77E-06 |
| Lymphatic Endothelial Cell | Tlr7 | 6 | 5.89E-09 | 5.09450396 | 0.579 | 0.026 | 1.18E-05 |
| Lymphatic Endothelial Cell | Hk3 | 6 | 7.38E-09 | 5.39192519 | 0.684 | 0.235 | 1.48E-05 |
| Lymphatic Endothelial Cell | Fgd2 | 6 | 8.57E-09 | 4.83922847 | 0.684 | 0.198 | 1.71E-05 |
| Lymphatic Endothelial Cell | Emilin21 | 6 | 8.65E-09 | 4.05845544 | 0.632 | 0.189 | 1.73E-05 |
| Lymphatic Endothelial Cell | Plek | 6 | 9.88E-09 | 5.87017619 | 0.474 | 0.014 | 1.98E-05 |
| Lymphatic Endothelial Cell | Nrg21 | 6 | 1.09E-08 | 3.4833534 | 0.368 | 0.056 | 2.19E-05 |
| Lymphatic Endothelial Cell | Mitf | 6 | 1.87E-08 | 2.82083909 | 0.895 | 0.333 | 3.73E-05 |
| Lymphatic Endothelial Cell | Camk1d | 6 | 1.97E-08 | 2.89039592 | 0.895 | 0.331 | 3.94E-05 |
| Lymphatic Endothelial Cell | Ptprj2 | 6 | 2.83E-08 | 2.77057 | 0.895 | 0.217 | 5.67E-05 |
| Lymphatic Endothelial Cell | Cdh7 | 6 | 3.05E-08 | 3.78850835 | 0.526 | 0.022 | 6.10E-05 |
| Lymphatic Endothelial Cell | Aldh1a2 | 6 | 7.56E-08 | 1.73851595 | 0.474 | 0.06 | 0.0001511 |
| Lymphatic Endothelial Cell | Arhgap241 | 6 | 8.47E-08 | 2.97801651 | 0.789 | 0.196 | 0.00016931 |
| Lymphatic Endothelial Cell | Cyfip21 | 6 | 9.57E-08 | 3.57693355 | 0.579 | 0.173 | 0.0001914 |
| Lymphatic Endothelial Cell | Tafa21 | 6 | 9.89E-08 | 1.37940504 | 0.474 | 0.091 | 0.00019772 |

| | | | | | | | |
|---|---|---|---|---|---|---|---|
| Lymphatic Endothelial Cell | Cd300lb | 6 | 1.00E-07 | 4.91126556 | 0.632 | 0.193 | 0.00020061 |
| Lymphatic Endothelial Cell | Dhrs7c | 6 | 1.14E-07 | 2.65646748 | 0.368 | 0.077 | 0.00022796 |
| Lymphatic Endothelial Cell | Itgb21 | 6 | 1.34E-07 | 5.76807547 | 0.632 | 0.254 | 0.00026874 |
| Lymphatic Endothelial Cell | Shtn1 | 6 | 1.43E-07 | 5.29179273 | 0.684 | 0.273 | 0.00028552 |
| Lymphatic Endothelial Cell | Iqgap2 | 6 | 1.47E-07 | 3.81147665 | 0.684 | 0.063 | 0.00029326 |
| Lymphatic Endothelial Cell | Otulinl | 6 | 2.06E-07 | 3.79121994 | 0.789 | 0.248 | 0.00041204 |
| Lymphatic Endothelial Cell | Cfh | 6 | 2.36E-07 | 3.61043816 | 0.789 | 0.191 | 0.0004723 |
| Lymphatic Endothelial Cell | Lilrb4 | 6 | 4.19E-07 | 5.23251369 | 0.737 | 0.241 | 0.00083828 |
| Lymphatic Endothelial Cell | Knl11 | 6 | 5.60E-07 | 3.65220771 | 0.316 | 0.055 | 0.00112014 |
| Lymphatic Endothelial Cell | Nlrp3 | 6 | 5.74E-07 | 4.65545181 | 0.684 | 0.313 | 0.0011484 |
| Lymphatic Endothelial Cell | Pik3r5 | 6 | 5.89E-07 | 4.75815567 | 0.684 | 0.24 | 0.00117836 |
| Lymphatic Endothelial Cell | Prkcb | 6 | 7.26E-07 | 3.6759305 | 0.789 | 0.344 | 0.00145122 |
| Lymphatic Endothelial Cell | Tshr | 6 | 7.39E-07 | 3.53262118 | 0.421 | 0.044 | 0.00147816 |
| Lymphatic Endothelial Cell | Rgs171 | 6 | 7.73E-07 | 2.09551449 | 0.474 | 0.073 | 0.00154604 |
| Lymphatic Endothelial Cell | Alcam | 6 | 9.05E-07 | 3.06828608 | 0.842 | 0.422 | 0.00180976 |
| Lymphatic Endothelial Cell | LOC690045 | 6 | 9.63E-07 | 5.12441378 | 0.737 | 0.241 | 0.00192544 |
| Lymphatic Endothelial Cell | Atp1b11 | 6 | 9.92E-07 | 2.41307257 | 0.684 | 0.107 | 0.00198316 |
| Lymphatic Endothelial Cell | Runx1 | 6 | 1.13E-06 | 3.29929506 | 0.789 | 0.24 | 0.00226335 |
| Lymphatic Endothelial Cell | Inpp5d | 6 | 1.32E-06 | 2.66732017 | 0.789 | 0.261 | 0.00264927 |
| Lymphatic Endothelial Cell | Treml4 | 6 | 2.17E-06 | 4.93226386 | 0.421 | 0.025 | 0.00434028 |
| Lymphatic Endothelial Cell | Pik3cd | 6 | 2.73E-06 | 3.02511869 | 0.737 | 0.232 | 0.00546455 |
| Lymphatic Endothelial Cell | Cass4 | 6 | 5.02E-06 | 4.92190453 | 0.579 | 0.232 | 0.01003063 |
| Lymphatic Endothelial Cell | Csf1r | 6 | 5.41E-06 | 4.36931378 | 0.684 | 0.199 | 0.01082428 |
| Lymphatic Endothelial Cell | Slc37a21 | 6 | 5.54E-06 | 4.86413886 | 0.526 | 0.082 | 0.01108732 |
| Lymphatic Endothelial Cell | Exoc3l21 | 6 | 6.01E-06 | 1.45155114 | 0.474 | 0.095 | 0.01201478 |
| Lymphatic Endothelial Cell | Atp8b4 | 6 | 6.54E-06 | 5.04263233 | 0.579 | 0.292 | 0.01307714 |
| Lymphatic Endothelial Cell | Il6r | 6 | 1.29E-05 | 2.86531701 | 0.737 | 0.152 | 0.02586348 |
| Lymphatic Endothelial Cell | Cd841 | 6 | 1.36E-05 | 4.49033682 | 0.632 | 0.193 | 0.02716214 |
| Lymphatic Endothelial Cell | Cd300a | 6 | 2.06E-05 | 4.931319 | 0.684 | 0.14 | 0.04115036 |
| Lymphatic Endothelial Cell | Tbxas1 | 6 | 2.40E-05 | 4.41048174 | 0.684 | 0.067 | 0.04802236 |
