## Supplemental Table 13 for "Uncovering the Regional and Cell Specific Bioactivity of Injectable Extracellular Matrix Biomaterials in Myocardial Infarction through Spatial and Single Nucleus Transcriptomics"

**Supplementary Table 13. Subcluster Marker Genes in Chronic MI Model**

| Cell Type | Gene | cluster | p_val | avg_log2FC | pct.1 | pct.2 | p_val_adj |
|---|---|---|---|---|---|---|---|
| Macrophage | AABR07072096.1 | 1 | 2.13E-13 | 6.11293331 | 0.667 | 0.044 | 4.25E-10 |
| Macrophage | AABR07043667.1 | 1 | 2.77E-12 | 4.11768706 | 0.625 | 0.044 | 5.54E-09 |
| Macrophage | Msln | 1 | 2.77E-12 | 4.4609343 | 0.625 | 0.044 | 5.54E-09 |
| Macrophage | AABR07035722.1 | 1 | 3.01E-12 | 3.65766859 | 0.646 | 0.044 | 6.03E-09 |
| Macrophage | Cfh | 1 | 6.02E-12 | 2.8984823 | 0.958 | 0.618 | 1.20E-08 |
| Macrophage | Phf24 | 1 | 1.74E-11 | 4.70856287 | 0.625 | 0.029 | 3.48E-08 |
| Macrophage | Tceal7 | 1 | 1.74E-11 | 4.56556843 | 0.625 | 0.074 | 3.48E-08 |
| Macrophage | Brinp2 | 1 | 3.82E-11 | 5.05309844 | 0.646 | 0.044 | 7.63E-08 |
| Macrophage | Rasgrf1 | 1 | 2.21E-10 | 5.02117654 | 0.625 | 0.044 | 4.41E-07 |
| Macrophage | AABR07006311.1 | 1 | 4.02E-10 | 2.6476543 | 0.625 | 0.088 | 8.03E-07 |
| Macrophage | C5ar1 | 1 | 7.04E-10 | 2.66848546 | 0.708 | 0.059 | 1.41E-06 |
| Macrophage | AABR07001573.2 | 1 | 7.53E-10 | 1.38623648 | 0.604 | 0.074 | 1.51E-06 |

| | | | | | | | |
|---|---|---|---|---|---|---|---|
| Macrophage | Nrg4 | 1 | 1.54E-09 | 5.03593693 | 0.625 | 0.162 | 3.08E-06 |
| Macrophage | Rspo3 | 1 | 2.28E-09 | 1.16576959 | 0.625 | 0.029 | 4.55E-06 |
| Macrophage | Adgrb3 | 1 | 4.19E-09 | 4.67321329 | 0.646 | 0.221 | 8.39E-06 |
| Macrophage | Nav3 | 0 | 6.30E-09 | 4.84384631 | 0.724 | 0.034 | 1.26E-05 |
| Macrophage | Irf4 | 1 | 8.97E-09 | 2.91760626 | 0.625 | 0.176 | 1.79E-05 |
| Macrophage | Setmar | 1 | 9.36E-09 | 4.39302173 | 0.625 | 0.103 | 1.87E-05 |
| Macrophage | Sfrp4 | 1 | 9.36E-09 | 4.5149885 | 0.625 | 0.147 | 1.87E-05 |
| Macrophage | Rgs3 | 0 | 1.00E-08 | 2.4276757 | 0.81 | 0.448 | 2.00E-05 |
| Macrophage | Ly49s6 | 0 | 1.94E-08 | 5.15595974 | 0.31 | 0.052 | 3.87E-05 |
| Macrophage | Dapk2 | 0 | 3.44E-08 | 4.45328781 | 0.707 | 0.034 | 6.87E-05 |
| Macrophage | Susd4 | 1 | 3.69E-08 | 4.08601877 | 0.625 | 0.221 | 7.38E-05 |
| Macrophage | Stab1 | 1 | 5.12E-08 | 3.85527935 | 0.792 | 0.324 | 0.0001024 |
| Macrophage | Stum | 1 | 6.08E-08 | 0.9396398 | 0.479 | 0.015 | 0.00012167 |
| Macrophage | Pex5l | 1 | 6.13E-08 | 1.33102571 | 0.542 | 0.029 | 0.00012265 |

| | | | | | | | |
|---|---|---|---|---|---|---|---|
| Macrophage | Slc16a10 | 1 | 8.60E-08 | 4.51389165 | 0.75 | 0.132 | 0.00017197 |
| Macrophage | Asgr2 | 1 | 8.69E-08 | 3.71351086 | 0.771 | 0.324 | 0.00017388 |
| Macrophage | RGD1560281 | 1 | 8.76E-08 | 3.94937264 | 0.646 | 0.029 | 0.00017523 |
| Macrophage | Inpp5d | 1 | 9.79E-08 | 2.4471584 | 0.875 | 0.456 | 0.00019572 |
| Macrophage | Il7 | 1 | 1.14E-07 | 3.78441879 | 0.625 | 0.221 | 0.00022799 |
| Macrophage | Selp | 1 | 1.24E-07 | 3.9379021 | 0.625 | 0.25 | 0.00024828 |
| Macrophage | AABR07029470.1 | 1 | 1.45E-07 | 0.92525922 | 0.542 | 0.074 | 0.00028973 |
| Macrophage | AABR07059258.1 | 0 | 1.52E-07 | 2.89352991 | 0.672 | 0.069 | 0.00030395 |
| Macrophage | C1qb | 1 | 1.67E-07 | 1.35636405 | 0.542 | 0.059 | 0.00033302 |
| Macrophage | Pipox | 1 | 1.81E-07 | 2.90362559 | 0.625 | 0.191 | 0.00036266 |
| Macrophage | Lrrc31 | 1 | 1.92E-07 | 3.22755481 | 0.625 | 0.147 | 0.00038425 |
| Macrophage | P2ry6 | 1 | 2.47E-07 | 3.57520187 | 0.708 | 0.162 | 0.00049448 |
| Macrophage | Myo10 | 0 | 2.78E-07 | 4.11647294 | 0.638 | 0.086 | 0.00055569 |
| Macrophage | Clec4e | 1 | 2.83E-07 | 4.71962226 | 0.625 | 0.147 | 0.00056502 |

| | | | | | | | |
|---|---|---|---|---|---|---|---|
| Macrophage | Etl4 | 0 | 3.08E-07 | 1.99826403 | 0.672 | 0.155 | 0.00061699 |
| Macrophage | Ccdc88b | 1 | 3.39E-07 | 1.97120984 | 0.604 | 0.029 | 0.00067771 |
| Macrophage | Cnksr2 | 1 | 4.31E-07 | 4.79174086 | 0.646 | 0.191 | 0.00086276 |
| Macrophage | Actg2 | 1 | 4.46E-07 | 4.53716217 | 0.625 | 0.221 | 0.00089144 |
| Macrophage | Cav1 | 0 | 4.59E-07 | 2.04905304 | 0.724 | 0.069 | 0.00091801 |
| Macrophage | Slc7a7 | 1 | 4.89E-07 | 3.37061598 | 0.729 | 0.382 | 0.00097798 |
| Macrophage | AABR07049292.1 | 1 | 4.90E-07 | 3.27952577 | 0.625 | 0.118 | 0.00098069 |
| Macrophage | Rnf150 | 1 | 5.35E-07 | 3.09375825 | 0.812 | 0.25 | 0.00107053 |
| Macrophage | Matk | 1 | 5.94E-07 | 1.52053832 | 0.667 | 0.118 | 0.00118891 |
| Macrophage | Col19a1 | 1 | 7.93E-07 | 2.66672814 | 0.562 | 0.147 | 0.00158698 |
| Macrophage | Plbd1 | 1 | 9.06E-07 | 3.29692412 | 0.708 | 0.221 | 0.00181186 |
| Macrophage | Cadm3 | 0 | 9.41E-07 | 2.3927922 | 0.638 | 0.052 | 0.00188218 |
| Macrophage | Arl15 | 0 | 9.93E-07 | 1.21886789 | 0.983 | 0.724 | 0.00198551 |
| Macrophage | Anks1b | 1 | 9.93E-07 | 4.37333791 | 0.625 | 0.147 | 0.00198575 |

| | | | | | | | |
|---|---|---|---|---|---|---|---|
| Macrophage | Cdkn1a | 1 | 9.93E-07 | 5.00349673 | 0.625 | 0.206 | 0.00198575 |
| Macrophage | Adamtsl3 | 0 | 1.03E-06 | 1.65263094 | 0.81 | 0.552 | 0.00206691 |
| Macrophage | Chrdl1 | 1 | 1.04E-06 | 5.63959185 | 0.583 | 0.059 | 0.00208748 |
| Macrophage | AABR07050449.1 | 0 | 1.42E-06 | 1.29143914 | 0.466 | 0.121 | 0.00283184 |
| Macrophage | Col22a1 | 0 | 1.46E-06 | 2.29653697 | 0.69 | 0.069 | 0.00291846 |
| Macrophage | Csf1r | 1 | 1.62E-06 | 2.57893653 | 0.771 | 0.338 | 0.00323546 |
| Macrophage | Gabrb2 | 1 | 1.69E-06 | 0.43692104 | 0.417 | 0.059 | 0.00337866 |
| Macrophage | Podxl | 0 | 1.83E-06 | 5.62161491 | 0.655 | 0.345 | 0.0036623 |
| Macrophage | Fam151a | 0 | 1.91E-06 | 0.44151737 | 0.362 | 0 | 0.00382213 |
| Macrophage | LOC100910636 | 1 | 1.91E-06 | 2.48974286 | 0.646 | 0.294 | 0.00382299 |
| Macrophage | RT1-Db1 | 1 | 2.01E-06 | 2.42003817 | 0.688 | 0.235 | 0.00402645 |
| Macrophage | Cyfip2 | 0 | 2.12E-06 | 7.26184897 | 0.603 | 0.034 | 0.00424463 |
| Macrophage | Srgap3 | 0 | 2.29E-06 | 4.19062214 | 0.448 | 0.069 | 0.00458656 |
| Macrophage | Mertk | 1 | 2.70E-06 | 3.23724301 | 0.729 | 0.368 | 0.00539927 |

| | | | | | | | |
|---|---|---|---|---|---|---|---|
| Macrophage | Cdh13 | 0 | 2.90E-06 | 2.28297476 | 0.793 | 0.259 | 0.00579103 |
| Macrophage | Rcan1 | 1 | 3.28E-06 | 4.17702435 | 0.646 | 0.324 | 0.00656015 |
| Macrophage | Spns2 | 0 | 3.59E-06 | 3.23437866 | 0.569 | 0.052 | 0.00717909 |
| Macrophage | Itgb8 | 0 | 3.99E-06 | 2.44682034 | 0.603 | 0.052 | 0.00798037 |
| Macrophage | AABR07003235.1 | 1 | 4.08E-06 | 1.60866305 | 0.625 | 0.265 | 0.0081556 |
| Macrophage | Hk2 | 1 | 4.23E-06 | 2.45539371 | 0.771 | 0.412 | 0.00846445 |
| Macrophage | Camk1d | 1 | 4.28E-06 | 1.83365321 | 0.812 | 0.471 | 0.00855772 |
| Macrophage | Kcnk13 | 1 | 4.48E-06 | 3.27408491 | 0.75 | 0.382 | 0.00895533 |
| Macrophage | Robo1 | 0 | 4.53E-06 | 2.31299047 | 0.534 | 0.086 | 0.0090552 |
| Macrophage | LOC690045 | 1 | 4.81E-06 | 1.72353855 | 0.833 | 0.412 | 0.00961955 |
| Macrophage | Rgs6 | 1 | 4.91E-06 | 1.464213 | 0.625 | 0.294 | 0.00982777 |
| Macrophage | Apba2 | 0 | 5.24E-06 | 2.44734578 | 0.052 | 0.448 | 0.01047193 |
| Macrophage | Gpr176 | 1 | 5.47E-06 | 3.83156919 | 0.646 | 0.368 | 0.0109496 |
| Macrophage | Sox5 | 0 | 5.62E-06 | 2.62936618 | 0.707 | 0.103 | 0.01124362 |

| | | | | | | | |
|---|---|---|---|---|---|---|---|
| Macrophage | Anp32b | 0 | 6.18E-06 | 1.30888647 | 0.569 | 0.069 | 0.01236249 |
| Macrophage | Ptgs1 | 1 | 6.82E-06 | 2.19475252 | 0.646 | 0.368 | 0.01364965 |
| Macrophage | Alox5 | 1 | 7.29E-06 | 1.84207619 | 0.708 | 0.147 | 0.01457854 |
| Macrophage | Timp3 | 0 | 7.54E-06 | 1.91849249 | 0.621 | 0.138 | 0.01507586 |
| Macrophage | Mctp1 | 1 | 7.62E-06 | 2.22181096 | 0.812 | 0.441 | 0.01523841 |
| Macrophage | Ngf | 0 | 7.84E-06 | 2.79123709 | 0.707 | 0.138 | 0.01568653 |
| Macrophage | Ccr1 | 1 | 7.91E-06 | 0.62396604 | 0.646 | 0.294 | 0.01582034 |
| Macrophage | Taldo1 | 0 | 8.12E-06 | 2.7577975 | 0.552 | 0.069 | 0.0162393 |
| Macrophage | Col8a1 | 0 | 8.17E-06 | 0.93015474 | 0.776 | 0.241 | 0.01634225 |
| Macrophage | Plaur | 0 | 8.17E-06 | 0.6499571 | 0.431 | 0.086 | 0.01634356 |
| Macrophage | Dach1 | 0 | 8.74E-06 | 2.48938209 | 0.741 | 0.31 | 0.01748663 |
| Macrophage | LOC103693323 | 1 | 8.85E-06 | 3.6850614 | 0.625 | 0.324 | 0.01770342 |
| Macrophage | Ninj2 | 0 | 9.20E-06 | 1.38959982 | 0.603 | 0.103 | 0.01840799 |
| Macrophage | Asb14 | 0 | 9.48E-06 | 0.82046932 | 0.448 | 0.103 | 0.01895941 |

| | | | | | | | |
|---|---|---|---|---|---|---|---|
| Macrophage | Slco2a1 | 1 | 9.89E-06 | 2.46366675 | 0.625 | 0.029 | 0.01978538 |
| Macrophage | Fhad1 | 1 | 1.02E-05 | 0.35784954 | 0.625 | 0.147 | 0.02036251 |
| Macrophage | Kitlg | 0 | 1.02E-05 | 1.62763115 | 0.552 | 0.121 | 0.0204796 |
| Macrophage | Klrd1 | 0 | 1.07E-05 | 5.78346954 | 0.379 | 0.069 | 0.02135259 |
| Macrophage | Smad6 | 0 | 1.11E-05 | 0.92332 | 0.672 | 0.121 | 0.0221137 |
| Macrophage | Cxcl12 | 0 | 1.13E-05 | 2.06109994 | 0.672 | 0.138 | 0.0226479 |
| Macrophage | Zfp366 | 0 | 1.16E-05 | 0.97174357 | 0.569 | 0.103 | 0.0232734 |
| Macrophage | Npr3 | 1 | 1.20E-05 | 4.40809803 | 0.646 | 0.279 | 0.0239596 |
| Macrophage | Pdpn | 1 | 1.20E-05 | 1.87041339 | 0.625 | 0.324 | 0.0240033 |
| Macrophage | Mrc1 | 1 | 1.24E-05 | 2.21090226 | 0.708 | 0.353 | 0.02472243 |
| Macrophage | Fbn1 | 0 | 1.26E-05 | 2.37217972 | 0.793 | 0.259 | 0.0251826 |
| Macrophage | AABR07007032.1 | 0 | 1.27E-05 | 1.48780198 | 0.845 | 0.397 | 0.02531457 |
| Macrophage | Ackr3 | 0 | 1.28E-05 | 3.98729915 | 0.586 | 0.052 | 0.02561814 |
| Macrophage | Sctr | 1 | 1.32E-05 | 2.98761722 | 0.625 | 0.25 | 0.02635285 |

| | | | | | | | |
|---|---|---|---|---|---|---|---|
| Macrophage | Clmp | 0 | 1.43E-05 | 1.53498876 | 0.655 | 0.138 | 0.02865941 |
| Macrophage | Gfod1 | 0 | 1.43E-05 | 0.5298178 | 0.5 | 0.103 | 0.02866266 |
| Macrophage | Dtx1 | 1 | 1.46E-05 | 4.7772318 | 0.542 | 0.088 | 0.02927946 |
| Macrophage | Scn7a | 0 | 1.47E-05 | 2.33387776 | 0.621 | 0.069 | 0.02940833 |
| Macrophage | Atp8b1 | 0 | 1.47E-05 | 2.49785901 | 0.672 | 0.121 | 0.0294109 |
| Macrophage | Ncr1 | 0 | 1.61E-05 | 4.64169287 | 0.362 | 0.052 | 0.03216489 |
| Macrophage | Adamts5 | 0 | 1.68E-05 | 1.20521906 | 0.69 | 0.172 | 0.03357371 |
| Macrophage | Myof | 1 | 1.69E-05 | 1.89968569 | 0.833 | 0.412 | 0.0337952 |
| Macrophage | Tenm3 | 0 | 1.70E-05 | 2.29731195 | 0.362 | 0.052 | 0.0340861 |
| Macrophage | Map1b | 0 | 1.71E-05 | 1.45855541 | 0.776 | 0.259 | 0.03418776 |
| Macrophage | Fabp4 | 0 | 1.72E-05 | 2.75556171 | 0.586 | 0.121 | 0.0343067 |
| Macrophage | AABR07007026.1 | 1 | 1.72E-05 | 2.83664262 | 0.625 | 0.162 | 0.03430721 |
| Macrophage | Flt1 | 0 | 1.85E-05 | 2.17009432 | 0.672 | 0.293 | 0.03706263 |
| Macrophage | Dpep2 | 1 | 1.91E-05 | 3.98975766 | 0.625 | 0.191 | 0.03811018 |

| | | | | | | | |
|---|---|---|---|---|---|---|---|
| Macrophage | Col24a1 | 0 | 1.96E-05 | 3.05516775 | 0.621 | 0.086 | 0.03920954 |
| Macrophage | Casq2 | 0 | 1.97E-05 | 2.65605183 | 0.586 | 0.069 | 0.03942398 |
| Macrophage | Ntn1 | 0 | 2.00E-05 | 1.53716487 | 0.707 | 0.259 | 0.04008347 |
| Macrophage | Tmeff2 | 0 | 2.01E-05 | 1.9291711 | 0.69 | 0.224 | 0.04022004 |
| Macrophage | Ptprn2 | 0 | 2.02E-05 | 0.69384299 | 0.448 | 0.172 | 0.04037209 |
| Macrophage | Vwf | 0 | 2.05E-05 | 2.49008902 | 0.586 | 0.069 | 0.04105003 |
| Macrophage | Sele | 1 | 2.14E-05 | 4.53390587 | 0.562 | 0.088 | 0.04288389 |
| Macrophage | Sh2d3c | 0 | 2.22E-05 | 1.80768408 | 0.534 | 0.172 | 0.04432573 |
| Macrophage | Ccdc80 | 0 | 2.22E-05 | 1.94698643 | 0.655 | 0.155 | 0.04445332 |
| Macrophage | Adamts17 | 0 | 2.25E-05 | 0.57344574 | 0.638 | 0.086 | 0.04492402 |
| Endothelial Cell | RT1-DOa | 0 | 2.98E-86 | 2.02459253 | 0.497 | 0.024 | 5.95E-83 |
| Endothelial Cell | AABR07068316.2 | 0 | 1.04E-85 | 1.88957911 | 0.433 | 0.011 | 2.08E-82 |
| Endothelial Cell | AABR07002677.2 | 0 | 1.35E-70 | 1.94462519 | 0.548 | 0.041 | 2.69E-67 |
| Endothelial Cell | Ggta1l1 | 0 | 2.61E-68 | 5.72993237 | 0.427 | 0.024 | 5.22E-65 |

| | | | | | | | |
|---|---|---|---|---|---|---|---|
| Endothelial Cell | Shc3 | 0 | 7.53E-65 | 2.79077183 | 0.535 | 0.032 | 1.51E-61 |
| Endothelial Cell | Wnt9b | 0 | 1.72E-64 | 3.34497954 | 0.503 | 0.055 | 3.45E-61 |
| Endothelial Cell | AABR07041411.1 | 0 | 5.11E-58 | 4.72781012 | 0.452 | 0.045 | 1.02E-54 |
| Endothelial Cell | Cpvl | 0 | 1.25E-57 | 3.30454775 | 0.497 | 0.031 | 2.51E-54 |
| Endothelial Cell | Cyp2e1 | 0 | 1.97E-56 | 5.186369 | 0.471 | 0.056 | 3.94E-53 |
| Endothelial Cell | Hydin | 0 | 7.76E-54 | 6.71785347 | 0.287 | 0.009 | 1.55E-50 |
| Endothelial Cell | Nlrp3 | 0 | 3.92E-51 | 1.99796077 | 0.433 | 0.045 | 7.83E-48 |
| Endothelial Cell | Slc15a1 | 0 | 7.13E-51 | 0.61930167 | 0.369 | 0.032 | 1.43E-47 |
| Endothelial Cell | Hmcn1 | 0 | 1.90E-49 | 1.9874693 | 0.955 | 0.693 | 3.81E-46 |
| Endothelial Cell | Edaradd | 0 | 6.88E-48 | 3.08518276 | 0.344 | 0.05 | 1.38E-44 |
| Endothelial Cell | Apba2 | 0 | 3.35E-46 | 1.99893471 | 0.325 | 0.01 | 6.70E-43 |
| Endothelial Cell | Wwc1 | 0 | 2.00E-44 | 2.73384455 | 0.561 | 0.14 | 4.00E-41 |
| Endothelial Cell | AABR07017268.1 | 0 | 8.42E-44 | 1.78555925 | 0.439 | 0.024 | 1.68E-40 |
| Endothelial Cell | Slc35g2 | 0 | 1.12E-43 | 2.67582184 | 0.395 | 0.092 | 2.23E-40 |

| | | | | | | | |
|---|---|---|---|---|---|---|---|
| Endothelial Cell | Kif27 | 0 | 6.26E-43 | 3.42123582 | 0.567 | 0.123 | 1.25E-39 |
| Endothelial Cell | Gria2 | 0 | 5.07E-42 | 4.19127879 | 0.452 | 0.031 | 1.01E-38 |
| Endothelial Cell | LOC690045 | 0 | 5.42E-42 | 1.47076567 | 0.312 | 0.02 | 1.08E-38 |
| Endothelial Cell | AABR07056605.1 | 0 | 1.10E-39 | 5.06460336 | 0.389 | 0.054 | 2.19E-36 |
| Endothelial Cell | RGD1564053 | 0 | 3.68E-38 | 2.30503862 | 0.567 | 0.075 | 7.35E-35 |
| Endothelial Cell | AABR07058170.1 | 0 | 1.18E-33 | 1.54552343 | 0.548 | 0.114 | 2.36E-30 |
| Endothelial Cell | Cpne7 | 0 | 1.28E-32 | 1.72531858 | 0.548 | 0.182 | 2.56E-29 |
| Endothelial Cell | Has2 | 0 | 1.94E-32 | 1.71741402 | 0.541 | 0.1 | 3.88E-29 |
| Endothelial Cell | Mcm5 | 0 | 1.49E-31 | 1.93929299 | 0.382 | 0.027 | 2.99E-28 |
| Endothelial Cell | Cyp4f18 | 0 | 2.02E-30 | 0.68214362 | 0.369 | 0.06 | 4.05E-27 |
| Endothelial Cell | Sell | 0 | 8.71E-30 | 2.3749825 | 0.369 | 0.037 | 1.74E-26 |
| Endothelial Cell | Luzp2 | 0 | 1.74E-29 | 1.98547133 | 0.471 | 0.1 | 3.49E-26 |
| Endothelial Cell | Thsd7b | 0 | 1.89E-29 | 1.73709929 | 0.567 | 0.155 | 3.78E-26 |
| Endothelial Cell | Kcnk5 | 0 | 6.87E-29 | 1.08984832 | 0.516 | 0.12 | 1.37E-25 |

| | | | | | | | |
|---|---|---|---|---|---|---|---|
| Endothelial Cell | Hdac9 | 0 | 2.47E-28 | 1.38978626 | 0.949 | 0.69 | 4.94E-25 |
| Endothelial Cell | Fam189a2 | 0 | 9.12E-28 | 2.02673528 | 0.446 | 0.096 | 1.82E-24 |
| Endothelial Cell | Csmd1 | 0 | 5.09E-27 | 1.81315291 | 0.561 | 0.222 | 1.02E-23 |
| Endothelial Cell | Vegfc | 0 | 7.06E-27 | 1.96230445 | 0.803 | 0.506 | 1.41E-23 |
| Endothelial Cell | Angptl8 | 0 | 9.39E-27 | 2.59424234 | 0.516 | 0.175 | 1.88E-23 |
| Endothelial Cell | Ntng1 | 0 | 1.22E-26 | 2.38753855 | 0.535 | 0.107 | 2.43E-23 |
| Endothelial Cell | Actg2 | 0 | 1.56E-25 | 3.39033588 | 0.51 | 0.128 | 3.12E-22 |
| Endothelial Cell | AC130940.1 | 0 | 1.10E-24 | 1.63117931 | 0.471 | 0.135 | 2.21E-21 |
| Endothelial Cell | Lpar3 | 0 | 2.94E-24 | 2.9196808 | 0.567 | 0.222 | 5.88E-21 |
| Endothelial Cell | Vwf | 0 | 7.40E-24 | 1.48342446 | 0.854 | 0.587 | 1.48E-20 |
| Endothelial Cell | LOC102551356 | 0 | 7.50E-23 | 2.40683421 | 0.484 | 0.064 | 1.50E-19 |
| Endothelial Cell | Lypd1 | 0 | 3.85E-22 | 2.82575358 | 0.395 | 0.109 | 7.70E-19 |
| Endothelial Cell | Cntn3 | 0 | 8.03E-22 | 4.82393039 | 0.478 | 0.203 | 1.61E-18 |
| Endothelial Cell | Pde1c | 0 | 2.39E-21 | 2.78633645 | 0.637 | 0.328 | 4.78E-18 |

| | | | | | | | |
|---|---|---|---|---|---|---|---|
| Endothelial Cell | Mt3 | 0 | 2.31E-20 | 3.0574353 | 0.35 | 0.073 | 4.63E-17 |
| Endothelial Cell | Gfra2 | 0 | 4.21E-20 | 0.40729876 | 0.478 | 0.114 | 8.43E-17 |
| Endothelial Cell | Coa4 | 0 | 6.50E-20 | 0.79837787 | 0.554 | 0.191 | 1.30E-16 |
| Endothelial Cell | Plekha7 | 0 | 2.38E-19 | 2.05150328 | 0.688 | 0.362 | 4.76E-16 |
| Endothelial Cell | Kntc1 | 0 | 5.76E-19 | 0.73427487 | 0.478 | 0.162 | 1.15E-15 |
| Endothelial Cell | Ptprf | 0 | 1.10E-18 | 1.75958284 | 0.331 | 0.052 | 2.20E-15 |
| Endothelial Cell | Tbxas1 | 0 | 1.20E-18 | 2.0083609 | 0.573 | 0.131 | 2.39E-15 |
| Endothelial Cell | Procr | 0 | 2.58E-18 | 0.73923352 | 0.541 | 0.171 | 5.17E-15 |
| Endothelial Cell | Igsf11 | 0 | 8.29E-18 | 0.98677702 | 0.503 | 0.062 | 1.66E-14 |
| Endothelial Cell | Dusp27 | 0 | 1.51E-17 | 1.80738799 | 0.554 | 0.207 | 3.03E-14 |
| Endothelial Cell | Fcer1g | 0 | 1.11E-16 | 2.32264984 | 0.452 | 0.082 | 2.23E-13 |
| Endothelial Cell | Coro1a | 0 | 1.85E-16 | 1.43848911 | 0.561 | 0.227 | 3.69E-13 |
| Endothelial Cell | Lsamp | 0 | 2.09E-16 | 1.06861567 | 0.529 | 0.066 | 4.18E-13 |
| Endothelial Cell | Smoc1 | 0 | 2.40E-16 | 1.14347888 | 0.427 | 0.103 | 4.80E-13 |

| | | | | | | | |
|---|---|---|---|---|---|---|---|
| Endothelial Cell | Ros1 | 0 | 4.50E-16 | 0.83440437 | 0.497 | 0.12 | 8.99E-13 |
| Endothelial Cell | Fos | 0 | 2.75E-15 | 1.34373867 | 0.446 | 0.168 | 5.50E-12 |
| Endothelial Cell | Meox2 | 0 | 7.78E-15 | 1.118516 | 0.898 | 0.637 | 1.56E-11 |
| Endothelial Cell | B3galt2 | 0 | 1.05E-14 | 1.78811685 | 0.541 | 0.197 | 2.10E-11 |
| Endothelial Cell | Zbtb7c | 0 | 1.39E-14 | 1.25045815 | 0.586 | 0.251 | 2.78E-11 |
| Endothelial Cell | Srgap3 | 0 | 1.80E-14 | 0.49708963 | 0.573 | 0.147 | 3.60E-11 |
| Endothelial Cell | Cobl | 0 | 2.56E-14 | 2.5225572 | 0.605 | 0.327 | 5.12E-11 |
| Endothelial Cell | LOC103693323 | 0 | 3.01E-14 | 1.38637656 | 0.427 | 0.097 | 6.02E-11 |
| Endothelial Cell | AABR07006724.1 | 0 | 6.45E-14 | 1.64126439 | 0.58 | 0.289 | 1.29E-10 |
| Endothelial Cell | Asb14 | 0 | 1.57E-13 | 0.30656175 | 0.051 | 0.303 | 3.15E-10 |
| Endothelial Cell | Ctnnd2 | 0 | 2.81E-13 | 1.0485974 | 0.529 | 0.262 | 5.63E-10 |
| Endothelial Cell | Efemp1 | 0 | 3.54E-13 | 1.0378817 | 0.554 | 0.277 | 7.09E-10 |
| Endothelial Cell | Cnnm2 | 0 | 9.05E-13 | 1.26264294 | 0.611 | 0.266 | 1.81E-09 |
| Endothelial Cell | Brca1 | 0 | 2.84E-12 | 1.23965294 | 0.484 | 0.099 | 5.68E-09 |

| | | | | | | | |
|---|---|---|---|---|---|---|---|
| Endothelial Cell | Dkk3 | 0 | 3.54E-12 | 0.40380432 | 0.548 | 0.189 | 7.09E-09 |
| Endothelial Cell | Sema3a | 0 | 3.73E-12 | 1.3011043 | 0.541 | 0.167 | 7.47E-09 |
| Endothelial Cell | Abo | 0 | 3.86E-12 | 0.7059878 | 0.452 | 0.138 | 7.73E-09 |
| Endothelial Cell | Sfxn5 | 0 | 2.05E-11 | 1.40177389 | 0.51 | 0.173 | 4.10E-08 |
| Endothelial Cell | Ppm1e | 0 | 2.59E-11 | 0.26800478 | 0.535 | 0.244 | 5.17E-08 |
| Endothelial Cell | Trabd2b | 0 | 6.78E-11 | 1.51303537 | 0.599 | 0.297 | 1.36E-07 |
| Endothelial Cell | Abcb4 | 0 | 7.80E-11 | 1.59399114 | 0.656 | 0.395 | 1.56E-07 |
| Endothelial Cell | Srpx | 0 | 1.06E-10 | 0.95813445 | 0.713 | 0.402 | 2.11E-07 |
| Endothelial Cell | Sned1 | 0 | 1.38E-10 | 0.63996854 | 0.611 | 0.207 | 2.76E-07 |
| Endothelial Cell | Adamts1 | 0 | 1.54E-10 | 0.54102324 | 0.586 | 0.294 | 3.08E-07 |
| Endothelial Cell | Pkhd1l1 | 0 | 1.82E-10 | 1.28189975 | 0.643 | 0.371 | 3.64E-07 |
| Endothelial Cell | Fcgr2b | 0 | 2.37E-10 | 1.45288992 | 0.376 | 0.029 | 4.75E-07 |
| Endothelial Cell | Ptpro | 0 | 3.17E-10 | 1.55876059 | 0.395 | 0.097 | 6.33E-07 |
| Endothelial Cell | Nkd2 | 0 | 7.28E-10 | 1.63400306 | 0.459 | 0.14 | 1.46E-06 |

| | | | | | | | |
|---|---|---|---|---|---|---|---|
| Endothelial Cell | Sdc2 | 0 | 8.18E-10 | 0.81185246 | 0.58 | 0.303 | 1.64E-06 |
| Endothelial Cell | Ccn2 | 0 | 1.80E-09 | 1.01000913 | 0.554 | 0.23 | 3.60E-06 |
| Endothelial Cell | Kif23 | 0 | 2.15E-09 | 2.16136075 | 0.503 | 0.227 | 4.31E-06 |
| Endothelial Cell | Dscaml1 | 0 | 3.55E-09 | 1.28511192 | 0.433 | 0.174 | 7.11E-06 |
| Endothelial Cell | Ednrb | 0 | 5.67E-09 | 1.72372751 | 0.618 | 0.34 | 1.13E-05 |
| Endothelial Cell | Pstpip1 | 0 | 6.05E-09 | 0.48745527 | 0.529 | 0.222 | 1.21E-05 |
| Endothelial Cell | Ca5b | 0 | 6.15E-09 | 1.4216223 | 0.573 | 0.253 | 1.23E-05 |
| Endothelial Cell | Mapt | 0 | 7.28E-09 | 1.99531603 | 0.567 | 0.296 | 1.46E-05 |
| Endothelial Cell | Frmd3 | 0 | 1.35E-08 | 0.62861467 | 0.503 | 0.236 | 2.69E-05 |
| Endothelial Cell | Art3 | 0 | 1.58E-08 | 0.47497618 | 0.471 | 0.137 | 3.16E-05 |
| Endothelial Cell | AABR07066861.1 | 0 | 2.31E-08 | 2.11906497 | 0.389 | 0.066 | 4.62E-05 |
| Endothelial Cell | Gpm6a | 0 | 3.10E-08 | 1.03382385 | 0.656 | 0.398 | 6.21E-05 |
| Endothelial Cell | Hlf | 0 | 3.36E-08 | 1.13813617 | 0.42 | 0.132 | 6.71E-05 |
| Endothelial Cell | Dtna | 0 | 3.78E-08 | 0.94350267 | 0.561 | 0.24 | 7.56E-05 |

| | | | | | | | |
|---|---|---|---|---|---|---|---|
| Endothelial Cell | Syt14 | 0 | 6.82E-08 | 0.94016301 | 0.529 | 0.241 | 0.00013633 |
| Endothelial Cell | Nrxn1 | 0 | 6.95E-08 | 0.37105649 | 0.675 | 0.343 | 0.00013898 |
| Endothelial Cell | Fgf1 | 0 | 4.95E-07 | 0.67984702 | 0.446 | 0.19 | 0.00098948 |
| Endothelial Cell | Mx2 | 0 | 1.38E-06 | 0.25445473 | 0.586 | 0.301 | 0.00276027 |
| Endothelial Cell | Hdx | 0 | 4.49E-06 | 0.50453519 | 0.49 | 0.219 | 0.0089869 |
| Endothelial Cell | Iqgap3 | 0 | 5.50E-06 | 0.38827349 | 0.006 | 0.282 | 0.01100399 |
| Endothelial Cell | Ppp2r3a | 0 | 2.14E-05 | 0.34464378 | 0.503 | 0.246 | 0.04288504 |
| Endothelial Cell | Mb | 1 | 5.78E-29 | 1.98898191 | 0.894 | 0.606 | 1.16E-25 |
| Endothelial Cell | Tnni3 | 1 | 2.08E-25 | 1.91726895 | 0.879 | 0.604 | 4.17E-22 |
| Endothelial Cell | Tmsb4x | 1 | 5.47E-23 | 1.86432195 | 0.844 | 0.593 | 1.09E-19 |
| Endothelial Cell | Sema3e | 1 | 8.12E-23 | 4.0700783 | 0.369 | 0.083 | 1.62E-19 |
| Endothelial Cell | P2rx1 | 1 | 1.17E-22 | 1.27719013 | 0.355 | 0.085 | 2.34E-19 |
| Endothelial Cell | Fbxl2 | 1 | 1.87E-22 | 1.94112341 | 0.333 | 0.068 | 3.75E-19 |
| Endothelial Cell | Dcx | 1 | 3.28E-20 | 0.50372177 | 0.333 | 0.075 | 6.56E-17 |

| | | | | | | | |
|---|---|---|---|---|---|---|---|
| Endothelial Cell | Csf3r | 1 | 1.28E-18 | 3.88086439 | 0.326 | 0.03 | 2.55E-15 |
| Endothelial Cell | Tpt1 | 1 | 6.39E-13 | 2.30391616 | 0.667 | 0.395 | 1.28E-09 |
| Endothelial Cell | Alk | 1 | 3.15E-11 | 1.72379023 | 0.326 | 0.036 | 6.30E-08 |
| Endothelial Cell | Pi15 | 1 | 8.85E-10 | 2.7238659 | 0.362 | 0.109 | 1.77E-06 |
| Endothelial Cell | Ighm | 1 | 1.30E-08 | 3.2610923 | 0.348 | 0.079 | 2.60E-05 |
| Endothelial Cell | Zfp385b | 1 | 1.16E-07 | 0.26266993 | 0.078 | 0.368 | 0.00023263 |
| Endothelial Cell | Actn2 | 1 | 2.30E-07 | 1.89352915 | 0.553 | 0.241 | 0.00045998 |
| Endothelial Cell | Trim55 | 1 | 4.29E-07 | 0.38313258 | 0.128 | 0.387 | 0.0008581 |
| Endothelial Cell | Nrg4 | 2 | 4.25E-40 | 5.18057857 | 0.375 | 0.047 | 8.51E-37 |
| Endothelial Cell | Nrp2 | 2 | 5.06E-32 | 2.39466213 | 0.891 | 0.596 | 1.01E-28 |
| Endothelial Cell | Mest | 2 | 3.44E-31 | 5.57223631 | 0.336 | 0.048 | 6.89E-28 |
| Endothelial Cell | Smc1b | 2 | 2.60E-28 | 3.34245714 | 0.297 | 0.035 | 5.20E-25 |
| Endothelial Cell | Kmo | 2 | 2.65E-22 | 3.88665939 | 0.305 | 0.004 | 5.30E-19 |
| Endothelial Cell | Neil3 | 2 | 6.46E-22 | 0.76195985 | 0.336 | 0.066 | 1.29E-18 |

| | | | | | | | |
|---|---|---|---|---|---|---|---|
| Endothelial Cell | Tp63 | 2 | 1.17E-20 | 4.95902365 | 0.266 | 0.003 | 2.35E-17 |
| Endothelial Cell | Rergl | 2 | 4.52E-20 | 4.04034246 | 0.328 | 0.043 | 9.04E-17 |
| Endothelial Cell | Nox4 | 2 | 2.24E-19 | 2.73299594 | 0.742 | 0.41 | 4.49E-16 |
| Endothelial Cell | Kit | 2 | 4.11E-11 | 3.34713393 | 0.555 | 0.28 | 8.22E-08 |
| Endothelial Cell | Tnc | 2 | 8.51E-10 | 3.66985377 | 0.398 | 0.107 | 1.70E-06 |
| Endothelial Cell | Myo1b | 2 | 1.30E-09 | 1.52142926 | 0.688 | 0.435 | 2.59E-06 |
| Endothelial Cell | Dgki | 2 | 4.44E-07 | 0.48197158 | 0.07 | 0.324 | 0.00088888 |
| Endothelial Cell | Mlph | 3 | 1.21E-63 | 3.0182542 | 0.514 | 0.045 | 2.43E-60 |
| Endothelial Cell | C1qb | 3 | 3.27E-59 | 4.18358675 | 0.624 | 0.071 | 6.53E-56 |
| Endothelial Cell | Erbb3 | 3 | 1.46E-54 | 3.8958424 | 0.615 | 0.094 | 2.92E-51 |
| Endothelial Cell | Ccl6 | 3 | 1.23E-53 | 3.01993299 | 0.532 | 0.065 | 2.46E-50 |
| Endothelial Cell | Trpc6 | 3 | 5.17E-52 | 2.12002686 | 0.624 | 0.101 | 1.03E-48 |
| Endothelial Cell | Klhdc8a | 3 | 1.09E-50 | 5.37160466 | 0.44 | 0.042 | 2.17E-47 |
| Endothelial Cell | Marchf10 | 3 | 6.71E-47 | 2.95857519 | 0.477 | 0.03 | 1.34E-43 |

| | | | | | | | |
|---|---|---|---|---|---|---|---|
| Endothelial Cell | Lilrb2 | 3 | 3.04E-46 | 1.50138367 | 0.587 | 0.096 | 6.08E-43 |
| Endothelial Cell | Btla | 3 | 1.25E-45 | 4.23410269 | 0.596 | 0.121 | 2.51E-42 |
| Endothelial Cell | Pik3r5 | 3 | 2.39E-44 | 2.70727977 | 0.339 | 0.061 | 4.77E-41 |
| Endothelial Cell | Nkain2 | 3 | 2.00E-41 | 5.30453665 | 0.312 | 0.012 | 4.00E-38 |
| Endothelial Cell | Gpc3 | 3 | 3.84E-41 | 4.51208094 | 0.624 | 0.102 | 7.68E-38 |
| Endothelial Cell | Ptger3 | 3 | 6.39E-40 | 3.28924163 | 0.404 | 0.043 | 1.28E-36 |
| Endothelial Cell | Olfml2b | 3 | 2.45E-33 | 2.65083816 | 0.376 | 0.067 | 4.91E-30 |
| Endothelial Cell | F3 | 3 | 1.18E-32 | 2.48750571 | 0.431 | 0.074 | 2.37E-29 |
| Endothelial Cell | Grip2 | 3 | 2.55E-29 | 2.96461195 | 0.477 | 0.102 | 5.11E-26 |
| Endothelial Cell | Cldn22 | 3 | 3.76E-28 | 3.22800045 | 0.624 | 0.189 | 7.51E-25 |
| Endothelial Cell | Gfra3 | 3 | 1.35E-27 | 2.114032 | 0.633 | 0.183 | 2.70E-24 |
| Endothelial Cell | Ubap1l | 3 | 2.10E-26 | 2.1522909 | 0.661 | 0.29 | 4.20E-23 |
| Endothelial Cell | Rp1 | 3 | 3.46E-26 | 2.14000892 | 0.56 | 0.129 | 6.93E-23 |
| Endothelial Cell | Kcnt2 | 3 | 1.15E-25 | 2.72308048 | 0.789 | 0.357 | 2.30E-22 |

| | | | | | | | |
|---|---|---|---|---|---|---|---|
| Endothelial Cell | AABR07007068.1 | 3 | 1.05E-23 | 2.09623268 | 0.596 | 0.216 | 2.10E-20 |
| Endothelial Cell | Pkmyt1 | 3 | 8.42E-23 | 4.05674188 | 0.468 | 0.047 | 1.68E-19 |
| Endothelial Cell | Anln | 3 | 1.16E-22 | 2.26259189 | 0.477 | 0.089 | 2.32E-19 |
| Endothelial Cell | RT1-Db1 | 3 | 1.99E-22 | 0.57411603 | 0.56 | 0.183 | 3.99E-19 |
| Endothelial Cell | St6galnac3 | 3 | 3.04E-22 | 1.74516739 | 0.917 | 0.425 | 6.09E-19 |
| Endothelial Cell | Depdc1b | 3 | 8.90E-22 | 1.85597234 | 0.303 | 0.05 | 1.78E-18 |
| Endothelial Cell | Cdh4 | 3 | 2.54E-21 | 1.44202461 | 0.596 | 0.244 | 5.08E-18 |
| Endothelial Cell | Myo10 | 3 | 6.10E-21 | 1.19140126 | 0.972 | 0.718 | 1.22E-17 |
| Endothelial Cell | Amd1 | 3 | 3.45E-20 | 2.02354383 | 0.761 | 0.378 | 6.90E-17 |
| Endothelial Cell | Slc44a5 | 3 | 3.74E-20 | 0.5762776 | 0.514 | 0.139 | 7.48E-17 |
| Endothelial Cell | Trib3 | 3 | 5.50E-20 | 2.06446931 | 0.55 | 0.163 | 1.10E-16 |
| Endothelial Cell | Agbl1 | 3 | 1.02E-19 | 1.65451915 | 0.642 | 0.215 | 2.05E-16 |
| Endothelial Cell | Dscaml11 | 3 | 3.70E-19 | 1.28937957 | 0.56 | 0.173 | 7.40E-16 |
| Endothelial Cell | Dcx1 | 3 | 4.12E-19 | 1.88587511 | 0.358 | 0.08 | 8.24E-16 |

| | | | | | | | |
|---|---|---|---|---|---|---|---|
| Endothelial Cell | Il34 | 3 | 4.78E-19 | 1.58840722 | 0.422 | 0.114 | 9.55E-16 |
| Endothelial Cell | Tmem100 | 3 | 6.14E-19 | 0.3376803 | 0.661 | 0.247 | 1.23E-15 |
| Endothelial Cell | Tceal7 | 3 | 8.58E-19 | 0.57062595 | 0.312 | 0.039 | 1.72E-15 |
| Endothelial Cell | Kif26b | 3 | 1.52E-18 | 1.73926131 | 0.624 | 0.198 | 3.05E-15 |
| Endothelial Cell | Col11a1 | 3 | 2.39E-18 | 1.69823254 | 0.45 | 0.11 | 4.79E-15 |
| Endothelial Cell | Rbp7 | 3 | 3.09E-18 | 1.33513876 | 0.725 | 0.316 | 6.18E-15 |
| Endothelial Cell | Csrnp1 | 3 | 4.93E-18 | 2.12357996 | 0.596 | 0.197 | 9.85E-15 |
| Endothelial Cell | Btnl9 | 3 | 4.93E-18 | 2.24623949 | 0.633 | 0.284 | 9.87E-15 |
| Endothelial Cell | Fgf13 | 3 | 6.03E-18 | 0.31387092 | 0.587 | 0.105 | 1.21E-14 |
| Endothelial Cell | Enox1 | 3 | 9.40E-18 | 1.27194 | 0.633 | 0.225 | 1.88E-14 |
| Endothelial Cell | Sorcs1 | 3 | 9.50E-18 | 0.5483024 | 0.642 | 0.29 | 1.90E-14 |
| Endothelial Cell | Efnb2 | 3 | 5.26E-17 | 1.42932067 | 0.89 | 0.579 | 1.05E-13 |
| Endothelial Cell | Col11a2 | 3 | 5.49E-17 | 1.89967664 | 0.394 | 0.084 | 1.10E-13 |
| Endothelial Cell | Nr4a3 | 3 | 6.02E-17 | 0.95452356 | 0.505 | 0.088 | 1.20E-13 |

| | | | | | | | |
|---|---|---|---|---|---|---|---|
| Endothelial Cell | Cfh | 3 | 6.21E-17 | 1.32107295 | 0.697 | 0.395 | 1.24E-13 |
| Endothelial Cell | Galnt15 | 3 | 7.35E-17 | 1.93777028 | 0.679 | 0.266 | 1.47E-13 |
| Endothelial Cell | Magi2 | 3 | 1.31E-16 | 1.77539291 | 0.67 | 0.355 | 2.63E-13 |
| Endothelial Cell | LOC685963 | 3 | 3.13E-16 | 1.11143929 | 0.706 | 0.396 | 6.26E-13 |
| Endothelial Cell | Ano1 | 3 | 3.67E-16 | 1.48456547 | 0.633 | 0.347 | 7.34E-13 |
| Endothelial Cell | AABR07031740.1 | 3 | 3.88E-16 | 1.32918392 | 0.679 | 0.397 | 7.76E-13 |
| Endothelial Cell | Runx2 | 3 | 4.73E-16 | 1.06226132 | 0.615 | 0.252 | 9.45E-13 |
| Endothelial Cell | Lamc3 | 3 | 1.60E-15 | 0.65743784 | 0.404 | 0.123 | 3.19E-12 |
| Endothelial Cell | Tgfb2 | 3 | 1.67E-15 | 1.27427943 | 0.743 | 0.357 | 3.33E-12 |
| Endothelial Cell | AABR07042840.1 | 3 | 1.91E-15 | 0.39811519 | 0.642 | 0.196 | 3.81E-12 |
| Endothelial Cell | Mamdc2 | 3 | 2.83E-15 | 0.64301029 | 0.651 | 0.345 | 5.65E-12 |
| Endothelial Cell | Cspg4 | 3 | 5.15E-15 | 0.36304807 | 0.587 | 0.233 | 1.03E-11 |
| Endothelial Cell | Rab11fip4 | 3 | 5.59E-15 | 0.40513874 | 0.633 | 0.183 | 1.12E-11 |
| Endothelial Cell | Nebl | 3 | 5.61E-15 | 0.84222376 | 0.927 | 0.576 | 1.12E-11 |

| | | | | | | | |
|---|---|---|---|---|---|---|---|
| Endothelial Cell | Pcdh17 | 3 | 7.75E-15 | 1.69617797 | 0.771 | 0.495 | 1.55E-11 |
| Endothelial Cell | Fam83b | 3 | 1.41E-14 | 1.54666164 | 0.615 | 0.236 | 2.82E-11 |
| Endothelial Cell | Postn | 3 | 2.74E-14 | 1.18313701 | 0.587 | 0.161 | 5.48E-11 |
| Endothelial Cell | Slc24a2 | 3 | 2.74E-14 | 1.84765387 | 0.587 | 0.231 | 5.48E-11 |
| Endothelial Cell | Ptafr | 3 | 3.23E-14 | 1.29916641 | 0.661 | 0.315 | 6.45E-11 |
| Endothelial Cell | Coa41 | 3 | 4.05E-14 | 1.41762228 | 0.486 | 0.214 | 8.10E-11 |
| Endothelial Cell | Esm1 | 3 | 6.58E-14 | 0.58687086 | 0.651 | 0.275 | 1.32E-10 |
| Endothelial Cell | Kcnk2 | 3 | 7.33E-14 | 2.11268228 | 0.523 | 0.265 | 1.47E-10 |
| Endothelial Cell | Crip1 | 3 | 7.76E-14 | 1.27973267 | 0.706 | 0.331 | 1.55E-10 |
| Endothelial Cell | Clybl | 3 | 1.33E-13 | 1.70090325 | 0.716 | 0.37 | 2.65E-10 |
| Endothelial Cell | Thbd | 3 | 2.64E-13 | 0.79560691 | 0.697 | 0.403 | 5.29E-10 |
| Endothelial Cell | Itgb8 | 3 | 3.35E-13 | 0.30692779 | 0.67 | 0.399 | 6.70E-10 |
| Endothelial Cell | Clmn | 3 | 3.89E-13 | 1.38491697 | 0.596 | 0.307 | 7.79E-10 |
| Endothelial Cell | Atp5f1e | 3 | 4.00E-13 | 1.20011869 | 0.752 | 0.41 | 8.00E-10 |

| | | | | | | | |
|---|---|---|---|---|---|---|---|
| Endothelial Cell | AABR07001519.1 | 3 | 4.60E-13 | 1.14181935 | 0.716 | 0.318 | 9.20E-10 |
| Endothelial Cell | St8sia4 | 3 | 4.87E-13 | 1.38634363 | 0.789 | 0.374 | 9.73E-10 |
| Endothelial Cell | Synm | 3 | 1.02E-12 | 0.8512944 | 0.56 | 0.183 | 2.04E-09 |
| Endothelial Cell | Tc2n | 3 | 1.72E-12 | 1.10716077 | 0.587 | 0.279 | 3.43E-09 |
| Endothelial Cell | Mfap5 | 3 | 1.78E-12 | 1.76347487 | 0.587 | 0.213 | 3.56E-09 |
| Endothelial Cell | Pdgfd | 3 | 2.27E-12 | 1.14147019 | 0.761 | 0.427 | 4.54E-09 |
| Endothelial Cell | Galnt17 | 3 | 2.61E-12 | 0.33110338 | 0.596 | 0.215 | 5.22E-09 |
| Endothelial Cell | Cacna2d3 | 3 | 4.24E-12 | 1.19747848 | 0.596 | 0.214 | 8.48E-09 |
| Endothelial Cell | Lrrc17 | 3 | 4.81E-12 | 1.72249478 | 0.578 | 0.241 | 9.62E-09 |
| Endothelial Cell | Zmat4 | 3 | 6.33E-12 | 0.65663358 | 0.055 | 0.307 | 1.27E-08 |
| Endothelial Cell | Iqub | 3 | 1.11E-11 | 0.63691562 | 0.67 | 0.351 | 2.22E-08 |
| Endothelial Cell | Ckmt2 | 3 | 2.07E-11 | 0.59210782 | 0.697 | 0.363 | 4.15E-08 |
| Endothelial Cell | Slc26a10 | 3 | 4.31E-11 | 1.00551927 | 0.798 | 0.542 | 8.61E-08 |
| Endothelial Cell | Tnfrsf1b | 3 | 6.71E-11 | 0.27832616 | 0.716 | 0.38 | 1.34E-07 |

| | | | | | | | |
|---|---|---|---|---|---|---|---|
| Endothelial Cell | Slc35f1 | 3 | 1.52E-10 | 1.10856542 | 0.688 | 0.386 | 3.04E-07 |
| Endothelial Cell | Megf91 | 3 | 1.52E-10 | 0.70353117 | 0.651 | 0.293 | 3.05E-07 |
| Endothelial Cell | Frmd5 | 3 | 2.44E-10 | 1.03580373 | 0.532 | 0.169 | 4.87E-07 |
| Endothelial Cell | Anks1b | 3 | 2.74E-10 | 0.6487806 | 0.505 | 0.213 | 5.48E-07 |
| Endothelial Cell | Npr3 | 3 | 3.00E-10 | 0.87841826 | 0.532 | 0.156 | 6.00E-07 |
| Endothelial Cell | AABR07026536.1 | 3 | 3.52E-10 | 1.1810648 | 0.523 | 0.25 | 7.03E-07 |
| Endothelial Cell | Ndufa4 | 3 | 4.06E-10 | 0.65845519 | 0.716 | 0.356 | 8.13E-07 |
| Endothelial Cell | Lilrb3a | 3 | 5.42E-10 | 1.41563555 | 0.514 | 0.225 | 1.08E-06 |
| Endothelial Cell | Gpm6b | 3 | 8.58E-10 | 0.87791422 | 0.587 | 0.279 | 1.72E-06 |
| Endothelial Cell | Jag1 | 3 | 9.14E-10 | 0.43377717 | 0.725 | 0.383 | 1.83E-06 |
| Endothelial Cell | Mpc1 | 3 | 1.37E-09 | 0.45267574 | 0.633 | 0.297 | 2.75E-06 |
| Endothelial Cell | Lancl3 | 3 | 1.45E-09 | 0.53548278 | 0.358 | 0.097 | 2.91E-06 |
| Endothelial Cell | Rnf152 | 3 | 1.60E-09 | 0.80101721 | 0.661 | 0.366 | 3.19E-06 |
| Endothelial Cell | RGD1563354 | 3 | 2.05E-09 | 0.43008687 | 0.642 | 0.269 | 4.09E-06 |

| | | | | | | | |
|---|---|---|---|---|---|---|---|
| Endothelial Cell | Rrad | 3 | 2.06E-09 | 1.05768038 | 0.624 | 0.344 | 4.12E-06 |
| Endothelial Cell | Unc5b | 3 | 2.09E-09 | 1.17828101 | 0.688 | 0.409 | 4.18E-06 |
| Endothelial Cell | Fign | 3 | 2.13E-09 | 1.116874 | 0.394 | 0.086 | 4.27E-06 |
| Endothelial Cell | Col6a2 | 3 | 3.82E-09 | 0.76365574 | 0.642 | 0.352 | 7.64E-06 |
| Endothelial Cell | Arhgap24 | 3 | 6.50E-09 | 0.45344855 | 0.67 | 0.365 | 1.30E-05 |
| Endothelial Cell | Sybu | 3 | 6.61E-09 | 1.11235511 | 0.706 | 0.401 | 1.32E-05 |
| Endothelial Cell | Plekha4 | 3 | 7.18E-09 | 0.59073058 | 0.596 | 0.308 | 1.44E-05 |
| Endothelial Cell | Enpp3 | 3 | 9.23E-09 | 0.83431669 | 0.642 | 0.303 | 1.85E-05 |
| Endothelial Cell | Sfxn51 | 3 | 1.26E-08 | 0.44384573 | 0.477 | 0.191 | 2.52E-05 |
| Endothelial Cell | AABR07035916.1 | 3 | 1.35E-08 | 0.90243371 | 0.761 | 0.467 | 2.71E-05 |
| Endothelial Cell | Dgkg | 3 | 1.38E-08 | 0.38185617 | 0.385 | 0.101 | 2.76E-05 |
| Endothelial Cell | Daam2 | 3 | 1.45E-08 | 1.42144587 | 0.404 | 0.103 | 2.91E-05 |
| Endothelial Cell | Alpl | 3 | 1.49E-08 | 0.92116585 | 0.67 | 0.377 | 2.98E-05 |
| Endothelial Cell | Gvin1 | 3 | 1.97E-08 | 0.94517333 | 0.56 | 0.24 | 3.93E-05 |

| | | | | | | | |
|---|---|---|---|---|---|---|---|
| Endothelial Cell | Arhgef37 | 3 | 2.07E-08 | 0.64438467 | 0.505 | 0.184 | 4.14E-05 |
| Endothelial Cell | AABR07044 900.1 | 3 | 2.45E-08 | 0.28742461 | 0.679 | 0.327 | 4.91E-05 |
| Endothelial Cell | Slc66a1 | 3 | 2.56E-08 | 0.46560049 | 0.477 | 0.118 | 5.11E-05 |
| Endothelial Cell | Abcc9 | 3 | 3.15E-08 | 0.31662686 | 0.679 | 0.339 | 6.31E-05 |
| Endothelial Cell | Aox1 | 3 | 4.09E-08 | 0.60852127 | 0.569 | 0.293 | 8.18E-05 |
| Endothelial Cell | Htra2 | 3 | 5.30E-08 | 1.05148918 | 0.532 | 0.217 | 0.00010591 |
| Endothelial Cell | Ppm1h | 3 | 5.58E-08 | 0.7102461 | 0.807 | 0.534 | 0.00011168 |
| Endothelial Cell | Limch1 | 3 | 8.60E-08 | 0.71147865 | 0.872 | 0.603 | 0.00017201 |
| Endothelial Cell | Sh3gl2 | 3 | 1.00E-07 | 0.86500031 | 0.578 | 0.246 | 0.00020028 |
| Endothelial Cell | Tpt11 | 3 | 1.23E-07 | 0.57239132 | 0.697 | 0.4 | 0.00024622 |
| Endothelial Cell | Inpp5d | 3 | 1.61E-07 | 0.63820175 | 0.716 | 0.401 | 0.00032157 |
| Endothelial Cell | Papss2 | 3 | 2.46E-07 | 0.57326345 | 0.78 | 0.46 | 0.00049192 |
| Endothelial Cell | Kcnd3 | 3 | 2.90E-07 | 0.51562088 | 0.56 | 0.291 | 0.00058068 |
| Endothelial Cell | Fbln5 | 3 | 3.56E-07 | 0.75802265 | 0.67 | 0.416 | 0.00071262 |

| | | | | | | | |
|---|---|---|---|---|---|---|---|
| Endothelial Cell | Bnc2 | 3 | 4.37E-07 | 0.40545486 | 0.606 | 0.329 | 0.00087451 |
| Endothelial Cell | Chn1 | 3 | 4.42E-07 | 0.44814596 | 0.761 | 0.422 | 0.00088466 |
| Endothelial Cell | Pygm | 3 | 5.29E-07 | 1.06118935 | 0.532 | 0.266 | 0.00105717 |
| Endothelial Cell | Nnt | 3 | 8.16E-07 | 0.46231752 | 0.734 | 0.431 | 0.00163128 |
| Endothelial Cell | Phyh | 3 | 8.88E-07 | 0.54770982 | 0.725 | 0.424 | 0.00177592 |
| Endothelial Cell | AABR07040864.1 | 3 | 1.10E-06 | 0.27633008 | 0.541 | 0.212 | 0.00219767 |
| Endothelial Cell | Cadps | 3 | 1.70E-06 | 1.50197623 | 0.569 | 0.304 | 0.00340279 |
| Endothelial Cell | Ca8 | 3 | 2.96E-06 | 0.52689722 | 0.514 | 0.239 | 0.00592451 |
| Endothelial Cell | Anp32b | 3 | 3.17E-06 | 0.56484678 | 0.633 | 0.352 | 0.00633918 |
| Endothelial Cell | Dgkz | 3 | 7.40E-06 | 0.50189796 | 0.706 | 0.421 | 0.01479658 |
| Endothelial Cell | Atp2a2 | 3 | 2.04E-05 | 0.43228747 | 0.826 | 0.563 | 0.0407516 |
| Endothelial Cell | Cacna1b | 4 | 2.46E-71 | 3.10995805 | 0.759 | 0.114 | 4.92E-68 |
| Endothelial Cell | Alox5 | 4 | 1.61E-66 | 3.9559657 | 0.704 | 0.073 | 3.22E-63 |
| Endothelial Cell | Ccdc88b | 4 | 2.51E-59 | 1.54639888 | 0.685 | 0.105 | 5.01E-56 |

| | | | | | | | |
|---|---|---|---|---|---|---|---|
| Endothelial Cell | Tmprss6 | 4 | 8.06E-47 | 5.26255434 | 0.509 | 0.068 | 1.61E-43 |
| Endothelial Cell | Rbm44 | 4 | 2.08E-46 | 1.60445963 | 0.574 | 0.096 | 4.16E-43 |
| Endothelial Cell | Abcc8 | 4 | 2.89E-45 | 3.34455268 | 0.546 | 0.079 | 5.77E-42 |
| Endothelial Cell | Gba3 | 4 | 5.73E-45 | 2.03077478 | 0.574 | 0.098 | 1.15E-41 |
| Endothelial Cell | AABR07065190.1 | 4 | 2.22E-44 | 0.60333607 | 0.574 | 0.085 | 4.44E-41 |
| Endothelial Cell | Olfm2 | 4 | 3.20E-44 | 1.70986886 | 0.694 | 0.113 | 6.40E-41 |
| Endothelial Cell | Adgre4 | 4 | 4.72E-44 | 3.75523108 | 0.63 | 0.125 | 9.44E-41 |
| Endothelial Cell | Setmar | 4 | 7.75E-44 | 3.29343302 | 0.704 | 0.148 | 1.55E-40 |
| Endothelial Cell | AABR07054716.1 | 4 | 9.44E-43 | 1.48332192 | 0.593 | 0.115 | 1.89E-39 |
| Endothelial Cell | LOC100910237 | 4 | 3.85E-42 | 2.40942098 | 0.685 | 0.158 | 7.71E-39 |
| Endothelial Cell | Kcp | 4 | 2.14E-39 | 1.8168884 | 0.519 | 0.09 | 4.28E-36 |
| Endothelial Cell | AABR07026614.1 | 4 | 2.04E-38 | 2.36445476 | 0.731 | 0.15 | 4.08E-35 |
| Endothelial Cell | Sh3bp2 | 4 | 5.69E-38 | 1.63919976 | 0.741 | 0.233 | 1.14E-34 |
| Endothelial Cell | AABR07032338.1 | 4 | 6.05E-38 | 0.7335744 | 0.481 | 0.059 | 1.21E-34 |

| | | | | | | | |
|---|---|---|---|---|---|---|---|
| Endothelial Cell | Asb141 | 4 | 1.82E-36 | 1.04958887 | 0.75 | 0.222 | 3.65E-33 |
| Endothelial Cell | Srpk3 | 4 | 3.01E-35 | 0.9018326 | 0.787 | 0.274 | 6.02E-32 |
| Endothelial Cell | AABR07007026.1 | 4 | 7.20E-35 | 1.35482646 | 0.713 | 0.181 | 1.44E-31 |
| Endothelial Cell | Gdf6 | 4 | 1.02E-34 | 0.42776914 | 0.593 | 0.135 | 2.03E-31 |
| Endothelial Cell | Nrg2 | 4 | 1.89E-34 | 2.08418709 | 0.62 | 0.165 | 3.78E-31 |
| Endothelial Cell | AABR07001068.1 | 4 | 7.20E-34 | 1.22196934 | 0.778 | 0.221 | 1.44E-30 |
| Endothelial Cell | AABR07040095.1 | 4 | 1.34E-32 | 1.69264298 | 0.75 | 0.242 | 2.69E-29 |
| Endothelial Cell | Slit3 | 4 | 6.19E-32 | 1.60281733 | 0.75 | 0.272 | 1.24E-28 |
| Endothelial Cell | Map3k7cl | 4 | 1.06E-31 | 1.11113652 | 0.667 | 0.173 | 2.11E-28 |
| Endothelial Cell | Matn2 | 4 | 3.80E-31 | 2.10696151 | 0.796 | 0.29 | 7.59E-28 |
| Endothelial Cell | Gap43 | 4 | 6.07E-31 | 1.89128068 | 0.685 | 0.18 | 1.21E-27 |
| Endothelial Cell | Serpine1 | 4 | 1.03E-30 | 0.8303609 | 0.787 | 0.241 | 2.06E-27 |
| Endothelial Cell | S100a4 | 4 | 1.38E-30 | 0.95704065 | 0.778 | 0.289 | 2.76E-27 |
| Endothelial Cell | Prlr | 4 | 1.95E-30 | 2.58319039 | 0.602 | 0.114 | 3.89E-27 |

| | | | | | | | |
|---|---|---|---|---|---|---|---|
| Endothelial Cell | Slc24a21 | 4 | 3.23E-30 | 2.42629395 | 0.694 | 0.221 | 6.46E-27 |
| Endothelial Cell | Slc27a2 | 4 | 5.23E-30 | 2.33023666 | 0.722 | 0.232 | 1.05E-26 |
| Endothelial Cell | Tc2n1 | 4 | 5.30E-30 | 1.36829935 | 0.769 | 0.261 | 1.06E-26 |
| Endothelial Cell | Vangl2 | 4 | 8.01E-30 | 1.50748782 | 0.676 | 0.171 | 1.60E-26 |
| Endothelial Cell | Slc38a3 | 4 | 1.07E-29 | 0.56121454 | 0.704 | 0.101 | 2.14E-26 |
| Endothelial Cell | Col24a1 | 4 | 1.29E-29 | 2.34289982 | 0.75 | 0.305 | 2.59E-26 |
| Endothelial Cell | Rassf4 | 4 | 1.51E-29 | 0.90575248 | 0.759 | 0.291 | 3.01E-26 |
| Endothelial Cell | Efhd1 | 4 | 3.25E-28 | 1.38323181 | 0.676 | 0.169 | 6.50E-25 |
| Endothelial Cell | Gmpr | 4 | 6.93E-28 | 0.90176945 | 0.731 | 0.239 | 1.39E-24 |
| Endothelial Cell | Hey2 | 4 | 1.14E-27 | 0.82403171 | 0.62 | 0.153 | 2.27E-24 |
| Endothelial Cell | Pla2g2a | 4 | 1.16E-27 | 1.89918712 | 0.694 | 0.21 | 2.32E-24 |
| Endothelial Cell | Xrra1 | 4 | 1.33E-27 | 1.053789 | 0.602 | 0.162 | 2.66E-24 |
| Endothelial Cell | AC111831.1 | 4 | 1.11E-26 | 2.25450458 | 0.722 | 0.208 | 2.21E-23 |
| Endothelial Cell | Gata3 | 4 | 6.84E-26 | 0.27501166 | 0.657 | 0.244 | 1.37E-22 |

| | | | | | | | |
|---|---|---|---|---|---|---|---|
| Endothelial Cell | Lrsam1 | 4 | 1.43E-25 | 0.74899536 | 0.769 | 0.274 | 2.86E-22 |
| Endothelial Cell | Gpr63 | 4 | 2.08E-25 | 1.75829765 | 0.704 | 0.185 | 4.16E-22 |
| Endothelial Cell | E2f7 | 4 | 2.57E-25 | 1.36824813 | 0.75 | 0.298 | 5.14E-22 |
| Endothelial Cell | Hcn1 | 4 | 6.82E-25 | 2.42328173 | 0.5 | 0.087 | 1.36E-21 |
| Endothelial Cell | Bdh1 | 4 | 1.47E-24 | 1.70697425 | 0.759 | 0.368 | 2.94E-21 |
| Endothelial Cell | Kif231 | 4 | 2.17E-24 | 1.15702008 | 0.694 | 0.221 | 4.33E-21 |
| Endothelial Cell | Lrrc171 | 4 | 2.57E-24 | 0.4892357 | 0.741 | 0.225 | 5.14E-21 |
| Endothelial Cell | Plekha41 | 4 | 3.28E-24 | 2.10916145 | 0.741 | 0.294 | 6.56E-21 |
| Endothelial Cell | Fbxl21 | 4 | 3.35E-24 | 3.22850016 | 0.398 | 0.07 | 6.70E-21 |
| Endothelial Cell | Pla2g7 | 4 | 6.85E-24 | 1.09369159 | 0.593 | 0.171 | 1.37E-20 |
| Endothelial Cell | Fam167b | 4 | 7.89E-24 | 1.86063347 | 0.602 | 0.147 | 1.58E-20 |
| Endothelial Cell | Kcnma1 | 4 | 9.20E-24 | 1.70826579 | 0.713 | 0.239 | 1.84E-20 |
| Endothelial Cell | Slc16a1 | 4 | 1.24E-23 | 0.71267025 | 0.713 | 0.158 | 2.47E-20 |
| Endothelial Cell | Rcan1 | 4 | 1.77E-23 | 1.39813198 | 0.759 | 0.327 | 3.53E-20 |

| | | | | | | | |
|---|---|---|---|---|---|---|---|
| Endothelial Cell | Susd5 | 4 | 2.00E-23 | 0.97974989 | 0.426 | 0.099 | 3.99E-20 |
| Endothelial Cell | Trim54 | 4 | 2.49E-23 | 0.34871322 | 0.741 | 0.313 | 4.98E-20 |
| Endothelial Cell | Itga7 | 4 | 6.09E-23 | 0.34959363 | 0.759 | 0.262 | 1.22E-19 |
| Endothelial Cell | Ddc | 4 | 1.36E-22 | 1.37351134 | 0.741 | 0.292 | 2.71E-19 |
| Endothelial Cell | Hap1 | 4 | 1.37E-22 | 1.1674432 | 0.667 | 0.253 | 2.73E-19 |
| Endothelial Cell | Slc4a3 | 4 | 2.49E-22 | 0.80272745 | 0.704 | 0.239 | 4.98E-19 |
| Endothelial Cell | Negr1 | 4 | 2.81E-22 | 0.52567736 | 0.722 | 0.262 | 5.62E-19 |
| Endothelial Cell | Pstpip2 | 4 | 4.45E-22 | 1.36137432 | 0.676 | 0.291 | 8.89E-19 |
| Endothelial Cell | Trpc3 | 4 | 8.53E-22 | 1.51272382 | 0.417 | 0.154 | 1.71E-18 |
| Endothelial Cell | Usp2 | 4 | 8.79E-22 | 0.46468465 | 0.741 | 0.277 | 1.76E-18 |
| Endothelial Cell | Ppm1l | 4 | 9.08E-22 | 1.05498542 | 0.796 | 0.391 | 1.82E-18 |
| Endothelial Cell | Popdc2 | 4 | 2.07E-21 | 1.32590812 | 0.75 | 0.273 | 4.15E-18 |
| Endothelial Cell | Ror2 | 4 | 4.56E-21 | 0.61025487 | 0.75 | 0.327 | 9.12E-18 |
| Endothelial Cell | Casz1 | 4 | 5.05E-21 | 1.49521806 | 0.778 | 0.351 | 1.01E-17 |

| | | | | | | | |
|---|---|---|---|---|---|---|---|
| Endothelial Cell | Coro1a1 | 4 | 7.96E-21 | 2.32418168 | 0.583 | 0.24 | 1.59E-17 |
| Endothelial Cell | Sgip1 | 4 | 1.12E-20 | 1.33938711 | 0.778 | 0.327 | 2.25E-17 |
| Endothelial Cell | Samd151 | 4 | 1.89E-20 | 0.99674181 | 0.481 | 0.11 | 3.77E-17 |
| Endothelial Cell | Antxr1 | 4 | 3.36E-20 | 1.02864948 | 0.769 | 0.36 | 6.72E-17 |
| Endothelial Cell | Neil31 | 4 | 3.65E-20 | 1.0893153 | 0.343 | 0.071 | 7.30E-17 |
| Endothelial Cell | Smyd1 | 4 | 3.71E-20 | 0.86931819 | 0.676 | 0.199 | 7.42E-17 |
| Endothelial Cell | Cpe | 4 | 8.79E-20 | 0.43979372 | 0.648 | 0.218 | 1.76E-16 |
| Endothelial Cell | Prrx1 | 4 | 1.11E-19 | 0.55075423 | 0.704 | 0.277 | 2.21E-16 |
| Endothelial Cell | Cdh19 | 4 | 2.05E-19 | 1.12888748 | 0.75 | 0.365 | 4.09E-16 |
| Endothelial Cell | Greb1l | 4 | 3.19E-19 | 1.90090558 | 0.537 | 0.143 | 6.38E-16 |
| Endothelial Cell | Tmem51 | 4 | 4.64E-19 | 0.75025703 | 0.75 | 0.311 | 9.28E-16 |
| Endothelial Cell | Sh3gl3 | 4 | 4.65E-19 | 2.27655537 | 0.454 | 0.158 | 9.31E-16 |
| Endothelial Cell | Cacna1d | 4 | 4.70E-19 | 0.84293371 | 0.537 | 0.173 | 9.40E-16 |
| Endothelial Cell | Galnt16 | 4 | 5.72E-19 | 0.25162594 | 0.713 | 0.302 | 1.14E-15 |

| | | | | | | | |
|---|---|---|---|---|---|---|---|
| Endothelial Cell | Lrg1 | 4 | 5.88E-19 | 1.30904381 | 0.704 | 0.351 | 1.18E-15 |
| Endothelial Cell | Mob3b | 4 | 9.04E-19 | 0.82586809 | 0.778 | 0.454 | 1.81E-15 |
| Endothelial Cell | Palm2 | 4 | 1.38E-18 | 0.97454896 | 0.593 | 0.185 | 2.75E-15 |
| Endothelial Cell | Bcat1 | 4 | 1.82E-18 | 1.14466479 | 0.583 | 0.187 | 3.63E-15 |
| Endothelial Cell | Uqcrq | 4 | 6.27E-18 | 0.37543084 | 0.787 | 0.37 | 1.25E-14 |
| Endothelial Cell | Glis3 | 4 | 7.45E-18 | 0.94075881 | 0.787 | 0.353 | 1.49E-14 |
| Endothelial Cell | Ca4 | 4 | 7.93E-18 | 1.30055891 | 0.815 | 0.353 | 1.59E-14 |
| Endothelial Cell | Pcdh7 | 4 | 9.48E-18 | 2.11719042 | 0.593 | 0.222 | 1.90E-14 |
| Endothelial Cell | Lilrb4 | 4 | 1.21E-17 | 1.39431307 | 0.556 | 0.189 | 2.42E-14 |
| Endothelial Cell | Setbp1 | 4 | 1.23E-17 | 0.26494028 | 0.787 | 0.34 | 2.46E-14 |
| Endothelial Cell | Ckm | 4 | 1.27E-17 | 0.88556761 | 0.769 | 0.398 | 2.53E-14 |
| Endothelial Cell | Slc39a8 | 4 | 1.31E-17 | 0.89942043 | 0.75 | 0.369 | 2.61E-14 |
| Endothelial Cell | Pcbp3 | 4 | 1.38E-17 | 0.73246213 | 0.759 | 0.301 | 2.76E-14 |
| Endothelial Cell | Fsd2 | 4 | 1.86E-17 | 0.31663374 | 0.611 | 0.198 | 3.72E-14 |

| | | | | | | | |
|---|---|---|---|---|---|---|---|
| Endothelial Cell | Dsp | 4 | 2.06E-17 | 0.50033682 | 0.843 | 0.35 | 4.11E-14 |
| Endothelial Cell | Bend6 | 4 | 2.11E-17 | 0.29817892 | 0.676 | 0.27 | 4.21E-14 |
| Endothelial Cell | Hk2 | 4 | 2.62E-17 | 0.38838096 | 0.602 | 0.248 | 5.23E-14 |
| Endothelial Cell | Alpk2 | 4 | 2.64E-17 | 0.34350203 | 0.759 | 0.359 | 5.28E-14 |
| Endothelial Cell | Smpx | 4 | 3.21E-17 | 1.10134509 | 0.787 | 0.446 | 6.42E-14 |
| Endothelial Cell | Rhpn2 | 4 | 3.43E-17 | 1.03959159 | 0.444 | 0.16 | 6.85E-14 |
| Endothelial Cell | Cd274 | 4 | 4.91E-17 | 0.75584486 | 0.787 | 0.322 | 9.81E-14 |
| Endothelial Cell | Tox3 | 4 | 5.27E-17 | 1.26180794 | 0.824 | 0.383 | 1.05E-13 |
| Endothelial Cell | Bmp6 | 4 | 5.39E-17 | 1.35682636 | 0.926 | 0.569 | 1.08E-13 |
| Endothelial Cell | Srl | 4 | 5.43E-17 | 1.42005346 | 0.63 | 0.249 | 1.09E-13 |
| Endothelial Cell | Efhc2 | 4 | 7.94E-17 | 1.1866913 | 0.463 | 0.151 | 1.59E-13 |
| Endothelial Cell | Masp1 | 4 | 8.09E-17 | 0.53927919 | 0.519 | 0.185 | 1.62E-13 |
| Endothelial Cell | Hmgcll1 | 4 | 1.10E-16 | 1.42623404 | 0.75 | 0.337 | 2.21E-13 |
| Endothelial Cell | Kcnk3 | 4 | 1.12E-16 | 0.41707731 | 0.435 | 0.096 | 2.24E-13 |

| | | | | | | | |
|---|---|---|---|---|---|---|---|
| Endothelial Cell | Dipk1a | 4 | 1.19E-16 | 0.92474287 | 0.796 | 0.417 | 2.38E-13 |
| Endothelial Cell | Abca8 | 4 | 1.22E-16 | 1.17741114 | 0.481 | 0.088 | 2.44E-13 |
| Endothelial Cell | Plet1 | 4 | 1.46E-16 | 0.785555 | 0.333 | 0.083 | 2.92E-13 |
| Endothelial Cell | Sgcg | 4 | 2.07E-16 | 1.08488342 | 0.528 | 0.168 | 4.13E-13 |
| Endothelial Cell | Hs3st5 | 4 | 2.08E-16 | 1.56247448 | 0.611 | 0.224 | 4.17E-13 |
| Endothelial Cell | Chrdl1 | 4 | 4.37E-16 | 2.28303763 | 0.472 | 0.135 | 8.73E-13 |
| Endothelial Cell | Rbp71 | 4 | 4.52E-16 | 1.54782014 | 0.657 | 0.323 | 9.05E-13 |
| Endothelial Cell | Il1rapl1 | 4 | 4.87E-16 | 0.28652116 | 0.648 | 0.26 | 9.73E-13 |
| Endothelial Cell | Adamts19 | 4 | 5.06E-16 | 0.58703978 | 0.574 | 0.242 | 1.01E-12 |
| Endothelial Cell | Pld5 | 4 | 5.48E-16 | 2.24007963 | 0.38 | 0.13 | 1.10E-12 |
| Endothelial Cell | Veph1 | 4 | 7.06E-16 | 0.67967107 | 0.759 | 0.344 | 1.41E-12 |
| Endothelial Cell | Prox1 | 4 | 8.41E-16 | 0.37012842 | 0.75 | 0.316 | 1.68E-12 |
| Endothelial Cell | Cd300lg | 4 | 8.45E-16 | 1.17630849 | 0.759 | 0.42 | 1.69E-12 |
| Endothelial Cell | Tmem168 | 4 | 9.26E-16 | 1.30875702 | 0.63 | 0.318 | 1.85E-12 |

| | | | | | | | |
|---|---|---|---|---|---|---|---|
| Endothelial Cell | Dpf3 | 4 | 1.05E-15 | 0.82501267 | 0.519 | 0.181 | 2.09E-12 |
| Endothelial Cell | C1qtnf1 | 4 | 1.08E-15 | 1.67579503 | 0.639 | 0.229 | 2.16E-12 |
| Endothelial Cell | Casq2 | 4 | 1.69E-15 | 0.42710067 | 0.583 | 0.217 | 3.39E-12 |
| Endothelial Cell | Ncam2 | 4 | 1.82E-15 | 1.46523104 | 0.583 | 0.212 | 3.64E-12 |
| Endothelial Cell | Pcdh12 | 4 | 2.44E-15 | 0.70266963 | 0.769 | 0.354 | 4.88E-12 |
| Endothelial Cell | Fam78b | 4 | 2.70E-15 | 0.81114512 | 0.806 | 0.443 | 5.40E-12 |
| Endothelial Cell | Rcan2 | 4 | 6.16E-15 | 0.6959557 | 0.759 | 0.286 | 1.23E-11 |
| Endothelial Cell | C1qtnf9 | 4 | 6.71E-15 | 1.47979551 | 0.593 | 0.284 | 1.34E-11 |
| Endothelial Cell | Pcsk6 | 4 | 7.57E-15 | 0.72029201 | 0.75 | 0.359 | 1.51E-11 |
| Endothelial Cell | Ngdn | 4 | 7.95E-15 | 1.00349069 | 0.741 | 0.357 | 1.59E-11 |
| Endothelial Cell | Papss21 | 4 | 9.42E-15 | 1.17640656 | 0.87 | 0.451 | 1.88E-11 |
| Endothelial Cell | Tmem132d | 4 | 1.07E-14 | 0.37220934 | 0.593 | 0.276 | 2.13E-11 |
| Endothelial Cell | AABR07001054.2 | 4 | 1.26E-14 | 1.24117255 | 0.593 | 0.267 | 2.52E-11 |
| Endothelial Cell | Myh7b | 4 | 1.86E-14 | 1.01416155 | 0.704 | 0.352 | 3.72E-11 |

| | | | | | | | |
|---|---|---|---|---|---|---|---|
| Endothelial Cell | Csrp1 | 4 | 2.44E-14 | 0.37351757 | 0.787 | 0.319 | 4.88E-11 |
| Endothelial Cell | Ptpn13 | 4 | 2.49E-14 | 0.50719485 | 0.537 | 0.206 | 4.98E-11 |
| Endothelial Cell | Angptl81 | 4 | 3.79E-14 | 0.29639592 | 0.519 | 0.19 | 7.59E-11 |
| Endothelial Cell | Ccl21 | 4 | 4.45E-14 | 1.3012168 | 0.583 | 0.237 | 8.90E-11 |
| Endothelial Cell | Mafb | 4 | 5.22E-14 | 0.6841897 | 0.444 | 0.14 | 1.04E-10 |
| Endothelial Cell | Mitf | 4 | 5.73E-14 | 0.78295995 | 0.509 | 0.207 | 1.15E-10 |
| Endothelial Cell | Fgf10 | 4 | 5.92E-14 | 1.87420701 | 0.593 | 0.301 | 1.18E-10 |
| Endothelial Cell | Sox5 | 4 | 6.04E-14 | 0.86395791 | 0.694 | 0.321 | 1.21E-10 |
| Endothelial Cell | Ppip5k1 | 4 | 6.90E-14 | 0.52719032 | 0.741 | 0.349 | 1.38E-10 |
| Endothelial Cell | Cadm2 | 4 | 7.71E-14 | 0.47243563 | 0.546 | 0.194 | 1.54E-10 |
| Endothelial Cell | Slc25a13 | 4 | 8.67E-14 | 0.37775123 | 0.731 | 0.429 | 1.73E-10 |
| Endothelial Cell | Myocd | 4 | 1.12E-13 | 1.33392981 | 0.602 | 0.258 | 2.23E-10 |
| Endothelial Cell | Smpd3 | 4 | 1.46E-13 | 0.56883515 | 0.426 | 0.138 | 2.92E-10 |
| Endothelial Cell | Pip5k1b | 4 | 1.58E-13 | 0.33167526 | 0.75 | 0.297 | 3.15E-10 |

| | | | | | | | |
|---|---|---|---|---|---|---|---|
| Endothelial Cell | Xirp2 | 4 | 2.03E-13 | 0.25077473 | 0.815 | 0.44 | 4.06E-10 |
| Endothelial Cell | Apoe | 4 | 2.22E-13 | 0.39939902 | 0.824 | 0.489 | 4.44E-10 |
| Endothelial Cell | Rxfp1 | 4 | 2.46E-13 | 1.06307474 | 0.602 | 0.244 | 4.93E-10 |
| Endothelial Cell | Rgcc | 4 | 2.75E-13 | 1.64706866 | 0.704 | 0.442 | 5.50E-10 |
| Endothelial Cell | Alpl1 | 4 | 2.96E-13 | 0.93872636 | 0.741 | 0.37 | 5.91E-10 |
| Endothelial Cell | Rhobtb1 | 4 | 3.57E-13 | 0.85727864 | 0.694 | 0.332 | 7.13E-10 |
| Endothelial Cell | Engase | 4 | 4.10E-13 | 1.67559108 | 0.528 | 0.239 | 8.20E-10 |
| Endothelial Cell | Dip2c | 4 | 5.66E-13 | 0.56012776 | 0.806 | 0.415 | 1.13E-09 |
| Endothelial Cell | Rem1 | 4 | 6.30E-13 | 0.46908792 | 0.657 | 0.315 | 1.26E-09 |
| Endothelial Cell | Dact2 | 4 | 7.87E-13 | 0.32460421 | 0.519 | 0.215 | 1.57E-09 |
| Endothelial Cell | Slc22a23 | 4 | 9.98E-13 | 0.58812122 | 0.713 | 0.366 | 2.00E-09 |
| Endothelial Cell | Prag1 | 4 | 1.34E-12 | 1.068917 | 0.861 | 0.546 | 2.69E-09 |
| Endothelial Cell | Lyve1 | 4 | 1.35E-12 | 1.40741341 | 0.352 | 0.071 | 2.69E-09 |
| Endothelial Cell | Nudt4 | 4 | 2.15E-12 | 0.40795687 | 0.713 | 0.373 | 4.29E-09 |

| | | | | | | | |
|---|---|---|---|---|---|---|---|
| Endothelial Cell | Rnf207 | 4 | 2.52E-12 | 0.78056633 | 0.491 | 0.238 | 5.03E-09 |
| Endothelial Cell | Pde4b | 4 | 2.71E-12 | 0.41641079 | 0.75 | 0.34 | 5.41E-09 |
| Endothelial Cell | Trim63 | 4 | 2.96E-12 | 0.25120848 | 0.574 | 0.259 | 5.92E-09 |
| Endothelial Cell | Coq8a | 4 | 3.49E-12 | 1.14031429 | 0.602 | 0.278 | 6.98E-09 |
| Endothelial Cell | Camk1d | 4 | 3.56E-12 | 0.28889945 | 0.704 | 0.304 | 7.13E-09 |
| Endothelial Cell | Adora2a | 4 | 4.18E-12 | 0.66861155 | 0.667 | 0.34 | 8.36E-09 |
| Endothelial Cell | Satb2 | 4 | 5.90E-12 | 1.27399965 | 0.537 | 0.247 | 1.18E-08 |
| Endothelial Cell | Aff2 | 4 | 5.91E-12 | 0.83895597 | 0.676 | 0.288 | 1.18E-08 |
| Endothelial Cell | Tpm2 | 4 | 9.75E-12 | 0.5084015 | 0.593 | 0.318 | 1.95E-08 |
| Endothelial Cell | Col4a3 | 4 | 1.24E-11 | 0.41946373 | 0.815 | 0.517 | 2.47E-08 |
| Endothelial Cell | C1qtnf7 | 4 | 1.41E-11 | 0.6339887 | 0.704 | 0.365 | 2.81E-08 |
| Endothelial Cell | Iqgap31 | 4 | 2.86E-11 | 1.39253539 | 0.528 | 0.217 | 5.72E-08 |
| Endothelial Cell | Fmo2 | 4 | 3.50E-11 | 0.94049473 | 0.648 | 0.353 | 7.00E-08 |
| Endothelial Cell | Itgb81 | 4 | 3.73E-11 | 0.34865439 | 0.769 | 0.39 | 7.47E-08 |

| | | | | | | | |
|---|---|---|---|---|---|---|---|
| Endothelial Cell | Arhgef26 | 4 | 4.41E-11 | 0.75906395 | 0.685 | 0.359 | 8.82E-08 |
| Endothelial Cell | Sod2 | 4 | 4.48E-11 | 1.19940473 | 0.398 | 0.136 | 8.95E-08 |
| Endothelial Cell | Sacs | 4 | 5.69E-11 | 0.51569139 | 0.667 | 0.289 | 1.14E-07 |
| Endothelial Cell | Tspan18 | 4 | 6.02E-11 | 0.78826927 | 0.741 | 0.338 | 1.20E-07 |
| Endothelial Cell | Krt1 | 4 | 7.32E-11 | 3.44564578 | 0.491 | 0.185 | 1.46E-07 |
| Endothelial Cell | Sema3c | 4 | 7.89E-11 | 0.51089897 | 0.5 | 0.243 | 1.58E-07 |
| Endothelial Cell | Smox | 4 | 8.25E-11 | 0.88717935 | 0.741 | 0.413 | 1.65E-07 |
| Endothelial Cell | Kcnq5 | 4 | 8.57E-11 | 1.06886748 | 0.37 | 0.062 | 1.71E-07 |
| Endothelial Cell | Ano5 | 4 | 8.71E-11 | 0.73868503 | 0.509 | 0.157 | 1.74E-07 |
| Endothelial Cell | Adam19 | 4 | 8.88E-11 | 0.38895831 | 0.769 | 0.39 | 1.78E-07 |
| Endothelial Cell | Gnao1 | 4 | 9.97E-11 | 1.24320456 | 0.806 | 0.493 | 1.99E-07 |
| Endothelial Cell | Pcdh19 | 4 | 1.23E-10 | 0.81941403 | 0.852 | 0.483 | 2.46E-07 |
| Endothelial Cell | Rec114 | 4 | 1.45E-10 | 0.59603487 | 0.676 | 0.328 | 2.91E-07 |
| Endothelial Cell | Efcab6 | 4 | 1.50E-10 | 1.92692688 | 0.5 | 0.183 | 3.01E-07 |

| | | | | | | | |
|---|---|---|---|---|---|---|---|
| Endothelial Cell | Notch3 | 4 | 2.16E-10 | 0.88701225 | 0.657 | 0.372 | 4.32E-07 |
| Endothelial Cell | Sh3kbp1 | 4 | 2.27E-10 | 0.30356471 | 0.63 | 0.281 | 4.54E-07 |
| Endothelial Cell | Rgs6 | 4 | 2.34E-10 | 1.21779939 | 0.463 | 0.108 | 4.69E-07 |
| Endothelial Cell | Ryr2 | 4 | 2.51E-10 | 0.63567309 | 0.926 | 0.597 | 5.01E-07 |
| Endothelial Cell | Tmsb4x1 | 4 | 3.18E-10 | 0.83090295 | 0.889 | 0.596 | 6.37E-07 |
| Endothelial Cell | RGD156335 41 | 4 | 3.64E-10 | 0.33302427 | 0.574 | 0.276 | 7.29E-07 |
| Endothelial Cell | Cxcl12 | 4 | 4.36E-10 | 0.70407598 | 0.907 | 0.609 | 8.72E-07 |
| Endothelial Cell | Acer2 | 4 | 4.46E-10 | 0.82826089 | 0.611 | 0.287 | 8.92E-07 |
| Endothelial Cell | Trdn | 4 | 4.88E-10 | 0.71973984 | 0.639 | 0.309 | 9.76E-07 |
| Endothelial Cell | Gda | 4 | 5.46E-10 | 0.47376133 | 0.815 | 0.473 | 1.09E-06 |
| Endothelial Cell | Spon1 | 4 | 5.70E-10 | 0.71296644 | 0.593 | 0.323 | 1.14E-06 |
| Endothelial Cell | Dcn | 4 | 7.60E-10 | 0.45634363 | 0.806 | 0.44 | 1.52E-06 |
| Endothelial Cell | AABR07049 085.1 | 4 | 1.22E-09 | 0.45412758 | 0.722 | 0.351 | 2.43E-06 |
| Endothelial Cell | Atp5f1e1 | 4 | 1.63E-09 | 0.26588047 | 0.815 | 0.404 | 3.25E-06 |

| | | | | | | | |
|---|---|---|---|---|---|---|---|
| Endothelial Cell | Ppp1r3a | 4 | 1.90E-09 | 0.44145162 | 0.639 | 0.336 | 3.80E-06 |
| Endothelial Cell | Lbh | 4 | 2.26E-09 | 0.37308248 | 0.694 | 0.381 | 4.52E-06 |
| Endothelial Cell | Reep1 | 4 | 2.34E-09 | 0.29182942 | 0.648 | 0.375 | 4.67E-06 |
| Endothelial Cell | Bnc21 | 4 | 2.60E-09 | 0.41404157 | 0.62 | 0.327 | 5.21E-06 |
| Endothelial Cell | Fabp4 | 4 | 3.02E-09 | 0.71277851 | 0.935 | 0.629 | 6.04E-06 |
| Endothelial Cell | Ubash3b | 4 | 3.03E-09 | 0.55579973 | 0.769 | 0.473 | 6.06E-06 |
| Endothelial Cell | Adamts2 | 4 | 3.80E-09 | 0.52234967 | 0.574 | 0.314 | 7.60E-06 |
| Endothelial Cell | Rimbp2 | 4 | 4.30E-09 | 0.3016881 | 0.676 | 0.413 | 8.59E-06 |
| Endothelial Cell | Atp5mc1 | 4 | 6.37E-09 | 0.2507214 | 0.722 | 0.417 | 1.27E-05 |
| Endothelial Cell | Pde7b | 4 | 6.89E-09 | 0.80987305 | 0.778 | 0.409 | 1.38E-05 |
| Endothelial Cell | Sparcl1 | 4 | 1.43E-08 | 0.6637312 | 0.759 | 0.46 | 2.87E-05 |
| Endothelial Cell | Pi16 | 4 | 1.67E-08 | 0.42742838 | 0.63 | 0.355 | 3.34E-05 |
| Endothelial Cell | Ttll7 | 4 | 1.71E-08 | 0.64415604 | 0.704 | 0.399 | 3.42E-05 |
| Endothelial Cell | Cmtm8 | 4 | 5.65E-08 | 0.93888389 | 0.787 | 0.523 | 0.00011303 |

| | | | | | | | |
|---|---|---|---|---|---|---|---|
| Endothelial Cell | Eya2 | 4 | 6.77E-08 | 0.76564886 | 0.704 | 0.423 | 0.00013542 |
| Endothelial Cell | Steap4 | 4 | 1.13E-07 | 0.61826645 | 0.528 | 0.268 | 0.00022551 |
| Endothelial Cell | Pparg | 4 | 2.01E-07 | 0.55050137 | 0.713 | 0.448 | 0.00040141 |
| Endothelial Cell | Cacnb2 | 4 | 2.49E-07 | 0.3549668 | 0.778 | 0.432 | 0.00049784 |
| Endothelial Cell | Clec1a | 4 | 5.37E-07 | 0.47405929 | 0.685 | 0.409 | 0.00107413 |
| Endothelial Cell | Myom2 | 4 | 6.24E-07 | 0.36332184 | 0.713 | 0.379 | 0.00124885 |
| Endothelial Cell | Med12l | 4 | 7.58E-07 | 0.54654105 | 0.389 | 0.098 | 0.00151571 |
| Endothelial Cell | Dipk2b | 4 | 8.32E-07 | 0.84625791 | 0.759 | 0.5 | 0.00166385 |
| Endothelial Cell | Mtss1 | 4 | 1.07E-06 | 0.76157303 | 0.852 | 0.552 | 0.00213075 |
| Endothelial Cell | Celf2 | 4 | 4.05E-06 | 0.31237112 | 0.731 | 0.423 | 0.00809342 |
| Endothelial Cell | Cav1 | 4 | 1.87E-05 | 0.35301565 | 0.87 | 0.575 | 0.03731928 |
| Endothelial Cell | Insl3 | 5 | 3.38E-131 | 7.15067552 | 0.543 | 0.002 | 6.75E-128 |
| Endothelial Cell | Kif20a | 5 | 5.04E-120 | 6.70012605 | 0.479 | 0 | 1.01E-116 |
| Endothelial Cell | Brinp1 | 5 | 2.34E-118 | 5.40333923 | 0.532 | 0.005 | 4.69E-115 |

| | | | | | | | |
|---|---|---|---|---|---|---|---|
| Endothelial Cell | Slco6c1 | 5 | 6.36E-112 | 3.96410435 | 0.5 | 0.005 | 1.27E-108 |
| Endothelial Cell | Clec4e | 5 | 5.81E-110 | 4.21412799 | 0.511 | 0.005 | 1.16E-106 |
| Endothelial Cell | Aim2 | 5 | 7.40E-107 | 0.39054886 | 0.5 | 0.006 | 1.48E-103 |
| Endothelial Cell | Trpc4 | 5 | 5.02E-103 | 3.85750703 | 0.511 | 0.008 | 1.00E-99 |
| Endothelial Cell | Kcna7 | 5 | 1.54E-102 | 6.55712175 | 0.543 | 0.015 | 3.08E-99 |
| Endothelial Cell | Iqcf1 | 5 | 1.27E-100 | 2.94433896 | 0.543 | 0.016 | 2.55E-97 |
| Endothelial Cell | Cdh3 | 5 | 1.43E-99 | 1.65915634 | 0.468 | 0.005 | 2.87E-96 |
| Endothelial Cell | Fam151a | 5 | 1.23E-98 | 0.59171567 | 0.5 | 0.01 | 2.47E-95 |
| Endothelial Cell | Ubash3a | 5 | 1.30E-92 | 3.57876774 | 0.543 | 0.02 | 2.60E-89 |
| Endothelial Cell | Elavl2 | 5 | 8.60E-84 | 5.78438677 | 0.553 | 0.027 | 1.72E-80 |
| Endothelial Cell | Spata31d1 | 5 | 4.04E-83 | 5.95507736 | 0.532 | 0.006 | 8.07E-80 |
| Endothelial Cell | Snx31 | 5 | 1.06E-81 | 5.76455428 | 0.553 | 0.029 | 2.11E-78 |
| Endothelial Cell | Adamts20 | 5 | 2.00E-79 | 3.43202033 | 0.553 | 0.03 | 4.01E-76 |
| Endothelial Cell | Selp | 5 | 1.33E-67 | 6.01396608 | 0.511 | 0.049 | 2.66E-64 |

| | | | | | | | |
|---|---|---|---|---|---|---|---|
| Endothelial Cell | S100a9 | 5 | 1.67E-61 | 0.39339407 | 0.5 | 0.015 | 3.34E-58 |
| Endothelial Cell | Pls1 | 5 | 1.89E-60 | 2.81547463 | 0.489 | 0.035 | 3.79E-57 |
| Endothelial Cell | Hyi | 5 | 1.00E-59 | 0.53011579 | 0.543 | 0.05 | 2.00E-56 |
| Endothelial Cell | Soat2 | 5 | 4.24E-59 | 3.36395975 | 0.553 | 0.049 | 8.48E-56 |
| Endothelial Cell | Cyp4f181 | 5 | 2.24E-51 | 4.595329 | 0.532 | 0.064 | 4.47E-48 |
| Endothelial Cell | AABR07033925.1 | 5 | 8.73E-51 | 1.00943233 | 0.532 | 0.053 | 1.75E-47 |
| Endothelial Cell | Syt15 | 5 | 2.39E-50 | 2.86020891 | 0.553 | 0.06 | 4.78E-47 |
| Endothelial Cell | Myoz3 | 5 | 4.44E-42 | 1.74036164 | 0.479 | 0.057 | 8.88E-39 |
| Endothelial Cell | Tmem163 | 5 | 5.52E-42 | 1.29101114 | 0.543 | 0.024 | 1.10E-38 |
| Endothelial Cell | Pdgfc | 5 | 3.02E-38 | 4.08811148 | 0.574 | 0.071 | 6.04E-35 |
| Endothelial Cell | Exoc3l2 | 5 | 3.51E-38 | 3.98609419 | 0.606 | 0.091 | 7.02E-35 |
| Endothelial Cell | Mt31 | 5 | 9.33E-38 | 1.24763218 | 0.543 | 0.072 | 1.87E-34 |
| Endothelial Cell | Prkcq | 5 | 1.94E-37 | 0.61523318 | 0.543 | 0.036 | 3.87E-34 |
| Endothelial Cell | Nmnat2 | 5 | 2.47E-37 | 2.30599847 | 0.564 | 0.098 | 4.94E-34 |

| | | | | | | | |
|---|---|---|---|---|---|---|---|
| Endothelial Cell | AABR07005844.1 | 5 | 2.29E-36 | 6.21382007 | 0.351 | 0.043 | 4.58E-33 |
| Endothelial Cell | Chst9 | 5 | 1.68E-35 | 3.01200835 | 0.553 | 0.11 | 3.35E-32 |
| Endothelial Cell | Gabre | 5 | 4.86E-35 | 0.76375082 | 0.511 | 0.063 | 9.71E-32 |
| Endothelial Cell | Cyp2e11 | 5 | 1.50E-34 | 0.96955701 | 0.511 | 0.077 | 3.01E-31 |
| Endothelial Cell | Sele | 5 | 1.19E-33 | 4.99652307 | 0.596 | 0.148 | 2.39E-30 |
| Endothelial Cell | Vwf1 | 5 | 5.92E-33 | 2.49996137 | 0.936 | 0.595 | 1.18E-29 |
| Endothelial Cell | Nalcn | 5 | 8.75E-31 | 1.6138453 | 0.553 | 0.033 | 1.75E-27 |
| Endothelial Cell | Mapk10 | 5 | 3.86E-29 | 2.27638845 | 0.532 | 0.066 | 7.71E-26 |
| Endothelial Cell | Fhad1 | 5 | 3.00E-28 | 2.14935852 | 0.543 | 0.119 | 6.00E-25 |
| Endothelial Cell | AABR07005821.1 | 5 | 3.00E-27 | 5.06581976 | 0.628 | 0.137 | 6.00E-24 |
| Endothelial Cell | Thbs1 | 5 | 3.37E-27 | 1.68215258 | 0.553 | 0.055 | 6.74E-24 |
| Endothelial Cell | Tnfrsf11b | 5 | 2.05E-26 | 2.60789423 | 0.479 | 0.011 | 4.10E-23 |
| Endothelial Cell | Kif22 | 5 | 9.83E-26 | 2.53706331 | 0.479 | 0.035 | 1.97E-22 |
| Endothelial Cell | Adam23 | 5 | 1.34E-24 | 0.95765171 | 0.564 | 0.125 | 2.69E-21 |

| | | | | | | | |
|---|---|---|---|---|---|---|---|
| Endothelial Cell | Cysltr1 | 5 | 1.35E-24 | 4.32802471 | 0.596 | 0.152 | 2.70E-21 |
| Endothelial Cell | Sh3gl31 | 5 | 1.49E-24 | 2.130125 | 0.553 | 0.153 | 2.98E-21 |
| Endothelial Cell | Ehd4 | 5 | 1.21E-23 | 2.58796421 | 0.872 | 0.402 | 2.42E-20 |
| Endothelial Cell | Tbx21 | 5 | 1.44E-23 | 3.30580075 | 0.543 | 0.047 | 2.87E-20 |
| Endothelial Cell | Ldb2 | 5 | 1.94E-23 | 2.55247475 | 0.809 | 0.408 | 3.89E-20 |
| Endothelial Cell | Col11a11 | 5 | 1.99E-23 | 1.84228427 | 0.553 | 0.105 | 3.98E-20 |
| Endothelial Cell | Csf1r | 5 | 3.21E-23 | 0.6177795 | 0.521 | 0.031 | 6.41E-20 |
| Endothelial Cell | Thsd7a | 5 | 5.48E-23 | 1.9706892 | 0.904 | 0.582 | 1.10E-19 |
| Endothelial Cell | Fbxo40 | 5 | 6.48E-23 | 1.21309677 | 0.479 | 0.05 | 1.30E-19 |
| Endothelial Cell | Ptprn2 | 5 | 7.70E-23 | 1.36840386 | 0.511 | 0.079 | 1.54E-19 |
| Endothelial Cell | Nuak1 | 5 | 9.28E-23 | 1.98057556 | 0.894 | 0.521 | 1.86E-19 |
| Endothelial Cell | Pgm5 | 5 | 7.42E-22 | 2.09657416 | 0.894 | 0.553 | 1.48E-18 |
| Endothelial Cell | Vcam1 | 5 | 1.38E-21 | 3.14211034 | 0.649 | 0.27 | 2.75E-18 |
| Endothelial Cell | Fbxw10 | 5 | 3.63E-21 | 0.92756741 | 0.543 | 0.162 | 7.27E-18 |

| | | | | | | | |
|---|---|---|---|---|---|---|---|
| Endothelial Cell | Ptn | 5 | 4.65E-21 | 3.46950522 | 0.702 | 0.239 | 9.30E-18 |
| Endothelial Cell | LOC1025513561 | 5 | 6.79E-21 | 2.19238818 | 0.543 | 0.083 | 1.36E-17 |
| Endothelial Cell | Adgrg6 | 5 | 1.24E-20 | 2.95063369 | 0.553 | 0.14 | 2.47E-17 |
| Endothelial Cell | AABR07034940.2 | 5 | 1.50E-20 | 1.01490585 | 0.553 | 0.146 | 3.00E-17 |
| Endothelial Cell | Syt9 | 5 | 2.88E-20 | 1.26655564 | 0.532 | 0.061 | 5.77E-17 |
| Endothelial Cell | Rasa4 | 5 | 7.27E-20 | 1.83033422 | 0.915 | 0.663 | 1.45E-16 |
| Endothelial Cell | Anln1 | 5 | 1.81E-19 | 3.22276646 | 0.479 | 0.094 | 3.62E-16 |
| Endothelial Cell | Col28a1 | 5 | 1.38E-18 | 1.96630761 | 0.553 | 0.171 | 2.75E-15 |
| Endothelial Cell | Krt11 | 5 | 6.00E-18 | 1.22327248 | 0.532 | 0.185 | 1.20E-14 |
| Endothelial Cell | Gabbr2 | 5 | 1.02E-17 | 3.32125011 | 0.489 | 0.008 | 2.03E-14 |
| Endothelial Cell | AABR07004228.1 | 5 | 3.25E-17 | 2.97622682 | 0.596 | 0.276 | 6.51E-14 |
| Endothelial Cell | Maob | 5 | 5.83E-17 | 2.21905625 | 0.723 | 0.406 | 1.17E-13 |
| Endothelial Cell | Eln | 5 | 1.09E-16 | 1.97623372 | 0.734 | 0.385 | 2.17E-13 |
| Endothelial Cell | Mctp1 | 5 | 2.44E-16 | 4.8550043 | 0.606 | 0.196 | 4.88E-13 |

| | | | | | | | |
|---|---|---|---|---|---|---|---|
| Endothelial Cell | AABR07050449.1 | 5 | 2.49E-16 | 1.84212069 | 0.532 | 0.156 | 4.98E-13 |
| Endothelial Cell | Adamts14 | 5 | 2.71E-16 | 1.2889535 | 0.543 | 0.225 | 5.41E-13 |
| Endothelial Cell | Dtx1 | 5 | 2.94E-16 | 2.57162143 | 0.553 | 0.166 | 5.87E-13 |
| Endothelial Cell | Smoc11 | 5 | 3.86E-16 | 1.5134352 | 0.532 | 0.113 | 7.72E-13 |
| Endothelial Cell | Npnt | 5 | 4.20E-16 | 2.46240202 | 0.543 | 0.201 | 8.41E-13 |
| Endothelial Cell | Txk | 5 | 7.59E-16 | 2.51778776 | 0.553 | 0.181 | 1.52E-12 |
| Endothelial Cell | Cux2 | 5 | 1.35E-15 | 1.87891829 | 0.553 | 0.241 | 2.69E-12 |
| Endothelial Cell | Stab1 | 5 | 3.65E-15 | 3.23623617 | 0.638 | 0.251 | 7.30E-12 |
| Endothelial Cell | Cttnbp2 | 5 | 1.22E-14 | 2.25252301 | 0.755 | 0.284 | 2.44E-11 |
| Endothelial Cell | Oprd1 | 5 | 1.58E-14 | 3.1985654 | 0.564 | 0.205 | 3.16E-11 |
| Endothelial Cell | Abca1 | 5 | 3.48E-14 | 2.38063011 | 0.681 | 0.282 | 6.96E-11 |
| Endothelial Cell | Ptk2b | 5 | 3.68E-14 | 3.13167948 | 0.628 | 0.353 | 7.35E-11 |
| Endothelial Cell | Abi3bp | 5 | 8.01E-14 | 2.40941766 | 0.66 | 0.337 | 1.60E-10 |
| Endothelial Cell | Dagla | 5 | 8.81E-14 | 1.98918637 | 0.564 | 0.218 | 1.76E-10 |

| | | | | | | | |
|---|---|---|---|---|---|---|---|
| Endothelial Cell | Trmt9b | 5 | 1.14E-13 | 2.99306796 | 0.628 | 0.357 | 2.28E-10 |
| Endothelial Cell | Cadm3 | 5 | 1.21E-13 | 2.87885578 | 0.606 | 0.174 | 2.41E-10 |
| Endothelial Cell | Rasl12 | 5 | 1.24E-13 | 1.90651181 | 0.553 | 0.141 | 2.48E-10 |
| Endothelial Cell | Ror11 | 5 | 1.25E-13 | 1.99649579 | 0.574 | 0.143 | 2.51E-10 |
| Endothelial Cell | Gnb3 | 5 | 1.93E-13 | 1.76785969 | 0.564 | 0.278 | 3.86E-10 |
| Endothelial Cell | Musk | 5 | 3.68E-13 | 0.83887239 | 0.553 | 0.234 | 7.36E-10 |
| Endothelial Cell | Susd4 | 5 | 4.17E-13 | 2.25167114 | 0.543 | 0.252 | 8.34E-10 |
| Endothelial Cell | Chn2 | 5 | 1.79E-12 | 1.54905577 | 0.755 | 0.388 | 3.57E-09 |
| Endothelial Cell | Ntn1 | 5 | 2.65E-12 | 1.11282552 | 0.872 | 0.505 | 5.30E-09 |
| Endothelial Cell | Skap1 | 5 | 3.27E-12 | 0.43038649 | 0.5 | 0.078 | 6.53E-09 |
| Endothelial Cell | Rgs17 | 5 | 3.97E-12 | 2.37080016 | 0.457 | 0.114 | 7.95E-09 |
| Endothelial Cell | Pde8b | 5 | 7.37E-12 | 0.94633524 | 0.553 | 0.165 | 1.47E-08 |
| Endothelial Cell | Ltbp2 | 5 | 8.54E-12 | 0.6615277 | 0.532 | 0.187 | 1.71E-08 |
| Endothelial Cell | Ubash3b1 | 5 | 9.45E-12 | 1.93684272 | 0.755 | 0.478 | 1.89E-08 |

| | | | | | | | |
|---|---|---|---|---|---|---|---|
| Endothelial Cell | Ifitm10 | 5 | 9.75E-12 | 1.85478358 | 0.617 | 0.204 | 1.95E-08 |
| Endothelial Cell | Dact21 | 5 | 1.11E-11 | 0.404791 | 0.521 | 0.219 | 2.23E-08 |
| Endothelial Cell | Plekha71 | 5 | 2.41E-11 | 1.53284711 | 0.66 | 0.383 | 4.82E-08 |
| Endothelial Cell | Abo1 | 5 | 3.40E-11 | 1.20792736 | 0.521 | 0.15 | 6.79E-08 |
| Endothelial Cell | Grid2 | 5 | 4.49E-11 | 1.27964145 | 0.436 | 0.091 | 8.98E-08 |
| Endothelial Cell | Pstpip11 | 5 | 6.20E-11 | 0.62198175 | 0.543 | 0.238 | 1.24E-07 |
| Endothelial Cell | Brip1 | 5 | 6.38E-11 | 0.91854199 | 0.553 | 0.252 | 1.28E-07 |
| Endothelial Cell | Actn1 | 5 | 1.52E-10 | 2.0121422 | 0.691 | 0.426 | 3.05E-07 |
| Endothelial Cell | Kcnk21 | 5 | 1.66E-10 | 1.98484357 | 0.553 | 0.266 | 3.32E-07 |
| Endothelial Cell | Mctp2 | 5 | 2.01E-10 | 2.01298993 | 0.543 | 0.162 | 4.02E-07 |
| Endothelial Cell | Sorcs11 | 5 | 2.54E-10 | 0.91478413 | 0.553 | 0.302 | 5.07E-07 |
| Endothelial Cell | Slit31 | 5 | 2.81E-10 | 0.93905107 | 0.021 | 0.341 | 5.61E-07 |
| Endothelial Cell | C1qtnf11 | 5 | 1.05E-09 | 1.28871246 | 0.564 | 0.241 | 2.10E-06 |
| Endothelial Cell | Cp | 5 | 1.20E-09 | 3.99114668 | 0.394 | 0.136 | 2.40E-06 |

| | | | | | | | |
|---|---|---|---|---|---|---|---|
| Endothelial Cell | Dcdc5 | 5 | 1.89E-09 | 1.24455546 | 0.404 | 0.014 | 3.78E-06 |
| Endothelial Cell | Slfn4 | 5 | 2.10E-09 | 2.26988527 | 0.691 | 0.374 | 4.19E-06 |
| Endothelial Cell | Prph | 5 | 2.79E-09 | 2.80464584 | 0.553 | 0.25 | 5.58E-06 |
| Endothelial Cell | Mid1 | 5 | 5.48E-09 | 2.44507856 | 0.521 | 0.127 | 1.10E-05 |
| Endothelial Cell | Itgb4 | 5 | 6.54E-09 | 3.18003422 | 0.468 | 0.194 | 1.31E-05 |
| Endothelial Cell | Hs3st1 | 5 | 6.91E-09 | 1.79370204 | 0.479 | 0.223 | 1.38E-05 |
| Endothelial Cell | Ifit3 | 5 | 8.67E-09 | 2.20691705 | 0.564 | 0.255 | 1.73E-05 |
| Endothelial Cell | Icam1 | 5 | 9.10E-09 | 1.77028496 | 0.585 | 0.234 | 1.82E-05 |
| Endothelial Cell | Gria4 | 5 | 1.09E-08 | 1.71597252 | 0.553 | 0.242 | 2.18E-05 |
| Endothelial Cell | Sv2c | 5 | 1.73E-08 | 0.66210876 | 0.34 | 0.066 | 3.46E-05 |
| Endothelial Cell | Art31 | 5 | 6.72E-08 | 1.08102621 | 0.543 | 0.15 | 0.00013449 |
| Endothelial Cell | Slc16a101 | 5 | 7.96E-08 | 1.00310223 | 0.543 | 0.119 | 0.00015921 |
| Endothelial Cell | Plod2 | 5 | 8.95E-08 | 1.20346615 | 0.66 | 0.383 | 0.00017904 |
| Endothelial Cell | PCOLCE2 | 5 | 9.30E-08 | 0.97362418 | 0.394 | 0.083 | 0.00018595 |

| | | | | | | | |
|---|---|---|---|---|---|---|---|
| Endothelial Cell | Manf | 5 | 9.64E-08 | 0.31898257 | 0.543 | 0.189 | 0.00019286 |
| Endothelial Cell | Fstl1 | 5 | 1.43E-07 | 1.52262397 | 0.649 | 0.355 | 0.00028671 |
| Endothelial Cell | Adgrb3 | 5 | 1.46E-07 | 1.63825997 | 0.543 | 0.291 | 0.00029282 |
| Endothelial Cell | Tmod1 | 5 | 2.43E-07 | 0.73031436 | 0.532 | 0.242 | 0.00048518 |
| Endothelial Cell | Pkhd1l11 | 5 | 4.09E-07 | 1.19025506 | 0.723 | 0.38 | 0.00081813 |
| Endothelial Cell | LOC103694210 | 5 | 4.22E-07 | 1.63642357 | 0.628 | 0.302 | 0.00084472 |
| Endothelial Cell | Acta2 | 5 | 4.65E-07 | 0.91298145 | 0.5 | 0.125 | 0.00092932 |
| Endothelial Cell | Tmcc3 | 5 | 7.70E-07 | 1.08214733 | 0.766 | 0.49 | 0.00154059 |
| Endothelial Cell | Pmepa1 | 5 | 8.24E-07 | 1.06009556 | 0.574 | 0.32 | 0.00164751 |
| Endothelial Cell | Clmp | 5 | 1.19E-06 | 0.53349036 | 0.585 | 0.311 | 0.00237128 |
| Endothelial Cell | Tango2 | 5 | 1.61E-06 | 0.82996592 | 0.606 | 0.283 | 0.00322646 |
| Endothelial Cell | Ngf | 5 | 4.94E-06 | 1.49912562 | 0.574 | 0.296 | 0.0098794 |
| Endothelial Cell | Trabd2b1 | 5 | 7.68E-06 | 1.17631321 | 0.617 | 0.313 | 0.01536248 |
| Endothelial Cell | Gpc6 | 5 | 1.02E-05 | 0.94509115 | 0.702 | 0.437 | 0.02047699 |

| | | | | | | | |
|---|---|---|---|---|---|---|---|
| Endothelial Cell | Corin | 5 | 1.04E-05 | 0.44133652 | 0.574 | 0.286 | 0.02073059 |
| Endothelial Cell | Arhgef371 | 5 | 1.18E-05 | 1.50463498 | 0.479 | 0.19 | 0.02364881 |
| Endothelial Cell | Slc9a9 | 5 | 1.32E-05 | 0.40402272 | 0.617 | 0.307 | 0.02646598 |
| Endothelial Cell | Fgf11 | 5 | 2.02E-05 | 0.99430915 | 0.532 | 0.198 | 0.04033536 |
| Endothelial Cell | Maoa | 5 | 2.08E-05 | 0.32749306 | 0.585 | 0.279 | 0.04169118 |
| Endothelial Cell | AABR07013085.1 | 6 | 1.32E-53 | 6.82258903 | 0.321 | 0.011 | 2.64E-50 |
| Endothelial Cell | AABR07044562.1 | 6 | 1.29E-52 | 4.34361562 | 0.393 | 0.005 | 2.57E-49 |
| Endothelial Cell | Ptchd4 | 6 | 5.23E-42 | 5.53951961 | 0.452 | 0.053 | 1.05E-38 |
| Endothelial Cell | Gadd45b | 6 | 7.09E-31 | 2.85955639 | 0.452 | 0.063 | 1.42E-27 |
| Endothelial Cell | Col9a1 | 6 | 1.10E-28 | 0.68728031 | 0.31 | 0.032 | 2.19E-25 |
| Endothelial Cell | Nebl1 | 6 | 5.56E-28 | 1.85739188 | 0.976 | 0.58 | 1.11E-24 |
| Endothelial Cell | Il1rn | 6 | 8.13E-27 | 1.4259999 | 0.417 | 0.028 | 1.63E-23 |
| Endothelial Cell | Klhdc8a1 | 6 | 5.71E-24 | 1.27016491 | 0.369 | 0.056 | 1.14E-20 |
| Endothelial Cell | Scube3 | 6 | 2.61E-22 | 6.52341706 | 0.286 | 0.035 | 5.23E-19 |

| | | | | | | | |
|---|---|---|---|---|---|---|---|
| Endothelial Cell | Sulf1 | 6 | 3.34E-22 | 2.70927556 | 0.845 | 0.509 | 6.67E-19 |
| Endothelial Cell | Tacr1 | 6 | 8.43E-22 | 3.34459123 | 0.345 | 0.049 | 1.69E-18 |
| Endothelial Cell | Chn21 | 6 | 2.12E-21 | 2.3017074 | 0.857 | 0.384 | 4.24E-18 |
| Endothelial Cell | Ccl61 | 6 | 2.32E-21 | 0.95624502 | 0.417 | 0.084 | 4.64E-18 |
| Endothelial Cell | Ptger31 | 6 | 5.42E-21 | 0.5115513 | 0.357 | 0.054 | 1.08E-17 |
| Endothelial Cell | St6galnac31 | 6 | 6.32E-20 | 2.12303023 | 0.857 | 0.441 | 1.26E-16 |
| Endothelial Cell | Tmem1001 | 6 | 7.73E-19 | 4.36427731 | 0.643 | 0.258 | 1.55E-15 |
| Endothelial Cell | Tceal71 | 6 | 2.59E-18 | 0.71642172 | 0.345 | 0.043 | 5.17E-15 |
| Endothelial Cell | Ramp1 | 6 | 2.07E-16 | 1.65918863 | 0.452 | 0.118 | 4.15E-13 |
| Endothelial Cell | Hck | 6 | 5.47E-16 | 3.07533182 | 0.381 | 0.098 | 1.09E-12 |
| Endothelial Cell | LOC100361087 | 6 | 1.29E-15 | 1.85521023 | 0.821 | 0.367 | 2.57E-12 |
| Endothelial Cell | Tgfb21 | 6 | 3.57E-14 | 3.05708272 | 0.738 | 0.367 | 7.14E-11 |
| Endothelial Cell | Epha4 | 6 | 5.15E-12 | 2.27811989 | 0.726 | 0.357 | 1.03E-08 |
| Endothelial Cell | Slc44a51 | 6 | 7.33E-12 | 1.46728566 | 0.44 | 0.153 | 1.47E-08 |

| | | | | | | | |
|---|---|---|---|---|---|---|---|
| Endothelial Cell | Dgkg1 | 6 | 3.82E-11 | 1.31594469 | 0.429 | 0.104 | 7.63E-08 |
| Endothelial Cell | Kcnt21 | 6 | 4.95E-11 | 2.52399272 | 0.667 | 0.377 | 9.90E-08 |
| Endothelial Cell | Rgs7 | 6 | 2.81E-10 | 3.14160688 | 0.393 | 0.133 | 5.62E-07 |
| Endothelial Cell | Kcnn4 | 6 | 2.96E-09 | 1.57737383 | 0.488 | 0.185 | 5.93E-06 |
| Endothelial Cell | Ccser1 | 6 | 4.25E-09 | 1.36286223 | 0.762 | 0.456 | 8.50E-06 |
| Endothelial Cell | Pcsk5 | 6 | 8.05E-09 | 2.92403557 | 0.583 | 0.323 | 1.61E-05 |
| Endothelial Cell | Nr4a31 | 6 | 1.93E-08 | 2.18331304 | 0.452 | 0.102 | 3.87E-05 |
| Endothelial Cell | Fbln51 | 6 | 6.27E-08 | 1.90062505 | 0.679 | 0.421 | 0.00012545 |
| Endothelial Cell | Eln1 | 6 | 2.73E-07 | 1.10381891 | 0.702 | 0.39 | 0.00054692 |
| Endothelial Cell | Ntn11 | 6 | 3.41E-07 | 1.12186035 | 0.774 | 0.515 | 0.00068124 |
| Endothelial Cell | Slc8a1 | 6 | 5.94E-07 | 1.25988433 | 0.821 | 0.558 | 0.00118808 |
| Endothelial Cell | Aoah | 6 | 6.99E-07 | 2.05090939 | 0.405 | 0.114 | 0.00139829 |
| Endothelial Cell | Lmod1 | 6 | 1.18E-06 | 0.2948145 | 0.429 | 0.167 | 0.00236024 |
| Endothelial Cell | Dkk2 | 6 | 4.45E-06 | 0.39219227 | 0.405 | 0.103 | 0.00889757 |

| | | | | | | | |
|---|---|---|---|---|---|---|---|
| Endothelial Cell | Fgf14 | 7 | 6.63E-40 | 6.03083079 | 0.459 | 0.05 | 1.33E-36 |
| Endothelial Cell | LOC102553338 | 7 | 6.63E-40 | 6.01878788 | 0.459 | 0.05 | 1.33E-36 |
| Endothelial Cell | RGD1311946 | 7 | 2.19E-38 | 5.69120067 | 0.459 | 0.05 | 4.37E-35 |
| Endothelial Cell | Swsap1 | 7 | 9.78E-38 | 5.15482322 | 0.446 | 0.045 | 1.96E-34 |
| Endothelial Cell | Cxcl9 | 7 | 4.10E-36 | 2.55542697 | 0.284 | 0.015 | 8.19E-33 |
| Endothelial Cell | Syt1 | 7 | 6.59E-33 | 4.73330323 | 0.446 | 0.058 | 1.32E-29 |
| Endothelial Cell | Ikzf3 | 7 | 1.19E-30 | 2.81795013 | 0.486 | 0.079 | 2.39E-27 |
| Endothelial Cell | Ndst3 | 7 | 1.86E-19 | 2.28543718 | 0.365 | 0.067 | 3.72E-16 |
| Endothelial Cell | Kcnj12 | 7 | 1.11E-18 | 0.36502998 | 0.338 | 0.049 | 2.21E-15 |
| Endothelial Cell | Chst91 | 7 | 4.62E-18 | 3.84523904 | 0.459 | 0.124 | 9.25E-15 |
| Endothelial Cell | Plch1 | 7 | 5.58E-17 | 4.00694555 | 0.405 | 0.069 | 1.12E-13 |
| Endothelial Cell | AABR07025140.1 | 7 | 6.68E-15 | 2.34826338 | 0.77 | 0.496 | 1.34E-11 |
| Endothelial Cell | Pmfbp1 | 7 | 4.08E-13 | 4.95354397 | 0.378 | 0.064 | 8.17E-10 |
| Endothelial Cell | Fbxw101 | 7 | 7.72E-13 | 3.75075466 | 0.459 | 0.174 | 1.54E-09 |

| | | | | | | | |
|---|---|---|---|---|---|---|---|
| Endothelial Cell | Pcp4l1 | 7 | 3.39E-12 | 4.89279361 | 0.351 | 0.096 | 6.79E-09 |
| Endothelial Cell | Ca41 | 7 | 5.28E-12 | 2.2307933 | 0.743 | 0.372 | 1.06E-08 |
| Endothelial Cell | Crlf1 | 7 | 2.24E-11 | 2.32886502 | 0.473 | 0.165 | 4.47E-08 |
| Endothelial Cell | Herc6 | 7 | 1.04E-10 | 2.02069039 | 0.703 | 0.397 | 2.07E-07 |
| Endothelial Cell | Rtp3 | 7 | 1.58E-10 | 1.45625858 | 0.378 | 0.059 | 3.15E-07 |
| Endothelial Cell | Klf5 | 7 | 1.14E-09 | 2.69051181 | 0.473 | 0.171 | 2.28E-06 |
| Endothelial Cell | Edil3 | 7 | 1.17E-09 | 3.47598702 | 0.419 | 0.158 | 2.34E-06 |
| Endothelial Cell | Fbxo401 | 7 | 2.13E-09 | 1.25315257 | 0.351 | 0.067 | 4.26E-06 |
| Endothelial Cell | Cenpe | 7 | 7.72E-09 | 2.64444209 | 0.392 | 0.098 | 1.54E-05 |
| Endothelial Cell | Rp11 | 7 | 8.63E-09 | 1.22829593 | 0.459 | 0.149 | 1.73E-05 |
| Endothelial Cell | Cldn221 | 7 | 8.81E-09 | 1.78801937 | 0.473 | 0.213 | 1.76E-05 |
| Endothelial Cell | Meox1 | 7 | 1.23E-08 | 2.04966589 | 0.432 | 0.143 | 2.46E-05 |
| Endothelial Cell | AABR07053470.1 | 7 | 2.34E-08 | 3.93500299 | 0.446 | 0.117 | 4.68E-05 |
| Endothelial Cell | Dagla1 | 7 | 6.56E-08 | 1.2876967 | 0.486 | 0.229 | 0.00013117 |

| | | | | | | | |
|---|---|---|---|---|---|---|---|
| Endothelial Cell | Adamts3 | 7 | 1.76E-06 | 0.91419827 | 0.419 | 0.098 | 0.00352786 |
| Endothelial Cell | Sphkap | 7 | 2.27E-06 | 2.22110329 | 0.365 | 0.06 | 0.0045401 |
| Endothelial Cell | Itga4 | 7 | 3.30E-06 | 1.7380219 | 0.486 | 0.174 | 0.00660197 |
| Endothelial Cell | Eif2ak2 | 7 | 8.71E-06 | 2.42294542 | 0.595 | 0.342 | 0.01742967 |
| Endothelial Cell | Tex14 | 7 | 1.36E-05 | 1.91168526 | 0.473 | 0.17 | 0.02714963 |
| Endothelial Cell | Dpep1 | 8 | 4.47E-118 | 7.30088738 | 0.493 | 0.003 | 8.93E-115 |
| Endothelial Cell | Tcp11x2 | 8 | 2.35E-114 | 7.4197673 | 0.493 | 0.003 | 4.71E-111 |
| Endothelial Cell | Plin1 | 8 | 7.67E-111 | 7.33637963 | 0.478 | 0.003 | 1.53E-107 |
| Endothelial Cell | Plekhs1 | 8 | 1.27E-109 | 8.07832646 | 0.507 | 0.005 | 2.54E-106 |
| Endothelial Cell | Atp6v1g2 | 8 | 5.76E-107 | 7.39278732 | 0.507 | 0.006 | 1.15E-103 |
| Endothelial Cell | Dusp2 | 8 | 1.33E-106 | 2.53255062 | 0.464 | 0.002 | 2.66E-103 |
| Endothelial Cell | Hs3st2 | 8 | 1.32E-101 | 5.0963427 | 0.522 | 0.01 | 2.63E-98 |
| Endothelial Cell | Tbx15 | 8 | 1.15E-100 | 6.43531087 | 0.493 | 0.007 | 2.30E-97 |
| Endothelial Cell | Flt3 | 8 | 1.21E-92 | 4.09302546 | 0.507 | 0.012 | 2.43E-89 |

| | | | | | | | |
|---|---|---|---|---|---|---|---|
| Endothelial Cell | Col7a1 | 8 | 3.30E-91 | 2.43789054 | 0.449 | 0.006 | 6.60E-88 |
| Endothelial Cell | Prc1 | 8 | 1.58E-88 | 3.0580882 | 0.507 | 0.013 | 3.15E-85 |
| Endothelial Cell | Fbxo16 | 8 | 2.34E-88 | 3.60118838 | 0.391 | 0.003 | 4.68E-85 |
| Endothelial Cell | Kcnh5 | 8 | 2.85E-88 | 6.82475436 | 0.493 | 0.012 | 5.70E-85 |
| Endothelial Cell | Cyp2d2 | 8 | 4.46E-80 | 6.71306499 | 0.493 | 0.016 | 8.92E-77 |
| Endothelial Cell | Hmga2 | 8 | 2.80E-75 | 4.35138335 | 0.507 | 0.008 | 5.59E-72 |
| Endothelial Cell | Adarb2 | 8 | 5.10E-75 | 6.38086481 | 0.507 | 0.021 | 1.02E-71 |
| Endothelial Cell | Sdc1 | 8 | 1.94E-65 | 6.19854807 | 0.609 | 0.051 | 3.87E-62 |
| Endothelial Cell | Col9a11 | 8 | 2.62E-63 | 7.04352554 | 0.493 | 0.024 | 5.24E-60 |
| Endothelial Cell | AABR07031168.1 | 8 | 3.08E-63 | 3.83626259 | 0.261 | 0.008 | 6.17E-60 |
| Endothelial Cell | Nexmif | 8 | 1.16E-56 | 2.89238372 | 0.507 | 0.011 | 2.32E-53 |
| Endothelial Cell | Myo5b | 8 | 2.41E-51 | 7.12500651 | 0.565 | 0.057 | 4.82E-48 |
| Endothelial Cell | Dusp5 | 8 | 4.98E-50 | 3.42897847 | 0.522 | 0.022 | 9.97E-47 |
| Endothelial Cell | Alas2 | 8 | 4.43E-49 | 5.4346886 | 0.493 | 0.019 | 8.85E-46 |

| | | | | | | | |
|---|---|---|---|---|---|---|---|
| Endothelial Cell | Pls11 | 8 | 2.22E-47 | 4.70631495 | 0.493 | 0.045 | 4.43E-44 |
| Endothelial Cell | Myoz31 | 8 | 8.67E-42 | 4.00582855 | 0.522 | 0.064 | 1.73E-38 |
| Endothelial Cell | Ndst31 | 8 | 6.28E-39 | 2.8047833 | 0.507 | 0.059 | 1.26E-35 |
| Endothelial Cell | Syt151 | 8 | 3.88E-38 | 3.76147979 | 0.522 | 0.073 | 7.76E-35 |
| Endothelial Cell | Gadd45b1 | 8 | 1.35E-37 | 3.58516141 | 0.522 | 0.064 | 2.71E-34 |
| Endothelial Cell | Acap1 | 8 | 5.63E-37 | 3.48079939 | 0.464 | 0.056 | 1.13E-33 |
| Endothelial Cell | Ramp11 | 8 | 1.45E-35 | 5.32764556 | 0.609 | 0.113 | 2.89E-32 |
| Endothelial Cell | AABR07033925.11 | 8 | 5.71E-35 | 5.01411927 | 0.42 | 0.071 | 1.14E-31 |
| Endothelial Cell | Dkk21 | 8 | 7.56E-34 | 6.15412393 | 0.681 | 0.09 | 1.51E-30 |
| Endothelial Cell | Itgal | 8 | 5.46E-31 | 2.32096886 | 0.493 | 0.03 | 1.09E-27 |
| Endothelial Cell | Rab27b | 8 | 7.74E-31 | 3.57543538 | 0.536 | 0.021 | 1.55E-27 |
| Endothelial Cell | Slc8a11 | 8 | 1.27E-29 | 2.45152643 | 0.971 | 0.552 | 2.54E-26 |
| Endothelial Cell | Nebl2 | 8 | 6.13E-29 | 2.73073364 | 0.971 | 0.585 | 1.23E-25 |
| Endothelial Cell | Drc3 | 8 | 1.30E-26 | 3.94821259 | 0.522 | 0.015 | 2.60E-23 |

| | | | | | | | |
|---|---|---|---|---|---|---|---|
| Endothelial Cell | Myo16 | 8 | 1.62E-26 | 5.37416178 | 0.522 | 0.009 | 3.24E-23 |
| Endothelial Cell | AABR07041096.1 | 8 | 2.33E-26 | 1.0193219 | 0.449 | 0.007 | 4.65E-23 |
| Endothelial Cell | Itgam | 8 | 1.79E-25 | 3.41987712 | 0.493 | 0.099 | 3.58E-22 |
| Endothelial Cell | Nlrp31 | 8 | 6.53E-25 | 2.74204529 | 0.435 | 0.075 | 1.31E-21 |
| Endothelial Cell | Hcn2 | 8 | 1.17E-24 | 2.12064638 | 0.507 | 0.018 | 2.34E-21 |
| Endothelial Cell | Sulf11 | 8 | 1.30E-24 | 2.66786955 | 0.884 | 0.511 | 2.61E-21 |
| Endothelial Cell | Edaradd1 | 8 | 2.41E-24 | 1.87238061 | 0.507 | 0.063 | 4.82E-21 |
| Endothelial Cell | Bmx | 8 | 4.12E-24 | 4.45385914 | 0.609 | 0.143 | 8.24E-21 |
| Endothelial Cell | Hck1 | 8 | 1.22E-23 | 0.49358769 | 0.478 | 0.096 | 2.45E-20 |
| Endothelial Cell | LOC100912195 | 8 | 9.22E-22 | 4.21159896 | 0.507 | 0.013 | 1.84E-18 |
| Endothelial Cell | Tm6sf2 | 8 | 1.29E-21 | 3.60712082 | 0.449 | 0.023 | 2.58E-18 |
| Endothelial Cell | Afap1l2 | 8 | 3.91E-20 | 3.04592093 | 0.826 | 0.533 | 7.81E-17 |
| Endothelial Cell | Sntg2 | 8 | 5.63E-20 | 1.52636251 | 0.391 | 0.049 | 1.13E-16 |
| Endothelial Cell | Fn11 | 8 | 1.25E-19 | 3.69489628 | 0.783 | 0.236 | 2.49E-16 |

| | | | | | | | |
|---|---|---|---|---|---|---|---|
| Endothelial Cell | Stk32b | 8 | 5.60E-19 | 0.28549476 | 0.435 | 0.047 | 1.12E-15 |
| Endothelial Cell | Hey21 | 8 | 1.07E-18 | 3.95292915 | 0.536 | 0.175 | 2.14E-15 |
| Endothelial Cell | Adcy3 | 8 | 1.81E-18 | 2.75686453 | 0.464 | 0.099 | 3.62E-15 |
| Endothelial Cell | LOC100361 0871 | 8 | 6.27E-18 | 2.72618679 | 0.797 | 0.374 | 1.25E-14 |
| Endothelial Cell | Kif271 | 8 | 9.95E-18 | 2.91549166 | 0.507 | 0.162 | 1.99E-14 |
| Endothelial Cell | Ldb21 | 8 | 1.68E-17 | 3.06536807 | 0.739 | 0.421 | 3.37E-14 |
| Endothelial Cell | Sgo2 | 8 | 5.94E-17 | 3.77140426 | 0.493 | 0.029 | 1.19E-13 |
| Endothelial Cell | Col12a1 | 8 | 1.19E-16 | 2.83050721 | 0.493 | 0.027 | 2.38E-13 |
| Endothelial Cell | Chdh | 8 | 1.37E-16 | 3.52568597 | 0.493 | 0.013 | 2.73E-13 |
| Endothelial Cell | Luzp21 | 8 | 5.16E-16 | 1.36409469 | 0.493 | 0.128 | 1.03E-12 |
| Endothelial Cell | P2ry6 | 8 | 7.38E-16 | 0.47467463 | 0.464 | 0.016 | 1.48E-12 |
| Endothelial Cell | Prune2 | 8 | 1.10E-15 | 3.6820549 | 0.754 | 0.38 | 2.21E-12 |
| Endothelial Cell | Ptchd41 | 8 | 2.83E-15 | 1.12677525 | 0.348 | 0.064 | 5.66E-12 |
| Endothelial Cell | Nrxn3 | 8 | 2.85E-15 | 2.75759865 | 0.536 | 0.125 | 5.69E-12 |

| | | | | | | | |
|---|---|---|---|---|---|---|---|
| Endothelial Cell | Dgkg2 | 8 | 4.32E-15 | 0.93244032 | 0.493 | 0.105 | 8.64E-12 |
| Endothelial Cell | Lmod11 | 8 | 8.77E-15 | 2.88226192 | 0.536 | 0.164 | 1.75E-11 |
| Endothelial Cell | Myof | 8 | 1.10E-14 | 2.99171348 | 0.681 | 0.348 | 2.21E-11 |
| Endothelial Cell | Plcg2 | 8 | 1.21E-14 | 4.19307888 | 0.667 | 0.38 | 2.42E-11 |
| Endothelial Cell | Has21 | 8 | 2.11E-14 | 4.44433924 | 0.507 | 0.137 | 4.22E-11 |
| Endothelial Cell | Hsf5 | 8 | 3.01E-14 | 0.62045521 | 0.449 | 0.063 | 6.01E-11 |
| Endothelial Cell | Diaph3 | 8 | 5.96E-14 | 3.94057046 | 0.522 | 0.03 | 1.19E-10 |
| Endothelial Cell | Sypl2 | 8 | 8.20E-14 | 0.61236943 | 0.449 | 0.018 | 1.64E-10 |
| Endothelial Cell | Tox | 8 | 1.18E-13 | 3.49001402 | 0.71 | 0.398 | 2.35E-10 |
| Endothelial Cell | Kcnn41 | 8 | 5.45E-13 | 3.72323465 | 0.565 | 0.184 | 1.09E-09 |
| Endothelial Cell | Gpnmb | 8 | 7.48E-13 | 2.32409011 | 0.464 | 0.034 | 1.50E-09 |
| Endothelial Cell | Olfml2a | 8 | 7.58E-13 | 4.28157081 | 0.565 | 0.226 | 1.52E-09 |
| Endothelial Cell | Carmil1 | 8 | 7.77E-13 | 3.00905374 | 0.696 | 0.293 | 1.55E-09 |
| Endothelial Cell | Jag11 | 8 | 8.57E-13 | 2.21767792 | 0.783 | 0.392 | 1.71E-09 |

| | | | | | | | |
|---|---|---|---|---|---|---|---|
| Endothelial Cell | Cldn222 | 8 | 1.79E-12 | 2.38765556 | 0.507 | 0.212 | 3.59E-09 |
| Endothelial Cell | Fbn1 | 8 | 2.22E-12 | 2.05222473 | 0.754 | 0.401 | 4.44E-09 |
| Endothelial Cell | Aoah1 | 8 | 3.22E-12 | 1.64891884 | 0.493 | 0.113 | 6.44E-09 |
| Endothelial Cell | Sarm1 | 8 | 4.82E-12 | 4.07350114 | 0.522 | 0.208 | 9.64E-09 |
| Endothelial Cell | Msr1 | 8 | 4.84E-12 | 2.70789888 | 0.507 | 0.197 | 9.68E-09 |
| Endothelial Cell | Ptprn21 | 8 | 5.68E-12 | 3.95199275 | 0.362 | 0.098 | 1.14E-08 |
| Endothelial Cell | Ncam21 | 8 | 1.08E-11 | 2.8030754 | 0.522 | 0.229 | 2.17E-08 |
| Endothelial Cell | Sox13 | 8 | 1.54E-11 | 1.67694056 | 0.855 | 0.591 | 3.09E-08 |
| Endothelial Cell | Smoc12 | 8 | 2.30E-11 | 1.39996816 | 0.493 | 0.124 | 4.60E-08 |
| Endothelial Cell | Thsd7a1 | 8 | 2.34E-11 | 1.35570943 | 0.855 | 0.593 | 4.68E-08 |
| Endothelial Cell | Cxcl1 | 8 | 6.75E-11 | 1.98871061 | 0.507 | 0.11 | 1.35E-07 |
| Endothelial Cell | Npnt1 | 8 | 7.73E-11 | 3.15533022 | 0.478 | 0.213 | 1.55E-07 |
| Endothelial Cell | Adam12 | 8 | 8.54E-11 | 3.98929624 | 0.551 | 0.208 | 1.71E-07 |
| Endothelial Cell | Kcnn3 | 8 | 1.03E-10 | 2.66668018 | 0.667 | 0.296 | 2.06E-07 |

| | | | | | | | |
|---|---|---|---|---|---|---|---|
| Endothelial Cell | Hlf1 | 8 | 1.11E-10 | 2.59561025 | 0.507 | 0.15 | 2.21E-07 |
| Endothelial Cell | Trib31 | 8 | 1.20E-10 | 0.67108982 | 0.464 | 0.183 | 2.41E-07 |
| Endothelial Cell | Tmem17 | 8 | 1.28E-10 | 1.57066935 | 0.493 | 0.213 | 2.56E-07 |
| Endothelial Cell | Pkhd1l12 | 8 | 1.33E-10 | 2.1346778 | 0.667 | 0.391 | 2.65E-07 |
| Endothelial Cell | Nuak11 | 8 | 1.50E-10 | 1.57194163 | 0.826 | 0.534 | 3.01E-07 |
| Endothelial Cell | Unc45b | 8 | 2.29E-10 | 2.15768169 | 0.522 | 0.133 | 4.57E-07 |
| Endothelial Cell | Eln2 | 8 | 2.42E-10 | 2.40834874 | 0.71 | 0.394 | 4.85E-07 |
| Endothelial Cell | Dmtn | 8 | 2.61E-10 | 2.59435158 | 0.58 | 0.278 | 5.22E-07 |
| Endothelial Cell | Kcng3 | 8 | 2.65E-10 | 3.07154586 | 0.464 | 0.146 | 5.29E-07 |
| Endothelial Cell | Eva1c | 8 | 3.10E-10 | 2.72033952 | 0.536 | 0.199 | 6.19E-07 |
| Endothelial Cell | Kif21a | 8 | 4.42E-10 | 2.48743736 | 0.478 | 0.146 | 8.84E-07 |
| Endothelial Cell | AABR07050449.11 | 8 | 9.85E-10 | 2.04375529 | 0.478 | 0.167 | 1.97E-06 |
| Endothelial Cell | Alcam1 | 8 | 1.38E-09 | 2.72804166 | 0.667 | 0.234 | 2.77E-06 |
| Endothelial Cell | Synpo | 8 | 2.37E-09 | 3.04701318 | 0.652 | 0.347 | 4.73E-06 |

| | | | | | | | |
|---|---|---|---|---|---|---|---|
| Endothelial Cell | Pcsk51 | 8 | 2.39E-09 | 3.09233218 | 0.638 | 0.323 | 4.78E-06 |
| Endothelial Cell | Sv2c1 | 8 | 2.56E-09 | 5.29108782 | 0.522 | 0.061 | 5.12E-06 |
| Endothelial Cell | Prrg4 | 8 | 2.77E-09 | 2.22823654 | 0.507 | 0.12 | 5.55E-06 |
| Endothelial Cell | Abo2 | 8 | 2.88E-09 | 1.30552924 | 0.493 | 0.16 | 5.76E-06 |
| Endothelial Cell | Tnc1 | 8 | 3.03E-09 | 2.03341533 | 0.507 | 0.115 | 6.06E-06 |
| Endothelial Cell | Rfx2 | 8 | 6.71E-09 | 2.49183759 | 0.522 | 0.257 | 1.34E-05 |
| Endothelial Cell | Cd55 | 8 | 8.65E-09 | 3.23348074 | 0.522 | 0.113 | 1.73E-05 |
| Endothelial Cell | Rhpn21 | 8 | 9.80E-09 | 1.96778651 | 0.478 | 0.168 | 1.96E-05 |
| Endothelial Cell | Hs3st11 | 8 | 1.29E-08 | 1.78282517 | 0.565 | 0.223 | 2.58E-05 |
| Endothelial Cell | Aacs | 8 | 2.66E-08 | 2.40952055 | 0.493 | 0.199 | 5.33E-05 |
| Endothelial Cell | Scn1a | 8 | 3.65E-08 | 1.14152606 | 0.029 | 0.351 | 7.30E-05 |
| Endothelial Cell | LOC1036933231 | 8 | 3.86E-08 | 4.25236793 | 0.493 | 0.119 | 7.72E-05 |
| Endothelial Cell | Srpk31 | 8 | 4.59E-08 | 0.99392133 | 0.043 | 0.338 | 9.18E-05 |
| Endothelial Cell | Gng10 | 8 | 6.37E-08 | 1.94639187 | 0.522 | 0.254 | 0.00012734 |

| | | | | | | | |
|---|---|---|---|---|---|---|---|
| Endothelial Cell | Fndc11 | 8 | 9.12E-08 | 2.43229744 | 0.609 | 0.272 | 0.00018238 |
| Endothelial Cell | Zmat41 | 8 | 1.07E-07 | 1.60747654 | 0.522 | 0.269 | 0.00021422 |
| Endothelial Cell | Gpr631 | 8 | 1.29E-07 | 2.71895719 | 0.493 | 0.216 | 0.00025813 |
| Endothelial Cell | Edn1 | 8 | 2.28E-07 | 2.10062204 | 0.522 | 0.216 | 0.00045517 |
| Endothelial Cell | Tmem178a | 8 | 3.06E-07 | 2.74064631 | 0.449 | 0.137 | 0.00061188 |
| Endothelial Cell | Flnc | 8 | 3.26E-07 | 0.27089486 | 0.029 | 0.285 | 0.00065164 |
| Endothelial Cell | Efhd11 | 8 | 3.97E-07 | 3.40435863 | 0.464 | 0.2 | 0.00079366 |
| Endothelial Cell | Slc25a21 | 8 | 6.86E-07 | 0.6250594 | 0.435 | 0.071 | 0.00137244 |
| Endothelial Cell | Bdnf | 8 | 6.92E-07 | 2.66696443 | 0.507 | 0.144 | 0.00138466 |
| Endothelial Cell | Adgrd1 | 8 | 7.14E-07 | 2.92936484 | 0.522 | 0.198 | 0.00142826 |
| Endothelial Cell | Enpp2 | 8 | 7.63E-07 | 1.84050138 | 0.464 | 0.115 | 0.00152605 |
| Endothelial Cell | Prox11 | 8 | 8.08E-07 | 0.33293374 | 0.029 | 0.376 | 0.00161514 |
| Endothelial Cell | AABR07007026.11 | 8 | 8.43E-07 | 2.0722573 | 0.522 | 0.211 | 0.00168586 |
| Endothelial Cell | Atp1a1 | 8 | 9.01E-07 | 1.3180796 | 0.652 | 0.382 | 0.00180155 |

| | | | | | | | |
|---|---|---|---|---|---|---|---|
| Endothelial Cell | Ltbp21 | 8 | 9.23E-07 | 2.34008193 | 0.493 | 0.197 | 0.00184552 |
| Endothelial Cell | Inhba1 | 8 | 9.89E-07 | 2.8860767 | 0.478 | 0.094 | 0.00197775 |
| Endothelial Cell | Crip11 | 8 | 1.21E-06 | 1.76146624 | 0.609 | 0.35 | 0.00242877 |
| Endothelial Cell | Chn22 | 8 | 1.22E-06 | 1.49066325 | 0.696 | 0.4 | 0.00243024 |
| Endothelial Cell | AABR07044049.1 | 8 | 1.22E-06 | 0.97523428 | 0.71 | 0.345 | 0.00244609 |
| Endothelial Cell | Aox3 | 8 | 1.45E-06 | 1.78661803 | 0.594 | 0.274 | 0.00289815 |
| Endothelial Cell | AABR07006275.1 | 8 | 1.64E-06 | 2.47305845 | 0.377 | 0.035 | 0.00327303 |
| Endothelial Cell | Itgbl11 | 8 | 1.71E-06 | 1.56814119 | 0.681 | 0.338 | 0.0034294 |
| Endothelial Cell | Ror21 | 8 | 1.85E-06 | 1.60323233 | 0.101 | 0.382 | 0.00370512 |
| Endothelial Cell | Col18a1 | 8 | 1.99E-06 | 3.11926917 | 0.594 | 0.279 | 0.00397684 |
| Endothelial Cell | Mbp | 8 | 2.03E-06 | 0.69964727 | 0.478 | 0.129 | 0.00405692 |
| Endothelial Cell | Frmd4b | 8 | 2.32E-06 | 1.02457041 | 0.754 | 0.41 | 0.0046314 |
| Endothelial Cell | Epha41 | 8 | 2.44E-06 | 1.35990285 | 0.696 | 0.363 | 0.00487016 |
| Endothelial Cell | Usp21 | 8 | 2.81E-06 | 1.27074989 | 0.043 | 0.337 | 0.00561744 |

| | | | | | | | |
|---|---|---|---|---|---|---|---|
| Endothelial Cell | Tmem1002 | 8 | 3.76E-06 | 1.89378054 | 0.551 | 0.269 | 0.00752239 |
| Endothelial Cell | Tbxas11 | 8 | 3.98E-06 | 1.78815243 | 0.522 | 0.169 | 0.00795334 |
| Endothelial Cell | AABR07031399.1 | 8 | 4.71E-06 | 1.29993196 | 0.536 | 0.12 | 0.00941843 |
| Endothelial Cell | Pip5k1b1 | 8 | 6.72E-06 | 3.20286077 | 0.594 | 0.322 | 0.01344033 |
| Endothelial Cell | Tgfb22 | 8 | 7.30E-06 | 1.75670972 | 0.638 | 0.378 | 0.01459161 |
| Endothelial Cell | Dpep2 | 8 | 9.91E-06 | 1.5201007 | 0.522 | 0.255 | 0.01981555 |
| Endothelial Cell | Nkd1 | 8 | 1.01E-05 | 1.24376029 | 0.464 | 0.182 | 0.02017 |
| Endothelial Cell | Bche1 | 8 | 1.23E-05 | 1.2762994 | 0.478 | 0.129 | 0.02456322 |
| Endothelial Cell | Creb5 | 8 | 1.44E-05 | 0.90481938 | 0.783 | 0.478 | 0.02877326 |
| Endothelial Cell | AABR07058158.1 | 8 | 1.69E-05 | 1.3437051 | 0.58 | 0.315 | 0.03386916 |
| Endothelial Cell | Nfatc21 | 8 | 2.04E-05 | 3.10568122 | 0.536 | 0.268 | 0.04085033 |
| Endothelial Cell | Cdh8 | 9 | 5.21E-28 | 1.61934969 | 0.417 | 0.051 | 1.04E-24 |
| Endothelial Cell | Ceacam16 | 9 | 2.08E-27 | 1.58996243 | 0.417 | 0.052 | 4.16E-24 |
| Endothelial Cell | Lilrb21 | 9 | 5.06E-21 | 5.06504708 | 0.533 | 0.12 | 1.01E-17 |

| | | | | | | | |
|---|---|---|---|---|---|---|---|
| Endothelial Cell | AC128789.1 | 9 | 2.24E-19 | 5.25515113 | 0.467 | 0.102 | 4.48E-16 |
| Endothelial Cell | Lancl31 | 9 | 8.39E-14 | 5.2595032 | 0.467 | 0.103 | 1.68E-10 |
| Endothelial Cell | Nlgn1 | 9 | 4.23E-13 | 3.19363485 | 0.317 | 0.044 | 8.46E-10 |
| Endothelial Cell | Lamc31 | 9 | 3.15E-12 | 5.70114614 | 0.45 | 0.132 | 6.29E-09 |
| Endothelial Cell | Erbb31 | 9 | 1.60E-10 | 0.82448818 | 0.433 | 0.126 | 3.19E-07 |
| Endothelial Cell | Vtn | 9 | 1.95E-10 | 2.36048326 | 0.45 | 0.143 | 3.90E-07 |
| Endothelial Cell | C1qb1 | 9 | 6.72E-10 | 1.04867089 | 0.4 | 0.107 | 1.34E-06 |
| Endothelial Cell | AABR07054565.1 | 9 | 2.67E-09 | 3.72419265 | 0.533 | 0.21 | 5.33E-06 |
| Endothelial Cell | Rgs71 | 9 | 1.20E-08 | 3.52029788 | 0.45 | 0.136 | 2.41E-05 |
| Endothelial Cell | Gfra31 | 9 | 1.42E-08 | 0.45020786 | 0.517 | 0.209 | 2.85E-05 |
| Endothelial Cell | Lrrc172 | 9 | 1.61E-08 | 1.08514511 | 0.55 | 0.258 | 3.22E-05 |
| Endothelial Cell | Ctnnd21 | 9 | 1.95E-08 | 3.15455523 | 0.55 | 0.284 | 3.90E-05 |
| Endothelial Cell | Trpm3 | 9 | 2.67E-08 | 1.64085561 | 0.467 | 0.181 | 5.35E-05 |
| Endothelial Cell | Fbln7 | 9 | 1.58E-07 | 0.63642109 | 0.417 | 0.16 | 0.00031535 |

| | | | | | | | |
|---|---|---|---|---|---|---|---|
| Endothelial Cell | Wdfy4 | 9 | 1.96E-07 | 1.80208627 | 0.317 | 0.064 | 0.00039181 |
| Endothelial Cell | Gpc31 | 9 | 4.90E-07 | 0.5510364 | 0.433 | 0.135 | 0.00097923 |
| Endothelial Cell | Sarm11 | 9 | 7.64E-07 | 2.82192176 | 0.533 | 0.21 | 0.00152793 |
| Endothelial Cell | Gabrb1 | 9 | 9.44E-06 | 1.63336291 | 0.4 | 0.084 | 0.01887644 |
| Endothelial Cell | Egflam | 10 | 4.78E-06 | 0.65436765 | 0.102 | 0.376 | 0.00956258 |
| Endothelial Cell | Cdh191 | 10 | 5.55E-06 | 0.54565586 | 0.153 | 0.413 | 0.01110732 |
| Endothelial Cell | Nckap5 | 10 | 1.06E-05 | 0.82321799 | 0.102 | 0.357 | 0.02112195 |
| Endothelial Cell | Col6a6 | 10 | 2.27E-05 | 0.61246671 | 0.102 | 0.366 | 0.04534616 |
| Endothelial Cell | Tmsb4x2 | 11 | 1.00E-06 | 1.06349871 | 0.863 | 0.612 | 0.00200625 |
| Endothelial Cell | Fabp41 | 11 | 1.28E-06 | 0.89790224 | 0.902 | 0.646 | 0.00255281 |
| Endothelial Cell | Plet11 | 12 | 1.47E-26 | 6.4564777 | 0.568 | 0.088 | 2.94E-23 |
| Endothelial Cell | Rufy4 | 12 | 6.01E-24 | 3.40512691 | 0.523 | 0.08 | 1.20E-20 |
| Endothelial Cell | Nlrp12 | 12 | 1.21E-23 | 1.73951888 | 0.523 | 0.081 | 2.42E-20 |
| Endothelial Cell | Pax3 | 12 | 2.40E-23 | 0.50939622 | 0.523 | 0.081 | 4.81E-20 |

| | | | | | | | |
|---|---|---|---|---|---|---|---|
| Endothelial Cell | Il11 | 12 | 8.07E-15 | 5.75927034 | 0.568 | 0.132 | 1.61E-11 |
| Endothelial Cell | Tll2 | 12 | 1.92E-14 | 3.26656483 | 0.659 | 0.233 | 3.84E-11 |
| Endothelial Cell | Sgpp2 | 12 | 3.87E-14 | 0.57535066 | 0.568 | 0.156 | 7.75E-11 |
| Endothelial Cell | Angptl82 | 12 | 8.10E-13 | 3.86112673 | 0.614 | 0.205 | 1.62E-09 |
| Endothelial Cell | Cacna1b1 | 12 | 1.00E-12 | 1.16548857 | 0.568 | 0.158 | 2.01E-09 |
| Endothelial Cell | Bend61 | 12 | 1.53E-12 | 3.06035618 | 0.682 | 0.292 | 3.05E-09 |
| Endothelial Cell | AABR07041411.11 | 12 | 2.44E-12 | 1.99922248 | 0.409 | 0.087 | 4.88E-09 |
| Endothelial Cell | Galnt13 | 12 | 1.07E-11 | 3.69743942 | 0.523 | 0.102 | 2.14E-08 |
| Endothelial Cell | Slc35g21 | 12 | 1.20E-11 | 0.28561157 | 0.455 | 0.12 | 2.40E-08 |
| Endothelial Cell | Dact22 | 12 | 2.95E-11 | 3.89212027 | 0.614 | 0.229 | 5.91E-08 |
| Endothelial Cell | Ttyh1 | 12 | 4.75E-11 | 3.29505849 | 0.591 | 0.171 | 9.50E-08 |
| Endothelial Cell | Hap11 | 12 | 1.17E-10 | 1.89981309 | 0.636 | 0.278 | 2.34E-07 |
| Endothelial Cell | Sctr | 12 | 1.79E-10 | 3.21893817 | 0.409 | 0.075 | 3.58E-07 |
| Endothelial Cell | Olfm21 | 12 | 1.96E-10 | 1.53192494 | 0.523 | 0.152 | 3.93E-07 |

| | | | | | | | |
|---|---|---|---|---|---|---|---|
| Endothelial Cell | Eya1 | 12 | 3.19E-10 | 0.86615206 | 0.364 | 0.06 | 6.39E-07 |
| Endothelial Cell | Adgre41 | 12 | 5.21E-10 | 0.8282203 | 0.545 | 0.157 | 1.04E-06 |
| Endothelial Cell | Kif21a1 | 12 | 6.07E-10 | 0.32799417 | 0.568 | 0.15 | 1.21E-06 |
| Endothelial Cell | Setmar1 | 12 | 7.30E-10 | 0.63666396 | 0.568 | 0.185 | 1.46E-06 |
| Endothelial Cell | Brinp3 | 12 | 5.30E-09 | 0.52511565 | 0.523 | 0.174 | 1.06E-05 |
| Endothelial Cell | Coro1a2 | 12 | 1.07E-08 | 0.89754293 | 0.614 | 0.258 | 2.14E-05 |
| Endothelial Cell | Slc16a7 | 12 | 2.66E-08 | 3.09707519 | 0.568 | 0.166 | 5.31E-05 |
| Endothelial Cell | Ppm1e1 | 12 | 5.44E-08 | 1.92918641 | 0.591 | 0.271 | 0.00010888 |
| Endothelial Cell | Trim7 | 12 | 8.71E-08 | 1.64497885 | 0.659 | 0.299 | 0.00017421 |
| Endothelial Cell | AABR07052585.2 | 12 | 1.56E-07 | 2.92238973 | 0.682 | 0.353 | 0.0003119 |
| Endothelial Cell | Lilrb41 | 12 | 1.86E-07 | 1.49796962 | 0.5 | 0.212 | 0.00037166 |
| Endothelial Cell | Gap431 | 12 | 2.03E-07 | 0.75390717 | 0.545 | 0.214 | 0.00040677 |
| Endothelial Cell | Aqp1 | 12 | 2.50E-07 | 1.03530984 | 0.886 | 0.606 | 0.00050045 |
| Endothelial Cell | Prkcz | 12 | 2.57E-07 | 2.32069689 | 0.455 | 0.088 | 0.0005138 |

| | | | | | | | |
|---|---|---|---|---|---|---|---|
| Endothelial Cell | Steap41 | 12 | 4.19E-07 | 3.154597 | 0.614 | 0.279 | 0.0008373 |
| Endothelial Cell | Trim50 | 12 | 5.36E-07 | 2.04059026 | 0.614 | 0.267 | 0.00107103 |
| Endothelial Cell | Rcan11 | 12 | 8.43E-07 | 1.70433061 | 0.659 | 0.355 | 0.00168623 |
| Endothelial Cell | Prss48 | 12 | 9.04E-07 | 1.81013178 | 0.636 | 0.337 | 0.00180715 |
| Endothelial Cell | Ntng11 | 12 | 1.44E-06 | 0.52142337 | 0.455 | 0.152 | 0.00288782 |
| Endothelial Cell | Gdf61 | 12 | 1.57E-06 | 0.71820422 | 0.455 | 0.165 | 0.00313923 |
| Endothelial Cell | Ccl212 | 12 | 1.81E-06 | 3.22197153 | 0.591 | 0.257 | 0.00362437 |
| Endothelial Cell | Sncaip | 12 | 2.60E-06 | 1.14829969 | 0.886 | 0.472 | 0.00520763 |
| Endothelial Cell | Aff21 | 12 | 2.84E-06 | 1.68238995 | 0.636 | 0.312 | 0.00567026 |
| Endothelial Cell | Lpar31 | 12 | 4.37E-06 | 0.29110235 | 0.568 | 0.256 | 0.00873503 |
| Endothelial Cell | Klf51 | 12 | 5.37E-06 | 2.24632355 | 0.477 | 0.179 | 0.01074048 |
| Endothelial Cell | Nova1 | 12 | 5.67E-06 | 2.22572571 | 0.364 | 0.097 | 0.011337 |
| Endothelial Cell | Itm2a | 12 | 6.63E-06 | 1.46316679 | 0.636 | 0.295 | 0.01325479 |
| Endothelial Cell | Speg | 12 | 7.70E-06 | 2.00123972 | 0.5 | 0.197 | 0.01539479 |

| | | | | | | | |
|---|---|---|---|---|---|---|---|
| Endothelial Cell | Rgs61 | 12 | 9.45E-06 | 2.5530376 | 0.409 | 0.13 | 0.01889544 |
| Endothelial Cell | Rassf9 | 12 | 1.09E-05 | 1.06875177 | 0.705 | 0.41 | 0.02174054 |
| Endothelial Cell | Errfi1 | 12 | 1.23E-05 | 1.20970816 | 0.409 | 0.125 | 0.02457781 |
| Endothelial Cell | Piezo2 | 12 | 1.33E-05 | 0.68287739 | 0.614 | 0.253 | 0.02669599 |
| Endothelial Cell | Scn1a1 | 12 | 1.46E-05 | 1.48999902 | 0.636 | 0.32 | 0.02915766 |
| Endothelial Cell | Ptn1 | 12 | 1.54E-05 | 0.28211384 | 0.659 | 0.261 | 0.0307567 |
| Endothelial Cell | Slco2a1 | 12 | 1.66E-05 | 0.39007092 | 0.341 | 0.057 | 0.03328006 |
| Endothelial Cell | Sema3d | 12 | 1.72E-05 | 1.33813025 | 0.705 | 0.434 | 0.03446714 |
| Endothelial Cell | Prg4 | 12 | 2.20E-05 | 2.3064549 | 0.455 | 0.064 | 0.0440436 |
| Endothelial Cell | Adamts17 | 12 | 2.31E-05 | 1.56477147 | 0.614 | 0.322 | 0.04614751 |
| Endothelial Cell | Trabd2b2 | 12 | 2.47E-05 | 0.98932181 | 0.705 | 0.323 | 0.04935754 |
| Endothelial Cell | Kif232 | 12 | 2.49E-05 | 2.02714724 | 0.568 | 0.252 | 0.0497346 |
| Endothelial Cell | LOC103690241 | 13 | 5.87E-110 | 7.21730821 | 0.875 | 0.008 | 1.17E-106 |
| Endothelial Cell | AC121209.1 | 13 | 2.38E-87 | 7.2715286 | 0.5 | 0.001 | 4.76E-84 |

| | | | | | | | |
|---|---|---|---|---|---|---|---|
| Endothelial Cell | Fcgr3a | 13 | 3.32E-87 | 7.26434294 | 0.5 | 0.002 | 6.65E-84 |
| Endothelial Cell | Crhr1 | 13 | 7.75E-82 | 7.15279861 | 0.625 | 0.004 | 1.55E-78 |
| Endothelial Cell | Hcn3 | 13 | 9.97E-82 | 7.16570453 | 0.625 | 0.003 | 1.99E-78 |
| Endothelial Cell | LOC691995 | 13 | 2.94E-68 | 4.39086299 | 0.625 | 0.006 | 5.88E-65 |
| Endothelial Cell | LOC100362366 | 13 | 4.02E-68 | 7.02200947 | 0.625 | 0.006 | 8.05E-65 |
| Endothelial Cell | Kcna6 | 13 | 1.45E-49 | 7.21431484 | 0.375 | 0.002 | 2.91E-46 |
| Endothelial Cell | Esco2 | 13 | 2.55E-36 | 3.23345648 | 0.625 | 0.022 | 5.10E-33 |
| Endothelial Cell | AABR07057997.1 | 13 | 7.54E-36 | 6.24349952 | 0.75 | 0.005 | 1.51E-32 |
| Endothelial Cell | Smoc2 | 13 | 1.10E-23 | 2.35907283 | 0.75 | 0.022 | 2.21E-20 |
| Endothelial Cell | Aspa | 13 | 1.12E-19 | 5.36333755 | 0.5 | 0.017 | 2.24E-16 |
| Endothelial Cell | Alox5ap | 13 | 2.41E-19 | 5.04394073 | 0.5 | 0.014 | 4.82E-16 |
| Endothelial Cell | Npffr2 | 13 | 3.11E-19 | 3.18646459 | 0.625 | 0.044 | 6.22E-16 |
| Endothelial Cell | Slit1 | 13 | 4.27E-18 | 4.44279103 | 0.625 | 0.014 | 8.53E-15 |
| Endothelial Cell | Cntfr | 13 | 5.84E-17 | 3.89432831 | 0.5 | 0.014 | 1.17E-13 |

| | | | | | | | |
|---|---|---|---|---|---|---|---|
| Endothelial Cell | Fam180a | 13 | 6.21E-16 | 4.55805326 | 0.5 | 0.003 | 1.24E-12 |
| Endothelial Cell | Ndst32 | 13 | 3.21E-15 | 4.62021625 | 0.875 | 0.08 | 6.43E-12 |
| Endothelial Cell | S100a91 | 13 | 2.06E-14 | 3.66078302 | 0.75 | 0.048 | 4.12E-11 |
| Endothelial Cell | Sntg21 | 13 | 3.29E-14 | 4.62852203 | 0.75 | 0.065 | 6.58E-11 |
| Endothelial Cell | AABR07029272.1 | 13 | 3.29E-13 | 1.3769637 | 0.875 | 0.081 | 6.59E-10 |
| Endothelial Cell | Cpvl1 | 13 | 3.37E-13 | 2.71514873 | 0.875 | 0.087 | 6.75E-10 |
| Endothelial Cell | Pak1 | 13 | 7.19E-13 | 3.12532309 | 0.375 | 0.012 | 1.44E-09 |
| Endothelial Cell | Plk4 | 13 | 8.55E-13 | 5.11638582 | 0.75 | 0.037 | 1.71E-09 |
| Endothelial Cell | Slamf8 | 13 | 1.42E-12 | 5.11582375 | 0.625 | 0.006 | 2.84E-09 |
| Endothelial Cell | Nrk | 13 | 1.63E-12 | 4.98392334 | 0.75 | 0.109 | 3.27E-09 |
| Endothelial Cell | Vgll3 | 13 | 2.37E-12 | 2.42419453 | 1 | 0.124 | 4.73E-09 |
| Endothelial Cell | Pdpn | 13 | 3.21E-12 | 2.00132124 | 0.5 | 0.063 | 6.41E-09 |
| Endothelial Cell | LOC691141 | 13 | 3.41E-12 | 2.48560307 | 0.5 | 0.006 | 6.82E-09 |
| Endothelial Cell | Ntrk2 | 13 | 7.66E-12 | 2.41295569 | 0.75 | 0.104 | 1.53E-08 |

| | | | | | | | |
|---|---|---|---|---|---|---|---|
| Endothelial Cell | Kif6 | 13 | 1.01E-10 | 4.1226957 | 0.75 | 0.083 | 2.03E-07 |
| Endothelial Cell | Syt11 | 13 | 1.50E-10 | 2.89533096 | 0.5 | 0.079 | 3.00E-07 |
| Endothelial Cell | Col11a21 | 13 | 3.10E-10 | 2.30512562 | 0.875 | 0.107 | 6.20E-07 |
| Endothelial Cell | AABR07034940.21 | 13 | 6.85E-10 | 2.01311335 | 1 | 0.172 | 1.37E-06 |
| Endothelial Cell | Nhs | 13 | 7.59E-10 | 1.22988934 | 0.375 | 0.08 | 1.52E-06 |
| Endothelial Cell | Adam231 | 13 | 2.82E-09 | 3.83700011 | 1 | 0.154 | 5.64E-06 |
| Endothelial Cell | Il1rapl2 | 13 | 6.37E-09 | 2.24801039 | 0.625 | 0.091 | 1.27E-05 |
| Endothelial Cell | Aldh1a2 | 13 | 9.30E-09 | 1.49232897 | 0.5 | 0.144 | 1.86E-05 |
| Endothelial Cell | Dusp51 | 13 | 1.13E-08 | 2.1200512 | 0.625 | 0.048 | 2.26E-05 |
| Endothelial Cell | Scube31 | 13 | 1.87E-08 | 3.49886882 | 0.5 | 0.05 | 3.74E-05 |
| Endothelial Cell | AABR07033925.12 | 13 | 1.94E-08 | 1.69240603 | 0.625 | 0.087 | 3.88E-05 |
| Endothelial Cell | P2rx11 | 13 | 2.20E-08 | 2.62249076 | 0.75 | 0.113 | 4.41E-05 |
| Endothelial Cell | Xirp1 | 13 | 3.39E-08 | 1.54708377 | 0.75 | 0.06 | 6.78E-05 |
| Endothelial Cell | Tmem178a1 | 13 | 4.16E-08 | 1.38310501 | 1 | 0.149 | 8.33E-05 |

| | | | | | | | |
|---|---|---|---|---|---|---|---|
| Endothelial Cell | Adcy31 | 13 | 4.50E-08 | 0.39501447 | 0.375 | 0.119 | 9.00E-05 |
| Endothelial Cell | Ptgfr | 13 | 5.75E-08 | 1.23871446 | 0.5 | 0.114 | 0.000115 |
| Endothelial Cell | Slc25a211 | 13 | 1.64E-07 | 1.10068247 | 0.75 | 0.087 | 0.00032716 |
| Endothelial Cell | Nlrp32 | 13 | 1.70E-07 | 2.66185736 | 0.625 | 0.093 | 0.0003404 |
| Endothelial Cell | Pdp2 | 13 | 2.36E-07 | 2.28915652 | 0.75 | 0.117 | 0.00047187 |
| Endothelial Cell | Has22 | 13 | 2.95E-07 | 2.44081415 | 0.625 | 0.155 | 0.00059068 |
| Endothelial Cell | Dab1 | 13 | 3.01E-07 | 3.39452662 | 0.625 | 0.008 | 0.00060252 |
| Endothelial Cell | Trpm31 | 13 | 3.14E-07 | 4.40678598 | 0.875 | 0.191 | 0.00062731 |
| Endothelial Cell | Med12l1 | 13 | 5.19E-07 | 0.79126923 | 0.75 | 0.121 | 0.00103816 |
| Endothelial Cell | Sema3e1 | 13 | 5.43E-07 | 2.42583149 | 0.5 | 0.115 | 0.00108628 |
| Endothelial Cell | Daam21 | 13 | 7.40E-07 | 3.76996359 | 0.875 | 0.126 | 0.00148068 |
| Endothelial Cell | Smarca1 | 13 | 7.50E-07 | 2.56859588 | 0.375 | 0.078 | 0.00150042 |
| Endothelial Cell | Lyz2 | 13 | 7.58E-07 | 2.67410526 | 0.75 | 0.131 | 0.00151604 |
| Endothelial Cell | Gucy1b1 | 13 | 8.23E-07 | 1.2543202 | 0.625 | 0.11 | 0.00164531 |

| | | | | | | | |
|---|---|---|---|---|---|---|---|
| Endothelial Cell | AABR07007068.11 | 13 | 9.04E-07 | 4.47362104 | 1 | 0.246 | 0.00180726 |
| Endothelial Cell | Adamts31 | 13 | 9.23E-07 | 0.55373361 | 0.75 | 0.114 | 0.00184637 |
| Endothelial Cell | Cspg41 | 13 | 1.22E-06 | 0.90985301 | 0.875 | 0.261 | 0.00243241 |
| Endothelial Cell | C1qb2 | 13 | 1.25E-06 | 3.1062413 | 0.375 | 0.121 | 0.00250325 |
| Endothelial Cell | Tenm4 | 13 | 1.29E-06 | 1.92229318 | 0.5 | 0.149 | 0.00258454 |
| Endothelial Cell | Plac8 | 13 | 1.34E-06 | 2.07023696 | 0.625 | 0.143 | 0.00267598 |
| Endothelial Cell | Cacnb4 | 13 | 1.52E-06 | 1.62136396 | 0.5 | 0.049 | 0.00303029 |
| Endothelial Cell | Enpp21 | 13 | 3.25E-06 | 1.11405756 | 0.625 | 0.132 | 0.00649942 |
| Endothelial Cell | Olfml2b1 | 13 | 3.29E-06 | 1.87557504 | 0.75 | 0.091 | 0.00658438 |
| Endothelial Cell | Musk1 | 13 | 3.47E-06 | 3.69696936 | 0.75 | 0.256 | 0.00693121 |
| Endothelial Cell | Zfp385b1 | 13 | 3.71E-06 | 4.32375322 | 1 | 0.329 | 0.00742919 |
| Endothelial Cell | Aox13 | 13 | 3.76E-06 | 3.20669402 | 1 | 0.314 | 0.00752343 |
| Endothelial Cell | Lgi4 | 13 | 3.81E-06 | 2.53219947 | 1 | 0.372 | 0.00761179 |
| Endothelial Cell | Robo2 | 13 | 3.94E-06 | 1.39445269 | 1 | 0.34 | 0.00788022 |

| | | | | | | | |
|---|---|---|---|---|---|---|---|
| Endothelial Cell | Crispld21 | 13 | 4.04E-06 | 4.80211641 | 0.875 | 0.337 | 0.00808118 |
| Endothelial Cell | Fosl1 | 13 | 4.57E-06 | 3.82413923 | 0.5 | 0.098 | 0.00914486 |
| Endothelial Cell | Cfap77 | 13 | 4.60E-06 | 2.82258726 | 0.5 | 0.099 | 0.00920575 |
| Endothelial Cell | Tmem132d1 | 13 | 5.57E-06 | 2.76685354 | 1 | 0.301 | 0.01114058 |
| Endothelial Cell | Trhde | 13 | 5.96E-06 | 0.95910039 | 0.625 | 0.135 | 0.01192696 |
| Endothelial Cell | Gpc32 | 13 | 6.48E-06 | 2.34817725 | 0.625 | 0.147 | 0.012955 |
| Endothelial Cell | Syt91 | 13 | 7.00E-06 | 1.0598066 | 0.5 | 0.096 | 0.01399433 |
| Endothelial Cell | Rtn4rl1 | 13 | 7.47E-06 | 2.09878045 | 1 | 0.202 | 0.01493105 |
| Endothelial Cell | Prlr1 | 13 | 7.70E-06 | 2.12304698 | 0.625 | 0.155 | 0.01540886 |
| Endothelial Cell | Blnk | 13 | 8.49E-06 | 4.09973287 | 0.625 | 0.227 | 0.01697412 |
| Endothelial Cell | Asic2 | 13 | 8.66E-06 | 0.46551308 | 0.5 | 0.107 | 0.01731619 |
| Endothelial Cell | Prkcz1 | 13 | 8.87E-06 | 1.53371906 | 0.375 | 0.1 | 0.01773898 |
| Endothelial Cell | Errfi11 | 13 | 9.25E-06 | 3.25485395 | 0.625 | 0.132 | 0.01849067 |
| Endothelial Cell | Abca81 | 13 | 1.06E-05 | 1.36476885 | 0.375 | 0.122 | 0.02112831 |

| | | | | | | | |
|---|---|---|---|---|---|---|---|
| Endothelial Cell | Agbl11 | 13 | 1.10E-05 | 1.40040898 | 0.75 | 0.251 | 0.02196854 |
| Endothelial Cell | Hydin1 | 13 | 1.32E-05 | 3.00176631 | 0.375 | 0.043 | 0.02648199 |
| Endothelial Cell | Inhba2 | 13 | 1.49E-05 | 1.79945602 | 0.5 | 0.114 | 0.02985557 |
| Endothelial Cell | Pappa1 | 13 | 1.96E-05 | 3.6257795 | 0.375 | 0.105 | 0.03925643 |
| Endothelial Cell | Kif26b1 | 13 | 2.17E-05 | 5.00680815 | 0.875 | 0.233 | 0.04348801 |
| Endothelial Cell | Spsb4 | 13 | 2.21E-05 | 0.48280575 | 0.5 | 0.126 | 0.04422167 |
| Endothelial Cell | Rab27b1 | 13 | 2.29E-05 | 3.2407841 | 0.375 | 0.049 | 0.04572176 |
| Cardiomyocyte | Tprn | 0 | 1.20E-124 | 4.85675045 | 0.83 | 0.148 | 2.40E-121 |
| Cardiomyocyte | AABR07041411.1 | 0 | 1.15E-113 | 4.49744843 | 0.613 | 0.048 | 2.31E-110 |
| Cardiomyocyte | Nlrp12 | 0 | 2.16E-113 | 3.9700848 | 0.736 | 0.087 | 4.33E-110 |
| Cardiomyocyte | AABR07035796.1 | 0 | 2.16E-113 | 3.9700848 | 0.736 | 0.087 | 4.33E-110 |
| Cardiomyocyte | Spock3 | 0 | 5.14E-112 | 2.87352568 | 0.613 | 0.049 | 1.03E-108 |
| Cardiomyocyte | AC121209.1 | 0 | 8.61E-112 | 3.2540083 | 0.792 | 0.138 | 1.72E-108 |
| Cardiomyocyte | Has2 | 0 | 8.56E-110 | 1.71705773 | 0.613 | 0.051 | 1.71E-106 |

| | | | | | | | |
|---|---|---|---|---|---|---|---|
| Cardiomyocyte | Stac | 0 | 3.74E-106 | 4.02039924 | 0.835 | 0.173 | 7.47E-103 |
| Cardiomyocyte | Nr5a2 | 0 | 9.69E-105 | 4.0215893 | 0.84 | 0.161 | 1.94E-101 |
| Cardiomyocyte | Mill1 | 0 | 7.27E-100 | 3.30079588 | 0.458 | 0.071 | 1.45E-96 |
| Cardiomyocyte | AABR07049156.1 | 0 | 2.34E-97 | 3.64169617 | 0.816 | 0.198 | 4.67E-94 |
| Cardiomyocyte | AABR07049918.1 | 0 | 1.83E-94 | 2.90400891 | 0.608 | 0.074 | 3.66E-91 |
| Cardiomyocyte | Tagln | 0 | 6.51E-92 | 2.64384607 | 0.608 | 0.078 | 1.30E-88 |
| Cardiomyocyte | Edar | 0 | 8.57E-89 | 2.00271882 | 0.792 | 0.164 | 1.71E-85 |
| Cardiomyocyte | AABR07032338.1 | 0 | 1.62E-79 | 1.23002837 | 0.84 | 0.175 | 3.25E-76 |
| Cardiomyocyte | Syt9 | 0 | 4.82E-77 | 1.24630463 | 0.783 | 0.162 | 9.64E-74 |
| Cardiomyocyte | Fyb1 | 0 | 1.05E-74 | 0.53407049 | 0.83 | 0.155 | 2.10E-71 |
| Cardiomyocyte | Cd300lg | 0 | 9.35E-74 | 1.08282015 | 0.75 | 0.149 | 1.87E-70 |
| Cardiomyocyte | Pik3r5 | 0 | 7.49E-73 | 1.39342632 | 0.83 | 0.202 | 1.50E-69 |
| Cardiomyocyte | Spp1 | 0 | 3.22E-72 | 2.92010142 | 0.66 | 0.088 | 6.44E-69 |
| Cardiomyocyte | Greb1l | 0 | 1.63E-71 | 1.57128639 | 0.825 | 0.27 | 3.26E-68 |

| | | | | | | | |
|---|---|---|---|---|---|---|---|
| Cardiomyocyte | Notch3 | 0 | 1.23E-70 | 0.5843181 | 0.354 | 0.093 | 2.46E-67 |
| Cardiomyocyte | Tbx1 | 0 | 3.49E-69 | 0.62560756 | 0.514 | 0.069 | 6.98E-66 |
| Cardiomyocyte | Acta2 | 0 | 2.88E-68 | 1.00697605 | 0.613 | 0.106 | 5.76E-65 |
| Cardiomyocyte | Txk | 0 | 2.39E-67 | 1.78296256 | 0.825 | 0.228 | 4.78E-64 |
| Cardiomyocyte | Nr2f2 | 0 | 1.07E-66 | 0.53952148 | 0.83 | 0.278 | 2.14E-63 |
| Cardiomyocyte | Nnat | 0 | 1.54E-66 | 1.27000165 | 0.811 | 0.298 | 3.08E-63 |
| Cardiomyocyte | Grid2 | 0 | 1.20E-64 | 1.74841127 | 0.769 | 0.219 | 2.41E-61 |
| Cardiomyocyte | Angpt2 | 0 | 2.35E-61 | 1.9851183 | 0.83 | 0.217 | 4.71E-58 |
| Cardiomyocyte | Oasl | 0 | 8.41E-61 | 0.6078761 | 0.717 | 0.181 | 1.68E-57 |
| Cardiomyocyte | RGD1564053 | 0 | 6.93E-60 | 1.12612457 | 0.66 | 0.183 | 1.39E-56 |
| Cardiomyocyte | Olfml2b | 0 | 2.96E-59 | 1.39901543 | 0.811 | 0.228 | 5.91E-56 |
| Cardiomyocyte | Galnt16 | 0 | 3.64E-59 | 1.15727649 | 0.854 | 0.438 | 7.28E-56 |
| Cardiomyocyte | Srgn | 0 | 2.19E-56 | 0.71303515 | 0.797 | 0.227 | 4.37E-53 |
| Cardiomyocyte | LOC100910978.1 | 0 | 4.10E-55 | 0.98103382 | 0.802 | 0.298 | 8.20E-52 |

| | | | | | | | |
|---|---|---|---|---|---|---|---|
| Cardiomyocyte | Sema3e | 0 | 6.28E-54 | 2.09331835 | 0.354 | 0.07 | 1.26E-50 |
| Cardiomyocyte | Lcp2 | 0 | 9.49E-53 | 1.63008575 | 0.84 | 0.364 | 1.90E-49 |
| Cardiomyocyte | Dclk1 | 0 | 1.41E-51 | 1.38611926 | 0.868 | 0.464 | 2.81E-48 |
| Cardiomyocyte | Fam151a | 0 | 1.06E-50 | 2.51417505 | 0.825 | 0.314 | 2.12E-47 |
| Cardiomyocyte | Shc3 | 0 | 2.84E-50 | 0.43017898 | 0.665 | 0.183 | 5.69E-47 |
| Cardiomyocyte | Musk | 0 | 3.33E-48 | 0.47358712 | 0.66 | 0.234 | 6.66E-45 |
| Cardiomyocyte | Nes | 0 | 3.96E-48 | 0.25978195 | 0.604 | 0.141 | 7.93E-45 |
| Cardiomyocyte | Entpd1 | 0 | 9.41E-46 | 0.60728563 | 0.835 | 0.421 | 1.88E-42 |
| Cardiomyocyte | Nrxn3 | 0 | 1.34E-45 | 0.39051703 | 0.844 | 0.455 | 2.69E-42 |
| Cardiomyocyte | Slc35f1 | 0 | 2.06E-45 | 0.63255355 | 0.769 | 0.232 | 4.13E-42 |
| Cardiomyocyte | RGD1563354 | 0 | 2.73E-45 | 0.61146712 | 0.802 | 0.224 | 5.47E-42 |
| Cardiomyocyte | Kcnab1 | 0 | 3.73E-45 | 0.80060066 | 0.816 | 0.342 | 7.47E-42 |
| Cardiomyocyte | Tmem163 | 0 | 1.07E-44 | 1.3848348 | 0.783 | 0.398 | 2.14E-41 |
| Cardiomyocyte | AABR07059258.1 | 0 | 1.73E-44 | 0.95835013 | 0.835 | 0.338 | 3.46E-41 |

| | | | | | | | |
|---|---|---|---|---|---|---|---|
| Cardiomyocyte | Hsf5 | 0 | 2.04E-44 | 0.74667203 | 0.722 | 0.188 | 4.08E-41 |
| Cardiomyocyte | Slfn13 | 0 | 3.35E-44 | 0.35302944 | 0.774 | 0.236 | 6.71E-41 |
| Cardiomyocyte | Cdh11 | 0 | 5.38E-44 | 0.62258216 | 0.849 | 0.417 | 1.08E-40 |
| Cardiomyocyte | Adamtsl3 | 0 | 7.99E-43 | 0.40156693 | 0.863 | 0.478 | 1.60E-39 |
| Cardiomyocyte | Ca5b | 0 | 9.21E-43 | 0.85965215 | 0.585 | 0.152 | 1.84E-39 |
| Cardiomyocyte | Vcan | 0 | 3.90E-42 | 0.41059154 | 0.844 | 0.44 | 7.81E-39 |
| Cardiomyocyte | Agmo | 0 | 8.11E-42 | 0.27709178 | 0.844 | 0.524 | 1.62E-38 |
| Cardiomyocyte | Rasgef1b | 0 | 1.18E-40 | 0.86909464 | 0.745 | 0.258 | 2.36E-37 |
| Cardiomyocyte | Ccl2 | 0 | 4.49E-40 | 3.67483489 | 0.373 | 0.065 | 8.98E-37 |
| Cardiomyocyte | Rnf152 | 0 | 5.31E-40 | 0.83578009 | 0.811 | 0.344 | 1.06E-36 |
| Cardiomyocyte | Zfhx4 | 0 | 5.48E-40 | 2.13602474 | 0.373 | 0.065 | 1.10E-36 |
| Cardiomyocyte | Tbc1d1 | 0 | 8.80E-40 | 0.26352982 | 0.854 | 0.393 | 1.76E-36 |
| Cardiomyocyte | Postn | 0 | 9.29E-40 | 0.73419985 | 0.708 | 0.249 | 1.86E-36 |
| Cardiomyocyte | Alk | 0 | 1.25E-39 | 0.91857399 | 0.689 | 0.221 | 2.50E-36 |

| | | | | | | | |
|---|---|---|---|---|---|---|---|
| Cardiomyocyte | Niban1 | 0 | 2.15E-39 | 0.79986023 | 0.854 | 0.398 | 4.30E-36 |
| Cardiomyocyte | Ppm1h | 0 | 1.22E-38 | 0.60311214 | 0.844 | 0.418 | 2.43E-35 |
| Cardiomyocyte | C3 | 0 | 6.88E-38 | 1.75260982 | 0.34 | 0.054 | 1.38E-34 |
| Cardiomyocyte | Dnajc6 | 0 | 9.29E-37 | 0.76552446 | 0.377 | 0.07 | 1.86E-33 |
| Cardiomyocyte | Cd55 | 0 | 1.31E-36 | 0.28013762 | 0.783 | 0.358 | 2.62E-33 |
| Cardiomyocyte | B3galt2 | 0 | 1.32E-36 | 0.4169696 | 0.778 | 0.309 | 2.64E-33 |
| Cardiomyocyte | Gadd45b | 0 | 3.08E-36 | 0.59759398 | 0.816 | 0.298 | 6.17E-33 |
| Cardiomyocyte | Tdrd12 | 0 | 5.83E-36 | 1.76529499 | 0.585 | 0.165 | 1.17E-32 |
| Cardiomyocyte | Robo2 | 0 | 7.15E-36 | 1.83391815 | 0.665 | 0.189 | 1.43E-32 |
| Cardiomyocyte | Adam23 | 0 | 2.67E-35 | 0.47506585 | 0.41 | 0.149 | 5.33E-32 |
| Cardiomyocyte | Cemip | 0 | 1.06E-34 | 0.81423247 | 0.547 | 0.145 | 2.11E-31 |
| Cardiomyocyte | Igf1 | 0 | 5.92E-34 | 0.3699469 | 0.84 | 0.455 | 1.18E-30 |
| Cardiomyocyte | Gabrb1 | 0 | 1.07E-33 | 1.35829836 | 0.585 | 0.199 | 2.13E-30 |
| Cardiomyocyte | Dpt | 0 | 1.23E-33 | 0.64122751 | 0.844 | 0.415 | 2.46E-30 |

| | | | | | | | |
|---|---|---|---|---|---|---|---|
| Cardiomyocyte | Rxrg | 0 | 1.77E-33 | 0.97265402 | 0.868 | 0.517 | 3.53E-30 |
| Cardiomyocyte | Opcml | 0 | 1.80E-33 | 0.52607511 | 0.755 | 0.392 | 3.61E-30 |
| Cardiomyocyte | Pdgfd | 0 | 2.32E-33 | 0.51477925 | 0.882 | 0.546 | 4.64E-30 |
| Cardiomyocyte | Mrc2 | 0 | 7.66E-33 | 0.54645111 | 0.825 | 0.471 | 1.53E-29 |
| Cardiomyocyte | Adamts5 | 0 | 8.25E-33 | 0.83317404 | 0.849 | 0.45 | 1.65E-29 |
| Cardiomyocyte | Bard1 | 0 | 5.83E-32 | 1.89252089 | 0.797 | 0.341 | 1.17E-28 |
| Cardiomyocyte | Nhs | 0 | 1.09E-31 | 0.84320757 | 0.571 | 0.22 | 2.18E-28 |
| Cardiomyocyte | Gria4 | 0 | 5.86E-31 | 0.72228568 | 0.755 | 0.36 | 1.17E-27 |
| Cardiomyocyte | Mrc1 | 0 | 1.23E-30 | 0.55684736 | 0.84 | 0.37 | 2.46E-27 |
| Cardiomyocyte | Gpc3 | 0 | 1.91E-30 | 3.02564434 | 0.373 | 0.092 | 3.81E-27 |
| Cardiomyocyte | Fam189a1 | 0 | 1.97E-30 | 1.76150263 | 0.825 | 0.485 | 3.93E-27 |
| Cardiomyocyte | Adam12 | 0 | 1.98E-30 | 0.42735804 | 0.797 | 0.365 | 3.96E-27 |
| Cardiomyocyte | Pde1a | 0 | 3.95E-30 | 1.08735617 | 0.835 | 0.437 | 7.89E-27 |
| Cardiomyocyte | Il1rapl2 | 0 | 4.52E-30 | 1.27008811 | 0.401 | 0.141 | 9.04E-27 |

| | | | | | | | |
|---|---|---|---|---|---|---|---|
| Cardiomyocyte | Xkr4 | 0 | 4.83E-30 | 0.65243335 | 0.811 | 0.438 | 9.67E-27 |
| Cardiomyocyte | Apoe | 0 | 8.16E-30 | 0.83710174 | 0.802 | 0.442 | 1.63E-26 |
| Cardiomyocyte | Runx2 | 0 | 1.11E-29 | 0.5943527 | 0.651 | 0.292 | 2.23E-26 |
| Cardiomyocyte | Nr4a1 | 0 | 1.82E-29 | 0.89223162 | 0.873 | 0.559 | 3.64E-26 |
| Cardiomyocyte | Sh3pxd2b | 0 | 2.35E-29 | 0.31032836 | 0.783 | 0.45 | 4.69E-26 |
| Cardiomyocyte | Neb | 0 | 8.91E-29 | 0.29400944 | 0.849 | 0.468 | 1.78E-25 |
| Cardiomyocyte | Atp5me | 0 | 4.42E-28 | 0.77718385 | 0.873 | 0.432 | 8.84E-25 |
| Cardiomyocyte | Mertk | 0 | 1.01E-27 | 0.41184387 | 0.618 | 0.164 | 2.02E-24 |
| Cardiomyocyte | Casc4 | 0 | 1.06E-27 | 0.62432198 | 0.792 | 0.383 | 2.12E-24 |
| Cardiomyocyte | Il7 | 0 | 1.97E-27 | 0.42339876 | 0.425 | 0.08 | 3.94E-24 |
| Cardiomyocyte | Eda | 0 | 2.75E-27 | 0.28247911 | 0.849 | 0.424 | 5.51E-24 |
| Cardiomyocyte | Uap1 | 0 | 2.87E-27 | 0.53865539 | 0.769 | 0.346 | 5.74E-24 |
| Cardiomyocyte | Sned1 | 0 | 4.75E-27 | 0.82040236 | 0.736 | 0.292 | 9.51E-24 |
| Cardiomyocyte | Epha3 | 0 | 5.20E-27 | 0.89489838 | 0.844 | 0.475 | 1.04E-23 |

| | | | | | | | |
|---|---|---|---|---|---|---|---|
| Cardiomyocyte | Magi2 | 0 | 7.07E-27 | 0.82264257 | 0.656 | 0.237 | 1.41E-23 |
| Cardiomyocyte | AABR07068046.1 | 0 | 8.11E-27 | 1.11395707 | 0.623 | 0.324 | 1.62E-23 |
| Cardiomyocyte | Galnt17 | 0 | 3.36E-26 | 0.9858427 | 0.83 | 0.389 | 6.72E-23 |
| Cardiomyocyte | Zfp366 | 0 | 9.96E-26 | 0.81578191 | 0.835 | 0.4 | 1.99E-22 |
| Cardiomyocyte | Col1a1 | 0 | 1.18E-25 | 0.5335409 | 0.858 | 0.502 | 2.35E-22 |
| Cardiomyocyte | Anks1b | 0 | 1.33E-25 | 0.29273263 | 0.854 | 0.439 | 2.66E-22 |
| Cardiomyocyte | Tmem50b | 0 | 1.70E-25 | 0.37576583 | 0.858 | 0.438 | 3.39E-22 |
| Cardiomyocyte | Plod2 | 0 | 6.87E-25 | 0.30171444 | 0.863 | 0.493 | 1.37E-21 |
| Cardiomyocyte | Camk1d | 0 | 7.45E-25 | 0.33695796 | 0.816 | 0.421 | 1.49E-21 |
| Cardiomyocyte | Adamts2 | 0 | 1.93E-24 | 0.59881827 | 0.797 | 0.431 | 3.86E-21 |
| Cardiomyocyte | Tpx2 | 0 | 3.17E-24 | 0.27008391 | 0.585 | 0.23 | 6.34E-21 |
| Cardiomyocyte | Hs3st3a1 | 0 | 5.46E-24 | 2.20285976 | 0.486 | 0.107 | 1.09E-20 |
| Cardiomyocyte | Hmga2 | 0 | 6.17E-24 | 2.53350513 | 0.373 | 0.117 | 1.23E-20 |
| Cardiomyocyte | Aff3 | 0 | 8.45E-24 | 0.3988619 | 0.854 | 0.492 | 1.69E-20 |

| | | | | | | | |
|---|---|---|---|---|---|---|---|
| Cardiomyocyte | AC123500.1 | 0 | 8.68E-24 | 1.18491216 | 0.637 | 0.193 | 1.74E-20 |
| Cardiomyocyte | Nek10 | 0 | 1.04E-23 | 0.9161926 | 0.524 | 0.224 | 2.08E-20 |
| Cardiomyocyte | Medag | 0 | 1.10E-23 | 0.69008556 | 0.623 | 0.309 | 2.19E-20 |
| Cardiomyocyte | Cdh6 | 0 | 1.84E-23 | 1.02369744 | 0.453 | 0.143 | 3.68E-20 |
| Cardiomyocyte | Hlf | 0 | 1.09E-22 | 0.37259839 | 0.675 | 0.272 | 2.18E-19 |
| Cardiomyocyte | Cox6a2 | 0 | 2.24E-22 | 0.46164165 | 0.858 | 0.531 | 4.49E-19 |
| Cardiomyocyte | LOC690045 | 0 | 2.54E-22 | 1.55862953 | 0.392 | 0.118 | 5.09E-19 |
| Cardiomyocyte | Ube2ql1 | 0 | 2.83E-22 | 0.59369758 | 0.854 | 0.424 | 5.67E-19 |
| Cardiomyocyte | AABR07040864.1 | 0 | 6.47E-22 | 0.84377352 | 0.788 | 0.434 | 1.29E-18 |
| Cardiomyocyte | Mamdc2 | 0 | 2.09E-21 | 0.48508352 | 0.816 | 0.438 | 4.18E-18 |
| Cardiomyocyte | Luzp2 | 0 | 2.56E-21 | 1.40620321 | 0.604 | 0.202 | 5.12E-18 |
| Cardiomyocyte | Kif21a | 0 | 1.15E-20 | 0.82906798 | 0.84 | 0.459 | 2.30E-17 |
| Cardiomyocyte | Drc3 | 0 | 1.22E-20 | 0.92307654 | 0.821 | 0.465 | 2.43E-17 |
| Cardiomyocyte | Map2 | 0 | 1.12E-19 | 1.61335481 | 0.575 | 0.262 | 2.24E-16 |

| | | | | | | | |
|---|---|---|---|---|---|---|---|
| Cardiomyocyte | AC130940.1 | 0 | 2.87E-18 | 1.13670716 | 0.613 | 0.333 | 5.74E-15 |
| Cardiomyocyte | Rgs5 | 0 | 6.09E-18 | 0.28418757 | 0.679 | 0.351 | 1.22E-14 |
| Cardiomyocyte | Apoo | 0 | 1.01E-17 | 0.68593973 | 0.684 | 0.352 | 2.03E-14 |
| Cardiomyocyte | Sctr | 0 | 1.32E-17 | 0.36660124 | 0.84 | 0.451 | 2.64E-14 |
| Cardiomyocyte | Svep1 | 0 | 1.58E-17 | 1.18121458 | 0.665 | 0.408 | 3.16E-14 |
| Cardiomyocyte | Nlgn1 | 0 | 2.04E-17 | 0.68952631 | 0.524 | 0.189 | 4.08E-14 |
| Cardiomyocyte | Egflam | 0 | 2.82E-17 | 0.64536157 | 0.83 | 0.445 | 5.64E-14 |
| Cardiomyocyte | Slfn4 | 0 | 4.50E-17 | 1.0173716 | 0.66 | 0.396 | 8.99E-14 |
| Cardiomyocyte | Hdx | 0 | 6.83E-17 | 0.61228167 | 0.495 | 0.192 | 1.37E-13 |
| Cardiomyocyte | Stab1 | 0 | 7.50E-17 | 0.29096841 | 0.443 | 0.125 | 1.50E-13 |
| Cardiomyocyte | Dmtn | 0 | 8.28E-17 | 0.44131327 | 0.024 | 0.297 | 1.66E-13 |
| Cardiomyocyte | AABR07052523.2 | 0 | 9.33E-17 | 1.0468468 | 0.731 | 0.404 | 1.87E-13 |
| Cardiomyocyte | Kcnq3 | 0 | 2.15E-16 | 1.1499265 | 0.458 | 0.198 | 4.31E-13 |
| Cardiomyocyte | Mbp | 0 | 6.09E-16 | 0.75921628 | 0.429 | 0.178 | 1.22E-12 |

| | | | | | | | |
|---|---|---|---|---|---|---|---|
| Cardiomyocyte | Ablim2 | 0 | 1.66E-15 | 0.25362148 | 0.835 | 0.452 | 3.33E-12 |
| Cardiomyocyte | Atp5mc3 | 0 | 2.94E-15 | 0.63858763 | 0.868 | 0.583 | 5.87E-12 |
| Cardiomyocyte | Slamf8 | 0 | 4.70E-14 | 0.34086844 | 0.425 | 0.172 | 9.39E-11 |
| Cardiomyocyte | Prph | 0 | 9.54E-14 | 0.73598003 | 0.415 | 0.141 | 1.91E-10 |
| Cardiomyocyte | Fcer1g | 0 | 9.83E-14 | 0.49020228 | 0.835 | 0.416 | 1.97E-10 |
| Cardiomyocyte | Pstpip2 | 0 | 2.19E-13 | 0.6335284 | 0.792 | 0.45 | 4.39E-10 |
| Cardiomyocyte | Adamtsl2 | 0 | 1.57E-12 | 0.26904233 | 0.759 | 0.406 | 3.15E-09 |
| Cardiomyocyte | Mtus2 | 0 | 1.65E-12 | 0.93904117 | 0.868 | 0.569 | 3.31E-09 |
| Cardiomyocyte | Sorcs2 | 0 | 1.75E-12 | 0.4716885 | 0.656 | 0.28 | 3.51E-09 |
| Cardiomyocyte | PCOLCE2 | 0 | 3.37E-12 | 0.45033194 | 0.788 | 0.456 | 6.73E-09 |
| Cardiomyocyte | Asb14 | 0 | 2.58E-11 | 0.26951652 | 0.825 | 0.551 | 5.17E-08 |
| Cardiomyocyte | Bicc1 | 0 | 4.21E-11 | 0.34792564 | 0.835 | 0.58 | 8.42E-08 |
| Cardiomyocyte | Esr1 | 0 | 1.31E-10 | 0.34218248 | 0.717 | 0.445 | 2.62E-07 |
| Cardiomyocyte | Unc45b | 0 | 2.04E-10 | 0.67317501 | 0.736 | 0.474 | 4.08E-07 |

| | | | | | | | |
|---|---|---|---|---|---|---|---|
| Cardiomyocyte | Fmod | 0 | 2.85E-09 | 1.28514003 | 0.387 | 0.094 | 5.71E-06 |
| Cardiomyocyte | AABR07052585.2 | 0 | 7.48E-08 | 0.49879969 | 0.882 | 0.588 | 0.00014959 |
| Cardiomyocyte | Col6a6 | 0 | 6.31E-07 | 1.06063106 | 0.448 | 0.185 | 0.0012615 |
| Cardiomyocyte | Fcgr2a | 1 | 2.72E-104 | 4.85029833 | 0.523 | 0.025 | 5.44E-101 |
| Cardiomyocyte | Plxnb3 | 1 | 1.78E-101 | 2.026318 | 0.671 | 0.07 | 3.57E-98 |
| Cardiomyocyte | Clec4e | 1 | 2.58E-101 | 3.49271148 | 0.671 | 0.07 | 5.15E-98 |
| Cardiomyocyte | AABR07034457.1 | 1 | 1.37E-98 | 3.46066407 | 0.497 | 0.024 | 2.74E-95 |
| Cardiomyocyte | Tsbp1 | 1 | 1.38E-98 | 1.4791772 | 0.497 | 0.024 | 2.76E-95 |
| Cardiomyocyte | Ttyh1 | 1 | 5.21E-96 | 1.90377972 | 0.497 | 0.025 | 1.04E-92 |
| Cardiomyocyte | Alox5ap | 1 | 4.25E-90 | 2.72173768 | 0.69 | 0.088 | 8.51E-87 |
| Cardiomyocyte | Sbspon | 1 | 4.51E-90 | 2.96781545 | 0.748 | 0.132 | 9.02E-87 |
| Cardiomyocyte | Stk31 | 1 | 4.21E-84 | 2.68500378 | 0.484 | 0.023 | 8.42E-81 |
| Cardiomyocyte | Insc | 1 | 2.80E-81 | 3.69037597 | 0.645 | 0.041 | 5.59E-78 |
| Cardiomyocyte | Cps1 | 1 | 1.72E-80 | 4.22485147 | 0.645 | 0.095 | 3.45E-77 |

| | | | | | | | |
|---|---|---|---|---|---|---|---|
| Cardiomyocyte | 5330417C22Rik | 1 | 2.83E-78 | 2.89460753 | 0.755 | 0.148 | 5.65E-75 |
| Cardiomyocyte | Arnt2 | 1 | 3.43E-72 | 0.84834749 | 0.671 | 0.108 | 6.86E-69 |
| Cardiomyocyte | Tc2n | 1 | 8.49E-71 | 3.41683085 | 0.755 | 0.133 | 1.70E-67 |
| Cardiomyocyte | Cenpe | 1 | 1.73E-69 | 1.39232437 | 0.394 | 0.031 | 3.47E-66 |
| Cardiomyocyte | Gcat | 1 | 1.17E-68 | 2.11919411 | 0.735 | 0.168 | 2.35E-65 |
| Cardiomyocyte | Susd5 | 1 | 4.49E-68 | 2.54896056 | 0.723 | 0.09 | 8.98E-65 |
| Cardiomyocyte | Hs6st2 | 1 | 1.50E-67 | 1.30987957 | 0.742 | 0.179 | 3.00E-64 |
| Cardiomyocyte | Itga10 | 1 | 1.98E-66 | 2.48067211 | 0.406 | 0.031 | 3.95E-63 |
| Cardiomyocyte | AABR07054000.1 | 1 | 3.46E-66 | 1.73791814 | 0.729 | 0.145 | 6.93E-63 |
| Cardiomyocyte | Cdh3 | 1 | 2.04E-63 | 1.83991581 | 0.613 | 0.1 | 4.07E-60 |
| Cardiomyocyte | Kcnn2 | 1 | 2.70E-63 | 2.17587866 | 0.497 | 0.027 | 5.40E-60 |
| Cardiomyocyte | Scube3 | 1 | 8.99E-63 | 2.81541554 | 0.626 | 0.135 | 1.80E-59 |
| Cardiomyocyte | Ubap1l | 1 | 3.57E-59 | 0.89227002 | 0.645 | 0.084 | 7.15E-56 |
| Cardiomyocyte | Lrrc63 | 1 | 4.39E-58 | 1.24457743 | 0.484 | 0.025 | 8.77E-55 |

| | | | | | | | |
|---|---|---|---|---|---|---|---|
| Cardiomyocyte | Esco2 | 1 | 4.86E-58 | 0.82226401 | 0.587 | 0.091 | 9.72E-55 |
| Cardiomyocyte | Chi3l1 | 1 | 3.64E-56 | 2.99890704 | 0.535 | 0.101 | 7.28E-53 |
| Cardiomyocyte | Col9a1 | 1 | 3.99E-56 | 2.05542766 | 0.645 | 0.142 | 7.98E-53 |
| Cardiomyocyte | AABR07030527.1 | 1 | 2.43E-55 | 3.35008657 | 0.761 | 0.286 | 4.86E-52 |
| Cardiomyocyte | Galnt14 | 1 | 7.54E-54 | 1.91895106 | 0.613 | 0.119 | 1.51E-50 |
| Cardiomyocyte | Tas2r121 | 1 | 3.55E-53 | 0.91970081 | 0.619 | 0.138 | 7.09E-50 |
| Cardiomyocyte | Mapk10 | 1 | 4.92E-53 | 1.47062485 | 0.671 | 0.09 | 9.84E-50 |
| Cardiomyocyte | Ckap2 | 1 | 2.98E-52 | 3.17982068 | 0.723 | 0.128 | 5.96E-49 |
| Cardiomyocyte | Tnc | 1 | 7.06E-52 | 3.83400093 | 0.626 | 0.121 | 1.41E-48 |
| Cardiomyocyte | Crlf1 | 1 | 1.48E-51 | 0.99199548 | 0.587 | 0.108 | 2.97E-48 |
| Cardiomyocyte | LOC680920 | 1 | 1.58E-50 | 1.17672168 | 0.535 | 0.114 | 3.16E-47 |
| Cardiomyocyte | Ndst3 | 1 | 1.06E-48 | 1.037749 | 0.619 | 0.09 | 2.13E-45 |
| Cardiomyocyte | Dscaml1 | 1 | 1.03E-46 | 2.31013517 | 0.6 | 0.128 | 2.05E-43 |
| Cardiomyocyte | Kcna2 | 1 | 1.58E-46 | 2.33402876 | 0.658 | 0.164 | 3.15E-43 |

| | | | | | | | |
|---|---|---|---|---|---|---|---|
| Cardiomyocyte | Ect2 | 1 | 1.65E-45 | 2.0580532 | 0.458 | 0.147 | 3.29E-42 |
| Cardiomyocyte | Aldh1a2 | 1 | 5.55E-45 | 1.41332551 | 0.529 | 0.059 | 1.11E-41 |
| Cardiomyocyte | Itgb4 | 1 | 7.35E-45 | 0.34015233 | 0.729 | 0.243 | 1.47E-41 |
| Cardiomyocyte | Trpc3 | 1 | 5.93E-44 | 0.83431875 | 0.639 | 0.148 | 1.19E-40 |
| Cardiomyocyte | AABR07026483.1 | 1 | 1.76E-43 | 1.51039031 | 0.613 | 0.135 | 3.53E-40 |
| Cardiomyocyte | Arrdc5 | 1 | 1.75E-42 | 3.10932693 | 0.381 | 0.055 | 3.50E-39 |
| Cardiomyocyte | Mafb | 1 | 2.47E-41 | 0.25964147 | 0.71 | 0.192 | 4.95E-38 |
| Cardiomyocyte | Fosl1 | 1 | 7.82E-41 | 1.63634454 | 0.742 | 0.365 | 1.56E-37 |
| Cardiomyocyte | Slc24a2 | 1 | 9.29E-40 | 2.81142115 | 0.6 | 0.154 | 1.86E-36 |
| Cardiomyocyte | AABR07001573.2 | 1 | 4.66E-38 | 1.12070406 | 0.645 | 0.247 | 9.33E-35 |
| Cardiomyocyte | Gng10 | 1 | 8.38E-38 | 0.93668913 | 0.748 | 0.208 | 1.68E-34 |
| Cardiomyocyte | Lepr | 1 | 1.44E-37 | 1.13261803 | 0.748 | 0.295 | 2.88E-34 |
| Cardiomyocyte | Wnt5b | 1 | 1.46E-36 | 1.3487676 | 0.574 | 0.224 | 2.91E-33 |
| Cardiomyocyte | Steap4 | 1 | 5.14E-36 | 0.70475629 | 0.716 | 0.281 | 1.03E-32 |

| | | | | | | | |
|---|---|---|---|---|---|---|---|
| Cardiomyocyte | Kcng3 | 1 | 3.95E-35 | 0.97422624 | 0.523 | 0.083 | 7.90E-32 |
| Cardiomyocyte | Col28a1 | 1 | 2.21E-34 | 0.83004877 | 0.729 | 0.294 | 4.43E-31 |
| Cardiomyocyte | Mki67 | 1 | 9.66E-34 | 0.41272122 | 0.555 | 0.141 | 1.93E-30 |
| Cardiomyocyte | Pax3 | 1 | 4.26E-33 | 0.75324328 | 0.387 | 0.074 | 8.53E-30 |
| Cardiomyocyte | AC118957.1 | 1 | 6.19E-33 | 0.53273537 | 0.645 | 0.206 | 1.24E-29 |
| Cardiomyocyte | Angptl1 | 1 | 1.29E-32 | 0.89987751 | 0.652 | 0.175 | 2.59E-29 |
| Cardiomyocyte | Edil3 | 1 | 1.45E-32 | 1.12053181 | 0.723 | 0.309 | 2.90E-29 |
| Cardiomyocyte | Aff31 | 1 | 5.34E-32 | 1.40260809 | 0.794 | 0.516 | 1.07E-28 |
| Cardiomyocyte | Olfm2 | 1 | 3.72E-31 | 2.40274638 | 0.329 | 0.049 | 7.44E-28 |
| Cardiomyocyte | Mctp2 | 1 | 3.89E-31 | 1.00354218 | 0.735 | 0.438 | 7.78E-28 |
| Cardiomyocyte | Hs3st2 | 1 | 1.20E-30 | 1.06019922 | 0.426 | 0.158 | 2.41E-27 |
| Cardiomyocyte | Aoah | 1 | 2.15E-30 | 1.23652934 | 0.529 | 0.125 | 4.31E-27 |
| Cardiomyocyte | Cd83 | 1 | 2.39E-30 | 1.64520205 | 0.697 | 0.378 | 4.78E-27 |
| Cardiomyocyte | Rd3l | 1 | 4.83E-30 | 1.69473932 | 0.742 | 0.241 | 9.65E-27 |

| | | | | | | | |
|---|---|---|---|---|---|---|---|
| Cardiomyocyte | AABR07032261.1 | 1 | 5.36E-30 | 1.48062852 | 0.458 | 0.11 | 1.07E-26 |
| Cardiomyocyte | AABR07037343.1 | 1 | 9.68E-30 | 1.72849036 | 0.432 | 0.103 | 1.94E-26 |
| Cardiomyocyte | Gba3 | 1 | 3.78E-29 | 0.84945602 | 0.755 | 0.504 | 7.57E-26 |
| Cardiomyocyte | Itga8 | 1 | 5.59E-29 | 0.8204237 | 0.755 | 0.457 | 1.12E-25 |
| Cardiomyocyte | Fcer1g1 | 1 | 4.52E-28 | 0.88654144 | 0.742 | 0.446 | 9.05E-25 |
| Cardiomyocyte | Kcnh8 | 1 | 8.30E-28 | 1.20601865 | 0.619 | 0.147 | 1.66E-24 |
| Cardiomyocyte | Lrsam1 | 1 | 3.53E-27 | 0.79063006 | 0.561 | 0.14 | 7.06E-24 |
| Cardiomyocyte | Cubn | 1 | 8.66E-27 | 0.95644419 | 0.548 | 0.16 | 1.73E-23 |
| Cardiomyocyte | Meox1 | 1 | 1.83E-26 | 0.28110369 | 0.613 | 0.204 | 3.66E-23 |
| Cardiomyocyte | Pappa2 | 1 | 2.10E-26 | 0.47682043 | 0.639 | 0.251 | 4.21E-23 |
| Cardiomyocyte | Grip2 | 1 | 3.33E-26 | 1.51311233 | 0.748 | 0.357 | 6.66E-23 |
| Cardiomyocyte | Nlgn11 | 1 | 5.80E-26 | 1.21468262 | 0.477 | 0.21 | 1.16E-22 |
| Cardiomyocyte | Cdon | 1 | 1.33E-25 | 0.58853313 | 0.748 | 0.342 | 2.66E-22 |
| Cardiomyocyte | AABR07027925.1 | 1 | 2.00E-25 | 3.23505305 | 0.381 | 0.017 | 3.99E-22 |

| | | | | | | | |
|---|---|---|---|---|---|---|---|
| Cardiomyocyte | Chrdl1 | 1 | 2.30E-25 | 0.26377644 | 0.626 | 0.218 | 4.59E-22 |
| Cardiomyocyte | Slc1a1 | 1 | 1.90E-24 | 0.72581569 | 0.755 | 0.397 | 3.80E-21 |
| Cardiomyocyte | Adamts3 | 1 | 3.55E-24 | 1.31973265 | 0.665 | 0.356 | 7.09E-21 |
| Cardiomyocyte | Lck | 1 | 6.72E-24 | 0.68217575 | 0.748 | 0.373 | 1.34E-20 |
| Cardiomyocyte | Fads6 | 1 | 9.66E-24 | 2.65582134 | 0.548 | 0.187 | 1.93E-20 |
| Cardiomyocyte | Crispld2 | 1 | 2.12E-23 | 0.50406586 | 0.755 | 0.379 | 4.24E-20 |
| Cardiomyocyte | Ifit1bl | 1 | 5.18E-23 | 1.05548731 | 0.723 | 0.39 | 1.04E-19 |
| Cardiomyocyte | Ptprt | 1 | 5.36E-23 | 0.44299989 | 0.748 | 0.334 | 1.07E-19 |
| Cardiomyocyte | Masp1 | 1 | 7.87E-23 | 0.88036152 | 0.787 | 0.445 | 1.57E-19 |
| Cardiomyocyte | Apeg3 | 1 | 1.03E-22 | 1.22325507 | 0.684 | 0.395 | 2.06E-19 |
| Cardiomyocyte | Klhl4 | 1 | 1.23E-22 | 0.62297086 | 0.748 | 0.328 | 2.45E-19 |
| Cardiomyocyte | Aox3 | 1 | 1.43E-22 | 0.3710629 | 0.735 | 0.395 | 2.87E-19 |
| Cardiomyocyte | Sgpp2 | 1 | 1.54E-22 | 1.78312185 | 0.374 | 0.093 | 3.07E-19 |
| Cardiomyocyte | Skap1 | 1 | 1.68E-22 | 1.98015784 | 0.594 | 0.326 | 3.36E-19 |

| | | | | | | | |
|---|---|---|---|---|---|---|---|
| Cardiomyocyte | Ngf | 1 | 2.07E-22 | 0.45172718 | 0.697 | 0.297 | 4.13E-19 |
| Cardiomyocyte | Cyp2e1 | 1 | 2.11E-22 | 0.49449682 | 0.658 | 0.161 | 4.22E-19 |
| Cardiomyocyte | AABR07052441.1 | 1 | 6.52E-22 | 1.60160822 | 0.594 | 0.3 | 1.30E-18 |
| Cardiomyocyte | Nrk | 1 | 8.06E-22 | 0.94203848 | 0.329 | 0.053 | 1.61E-18 |
| Cardiomyocyte | Map6 | 1 | 1.35E-21 | 0.69678786 | 0.761 | 0.305 | 2.69E-18 |
| Cardiomyocyte | Brca1 | 1 | 1.67E-21 | 1.22153205 | 0.445 | 0.173 | 3.34E-18 |
| Cardiomyocyte | Lrrtm3 | 1 | 5.66E-21 | 0.35911374 | 0.806 | 0.515 | 1.13E-17 |
| Cardiomyocyte | Ephb1 | 1 | 8.96E-21 | 0.47552899 | 0.432 | 0.155 | 1.79E-17 |
| Cardiomyocyte | Ikzf3 | 1 | 1.83E-19 | 1.31433174 | 0.703 | 0.29 | 3.67E-16 |
| Cardiomyocyte | Adgrl3 | 1 | 2.17E-19 | 0.51019893 | 0.755 | 0.387 | 4.33E-16 |
| Cardiomyocyte | Dkk3 | 1 | 4.37E-19 | 0.93898158 | 0.735 | 0.38 | 8.74E-16 |
| Cardiomyocyte | Tnfrsf19 | 1 | 5.33E-19 | 1.04637208 | 0.768 | 0.457 | 1.07E-15 |
| Cardiomyocyte | Ntrk2 | 1 | 5.71E-19 | 0.34311508 | 0.594 | 0.209 | 1.14E-15 |
| Cardiomyocyte | Blnk | 1 | 6.83E-19 | 0.84456163 | 0.723 | 0.42 | 1.37E-15 |

| | | | | | | | |
|---|---|---|---|---|---|---|---|
| Cardiomyocyte | Aldh1a1 | 1 | 7.02E-19 | 0.52555456 | 0.755 | 0.411 | 1.40E-15 |
| Cardiomyocyte | Fmo2 | 1 | 7.84E-19 | 0.32057347 | 0.542 | 0.215 | 1.57E-15 |
| Cardiomyocyte | Lekr1 | 1 | 2.33E-18 | 0.63438026 | 0.839 | 0.547 | 4.66E-15 |
| Cardiomyocyte | Prss48 | 1 | 4.96E-18 | 0.491709 | 0.387 | 0.126 | 9.92E-15 |
| Cardiomyocyte | Slc44a5 | 1 | 5.26E-18 | 0.50083402 | 0.626 | 0.279 | 1.05E-14 |
| Cardiomyocyte | Serf1 | 1 | 1.40E-17 | 1.33334326 | 0.703 | 0.344 | 2.80E-14 |
| Cardiomyocyte | Atp10a | 1 | 1.44E-17 | 1.57240079 | 0.729 | 0.466 | 2.88E-14 |
| Cardiomyocyte | Lsamp | 1 | 1.67E-17 | 1.1389838 | 0.852 | 0.497 | 3.33E-14 |
| Cardiomyocyte | Matn2 | 1 | 1.87E-17 | 0.31469503 | 0.665 | 0.296 | 3.73E-14 |
| Cardiomyocyte | Ano5 | 1 | 2.20E-17 | 1.12164719 | 0.761 | 0.364 | 4.40E-14 |
| Cardiomyocyte | Nudt4 | 1 | 2.30E-17 | 0.88934506 | 0.671 | 0.253 | 4.60E-14 |
| Cardiomyocyte | Iqgap2 | 1 | 4.09E-17 | 1.06332502 | 0.839 | 0.547 | 8.19E-14 |
| Cardiomyocyte | Pcp4l1 | 1 | 6.10E-17 | 1.0765079 | 0.626 | 0.214 | 1.22E-13 |
| Cardiomyocyte | Zfp536 | 1 | 6.22E-17 | 0.55751216 | 0.723 | 0.329 | 1.24E-13 |

| | | | | | | | |
|---|---|---|---|---|---|---|---|
| Cardiomyocyte | Grid1 | 1 | 1.04E-16 | 0.56334881 | 0.723 | 0.312 | 2.07E-13 |
| Cardiomyocyte | Dpt1 | 1 | 3.70E-16 | 0.53098771 | 0.768 | 0.444 | 7.40E-13 |
| Cardiomyocyte | Slc25a21 | 1 | 5.50E-16 | 0.56584767 | 0.819 | 0.454 | 1.10E-12 |
| Cardiomyocyte | Abca1 | 1 | 5.86E-16 | 0.76252897 | 0.787 | 0.522 | 1.17E-12 |
| Cardiomyocyte | Vim | 1 | 1.49E-15 | 0.69547416 | 0.632 | 0.291 | 2.98E-12 |
| Cardiomyocyte | Gsn | 1 | 2.16E-15 | 0.3894709 | 0.787 | 0.439 | 4.32E-12 |
| Cardiomyocyte | Tafa2 | 1 | 3.17E-15 | 0.54423029 | 0.671 | 0.41 | 6.34E-12 |
| Cardiomyocyte | Ccser1 | 1 | 5.70E-15 | 0.2627287 | 0.677 | 0.279 | 1.14E-11 |
| Cardiomyocyte | Myo3b | 1 | 7.00E-15 | 0.88406359 | 0.697 | 0.404 | 1.40E-11 |
| Cardiomyocyte | Itga4 | 1 | 1.08E-14 | 1.24892626 | 0.465 | 0.146 | 2.16E-11 |
| Cardiomyocyte | Rhpn2 | 1 | 1.09E-14 | 0.55835007 | 0.432 | 0.127 | 2.18E-11 |
| Cardiomyocyte | Ppp1r3a | 1 | 1.73E-14 | 0.83574981 | 0.871 | 0.52 | 3.46E-11 |
| Cardiomyocyte | Adhfe1 | 1 | 3.26E-14 | 0.88101543 | 0.819 | 0.484 | 6.51E-11 |
| Cardiomyocyte | Ralgps2 | 1 | 1.06E-13 | 0.4696633 | 0.735 | 0.387 | 2.12E-10 |

| | | | | | | | |
|---|---|---|---|---|---|---|---|
| Cardiomyocyte | Itgal | 1 | 2.78E-13 | 1.08282391 | 0.703 | 0.303 | 5.55E-10 |
| Cardiomyocyte | Fign | 1 | 2.87E-13 | 0.62497016 | 0.897 | 0.538 | 5.74E-10 |
| Cardiomyocyte | Egr1 | 1 | 3.25E-13 | 0.89212035 | 0.548 | 0.298 | 6.50E-10 |
| Cardiomyocyte | Ngdn | 1 | 3.42E-13 | 0.52719183 | 0.787 | 0.366 | 6.84E-10 |
| Cardiomyocyte | Slc25a20 | 1 | 4.78E-13 | 0.50154432 | 0.884 | 0.621 | 9.55E-10 |
| Cardiomyocyte | Xirp1 | 1 | 5.40E-13 | 0.3097961 | 0.716 | 0.304 | 1.08E-09 |
| Cardiomyocyte | Pparg | 1 | 5.80E-13 | 0.51642527 | 0.768 | 0.465 | 1.16E-09 |
| Cardiomyocyte | Anxa1 | 1 | 5.91E-13 | 0.61960175 | 0.626 | 0.333 | 1.18E-09 |
| Cardiomyocyte | Gal3st3 | 1 | 6.23E-13 | 0.55571963 | 0.794 | 0.462 | 1.25E-09 |
| Cardiomyocyte | Nr4a3 | 1 | 8.38E-13 | 2.33669257 | 0.535 | 0.262 | 1.68E-09 |
| Cardiomyocyte | Npas2 | 1 | 1.43E-12 | 0.36493065 | 0.652 | 0.331 | 2.85E-09 |
| Cardiomyocyte | Glb1l2 | 1 | 6.20E-12 | 0.68743962 | 0.794 | 0.445 | 1.24E-08 |
| Cardiomyocyte | Cryab | 1 | 6.29E-12 | 0.61033815 | 0.748 | 0.362 | 1.26E-08 |
| Cardiomyocyte | Prox1 | 1 | 1.37E-11 | 0.64161119 | 0.865 | 0.589 | 2.74E-08 |

| | | | | | | | |
|---|---|---|---|---|---|---|---|
| Cardiomyocyte | Lrmp | 1 | 1.48E-11 | 1.48602915 | 0.445 | 0.178 | 2.97E-08 |
| Cardiomyocyte | Fam228b | 1 | 2.08E-11 | 0.88771313 | 0.51 | 0.245 | 4.15E-08 |
| Cardiomyocyte | Pik3ap1 | 1 | 2.23E-11 | 0.71358032 | 0.852 | 0.586 | 4.46E-08 |
| Cardiomyocyte | C7 | 1 | 2.69E-11 | 0.37587965 | 0.703 | 0.386 | 5.38E-08 |
| Cardiomyocyte | Pla2g7 | 1 | 2.88E-11 | 0.57538536 | 0.748 | 0.488 | 5.76E-08 |
| Cardiomyocyte | Hk2 | 1 | 6.16E-11 | 0.45264943 | 0.852 | 0.496 | 1.23E-07 |
| Cardiomyocyte | Epha4 | 1 | 6.81E-11 | 0.33618885 | 0.742 | 0.471 | 1.36E-07 |
| Cardiomyocyte | Myh11 | 1 | 7.29E-11 | 0.28954115 | 0.781 | 0.474 | 1.46E-07 |
| Cardiomyocyte | Nrxn31 | 1 | 9.79E-11 | 0.93467042 | 0.748 | 0.484 | 1.96E-07 |
| Cardiomyocyte | Tp53inp2 | 1 | 1.44E-10 | 0.57672417 | 0.787 | 0.537 | 2.88E-07 |
| Cardiomyocyte | Coq8a | 1 | 1.75E-10 | 0.35593986 | 0.877 | 0.539 | 3.49E-07 |
| Cardiomyocyte | Ppp1r9a | 1 | 3.52E-10 | 0.31826476 | 0.787 | 0.508 | 7.04E-07 |
| Cardiomyocyte | Mpc1 | 1 | 9.72E-10 | 0.48401121 | 0.781 | 0.439 | 1.94E-06 |
| Cardiomyocyte | Me3 | 1 | 1.02E-09 | 0.61308799 | 0.884 | 0.632 | 2.04E-06 |

| | | | | | | | |
|---|---|---|---|---|---|---|---|
| Cardiomyocyte | Cobl | 1 | 3.19E-09 | 0.37201724 | 0.69 | 0.352 | 6.38E-06 |
| Cardiomyocyte | Gja1 | 1 | 3.35E-09 | 0.44657425 | 0.91 | 0.587 | 6.70E-06 |
| Cardiomyocyte | Sgca | 1 | 1.20E-08 | 0.47860164 | 0.832 | 0.531 | 2.40E-05 |
| Cardiomyocyte | Kcnma1 | 1 | 1.31E-08 | 0.26527157 | 0.497 | 0.247 | 2.63E-05 |
| Cardiomyocyte | Stac3 | 1 | 1.83E-08 | 0.87298502 | 0.039 | 0.306 | 3.66E-05 |
| Cardiomyocyte | Ky | 1 | 2.01E-08 | 0.68029814 | 0.697 | 0.433 | 4.02E-05 |
| Cardiomyocyte | Atp1a1 | 1 | 2.51E-08 | 0.50097504 | 0.858 | 0.567 | 5.02E-05 |
| Cardiomyocyte | Usp2 | 1 | 2.69E-08 | 0.32857593 | 0.768 | 0.467 | 5.39E-05 |
| Cardiomyocyte | Tmem17 | 1 | 3.13E-08 | 1.27844886 | 0.465 | 0.119 | 6.26E-05 |
| Cardiomyocyte | Rnf150 | 1 | 8.72E-08 | 0.45320887 | 0.716 | 0.41 | 0.00017444 |
| Cardiomyocyte | Pcbp3 | 1 | 2.34E-07 | 0.33517529 | 0.871 | 0.601 | 0.00046785 |
| Cardiomyocyte | Hecw2 | 1 | 1.87E-05 | 0.25615324 | 0.839 | 0.582 | 0.03748707 |
| Cardiomyocyte | Melk | 2 | 6.77E-40 | 2.04741606 | 0.288 | 0.029 | 1.35E-36 |
| Cardiomyocyte | Itk | 2 | 8.15E-31 | 2.0688081 | 0.346 | 0.067 | 1.63E-27 |

| | | | | | | | |
|---|---|---|---|---|---|---|---|
| Cardiomyocyte | Tnni2 | 2 | 1.40E-30 | 2.41712091 | 0.32 | 0.056 | 2.79E-27 |
| Cardiomyocyte | Kcna7 | 2 | 9.98E-27 | 4.46077099 | 0.386 | 0.091 | 2.00E-23 |
| Cardiomyocyte | Smc1b | 2 | 3.59E-22 | 1.37339726 | 0.399 | 0.125 | 7.18E-19 |
| Cardiomyocyte | Lhfpl4 | 2 | 8.94E-21 | 1.63746675 | 0.359 | 0.106 | 1.79E-17 |
| Cardiomyocyte | LOC100302465 | 2 | 5.03E-13 | 0.95518067 | 0.418 | 0.167 | 1.01E-09 |
| Cardiomyocyte | LOC6809201 | 2 | 9.07E-13 | 1.32802706 | 0.386 | 0.133 | 1.81E-09 |
| Cardiomyocyte | Hmmr | 2 | 1.60E-12 | 2.1428739 | 0.386 | 0.094 | 3.19E-09 |
| Cardiomyocyte | Cyp26b1 | 2 | 3.38E-12 | 1.02937093 | 0.013 | 0.275 | 6.76E-09 |
| Cardiomyocyte | Cilp | 2 | 3.90E-12 | 3.45733562 | 0.078 | 0.38 | 7.80E-09 |
| Cardiomyocyte | AABR07057510.3 | 2 | 2.84E-11 | 3.12539159 | 0.327 | 0.056 | 5.69E-08 |
| Cardiomyocyte | Rhpn21 | 2 | 2.86E-11 | 2.48826779 | 0.399 | 0.132 | 5.72E-08 |
| Cardiomyocyte | Daam2 | 2 | 7.64E-10 | 0.93842539 | 0.078 | 0.433 | 1.53E-06 |
| Cardiomyocyte | Skap11 | 2 | 8.69E-09 | 1.1820748 | 0.111 | 0.385 | 1.74E-05 |
| Cardiomyocyte | Fam92b | 3 | 1.95E-34 | 1.09132037 | 0.37 | 0.064 | 3.90E-31 |

| | | | | | | | |
|---|---|---|---|---|---|---|---|
| Cardiomyocyte | Igfbp4 | 3 | 4.37E-26 | 1.6744271 | 0.333 | 0.075 | 8.75E-23 |
| Cardiomyocyte | Kcng31 | 3 | 1.32E-20 | 2.34076914 | 0.406 | 0.101 | 2.63E-17 |
| Cardiomyocyte | Shank3 | 3 | 3.59E-20 | 3.49483263 | 0.667 | 0.398 | 7.19E-17 |
| Cardiomyocyte | Lurap1l | 3 | 3.54E-10 | 2.16856868 | 0.391 | 0.108 | 7.09E-07 |
| Cardiomyocyte | Sfrp4 | 3 | 2.02E-09 | 0.40860352 | 0.341 | 0.087 | 4.03E-06 |
| Cardiomyocyte | Ntng1 | 3 | 3.01E-08 | 1.28817961 | 0.384 | 0.07 | 6.02E-05 |
| Cardiomyocyte | Rasgrp1 | 3 | 2.33E-07 | 1.49890073 | 0.109 | 0.359 | 0.00046533 |
| Cardiomyocyte | Tbx21 | 4 | 7.50E-27 | 2.09812674 | 0.489 | 0.143 | 1.50E-23 |
| Cardiomyocyte | Alox5ap1 | 4 | 5.38E-21 | 2.72314282 | 0.431 | 0.125 | 1.08E-17 |
| Cardiomyocyte | Prss481 | 4 | 2.77E-20 | 0.54660486 | 0.431 | 0.125 | 5.54E-17 |
| Cardiomyocyte | Dscaml11 | 4 | 2.48E-17 | 1.74479656 | 0.431 | 0.153 | 4.96E-14 |
| Cardiomyocyte | Oasl1 | 4 | 4.85E-17 | 2.36979157 | 0.504 | 0.235 | 9.71E-14 |
| Cardiomyocyte | 5330417C22Rik1 | 4 | 2.86E-16 | 0.6277573 | 0.467 | 0.187 | 5.72E-13 |
| Cardiomyocyte | Esm1 | 4 | 1.73E-15 | 3.06676009 | 0.467 | 0.149 | 3.46E-12 |

| | | | | | | | |
|---|---|---|---|---|---|---|---|
| Cardiomyocyte | Serpine2 | 4 | 2.16E-15 | 0.4428987 | 0.095 | 0.456 | 4.32E-12 |
| Cardiomyocyte | Slc24a21 | 4 | 2.33E-15 | 0.63625887 | 0.445 | 0.177 | 4.66E-12 |
| Cardiomyocyte | Il71 | 4 | 6.46E-15 | 2.6309434 | 0.38 | 0.105 | 1.29E-11 |
| Cardiomyocyte | Stac31 | 4 | 1.34E-13 | 0.50162367 | 0.015 | 0.305 | 2.68E-10 |
| Cardiomyocyte | Lrrn2 | 4 | 2.36E-11 | 2.15117916 | 0.431 | 0.14 | 4.71E-08 |
| Cardiomyocyte | Inhba | 4 | 8.51E-11 | 0.25647663 | 0.015 | 0.269 | 1.70E-07 |
| Cardiomyocyte | Chi3l11 | 4 | 5.44E-09 | 1.01576026 | 0.401 | 0.122 | 1.09E-05 |
| Cardiomyocyte | Prph1 | 4 | 1.35E-08 | 0.62391863 | 0.445 | 0.154 | 2.70E-05 |
| Cardiomyocyte | Fam151a1 | 4 | 2.32E-08 | 0.43205174 | 0.131 | 0.419 | 4.64E-05 |
| Cardiomyocyte | Rfx2 | 4 | 3.42E-07 | 0.7720378 | 0.073 | 0.368 | 0.00068358 |
| Cardiomyocyte | Brca11 | 4 | 1.02E-05 | 0.84810985 | 0.438 | 0.178 | 0.02033634 |
| Cardiomyocyte | Saxo1 | 5 | 3.32E-128 | 2.83500512 | 0.555 | 0.014 | 6.64E-125 |
| Cardiomyocyte | P2rx1 | 5 | 4.91E-127 | 5.61016543 | 0.555 | 0.015 | 9.81E-124 |
| Cardiomyocyte | Pla2g2a | 5 | 4.88E-126 | 3.3048891 | 0.461 | 0.012 | 9.76E-123 |

| | | | | | | | |
|---|---|---|---|---|---|---|---|
| Cardiomyocyte | Cass4 | 5 | 4.99E-125 | 2.40899471 | 0.539 | 0.013 | 9.99E-122 |
| Cardiomyocyte | Aim2 | 5 | 1.54E-122 | 3.38576294 | 0.57 | 0.019 | 3.07E-119 |
| Cardiomyocyte | Ly49s6 | 5 | 1.96E-112 | 0.48609407 | 0.469 | 0.009 | 3.91E-109 |
| Cardiomyocyte | Hapln3 | 5 | 2.48E-112 | 3.37767471 | 0.531 | 0.019 | 4.95E-109 |
| Cardiomyocyte | Lef1 | 5 | 3.06E-109 | 3.27800017 | 0.516 | 0.017 | 6.12E-106 |
| Cardiomyocyte | RGD1559482 | 5 | 1.49E-108 | 1.27019164 | 0.508 | 0.016 | 2.97E-105 |
| Cardiomyocyte | Spata31d1 | 5 | 3.61E-98 | 1.72668414 | 0.328 | 0.026 | 7.21E-95 |
| Cardiomyocyte | Gata3 | 5 | 3.20E-95 | 1.10830216 | 0.438 | 0.012 | 6.40E-92 |
| Cardiomyocyte | Cd226 | 5 | 1.00E-92 | 3.95080408 | 0.531 | 0.044 | 2.01E-89 |
| Cardiomyocyte | Fut4 | 5 | 1.59E-87 | 2.34218687 | 0.312 | 0.023 | 3.19E-84 |
| Cardiomyocyte | Prkcz | 5 | 2.62E-87 | 4.07857264 | 0.602 | 0.048 | 5.24E-84 |
| Cardiomyocyte | Chl1 | 5 | 9.67E-87 | 1.9148572 | 0.539 | 0.039 | 1.93E-83 |
| Cardiomyocyte | Duox2 | 5 | 3.84E-84 | 1.67616205 | 0.555 | 0.045 | 7.69E-81 |
| Cardiomyocyte | Fam180a | 5 | 2.53E-82 | 3.12329257 | 0.531 | 0.033 | 5.07E-79 |

| | | | | | | | |
|---|---|---|---|---|---|---|---|
| Cardiomyocyte | AABR07030386.2 | 5 | 5.38E-82 | 2.44005496 | 0.57 | 0.039 | 1.08E-78 |
| Cardiomyocyte | C1qb | 5 | 1.16E-81 | 1.67543032 | 0.555 | 0.048 | 2.33E-78 |
| Cardiomyocyte | Pdpn | 5 | 2.37E-81 | 2.00073847 | 0.555 | 0.044 | 4.74E-78 |
| Cardiomyocyte | Ndc80 | 5 | 5.04E-81 | 1.94099377 | 0.555 | 0.047 | 1.01E-77 |
| Cardiomyocyte | Dll1 | 5 | 2.08E-80 | 1.75302079 | 0.531 | 0.04 | 4.15E-77 |
| Cardiomyocyte | Ect2l | 5 | 2.25E-77 | 1.68604889 | 0.336 | 0.008 | 4.50E-74 |
| Cardiomyocyte | Gria2 | 5 | 6.71E-76 | 1.28322088 | 0.336 | 0.008 | 1.34E-72 |
| Cardiomyocyte | Brinp1 | 5 | 5.11E-74 | 4.40078353 | 0.508 | 0.041 | 1.02E-70 |
| Cardiomyocyte | Pi15 | 5 | 5.55E-71 | 2.69939072 | 0.438 | 0.041 | 1.11E-67 |
| Cardiomyocyte | Opn4 | 5 | 2.09E-70 | 3.27937845 | 0.508 | 0.044 | 4.18E-67 |
| Cardiomyocyte | LOC103693323 | 5 | 6.59E-67 | 0.76345503 | 0.539 | 0.026 | 1.32E-63 |
| Cardiomyocyte | Kif27 | 5 | 1.45E-65 | 1.50632695 | 0.289 | 0.007 | 2.90E-62 |
| Cardiomyocyte | Cntn3 | 5 | 1.85E-65 | 0.72720872 | 0.609 | 0.04 | 3.70E-62 |
| Cardiomyocyte | Efhc2 | 5 | 2.49E-65 | 0.77740364 | 0.562 | 0.034 | 4.98E-62 |

| | | | | | | | |
|---|---|---|---|---|---|---|---|
| Cardiomyocyte | AABR07054565.1 | 5 | 6.48E-63 | 3.95074316 | 0.453 | 0.038 | 1.30E-59 |
| Cardiomyocyte | S100a9 | 5 | 1.79E-62 | 3.91065278 | 0.469 | 0.012 | 3.59E-59 |
| Cardiomyocyte | Lcp1 | 5 | 5.12E-60 | 0.59111631 | 0.508 | 0.048 | 1.02E-56 |
| Cardiomyocyte | Edaradd | 5 | 1.71E-50 | 1.97159039 | 0.273 | 0.006 | 3.41E-47 |
| Cardiomyocyte | Cnksr2 | 5 | 1.36E-47 | 0.81595621 | 0.43 | 0.114 | 2.71E-44 |
| Cardiomyocyte | Tmem63c | 5 | 5.60E-47 | 1.16183174 | 0.633 | 0.139 | 1.12E-43 |
| Cardiomyocyte | Enc1 | 5 | 3.53E-45 | 1.11512096 | 0.32 | 0.036 | 7.07E-42 |
| Cardiomyocyte | Rab11fip1 | 5 | 1.06E-44 | 2.00137825 | 0.359 | 0.065 | 2.11E-41 |
| Cardiomyocyte | Gsg1 | 5 | 1.99E-41 | 4.80845579 | 0.297 | 0.024 | 3.98E-38 |
| Cardiomyocyte | Esco21 | 5 | 2.33E-39 | 2.94555453 | 0.516 | 0.109 | 4.66E-36 |
| Cardiomyocyte | Crlf11 | 5 | 2.58E-39 | 1.08698507 | 0.555 | 0.121 | 5.16E-36 |
| Cardiomyocyte | RGD1311946 | 5 | 4.24E-38 | 1.15085694 | 0.398 | 0.117 | 8.48E-35 |
| Cardiomyocyte | Brip1 | 5 | 1.43E-37 | 1.09482699 | 0.602 | 0.158 | 2.86E-34 |
| Cardiomyocyte | Dab1 | 5 | 3.22E-37 | 1.7490229 | 0.547 | 0.024 | 6.43E-34 |

| | | | | | | | |
|---|---|---|---|---|---|---|---|
| Cardiomyocyte | Antxr1 | 5 | 1.70E-35 | 0.59425815 | 0.625 | 0.123 | 3.41E-32 |
| Cardiomyocyte | Lrrc31 | 5 | 1.72E-34 | 0.95611922 | 0.617 | 0.141 | 3.44E-31 |
| Cardiomyocyte | AABR07032261.11 | 5 | 3.70E-34 | 1.47585359 | 0.508 | 0.112 | 7.41E-31 |
| Cardiomyocyte | Catsper3 | 5 | 3.87E-33 | 1.92163885 | 0.336 | 0.033 | 7.74E-30 |
| Cardiomyocyte | Erbb3 | 5 | 2.58E-32 | 1.86649849 | 0.555 | 0.18 | 5.16E-29 |
| Cardiomyocyte | Samd15 | 5 | 3.63E-32 | 1.71662835 | 0.508 | 0.066 | 7.25E-29 |
| Cardiomyocyte | LOC100910636 | 5 | 2.16E-31 | 1.33880207 | 0.414 | 0.012 | 4.33E-28 |
| Cardiomyocyte | AC131411.1 | 5 | 2.74E-31 | 2.40546621 | 0.617 | 0.195 | 5.49E-28 |
| Cardiomyocyte | LOC682419 | 5 | 9.20E-31 | 1.97824895 | 0.641 | 0.236 | 1.84E-27 |
| Cardiomyocyte | Plaur | 5 | 9.71E-31 | 0.69249874 | 0.352 | 0.048 | 1.94E-27 |
| Cardiomyocyte | Lrmp1 | 5 | 2.25E-29 | 1.88613737 | 0.555 | 0.173 | 4.50E-26 |
| Cardiomyocyte | Inhba1 | 5 | 1.00E-28 | 0.64459503 | 0.602 | 0.209 | 2.01E-25 |
| Cardiomyocyte | Lancl3 | 5 | 1.02E-28 | 0.60931955 | 0.438 | 0.121 | 2.03E-25 |
| Cardiomyocyte | Ptafr | 5 | 1.81E-28 | 3.10582994 | 0.508 | 0.226 | 3.62E-25 |

| | | | | | | | |
|---|---|---|---|---|---|---|---|
| Cardiomyocyte | AC118957.11 | 5 | 3.86E-28 | 1.12800438 | 0.609 | 0.219 | 7.72E-25 |
| Cardiomyocyte | Chrdl11 | 5 | 1.10E-27 | 0.44661819 | 0.602 | 0.229 | 2.19E-24 |
| Cardiomyocyte | Sorcs1 | 5 | 1.11E-26 | 1.44419202 | 0.633 | 0.193 | 2.22E-23 |
| Cardiomyocyte | AABR07017902.1 | 5 | 4.04E-26 | 1.01865125 | 0.609 | 0.197 | 8.08E-23 |
| Cardiomyocyte | Col9a11 | 5 | 6.37E-25 | 2.1934544 | 0.602 | 0.157 | 1.27E-21 |
| Cardiomyocyte | Chrna1 | 5 | 7.33E-25 | 2.62532422 | 0.609 | 0.291 | 1.47E-21 |
| Cardiomyocyte | Dagla | 5 | 1.31E-24 | 0.95709839 | 0.57 | 0.223 | 2.62E-21 |
| Cardiomyocyte | Unc13a | 5 | 3.37E-24 | 3.298501 | 0.531 | 0.202 | 6.75E-21 |
| Cardiomyocyte | AABR07001477.1 | 5 | 6.06E-24 | 1.39938179 | 0.414 | 0.097 | 1.21E-20 |
| Cardiomyocyte | Cdh31 | 5 | 3.04E-23 | 0.39321714 | 0.453 | 0.127 | 6.09E-20 |
| Cardiomyocyte | AABR07026483.11 | 5 | 4.18E-23 | 0.42183861 | 0.523 | 0.154 | 8.36E-20 |
| Cardiomyocyte | Sv2c | 5 | 1.26E-22 | 0.86822363 | 0.523 | 0.141 | 2.53E-19 |
| Cardiomyocyte | Nkain2 | 5 | 1.55E-22 | 1.2324183 | 0.5 | 0.171 | 3.10E-19 |
| Cardiomyocyte | Serpine21 | 5 | 4.74E-22 | 0.9513912 | 0.648 | 0.398 | 9.47E-19 |

| | | | | | | | |
|---|---|---|---|---|---|---|---|
| Cardiomyocyte | Mafb1 | 5 | 6.57E-22 | 1.56993729 | 0.617 | 0.212 | 1.31E-18 |
| Cardiomyocyte | Dmp1 | 5 | 9.55E-22 | 1.73632474 | 0.305 | 0.045 | 1.91E-18 |
| Cardiomyocyte | Bace2 | 5 | 1.00E-20 | 0.30457647 | 0.445 | 0.139 | 2.00E-17 |
| Cardiomyocyte | Reln | 5 | 8.10E-20 | 1.16251748 | 0.664 | 0.311 | 1.62E-16 |
| Cardiomyocyte | Cd44 | 5 | 1.91E-19 | 1.40352058 | 0.672 | 0.372 | 3.83E-16 |
| Cardiomyocyte | Aurkb | 5 | 2.96E-19 | 1.24806695 | 0.594 | 0.225 | 5.93E-16 |
| Cardiomyocyte | Lrsam11 | 5 | 1.47E-18 | 1.01715091 | 0.547 | 0.15 | 2.94E-15 |
| Cardiomyocyte | Ankrd6 | 5 | 1.70E-18 | 1.07615529 | 0.797 | 0.39 | 3.40E-15 |
| Cardiomyocyte | Tceal7 | 5 | 1.76E-18 | 1.16283267 | 0.523 | 0.164 | 3.51E-15 |
| Cardiomyocyte | Mcam | 5 | 1.89E-17 | 0.80843661 | 0.641 | 0.232 | 3.77E-14 |
| Cardiomyocyte | Piwil2 | 5 | 3.68E-16 | 0.97212948 | 0.695 | 0.269 | 7.35E-13 |
| Cardiomyocyte | AABR07027581.1 | 5 | 7.76E-16 | 0.87958807 | 0.625 | 0.255 | 1.55E-12 |
| Cardiomyocyte | Scara5 | 5 | 8.99E-16 | 1.11440458 | 0.602 | 0.157 | 1.80E-12 |
| Cardiomyocyte | Carmil1 | 5 | 1.06E-15 | 0.81000506 | 0.844 | 0.546 | 2.12E-12 |

| | | | | | | | |
|---|---|---|---|---|---|---|---|
| Cardiomyocyte | Sdk2 | 5 | 1.17E-15 | 0.79198141 | 0.508 | 0.083 | 2.35E-12 |
| Cardiomyocyte | Nalcn | 5 | 2.14E-15 | 1.20025668 | 0.43 | 0.058 | 4.28E-12 |
| Cardiomyocyte | Pip5k1b | 5 | 2.57E-15 | 0.907976 | 0.844 | 0.519 | 5.15E-12 |
| Cardiomyocyte | AABR07054490.1 | 5 | 7.38E-15 | 1.24750723 | 0.617 | 0.135 | 1.48E-11 |
| Cardiomyocyte | Ptpn18 | 5 | 1.23E-14 | 1.54187991 | 0.578 | 0.262 | 2.46E-11 |
| Cardiomyocyte | Sntb1 | 5 | 7.45E-14 | 1.04682388 | 0.859 | 0.54 | 1.49E-10 |
| Cardiomyocyte | Slc25a13 | 5 | 8.32E-14 | 0.76605723 | 0.883 | 0.555 | 1.66E-10 |
| Cardiomyocyte | RGD15640531 | 5 | 1.52E-13 | 1.05470832 | 0.594 | 0.221 | 3.04E-10 |
| Cardiomyocyte | Slc25a211 | 5 | 1.59E-13 | 0.45750537 | 0.75 | 0.469 | 3.19E-10 |
| Cardiomyocyte | Tm4sf4 | 5 | 2.03E-13 | 0.7567693 | 0.562 | 0.261 | 4.05E-10 |
| Cardiomyocyte | Itgam | 5 | 2.49E-13 | 0.35274065 | 0.062 | 0.397 | 4.99E-10 |
| Cardiomyocyte | Fhl1 | 5 | 2.97E-13 | 0.87591093 | 0.773 | 0.415 | 5.93E-10 |
| Cardiomyocyte | Pcsk5 | 5 | 3.26E-13 | 1.04732484 | 0.664 | 0.406 | 6.51E-10 |
| Cardiomyocyte | Sphkap | 5 | 6.56E-13 | 0.87137578 | 0.734 | 0.44 | 1.31E-09 |

| | | | | | | | |
|---|---|---|---|---|---|---|---|
| Cardiomyocyte | Scn1a | 5 | 1.57E-12 | 0.80083832 | 0.789 | 0.508 | 3.14E-09 |
| Cardiomyocyte | Mpc11 | 5 | 1.84E-12 | 0.6978766 | 0.859 | 0.438 | 3.67E-09 |
| Cardiomyocyte | Lrrc4b | 5 | 4.76E-12 | 0.69514116 | 0.844 | 0.478 | 9.53E-09 |
| Cardiomyocyte | Col6a2 | 5 | 7.07E-12 | 0.31333193 | 0.602 | 0.189 | 1.41E-08 |
| Cardiomyocyte | Filip1l | 5 | 8.39E-12 | 0.68133408 | 0.953 | 0.593 | 1.68E-08 |
| Cardiomyocyte | Enpp2 | 5 | 8.41E-12 | 2.00570519 | 0.453 | 0.108 | 1.68E-08 |
| Cardiomyocyte | Sema3a | 5 | 1.16E-11 | 1.35074892 | 0.609 | 0.302 | 2.31E-08 |
| Cardiomyocyte | Bdh1 | 5 | 1.27E-11 | 0.79173929 | 0.742 | 0.458 | 2.54E-08 |
| Cardiomyocyte | Scube31 | 5 | 1.55E-11 | 1.27590374 | 0.453 | 0.163 | 3.11E-08 |
| Cardiomyocyte | Ifit3 | 5 | 1.78E-11 | 1.14097178 | 0.008 | 0.289 | 3.55E-08 |
| Cardiomyocyte | Rasgef1b1 | 5 | 2.41E-11 | 2.57442116 | 0.586 | 0.306 | 4.81E-08 |
| Cardiomyocyte | Cemip1 | 5 | 2.42E-11 | 0.55452031 | 0.484 | 0.178 | 4.83E-08 |
| Cardiomyocyte | Pola2 | 5 | 2.85E-11 | 0.65795159 | 0.664 | 0.221 | 5.70E-08 |
| Cardiomyocyte | Pkia | 5 | 3.36E-11 | 0.77742308 | 0.836 | 0.426 | 6.73E-08 |

| | | | | | | | |
|---|---|---|---|---|---|---|---|
| Cardiomyocyte | Tanc2 | 5 | 4.64E-11 | 0.31182963 | 0.758 | 0.424 | 9.29E-08 |
| Cardiomyocyte | Dusp27 | 5 | 1.63E-10 | 0.35783091 | 0.773 | 0.448 | 3.25E-07 |
| Cardiomyocyte | Glb1l21 | 5 | 1.79E-10 | 0.59347942 | 0.82 | 0.449 | 3.58E-07 |
| Cardiomyocyte | Dcx | 5 | 1.92E-10 | 0.28233384 | 0.008 | 0.259 | 3.84E-07 |
| Cardiomyocyte | AABR07034767.1 | 5 | 2.85E-10 | 0.71583715 | 0.844 | 0.493 | 5.70E-07 |
| Cardiomyocyte | Cxadr | 5 | 3.37E-10 | 0.27467264 | 0.844 | 0.468 | 6.75E-07 |
| Cardiomyocyte | Fign1 | 5 | 5.27E-10 | 0.71231113 | 0.844 | 0.551 | 1.05E-06 |
| Cardiomyocyte | Myo16 | 5 | 2.26E-09 | 0.26305043 | 0.594 | 0.241 | 4.52E-06 |
| Cardiomyocyte | Adamts1 | 5 | 3.11E-09 | 0.3710118 | 0.664 | 0.396 | 6.22E-06 |
| Cardiomyocyte | Gja11 | 5 | 7.01E-09 | 0.58772783 | 0.898 | 0.595 | 1.40E-05 |
| Cardiomyocyte | Stk38l | 5 | 7.04E-09 | 0.25741058 | 0.719 | 0.461 | 1.41E-05 |
| Cardiomyocyte | Ppp1r3a1 | 5 | 7.26E-09 | 0.54171047 | 0.828 | 0.532 | 1.45E-05 |
| Cardiomyocyte | Plbd1 | 5 | 1.33E-08 | 0.46910704 | 0.719 | 0.417 | 2.66E-05 |
| Cardiomyocyte | Syt14 | 5 | 1.91E-08 | 0.89587146 | 0.609 | 0.332 | 3.83E-05 |

| | | | | | | | |
|---|---|---|---|---|---|---|---|
| Cardiomyocyte | Slc7a1 | 5 | 2.39E-08 | 0.39606815 | 0.695 | 0.418 | 4.78E-05 |
| Cardiomyocyte | LOC100911847 | 5 | 2.56E-08 | 0.860403 | 0.641 | 0.385 | 5.12E-05 |
| Cardiomyocyte | Hk21 | 5 | 3.74E-08 | 0.44078547 | 0.852 | 0.503 | 7.48E-05 |
| Cardiomyocyte | Slit2 | 5 | 7.64E-08 | 1.21138897 | 0.414 | 0.158 | 0.00015279 |
| Cardiomyocyte | Efcab6 | 5 | 1.90E-07 | 0.28060595 | 0.695 | 0.4 | 0.00037986 |
| Cardiomyocyte | Mrvi1 | 5 | 2.95E-07 | 0.34456527 | 0.586 | 0.25 | 0.00059042 |
| Cardiomyocyte | Ac1576 | 5 | 4.54E-07 | 0.73319673 | 0.516 | 0.232 | 0.00090832 |
| Cardiomyocyte | Lmx1a | 5 | 7.48E-07 | 0.31056961 | 0.586 | 0.251 | 0.00149512 |
| Cardiomyocyte | Matk | 5 | 1.37E-06 | 3.36538573 | 0.273 | 0.007 | 0.00273859 |
| Cardiomyocyte | Myoz2 | 5 | 1.47E-06 | 0.39591027 | 0.898 | 0.604 | 0.00294946 |
| Cardiomyocyte | Ros1 | 5 | 3.23E-06 | 0.46308886 | 0.703 | 0.409 | 0.00645286 |
| Cardiomyocyte | AABR07051518.1 | 5 | 8.82E-06 | 0.3387741 | 0.648 | 0.384 | 0.01763277 |
| Cardiomyocyte | Iqub | 5 | 1.01E-05 | 0.33356795 | 0.781 | 0.445 | 0.02015907 |
| Cardiomyocyte | Bzw2 | 5 | 1.03E-05 | 0.27097912 | 0.969 | 0.699 | 0.02062916 |

| | | | | | | | |
|---|---|---|---|---|---|---|---|
| Cardiomyocyte | Trpc6 | 6 | 1.13E-86 | 4.09705371 | 0.773 | 0.121 | 2.27E-83 |
| Cardiomyocyte | Fgr | 6 | 3.24E-85 | 1.56118342 | 0.773 | 0.123 | 6.49E-82 |
| Cardiomyocyte | Hrh2 | 6 | 2.02E-82 | 3.21542785 | 0.734 | 0.114 | 4.04E-79 |
| Cardiomyocyte | Cmtm5 | 6 | 5.09E-76 | 4.27650615 | 0.633 | 0.078 | 1.02E-72 |
| Cardiomyocyte | Siah3 | 6 | 2.43E-75 | 2.56754238 | 0.633 | 0.079 | 4.87E-72 |
| Cardiomyocyte | Spetex-2F | 6 | 1.23E-73 | 3.23639153 | 0.531 | 0.051 | 2.46E-70 |
| Cardiomyocyte | Hao1 | 6 | 5.92E-61 | 3.83035322 | 0.797 | 0.207 | 1.18E-57 |
| Cardiomyocyte | Matn3 | 6 | 1.92E-59 | 3.54696977 | 0.531 | 0.074 | 3.85E-56 |
| Cardiomyocyte | Ctss | 6 | 1.96E-57 | 1.66232689 | 0.781 | 0.189 | 3.93E-54 |
| Cardiomyocyte | Bdkrb2 | 6 | 5.15E-56 | 0.93191356 | 0.773 | 0.124 | 1.03E-52 |
| Cardiomyocyte | Vit | 6 | 2.64E-55 | 1.0541421 | 0.625 | 0.107 | 5.27E-52 |
| Cardiomyocyte | Mt2A | 6 | 6.24E-55 | 0.41993783 | 0.75 | 0.182 | 1.25E-51 |
| Cardiomyocyte | Dcx1 | 6 | 8.87E-54 | 1.41256644 | 0.75 | 0.185 | 1.77E-50 |
| Cardiomyocyte | Epsti1 | 6 | 7.39E-53 | 0.39387072 | 0.711 | 0.157 | 1.48E-49 |

| | | | | | | | |
|---|---|---|---|---|---|---|---|
| Cardiomyocyte | AC123500.11 | 6 | 7.50E-51 | 2.04581034 | 0.812 | 0.205 | 1.50E-47 |
| Cardiomyocyte | Hyi | 6 | 1.48E-49 | 0.35622144 | 0.812 | 0.243 | 2.96E-46 |
| Cardiomyocyte | Tspan33 | 6 | 6.07E-48 | 4.16376793 | 0.656 | 0.111 | 1.21E-44 |
| Cardiomyocyte | Atf3 | 6 | 1.96E-47 | 2.705117 | 0.734 | 0.081 | 3.92E-44 |
| Cardiomyocyte | St14 | 6 | 5.12E-47 | 3.44375001 | 0.633 | 0.077 | 1.02E-43 |
| Cardiomyocyte | Cyfip2 | 6 | 1.47E-46 | 1.84682643 | 0.797 | 0.244 | 2.93E-43 |
| Cardiomyocyte | Slamf81 | 6 | 1.20E-45 | 3.96947485 | 0.695 | 0.161 | 2.39E-42 |
| Cardiomyocyte | Galnt15 | 6 | 4.69E-45 | 1.20459579 | 0.797 | 0.228 | 9.39E-42 |
| Cardiomyocyte | Hap1 | 6 | 6.13E-45 | 2.73554324 | 0.805 | 0.319 | 1.23E-41 |
| Cardiomyocyte | Dnajc61 | 6 | 3.64E-44 | 4.04497239 | 0.484 | 0.079 | 7.27E-41 |
| Cardiomyocyte | Sh3bp2 | 6 | 3.67E-42 | 1.71724373 | 0.82 | 0.192 | 7.34E-39 |
| Cardiomyocyte | Ca4 | 6 | 5.92E-42 | 1.22501778 | 0.703 | 0.159 | 1.18E-38 |
| Cardiomyocyte | Itgam1 | 6 | 1.95E-41 | 2.17718628 | 0.82 | 0.322 | 3.90E-38 |
| Cardiomyocyte | Slc7a7 | 6 | 2.75E-39 | 0.82127875 | 0.742 | 0.245 | 5.49E-36 |

| | | | | | | | |
|---|---|---|---|---|---|---|---|
| Cardiomyocyte | LOC102553018 | 6 | 6.86E-39 | 0.44520642 | 0.711 | 0.217 | 1.37E-35 |
| Cardiomyocyte | Gria41 | 6 | 4.02E-38 | 1.78157363 | 0.836 | 0.378 | 8.04E-35 |
| Cardiomyocyte | Shc31 | 6 | 1.24E-36 | 2.75455323 | 0.711 | 0.21 | 2.49E-33 |
| Cardiomyocyte | Mamdc21 | 6 | 7.23E-36 | 0.25115001 | 0.828 | 0.461 | 1.45E-32 |
| Cardiomyocyte | Lyz2 | 6 | 8.91E-36 | 2.61431888 | 0.711 | 0.12 | 1.78E-32 |
| Cardiomyocyte | Nr5a21 | 6 | 3.70E-35 | 0.32959337 | 0.742 | 0.215 | 7.39E-32 |
| Cardiomyocyte | Gxylt2 | 6 | 8.82E-35 | 1.16419663 | 0.844 | 0.423 | 1.76E-31 |
| Cardiomyocyte | Igf11 | 6 | 9.29E-35 | 0.83944091 | 0.828 | 0.481 | 1.86E-31 |
| Cardiomyocyte | Arhgef26 | 6 | 5.89E-34 | 2.14780707 | 0.75 | 0.266 | 1.18E-30 |
| Cardiomyocyte | Dpysl3 | 6 | 2.11E-33 | 1.12711136 | 0.836 | 0.349 | 4.22E-30 |
| Cardiomyocyte | Tdrd121 | 6 | 4.95E-33 | 0.43135605 | 0.68 | 0.183 | 9.89E-30 |
| Cardiomyocyte | AABR07049033.1 | 6 | 6.80E-33 | 1.2482801 | 0.828 | 0.338 | 1.36E-29 |
| Cardiomyocyte | Smpd3 | 6 | 7.35E-33 | 2.83585282 | 0.648 | 0.195 | 1.47E-29 |
| Cardiomyocyte | AABR07017902.11 | 6 | 8.31E-33 | 2.35338272 | 0.656 | 0.192 | 1.66E-29 |

| | | | | | | | |
|---|---|---|---|---|---|---|---|
| Cardiomyocyte | Srpx | 6 | 7.27E-32 | 0.90747143 | 0.828 | 0.389 | 1.45E-28 |
| Cardiomyocyte | Zmat4 | 6 | 5.82E-31 | 3.43624204 | 0.578 | 0.206 | 1.16E-27 |
| Cardiomyocyte | Bard11 | 6 | 6.00E-31 | 0.98989212 | 0.812 | 0.369 | 1.20E-27 |
| Cardiomyocyte | Lmod1 | 6 | 1.34E-30 | 0.37173177 | 0.828 | 0.469 | 2.68E-27 |
| Cardiomyocyte | Ctse | 6 | 1.56E-30 | 0.79665444 | 0.398 | 0.075 | 3.12E-27 |
| Cardiomyocyte | Myl4 | 6 | 3.55E-30 | 0.82353618 | 0.781 | 0.307 | 7.10E-27 |
| Cardiomyocyte | Itgal1 | 6 | 4.62E-30 | 2.04567426 | 0.781 | 0.304 | 9.24E-27 |
| Cardiomyocyte | Svep11 | 6 | 8.90E-30 | 1.73746685 | 0.758 | 0.415 | 1.78E-26 |
| Cardiomyocyte | Clmn | 6 | 2.23E-29 | 0.79262685 | 0.789 | 0.34 | 4.45E-26 |
| Cardiomyocyte | Rbm44 | 6 | 5.98E-29 | 1.6187301 | 0.5 | 0.178 | 1.20E-25 |
| Cardiomyocyte | Fam151a2 | 6 | 1.06E-28 | 0.91263094 | 0.797 | 0.35 | 2.12E-25 |
| Cardiomyocyte | Lrrtm4 | 6 | 2.45E-28 | 2.2428113 | 0.461 | 0.145 | 4.89E-25 |
| Cardiomyocyte | Jag1 | 6 | 3.41E-28 | 0.66763887 | 0.797 | 0.4 | 6.83E-25 |
| Cardiomyocyte | Samd141 | 6 | 7.53E-28 | 1.03036228 | 0.828 | 0.406 | 1.51E-24 |

| | | | | | | | |
|---|---|---|---|---|---|---|---|
| Cardiomyocyte | Frem1 | 6 | 1.07E-27 | 0.35860789 | 0.836 | 0.485 | 2.13E-24 |
| Cardiomyocyte | Bnc2 | 6 | 3.43E-27 | 1.00994258 | 0.836 | 0.455 | 6.85E-24 |
| Cardiomyocyte | Marchf10 | 6 | 7.11E-27 | 1.91417362 | 0.68 | 0.378 | 1.42E-23 |
| Cardiomyocyte | Slc8a2 | 6 | 1.41E-26 | 0.75341943 | 0.82 | 0.33 | 2.83E-23 |
| Cardiomyocyte | C1qtnf1 | 6 | 1.64E-26 | 1.54736222 | 0.406 | 0.079 | 3.29E-23 |
| Cardiomyocyte | Ccdc3 | 6 | 8.89E-26 | 0.82288892 | 0.852 | 0.498 | 1.78E-22 |
| Cardiomyocyte | Alpl | 6 | 1.14E-25 | 0.53073103 | 0.828 | 0.35 | 2.29E-22 |
| Cardiomyocyte | Plcg2 | 6 | 1.17E-25 | 0.4887796 | 0.812 | 0.362 | 2.34E-22 |
| Cardiomyocyte | Map3k7cl | 6 | 1.27E-25 | 3.6346863 | 0.516 | 0.248 | 2.53E-22 |
| Cardiomyocyte | Popdc31 | 6 | 6.04E-25 | 0.72761524 | 0.812 | 0.368 | 1.21E-21 |
| Cardiomyocyte | Gpm6b | 6 | 8.75E-25 | 0.43774721 | 0.781 | 0.295 | 1.75E-21 |
| Cardiomyocyte | Trhde | 6 | 1.17E-24 | 1.91505494 | 0.766 | 0.347 | 2.34E-21 |
| Cardiomyocyte | Uqcrq | 6 | 1.49E-24 | 1.63935959 | 0.844 | 0.371 | 2.98E-21 |
| Cardiomyocyte | Pfkfb3 | 6 | 2.13E-24 | 0.71612764 | 0.828 | 0.474 | 4.27E-21 |

| | | | | | | | |
|---|---|---|---|---|---|---|---|
| Cardiomyocyte | Smad6 | 6 | 8.43E-24 | 0.50501963 | 0.859 | 0.48 | 1.69E-20 |
| Cardiomyocyte | AABR07068046.11 | 6 | 9.61E-24 | 0.92281426 | 0.805 | 0.326 | 1.92E-20 |
| Cardiomyocyte | Tpx21 | 6 | 1.64E-23 | 0.73310803 | 0.68 | 0.244 | 3.29E-20 |
| Cardiomyocyte | Tmem1631 | 6 | 6.08E-23 | 1.14873434 | 0.797 | 0.421 | 1.22E-19 |
| Cardiomyocyte | Ikzf31 | 6 | 9.32E-23 | 2.15313647 | 0.727 | 0.297 | 1.86E-19 |
| Cardiomyocyte | Hey2 | 6 | 1.63E-22 | 1.33814125 | 0.594 | 0.276 | 3.25E-19 |
| Cardiomyocyte | Chsy3 | 6 | 4.95E-22 | 0.61297864 | 0.797 | 0.365 | 9.91E-19 |
| Cardiomyocyte | Mt1 | 6 | 6.87E-22 | 1.7004348 | 0.812 | 0.301 | 1.37E-18 |
| Cardiomyocyte | E2f7 | 6 | 1.29E-21 | 2.45003024 | 0.484 | 0.119 | 2.58E-18 |
| Cardiomyocyte | Nox4 | 6 | 1.44E-21 | 0.31856377 | 0.805 | 0.419 | 2.87E-18 |
| Cardiomyocyte | Fam189a11 | 6 | 1.44E-21 | 0.77743524 | 0.82 | 0.508 | 2.88E-18 |
| Cardiomyocyte | Hs3st1 | 6 | 2.71E-21 | 0.68400147 | 0.766 | 0.344 | 5.42E-18 |
| Cardiomyocyte | Grin2b | 6 | 3.01E-21 | 2.78985935 | 0.43 | 0.18 | 6.03E-18 |
| Cardiomyocyte | Tgfb2 | 6 | 5.05E-21 | 0.67016545 | 0.859 | 0.542 | 1.01E-17 |

| | | | | | | | |
|---|---|---|---|---|---|---|---|
| Cardiomyocyte | Snx31 | 6 | 1.70E-20 | 1.99982764 | 0.75 | 0.331 | 3.40E-17 |
| Cardiomyocyte | RGD15633541 | 6 | 3.74E-20 | 0.5753776 | 0.703 | 0.272 | 7.47E-17 |
| Cardiomyocyte | Cxcl12 | 6 | 4.53E-20 | 0.59189575 | 0.758 | 0.399 | 9.06E-17 |
| Cardiomyocyte | Rgs7 | 6 | 9.50E-20 | 1.5402351 | 0.836 | 0.546 | 1.90E-16 |
| Cardiomyocyte | Tp53inp21 | 6 | 1.97E-19 | 0.99078499 | 0.867 | 0.534 | 3.94E-16 |
| Cardiomyocyte | Spp11 | 6 | 3.18E-19 | 1.37300251 | 0.523 | 0.139 | 6.37E-16 |
| Cardiomyocyte | Ptgs2 | 6 | 2.03E-18 | 0.29006195 | 0.547 | 0.203 | 4.06E-15 |
| Cardiomyocyte | Errfi1 | 6 | 2.27E-18 | 0.6459706 | 0.734 | 0.289 | 4.54E-15 |
| Cardiomyocyte | Ptk2b | 6 | 2.70E-18 | 0.85227802 | 0.477 | 0.091 | 5.41E-15 |
| Cardiomyocyte | Bcl6b | 6 | 4.17E-18 | 0.74783237 | 0.539 | 0.111 | 8.33E-15 |
| Cardiomyocyte | Casc41 | 6 | 4.34E-18 | 1.21436922 | 0.734 | 0.416 | 8.69E-15 |
| Cardiomyocyte | Serpine1 | 6 | 4.69E-18 | 2.14254563 | 0.633 | 0.291 | 9.38E-15 |
| Cardiomyocyte | Tex14 | 6 | 4.96E-18 | 0.93519157 | 0.656 | 0.292 | 9.92E-15 |
| Cardiomyocyte | Cdh111 | 6 | 5.43E-18 | 0.64263606 | 0.828 | 0.447 | 1.09E-14 |

| | | | | | | | |
|---|---|---|---|---|---|---|---|
| Cardiomyocyte | Bend6 | 6 | 9.29E-18 | 1.13264416 | 0.656 | 0.329 | 1.86E-14 |
| Cardiomyocyte | Plod21 | 6 | 2.96E-17 | 0.3400245 | 0.836 | 0.52 | 5.92E-14 |
| Cardiomyocyte | Ccl21 | 6 | 3.51E-17 | 0.50526937 | 0.344 | 0.088 | 7.01E-14 |
| Cardiomyocyte | Nxnl2 | 6 | 4.17E-17 | 0.63734219 | 0.641 | 0.252 | 8.33E-14 |
| Cardiomyocyte | Tshr | 6 | 5.26E-17 | 1.64994574 | 0.742 | 0.319 | 1.05E-13 |
| Cardiomyocyte | Flna | 6 | 5.47E-17 | 0.51209255 | 0.852 | 0.501 | 1.09E-13 |
| Cardiomyocyte | AABR07006724.1 | 6 | 9.53E-17 | 1.23989458 | 0.812 | 0.465 | 1.91E-13 |
| Cardiomyocyte | Fgf10 | 6 | 1.56E-16 | 2.09938648 | 0.617 | 0.321 | 3.12E-13 |
| Cardiomyocyte | Luzp21 | 6 | 3.51E-16 | 1.16228759 | 0.633 | 0.225 | 7.01E-13 |
| Cardiomyocyte | Flnc | 6 | 4.86E-16 | 1.02476688 | 0.891 | 0.586 | 9.73E-13 |
| Cardiomyocyte | Egfr | 6 | 6.38E-16 | 1.06405617 | 0.672 | 0.325 | 1.28E-12 |
| Cardiomyocyte | AABR07052441.11 | 6 | 1.81E-15 | 1.75547643 | 0.586 | 0.307 | 3.62E-12 |
| Cardiomyocyte | Mrc21 | 6 | 4.73E-15 | 0.63875733 | 0.812 | 0.495 | 9.45E-12 |
| Cardiomyocyte | Pmepa1 | 6 | 4.82E-15 | 0.55269693 | 0.844 | 0.46 | 9.64E-12 |

| | | | | | | | |
|---|---|---|---|---|---|---|---|
| Cardiomyocyte | Pdp2 | 6 | 8.70E-15 | 0.40935736 | 0.539 | 0.275 | 1.74E-11 |
| Cardiomyocyte | Nr4a31 | 6 | 9.22E-15 | 0.98193798 | 0.664 | 0.255 | 1.84E-11 |
| Cardiomyocyte | Slc9a3r2 | 6 | 3.27E-14 | 0.41158812 | 0.656 | 0.347 | 6.54E-11 |
| Cardiomyocyte | LOC685963 | 6 | 8.11E-14 | 1.01557038 | 0.805 | 0.421 | 1.62E-10 |
| Cardiomyocyte | Stac1 | 6 | 2.73E-13 | 0.70913441 | 0.586 | 0.241 | 5.47E-10 |
| Cardiomyocyte | Npffr2 | 6 | 3.34E-13 | 0.56310597 | 0.547 | 0.188 | 6.67E-10 |
| Cardiomyocyte | PCOLCE21 | 6 | 3.98E-13 | 1.32530167 | 0.766 | 0.48 | 7.97E-10 |
| Cardiomyocyte | Adamtsl31 | 6 | 5.00E-13 | 0.54137668 | 0.836 | 0.505 | 1.00E-09 |
| Cardiomyocyte | Jag2 | 6 | 6.04E-13 | 0.4888844 | 0.625 | 0.276 | 1.21E-09 |
| Cardiomyocyte | Nmnat2 | 6 | 7.76E-13 | 0.86620703 | 0.773 | 0.459 | 1.55E-09 |
| Cardiomyocyte | Dock8 | 6 | 8.33E-13 | 0.34635497 | 0.82 | 0.484 | 1.67E-09 |
| Cardiomyocyte | Bdnf | 6 | 1.43E-12 | 1.16715758 | 0.867 | 0.518 | 2.86E-09 |
| Cardiomyocyte | Lrrc10 | 6 | 1.67E-12 | 0.56767757 | 0.727 | 0.365 | 3.34E-09 |
| Cardiomyocyte | Grip21 | 6 | 4.15E-12 | 0.56736918 | 0.727 | 0.368 | 8.29E-09 |

| | | | | | | | |
|---|---|---|---|---|---|---|---|
| Cardiomyocyte | Gadd45b1 | 6 | 4.25E-12 | 1.79176198 | 0.688 | 0.344 | 8.50E-09 |
| Cardiomyocyte | Scn1a1 | 6 | 6.70E-12 | 0.54654984 | 0.805 | 0.506 | 1.34E-08 |
| Cardiomyocyte | Acta21 | 6 | 1.12E-11 | 1.27715789 | 0.438 | 0.157 | 2.25E-08 |
| Cardiomyocyte | Sctr1 | 6 | 1.16E-11 | 1.12103014 | 0.812 | 0.479 | 2.32E-08 |
| Cardiomyocyte | Astn2 | 6 | 1.20E-11 | 0.32905631 | 0.836 | 0.558 | 2.40E-08 |
| Cardiomyocyte | Anp32e | 6 | 1.31E-11 | 0.42046828 | 0.789 | 0.452 | 2.63E-08 |
| Cardiomyocyte | Ddc | 6 | 1.55E-11 | 1.11943591 | 0.789 | 0.525 | 3.10E-08 |
| Cardiomyocyte | Prlr | 6 | 4.03E-11 | 2.82770855 | 0.453 | 0.148 | 8.07E-08 |
| Cardiomyocyte | Nnat1 | 6 | 5.47E-11 | 1.71938261 | 0.656 | 0.347 | 1.09E-07 |
| Cardiomyocyte | Map1b | 6 | 6.39E-11 | 0.58335114 | 0.859 | 0.479 | 1.28E-07 |
| Cardiomyocyte | Abcb4 | 6 | 8.40E-11 | 1.11985606 | 0.898 | 0.597 | 1.68E-07 |
| Cardiomyocyte | Rxrg1 | 6 | 9.28E-11 | 0.32173272 | 0.852 | 0.542 | 1.86E-07 |
| Cardiomyocyte | Mrc11 | 6 | 1.07E-10 | 0.49638254 | 0.734 | 0.411 | 2.13E-07 |
| Cardiomyocyte | Mctp1 | 6 | 1.16E-10 | 0.2872024 | 0.781 | 0.398 | 2.32E-07 |

| | | | | | | | |
|---|---|---|---|---|---|---|---|
| Cardiomyocyte | Hhatl | 6 | 3.25E-10 | 0.52926772 | 0.703 | 0.33 | 6.50E-07 |
| Cardiomyocyte | Art11 | 6 | 7.74E-10 | 0.3639087 | 0.852 | 0.564 | 1.55E-06 |
| Cardiomyocyte | AABR07049156.11 | 6 | 9.31E-10 | 0.40501383 | 0.578 | 0.262 | 1.86E-06 |
| Cardiomyocyte | Synm | 6 | 1.76E-09 | 0.86141585 | 0.852 | 0.508 | 3.52E-06 |
| Cardiomyocyte | Cdk19 | 6 | 2.26E-09 | 0.48194363 | 0.812 | 0.465 | 4.51E-06 |
| Cardiomyocyte | Mtus21 | 6 | 5.85E-09 | 0.35109122 | 0.859 | 0.59 | 1.17E-05 |
| Cardiomyocyte | Dtnb | 6 | 1.51E-08 | 0.64300397 | 0.797 | 0.495 | 3.03E-05 |
| Cardiomyocyte | Slc12a2 | 6 | 1.72E-08 | 0.33059945 | 0.758 | 0.469 | 3.44E-05 |
| Cardiomyocyte | Gmpr | 6 | 4.97E-08 | 0.27870036 | 0.883 | 0.627 | 9.94E-05 |
| Cardiomyocyte | Atp5f1e | 6 | 6.21E-08 | 0.73363037 | 0.742 | 0.468 | 0.00012425 |
| Cardiomyocyte | Zbtb7c | 6 | 6.99E-08 | 0.72653171 | 0.766 | 0.42 | 0.00013975 |
| Cardiomyocyte | Arhgap44 | 6 | 1.41E-07 | 0.60551575 | 0.898 | 0.646 | 0.00028113 |
| Cardiomyocyte | Enox1 | 6 | 1.42E-07 | 0.59845215 | 0.797 | 0.481 | 0.00028468 |
| Cardiomyocyte | Dgki | 6 | 1.81E-07 | 0.40478425 | 0.828 | 0.51 | 0.0003615 |

| | | | | | | | |
|---|---|---|---|---|---|---|---|
| Cardiomyocyte | AABR07040864.11 | 6 | 5.64E-07 | 0.37440312 | 0.742 | 0.462 | 0.00112795 |
| Cardiomyocyte | Ube2ql11 | 6 | 9.23E-07 | 0.7215375 | 0.719 | 0.466 | 0.00184569 |
| Cardiomyocyte | Rbm24 | 6 | 1.04E-06 | 0.35880036 | 0.742 | 0.44 | 0.00208111 |
| Cardiomyocyte | Fyb11 | 6 | 2.12E-06 | 1.97862391 | 0.516 | 0.231 | 0.00423903 |
| Cardiomyocyte | Pla2g71 | 6 | 2.45E-06 | 0.45927601 | 0.812 | 0.488 | 0.00489515 |
| Cardiomyocyte | P3h2 | 6 | 3.31E-06 | 0.5961772 | 0.836 | 0.534 | 0.00662141 |
| Cardiomyocyte | Phf24 | 6 | 3.95E-06 | 0.66474359 | 0.836 | 0.561 | 0.00790865 |
| Cardiomyocyte | Ndrg4 | 6 | 6.20E-06 | 0.64744445 | 0.805 | 0.555 | 0.0124068 |
| Cardiomyocyte | Actn1 | 6 | 7.16E-06 | 0.95775755 | 0.75 | 0.476 | 0.01432125 |
| Cardiomyocyte | Klhl41 | 6 | 7.18E-06 | 0.710924 | 0.617 | 0.35 | 0.01435944 |
| Cardiomyocyte | Pkia1 | 6 | 1.35E-05 | 0.37002109 | 0.703 | 0.439 | 0.0269052 |
| Cardiomyocyte | AABR07016845.1 | 7 | 3.02E-29 | 0.45408347 | 0.047 | 0.485 | 6.04E-26 |
| Cardiomyocyte | Dipk2b | 7 | 1.24E-28 | 0.46374114 | 0.039 | 0.412 | 2.48E-25 |
| Cardiomyocyte | Dgkg | 7 | 2.87E-28 | 0.65731689 | 0.047 | 0.431 | 5.74E-25 |

| | | | | | | | |
|---|---|---|---|---|---|---|---|
| Cardiomyocyte | Zfp385b | 7 | 1.80E-27 | 0.29463745 | 0.055 | 0.495 | 3.60E-24 |
| Cardiomyocyte | Medag1 | 7 | 1.90E-27 | 0.32915013 | 0.039 | 0.388 | 3.80E-24 |
| Cardiomyocyte | Grip22 | 7 | 2.28E-27 | 0.62413583 | 0.047 | 0.435 | 4.55E-24 |
| Cardiomyocyte | AABR07065190.1 | 7 | 3.88E-27 | 0.50745322 | 0.078 | 0.586 | 7.76E-24 |
| Cardiomyocyte | Lrrc4c | 7 | 7.57E-27 | 0.25006791 | 0.07 | 0.632 | 1.51E-23 |
| Cardiomyocyte | Col24a1 | 7 | 9.97E-27 | 0.33836379 | 0.094 | 0.64 | 1.99E-23 |
| Cardiomyocyte | Abi3bp | 7 | 1.86E-26 | 0.3596709 | 0.07 | 0.569 | 3.72E-23 |
| Cardiomyocyte | Cers6 | 7 | 1.96E-26 | 0.65581951 | 0.062 | 0.51 | 3.91E-23 |
| Cardiomyocyte | Sat1 | 7 | 2.23E-26 | 0.27258312 | 0.086 | 0.497 | 4.46E-23 |
| Cardiomyocyte | Dlg2 | 7 | 3.19E-26 | 0.35831238 | 0.07 | 0.561 | 6.38E-23 |
| Cardiomyocyte | Chsy31 | 7 | 5.44E-26 | 0.90836305 | 0.055 | 0.439 | 1.09E-22 |
| Cardiomyocyte | Specc1 | 7 | 5.91E-26 | 0.60119318 | 0.047 | 0.407 | 1.18E-22 |
| Cardiomyocyte | LOC6859631 | 7 | 8.19E-26 | 0.50981365 | 0.039 | 0.498 | 1.64E-22 |
| Cardiomyocyte | Kcnk2 | 7 | 9.09E-26 | 0.83959922 | 0.055 | 0.396 | 1.82E-22 |

| | | | | | | | |
|---|---|---|---|---|---|---|---|
| Cardiomyocyte | Plekha7 | 7 | 9.41E-26 | 0.56159768 | 0.055 | 0.565 | 1.88E-22 |
| Cardiomyocyte | Frem11 | 7 | 1.05E-25 | 0.59774274 | 0.078 | 0.561 | 2.10E-22 |
| Cardiomyocyte | Vcan1 | 7 | 1.21E-25 | 0.31945551 | 0.078 | 0.543 | 2.43E-22 |
| Cardiomyocyte | Rnf1521 | 7 | 1.39E-25 | 0.49709846 | 0.07 | 0.449 | 2.78E-22 |
| Cardiomyocyte | Apba1 | 7 | 1.45E-25 | 0.56534938 | 0.07 | 0.533 | 2.90E-22 |
| Cardiomyocyte | Dusp5 | 7 | 1.77E-25 | 2.15618909 | 0.07 | 0.442 | 3.53E-22 |
| Cardiomyocyte | Rfx21 | 7 | 2.15E-25 | 0.7228001 | 0.055 | 0.368 | 4.29E-22 |
| Cardiomyocyte | Zfp5361 | 7 | 2.25E-25 | 0.40516872 | 0.07 | 0.403 | 4.50E-22 |
| Cardiomyocyte | Slco2b1 | 7 | 2.27E-25 | 0.28618685 | 0.047 | 0.486 | 4.54E-22 |
| Cardiomyocyte | Ninj2 | 7 | 2.53E-25 | 0.45811968 | 0.078 | 0.429 | 5.06E-22 |
| Cardiomyocyte | Adamts51 | 7 | 2.85E-25 | 0.40817084 | 0.07 | 0.554 | 5.69E-22 |
| Cardiomyocyte | Sparcl1 | 7 | 3.06E-25 | 0.33126608 | 0.07 | 0.432 | 6.13E-22 |
| Cardiomyocyte | Ppp1r3c | 7 | 4.73E-25 | 0.75472658 | 0.078 | 0.508 | 9.47E-22 |
| Cardiomyocyte | Cspg4 | 7 | 5.44E-25 | 0.32914751 | 0.086 | 0.519 | 1.09E-21 |

| | | | | | | | |
|---|---|---|---|---|---|---|---|
| Cardiomyocyte | Ackr3 | 7 | 5.90E-25 | 0.79779098 | 0.062 | 0.563 | 1.18E-21 |
| Cardiomyocyte | LOC100910978.11 | 7 | 6.09E-25 | 0.32702237 | 0.031 | 0.407 | 1.22E-21 |
| Cardiomyocyte | AABR07003030.2 | 7 | 6.21E-25 | 0.42774363 | 0.062 | 0.388 | 1.24E-21 |
| Cardiomyocyte | Cdkn1a | 7 | 7.95E-25 | 0.67909902 | 0.062 | 0.605 | 1.59E-21 |
| Cardiomyocyte | Pcdh17 | 7 | 8.02E-25 | 0.3202943 | 0.078 | 0.492 | 1.60E-21 |
| Cardiomyocyte | Serpine11 | 7 | 1.10E-24 | 0.43796961 | 0.031 | 0.351 | 2.20E-21 |
| Cardiomyocyte | Gxylt21 | 7 | 1.41E-24 | 0.44681485 | 0.07 | 0.5 | 2.82E-21 |
| Cardiomyocyte | Gja3 | 7 | 1.81E-24 | 0.35646729 | 0.086 | 0.518 | 3.62E-21 |
| Cardiomyocyte | Map61 | 7 | 1.99E-24 | 0.7012174 | 0.078 | 0.382 | 3.99E-21 |
| Cardiomyocyte | Dpysl31 | 7 | 2.64E-24 | 0.85783954 | 0.078 | 0.424 | 5.28E-21 |
| Cardiomyocyte | Col27a1 | 7 | 2.89E-24 | 0.52125596 | 0.078 | 0.474 | 5.79E-21 |
| Cardiomyocyte | Cacna1h | 7 | 3.18E-24 | 0.6071064 | 0.094 | 0.569 | 6.35E-21 |
| Cardiomyocyte | Galnt161 | 7 | 3.71E-24 | 1.17208244 | 0.078 | 0.542 | 7.42E-21 |
| Cardiomyocyte | Igf12 | 7 | 4.17E-24 | 0.40171283 | 0.07 | 0.557 | 8.33E-21 |

| | | | | | | | |
|---|---|---|---|---|---|---|---|
| Cardiomyocyte | Pdgfc | 7 | 5.06E-24 | 1.22293951 | 0.047 | 0.393 | 1.01E-20 |
| Cardiomyocyte | Mamdc22 | 7 | 5.16E-24 | 0.78242394 | 0.055 | 0.538 | 1.03E-20 |
| Cardiomyocyte | Slc43a2 | 7 | 5.27E-24 | 0.38965487 | 0.086 | 0.57 | 1.05E-20 |
| Cardiomyocyte | Atp8b1 | 7 | 5.63E-24 | 0.35439832 | 0.086 | 0.338 | 1.13E-20 |
| Cardiomyocyte | Nrg1 | 7 | 5.70E-24 | 0.39579711 | 0.094 | 0.587 | 1.14E-20 |
| Cardiomyocyte | St6galnac3 | 7 | 6.00E-24 | 0.267706 | 0.062 | 0.403 | 1.20E-20 |
| Cardiomyocyte | Taldo1 | 7 | 6.29E-24 | 0.55535851 | 0.07 | 0.553 | 1.26E-20 |
| Cardiomyocyte | Mob3b | 7 | 6.47E-24 | 0.42847819 | 0.07 | 0.377 | 1.29E-20 |
| Cardiomyocyte | Lrrtm31 | 7 | 7.31E-24 | 0.45170567 | 0.117 | 0.59 | 1.46E-20 |
| Cardiomyocyte | Anxa11 | 7 | 7.33E-24 | 1.00038744 | 0.047 | 0.397 | 1.47E-20 |
| Cardiomyocyte | Kcnn3 | 7 | 7.49E-24 | 1.06201751 | 0.062 | 0.316 | 1.50E-20 |
| Cardiomyocyte | Sybu | 7 | 8.13E-24 | 0.52606713 | 0.07 | 0.364 | 1.63E-20 |
| Cardiomyocyte | Hivep3 | 7 | 8.64E-24 | 0.41549035 | 0.055 | 0.351 | 1.73E-20 |
| Cardiomyocyte | Arhgap39 | 7 | 1.12E-23 | 0.42835117 | 0.07 | 0.412 | 2.25E-20 |

| | | | | | | | |
|---|---|---|---|---|---|---|---|
| Cardiomyocyte | Gabrb2 | 7 | 1.29E-23 | 0.90749259 | 0.086 | 0.495 | 2.58E-20 |
| Cardiomyocyte | Aox31 | 7 | 1.56E-23 | 0.33270709 | 0.055 | 0.47 | 3.11E-20 |
| Cardiomyocyte | Syn3 | 7 | 1.65E-23 | 0.9055839 | 0.078 | 0.361 | 3.31E-20 |
| Cardiomyocyte | Tspan5 | 7 | 1.72E-23 | 0.42027009 | 0.062 | 0.407 | 3.44E-20 |
| Cardiomyocyte | Gucy1a1 | 7 | 1.92E-23 | 0.6529346 | 0.094 | 0.572 | 3.83E-20 |
| Cardiomyocyte | Klhl40 | 7 | 2.05E-23 | 2.69179306 | 0.055 | 0.47 | 4.09E-20 |
| Cardiomyocyte | Ar | 7 | 2.07E-23 | 1.17388056 | 0.07 | 0.517 | 4.15E-20 |
| Cardiomyocyte | Tenm3 | 7 | 2.30E-23 | 1.02793878 | 0.07 | 0.442 | 4.60E-20 |
| Cardiomyocyte | Kcnt2 | 7 | 2.63E-23 | 0.30874473 | 0.07 | 0.504 | 5.25E-20 |
| Cardiomyocyte | Crispld21 | 7 | 2.84E-23 | 0.87251488 | 0.086 | 0.453 | 5.68E-20 |
| Cardiomyocyte | Gask1b | 7 | 3.07E-23 | 1.12591259 | 0.07 | 0.435 | 6.15E-20 |
| Cardiomyocyte | Mylk | 7 | 3.15E-23 | 0.35570991 | 0.102 | 0.424 | 6.31E-20 |
| Cardiomyocyte | Samd5 | 7 | 3.49E-23 | 0.57449582 | 0.055 | 0.41 | 6.98E-20 |
| Cardiomyocyte | Hcn2 | 7 | 3.57E-23 | 0.47245354 | 0.086 | 0.463 | 7.13E-20 |

| | | | | | | | |
|---|---|---|---|---|---|---|---|
| Cardiomyocyte | Cacna2d3 | 7 | 4.58E-23 | 0.87043102 | 0.078 | 0.461 | 9.17E-20 |
| Cardiomyocyte | Ntf3 | 7 | 5.17E-23 | 0.57960945 | 0.086 | 0.519 | 1.03E-19 |
| Cardiomyocyte | Ptprn2 | 7 | 5.70E-23 | 1.18410362 | 0.031 | 0.291 | 1.14E-19 |
| Cardiomyocyte | AABR07059258.11 | 7 | 5.75E-23 | 0.70359866 | 0.062 | 0.447 | 1.15E-19 |
| Cardiomyocyte | Anks1b1 | 7 | 6.32E-23 | 0.30892432 | 0.102 | 0.541 | 1.26E-19 |
| Cardiomyocyte | Cobl1 | 7 | 6.54E-23 | 0.84885556 | 0.102 | 0.417 | 1.31E-19 |
| Cardiomyocyte | AABR07057997.1 | 7 | 7.18E-23 | 1.61332935 | 0.055 | 0.325 | 1.44E-19 |
| Cardiomyocyte | AABR07058170.1 | 7 | 8.46E-23 | 0.50302098 | 0.109 | 0.611 | 1.69E-19 |
| Cardiomyocyte | Sema3a1 | 7 | 8.63E-23 | 0.66320967 | 0.078 | 0.355 | 1.73E-19 |
| Cardiomyocyte | Cdh112 | 7 | 8.83E-23 | 0.43105963 | 0.094 | 0.52 | 1.77E-19 |
| Cardiomyocyte | Errfi11 | 7 | 9.81E-23 | 0.81926892 | 0.055 | 0.357 | 1.96E-19 |
| Cardiomyocyte | Fstl1 | 7 | 9.87E-23 | 0.66185582 | 0.102 | 0.55 | 1.97E-19 |
| Cardiomyocyte | Dkk31 | 7 | 1.01E-22 | 0.48044754 | 0.094 | 0.452 | 2.02E-19 |
| Cardiomyocyte | AABR07058158.1 | 7 | 1.16E-22 | 0.40910774 | 0.125 | 0.574 | 2.31E-19 |

| | | | | | | | |
|---|---|---|---|---|---|---|---|
| Cardiomyocyte | Tmem132d | 7 | 1.62E-22 | 0.82383972 | 0.07 | 0.368 | 3.24E-19 |
| Cardiomyocyte | Egflam1 | 7 | 1.64E-22 | 1.0787163 | 0.094 | 0.544 | 3.29E-19 |
| Cardiomyocyte | AABR07068046.12 | 7 | 1.66E-22 | 1.14020659 | 0.055 | 0.4 | 3.32E-19 |
| Cardiomyocyte | Cnnm2 | 7 | 1.81E-22 | 0.46062727 | 0.109 | 0.514 | 3.62E-19 |
| Cardiomyocyte | Alcam | 7 | 1.82E-22 | 0.39919379 | 0.094 | 0.505 | 3.64E-19 |
| Cardiomyocyte | Pde1a1 | 7 | 1.94E-22 | 0.97772144 | 0.109 | 0.535 | 3.88E-19 |
| Cardiomyocyte | Ddc1 | 7 | 2.34E-22 | 0.28116377 | 0.125 | 0.591 | 4.68E-19 |
| Cardiomyocyte | Aldh1a11 | 7 | 2.72E-22 | 0.69172909 | 0.102 | 0.484 | 5.43E-19 |
| Cardiomyocyte | Glis3 | 7 | 2.80E-22 | 1.00727485 | 0.055 | 0.431 | 5.60E-19 |
| Cardiomyocyte | Fbxl2 | 7 | 3.03E-22 | 0.42429768 | 0.117 | 0.644 | 6.06E-19 |
| Cardiomyocyte | Nr4a11 | 7 | 3.32E-22 | 0.28782059 | 0.117 | 0.655 | 6.64E-19 |
| Cardiomyocyte | Esr11 | 7 | 3.60E-22 | 0.38837757 | 0.109 | 0.523 | 7.21E-19 |
| Cardiomyocyte | Nkd2 | 7 | 3.65E-22 | 0.95208985 | 0.031 | 0.426 | 7.30E-19 |
| Cardiomyocyte | Kcnj12 | 7 | 3.70E-22 | 1.53872277 | 0.086 | 0.397 | 7.39E-19 |

| | | | | | | | |
|---|---|---|---|---|---|---|---|
| Cardiomyocyte | Astn21 | 7 | 3.91E-22 | 1.07541474 | 0.102 | 0.632 | 7.83E-19 |
| Cardiomyocyte | Anp32b | 7 | 4.88E-22 | 0.77564893 | 0.094 | 0.38 | 9.75E-19 |
| Cardiomyocyte | Uap11 | 7 | 5.31E-22 | 0.6539067 | 0.094 | 0.441 | 1.06E-18 |
| Cardiomyocyte | Chn2 | 7 | 5.98E-22 | 1.02753707 | 0.094 | 0.373 | 1.20E-18 |
| Cardiomyocyte | Cacna1d | 7 | 6.06E-22 | 1.27089625 | 0.086 | 0.548 | 1.21E-18 |
| Cardiomyocyte | Rerg | 7 | 6.99E-22 | 1.39409347 | 0.078 | 0.426 | 1.40E-18 |
| Cardiomyocyte | Npr3 | 7 | 7.21E-22 | 1.22952817 | 0.102 | 0.469 | 1.44E-18 |
| Cardiomyocyte | Med12l | 7 | 7.77E-22 | 1.44829573 | 0.086 | 0.406 | 1.55E-18 |
| Cardiomyocyte | Csrnp1 | 7 | 8.60E-22 | 0.85099759 | 0.055 | 0.333 | 1.72E-18 |
| Cardiomyocyte | AABR07049033.11 | 7 | 9.39E-22 | 0.88185105 | 0.062 | 0.414 | 1.88E-18 |
| Cardiomyocyte | Cd551 | 7 | 1.02E-21 | 1.73870073 | 0.055 | 0.459 | 2.05E-18 |
| Cardiomyocyte | RGD15633542 | 7 | 1.11E-21 | 0.50668095 | 0.055 | 0.336 | 2.22E-18 |
| Cardiomyocyte | Svep12 | 7 | 1.17E-21 | 0.26215349 | 0.023 | 0.488 | 2.33E-18 |
| Cardiomyocyte | Bcar3 | 7 | 1.21E-21 | 0.26997084 | 0.125 | 0.63 | 2.42E-18 |

| | | | | | | | |
|---|---|---|---|---|---|---|---|
| Cardiomyocyte | Il1rapl1 | 7 | 1.28E-21 | 0.73960933 | 0.047 | 0.435 | 2.56E-18 |
| Cardiomyocyte | Aff32 | 7 | 1.42E-21 | 0.82582724 | 0.102 | 0.59 | 2.84E-18 |
| Cardiomyocyte | Sgip1 | 7 | 1.48E-21 | 1.62048164 | 0.055 | 0.484 | 2.97E-18 |
| Cardiomyocyte | Col18a1 | 7 | 3.37E-21 | 1.15341119 | 0.086 | 0.466 | 6.75E-18 |
| Cardiomyocyte | Megf9 | 7 | 4.47E-21 | 0.51743691 | 0.055 | 0.398 | 8.93E-18 |
| Cardiomyocyte | Bdh11 | 7 | 5.86E-21 | 0.25259805 | 0.133 | 0.519 | 1.17E-17 |
| Cardiomyocyte | AABR07034940.2 | 7 | 7.15E-21 | 1.36389579 | 0.047 | 0.354 | 1.43E-17 |
| Cardiomyocyte | Angpt21 | 7 | 8.31E-21 | 0.87954959 | 0.031 | 0.336 | 1.66E-17 |
| Cardiomyocyte | Aox1 | 7 | 8.36E-21 | 0.34639202 | 0.055 | 0.333 | 1.67E-17 |
| Cardiomyocyte | Slit3 | 7 | 8.44E-21 | 1.53011967 | 0.109 | 0.623 | 1.69E-17 |
| Cardiomyocyte | Runx21 | 7 | 8.73E-21 | 1.71643455 | 0.062 | 0.374 | 1.75E-17 |
| Cardiomyocyte | Prkcq | 7 | 9.04E-21 | 1.14438782 | 0.055 | 0.429 | 1.81E-17 |
| Cardiomyocyte | Flrt2 | 7 | 1.20E-20 | 0.59928825 | 0.055 | 0.454 | 2.40E-17 |
| Cardiomyocyte | Cp | 7 | 1.38E-20 | 0.44819885 | 0.055 | 0.482 | 2.75E-17 |

| | | | | | | | |
|---|---|---|---|---|---|---|---|
| Cardiomyocyte | Xkr41 | 7 | 1.61E-20 | 0.35236147 | 0.133 | 0.53 | 3.23E-17 |
| Cardiomyocyte | Ptprt1 | 7 | 1.67E-20 | 1.80120396 | 0.078 | 0.41 | 3.33E-17 |
| Cardiomyocyte | Kcnab11 | 7 | 2.25E-20 | 2.14704866 | 0.062 | 0.448 | 4.49E-17 |
| Cardiomyocyte | Cdon1 | 7 | 2.30E-20 | 1.4183813 | 0.102 | 0.415 | 4.61E-17 |
| Cardiomyocyte | Sctr2 | 7 | 2.31E-20 | 0.93338107 | 0.125 | 0.548 | 4.62E-17 |
| Cardiomyocyte | Lmod11 | 7 | 2.38E-20 | 1.51080939 | 0.07 | 0.544 | 4.77E-17 |
| Cardiomyocyte | Ntrk3 | 7 | 2.49E-20 | 0.67622937 | 0.07 | 0.517 | 4.98E-17 |
| Cardiomyocyte | Ncam1 | 7 | 2.65E-20 | 0.30382635 | 0.133 | 0.621 | 5.30E-17 |
| Cardiomyocyte | Sox5 | 7 | 2.68E-20 | 2.05655016 | 0.109 | 0.572 | 5.36E-17 |
| Cardiomyocyte | Bmp6 | 7 | 3.16E-20 | 0.30924545 | 0.125 | 0.538 | 6.31E-17 |
| Cardiomyocyte | LOC103694210 | 7 | 3.16E-20 | 1.88035081 | 0.078 | 0.498 | 6.33E-17 |
| Cardiomyocyte | Myo1b | 7 | 4.17E-20 | 0.40971864 | 0.133 | 0.612 | 8.35E-17 |
| Cardiomyocyte | Enox2 | 7 | 4.23E-20 | 0.49497617 | 0.125 | 0.59 | 8.47E-17 |
| Cardiomyocyte | Plod22 | 7 | 4.37E-20 | 0.6798487 | 0.125 | 0.591 | 8.75E-17 |

| | | | | | | | |
|---|---|---|---|---|---|---|---|
| Cardiomyocyte | Gpm6b1 | 7 | 4.59E-20 | 1.10304649 | 0.031 | 0.37 | 9.17E-17 |
| Cardiomyocyte | Flna1 | 7 | 5.08E-20 | 0.37502938 | 0.141 | 0.572 | 1.02E-16 |
| Cardiomyocyte | Dapk1 | 7 | 6.42E-20 | 0.81543428 | 0.109 | 0.491 | 1.28E-16 |
| Cardiomyocyte | Zfp385d | 7 | 7.33E-20 | 0.32603949 | 0.125 | 0.571 | 1.47E-16 |
| Cardiomyocyte | Tmtc2 | 7 | 8.87E-20 | 0.29160011 | 0.125 | 0.58 | 1.77E-16 |
| Cardiomyocyte | Efhd1 | 7 | 9.28E-20 | 0.97019455 | 0.133 | 0.564 | 1.86E-16 |
| Cardiomyocyte | AABR07026021.1 | 7 | 1.04E-19 | 1.49711007 | 0.055 | 0.306 | 2.08E-16 |
| Cardiomyocyte | Myo5b | 7 | 1.07E-19 | 1.58058451 | 0.047 | 0.301 | 2.14E-16 |
| Cardiomyocyte | Csrp1 | 7 | 1.09E-19 | 0.52068577 | 0.07 | 0.479 | 2.19E-16 |
| Cardiomyocyte | Npas21 | 7 | 1.18E-19 | 0.27137991 | 0.133 | 0.389 | 2.37E-16 |
| Cardiomyocyte | Dcn | 7 | 1.22E-19 | 0.82351673 | 0.125 | 0.591 | 2.45E-16 |
| Cardiomyocyte | Ngf1 | 7 | 1.50E-19 | 1.597126 | 0.078 | 0.367 | 3.01E-16 |
| Cardiomyocyte | Csgalnact1 | 7 | 1.53E-19 | 0.33319114 | 0.031 | 0.349 | 3.05E-16 |
| Cardiomyocyte | Vgll3 | 7 | 1.71E-19 | 1.58612623 | 0.039 | 0.311 | 3.41E-16 |

| | | | | | | | |
|---|---|---|---|---|---|---|---|
| Cardiomyocyte | Adamts31 | 7 | 1.77E-19 | 1.12107323 | 0.086 | 0.42 | 3.54E-16 |
| Cardiomyocyte | Maoa | 7 | 1.80E-19 | 0.34691899 | 0.148 | 0.609 | 3.60E-16 |
| Cardiomyocyte | Pnpla3 | 7 | 3.35E-19 | 0.4638203 | 0.023 | 0.301 | 6.70E-16 |
| Cardiomyocyte | Synpo | 7 | 3.56E-19 | 0.25473004 | 0.133 | 0.6 | 7.12E-16 |
| Cardiomyocyte | Tmeff2 | 7 | 3.95E-19 | 0.83769721 | 0.133 | 0.569 | 7.90E-16 |
| Cardiomyocyte | Stk17b | 7 | 5.14E-19 | 1.84364057 | 0.078 | 0.438 | 1.03E-15 |
| Cardiomyocyte | Scn7a | 7 | 5.20E-19 | 1.07103776 | 0.109 | 0.516 | 1.04E-15 |
| Cardiomyocyte | Cacna1g | 7 | 6.09E-19 | 0.37783881 | 0.133 | 0.445 | 1.22E-15 |
| Cardiomyocyte | Serf11 | 7 | 8.21E-19 | 1.68225331 | 0.047 | 0.417 | 1.64E-15 |
| Cardiomyocyte | Myl41 | 7 | 9.78E-19 | 0.38574901 | 0.023 | 0.382 | 1.96E-15 |
| Cardiomyocyte | Gucy1a2 | 7 | 1.41E-18 | 1.42103073 | 0.125 | 0.471 | 2.81E-15 |
| Cardiomyocyte | C1qtnf7 | 7 | 1.85E-18 | 0.6007939 | 0.125 | 0.498 | 3.71E-15 |
| Cardiomyocyte | Ltbp1 | 7 | 2.24E-18 | 0.25226964 | 0.164 | 0.665 | 4.49E-15 |
| Cardiomyocyte | Col8a1 | 7 | 2.68E-18 | 0.27493615 | 0.164 | 0.631 | 5.36E-15 |

| | | | | | | | |
|---|---|---|---|---|---|---|---|
| Cardiomyocyte | Abca11 | 7 | 2.70E-18 | 0.40797374 | 0.141 | 0.592 | 5.41E-15 |
| Cardiomyocyte | Srgap1 | 7 | 2.70E-18 | 0.89796917 | 0.148 | 0.673 | 5.41E-15 |
| Cardiomyocyte | Tanc21 | 7 | 2.73E-18 | 0.38314711 | 0.156 | 0.484 | 5.47E-15 |
| Cardiomyocyte | Fam189a2 | 7 | 3.34E-18 | 0.26986176 | 0.156 | 0.58 | 6.69E-15 |
| Cardiomyocyte | Bche | 7 | 3.48E-18 | 0.76105117 | 0.141 | 0.621 | 6.96E-15 |
| Cardiomyocyte | Boc | 7 | 3.59E-18 | 0.2757992 | 0.188 | 0.713 | 7.18E-15 |
| Cardiomyocyte | Myof | 7 | 3.62E-18 | 0.40426944 | 0.156 | 0.505 | 7.23E-15 |
| Cardiomyocyte | Mgp | 7 | 5.74E-18 | 0.92178322 | 0.133 | 0.439 | 1.15E-14 |
| Cardiomyocyte | Sdc2 | 7 | 7.47E-18 | 0.96562201 | 0.148 | 0.463 | 1.49E-14 |
| Cardiomyocyte | Palmd | 7 | 1.16E-17 | 0.47970862 | 0.148 | 0.447 | 2.33E-14 |
| Cardiomyocyte | Kcnma11 | 7 | 1.24E-17 | 0.84992173 | 0.047 | 0.297 | 2.48E-14 |
| Cardiomyocyte | Kntc1 | 7 | 1.49E-17 | 1.46870435 | 0.055 | 0.468 | 2.98E-14 |
| Cardiomyocyte | Lck1 | 7 | 2.04E-17 | 0.84004267 | 0.047 | 0.451 | 4.07E-14 |
| Cardiomyocyte | Ednrb | 7 | 3.19E-17 | 0.58672207 | 0.031 | 0.403 | 6.38E-14 |

| | | | | | | | |
|---|---|---|---|---|---|---|---|
| Cardiomyocyte | AABR07052441.12 | 7 | 3.24E-17 | 0.97497171 | 0.023 | 0.363 | 6.49E-14 |
| Cardiomyocyte | Ppp1r9a1 | 7 | 3.38E-17 | 0.56842563 | 0.164 | 0.576 | 6.77E-14 |
| Cardiomyocyte | Fgf101 | 7 | 4.47E-17 | 1.2677729 | 0.031 | 0.379 | 8.94E-14 |
| Cardiomyocyte | Slc7a11 | 7 | 4.55E-17 | 0.61084753 | 0.156 | 0.472 | 9.10E-14 |
| Cardiomyocyte | Pik3r51 | 7 | 5.63E-17 | 0.81118884 | 0.039 | 0.322 | 1.13E-13 |
| Cardiomyocyte | Satb2 | 7 | 6.63E-17 | 0.25618978 | 0.023 | 0.318 | 1.33E-13 |
| Cardiomyocyte | Tmem47 | 7 | 9.92E-17 | 0.60525626 | 0.047 | 0.433 | 1.98E-13 |
| Cardiomyocyte | Plcxd3 | 7 | 1.15E-16 | 1.90178547 | 0.039 | 0.372 | 2.31E-13 |
| Cardiomyocyte | Ikzf2 | 7 | 1.19E-16 | 0.55563432 | 0.172 | 0.646 | 2.37E-13 |
| Cardiomyocyte | Gmds | 7 | 1.21E-16 | 0.58048326 | 0.164 | 0.527 | 2.42E-13 |
| Cardiomyocyte | Nnat2 | 7 | 1.26E-16 | 1.00956306 | 0.023 | 0.41 | 2.51E-13 |
| Cardiomyocyte | Tshr1 | 7 | 1.31E-16 | 1.60995094 | 0.047 | 0.389 | 2.61E-13 |
| Cardiomyocyte | Edil31 | 7 | 1.43E-16 | 0.33551019 | 0.055 | 0.384 | 2.86E-13 |
| Cardiomyocyte | Efna5 | 7 | 1.54E-16 | 1.41684361 | 0.062 | 0.399 | 3.09E-13 |

| | | | | | | | |
|---|---|---|---|---|---|---|---|
| Cardiomyocyte | Rem1 | 7 | 2.03E-16 | 1.02764227 | 0.023 | 0.308 | 4.07E-13 |
| Cardiomyocyte | Stk38l1 | 7 | 2.06E-16 | 0.31297746 | 0.172 | 0.516 | 4.13E-13 |
| Cardiomyocyte | Tmcc3 | 7 | 2.25E-16 | 0.68603286 | 0.164 | 0.449 | 4.49E-13 |
| Cardiomyocyte | Plbd11 | 7 | 3.62E-16 | 0.42109722 | 0.172 | 0.471 | 7.25E-13 |
| Cardiomyocyte | Flvcr2 | 7 | 3.75E-16 | 1.20967389 | 0.047 | 0.324 | 7.50E-13 |
| Cardiomyocyte | Rtn4rl1 | 7 | 4.17E-16 | 0.78886434 | 0.172 | 0.642 | 8.34E-13 |
| Cardiomyocyte | Adgrd1 | 7 | 5.33E-16 | 1.16637872 | 0.031 | 0.296 | 1.07E-12 |
| Cardiomyocyte | Blnk1 | 7 | 5.56E-16 | 2.38509383 | 0.07 | 0.491 | 1.11E-12 |
| Cardiomyocyte | Mctp11 | 7 | 6.15E-16 | 0.43881714 | 0.156 | 0.46 | 1.23E-12 |
| Cardiomyocyte | Ndufa4 | 7 | 6.38E-16 | 0.86347337 | 0.164 | 0.626 | 1.28E-12 |
| Cardiomyocyte | Prrx1 | 7 | 9.97E-16 | 1.25150259 | 0.086 | 0.403 | 1.99E-12 |
| Cardiomyocyte | Itgbl1 | 7 | 1.51E-15 | 0.69187156 | 0.18 | 0.59 | 3.02E-12 |
| Cardiomyocyte | Mt11 | 7 | 1.83E-15 | 0.96411687 | 0.023 | 0.38 | 3.67E-12 |
| Cardiomyocyte | Pcdh12 | 7 | 2.13E-15 | 1.5177093 | 0.039 | 0.357 | 4.26E-12 |

| | | | | | | | |
|---|---|---|---|---|---|---|---|
| Cardiomyocyte | Carmil11 | 7 | 3.41E-15 | 0.29032638 | 0.203 | 0.61 | 6.81E-12 |
| Cardiomyocyte | Bcl11a | 7 | 3.54E-15 | 0.43273348 | 0.203 | 0.599 | 7.08E-12 |
| Cardiomyocyte | Nxn | 7 | 3.78E-15 | 0.93788468 | 0.18 | 0.661 | 7.57E-12 |
| Cardiomyocyte | AABR07031164.1 | 7 | 3.93E-15 | 3.41451366 | 0.062 | 0.434 | 7.85E-12 |
| Cardiomyocyte | Slfn131 | 7 | 5.88E-15 | 1.74188649 | 0.055 | 0.343 | 1.18E-11 |
| Cardiomyocyte | Gal3st31 | 7 | 7.13E-15 | 0.38892656 | 0.195 | 0.529 | 1.43E-11 |
| Cardiomyocyte | Postn1 | 7 | 7.52E-15 | 0.45416538 | 0.055 | 0.344 | 1.50E-11 |
| Cardiomyocyte | Kit | 7 | 7.53E-15 | 2.51304282 | 0.109 | 0.477 | 1.51E-11 |
| Cardiomyocyte | Unc5c | 7 | 8.06E-15 | 0.69298764 | 0.062 | 0.387 | 1.61E-11 |
| Cardiomyocyte | Col4a3 | 7 | 9.60E-15 | 0.33913804 | 0.203 | 0.588 | 1.92E-11 |
| Cardiomyocyte | Col3a1 | 7 | 9.65E-15 | 0.73212791 | 0.188 | 0.583 | 1.93E-11 |
| Cardiomyocyte | Adgrl31 | 7 | 1.59E-14 | 2.10295685 | 0.102 | 0.46 | 3.19E-11 |
| Cardiomyocyte | Osbp2 | 7 | 3.25E-14 | 0.90238037 | 0.172 | 0.465 | 6.50E-11 |
| Cardiomyocyte | Tex141 | 7 | 4.13E-14 | 1.65745923 | 0.031 | 0.354 | 8.25E-11 |

| | | | | | | | |
|---|---|---|---|---|---|---|---|
| Cardiomyocyte | Timp3 | 7 | 5.73E-14 | 0.54278354 | 0.219 | 0.653 | 1.15E-10 |
| Cardiomyocyte | Chst15 | 7 | 6.68E-14 | 0.68046583 | 0.211 | 0.607 | 1.34E-10 |
| Cardiomyocyte | Trps1 | 7 | 7.88E-14 | 0.42141962 | 0.211 | 0.653 | 1.58E-10 |
| Cardiomyocyte | Cobll1 | 7 | 8.74E-14 | 0.31860569 | 0.234 | 0.664 | 1.75E-10 |
| Cardiomyocyte | Sema5a | 7 | 1.06E-13 | 0.4762152 | 0.227 | 0.71 | 2.11E-10 |
| Cardiomyocyte | Rxfp1 | 7 | 1.18E-13 | 0.82094447 | 0.195 | 0.538 | 2.37E-10 |
| Cardiomyocyte | Fbn1 | 7 | 1.38E-13 | 0.89965933 | 0.188 | 0.541 | 2.76E-10 |
| Cardiomyocyte | Rhoh | 7 | 1.85E-13 | 2.24290095 | 0.055 | 0.322 | 3.70E-10 |
| Cardiomyocyte | Ltbp2 | 7 | 1.99E-13 | 1.36013512 | 0.031 | 0.318 | 3.99E-10 |
| Cardiomyocyte | Sfxn5 | 7 | 2.05E-13 | 0.26179342 | 0.227 | 0.596 | 4.10E-10 |
| Cardiomyocyte | Ptpn181 | 7 | 2.05E-13 | 1.07714241 | 0.047 | 0.315 | 4.10E-10 |
| Cardiomyocyte | Lepr1 | 7 | 2.10E-13 | 0.55367271 | 0.062 | 0.373 | 4.20E-10 |
| Cardiomyocyte | Greb1l1 | 7 | 2.10E-13 | 0.74288737 | 0.047 | 0.384 | 4.21E-10 |
| Cardiomyocyte | Pip5k1b1 | 7 | 2.61E-13 | 0.34351405 | 0.219 | 0.582 | 5.21E-10 |

| | | | | | | | |
|---|---|---|---|---|---|---|---|
| Cardiomyoc yte | Cenpf | 7 | 6.88E-13 | 1.8815503 | 0.031 | 0.336 | 1.38E-09 |
| Cardiomyoc yte | Slc44a51 | 7 | 7.44E-13 | 0.51796395 | 0.055 | 0.343 | 1.49E-09 |
| Cardiomyoc yte | Cadm2 | 7 | 9.42E-13 | 1.00411634 | 0.039 | 0.35 | 1.88E-09 |
| Cardiomyoc yte | Camk4 | 7 | 1.03E-12 | 2.29233418 | 0.062 | 0.387 | 2.06E-09 |
| Cardiomyoc yte | Slc16a10 | 7 | 1.12E-12 | 0.39669237 | 0.242 | 0.745 | 2.25E-09 |
| Cardiomyoc yte | Map1b1 | 7 | 1.35E-12 | 0.31181006 | 0.234 | 0.541 | 2.70E-09 |
| Cardiomyoc yte | Ebf2 | 7 | 1.36E-12 | 1.15989289 | 0.203 | 0.617 | 2.72E-09 |
| Cardiomyoc yte | Neurod4 | 7 | 2.05E-12 | 1.19326932 | 0.031 | 0.291 | 4.10E-09 |
| Cardiomyoc yte | Plxdc2 | 7 | 3.18E-12 | 1.10647035 | 0.195 | 0.566 | 6.36E-09 |
| Cardiomyoc yte | Grid21 | 7 | 3.51E-12 | 1.57366432 | 0.055 | 0.326 | 7.02E-09 |
| Cardiomyoc yte | Zeb2 | 7 | 3.55E-12 | 0.40087536 | 0.25 | 0.664 | 7.10E-09 |
| Cardiomyoc yte | Bicc11 | 7 | 5.16E-12 | 0.69090049 | 0.219 | 0.658 | 1.03E-08 |
| Cardiomyoc yte | Slc16a1 | 7 | 6.78E-12 | 0.3930911 | 0.234 | 0.674 | 1.36E-08 |
| Cardiomyoc yte | Itgb41 | 7 | 1.48E-11 | 1.79938678 | 0.055 | 0.321 | 2.96E-08 |

| | | | | | | | |
|---|---|---|---|---|---|---|---|
| Cardiomyocyte | Art3 | 7 | 1.54E-11 | 0.25199293 | 0.258 | 0.641 | 3.09E-08 |
| Cardiomyocyte | St3gal5 | 7 | 1.92E-11 | 0.32341947 | 0.242 | 0.601 | 3.84E-08 |
| Cardiomyocyte | Lgr6 | 7 | 2.55E-11 | 0.32399193 | 0.273 | 0.695 | 5.10E-08 |
| Cardiomyocyte | Hyi1 | 7 | 2.59E-11 | 3.5121295 | 0.031 | 0.321 | 5.19E-08 |
| Cardiomyocyte | Galnt151 | 7 | 5.24E-11 | 0.90342539 | 0.023 | 0.305 | 1.05E-07 |
| Cardiomyocyte | Adamts6 | 7 | 1.36E-10 | 0.29905098 | 0.273 | 0.692 | 2.72E-07 |
| Cardiomyocyte | Nrxn1 | 7 | 1.57E-10 | 1.21735069 | 0.242 | 0.695 | 3.14E-07 |
| Cardiomyocyte | Slc6a6 | 7 | 1.76E-10 | 0.79747031 | 0.289 | 0.745 | 3.52E-07 |
| Cardiomyocyte | Oasl2 | 7 | 2.32E-10 | 1.86346909 | 0.023 | 0.285 | 4.65E-07 |
| Cardiomyocyte | Maob | 7 | 2.42E-10 | 2.62470361 | 0.062 | 0.326 | 4.85E-07 |
| Cardiomyocyte | Kcnip1 | 7 | 2.72E-10 | 1.53603725 | 0.039 | 0.308 | 5.44E-07 |
| Cardiomyocyte | Dcx2 | 7 | 2.77E-10 | 0.29174948 | 0.008 | 0.259 | 5.53E-07 |
| Cardiomyocyte | Arhgap22 | 7 | 2.93E-10 | 1.07583335 | 0.031 | 0.298 | 5.85E-07 |
| Cardiomyocyte | Gpr176 | 7 | 3.16E-10 | 2.86368823 | 0.07 | 0.331 | 6.32E-07 |

| | | | | | | | |
|---|---|---|---|---|---|---|---|
| Cardiomyocyte | Dpf3 | 7 | 4.79E-10 | 0.27360372 | 0.289 | 0.739 | 9.59E-07 |
| Cardiomyocyte | Fam78b | 7 | 6.05E-10 | 0.53904917 | 0.242 | 0.533 | 1.21E-06 |
| Cardiomyocyte | Nceh1 | 7 | 9.05E-10 | 0.90479003 | 0.258 | 0.535 | 1.81E-06 |
| Cardiomyocyte | Epb41l4b | 7 | 1.71E-09 | 0.48955178 | 0.289 | 0.722 | 3.42E-06 |
| Cardiomyocyte | Pappa21 | 7 | 2.24E-09 | 2.66388044 | 0.055 | 0.318 | 4.47E-06 |
| Cardiomyocyte | Ntn4 | 7 | 2.40E-09 | 0.42152684 | 0.281 | 0.74 | 4.80E-06 |
| Cardiomyocyte | Ppp1r3a2 | 7 | 2.82E-09 | 0.30411312 | 0.289 | 0.586 | 5.64E-06 |
| Cardiomyocyte | Slco5a1 | 7 | 4.81E-09 | 0.30758623 | 0.32 | 0.768 | 9.61E-06 |
| Cardiomyocyte | Slc25a131 | 7 | 1.02E-08 | 0.59200852 | 0.297 | 0.614 | 2.04E-05 |
| Cardiomyocyte | Atp1a11 | 7 | 2.02E-08 | 0.44742464 | 0.305 | 0.629 | 4.04E-05 |
| Cardiomyocyte | Atp2b2 | 7 | 3.37E-08 | 0.71505877 | 0.305 | 0.727 | 6.73E-05 |
| Cardiomyocyte | Smpx | 7 | 5.73E-08 | 0.38828827 | 0.328 | 0.738 | 0.00011452 |
| Cardiomyocyte | Fign2 | 7 | 7.37E-08 | 0.48584773 | 0.305 | 0.604 | 0.00014744 |
| Cardiomyocyte | Tmem182 | 7 | 1.24E-07 | 0.39174512 | 0.336 | 0.769 | 0.00024806 |

| | | | | | | | |
|---|---|---|---|---|---|---|---|
| Cardiomyocyte | Ntn1 | 7 | 5.08E-07 | 0.44771809 | 0.336 | 0.726 | 0.00101534 |
| Cardiomyocyte | Acyp2 | 7 | 8.52E-07 | 0.48834509 | 0.328 | 0.657 | 0.00170415 |
| Cardiomyocyte | Me31 | 7 | 1.54E-05 | 0.42572273 | 0.375 | 0.688 | 0.03086763 |
| Cardiomyocyte | Smyd1 | 7 | 2.08E-05 | 0.26311004 | 0.375 | 0.777 | 0.04166163 |
| Cardiomyocyte | Nsg2 | 8 | 2.21E-105 | 2.98646587 | 0.508 | 0.018 | 4.43E-102 |
| Cardiomyocyte | Tacr1 | 8 | 8.67E-103 | 3.95776252 | 0.508 | 0.019 | 1.73E-99 |
| Cardiomyocyte | Cpne7 | 8 | 6.90E-80 | 2.79919667 | 0.525 | 0.042 | 1.38E-76 |
| Cardiomyocyte | Cdkn3 | 8 | 6.22E-77 | 1.70055204 | 0.425 | 0.021 | 1.24E-73 |
| Cardiomyocyte | AABR07048321.1 | 8 | 6.22E-77 | 1.70055204 | 0.425 | 0.021 | 1.24E-73 |
| Cardiomyocyte | Marchf4 | 8 | 1.24E-75 | 0.43231247 | 0.425 | 0.022 | 2.48E-72 |
| Cardiomyocyte | Cdh1 | 8 | 3.19E-56 | 4.56644246 | 0.525 | 0.069 | 6.38E-53 |
| Cardiomyocyte | AABR07033925.1 | 8 | 1.84E-53 | 3.44154593 | 0.425 | 0.052 | 3.67E-50 |
| Cardiomyocyte | Fosb | 8 | 1.08E-52 | 3.30364137 | 0.442 | 0.05 | 2.17E-49 |
| Cardiomyocyte | Lypd1 | 8 | 3.98E-52 | 2.76008329 | 0.5 | 0.015 | 7.95E-49 |

| | | | | | | | |
|---|---|---|---|---|---|---|---|
| Cardiomyocyte | Fcgr3a | 8 | 1.10E-51 | 4.10842561 | 0.267 | 0.011 | 2.20E-48 |
| Cardiomyocyte | C5ar1 | 8 | 5.48E-46 | 1.69339405 | 0.475 | 0.013 | 1.10E-42 |
| Cardiomyocyte | Prima1 | 8 | 6.26E-46 | 1.84448727 | 0.575 | 0.089 | 1.25E-42 |
| Cardiomyocyte | Ildr2 | 8 | 1.26E-43 | 4.38167934 | 0.725 | 0.18 | 2.52E-40 |
| Cardiomyocyte | Otulinl | 8 | 1.79E-42 | 0.40684865 | 0.517 | 0.091 | 3.58E-39 |
| Cardiomyocyte | Trmt9b | 8 | 1.31E-38 | 2.5874951 | 0.525 | 0.046 | 2.61E-35 |
| Cardiomyocyte | Aoah1 | 8 | 2.98E-37 | 1.25983502 | 0.542 | 0.135 | 5.96E-34 |
| Cardiomyocyte | AABR07003304.2 | 8 | 1.18E-35 | 1.82109992 | 0.4 | 0.059 | 2.36E-32 |
| Cardiomyocyte | Ptk2b1 | 8 | 1.71E-34 | 2.63438145 | 0.583 | 0.084 | 3.43E-31 |
| Cardiomyocyte | Tex22 | 8 | 5.83E-30 | 1.1185037 | 0.358 | 0.037 | 1.17E-26 |
| Cardiomyocyte | Cpa6 | 8 | 2.21E-29 | 2.32249045 | 0.583 | 0.149 | 4.42E-26 |
| Cardiomyocyte | Ly49s5 | 8 | 8.51E-29 | 0.8726009 | 0.558 | 0.161 | 1.70E-25 |
| Cardiomyocyte | Tspan331 | 8 | 2.45E-28 | 1.57688244 | 0.517 | 0.127 | 4.90E-25 |
| Cardiomyocyte | AABR07054716.1 | 8 | 1.05E-27 | 3.01186165 | 0.492 | 0.141 | 2.10E-24 |

| | | | | | | | |
|---|---|---|---|---|---|---|---|
| Cardiomyocyte | Cmtm51 | 8 | 2.87E-27 | 1.3367028 | 0.442 | 0.099 | 5.73E-24 |
| Cardiomyocyte | Siah31 | 8 | 5.54E-27 | 0.36914048 | 0.442 | 0.1 | 1.11E-23 |
| Cardiomyocyte | Adora2a | 8 | 8.54E-26 | 0.37533927 | 0.533 | 0.159 | 1.71E-22 |
| Cardiomyocyte | Alox5 | 8 | 1.41E-24 | 1.48055157 | 0.508 | 0.153 | 2.83E-21 |
| Cardiomyocyte | Myof1 | 8 | 6.04E-24 | 1.912595 | 0.833 | 0.44 | 1.21E-20 |
| Cardiomyocyte | Gabre | 8 | 1.71E-23 | 3.13889769 | 0.292 | 0.017 | 3.42E-20 |
| Cardiomyocyte | Tgfb21 | 8 | 1.46E-22 | 2.96931092 | 0.8 | 0.55 | 2.91E-19 |
| Cardiomyocyte | St141 | 8 | 8.09E-22 | 0.78698769 | 0.425 | 0.1 | 1.62E-18 |
| Cardiomyocyte | Mboat2 | 8 | 9.02E-22 | 1.70218149 | 0.525 | 0.156 | 1.80E-18 |
| Cardiomyocyte | Spetex-2F1 | 8 | 1.60E-21 | 0.57512342 | 0.342 | 0.071 | 3.20E-18 |
| Cardiomyocyte | Has1 | 8 | 1.72E-20 | 1.42855984 | 0.417 | 0.033 | 3.45E-17 |
| Cardiomyocyte | Cacna1g1 | 8 | 1.76E-20 | 2.64052434 | 0.75 | 0.386 | 3.51E-17 |
| Cardiomyocyte | Pak1 | 8 | 2.84E-20 | 1.26062081 | 0.55 | 0.126 | 5.68E-17 |
| Cardiomyocyte | Abca8 | 8 | 8.87E-20 | 1.51038848 | 0.592 | 0.19 | 1.77E-16 |

| | | | | | | | |
|---|---|---|---|---|---|---|---|
| Cardiomyocyte | Ikzf1 | 8 | 2.96E-19 | 1.02053504 | 0.542 | 0.047 | 5.92E-16 |
| Cardiomyocyte | C1qtnf11 | 8 | 1.08E-18 | 1.14353506 | 0.342 | 0.087 | 2.16E-15 |
| Cardiomyocyte | Ctse1 | 8 | 1.88E-18 | 2.98359571 | 0.342 | 0.082 | 3.76E-15 |
| Cardiomyocyte | Adgrg3 | 8 | 6.47E-18 | 0.87826163 | 0.35 | 0.097 | 1.29E-14 |
| Cardiomyocyte | Rbm441 | 8 | 1.55E-17 | 2.6297503 | 0.567 | 0.174 | 3.10E-14 |
| Cardiomyocyte | Ighm | 8 | 3.85E-17 | 1.75393936 | 0.55 | 0.27 | 7.70E-14 |
| Cardiomyocyte | Bdkrb21 | 8 | 1.32E-16 | 2.92659956 | 0.508 | 0.152 | 2.65E-13 |
| Cardiomyocyte | Carmil12 | 8 | 4.18E-16 | 1.70448365 | 0.825 | 0.55 | 8.36E-13 |
| Cardiomyocyte | Arhgef261 | 8 | 4.48E-16 | 1.77959464 | 0.55 | 0.288 | 8.96E-13 |
| Cardiomyocyte | Acta1 | 8 | 6.88E-16 | 3.23453162 | 0.667 | 0.323 | 1.38E-12 |
| Cardiomyocyte | AABR07017902.12 | 8 | 1.49E-15 | 0.59631261 | 0.542 | 0.206 | 2.98E-12 |
| Cardiomyocyte | Epsti11 | 8 | 1.75E-15 | 1.85142953 | 0.458 | 0.184 | 3.51E-12 |
| Cardiomyocyte | Tex142 | 8 | 2.94E-15 | 0.66190334 | 0.558 | 0.303 | 5.89E-12 |
| Cardiomyocyte | Ankrd55 | 8 | 3.15E-14 | 1.23471326 | 0.533 | 0.25 | 6.30E-11 |

| | | | | | | | |
|---|---|---|---|---|---|---|---|
| Cardiomyocyte | Vit1 | 8 | 6.47E-14 | 2.89070659 | 0.392 | 0.132 | 1.29E-10 |
| Cardiomyocyte | Kcnma12 | 8 | 7.74E-14 | 1.99650895 | 0.592 | 0.245 | 1.55E-10 |
| Cardiomyocyte | Ptger3 | 8 | 1.65E-13 | 3.65159321 | 0.492 | 0.228 | 3.31E-10 |
| Cardiomyocyte | Spag17 | 8 | 1.32E-12 | 1.06452876 | 0.433 | 0.054 | 2.65E-09 |
| Cardiomyocyte | Cyfip21 | 8 | 1.53E-12 | 1.14516168 | 0.558 | 0.269 | 3.06E-09 |
| Cardiomyocyte | Frmd3 | 8 | 1.84E-12 | 2.98591952 | 0.592 | 0.105 | 3.67E-09 |
| Cardiomyocyte | Ntng2 | 8 | 2.18E-12 | 0.46841862 | 0.542 | 0.159 | 4.35E-09 |
| Cardiomyocyte | Wt1 | 8 | 2.23E-12 | 1.79889683 | 0.55 | 0.135 | 4.46E-09 |
| Cardiomyocyte | Ndrg41 | 8 | 2.67E-12 | 1.30157002 | 0.85 | 0.552 | 5.35E-09 |
| Cardiomyocyte | Fcer1g2 | 8 | 3.83E-12 | 0.5413719 | 0.15 | 0.509 | 7.65E-09 |
| Cardiomyocyte | Smox | 8 | 4.72E-12 | 1.85702222 | 0.583 | 0.223 | 9.45E-09 |
| Cardiomyocyte | Kank11 | 8 | 6.18E-12 | 1.0713547 | 0.792 | 0.521 | 1.24E-08 |
| Cardiomyocyte | Gjc1 | 8 | 6.32E-12 | 1.84259034 | 0.642 | 0.258 | 1.26E-08 |
| Cardiomyocyte | Hao11 | 8 | 6.40E-12 | 0.83699241 | 0.525 | 0.236 | 1.28E-08 |

| | | | | | | | |
|---|---|---|---|---|---|---|---|
| Cardiomyocyte | AABR07016779.1 | 8 | 9.99E-12 | 2.15324482 | 0.492 | 0.143 | 2.00E-08 |
| Cardiomyocyte | Slc24a3 | 8 | 1.99E-11 | 3.10708519 | 0.575 | 0.165 | 3.98E-08 |
| Cardiomyocyte | Map1b2 | 8 | 5.04E-11 | 1.21128835 | 0.758 | 0.491 | 1.01E-07 |
| Cardiomyocyte | Sh3bp21 | 8 | 6.08E-11 | 0.75973587 | 0.558 | 0.22 | 1.22E-07 |
| Cardiomyocyte | Stk38l2 | 8 | 7.13E-11 | 1.40371598 | 0.717 | 0.463 | 1.43E-07 |
| Cardiomyocyte | Bcl6b1 | 8 | 5.34E-10 | 0.30874088 | 0.55 | 0.113 | 1.07E-06 |
| Cardiomyocyte | Syn31 | 8 | 1.39E-09 | 1.147177 | 0.633 | 0.307 | 2.77E-06 |
| Cardiomyocyte | Serpine12 | 8 | 2.57E-09 | 1.60406754 | 0.575 | 0.299 | 5.15E-06 |
| Cardiomyocyte | Magi21 | 8 | 2.89E-09 | 1.58084967 | 0.533 | 0.278 | 5.77E-06 |
| Cardiomyocyte | Glis31 | 8 | 4.32E-09 | 1.00942633 | 0.625 | 0.375 | 8.63E-06 |
| Cardiomyocyte | Ddah1 | 8 | 6.24E-09 | 0.96896141 | 0.742 | 0.464 | 1.25E-05 |
| Cardiomyocyte | Hivep31 | 8 | 6.63E-09 | 0.26996761 | 0.083 | 0.347 | 1.33E-05 |
| Cardiomyocyte | Aqp1 | 8 | 1.50E-08 | 0.27348904 | 0.55 | 0.195 | 2.99E-05 |
| Cardiomyocyte | Mgmt | 8 | 2.34E-08 | 1.07311767 | 0.658 | 0.376 | 4.67E-05 |

| | | | | | | | |
|---|---|---|---|---|---|---|---|
| Cardiomyocyte | Satb21 | 8 | 4.53E-08 | 0.42937541 | 0.533 | 0.269 | 9.06E-05 |
| Cardiomyocyte | Lyz21 | 8 | 4.87E-08 | 0.69115272 | 0.433 | 0.149 | 9.73E-05 |
| Cardiomyocyte | Pdp21 | 8 | 2.23E-07 | 0.41303992 | 0.567 | 0.274 | 0.00044618 |
| Cardiomyocyte | AABR07034940.21 | 8 | 3.19E-07 | 0.35163651 | 0.583 | 0.303 | 0.00063725 |
| Cardiomyocyte | Sod2 | 8 | 5.06E-07 | 0.67161531 | 0.575 | 0.324 | 0.00101108 |
| Cardiomyocyte | Manf | 8 | 7.50E-07 | 0.29397117 | 0.542 | 0.176 | 0.00150093 |
| Cardiomyocyte | Prrg4 | 8 | 8.15E-07 | 0.89879391 | 0.058 | 0.464 | 0.00163077 |
| Cardiomyocyte | Prg4 | 8 | 1.30E-06 | 0.88265955 | 0.458 | 0.191 | 0.0025937 |
| Cardiomyocyte | Rcan1 | 8 | 1.79E-06 | 2.14262505 | 0.492 | 0.232 | 0.00357527 |
| Cardiomyocyte | Fam227b | 8 | 2.95E-06 | 0.65169851 | 0.692 | 0.339 | 0.00590271 |
| Cardiomyocyte | Rem11 | 8 | 3.26E-06 | 0.82625655 | 0.517 | 0.26 | 0.00651306 |
| Cardiomyocyte | Fbln1 | 8 | 3.56E-06 | 1.22466658 | 0.533 | 0.269 | 0.00712101 |
| Cardiomyocyte | Slc66a1 | 8 | 4.55E-06 | 0.8462904 | 0.417 | 0.125 | 0.00909917 |
| Cardiomyocyte | Fndc1 | 8 | 5.14E-06 | 0.71040759 | 0.592 | 0.324 | 0.01027371 |

| | | | | | | | |
|---|---|---|---|---|---|---|---|
| Cardiomyocyte | Gucy1b1 | 8 | 1.45E-05 | 1.07016396 | 0.317 | 0.037 | 0.02890257 |
| Cardiomyocyte | Rasl10b | 9 | 1.35E-23 | 2.12364372 | 0.361 | 0.05 | 2.70E-20 |
| Cardiomyocyte | Catip | 9 | 1.10E-11 | 1.43401248 | 0.443 | 0.116 | 2.21E-08 |
| Cardiomyocyte | Trib3 | 9 | 3.08E-11 | 2.97570899 | 0.377 | 0.109 | 6.17E-08 |
| Cardiomyocyte | Otulinl1 | 9 | 4.30E-11 | 2.8258064 | 0.393 | 0.115 | 8.59E-08 |
| Cardiomyocyte | Prss482 | 9 | 1.13E-10 | 1.38800804 | 0.426 | 0.142 | 2.25E-07 |
| Cardiomyocyte | Trpc31 | 9 | 1.12E-09 | 1.81473889 | 0.492 | 0.189 | 2.23E-06 |
| Cardiomyocyte | AABR07007068.1 | 9 | 1.38E-09 | 0.50947222 | 0.426 | 0.15 | 2.77E-06 |
| Cardiomyocyte | Gcat1 | 9 | 3.23E-09 | 2.81638346 | 0.525 | 0.217 | 6.45E-06 |
| Cardiomyocyte | Bmpr1b | 9 | 4.47E-08 | 1.01515228 | 0.443 | 0.175 | 8.94E-05 |
| Cardiomyocyte | Lurap1l1 | 9 | 5.91E-08 | 1.51601693 | 0.377 | 0.124 | 0.00011813 |
| Cardiomyocyte | Gpr63 | 9 | 5.70E-07 | 2.13166591 | 0.525 | 0.238 | 0.00114066 |
| Cardiomyocyte | LOC1025530181 | 9 | 1.20E-06 | 1.99085655 | 0.508 | 0.25 | 0.00239512 |
| Cardiomyocyte | Tmem178a | 10 | 9.34E-51 | 1.3035441 | 0.327 | 0.029 | 1.87E-47 |

| | | | | | | | |
|---|---|---|---|---|---|---|---|
| Cardiomyocyte | Igfbp6 | 10 | 2.21E-22 | 4.26170239 | 0.538 | 0.082 | 4.42E-19 |
| Cardiomyocyte | Igfbp41 | 10 | 1.64E-19 | 3.9956038 | 0.385 | 0.089 | 3.28E-16 |
| Cardiomyocyte | Nostrin | 10 | 5.49E-16 | 1.74943208 | 0.385 | 0.059 | 1.10E-12 |
| Cardiomyocyte | Aldh1a21 | 10 | 7.24E-16 | 0.5419663 | 0.462 | 0.097 | 1.45E-12 |
| Cardiomyocyte | Antxr11 | 10 | 1.09E-13 | 3.86420641 | 0.5 | 0.156 | 2.17E-10 |
| Cardiomyocyte | Il18r1 | 10 | 2.99E-13 | 2.96886212 | 0.365 | 0.08 | 5.99E-10 |
| Cardiomyocyte | Loxl2 | 10 | 9.45E-13 | 2.41590203 | 0.577 | 0.215 | 1.89E-09 |
| Cardiomyocyte | Kcna71 | 10 | 4.89E-11 | 1.34612363 | 0.423 | 0.112 | 9.78E-08 |
| Cardiomyocyte | Ror2 | 10 | 6.53E-11 | 3.39241941 | 0.558 | 0.16 | 1.31E-07 |
| Cardiomyocyte | Fbn11 | 10 | 6.75E-10 | 2.42261808 | 0.808 | 0.498 | 1.35E-06 |
| Cardiomyocyte | Rassf4 | 10 | 4.14E-09 | 1.35435569 | 0.442 | 0.149 | 8.27E-06 |
| Cardiomyocyte | Sorcs11 | 10 | 5.22E-09 | 2.78833038 | 0.538 | 0.221 | 1.04E-05 |
| Cardiomyocyte | Smc1b1 | 10 | 1.00E-08 | 0.94187733 | 0.423 | 0.144 | 2.00E-05 |
| Cardiomyocyte | Serpine22 | 10 | 1.10E-08 | 2.29637196 | 0.712 | 0.41 | 2.21E-05 |

| | | | | | | | |
|---|---|---|---|---|---|---|---|
| Cardiomyocyte | RGD15633543 | 10 | 5.89E-08 | 3.48427403 | 0.577 | 0.301 | 0.00011774 |
| Cardiomyocyte | Pdgfra | 10 | 8.54E-08 | 3.46553209 | 0.673 | 0.349 | 0.00017073 |
| Cardiomyocyte | Hs6st21 | 10 | 1.04E-07 | 1.73620521 | 0.538 | 0.229 | 0.0002084 |
| Cardiomyocyte | Mafb2 | 10 | 4.20E-07 | 2.56132808 | 0.5 | 0.239 | 0.00084012 |
| Cardiomyocyte | Mbp1 | 10 | 8.43E-07 | 0.64687848 | 0.5 | 0.205 | 0.00168679 |
| Cardiomyocyte | Cpq | 10 | 9.18E-07 | 2.03806017 | 0.635 | 0.361 | 0.001835 |
| Cardiomyocyte | Lrrc17 | 10 | 1.00E-06 | 3.34088189 | 0.577 | 0.306 | 0.00200468 |
| Cardiomyocyte | Gucy1a21 | 10 | 1.01E-06 | 3.15406002 | 0.712 | 0.429 | 0.00202781 |
| Cardiomyocyte | Tspan51 | 10 | 5.06E-06 | 1.97361218 | 0.692 | 0.364 | 0.01011264 |
| Cardiomyocyte | Lrrtm32 | 10 | 5.67E-06 | 0.97164913 | 0.788 | 0.537 | 0.01133171 |
| Cardiomyocyte | Afap1l2 | 10 | 1.88E-05 | 2.38999453 | 0.423 | 0.112 | 0.03761774 |
| Cardiomyocyte | Gda | 10 | 2.15E-05 | 2.78647017 | 0.538 | 0.23 | 0.04305256 |
| Fibroblast | Tp53inp2 | 0 | 1.51E-14 | 0.58339301 | 0.133 | 0.428 | 3.02E-11 |
| Fibroblast | RGD1307916 | 2 | 1.34E-117 | 6.86658531 | 0.296 | 0.001 | 2.67E-114 |
| Fibroblast | Grm3 | 2 | 4.09E-117 | 5.03980545 | 0.382 | 0.017 | 8.19E-114 |
| Fibroblast | Edaradd | 2 | 1.03E-99 | 1.35646633 | 0.276 | 0.003 | 2.06E-96 |

| | | | | | | | |
|---|---|---|---|---|---|---|---|
| Fibroblast | Irf4 | 2 | 3.76E-92 | 4.5314082 | 0.503 | 0.08 | 7.53E-89 |
| Fibroblast | Myl4 | 2 | 2.81E-90 | 3.06328817 | 0.444 | 0.053 | 5.63E-87 |
| Fibroblast | Khdrbs3 | 2 | 3.74E-79 | 1.07926261 | 0.434 | 0.058 | 7.49E-76 |
| Fibroblast | LOC102553338 | 2 | 1.53E-73 | 1.55261846 | 0.434 | 0.043 | 3.07E-70 |
| Fibroblast | AABR07002677.2 | 2 | 3.71E-72 | 2.50676636 | 0.368 | 0.045 | 7.42E-69 |
| Fibroblast | LOC103690241 | 2 | 2.20E-67 | 1.06647062 | 0.5 | 0.12 | 4.40E-64 |
| Fibroblast | Rad51 | 2 | 1.26E-66 | 3.01062672 | 0.592 | 0.189 | 2.52E-63 |
| Fibroblast | Slc38a4 | 2 | 2.06E-65 | 3.00642214 | 0.438 | 0.042 | 4.12E-62 |
| Fibroblast | Sell | 2 | 2.17E-63 | 0.92669191 | 0.309 | 0.033 | 4.34E-60 |
| Fibroblast | Eya1 | 2 | 1.42E-61 | 2.21720366 | 0.533 | 0.099 | 2.83E-58 |
| Fibroblast | Tmem132d | 2 | 1.92E-52 | 2.00249951 | 0.845 | 0.487 | 3.83E-49 |
| Fibroblast | Myo3a | 2 | 3.65E-51 | 1.55662619 | 0.503 | 0.091 | 7.31E-48 |
| Fibroblast | Abca9 | 2 | 9.85E-47 | 1.93124112 | 0.753 | 0.35 | 1.97E-43 |
| Fibroblast | Mme | 2 | 3.25E-46 | 2.8483672 | 0.589 | 0.226 | 6.51E-43 |
| Fibroblast | St6galnac1 | 2 | 7.64E-46 | 4.00466576 | 0.438 | 0.085 | 1.53E-42 |
| Fibroblast | Bard1 | 2 | 3.19E-44 | 2.23632733 | 0.553 | 0.226 | 6.38E-41 |
| Fibroblast | Kcnc2 | 2 | 7.24E-42 | 2.2917173 | 0.684 | 0.33 | 1.45E-38 |
| Fibroblast | Wnt5b | 2 | 1.65E-39 | 0.47430994 | 0.497 | 0.162 | 3.30E-36 |
| Fibroblast | Pde8b | 2 | 1.25E-38 | 2.59271453 | 0.536 | 0.128 | 2.50E-35 |
| Fibroblast | Frem1 | 2 | 4.39E-38 | 1.56418796 | 0.839 | 0.558 | 8.77E-35 |
| Fibroblast | Slco2b1 | 2 | 5.26E-37 | 1.4466956 | 0.806 | 0.506 | 1.05E-33 |
| Fibroblast | Zdbf2 | 2 | 5.47E-37 | 1.61208516 | 0.352 | 0.074 | 1.09E-33 |
| Fibroblast | Fmod | 2 | 1.47E-36 | 1.6870306 | 0.352 | 0.062 | 2.94E-33 |
| Fibroblast | Prss48 | 2 | 1.74E-36 | 0.62892977 | 0.536 | 0.22 | 3.49E-33 |
| Fibroblast | Xirp1 | 2 | 2.49E-36 | 2.18542337 | 0.322 | 0.055 | 4.98E-33 |
| Fibroblast | Sphkap | 2 | 6.54E-36 | 2.38110542 | 0.576 | 0.185 | 1.31E-32 |

| | | | | | | | |
|---|---|---|---|---|---|---|---|
| Fibroblast | Pcdh12 | 2 | 9.28E-36 | 1.01732616 | 0.48 | 0.175 | 1.86E-32 |
| Fibroblast | Cacna1d | 2 | 3.05E-35 | 1.40697775 | 0.576 | 0.215 | 6.10E-32 |
| Fibroblast | Brip1 | 2 | 1.03E-34 | 0.97944196 | 0.618 | 0.315 | 2.06E-31 |
| Fibroblast | Aldh1a2 | 2 | 2.01E-34 | 1.67556164 | 0.638 | 0.258 | 4.02E-31 |
| Fibroblast | Alox5 | 2 | 2.09E-34 | 1.01799556 | 0.362 | 0.096 | 4.19E-31 |
| Fibroblast | Gria4 | 2 | 5.97E-34 | 1.38950909 | 0.711 | 0.397 | 1.19E-30 |
| Fibroblast | Astn2 | 2 | 7.57E-34 | 1.45326826 | 0.707 | 0.402 | 1.51E-30 |
| Fibroblast | Vash2 | 2 | 1.45E-33 | 2.38792241 | 0.477 | 0.09 | 2.91E-30 |
| Fibroblast | Bmper | 2 | 1.90E-33 | 1.30063537 | 0.783 | 0.471 | 3.81E-30 |
| Fibroblast | Ntf3 | 2 | 7.02E-33 | 1.64340579 | 0.684 | 0.346 | 1.40E-29 |
| Fibroblast | Lepr | 2 | 1.78E-32 | 1.57416616 | 0.681 | 0.409 | 3.57E-29 |
| Fibroblast | Gal3st3 | 2 | 3.71E-32 | 1.10010424 | 0.477 | 0.204 | 7.43E-29 |
| Fibroblast | Ly49s6 | 2 | 5.55E-32 | 1.43129474 | 0.408 | 0.129 | 1.11E-28 |
| Fibroblast | Mrvi1 | 2 | 5.83E-32 | 1.55354786 | 0.691 | 0.36 | 1.17E-28 |
| Fibroblast | AABR07007068.1 | 2 | 2.49E-31 | 2.36203726 | 0.632 | 0.342 | 4.98E-28 |
| Fibroblast | Eya2 | 2 | 2.54E-31 | 1.39822599 | 0.799 | 0.53 | 5.09E-28 |
| Fibroblast | Sgip1 | 2 | 6.93E-31 | 1.30733736 | 0.776 | 0.49 | 1.39E-27 |
| Fibroblast | Dusp27 | 2 | 8.33E-31 | 1.71240949 | 0.523 | 0.22 | 1.67E-27 |
| Fibroblast | Nlrp3 | 2 | 2.65E-30 | 0.72147064 | 0.405 | 0.131 | 5.30E-27 |
| Fibroblast | AABR07038986.1 | 2 | 3.68E-30 | 0.96320784 | 0.539 | 0.226 | 7.36E-27 |
| Fibroblast | Dtnb | 2 | 4.83E-30 | 1.19380829 | 0.72 | 0.402 | 9.66E-27 |
| Fibroblast | Ndst3 | 2 | 1.27E-29 | 0.87326618 | 0.648 | 0.377 | 2.54E-26 |
| Fibroblast | Timp4 | 2 | 5.74E-29 | 0.43586222 | 0.5 | 0.171 | 1.15E-25 |
| Fibroblast | Ano4 | 2 | 1.16E-28 | 1.61792674 | 0.559 | 0.282 | 2.32E-25 |
| Fibroblast | Vwa5b1 | 2 | 1.30E-28 | 1.64486879 | 0.359 | 0.068 | 2.61E-25 |
| Fibroblast | Cxcl14 | 2 | 1.71E-28 | 3.56932706 | 0.414 | 0.096 | 3.43E-25 |
| Fibroblast | Oprd1 | 2 | 5.86E-28 | 1.60612012 | 0.576 | 0.322 | 1.17E-24 |

| | | | | | | | |
|---|---|---|---|---|---|---|---|
| Fibroblast | Hs6st2 | 2 | 7.71E-28 | 1.25460186 | 0.684 | 0.337 | 1.54E-24 |
| Fibroblast | AABR07025140.1 | 2 | 1.27E-27 | 0.65304318 | 0.576 | 0.239 | 2.54E-24 |
| Fibroblast | Rec114 | 2 | 2.66E-27 | 1.25407845 | 0.582 | 0.32 | 5.32E-24 |
| Fibroblast | AABR07050449.1 | 2 | 3.46E-27 | 1.38838237 | 0.52 | 0.192 | 6.91E-24 |
| Fibroblast | Asic2 | 2 | 1.12E-26 | 1.75604717 | 0.592 | 0.241 | 2.23E-23 |
| Fibroblast | Shc3 | 2 | 1.99E-26 | 1.79266806 | 0.562 | 0.25 | 3.98E-23 |
| Fibroblast | Atp2b2 | 2 | 5.26E-26 | 0.4610157 | 0.431 | 0.167 | 1.05E-22 |
| Fibroblast | Grid1 | 2 | 1.06E-25 | 1.1067998 | 0.428 | 0.151 | 2.11E-22 |
| Fibroblast | Nostrin | 2 | 2.02E-25 | 0.38910449 | 0.461 | 0.106 | 4.03E-22 |
| Fibroblast | LOC100910978.1 | 2 | 2.65E-25 | 1.05358484 | 0.697 | 0.301 | 5.30E-22 |
| Fibroblast | Gpm6b | 2 | 6.14E-25 | 1.15114445 | 0.691 | 0.38 | 1.23E-21 |
| Fibroblast | Hsf5 | 2 | 6.36E-25 | 0.33320914 | 0.569 | 0.201 | 1.27E-21 |
| Fibroblast | Galnt15 | 2 | 7.98E-24 | 1.03028916 | 0.655 | 0.366 | 1.60E-20 |
| Fibroblast | Ramp1 | 2 | 1.21E-23 | 1.12959525 | 0.53 | 0.275 | 2.42E-20 |
| Fibroblast | AABR07003030.2 | 2 | 2.45E-23 | 1.80186069 | 0.536 | 0.214 | 4.91E-20 |
| Fibroblast | Ror2 | 2 | 6.13E-23 | 0.65566818 | 0.671 | 0.321 | 1.23E-19 |
| Fibroblast | Tafa5 | 2 | 1.42E-22 | 0.52836299 | 0.625 | 0.235 | 2.83E-19 |
| Fibroblast | Trpc3 | 2 | 1.74E-22 | 0.74603156 | 0.582 | 0.246 | 3.48E-19 |
| Fibroblast | Cgnl1 | 2 | 5.91E-22 | 0.32950432 | 0.599 | 0.294 | 1.18E-18 |
| Fibroblast | Trim7 | 2 | 1.10E-21 | 1.1272204 | 0.562 | 0.297 | 2.21E-18 |
| Fibroblast | Flnc | 2 | 1.39E-21 | 0.98159952 | 0.592 | 0.322 | 2.77E-18 |
| Fibroblast | Kif21a | 2 | 1.57E-21 | 0.31073818 | 0.48 | 0.221 | 3.13E-18 |
| Fibroblast | Zmat4 | 2 | 2.05E-21 | 1.57123772 | 0.464 | 0.17 | 4.10E-18 |
| Fibroblast | Cst3 | 2 | 4.65E-21 | 0.8024657 | 0.543 | 0.237 | 9.31E-18 |

| | | | | | | | |
|---|---|---|---|---|---|---|---|
| Fibroblast | AABR07026483.1 | 2 | 1.24E-20 | 1.65757067 | 0.365 | 0.065 | 2.48E-17 |
| Fibroblast | Ankrd6 | 2 | 1.79E-20 | 0.36075018 | 0.586 | 0.289 | 3.59E-17 |
| Fibroblast | Vegfd | 2 | 2.48E-20 | 0.78976614 | 0.641 | 0.304 | 4.95E-17 |
| Fibroblast | Pdcd1lg2 | 2 | 1.56E-19 | 0.93459558 | 0.559 | 0.26 | 3.13E-16 |
| Fibroblast | Gpm6a | 2 | 2.23E-19 | 0.92143249 | 0.809 | 0.542 | 4.46E-16 |
| Fibroblast | Sox13 | 2 | 2.57E-19 | 0.72489151 | 0.605 | 0.253 | 5.14E-16 |
| Fibroblast | Capn6 | 2 | 4.41E-19 | 1.37675873 | 0.444 | 0.147 | 8.82E-16 |
| Fibroblast | Pip5k1b | 2 | 6.01E-19 | 0.3412204 | 0.562 | 0.293 | 1.20E-15 |
| Fibroblast | Ccser1 | 2 | 1.12E-18 | 0.60854361 | 0.562 | 0.265 | 2.24E-15 |
| Fibroblast | Xkr4 | 2 | 1.66E-18 | 0.56459252 | 0.582 | 0.303 | 3.33E-15 |
| Fibroblast | Vegfc | 2 | 3.07E-18 | 0.7430591 | 0.651 | 0.326 | 6.13E-15 |
| Fibroblast | Gbe1 | 2 | 3.67E-18 | 1.0109856 | 0.602 | 0.341 | 7.34E-15 |
| Fibroblast | AABR07001054.2 | 2 | 4.89E-18 | 0.90102334 | 0.431 | 0.143 | 9.77E-15 |
| Fibroblast | Trim54 | 2 | 5.61E-18 | 1.00092986 | 0.559 | 0.305 | 1.12E-14 |
| Fibroblast | Cntfr | 2 | 5.69E-18 | 0.37171255 | 0.497 | 0.239 | 1.14E-14 |
| Fibroblast | Dkk2 | 2 | 6.49E-18 | 0.84473868 | 0.576 | 0.279 | 1.30E-14 |
| Fibroblast | Magi2 | 2 | 1.15E-17 | 0.53796503 | 0.684 | 0.384 | 2.31E-14 |
| Fibroblast | Acsl1 | 2 | 1.69E-17 | 0.36208363 | 0.609 | 0.336 | 3.37E-14 |
| Fibroblast | Casq2 | 2 | 5.04E-16 | 0.43570647 | 0.632 | 0.339 | 1.01E-12 |
| Fibroblast | Naca | 2 | 8.19E-16 | 0.55814641 | 0.507 | 0.249 | 1.64E-12 |
| Fibroblast | Rnf144b | 2 | 2.10E-15 | 0.66826501 | 0.507 | 0.254 | 4.21E-12 |
| Fibroblast | Ddc | 2 | 4.12E-15 | 0.54344059 | 0.454 | 0.178 | 8.23E-12 |
| Fibroblast | Csgalnact1 | 2 | 1.58E-14 | 0.34852553 | 0.648 | 0.37 | 3.16E-11 |
| Fibroblast | Fndc1 | 2 | 1.61E-14 | 0.61214463 | 0.806 | 0.493 | 3.22E-11 |
| Fibroblast | Slit2 | 2 | 6.54E-14 | 0.64501624 | 0.609 | 0.333 | 1.31E-10 |
| Fibroblast | Cox6a2 | 2 | 3.16E-13 | 0.26147081 | 0.589 | 0.321 | 6.32E-10 |
| Fibroblast | Angptl1 | 2 | 3.62E-13 | 0.44309425 | 0.559 | 0.304 | 7.23E-10 |

| | | | | | | | |
|---|---|---|---|---|---|---|---|
| Fibroblast | Il34 | 2 | 3.75E-13 | 0.38261944 | 0.586 | 0.32 | 7.50E-10 |
| Fibroblast | Nnt | 2 | 1.92E-12 | 0.51659756 | 0.612 | 0.362 | 3.84E-09 |
| Fibroblast | Insc | 2 | 2.82E-12 | 0.84332184 | 0.576 | 0.321 | 5.64E-09 |
| Fibroblast | Ppip5k1 | 2 | 2.90E-12 | 0.58511444 | 0.612 | 0.351 | 5.80E-09 |
| Fibroblast | Tbc1d4 | 2 | 3.59E-12 | 0.30807375 | 0.576 | 0.315 | 7.18E-09 |
| Fibroblast | Oxct1 | 2 | 4.77E-11 | 0.36948075 | 0.612 | 0.339 | 9.53E-08 |
| Fibroblast | Rbm44 | 4 | 6.13E-94 | 5.66110757 | 0.37 | 0.022 | 1.23E-90 |
| Fibroblast | Prkcq | 4 | 4.75E-72 | 1.55475451 | 0.457 | 0.049 | 9.51E-69 |
| Fibroblast | Smc1b | 4 | 6.40E-71 | 3.96250358 | 0.409 | 0.042 | 1.28E-67 |
| Fibroblast | Tmem236 | 4 | 2.91E-69 | 4.73572742 | 0.418 | 0.043 | 5.81E-66 |
| Fibroblast | AABR07054716.1 | 4 | 8.17E-68 | 1.55133071 | 0.486 | 0.08 | 1.63E-64 |
| Fibroblast | Cysltr1 | 4 | 1.18E-64 | 2.65635523 | 0.399 | 0.052 | 2.35E-61 |
| Fibroblast | Nrg4 | 4 | 7.48E-62 | 2.55581369 | 0.399 | 0.053 | 1.50E-58 |
| Fibroblast | AABR07057510.3 | 4 | 3.41E-56 | 2.20294597 | 0.462 | 0.092 | 6.82E-53 |
| Fibroblast | AC127756.1 | 4 | 1.10E-55 | 1.49260751 | 0.49 | 0.109 | 2.20E-52 |
| Fibroblast | Kif20a | 4 | 2.64E-46 | 1.37526935 | 0.365 | 0.061 | 5.27E-43 |
| Fibroblast | Nckap1l | 4 | 1.83E-45 | 0.61804449 | 0.37 | 0.063 | 3.66E-42 |
| Fibroblast | Atp6ap1l | 4 | 3.42E-44 | 2.93989754 | 0.365 | 0.02 | 6.84E-41 |
| Fibroblast | Tacr1 | 4 | 1.38E-42 | 1.21803278 | 0.51 | 0.14 | 2.76E-39 |
| Fibroblast | Stac3 | 4 | 9.82E-40 | 3.31210861 | 0.341 | 0.068 | 1.96E-36 |
| Fibroblast | Irf41 | 4 | 8.76E-39 | 0.70483795 | 0.457 | 0.107 | 1.75E-35 |
| Fibroblast | AABR07027925.1 | 4 | 3.93E-34 | 1.31387902 | 0.442 | 0.053 | 7.86E-31 |
| Fibroblast | Col9a1 | 4 | 7.35E-33 | 0.62972681 | 0.322 | 0.067 | 1.47E-29 |
| Fibroblast | LOC1036902411 | 4 | 1.33E-30 | 1.74629472 | 0.486 | 0.142 | 2.66E-27 |

| | | | | | | | |
|---|---|---|---|---|---|---|---|
| Fibroblast | Serpine2 | 4 | 2.28E-30 | 1.62316349 | 0.774 | 0.509 | 4.55E-27 |
| Fibroblast | Slc44a5 | 4 | 2.74E-30 | 2.45660995 | 0.644 | 0.252 | 5.48E-27 |
| Fibroblast | Prkg2 | 4 | 1.36E-28 | 1.56150117 | 0.466 | 0.145 | 2.72E-25 |
| Fibroblast | Kcnn4 | 4 | 2.14E-28 | 1.08838983 | 0.447 | 0.146 | 4.27E-25 |
| Fibroblast | Egr3 | 4 | 4.39E-28 | 2.06506329 | 0.577 | 0.279 | 8.78E-25 |
| Fibroblast | Dync1i1 | 4 | 9.14E-27 | 1.32897635 | 0.505 | 0.191 | 1.83E-23 |
| Fibroblast | Mcm5 | 4 | 1.13E-24 | 1.57751652 | 0.514 | 0.239 | 2.25E-21 |
| Fibroblast | Trmt9b | 4 | 2.22E-24 | 1.64034195 | 0.351 | 0.079 | 4.43E-21 |
| Fibroblast | Sorcs1 | 4 | 1.34E-23 | 1.46197294 | 0.567 | 0.215 | 2.68E-20 |
| Fibroblast | Gfra3 | 4 | 1.88E-23 | 1.04288723 | 0.452 | 0.154 | 3.76E-20 |
| Fibroblast | Ube2ql1 | 4 | 1.98E-23 | 3.87122501 | 0.356 | 0.03 | 3.97E-20 |
| Fibroblast | AC111804.2 | 4 | 2.26E-23 | 0.96753504 | 0.567 | 0.281 | 4.52E-20 |
| Fibroblast | Tex22 | 4 | 6.34E-22 | 1.05901798 | 0.582 | 0.208 | 1.27E-18 |
| Fibroblast | Sema3a | 4 | 3.28E-21 | 1.11475771 | 0.51 | 0.165 | 6.57E-18 |
| Fibroblast | Taldo1 | 4 | 1.39E-20 | 1.16329003 | 0.567 | 0.199 | 2.78E-17 |
| Fibroblast | Opcml | 4 | 1.47E-20 | 1.26939437 | 0.889 | 0.636 | 2.94E-17 |
| Fibroblast | Nrk | 4 | 5.37E-20 | 2.26494832 | 0.548 | 0.185 | 1.07E-16 |
| Fibroblast | Adamts14 | 4 | 1.19E-19 | 1.54330894 | 0.635 | 0.348 | 2.37E-16 |
| Fibroblast | Tnc | 4 | 2.01E-19 | 1.49350235 | 0.476 | 0.154 | 4.02E-16 |
| Fibroblast | LOC682419 | 4 | 2.92E-19 | 0.89071263 | 0.418 | 0.067 | 5.84E-16 |
| Fibroblast | Sgpp2 | 4 | 4.02E-19 | 0.70352544 | 0.452 | 0.189 | 8.04E-16 |
| Fibroblast | Prrg4 | 4 | 7.45E-19 | 1.66060312 | 0.582 | 0.295 | 1.49E-15 |
| Fibroblast | Klhl4 | 4 | 2.15E-18 | 0.87408437 | 0.577 | 0.323 | 4.30E-15 |
| Fibroblast | Rnf150 | 4 | 3.01E-18 | 1.36032369 | 0.692 | 0.363 | 6.01E-15 |
| Fibroblast | Cdkn1a | 4 | 5.36E-18 | 1.23155506 | 0.514 | 0.239 | 1.07E-14 |
| Fibroblast | Chst15 | 4 | 5.38E-18 | 0.38936158 | 0.471 | 0.19 | 1.08E-14 |
| Fibroblast | Adhfe1 | 4 | 5.72E-18 | 0.26915177 | 0.606 | 0.32 | 1.14E-14 |
| Fibroblast | Ano41 | 4 | 8.35E-18 | 1.61182402 | 0.611 | 0.291 | 1.67E-14 |

| | | | | | | | |
|---|---|---|---|---|---|---|---|
| Fibroblast | Dgkg | 4 | 1.29E-17 | 0.76774431 | 0.438 | 0.147 | 2.57E-14 |
| Fibroblast | Adamts17 | 4 | 1.51E-17 | 1.17311361 | 0.726 | 0.453 | 3.01E-14 |
| Fibroblast | Kif26b | 4 | 2.07E-17 | 0.7640217 | 0.548 | 0.226 | 4.14E-14 |
| Fibroblast | Tbc1d9 | 4 | 3.07E-17 | 1.10297631 | 0.534 | 0.241 | 6.13E-14 |
| Fibroblast | Unc5b | 4 | 3.77E-17 | 1.24492177 | 0.76 | 0.485 | 7.54E-14 |
| Fibroblast | AABR07001054.21 | 4 | 9.83E-17 | 0.75907278 | 0.524 | 0.148 | 1.97E-13 |
| Fibroblast | Sncaip | 4 | 1.26E-16 | 1.22603235 | 0.553 | 0.261 | 2.52E-13 |
| Fibroblast | Trim50 | 4 | 2.05E-16 | 0.44651718 | 0.389 | 0.034 | 4.10E-13 |
| Fibroblast | Cmahp | 4 | 2.15E-16 | 0.44463713 | 0.466 | 0.173 | 4.30E-13 |
| Fibroblast | Rem1 | 4 | 3.16E-16 | 1.19450529 | 0.697 | 0.423 | 6.32E-13 |
| Fibroblast | AABR07017268.1 | 4 | 3.74E-16 | 1.22555184 | 0.466 | 0.21 | 7.47E-13 |
| Fibroblast | Ablim2 | 4 | 4.06E-16 | 0.64848038 | 0.466 | 0.193 | 8.13E-13 |
| Fibroblast | Pamr1 | 4 | 6.45E-16 | 1.67986476 | 0.572 | 0.246 | 1.29E-12 |
| Fibroblast | Maoa | 4 | 7.40E-16 | 0.80147086 | 0.635 | 0.336 | 1.48E-12 |
| Fibroblast | Mbp | 4 | 1.26E-15 | 0.4009116 | 0.606 | 0.301 | 2.51E-12 |
| Fibroblast | Sfxn5 | 4 | 1.29E-15 | 0.32953931 | 0.548 | 0.243 | 2.58E-12 |
| Fibroblast | Prkaa2 | 4 | 1.52E-15 | 1.09030889 | 0.519 | 0.173 | 3.04E-12 |
| Fibroblast | Lyve1 | 4 | 1.70E-15 | 2.2333661 | 0.543 | 0.218 | 3.39E-12 |
| Fibroblast | Rp1 | 4 | 2.53E-15 | 0.93699197 | 0.462 | 0.171 | 5.07E-12 |
| Fibroblast | Tex14 | 4 | 6.18E-15 | 1.11561613 | 0.587 | 0.325 | 1.24E-11 |
| Fibroblast | Coq8a | 4 | 6.21E-15 | 0.64070107 | 0.505 | 0.23 | 1.24E-11 |
| Fibroblast | Gbp6 | 4 | 9.78E-15 | 1.34386979 | 0.514 | 0.23 | 1.96E-11 |
| Fibroblast | Alpk2 | 4 | 1.60E-14 | 0.34846157 | 0.514 | 0.222 | 3.20E-11 |
| Fibroblast | Ntrk2 | 4 | 1.85E-14 | 0.65460553 | 0.558 | 0.249 | 3.71E-11 |
| Fibroblast | Meox1 | 4 | 2.49E-14 | 0.55224062 | 0.481 | 0.21 | 4.99E-11 |
| Fibroblast | Cxcl1 | 4 | 2.64E-14 | 0.82840662 | 0.587 | 0.322 | 5.28E-11 |
| Fibroblast | Rnf157 | 4 | 3.01E-14 | 1.06259012 | 0.587 | 0.297 | 6.01E-11 |

| | | | | | | | |
|---|---|---|---|---|---|---|---|
| Fibroblast | Atf3 | 4 | 3.11E-14 | 1.33739859 | 0.341 | 0.085 | 6.22E-11 |
| Fibroblast | Veph1 | 4 | 4.45E-14 | 1.66067529 | 0.447 | 0.173 | 8.91E-11 |
| Fibroblast | Gask1b | 4 | 5.99E-14 | 0.91689995 | 0.779 | 0.52 | 1.20E-10 |
| Fibroblast | Capn61 | 4 | 1.81E-13 | 1.03692243 | 0.428 | 0.165 | 3.62E-10 |
| Fibroblast | Ednra | 4 | 2.19E-13 | 0.8566757 | 0.63 | 0.366 | 4.38E-10 |
| Fibroblast | Reep1 | 4 | 2.28E-13 | 0.66540728 | 0.514 | 0.222 | 4.56E-10 |
| Fibroblast | Ppm1l | 4 | 3.43E-13 | 0.26707257 | 0.51 | 0.193 | 6.87E-10 |
| Fibroblast | Pgf | 4 | 4.66E-13 | 1.54325848 | 0.337 | 0.057 | 9.32E-10 |
| Fibroblast | Tmem26 | 4 | 7.21E-13 | 0.53886726 | 0.548 | 0.25 | 1.44E-09 |
| Fibroblast | Lgr6 | 4 | 1.35E-12 | 0.93288192 | 0.582 | 0.291 | 2.71E-09 |
| Fibroblast | Popdc2 | 4 | 1.88E-12 | 0.27253013 | 0.558 | 0.227 | 3.75E-09 |
| Fibroblast | Fsd2 | 4 | 5.24E-12 | 0.68603127 | 0.524 | 0.243 | 1.05E-08 |
| Fibroblast | Bmp2 | 4 | 7.94E-12 | 2.00711205 | 0.385 | 0.09 | 1.59E-08 |
| Fibroblast | Rhobtb1 | 4 | 8.41E-12 | 0.65287122 | 0.587 | 0.303 | 1.68E-08 |
| Fibroblast | Gucy1a1 | 4 | 1.47E-11 | 0.93241535 | 0.688 | 0.435 | 2.95E-08 |
| Fibroblast | Fhl2 | 4 | 1.75E-11 | 0.33866191 | 0.63 | 0.322 | 3.51E-08 |
| Fibroblast | Cacna1a | 4 | 2.07E-11 | 0.697978 | 0.692 | 0.43 | 4.15E-08 |
| Fibroblast | Brca1 | 4 | 5.25E-11 | 0.65486404 | 0.567 | 0.292 | 1.05E-07 |
| Fibroblast | Aqp1 | 4 | 8.43E-11 | 0.46604228 | 0.519 | 0.245 | 1.69E-07 |
| Fibroblast | Shroom4 | 4 | 9.88E-11 | 0.41451972 | 0.649 | 0.361 | 1.98E-07 |
| Fibroblast | Iqub | 4 | 1.48E-10 | 0.51857032 | 0.486 | 0.21 | 2.97E-07 |
| Fibroblast | Sh3kbp1 | 4 | 3.46E-10 | 0.6041758 | 0.644 | 0.367 | 6.93E-07 |
| Fibroblast | Actn1 | 4 | 7.90E-10 | 0.33770755 | 0.668 | 0.412 | 1.58E-06 |
| Fibroblast | Tenm3 | 4 | 1.50E-09 | 0.6920596 | 0.663 | 0.41 | 3.00E-06 |
| Fibroblast | Flt1 | 4 | 2.40E-09 | 0.273697 | 0.582 | 0.319 | 4.81E-06 |
| Fibroblast | Rasgef1c | 4 | 5.28E-09 | 0.49642683 | 0.389 | 0.133 | 1.06E-05 |
| Fibroblast | AABR07001519.1 | 4 | 1.70E-08 | 0.61258777 | 0.702 | 0.446 | 3.40E-05 |
| Fibroblast | Ddc1 | 4 | 1.84E-08 | 0.56351937 | 0.447 | 0.193 | 3.67E-05 |

| | | | | | | | |
|---|---|---|---|---|---|---|---|
| Fibroblast | Slc20a2 | 4 | 2.54E-08 | 0.35265407 | 0.587 | 0.298 | 5.09E-05 |
| Fibroblast | Adk | 4 | 9.00E-08 | 0.37591141 | 0.678 | 0.405 | 0.00018003 |
| Fibroblast | Rbm20 | 4 | 5.26E-06 | 0.61746467 | 0.683 | 0.41 | 0.01052087 |
| Fibroblast | Vstm5 | 5 | 1.42E-199 | 4.64494248 | 0.51 | 0.004 | 2.85E-196 |
| Fibroblast | Clspn | 5 | 1.42E-141 | 4.6356655 | 0.495 | 0.025 | 2.84E-138 |
| Fibroblast | Adra2b | 5 | 1.86E-133 | 6.3382212 | 0.47 | 0.021 | 3.71E-130 |
| Fibroblast | Reg4 | 5 | 2.85E-119 | 6.3559219 | 0.3 | 0.002 | 5.71E-116 |
| Fibroblast | LOC691995 | 5 | 1.38E-116 | 1.90235479 | 0.47 | 0.029 | 2.77E-113 |
| Fibroblast | Gria2 | 5 | 3.68E-116 | 3.87123869 | 0.435 | 0.037 | 7.36E-113 |
| Fibroblast | Slc22a2 | 5 | 2.83E-113 | 7.40052715 | 0.31 | 0.004 | 5.65E-110 |
| Fibroblast | AABR07065282.1 | 5 | 5.64E-112 | 6.33130215 | 0.43 | 0.024 | 1.13E-108 |
| Fibroblast | Galnt14 | 5 | 1.53E-110 | 1.73811614 | 0.55 | 0.052 | 3.06E-107 |
| Fibroblast | AABR07032787.1 | 5 | 3.57E-105 | 3.96424542 | 0.34 | 0.011 | 7.13E-102 |
| Fibroblast | Zc3h12d | 5 | 7.13E-101 | 4.0588147 | 0.54 | 0.051 | 1.43E-97 |
| Fibroblast | Abo | 5 | 9.66E-101 | 1.55572161 | 0.525 | 0.025 | 1.93E-97 |
| Fibroblast | Rasgrf1 | 5 | 2.45E-99 | 1.98671723 | 0.55 | 0.054 | 4.91E-96 |
| Fibroblast | AC118957.1 | 5 | 9.17E-98 | 0.65000228 | 0.49 | 0.025 | 1.83E-94 |
| Fibroblast | Cpa6 | 5 | 3.15E-96 | 2.79640567 | 0.565 | 0.078 | 6.31E-93 |
| Fibroblast | AC128789.1 | 5 | 1.75E-95 | 5.71698006 | 0.385 | 0.023 | 3.49E-92 |
| Fibroblast | Csf3r | 5 | 2.25E-94 | 3.17804603 | 0.385 | 0.024 | 4.50E-91 |
| Fibroblast | Kcp | 5 | 7.45E-79 | 3.59307166 | 0.31 | 0.007 | 1.49E-75 |
| Fibroblast | Stk31 | 5 | 1.16E-74 | 0.76655796 | 0.47 | 0.062 | 2.33E-71 |
| Fibroblast | Ikzf3 | 5 | 6.50E-61 | 1.88515737 | 0.575 | 0.069 | 1.30E-57 |
| Fibroblast | Msr1 | 5 | 1.34E-57 | 2.03521471 | 0.295 | 0.017 | 2.68E-54 |
| Fibroblast | Epha3 | 5 | 2.76E-57 | 3.52057947 | 0.63 | 0.142 | 5.53E-54 |

| | | | | | | | |
|---|---|---|---|---|---|---|---|
| Fibroblast | Ceacam16 | 5 | 8.79E-56 | 5.17638454 | 0.34 | 0.011 | 1.76E-52 |
| Fibroblast | Cdh6 | 5 | 4.53E-52 | 1.99902682 | 0.38 | 0.018 | 9.07E-49 |
| Fibroblast | Lilrb4 | 5 | 3.23E-51 | 0.51159615 | 0.515 | 0.027 | 6.45E-48 |
| Fibroblast | Cacna1b | 5 | 6.02E-50 | 2.12612601 | 0.64 | 0.214 | 1.20E-46 |
| Fibroblast | Myo5b | 5 | 1.18E-49 | 1.75532746 | 0.48 | 0.09 | 2.36E-46 |
| Fibroblast | Hapln3 | 5 | 5.22E-48 | 3.85815839 | 0.41 | 0.015 | 1.04E-44 |
| Fibroblast | Thsd7b | 5 | 2.34E-46 | 2.64760767 | 0.315 | 0.063 | 4.68E-43 |
| Fibroblast | Ccn5 | 5 | 1.94E-45 | 3.86872707 | 0.355 | 0.088 | 3.88E-42 |
| Fibroblast | Arnt2 | 5 | 3.40E-44 | 4.1709409 | 0.385 | 0.057 | 6.80E-41 |
| Fibroblast | S100a4 | 5 | 4.30E-44 | 1.61870817 | 0.59 | 0.216 | 8.60E-41 |
| Fibroblast | St14 | 5 | 3.54E-42 | 1.63810994 | 0.425 | 0.056 | 7.09E-39 |
| Fibroblast | Fabp12 | 5 | 3.70E-42 | 2.62588 | 0.4 | 0.028 | 7.40E-39 |
| Fibroblast | Tnfrsf11b | 5 | 5.29E-42 | 0.93483607 | 0.585 | 0.101 | 1.06E-38 |
| Fibroblast | Mctp2 | 5 | 1.33E-41 | 1.92040312 | 0.465 | 0.109 | 2.66E-38 |
| Fibroblast | LOC100911486 | 5 | 1.49E-41 | 3.34883964 | 0.375 | 0.122 | 2.99E-38 |
| Fibroblast | Ky | 5 | 2.74E-39 | 2.72031854 | 0.55 | 0.119 | 5.49E-36 |
| Fibroblast | Tshr | 5 | 4.45E-39 | 3.63609926 | 0.375 | 0.021 | 8.90E-36 |
| Fibroblast | Col6a5 | 5 | 2.54E-38 | 0.82265293 | 0.365 | 0.1 | 5.08E-35 |
| Fibroblast | LOC103693323 | 5 | 4.50E-38 | 2.13430383 | 0.435 | 0.131 | 8.99E-35 |
| Fibroblast | Rtn4rl1 | 5 | 4.63E-37 | 1.07113206 | 0.65 | 0.239 | 9.26E-34 |
| Fibroblast | Frmd3 | 5 | 1.36E-36 | 1.76447822 | 0.515 | 0.104 | 2.73E-33 |
| Fibroblast | Unc5c | 5 | 1.58E-36 | 1.18028857 | 0.41 | 0.085 | 3.17E-33 |
| Fibroblast | Gvin1 | 5 | 1.35E-35 | 1.64164388 | 0.545 | 0.111 | 2.71E-32 |
| Fibroblast | Itgb2 | 5 | 3.57E-35 | 1.61915158 | 0.48 | 0.128 | 7.13E-32 |
| Fibroblast | Cnksr2 | 5 | 8.17E-34 | 0.62324301 | 0.545 | 0.174 | 1.63E-30 |
| Fibroblast | Ms4a6bl | 5 | 1.38E-33 | 2.12567866 | 0.4 | 0.04 | 2.77E-30 |
| Fibroblast | Mcm6 | 5 | 1.96E-33 | 1.41229677 | 0.31 | 0.037 | 3.92E-30 |

| | | | | | | | |
|---|---|---|---|---|---|---|---|
| Fibroblast | Rbp7 | 5 | 1.99E-33 | 3.23476065 | 0.38 | 0.039 | 3.98E-30 |
| Fibroblast | AABR07031740.1 | 5 | 3.14E-33 | 1.65327113 | 0.455 | 0.065 | 6.27E-30 |
| Fibroblast | Plbd1 | 5 | 1.25E-32 | 2.11066822 | 0.375 | 0.109 | 2.49E-29 |
| Fibroblast | Mt1 | 5 | 3.78E-32 | 1.70332255 | 0.415 | 0.062 | 7.55E-29 |
| Fibroblast | Icam1 | 5 | 4.24E-32 | 0.89010637 | 0.61 | 0.219 | 8.49E-29 |
| Fibroblast | Adamts15 | 5 | 1.65E-31 | 2.52822917 | 0.605 | 0.261 | 3.31E-28 |
| Fibroblast | Kntc1 | 5 | 1.95E-31 | 0.30087302 | 0.535 | 0.155 | 3.91E-28 |
| Fibroblast | Mx1 | 5 | 5.88E-31 | 2.32092507 | 0.395 | 0.109 | 1.18E-27 |
| Fibroblast | Plcxd3 | 5 | 1.60E-30 | 3.43182945 | 0.56 | 0.26 | 3.20E-27 |
| Fibroblast | Nkd1 | 5 | 5.08E-29 | 2.80299174 | 0.655 | 0.309 | 1.02E-25 |
| Fibroblast | Tmem17 | 5 | 5.18E-29 | 3.05273448 | 0.645 | 0.27 | 1.04E-25 |
| Fibroblast | Gpnmb | 5 | 1.68E-28 | 2.03209532 | 0.45 | 0.15 | 3.35E-25 |
| Fibroblast | Asb15 | 5 | 2.54E-28 | 1.55421793 | 0.425 | 0.08 | 5.07E-25 |
| Fibroblast | Rcan2 | 5 | 4.71E-28 | 1.42097991 | 0.81 | 0.481 | 9.43E-25 |
| Fibroblast | Prodh1 | 5 | 1.42E-27 | 0.97742558 | 0.675 | 0.337 | 2.84E-24 |
| Fibroblast | Sod2 | 5 | 1.80E-27 | 0.61833328 | 0.635 | 0.342 | 3.60E-24 |
| Fibroblast | Cyp26b1 | 5 | 2.51E-27 | 0.81806228 | 0.445 | 0.131 | 5.01E-24 |
| Fibroblast | Nkain3 | 5 | 5.35E-27 | 0.35665376 | 0.54 | 0.174 | 1.07E-23 |
| Fibroblast | Apoo | 5 | 5.63E-27 | 1.08418472 | 0.65 | 0.198 | 1.13E-23 |
| Fibroblast | Gabrb1 | 5 | 3.03E-26 | 2.80036045 | 0.325 | 0.075 | 6.06E-23 |
| Fibroblast | Jag2 | 5 | 3.09E-26 | 0.76318606 | 0.55 | 0.258 | 6.19E-23 |
| Fibroblast | Grin3a | 5 | 3.27E-26 | 1.72119375 | 0.465 | 0.182 | 6.54E-23 |
| Fibroblast | Tec | 5 | 3.73E-26 | 0.8350214 | 0.455 | 0.131 | 7.46E-23 |
| Fibroblast | Rbm24 | 5 | 4.16E-26 | 0.29168273 | 0.43 | 0.09 | 8.32E-23 |
| Fibroblast | Cav1 | 5 | 7.74E-26 | 1.07402032 | 0.695 | 0.363 | 1.55E-22 |
| Fibroblast | Lmod1 | 5 | 1.14E-25 | 2.2393188 | 0.63 | 0.321 | 2.28E-22 |
| Fibroblast | Ptpro | 5 | 1.89E-25 | 0.82619188 | 0.575 | 0.281 | 3.78E-22 |
| Fibroblast | Scube3 | 5 | 1.07E-24 | 0.77527605 | 0.565 | 0.248 | 2.13E-21 |

| | | | | | | | |
|---|---|---|---|---|---|---|---|
| Fibroblast | Mctp1 | 5 | 1.58E-24 | 1.0610763 | 0.675 | 0.31 | 3.17E-21 |
| Fibroblast | Efna5 | 5 | 1.84E-24 | 0.25866838 | 0.645 | 0.308 | 3.68E-21 |
| Fibroblast | AABR07003304.2 | 5 | 2.11E-24 | 1.19252001 | 0.58 | 0.308 | 4.23E-21 |
| Fibroblast | Cabcoco1 | 5 | 5.85E-24 | 1.77018397 | 0.485 | 0.092 | 1.17E-20 |
| Fibroblast | Ppp2r2b | 5 | 6.23E-24 | 0.92435065 | 0.705 | 0.328 | 1.25E-20 |
| Fibroblast | P2rx7 | 5 | 6.35E-24 | 1.01328005 | 0.6 | 0.228 | 1.27E-20 |
| Fibroblast | Hs3st1 | 5 | 2.67E-23 | 0.83879429 | 0.655 | 0.3 | 5.34E-20 |
| Fibroblast | Cdh19 | 5 | 7.01E-23 | 1.27005135 | 0.51 | 0.136 | 1.40E-19 |
| Fibroblast | Tenm4 | 5 | 9.02E-23 | 1.42760207 | 0.47 | 0.187 | 1.80E-19 |
| Fibroblast | Slc16a10 | 5 | 9.37E-23 | 2.04258496 | 0.47 | 0.173 | 1.87E-19 |
| Fibroblast | Mertk | 5 | 1.54E-22 | 1.60103921 | 0.465 | 0.144 | 3.08E-19 |
| Fibroblast | Adamtsl2 | 5 | 1.84E-22 | 1.53194701 | 0.755 | 0.466 | 3.68E-19 |
| Fibroblast | Gfra31 | 5 | 3.54E-22 | 0.34700782 | 0.42 | 0.159 | 7.08E-19 |
| Fibroblast | Slc38a3 | 5 | 8.04E-22 | 1.37934849 | 0.39 | 0.137 | 1.61E-18 |
| Fibroblast | Akr1c15 | 5 | 8.53E-22 | 1.89369432 | 0.58 | 0.238 | 1.71E-18 |
| Fibroblast | Abcc8 | 5 | 8.73E-22 | 1.99520501 | 0.45 | 0.114 | 1.75E-18 |
| Fibroblast | Pfkfb3 | 5 | 1.04E-21 | 0.96407236 | 0.645 | 0.299 | 2.07E-18 |
| Fibroblast | Slc9a3r2 | 5 | 2.52E-21 | 0.37564192 | 0.665 | 0.266 | 5.03E-18 |
| Fibroblast | Dlg2 | 5 | 2.90E-21 | 1.36951541 | 0.815 | 0.563 | 5.80E-18 |
| Fibroblast | Ifitm10 | 5 | 3.21E-21 | 1.08371372 | 0.58 | 0.328 | 6.43E-18 |
| Fibroblast | Coro6 | 5 | 3.69E-21 | 0.96356956 | 0.56 | 0.152 | 7.37E-18 |
| Fibroblast | Ano5 | 5 | 4.21E-21 | 1.03283654 | 0.49 | 0.167 | 8.43E-18 |
| Fibroblast | Lamc3 | 5 | 1.08E-20 | 3.08771607 | 0.43 | 0.173 | 2.15E-17 |
| Fibroblast | Gja1 | 5 | 1.56E-20 | 1.10235644 | 0.57 | 0.214 | 3.12E-17 |
| Fibroblast | Hmgcll1 | 5 | 2.34E-20 | 0.634398 | 0.635 | 0.342 | 4.68E-17 |
| Fibroblast | Ablim21 | 5 | 2.70E-20 | 1.03886454 | 0.57 | 0.183 | 5.40E-17 |
| Fibroblast | Dmpk | 5 | 2.89E-20 | 1.03756876 | 0.66 | 0.284 | 5.78E-17 |
| Fibroblast | Nav2 | 5 | 3.25E-20 | 0.80284448 | 0.625 | 0.262 | 6.49E-17 |

| | | | | | | | |
|---|---|---|---|---|---|---|---|
| Fibroblast | Acyp2 | 5 | 3.99E-20 | 0.89389244 | 0.74 | 0.417 | 7.98E-17 |
| Fibroblast | Rcan1 | 5 | 4.65E-20 | 1.20346042 | 0.49 | 0.221 | 9.30E-17 |
| Fibroblast | Srl | 5 | 5.43E-20 | 0.58541088 | 0.395 | 0.103 | 1.09E-16 |
| Fibroblast | Frmd5 | 5 | 1.15E-19 | 0.4518096 | 0.525 | 0.188 | 2.29E-16 |
| Fibroblast | Gjc1 | 5 | 1.16E-19 | 0.45323375 | 0.63 | 0.307 | 2.33E-16 |
| Fibroblast | Olfml2b | 5 | 1.19E-19 | 1.21006179 | 0.515 | 0.245 | 2.39E-16 |
| Fibroblast | Sgcg | 5 | 1.24E-19 | 0.78278481 | 0.625 | 0.332 | 2.48E-16 |
| Fibroblast | Efnb2 | 5 | 2.85E-19 | 0.9625042 | 0.615 | 0.338 | 5.70E-16 |
| Fibroblast | Rnf207 | 5 | 2.92E-19 | 0.96434683 | 0.525 | 0.207 | 5.84E-16 |
| Fibroblast | Pola2 | 5 | 2.94E-19 | 0.50406055 | 0.535 | 0.214 | 5.88E-16 |
| Fibroblast | Zfp366 | 5 | 2.98E-19 | 0.70341111 | 0.61 | 0.311 | 5.96E-16 |
| Fibroblast | Lrsam1 | 5 | 3.86E-19 | 1.16808928 | 0.56 | 0.259 | 7.72E-16 |
| Fibroblast | Alox5ap | 5 | 1.66E-18 | 0.44004368 | 0.425 | 0.175 | 3.32E-15 |
| Fibroblast | Plcb1 | 5 | 2.90E-18 | 0.98393787 | 0.56 | 0.256 | 5.80E-15 |
| Fibroblast | Nrp2 | 5 | 3.03E-18 | 1.72507244 | 0.535 | 0.252 | 6.05E-15 |
| Fibroblast | Slc39a8 | 5 | 3.73E-18 | 0.72780475 | 0.55 | 0.298 | 7.47E-15 |
| Fibroblast | Stk32b | 5 | 5.01E-18 | 0.53434217 | 0.665 | 0.357 | 1.00E-14 |
| Fibroblast | Tmem168 | 5 | 7.37E-18 | 0.25416904 | 0.52 | 0.258 | 1.47E-14 |
| Fibroblast | Tox3 | 5 | 9.55E-18 | 1.00097942 | 0.51 | 0.255 | 1.91E-14 |
| Fibroblast | AABR07040864.1 | 5 | 1.26E-17 | 0.85197094 | 0.675 | 0.289 | 2.52E-14 |
| Fibroblast | RGD1564053 | 5 | 1.71E-17 | 1.84235788 | 0.405 | 0.063 | 3.43E-14 |
| Fibroblast | Pappa1 | 5 | 3.22E-17 | 0.46749305 | 0.67 | 0.366 | 6.44E-14 |
| Fibroblast | Rnf213 | 5 | 9.41E-17 | 1.05182994 | 0.715 | 0.368 | 1.88E-13 |
| Fibroblast | Rasa4 | 5 | 1.43E-16 | 0.77585532 | 0.55 | 0.29 | 2.87E-13 |
| Fibroblast | Fam151a | 5 | 1.54E-16 | 0.80040352 | 0.405 | 0.148 | 3.08E-13 |
| Fibroblast | Trdn | 5 | 1.67E-16 | 0.81710652 | 0.58 | 0.265 | 3.33E-13 |
| Fibroblast | Arhgap44 | 5 | 2.51E-16 | 1.2770351 | 0.515 | 0.205 | 5.02E-13 |

| | | | | | | | |
|---|---|---|---|---|---|---|---|
| Fibroblast | Slc25a21 | 5 | 2.99E-16 | 1.01812739 | 0.62 | 0.366 | 5.98E-13 |
| Fibroblast | Apoe | 5 | 3.56E-16 | 0.94895659 | 0.56 | 0.278 | 7.11E-13 |
| Fibroblast | Thsd7a | 5 | 3.61E-16 | 0.76169475 | 0.565 | 0.245 | 7.21E-13 |
| Fibroblast | Pfkfb2 | 5 | 4.87E-16 | 0.4216839 | 0.595 | 0.295 | 9.75E-13 |
| Fibroblast | Fgf10 | 5 | 6.30E-16 | 0.68135562 | 0.66 | 0.361 | 1.26E-12 |
| Fibroblast | Scn5a | 5 | 6.78E-16 | 0.57447432 | 0.63 | 0.38 | 1.36E-12 |
| Fibroblast | Spns2 | 5 | 7.14E-16 | 0.36990924 | 0.535 | 0.261 | 1.43E-12 |
| Fibroblast | Twf2 | 5 | 1.36E-15 | 0.29036309 | 0.64 | 0.365 | 2.72E-12 |
| Fibroblast | Itgb8 | 5 | 1.55E-15 | 1.2764086 | 0.62 | 0.344 | 3.11E-12 |
| Fibroblast | Gpc3 | 5 | 2.16E-15 | 0.60470307 | 0.48 | 0.207 | 4.33E-12 |
| Fibroblast | Flnb | 5 | 2.65E-15 | 0.75852569 | 0.68 | 0.341 | 5.29E-12 |
| Fibroblast | Iqub1 | 5 | 8.47E-15 | 0.36268797 | 0.47 | 0.213 | 1.69E-11 |
| Fibroblast | Kdr | 5 | 1.36E-14 | 0.4471277 | 0.58 | 0.308 | 2.73E-11 |
| Fibroblast | Mybpc3 | 5 | 1.52E-14 | 0.70954037 | 0.695 | 0.406 | 3.05E-11 |
| Fibroblast | Acacb | 5 | 1.55E-14 | 0.89874437 | 0.595 | 0.295 | 3.10E-11 |
| Fibroblast | Ptprn2 | 5 | 6.90E-14 | 1.01064087 | 0.53 | 0.255 | 1.38E-10 |
| Fibroblast | Hlf | 5 | 9.39E-14 | 0.8269931 | 0.695 | 0.443 | 1.88E-10 |
| Fibroblast | Lrrc4b | 5 | 1.26E-13 | 0.66842078 | 0.52 | 0.245 | 2.52E-10 |
| Fibroblast | Greb1l | 5 | 1.51E-13 | 0.51319255 | 0.5 | 0.235 | 3.01E-10 |
| Fibroblast | Ntrk21 | 5 | 1.81E-13 | 0.31464886 | 0.555 | 0.251 | 3.63E-10 |
| Fibroblast | Amd1 | 5 | 3.41E-13 | 0.56908429 | 0.57 | 0.318 | 6.82E-10 |
| Fibroblast | Naca1 | 5 | 7.57E-13 | 0.30331566 | 0.52 | 0.262 | 1.51E-09 |
| Fibroblast | Ccn1 | 5 | 2.50E-12 | 0.65499918 | 0.585 | 0.317 | 5.00E-09 |
| Fibroblast | Cnnm2 | 5 | 2.53E-12 | 0.62465955 | 0.59 | 0.294 | 5.05E-09 |
| Fibroblast | Fgf1 | 5 | 2.54E-12 | 0.44763034 | 0.625 | 0.301 | 5.07E-09 |
| Fibroblast | Aox1 | 5 | 4.61E-12 | 0.66357615 | 0.83 | 0.501 | 9.22E-09 |
| Fibroblast | Ldb3 | 5 | 5.14E-12 | 0.93122299 | 0.58 | 0.316 | 1.03E-08 |
| Fibroblast | Rmdn1 | 5 | 9.58E-12 | 0.61831171 | 0.65 | 0.375 | 1.92E-08 |
| Fibroblast | Kalrn | 5 | 1.95E-11 | 0.73237616 | 0.675 | 0.4 | 3.90E-08 |

| | | | | | | | |
|---|---|---|---|---|---|---|---|
| Fibroblast | AABR07054000.1 | 5 | 3.06E-11 | 0.45413067 | 0.52 | 0.25 | 6.13E-08 |
| Fibroblast | Antxr1 | 5 | 1.24E-10 | 0.65863694 | 0.77 | 0.511 | 2.48E-07 |
| Fibroblast | Tbc1d1 | 5 | 2.32E-10 | 0.49712416 | 0.73 | 0.474 | 4.64E-07 |
| Fibroblast | Baiap2l2 | 6 | 2.34E-276 | 7.55088084 | 0.689 | 0.004 | 4.68E-273 |
| Fibroblast | Cd27 | 6 | 5.86E-256 | 7.34860102 | 0.689 | 0.007 | 1.17E-252 |
| Fibroblast | Egfl6 | 6 | 5.28E-245 | 3.30561603 | 0.66 | 0.006 | 1.06E-241 |
| Fibroblast | Ccn6 | 6 | 1.63E-241 | 3.52670726 | 0.66 | 0.007 | 3.26E-238 |
| Fibroblast | Scart1 | 6 | 2.45E-235 | 6.83242561 | 0.65 | 0.007 | 4.89E-232 |
| Fibroblast | Tcerg1l | 6 | 3.77E-171 | 4.5296512 | 0.515 | 0.008 | 7.53E-168 |
| Fibroblast | Dsc3 | 6 | 6.31E-150 | 3.84233235 | 0.68 | 0.035 | 1.26E-146 |
| Fibroblast | Gata3 | 6 | 5.68E-140 | 5.56380226 | 0.699 | 0.04 | 1.14E-136 |
| Fibroblast | Tmem156 | 6 | 2.88E-118 | 4.17351391 | 0.553 | 0.005 | 5.76E-115 |
| Fibroblast | Edil3 | 6 | 5.87E-114 | 3.07347858 | 0.68 | 0.008 | 1.17E-110 |
| Fibroblast | Kif11 | 6 | 1.52E-92 | 3.85165707 | 0.68 | 0.055 | 3.04E-89 |
| Fibroblast | Procr | 6 | 3.79E-75 | 2.25488114 | 0.641 | 0.02 | 7.57E-72 |
| Fibroblast | Adamts20 | 6 | 6.99E-69 | 2.46585543 | 0.699 | 0.044 | 1.40E-65 |
| Fibroblast | Tk1 | 6 | 9.89E-66 | 6.48081554 | 0.66 | 0.025 | 1.98E-62 |
| Fibroblast | Hs3st3b1 | 6 | 7.17E-61 | 3.14056867 | 0.709 | 0.132 | 1.43E-57 |
| Fibroblast | Pkmyt1 | 6 | 3.37E-56 | 2.24459429 | 0.272 | 0.007 | 6.74E-53 |
| Fibroblast | Samd15 | 6 | 9.76E-54 | 3.35210083 | 0.592 | 0.017 | 1.95E-50 |
| Fibroblast | Scn9a | 6 | 1.92E-51 | 5.34344687 | 0.592 | 0.103 | 3.83E-48 |
| Fibroblast | Cd55 | 6 | 3.04E-51 | 4.81896815 | 0.883 | 0.168 | 6.08E-48 |
| Fibroblast | Kif27 | 6 | 3.53E-51 | 2.50975551 | 0.583 | 0.053 | 7.06E-48 |
| Fibroblast | Cadm1 | 6 | 3.13E-49 | 2.20938497 | 0.466 | 0.04 | 6.27E-46 |
| Fibroblast | Dpp6 | 6 | 5.91E-47 | 2.05603568 | 0.553 | 0.063 | 1.18E-43 |
| Fibroblast | AABR07058170.1 | 6 | 1.39E-45 | 1.84680679 | 0.709 | 0.102 | 2.78E-42 |
| Fibroblast | Troap | 6 | 1.73E-45 | 0.73260555 | 0.398 | 0.023 | 3.45E-42 |

| | | | | | | | |
|---|---|---|---|---|---|---|---|
| Fibroblast | Flt3 | 6 | 3.01E-45 | 3.52398656 | 0.621 | 0.013 | 6.02E-42 |
| Fibroblast | Kcnk3 | 6 | 4.98E-45 | 2.03121452 | 0.592 | 0.035 | 9.96E-42 |
| Fibroblast | Fhad1 | 6 | 9.29E-45 | 3.81313265 | 0.427 | 0.033 | 1.86E-41 |
| Fibroblast | Dact2 | 6 | 3.72E-43 | 5.53478633 | 0.786 | 0.262 | 7.43E-40 |
| Fibroblast | Soat2 | 6 | 2.58E-41 | 3.44278317 | 0.476 | 0.003 | 5.16E-38 |
| Fibroblast | Ildr2 | 6 | 3.20E-41 | 4.04987555 | 0.806 | 0.25 | 6.40E-38 |
| Fibroblast | Gfpt2 | 6 | 1.09E-39 | 3.22108685 | 0.951 | 0.502 | 2.17E-36 |
| Fibroblast | Aurkb | 6 | 4.95E-38 | 0.38324617 | 0.408 | 0.06 | 9.90E-35 |
| Fibroblast | Ca8 | 6 | 1.15E-36 | 2.58756538 | 0.67 | 0.068 | 2.30E-33 |
| Fibroblast | Pmfbp1 | 6 | 2.17E-36 | 3.55547891 | 0.612 | 0.193 | 4.34E-33 |
| Fibroblast | Pnpla3 | 6 | 2.41E-36 | 1.50891488 | 0.709 | 0.159 | 4.82E-33 |
| Fibroblast | Thsd7b1 | 6 | 5.56E-36 | 1.98465026 | 0.621 | 0.06 | 1.11E-32 |
| Fibroblast | Nkain31 | 6 | 3.51E-35 | 1.29435779 | 0.699 | 0.184 | 7.02E-32 |
| Fibroblast | Edn1 | 6 | 5.93E-35 | 0.36628413 | 0.398 | 0.027 | 1.19E-31 |
| Fibroblast | Nkain2 | 6 | 2.02E-34 | 1.25849614 | 0.602 | 0.089 | 4.05E-31 |
| Fibroblast | Lurap1l | 6 | 4.27E-34 | 3.46506979 | 0.825 | 0.293 | 8.53E-31 |
| Fibroblast | Lyz2 | 6 | 7.16E-34 | 1.66672595 | 0.544 | 0.063 | 1.43E-30 |
| Fibroblast | Aldh1a3 | 6 | 1.02E-33 | 3.4420113 | 0.806 | 0.401 | 2.04E-30 |
| Fibroblast | Uap1 | 6 | 1.91E-33 | 2.05708532 | 0.922 | 0.611 | 3.82E-30 |
| Fibroblast | F13a1 | 6 | 2.74E-33 | 0.99264269 | 0.67 | 0.129 | 5.48E-30 |
| Fibroblast | Limch1 | 6 | 7.68E-33 | 2.94614864 | 0.903 | 0.474 | 1.54E-29 |
| Fibroblast | Marchf10 | 6 | 1.39E-32 | 1.01968072 | 0.534 | 0.135 | 2.78E-29 |
| Fibroblast | Phf24 | 6 | 1.73E-32 | 2.97866412 | 0.524 | 0.075 | 3.46E-29 |
| Fibroblast | Cnksr21 | 6 | 6.35E-32 | 0.81751366 | 0.68 | 0.186 | 1.27E-28 |
| Fibroblast | Plaur | 6 | 1.51E-31 | 3.03721703 | 0.806 | 0.25 | 3.02E-28 |
| Fibroblast | Nxnl2 | 6 | 2.18E-31 | 0.36446101 | 0.485 | 0.098 | 4.37E-28 |
| Fibroblast | Myo16 | 6 | 2.66E-31 | 0.69563939 | 0.641 | 0.217 | 5.31E-28 |
| Fibroblast | Klhl40 | 6 | 3.03E-31 | 2.78719277 | 0.592 | 0.172 | 6.06E-28 |
| Fibroblast | Cdh13 | 6 | 5.59E-31 | 2.06277268 | 0.942 | 0.604 | 1.12E-27 |

| | | | | | | | |
|---|---|---|---|---|---|---|---|
| Fibroblast | LOC100910237 | 6 | 7.79E-31 | 0.63701694 | 0.476 | 0.042 | 1.56E-27 |
| Fibroblast | Gap43 | 6 | 2.40E-30 | 4.98738174 | 0.456 | 0.07 | 4.81E-27 |
| Fibroblast | Rhpn2 | 6 | 9.74E-30 | 1.83074134 | 0.699 | 0.217 | 1.95E-26 |
| Fibroblast | Lilrb3a | 6 | 1.49E-29 | 1.59379741 | 0.583 | 0.141 | 2.97E-26 |
| Fibroblast | Lsamp | 6 | 1.70E-29 | 1.8723497 | 0.67 | 0.151 | 3.39E-26 |
| Fibroblast | Unc45b | 6 | 2.02E-29 | 0.56857286 | 0.631 | 0.145 | 4.05E-26 |
| Fibroblast | Samd5 | 6 | 5.10E-29 | 2.02944009 | 0.728 | 0.138 | 1.02E-25 |
| Fibroblast | Cubn | 6 | 5.50E-29 | 1.29344196 | 0.68 | 0.263 | 1.10E-25 |
| Fibroblast | Tagln | 6 | 6.29E-29 | 0.74912873 | 0.524 | 0.133 | 1.26E-25 |
| Fibroblast | Rab11fip4 | 6 | 8.41E-29 | 1.03787647 | 0.68 | 0.218 | 1.68E-25 |
| Fibroblast | Lrrtm4 | 6 | 2.77E-28 | 1.53017514 | 0.641 | 0.136 | 5.54E-25 |
| Fibroblast | Axl | 6 | 6.11E-28 | 1.44751643 | 0.951 | 0.667 | 1.22E-24 |
| Fibroblast | Dmtn | 6 | 6.64E-28 | 1.07747695 | 0.456 | 0.068 | 1.33E-24 |
| Fibroblast | Spock3 | 6 | 9.01E-28 | 2.56588296 | 0.515 | 0.13 | 1.80E-24 |
| Fibroblast | Ect2l | 6 | 9.45E-28 | 2.55548679 | 0.485 | 0.1 | 1.89E-24 |
| Fibroblast | Nppb | 6 | 1.73E-27 | 1.58382837 | 0.718 | 0.207 | 3.46E-24 |
| Fibroblast | Wnt5b1 | 6 | 2.09E-27 | 1.25810689 | 0.544 | 0.194 | 4.18E-24 |
| Fibroblast | Drc3 | 6 | 4.13E-27 | 1.24387012 | 0.553 | 0.03 | 8.26E-24 |
| Fibroblast | Sema3c | 6 | 6.68E-27 | 2.58286269 | 0.767 | 0.345 | 1.34E-23 |
| Fibroblast | Treml4 | 6 | 9.38E-27 | 0.6972045 | 0.485 | 0.126 | 1.88E-23 |
| Fibroblast | Sh3bp2 | 6 | 9.79E-27 | 2.47779325 | 0.748 | 0.293 | 1.96E-23 |
| Fibroblast | Fam189a2 | 6 | 1.15E-26 | 1.54845158 | 0.592 | 0.036 | 2.31E-23 |
| Fibroblast | Stab1 | 6 | 1.47E-26 | 2.43225195 | 0.631 | 0.146 | 2.95E-23 |
| Fibroblast | Galnt151 | 6 | 1.86E-26 | 1.9956061 | 0.767 | 0.39 | 3.72E-23 |
| Fibroblast | Pi16 | 6 | 3.62E-26 | 2.09820445 | 0.913 | 0.564 | 7.24E-23 |
| Fibroblast | Fbln2 | 6 | 7.68E-26 | 2.01042927 | 0.816 | 0.386 | 1.54E-22 |
| Fibroblast | Zdbf21 | 6 | 8.78E-26 | 2.22402655 | 0.417 | 0.099 | 1.76E-22 |
| Fibroblast | Esyt3 | 6 | 9.57E-26 | 1.25641544 | 0.68 | 0.18 | 1.91E-22 |

| | | | | | | | |
|---|---|---|---|---|---|---|---|
| Fibroblast | Ddc2 | 6 | 2.33E-25 | 1.01344687 | 0.631 | 0.197 | 4.66E-22 |
| Fibroblast | Diaph3 | 6 | 3.17E-25 | 4.35261061 | 0.495 | 0.209 | 6.34E-22 |
| Fibroblast | Cd44 | 6 | 5.20E-25 | 1.73701444 | 0.786 | 0.454 | 1.04E-21 |
| Fibroblast | Cd74 | 6 | 6.04E-25 | 1.71666927 | 0.699 | 0.275 | 1.21E-21 |
| Fibroblast | Smpd3 | 6 | 6.63E-25 | 0.39465547 | 0.68 | 0.183 | 1.33E-21 |
| Fibroblast | Csrp2 | 6 | 8.54E-25 | 1.85086285 | 0.602 | 0.174 | 1.71E-21 |
| Fibroblast | Spsb4 | 6 | 3.11E-24 | 2.9324318 | 0.728 | 0.297 | 6.22E-21 |
| Fibroblast | Acta2 | 6 | 4.01E-24 | 1.32826518 | 0.738 | 0.254 | 8.01E-21 |
| Fibroblast | Syt9 | 6 | 4.23E-24 | 0.91051464 | 0.544 | 0.113 | 8.46E-21 |
| Fibroblast | Ppp1r14c | 6 | 5.48E-24 | 2.06006732 | 0.718 | 0.185 | 1.10E-20 |
| Fibroblast | Nova1 | 6 | 7.77E-24 | 2.16869515 | 0.796 | 0.383 | 1.55E-20 |
| Fibroblast | Kcnj8 | 6 | 9.48E-24 | 1.33369194 | 0.515 | 0.123 | 1.90E-20 |
| Fibroblast | Zbtb7c | 6 | 1.27E-23 | 1.74929485 | 0.796 | 0.393 | 2.54E-20 |
| Fibroblast | Gabrb11 | 6 | 1.63E-23 | 1.3156928 | 0.398 | 0.083 | 3.25E-20 |
| Fibroblast | Hs3st5 | 6 | 6.15E-23 | 2.28488144 | 0.699 | 0.219 | 1.23E-19 |
| Fibroblast | Art3 | 6 | 6.39E-23 | 1.46067652 | 0.738 | 0.269 | 1.28E-19 |
| Fibroblast | Nr4a3 | 6 | 7.92E-23 | 2.28551599 | 0.524 | 0.08 | 1.58E-19 |
| Fibroblast | Lama5 | 6 | 1.02E-22 | 0.86940229 | 0.718 | 0.342 | 2.04E-19 |
| Fibroblast | Scara5 | 6 | 1.29E-22 | 1.85770835 | 0.874 | 0.48 | 2.58E-19 |
| Fibroblast | Sh3gl2 | 6 | 1.41E-22 | 1.74658562 | 0.728 | 0.194 | 2.83E-19 |
| Fibroblast | Has1 | 6 | 1.46E-22 | 2.5674741 | 0.709 | 0.265 | 2.92E-19 |
| Fibroblast | Flnb1 | 6 | 2.03E-22 | 2.26432689 | 0.806 | 0.352 | 4.06E-19 |
| Fibroblast | Hspb7 | 6 | 3.53E-22 | 0.37793498 | 0.689 | 0.233 | 7.06E-19 |
| Fibroblast | Optn | 6 | 4.30E-22 | 1.75832488 | 0.786 | 0.276 | 8.59E-19 |
| Fibroblast | Rap1gap2 | 6 | 6.14E-22 | 1.11221283 | 0.757 | 0.321 | 1.23E-18 |
| Fibroblast | Ripor2 | 6 | 9.58E-22 | 0.74122153 | 0.621 | 0.175 | 1.92E-18 |
| Fibroblast | Bcar3 | 6 | 9.64E-22 | 0.48604623 | 0.699 | 0.266 | 1.93E-18 |
| Fibroblast | LOC100911847 | 6 | 1.14E-21 | 1.34987324 | 0.728 | 0.41 | 2.28E-18 |

| | | | | | | | |
|---|---|---|---|---|---|---|---|
| Fibroblast | Lcp1 | 6 | 1.36E-21 | 0.80013463 | 0.427 | 0.148 | 2.73E-18 |
| Fibroblast | Mfap5 | 6 | 1.59E-21 | 1.95461682 | 0.738 | 0.311 | 3.17E-18 |
| Fibroblast | Atp5f1e | 6 | 2.10E-21 | 0.98548649 | 0.748 | 0.387 | 4.21E-18 |
| Fibroblast | Myo18b | 6 | 3.77E-21 | 1.97116662 | 0.699 | 0.325 | 7.55E-18 |
| Fibroblast | Tc2n | 6 | 4.87E-21 | 0.52269614 | 0.718 | 0.331 | 9.74E-18 |
| Fibroblast | Nr4a1 | 6 | 1.15E-20 | 0.41902702 | 0.689 | 0.362 | 2.30E-17 |
| Fibroblast | Odc1 | 6 | 1.79E-20 | 1.20561305 | 0.728 | 0.338 | 3.58E-17 |
| Fibroblast | Pde10a | 6 | 2.04E-20 | 1.66393668 | 0.854 | 0.485 | 4.08E-17 |
| Fibroblast | LOC103693 3231 | 6 | 2.30E-20 | 0.65488617 | 0.583 | 0.139 | 4.60E-17 |
| Fibroblast | Bcl11a | 6 | 3.54E-20 | 2.69245433 | 0.602 | 0.194 | 7.09E-17 |
| Fibroblast | Nptxr | 6 | 5.24E-20 | 0.50948304 | 0.68 | 0.231 | 1.05E-16 |
| Fibroblast | Sh3pxd2b | 6 | 5.76E-20 | 1.38300308 | 0.816 | 0.396 | 1.15E-16 |
| Fibroblast | Fstl1 | 6 | 7.94E-20 | 1.48262278 | 0.874 | 0.573 | 1.59E-16 |
| Fibroblast | Iqgap2 | 6 | 8.16E-20 | 0.4650832 | 0.67 | 0.22 | 1.63E-16 |
| Fibroblast | Crip1 | 6 | 1.26E-19 | 1.13912655 | 0.748 | 0.369 | 2.52E-16 |
| Fibroblast | Pcdh7 | 6 | 1.87E-19 | 0.55299787 | 0.699 | 0.295 | 3.74E-16 |
| Fibroblast | Anxa3 | 6 | 2.05E-19 | 1.10791317 | 0.709 | 0.358 | 4.09E-16 |
| Fibroblast | Gpnmb1 | 6 | 2.09E-19 | 1.24934769 | 0.563 | 0.159 | 4.18E-16 |
| Fibroblast | Rnf152 | 6 | 2.58E-19 | 2.28984849 | 0.485 | 0.204 | 5.16E-16 |
| Fibroblast | Mpp7 | 6 | 2.73E-19 | 0.77350214 | 0.748 | 0.21 | 5.47E-16 |
| Fibroblast | Gbp1 | 6 | 3.23E-19 | 1.33256381 | 0.612 | 0.186 | 6.47E-16 |
| Fibroblast | Ttll7 | 6 | 3.37E-19 | 0.88944959 | 0.738 | 0.217 | 6.75E-16 |
| Fibroblast | Jag1 | 6 | 3.51E-19 | 2.11916036 | 0.68 | 0.246 | 7.01E-16 |
| Fibroblast | Dnajc6 | 6 | 3.92E-19 | 1.65734747 | 0.388 | 0.074 | 7.85E-16 |
| Fibroblast | Ptpro1 | 6 | 5.25E-19 | 0.63089295 | 0.68 | 0.29 | 1.05E-15 |
| Fibroblast | Mgll | 6 | 6.64E-19 | 1.06238746 | 0.786 | 0.445 | 1.33E-15 |
| Fibroblast | Fgf18 | 6 | 8.52E-19 | 2.68460303 | 0.505 | 0.187 | 1.70E-15 |
| Fibroblast | Camk1d | 6 | 9.15E-19 | 1.46501049 | 0.748 | 0.368 | 1.83E-15 |

| | | | | | | | |
|---|---|---|---|---|---|---|---|
| Fibroblast | Negr1 | 6 | 1.87E-18 | 1.87057095 | 0.68 | 0.307 | 3.74E-15 |
| Fibroblast | Cavin4 | 6 | 2.40E-18 | 1.50809404 | 0.553 | 0.117 | 4.80E-15 |
| Fibroblast | Ston2 | 6 | 2.48E-18 | 1.4288938 | 0.903 | 0.533 | 4.96E-15 |
| Fibroblast | Timp41 | 6 | 3.37E-18 | 1.21050336 | 0.583 | 0.201 | 6.75E-15 |
| Fibroblast | Prom1 | 6 | 3.57E-18 | 0.2845916 | 0.65 | 0.285 | 7.14E-15 |
| Fibroblast | Pik3ap1 | 6 | 4.46E-18 | 2.2061576 | 0.592 | 0.164 | 8.92E-15 |
| Fibroblast | Pcdh17 | 6 | 4.89E-18 | 0.64152662 | 0.699 | 0.365 | 9.77E-15 |
| Fibroblast | Fgf14 | 6 | 5.87E-18 | 2.43562009 | 0.621 | 0.238 | 1.17E-14 |
| Fibroblast | Ptprf | 6 | 5.92E-18 | 1.48407476 | 0.689 | 0.266 | 1.18E-14 |
| Fibroblast | Has2 | 6 | 6.77E-18 | 1.91973713 | 0.748 | 0.395 | 1.35E-14 |
| Fibroblast | Ccdc3 | 6 | 7.10E-18 | 1.14276797 | 0.592 | 0.224 | 1.42E-14 |
| Fibroblast | Cobl | 6 | 7.55E-18 | 0.83562143 | 0.699 | 0.332 | 1.51E-14 |
| Fibroblast | Adam23 | 6 | 8.17E-18 | 1.25697147 | 0.709 | 0.276 | 1.63E-14 |
| Fibroblast | Adamts1 | 6 | 1.16E-17 | 1.78588284 | 0.612 | 0.2 | 2.32E-14 |
| Fibroblast | AABR07027581.1 | 6 | 1.21E-17 | 1.2791795 | 0.718 | 0.31 | 2.43E-14 |
| Fibroblast | Ralgps2 | 6 | 1.85E-17 | 2.70838455 | 0.612 | 0.288 | 3.70E-14 |
| Fibroblast | Btn2a2 | 6 | 1.87E-17 | 1.66407518 | 0.495 | 0.157 | 3.74E-14 |
| Fibroblast | Adgrd1 | 6 | 1.90E-17 | 1.64978878 | 0.825 | 0.55 | 3.80E-14 |
| Fibroblast | Ntn4 | 6 | 1.99E-17 | 1.66594065 | 0.816 | 0.431 | 3.97E-14 |
| Fibroblast | Col24a1 | 6 | 2.26E-17 | 0.4841424 | 0.709 | 0.334 | 4.52E-14 |
| Fibroblast | Smyd1 | 6 | 3.45E-17 | 0.88510256 | 0.544 | 0.12 | 6.90E-14 |
| Fibroblast | Hsf51 | 6 | 3.89E-17 | 0.92767226 | 0.583 | 0.239 | 7.77E-14 |
| Fibroblast | Pdcd1lg21 | 6 | 5.00E-17 | 0.50544038 | 0.68 | 0.285 | 1.00E-13 |
| Fibroblast | Adcy3 | 6 | 5.08E-17 | 0.54064047 | 0.748 | 0.396 | 1.02E-13 |
| Fibroblast | Pde7a | 6 | 5.34E-17 | 0.98285592 | 0.874 | 0.424 | 1.07E-13 |
| Fibroblast | Htra2 | 6 | 5.43E-17 | 0.89356718 | 0.709 | 0.35 | 1.09E-13 |
| Fibroblast | Itgb4 | 6 | 5.90E-17 | 0.87745172 | 0.67 | 0.351 | 1.18E-13 |
| Fibroblast | Abcc81 | 6 | 8.14E-17 | 0.42413534 | 0.583 | 0.123 | 1.63E-13 |

| | | | | | | | |
|---|---|---|---|---|---|---|---|
| Fibroblast | Me3 | 6 | 1.03E-16 | 0.62046163 | 0.641 | 0.283 | 2.05E-13 |
| Fibroblast | B2m | 6 | 1.50E-16 | 0.64035172 | 0.728 | 0.27 | 2.99E-13 |
| Fibroblast | Csrp3 | 6 | 1.60E-16 | 1.22648815 | 0.67 | 0.31 | 3.20E-13 |
| Fibroblast | AABR07057997.1 | 6 | 2.26E-16 | 0.98214147 | 0.544 | 0.209 | 4.52E-13 |
| Fibroblast | Ankrd61 | 6 | 2.57E-16 | 0.70150824 | 0.699 | 0.314 | 5.15E-13 |
| Fibroblast | Mob3b | 6 | 2.87E-16 | 0.29548381 | 0.67 | 0.194 | 5.75E-13 |
| Fibroblast | Ptprn21 | 6 | 3.50E-16 | 1.31315426 | 0.612 | 0.265 | 7.01E-13 |
| Fibroblast | Ntrk22 | 6 | 3.90E-16 | 1.2409591 | 0.68 | 0.259 | 7.81E-13 |
| Fibroblast | Cd274 | 6 | 4.27E-16 | 1.29635801 | 0.66 | 0.249 | 8.53E-13 |
| Fibroblast | Sgcd | 6 | 4.94E-16 | 0.38571776 | 0.738 | 0.267 | 9.88E-13 |
| Fibroblast | AABR07052585.1 | 6 | 5.25E-16 | 0.53751374 | 0.709 | 0.277 | 1.05E-12 |
| Fibroblast | Tm4sf4 | 6 | 5.32E-16 | 0.85143437 | 0.631 | 0.202 | 1.06E-12 |
| Fibroblast | Plcb11 | 6 | 9.34E-16 | 1.33059844 | 0.689 | 0.265 | 1.87E-12 |
| Fibroblast | PCOLCE2 | 6 | 1.15E-15 | 1.77235033 | 0.718 | 0.315 | 2.30E-12 |
| Fibroblast | Usp2 | 6 | 1.16E-15 | 1.43193828 | 0.553 | 0.212 | 2.33E-12 |
| Fibroblast | Cacna2d3 | 6 | 1.52E-15 | 1.27802604 | 0.825 | 0.519 | 3.03E-12 |
| Fibroblast | Egflam | 6 | 1.63E-15 | 1.08577061 | 0.602 | 0.241 | 3.26E-12 |
| Fibroblast | Sgpp21 | 6 | 2.54E-15 | 2.18237368 | 0.476 | 0.202 | 5.08E-12 |
| Fibroblast | Prg4 | 6 | 2.57E-15 | 0.44805965 | 0.718 | 0.371 | 5.15E-12 |
| Fibroblast | AABR07044900.1 | 6 | 3.62E-15 | 0.70067158 | 0.718 | 0.277 | 7.25E-12 |
| Fibroblast | Rgcc | 6 | 4.11E-15 | 1.13952547 | 0.544 | 0.201 | 8.22E-12 |
| Fibroblast | Pde1c | 6 | 4.12E-15 | 0.94591383 | 0.631 | 0.247 | 8.25E-12 |
| Fibroblast | AABR07026536.1 | 6 | 4.42E-15 | 0.57428395 | 0.65 | 0.269 | 8.85E-12 |
| Fibroblast | Tango2 | 6 | 4.52E-15 | 0.66583944 | 0.738 | 0.272 | 9.03E-12 |
| Fibroblast | Sema3a1 | 6 | 4.91E-15 | 0.83929994 | 0.495 | 0.185 | 9.82E-12 |

| | | | | | | | |
|---|---|---|---|---|---|---|---|
| Fibroblast | Dkk21 | 6 | 5.31E-15 | 1.45275552 | 0.728 | 0.302 | 1.06E-11 |
| Fibroblast | Chst14 | 6 | 5.71E-15 | 2.0605136 | 0.476 | 0.143 | 1.14E-11 |
| Fibroblast | Hspb8 | 6 | 5.99E-15 | 1.56663802 | 0.602 | 0.281 | 1.20E-11 |
| Fibroblast | Fn1 | 6 | 6.80E-15 | 1.5982577 | 0.592 | 0.22 | 1.36E-11 |
| Fibroblast | Epha4 | 6 | 9.07E-15 | 0.61969389 | 0.631 | 0.323 | 1.81E-11 |
| Fibroblast | Nuak1 | 6 | 1.53E-14 | 0.43899369 | 0.738 | 0.401 | 3.06E-11 |
| Fibroblast | Gja11 | 6 | 1.59E-14 | 0.77357096 | 0.573 | 0.232 | 3.18E-11 |
| Fibroblast | Tmem51 | 6 | 2.04E-14 | 1.7428995 | 0.495 | 0.211 | 4.08E-11 |
| Fibroblast | Dnah5 | 6 | 2.26E-14 | 1.56501876 | 0.67 | 0.302 | 4.51E-11 |
| Fibroblast | Mylk3 | 6 | 2.53E-14 | 0.59595251 | 0.699 | 0.325 | 5.05E-11 |
| Fibroblast | AABR07017268.11 | 6 | 3.88E-14 | 1.18343803 | 0.602 | 0.216 | 7.77E-11 |
| Fibroblast | Kitlg | 6 | 4.40E-14 | 1.07324655 | 0.777 | 0.385 | 8.80E-11 |
| Fibroblast | Emcn | 6 | 4.64E-14 | 0.29375118 | 0.699 | 0.351 | 9.29E-11 |
| Fibroblast | Slco3a1 | 6 | 4.79E-14 | 1.19668445 | 0.806 | 0.531 | 9.58E-11 |
| Fibroblast | Srpx | 6 | 5.66E-14 | 0.60559813 | 0.699 | 0.276 | 1.13E-10 |
| Fibroblast | Flrt2 | 6 | 7.40E-14 | 1.3390112 | 0.728 | 0.312 | 1.48E-10 |
| Fibroblast | Slc25a20 | 6 | 7.66E-14 | 0.92049709 | 0.68 | 0.383 | 1.53E-10 |
| Fibroblast | Flnc1 | 6 | 8.52E-14 | 0.69901722 | 0.631 | 0.348 | 1.70E-10 |
| Fibroblast | AABR07025140.11 | 6 | 1.31E-13 | 1.05578842 | 0.621 | 0.271 | 2.61E-10 |
| Fibroblast | Nalcn | 6 | 1.58E-13 | 1.03660441 | 0.612 | 0.207 | 3.15E-10 |
| Fibroblast | Anp32b | 6 | 2.14E-13 | 0.66881128 | 0.728 | 0.386 | 4.28E-10 |
| Fibroblast | Prkch | 6 | 2.65E-13 | 0.64543011 | 0.699 | 0.285 | 5.29E-10 |
| Fibroblast | Plk2 | 6 | 3.03E-13 | 0.5766977 | 0.641 | 0.342 | 6.07E-10 |
| Fibroblast | Pdlim5 | 6 | 3.29E-13 | 1.26288439 | 0.903 | 0.585 | 6.58E-10 |
| Fibroblast | Rimbp2 | 6 | 3.92E-13 | 1.81576585 | 0.544 | 0.266 | 7.84E-10 |
| Fibroblast | Ccn11 | 6 | 4.48E-13 | 0.28478462 | 0.631 | 0.328 | 8.95E-10 |
| Fibroblast | Kdr1 | 6 | 5.93E-13 | 0.54830766 | 0.709 | 0.315 | 1.19E-09 |

| | | | | | | | |
|---|---|---|---|---|---|---|---|
| Fibroblast | Vcl | 6 | 6.28E-13 | 1.03680823 | 0.748 | 0.311 | 1.26E-09 |
| Fibroblast | Ncald | 6 | 6.37E-13 | 1.15255126 | 0.728 | 0.34 | 1.27E-09 |
| Fibroblast | Ptpn18 | 6 | 6.99E-13 | 1.86792457 | 0.398 | 0.126 | 1.40E-09 |
| Fibroblast | LOC691083 | 6 | 7.67E-13 | 0.63232399 | 0.68 | 0.313 | 1.53E-09 |
| Fibroblast | Fbxo40 | 6 | 7.87E-13 | 1.32647578 | 0.505 | 0.193 | 1.57E-09 |
| Fibroblast | Dipk1a | 6 | 9.44E-13 | 0.40787403 | 0.67 | 0.314 | 1.89E-09 |
| Fibroblast | Col22a1 | 6 | 1.08E-12 | 0.68005271 | 0.641 | 0.344 | 2.16E-09 |
| Fibroblast | Prkcz | 6 | 1.38E-12 | 0.3693408 | 0.65 | 0.264 | 2.76E-09 |
| Fibroblast | Csrp1 | 6 | 2.35E-12 | 0.67953877 | 0.738 | 0.465 | 4.71E-09 |
| Fibroblast | Pcdh19 | 6 | 2.46E-12 | 0.76830509 | 0.631 | 0.301 | 4.93E-09 |
| Fibroblast | Gfod1 | 6 | 2.52E-12 | 0.34642608 | 0.738 | 0.321 | 5.05E-09 |
| Fibroblast | Arhgap11a | 6 | 3.16E-12 | 0.37177489 | 0.515 | 0.208 | 6.33E-09 |
| Fibroblast | Pip5k1b1 | 6 | 3.23E-12 | 0.6195646 | 0.709 | 0.313 | 6.45E-09 |
| Fibroblast | Itga6 | 6 | 3.30E-12 | 0.59233133 | 0.728 | 0.386 | 6.60E-09 |
| Fibroblast | Itga9 | 6 | 3.37E-12 | 1.28743808 | 0.903 | 0.637 | 6.74E-09 |
| Fibroblast | LOC691141 | 6 | 4.16E-12 | 1.21865327 | 0.485 | 0.151 | 8.31E-09 |
| Fibroblast | Adamts5 | 6 | 4.42E-12 | 1.44407708 | 0.883 | 0.574 | 8.85E-09 |
| Fibroblast | Medag | 6 | 5.11E-12 | 1.02218659 | 0.718 | 0.437 | 1.02E-08 |
| Fibroblast | Kcnab1 | 6 | 6.32E-12 | 0.87558029 | 0.621 | 0.342 | 1.26E-08 |
| Fibroblast | Slit3 | 6 | 7.73E-12 | 0.97326087 | 0.67 | 0.406 | 1.55E-08 |
| Fibroblast | Cfh | 6 | 1.06E-11 | 0.45592249 | 0.718 | 0.368 | 2.12E-08 |
| Fibroblast | Musk | 6 | 1.11E-11 | 0.35501112 | 0.65 | 0.279 | 2.22E-08 |
| Fibroblast | Ptgs2 | 6 | 1.25E-11 | 0.66565819 | 0.553 | 0.181 | 2.50E-08 |
| Fibroblast | Rasa41 | 6 | 1.32E-11 | 0.34373774 | 0.641 | 0.298 | 2.64E-08 |
| Fibroblast | Cdk19 | 6 | 1.98E-11 | 0.60222762 | 0.786 | 0.379 | 3.95E-08 |
| Fibroblast | Fsd21 | 6 | 2.09E-11 | 0.29489454 | 0.602 | 0.254 | 4.19E-08 |
| Fibroblast | Taldo11 | 6 | 3.30E-11 | 1.03679431 | 0.621 | 0.216 | 6.60E-08 |
| Fibroblast | Kcnt2 | 6 | 3.50E-11 | 0.49518294 | 0.505 | 0.178 | 7.00E-08 |
| Fibroblast | Nnt1 | 6 | 4.82E-11 | 0.63546275 | 0.709 | 0.383 | 9.64E-08 |

| | | | | | | | |
|---|---|---|---|---|---|---|---|
| Fibroblast | Ldb31 | 6 | 6.43E-11 | 0.2699799 | 0.709 | 0.322 | 1.29E-07 |
| Fibroblast | Gpr63 | 6 | 7.08E-11 | 1.07600836 | 0.66 | 0.284 | 1.42E-07 |
| Fibroblast | Fcgr2b | 6 | 8.50E-11 | 1.23305411 | 0.437 | 0.148 | 1.70E-07 |
| Fibroblast | Tmsb4x | 6 | 1.46E-10 | 0.88981739 | 0.68 | 0.312 | 2.92E-07 |
| Fibroblast | Plekha4 | 6 | 1.66E-10 | 0.8911324 | 0.699 | 0.38 | 3.32E-07 |
| Fibroblast | Mgmt | 6 | 2.02E-10 | 0.56626074 | 0.767 | 0.404 | 4.04E-07 |
| Fibroblast | Prrx1 | 6 | 2.33E-10 | 1.03622715 | 0.864 | 0.591 | 4.66E-07 |
| Fibroblast | Ptprj | 6 | 2.40E-10 | 0.84343877 | 0.922 | 0.617 | 4.80E-07 |
| Fibroblast | Fbxo32 | 6 | 2.61E-10 | 0.84539177 | 0.65 | 0.399 | 5.21E-07 |
| Fibroblast | Ptpn3 | 6 | 4.73E-10 | 0.42496813 | 0.709 | 0.348 | 9.47E-07 |
| Fibroblast | Rcan21 | 6 | 7.89E-10 | 1.10812157 | 0.835 | 0.496 | 1.58E-06 |
| Fibroblast | P2rx71 | 6 | 8.00E-10 | 0.91743829 | 0.544 | 0.249 | 1.60E-06 |
| Fibroblast | Mcf2l | 6 | 9.07E-10 | 1.35561272 | 0.495 | 0.213 | 1.81E-06 |
| Fibroblast | Cadps2 | 6 | 1.07E-09 | 0.65678732 | 0.641 | 0.296 | 2.14E-06 |
| Fibroblast | Slfn13 | 6 | 1.88E-09 | 0.39843112 | 0.709 | 0.275 | 3.76E-06 |
| Fibroblast | Antxr11 | 6 | 1.88E-09 | 0.84254721 | 0.825 | 0.521 | 3.76E-06 |
| Fibroblast | Afap1l2 | 6 | 2.31E-09 | 0.25227017 | 0.757 | 0.45 | 4.63E-06 |
| Fibroblast | Tspan5 | 6 | 2.70E-09 | 0.76921598 | 0.825 | 0.466 | 5.40E-06 |
| Fibroblast | Cdkn1a1 | 6 | 2.72E-09 | 0.77647273 | 0.524 | 0.253 | 5.44E-06 |
| Fibroblast | Slc38a1 | 6 | 5.47E-09 | 0.39079544 | 0.563 | 0.226 | 1.09E-05 |
| Fibroblast | Slc1a1 | 6 | 1.10E-08 | 1.16153189 | 0.524 | 0.174 | 2.20E-05 |
| Fibroblast | AABR07058158.1 | 6 | 1.31E-08 | 0.63223652 | 0.777 | 0.398 | 2.63E-05 |
| Fibroblast | Ralgapa2 | 6 | 1.41E-08 | 0.32533952 | 0.786 | 0.437 | 2.83E-05 |
| Fibroblast | Hydin | 6 | 3.63E-08 | 0.80250463 | 0.495 | 0.207 | 7.25E-05 |
| Fibroblast | Kif22 | 6 | 4.81E-08 | 2.80793978 | 0.447 | 0.187 | 9.62E-05 |
| Fibroblast | Rnls | 6 | 5.64E-08 | 0.5846403 | 0.689 | 0.38 | 0.00011275 |
| Fibroblast | Col16a1 | 6 | 6.13E-08 | 0.40746162 | 0.641 | 0.298 | 0.00012258 |
| Fibroblast | Polr2m | 6 | 7.14E-08 | 0.38918299 | 0.738 | 0.399 | 0.00014286 |

| | | | | | | | |
|---|---|---|---|---|---|---|---|
| Fibroblast | Aacs | 6 | 1.11E-07 | 0.68345325 | 0.553 | 0.296 | 0.00022132 |
| Fibroblast | Phyh | 6 | 1.11E-07 | 0.42790696 | 0.738 | 0.41 | 0.00022243 |
| Fibroblast | AABR07025295.1 | 6 | 2.16E-07 | 0.64812693 | 0.718 | 0.393 | 0.00043288 |
| Fibroblast | Actn2 | 6 | 2.79E-07 | 0.37750829 | 0.631 | 0.322 | 0.0005576 |
| Fibroblast | Ncam2 | 6 | 4.80E-07 | 1.2048839 | 0.456 | 0.198 | 0.0009595 |
| Fibroblast | Nox4 | 6 | 1.02E-06 | 0.32187903 | 0.66 | 0.353 | 0.00204253 |
| Fibroblast | Slc25a13 | 6 | 1.12E-06 | 0.30648423 | 0.699 | 0.365 | 0.00224071 |
| Fibroblast | Gnao1 | 6 | 1.33E-06 | 0.38743549 | 0.621 | 0.359 | 0.00266593 |
| Fibroblast | Sat1 | 6 | 1.39E-06 | 0.29179021 | 0.709 | 0.354 | 0.00278301 |
| Fibroblast | Sorbs1 | 6 | 1.48E-06 | 0.60875855 | 0.835 | 0.553 | 0.00295646 |
| Fibroblast | Efemp1 | 6 | 3.37E-06 | 0.3946889 | 0.68 | 0.428 | 0.00674064 |
| Fibroblast | Ntn1 | 6 | 3.96E-06 | 0.36033014 | 0.796 | 0.54 | 0.00792602 |
| Fibroblast | Ckm | 6 | 7.28E-06 | 0.49885788 | 0.68 | 0.385 | 0.01456423 |
| Fibroblast | Zfp622 | 6 | 9.40E-06 | 0.34672215 | 0.718 | 0.406 | 0.01880331 |
| Fibroblast | Lrg1 | 7 | 1.14E-23 | 0.95625981 | 0.261 | 0.011 | 2.27E-20 |
| Fibroblast | Vil1 | 7 | 1.47E-19 | 1.61999706 | 0.304 | 0.049 | 2.94E-16 |
| Fibroblast | AABR07005983.1 | 7 | 4.22E-19 | 0.49728316 | 0.304 | 0.05 | 8.43E-16 |
| Fibroblast | Crhr2 | 7 | 9.06E-17 | 0.25782064 | 0.42 | 0.129 | 1.81E-13 |
| Fibroblast | Grik4 | 7 | 1.17E-13 | 2.51091344 | 0.377 | 0.1 | 2.34E-10 |
| Fibroblast | Etl4 | 7 | 5.45E-12 | 2.00102262 | 0.696 | 0.373 | 1.09E-08 |
| Fibroblast | Cyyr1 | 7 | 9.92E-12 | 2.0452332 | 0.681 | 0.344 | 1.98E-08 |
| Fibroblast | Epas1 | 7 | 9.56E-11 | 1.58308563 | 0.797 | 0.505 | 1.91E-07 |
| Fibroblast | Ptprb | 7 | 2.96E-10 | 1.95332609 | 0.652 | 0.297 | 5.91E-07 |
| Fibroblast | Adgrf5 | 7 | 4.72E-10 | 2.06686621 | 0.652 | 0.392 | 9.45E-07 |
| Fibroblast | Adgrl4 | 7 | 1.31E-09 | 2.03668368 | 0.609 | 0.346 | 2.63E-06 |
| Fibroblast | Unc5c1 | 7 | 2.69E-09 | 2.17626447 | 0.377 | 0.108 | 5.38E-06 |
| Fibroblast | Flvcr2 | 7 | 1.83E-08 | 0.746626 | 0.348 | 0.089 | 3.66E-05 |

| | | | | | | | |
|---|---|---|---|---|---|---|---|
| Fibroblast | Herc6 | 7 | 5.39E-08 | 2.06495194 | 0.609 | 0.253 | 0.00010786 |
| Fibroblast | Mx2 | 7 | 1.21E-07 | 3.49269604 | 0.42 | 0.133 | 0.00024235 |
| Fibroblast | AC123500.1 | 7 | 1.22E-07 | 3.16015426 | 0.377 | 0.1 | 0.00024393 |
| Fibroblast | Mcf2l1 | 7 | 1.61E-07 | 1.97144667 | 0.522 | 0.217 | 0.00032104 |
| Fibroblast | Crim1 | 7 | 8.81E-07 | 1.48424602 | 0.725 | 0.458 | 0.00176183 |
| Fibroblast | Slc1a11 | 7 | 9.14E-07 | 1.85170946 | 0.449 | 0.183 | 0.00182773 |
| Fibroblast | Rnf2131 | 7 | 1.50E-06 | 1.70114302 | 0.652 | 0.393 | 0.00299686 |
| Fibroblast | Cadm2 | 7 | 5.59E-06 | 2.638065 | 0.594 | 0.328 | 0.01118181 |
| Fibroblast | Smyd11 | 7 | 9.88E-06 | 0.86996481 | 0.391 | 0.133 | 0.01975225 |
| Fibroblast | Cdk191 | 7 | 2.43E-05 | 1.01127739 | 0.667 | 0.391 | 0.04864693 |
| Fibroblast | Tyrobp | 8 | 3.62E-72 | 3.7472018 | 0.373 | 0.015 | 7.23E-69 |
| Fibroblast | Apobec1 | 8 | 7.80E-69 | 4.29844184 | 0.343 | 0.013 | 1.56E-65 |
| Fibroblast | Hdc | 8 | 9.21E-66 | 4.43153178 | 0.284 | 0.008 | 1.84E-62 |
| Fibroblast | Tbx21 | 8 | 8.64E-63 | 3.60345105 | 0.313 | 0.013 | 1.73E-59 |
| Fibroblast | Sdc1 | 8 | 3.60E-62 | 3.3975462 | 0.343 | 0.013 | 7.19E-59 |
| Fibroblast | Fgf5 | 8 | 2.09E-42 | 6.6669807 | 0.313 | 0.021 | 4.18E-39 |
| Fibroblast | Hao1 | 8 | 1.13E-37 | 5.9343173 | 0.358 | 0.034 | 2.25E-34 |
| Fibroblast | AABR07068046.1 | 8 | 3.67E-33 | 8.23549005 | 0.313 | 0.012 | 7.35E-30 |
| Fibroblast | AABR07067469.1 | 8 | 2.75E-31 | 6.53725178 | 0.522 | 0.096 | 5.50E-28 |
| Fibroblast | Map3k7cl | 8 | 1.19E-24 | 3.287507 | 0.373 | 0.043 | 2.39E-21 |
| Fibroblast | Ltbp2 | 8 | 2.89E-24 | 3.38489462 | 0.851 | 0.507 | 5.77E-21 |
| Fibroblast | Arhgef39 | 8 | 4.77E-24 | 6.27624201 | 0.328 | 0.012 | 9.55E-21 |
| Fibroblast | Ptprv | 8 | 4.75E-23 | 4.58054803 | 0.493 | 0.101 | 9.50E-20 |
| Fibroblast | Galnt141 | 8 | 2.01E-22 | 5.09741095 | 0.433 | 0.089 | 4.01E-19 |
| Fibroblast | Il11 | 8 | 4.36E-22 | 2.44219412 | 0.328 | 0.013 | 8.72E-19 |
| Fibroblast | Tpx2 | 8 | 5.41E-17 | 4.61721272 | 0.433 | 0.018 | 1.08E-13 |

| | | | | | | | |
|---|---|---|---|---|---|---|---|
| Fibroblast | Rergl | 8 | 3.58E-16 | 2.45155085 | 0.358 | 0.072 | 7.17E-13 |
| Fibroblast | Klhl29 | 8 | 1.80E-14 | 2.30540475 | 0.806 | 0.451 | 3.60E-11 |
| Fibroblast | Prc1 | 8 | 1.88E-14 | 3.26214878 | 0.284 | 0.005 | 3.76E-11 |
| Fibroblast | Fn11 | 8 | 2.51E-14 | 3.44656436 | 0.716 | 0.223 | 5.02E-11 |
| Fibroblast | Pdgfc | 8 | 5.43E-14 | 2.03352064 | 0.433 | 0.036 | 1.09E-10 |
| Fibroblast | Itgal | 8 | 6.60E-14 | 4.58806033 | 0.313 | 0.013 | 1.32E-10 |
| Fibroblast | Cdh1 | 8 | 9.56E-14 | 3.04398954 | 0.463 | 0.12 | 1.91E-10 |
| Fibroblast | Tagln1 | 8 | 1.40E-13 | 3.41881001 | 0.463 | 0.142 | 2.79E-10 |
| Fibroblast | Itga10 | 8 | 1.47E-13 | 4.6136898 | 0.358 | 0.038 | 2.94E-10 |
| Fibroblast | Crlf1 | 8 | 4.15E-13 | 2.95267119 | 0.716 | 0.444 | 8.31E-10 |
| Fibroblast | Runx1 | 8 | 4.49E-13 | 2.67672954 | 0.761 | 0.475 | 8.98E-10 |
| Fibroblast | Kcnh5 | 8 | 1.35E-12 | 2.4323585 | 0.343 | 0.06 | 2.70E-09 |
| Fibroblast | Itgb21 | 8 | 1.66E-12 | 1.78867637 | 0.448 | 0.153 | 3.33E-09 |
| Fibroblast | Tceal7 | 8 | 1.79E-12 | 0.79738336 | 0.343 | 0.079 | 3.57E-09 |
| Fibroblast | Hey2 | 8 | 2.01E-12 | 3.12328942 | 0.343 | 0.09 | 4.03E-09 |
| Fibroblast | Cobl1 | 8 | 3.16E-11 | 3.21725237 | 0.657 | 0.34 | 6.33E-08 |
| Fibroblast | Spp1 | 8 | 2.81E-10 | 2.03782198 | 0.403 | 0.092 | 5.61E-07 |
| Fibroblast | Kif111 | 8 | 3.71E-10 | 2.56847998 | 0.328 | 0.078 | 7.42E-07 |
| Fibroblast | Trpc6 | 8 | 7.75E-10 | 3.78602508 | 0.343 | 0.052 | 1.55E-06 |
| Fibroblast | Actg2 | 8 | 1.91E-09 | 3.87488751 | 0.343 | 0.085 | 3.82E-06 |
| Fibroblast | Col1a1 | 8 | 6.91E-09 | 1.70194879 | 0.806 | 0.485 | 1.38E-05 |
| Fibroblast | Ifit1bl | 8 | 1.17E-08 | 2.99504052 | 0.418 | 0.161 | 2.35E-05 |
| Fibroblast | Pmepa1 | 8 | 1.21E-08 | 1.35701062 | 0.776 | 0.432 | 2.42E-05 |
| Fibroblast | Col27a1 | 8 | 1.56E-08 | 2.43003041 | 0.627 | 0.324 | 3.12E-05 |
| Fibroblast | Anks1b | 8 | 2.20E-08 | 4.87587709 | 0.507 | 0.082 | 4.40E-05 |
| Fibroblast | Olfm2 | 8 | 2.92E-08 | 3.46184915 | 0.463 | 0.184 | 5.84E-05 |
| Fibroblast | Bcat1 | 8 | 5.21E-08 | 1.91706023 | 0.657 | 0.327 | 0.00010424 |
| Fibroblast | Col19a1 | 8 | 2.10E-07 | 1.22233796 | 0.418 | 0.123 | 0.00042059 |
| Fibroblast | Dkk3 | 8 | 2.23E-07 | 1.31076001 | 0.687 | 0.416 | 0.00044659 |

| | | | | | | | |
|---|---|---|---|---|---|---|---|
| Fibroblast | Chst11 | 8 | 2.66E-07 | 1.7538915 | 0.731 | 0.4 | 0.00053282 |
| Fibroblast | Ccl2 | 8 | 1.03E-06 | 3.0685483 | 0.433 | 0.183 | 0.0020612 |
| Fibroblast | Flna | 8 | 1.46E-06 | 1.38374109 | 0.701 | 0.379 | 0.00292417 |
| Fibroblast | Phf241 | 8 | 1.74E-06 | 2.93831892 | 0.343 | 0.089 | 0.00347239 |
| Fibroblast | AC134204.1 | 8 | 3.00E-06 | 0.90729759 | 0.701 | 0.388 | 0.00599707 |
| Fibroblast | Atp10a | 8 | 8.00E-06 | 0.83745894 | 0.761 | 0.484 | 0.01600584 |
| Fibroblast | Fgf11 | 8 | 1.34E-05 | 1.99751061 | 0.582 | 0.325 | 0.02688759 |
| Fibroblast | Bgn | 8 | 1.47E-05 | 1.46286063 | 0.627 | 0.337 | 0.02939868 |
| Fibroblast | Vim | 8 | 1.86E-05 | 1.28620694 | 0.582 | 0.31 | 0.03716221 |
| Fibroblast | Vcam1 | 8 | 2.40E-05 | 2.80310447 | 0.522 | 0.194 | 0.04806478 |
| Fibroblast | Spdef | 9 | 2.99E-12 | 3.07887111 | 0.4 | 0.028 | 5.97E-09 |
| Fibroblast | Tmem163 | 9 | 2.79E-10 | 6.62495542 | 0.6 | 0.012 | 5.59E-07 |
| Fibroblast | Brinp1 | 9 | 5.51E-08 | 6.35396124 | 0.3 | 0.002 | 0.00011028 |
| Fibroblast | Neto1 | 9 | 5.67E-08 | 2.69827346 | 0.5 | 0.072 | 0.00011341 |
| Fibroblast | Ifit3 | 9 | 5.37E-07 | 2.69934897 | 0.4 | 0.03 | 0.00107346 |
| Fibroblast | Tnni3 | 9 | 1.16E-06 | 2.62424006 | 1 | 0.602 | 0.00232269 |
| Fibroblast | Nppa | 9 | 1.93E-06 | 4.62363122 | 0.9 | 0.402 | 0.00385883 |
| Fibroblast | Tmem2361 | 9 | 2.46E-06 | 3.43199055 | 0.5 | 0.079 | 0.00492866 |
| Fibroblast | Mb | 9 | 2.93E-06 | 4.22731393 | 0.9 | 0.52 | 0.00586855 |
| Fibroblast | Adgre4 | 9 | 3.97E-06 | 5.33722495 | 0.6 | 0.06 | 0.00794603 |
| Fibroblast | Selp | 9 | 6.56E-06 | 3.17667697 | 0.3 | 0.047 | 0.01312022 |
| Fibroblast | Smc1b1 | 9 | 8.36E-06 | 3.39402655 | 0.4 | 0.078 | 0.01672344 |
| Fibroblast | Ankrd1 | 9 | 1.29E-05 | 2.7668679 | 0.9 | 0.579 | 0.02579052 |
| Fibroblast | LOC100910636 | 9 | 1.99E-05 | 3.18120563 | 0.5 | 0.068 | 0.0397211 |
| Fibroblast | LOC308990 | 9 | 2.42E-05 | 3.21801719 | 0.4 | 0.061 | 0.04840792 |

| | | | | | | | |
|---|---|---|---|---|---|---|---|
| Lymphatic Endothelial Cell | Lyn | 0 | 1.57E-05 | 1.19881816 | 0.797 | 0.528 | 0.03132908 |
| Lymphatic Endothelial Cell | Klhl29 | 1 | 9.42E-08 | 2.38051125 | 0.84 | 0.568 | 0.00018841 |
| Lymphatic Endothelial Cell | Gpm6a | 1 | 1.57E-05 | 1.58668309 | 0.86 | 0.605 | 0.03135142 |
| Lymphatic Endothelial Cell | Sgca | 2 | 1.25E-14 | 2.64034859 | 0.455 | 0.028 | 2.49E-11 |
| Lymphatic Endothelial Cell | Aacs | 2 | 4.18E-12 | 2.40963123 | 0.409 | 0.018 | 8.36E-09 |
| Lymphatic Endothelial Cell | Prg4 | 2 | 7.81E-12 | 2.35751697 | 0.273 | 0.009 | 1.56E-08 |
| Lymphatic Endothelial Cell | Ntrk2 | 2 | 5.74E-11 | 2.42357127 | 0.455 | 0.028 | 1.15E-07 |
| Lymphatic Endothelial Cell | Popdc2 | 2 | 7.30E-11 | 6.45002022 | 0.364 | 0.009 | 1.46E-07 |
| Lymphatic Endothelial Cell | Pla2g2a | 2 | 1.20E-10 | 4.65392528 | 0.318 | 0 | 2.40E-07 |

| | | | | | | | |
|---|---|---|---|---|---|---|---|
| Lymphatic Endothelial Cell | Rab11fip4 | 2 | 2.26E-10 | 2.28416066 | 0.318 | 0.028 | 4.53E-07 |
| Lymphatic Endothelial Cell | Ccdc3 | 2 | 1.78E-09 | 7.23327879 | 0.273 | 0.009 | 3.57E-06 |
| Lymphatic Endothelial Cell | Fgf10 | 2 | 2.85E-09 | 5.63562721 | 0.5 | 0.119 | 5.70E-06 |
| Lymphatic Endothelial Cell | Frmd3 | 2 | 5.27E-09 | 2.37927149 | 0.364 | 0.037 | 1.05E-05 |
| Lymphatic Endothelial Cell | Sorcs1 | 2 | 1.17E-08 | 2.44406596 | 0.409 | 0.046 | 2.34E-05 |
| Lymphatic Endothelial Cell | AABR07007068.1 | 2 | 1.81E-08 | 5.83330113 | 0.409 | 0.046 | 3.62E-05 |
| Lymphatic Endothelial Cell | Col11a2 | 2 | 1.81E-08 | 2.08827036 | 0.318 | 0.028 | 3.62E-05 |
| Lymphatic Endothelial Cell | AABR07038986.1 | 2 | 2.36E-08 | 2.22665862 | 0.273 | 0.018 | 4.72E-05 |
| Lymphatic Endothelial Cell | Prss48 | 2 | 2.86E-08 | 2.35947035 | 0.273 | 0 | 5.72E-05 |

| | | | | | | | |
|---|---|---|---|---|---|---|---|
| Lymphatic Endothelial Cell | Sctr | 2 | 2.97E-08 | 4.33027643 | 0.273 | 0.009 | 5.94E-05 |
| Lymphatic Endothelial Cell | Bdkrb2 | 2 | 3.10E-08 | 2.36255968 | 0.409 | 0.046 | 6.21E-05 |
| Lymphatic Endothelial Cell | Astn2 | 2 | 4.07E-08 | 0.65334337 | 0.318 | 0.046 | 8.14E-05 |
| Lymphatic Endothelial Cell | Hs6st2 | 2 | 6.71E-08 | 6.19483995 | 0.318 | 0.028 | 0.00013429 |
| Lymphatic Endothelial Cell | Prima1 | 2 | 1.59E-07 | 4.14298491 | 0.5 | 0.101 | 0.00031803 |
| Lymphatic Endothelial Cell | Nlrp12 | 2 | 2.76E-07 | 2.37997128 | 0.318 | 0.018 | 0.00055184 |
| Lymphatic Endothelial Cell | Pdzd2 | 2 | 2.84E-07 | 1.95870348 | 0.773 | 0.229 | 0.0005686 |
| Lymphatic Endothelial Cell | Slc16a10 | 2 | 2.96E-07 | 1.82392616 | 0.364 | 0.046 | 0.00059238 |
| Lymphatic Endothelial Cell | Trim7 | 2 | 2.97E-07 | 4.32534942 | 0.318 | 0.018 | 0.0005946 |

| | | | | | | | |
|---|---|---|---|---|---|---|---|
| Lymphatic Endothelial Cell | Tll2 | 2 | 3.71E-07 | 5.67352037 | 0.273 | 0.018 | 0.00074293 |
| Lymphatic Endothelial Cell | Vit | 2 | 4.86E-07 | 2.34200973 | 0.273 | 0.009 | 0.00097241 |
| Lymphatic Endothelial Cell | Enpp3 | 2 | 5.54E-07 | 5.07297946 | 0.409 | 0.083 | 0.00110709 |
| Lymphatic Endothelial Cell | Antxr1 | 2 | 5.54E-07 | 3.78668606 | 0.364 | 0.046 | 0.00110709 |
| Lymphatic Endothelial Cell | Steap4 | 2 | 5.76E-07 | 2.33611794 | 0.364 | 0.11 | 0.00115106 |
| Lymphatic Endothelial Cell | Lef1 | 2 | 1.07E-06 | 0.94989084 | 0.364 | 0.092 | 0.00214568 |
| Lymphatic Endothelial Cell | Crlf1 | 2 | 1.14E-06 | 4.78554669 | 0.455 | 0.101 | 0.0022865 |
| Lymphatic Endothelial Cell | Magi2 | 2 | 1.93E-06 | 3.37084808 | 0.5 | 0.018 | 0.00386791 |
| Lymphatic Endothelial Cell | Asb14 | 2 | 1.99E-06 | 5.30373802 | 0.273 | 0.018 | 0.00398577 |

| | | | | | | | |
|---|---|---|---|---|---|---|---|
| Lymphatic Endothelial Cell | Plac8 | 2 | 2.13E-06 | 2.43191612 | 0.318 | 0.028 | 0.00425812 |
| Lymphatic Endothelial Cell | Col12a1 | 2 | 2.21E-06 | 1.12776564 | 0.364 | 0.046 | 0.00442676 |
| Lymphatic Endothelial Cell | Ckap2 | 2 | 4.10E-06 | 2.31685042 | 0.273 | 0.018 | 0.00819539 |
| Lymphatic Endothelial Cell | Cntn3 | 2 | 4.25E-06 | 2.44513126 | 0.318 | 0.037 | 0.00849589 |
| Lymphatic Endothelial Cell | Adhfe1 | 2 | 4.37E-06 | 2.17905222 | 0.409 | 0.119 | 0.00874956 |
| Lymphatic Endothelial Cell | Rnls | 2 | 4.52E-06 | 1.25885016 | 0.727 | 0.174 | 0.00904322 |
| Lymphatic Endothelial Cell | Nkd2 | 2 | 4.58E-06 | 2.07766556 | 0.5 | 0.193 | 0.00915003 |
| Lymphatic Endothelial Cell | Cdon | 2 | 8.70E-06 | 1.3235869 | 0.636 | 0.055 | 0.01740374 |
| Lymphatic Endothelial Cell | Col14a1 | 2 | 9.52E-06 | 2.32535849 | 0.5 | 0.046 | 0.01904261 |

| | | | | | | | |
|---|---|---|---|---|---|---|---|
| Lymphatic Endothelial Cell | Kit | 2 | 1.07E-05 | 3.85341976 | 0.5 | 0.147 | 0.02146993 |
| Lymphatic Endothelial Cell | Trhde | 2 | 1.25E-05 | 4.84813706 | 0.455 | 0.018 | 0.02490723 |
| Lymphatic Endothelial Cell | Atp8b1 | 2 | 1.26E-05 | 2.96378675 | 0.864 | 0.495 | 0.02513673 |
| Lymphatic Endothelial Cell | Fabp4 | 2 | 1.26E-05 | 1.68497642 | 0.682 | 0.119 | 0.02520453 |
| Lymphatic Endothelial Cell | Ptprt | 2 | 1.55E-05 | 0.62965672 | 0.364 | 0.092 | 0.03097217 |
| Lymphatic Endothelial Cell | Bmp6 | 2 | 1.62E-05 | 0.50207455 | 0.727 | 0.183 | 0.03235937 |
| Lymphatic Endothelial Cell | Col1a1 | 2 | 2.45E-05 | 1.83857353 | 0.682 | 0.11 | 0.04891758 |
| Lymphatic Endothelial Cell | Adgrl3 | 2 | 2.45E-05 | 5.88393887 | 0.727 | 0.477 | 0.04892169 |
| Neuronal Cell | Cdh19 | 0 | 6.11E-08 | 1.55065739 | 0.875 | 0.564 | 0.00012219 |
| Neuronal Cell | RGD1306750 | 0 | 6.00E-07 | 4.43975748 | 0.475 | 0.091 | 0.00119961 |

| | | | | | | | |
|---|---|---|---|---|---|---|---|
| Neuronal Cell | Slc35f1 | 0 | 1.38E-05 | 1.58519255 | 0.8 | 0.473 | 0.02751322 |
| Neuronal Cell | Adgrl3 | 0 | 2.30E-05 | 1.6364685 | 0.863 | 0.6 | 0.0459728 |
| Neuronal Cell | Nav3 | 1 | 1.16E-07 | 2.62543819 | 0.745 | 0.225 | 0.00023295 |
| Neuronal Cell | Ablim3 | 1 | 1.44E-07 | 2.73845673 | 0.655 | 0.138 | 0.00028827 |
| Neuronal Cell | Adgrf5 | 1 | 2.94E-06 | 2.7954749 | 0.727 | 0.475 | 0.00587168 |
| Smooth Muscle Cell | Tp53inp2 | 0 | 1.51E-14 | 0.58339301 | 0.133 | 0.428 | 3.02E-11 |
| Smooth Muscle Cell | RGD1307916 | 2 | 1.34E-117 | 6.86658531 | 0.296 | 0.001 | 2.67E-114 |
| Smooth Muscle Cell | Grm3 | 2 | 4.09E-117 | 5.03980545 | 0.382 | 0.017 | 8.19E-114 |
| Smooth Muscle Cell | Edaradd | 2 | 1.03E-99 | 1.35646633 | 0.276 | 0.003 | 2.06E-96 |
| Smooth Muscle Cell | Irf4 | 2 | 3.76E-92 | 4.5314082 | 0.503 | 0.08 | 7.53E-89 |
| Smooth Muscle Cell | Myl4 | 2 | 2.81E-90 | 3.06328817 | 0.444 | 0.053 | 5.63E-87 |
| Smooth Muscle Cell | Khdrbs3 | 2 | 3.74E-79 | 1.07926261 | 0.434 | 0.058 | 7.49E-76 |
| Smooth Muscle Cell | LOC102553338 | 2 | 1.53E-73 | 1.55261846 | 0.434 | 0.043 | 3.07E-70 |
| Smooth Muscle Cell | AABR07002677.2 | 2 | 3.71E-72 | 2.50676636 | 0.368 | 0.045 | 7.42E-69 |

| | | | | | | | |
|---|---|---|---|---|---|---|---|
| Smooth Muscle Cell | LOC103690241 | 2 | 2.20E-67 | 1.06647062 | 0.5 | 0.12 | 4.40E-64 |
| Smooth Muscle Cell | Rad51 | 2 | 1.26E-66 | 3.01062672 | 0.592 | 0.189 | 2.52E-63 |
| Smooth Muscle Cell | Slc38a4 | 2 | 2.06E-65 | 3.00642214 | 0.438 | 0.042 | 4.12E-62 |
| Smooth Muscle Cell | Sell | 2 | 2.17E-63 | 0.92669191 | 0.309 | 0.033 | 4.34E-60 |
| Smooth Muscle Cell | Eya1 | 2 | 1.42E-61 | 2.21720366 | 0.533 | 0.099 | 2.83E-58 |
| Smooth Muscle Cell | Tmem132d | 2 | 1.92E-52 | 2.00249951 | 0.845 | 0.487 | 3.83E-49 |
| Smooth Muscle Cell | Myo3a | 2 | 3.65E-51 | 1.55662619 | 0.503 | 0.091 | 7.31E-48 |
| Smooth Muscle Cell | Abca9 | 2 | 9.85E-47 | 1.93124112 | 0.753 | 0.35 | 1.97E-43 |
| Smooth Muscle Cell | Mme | 2 | 3.25E-46 | 2.8483672 | 0.589 | 0.226 | 6.51E-43 |
| Smooth Muscle Cell | St6galnac1 | 2 | 7.64E-46 | 4.00466576 | 0.438 | 0.085 | 1.53E-42 |
| Smooth Muscle Cell | Bard1 | 2 | 3.19E-44 | 2.23632733 | 0.553 | 0.226 | 6.38E-41 |
| Smooth Muscle Cell | Kcnc2 | 2 | 7.24E-42 | 2.2917173 | 0.684 | 0.33 | 1.45E-38 |
| Smooth Muscle Cell | Wnt5b | 2 | 1.65E-39 | 0.47430994 | 0.497 | 0.162 | 3.30E-36 |
| Smooth Muscle Cell | Pde8b | 2 | 1.25E-38 | 2.59271453 | 0.536 | 0.128 | 2.50E-35 |

| | | | | | | | |
|---|---|---|---|---|---|---|---|
| Smooth Muscle Cell | Frem1 | 2 | 4.39E-38 | 1.56418796 | 0.839 | 0.558 | 8.77E-35 |
| Smooth Muscle Cell | Slco2b1 | 2 | 5.26E-37 | 1.4466956 | 0.806 | 0.506 | 1.05E-33 |
| Smooth Muscle Cell | Zdbf2 | 2 | 5.47E-37 | 1.61208516 | 0.352 | 0.074 | 1.09E-33 |
| Smooth Muscle Cell | Fmod | 2 | 1.47E-36 | 1.6870306 | 0.352 | 0.062 | 2.94E-33 |
| Smooth Muscle Cell | Prss48 | 2 | 1.74E-36 | 0.62892977 | 0.536 | 0.22 | 3.49E-33 |
| Smooth Muscle Cell | Xirp1 | 2 | 2.49E-36 | 2.18542337 | 0.322 | 0.055 | 4.98E-33 |
| Smooth Muscle Cell | Sphkap | 2 | 6.54E-36 | 2.38110542 | 0.576 | 0.185 | 1.31E-32 |
| Smooth Muscle Cell | Pcdh12 | 2 | 9.28E-36 | 1.01732616 | 0.48 | 0.175 | 1.86E-32 |
| Smooth Muscle Cell | Cacna1d | 2 | 3.05E-35 | 1.40697775 | 0.576 | 0.215 | 6.10E-32 |
| Smooth Muscle Cell | Brip1 | 2 | 1.03E-34 | 0.97944196 | 0.618 | 0.315 | 2.06E-31 |
| Smooth Muscle Cell | Aldh1a2 | 2 | 2.01E-34 | 1.67556164 | 0.638 | 0.258 | 4.02E-31 |
| Smooth Muscle Cell | Alox5 | 2 | 2.09E-34 | 1.01799556 | 0.362 | 0.096 | 4.19E-31 |
| Smooth Muscle Cell | Gria4 | 2 | 5.97E-34 | 1.38950909 | 0.711 | 0.397 | 1.19E-30 |
| Smooth Muscle Cell | Astn2 | 2 | 7.57E-34 | 1.45326826 | 0.707 | 0.402 | 1.51E-30 |

| | | | | | | | |
|---|---|---|---|---|---|---|---|
| Smooth Muscle Cell | Vash2 | 2 | 1.45E-33 | 2.38792241 | 0.477 | 0.09 | 2.91E-30 |
| Smooth Muscle Cell | Bmper | 2 | 1.90E-33 | 1.30063537 | 0.783 | 0.471 | 3.81E-30 |
| Smooth Muscle Cell | Ntf3 | 2 | 7.02E-33 | 1.64340579 | 0.684 | 0.346 | 1.40E-29 |
| Smooth Muscle Cell | Lepr | 2 | 1.78E-32 | 1.57416616 | 0.681 | 0.409 | 3.57E-29 |
| Smooth Muscle Cell | Gal3st3 | 2 | 3.71E-32 | 1.10010424 | 0.477 | 0.204 | 7.43E-29 |
| Smooth Muscle Cell | Ly49s6 | 2 | 5.55E-32 | 1.43129474 | 0.408 | 0.129 | 1.11E-28 |
| Smooth Muscle Cell | Mrvi1 | 2 | 5.83E-32 | 1.55354786 | 0.691 | 0.36 | 1.17E-28 |
| Smooth Muscle Cell | AABR07007068.1 | 2 | 2.49E-31 | 2.36203726 | 0.632 | 0.342 | 4.98E-28 |
| Smooth Muscle Cell | Eya2 | 2 | 2.54E-31 | 1.39822599 | 0.799 | 0.53 | 5.09E-28 |
| Smooth Muscle Cell | Sgip1 | 2 | 6.93E-31 | 1.30733736 | 0.776 | 0.49 | 1.39E-27 |
| Smooth Muscle Cell | Dusp27 | 2 | 8.33E-31 | 1.71240949 | 0.523 | 0.22 | 1.67E-27 |
| Smooth Muscle Cell | Nlrp3 | 2 | 2.65E-30 | 0.72147064 | 0.405 | 0.131 | 5.30E-27 |
| Smooth Muscle Cell | AABR07038986.1 | 2 | 3.68E-30 | 0.96320784 | 0.539 | 0.226 | 7.36E-27 |
| Smooth Muscle Cell | Dtnb | 2 | 4.83E-30 | 1.19380829 | 0.72 | 0.402 | 9.66E-27 |

| | | | | | | | |
|---|---|---|---|---|---|---|---|
| Smooth Muscle Cell | Ndst3 | 2 | 1.27E-29 | 0.87326618 | 0.648 | 0.377 | 2.54E-26 |
| Smooth Muscle Cell | Timp4 | 2 | 5.74E-29 | 0.43586222 | 0.5 | 0.171 | 1.15E-25 |
| Smooth Muscle Cell | Ano4 | 2 | 1.16E-28 | 1.61792674 | 0.559 | 0.282 | 2.32E-25 |
| Smooth Muscle Cell | Vwa5b1 | 2 | 1.30E-28 | 1.64486879 | 0.359 | 0.068 | 2.61E-25 |
| Smooth Muscle Cell | Cxcl14 | 2 | 1.71E-28 | 3.56932706 | 0.414 | 0.096 | 3.43E-25 |
| Smooth Muscle Cell | Oprd1 | 2 | 5.86E-28 | 1.60612012 | 0.576 | 0.322 | 1.17E-24 |
| Smooth Muscle Cell | Hs6st2 | 2 | 7.71E-28 | 1.25460186 | 0.684 | 0.337 | 1.54E-24 |
| Smooth Muscle Cell | AABR07025140.1 | 2 | 1.27E-27 | 0.65304318 | 0.576 | 0.239 | 2.54E-24 |
| Smooth Muscle Cell | Rec114 | 2 | 2.66E-27 | 1.25407845 | 0.582 | 0.32 | 5.32E-24 |
| Smooth Muscle Cell | AABR07050449.1 | 2 | 3.46E-27 | 1.38838237 | 0.52 | 0.192 | 6.91E-24 |
| Smooth Muscle Cell | Asic2 | 2 | 1.12E-26 | 1.75604717 | 0.592 | 0.241 | 2.23E-23 |
| Smooth Muscle Cell | Shc3 | 2 | 1.99E-26 | 1.79266806 | 0.562 | 0.25 | 3.98E-23 |
| Smooth Muscle Cell | Atp2b2 | 2 | 5.26E-26 | 0.4610157 | 0.431 | 0.167 | 1.05E-22 |
| Smooth Muscle Cell | Grid1 | 2 | 1.06E-25 | 1.1067998 | 0.428 | 0.151 | 2.11E-22 |

| | | | | | | | |
|---|---|---|---|---|---|---|---|
| Smooth Muscle Cell | Nostrin | 2 | 2.02E-25 | 0.38910449 | 0.461 | 0.106 | 4.03E-22 |
| Smooth Muscle Cell | LOC100910978.1 | 2 | 2.65E-25 | 1.05358484 | 0.697 | 0.301 | 5.30E-22 |
| Smooth Muscle Cell | Gpm6b | 2 | 6.14E-25 | 1.15114445 | 0.691 | 0.38 | 1.23E-21 |
| Smooth Muscle Cell | Hsf5 | 2 | 6.36E-25 | 0.33320914 | 0.569 | 0.201 | 1.27E-21 |
| Smooth Muscle Cell | Galnt15 | 2 | 7.98E-24 | 1.03028916 | 0.655 | 0.366 | 1.60E-20 |
| Smooth Muscle Cell | Ramp1 | 2 | 1.21E-23 | 1.12959525 | 0.53 | 0.275 | 2.42E-20 |
| Smooth Muscle Cell | AABR07003030.2 | 2 | 2.45E-23 | 1.80186069 | 0.536 | 0.214 | 4.91E-20 |
| Smooth Muscle Cell | Ror2 | 2 | 6.13E-23 | 0.65566818 | 0.671 | 0.321 | 1.23E-19 |
| Smooth Muscle Cell | Tafa5 | 2 | 1.42E-22 | 0.52836299 | 0.625 | 0.235 | 2.83E-19 |
| Smooth Muscle Cell | Trpc3 | 2 | 1.74E-22 | 0.74603156 | 0.582 | 0.246 | 3.48E-19 |
| Smooth Muscle Cell | Cgnl1 | 2 | 5.91E-22 | 0.32950432 | 0.599 | 0.294 | 1.18E-18 |
| Smooth Muscle Cell | Trim7 | 2 | 1.10E-21 | 1.1272204 | 0.562 | 0.297 | 2.21E-18 |
| Smooth Muscle Cell | Flnc | 2 | 1.39E-21 | 0.98159952 | 0.592 | 0.322 | 2.77E-18 |
| Smooth Muscle Cell | Kif21a | 2 | 1.57E-21 | 0.31073818 | 0.48 | 0.221 | 3.13E-18 |

| | | | | | | | |
|---|---|---|---|---|---|---|---|
| Smooth Muscle Cell | Zmat4 | 2 | 2.05E-21 | 1.57123772 | 0.464 | 0.17 | 4.10E-18 |
| Smooth Muscle Cell | Cst3 | 2 | 4.65E-21 | 0.8024657 | 0.543 | 0.237 | 9.31E-18 |
| Smooth Muscle Cell | AABR07026483.1 | 2 | 1.24E-20 | 1.65757067 | 0.365 | 0.065 | 2.48E-17 |
| Smooth Muscle Cell | Ankrd6 | 2 | 1.79E-20 | 0.36075018 | 0.586 | 0.289 | 3.59E-17 |
| Smooth Muscle Cell | Vegfd | 2 | 2.48E-20 | 0.78976614 | 0.641 | 0.304 | 4.95E-17 |
| Smooth Muscle Cell | Pdcd1lg2 | 2 | 1.56E-19 | 0.93459558 | 0.559 | 0.26 | 3.13E-16 |
| Smooth Muscle Cell | Gpm6a | 2 | 2.23E-19 | 0.92143249 | 0.809 | 0.542 | 4.46E-16 |
| Smooth Muscle Cell | Sox13 | 2 | 2.57E-19 | 0.72489151 | 0.605 | 0.253 | 5.14E-16 |
| Smooth Muscle Cell | Capn6 | 2 | 4.41E-19 | 1.37675873 | 0.444 | 0.147 | 8.82E-16 |
| Smooth Muscle Cell | Pip5k1b | 2 | 6.01E-19 | 0.3412204 | 0.562 | 0.293 | 1.20E-15 |
| Smooth Muscle Cell | Ccser1 | 2 | 1.12E-18 | 0.60854361 | 0.562 | 0.265 | 2.24E-15 |
| Smooth Muscle Cell | Xkr4 | 2 | 1.66E-18 | 0.56459252 | 0.582 | 0.303 | 3.33E-15 |
| Smooth Muscle Cell | Vegfc | 2 | 3.07E-18 | 0.7430591 | 0.651 | 0.326 | 6.13E-15 |
| Smooth Muscle Cell | Gbe1 | 2 | 3.67E-18 | 1.0109856 | 0.602 | 0.341 | 7.34E-15 |

| | | | | | | | |
|---|---|---|---|---|---|---|---|
| Smooth Muscle Cell | AABR07001054.2 | 2 | 4.89E-18 | 0.90102334 | 0.431 | 0.143 | 9.77E-15 |
| Smooth Muscle Cell | Trim54 | 2 | 5.61E-18 | 1.00092986 | 0.559 | 0.305 | 1.12E-14 |
| Smooth Muscle Cell | Cntfr | 2 | 5.69E-18 | 0.37171255 | 0.497 | 0.239 | 1.14E-14 |
| Smooth Muscle Cell | Dkk2 | 2 | 6.49E-18 | 0.84473868 | 0.576 | 0.279 | 1.30E-14 |
| Smooth Muscle Cell | Magi2 | 2 | 1.15E-17 | 0.53796503 | 0.684 | 0.384 | 2.31E-14 |
| Smooth Muscle Cell | Acsl1 | 2 | 1.69E-17 | 0.36208363 | 0.609 | 0.336 | 3.37E-14 |
| Smooth Muscle Cell | Casq2 | 2 | 5.04E-16 | 0.43570647 | 0.632 | 0.339 | 1.01E-12 |
| Smooth Muscle Cell | Naca | 2 | 8.19E-16 | 0.55814641 | 0.507 | 0.249 | 1.64E-12 |
| Smooth Muscle Cell | Rnf144b | 2 | 2.10E-15 | 0.66826501 | 0.507 | 0.254 | 4.21E-12 |
| Smooth Muscle Cell | Ddc | 2 | 4.12E-15 | 0.54344059 | 0.454 | 0.178 | 8.23E-12 |
| Smooth Muscle Cell | Csgalnact1 | 2 | 1.58E-14 | 0.34852553 | 0.648 | 0.37 | 3.16E-11 |
| Smooth Muscle Cell | Fndc1 | 2 | 1.61E-14 | 0.61214463 | 0.806 | 0.493 | 3.22E-11 |
| Smooth Muscle Cell | Slit2 | 2 | 6.54E-14 | 0.64501624 | 0.609 | 0.333 | 1.31E-10 |
| Smooth Muscle Cell | Cox6a2 | 2 | 3.16E-13 | 0.26147081 | 0.589 | 0.321 | 6.32E-10 |

| | | | | | | | |
|---|---|---|---|---|---|---|---|
| Smooth Muscle Cell | Angptl1 | 2 | 3.62E-13 | 0.44309425 | 0.559 | 0.304 | 7.23E-10 |
| Smooth Muscle Cell | Il34 | 2 | 3.75E-13 | 0.38261944 | 0.586 | 0.32 | 7.50E-10 |
| Smooth Muscle Cell | Nnt | 2 | 1.92E-12 | 0.51659756 | 0.612 | 0.362 | 3.84E-09 |
| Smooth Muscle Cell | Insc | 2 | 2.82E-12 | 0.84332184 | 0.576 | 0.321 | 5.64E-09 |
| Smooth Muscle Cell | Ppip5k1 | 2 | 2.90E-12 | 0.58511444 | 0.612 | 0.351 | 5.80E-09 |
| Smooth Muscle Cell | Tbc1d4 | 2 | 3.59E-12 | 0.30807375 | 0.576 | 0.315 | 7.18E-09 |
| Smooth Muscle Cell | Oxct1 | 2 | 4.77E-11 | 0.36948075 | 0.612 | 0.339 | 9.53E-08 |
| Smooth Muscle Cell | Rbm44 | 4 | 6.13E-94 | 5.66110757 | 0.37 | 0.022 | 1.23E-90 |
| Smooth Muscle Cell | Prkcq | 4 | 4.75E-72 | 1.55475451 | 0.457 | 0.049 | 9.51E-69 |
| Smooth Muscle Cell | Smc1b | 4 | 6.40E-71 | 3.96250358 | 0.409 | 0.042 | 1.28E-67 |
| Smooth Muscle Cell | Tmem236 | 4 | 2.91E-69 | 4.73572742 | 0.418 | 0.043 | 5.81E-66 |
| Smooth Muscle Cell | AABR07054716.1 | 4 | 8.17E-68 | 1.55133071 | 0.486 | 0.08 | 1.63E-64 |
| Smooth Muscle Cell | Cysltr1 | 4 | 1.18E-64 | 2.65635523 | 0.399 | 0.052 | 2.35E-61 |
| Smooth Muscle Cell | Nrg4 | 4 | 7.48E-62 | 2.55581369 | 0.399 | 0.053 | 1.50E-58 |

| | | | | | | | |
|---|---|---|---|---|---|---|---|
| Smooth Muscle Cell | AABR07057510.3 | 4 | 3.41E-56 | 2.20294597 | 0.462 | 0.092 | 6.82E-53 |
| Smooth Muscle Cell | AC127756.1 | 4 | 1.10E-55 | 1.49260751 | 0.49 | 0.109 | 2.20E-52 |
| Smooth Muscle Cell | Kif20a | 4 | 2.64E-46 | 1.37526935 | 0.365 | 0.061 | 5.27E-43 |
| Smooth Muscle Cell | Nckap1l | 4 | 1.83E-45 | 0.61804449 | 0.37 | 0.063 | 3.66E-42 |
| Smooth Muscle Cell | Atp6ap1l | 4 | 3.42E-44 | 2.93989754 | 0.365 | 0.02 | 6.84E-41 |
| Smooth Muscle Cell | Tacr1 | 4 | 1.38E-42 | 1.21803278 | 0.51 | 0.14 | 2.76E-39 |
| Smooth Muscle Cell | Stac3 | 4 | 9.82E-40 | 3.31210861 | 0.341 | 0.068 | 1.96E-36 |
| Smooth Muscle Cell | Irf41 | 4 | 8.76E-39 | 0.70483795 | 0.457 | 0.107 | 1.75E-35 |
| Smooth Muscle Cell | AABR07027925.1 | 4 | 3.93E-34 | 1.31387902 | 0.442 | 0.053 | 7.86E-31 |
| Smooth Muscle Cell | Col9a1 | 4 | 7.35E-33 | 0.62972681 | 0.322 | 0.067 | 1.47E-29 |
| Smooth Muscle Cell | LOC1036902411 | 4 | 1.33E-30 | 1.74629472 | 0.486 | 0.142 | 2.66E-27 |
| Smooth Muscle Cell | Serpine2 | 4 | 2.28E-30 | 1.62316349 | 0.774 | 0.509 | 4.55E-27 |
| Smooth Muscle Cell | Slc44a5 | 4 | 2.74E-30 | 2.45660995 | 0.644 | 0.252 | 5.48E-27 |
| Smooth Muscle Cell | Prkg2 | 4 | 1.36E-28 | 1.56150117 | 0.466 | 0.145 | 2.72E-25 |

| | | | | | | | |
|---|---|---|---|---|---|---|---|
| Smooth Muscle Cell | Kcnn4 | 4 | 2.14E-28 | 1.08838983 | 0.447 | 0.146 | 4.27E-25 |
| Smooth Muscle Cell | Egr3 | 4 | 4.39E-28 | 2.06506329 | 0.577 | 0.279 | 8.78E-25 |
| Smooth Muscle Cell | Dync1i1 | 4 | 9.14E-27 | 1.32897635 | 0.505 | 0.191 | 1.83E-23 |
| Smooth Muscle Cell | Mcm5 | 4 | 1.13E-24 | 1.57751652 | 0.514 | 0.239 | 2.25E-21 |
| Smooth Muscle Cell | Trmt9b | 4 | 2.22E-24 | 1.64034195 | 0.351 | 0.079 | 4.43E-21 |
| Smooth Muscle Cell | Sorcs1 | 4 | 1.34E-23 | 1.46197294 | 0.567 | 0.215 | 2.68E-20 |
| Smooth Muscle Cell | Gfra3 | 4 | 1.88E-23 | 1.04288723 | 0.452 | 0.154 | 3.76E-20 |
| Smooth Muscle Cell | Ube2ql1 | 4 | 1.98E-23 | 3.87122501 | 0.356 | 0.03 | 3.97E-20 |
| Smooth Muscle Cell | AC111804.2 | 4 | 2.26E-23 | 0.96753504 | 0.567 | 0.281 | 4.52E-20 |
| Smooth Muscle Cell | Tex22 | 4 | 6.34E-22 | 1.05901798 | 0.582 | 0.208 | 1.27E-18 |
| Smooth Muscle Cell | Sema3a | 4 | 3.28E-21 | 1.11475771 | 0.51 | 0.165 | 6.57E-18 |
| Smooth Muscle Cell | Taldo1 | 4 | 1.39E-20 | 1.16329003 | 0.567 | 0.199 | 2.78E-17 |
| Smooth Muscle Cell | Opcml | 4 | 1.47E-20 | 1.26939437 | 0.889 | 0.636 | 2.94E-17 |
| Smooth Muscle Cell | Nrk | 4 | 5.37E-20 | 2.26494832 | 0.548 | 0.185 | 1.07E-16 |

| | | | | | | | |
|---|---|---|---|---|---|---|---|
| Smooth Muscle Cell | Adamts14 | 4 | 1.19E-19 | 1.54330894 | 0.635 | 0.348 | 2.37E-16 |
| Smooth Muscle Cell | Tnc | 4 | 2.01E-19 | 1.49350235 | 0.476 | 0.154 | 4.02E-16 |
| Smooth Muscle Cell | LOC682419 | 4 | 2.92E-19 | 0.89071263 | 0.418 | 0.067 | 5.84E-16 |
| Smooth Muscle Cell | Sgpp2 | 4 | 4.02E-19 | 0.70352544 | 0.452 | 0.189 | 8.04E-16 |
| Smooth Muscle Cell | Prrg4 | 4 | 7.45E-19 | 1.66060312 | 0.582 | 0.295 | 1.49E-15 |
| Smooth Muscle Cell | Klhl4 | 4 | 2.15E-18 | 0.87408437 | 0.577 | 0.323 | 4.30E-15 |
| Smooth Muscle Cell | Rnf150 | 4 | 3.01E-18 | 1.36032369 | 0.692 | 0.363 | 6.01E-15 |
| Smooth Muscle Cell | Cdkn1a | 4 | 5.36E-18 | 1.23155506 | 0.514 | 0.239 | 1.07E-14 |
| Smooth Muscle Cell | Chst15 | 4 | 5.38E-18 | 0.38936158 | 0.471 | 0.19 | 1.08E-14 |
| Smooth Muscle Cell | Adhfe1 | 4 | 5.72E-18 | 0.26915177 | 0.606 | 0.32 | 1.14E-14 |
| Smooth Muscle Cell | Ano41 | 4 | 8.35E-18 | 1.61182402 | 0.611 | 0.291 | 1.67E-14 |
| Smooth Muscle Cell | Dgkg | 4 | 1.29E-17 | 0.76774431 | 0.438 | 0.147 | 2.57E-14 |
| Smooth Muscle Cell | Adamts17 | 4 | 1.51E-17 | 1.17311361 | 0.726 | 0.453 | 3.01E-14 |
| Smooth Muscle Cell | Kif26b | 4 | 2.07E-17 | 0.7640217 | 0.548 | 0.226 | 4.14E-14 |

| | | | | | | | |
|---|---|---|---|---|---|---|---|
| Smooth Muscle Cell | Tbc1d9 | 4 | 3.07E-17 | 1.10297631 | 0.534 | 0.241 | 6.13E-14 |
| Smooth Muscle Cell | Unc5b | 4 | 3.77E-17 | 1.24492177 | 0.76 | 0.485 | 7.54E-14 |
| Smooth Muscle Cell | AABR07001054.21 | 4 | 9.83E-17 | 0.75907278 | 0.524 | 0.148 | 1.97E-13 |
| Smooth Muscle Cell | Sncaip | 4 | 1.26E-16 | 1.22603235 | 0.553 | 0.261 | 2.52E-13 |
| Smooth Muscle Cell | Trim50 | 4 | 2.05E-16 | 0.44651718 | 0.389 | 0.034 | 4.10E-13 |
| Smooth Muscle Cell | Cmahp | 4 | 2.15E-16 | 0.44463713 | 0.466 | 0.173 | 4.30E-13 |
| Smooth Muscle Cell | Rem1 | 4 | 3.16E-16 | 1.19450529 | 0.697 | 0.423 | 6.32E-13 |
| Smooth Muscle Cell | AABR07017268.1 | 4 | 3.74E-16 | 1.22555184 | 0.466 | 0.21 | 7.47E-13 |
| Smooth Muscle Cell | Ablim2 | 4 | 4.06E-16 | 0.64848038 | 0.466 | 0.193 | 8.13E-13 |
| Smooth Muscle Cell | Pamr1 | 4 | 6.45E-16 | 1.67986476 | 0.572 | 0.246 | 1.29E-12 |
| Smooth Muscle Cell | Maoa | 4 | 7.40E-16 | 0.80147086 | 0.635 | 0.336 | 1.48E-12 |
| Smooth Muscle Cell | Mbp | 4 | 1.26E-15 | 0.4009116 | 0.606 | 0.301 | 2.51E-12 |
| Smooth Muscle Cell | Sfxn5 | 4 | 1.29E-15 | 0.32953931 | 0.548 | 0.243 | 2.58E-12 |
| Smooth Muscle Cell | Prkaa2 | 4 | 1.52E-15 | 1.09030889 | 0.519 | 0.173 | 3.04E-12 |

| | | | | | | | |
|---|---|---|---|---|---|---|---|
| Smooth Muscle Cell | Lyve1 | 4 | 1.70E-15 | 2.2333661 | 0.543 | 0.218 | 3.39E-12 |
| Smooth Muscle Cell | Rp1 | 4 | 2.53E-15 | 0.93699197 | 0.462 | 0.171 | 5.07E-12 |
| Smooth Muscle Cell | Tex14 | 4 | 6.18E-15 | 1.11561613 | 0.587 | 0.325 | 1.24E-11 |
| Smooth Muscle Cell | Coq8a | 4 | 6.21E-15 | 0.64070107 | 0.505 | 0.23 | 1.24E-11 |
| Smooth Muscle Cell | Gbp6 | 4 | 9.78E-15 | 1.34386979 | 0.514 | 0.23 | 1.96E-11 |
| Smooth Muscle Cell | Alpk2 | 4 | 1.60E-14 | 0.34846157 | 0.514 | 0.222 | 3.20E-11 |
| Smooth Muscle Cell | Ntrk2 | 4 | 1.85E-14 | 0.65460553 | 0.558 | 0.249 | 3.71E-11 |
| Smooth Muscle Cell | Meox1 | 4 | 2.49E-14 | 0.55224062 | 0.481 | 0.21 | 4.99E-11 |
| Smooth Muscle Cell | Cxcl1 | 4 | 2.64E-14 | 0.82840662 | 0.587 | 0.322 | 5.28E-11 |
| Smooth Muscle Cell | Rnf157 | 4 | 3.01E-14 | 1.06259012 | 0.587 | 0.297 | 6.01E-11 |
| Smooth Muscle Cell | Atf3 | 4 | 3.11E-14 | 1.33739859 | 0.341 | 0.085 | 6.22E-11 |
| Smooth Muscle Cell | Veph1 | 4 | 4.45E-14 | 1.66067529 | 0.447 | 0.173 | 8.91E-11 |
| Smooth Muscle Cell | Gask1b | 4 | 5.99E-14 | 0.91689995 | 0.779 | 0.52 | 1.20E-10 |
| Smooth Muscle Cell | Capn61 | 4 | 1.81E-13 | 1.03692243 | 0.428 | 0.165 | 3.62E-10 |

| | | | | | | | |
|---|---|---|---|---|---|---|---|
| Smooth Muscle Cell | Ednra | 4 | 2.19E-13 | 0.8566757 | 0.63 | 0.366 | 4.38E-10 |
| Smooth Muscle Cell | Reep1 | 4 | 2.28E-13 | 0.66540728 | 0.514 | 0.222 | 4.56E-10 |
| Smooth Muscle Cell | Ppm1l | 4 | 3.43E-13 | 0.26707257 | 0.51 | 0.193 | 6.87E-10 |
| Smooth Muscle Cell | Pgf | 4 | 4.66E-13 | 1.54325848 | 0.337 | 0.057 | 9.32E-10 |
| Smooth Muscle Cell | Tmem26 | 4 | 7.21E-13 | 0.53886726 | 0.548 | 0.25 | 1.44E-09 |
| Smooth Muscle Cell | Lgr6 | 4 | 1.35E-12 | 0.93288192 | 0.582 | 0.291 | 2.71E-09 |
| Smooth Muscle Cell | Popdc2 | 4 | 1.88E-12 | 0.27253013 | 0.558 | 0.227 | 3.75E-09 |
| Smooth Muscle Cell | Fsd2 | 4 | 5.24E-12 | 0.68603127 | 0.524 | 0.243 | 1.05E-08 |
| Smooth Muscle Cell | Bmp2 | 4 | 7.94E-12 | 2.00711205 | 0.385 | 0.09 | 1.59E-08 |
| Smooth Muscle Cell | Rhobtb1 | 4 | 8.41E-12 | 0.65287122 | 0.587 | 0.303 | 1.68E-08 |
| Smooth Muscle Cell | Gucy1a1 | 4 | 1.47E-11 | 0.93241535 | 0.688 | 0.435 | 2.95E-08 |
| Smooth Muscle Cell | Fhl2 | 4 | 1.75E-11 | 0.33866191 | 0.63 | 0.322 | 3.51E-08 |
| Smooth Muscle Cell | Cacna1a | 4 | 2.07E-11 | 0.697978 | 0.692 | 0.43 | 4.15E-08 |
| Smooth Muscle Cell | Brca1 | 4 | 5.25E-11 | 0.65486404 | 0.567 | 0.292 | 1.05E-07 |

| | | | | | | | |
|---|---|---|---|---|---|---|---|
| Smooth Muscle Cell | Aqp1 | 4 | 8.43E-11 | 0.46604228 | 0.519 | 0.245 | 1.69E-07 |
| Smooth Muscle Cell | Shroom4 | 4 | 9.88E-11 | 0.41451972 | 0.649 | 0.361 | 1.98E-07 |
| Smooth Muscle Cell | Iqub | 4 | 1.48E-10 | 0.51857032 | 0.486 | 0.21 | 2.97E-07 |
| Smooth Muscle Cell | Sh3kbp1 | 4 | 3.46E-10 | 0.6041758 | 0.644 | 0.367 | 6.93E-07 |
| Smooth Muscle Cell | Actn1 | 4 | 7.90E-10 | 0.33770755 | 0.668 | 0.412 | 1.58E-06 |
| Smooth Muscle Cell | Tenm3 | 4 | 1.50E-09 | 0.6920596 | 0.663 | 0.41 | 3.00E-06 |
| Smooth Muscle Cell | Flt1 | 4 | 2.40E-09 | 0.273697 | 0.582 | 0.319 | 4.81E-06 |
| Smooth Muscle Cell | Rasgef1c | 4 | 5.28E-09 | 0.49642683 | 0.389 | 0.133 | 1.06E-05 |
| Smooth Muscle Cell | AABR07001519.1 | 4 | 1.70E-08 | 0.61258777 | 0.702 | 0.446 | 3.40E-05 |
| Smooth Muscle Cell | Ddc1 | 4 | 1.84E-08 | 0.56351937 | 0.447 | 0.193 | 3.67E-05 |
| Smooth Muscle Cell | Slc20a2 | 4 | 2.54E-08 | 0.35265407 | 0.587 | 0.298 | 5.09E-05 |
| Smooth Muscle Cell | Adk | 4 | 9.00E-08 | 0.37591141 | 0.678 | 0.405 | 0.00018003 |
| Smooth Muscle Cell | Rbm20 | 4 | 5.26E-06 | 0.61746467 | 0.683 | 0.41 | 0.01052087 |
| Smooth Muscle Cell | Vstm5 | 5 | 1.42E-199 | 4.64494248 | 0.51 | 0.004 | 2.85E-196 |

| | | | | | | | |
|---|---|---|---|---|---|---|---|
| Smooth Muscle Cell | Clspn | 5 | 1.42E-141 | 4.6356655 | 0.495 | 0.025 | 2.84E-138 |
| Smooth Muscle Cell | Adra2b | 5 | 1.86E-133 | 6.3382212 | 0.47 | 0.021 | 3.71E-130 |
| Smooth Muscle Cell | Reg4 | 5 | 2.85E-119 | 6.3559219 | 0.3 | 0.002 | 5.71E-116 |
| Smooth Muscle Cell | LOC691995 | 5 | 1.38E-116 | 1.90235479 | 0.47 | 0.029 | 2.77E-113 |
| Smooth Muscle Cell | Gria2 | 5 | 3.68E-116 | 3.87123869 | 0.435 | 0.037 | 7.36E-113 |
| Smooth Muscle Cell | Slc22a2 | 5 | 2.83E-113 | 7.40052715 | 0.31 | 0.004 | 5.65E-110 |
| Smooth Muscle Cell | AABR07065282.1 | 5 | 5.64E-112 | 6.33130215 | 0.43 | 0.024 | 1.13E-108 |
| Smooth Muscle Cell | Galnt14 | 5 | 1.53E-110 | 1.73811614 | 0.55 | 0.052 | 3.06E-107 |
| Smooth Muscle Cell | AABR07032787.1 | 5 | 3.57E-105 | 3.96424542 | 0.34 | 0.011 | 7.13E-102 |
| Smooth Muscle Cell | Zc3h12d | 5 | 7.13E-101 | 4.0588147 | 0.54 | 0.051 | 1.43E-97 |
| Smooth Muscle Cell | Abo | 5 | 9.66E-101 | 1.55572161 | 0.525 | 0.025 | 1.93E-97 |
| Smooth Muscle Cell | Rasgrf1 | 5 | 2.45E-99 | 1.98671723 | 0.55 | 0.054 | 4.91E-96 |
| Smooth Muscle Cell | AC118957.1 | 5 | 9.17E-98 | 0.65000228 | 0.49 | 0.025 | 1.83E-94 |
| Smooth Muscle Cell | Cpa6 | 5 | 3.15E-96 | 2.79640567 | 0.565 | 0.078 | 6.31E-93 |

| | | | | | | | |
|---|---|---|---|---|---|---|---|
| Smooth Muscle Cell | AC128789.1 | 5 | 1.75E-95 | 5.71698006 | 0.385 | 0.023 | 3.49E-92 |
| Smooth Muscle Cell | Csf3r | 5 | 2.25E-94 | 3.17804603 | 0.385 | 0.024 | 4.50E-91 |
| Smooth Muscle Cell | Kcp | 5 | 7.45E-79 | 3.59307166 | 0.31 | 0.007 | 1.49E-75 |
| Smooth Muscle Cell | Stk31 | 5 | 1.16E-74 | 0.76655796 | 0.47 | 0.062 | 2.33E-71 |
| Smooth Muscle Cell | Ikzf3 | 5 | 6.50E-61 | 1.88515737 | 0.575 | 0.069 | 1.30E-57 |
| Smooth Muscle Cell | Msr1 | 5 | 1.34E-57 | 2.03521471 | 0.295 | 0.017 | 2.68E-54 |
| Smooth Muscle Cell | Epha3 | 5 | 2.76E-57 | 3.52057947 | 0.63 | 0.142 | 5.53E-54 |
| Smooth Muscle Cell | Ceacam16 | 5 | 8.79E-56 | 5.17638454 | 0.34 | 0.011 | 1.76E-52 |
| Smooth Muscle Cell | Cdh6 | 5 | 4.53E-52 | 1.99902682 | 0.38 | 0.018 | 9.07E-49 |
| Smooth Muscle Cell | Lilrb4 | 5 | 3.23E-51 | 0.51159615 | 0.515 | 0.027 | 6.45E-48 |
| Smooth Muscle Cell | Cacna1b | 5 | 6.02E-50 | 2.12612601 | 0.64 | 0.214 | 1.20E-46 |
| Smooth Muscle Cell | Myo5b | 5 | 1.18E-49 | 1.75532746 | 0.48 | 0.09 | 2.36E-46 |
| Smooth Muscle Cell | Hapln3 | 5 | 5.22E-48 | 3.85815839 | 0.41 | 0.015 | 1.04E-44 |
| Smooth Muscle Cell | Thsd7b | 5 | 2.34E-46 | 2.64760767 | 0.315 | 0.063 | 4.68E-43 |

| | | | | | | | |
|---|---|---|---|---|---|---|---|
| Smooth Muscle Cell | Ccn5 | 5 | 1.94E-45 | 3.86872707 | 0.355 | 0.088 | 3.88E-42 |
| Smooth Muscle Cell | Arnt2 | 5 | 3.40E-44 | 4.1709409 | 0.385 | 0.057 | 6.80E-41 |
| Smooth Muscle Cell | S100a4 | 5 | 4.30E-44 | 1.61870817 | 0.59 | 0.216 | 8.60E-41 |
| Smooth Muscle Cell | St14 | 5 | 3.54E-42 | 1.63810994 | 0.425 | 0.056 | 7.09E-39 |
| Smooth Muscle Cell | Fabp12 | 5 | 3.70E-42 | 2.62588 | 0.4 | 0.028 | 7.40E-39 |
| Smooth Muscle Cell | Tnfrsf11b | 5 | 5.29E-42 | 0.93483607 | 0.585 | 0.101 | 1.06E-38 |
| Smooth Muscle Cell | Mctp2 | 5 | 1.33E-41 | 1.92040312 | 0.465 | 0.109 | 2.66E-38 |
| Smooth Muscle Cell | LOC100911486 | 5 | 1.49E-41 | 3.34883964 | 0.375 | 0.122 | 2.99E-38 |
| Smooth Muscle Cell | Ky | 5 | 2.74E-39 | 2.72031854 | 0.55 | 0.119 | 5.49E-36 |
| Smooth Muscle Cell | Tshr | 5 | 4.45E-39 | 3.63609926 | 0.375 | 0.021 | 8.90E-36 |
| Smooth Muscle Cell | Col6a5 | 5 | 2.54E-38 | 0.82265293 | 0.365 | 0.1 | 5.08E-35 |
| Smooth Muscle Cell | LOC103693323 | 5 | 4.50E-38 | 2.13430383 | 0.435 | 0.131 | 8.99E-35 |
| Smooth Muscle Cell | Rtn4rl1 | 5 | 4.63E-37 | 1.07113206 | 0.65 | 0.239 | 9.26E-34 |
| Smooth Muscle Cell | Frmd3 | 5 | 1.36E-36 | 1.76447822 | 0.515 | 0.104 | 2.73E-33 |

| | | | | | | | |
|---|---|---|---|---|---|---|---|
| Smooth Muscle Cell | Unc5c | 5 | 1.58E-36 | 1.18028857 | 0.41 | 0.085 | 3.17E-33 |
| Smooth Muscle Cell | Gvin1 | 5 | 1.35E-35 | 1.64164388 | 0.545 | 0.111 | 2.71E-32 |
| Smooth Muscle Cell | Itgb2 | 5 | 3.57E-35 | 1.61915158 | 0.48 | 0.128 | 7.13E-32 |
| Smooth Muscle Cell | Cnksr2 | 5 | 8.17E-34 | 0.62324301 | 0.545 | 0.174 | 1.63E-30 |
| Smooth Muscle Cell | Ms4a6bl | 5 | 1.38E-33 | 2.12567866 | 0.4 | 0.04 | 2.77E-30 |
| Smooth Muscle Cell | Mcm6 | 5 | 1.96E-33 | 1.41229677 | 0.31 | 0.037 | 3.92E-30 |
| Smooth Muscle Cell | Rbp7 | 5 | 1.99E-33 | 3.23476065 | 0.38 | 0.039 | 3.98E-30 |
| Smooth Muscle Cell | AABR07031740.1 | 5 | 3.14E-33 | 1.65327113 | 0.455 | 0.065 | 6.27E-30 |
| Smooth Muscle Cell | Plbd1 | 5 | 1.25E-32 | 2.11066822 | 0.375 | 0.109 | 2.49E-29 |
| Smooth Muscle Cell | Mt1 | 5 | 3.78E-32 | 1.70332255 | 0.415 | 0.062 | 7.55E-29 |
| Smooth Muscle Cell | Icam1 | 5 | 4.24E-32 | 0.89010637 | 0.61 | 0.219 | 8.49E-29 |
| Smooth Muscle Cell | Adamts15 | 5 | 1.65E-31 | 2.52822917 | 0.605 | 0.261 | 3.31E-28 |
| Smooth Muscle Cell | Kntc1 | 5 | 1.95E-31 | 0.30087302 | 0.535 | 0.155 | 3.91E-28 |
| Smooth Muscle Cell | Mx1 | 5 | 5.88E-31 | 2.32092507 | 0.395 | 0.109 | 1.18E-27 |

| | | | | | | | |
|---|---|---|---|---|---|---|---|
| Smooth Muscle Cell | Plcxd3 | 5 | 1.60E-30 | 3.43182945 | 0.56 | 0.26 | 3.20E-27 |
| Smooth Muscle Cell | Nkd1 | 5 | 5.08E-29 | 2.80299174 | 0.655 | 0.309 | 1.02E-25 |
| Smooth Muscle Cell | Tmem17 | 5 | 5.18E-29 | 3.05273448 | 0.645 | 0.27 | 1.04E-25 |
| Smooth Muscle Cell | Gpnmb | 5 | 1.68E-28 | 2.03209532 | 0.45 | 0.15 | 3.35E-25 |
| Smooth Muscle Cell | Asb15 | 5 | 2.54E-28 | 1.55421793 | 0.425 | 0.08 | 5.07E-25 |
| Smooth Muscle Cell | Rcan2 | 5 | 4.71E-28 | 1.42097991 | 0.81 | 0.481 | 9.43E-25 |
| Smooth Muscle Cell | Prodh1 | 5 | 1.42E-27 | 0.97742558 | 0.675 | 0.337 | 2.84E-24 |
| Smooth Muscle Cell | Sod2 | 5 | 1.80E-27 | 0.61833328 | 0.635 | 0.342 | 3.60E-24 |
| Smooth Muscle Cell | Cyp26b1 | 5 | 2.51E-27 | 0.81806228 | 0.445 | 0.131 | 5.01E-24 |
| Smooth Muscle Cell | Nkain3 | 5 | 5.35E-27 | 0.35665376 | 0.54 | 0.174 | 1.07E-23 |
| Smooth Muscle Cell | Apoo | 5 | 5.63E-27 | 1.08418472 | 0.65 | 0.198 | 1.13E-23 |
| Smooth Muscle Cell | Gabrb1 | 5 | 3.03E-26 | 2.80036045 | 0.325 | 0.075 | 6.06E-23 |
| Smooth Muscle Cell | Jag2 | 5 | 3.09E-26 | 0.76318606 | 0.55 | 0.258 | 6.19E-23 |
| Smooth Muscle Cell | Grin3a | 5 | 3.27E-26 | 1.72119375 | 0.465 | 0.182 | 6.54E-23 |

| | | | | | | | |
|---|---|---|---|---|---|---|---|
| Smooth Muscle Cell | Tec | 5 | 3.73E-26 | 0.8350214 | 0.455 | 0.131 | 7.46E-23 |
| Smooth Muscle Cell | Rbm24 | 5 | 4.16E-26 | 0.29168273 | 0.43 | 0.09 | 8.32E-23 |
| Smooth Muscle Cell | Cav1 | 5 | 7.74E-26 | 1.07402032 | 0.695 | 0.363 | 1.55E-22 |
| Smooth Muscle Cell | Lmod1 | 5 | 1.14E-25 | 2.2393188 | 0.63 | 0.321 | 2.28E-22 |
| Smooth Muscle Cell | Ptpro | 5 | 1.89E-25 | 0.82619188 | 0.575 | 0.281 | 3.78E-22 |
| Smooth Muscle Cell | Scube3 | 5 | 1.07E-24 | 0.77527605 | 0.565 | 0.248 | 2.13E-21 |
| Smooth Muscle Cell | Mctp1 | 5 | 1.58E-24 | 1.0610763 | 0.675 | 0.31 | 3.17E-21 |
| Smooth Muscle Cell | Efna5 | 5 | 1.84E-24 | 0.25866838 | 0.645 | 0.308 | 3.68E-21 |
| Smooth Muscle Cell | AABR07003304.2 | 5 | 2.11E-24 | 1.19252001 | 0.58 | 0.308 | 4.23E-21 |
| Smooth Muscle Cell | Cabcoco1 | 5 | 5.85E-24 | 1.77018397 | 0.485 | 0.092 | 1.17E-20 |
| Smooth Muscle Cell | Ppp2r2b | 5 | 6.23E-24 | 0.92435065 | 0.705 | 0.328 | 1.25E-20 |
| Smooth Muscle Cell | P2rx7 | 5 | 6.35E-24 | 1.01328005 | 0.6 | 0.228 | 1.27E-20 |
| Smooth Muscle Cell | Hs3st1 | 5 | 2.67E-23 | 0.83879429 | 0.655 | 0.3 | 5.34E-20 |
| Smooth Muscle Cell | Cdh19 | 5 | 7.01E-23 | 1.27005135 | 0.51 | 0.136 | 1.40E-19 |

| | | | | | | | |
|---|---|---|---|---|---|---|---|
| Smooth Muscle Cell | Tenm4 | 5 | 9.02E-23 | 1.42760207 | 0.47 | 0.187 | 1.80E-19 |
| Smooth Muscle Cell | Slc16a10 | 5 | 9.37E-23 | 2.04258496 | 0.47 | 0.173 | 1.87E-19 |
| Smooth Muscle Cell | Mertk | 5 | 1.54E-22 | 1.60103921 | 0.465 | 0.144 | 3.08E-19 |
| Smooth Muscle Cell | Adamtsl2 | 5 | 1.84E-22 | 1.53194701 | 0.755 | 0.466 | 3.68E-19 |
| Smooth Muscle Cell | Gfra31 | 5 | 3.54E-22 | 0.34700782 | 0.42 | 0.159 | 7.08E-19 |
| Smooth Muscle Cell | Slc38a3 | 5 | 8.04E-22 | 1.37934849 | 0.39 | 0.137 | 1.61E-18 |
| Smooth Muscle Cell | Akr1c15 | 5 | 8.53E-22 | 1.89369432 | 0.58 | 0.238 | 1.71E-18 |
| Smooth Muscle Cell | Abcc8 | 5 | 8.73E-22 | 1.99520501 | 0.45 | 0.114 | 1.75E-18 |
| Smooth Muscle Cell | Pfkfb3 | 5 | 1.04E-21 | 0.96407236 | 0.645 | 0.299 | 2.07E-18 |
| Smooth Muscle Cell | Slc9a3r2 | 5 | 2.52E-21 | 0.37564192 | 0.665 | 0.266 | 5.03E-18 |
| Smooth Muscle Cell | Dlg2 | 5 | 2.90E-21 | 1.36951541 | 0.815 | 0.563 | 5.80E-18 |
| Smooth Muscle Cell | Ifitm10 | 5 | 3.21E-21 | 1.08371372 | 0.58 | 0.328 | 6.43E-18 |
| Smooth Muscle Cell | Coro6 | 5 | 3.69E-21 | 0.96356956 | 0.56 | 0.152 | 7.37E-18 |
| Smooth Muscle Cell | Ano5 | 5 | 4.21E-21 | 1.03283654 | 0.49 | 0.167 | 8.43E-18 |

| | | | | | | | |
|---|---|---|---|---|---|---|---|
| Smooth Muscle Cell | Lamc3 | 5 | 1.08E-20 | 3.08771607 | 0.43 | 0.173 | 2.15E-17 |
| Smooth Muscle Cell | Gja1 | 5 | 1.56E-20 | 1.10235644 | 0.57 | 0.214 | 3.12E-17 |
| Smooth Muscle Cell | Hmgcll1 | 5 | 2.34E-20 | 0.634398 | 0.635 | 0.342 | 4.68E-17 |
| Smooth Muscle Cell | Ablim21 | 5 | 2.70E-20 | 1.03886454 | 0.57 | 0.183 | 5.40E-17 |
| Smooth Muscle Cell | Dmpk | 5 | 2.89E-20 | 1.03756876 | 0.66 | 0.284 | 5.78E-17 |
| Smooth Muscle Cell | Nav2 | 5 | 3.25E-20 | 0.80284448 | 0.625 | 0.262 | 6.49E-17 |
| Smooth Muscle Cell | Acyp2 | 5 | 3.99E-20 | 0.89389244 | 0.74 | 0.417 | 7.98E-17 |
| Smooth Muscle Cell | Rcan1 | 5 | 4.65E-20 | 1.20346042 | 0.49 | 0.221 | 9.30E-17 |
| Smooth Muscle Cell | Srl | 5 | 5.43E-20 | 0.58541088 | 0.395 | 0.103 | 1.09E-16 |
| Smooth Muscle Cell | Frmd5 | 5 | 1.15E-19 | 0.4518096 | 0.525 | 0.188 | 2.29E-16 |
| Smooth Muscle Cell | Gjc1 | 5 | 1.16E-19 | 0.45323375 | 0.63 | 0.307 | 2.33E-16 |
| Smooth Muscle Cell | Olfml2b | 5 | 1.19E-19 | 1.21006179 | 0.515 | 0.245 | 2.39E-16 |
| Smooth Muscle Cell | Sgcg | 5 | 1.24E-19 | 0.78278481 | 0.625 | 0.332 | 2.48E-16 |
| Smooth Muscle Cell | Efnb2 | 5 | 2.85E-19 | 0.9625042 | 0.615 | 0.338 | 5.70E-16 |

| | | | | | | | |
|---|---|---|---|---|---|---|---|
| Smooth Muscle Cell | Rnf207 | 5 | 2.92E-19 | 0.96434683 | 0.525 | 0.207 | 5.84E-16 |
| Smooth Muscle Cell | Pola2 | 5 | 2.94E-19 | 0.50406055 | 0.535 | 0.214 | 5.88E-16 |
| Smooth Muscle Cell | Zfp366 | 5 | 2.98E-19 | 0.70341111 | 0.61 | 0.311 | 5.96E-16 |
| Smooth Muscle Cell | Lrsam1 | 5 | 3.86E-19 | 1.16808928 | 0.56 | 0.259 | 7.72E-16 |
| Smooth Muscle Cell | Alox5ap | 5 | 1.66E-18 | 0.44004368 | 0.425 | 0.175 | 3.32E-15 |
| Smooth Muscle Cell | Plcb1 | 5 | 2.90E-18 | 0.98393787 | 0.56 | 0.256 | 5.80E-15 |
| Smooth Muscle Cell | Nrp2 | 5 | 3.03E-18 | 1.72507244 | 0.535 | 0.252 | 6.05E-15 |
| Smooth Muscle Cell | Slc39a8 | 5 | 3.73E-18 | 0.72780475 | 0.55 | 0.298 | 7.47E-15 |
| Smooth Muscle Cell | Stk32b | 5 | 5.01E-18 | 0.53434217 | 0.665 | 0.357 | 1.00E-14 |
| Smooth Muscle Cell | Tmem168 | 5 | 7.37E-18 | 0.25416904 | 0.52 | 0.258 | 1.47E-14 |
| Smooth Muscle Cell | Tox3 | 5 | 9.55E-18 | 1.00097942 | 0.51 | 0.255 | 1.91E-14 |
| Smooth Muscle Cell | AABR07040864.1 | 5 | 1.26E-17 | 0.85197094 | 0.675 | 0.289 | 2.52E-14 |
| Smooth Muscle Cell | RGD1564053 | 5 | 1.71E-17 | 1.84235788 | 0.405 | 0.063 | 3.43E-14 |
| Smooth Muscle Cell | Pappa1 | 5 | 3.22E-17 | 0.46749305 | 0.67 | 0.366 | 6.44E-14 |

| | | | | | | | |
|---|---|---|---|---|---|---|---|
| Smooth Muscle Cell | Rnf213 | 5 | 9.41E-17 | 1.05182994 | 0.715 | 0.368 | 1.88E-13 |
| Smooth Muscle Cell | Rasa4 | 5 | 1.43E-16 | 0.77585532 | 0.55 | 0.29 | 2.87E-13 |
| Smooth Muscle Cell | Fam151a | 5 | 1.54E-16 | 0.80040352 | 0.405 | 0.148 | 3.08E-13 |
| Smooth Muscle Cell | Trdn | 5 | 1.67E-16 | 0.81710652 | 0.58 | 0.265 | 3.33E-13 |
| Smooth Muscle Cell | Arhgap44 | 5 | 2.51E-16 | 1.2770351 | 0.515 | 0.205 | 5.02E-13 |
| Smooth Muscle Cell | Slc25a21 | 5 | 2.99E-16 | 1.01812739 | 0.62 | 0.366 | 5.98E-13 |
| Smooth Muscle Cell | Apoe | 5 | 3.56E-16 | 0.94895659 | 0.56 | 0.278 | 7.11E-13 |
| Smooth Muscle Cell | Thsd7a | 5 | 3.61E-16 | 0.76169475 | 0.565 | 0.245 | 7.21E-13 |
| Smooth Muscle Cell | Pfkfb2 | 5 | 4.87E-16 | 0.4216839 | 0.595 | 0.295 | 9.75E-13 |
| Smooth Muscle Cell | Fgf10 | 5 | 6.30E-16 | 0.68135562 | 0.66 | 0.361 | 1.26E-12 |
| Smooth Muscle Cell | Scn5a | 5 | 6.78E-16 | 0.57447432 | 0.63 | 0.38 | 1.36E-12 |
| Smooth Muscle Cell | Spns2 | 5 | 7.14E-16 | 0.36990924 | 0.535 | 0.261 | 1.43E-12 |
| Smooth Muscle Cell | Twf2 | 5 | 1.36E-15 | 0.29036309 | 0.64 | 0.365 | 2.72E-12 |
| Smooth Muscle Cell | Itgb8 | 5 | 1.55E-15 | 1.2764086 | 0.62 | 0.344 | 3.11E-12 |

| | | | | | | | |
|---|---|---|---|---|---|---|---|
| Smooth Muscle Cell | Gpc3 | 5 | 2.16E-15 | 0.60470307 | 0.48 | 0.207 | 4.33E-12 |
| Smooth Muscle Cell | Flnb | 5 | 2.65E-15 | 0.75852569 | 0.68 | 0.341 | 5.29E-12 |
| Smooth Muscle Cell | Iqub1 | 5 | 8.47E-15 | 0.36268797 | 0.47 | 0.213 | 1.69E-11 |
| Smooth Muscle Cell | Kdr | 5 | 1.36E-14 | 0.4471277 | 0.58 | 0.308 | 2.73E-11 |
| Smooth Muscle Cell | Mybpc3 | 5 | 1.52E-14 | 0.70954037 | 0.695 | 0.406 | 3.05E-11 |
| Smooth Muscle Cell | Acacb | 5 | 1.55E-14 | 0.89874437 | 0.595 | 0.295 | 3.10E-11 |
| Smooth Muscle Cell | Ptprn2 | 5 | 6.90E-14 | 1.01064087 | 0.53 | 0.255 | 1.38E-10 |
| Smooth Muscle Cell | Hlf | 5 | 9.39E-14 | 0.8269931 | 0.695 | 0.443 | 1.88E-10 |
| Smooth Muscle Cell | Lrrc4b | 5 | 1.26E-13 | 0.66842078 | 0.52 | 0.245 | 2.52E-10 |
| Smooth Muscle Cell | Greb1l | 5 | 1.51E-13 | 0.51319255 | 0.5 | 0.235 | 3.01E-10 |
| Smooth Muscle Cell | Ntrk21 | 5 | 1.81E-13 | 0.31464886 | 0.555 | 0.251 | 3.63E-10 |
| Smooth Muscle Cell | Amd1 | 5 | 3.41E-13 | 0.56908429 | 0.57 | 0.318 | 6.82E-10 |
| Smooth Muscle Cell | Naca1 | 5 | 7.57E-13 | 0.30331566 | 0.52 | 0.262 | 1.51E-09 |
| Smooth Muscle Cell | Ccn1 | 5 | 2.50E-12 | 0.65499918 | 0.585 | 0.317 | 5.00E-09 |

| | | | | | | | |
|---|---|---|---|---|---|---|---|
| Smooth Muscle Cell | Cnnm2 | 5 | 2.53E-12 | 0.62465955 | 0.59 | 0.294 | 5.05E-09 |
| Smooth Muscle Cell | Fgf1 | 5 | 2.54E-12 | 0.44763034 | 0.625 | 0.301 | 5.07E-09 |
| Smooth Muscle Cell | Aox1 | 5 | 4.61E-12 | 0.66357615 | 0.83 | 0.501 | 9.22E-09 |
| Smooth Muscle Cell | Ldb3 | 5 | 5.14E-12 | 0.93122299 | 0.58 | 0.316 | 1.03E-08 |
| Smooth Muscle Cell | Rmdn1 | 5 | 9.58E-12 | 0.61831171 | 0.65 | 0.375 | 1.92E-08 |
| Smooth Muscle Cell | Kalrn | 5 | 1.95E-11 | 0.73237616 | 0.675 | 0.4 | 3.90E-08 |
| Smooth Muscle Cell | AABR07054000.1 | 5 | 3.06E-11 | 0.45413067 | 0.52 | 0.25 | 6.13E-08 |
| Smooth Muscle Cell | Antxr1 | 5 | 1.24E-10 | 0.65863694 | 0.77 | 0.511 | 2.48E-07 |
| Smooth Muscle Cell | Tbc1d1 | 5 | 2.32E-10 | 0.49712416 | 0.73 | 0.474 | 4.64E-07 |
| Smooth Muscle Cell | Baiap2l2 | 6 | 2.34E-276 | 7.55088084 | 0.689 | 0.004 | 4.68E-273 |
| Smooth Muscle Cell | Cd27 | 6 | 5.86E-256 | 7.34860102 | 0.689 | 0.007 | 1.17E-252 |
| Smooth Muscle Cell | Egfl6 | 6 | 5.28E-245 | 3.30561603 | 0.66 | 0.006 | 1.06E-241 |
| Smooth Muscle Cell | Ccn6 | 6 | 1.63E-241 | 3.52670726 | 0.66 | 0.007 | 3.26E-238 |
| Smooth Muscle Cell | Scart1 | 6 | 2.45E-235 | 6.83242561 | 0.65 | 0.007 | 4.89E-232 |

| | | | | | | | |
|---|---|---|---|---|---|---|---|
| Smooth Muscle Cell | Tcerg1l | 6 | 3.77E-171 | 4.5296512 | 0.515 | 0.008 | 7.53E-168 |
| Smooth Muscle Cell | Dsc3 | 6 | 6.31E-150 | 3.84233235 | 0.68 | 0.035 | 1.26E-146 |
| Smooth Muscle Cell | Gata3 | 6 | 5.68E-140 | 5.56380226 | 0.699 | 0.04 | 1.14E-136 |
| Smooth Muscle Cell | Tmem156 | 6 | 2.88E-118 | 4.17351391 | 0.553 | 0.005 | 5.76E-115 |
| Smooth Muscle Cell | Edil3 | 6 | 5.87E-114 | 3.07347858 | 0.68 | 0.008 | 1.17E-110 |
| Smooth Muscle Cell | Kif11 | 6 | 1.52E-92 | 3.85165707 | 0.68 | 0.055 | 3.04E-89 |
| Smooth Muscle Cell | Procr | 6 | 3.79E-75 | 2.25488114 | 0.641 | 0.02 | 7.57E-72 |
| Smooth Muscle Cell | Adamts20 | 6 | 6.99E-69 | 2.46585543 | 0.699 | 0.044 | 1.40E-65 |
| Smooth Muscle Cell | Tk1 | 6 | 9.89E-66 | 6.48081554 | 0.66 | 0.025 | 1.98E-62 |
| Smooth Muscle Cell | Hs3st3b1 | 6 | 7.17E-61 | 3.14056867 | 0.709 | 0.132 | 1.43E-57 |
| Smooth Muscle Cell | Pkmyt1 | 6 | 3.37E-56 | 2.24459429 | 0.272 | 0.007 | 6.74E-53 |
| Smooth Muscle Cell | Samd15 | 6 | 9.76E-54 | 3.35210083 | 0.592 | 0.017 | 1.95E-50 |
| Smooth Muscle Cell | Scn9a | 6 | 1.92E-51 | 5.34344687 | 0.592 | 0.103 | 3.83E-48 |
| Smooth Muscle Cell | Cd55 | 6 | 3.04E-51 | 4.81896815 | 0.883 | 0.168 | 6.08E-48 |

| | | | | | | | |
|---|---|---|---|---|---|---|---|
| Smooth Muscle Cell | Kif27 | 6 | 3.53E-51 | 2.50975551 | 0.583 | 0.053 | 7.06E-48 |
| Smooth Muscle Cell | Cadm1 | 6 | 3.13E-49 | 2.20938497 | 0.466 | 0.04 | 6.27E-46 |
| Smooth Muscle Cell | Dpp6 | 6 | 5.91E-47 | 2.05603568 | 0.553 | 0.063 | 1.18E-43 |
| Smooth Muscle Cell | AABR07058 170.1 | 6 | 1.39E-45 | 1.84680679 | 0.709 | 0.102 | 2.78E-42 |
| Smooth Muscle Cell | Troap | 6 | 1.73E-45 | 0.73260555 | 0.398 | 0.023 | 3.45E-42 |
| Smooth Muscle Cell | Flt3 | 6 | 3.01E-45 | 3.52398656 | 0.621 | 0.013 | 6.02E-42 |
| Smooth Muscle Cell | Kcnk3 | 6 | 4.98E-45 | 2.03121452 | 0.592 | 0.035 | 9.96E-42 |
| Smooth Muscle Cell | Fhad1 | 6 | 9.29E-45 | 3.81313265 | 0.427 | 0.033 | 1.86E-41 |
| Smooth Muscle Cell | Dact2 | 6 | 3.72E-43 | 5.53478633 | 0.786 | 0.262 | 7.43E-40 |
| Smooth Muscle Cell | Soat2 | 6 | 2.58E-41 | 3.44278317 | 0.476 | 0.003 | 5.16E-38 |
| Smooth Muscle Cell | Ildr2 | 6 | 3.20E-41 | 4.04987555 | 0.806 | 0.25 | 6.40E-38 |
| Smooth Muscle Cell | Gfpt2 | 6 | 1.09E-39 | 3.22108685 | 0.951 | 0.502 | 2.17E-36 |
| Smooth Muscle Cell | Aurkb | 6 | 4.95E-38 | 0.38324617 | 0.408 | 0.06 | 9.90E-35 |
| Smooth Muscle Cell | Ca8 | 6 | 1.15E-36 | 2.58756538 | 0.67 | 0.068 | 2.30E-33 |

| | | | | | | | |
|---|---|---|---|---|---|---|---|
| Smooth Muscle Cell | Pmfbp1 | 6 | 2.17E-36 | 3.55547891 | 0.612 | 0.193 | 4.34E-33 |
| Smooth Muscle Cell | Pnpla3 | 6 | 2.41E-36 | 1.50891488 | 0.709 | 0.159 | 4.82E-33 |
| Smooth Muscle Cell | Thsd7b1 | 6 | 5.56E-36 | 1.98465026 | 0.621 | 0.06 | 1.11E-32 |
| Smooth Muscle Cell | Nkain31 | 6 | 3.51E-35 | 1.29435779 | 0.699 | 0.184 | 7.02E-32 |
| Smooth Muscle Cell | Edn1 | 6 | 5.93E-35 | 0.36628413 | 0.398 | 0.027 | 1.19E-31 |
| Smooth Muscle Cell | Nkain2 | 6 | 2.02E-34 | 1.25849614 | 0.602 | 0.089 | 4.05E-31 |
| Smooth Muscle Cell | Lurap1l | 6 | 4.27E-34 | 3.46506979 | 0.825 | 0.293 | 8.53E-31 |
| Smooth Muscle Cell | Lyz2 | 6 | 7.16E-34 | 1.66672595 | 0.544 | 0.063 | 1.43E-30 |
| Smooth Muscle Cell | Aldh1a3 | 6 | 1.02E-33 | 3.4420113 | 0.806 | 0.401 | 2.04E-30 |
| Smooth Muscle Cell | Uap1 | 6 | 1.91E-33 | 2.05708532 | 0.922 | 0.611 | 3.82E-30 |
| Smooth Muscle Cell | F13a1 | 6 | 2.74E-33 | 0.99264269 | 0.67 | 0.129 | 5.48E-30 |
| Smooth Muscle Cell | Limch1 | 6 | 7.68E-33 | 2.94614864 | 0.903 | 0.474 | 1.54E-29 |
| Smooth Muscle Cell | Marchf10 | 6 | 1.39E-32 | 1.01968072 | 0.534 | 0.135 | 2.78E-29 |
| Smooth Muscle Cell | Phf24 | 6 | 1.73E-32 | 2.97866412 | 0.524 | 0.075 | 3.46E-29 |

| | | | | | | | |
|---|---|---|---|---|---|---|---|
| Smooth Muscle Cell | Cnksr21 | 6 | 6.35E-32 | 0.81751366 | 0.68 | 0.186 | 1.27E-28 |
| Smooth Muscle Cell | Plaur | 6 | 1.51E-31 | 3.03721703 | 0.806 | 0.25 | 3.02E-28 |
| Smooth Muscle Cell | Nxnl2 | 6 | 2.18E-31 | 0.36446101 | 0.485 | 0.098 | 4.37E-28 |
| Smooth Muscle Cell | Myo16 | 6 | 2.66E-31 | 0.69563939 | 0.641 | 0.217 | 5.31E-28 |
| Smooth Muscle Cell | Klhl40 | 6 | 3.03E-31 | 2.78719277 | 0.592 | 0.172 | 6.06E-28 |
| Smooth Muscle Cell | Cdh13 | 6 | 5.59E-31 | 2.06277268 | 0.942 | 0.604 | 1.12E-27 |
| Smooth Muscle Cell | LOC100910237 | 6 | 7.79E-31 | 0.63701694 | 0.476 | 0.042 | 1.56E-27 |
| Smooth Muscle Cell | Gap43 | 6 | 2.40E-30 | 4.98738174 | 0.456 | 0.07 | 4.81E-27 |
| Smooth Muscle Cell | Rhpn2 | 6 | 9.74E-30 | 1.83074134 | 0.699 | 0.217 | 1.95E-26 |
| Smooth Muscle Cell | Lilrb3a | 6 | 1.49E-29 | 1.59379741 | 0.583 | 0.141 | 2.97E-26 |
| Smooth Muscle Cell | Lsamp | 6 | 1.70E-29 | 1.8723497 | 0.67 | 0.151 | 3.39E-26 |
| Smooth Muscle Cell | Unc45b | 6 | 2.02E-29 | 0.56857286 | 0.631 | 0.145 | 4.05E-26 |
| Smooth Muscle Cell | Samd5 | 6 | 5.10E-29 | 2.02944009 | 0.728 | 0.138 | 1.02E-25 |
| Smooth Muscle Cell | Cubn | 6 | 5.50E-29 | 1.29344196 | 0.68 | 0.263 | 1.10E-25 |

| | | | | | | | |
|---|---|---|---|---|---|---|---|
| Smooth Muscle Cell | Tagln | 6 | 6.29E-29 | 0.74912873 | 0.524 | 0.133 | 1.26E-25 |
| Smooth Muscle Cell | Rab11fip4 | 6 | 8.41E-29 | 1.03787647 | 0.68 | 0.218 | 1.68E-25 |
| Smooth Muscle Cell | Lrrtm4 | 6 | 2.77E-28 | 1.53017514 | 0.641 | 0.136 | 5.54E-25 |
| Smooth Muscle Cell | Axl | 6 | 6.11E-28 | 1.44751643 | 0.951 | 0.667 | 1.22E-24 |
| Smooth Muscle Cell | Dmtn | 6 | 6.64E-28 | 1.07747695 | 0.456 | 0.068 | 1.33E-24 |
| Smooth Muscle Cell | Spock3 | 6 | 9.01E-28 | 2.56588296 | 0.515 | 0.13 | 1.80E-24 |
| Smooth Muscle Cell | Ect2l | 6 | 9.45E-28 | 2.55548679 | 0.485 | 0.1 | 1.89E-24 |
| Smooth Muscle Cell | Nppb | 6 | 1.73E-27 | 1.58382837 | 0.718 | 0.207 | 3.46E-24 |
| Smooth Muscle Cell | Wnt5b1 | 6 | 2.09E-27 | 1.25810689 | 0.544 | 0.194 | 4.18E-24 |
| Smooth Muscle Cell | Drc3 | 6 | 4.13E-27 | 1.24387012 | 0.553 | 0.03 | 8.26E-24 |
| Smooth Muscle Cell | Sema3c | 6 | 6.68E-27 | 2.58286269 | 0.767 | 0.345 | 1.34E-23 |
| Smooth Muscle Cell | Treml4 | 6 | 9.38E-27 | 0.6972045 | 0.485 | 0.126 | 1.88E-23 |
| Smooth Muscle Cell | Sh3bp2 | 6 | 9.79E-27 | 2.47779325 | 0.748 | 0.293 | 1.96E-23 |
| Smooth Muscle Cell | Fam189a2 | 6 | 1.15E-26 | 1.54845158 | 0.592 | 0.036 | 2.31E-23 |

| | | | | | | | |
|---|---|---|---|---|---|---|---|
| Smooth Muscle Cell | Stab1 | 6 | 1.47E-26 | 2.43225195 | 0.631 | 0.146 | 2.95E-23 |
| Smooth Muscle Cell | Galnt151 | 6 | 1.86E-26 | 1.9956061 | 0.767 | 0.39 | 3.72E-23 |
| Smooth Muscle Cell | Pi16 | 6 | 3.62E-26 | 2.09820445 | 0.913 | 0.564 | 7.24E-23 |
| Smooth Muscle Cell | Fbln2 | 6 | 7.68E-26 | 2.01042927 | 0.816 | 0.386 | 1.54E-22 |
| Smooth Muscle Cell | Zdbf21 | 6 | 8.78E-26 | 2.22402655 | 0.417 | 0.099 | 1.76E-22 |
| Smooth Muscle Cell | Esyt3 | 6 | 9.57E-26 | 1.25641544 | 0.68 | 0.18 | 1.91E-22 |
| Smooth Muscle Cell | Ddc2 | 6 | 2.33E-25 | 1.01344687 | 0.631 | 0.197 | 4.66E-22 |
| Smooth Muscle Cell | Diaph3 | 6 | 3.17E-25 | 4.35261061 | 0.495 | 0.209 | 6.34E-22 |
| Smooth Muscle Cell | Cd44 | 6 | 5.20E-25 | 1.73701444 | 0.786 | 0.454 | 1.04E-21 |
| Smooth Muscle Cell | Cd74 | 6 | 6.04E-25 | 1.71666927 | 0.699 | 0.275 | 1.21E-21 |
| Smooth Muscle Cell | Smpd3 | 6 | 6.63E-25 | 0.39465547 | 0.68 | 0.183 | 1.33E-21 |
| Smooth Muscle Cell | Csrp2 | 6 | 8.54E-25 | 1.85086285 | 0.602 | 0.174 | 1.71E-21 |
| Smooth Muscle Cell | Spsb4 | 6 | 3.11E-24 | 2.9324318 | 0.728 | 0.297 | 6.22E-21 |
| Smooth Muscle Cell | Acta2 | 6 | 4.01E-24 | 1.32826518 | 0.738 | 0.254 | 8.01E-21 |

| | | | | | | | |
|---|---|---|---|---|---|---|---|
| Smooth Muscle Cell | Syt9 | 6 | 4.23E-24 | 0.91051464 | 0.544 | 0.113 | 8.46E-21 |
| Smooth Muscle Cell | Ppp1r14c | 6 | 5.48E-24 | 2.06006732 | 0.718 | 0.185 | 1.10E-20 |
| Smooth Muscle Cell | Nova1 | 6 | 7.77E-24 | 2.16869515 | 0.796 | 0.383 | 1.55E-20 |
| Smooth Muscle Cell | Kcnj8 | 6 | 9.48E-24 | 1.33369194 | 0.515 | 0.123 | 1.90E-20 |
| Smooth Muscle Cell | Zbtb7c | 6 | 1.27E-23 | 1.74929485 | 0.796 | 0.393 | 2.54E-20 |
| Smooth Muscle Cell | Gabrb11 | 6 | 1.63E-23 | 1.3156928 | 0.398 | 0.083 | 3.25E-20 |
| Smooth Muscle Cell | Hs3st5 | 6 | 6.15E-23 | 2.28488144 | 0.699 | 0.219 | 1.23E-19 |
| Smooth Muscle Cell | Art3 | 6 | 6.39E-23 | 1.46067652 | 0.738 | 0.269 | 1.28E-19 |
| Smooth Muscle Cell | Nr4a3 | 6 | 7.92E-23 | 2.28551599 | 0.524 | 0.08 | 1.58E-19 |
| Smooth Muscle Cell | Lama5 | 6 | 1.02E-22 | 0.86940229 | 0.718 | 0.342 | 2.04E-19 |
| Smooth Muscle Cell | Scara5 | 6 | 1.29E-22 | 1.85770835 | 0.874 | 0.48 | 2.58E-19 |
| Smooth Muscle Cell | Sh3gl2 | 6 | 1.41E-22 | 1.74658562 | 0.728 | 0.194 | 2.83E-19 |
| Smooth Muscle Cell | Has1 | 6 | 1.46E-22 | 2.5674741 | 0.709 | 0.265 | 2.92E-19 |
| Smooth Muscle Cell | Flnb1 | 6 | 2.03E-22 | 2.26432689 | 0.806 | 0.352 | 4.06E-19 |

| | | | | | | | |
|---|---|---|---|---|---|---|---|
| Smooth Muscle Cell | Hspb7 | 6 | 3.53E-22 | 0.37793498 | 0.689 | 0.233 | 7.06E-19 |
| Smooth Muscle Cell | Optn | 6 | 4.30E-22 | 1.75832488 | 0.786 | 0.276 | 8.59E-19 |
| Smooth Muscle Cell | Rap1gap2 | 6 | 6.14E-22 | 1.11221283 | 0.757 | 0.321 | 1.23E-18 |
| Smooth Muscle Cell | Ripor2 | 6 | 9.58E-22 | 0.74122153 | 0.621 | 0.175 | 1.92E-18 |
| Smooth Muscle Cell | Bcar3 | 6 | 9.64E-22 | 0.48604623 | 0.699 | 0.266 | 1.93E-18 |
| Smooth Muscle Cell | LOC100911847 | 6 | 1.14E-21 | 1.34987324 | 0.728 | 0.41 | 2.28E-18 |
| Smooth Muscle Cell | Lcp1 | 6 | 1.36E-21 | 0.80013463 | 0.427 | 0.148 | 2.73E-18 |
| Smooth Muscle Cell | Mfap5 | 6 | 1.59E-21 | 1.95461682 | 0.738 | 0.311 | 3.17E-18 |
| Smooth Muscle Cell | Atp5f1e | 6 | 2.10E-21 | 0.98548649 | 0.748 | 0.387 | 4.21E-18 |
| Smooth Muscle Cell | Myo18b | 6 | 3.77E-21 | 1.97116662 | 0.699 | 0.325 | 7.55E-18 |
| Smooth Muscle Cell | Tc2n | 6 | 4.87E-21 | 0.52269614 | 0.718 | 0.331 | 9.74E-18 |
| Smooth Muscle Cell | Nr4a1 | 6 | 1.15E-20 | 0.41902702 | 0.689 | 0.362 | 2.30E-17 |
| Smooth Muscle Cell | Odc1 | 6 | 1.79E-20 | 1.20561305 | 0.728 | 0.338 | 3.58E-17 |
| Smooth Muscle Cell | Pde10a | 6 | 2.04E-20 | 1.66393668 | 0.854 | 0.485 | 4.08E-17 |

| | | | | | | | |
|---|---|---|---|---|---|---|---|
| Smooth Muscle Cell | LOC1036933231 | 6 | 2.30E-20 | 0.65488617 | 0.583 | 0.139 | 4.60E-17 |
| Smooth Muscle Cell | Bcl11a | 6 | 3.54E-20 | 2.69245433 | 0.602 | 0.194 | 7.09E-17 |
| Smooth Muscle Cell | Nptxr | 6 | 5.24E-20 | 0.50948304 | 0.68 | 0.231 | 1.05E-16 |
| Smooth Muscle Cell | Sh3pxd2b | 6 | 5.76E-20 | 1.38300308 | 0.816 | 0.396 | 1.15E-16 |
| Smooth Muscle Cell | Fstl1 | 6 | 7.94E-20 | 1.48262278 | 0.874 | 0.573 | 1.59E-16 |
| Smooth Muscle Cell | Iqgap2 | 6 | 8.16E-20 | 0.4650832 | 0.67 | 0.22 | 1.63E-16 |
| Smooth Muscle Cell | Crip1 | 6 | 1.26E-19 | 1.13912655 | 0.748 | 0.369 | 2.52E-16 |
| Smooth Muscle Cell | Pcdh7 | 6 | 1.87E-19 | 0.55299787 | 0.699 | 0.295 | 3.74E-16 |
| Smooth Muscle Cell | Anxa3 | 6 | 2.05E-19 | 1.10791317 | 0.709 | 0.358 | 4.09E-16 |
| Smooth Muscle Cell | Gpnmb1 | 6 | 2.09E-19 | 1.24934769 | 0.563 | 0.159 | 4.18E-16 |
| Smooth Muscle Cell | Rnf152 | 6 | 2.58E-19 | 2.28984849 | 0.485 | 0.204 | 5.16E-16 |
| Smooth Muscle Cell | Mpp7 | 6 | 2.73E-19 | 0.77350214 | 0.748 | 0.21 | 5.47E-16 |
| Smooth Muscle Cell | Gbp1 | 6 | 3.23E-19 | 1.33256381 | 0.612 | 0.186 | 6.47E-16 |
| Smooth Muscle Cell | Ttll7 | 6 | 3.37E-19 | 0.88944959 | 0.738 | 0.217 | 6.75E-16 |

| | | | | | | | |
|---|---|---|---|---|---|---|---|
| Smooth Muscle Cell | Jag1 | 6 | 3.51E-19 | 2.11916036 | 0.68 | 0.246 | 7.01E-16 |
| Smooth Muscle Cell | Dnajc6 | 6 | 3.92E-19 | 1.65734747 | 0.388 | 0.074 | 7.85E-16 |
| Smooth Muscle Cell | Ptpro1 | 6 | 5.25E-19 | 0.63089295 | 0.68 | 0.29 | 1.05E-15 |
| Smooth Muscle Cell | Mgll | 6 | 6.64E-19 | 1.06238746 | 0.786 | 0.445 | 1.33E-15 |
| Smooth Muscle Cell | Fgf18 | 6 | 8.52E-19 | 2.68460303 | 0.505 | 0.187 | 1.70E-15 |
| Smooth Muscle Cell | Camk1d | 6 | 9.15E-19 | 1.46501049 | 0.748 | 0.368 | 1.83E-15 |
| Smooth Muscle Cell | Negr1 | 6 | 1.87E-18 | 1.87057095 | 0.68 | 0.307 | 3.74E-15 |
| Smooth Muscle Cell | Cavin4 | 6 | 2.40E-18 | 1.50809404 | 0.553 | 0.117 | 4.80E-15 |
| Smooth Muscle Cell | Ston2 | 6 | 2.48E-18 | 1.4288938 | 0.903 | 0.533 | 4.96E-15 |
| Smooth Muscle Cell | Timp41 | 6 | 3.37E-18 | 1.21050336 | 0.583 | 0.201 | 6.75E-15 |
| Smooth Muscle Cell | Prom1 | 6 | 3.57E-18 | 0.2845916 | 0.65 | 0.285 | 7.14E-15 |
| Smooth Muscle Cell | Pik3ap1 | 6 | 4.46E-18 | 2.2061576 | 0.592 | 0.164 | 8.92E-15 |
| Smooth Muscle Cell | Pcdh17 | 6 | 4.89E-18 | 0.64152662 | 0.699 | 0.365 | 9.77E-15 |
| Smooth Muscle Cell | Fgf14 | 6 | 5.87E-18 | 2.43562009 | 0.621 | 0.238 | 1.17E-14 |

| | | | | | | | |
|---|---|---|---|---|---|---|---|
| Smooth Muscle Cell | Ptprf | 6 | 5.92E-18 | 1.48407476 | 0.689 | 0.266 | 1.18E-14 |
| Smooth Muscle Cell | Has2 | 6 | 6.77E-18 | 1.91973713 | 0.748 | 0.395 | 1.35E-14 |
| Smooth Muscle Cell | Ccdc3 | 6 | 7.10E-18 | 1.14276797 | 0.592 | 0.224 | 1.42E-14 |
| Smooth Muscle Cell | Cobl | 6 | 7.55E-18 | 0.83562143 | 0.699 | 0.332 | 1.51E-14 |
| Smooth Muscle Cell | Adam23 | 6 | 8.17E-18 | 1.25697147 | 0.709 | 0.276 | 1.63E-14 |
| Smooth Muscle Cell | Adamts1 | 6 | 1.16E-17 | 1.78588284 | 0.612 | 0.2 | 2.32E-14 |
| Smooth Muscle Cell | AABR07027581.1 | 6 | 1.21E-17 | 1.2791795 | 0.718 | 0.31 | 2.43E-14 |
| Smooth Muscle Cell | Ralgps2 | 6 | 1.85E-17 | 2.70838455 | 0.612 | 0.288 | 3.70E-14 |
| Smooth Muscle Cell | Btn2a2 | 6 | 1.87E-17 | 1.66407518 | 0.495 | 0.157 | 3.74E-14 |
| Smooth Muscle Cell | Adgrd1 | 6 | 1.90E-17 | 1.64978878 | 0.825 | 0.55 | 3.80E-14 |
| Smooth Muscle Cell | Ntn4 | 6 | 1.99E-17 | 1.66594065 | 0.816 | 0.431 | 3.97E-14 |
| Smooth Muscle Cell | Col24a1 | 6 | 2.26E-17 | 0.4841424 | 0.709 | 0.334 | 4.52E-14 |
| Smooth Muscle Cell | Smyd1 | 6 | 3.45E-17 | 0.88510256 | 0.544 | 0.12 | 6.90E-14 |
| Smooth Muscle Cell | Hsf51 | 6 | 3.89E-17 | 0.92767226 | 0.583 | 0.239 | 7.77E-14 |

| | | | | | | | |
|---|---|---|---|---|---|---|---|
| Smooth Muscle Cell | Pdcd1lg21 | 6 | 5.00E-17 | 0.50544038 | 0.68 | 0.285 | 1.00E-13 |
| Smooth Muscle Cell | Adcy3 | 6 | 5.08E-17 | 0.54064047 | 0.748 | 0.396 | 1.02E-13 |
| Smooth Muscle Cell | Pde7a | 6 | 5.34E-17 | 0.98285592 | 0.874 | 0.424 | 1.07E-13 |
| Smooth Muscle Cell | Htra2 | 6 | 5.43E-17 | 0.89356718 | 0.709 | 0.35 | 1.09E-13 |
| Smooth Muscle Cell | Itgb4 | 6 | 5.90E-17 | 0.87745172 | 0.67 | 0.351 | 1.18E-13 |
| Smooth Muscle Cell | Abcc81 | 6 | 8.14E-17 | 0.42413534 | 0.583 | 0.123 | 1.63E-13 |
| Smooth Muscle Cell | Me3 | 6 | 1.03E-16 | 0.62046163 | 0.641 | 0.283 | 2.05E-13 |
| Smooth Muscle Cell | B2m | 6 | 1.50E-16 | 0.64035172 | 0.728 | 0.27 | 2.99E-13 |
| Smooth Muscle Cell | Csrp3 | 6 | 1.60E-16 | 1.22648815 | 0.67 | 0.31 | 3.20E-13 |
| Smooth Muscle Cell | AABR07057997.1 | 6 | 2.26E-16 | 0.98214147 | 0.544 | 0.209 | 4.52E-13 |
| Smooth Muscle Cell | Ankrd61 | 6 | 2.57E-16 | 0.70150824 | 0.699 | 0.314 | 5.15E-13 |
| Smooth Muscle Cell | Mob3b | 6 | 2.87E-16 | 0.29548381 | 0.67 | 0.194 | 5.75E-13 |
| Smooth Muscle Cell | Ptprn21 | 6 | 3.50E-16 | 1.31315426 | 0.612 | 0.265 | 7.01E-13 |
| Smooth Muscle Cell | Ntrk22 | 6 | 3.90E-16 | 1.2409591 | 0.68 | 0.259 | 7.81E-13 |

| | | | | | | | |
|---|---|---|---|---|---|---|---|
| Smooth Muscle Cell | Cd274 | 6 | 4.27E-16 | 1.29635801 | 0.66 | 0.249 | 8.53E-13 |
| Smooth Muscle Cell | Sgcd | 6 | 4.94E-16 | 0.38571776 | 0.738 | 0.267 | 9.88E-13 |
| Smooth Muscle Cell | AABR07052585.1 | 6 | 5.25E-16 | 0.53751374 | 0.709 | 0.277 | 1.05E-12 |
| Smooth Muscle Cell | Tm4sf4 | 6 | 5.32E-16 | 0.85143437 | 0.631 | 0.202 | 1.06E-12 |
| Smooth Muscle Cell | Plcb11 | 6 | 9.34E-16 | 1.33059844 | 0.689 | 0.265 | 1.87E-12 |
| Smooth Muscle Cell | PCOLCE2 | 6 | 1.15E-15 | 1.77235033 | 0.718 | 0.315 | 2.30E-12 |
| Smooth Muscle Cell | Usp2 | 6 | 1.16E-15 | 1.43193828 | 0.553 | 0.212 | 2.33E-12 |
| Smooth Muscle Cell | Cacna2d3 | 6 | 1.52E-15 | 1.27802604 | 0.825 | 0.519 | 3.03E-12 |
| Smooth Muscle Cell | Egflam | 6 | 1.63E-15 | 1.08577061 | 0.602 | 0.241 | 3.26E-12 |
| Smooth Muscle Cell | Sgpp21 | 6 | 2.54E-15 | 2.18237368 | 0.476 | 0.202 | 5.08E-12 |
| Smooth Muscle Cell | Prg4 | 6 | 2.57E-15 | 0.44805965 | 0.718 | 0.371 | 5.15E-12 |
| Smooth Muscle Cell | AABR07044900.1 | 6 | 3.62E-15 | 0.70067158 | 0.718 | 0.277 | 7.25E-12 |
| Smooth Muscle Cell | Rgcc | 6 | 4.11E-15 | 1.13952547 | 0.544 | 0.201 | 8.22E-12 |
| Smooth Muscle Cell | Pde1c | 6 | 4.12E-15 | 0.94591383 | 0.631 | 0.247 | 8.25E-12 |

| | | | | | | | |
|---|---|---|---|---|---|---|---|
| Smooth Muscle Cell | AABR07026536.1 | 6 | 4.42E-15 | 0.57428395 | 0.65 | 0.269 | 8.85E-12 |
| Smooth Muscle Cell | Tango2 | 6 | 4.52E-15 | 0.66583944 | 0.738 | 0.272 | 9.03E-12 |
| Smooth Muscle Cell | Sema3a1 | 6 | 4.91E-15 | 0.83929994 | 0.495 | 0.185 | 9.82E-12 |
| Smooth Muscle Cell | Dkk21 | 6 | 5.31E-15 | 1.45275552 | 0.728 | 0.302 | 1.06E-11 |
| Smooth Muscle Cell | Chst14 | 6 | 5.71E-15 | 2.0605136 | 0.476 | 0.143 | 1.14E-11 |
| Smooth Muscle Cell | Hspb8 | 6 | 5.99E-15 | 1.56663802 | 0.602 | 0.281 | 1.20E-11 |
| Smooth Muscle Cell | Fn1 | 6 | 6.80E-15 | 1.5982577 | 0.592 | 0.22 | 1.36E-11 |
| Smooth Muscle Cell | Epha4 | 6 | 9.07E-15 | 0.61969389 | 0.631 | 0.323 | 1.81E-11 |
| Smooth Muscle Cell | Nuak1 | 6 | 1.53E-14 | 0.43899369 | 0.738 | 0.401 | 3.06E-11 |
| Smooth Muscle Cell | Gja11 | 6 | 1.59E-14 | 0.77357096 | 0.573 | 0.232 | 3.18E-11 |
| Smooth Muscle Cell | Tmem51 | 6 | 2.04E-14 | 1.7428995 | 0.495 | 0.211 | 4.08E-11 |
| Smooth Muscle Cell | Dnah5 | 6 | 2.26E-14 | 1.56501876 | 0.67 | 0.302 | 4.51E-11 |
| Smooth Muscle Cell | Mylk3 | 6 | 2.53E-14 | 0.59595251 | 0.699 | 0.325 | 5.05E-11 |
| Smooth Muscle Cell | AABR07017268.11 | 6 | 3.88E-14 | 1.18343803 | 0.602 | 0.216 | 7.77E-11 |

| | | | | | | | |
|---|---|---|---|---|---|---|---|
| Smooth Muscle Cell | Kitlg | 6 | 4.40E-14 | 1.07324655 | 0.777 | 0.385 | 8.80E-11 |
| Smooth Muscle Cell | Emcn | 6 | 4.64E-14 | 0.29375118 | 0.699 | 0.351 | 9.29E-11 |
| Smooth Muscle Cell | Slco3a1 | 6 | 4.79E-14 | 1.19668445 | 0.806 | 0.531 | 9.58E-11 |
| Smooth Muscle Cell | Srpx | 6 | 5.66E-14 | 0.60559813 | 0.699 | 0.276 | 1.13E-10 |
| Smooth Muscle Cell | Flrt2 | 6 | 7.40E-14 | 1.3390112 | 0.728 | 0.312 | 1.48E-10 |
| Smooth Muscle Cell | Slc25a20 | 6 | 7.66E-14 | 0.92049709 | 0.68 | 0.383 | 1.53E-10 |
| Smooth Muscle Cell | Flnc1 | 6 | 8.52E-14 | 0.69901722 | 0.631 | 0.348 | 1.70E-10 |
| Smooth Muscle Cell | AABR07025 140.11 | 6 | 1.31E-13 | 1.05578842 | 0.621 | 0.271 | 2.61E-10 |
| Smooth Muscle Cell | Nalcn | 6 | 1.58E-13 | 1.03660441 | 0.612 | 0.207 | 3.15E-10 |
| Smooth Muscle Cell | Anp32b | 6 | 2.14E-13 | 0.66881128 | 0.728 | 0.386 | 4.28E-10 |
| Smooth Muscle Cell | Prkch | 6 | 2.65E-13 | 0.64543011 | 0.699 | 0.285 | 5.29E-10 |
| Smooth Muscle Cell | Plk2 | 6 | 3.03E-13 | 0.5766977 | 0.641 | 0.342 | 6.07E-10 |
| Smooth Muscle Cell | Pdlim5 | 6 | 3.29E-13 | 1.26288439 | 0.903 | 0.585 | 6.58E-10 |
| Smooth Muscle Cell | Rimbp2 | 6 | 3.92E-13 | 1.81576585 | 0.544 | 0.266 | 7.84E-10 |

| | | | | | | | |
|---|---|---|---|---|---|---|---|
| Smooth Muscle Cell | Ccn11 | 6 | 4.48E-13 | 0.28478462 | 0.631 | 0.328 | 8.95E-10 |
| Smooth Muscle Cell | Kdr1 | 6 | 5.93E-13 | 0.54830766 | 0.709 | 0.315 | 1.19E-09 |
| Smooth Muscle Cell | Vcl | 6 | 6.28E-13 | 1.03680823 | 0.748 | 0.311 | 1.26E-09 |
| Smooth Muscle Cell | Ncald | 6 | 6.37E-13 | 1.15255126 | 0.728 | 0.34 | 1.27E-09 |
| Smooth Muscle Cell | Ptpn18 | 6 | 6.99E-13 | 1.86792457 | 0.398 | 0.126 | 1.40E-09 |
| Smooth Muscle Cell | LOC691083 | 6 | 7.67E-13 | 0.63232399 | 0.68 | 0.313 | 1.53E-09 |
| Smooth Muscle Cell | Fbxo40 | 6 | 7.87E-13 | 1.32647578 | 0.505 | 0.193 | 1.57E-09 |
| Smooth Muscle Cell | Dipk1a | 6 | 9.44E-13 | 0.40787403 | 0.67 | 0.314 | 1.89E-09 |
| Smooth Muscle Cell | Col22a1 | 6 | 1.08E-12 | 0.68005271 | 0.641 | 0.344 | 2.16E-09 |
| Smooth Muscle Cell | Prkcz | 6 | 1.38E-12 | 0.3693408 | 0.65 | 0.264 | 2.76E-09 |
| Smooth Muscle Cell | Csrp1 | 6 | 2.35E-12 | 0.67953877 | 0.738 | 0.465 | 4.71E-09 |
| Smooth Muscle Cell | Pcdh19 | 6 | 2.46E-12 | 0.76830509 | 0.631 | 0.301 | 4.93E-09 |
| Smooth Muscle Cell | Gfod1 | 6 | 2.52E-12 | 0.34642608 | 0.738 | 0.321 | 5.05E-09 |
| Smooth Muscle Cell | Arhgap11a | 6 | 3.16E-12 | 0.37177489 | 0.515 | 0.208 | 6.33E-09 |

| | | | | | | | |
|---|---|---|---|---|---|---|---|
| Smooth Muscle Cell | Pip5k1b1 | 6 | 3.23E-12 | 0.6195646 | 0.709 | 0.313 | 6.45E-09 |
| Smooth Muscle Cell | Itga6 | 6 | 3.30E-12 | 0.59233133 | 0.728 | 0.386 | 6.60E-09 |
| Smooth Muscle Cell | Itga9 | 6 | 3.37E-12 | 1.28743808 | 0.903 | 0.637 | 6.74E-09 |
| Smooth Muscle Cell | LOC691141 | 6 | 4.16E-12 | 1.21865327 | 0.485 | 0.151 | 8.31E-09 |
| Smooth Muscle Cell | Adamts5 | 6 | 4.42E-12 | 1.44407708 | 0.883 | 0.574 | 8.85E-09 |
| Smooth Muscle Cell | Medag | 6 | 5.11E-12 | 1.02218659 | 0.718 | 0.437 | 1.02E-08 |
| Smooth Muscle Cell | Kcnab1 | 6 | 6.32E-12 | 0.87558029 | 0.621 | 0.342 | 1.26E-08 |
| Smooth Muscle Cell | Slit3 | 6 | 7.73E-12 | 0.97326087 | 0.67 | 0.406 | 1.55E-08 |
| Smooth Muscle Cell | Cfh | 6 | 1.06E-11 | 0.45592249 | 0.718 | 0.368 | 2.12E-08 |
| Smooth Muscle Cell | Musk | 6 | 1.11E-11 | 0.35501112 | 0.65 | 0.279 | 2.22E-08 |
| Smooth Muscle Cell | Ptgs2 | 6 | 1.25E-11 | 0.66565819 | 0.553 | 0.181 | 2.50E-08 |
| Smooth Muscle Cell | Rasa41 | 6 | 1.32E-11 | 0.34373774 | 0.641 | 0.298 | 2.64E-08 |
| Smooth Muscle Cell | Cdk19 | 6 | 1.98E-11 | 0.60222762 | 0.786 | 0.379 | 3.95E-08 |
| Smooth Muscle Cell | Fsd21 | 6 | 2.09E-11 | 0.29489454 | 0.602 | 0.254 | 4.19E-08 |

| | | | | | | | |
|---|---|---|---|---|---|---|---|
| Smooth Muscle Cell | Taldo11 | 6 | 3.30E-11 | 1.03679431 | 0.621 | 0.216 | 6.60E-08 |
| Smooth Muscle Cell | Kcnt2 | 6 | 3.50E-11 | 0.49518294 | 0.505 | 0.178 | 7.00E-08 |
| Smooth Muscle Cell | Nnt1 | 6 | 4.82E-11 | 0.63546275 | 0.709 | 0.383 | 9.64E-08 |
| Smooth Muscle Cell | Ldb31 | 6 | 6.43E-11 | 0.2699799 | 0.709 | 0.322 | 1.29E-07 |
| Smooth Muscle Cell | Gpr63 | 6 | 7.08E-11 | 1.07600836 | 0.66 | 0.284 | 1.42E-07 |
| Smooth Muscle Cell | Fcgr2b | 6 | 8.50E-11 | 1.23305411 | 0.437 | 0.148 | 1.70E-07 |
| Smooth Muscle Cell | Tmsb4x | 6 | 1.46E-10 | 0.88981739 | 0.68 | 0.312 | 2.92E-07 |
| Smooth Muscle Cell | Plekha4 | 6 | 1.66E-10 | 0.8911324 | 0.699 | 0.38 | 3.32E-07 |
| Smooth Muscle Cell | Mgmt | 6 | 2.02E-10 | 0.56626074 | 0.767 | 0.404 | 4.04E-07 |
| Smooth Muscle Cell | Prrx1 | 6 | 2.33E-10 | 1.03622715 | 0.864 | 0.591 | 4.66E-07 |
| Smooth Muscle Cell | Ptprj | 6 | 2.40E-10 | 0.84343877 | 0.922 | 0.617 | 4.80E-07 |
| Smooth Muscle Cell | Fbxo32 | 6 | 2.61E-10 | 0.84539177 | 0.65 | 0.399 | 5.21E-07 |
| Smooth Muscle Cell | Ptpn3 | 6 | 4.73E-10 | 0.42496813 | 0.709 | 0.348 | 9.47E-07 |
| Smooth Muscle Cell | Rcan21 | 6 | 7.89E-10 | 1.10812157 | 0.835 | 0.496 | 1.58E-06 |

| | | | | | | | |
|---|---|---|---|---|---|---|---|
| Smooth Muscle Cell | P2rx71 | 6 | 8.00E-10 | 0.91743829 | 0.544 | 0.249 | 1.60E-06 |
| Smooth Muscle Cell | Mcf2l | 6 | 9.07E-10 | 1.35561272 | 0.495 | 0.213 | 1.81E-06 |
| Smooth Muscle Cell | Cadps2 | 6 | 1.07E-09 | 0.65678732 | 0.641 | 0.296 | 2.14E-06 |
| Smooth Muscle Cell | Slfn13 | 6 | 1.88E-09 | 0.39843112 | 0.709 | 0.275 | 3.76E-06 |
| Smooth Muscle Cell | Antxr11 | 6 | 1.88E-09 | 0.84254721 | 0.825 | 0.521 | 3.76E-06 |
| Smooth Muscle Cell | Afap1l2 | 6 | 2.31E-09 | 0.25227017 | 0.757 | 0.45 | 4.63E-06 |
| Smooth Muscle Cell | Tspan5 | 6 | 2.70E-09 | 0.76921598 | 0.825 | 0.466 | 5.40E-06 |
| Smooth Muscle Cell | Cdkn1a1 | 6 | 2.72E-09 | 0.77647273 | 0.524 | 0.253 | 5.44E-06 |
| Smooth Muscle Cell | Slc38a1 | 6 | 5.47E-09 | 0.39079544 | 0.563 | 0.226 | 1.09E-05 |
| Smooth Muscle Cell | Slc1a1 | 6 | 1.10E-08 | 1.16153189 | 0.524 | 0.174 | 2.20E-05 |
| Smooth Muscle Cell | AABR07058158.1 | 6 | 1.31E-08 | 0.63223652 | 0.777 | 0.398 | 2.63E-05 |
| Smooth Muscle Cell | Ralgapa2 | 6 | 1.41E-08 | 0.32533952 | 0.786 | 0.437 | 2.83E-05 |
| Smooth Muscle Cell | Hydin | 6 | 3.63E-08 | 0.80250463 | 0.495 | 0.207 | 7.25E-05 |
| Smooth Muscle Cell | Kif22 | 6 | 4.81E-08 | 2.80793978 | 0.447 | 0.187 | 9.62E-05 |

| | | | | | | | |
|---|---|---|---|---|---|---|---|
| Smooth Muscle Cell | Rnls | 6 | 5.64E-08 | 0.5846403 | 0.689 | 0.38 | 0.00011275 |
| Smooth Muscle Cell | Col16a1 | 6 | 6.13E-08 | 0.40746162 | 0.641 | 0.298 | 0.00012258 |
| Smooth Muscle Cell | Polr2m | 6 | 7.14E-08 | 0.38918299 | 0.738 | 0.399 | 0.00014286 |
| Smooth Muscle Cell | Aacs | 6 | 1.11E-07 | 0.68345325 | 0.553 | 0.296 | 0.00022132 |
| Smooth Muscle Cell | Phyh | 6 | 1.11E-07 | 0.42790696 | 0.738 | 0.41 | 0.00022243 |
| Smooth Muscle Cell | AABR07025295.1 | 6 | 2.16E-07 | 0.64812693 | 0.718 | 0.393 | 0.00043288 |
| Smooth Muscle Cell | Actn2 | 6 | 2.79E-07 | 0.37750829 | 0.631 | 0.322 | 0.0005576 |
| Smooth Muscle Cell | Ncam2 | 6 | 4.80E-07 | 1.2048839 | 0.456 | 0.198 | 0.0009595 |
| Smooth Muscle Cell | Nox4 | 6 | 1.02E-06 | 0.32187903 | 0.66 | 0.353 | 0.00204253 |
| Smooth Muscle Cell | Slc25a13 | 6 | 1.12E-06 | 0.30648423 | 0.699 | 0.365 | 0.00224071 |
| Smooth Muscle Cell | Gnao1 | 6 | 1.33E-06 | 0.38743549 | 0.621 | 0.359 | 0.00266593 |
| Smooth Muscle Cell | Sat1 | 6 | 1.39E-06 | 0.29179021 | 0.709 | 0.354 | 0.00278301 |
| Smooth Muscle Cell | Sorbs1 | 6 | 1.48E-06 | 0.60875855 | 0.835 | 0.553 | 0.00295646 |
| Smooth Muscle Cell | Efemp1 | 6 | 3.37E-06 | 0.3946889 | 0.68 | 0.428 | 0.00674064 |

| | | | | | | | |
|---|---|---|---|---|---|---|---|
| Smooth Muscle Cell | Ntn1 | 6 | 3.96E-06 | 0.36033014 | 0.796 | 0.54 | 0.00792602 |
| Smooth Muscle Cell | Ckm | 6 | 7.28E-06 | 0.49885788 | 0.68 | 0.385 | 0.01456423 |
| Smooth Muscle Cell | Zfp622 | 6 | 9.40E-06 | 0.34672215 | 0.718 | 0.406 | 0.01880331 |
| Smooth Muscle Cell | Lrg1 | 7 | 1.14E-23 | 0.95625981 | 0.261 | 0.011 | 2.27E-20 |
| Smooth Muscle Cell | Vil1 | 7 | 1.47E-19 | 1.61999706 | 0.304 | 0.049 | 2.94E-16 |
| Smooth Muscle Cell | AABR07005983.1 | 7 | 4.22E-19 | 0.49728316 | 0.304 | 0.05 | 8.43E-16 |
| Smooth Muscle Cell | Crhr2 | 7 | 9.06E-17 | 0.25782064 | 0.42 | 0.129 | 1.81E-13 |
| Smooth Muscle Cell | Grik4 | 7 | 1.17E-13 | 2.51091344 | 0.377 | 0.1 | 2.34E-10 |
| Smooth Muscle Cell | Etl4 | 7 | 5.45E-12 | 2.00102262 | 0.696 | 0.373 | 1.09E-08 |
| Smooth Muscle Cell | Cyyr1 | 7 | 9.92E-12 | 2.0452332 | 0.681 | 0.344 | 1.98E-08 |
| Smooth Muscle Cell | Epas1 | 7 | 9.56E-11 | 1.58308563 | 0.797 | 0.505 | 1.91E-07 |
| Smooth Muscle Cell | Ptprb | 7 | 2.96E-10 | 1.95332609 | 0.652 | 0.297 | 5.91E-07 |
| Smooth Muscle Cell | Adgrf5 | 7 | 4.72E-10 | 2.06686621 | 0.652 | 0.392 | 9.45E-07 |
| Smooth Muscle Cell | Adgrl4 | 7 | 1.31E-09 | 2.03668368 | 0.609 | 0.346 | 2.63E-06 |

| | | | | | | | |
|---|---|---|---|---|---|---|---|
| Smooth Muscle Cell | Unc5c1 | 7 | 2.69E-09 | 2.17626447 | 0.377 | 0.108 | 5.38E-06 |
| Smooth Muscle Cell | Flvcr2 | 7 | 1.83E-08 | 0.746626 | 0.348 | 0.089 | 3.66E-05 |
| Smooth Muscle Cell | Herc6 | 7 | 5.39E-08 | 2.06495194 | 0.609 | 0.253 | 0.00010786 |
| Smooth Muscle Cell | Mx2 | 7 | 1.21E-07 | 3.49269604 | 0.42 | 0.133 | 0.00024235 |
| Smooth Muscle Cell | AC123500.1 | 7 | 1.22E-07 | 3.16015426 | 0.377 | 0.1 | 0.00024393 |
| Smooth Muscle Cell | Mcf2l1 | 7 | 1.61E-07 | 1.97144667 | 0.522 | 0.217 | 0.00032104 |
| Smooth Muscle Cell | Crim1 | 7 | 8.81E-07 | 1.48424602 | 0.725 | 0.458 | 0.00176183 |
| Smooth Muscle Cell | Slc1a11 | 7 | 9.14E-07 | 1.85170946 | 0.449 | 0.183 | 0.00182773 |
| Smooth Muscle Cell | Rnf2131 | 7 | 1.50E-06 | 1.70114302 | 0.652 | 0.393 | 0.00299686 |
| Smooth Muscle Cell | Cadm2 | 7 | 5.59E-06 | 2.638065 | 0.594 | 0.328 | 0.01118181 |
| Smooth Muscle Cell | Smyd11 | 7 | 9.88E-06 | 0.86996481 | 0.391 | 0.133 | 0.01975225 |
| Smooth Muscle Cell | Cdk191 | 7 | 2.43E-05 | 1.01127739 | 0.667 | 0.391 | 0.04864693 |
| Smooth Muscle Cell | Tyrobp | 8 | 3.62E-72 | 3.7472018 | 0.373 | 0.015 | 7.23E-69 |
| Smooth Muscle Cell | Apobec1 | 8 | 7.80E-69 | 4.29844184 | 0.343 | 0.013 | 1.56E-65 |

| | | | | | | | |
|---|---|---|---|---|---|---|---|
| Smooth Muscle Cell | Hdc | 8 | 9.21E-66 | 4.43153178 | 0.284 | 0.008 | 1.84E-62 |
| Smooth Muscle Cell | Tbx21 | 8 | 8.64E-63 | 3.60345105 | 0.313 | 0.013 | 1.73E-59 |
| Smooth Muscle Cell | Sdc1 | 8 | 3.60E-62 | 3.3975462 | 0.343 | 0.013 | 7.19E-59 |
| Smooth Muscle Cell | Fgf5 | 8 | 2.09E-42 | 6.6669807 | 0.313 | 0.021 | 4.18E-39 |
| Smooth Muscle Cell | Hao1 | 8 | 1.13E-37 | 5.9343173 | 0.358 | 0.034 | 2.25E-34 |
| Smooth Muscle Cell | AABR07068046.1 | 8 | 3.67E-33 | 8.23549005 | 0.313 | 0.012 | 7.35E-30 |
| Smooth Muscle Cell | AABR07067469.1 | 8 | 2.75E-31 | 6.53725178 | 0.522 | 0.096 | 5.50E-28 |
| Smooth Muscle Cell | Map3k7cl | 8 | 1.19E-24 | 3.287507 | 0.373 | 0.043 | 2.39E-21 |
| Smooth Muscle Cell | Ltbp2 | 8 | 2.89E-24 | 3.38489462 | 0.851 | 0.507 | 5.77E-21 |
| Smooth Muscle Cell | Arhgef39 | 8 | 4.77E-24 | 6.27624201 | 0.328 | 0.012 | 9.55E-21 |
| Smooth Muscle Cell | Ptprv | 8 | 4.75E-23 | 4.58054803 | 0.493 | 0.101 | 9.50E-20 |
| Smooth Muscle Cell | Galnt141 | 8 | 2.01E-22 | 5.09741095 | 0.433 | 0.089 | 4.01E-19 |
| Smooth Muscle Cell | Il11 | 8 | 4.36E-22 | 2.44219412 | 0.328 | 0.013 | 8.72E-19 |
| Smooth Muscle Cell | Tpx2 | 8 | 5.41E-17 | 4.61721272 | 0.433 | 0.018 | 1.08E-13 |

| | | | | | | | |
|---|---|---|---|---|---|---|---|
| Smooth Muscle Cell | Rergl | 8 | 3.58E-16 | 2.45155085 | 0.358 | 0.072 | 7.17E-13 |
| Smooth Muscle Cell | Klhl29 | 8 | 1.80E-14 | 2.30540475 | 0.806 | 0.451 | 3.60E-11 |
| Smooth Muscle Cell | Prc1 | 8 | 1.88E-14 | 3.26214878 | 0.284 | 0.005 | 3.76E-11 |
| Smooth Muscle Cell | Fn11 | 8 | 2.51E-14 | 3.44656436 | 0.716 | 0.223 | 5.02E-11 |
| Smooth Muscle Cell | Pdgfc | 8 | 5.43E-14 | 2.03352064 | 0.433 | 0.036 | 1.09E-10 |
| Smooth Muscle Cell | Itgal | 8 | 6.60E-14 | 4.58806033 | 0.313 | 0.013 | 1.32E-10 |
| Smooth Muscle Cell | Cdh1 | 8 | 9.56E-14 | 3.04398954 | 0.463 | 0.12 | 1.91E-10 |
| Smooth Muscle Cell | Tagln1 | 8 | 1.40E-13 | 3.41881001 | 0.463 | 0.142 | 2.79E-10 |
| Smooth Muscle Cell | Itga10 | 8 | 1.47E-13 | 4.6136898 | 0.358 | 0.038 | 2.94E-10 |
| Smooth Muscle Cell | Crlf1 | 8 | 4.15E-13 | 2.95267119 | 0.716 | 0.444 | 8.31E-10 |
| Smooth Muscle Cell | Runx1 | 8 | 4.49E-13 | 2.67672954 | 0.761 | 0.475 | 8.98E-10 |
| Smooth Muscle Cell | Kcnh5 | 8 | 1.35E-12 | 2.4323585 | 0.343 | 0.06 | 2.70E-09 |
| Smooth Muscle Cell | Itgb21 | 8 | 1.66E-12 | 1.78867637 | 0.448 | 0.153 | 3.33E-09 |
| Smooth Muscle Cell | Tceal7 | 8 | 1.79E-12 | 0.79738336 | 0.343 | 0.079 | 3.57E-09 |

| | | | | | | | |
|---|---|---|---|---|---|---|---|
| Smooth Muscle Cell | Hey2 | 8 | 2.01E-12 | 3.12328942 | 0.343 | 0.09 | 4.03E-09 |
| Smooth Muscle Cell | Cobl1 | 8 | 3.16E-11 | 3.21725237 | 0.657 | 0.34 | 6.33E-08 |
| Smooth Muscle Cell | Spp1 | 8 | 2.81E-10 | 2.03782198 | 0.403 | 0.092 | 5.61E-07 |
| Smooth Muscle Cell | Kif111 | 8 | 3.71E-10 | 2.56847998 | 0.328 | 0.078 | 7.42E-07 |
| Smooth Muscle Cell | Trpc6 | 8 | 7.75E-10 | 3.78602508 | 0.343 | 0.052 | 1.55E-06 |
| Smooth Muscle Cell | Actg2 | 8 | 1.91E-09 | 3.87488751 | 0.343 | 0.085 | 3.82E-06 |
| Smooth Muscle Cell | Col1a1 | 8 | 6.91E-09 | 1.70194879 | 0.806 | 0.485 | 1.38E-05 |
| Smooth Muscle Cell | Ifit1bl | 8 | 1.17E-08 | 2.99504052 | 0.418 | 0.161 | 2.35E-05 |
| Smooth Muscle Cell | Pmepa1 | 8 | 1.21E-08 | 1.35701062 | 0.776 | 0.432 | 2.42E-05 |
| Smooth Muscle Cell | Col27a1 | 8 | 1.56E-08 | 2.43003041 | 0.627 | 0.324 | 3.12E-05 |
| Smooth Muscle Cell | Anks1b | 8 | 2.20E-08 | 4.87587709 | 0.507 | 0.082 | 4.40E-05 |
| Smooth Muscle Cell | Olfm2 | 8 | 2.92E-08 | 3.46184915 | 0.463 | 0.184 | 5.84E-05 |
| Smooth Muscle Cell | Bcat1 | 8 | 5.21E-08 | 1.91706023 | 0.657 | 0.327 | 0.00010424 |
| Smooth Muscle Cell | Col19a1 | 8 | 2.10E-07 | 1.22233796 | 0.418 | 0.123 | 0.00042059 |

| | | | | | | | |
|---|---|---|---|---|---|---|---|
| Smooth Muscle Cell | Dkk3 | 8 | 2.23E-07 | 1.31076001 | 0.687 | 0.416 | 0.00044659 |
| Smooth Muscle Cell | Chst11 | 8 | 2.66E-07 | 1.7538915 | 0.731 | 0.4 | 0.00053282 |
| Smooth Muscle Cell | Ccl2 | 8 | 1.03E-06 | 3.0685483 | 0.433 | 0.183 | 0.0020612 |
| Smooth Muscle Cell | Flna | 8 | 1.46E-06 | 1.38374109 | 0.701 | 0.379 | 0.00292417 |
| Smooth Muscle Cell | Phf241 | 8 | 1.74E-06 | 2.93831892 | 0.343 | 0.089 | 0.00347239 |
| Smooth Muscle Cell | AC134204.1 | 8 | 3.00E-06 | 0.90729759 | 0.701 | 0.388 | 0.00599707 |
| Smooth Muscle Cell | Atp10a | 8 | 8.00E-06 | 0.83745894 | 0.761 | 0.484 | 0.01600584 |
| Smooth Muscle Cell | Fgf11 | 8 | 1.34E-05 | 1.99751061 | 0.582 | 0.325 | 0.02688759 |
| Smooth Muscle Cell | Bgn | 8 | 1.47E-05 | 1.46286063 | 0.627 | 0.337 | 0.02939868 |
| Smooth Muscle Cell | Vim | 8 | 1.86E-05 | 1.28620694 | 0.582 | 0.31 | 0.03716221 |
| Smooth Muscle Cell | Vcam1 | 8 | 2.40E-05 | 2.80310447 | 0.522 | 0.194 | 0.04806478 |
| Smooth Muscle Cell | Spdef | 9 | 2.99E-12 | 3.07887111 | 0.4 | 0.028 | 5.97E-09 |
| Smooth Muscle Cell | Tmem163 | 9 | 2.79E-10 | 6.62495542 | 0.6 | 0.012 | 5.59E-07 |
| Smooth Muscle Cell | Brinp1 | 9 | 5.51E-08 | 6.35396124 | 0.3 | 0.002 | 0.00011028 |

| | | | | | | | |
|---|---|---|---|---|---|---|---|
| Smooth Muscle Cell | Neto1 | 9 | 5.67E-08 | 2.69827346 | 0.5 | 0.072 | 0.00011341 |
| Smooth Muscle Cell | Ifit3 | 9 | 5.37E-07 | 2.69934897 | 0.4 | 0.03 | 0.00107346 |
| Smooth Muscle Cell | Tnni3 | 9 | 1.16E-06 | 2.62424006 | 1 | 0.602 | 0.00232269 |
| Smooth Muscle Cell | Nppa | 9 | 1.93E-06 | 4.62363122 | 0.9 | 0.402 | 0.00385883 |
| Smooth Muscle Cell | Tmem2361 | 9 | 2.46E-06 | 3.43199055 | 0.5 | 0.079 | 0.00492866 |
| Smooth Muscle Cell | Mb | 9 | 2.93E-06 | 4.22731393 | 0.9 | 0.52 | 0.00586855 |
| Smooth Muscle Cell | Adgre4 | 9 | 3.97E-06 | 5.33722495 | 0.6 | 0.06 | 0.00794603 |
| Smooth Muscle Cell | Selp | 9 | 6.56E-06 | 3.17667697 | 0.3 | 0.047 | 0.01312022 |
| Smooth Muscle Cell | Smc1b1 | 9 | 8.36E-06 | 3.39402655 | 0.4 | 0.078 | 0.01672344 |
| Smooth Muscle Cell | Ankrd1 | 9 | 1.29E-05 | 2.7668679 | 0.9 | 0.579 | 0.02579052 |
| Smooth Muscle Cell | LOC100910636 | 9 | 1.99E-05 | 3.18120563 | 0.5 | 0.068 | 0.0397211 |
| Smooth Muscle Cell | LOC308990 | 9 | 2.42E-05 | 3.21801719 | 0.4 | 0.061 | 0.04840792 |
