## Supplemental Table 5 for "Uncovering the Regional and Cell Specific Bioactivity of Injectable Extracellular Matrix Biomaterials in Myocardial Infarction through Spatial and Single Nucleus Transcriptomics"

**Supplementary Table 5. Coarse Cluster Markers for Subacute Model**

| Cell Type | Gene | p_val | avg_log2FC | pct.1 | pct.2 | p_val_adj |
|---|---|---|---|---|---|---|
| Proliferating Cells | Top2a | 0 | 6.92624874 | 0.852 | 0.199 | 0 |
| Proliferating Cells | Sgo2 | 0 | 6.66196305 | 0.784 | 0.176 | 0 |
| Proliferating Cells | Diaph3 | 0 | 5.26160024 | 0.866 | 0.261 | 0 |
| Proliferating Cells | Kif4a | 0 | 6.81140334 | 0.72 | 0.119 | 0 |
| Proliferating Cells | Kif11 | 0 | 6.71892744 | 0.766 | 0.166 | 0 |
| Proliferating Cells | Ect2 | 0 | 6.531051 | 0.755 | 0.156 | 0 |
| Proliferating Cells | Prc1 | 0 | 6.8438098 | 0.737 | 0.144 | 0 |
| Proliferating Cells | Cenpf | 0 | 6.99066164 | 0.754 | 0.163 | 0 |
| Proliferating Cells | Kif14 | 0 | 7.36740601 | 0.643 | 0.059 | 0 |
| Proliferating Cells | Melk | 0 | 6.68253691 | 0.63 | 0.051 | 0 |
| Proliferating Cells | Iqgap3 | 0 | 6.80496477 | 0.71 | 0.137 | 0 |
| Proliferating Cells | Bub1b | 0 | 6.96735965 | 0.658 | 0.087 | 0 |

| | | | | | | |
|---|---|---|---|---|---|---|
| Proliferating Cells | Kif23 | 0 | 6.72281605 | 0.75 | 0.179 | 0 |
| Proliferating Cells | Kifc1 | 0 | 6.61977189 | 0.646 | 0.082 | 0 |
| Proliferating Cells | Cdca2 | 0 | 7.26518678 | 0.664 | 0.102 | 0 |
| Proliferating Cells | Ska1 | 0 | 7.38669518 | 0.654 | 0.092 | 0 |
| Proliferating Cells | Troap | 0 | 7.23799134 | 0.669 | 0.109 | 0 |
| Proliferating Cells | Ttk | 0 | 6.97525248 | 0.63 | 0.092 | 0 |
| Proliferating Cells | Ccnf | 0 | 6.47419305 | 0.608 | 0.086 | 0 |
| Proliferating Cells | Espl1 | 0 | 6.95697064 | 0.585 | 0.078 | 0 |
| Proliferating Cells | E2f7 | 0 | 5.94672657 | 0.674 | 0.168 | 0 |
| Proliferating Cells | Ncapg | 0 | 6.62838927 | 0.57 | 0.072 | 0 |
| Proliferating Cells | Hjurp | 0 | 7.14800029 | 0.581 | 0.099 | 0 |
| Proliferating Cells | Kif18b | 0 | 6.77884551 | 0.594 | 0.15 | 0 |
| Proliferating Cells | Knstrn | 0 | 6.54650302 | 0.509 | 0.102 | 0 |
| Proliferating Cells | Depdc1 | 0 | 6.11664085 | 0.449 | 0.075 | 0 |

| | | | | | | |
|---|---|---|---|---|---|---|
| Proliferating Cells | Timd2 | 0 | 3.31806065 | 0.368 | 0.011 | 0 |
| Proliferating Cells | Polq | 1.19E-307 | 6.36254731 | 0.615 | 0.125 | 2.38E-304 |
| Proliferating Cells | Mki67 | 1.06E-304 | 5.8114429 | 0.792 | 0.254 | 2.11E-301 |
| Proliferating Cells | Cep55 | 3.57E-297 | 7.22509801 | 0.615 | 0.067 | 7.15E-294 |
| Proliferating Cells | Depdc1b | 1.04E-296 | 6.48778555 | 0.536 | 0.09 | 2.08E-293 |
| Proliferating Cells | Cdkn3 | 1.25E-291 | 6.25974748 | 0.4 | 0.061 | 2.50E-288 |
| Proliferating Cells | Cenpe | 5.77E-287 | 6.28322794 | 0.767 | 0.233 | 1.15E-283 |
| Proliferating Cells | Nusap1 | 4.22E-276 | 6.41665131 | 0.725 | 0.221 | 8.44E-273 |
| Proliferating Cells | Ncaph | 1.53E-275 | 5.98316615 | 0.716 | 0.194 | 3.07E-272 |
| Proliferating Cells | Ckap2l | 2.15E-275 | 6.57275389 | 0.533 | 0.13 | 4.30E-272 |
| Proliferating Cells | Kif2c | 1.07E-272 | 6.86551164 | 0.452 | 0.05 | 2.15E-269 |
| Proliferating Cells | Neil3 | 9.57E-269 | 6.10577122 | 0.594 | 0.133 | 1.91E-265 |
| Proliferating Cells | Tpx2 | 1.17E-262 | 6.20294852 | 0.733 | 0.231 | 2.34E-259 |
| Proliferating Cells | Esco2 | 8.89E-261 | 6.54526074 | 0.462 | 0.12 | 1.78E-257 |

| | | | | | | |
|---|---|---|---|---|---|---|
| Proliferating Cells | Aspm | 7.52E-255 | 6.31900215 | 0.667 | 0.196 | 1.50E-251 |
| Proliferating Cells | Ckap2 | 3.61E-254 | 6.20820737 | 0.581 | 0.134 | 7.22E-251 |
| Proliferating Cells | Ccnb1 | 4.39E-253 | 6.1049984 | 0.461 | 0.07 | 8.77E-250 |
| Proliferating Cells | Anln | 2.98E-247 | 6.1223307 | 0.664 | 0.188 | 5.96E-244 |
| Proliferating Cells | Parpbp | 4.06E-246 | 6.96727489 | 0.5 | 0.089 | 8.12E-243 |
| Proliferating Cells | Efcab11 | 5.21E-244 | 5.52465505 | 0.454 | 0.062 | 1.04E-240 |
| Proliferating Cells | Ndc80 | 1.49E-230 | 5.85809423 | 0.69 | 0.166 | 2.97E-227 |
| Proliferating Cells | Cdk1 | 5.12E-228 | 5.96067871 | 0.535 | 0.095 | 1.02E-224 |
| Proliferating Cells | Knl1 | 8.33E-221 | 5.43521343 | 0.682 | 0.226 | 1.67E-217 |
| Proliferating Cells | Nav2 | 3.19E-213 | 3.36416765 | 0.826 | 0.422 | 6.38E-210 |
| Proliferating Cells | Fscn2 | 1.72E-212 | 1.98518177 | 0.366 | 0.06 | 3.44E-209 |
| Proliferating Cells | Bard1 | 1.93E-212 | 5.90397058 | 0.596 | 0.126 | 3.86E-209 |
| Proliferating Cells | Ezh2 | 2.63E-210 | 3.56265713 | 0.794 | 0.397 | 5.25E-207 |
| Proliferating Cells | Kif22 | 6.80E-208 | 5.77812219 | 0.663 | 0.18 | 1.36E-204 |

| | | | | | | |
|---|---|---|---|---|---|---|
| Proliferating Cells | Cit | 6.24E-202 | 4.6781979 | 0.725 | 0.319 | 1.25E-198 |
| Proliferating Cells | Kif20b | 1.43E-200 | 6.18399179 | 0.555 | 0.198 | 2.86E-197 |
| Proliferating Cells | Hmmr | 2.22E-200 | 5.98830651 | 0.6 | 0.149 | 4.45E-197 |
| Proliferating Cells | Arhgef39 | 1.95E-190 | 6.21098547 | 0.439 | 0.095 | 3.90E-187 |
| Proliferating Cells | Kntc1 | 1.10E-179 | 5.13977306 | 0.628 | 0.209 | 2.20E-176 |
| Proliferating Cells | Dlgap5 | 6.98E-169 | 5.33172812 | 0.682 | 0.229 | 1.40E-165 |
| Proliferating Cells | Mis18bp1 | 2.08E-166 | 4.93163708 | 0.661 | 0.228 | 4.16E-163 |
| Proliferating Cells | Brca1 | 1.26E-160 | 4.51117247 | 0.691 | 0.265 | 2.52E-157 |
| Proliferating Cells | Gen1 | 1.23E-158 | 5.06074582 | 0.633 | 0.276 | 2.46E-155 |
| Proliferating Cells | Ttll9 | 3.76E-157 | 4.36047943 | 0.486 | 0.123 | 7.51E-154 |
| Proliferating Cells | Fli1 | 6.40E-87 | 0.9086007 | 0.905 | 0.623 | 1.28E-83 |
| Proliferating Cells | Tp63 | 3.82E-84 | 1.07097003 | 0.363 | 0.104 | 7.65E-81 |
| Proliferating Cells | Pcdh19 | 9.69E-73 | 1.71451429 | 0.699 | 0.413 | 1.94E-69 |
| Proliferating Cells | Adamts9 | 8.52E-68 | 1.51641327 | 0.693 | 0.382 | 1.70E-64 |

| | | | | | | |
|---|---|---|---|---|---|---|
| Proliferating Cells | Tmem150c | 1.39E-64 | 1.28687685 | 0.474 | 0.183 | 2.78E-61 |
| Proliferating Cells | Dok6 | 2.29E-58 | 0.47029456 | 0.413 | 0.137 | 4.57E-55 |
| Proliferating Cells | Meox2 | 4.52E-57 | 1.18908171 | 0.749 | 0.486 | 9.05E-54 |
| Proliferating Cells | Nkain2 | 3.28E-55 | 0.56444482 | 0.411 | 0.123 | 6.57E-52 |
| Proliferating Cells | Ptprb | 1.33E-54 | 0.71113071 | 0.816 | 0.487 | 2.66E-51 |
| Proliferating Cells | Mcf2l | 9.27E-52 | 1.12605857 | 0.677 | 0.381 | 1.85E-48 |
| Proliferating Cells | Dach1 | 2.50E-51 | 0.69018882 | 0.766 | 0.515 | 5.00E-48 |
| Proliferating Cells | Ptpn22 | 2.84E-48 | 0.50524038 | 0.41 | 0.152 | 5.68E-45 |
| Proliferating Cells | Wt1 | 1.25E-46 | 1.27061441 | 0.6 | 0.309 | 2.51E-43 |
| Proliferating Cells | Zfp366 | 2.77E-44 | 0.9267994 | 0.647 | 0.372 | 5.54E-41 |
| Proliferating Cells | Col18a1 | 1.75E-42 | 1.14526321 | 0.633 | 0.371 | 3.50E-39 |
| Proliferating Cells | Adgrf5 | 7.32E-41 | 0.60641723 | 0.757 | 0.46 | 1.46E-37 |
| Proliferating Cells | Cyyr1 | 1.71E-38 | 0.44868686 | 0.729 | 0.451 | 3.43E-35 |
| Proliferating Cells | Rassf4 | 5.63E-33 | 0.45715764 | 0.438 | 0.174 | 1.13E-29 |

| | | | | | | |
|---|---|---|---|---|---|---|
| Proliferating Cells | Flt1 | 1.29E-32 | 0.29445472 | 0.779 | 0.498 | 2.58E-29 |
| CM | AABR07052585.1 | 0 | 5.20585387 | 0.968 | 0.292 | 0 |
| CM | Obscn | 0 | 4.44600897 | 0.956 | 0.287 | 0 |
| CM | Rnf207 | 0 | 4.30772059 | 0.942 | 0.277 | 0 |
| CM | Asb2 | 0 | 4.2915272 | 0.936 | 0.281 | 0 |
| CM | Trim63 | 0 | 4.17258169 | 0.925 | 0.272 | 0 |
| CM | Tnni3k | 0 | 4.56323291 | 0.961 | 0.311 | 0 |
| CM | Mlip | 0 | 4.22957642 | 0.941 | 0.294 | 0 |
| CM | Speg | 0 | 4.01131783 | 0.92 | 0.277 | 0 |
| CM | Acacb | 0 | 4.0021746 | 0.96 | 0.326 | 0 |
| CM | Casq2 | 0 | 4.10710478 | 0.953 | 0.326 | 0 |
| CM | Myh7b | 0 | 4.63899618 | 0.901 | 0.279 | 0 |
| CM | Trdn | 0 | 3.64595958 | 0.917 | 0.298 | 0 |
| CM | Myocd | 0 | 4.12123738 | 0.908 | 0.292 | 0 |
| CM | Kcnd3 | 0 | 4.84051153 | 0.895 | 0.279 | 0 |
| CM | Fgf1 | 0 | 4.01745219 | 0.899 | 0.285 | 0 |
| CM | Mylk3 | 0 | 4.22568706 | 0.945 | 0.338 | 0 |
| CM | Coro6 | 0 | 4.26629621 | 0.851 | 0.25 | 0 |
| CM | Macrod1 | 0 | 4.22136684 | 0.899 | 0.3 | 0 |
| CM | Scn5a | 0 | 4.45915153 | 0.861 | 0.263 | 0 |
| CM | Fsd2 | 0 | 4.69166085 | 0.904 | 0.313 | 0 |
| CM | Myom1 | 0 | 3.7202716 | 0.93 | 0.344 | 0 |
| CM | Mybpc3 | 0 | 4.11615315 | 0.991 | 0.408 | 0 |
| CM | Wnk2 | 0 | 4.83142742 | 0.865 | 0.282 | 0 |
| CM | Shroom3 | 0 | 4.03618133 | 0.925 | 0.345 | 0 |
| CM | Myoz2 | 0 | 3.73590086 | 0.872 | 0.296 | 0 |
| CM | Ppargc1b | 0 | 3.20328597 | 0.929 | 0.354 | 0 |

| | | | | | | |
|---|---|---|---|---|---|---|
| CM | Ldb3 | 0 | 4.40788179 | 0.996 | 0.421 | 0 |
| CM | Fhl2 | 0 | 3.95027816 | 0.962 | 0.387 | 0 |
| CM | Myom2 | 0 | 4.30402378 | 0.866 | 0.293 | 0 |
| CM | Nebl | 0 | 2.01829666 | 0.945 | 0.373 | 0 |
| CM | Kcnq1 | 0 | 4.23491648 | 0.965 | 0.394 | 0 |
| CM | Nrap | 0 | 4.21693424 | 0.839 | 0.268 | 0 |
| CM | Rbfox1 | 0 | 3.74282376 | 0.889 | 0.322 | 0 |
| CM | Sgca | 0 | 4.20039487 | 0.832 | 0.266 | 0 |
| CM | Nav21 | 0 | 1.94325048 | 0.906 | 0.344 | 0 |
| CM | Actn2 | 0 | 3.34666725 | 0.933 | 0.376 | 0 |
| CM | Jph2 | 0 | 4.01354213 | 0.882 | 0.329 | 0 |
| CM | Cap2 | 0 | 3.41333216 | 0.868 | 0.317 | 0 |
| CM | Slc38a3 | 0 | 5.15528659 | 0.776 | 0.227 | 0 |
| CM | Ppara | 0 | 4.41427044 | 0.844 | 0.296 | 0 |
| CM | Ptpn3 | 0 | 3.09051321 | 0.909 | 0.364 | 0 |
| CM | AABR07034767.1 | 0 | 3.89244528 | 0.846 | 0.301 | 0 |
| CM | Ppargc1a | 0 | 4.03849137 | 0.789 | 0.245 | 0 |
| CM | Mapt | 0 | 3.4152637 | 0.831 | 0.287 | 0 |
| CM | Ccdc141 | 0 | 3.15588039 | 0.972 | 0.429 | 0 |
| CM | Veph1 | 0 | 3.97455182 | 0.79 | 0.248 | 0 |
| CM | Trim55 | 0 | 4.17837351 | 0.847 | 0.306 | 0 |
| CM | Nexn | 0 | 3.99899687 | 0.992 | 0.452 | 0 |
| CM | AABR07044900.1 | 0 | 2.78023782 | 0.902 | 0.364 | 0 |
| CM | Ppip5k2 | 0 | 2.7361884 | 0.93 | 0.392 | 0 |
| CM | Srl | 0 | 4.2910456 | 0.793 | 0.257 | 0 |
| CM | Mypn | 0 | 4.1998337 | 0.841 | 0.308 | 0 |
| CM | Slc38a1 | 0 | 3.54272806 | 0.856 | 0.324 | 0 |

| | | | | | | |
|---|---|---|---|---|---|---|
| CM | Alpk3 | 0 | 4.13193583 | 0.8 | 0.27 | 0 |
| CM | Ank3 | 0 | 2.09637763 | 0.944 | 0.415 | 0 |
| CM | Trim54 | 0 | 3.84701081 | 0.794 | 0.267 | 0 |
| CM | Akap6 | 0 | 3.29488007 | 0.965 | 0.438 | 0 |
| CM | Cux2 | 0 | 4.9425401 | 0.804 | 0.283 | 0 |
| CM | Sgcd | 0 | 3.03482776 | 0.945 | 0.424 | 0 |
| CM | Synpo2 | 0 | 3.1824987 | 0.959 | 0.439 | 0 |
| CM | Kcnj3 | 0 | 3.46570549 | 0.746 | 0.227 | 0 |
| CM | Tnni3 | 0 | 3.32459025 | 0.991 | 0.473 | 0 |
| CM | Txlnb | 0 | 4.38551876 | 0.842 | 0.324 | 0 |
| CM | Adra1a | 0 | 3.39815128 | 0.819 | 0.303 | 0 |
| CM | Dmpk | 0 | 3.3140061 | 0.976 | 0.465 | 0 |
| CM | Lrrc2 | 0 | 3.95603437 | 0.768 | 0.258 | 0 |
| CM | Ryr3 | 0 | 3.61445909 | 0.766 | 0.256 | 0 |
| CM | Myo18b | 0 | 4.00071839 | 0.802 | 0.292 | 0 |
| CM | Cacnb2 | 0 | 3.030434 | 0.963 | 0.455 | 0 |
| CM | Rilpl1 | 0 | 3.00190839 | 0.944 | 0.436 | 0 |
| CM | Fhod3 | 0 | 3.79563358 | 0.982 | 0.476 | 0 |
| CM | Phyh | 0 | 2.31378111 | 0.809 | 0.306 | 0 |
| CM | Cabcoco1 | 0 | 3.87633797 | 0.781 | 0.278 | 0 |
| CM | Dgkz | 0 | 2.26184412 | 0.886 | 0.383 | 0 |
| CM | Cacna1c | 0 | 2.84507983 | 0.96 | 0.46 | 0 |
| CM | Asb18 | 0 | 4.49211717 | 0.805 | 0.306 | 0 |
| CM | Unc45b | 0 | 4.6845645 | 0.728 | 0.229 | 0 |
| CM | Pdlim3 | 0 | 2.33231008 | 0.919 | 0.421 | 0 |
| CM | Cdh2 | 0 | 3.07947279 | 0.951 | 0.453 | 0 |
| CM | Me3 | 0 | 3.70355292 | 0.793 | 0.296 | 0 |
| CM | Bzw2 | 0 | 2.57203629 | 0.872 | 0.376 | 0 |
| CM | Naca | 0 | 2.52093402 | 0.835 | 0.339 | 0 |

| | | | | | | |
|---|---|---|---|---|---|---|
| CM | RGD1565355 | 0 | 2.0274303 | 0.924 | 0.43 | 0 |
| CM | Vegfa | 0 | 3.47482048 | 0.997 | 0.503 | 0 |
| CM | AC096301.1 | 0 | 4.50243451 | 0.748 | 0.259 | 0 |
| CM | Smyd1 | 0 | 4.27785465 | 0.757 | 0.275 | 0 |
| CM | Cmya5 | 0 | 2.82626092 | 0.865 | 0.386 | 0 |
| CM | Oxr1 | 0 | 2.59331827 | 0.938 | 0.459 | 0 |
| CM | Dtna | 0 | 2.90238925 | 0.775 | 0.298 | 0 |
| CM | Pde4dip | 0 | 3.8077228 | 0.994 | 0.517 | 0 |
| CM | Adcy5 | 0 | 2.45943634 | 0.838 | 0.363 | 0 |
| CM | Pln | 0 | 2.82938378 | 0.942 | 0.467 | 0 |
| CM | Rnf144b | 0 | 2.48075705 | 0.868 | 0.394 | 0 |
| CM | Hs3st5 | 0 | 3.91508594 | 0.736 | 0.263 | 0 |
| CM | Rbm24 | 0 | 3.87287504 | 0.739 | 0.267 | 0 |
| CM | Polr2m | 0 | 2.45359475 | 0.925 | 0.455 | 0 |
| CM | Fbxo40 | 0 | 5.00508408 | 0.627 | 0.157 | 0 |
| CM | Popdc2 | 0 | 3.88472447 | 0.772 | 0.303 | 0 |
| CM | Prkaa2 | 0 | 3.31671002 | 0.77 | 0.302 | 0 |
| CM | Cluh | 0 | 2.99392599 | 0.746 | 0.278 | 0 |
| CM | Abcc9 | 0 | 0.80429399 | 0.944 | 0.477 | 0 |
| CM | Limch1 | 0 | 1.77579127 | 0.911 | 0.445 | 0 |
| CM | Tnik | 0 | 2.31499403 | 0.834 | 0.368 | 0 |
| CM | Lmo7 | 0 | 2.29356512 | 0.977 | 0.513 | 0 |
| CM | Acsl1 | 0 | 3.02906055 | 0.773 | 0.312 | 0 |
| CM | Miga2 | 0 | 2.71451471 | 0.774 | 0.313 | 0 |
| CM | Nckap5 | 0 | 2.5954138 | 0.81 | 0.35 | 0 |
| CM | Sorbs2 | 0 | 2.2181435 | 0.972 | 0.513 | 0 |
| CM | Sox6 | 0 | 1.46921063 | 0.869 | 0.411 | 0 |

| | | | | | | |
|---|---|---|---|---|---|---|
| CM | Rap1gap2 | 0 | 2.5201437 | 0.767 | 0.313 | 0 |
| CM | Ky | 0 | 4.71424443 | 0.649 | 0.196 | 0 |
| CM | Xirp2 | 0 | 3.39702772 | 0.775 | 0.323 | 0 |
| CM | Iqub | 0 | 3.6440751 | 0.758 | 0.307 | 0 |
| CM | Nt5e | 0 | 1.85834071 | 0.765 | 0.314 | 0 |
| CM | Adk | 0 | 2.1533294 | 0.896 | 0.445 | 0 |
| CM | Rbm20 | 0 | 4.91932496 | 0.988 | 0.537 | 0 |
| CM | Ppp1r14c | 0 | 3.43480376 | 0.74 | 0.29 | 0 |
| CM | Inpp4b | 0 | 1.33924974 | 0.806 | 0.359 | 0 |
| CM | Gal3st3 | 0 | 5.11442026 | 0.658 | 0.211 | 0 |
| CM | Tbc1d4 | 0 | 1.99718241 | 0.908 | 0.461 | 0 |
| CM | Slc4a3 | 0 | 3.31276254 | 0.755 | 0.309 | 0 |
| CM | AABR07031740.1 | 0 | 3.66398343 | 0.7 | 0.254 | 0 |
| CM | Prodh1 | 0 | 4.17277266 | 0.674 | 0.229 | 0 |
| CM | Erbb4 | 0 | 3.71350844 | 0.657 | 0.213 | 0 |
| CM | Alpk2 | 0 | 4.44937087 | 0.683 | 0.239 | 0 |
| CM | LOC100361457 | 0 | 1.8890747 | 0.802 | 0.358 | 0 |
| CM | Tmem196 | 0 | 2.47795048 | 0.81 | 0.367 | 0 |
| CM | Actc1 | 0 | 1.98151672 | 0.918 | 0.476 | 0 |
| CM | Sptb | 0 | 4.39740233 | 0.739 | 0.297 | 0 |
| CM | Ivns1abp | 0 | 2.54648167 | 0.842 | 0.4 | 0 |
| CM | Pde3a | 0 | 1.32929882 | 0.962 | 0.521 | 0 |
| CM | AABR07026536.1 | 0 | 4.24478765 | 0.692 | 0.252 | 0 |
| CM | Sacs | 0 | 2.39037899 | 0.736 | 0.3 | 0 |
| CM | Pde4b | 0 | 2.28002637 | 0.885 | 0.449 | 0 |
| CM | Tmem182 | 0 | 3.98264393 | 0.665 | 0.23 | 0 |

| | | | | | | |
|---|---|---|---|---|---|---|
| CM | Spink8 | 0 | 3.34246037 | 0.694 | 0.259 | 0 |
| CM | Unc13b | 0 | 1.82344474 | 0.824 | 0.391 | 0 |
| CM | Abcc8 | 0 | 4.89331457 | 0.637 | 0.205 | 0 |
| CM | Usp13 | 0 | 2.57949814 | 0.774 | 0.343 | 0 |
| CM | Rgs6 | 0 | 3.79545807 | 0.647 | 0.219 | 0 |
| CM | Trim50 | 0 | 4.74563574 | 0.653 | 0.226 | 0 |
| CM | Ppp2r3a | 0 | 2.14732371 | 0.927 | 0.5 | 0 |
| CM | Ptprd | 0 | 1.72310098 | 0.899 | 0.472 | 0 |
| CM | Ndrg4 | 0 | 3.56339152 | 0.736 | 0.311 | 0 |
| CM | Myl3 | 0 | 2.17680945 | 0.963 | 0.539 | 0 |
| CM | Usp2 | 0 | 3.69317762 | 0.71 | 0.287 | 0 |
| CM | AABR07001054.2 | 0 | 5.07122726 | 0.684 | 0.262 | 0 |
| CM | Srpk3 | 0 | 4.7581117 | 0.647 | 0.225 | 0 |
| CM | Tango2 | 0 | 2.32348871 | 0.788 | 0.366 | 0 |
| CM | Parm1 | 0 | 3.40120622 | 0.692 | 0.27 | 0 |
| CM | Dsp | 0 | 2.67265861 | 0.738 | 0.317 | 0 |
| CM | Kcng2 | 0 | 4.16472278 | 0.723 | 0.303 | 0 |
| CM | Corin | 0 | 3.34159638 | 0.674 | 0.257 | 0 |
| CM | Lsamp | 0 | 3.4650089 | 0.666 | 0.251 | 0 |
| CM | Enah | 0 | 2.08143572 | 0.831 | 0.417 | 0 |
| CM | Ankrd9 | 0 | 4.91777029 | 0.622 | 0.209 | 0 |
| CM | AABR07001519.1 | 0 | 1.77346888 | 0.795 | 0.382 | 0 |
| CM | Kcnk3 | 0 | 4.20203545 | 0.608 | 0.196 | 0 |
| CM | Art3 | 0 | 1.86750801 | 0.722 | 0.31 | 0 |
| CM | Rcan2 | 0 | 2.41161135 | 0.966 | 0.554 | 0 |
| CM | AABR07044049.1 | 0 | 1.97496316 | 0.783 | 0.372 | 0 |

| | | | | | | |
|---|---|---|---|---|---|---|
| CM | Pcbp3 | 0 | 2.44078992 | 0.732 | 0.321 | 0 |
| CM | Svil | 0 | 1.73368927 | 0.916 | 0.506 | 0 |
| CM | Lims2 | 0 | 1.54242871 | 0.859 | 0.45 | 0 |
| CM | Sntb1 | 0 | 1.83583722 | 0.804 | 0.396 | 0 |
| CM | Mitf | 0 | 1.55377364 | 0.883 | 0.476 | 0 |
| CM | Oxct1 | 0 | 2.21182465 | 0.879 | 0.473 | 0 |
| CM | Grb14 | 0 | 2.21284118 | 0.773 | 0.368 | 0 |
| CM | Wipf3 | 0 | 1.90203423 | 0.808 | 0.403 | 0 |
| CM | Tesc | 0 | 3.53047893 | 0.692 | 0.287 | 0 |
| CM | Pygm | 0 | 3.80657446 | 0.69 | 0.286 | 0 |
| CM | Glb1l2 | 0 | 4.36343124 | 0.669 | 0.266 | 0 |
| CM | Pfkfb2 | 0 | 2.89818097 | 0.721 | 0.318 | 0 |
| CM | Esrrg | 0 | 2.05493659 | 0.824 | 0.425 | 0 |
| CM | AABR07017268.1 | 0 | 4.66310094 | 0.593 | 0.194 | 0 |
| CM | Ankrd1 | 0 | 3.22854889 | 0.97 | 0.572 | 0 |
| CM | Cryab | 0 | 1.8736238 | 0.788 | 0.394 | 0 |
| CM | Cadps | 0 | 3.14414581 | 0.667 | 0.273 | 0 |
| CM | Epb41l4b | 0 | 3.00978223 | 0.669 | 0.275 | 0 |
| CM | Tnnt2 | 0 | 3.40241056 | 0.997 | 0.604 | 0 |
| CM | Mpp7 | 0 | 1.39916182 | 0.73 | 0.34 | 0 |
| CM | Sh3kbp1 | 0 | 1.97190759 | 0.835 | 0.446 | 0 |
| CM | Trabd2b | 0 | 1.84951964 | 0.775 | 0.387 | 0 |
| CM | Des | 0 | 1.37253245 | 0.745 | 0.358 | 0 |
| CM | Tmem116 | 0 | 4.30317572 | 0.595 | 0.208 | 0 |
| CM | Lrrc4b | 0 | 3.99775469 | 0.601 | 0.215 | 0 |
| CM | Zbtb16 | 0 | 1.74926347 | 0.872 | 0.486 | 0 |
| CM | Gata4 | 0 | 1.35074843 | 0.862 | 0.477 | 0 |
| CM | Dgkb | 0 | 1.4322086 | 0.803 | 0.419 | 0 |

| | | | | | | |
|---|---|---|---|---|---|---|
| CM | Slco5a1 | 0 | 4.2006273 | 0.628 | 0.246 | 0 |
| CM | Adhfe1 | 0 | 3.86242981 | 0.65 | 0.269 | 0 |
| CM | Slc25a13 | 0 | 2.46603061 | 0.696 | 0.316 | 0 |
| CM | AABR07025 295.1 | 0 | 1.48746814 | 0.922 | 0.543 | 0 |
| CM | AABR07052 523.2 | 0 | 4.54854933 | 0.614 | 0.238 | 0 |
| CM | Ckmt2 | 0 | 1.84688149 | 0.749 | 0.373 | 0 |
| CM | LOC691485 | 0 | 3.73135877 | 0.57 | 0.194 | 0 |
| CM | Adra1b | 0 | 4.13868427 | 0.658 | 0.283 | 0 |
| CM | Dpf3 | 0 | 3.33266827 | 0.67 | 0.295 | 0 |
| CM | Gria3 | 0 | 1.27862737 | 0.807 | 0.432 | 0 |
| CM | Trim7 | 0 | 3.70307047 | 0.622 | 0.249 | 0 |
| CM | Snta1 | 0 | 2.17113588 | 0.721 | 0.348 | 0 |
| CM | Kank1 | 0 | 1.37691851 | 0.766 | 0.394 | 0 |
| CM | Trak2 | 0 | 1.89621292 | 0.876 | 0.504 | 0 |
| CM | Myl2 | 0 | 1.77776765 | 0.952 | 0.581 | 0 |
| CM | Nuak1 | 0 | 1.18755206 | 0.836 | 0.465 | 0 |
| CM | Tbx20 | 0 | 1.49340234 | 0.792 | 0.422 | 0 |
| CM | Ldhb | 0 | 1.8795536 | 0.791 | 0.421 | 0 |
| CM | Ros1 | 0 | 4.56578717 | 0.626 | 0.257 | 0 |
| CM | AABR07007 026.1 | 0 | 4.36544652 | 0.567 | 0.199 | 0 |
| CM | Lgr6 | 0 | 2.47886392 | 0.704 | 0.337 | 0 |
| CM | Mtss1 | 0 | 1.30040644 | 0.926 | 0.56 | 0 |
| CM | Ryr2 | 0 | 4.40620221 | 0.994 | 0.629 | 0 |
| CM | Myh7 | 0 | 2.78153904 | 0.919 | 0.554 | 0 |
| CM | Cpe | 0 | 2.50898534 | 0.691 | 0.326 | 0 |
| CM | Ckm | 0 | 1.93225843 | 0.7 | 0.336 | 0 |

| | | | | | | |
|---|---|---|---|---|---|---|
| CM | Cpt1b | 0 | 3.51248071 | 0.567 | 0.203 | 0 |
| CM | Slc20a2 | 0 | 1.82800158 | 0.759 | 0.396 | 0 |
| CM | Asb15 | 0 | 4.50290557 | 0.626 | 0.264 | 0 |
| CM | AC110709.2 | 0 | 4.92981236 | 0.58 | 0.219 | 0 |
| CM | Foxp2 | 0 | 1.12078469 | 0.749 | 0.388 | 0 |
| CM | Arhgap21 | 0 | 1.79959061 | 0.928 | 0.568 | 0 |
| CM | Pla2g5 | 0 | 4.41241144 | 0.553 | 0.194 | 0 |
| CM | Slc4a4 | 0 | 1.1456325 | 0.725 | 0.366 | 0 |
| CM | Ank2 | 0 | 1.26592327 | 0.946 | 0.588 | 0 |
| CM | Clu | 0 | 1.65190112 | 0.658 | 0.302 | 0 |
| CM | Arhgef37 | 0 | 3.05737535 | 0.643 | 0.287 | 0 |
| CM | Coq8a | 0 | 3.32243644 | 0.592 | 0.236 | 0 |
| CM | AABR07049292.1 | 0 | 4.06726439 | 0.561 | 0.208 | 0 |
| CM | Acot11 | 0 | 3.24558751 | 0.665 | 0.312 | 0 |
| CM | Shb | 0 | 1.59793743 | 0.772 | 0.42 | 0 |
| CM | Gja1 | 0 | 2.07498445 | 0.744 | 0.394 | 0 |
| CM | Fbxo32 | 0 | 2.40166797 | 0.684 | 0.335 | 0 |
| CM | Asb14 | 0 | 4.42563427 | 0.57 | 0.221 | 0 |
| CM | Atp2b2 | 0 | 3.88171911 | 0.62 | 0.272 | 0 |
| CM | AC134204.1 | 0 | 0.89690274 | 0.782 | 0.435 | 0 |
| CM | Vcl | 0 | 1.33665331 | 0.904 | 0.557 | 0 |
| CM | Amd1 | 0 | 1.47197571 | 0.749 | 0.402 | 0 |
| CM | Asb9 | 0 | 3.83237977 | 0.549 | 0.203 | 0 |
| CM | Shisal1 | 0 | 2.30321215 | 0.456 | 0.111 | 0 |
| CM | Gpcpd1 | 0 | 2.05839355 | 0.747 | 0.402 | 0 |
| CM | Rmdn1 | 0 | 1.65178512 | 0.814 | 0.469 | 0 |

| | | | | | | |
|---|---|---|---|---|---|---|
| CM | Dgki | 0 | 2.342632 | 0.609 | 0.265 | 0 |
| CM | Arhgap44 | 0 | 2.13747672 | 0.662 | 0.319 | 0 |
| CM | Fbxw10 | 0 | 4.58812624 | 0.466 | 0.124 | 0 |
| CM | Sfxn5 | 0 | 3.18909029 | 0.622 | 0.281 | 0 |
| CM | Ppp1r12b | 0 | 1.79977082 | 0.953 | 0.612 | 0 |
| CM | Synpo2l | 0 | 4.21317524 | 0.589 | 0.249 | 0 |
| CM | AABR07035916.1 | 0 | 1.31614446 | 0.961 | 0.621 | 0 |
| CM | Pip5k1b | 0 | 1.05522982 | 0.613 | 0.274 | 0 |
| CM | Btbd11 | 0 | 1.42459424 | 0.633 | 0.297 | 0 |
| CM | Ddc | 0 | 3.90513853 | 0.5 | 0.164 | 0 |
| CM | Fabp3 | 0 | 2.02436077 | 0.63 | 0.295 | 0 |
| CM | Acyp2 | 0 | 1.91121436 | 0.712 | 0.377 | 0 |
| CM | Gpm6a | 0 | 1.0702595 | 0.804 | 0.47 | 0 |
| CM | Pkia | 0 | 1.95088659 | 0.68 | 0.346 | 0 |
| CM | Sphkap | 0 | 4.12793495 | 0.614 | 0.28 | 0 |
| CM | Tecrl | 0 | 3.80762398 | 0.595 | 0.261 | 0 |
| CM | Mlf1 | 0 | 3.80086677 | 0.599 | 0.265 | 0 |
| CM | Slc16a1 | 0 | 2.82984321 | 0.617 | 0.284 | 0 |
| CM | Lpl | 0 | 1.50897026 | 0.754 | 0.421 | 0 |
| CM | Pam | 0 | 1.26612383 | 0.91 | 0.579 | 0 |
| CM | Flnc | 0 | 2.9305607 | 0.582 | 0.254 | 0 |
| CM | Bcl11a | 0 | 3.58585776 | 0.543 | 0.217 | 0 |
| CM | Sgcg | 0 | 3.24328917 | 0.595 | 0.27 | 0 |
| CM | Fign | 0 | 1.16313855 | 0.644 | 0.32 | 0 |
| CM | Slc16a10 | 0 | 2.44172653 | 0.599 | 0.275 | 0 |
| CM | Gnao1 | 0 | 1.81748428 | 0.697 | 0.373 | 0 |
| CM | Acad10 | 0 | 2.18925647 | 0.622 | 0.303 | 0 |
| CM | Nppb | 0 | 2.59788687 | 0.569 | 0.25 | 0 |

| | | | | | | |
|---|---|---|---|---|---|---|
| CM | AC111831.1 | 0 | 4.38402205 | 0.417 | 0.099 | 0 |
| CM | Ankh | 0 | 1.60210786 | 0.719 | 0.401 | 0 |
| CM | Ppp1r3a | 0 | 3.60491666 | 0.523 | 0.207 | 0 |
| CM | Pcdh7 | 0 | 2.96319671 | 0.632 | 0.317 | 0 |
| CM | Hk2 | 0 | 1.15615205 | 0.769 | 0.455 | 0 |
| CM | Pdzrn3 | 0 | 0.39227743 | 0.943 | 0.629 | 0 |
| CM | Rbm38 | 0 | 3.84988166 | 0.557 | 0.244 | 0 |
| CM | Csrp3 | 0 | 2.30134 | 0.595 | 0.283 | 0 |
| CM | Ano5 | 0 | 3.34017657 | 0.524 | 0.213 | 0 |
| CM | Dcdc5 | 0 | 3.69691509 | 0.547 | 0.237 | 0 |
| CM | Pde7b | 0 | 0.89741348 | 0.922 | 0.615 | 0 |
| CM | Syt14 | 0 | 3.91951469 | 0.532 | 0.227 | 0 |
| CM | Sorbs1 | 0 | 2.83465317 | 0.99 | 0.685 | 0 |
| CM | Myom3 | 0 | 2.79328381 | 0.39 | 0.086 | 0 |
| CM | Clybl | 0 | 1.57451477 | 0.671 | 0.368 | 0 |
| CM | Itga7 | 0 | 2.06735709 | 0.579 | 0.276 | 0 |
| CM | Ptpn13 | 0 | 1.14943192 | 0.672 | 0.371 | 0 |
| CM | Fgf13 | 0 | 3.81786256 | 0.627 | 0.326 | 0 |
| CM | AABR07007032.1 | 0 | 1.92330897 | 0.987 | 0.689 | 0 |
| CM | Rimbp2 | 0 | 2.01411057 | 0.645 | 0.348 | 0 |
| CM | Fam189a2 | 0 | 3.16747966 | 0.485 | 0.188 | 0 |
| CM | AC130940.1 | 0 | 5.44384823 | 0.456 | 0.161 | 0 |
| CM | Sypl2 | 0 | 2.81753966 | 0.311 | 0.018 | 0 |
| CM | Smpx | 0 | 2.81538559 | 0.581 | 0.29 | 0 |
| CM | Ablim2 | 0 | 3.46442184 | 0.56 | 0.27 | 0 |
| CM | Ralgapa2 | 0 | 0.61269069 | 0.883 | 0.597 | 0 |

| | | | | | | |
|---|---|---|---|---|---|---|
| CM | Prox1 | 0 | 1.61121716 | 0.548 | 0.263 | 0 |
| CM | Eva1c | 0 | 1.15180199 | 0.665 | 0.38 | 0 |
| CM | Slc30a3 | 0 | 4.66784136 | 0.439 | 0.155 | 0 |
| CM | Dsg2 | 0 | 3.02555983 | 0.539 | 0.255 | 0 |
| CM | Homer1 | 0 | 1.99300096 | 0.611 | 0.327 | 0 |
| CM | Tox3 | 0 | 1.68943637 | 0.646 | 0.363 | 0 |
| CM | Bdnf | 0 | 3.68112096 | 0.516 | 0.234 | 0 |
| CM | Armc4 | 0 | 2.30468902 | 0.335 | 0.053 | 0 |
| CM | Myo3b | 0 | 4.206509 | 0.4 | 0.12 | 0 |
| CM | Atp2a2 | 0 | 2.56670229 | 0.988 | 0.709 | 0 |
| CM | Nmrk2 | 0 | 3.28779283 | 0.444 | 0.165 | 0 |
| CM | Bcl11b | 0 | 1.10722425 | 0.393 | 0.115 | 0 |
| CM | Tpd52l1 | 0 | 3.33487115 | 0.503 | 0.225 | 0 |
| CM | Dusp27 | 0 | 3.86086597 | 0.519 | 0.242 | 0 |
| CM | Myo16 | 0 | 3.90394853 | 0.415 | 0.138 | 0 |
| CM | Rxfp1 | 0 | 3.55684867 | 0.466 | 0.19 | 0 |
| CM | Lamc2 | 0 | 3.21898183 | 0.499 | 0.223 | 0 |
| CM | Gbp1 | 0 | 4.03353271 | 0.525 | 0.251 | 0 |
| CM | Tpm1 | 0 | 2.41237094 | 0.992 | 0.72 | 0 |
| CM | Mtus2 | 0 | 3.22494154 | 0.554 | 0.283 | 0 |
| CM | Frmd5 | 0 | 3.07875604 | 0.569 | 0.298 | 0 |
| CM | Pank1 | 0 | 3.00851117 | 0.593 | 0.323 | 0 |
| CM | Pdlim5 | 0 | 1.38349818 | 0.941 | 0.671 | 0 |
| CM | Hpd | 0 | 4.19436618 | 0.343 | 0.074 | 0 |
| CM | Reep1 | 0 | 2.50920069 | 0.591 | 0.322 | 0 |
| CM | Fgf12 | 0 | 1.7779769 | 0.597 | 0.329 | 0 |
| CM | Lrrc7 | 0 | 4.09352225 | 0.506 | 0.239 | 0 |
| CM | Col28a1 | 0 | 1.79042794 | 0.311 | 0.044 | 0 |
| CM | Kcnj5 | 0 | 4.24470043 | 0.455 | 0.195 | 0 |

| | | | | | | |
|---|---|---|---|---|---|---|
| CM | Fgf7 | 0 | 1.61347792 | 0.365 | 0.106 | 0 |
| CM | Etv3l | 0 | 2.91898019 | 0.26 | 0.002 | 0 |
| CM | Tspan18 | 0 | 0.70878806 | 0.794 | 0.537 | 0 |
| CM | Efhd1 | 0 | 1.45147776 | 0.573 | 0.316 | 0 |
| CM | Dmd | 0 | 3.43285956 | 0.986 | 0.733 | 0 |
| CM | Ntn4 | 6.90E-305 | 1.05597052 | 0.656 | 0.346 | 1.38E-301 |
| CM | Clic5 | 6.21E-304 | 0.59203182 | 0.724 | 0.373 | 1.24E-300 |
| CM | Tbx5 | 1.06E-300 | 2.98293817 | 0.476 | 0.207 | 2.12E-297 |
| CM | Hspb6 | 5.50E-297 | 1.71815432 | 0.583 | 0.266 | 1.10E-293 |
| CM | Insr | 1.23E-293 | 0.57992373 | 0.753 | 0.488 | 2.46E-290 |
| CM | Zfp536 | 2.88E-292 | 0.41802756 | 0.383 | 0.133 | 5.76E-289 |
| CM | Rtn4rl1 | 1.75E-287 | 1.86090488 | 0.54 | 0.26 | 3.49E-284 |
| CM | Atp1b1 | 1.18E-283 | 1.46713456 | 0.593 | 0.339 | 2.36E-280 |
| CM | Fam78b | 1.42E-280 | 1.49284377 | 0.554 | 0.254 | 2.84E-277 |
| CM | Slco3a1 | 4.74E-280 | 0.48148602 | 0.824 | 0.529 | 9.49E-277 |
| CM | Jph1 | 1.99E-277 | 2.33254059 | 0.534 | 0.266 | 3.98E-274 |
| CM | Sema5a | 2.46E-276 | 1.3322471 | 0.568 | 0.257 | 4.92E-273 |
| CM | Mctp1 | 7.47E-271 | 0.66629642 | 0.635 | 0.377 | 1.49E-267 |
| CM | Tnnc1 | 2.26E-263 | 1.27685287 | 0.573 | 0.307 | 4.51E-260 |
| CM | Abcb4 | 6.19E-262 | 0.9416841 | 0.62 | 0.356 | 1.24E-258 |
| CM | Baiap2l1 | 1.26E-261 | 1.38325097 | 0.542 | 0.252 | 2.53E-258 |
| CM | Lrrc10 | 1.30E-246 | 2.41114404 | 0.513 | 0.26 | 2.60E-243 |
| CM | Crim1 | 8.76E-220 | 0.26884404 | 0.839 | 0.542 | 1.75E-216 |
| CM | Mb | 2.53E-210 | 0.61126299 | 0.829 | 0.56 | 5.06E-207 |
| CM | Slc25a4 | 3.15E-205 | 0.61744583 | 0.757 | 0.501 | 6.29E-202 |
| CM | Acta1 | 4.26E-200 | 1.43072273 | 0.549 | 0.267 | 8.52E-197 |
| CM | Stxbp6 | 5.93E-179 | 0.40840482 | 0.776 | 0.525 | 1.19E-175 |
| CM | Serf2 | 7.46E-152 | 0.59767468 | 0.638 | 0.383 | 1.49E-148 |
| EC | Cyyr11 | 0 | 3.58018881 | 0.94 | 0.3 | 0 |

| | | | | | | |
|---|---|---|---|---|---|---|
| EC | Ptprb1 | 0 | 3.14126969 | 0.949 | 0.347 | 0 |
| EC | Flt11 | 0 | 3.63091086 | 0.943 | 0.362 | 0 |
| EC | Shank3 | 0 | 3.04683661 | 0.958 | 0.386 | 0 |
| EC | Adgrf51 | 0 | 3.13614693 | 0.898 | 0.327 | 0 |
| EC | Adgrl4 | 0 | 3.09232114 | 0.786 | 0.276 | 0 |
| EC | Prkch | 0 | 2.65928472 | 0.881 | 0.377 | 0 |
| EC | Emcn | 0 | 2.76589136 | 0.804 | 0.305 | 0 |
| EC | Dach11 | 0 | 2.8240626 | 0.895 | 0.399 | 0 |
| EC | Ccdc85a | 0 | 2.94800429 | 0.802 | 0.349 | 0 |
| EC | Ablim3 | 0 | 3.42768685 | 0.765 | 0.319 | 0 |
| EC | Zfp3661 | 0 | 3.66233218 | 0.715 | 0.269 | 0 |
| EC | Myrip | 0 | 3.45775188 | 0.766 | 0.326 | 0 |
| EC | Etl4 | 0 | 2.85080444 | 0.959 | 0.519 | 0 |
| EC | Kitlg | 0 | 2.8524877 | 0.828 | 0.391 | 0 |
| EC | Cadps2 | 0 | 3.00144757 | 0.702 | 0.292 | 0 |
| EC | Meox21 | 0 | 2.72059688 | 0.799 | 0.392 | 0 |
| EC | Mcf2l1 | 0 | 3.20467985 | 0.69 | 0.29 | 0 |
| EC | Fli11 | 0 | 2.20587087 | 0.929 | 0.532 | 0 |
| EC | Thsd7a | 0 | 3.30290605 | 0.703 | 0.306 | 0 |
| EC | Dnm3 | 0 | 2.65491891 | 0.854 | 0.459 | 0 |
| EC | Anxa3 | 0 | 2.93294377 | 0.663 | 0.273 | 0 |
| EC | Slc26a10 | 0 | 3.81886179 | 0.684 | 0.296 | 0 |
| EC | Podxl | 0 | 3.00805561 | 0.673 | 0.287 | 0 |
| EC | Spns2 | 0 | 2.74103261 | 0.636 | 0.263 | 0 |
| EC | Wt11 | 0 | 2.62416464 | 0.594 | 0.226 | 0 |
| EC | Mgll | 0 | 2.77714771 | 0.799 | 0.433 | 0 |
| EC | Cdh13 | 0 | 2.61477812 | 0.985 | 0.621 | 0 |
| EC | AABR07007642.1 | 0 | 3.79707512 | 0.642 | 0.283 | 0 |

| | | | | | | |
|---|---|---|---|---|---|---|
| EC | Itpkb | 0 | 2.63806856 | 0.893 | 0.534 | 0 |
| EC | Cmtm8 | 0 | 3.34837358 | 0.598 | 0.241 | 0 |
| EC | Efnb2 | 0 | 2.76299669 | 0.66 | 0.305 | 0 |
| EC | Plcb1 | 0 | 2.85154788 | 0.726 | 0.374 | 0 |
| EC | Dipk2b | 0 | 3.54541237 | 0.609 | 0.262 | 0 |
| EC | Nav3 | 0 | 2.72354917 | 0.747 | 0.4 | 0 |
| EC | Adamts91 | 0 | 2.22617799 | 0.636 | 0.309 | 0 |
| EC | Vwf | 0 | 2.0017944 | 0.548 | 0.224 | 0 |
| EC | Ppp1r16b | 0 | 2.88565188 | 0.631 | 0.315 | 0 |
| EC | Hmcn1 | 0 | 1.25760309 | 0.835 | 0.521 | 0 |
| EC | Clic51 | 0 | 2.15695456 | 0.658 | 0.353 | 0 |
| EC | Akr1c15 | 0 | 2.5295808 | 0.657 | 0.355 | 0 |
| EC | St6galnac3 | 0 | 2.53653868 | 0.627 | 0.325 | 0 |
| EC | Rasa4 | 0 | 2.82682556 | 0.615 | 0.317 | 0 |
| EC | Myo10 | 0 | 2.64448489 | 0.699 | 0.407 | 0 |
| EC | Chn1 | 0 | 1.57954838 | 0.613 | 0.325 | 0 |
| EC | Pcdh17 | 0 | 3.13445118 | 0.523 | 0.241 | 0 |
| EC | LOC103694210 | 0 | 2.37572573 | 0.463 | 0.185 | 0 |
| EC | Gabra4 | 0 | 0.9944225 | 0.374 | 0.097 | 0 |
| EC | Prom1 | 0 | 3.55307722 | 0.548 | 0.273 | 0 |
| EC | Aqp1 | 0 | 2.4060888 | 0.511 | 0.24 | 0 |
| EC | Rgcc | 0 | 3.09807081 | 0.531 | 0.265 | 0 |
| EC | Palmd | 0 | 2.50662788 | 0.637 | 0.372 | 0 |
| EC | Fabp4 | 0 | 2.15308774 | 0.571 | 0.306 | 0 |
| EC | Tmtc2 | 0 | 1.9749459 | 0.609 | 0.349 | 0 |
| EC | Ccser1 | 0 | 3.12305208 | 0.537 | 0.279 | 0 |
| EC | Exoc3l2 | 0 | 3.67428257 | 0.309 | 0.054 | 0 |

| | | | | | | |
|---|---|---|---|---|---|---|
| Endocardial ECs | Pkhd1l1 | 0 | 3.86199117 | 0.934 | 0.255 | 0 |
| Endocardial ECs | Cgnl1 | 0 | 5.75533757 | 0.977 | 0.335 | 0 |
| Endocardial ECs | Vwf1 | 0 | 3.74648812 | 0.882 | 0.286 | 0 |
| Endocardial ECs | Chn2 | 0 | 3.31692925 | 0.935 | 0.387 | 0 |
| Endocardial ECs | Mmp24 | 0 | 5.46404351 | 0.586 | 0.049 | 0 |
| Endocardial ECs | Il1a | 0 | 6.18400656 | 0.543 | 0.015 | 0 |
| Endocardial ECs | Sncaip | 0 | 4.20811185 | 0.864 | 0.352 | 0 |
| Endocardial ECs | Cdh11 | 0 | 3.76598023 | 0.94 | 0.432 | 0 |
| Endocardial ECs | Reg3g | 0 | 6.6084606 | 0.605 | 0.098 | 0 |
| Endocardial ECs | Reg3b | 0 | 4.70975804 | 0.553 | 0.07 | 0 |
| Endocardial ECs | Pgm5 | 0 | 3.83877688 | 0.879 | 0.415 | 0 |
| Endocardial ECs | Hecw2 | 0 | 2.9100465 | 0.931 | 0.467 | 0 |
| Endocardial ECs | Zmat4 | 0 | 7.38328048 | 0.591 | 0.129 | 0 |
| Endocardial ECs | Cemip2 | 0 | 3.51994198 | 0.928 | 0.474 | 0 |

| | | | | | | |
|---|---|---|---|---|---|---|
| Endocardia c ECs | Cpvl | 0 | 5.85622581 | 0.502 | 0.051 | 0 |
| Endocardia c ECs | Nkain4 | 0 | 1.26821991 | 0.495 | 0.051 | 0 |
| Endocardia c ECs | Ltbp1 | 0 | 2.702852 | 0.942 | 0.562 | 0 |
| Endocardia c ECs | LOC499542 | 0 | 4.84437463 | 0.38 | 0.002 | 0 |
| Endocardia c ECs | Shisa6 | 0 | 1.3557949 | 0.425 | 0.053 | 0 |
| Endocardia c ECs | Hmcn11 | 0 | 3.39113231 | 0.947 | 0.587 | 0 |
| Endocardia c ECs | Hapln1 | 0 | 6.42489223 | 0.376 | 0.023 | 0 |
| Endocardia c ECs | Chst9 | 0 | 4.84800499 | 0.435 | 0.089 | 0 |
| Endocardia c ECs | Trpc5 | 3.44E-302 | 6.1097877 | 0.384 | 0.099 | 6.88E-299 |
| Endocardia c ECs | Susd4 | 2.82E-290 | 4.56886934 | 0.438 | 0.084 | 5.63E-287 |
| Endocardia c ECs | Nrg1 | 7.92E-258 | 5.58297123 | 0.777 | 0.285 | 1.58E-254 |
| Endocardia c ECs | Erc2 | 5.51E-256 | 4.04528772 | 0.396 | 0.08 | 1.10E-252 |
| Endocardia c ECs | Dscaml1 | 1.12E-253 | 1.93338175 | 0.402 | 0.074 | 2.23E-250 |
| Endocardia c ECs | Bmx | 6.80E-252 | 3.82704924 | 0.527 | 0.142 | 1.36E-248 |

| | | | | | | |
|---|---|---|---|---|---|---|
| Endocardia c ECs | Slc9a9 | 5.77E-251 | 1.99911957 | 0.91 | 0.556 | 1.15E-247 |
| Endocardia c ECs | Ctnnd2 | 5.89E-220 | 3.03073033 | 0.518 | 0.11 | 1.18E-216 |
| Endocardia c ECs | Plvap | 2.07E-219 | 3.30987783 | 0.396 | 0.074 | 4.14E-216 |
| Endocardia c ECs | Tnni2 | 2.57E-215 | 2.09497312 | 0.356 | 0.065 | 5.13E-212 |
| Endocardia c ECs | Dok61 | 4.62E-208 | 2.37957593 | 0.532 | 0.132 | 9.24E-205 |
| Endocardia c ECs | Lancl3 | 1.38E-206 | 2.72254008 | 0.499 | 0.14 | 2.77E-203 |
| Endocardia c ECs | Npr3 | 2.27E-204 | 4.70978274 | 0.664 | 0.297 | 4.54E-201 |
| Endocardia c ECs | Ass1 | 3.73E-199 | 1.52744707 | 0.492 | 0.137 | 7.47E-196 |
| Endocardia c ECs | AABR07005 821.1 | 3.52E-191 | 1.24626052 | 0.371 | 0.055 | 7.04E-188 |
| Endocardia c ECs | Rgs20 | 1.34E-188 | 2.31068902 | 0.428 | 0.09 | 2.68E-185 |
| Endocardia c ECs | Hmga2 | 3.15E-188 | 2.72167913 | 0.385 | 0.081 | 6.30E-185 |
| Endocardia c ECs | Acer2 | 1.19E-184 | 3.00429309 | 0.754 | 0.319 | 2.38E-181 |
| Endocardia c ECs | Edil3 | 5.50E-184 | 4.56156215 | 0.506 | 0.15 | 1.10E-180 |
| Endocardia c ECs | LOC100361 087 | 8.02E-181 | 2.83786536 | 0.761 | 0.388 | 1.60E-177 |

| | | | | | | |
|---|---|---|---|---|---|---|
| Endocardiac ECs | Ntn1 | 2.93E-174 | 2.02408491 | 0.839 | 0.532 | 5.86E-171 |
| Endocardiac ECs | Itgb4 | 4.23E-169 | 2.8578184 | 0.658 | 0.253 | 8.45E-166 |
| Endocardiac ECs | Emcn1 | 8.89E-166 | 1.8064563 | 0.792 | 0.416 | 1.78E-162 |
| Endocardiac ECs | Opcml | 3.60E-165 | 2.85673065 | 0.686 | 0.255 | 7.19E-162 |
| Endocardiac ECs | Mettl24 | 1.12E-161 | 3.70744882 | 0.459 | 0.169 | 2.24E-158 |
| Endocardiac ECs | St6galnac31 | 6.67E-161 | 1.85610157 | 0.719 | 0.389 | 1.33E-157 |
| Endocardiac ECs | Bmp6 | 3.02E-160 | 3.1334581 | 0.73 | 0.396 | 6.05E-157 |
| Endocardiac ECs | Plxnc1 | 3.20E-159 | 3.18854968 | 0.631 | 0.222 | 6.39E-156 |
| Endocardiac ECs | Vcam1 | 3.85E-157 | 3.71549834 | 0.643 | 0.232 | 7.70E-154 |
| Endocardiac ECs | Ifitm10 | 2.82E-152 | 2.42072731 | 0.635 | 0.217 | 5.64E-149 |
| Endocardiac ECs | Klhl29 | 2.83E-152 | 2.27434016 | 0.69 | 0.364 | 5.65E-149 |
| Endocardiac ECs | Selp | 1.87E-148 | 1.39014865 | 0.339 | 0.068 | 3.74E-145 |
| Endocardiac ECs | Nckap51 | 3.62E-146 | 2.1663851 | 0.787 | 0.411 | 7.23E-143 |
| Endocardiac ECs | Plxna4 | 9.33E-144 | 1.48121382 | 0.829 | 0.521 | 1.87E-140 |

| | | | | | | |
|---|---|---|---|---|---|---|
| Endocardia c ECs | Pawr | 5.50E-140 | 2.89414926 | 0.631 | 0.305 | 1.10E-136 |
| Endocardia c ECs | Bace2 | 3.48E-139 | 2.57496212 | 0.649 | 0.369 | 6.95E-136 |
| Endocardia c ECs | Ralgapa21 | 3.63E-139 | 1.38396586 | 0.889 | 0.634 | 7.26E-136 |
| Endocardia c ECs | Smad6 | 1.17E-131 | 2.39372441 | 0.708 | 0.402 | 2.34E-128 |
| Endocardia c ECs | Eva1c1 | 2.34E-130 | 1.9321873 | 0.756 | 0.415 | 4.67E-127 |
| Endocardia c ECs | Cntn3 | 1.17E-128 | 1.70926455 | 0.41 | 0.101 | 2.35E-125 |
| Endocardia c ECs | Ano2 | 1.31E-128 | 2.3121355 | 0.608 | 0.24 | 2.63E-125 |
| Endocardia c ECs | Mt3 | 4.88E-128 | 0.47799898 | 0.347 | 0.04 | 9.76E-125 |
| Endocardia c ECs | Thsd7b | 6.82E-126 | 3.6353088 | 0.503 | 0.174 | 1.36E-122 |
| Endocardia c ECs | Pde3a1 | 9.08E-124 | 1.05137994 | 0.873 | 0.581 | 1.82E-120 |
| Endocardia c ECs | Hamp | 2.10E-121 | 1.53032482 | 0.415 | 0.133 | 4.20E-118 |
| Endocardia c ECs | Slc16a7 | 3.52E-121 | 2.12151793 | 0.657 | 0.273 | 7.03E-118 |
| Endocardia c ECs | Fli12 | 8.90E-119 | 1.08739706 | 0.917 | 0.621 | 1.78E-115 |
| Endocardia c ECs | Ednrb | 1.08E-116 | 2.3443285 | 0.609 | 0.285 | 2.16E-113 |

| | | | | | | |
|---|---|---|---|---|---|---|
| Endocardiac ECs | Ptprb2 | 1.19E-112 | 0.84834979 | 0.904 | 0.482 | 2.38E-109 |
| Endocardiac ECs | LOC100911486 | 1.67E-110 | 2.23178411 | 0.589 | 0.25 | 3.33E-107 |
| Endocardiac ECs | LOC1036942101 | 1.31E-109 | 1.56116774 | 0.586 | 0.243 | 2.62E-106 |
| Endocardiac ECs | Gask1b | 1.19E-105 | 1.84002344 | 0.707 | 0.386 | 2.37E-102 |
| Endocardiac ECs | Zdbf2 | 1.34E-105 | 0.839975 | 0.331 | 0.047 | 2.68E-102 |
| Endocardiac ECs | P3h2 | 7.28E-104 | 1.66938623 | 0.729 | 0.384 | 1.46E-100 |
| Endocardiac ECs | Shank31 | 2.13E-101 | 0.70130603 | 0.859 | 0.516 | 4.27E-98 |
| Endocardiac ECs | Lifr | 3.88E-100 | 1.89871195 | 0.725 | 0.377 | 7.75E-97 |
| Endocardiac ECs | AABR07041096.1 | 5.69E-100 | 1.60955291 | 0.556 | 0.203 | 1.14E-96 |
| Endocardiac ECs | Adgrf52 | 4.77E-99 | 0.88659801 | 0.797 | 0.457 | 9.53E-96 |
| Endocardiac ECs | Tmem63c | 4.01E-98 | 0.60300047 | 0.477 | 0.202 | 8.02E-95 |
| Endocardiac ECs | Tacr1 | 1.63E-96 | 2.09786457 | 0.361 | 0.111 | 3.25E-93 |
| Endocardiac ECs | Slco2a1 | 1.24E-94 | 1.4635431 | 0.46 | 0.208 | 2.48E-91 |
| Endocardiac ECs | Hs3st2 | 7.75E-93 | 1.32922829 | 0.47 | 0.157 | 1.55E-89 |

| | | | | | | |
|---|---|---|---|---|---|---|
| Endocardiac ECs | Pdgfc | 1.96E-90 | 2.20096891 | 0.541 | 0.232 | 3.91E-87 |
| Endocardiac ECs | Sh3gl3 | 1.41E-86 | 0.89292986 | 0.469 | 0.185 | 2.81E-83 |
| Endocardiac ECs | Prkch1 | 7.06E-84 | 0.77646387 | 0.81 | 0.491 | 1.41E-80 |
| Endocardiac ECs | Grid2 | 5.62E-83 | 1.52660196 | 0.464 | 0.152 | 1.12E-79 |
| Endocardiac ECs | Megf11 | 6.53E-81 | 1.95752827 | 0.516 | 0.237 | 1.31E-77 |
| Endocardiac ECs | Myrip1 | 3.33E-79 | 1.12728205 | 0.678 | 0.426 | 6.66E-76 |
| Endocardiac ECs | Plcg2 | 3.05E-78 | 0.58696435 | 0.538 | 0.243 | 6.10E-75 |
| Endocardiac ECs | Adamts17 | 8.86E-76 | 1.72990095 | 0.631 | 0.38 | 1.77E-72 |
| Endocardiac ECs | Flt12 | 3.48E-75 | 0.30393497 | 0.859 | 0.494 | 6.96E-72 |
| Endocardiac ECs | Tmem100 | 1.46E-74 | 2.48527276 | 0.523 | 0.267 | 2.92E-71 |
| Endocardiac ECs | Slc25a21 | 8.67E-72 | 0.38360268 | 0.503 | 0.177 | 1.73E-68 |
| Endocardiac ECs | Slc7a1 | 3.78E-69 | 1.53371028 | 0.68 | 0.403 | 7.56E-66 |
| Endocardiac ECs | Fam189a21 | 9.66E-69 | 1.19845916 | 0.493 | 0.226 | 1.93E-65 |
| Endocardiac ECs | Vcan | 5.41E-68 | 1.75000149 | 0.698 | 0.403 | 1.08E-64 |

| | | | | | | |
|---|---|---|---|---|---|---|
| Endocardiac ECs | AABR07006724.1 | 4.62E-67 | 1.30797009 | 0.593 | 0.283 | 9.24E-64 |
| Endocardiac ECs | Cyyr12 | 1.37E-64 | 0.28710548 | 0.811 | 0.446 | 2.73E-61 |
| Endocardiac ECs | Icam1 | 2.94E-63 | 1.88005392 | 0.527 | 0.25 | 5.88E-60 |
| Endocardiac ECs | Tec | 1.80E-62 | 1.0642896 | 0.645 | 0.378 | 3.59E-59 |
| Endocardiac ECs | Aldh1a1 | 1.87E-55 | 0.91504639 | 0.534 | 0.272 | 3.74E-52 |
| Endocardiac ECs | Grm1 | 6.31E-51 | 0.52395214 | 0.378 | 0.113 | 1.26E-47 |
| Endocardiac ECs | Aqp11 | 6.29E-49 | 0.77588883 | 0.587 | 0.298 | 1.26E-45 |
| Endocardiac ECs | Gcnt2 | 1.20E-45 | 0.9127794 | 0.6 | 0.344 | 2.40E-42 |
| Endocardiac ECs | Nebl1 | 2.07E-45 | 0.51092759 | 0.723 | 0.454 | 4.14E-42 |
| Endocardiac ECs | Ctsk | 1.04E-44 | 0.73618701 | 0.536 | 0.273 | 2.09E-41 |
| Endocardiac ECs | Sema5a1 | 4.34E-44 | 0.59208924 | 0.61 | 0.296 | 8.67E-41 |
| Endocardiac ECs | Spp1 | 1.07E-42 | 0.4507356 | 0.565 | 0.255 | 2.14E-39 |
| Endocardiac ECs | Lsamp1 | 1.36E-34 | 0.88043853 | 0.572 | 0.308 | 2.71E-31 |
| Endocardiac ECs | Parm11 | 4.21E-34 | 0.31043734 | 0.586 | 0.328 | 8.42E-31 |

| | | | | | | |
|---|---|---|---|---|---|---|
| Lymphatic ECs | Reln | 0 | 7.12653422 | 0.969 | 0.207 | 0 |
| Lymphatic ECs | Dtx1 | 0 | 9.05191442 | 0.773 | 0.042 | 0 |
| Lymphatic ECs | Mmrn1 | 0 | 7.44443953 | 0.832 | 0.132 | 0 |
| Lymphatic ECs | Flt4 | 0 | 5.52913094 | 0.922 | 0.241 | 0 |
| Lymphatic ECs | Ccl21 | 0 | 6.75588768 | 0.87 | 0.233 | 0 |
| Lymphatic ECs | Ldb2 | 0 | 3.19730446 | 0.929 | 0.343 | 0 |
| Lymphatic ECs | Sh3gl31 | 0 | 6.82318968 | 0.749 | 0.181 | 0 |
| Lymphatic ECs | Tbx1 | 0 | 7.39565849 | 0.6 | 0.032 | 0 |
| Lymphatic ECs | Mpp71 | 0 | 4.19522728 | 0.951 | 0.388 | 0 |
| Lymphatic ECs | Apba2 | 0 | 7.11048081 | 0.636 | 0.093 | 0 |
| Lymphatic ECs | Pex5l | 0 | 6.07096134 | 0.586 | 0.085 | 0 |
| Lymphatic ECs | Cst6 | 0 | 6.11116267 | 0.475 | 0.019 | 0 |
| Lymphatic ECs | Fam205a | 0 | 8.55787211 | 0.46 | 0.011 | 0 |
| Lymphatic ECs | Rspo3 | 0 | 4.26857133 | 0.479 | 0.041 | 0 |

| | | | | | | |
|---|---|---|---|---|---|---|
| Lymphatic ECs | Elavl2 | 0 | 2.01241061 | 0.365 | 0.03 | 0 |
| Lymphatic ECs | AABR07058412.1 | 0 | 4.10767293 | 0.32 | 0.009 | 0 |
| Lymphatic ECs | Nsg2 | 0 | 6.1573432 | 0.329 | 0.052 | 0 |
| Lymphatic ECs | Grin2b | 2.37E-307 | 6.02801537 | 0.595 | 0.087 | 4.74E-304 |
| Lymphatic ECs | Sema3a | 1.44E-298 | 5.9078832 | 0.837 | 0.25 | 2.88E-295 |
| Lymphatic ECs | Nrp2 | 1.66E-289 | 2.81905026 | 0.949 | 0.479 | 3.31E-286 |
| Lymphatic ECs | Pde7b1 | 1.79E-270 | 2.53658169 | 0.956 | 0.656 | 3.57E-267 |
| Lymphatic ECs | Prox11 | 2.20E-264 | 4.89171044 | 0.789 | 0.296 | 4.39E-261 |
| Lymphatic ECs | Dmtn | 7.64E-245 | 5.15399758 | 0.717 | 0.155 | 1.53E-241 |
| Lymphatic ECs | Sema3d | 4.20E-242 | 3.67944617 | 0.834 | 0.328 | 8.39E-239 |
| Lymphatic ECs | Nxn | 6.03E-232 | 2.35512395 | 0.908 | 0.55 | 1.21E-228 |
| Lymphatic ECs | Pkhd1l11 | 8.60E-230 | 2.92932182 | 0.817 | 0.262 | 1.72E-226 |
| Lymphatic ECs | Fcgr3a | 1.52E-220 | 2.28703506 | 0.357 | 0.022 | 3.04E-217 |
| Lymphatic ECs | Adgrg3 | 4.91E-217 | 5.31489106 | 0.67 | 0.258 | 9.83E-214 |

| | | | | | | |
|---|---|---|---|---|---|---|
| Lymphatic ECs | Shisal11 | 8.00E-210 | 4.49038974 | 0.533 | 0.156 | 1.60E-206 |
| Lymphatic ECs | Prkcz | 2.23E-208 | 4.38134055 | 0.73 | 0.283 | 4.47E-205 |
| Lymphatic ECs | Ralgapa22 | 3.57E-194 | 2.11429796 | 0.904 | 0.635 | 7.14E-191 |
| Lymphatic ECs | Spns21 | 5.27E-193 | 2.62118858 | 0.811 | 0.344 | 1.05E-189 |
| Lymphatic ECs | Abi3bp | 1.25E-187 | 1.95874693 | 0.92 | 0.553 | 2.50E-184 |
| Lymphatic ECs | Dapk2 | 1.94E-182 | 3.30688924 | 0.791 | 0.371 | 3.89E-179 |
| Lymphatic ECs | Tspan5 | 2.73E-181 | 2.23770027 | 0.873 | 0.457 | 5.46E-178 |
| Lymphatic ECs | AABR07049085.1 | 4.26E-180 | 2.58926803 | 0.825 | 0.334 | 8.52E-177 |
| Lymphatic ECs | Tspan181 | 4.85E-180 | 2.33666341 | 0.88 | 0.57 | 9.70E-177 |
| Lymphatic ECs | Dapk1 | 2.67E-179 | 2.06735983 | 0.85 | 0.5 | 5.35E-176 |
| Lymphatic ECs | Lyve1 | 4.39E-176 | 4.2164313 | 0.672 | 0.226 | 8.79E-173 |
| Lymphatic ECs | Unc5b | 1.24E-175 | 2.5236419 | 0.789 | 0.42 | 2.49E-172 |
| Lymphatic ECs | Mctp11 | 2.76E-160 | 2.86032178 | 0.791 | 0.409 | 5.52E-157 |
| Lymphatic ECs | Shank32 | 4.71E-159 | 1.2168378 | 0.957 | 0.516 | 9.42E-156 |

| | | | | | | |
|---|---|---|---|---|---|---|
| Lymphatic ECs | Slc45a3 | 2.74E-149 | 4.41378221 | 0.547 | 0.183 | 5.48E-146 |
| Lymphatic ECs | Pdpn | 8.55E-143 | 4.02036331 | 0.631 | 0.192 | 1.71E-139 |
| Lymphatic ECs | Lyn | 4.01E-134 | 1.5029194 | 0.885 | 0.554 | 8.02E-131 |
| Lymphatic ECs | Adgrl41 | 7.49E-131 | 1.52188026 | 0.848 | 0.391 | 1.50E-127 |
| Lymphatic ECs | Flrt2 | 1.09E-125 | 2.0834932 | 0.777 | 0.428 | 2.18E-122 |
| Lymphatic ECs | Prss12 | 2.03E-121 | 3.98784377 | 0.479 | 0.131 | 4.05E-118 |
| Lymphatic ECs | Ston2 | 1.36E-118 | 1.82547937 | 0.779 | 0.471 | 2.71E-115 |
| Lymphatic ECs | Rgs201 | 3.79E-118 | 2.56501268 | 0.452 | 0.092 | 7.57E-115 |
| Lymphatic ECs | Mfsd4a | 1.81E-117 | 5.33921911 | 0.529 | 0.181 | 3.62E-114 |
| Lymphatic ECs | Plxdc2 | 6.27E-111 | 1.15898801 | 0.913 | 0.63 | 1.25E-107 |
| Lymphatic ECs | Podnl1 | 1.82E-110 | 0.37268038 | 0.366 | 0.082 | 3.64E-107 |
| Lymphatic ECs | Polr2m1 | 1.49E-109 | 1.55061939 | 0.822 | 0.521 | 2.98E-106 |
| Lymphatic ECs | Col18a11 | 7.80E-109 | 2.2510565 | 0.702 | 0.37 | 1.56E-105 |
| Lymphatic ECs | Klhl4 | 4.76E-106 | 3.33846204 | 0.656 | 0.249 | 9.52E-103 |

| | | | | | | |
|---|---|---|---|---|---|---|
| Lymphatic ECs | Frmd4b | 1.78E-103 | 1.38301578 | 0.828 | 0.545 | 3.56E-100 |
| Lymphatic ECs | Mdga2 | 9.17E-103 | 4.38257845 | 0.532 | 0.18 | 1.83E-99 |
| Lymphatic ECs | Ephb1 | 2.49E-102 | 3.22212494 | 0.628 | 0.295 | 4.98E-99 |
| Lymphatic ECs | Camk4 | 4.18E-101 | 0.89320255 | 0.428 | 0.165 | 8.37E-98 |
| Lymphatic ECs | Ptprb3 | 5.84E-101 | 0.76692022 | 0.908 | 0.485 | 1.17E-97 |
| Lymphatic ECs | Ntn11 | 7.28E-100 | 1.82037271 | 0.79 | 0.535 | 1.46E-96 |
| Lymphatic ECs | Chst15 | 3.60E-98 | 2.28013843 | 0.68 | 0.33 | 7.21E-95 |
| Lymphatic ECs | Npnt | 6.14E-91 | 2.5257207 | 0.504 | 0.208 | 1.23E-87 |
| Lymphatic ECs | Il1rl1 | 6.43E-91 | 1.99374791 | 0.388 | 0.125 | 1.29E-87 |
| Lymphatic ECs | Hecw21 | 1.99E-90 | 1.36226086 | 0.793 | 0.474 | 3.99E-87 |
| Lymphatic ECs | Gria2 | 2.69E-83 | 3.13966011 | 0.404 | 0.151 | 5.39E-80 |
| Lymphatic ECs | Rai14 | 2.95E-83 | 1.38208448 | 0.806 | 0.472 | 5.90E-80 |
| Lymphatic ECs | Cgnl11 | 1.39E-81 | 1.16750965 | 0.626 | 0.347 | 2.77E-78 |
| Lymphatic ECs | Dnm31 | 3.44E-81 | 0.95107189 | 0.826 | 0.55 | 6.88E-78 |

| | | | | | | |
|---|---|---|---|---|---|---|
| Lymphatic ECs | Tnfaip8l3 | 1.47E-77 | 2.95854612 | 0.508 | 0.229 | 2.95E-74 |
| Lymphatic ECs | Ptgs1 | 1.66E-76 | 2.54838616 | 0.594 | 0.269 | 3.33E-73 |
| Lymphatic ECs | Ube2ql1 | 4.30E-75 | 2.59200782 | 0.448 | 0.173 | 8.60E-72 |
| Lymphatic ECs | AABR07027581.1 | 2.13E-73 | 1.77829082 | 0.642 | 0.385 | 4.26E-70 |
| Lymphatic ECs | Emid1 | 2.34E-73 | 3.12057166 | 0.513 | 0.211 | 4.67E-70 |
| Lymphatic ECs | Stard10 | 3.62E-72 | 3.95784857 | 0.437 | 0.115 | 7.24E-69 |
| Lymphatic ECs | LOC1009114861 | 1.31E-71 | 3.12429726 | 0.523 | 0.253 | 2.62E-68 |
| Lymphatic ECs | Olfml2a | 2.02E-70 | 3.03588153 | 0.524 | 0.207 | 4.04E-67 |
| Lymphatic ECs | Prkch2 | 4.61E-69 | 0.79341701 | 0.84 | 0.493 | 9.21E-66 |
| Lymphatic ECs | P3h21 | 5.94E-69 | 1.76304107 | 0.697 | 0.387 | 1.19E-65 |
| Lymphatic ECs | Mapk10 | 2.95E-68 | 2.06956176 | 0.459 | 0.184 | 5.90E-65 |
| Lymphatic ECs | Trpc6 | 1.63E-67 | 1.1483944 | 0.488 | 0.207 | 3.26E-64 |
| Lymphatic ECs | Fam189a1 | 2.81E-67 | 3.61717919 | 0.524 | 0.234 | 5.63E-64 |
| Lymphatic ECs | Osbp2 | 2.08E-65 | 2.40075185 | 0.604 | 0.342 | 4.16E-62 |

| | | | | | | |
|---|---|---|---|---|---|---|
| Lymphatic ECs | St6galnac32 | 1.91E-57 | 1.26322669 | 0.651 | 0.393 | 3.83E-54 |
| Lymphatic ECs | Ankh1 | 2.98E-55 | 1.66181434 | 0.714 | 0.444 | 5.97E-52 |
| Lymphatic ECs | Inpp4b1 | 3.69E-44 | 1.05130421 | 0.694 | 0.422 | 7.39E-41 |
| Lymphatic ECs | Thsd7a1 | 1.69E-43 | 0.67706404 | 0.671 | 0.397 | 3.38E-40 |
| Lymphatic ECs | Aqp12 | 6.40E-41 | 1.07860312 | 0.618 | 0.299 | 1.28E-37 |
| Lymphatic ECs | Podxl1 | 6.91E-41 | 0.84281346 | 0.67 | 0.375 | 1.38E-37 |
| Lymphatic ECs | Ptprv | 3.22E-37 | 0.60352612 | 0.488 | 0.212 | 6.43E-34 |
| Lymphatic ECs | Cp | 3.03E-36 | 1.15396596 | 0.588 | 0.312 | 6.05E-33 |
| Lymphatic ECs | Ahr | 1.92E-35 | 1.03861656 | 0.65 | 0.384 | 3.83E-32 |
| Lymphatic ECs | Aff3 | 9.65E-33 | 0.72560486 | 0.678 | 0.426 | 1.93E-29 |
| Lymphatic ECs | Ccdc1411 | 6.28E-27 | 0.37963736 | 0.778 | 0.508 | 1.26E-23 |
| Fibs | Clmp | 0 | 3.29514724 | 0.807 | 0.269 | 0 |
| Fibs | C1qtnf7 | 0 | 3.04921651 | 0.894 | 0.373 | 0 |
| Fibs | Ebf2 | 0 | 2.47575707 | 0.863 | 0.373 | 0 |
| Fibs | Adamts2 | 0 | 2.47473931 | 0.883 | 0.406 | 0 |
| Fibs | Pdgfra | 0 | 3.2063929 | 0.769 | 0.301 | 0 |
| Fibs | Fndc1 | 0 | 2.74254313 | 0.852 | 0.385 | 0 |
| Fibs | Itgbl1 | 0 | 2.94640697 | 0.879 | 0.413 | 0 |

| | | | | | | |
|---|---|---|---|---|---|---|
| Fibs | Dlg2 | 0 | 3.18990676 | 0.752 | 0.29 | 0 |
| Fibs | C7 | 0 | 3.08795438 | 0.899 | 0.441 | 0 |
| Fibs | Zfp385d | 0 | 2.13075362 | 0.9 | 0.444 | 0 |
| Fibs | Bnc2 | 0 | 2.66606703 | 0.747 | 0.292 | 0 |
| Fibs | Mrc2 | 0 | 3.21027095 | 0.732 | 0.293 | 0 |
| Fibs | Ccdc80 | 0 | 2.50686111 | 0.895 | 0.474 | 0 |
| Fibs | Dcn | 0 | 2.46904355 | 0.795 | 0.375 | 0 |
| Fibs | Col14a1 | 0 | 2.86449052 | 0.722 | 0.306 | 0 |
| Fibs | Postn | 0 | 2.64494409 | 0.776 | 0.363 | 0 |
| Fibs | Glis3 | 0 | 2.42171391 | 0.77 | 0.36 | 0 |
| Fibs | Antxr1 | 0 | 2.67161878 | 0.791 | 0.386 | 0 |
| Fibs | Tmeff2 | 0 | 3.06129159 | 0.69 | 0.289 | 0 |
| Fibs | Dclk1 | 0 | 2.403513 | 0.723 | 0.326 | 0 |
| Fibs | Gpc6 | 0 | 2.59610523 | 0.98 | 0.587 | 0 |
| Fibs | Col6a6 | 0 | 3.15934224 | 0.737 | 0.348 | 0 |
| Fibs | Dpt | 0 | 2.39164432 | 0.722 | 0.335 | 0 |
| Fibs | Col1a1 | 0 | 2.02475533 | 0.876 | 0.491 | 0 |
| Fibs | Loxl1 | 0 | 2.89226783 | 0.69 | 0.305 | 0 |
| Fibs | Cacna2d3 | 0 | 2.89741395 | 0.645 | 0.262 | 0 |
| Fibs | Mfap5 | 0 | 2.88180449 | 0.7 | 0.319 | 0 |
| Fibs | Vcan1 | 0 | 2.1445457 | 0.674 | 0.301 | 0 |
| Fibs | Col8a1 | 0 | 2.79959395 | 0.874 | 0.502 | 0 |
| Fibs | Ston21 | 0 | 2.18469404 | 0.74 | 0.368 | 0 |
| Fibs | Aff31 | 0 | 2.09933645 | 0.694 | 0.322 | 0 |
| Fibs | Abi3bp1 | 0 | 1.86913857 | 0.822 | 0.452 | 0 |
| Fibs | Egfr | 0 | 2.4510072 | 0.714 | 0.347 | 0 |
| Fibs | Medag | 0 | 3.12352154 | 0.626 | 0.263 | 0 |
| Fibs | Adamtsl3 | 0 | 2.67383453 | 0.735 | 0.375 | 0 |
| Fibs | Plekha7 | 0 | 2.70364751 | 0.667 | 0.31 | 0 |

| | | | | | | |
|---|---|---|---|---|---|---|
| Fibs | Timp2 | 0 | 1.91397172 | 0.792 | 0.436 | 0 |
| Fibs | Col6a2 | 0 | 1.88110266 | 0.676 | 0.324 | 0 |
| Fibs | Plod2 | 0 | 1.99857146 | 0.754 | 0.402 | 0 |
| Fibs | Adgrd1 | 0 | 2.75091316 | 0.632 | 0.283 | 0 |
| Fibs | Col5a2 | 0 | 1.7234427 | 0.822 | 0.473 | 0 |
| Fibs | Cdh111 | 0 | 0.80599892 | 0.69 | 0.344 | 0 |
| Fibs | Boc | 0 | 2.22975437 | 0.76 | 0.416 | 0 |
| Fibs | Cpq | 0 | 1.99631593 | 0.757 | 0.413 | 0 |
| Fibs | Prrx1 | 0 | 1.83883497 | 0.687 | 0.345 | 0 |
| Fibs | Pdzrn31 | 0 | 1.83363732 | 0.918 | 0.576 | 0 |
| Fibs | Gda | 0 | 2.03536947 | 0.76 | 0.418 | 0 |
| Fibs | Fbn1 | 0 | 2.57519688 | 0.939 | 0.6 | 0 |
| Fibs | Fstl1 | 0 | 1.94900746 | 0.821 | 0.484 | 0 |
| Fibs | Fbln1 | 0 | 2.87277269 | 0.616 | 0.285 | 0 |
| Fibs | Axl | 0 | 2.41416357 | 0.684 | 0.356 | 0 |
| Fibs | Galnt17 | 0 | 2.57459265 | 0.669 | 0.346 | 0 |
| Fibs | Pdgfrb | 0 | 1.05299058 | 0.712 | 0.394 | 0 |
| Fibs | Gxylt2 | 0 | 2.63843014 | 0.721 | 0.406 | 0 |
| Fibs | Eda | 0 | 2.1264304 | 0.668 | 0.355 | 0 |
| Fibs | Robo1 | 0 | 2.36170115 | 0.665 | 0.354 | 0 |
| Fibs | Sdk2 | 0 | 2.60731922 | 0.58 | 0.271 | 0 |
| Fibs | Pcdha13 | 0 | 2.25705926 | 0.629 | 0.321 | 0 |
| Fibs | Pde1a | 0 | 2.47291892 | 0.66 | 0.354 | 0 |
| Fibs | Galnt16 | 0 | 2.60751119 | 0.624 | 0.318 | 0 |
| Fibs | Ank21 | 0 | 1.13151419 | 0.857 | 0.553 | 0 |
| Fibs | Pid1 | 0 | 0.70025269 | 0.803 | 0.503 | 0 |
| Fibs | RGD1563354 | 0 | 3.32630808 | 0.537 | 0.24 | 0 |
| Fibs | Asap3 | 0 | 2.26872227 | 0.678 | 0.386 | 0 |

| | | | | | | |
|---|---|---|---|---|---|---|
| Fibs | Spon1 | 0 | 2.46879392 | 0.54 | 0.249 | 0 |
| Fibs | Npas2 | 0 | 2.61874949 | 0.607 | 0.319 | 0 |
| Fibs | Tenm3 | 0 | 1.98262925 | 0.574 | 0.288 | 0 |
| Fibs | Gli2 | 0 | 1.98384351 | 0.596 | 0.313 | 0 |
| Fibs | Il16 | 0 | 1.99540097 | 0.533 | 0.252 | 0 |
| Fibs | Mgp | 0 | 1.66167252 | 0.802 | 0.525 | 0 |
| Fibs | Bmper | 0 | 3.18462968 | 0.545 | 0.268 | 0 |
| Fibs | Gfpt2 | 0 | 3.1651644 | 0.58 | 0.306 | 0 |
| Fibs | AABR07054 614.1 | 0 | 2.30262438 | 0.571 | 0.299 | 0 |
| Fibs | Adamts19 | 0 | 2.23340759 | 0.649 | 0.381 | 0 |
| Fibs | Srgap3 | 0 | 2.16026668 | 0.619 | 0.352 | 0 |
| Fibs | Ror2 | 0 | 2.42412075 | 0.513 | 0.247 | 0 |
| Fibs | Heyl | 0 | 2.12046419 | 0.544 | 0.278 | 0 |
| Fibs | Col4a5 | 0 | 1.85295356 | 0.732 | 0.467 | 0 |
| Fibs | Pcsk6 | 0 | 2.06810145 | 0.691 | 0.426 | 0 |
| Fibs | AABR07059 258.1 | 0 | 2.09964677 | 0.553 | 0.29 | 0 |
| Fibs | Col3a1 | 0 | 2.06745953 | 0.968 | 0.705 | 0 |
| Fibs | Tgfb3 | 0 | 2.51720081 | 0.515 | 0.252 | 0 |
| Fibs | Plxdc21 | 0 | 1.26934313 | 0.819 | 0.559 | 0 |
| Fibs | Fn1 | 0 | 1.49365691 | 0.629 | 0.373 | 0 |
| Fibs | Itga8 | 0 | 2.06962856 | 0.6 | 0.344 | 0 |
| Fibs | Mid1 | 0 | 1.85365792 | 0.595 | 0.341 | 0 |
| Fibs | Has1 | 0 | 3.11330184 | 0.396 | 0.146 | 0 |
| Fibs | Gpm6b | 0 | 2.44063438 | 0.531 | 0.281 | 0 |
| IC | Ptprc | 0 | 4.2613731 | 0.843 | 0.278 | 0 |
| IC | Tbxas1 | 0 | 5.48832093 | 0.796 | 0.255 | 0 |
| IC | Fyb1 | 0 | 4.76250859 | 0.807 | 0.283 | 0 |

| | | | | | | |
|---|---|---|---|---|---|---|
| IC | Cfh | 0 | 3.88390521 | 0.809 | 0.324 | 0 |
| IC | LOC690045 | 0 | 5.09787217 | 0.783 | 0.3 | 0 |
| IC | Dock8 | 0 | 3.74711314 | 0.837 | 0.354 | 0 |
| IC | Arhgap15 | 0 | 3.69629352 | 0.872 | 0.418 | 0 |
| IC | Inpp5d | 0 | 3.68454147 | 0.809 | 0.364 | 0 |
| IC | Mrc1 | 0 | 4.88794866 | 0.71 | 0.265 | 0 |
| IC | Csf1r | 0 | 5.19467878 | 0.688 | 0.248 | 0 |
| IC | Il6r | 0 | 3.21431434 | 0.797 | 0.362 | 0 |
| IC | Hck | 0 | 5.55824097 | 0.63 | 0.202 | 0 |
| IC | Aoah | 0 | 5.51315373 | 0.702 | 0.276 | 0 |
| IC | Ly86 | 0 | 5.43922793 | 0.635 | 0.211 | 0 |
| IC | Arhgap22 | 0 | 4.02398488 | 0.775 | 0.353 | 0 |
| IC | Ikzf1 | 0 | 4.48349406 | 0.667 | 0.255 | 0 |
| IC | Cd300a | 0 | 5.30048387 | 0.619 | 0.209 | 0 |
| IC | Lilrb4 | 0 | 4.96898375 | 0.633 | 0.224 | 0 |
| IC | Sh3bp2 | 0 | 4.49394281 | 0.709 | 0.302 | 0 |
| IC | Camk1d | 0 | 3.34614159 | 0.733 | 0.332 | 0 |
| IC | Pik3r5 | 0 | 5.03278254 | 0.591 | 0.199 | 0 |
| IC | Lgmn | 0 | 3.73588724 | 0.718 | 0.326 | 0 |
| IC | Cd4 | 0 | 5.43671012 | 0.608 | 0.232 | 0 |
| IC | Ptprj | 0 | 2.21014448 | 0.835 | 0.459 | 0 |
| IC | Fermt3 | 0 | 4.22292223 | 0.614 | 0.243 | 0 |
| IC | Myo1f | 0 | 3.52656364 | 0.673 | 0.303 | 0 |
| IC | Syk | 0 | 4.40350501 | 0.609 | 0.242 | 0 |
| IC | Ptpro | 0 | 5.07217403 | 0.605 | 0.24 | 0 |
| IC | Ctss | 0 | 5.25709269 | 0.662 | 0.299 | 0 |
| IC | Frmd4b1 | 0 | 2.75602086 | 0.88 | 0.517 | 0 |
| IC | Lyn1 | 0 | 2.24334368 | 0.888 | 0.527 | 0 |
| IC | LOC690097 | 0 | 5.29108504 | 0.551 | 0.191 | 0 |

| | | | | | | |
|---|---|---|---|---|---|---|
| IC | Cd84 | 0 | 5.12654364 | 0.472 | 0.113 | 0 |
| IC | Mertk | 0 | 3.16009192 | 0.633 | 0.275 | 0 |
| IC | Kcnk13 | 0 | 5.17652279 | 0.651 | 0.3 | 0 |
| IC | Slc9a91 | 0 | 3.03868598 | 0.88 | 0.533 | 0 |
| IC | Gna15 | 0 | 5.26261811 | 0.599 | 0.252 | 0 |
| IC | Adgre1 | 0 | 5.50862447 | 0.491 | 0.146 | 0 |
| IC | Pid11 | 0 | 1.91178944 | 0.902 | 0.559 | 0 |
| IC | Otulinl | 0 | 4.86337687 | 0.593 | 0.25 | 0 |
| IC | Itgb2 | 0 | 4.71319506 | 0.481 | 0.139 | 0 |
| IC | Nckap1l | 0 | 4.82698961 | 0.578 | 0.241 | 0 |
| IC | Cass4 | 0 | 5.1481722 | 0.424 | 0.088 | 0 |
| IC | Lcp2 | 0 | 3.2591054 | 0.605 | 0.27 | 0 |
| IC | Cd300lb | 0 | 5.62880291 | 0.374 | 0.04 | 0 |
| IC | Lcp1 | 0 | 4.32462519 | 0.615 | 0.286 | 0 |
| IC | Alox5 | 0 | 5.26583714 | 0.48 | 0.153 | 0 |
| IC | Cyth4 | 0 | 4.68673324 | 0.58 | 0.255 | 0 |
| IC | Abca1 | 0 | 2.45525684 | 0.765 | 0.441 | 0 |
| IC | Cers6 | 0 | 2.68307469 | 0.738 | 0.416 | 0 |
| IC | Runx1 | 0 | 2.52975162 | 0.758 | 0.437 | 0 |
| IC | Stab1 | 0 | 4.12063604 | 0.655 | 0.334 | 0 |
| IC | Marchf1 | 0 | 4.92454615 | 0.518 | 0.203 | 0 |
| IC | Epsti1 | 0 | 4.93011036 | 0.476 | 0.168 | 0 |
| IC | Fgd2 | 0 | 4.98496324 | 0.537 | 0.23 | 0 |
| IC | Colec12 | 0 | 2.23573755 | 0.749 | 0.442 | 0 |
| IC | Blnk | 0 | 4.02273483 | 0.643 | 0.337 | 0 |
| IC | Prkcb | 0 | 4.61782525 | 0.594 | 0.293 | 0 |
| IC | Bin2 | 0 | 4.38296783 | 0.57 | 0.27 | 0 |
| IC | AABR07001573.2 | 0 | 5.19224789 | 0.385 | 0.086 | 0 |

| | | | | | | |
|---|---|---|---|---|---|---|
| IC | Kcnt2 | 0 | 3.4389913 | 0.662 | 0.367 | 0 |
| IC | Ciita | 0 | 4.79327268 | 0.405 | 0.11 | 0 |
| IC | Grk3 | 0 | 4.63037466 | 0.516 | 0.225 | 0 |
| IC | Lilrb3 | 0 | 5.7807703 | 0.303 | 0.018 | 0 |
| IC | Apba1 | 0 | 2.64626574 | 0.735 | 0.451 | 0 |
| IC | Mtss11 | 0 | 1.52700968 | 0.867 | 0.591 | 0 |
| IC | Zeb2 | 0 | 1.82255335 | 0.961 | 0.687 | 0 |
| IC | Alcam | 0 | 2.67778988 | 0.662 | 0.39 | 0 |
| IC | Hk3 | 0 | 5.26689443 | 0.332 | 0.063 | 0 |
| IC | Hk21 | 0 | 2.34861371 | 0.743 | 0.479 | 0 |
| IC | Abca17 | 8.80E-303 | 5.33120392 | 0.534 | 0.247 | 1.76E-299 |
| IC | Asgr2 | 6.19E-300 | 4.76697343 | 0.464 | 0.162 | 1.24E-296 |
| IC | Slc43a2 | 1.88E-298 | 3.83100498 | 0.602 | 0.287 | 3.76E-295 |
| IC | Plek | 4.57E-297 | 4.61475114 | 0.348 | 0.077 | 9.14E-294 |
| IC | Atp8b4 | 1.05E-291 | 4.33782324 | 0.545 | 0.25 | 2.11E-288 |
| IC | Apobec1 | 2.88E-289 | 5.2532092 | 0.513 | 0.233 | 5.76E-286 |
| IC | Msr1 | 8.55E-288 | 4.83724875 | 0.391 | 0.133 | 1.71E-284 |
| IC | Slco2b1 | 1.57E-282 | 2.29127102 | 0.668 | 0.388 | 3.15E-279 |
| IC | F13a1 | 2.27E-265 | 5.76551506 | 0.504 | 0.214 | 4.55E-262 |
| IC | Pltp | 3.29E-249 | 3.26126811 | 0.588 | 0.319 | 6.58E-246 |
| IC | Ptafr | 3.58E-227 | 3.97927343 | 0.519 | 0.245 | 7.16E-224 |
| IC | Rnf150 | 5.77E-224 | 1.97508434 | 0.657 | 0.377 | 1.15E-220 |
| IC | Tgfbi | 1.38E-218 | 2.92055592 | 0.585 | 0.325 | 2.75E-215 |
| IC | Cybb | 1.58E-202 | 4.50480961 | 0.499 | 0.245 | 3.15E-199 |
| IC | Pstpip2 | 9.90E-201 | 3.15362212 | 0.557 | 0.293 | 1.98E-197 |
| IC | Mctp2 | 2.67E-160 | 3.78473116 | 0.483 | 0.214 | 5.34E-157 |
| IC | Slc16a71 | 6.83E-158 | 2.72712673 | 0.515 | 0.26 | 1.37E-154 |
| IC | Lgals3 | 1.45E-92 | 2.53488223 | 0.488 | 0.23 | 2.91E-89 |
| T-cell | Itk | 0 | 7.96348679 | 0.586 | 0.024 | 0 |

| | | | | | | |
|---|---|---|---|---|---|---|
| T-cell | Ncr1 | 0 | 7.95406958 | 0.485 | 0.011 | 0 |
| T-cell | Cd3e | 0 | 8.33143267 | 0.481 | 0.026 | 0 |
| T-cell | Clec2d2 | 0 | 6.21387078 | 0.31 | 0.007 | 0 |
| T-cell | Gzma | 0 | 7.11942511 | 0.299 | 0.005 | 0 |
| T-cell | Kir3dl1 | 1.14E-293 | 7.57732381 | 0.388 | 0.023 | 2.28E-290 |
| T-cell | Cd247 | 4.25E-275 | 8.52389114 | 0.687 | 0.068 | 8.51E-272 |
| T-cell | Skap1 | 8.40E-249 | 8.31045295 | 0.974 | 0.206 | 1.68E-245 |
| T-cell | Il2rb | 7.81E-196 | 8.21269633 | 0.455 | 0.042 | 1.56E-192 |
| T-cell | Ikzf3 | 4.69E-165 | 6.77649601 | 0.627 | 0.107 | 9.37E-162 |
| T-cell | Tigit | 3.73E-153 | 4.71712156 | 0.313 | 0.026 | 7.46E-150 |
| T-cell | Cd96 | 2.69E-152 | 7.83425137 | 0.526 | 0.059 | 5.38E-149 |
| T-cell | Gpr174 | 5.14E-152 | 6.92523187 | 0.354 | 0.032 | 1.03E-148 |
| T-cell | Ptprc1 | 5.37E-144 | 4.14062805 | 0.951 | 0.327 | 1.07E-140 |
| T-cell | Scml4 | 2.24E-140 | 7.67556585 | 0.515 | 0.071 | 4.48E-137 |
| T-cell | Tbx21 | 3.62E-136 | 7.57622911 | 0.5 | 0.095 | 7.24E-133 |
| T-cell | AABR07017902.1 | 9.31E-129 | 6.94649247 | 0.817 | 0.255 | 1.86E-125 |
| T-cell | Ly49s6 | 2.98E-128 | 8.17011527 | 0.459 | 0.078 | 5.97E-125 |
| T-cell | Camk41 | 3.13E-127 | 6.8183454 | 0.683 | 0.167 | 6.27E-124 |
| T-cell | AABR07027872.1 | 2.16E-124 | 7.56253356 | 0.44 | 0.06 | 4.32E-121 |
| T-cell | Ikzf11 | 3.55E-120 | 4.28263343 | 0.854 | 0.29 | 7.11E-117 |
| T-cell | Ly49i4 | 1.42E-116 | 6.74558132 | 0.332 | 0.02 | 2.84E-113 |
| T-cell | Arhgap151 | 1.65E-113 | 3.26261646 | 0.918 | 0.458 | 3.30E-110 |
| T-cell | Olr292 | 1.06E-112 | 5.38497759 | 0.34 | 0.03 | 2.12E-109 |
| T-cell | Clnk | 4.10E-108 | 7.48740826 | 0.582 | 0.081 | 8.19E-105 |
| T-cell | Themis | 1.70E-97 | 7.13640848 | 0.541 | 0.132 | 3.40E-94 |
| T-cell | Bcl11b1 | 1.16E-95 | 6.77659379 | 0.522 | 0.156 | 2.33E-92 |
| T-cell | Il18r1 | 3.28E-95 | 5.71643071 | 0.466 | 0.066 | 6.55E-92 |

| | | | | | | |
|---|---|---|---|---|---|---|
| T-cell | Fyb11 | 3.66E-91 | 3.09716175 | 0.847 | 0.329 | 7.32E-88 |
| T-cell | Slamf6 | 4.05E-89 | 5.2061244 | 0.418 | 0.068 | 8.11E-86 |
| T-cell | Klri1 | 7.73E-82 | 7.0412007 | 0.507 | 0.111 | 1.55E-78 |
| T-cell | Cyfip2 | 1.54E-77 | 4.84061749 | 0.631 | 0.175 | 3.09E-74 |
| T-cell | Ly49s5 | 1.72E-68 | 6.33863656 | 0.392 | 0.042 | 3.43E-65 |
| T-cell | Hspa1b | 8.98E-67 | 2.20125377 | 0.313 | 0.054 | 1.80E-63 |
| T-cell | Prkcq | 6.96E-66 | 4.50143541 | 0.597 | 0.189 | 1.39E-62 |
| T-cell | Gfra3 | 1.33E-65 | 3.40647863 | 0.369 | 0.048 | 2.66E-62 |
| T-cell | Txk | 1.97E-62 | 6.35103704 | 0.422 | 0.107 | 3.94E-59 |
| T-cell | Rbm44 | 2.62E-62 | 3.45428256 | 0.328 | 0.054 | 5.24E-59 |
| T-cell | Ptpn221 | 6.08E-58 | 5.84032816 | 0.526 | 0.155 | 1.22E-54 |
| T-cell | Mcoln2 | 1.81E-57 | 5.04727474 | 0.325 | 0.064 | 3.63E-54 |
| T-cell | Itgal | 2.47E-55 | 3.97847487 | 0.638 | 0.277 | 4.94E-52 |
| T-cell | AABR07063 740.1 | 4.66E-54 | 4.02188636 | 0.369 | 0.055 | 9.33E-51 |
| T-cell | Rasgrp1 | 6.37E-52 | 4.81286221 | 0.455 | 0.085 | 1.27E-48 |
| T-cell | St8sia1 | 3.43E-49 | 5.83331315 | 0.455 | 0.139 | 6.86E-46 |
| T-cell | Pik3cd | 1.79E-47 | 3.41324755 | 0.668 | 0.32 | 3.58E-44 |
| T-cell | St6galnac33 | 1.50E-46 | 2.27627487 | 0.765 | 0.396 | 3.01E-43 |
| T-cell | Ovol1 | 3.72E-44 | 4.05845171 | 0.269 | 0.008 | 7.44E-41 |
| T-cell | Prkch3 | 1.27E-40 | 1.6304821 | 0.847 | 0.498 | 2.54E-37 |
| T-cell | Lcp21 | 1.36E-39 | 2.79640406 | 0.653 | 0.3 | 2.73E-36 |
| T-cell | Lamc3 | 3.53E-39 | 0.58742341 | 0.306 | 0.038 | 7.06E-36 |
| T-cell | Chn21 | 6.05E-39 | 2.40081019 | 0.743 | 0.4 | 1.21E-35 |
| T-cell | Scel | 4.23E-37 | 0.52921398 | 0.358 | 0.091 | 8.45E-34 |
| T-cell | Runx11 | 2.73E-36 | 1.96091181 | 0.761 | 0.465 | 5.46E-33 |
| T-cell | Prkcb1 | 1.07E-35 | 2.92789719 | 0.619 | 0.319 | 2.14E-32 |
| T-cell | Inpp4b2 | 4.10E-35 | 1.7650605 | 0.784 | 0.425 | 8.20E-32 |

| | | | | | | |
|---|---|---|---|---|---|---|
| T-cell | Cdh15 | 7.54E-35 | 1.9802754 | 0.306 | 0.049 | 1.51E-31 |
| T-cell | Dock81 | 4.28E-34 | 1.77116208 | 0.709 | 0.398 | 8.55E-31 |
| T-cell | Pik3r51 | 4.90E-34 | 3.32006291 | 0.493 | 0.235 | 9.79E-31 |
| T-cell | Plcg21 | 1.49E-31 | 2.75927657 | 0.582 | 0.249 | 2.97E-28 |
| T-cell | Stap1 | 1.25E-30 | 3.24345026 | 0.384 | 0.131 | 2.50E-27 |
| T-cell | Inpp5d1 | 8.60E-30 | 1.89680749 | 0.683 | 0.405 | 1.72E-26 |
| T-cell | Rnf43 | 2.82E-29 | 4.83629915 | 0.493 | 0.173 | 5.63E-26 |
| T-cell | Atp8b41 | 8.72E-28 | 3.58339724 | 0.56 | 0.276 | 1.74E-24 |
| T-cell | Runx2 | 1.59E-26 | 2.44082052 | 0.653 | 0.389 | 3.17E-23 |
| T-cell | Iqgap2 | 8.10E-25 | 2.4390274 | 0.612 | 0.35 | 1.62E-21 |
| T-cell | Itga4 | 2.73E-20 | 3.21006429 | 0.522 | 0.233 | 5.45E-17 |
| T-cell | Mctp21 | 5.81E-16 | 2.36901707 | 0.519 | 0.237 | 1.16E-12 |
| T-cell | Plxnc11 | 1.65E-14 | 2.37873818 | 0.493 | 0.232 | 3.29E-11 |
| T-cell | Stard101 | 1.24E-13 | 3.56813293 | 0.392 | 0.12 | 2.48E-10 |
| SMC | Notch3 | 0 | 5.03100616 | 0.834 | 0.257 | 0 |
| SMC | Rgs5 | 0 | 5.57428454 | 0.831 | 0.279 | 0 |
| SMC | Mrvi1 | 0 | 4.25348577 | 0.867 | 0.316 | 0 |
| SMC | Gucy1a2 | 0 | 3.87255001 | 0.921 | 0.387 | 0 |
| SMC | Ano1 | 0 | 5.53239956 | 0.731 | 0.206 | 0 |
| SMC | Adgrl3 | 0 | 4.62955104 | 0.758 | 0.246 | 0 |
| SMC | Egflam | 0 | 4.16219186 | 0.775 | 0.271 | 0 |
| SMC | Gucy1a1 | 0 | 3.28478238 | 0.872 | 0.385 | 0 |
| SMC | Pdgfrb1 | 0 | 2.48277557 | 0.914 | 0.446 | 0 |
| SMC | Il34 | 0 | 5.17037341 | 0.673 | 0.234 | 0 |
| SMC | Lrrc4c | 0 | 4.71929836 | 0.693 | 0.275 | 0 |
| SMC | Rasl12 | 0 | 5.30536469 | 0.635 | 0.219 | 0 |
| SMC | Pde3a2 | 0 | 2.74440705 | 0.968 | 0.552 | 0 |
| SMC | Ldb21 | 0 | 1.84792352 | 0.72 | 0.321 | 0 |
| SMC | Cspg4 | 0 | 3.77472435 | 0.664 | 0.271 | 0 |

| | | | | | | |
|---|---|---|---|---|---|---|
| SMC | Trpc3 | 0 | 5.04264997 | 0.612 | 0.224 | 0 |
| SMC | Abcc91 | 0 | 3.19447128 | 0.887 | 0.516 | 0 |
| SMC | AC134204.1<br>1 | 0 | 2.40909753 | 0.825 | 0.455 | 0 |
| SMC | Pde8b | 0 | 4.72972787 | 0.577 | 0.211 | 0 |
| SMC | Mylk | 0 | 2.30115272 | 0.83 | 0.478 | 0 |
| SMC | Mcam | 0 | 2.11396319 | 0.684 | 0.333 | 0 |
| SMC | P2ry14 | 0 | 3.90399727 | 0.55 | 0.205 | 0 |
| SMC | Rerg | 0 | 3.49297051 | 0.72 | 0.381 | 0 |
| SMC | Ebf21 | 0 | 1.01548173 | 0.826 | 0.487 | 0 |
| SMC | Trpc4 | 0 | 5.35864174 | 0.376 | 0.045 | 0 |
| SMC | Sgip1 | 0 | 2.99610147 | 0.625 | 0.298 | 0 |
| SMC | Zfp385d1 | 0 | 1.22291001 | 0.876 | 0.55 | 0 |
| SMC | Mark1 | 0 | 4.09145523 | 0.588 | 0.262 | 0 |
| SMC | Cobll1 | 0 | 2.47612986 | 0.73 | 0.408 | 0 |
| SMC | Mob3b | 0 | 3.75434808 | 0.601 | 0.281 | 0 |
| SMC | Palm2 | 0 | 2.59125662 | 0.701 | 0.385 | 0 |
| SMC | Olr59 | 0 | 5.49625609 | 0.415 | 0.101 | 0 |
| SMC | Kcnip1 | 0 | 3.26140387 | 0.414 | 0.102 | 0 |
| SMC | Dmp1 | 0 | 5.72906278 | 0.34 | 0.04 | 0 |
| SMC | Trpc61 | 0 | 3.81875385 | 0.483 | 0.187 | 0 |
| SMC | Ednra | 0 | 2.42940441 | 0.764 | 0.469 | 0 |
| SMC | Inpp4b3 | 0 | 2.33995139 | 0.696 | 0.402 | 0 |
| SMC | Slco3a11 | 0 | 1.93909499 | 0.834 | 0.549 | 0 |
| SMC | Pmfbp1 | 0 | 5.45831608 | 0.331 | 0.048 | 0 |
| SMC | Stac | 0 | 5.14069822 | 0.372 | 0.095 | 0 |
| SMC | Agap2 | 0 | 5.15558517 | 0.415 | 0.14 | 0 |
| SMC | Grm3 | 0 | 3.87461682 | 0.28 | 0.013 | 0 |
| SMC | Cald1 | 0 | 1.91215478 | 0.879 | 0.612 | 0 |

| | | | | | | |
|---|---|---|---|---|---|---|
| SMC | Col25a1 | 0 | 4.25490933 | 0.378 | 0.119 | 0 |
| SMC | Fhl5 | 0 | 3.56861149 | 0.334 | 0.077 | 0 |
| SMC | Zeb21 | 0 | 1.41337766 | 0.945 | 0.69 | 0 |
| SMC | Cdh6 | 0 | 5.06670266 | 0.337 | 0.085 | 0 |
| SMC | Cyp4f18 | 7.36E-301 | 4.43037126 | 0.379 | 0.108 | 1.47E-297 |
| SMC | Ntng1 | 8.37E-298 | 4.12754458 | 0.518 | 0.212 | 1.67E-294 |
| SMC | Kcnj8 | 5.64E-282 | 4.85181336 | 0.554 | 0.21 | 1.13E-278 |
| SMC | Prrx11 | 8.40E-280 | 1.66507418 | 0.738 | 0.418 | 1.68E-276 |
| SMC | AABR07006724.11 | 8.76E-272 | 3.60033452 | 0.583 | 0.264 | 1.75E-268 |
| SMC | Myh11 | 6.81E-271 | 5.65575573 | 0.547 | 0.257 | 1.36E-267 |
| SMC | Akap61 | 2.70E-263 | 1.04479526 | 0.789 | 0.493 | 5.40E-260 |
| SMC | Xkr4 | 1.57E-219 | 3.75200969 | 0.533 | 0.271 | 3.15E-216 |
| SMC | Lmod1 | 3.75E-211 | 3.9088964 | 0.568 | 0.297 | 7.51E-208 |
| SMC | P3h22 | 1.93E-207 | 1.74466488 | 0.632 | 0.371 | 3.87E-204 |
| SMC | Med12l | 8.00E-201 | 3.12627378 | 0.541 | 0.265 | 1.60E-197 |
| SMC | Agmo | 1.59E-184 | 3.42814597 | 0.531 | 0.266 | 3.17E-181 |
| SMC | Pip5k1b1 | 9.81E-182 | 2.7133413 | 0.564 | 0.303 | 1.96E-178 |
| SMC | Chn11 | 1.56E-174 | 1.62554789 | 0.628 | 0.373 | 3.12E-171 |
| SMC | Slc16a72 | 4.34E-160 | 1.9309975 | 0.54 | 0.259 | 8.68E-157 |
| Epicardium | AABR07069371.1 | 0 | 8.67879501 | 0.778 | 0.035 | 0 |
| Epicardium | Nkain41 | 0 | 7.13689319 | 0.774 | 0.053 | 0 |
| Epicardium | Ly6k | 0 | 8.6092892 | 0.694 | 0.003 | 0 |
| Epicardium | Scel1 | 0 | 7.23570283 | 0.759 | 0.084 | 0 |
| Epicardium | Cgn | 0 | 8.15796426 | 0.703 | 0.049 | 0 |
| Epicardium | Bicdl1 | 0 | 6.72712714 | 0.709 | 0.087 | 0 |
| Epicardium | Upk1b | 0 | 7.19923634 | 0.677 | 0.059 | 0 |

| | | | | | | |
|---|---|---|---|---|---|---|
| Epicardium | AABR07065531.26 | 0 | 4.77469649 | 0.694 | 0.084 | 0 |
| Epicardium | AABR07063274.1 | 0 | 8.12806043 | 0.651 | 0.046 | 0 |
| Epicardium | Chst4 | 0 | 9.39578946 | 0.603 | 0 | 0 |
| Epicardium | Tmem233 | 0 | 8.28516457 | 0.597 | 0.002 | 0 |
| Epicardium | Il22ra1 | 0 | 7.16255563 | 0.64 | 0.055 | 0 |
| Epicardium | Bnc1 | 0 | 8.12821426 | 0.601 | 0.023 | 0 |
| Epicardium | Krt7 | 0 | 7.41161563 | 0.565 | 0.003 | 0 |
| Epicardium | Entpd3 | 0 | 5.93087805 | 0.56 | 0.008 | 0 |
| Epicardium | Pnoc | 0 | 5.30686394 | 0.619 | 0.069 | 0 |
| Epicardium | Vtcn1 | 0 | 6.00615158 | 0.595 | 0.045 | 0 |
| Epicardium | Wnt7b | 0 | 5.89998419 | 0.558 | 0.009 | 0 |
| Epicardium | Isl1 | 0 | 6.42454144 | 0.554 | 0.006 | 0 |
| Epicardium | Prkg2 | 0 | 3.74604317 | 0.56 | 0.013 | 0 |
| Epicardium | Fgf8 | 0 | 4.74964245 | 0.552 | 0.009 | 0 |
| Epicardium | Cela3b | 0 | 4.10118354 | 0.606 | 0.065 | 0 |
| Epicardium | Krt18 | 0 | 6.34052695 | 0.586 | 0.05 | 0 |
| Epicardium | Ribc2 | 0 | 3.23231107 | 0.545 | 0.013 | 0 |
| Epicardium | Lypd6b | 0 | 4.70549895 | 0.554 | 0.023 | 0 |
| Epicardium | Egfl6 | 0 | 5.95233327 | 0.582 | 0.054 | 0 |
| Epicardium | Atp6v0a4 | 0 | 4.07171229 | 0.55 | 0.026 | 0 |
| Epicardium | AABR07003304.2 | 0 | 3.81326024 | 0.558 | 0.035 | 0 |
| Epicardium | AABR07029164.1 | 0 | 1.72951041 | 0.53 | 0.007 | 0 |
| Epicardium | Stmn2 | 0 | 6.81753243 | 0.539 | 0.017 | 0 |
| Epicardium | Gpx2 | 0 | 4.47661161 | 0.513 | 0.007 | 0 |

| | | | | | | |
|---|---|---|---|---|---|---|
| Epicardium | AABR07056415.1 | 0 | 3.27085076 | 0.511 | 0.005 | 0 |
| Epicardium | AABR07005949.1 | 0 | 3.07967265 | 0.539 | 0.044 | 0 |
| Epicardium | Anxa8 | 0 | 3.70549546 | 0.511 | 0.026 | 0 |
| Epicardium | AABR07035486.1 | 0 | 4.11413357 | 0.494 | 0.022 | 0 |
| Epicardium | LOC100909857 | 0 | 2.22027942 | 0.487 | 0.038 | 0 |
| Epicardium | Col6a5 | 0 | 4.62489509 | 0.474 | 0.048 | 0 |
| Epicardium | Upk3b | 0 | 7.81848375 | 0.416 | 0.018 | 0 |
| Epicardium | RGD1564614 | 0 | 1.87654728 | 0.425 | 0.027 | 0 |
| Epicardium | Elavl21 | 0 | 5.6508746 | 0.429 | 0.032 | 0 |
| Epicardium | AABR07004783.1 | 0 | 3.58554861 | 0.36 | 0.017 | 0 |
| Epicardium | AABR07010985.1 | 0 | 7.1332994 | 0.39 | 0.054 | 0 |
| Epicardium | Mmp241 | 0 | 3.03754375 | 0.375 | 0.06 | 0 |
| Epicardium | Krt17 | 0 | 7.41570666 | 0.375 | 0.066 | 0 |
| Epicardium | AABR07058498.1 | 0 | 4.9082502 | 0.256 | 0.002 | 0 |
| Epicardium | Uchl1 | 7.79E-305 | 3.86185442 | 0.629 | 0.064 | 1.56E-301 |
| Epicardium | Hrh2 | 2.82E-304 | 0.49340815 | 0.481 | 0.072 | 5.63E-301 |
| Epicardium | Cdh3 | 2.86E-304 | 6.46420133 | 0.789 | 0.093 | 5.71E-301 |
| Epicardium | Stk26 | 1.45E-297 | 5.62996106 | 0.694 | 0.112 | 2.90E-294 |
| Epicardium | Il18r11 | 1.64E-269 | 1.64204755 | 0.562 | 0.062 | 3.29E-266 |
| Epicardium | Ptprz1 | 3.40E-269 | 2.07137043 | 0.696 | 0.061 | 6.81E-266 |
| Epicardium | Erc21 | 1.57E-265 | 4.50630726 | 0.625 | 0.082 | 3.14E-262 |

| | | | | | | |
|---|---|---|---|---|---|---|
| Epicardium | Slc26a3 | 6.77E-264 | 6.73957449 | 0.425 | 0.077 | 1.35E-260 |
| Epicardium | Dhrs9 | 2.00E-261 | 6.39624991 | 0.42 | 0.08 | 4.00E-258 |
| Epicardium | Tnfrsf9 | 1.24E-258 | 5.36069735 | 0.593 | 0.062 | 2.48E-255 |
| Epicardium | Myo5b | 3.22E-251 | 5.99239599 | 0.534 | 0.13 | 6.45E-248 |
| Epicardium | Rims1 | 3.06E-244 | 3.26069673 | 0.468 | 0.083 | 6.11E-241 |
| Epicardium | Msln | 4.99E-244 | 6.78334091 | 0.616 | 0.047 | 9.98E-241 |
| Epicardium | Efna5 | 4.80E-242 | 4.63055073 | 0.875 | 0.27 | 9.60E-239 |
| Epicardium | Chst91 | 1.10E-237 | 1.11272999 | 0.636 | 0.091 | 2.20E-234 |
| Epicardium | Baiap2l11 | 4.45E-230 | 4.67192517 | 0.897 | 0.289 | 8.91E-227 |
| Epicardium | Unc5d | 2.61E-229 | 4.28969414 | 0.571 | 0.058 | 5.22E-226 |
| Epicardium | Sema3c | 1.08E-227 | 4.28696063 | 0.927 | 0.358 | 2.16E-224 |
| Epicardium | Il13ra2 | 6.77E-224 | 4.35214031 | 0.655 | 0.097 | 1.35E-220 |
| Epicardium | Htr4 | 5.87E-223 | 6.6613404 | 0.606 | 0.142 | 1.17E-219 |
| Epicardium | Fmod | 9.63E-223 | 3.77444066 | 0.7 | 0.11 | 1.93E-219 |
| Epicardium | Mybphl | 5.35E-221 | 1.95193099 | 0.526 | 0.031 | 1.07E-217 |
| Epicardium | Pkhd1l12 | 1.52E-216 | 3.9735694 | 0.86 | 0.266 | 3.04E-213 |
| Epicardium | Rspo1 | 2.66E-216 | 4.58900181 | 0.836 | 0.212 | 5.32E-213 |
| Epicardium | Cdkn2b | 3.30E-208 | 2.90469532 | 0.662 | 0.134 | 6.61E-205 |
| Epicardium | Prss121 | 1.04E-204 | 4.12083764 | 0.7 | 0.131 | 2.07E-201 |
| Epicardium | Aldh1a2 | 1.92E-203 | 4.40802725 | 0.834 | 0.286 | 3.83E-200 |
| Epicardium | Cmahp | 1.05E-201 | 5.29967254 | 0.558 | 0.118 | 2.11E-198 |
| Epicardium | Pla2g2a | 3.93E-200 | 4.21696646 | 0.664 | 0.112 | 7.86E-197 |
| Epicardium | Hpgd | 5.77E-199 | 3.49251862 | 0.595 | 0.139 | 1.15E-195 |
| Epicardium | Sema3e | 2.21E-198 | 3.86636396 | 0.75 | 0.17 | 4.42E-195 |
| Epicardium | P4ha3 | 3.36E-198 | 4.45306418 | 0.711 | 0.145 | 6.72E-195 |
| Epicardium | Cpvl1 | 2.27E-194 | 1.98507324 | 0.528 | 0.058 | 4.54E-191 |
| Epicardium | Apba21 | 3.66E-194 | 1.30594252 | 0.513 | 0.099 | 7.31E-191 |
| Epicardium | Acoxl | 4.71E-193 | 2.26884462 | 0.619 | 0.103 | 9.41E-190 |
| Epicardium | Gpm6a1 | 3.51E-192 | 3.92891732 | 0.927 | 0.517 | 7.03E-189 |

| | | | | | | |
|---|---|---|---|---|---|---|
| Epicardium | Cobl | 4.10E-192 | 4.34803979 | 0.804 | 0.274 | 8.20E-189 |
| Epicardium | AABR07031193.1 | 5.35E-191 | 3.62794832 | 0.711 | 0.166 | 1.07E-187 |
| Epicardium | Tmem63c1 | 2.02E-189 | 3.03018156 | 0.759 | 0.202 | 4.04E-186 |
| Epicardium | Rhpn2 | 6.41E-189 | 4.76891337 | 0.769 | 0.192 | 1.28E-185 |
| Epicardium | Cdh112 | 1.12E-184 | 2.70502267 | 0.946 | 0.44 | 2.25E-181 |
| Epicardium | Neurl1 | 3.58E-183 | 3.58962333 | 0.655 | 0.16 | 7.16E-180 |
| Epicardium | Cdon | 1.38E-182 | 3.433402 | 0.871 | 0.388 | 2.76E-179 |
| Epicardium | RGD1564053 | 2.28E-182 | 2.97727569 | 0.582 | 0.113 | 4.55E-179 |
| Epicardium | Sema5a2 | 1.34E-181 | 3.33567977 | 0.869 | 0.297 | 2.68E-178 |
| Epicardium | Bnc21 | 1.26E-180 | 2.34523345 | 0.948 | 0.419 | 2.51E-177 |
| Epicardium | Rab11fip1 | 2.22E-180 | 3.55760359 | 0.53 | 0.105 | 4.44E-177 |
| Epicardium | Mybpc2 | 2.08E-178 | 2.19819974 | 0.601 | 0.136 | 4.16E-175 |
| Epicardium | Adamts20 | 4.56E-177 | 1.81109788 | 0.644 | 0.122 | 9.12E-174 |
| Epicardium | AABR07029470.1 | 4.99E-176 | 4.46486157 | 0.483 | 0.131 | 9.99E-173 |
| Epicardium | Flt3 | 1.40E-174 | 4.55250638 | 0.709 | 0.148 | 2.81E-171 |
| Epicardium | Gpc3 | 1.17E-172 | 5.46713074 | 0.616 | 0.163 | 2.34E-169 |
| Epicardium | Galnt14 | 2.74E-172 | 1.18956536 | 0.586 | 0.087 | 5.48E-169 |
| Epicardium | Pak3 | 1.80E-171 | 2.19786776 | 0.685 | 0.174 | 3.59E-168 |
| Epicardium | Tll2 | 2.79E-171 | 3.33063024 | 0.666 | 0.133 | 5.59E-168 |
| Epicardium | Sh3gl32 | 2.99E-171 | 1.15135231 | 0.722 | 0.186 | 5.99E-168 |
| Epicardium | Enpp2 | 3.96E-171 | 4.61132275 | 0.778 | 0.234 | 7.93E-168 |
| Epicardium | Nrg4 | 4.35E-171 | 4.3360569 | 0.388 | 0.093 | 8.70E-168 |
| Epicardium | Ptpn131 | 5.42E-171 | 2.90474805 | 0.869 | 0.411 | 1.08E-167 |
| Epicardium | Lancl31 | 2.30E-170 | 0.76303273 | 0.659 | 0.143 | 4.60E-167 |
| Epicardium | Cdh61 | 1.27E-168 | 0.29397724 | 0.545 | 0.102 | 2.54E-165 |
| Epicardium | Dok62 | 1.16E-167 | 4.45026053 | 0.56 | 0.138 | 2.31E-164 |

| | | | | | | |
|---|---|---|---|---|---|---|
| Epicardium | Katnal2 | 7.45E-167 | 1.58974287 | 0.528 | 0.09 | 1.49E-163 |
| Epicardium | Ezr | 7.43E-164 | 4.24833066 | 0.754 | 0.301 | 1.49E-160 |
| Epicardium | Pdpn1 | 8.40E-164 | 4.12971757 | 0.709 | 0.194 | 1.68E-160 |
| Epicardium | Ano3 | 7.41E-163 | 2.59826311 | 0.64 | 0.133 | 1.48E-159 |
| Epicardium | Cacna1d | 1.24E-162 | 3.25355575 | 0.677 | 0.167 | 2.48E-159 |
| Epicardium | Fam180a | 8.68E-161 | 3.03124411 | 0.698 | 0.175 | 1.74E-157 |
| Epicardium | Kcnq5 | 1.68E-160 | 4.09918204 | 0.731 | 0.191 | 3.37E-157 |
| Epicardium | Il161 | 3.73E-160 | 3.08636589 | 0.83 | 0.328 | 7.46E-157 |
| Epicardium | Nnat | 1.08E-159 | 0.64956503 | 0.506 | 0.108 | 2.16E-156 |
| Epicardium | Prlr | 3.19E-158 | 1.16438005 | 0.588 | 0.147 | 6.38E-155 |
| Epicardium | Fras1 | 1.26E-157 | 5.38603275 | 0.644 | 0.197 | 2.52E-154 |
| Epicardium | Grem2 | 1.39E-157 | 1.18170915 | 0.616 | 0.139 | 2.79E-154 |
| Epicardium | Bcar3 | 8.14E-157 | 2.96976035 | 0.83 | 0.382 | 1.63E-153 |
| Epicardium | Rgs7bp | 1.75E-156 | 3.59934175 | 0.634 | 0.183 | 3.50E-153 |
| Epicardium | Sbspon | 2.19E-156 | 0.34285341 | 0.489 | 0.067 | 4.39E-153 |
| Epicardium | Cpxm2 | 4.48E-155 | 1.80529468 | 0.724 | 0.18 | 8.96E-152 |
| Epicardium | Efemp1 | 1.07E-154 | 3.72918859 | 0.772 | 0.272 | 2.14E-151 |
| Epicardium | Ptger3 | 4.54E-153 | 1.43755415 | 0.653 | 0.167 | 9.09E-150 |
| Epicardium | Car10 | 1.10E-151 | 4.5483918 | 0.591 | 0.156 | 2.19E-148 |
| Epicardium | Sulf1 | 2.30E-151 | 2.21064771 | 0.922 | 0.477 | 4.59E-148 |
| Epicardium | Flrt21 | 4.29E-151 | 2.77251018 | 0.881 | 0.43 | 8.57E-148 |
| Epicardium | Plxna41 | 5.95E-151 | 2.42304016 | 0.925 | 0.525 | 1.19E-147 |
| Epicardium | Kcnk5 | 1.61E-150 | 1.8957382 | 0.711 | 0.178 | 3.22E-147 |
| Epicardium | Pak7 | 3.03E-150 | 2.821518 | 0.483 | 0.127 | 6.07E-147 |
| Epicardium | Adgrd11 | 4.65E-150 | 2.74640427 | 0.881 | 0.379 | 9.31E-147 |
| Epicardium | Zbtb7c | 8.38E-149 | 2.76936084 | 0.825 | 0.339 | 1.68E-145 |
| Epicardium | Slc39a8 | 1.20E-148 | 4.50367058 | 0.724 | 0.227 | 2.40E-145 |
| Epicardium | Ptk2b | 2.36E-148 | 2.51478777 | 0.879 | 0.392 | 4.73E-145 |
| Epicardium | Dsg21 | 1.63E-147 | 3.67897247 | 0.72 | 0.293 | 3.27E-144 |

| | | | | | | |
|---|---|---|---|---|---|---|
| Epicardium | Prkcz1 | 5.21E-147 | 3.16623956 | 0.754 | 0.286 | 1.04E-143 |
| Epicardium | Il1rn | 5.48E-147 | 1.53415678 | 0.496 | 0.073 | 1.10E-143 |
| Epicardium | Fgfr2 | 8.09E-145 | 3.52474536 | 0.739 | 0.27 | 1.62E-141 |
| Epicardium | Hs3st3a1 | 1.25E-142 | 3.01729127 | 0.685 | 0.185 | 2.50E-139 |
| Epicardium | Fstl4 | 1.48E-142 | 0.79585531 | 0.547 | 0.098 | 2.95E-139 |
| Epicardium | Fgf9 | 5.85E-142 | 3.69313534 | 0.707 | 0.21 | 1.17E-138 |
| Epicardium | Tuft1 | 2.16E-139 | 4.09904902 | 0.765 | 0.263 | 4.32E-136 |
| Epicardium | AABR07059258.11 | 3.79E-139 | 2.69914117 | 0.879 | 0.36 | 7.57E-136 |
| Epicardium | Erbb3 | 2.11E-138 | 0.53080534 | 0.575 | 0.097 | 4.23E-135 |
| Epicardium | Adamts16 | 2.93E-138 | 3.98496313 | 0.694 | 0.156 | 5.87E-135 |
| Epicardium | Kank11 | 1.04E-136 | 2.46353075 | 0.875 | 0.446 | 2.07E-133 |
| Epicardium | Caln1 | 1.34E-136 | 1.63401235 | 0.603 | 0.138 | 2.67E-133 |
| Epicardium | Pamr1 | 1.43E-136 | 2.05818828 | 0.726 | 0.217 | 2.85E-133 |
| Epicardium | AABR07054614.11 | 1.41E-134 | 2.23600902 | 0.877 | 0.372 | 2.81E-131 |
| Epicardium | Pappa2 | 2.86E-134 | 1.93982159 | 0.647 | 0.175 | 5.72E-131 |
| Epicardium | Abcd2 | 4.74E-134 | 3.76906438 | 0.69 | 0.208 | 9.48E-131 |
| Epicardium | Dpp4 | 5.21E-134 | 2.89835327 | 0.767 | 0.286 | 1.04E-130 |
| Epicardium | Tm4sf19 | 9.44E-134 | 0.52720929 | 0.56 | 0.124 | 1.89E-130 |
| Epicardium | Prph | 3.79E-133 | 3.23695923 | 0.733 | 0.196 | 7.59E-130 |
| Epicardium | Wt12 | 7.41E-133 | 2.96528149 | 0.752 | 0.31 | 1.48E-129 |
| Epicardium | Cacna2d31 | 2.23E-131 | 2.46265659 | 0.89 | 0.368 | 4.46E-128 |
| Epicardium | Has11 | 4.68E-131 | 1.80446897 | 0.726 | 0.213 | 9.35E-128 |
| Epicardium | Gria1 | 7.83E-131 | 0.33965114 | 0.636 | 0.137 | 1.57E-127 |
| Epicardium | St8sia11 | 9.99E-131 | 0.4087951 | 0.612 | 0.135 | 2.00E-127 |
| Epicardium | Rbfox11 | 2.20E-130 | 2.41776861 | 0.886 | 0.403 | 4.40E-127 |
| Epicardium | Plod21 | 7.37E-130 | 2.01257477 | 0.959 | 0.5 | 1.47E-126 |
| Epicardium | Slc4a41 | 1.94E-128 | 2.43677303 | 0.838 | 0.416 | 3.88E-125 |

| | | | | | | |
|---|---|---|---|---|---|---|
| Epicardium | Ppl | 2.98E-128 | 3.66746481 | 0.653 | 0.24 | 5.97E-125 |
| Epicardium | LOC103689965 | 1.35E-127 | 2.92777475 | 0.769 | 0.398 | 2.70E-124 |
| Epicardium | Shroom2 | 5.38E-127 | 2.92673996 | 0.731 | 0.241 | 1.08E-123 |
| Epicardium | Anxa31 | 1.41E-124 | 2.22573416 | 0.832 | 0.362 | 2.81E-121 |
| Epicardium | Tnfrsf11b | 5.69E-124 | 2.27181716 | 0.767 | 0.294 | 1.14E-120 |
| Epicardium | Arnt2 | 4.01E-123 | 2.1475083 | 0.694 | 0.231 | 8.02E-120 |
| Epicardium | Gcnt21 | 3.85E-122 | 2.74387946 | 0.724 | 0.346 | 7.70E-119 |
| Epicardium | Ltk | 2.16E-121 | 0.78508717 | 0.459 | 0.135 | 4.32E-118 |
| Epicardium | Ildr2 | 1.14E-120 | 3.88125784 | 0.603 | 0.254 | 2.28E-117 |
| Epicardium | Igsf11 | 1.28E-120 | 2.72306072 | 0.517 | 0.144 | 2.57E-117 |
| Epicardium | Ccdc801 | 2.18E-120 | 1.52955711 | 0.957 | 0.593 | 4.36E-117 |
| Epicardium | Sctr | 7.71E-120 | 0.3413705 | 0.593 | 0.146 | 1.54E-116 |
| Epicardium | Adgrb3 | 8.39E-120 | 4.5095931 | 0.569 | 0.145 | 1.68E-116 |
| Epicardium | AABR07065190.1 | 6.69E-118 | 0.55340321 | 0.599 | 0.124 | 1.34E-114 |
| Epicardium | AABR07041096.11 | 1.56E-115 | 2.78812975 | 0.666 | 0.207 | 3.11E-112 |
| Epicardium | Ephb2 | 1.60E-115 | 2.09574805 | 0.679 | 0.224 | 3.20E-112 |
| Epicardium | Grid21 | 2.45E-115 | 1.20597034 | 0.489 | 0.156 | 4.89E-112 |
| Epicardium | Vgll3 | 4.89E-114 | 2.0019184 | 0.735 | 0.265 | 9.78E-111 |
| Epicardium | Sox61 | 2.41E-112 | 1.98029159 | 0.841 | 0.478 | 4.81E-109 |
| Epicardium | Syndig1 | 4.34E-112 | 2.11529397 | 0.793 | 0.371 | 8.68E-109 |
| Epicardium | Anln1 | 5.10E-112 | 1.87665118 | 0.623 | 0.193 | 1.02E-108 |
| Epicardium | Cldn15 | 6.07E-112 | 5.1360591 | 0.513 | 0.212 | 1.21E-108 |
| Epicardium | Egfr1 | 7.18E-112 | 2.12495089 | 0.838 | 0.45 | 1.44E-108 |
| Epicardium | Hs3st21 | 1.53E-111 | 0.64260441 | 0.599 | 0.16 | 3.06E-108 |
| Epicardium | Asic2 | 2.88E-111 | 3.07178773 | 0.651 | 0.218 | 5.77E-108 |
| Epicardium | Tex22 | 9.59E-111 | 2.46190768 | 0.614 | 0.213 | 1.92E-107 |

| | | | | | | |
|---|---|---|---|---|---|---|
| Epicardium | Gli21 | 2.31E-110 | 1.98781681 | 0.838 | 0.39 | 4.61E-107 |
| Epicardium | Grm11 | 1.35E-109 | 1.44366248 | 0.429 | 0.116 | 2.70E-106 |
| Epicardium | Tll1 | 5.07E-109 | 3.33649002 | 0.675 | 0.241 | 1.01E-105 |
| Epicardium | Tmem200a | 2.32E-108 | 1.22217126 | 0.498 | 0.172 | 4.63E-105 |
| Epicardium | Penk | 1.02E-107 | 0.5160867 | 0.634 | 0.196 | 2.03E-104 |
| Epicardium | Tbx18 | 2.54E-107 | 3.08681595 | 0.692 | 0.328 | 5.08E-104 |
| Epicardium | Galnt13 | 2.61E-107 | 6.09498548 | 0.47 | 0.167 | 5.23E-104 |
| Epicardium | Arhgef26 | 2.26E-104 | 2.05970548 | 0.726 | 0.26 | 4.52E-101 |
| Epicardium | Ptgs11 | 1.04E-103 | 2.12851778 | 0.705 | 0.27 | 2.09E-100 |
| Epicardium | AABR07027581.11 | 1.58E-103 | 2.14299642 | 0.782 | 0.385 | 3.16E-100 |
| Epicardium | Crip1 | 2.78E-102 | 2.44193842 | 0.802 | 0.365 | 5.57E-99 |
| Epicardium | Fgf121 | 8.52E-102 | 2.57726406 | 0.735 | 0.365 | 1.70E-98 |
| Epicardium | Adgrg2 | 1.28E-101 | 3.21385597 | 0.597 | 0.22 | 2.57E-98 |
| Epicardium | Rnf431 | 2.98E-101 | 2.2449688 | 0.53 | 0.17 | 5.96E-98 |
| Epicardium | Itgb41 | 4.32E-101 | 1.55316515 | 0.731 | 0.258 | 8.65E-98 |
| Epicardium | Arhgap441 | 4.92E-100 | 2.36764749 | 0.776 | 0.367 | 9.84E-97 |
| Epicardium | Pawr1 | 1.76E-99 | 1.72419385 | 0.774 | 0.308 | 3.53E-96 |
| Epicardium | Cttnbp2 | 1.28E-98 | 1.641921 | 0.8 | 0.349 | 2.56E-95 |
| Epicardium | Shc4 | 2.39E-97 | 1.24573047 | 0.47 | 0.116 | 4.79E-94 |
| Epicardium | AABR07034940.2 | 1.04E-96 | 0.90917848 | 0.666 | 0.184 | 2.08E-93 |
| Epicardium | Esr1 | 1.04E-96 | 2.95993643 | 0.737 | 0.294 | 2.08E-93 |
| Epicardium | Nova1 | 1.11E-96 | 1.79349178 | 0.722 | 0.307 | 2.23E-93 |
| Epicardium | Osr1 | 2.04E-96 | 1.37943286 | 0.675 | 0.276 | 4.08E-93 |
| Epicardium | Crim11 | 2.39E-96 | 1.97426458 | 0.862 | 0.584 | 4.77E-93 |
| Epicardium | Slc1a3 | 3.21E-96 | 2.77676813 | 0.631 | 0.256 | 6.41E-93 |
| Epicardium | Hs6st2 | 1.16E-95 | 2.04463907 | 0.724 | 0.306 | 2.32E-92 |
| Epicardium | Tmtc21 | 3.00E-95 | 1.64531012 | 0.847 | 0.407 | 5.99E-92 |

| | | | | | | |
|---|---|---|---|---|---|---|
| Epicardium | Lgals1 | 3.87E-95 | 2.68513465 | 0.769 | 0.354 | 7.75E-92 |
| Epicardium | Icam11 | 1.59E-94 | 0.9601006 | 0.705 | 0.251 | 3.17E-91 |
| Epicardium | Epha7 | 5.76E-94 | 2.61662173 | 0.631 | 0.229 | 1.15E-90 |
| Epicardium | Prelp | 2.93E-93 | 1.58418265 | 0.685 | 0.197 | 5.85E-90 |
| Epicardium | Cpq1 | 5.19E-93 | 1.44288284 | 0.933 | 0.509 | 1.04E-89 |
| Epicardium | Pou2f2 | 5.89E-93 | 0.9775585 | 0.625 | 0.2 | 1.18E-89 |
| Epicardium | Kif221 | 2.45E-92 | 0.70896269 | 0.586 | 0.185 | 4.89E-89 |
| Epicardium | AABR07049 085.11 | 8.83E-91 | 1.55142269 | 0.765 | 0.339 | 1.77E-87 |
| Epicardium | Mctp22 | 1.32E-89 | 3.11332073 | 0.578 | 0.235 | 2.63E-86 |
| Epicardium | Tspan51 | 3.46E-88 | 1.78498824 | 0.804 | 0.462 | 6.92E-85 |
| Epicardium | Slc1a7 | 4.99E-88 | 0.25980666 | 0.754 | 0.332 | 9.97E-85 |
| Epicardium | Piezo2 | 9.58E-87 | 2.20577497 | 0.802 | 0.348 | 1.92E-83 |
| Epicardium | Cit1 | 3.32E-84 | 1.31588469 | 0.653 | 0.324 | 6.65E-81 |
| Epicardium | Kif20a | 3.59E-84 | 0.48191389 | 0.584 | 0.185 | 7.18E-81 |
| Epicardium | Tnfrsf12a | 4.35E-84 | 1.77980927 | 0.575 | 0.2 | 8.70E-81 |
| Epicardium | Csrp2 | 8.60E-84 | 2.03502762 | 0.744 | 0.33 | 1.72E-80 |
| Epicardium | Slc16a11 | 1.46E-83 | 1.52555991 | 0.774 | 0.329 | 2.93E-80 |
| Epicardium | Ap1s3 | 1.01E-82 | 1.70554597 | 0.683 | 0.333 | 2.02E-79 |
| Epicardium | Fam189a22 | 1.22E-82 | 1.16916406 | 0.616 | 0.228 | 2.43E-79 |
| Epicardium | Agtr1a | 1.27E-80 | 1.58539625 | 0.737 | 0.341 | 2.53E-77 |
| Epicardium | Diaph31 | 6.31E-80 | 0.67308226 | 0.698 | 0.269 | 1.26E-76 |
| Epicardium | Ccdc148 | 4.53E-79 | 0.94094734 | 0.522 | 0.162 | 9.06E-76 |
| Epicardium | Magi2 | 1.95E-78 | 2.39748279 | 0.612 | 0.298 | 3.90E-75 |
| Epicardium | Bmp61 | 4.61E-78 | 1.16253162 | 0.845 | 0.4 | 9.22E-75 |
| Epicardium | Stard102 | 5.99E-77 | 1.16967184 | 0.466 | 0.117 | 1.20E-73 |
| Epicardium | S100a6 | 2.40E-76 | 1.95293875 | 0.685 | 0.262 | 4.81E-73 |
| Epicardium | Steap4 | 9.48E-76 | 0.54736332 | 0.61 | 0.204 | 1.90E-72 |
| Epicardium | Tenm31 | 2.07E-75 | 1.67887095 | 0.7 | 0.368 | 4.14E-72 |

| | | | | | | |
|---|---|---|---|---|---|---|
| Epicardium | Ppp2r2b | 5.02E-75 | 2.91292331 | 0.565 | 0.24 | 1.00E-71 |
| Epicardium | Slc24a3 | 1.12E-74 | 0.90998425 | 0.692 | 0.279 | 2.23E-71 |
| Epicardium | AABR07034767.11 | 1.66E-74 | 1.18343624 | 0.765 | 0.38 | 3.31E-71 |
| Epicardium | Tnfaip8l31 | 1.85E-74 | 1.16466928 | 0.603 | 0.23 | 3.70E-71 |
| Epicardium | Cdh151 | 6.80E-74 | 0.61235836 | 0.343 | 0.047 | 1.36E-70 |
| Epicardium | LOC1009114862 | 1.13E-73 | 3.10689285 | 0.506 | 0.256 | 2.27E-70 |
| Epicardium | AABR07006724.12 | 1.22E-73 | 1.60170538 | 0.642 | 0.287 | 2.44E-70 |
| Epicardium | Klhl291 | 6.77E-73 | 1.49081066 | 0.726 | 0.368 | 1.35E-69 |
| Epicardium | Serping1 | 7.78E-73 | 1.17143886 | 0.675 | 0.254 | 1.56E-69 |
| Epicardium | Zfp385b | 2.99E-72 | 0.97940786 | 0.748 | 0.313 | 5.99E-69 |
| Epicardium | Slit2 | 3.46E-72 | 2.12672505 | 0.711 | 0.356 | 6.91E-69 |
| Epicardium | Fn11 | 6.77E-72 | 1.36810684 | 0.864 | 0.443 | 1.35E-68 |
| Epicardium | Itgb8 | 3.23E-71 | 2.50171425 | 0.653 | 0.36 | 6.46E-68 |
| Epicardium | Cd55 | 5.07E-71 | 1.08010905 | 0.649 | 0.293 | 1.01E-67 |
| Epicardium | Cobll11 | 5.51E-71 | 1.57483006 | 0.728 | 0.433 | 1.10E-67 |
| Epicardium | Tmsb4x | 7.19E-70 | 1.61927356 | 0.843 | 0.464 | 1.44E-66 |
| Epicardium | Spon11 | 9.36E-70 | 1.41341412 | 0.681 | 0.33 | 1.87E-66 |
| Epicardium | Trpm3 | 1.05E-69 | 0.57729825 | 0.657 | 0.217 | 2.10E-66 |
| Epicardium | Lurap1l | 1.52E-69 | 0.97001535 | 0.769 | 0.333 | 3.03E-66 |
| Epicardium | Col5a21 | 4.11E-69 | 1.15485181 | 0.931 | 0.571 | 8.22E-66 |
| Epicardium | Ddah1 | 6.74E-69 | 2.02162293 | 0.69 | 0.374 | 1.35E-65 |
| Epicardium | Olfml2a1 | 1.35E-68 | 0.34806344 | 0.58 | 0.209 | 2.70E-65 |
| Epicardium | Apoe | 1.79E-68 | 1.40315113 | 0.789 | 0.327 | 3.59E-65 |
| Epicardium | Abi3bp2 | 1.53E-67 | 1.0841931 | 0.916 | 0.556 | 3.06E-64 |
| Epicardium | Syn3 | 2.12E-67 | 2.7105312 | 0.625 | 0.357 | 4.23E-64 |
| Epicardium | Flvcr2 | 3.90E-67 | 1.55828334 | 0.647 | 0.267 | 7.79E-64 |

| | | | | | | |
|---|---|---|---|---|---|---|
| Epicardium | Prdm6 | 5.90E-67 | 1.55778142 | 0.716 | 0.343 | 1.18E-63 |
| Epicardium | Colec121 | 2.76E-66 | 1.27703579 | 0.765 | 0.467 | 5.53E-63 |
| Epicardium | S100a4 | 2.56E-65 | 0.32541067 | 0.606 | 0.227 | 5.11E-62 |
| Epicardium | Dsp1 | 3.00E-65 | 1.55041939 | 0.761 | 0.377 | 6.01E-62 |
| Epicardium | Afap1l2 | 4.13E-65 | 1.09478871 | 0.754 | 0.37 | 8.26E-62 |
| Epicardium | Col11a1 | 1.51E-64 | 0.32881685 | 0.696 | 0.313 | 3.03E-61 |
| Epicardium | Tmem1001 | 4.74E-64 | 1.62790676 | 0.638 | 0.27 | 9.47E-61 |
| Epicardium | Slit3 | 1.81E-63 | 1.75939909 | 0.692 | 0.363 | 3.61E-60 |
| Epicardium | Ugdh | 6.87E-63 | 0.74952456 | 0.688 | 0.334 | 1.37E-59 |
| Epicardium | Lgals31 | 1.71E-62 | 1.1687685 | 0.634 | 0.249 | 3.42E-59 |
| Epicardium | Col16a1 | 2.07E-62 | 1.36771929 | 0.675 | 0.32 | 4.14E-59 |
| Epicardium | Atp5f1e | 3.20E-62 | 0.79759781 | 0.726 | 0.333 | 6.39E-59 |
| Epicardium | Eda2r | 4.48E-62 | 0.44524275 | 0.705 | 0.309 | 8.95E-59 |
| Epicardium | Tbx51 | 2.56E-61 | 1.60056438 | 0.545 | 0.245 | 5.13E-58 |
| Epicardium | Dock82 | 1.83E-59 | 0.74100062 | 0.772 | 0.395 | 3.65E-56 |
| Epicardium | Pcdh71 | 4.51E-59 | 0.26372253 | 0.746 | 0.36 | 9.01E-56 |
| Epicardium | Itgbl11 | 5.31E-59 | 0.89895415 | 0.925 | 0.546 | 1.06E-55 |
| Epicardium | Arhgap221 | 8.32E-59 | 0.43663042 | 0.731 | 0.389 | 1.66E-55 |
| Epicardium | Nav22 | 5.76E-58 | 0.97767951 | 0.726 | 0.427 | 1.15E-54 |
| Epicardium | Btbd111 | 7.19E-58 | 0.95204114 | 0.726 | 0.343 | 1.44E-54 |
| Epicardium | Ror1 | 3.72E-56 | 1.02856556 | 0.748 | 0.366 | 7.44E-53 |
| Epicardium | Alcam1 | 4.54E-56 | 1.11813647 | 0.752 | 0.411 | 9.09E-53 |
| Epicardium | Tp631 | 1.01E-55 | 0.68193613 | 0.379 | 0.106 | 2.03E-52 |
| Epicardium | LOC100909 595 | 4.25E-55 | 0.79811049 | 0.612 | 0.279 | 8.51E-52 |
| Epicardium | Cox6a1 | 8.86E-55 | 0.84038486 | 0.655 | 0.299 | 1.77E-51 |
| Epicardium | Plcxd3 | 1.82E-54 | 1.97266451 | 0.69 | 0.277 | 3.64E-51 |
| Epicardium | P2rx7 | 2.08E-54 | 0.87626643 | 0.631 | 0.357 | 4.17E-51 |
| Epicardium | Gulp1 | 3.32E-54 | 0.88579675 | 0.787 | 0.445 | 6.64E-51 |

| | | | | | | |
|---|---|---|---|---|---|---|
| Epicardium | Bgn | 2.10E-53 | 1.1223603 | 0.778 | 0.385 | 4.20E-50 |
| Epicardium | Adk1 | 5.40E-52 | 0.93228666 | 0.853 | 0.51 | 1.08E-48 |
| Epicardium | Gm2a | 9.41E-52 | 0.95625976 | 0.545 | 0.261 | 1.88E-48 |
| Epicardium | Ccn1 | 1.08E-51 | 0.28400823 | 0.601 | 0.274 | 2.17E-48 |
| Epicardium | Enox1 | 3.64E-51 | 1.04858581 | 0.675 | 0.294 | 7.27E-48 |
| Epicardium | Adgrg6 | 1.20E-50 | 0.89622605 | 0.562 | 0.276 | 2.40E-47 |
| Epicardium | LOC100911932 | 2.41E-50 | 0.66473896 | 0.595 | 0.244 | 4.82E-47 |
| Epicardium | Brca11 | 1.10E-49 | 0.42477837 | 0.677 | 0.269 | 2.21E-46 |
| Epicardium | Eva1c2 | 2.04E-49 | 0.909248 | 0.744 | 0.42 | 4.08E-46 |
| Epicardium | Cers61 | 6.99E-49 | 0.82415515 | 0.789 | 0.442 | 1.40E-45 |
| Epicardium | Adamts21 | 4.15E-48 | 0.81245179 | 0.901 | 0.542 | 8.30E-45 |
| Epicardium | Clu1 | 5.29E-48 | 0.85500381 | 0.711 | 0.352 | 1.06E-44 |
| Epicardium | Csrp1 | 1.40E-47 | 0.43928909 | 0.778 | 0.373 | 2.79E-44 |
| Epicardium | Rnd3 | 1.98E-47 | 0.79012652 | 0.81 | 0.408 | 3.96E-44 |
| Epicardium | Il17rd | 3.80E-47 | 0.90776567 | 0.612 | 0.292 | 7.60E-44 |
| Epicardium | LOC100364435 | 6.01E-47 | 0.97940462 | 0.7 | 0.359 | 1.20E-43 |
| Epicardium | Abcc4 | 2.25E-46 | 0.7375144 | 0.728 | 0.397 | 4.50E-43 |
| Epicardium | Antxr11 | 3.49E-46 | 1.03913203 | 0.761 | 0.502 | 6.97E-43 |
| Epicardium | Ptn | 9.51E-46 | 0.57694561 | 0.718 | 0.354 | 1.90E-42 |
| Epicardium | Efnb21 | 1.10E-45 | 0.49850131 | 0.744 | 0.387 | 2.19E-42 |
| Epicardium | Sdc2 | 5.55E-45 | 0.78202885 | 0.722 | 0.364 | 1.11E-41 |
| Epicardium | Plekha71 | 1.43E-44 | 0.58794203 | 0.787 | 0.41 | 2.85E-41 |
| Epicardium | Sgcd1 | 5.53E-44 | 0.5800362 | 0.864 | 0.5 | 1.11E-40 |
| Epicardium | Ncam1 | 1.04E-43 | 0.89840785 | 0.692 | 0.405 | 2.09E-40 |
| Epicardium | Shb1 | 2.17E-42 | 0.92277452 | 0.832 | 0.47 | 4.35E-39 |
| Epicardium | Negr1 | 3.03E-42 | 1.2540677 | 0.627 | 0.353 | 6.05E-39 |
| Epicardium | Chn22 | 5.66E-42 | 0.6771545 | 0.754 | 0.398 | 1.13E-38 |

| | | | | | | |
|---|---|---|---|---|---|---|
| Epicardium | Gpcpd11 | 2.03E-41 | 0.79193749 | 0.761 | 0.452 | 4.06E-38 |
| Epicardium | Myo1f1 | 2.68E-41 | 0.25187335 | 0.642 | 0.334 | 5.37E-38 |
| Epicardium | Rxfp11 | 1.21E-40 | 0.74402293 | 0.565 | 0.228 | 2.41E-37 |
| Epicardium | Fstl5 | 2.03E-40 | 1.45070789 | 0.543 | 0.181 | 4.07E-37 |
| Epicardium | Tpt1 | 2.78E-40 | 0.52055915 | 0.739 | 0.32 | 5.56E-37 |
| Epicardium | Lepr | 1.05E-39 | 1.0931769 | 0.575 | 0.272 | 2.11E-36 |
| Epicardium | Mtss12 | 1.24E-39 | 0.84080054 | 0.916 | 0.613 | 2.48E-36 |
| Epicardium | Pcsk5 | 4.82E-38 | 0.74802079 | 0.72 | 0.444 | 9.64E-35 |
| Epicardium | Pcbp31 | 6.44E-38 | 0.8475407 | 0.72 | 0.38 | 1.29E-34 |
| Epicardium | Mnda | 1.12E-37 | 0.98785393 | 0.64 | 0.364 | 2.24E-34 |
| Epicardium | Nr4a3 | 1.20E-37 | 0.83622783 | 0.554 | 0.275 | 2.40E-34 |
| Epicardium | LOC108352650 | 1.97E-37 | 0.46765457 | 0.644 | 0.32 | 3.95E-34 |
| Epicardium | Atp5me | 2.73E-37 | 0.27385605 | 0.744 | 0.405 | 5.46E-34 |
| Epicardium | Mrvi11 | 9.68E-37 | 0.74959689 | 0.659 | 0.361 | 1.94E-33 |
| Epicardium | Fth1 | 4.76E-36 | 0.51189692 | 0.659 | 0.359 | 9.52E-33 |
| Epicardium | Trim71 | 4.81E-36 | 0.56885128 | 0.571 | 0.303 | 9.62E-33 |
| Epicardium | Bend6 | 1.02E-35 | 0.44614593 | 0.612 | 0.287 | 2.03E-32 |
| Epicardium | Ryr31 | 1.52E-35 | 0.32300242 | 0.659 | 0.331 | 3.04E-32 |
| Epicardium | Usp131 | 5.88E-35 | 0.60732978 | 0.724 | 0.405 | 1.18E-31 |
| Epicardium | Sparc | 6.99E-35 | 0.6842464 | 0.948 | 0.649 | 1.40E-31 |
| Epicardium | Sned1 | 3.25E-34 | 0.76992397 | 0.599 | 0.338 | 6.49E-31 |
| Epicardium | Myh10 | 3.54E-34 | 1.13790277 | 0.692 | 0.428 | 7.08E-31 |
| Epicardium | Adam12 | 7.56E-34 | 1.4762385 | 0.647 | 0.33 | 1.51E-30 |
| Epicardium | Loxl11 | 8.27E-34 | 0.30803721 | 0.75 | 0.414 | 1.65E-30 |
| Epicardium | Loxl2 | 2.39E-33 | 0.53169135 | 0.675 | 0.379 | 4.77E-30 |
| Epicardium | Hspb8 | 2.40E-33 | 0.64206126 | 0.657 | 0.387 | 4.80E-30 |
| Epicardium | Gja11 | 7.88E-33 | 0.74945818 | 0.716 | 0.444 | 1.58E-29 |
| Epicardium | Lsamp2 | 1.62E-32 | 1.23386405 | 0.567 | 0.312 | 3.24E-29 |

| | | | | | | |
|---|---|---|---|---|---|---|
| Epicardium | Asap31 | 2.23E-32 | 0.49430196 | 0.774 | 0.468 | 4.47E-29 |
| Epicardium | Pcdha131 | 1.16E-30 | 0.53534618 | 0.713 | 0.408 | 2.33E-27 |
| Epicardium | Dclk11 | 1.68E-30 | 0.4469638 | 0.761 | 0.439 | 3.35E-27 |
| Epicardium | Abca11 | 1.56E-28 | 0.28223867 | 0.804 | 0.467 | 3.12E-25 |
| Epicardium | Nckap52 | 4.73E-28 | 0.42862409 | 0.709 | 0.418 | 9.47E-25 |
| Epicardium | Col4a51 | 2.28E-27 | 0.40907104 | 0.819 | 0.542 | 4.56E-24 |
| Epicardium | Plxdc22 | 8.00E-27 | 0.65028225 | 0.916 | 0.632 | 1.60E-23 |
| Epicardium | Col14a11 | 3.24E-26 | 0.32710019 | 0.752 | 0.424 | 6.49E-23 |
| Epicardium | Srgap1 | 6.73E-26 | 0.39524859 | 0.838 | 0.524 | 1.35E-22 |
| Epicardium | Slc38a11 | 1.75E-25 | 0.28101823 | 0.655 | 0.403 | 3.50E-22 |
| Epicardium | B2m | 4.56E-25 | 0.38993607 | 0.772 | 0.484 | 9.12E-22 |
| Epicardium | Boc1 | 1.39E-24 | 0.6091071 | 0.808 | 0.514 | 2.77E-21 |
| Epicardium | Il1r1 | 7.19E-17 | 0.69856006 | 0.873 | 0.623 | 1.44E-13 |
| Neuronal Cells | Cdh19 | 0 | 7.84118655 | 0.845 | 0.079 | 0 |
| Neuronal Cells | Grik2 | 0 | 5.79624104 | 0.962 | 0.243 | 0 |
| Neuronal Cells | Sox10 | 0 | 8.48631694 | 0.704 | 0.008 | 0 |
| Neuronal Cells | Gldn | 0 | 8.59287897 | 0.748 | 0.06 | 0 |
| Neuronal Cells | Igsf111 | 0 | 7.14166885 | 0.809 | 0.136 | 0 |
| Neuronal Cells | Adam23 | 0 | 5.64867802 | 0.886 | 0.217 | 0 |
| Neuronal Cells | Kcnq3 | 0 | 8.43573233 | 0.818 | 0.164 | 0 |
| Neuronal Cells | Scn7a | 0 | 6.03571026 | 0.907 | 0.254 | 0 |

| | | | | | | |
|---|---|---|---|---|---|---|
| Neuronal Cells | Csmd1 | 0 | 7.46958499 | 0.736 | 0.091 | 0 |
| Neuronal Cells | Rasgef1c | 0 | 7.63384258 | 0.696 | 0.084 | 0 |
| Neuronal Cells | Ttyh1 | 0 | 7.24195602 | 0.69 | 0.085 | 0 |
| Neuronal Cells | Lgi4 | 0 | 5.77636706 | 0.829 | 0.227 | 0 |
| Neuronal Cells | Cadm2 | 0 | 7.67199062 | 0.707 | 0.106 | 0 |
| Neuronal Cells | Fign1 | 0 | 4.51795163 | 0.946 | 0.359 | 0 |
| Neuronal Cells | Chl1 | 0 | 7.19237073 | 0.725 | 0.146 | 0 |
| Neuronal Cells | Mlph | 0 | 7.91604611 | 0.612 | 0.036 | 0 |
| Neuronal Cells | Dab1 | 0 | 7.27644888 | 0.606 | 0.039 | 0 |
| Neuronal Cells | Erbb31 | 0 | 6.77050349 | 0.646 | 0.093 | 0 |
| Neuronal Cells | Tdrd12 | 0 | 8.43799567 | 0.562 | 0.015 | 0 |
| Neuronal Cells | Fgf5 | 0 | 8.96872235 | 0.533 | 0.001 | 0 |
| Neuronal Cells | L1cam | 0 | 7.19603397 | 0.6 | 0.07 | 0 |
| Neuronal Cells | Gfra31 | 0 | 7.10618433 | 0.555 | 0.04 | 0 |

| | | | | | | |
|---|---|---|---|---|---|---|
| Neuronal Cells | Tmprss11f | 0 | 7.24365359 | 0.519 | 0.013 | 0 |
| Neuronal Cells | Tfap2b | 0 | 7.33686518 | 0.549 | 0.056 | 0 |
| Neuronal Cells | Ngfr | 0 | 6.97958381 | 0.533 | 0.048 | 0 |
| Neuronal Cells | S100a3 | 0 | 2.32413284 | 0.53 | 0.053 | 0 |
| Neuronal Cells | Lhx3 | 0 | 6.6144021 | 0.472 | 0.005 | 0 |
| Neuronal Cells | Ptprz11 | 0 | 6.72574045 | 0.506 | 0.061 | 0 |
| Neuronal Cells | Cmtm5 | 0 | 7.81681759 | 0.441 | 0.02 | 0 |
| Neuronal Cells | Snca | 0 | 5.10463071 | 0.424 | 0.015 | 0 |
| Neuronal Cells | Il1a1 | 0 | 1.57385802 | 0.429 | 0.023 | 0 |
| Neuronal Cells | Apod | 0 | 4.28679966 | 0.443 | 0.039 | 0 |
| Neuronal Cells | Ptchd1 | 0 | 2.99868752 | 0.457 | 0.058 | 0 |
| Neuronal Cells | Hepacam | 0 | 8.10643919 | 0.411 | 0.017 | 0 |
| Neuronal Cells | Stum | 0 | 3.17515663 | 0.476 | 0.084 | 0 |
| Neuronal Cells | Slc6a15 | 0 | 6.17169882 | 0.4 | 0.025 | 0 |

| | | | | | | |
|---|---|---|---|---|---|---|
| Neuronal Cells | Col28a11 | 0 | 3.94659112 | 0.44 | 0.079 | 0 |
| Neuronal Cells | AC128394.2 | 0 | 2.86400106 | 0.37 | 0.015 | 0 |
| Neuronal Cells | Zfhx4 | 0 | 5.9561426 | 0.369 | 0.034 | 0 |
| Neuronal Cells | Slc18a2 | 0 | 5.45736388 | 0.373 | 0.04 | 0 |
| Neuronal Cells | NEWGENE-2134 | 0 | 3.226046 | 0.345 | 0.015 | 0 |
| Neuronal Cells | AABR07026483.1 | 0 | 7.2205582 | 0.324 | 0.048 | 0 |
| Neuronal Cells | Il1rapl1 | 5.36E-303 | 5.25333329 | 0.888 | 0.234 | 1.07E-299 |
| Neuronal Cells | Krt181 | 4.34E-298 | 1.30212529 | 0.472 | 0.05 | 8.69E-295 |
| Neuronal Cells | Cpvl2 | 1.28E-297 | 0.9245498 | 0.532 | 0.055 | 2.55E-294 |
| Neuronal Cells | Rassf41 | 1.74E-294 | 5.44592584 | 0.809 | 0.167 | 3.47E-291 |
| Neuronal Cells | Metrn | 5.95E-291 | 4.9638168 | 0.454 | 0.057 | 1.19E-287 |
| Neuronal Cells | Itgb42 | 2.66E-286 | 4.83637624 | 0.881 | 0.253 | 5.32E-283 |
| Neuronal Cells | AABR07026924.1 | 2.14E-279 | 4.24172373 | 0.302 | 0.025 | 4.28E-276 |
| Neuronal Cells | Pex5l1 | 2.30E-277 | 3.11652257 | 0.563 | 0.088 | 4.60E-274 |

| | | | | | | |
|---|---|---|---|---|---|---|
| Neuronal Cells | Col18a12 | 1.50E-274 | 3.66020578 | 0.921 | 0.367 | 3.01E-271 |
| Neuronal Cells | Fa2h | 1.83E-272 | 5.5605635 | 0.441 | 0.091 | 3.66E-269 |
| Neuronal Cells | Gfra2 | 5.63E-262 | 5.86526952 | 0.672 | 0.146 | 1.13E-258 |
| Neuronal Cells | Trhde | 1.76E-259 | 5.75746704 | 0.745 | 0.169 | 3.51E-256 |
| Neuronal Cells | Sorcs1 | 8.15E-257 | 5.64924799 | 0.799 | 0.209 | 1.63E-253 |
| Neuronal Cells | Hoxd3 | 1.44E-254 | 2.82821783 | 0.402 | 0.014 | 2.89E-251 |
| Neuronal Cells | Cnksr2 | 1.61E-251 | 6.62148499 | 0.682 | 0.153 | 3.22E-248 |
| Neuronal Cells | Insc | 8.20E-249 | 6.24605491 | 0.604 | 0.125 | 1.64E-245 |
| Neuronal Cells | Arpp21 | 1.61E-240 | 2.92040175 | 0.445 | 0.05 | 3.21E-237 |
| Neuronal Cells | AABR07003 030.2 | 4.82E-234 | 5.90699469 | 0.646 | 0.168 | 9.64E-231 |
| Neuronal Cells | Tenm32 | 8.23E-233 | 3.39183843 | 0.897 | 0.363 | 1.65E-229 |
| Neuronal Cells | Zfp5361 | 3.27E-225 | 6.20480885 | 0.579 | 0.164 | 6.54E-222 |
| Neuronal Cells | Slamf61 | 1.36E-224 | 0.71431946 | 0.457 | 0.063 | 2.73E-221 |
| Neuronal Cells | Pax5 | 2.06E-223 | 0.350128 | 0.541 | 0.085 | 4.11E-220 |

| | | | | | | |
|---|---|---|---|---|---|---|
| Neuronal Cells | AABR07049085.12 | 2.95E-217 | 3.80747025 | 0.866 | 0.335 | 5.91E-214 |
| Neuronal Cells | Aatk | 2.96E-215 | 4.69799647 | 0.796 | 0.281 | 5.92E-212 |
| Neuronal Cells | Atp10b | 4.73E-214 | 5.89126602 | 0.43 | 0.097 | 9.45E-211 |
| Neuronal Cells | Sema3e1 | 5.49E-214 | 6.07872123 | 0.671 | 0.169 | 1.10E-210 |
| Neuronal Cells | Sox62 | 3.32E-213 | 2.50431484 | 0.9 | 0.474 | 6.65E-210 |
| Neuronal Cells | Megf10 | 1.35E-212 | 5.78043797 | 0.598 | 0.172 | 2.69E-209 |
| Neuronal Cells | Afap1l21 | 2.11E-209 | 3.46754275 | 0.81 | 0.367 | 4.21E-206 |
| Neuronal Cells | Cpa6 | 6.46E-208 | 2.656211 | 0.557 | 0.094 | 1.29E-204 |
| Neuronal Cells | P2rx71 | 3.40E-207 | 4.46407898 | 0.791 | 0.353 | 6.81E-204 |
| Neuronal Cells | Zeb22 | 4.55E-203 | 1.70811902 | 0.981 | 0.708 | 9.10E-200 |
| Neuronal Cells | Nkain21 | 1.70E-202 | 5.32073739 | 0.652 | 0.12 | 3.40E-199 |
| Neuronal Cells | Adamts201 | 1.07E-201 | 4.73148542 | 0.582 | 0.12 | 2.15E-198 |
| Neuronal Cells | Alcam2 | 4.16E-201 | 2.84417084 | 0.854 | 0.407 | 8.32E-198 |
| Neuronal Cells | Gulp11 | 6.23E-200 | 3.07635681 | 0.884 | 0.441 | 1.25E-196 |

| | | | | | | |
|---|---|---|---|---|---|---|
| Neuronal Cells | Slco4a1 | 2.88E-199 | 1.36129926 | 0.294 | 0.029 | 5.75E-196 |
| Neuronal Cells | Ank31 | 5.82E-199 | 2.59848391 | 0.911 | 0.489 | 1.16E-195 |
| Neuronal Cells | Mboat2 | 1.06E-197 | 4.25087344 | 0.775 | 0.329 | 2.12E-194 |
| Neuronal Cells | Bcar31 | 1.10E-194 | 3.07149843 | 0.858 | 0.379 | 2.19E-191 |
| Neuronal Cells | Dclk12 | 1.32E-191 | 2.58676936 | 0.878 | 0.435 | 2.65E-188 |
| Neuronal Cells | Ntng11 | 2.18E-191 | 3.44923115 | 0.748 | 0.23 | 4.35E-188 |
| Neuronal Cells | Niban1 | 5.14E-190 | 2.85852573 | 0.883 | 0.455 | 1.03E-186 |
| Neuronal Cells | Matn2 | 2.33E-189 | 3.76156997 | 0.767 | 0.35 | 4.66E-186 |
| Neuronal Cells | Col16a11 | 2.39E-183 | 3.62712206 | 0.756 | 0.317 | 4.78E-180 |
| Neuronal Cells | Pde8b1 | 4.94E-182 | 2.66900724 | 0.734 | 0.235 | 9.89E-179 |
| Neuronal Cells | Tspan11 | 2.09E-181 | 4.42532551 | 0.672 | 0.239 | 4.19E-178 |
| Neuronal Cells | Adgrl31 | 7.38E-181 | 2.64418033 | 0.799 | 0.282 | 1.48E-177 |
| Neuronal Cells | Lrrtm4 | 8.43E-181 | 6.01184465 | 0.628 | 0.205 | 1.69E-177 |
| Neuronal Cells | Grin2b1 | 2.36E-180 | 3.46718615 | 0.367 | 0.093 | 4.72E-177 |

| | | | | | | |
|---|---|---|---|---|---|---|
| Neuronal Cells | Ephb21 | 6.32E-178 | 3.63783 | 0.704 | 0.222 | 1.26E-174 |
| Neuronal Cells | Shc41 | 1.05E-177 | 5.52558323 | 0.521 | 0.114 | 2.10E-174 |
| Neuronal Cells | Alk | 7.37E-172 | 5.78374542 | 0.633 | 0.218 | 1.47E-168 |
| Neuronal Cells | Ltk1 | 2.23E-170 | 2.82100183 | 0.472 | 0.133 | 4.46E-167 |
| Neuronal Cells | AABR07065531.261 | 1.64E-168 | 1.9857611 | 0.373 | 0.087 | 3.27E-165 |
| Neuronal Cells | Nlgn1 | 1.77E-167 | 6.36677699 | 0.476 | 0.134 | 3.54E-164 |
| Neuronal Cells | Gas7 | 1.85E-167 | 3.03047241 | 0.783 | 0.36 | 3.69E-164 |
| Neuronal Cells | Il1rapl2 | 8.58E-162 | 5.97010938 | 0.663 | 0.206 | 1.72E-158 |
| Neuronal Cells | Nrxn1 | 1.66E-161 | 3.18214337 | 0.839 | 0.409 | 3.33E-158 |
| Neuronal Cells | Fam78b1 | 1.95E-160 | 3.44932794 | 0.725 | 0.293 | 3.91E-157 |
| Neuronal Cells | Iqgap21 | 2.85E-160 | 3.49698193 | 0.723 | 0.345 | 5.70E-157 |
| Neuronal Cells | Adgrg61 | 1.31E-156 | 3.33433967 | 0.739 | 0.271 | 2.63E-153 |
| Neuronal Cells | Fam135b | 4.22E-155 | 2.47315182 | 0.367 | 0.087 | 8.45E-152 |
| Neuronal Cells | Fstl41 | 3.95E-153 | 2.64912423 | 0.351 | 0.099 | 7.89E-150 |

| | | | | | | |
|---|---|---|---|---|---|---|
| Neuronal Cells | Neurl11 | 1.96E-151 | 4.29430364 | 0.54 | 0.159 | 3.91E-148 |
| Neuronal Cells | Lef1 | 6.02E-150 | 3.27104159 | 0.641 | 0.202 | 1.20E-146 |
| Neuronal Cells | AABR07027581.12 | 2.91E-147 | 2.54385396 | 0.802 | 0.382 | 5.81E-144 |
| Neuronal Cells | Tgfa | 2.42E-144 | 2.40198932 | 0.541 | 0.144 | 4.84E-141 |
| Neuronal Cells | Ptprt | 4.45E-143 | 4.84464366 | 0.595 | 0.173 | 8.91E-140 |
| Neuronal Cells | Hmga21 | 8.00E-142 | 4.56733137 | 0.399 | 0.084 | 1.60E-138 |
| Neuronal Cells | Nav23 | 9.54E-142 | 2.06228669 | 0.856 | 0.423 | 1.91E-138 |
| Neuronal Cells | Fhl51 | 7.06E-141 | 1.2848722 | 0.521 | 0.092 | 1.41E-137 |
| Neuronal Cells | Slc35f1 | 8.37E-140 | 3.77713821 | 0.593 | 0.249 | 1.67E-136 |
| Neuronal Cells | Sorcs2 | 5.08E-139 | 4.15653307 | 0.62 | 0.266 | 1.02E-135 |
| Neuronal Cells | Sv2c | 3.62E-132 | 2.65670911 | 0.752 | 0.381 | 7.24E-129 |
| Neuronal Cells | Agmo1 | 5.93E-131 | 3.32720921 | 0.649 | 0.283 | 1.19E-127 |
| Neuronal Cells | Grid22 | 7.67E-127 | 5.50686037 | 0.421 | 0.156 | 1.53E-123 |
| Neuronal Cells | Abca8a | 1.22E-126 | 2.32739927 | 0.764 | 0.391 | 2.44E-123 |

| | | | | | | |
|---|---|---|---|---|---|---|
| Neuronal Cells | Arhgef261 | 5.61E-125 | 4.22336684 | 0.612 | 0.26 | 1.12E-121 |
| Neuronal Cells | Mctp12 | 1.19E-124 | 2.23525378 | 0.797 | 0.41 | 2.39E-121 |
| Neuronal Cells | Mcam1 | 2.14E-120 | 2.08930756 | 0.796 | 0.357 | 4.29E-117 |
| Neuronal Cells | Tnfrsf19 | 1.59E-117 | 4.20078952 | 0.456 | 0.175 | 3.17E-114 |
| Neuronal Cells | Gpm6b1 | 2.26E-117 | 2.38185333 | 0.769 | 0.347 | 4.53E-114 |
| Neuronal Cells | Egflam1 | 3.66E-116 | 2.33950555 | 0.714 | 0.308 | 7.32E-113 |
| Neuronal Cells | Tnc | 4.97E-116 | 4.01206651 | 0.641 | 0.25 | 9.94E-113 |
| Neuronal Cells | Sorbs21 | 1.27E-115 | 1.2530452 | 0.959 | 0.577 | 2.54E-112 |
| Neuronal Cells | Dync1i1 | 6.84E-115 | 1.60237483 | 0.522 | 0.135 | 1.37E-111 |
| Neuronal Cells | Sfrp4 | 3.50E-114 | 1.95258365 | 0.608 | 0.204 | 7.00E-111 |
| Neuronal Cells | Mapk101 | 4.55E-114 | 3.40643911 | 0.46 | 0.184 | 9.11E-111 |
| Neuronal Cells | Ptprj1 | 6.96E-114 | 1.74940249 | 0.837 | 0.488 | 1.39E-110 |
| Neuronal Cells | Ppp1r9a | 1.14E-113 | 2.60918874 | 0.696 | 0.288 | 2.29E-110 |
| Neuronal Cells | Gpd1 | 1.73E-113 | 1.31306375 | 0.603 | 0.206 | 3.47E-110 |

| | | | | | | |
|---|---|---|---|---|---|---|
| Neuronal Cells | Reln1 | 8.45E-113 | 1.88065451 | 0.669 | 0.216 | 1.69E-109 |
| Neuronal Cells | Art31 | 8.99E-112 | 2.17665462 | 0.778 | 0.366 | 1.80E-108 |
| Neuronal Cells | Nrcam | 1.54E-110 | 2.65601398 | 0.638 | 0.27 | 3.09E-107 |
| Neuronal Cells | Trpm31 | 2.12E-110 | 4.1216061 | 0.579 | 0.217 | 4.23E-107 |
| Neuronal Cells | Dok5 | 8.57E-109 | 1.82144243 | 0.582 | 0.178 | 1.71E-105 |
| Neuronal Cells | Cyfip21 | 3.15E-108 | 1.76190925 | 0.495 | 0.173 | 6.30E-105 |
| Neuronal Cells | Cobl1 | 5.54E-105 | 2.9217652 | 0.603 | 0.275 | 1.11E-101 |
| Neuronal Cells | Prss122 | 2.84E-103 | 2.01184273 | 0.451 | 0.133 | 5.67E-100 |
| Neuronal Cells | Lrrtm3 | 8.76E-99 | 2.26941931 | 0.691 | 0.352 | 1.75E-95 |
| Neuronal Cells | Sntb11 | 7.42E-98 | 1.80588503 | 0.78 | 0.453 | 1.48E-94 |
| Neuronal Cells | Eda1 | 7.95E-97 | 1.86895489 | 0.774 | 0.442 | 1.59E-93 |
| Neuronal Cells | Aff32 | 1.94E-96 | 1.78940751 | 0.793 | 0.425 | 3.87E-93 |
| Neuronal Cells | Hs3st3a11 | 5.63E-96 | 0.68221983 | 0.568 | 0.184 | 1.13E-92 |
| Neuronal Cells | Mettl241 | 6.78E-95 | 2.10759209 | 0.464 | 0.172 | 1.36E-91 |

| | | | | | | |
|---|---|---|---|---|---|---|
| Neuronal Cells | Col27a1 | 1.27E-94 | 2.33470585 | 0.709 | 0.371 | 2.54E-91 |
| Neuronal Cells | Sema3c1 | 2.85E-94 | 1.71788101 | 0.767 | 0.358 | 5.70E-91 |
| Neuronal Cells | Pfkfb1 | 1.34E-93 | 1.07041814 | 0.604 | 0.225 | 2.68E-90 |
| Neuronal Cells | Pak31 | 4.38E-91 | 0.2687129 | 0.451 | 0.176 | 8.76E-88 |
| Neuronal Cells | Lpcat2 | 9.02E-91 | 1.848266 | 0.668 | 0.291 | 1.80E-87 |
| Neuronal Cells | Sgcd2 | 3.32E-89 | 1.48168817 | 0.777 | 0.5 | 6.64E-86 |
| Neuronal Cells | Ildr21 | 1.80E-88 | 2.32644916 | 0.597 | 0.252 | 3.61E-85 |
| Neuronal Cells | Crlf1 | 2.13E-88 | 2.10035262 | 0.468 | 0.175 | 4.26E-85 |
| Neuronal Cells | Lrrc4c1 | 4.47E-88 | 1.34185538 | 0.679 | 0.305 | 8.95E-85 |
| Neuronal Cells | Neb | 5.96E-88 | 1.95917648 | 0.566 | 0.202 | 1.19E-84 |
| Neuronal Cells | Mx1 | 1.39E-85 | 0.39221356 | 0.589 | 0.236 | 2.77E-82 |
| Neuronal Cells | Olfml2a2 | 4.53E-85 | 3.92679981 | 0.464 | 0.21 | 9.07E-82 |
| Neuronal Cells | Nrg41 | 1.77E-84 | 0.51697732 | 0.411 | 0.091 | 3.55E-81 |
| Neuronal Cells | Abcb41 | 2.02E-84 | 2.1632143 | 0.695 | 0.391 | 4.03E-81 |

| | | | | | | |
|---|---|---|---|---|---|---|
| Neuronal Cells | Kcnh8 | 2.50E-83 | 5.61065285 | 0.476 | 0.208 | 5.00E-80 |
| Neuronal Cells | Dok63 | 7.79E-79 | 2.04259748 | 0.5 | 0.136 | 1.56E-75 |
| Neuronal Cells | Col5a3 | 6.22E-78 | 1.85451843 | 0.71 | 0.379 | 1.24E-74 |
| Neuronal Cells | P4ha31 | 7.01E-78 | 1.41905728 | 0.491 | 0.146 | 1.40E-74 |
| Neuronal Cells | Enox11 | 4.81E-77 | 2.04739652 | 0.65 | 0.292 | 9.61E-74 |
| Neuronal Cells | Slc10a6 | 2.18E-76 | 0.47145971 | 0.587 | 0.257 | 4.35E-73 |
| Neuronal Cells | Pappa1 | 5.16E-76 | 1.61110842 | 0.672 | 0.274 | 1.03E-72 |
| Neuronal Cells | Mapt1 | 3.19E-75 | 1.60669684 | 0.742 | 0.364 | 6.38E-72 |
| Neuronal Cells | Aspa | 8.26E-75 | 2.83065268 | 0.581 | 0.271 | 1.65E-71 |
| Neuronal Cells | Jph11 | 4.59E-73 | 2.04229495 | 0.625 | 0.302 | 9.17E-70 |
| Neuronal Cells | Hspb3 | 2.37E-72 | 0.25713793 | 0.533 | 0.194 | 4.74E-69 |
| Neuronal Cells | Megf111 | 1.19E-70 | 0.28268855 | 0.563 | 0.239 | 2.39E-67 |
| Neuronal Cells | Sntg2 | 2.54E-68 | 0.91490694 | 0.6 | 0.204 | 5.07E-65 |
| Neuronal Cells | Ctsz | 2.86E-67 | 0.88293494 | 0.498 | 0.194 | 5.73E-64 |

| | | | | | | |
|---|---|---|---|---|---|---|
| Neuronal Cells | Baiap2l12 | 1.69E-66 | 1.94302239 | 0.634 | 0.291 | 3.39E-63 |
| Neuronal Cells | Olfml2b | 8.92E-66 | 2.03849635 | 0.604 | 0.313 | 1.78E-62 |
| Neuronal Cells | Ldb22 | 5.14E-64 | 1.36073818 | 0.63 | 0.351 | 1.03E-60 |
| Neuronal Cells | Cdh8 | 2.80E-62 | 0.51931496 | 0.37 | 0.11 | 5.60E-59 |
| Neuronal Cells | Sh3bp21 | 4.41E-62 | 0.8761605 | 0.663 | 0.335 | 8.83E-59 |
| Neuronal Cells | Chn23 | 8.57E-62 | 1.32612867 | 0.755 | 0.396 | 1.71E-58 |
| Neuronal Cells | LOC103693323 | 7.30E-61 | 0.86082229 | 0.574 | 0.249 | 1.46E-57 |
| Neuronal Cells | Tmtc22 | 2.82E-59 | 1.17214919 | 0.755 | 0.406 | 5.63E-56 |
| Neuronal Cells | Apba11 | 1.16E-58 | 1.29620667 | 0.744 | 0.473 | 2.32E-55 |
| Neuronal Cells | Kcnc2 | 6.18E-56 | 0.73234227 | 0.633 | 0.305 | 1.24E-52 |
| Neuronal Cells | Grm8 | 2.68E-55 | 0.72813758 | 0.495 | 0.206 | 5.37E-52 |
| Neuronal Cells | Ano4 | 3.38E-53 | 0.80177103 | 0.603 | 0.268 | 6.75E-50 |
| Neuronal Cells | Myo101 | 1.53E-50 | 0.79179555 | 0.782 | 0.472 | 3.06E-47 |
| Neuronal Cells | Pclo | 5.02E-50 | 0.34767971 | 0.495 | 0.197 | 1.00E-46 |

| | | | | | | |
|---|---|---|---|---|---|---|
| Neuronal Cells | Mmp16 | 1.60E-47 | 1.45798886 | 0.541 | 0.272 | 3.21E-44 |
| Neuronal Cells | Slc44a5 | 3.01E-47 | 0.25905441 | 0.592 | 0.239 | 6.03E-44 |
| Neuronal Cells | Rcan1 | 2.49E-46 | 1.31336523 | 0.592 | 0.287 | 4.97E-43 |
| Neuronal Cells | Tnik1 | 5.33E-46 | 0.98502896 | 0.739 | 0.435 | 1.07E-42 |
| Neuronal Cells | Xkr41 | 3.13E-45 | 0.60311246 | 0.611 | 0.288 | 6.26E-42 |
| Neuronal Cells | Dpyd | 1.59E-44 | 0.94379769 | 0.616 | 0.291 | 3.18E-41 |
| Neuronal Cells | Irs2 | 1.88E-44 | 0.70584214 | 0.668 | 0.409 | 3.76E-41 |
| Neuronal Cells | Igfbp3 | 2.38E-42 | 0.30636141 | 0.59 | 0.286 | 4.76E-39 |
| Neuronal Cells | Plxdc23 | 3.04E-40 | 0.52571613 | 0.891 | 0.631 | 6.07E-37 |
| Neuronal Cells | Ddah11 | 5.40E-39 | 0.59982958 | 0.644 | 0.373 | 1.08E-35 |
| Neuronal Cells | Tnfaip6 | 5.44E-39 | 0.6938903 | 0.538 | 0.277 | 1.09E-35 |
| Neuronal Cells | Pla2r1 | 3.46E-37 | 0.38919853 | 0.559 | 0.269 | 6.91E-34 |
| Neuronal Cells | Gpx1 | 1.09E-36 | 0.4617659 | 0.601 | 0.311 | 2.18E-33 |
| Neuronal Cells | Prelp1 | 4.54E-36 | 0.4587013 | 0.478 | 0.199 | 9.07E-33 |

| | | | | | | |
|---|---|---|---|---|---|---|
| Neuronal Cells | Pcdh72 | 8.98E-34 | 0.41745927 | 0.633 | 0.36 | 1.80E-30 |
| Neuronal Cells | Ppp1r14c1 | 6.67E-33 | 0.44839566 | 0.617 | 0.355 | 1.33E-29 |
| Neuronal Cells | Atp5f1e1 | 1.06E-31 | 0.82407465 | 0.589 | 0.333 | 2.13E-28 |
| Neuronal Cells | Cd63 | 1.61E-30 | 0.86584317 | 0.544 | 0.263 | 3.21E-27 |
| Neuronal Cells | Maoa | 5.80E-29 | 0.31388014 | 0.608 | 0.343 | 1.16E-25 |
| Neuronal Cells | Pde10a | 1.23E-27 | 0.56074573 | 0.704 | 0.435 | 2.47E-24 |
| Neuronal Cells | Spon12 | 7.48E-27 | 0.34572131 | 0.59 | 0.33 | 1.50E-23 |
| Neuronal Cells | Cox8b | 2.66E-23 | 0.59023732 | 0.682 | 0.372 | 5.33E-20 |
| Neuronal Cells | Atp5mc3 | 2.44E-21 | 0.45677242 | 0.646 | 0.382 | 4.89E-18 |
| Neuronal Cells | Cst3 | 6.92E-21 | 0.41689691 | 0.619 | 0.319 | 1.38E-17 |
| Neuronal Cells | Ckm1 | 7.24E-21 | 0.40652633 | 0.682 | 0.387 | 1.45E-17 |
| Neuronal Cells | Glis31 | 6.65E-20 | 0.51536445 | 0.729 | 0.477 | 1.33E-16 |
