## Supplemental Table 7 for "Uncovering the Regional and Cell Specific Bioactivity of Injectable Extracellular Matrix Biomaterials in Myocardial Infarction through Spatial and Single Nucleus Transcriptomics"

| Supplementary Table 7. Cell-Specific Differentially Expressed Genes with ECM (Up) or Saline (Down) Treatment in the Subacute MI Model | | | | | | | |
|---|---|---|---|---|---|---|---|
| **Subsetted Cell Type** | **Direction** | **Gene** | **p_val** | **avg_log2FC** | **pct.1** | **pct.2** | **p_val_adj** |
| Macrophages | Up | Rnf207 | 5.52E-07 | 1.31061653 | 0.808 | 0.07 | 0.00110371 |
| Macrophages | Up | Speg | 2.15E-14 | 1.12351317 | 0.942 | 0.053 | 4.30E-11 |
| Macrophages | Up | Trabd2b | 1.75E-09 | 0.96310672 | 0.865 | 0.07 | 3.49E-06 |
| Macrophages | Up | Hspb7 | 2.30E-08 | 0.94453252 | 0.846 | 0.07 | 4.60E-05 |
| Macrophages | Up | Corin | 4.78E-10 | 0.89557563 | 0.885 | 0.07 | 9.56E-07 |
| Macrophages | Up | Mlip | 1.05E-14 | 0.87085707 | 0.981 | 0.088 | 2.10E-11 |
| Macrophages | Up | Uqcrq | 3.30E-08 | 0.87004015 | 0.827 | 0.053 | 6.60E-05 |
| Macrophages | Up | Lgals3 | 2.22E-08 | 0.81536455 | 0.846 | 0.07 | 4.43E-05 |
| Macrophages | Up | Blnk | 1.66E-16 | 0.7922847 | 0.942 | 0.018 | 3.32E-13 |
| Macrophages | Up | Cmya5 | 1.93E-12 | 0.78207762 | 0.942 | 0.088 | 3.87E-09 |
| Macrophages | Up | Acacb | 1.53E-05 | 0.76109831 | 0.808 | 0.123 | 0.03062721 |
| Macrophages | Up | Slc20a2 | 1.05E-09 | 0.72456413 | 0.885 | 0.088 | 2.11E-06 |
| Macrophages | Up | Cox4i1 | 7.58E-06 | 0.7015148 | 0.885 | 0.211 | 0.01515442 |
| Macrophages | Up | Tgfb3 | 6.11E-10 | 0.68706574 | 0.942 | 0.14 | 1.22E-06 |
| Macrophages | Up | Myh7b | 1.43E-09 | 0.65351312 | 0.827 | 0.018 | 2.86E-06 |
| Macrophages | Up | Ptgs1 | 5.50E-06 | 0.64743771 | 0.269 | 0.018 | 0.01099852 |
| Macrophages | Up | Atp1b1 | 3.93E-11 | 0.62695038 | 0.904 | 0.07 | 7.86E-08 |
| Macrophages | Up | Otulinl | 5.32E-06 | 0.61370803 | 0.981 | 0.298 | 0.01064529 |
| Macrophages | Up | Ptpn3 | 1.20E-09 | 0.60839239 | 0.904 | 0.105 | 2.40E-06 |
| Macrophages | Up | AABR070525 | 2.57E-08 | 0.59643917 | 0.846 | 0.07 | 5.14E-05 |
| Macrophages | Up | Alcam | 6.27E-06 | 0.59604479 | 0.923 | 0.281 | 0.0125366 |
| Macrophages | Up | Hs3st5 | 1.44E-05 | 0.59469474 | 0.75 | 0.035 | 0.02879841 |
| Macrophages | Up | Ccn5 | 3.97E-08 | 0.5801524 | 0.827 | 0.053 | 7.94E-05 |
| Macrophages | Up | Wnk2 | 5.08E-12 | 0.57611521 | 0.904 | 0.053 | 1.02E-08 |
| Macrophages | Up | Tyrobp | 7.73E-11 | 0.56886907 | 0.962 | 0.14 | 1.55E-07 |
| Macrophages | Up | Ezh2 | 1.45E-11 | 0.56648856 | 0.942 | 0.105 | 2.91E-08 |
| Macrophages | Up | Pygm | 5.50E-06 | 0.55985928 | 0.269 | 0.018 | 0.01099852 |
| Macrophages | Up | Mrc1 | 5.06E-10 | 0.55527159 | 0.865 | 0.053 | 1.01E-06 |
| Macrophages | Up | Creb5 | 3.01E-06 | 0.55238249 | 0.942 | 0.263 | 0.0060128 |
| Macrophages | Up | Trdn | 7.80E-09 | 0.54936934 | 0.885 | 0.105 | 1.56E-05 |
| Macrophages | Up | Hk2 | 3.40E-09 | 0.53740069 | 0.962 | 0.175 | 6.79E-06 |
| Macrophages | Up | Lyve1 | 1.22E-06 | 0.53661058 | 0.25 | 0 | 0.00243597 |
| Macrophages | Up | Ppm1e | 1.30E-12 | 0.53183464 | 0.865 | 0 | 2.61E-09 |
| Macrophages | Up | AABR070265 | 7.45E-07 | 0.50444925 | 0.769 | 0.018 | 0.00148931 |
| Macrophages | Up | S100a4 | 3.97E-08 | 0.49796325 | 0.827 | 0.053 | 7.94E-05 |
| Macrophages | Up | Cd4 | 1.86E-07 | 0.49646056 | 0.981 | 0.246 | 0.00037154 |
| Macrophages | Up | Cadm3 | 1.65E-05 | 0.49201331 | 0.788 | 0.088 | 0.03290562 |
| Macrophages | Up | Adamts9 | 9.17E-07 | 0.48946399 | 0.865 | 0.14 | 0.00183377 |

| | | | | | | | |
|---|---|---|---|---|---|---|---|
| Macrophages | Up | Alpl | 7.94E-12 | 0.46949405 | 0.885 | 0.035 | 1.59E-08 |
| Macrophages | Up | LOC687780 | 9.51E-06 | 0.45765554 | 0.808 | 0.105 | 0.01902365 |
| Macrophages | Up | Cotl1 | 1.17E-06 | 0.45494506 | 0.923 | 0.211 | 0.00234049 |
| Macrophages | Up | Coro6 | 4.05E-10 | 0.44152899 | 0.885 | 0.07 | 8.11E-07 |
| Macrophages | Up | Igf1 | 1.65E-10 | 0.43931238 | 0.981 | 0.158 | 3.29E-07 |
| Macrophages | Up | Nav2 | 5.11E-08 | 0.43248946 | 0.885 | 0.123 | 0.0001022 |
| Macrophages | Up | Sdk2 | 3.09E-13 | 0.42926044 | 0.981 | 0.105 | 6.18E-10 |
| Macrophages | Up | Cyth4 | 1.74E-06 | 0.42914091 | 0.942 | 0.246 | 0.00348902 |
| Macrophages | Up | Diaph3 | 5.33E-11 | 0.42643735 | 0.904 | 0.07 | 1.07E-07 |
| Macrophages | Up | Fam227b | 5.47E-14 | 0.42511645 | 0.904 | 0.018 | 1.09E-10 |
| Macrophages | Up | Cadps2 | 1.36E-09 | 0.42353343 | 0.942 | 0.14 | 2.72E-06 |
| Macrophages | Up | Ptafr | 4.43E-12 | 0.41894831 | 0.923 | 0.07 | 8.87E-09 |
| Macrophages | Up | Abca1 | 1.73E-11 | 0.41655088 | 1 | 0.158 | 3.46E-08 |
| Macrophages | Up | Unc5b | 5.43E-08 | 0.40937041 | 0.904 | 0.14 | 0.00010851 |
| Macrophages | Up | Ndrg4 | 6.46E-13 | 0.40712745 | 0.904 | 0.035 | 1.29E-09 |
| Macrophages | Up | Dsp | 4.99E-06 | 0.39939268 | 0.846 | 0.14 | 0.00998463 |
| Macrophages | Up | Tox3 | 1.22E-06 | 0.39771098 | 0.25 | 0 | 0.00243597 |
| Macrophages | Up | Fgd2 | 1.05E-08 | 0.39447485 | 0.904 | 0.123 | 2.11E-05 |
| Macrophages | Up | Jph2 | 2.98E-08 | 0.39278687 | 0.846 | 0.07 | 5.96E-05 |
| Macrophages | Up | Ckb | 3.06E-14 | 0.38198521 | 0.942 | 0.053 | 6.11E-11 |
| Macrophages | Up | Sdc2 | 7.23E-06 | 0.38060695 | 0.942 | 0.246 | 0.01446963 |
| Macrophages | Up | Ptcd3 | 2.10E-10 | 0.37716795 | 0.942 | 0.123 | 4.20E-07 |
| Macrophages | Up | AABR070544 | 2.15E-15 | 0.36884622 | 0.962 | 0.053 | 4.29E-12 |
| Macrophages | Up | Col22a1 | 1.81E-11 | 0.36367203 | 0.846 | 0 | 3.63E-08 |
| Macrophages | Up | Tgfb2 | 6.04E-08 | 0.36077575 | 0.904 | 0.14 | 0.00012075 |
| Macrophages | Up | Prkcq | 2.20E-10 | 0.35629184 | 0.827 | 0 | 4.40E-07 |
| Macrophages | Up | Dtna | 4.07E-07 | 0.3514606 | 0.788 | 0.035 | 0.00081464 |
| Macrophages | Up | Sh3gl2 | 6.75E-10 | 0.35052482 | 0.865 | 0.053 | 1.35E-06 |
| Macrophages | Up | Arhgef37 | 2.47E-12 | 0.34614208 | 0.827 | 0 | 4.94E-09 |
| Macrophages | Up | Atp5mg | 1.56E-05 | 0.34250757 | 0.885 | 0.211 | 0.03127757 |
| Macrophages | Up | Msr1 | 1.45E-10 | 0.34115029 | 0.846 | 0.018 | 2.89E-07 |
| Macrophages | Up | Ankh | 7.79E-09 | 0.31967177 | 0.904 | 0.123 | 1.56E-05 |
| Macrophages | Up | Ckmt2 | 1.37E-05 | 0.31689077 | 0.923 | 0.246 | 0.02736459 |
| Macrophages | Up | Apobec1 | 6.54E-08 | 0.31649569 | 0.808 | 0.035 | 0.00013076 |
| Macrophages | Up | Cass4 | 2.13E-16 | 0.31538429 | 0.923 | 0 | 4.26E-13 |
| Macrophages | Up | Cyba | 1.51E-05 | 0.31147789 | 0.904 | 0.228 | 0.03013412 |
| Macrophages | Up | Trim55 | 2.96E-13 | 0.31005968 | 0.962 | 0.088 | 5.92E-10 |
| Macrophages | Up | Slc43a2 | 3.09E-09 | 0.3092022 | 0.981 | 0.193 | 6.18E-06 |
| Macrophages | Up | Lmln | 7.74E-18 | 0.30848317 | 0.962 | 0.018 | 1.55E-14 |
| Macrophages | Up | Slc20a1 | 2.19E-06 | 0.30590723 | 0.885 | 0.175 | 0.00438904 |
| Macrophages | Up | Fras1 | 4.42E-06 | 0.29848683 | 0.75 | 0.018 | 0.00884556 |

| | | | | | | | |
|---|---|---|---|---|---|---|---|
| Macrophages | Up | LOC1009095 | 4.68E-11 | 0.29822668 | 0.904 | 0.07 | 9.36E-08 |
| Macrophages | Up | Srpk3 | 1.59E-16 | 0.29768725 | 0.962 | 0.035 | 3.18E-13 |
| Macrophages | Up | Abca17 | 7.45E-07 | 0.29730787 | 0.769 | 0.018 | 0.00148931 |
| Macrophages | Up | Syn3 | 5.54E-06 | 0.29493741 | 0.788 | 0.07 | 0.01108942 |
| Macrophages | Up | Tspan5 | 2.46E-05 | 0.2831481 | 0.885 | 0.211 | 0.04911674 |
| Macrophages | Up | Wipf3 | 2.48E-06 | 0.28221015 | 0.827 | 0.105 | 0.00496078 |
| Macrophages | Up | Acyp2 | 1.19E-06 | 0.28171089 | 0.865 | 0.14 | 0.00237411 |
| Macrophages | Up | Pln | 2.06E-07 | 0.28063649 | 0.962 | 0.228 | 0.00041192 |
| Macrophages | Up | Prdx5 | 3.85E-12 | 0.28038724 | 0.942 | 0.088 | 7.69E-09 |
| Macrophages | Up | Epha4 | 2.46E-07 | 0.27823248 | 0.904 | 0.158 | 0.00049266 |
| Macrophages | Up | Itga7 | 1.57E-05 | 0.27799481 | 0.75 | 0.035 | 0.03146299 |
| Macrophages | Up | Tmem196 | 7.03E-10 | 0.2773047 | 0.865 | 0.053 | 1.41E-06 |
| Macrophages | Up | LOC1036933 | 9.21E-13 | 0.27612018 | 0.885 | 0.018 | 1.84E-09 |
| Macrophages | Up | Ncam1 | 4.64E-12 | 0.27232107 | 0.923 | 0.07 | 9.28E-09 |
| Macrophages | Up | PCOLCE2 | 1.27E-06 | 0.27107171 | 0.731 | 0.018 | 0.00253837 |
| Macrophages | Up | Slit2 | 1.25E-05 | 0.26615743 | 0.769 | 0.07 | 0.02493949 |
| Macrophages | Up | Tmem51 | 6.77E-13 | 0.26358114 | 0.904 | 0.035 | 1.35E-09 |
| Macrophages | Up | Pde7a | 1.32E-07 | 0.26321658 | 0.981 | 0.228 | 0.00026483 |
| Macrophages | Up | Olr1 | 4.56E-06 | 0.25661548 | 0.75 | 0.018 | 0.00912629 |
| Macrophages | Up | Ighm | 5.27E-11 | 0.25125097 | 0.827 | 0 | 1.05E-07 |
| Macrophages | Down | Mpped2 | 2.43E-09 | -0.2524549 | 0.962 | 0.158 | 4.86E-06 |
| Macrophages | Down | Dmtn | 5.11E-10 | -0.2542173 | 0.865 | 0.07 | 1.02E-06 |
| Macrophages | Down | Gbp1 | 6.31E-10 | -0.2544123 | 0.846 | 0.053 | 1.26E-06 |
| Macrophages | Down | Pde5a | 8.00E-12 | -0.2550289 | 0.904 | 0.053 | 1.60E-08 |
| Macrophages | Down | Adgrl3 | 1.19E-07 | -0.2582537 | 0.885 | 0.123 | 0.00023899 |
| Macrophages | Down | Il1b | 3.36E-10 | -0.2591407 | 0.846 | 0.07 | 6.72E-07 |
| Macrophages | Down | Eva1c | 4.08E-06 | -0.260045 | 0.827 | 0.105 | 0.00815942 |
| Macrophages | Down | Ano2 | 1.80E-05 | -0.2624122 | 0.635 | 0.053 | 0.03599704 |
| Macrophages | Down | AABR070428 | 3.36E-06 | -0.263998 | 0.558 | 0.035 | 0.0067224 |
| Macrophages | Down | LOC691485 | 1.02E-14 | -0.2651291 | 0.942 | 0.053 | 2.05E-11 |
| Macrophages | Down | Enpp1 | 4.73E-07 | -0.2698992 | 0.962 | 0.211 | 0.00094682 |
| Macrophages | Down | Fcgr2b | 1.45E-05 | -0.2747418 | 0.827 | 0.123 | 0.02890213 |
| Macrophages | Down | Ano1 | 7.20E-10 | -0.2789966 | 0.885 | 0.07 | 1.44E-06 |
| Macrophages | Down | Adam12 | 4.61E-06 | -0.2836088 | 0.827 | 0.105 | 0.00922572 |
| Macrophages | Down | Mertk | 4.37E-10 | -0.2836907 | 0.923 | 0.105 | 8.74E-07 |
| Macrophages | Down | Scara5 | 6.11E-11 | -0.294233 | 0.942 | 0.105 | 1.22E-07 |
| Macrophages | Down | Hmox1 | 7.56E-08 | -0.2958344 | 0.808 | 0.035 | 0.00015111 |
| Macrophages | Down | Ninj2 | 7.46E-08 | -0.2991027 | 0.942 | 0.175 | 0.00014911 |
| Macrophages | Down | Kcnt2 | 4.22E-11 | -0.3009664 | 0.962 | 0.123 | 8.43E-08 |
| Macrophages | Down | Ros1 | 7.31E-09 | -0.3023559 | 0.846 | 0.053 | 1.46E-05 |
| Macrophages | Down | Aff2 | 2.22E-08 | -0.3070635 | 0.904 | 0.123 | 4.43E-05 |

| | | | | | | | |
|---|---|---|---|---|---|---|---|
| Macrophages | Down | Pltp | 5.78E-12 | -0.310865 | 0.942 | 0.088 | 1.16E-08 |
| Macrophages | Down | Csrp1 | 1.12E-05 | -0.3119451 | 0.865 | 0.158 | 0.02236939 |
| Macrophages | Down | Dipk2b | 2.78E-08 | -0.3163581 | 0.885 | 0.105 | 5.55E-05 |
| Macrophages | Down | Fam180a | 5.01E-13 | -0.317405 | 0.885 | 0.035 | 1.00E-09 |
| Macrophages | Down | Pla2r1 | 4.50E-13 | -0.3178126 | 0.904 | 0.053 | 9.00E-10 |
| Macrophages | Down | Cacna1e | 3.10E-08 | -0.3181559 | 0.865 | 0.088 | 6.21E-05 |
| Macrophages | Down | Tnc | 8.06E-10 | -0.3435861 | 0.827 | 0.035 | 1.61E-06 |
| Macrophages | Down | Gna15 | 4.42E-10 | -0.356595 | 0.962 | 0.14 | 8.84E-07 |
| Macrophages | Down | Meox1 | 1.28E-08 | -0.3571141 | 0.769 | 0.035 | 2.55E-05 |
| Macrophages | Down | Pappa1 | 1.11E-09 | -0.357942 | 0.846 | 0.035 | 2.23E-06 |
| Macrophages | Down | Frem1 | 2.54E-08 | -0.3588888 | 0.923 | 0.14 | 5.08E-05 |
| Macrophages | Down | Dcdc5 | 1.26E-11 | -0.3828705 | 0.885 | 0.07 | 2.52E-08 |
| Macrophages | Down | Ckap2 | 3.35E-11 | -0.3836953 | 0.808 | 0.035 | 6.71E-08 |
| Macrophages | Down | Itgb8 | 3.88E-06 | -0.389382 | 0.846 | 0.123 | 0.0077595 |
| Macrophages | Down | Cd163 | 3.25E-15 | -0.412761 | 0.962 | 0.053 | 6.51E-12 |
| Macrophages | Down | Dlg2 | 1.71E-08 | -0.4191306 | 0.962 | 0.175 | 3.42E-05 |
| Macrophages | Down | Pde8b | 2.07E-10 | -0.424878 | 0.885 | 0.07 | 4.13E-07 |
| Macrophages | Down | Plxnc1 | 8.37E-12 | -0.4265165 | 0.904 | 0.053 | 1.67E-08 |
| Macrophages | Down | Pla2g7 | 2.48E-06 | -0.4286349 | 0.942 | 0.211 | 0.00496406 |
| Macrophages | Down | Errfi1 | 5.54E-07 | -0.4322321 | 0.923 | 0.175 | 0.00110783 |
| Macrophages | Down | Cenpf | 1.02E-14 | -0.4339788 | 0.942 | 0.053 | 2.05E-11 |
| Macrophages | Down | Lbh | 7.81E-06 | -0.4359374 | 0.981 | 0.263 | 0.01561132 |
| Macrophages | Down | AC134204.1 | 1.48E-07 | -0.4472458 | 0.923 | 0.158 | 0.00029505 |
| Macrophages | Down | Adamts17 | 4.41E-10 | -0.4770588 | 0.942 | 0.123 | 8.81E-07 |
| Macrophages | Down | Cd300lb | 1.60E-07 | -0.4850643 | 0.865 | 0.105 | 0.00031903 |
| Macrophages | Down | Art3 | 8.20E-06 | -0.4903199 | 0.731 | 0.053 | 0.0164067 |
| Macrophages | Down | Egflam | 3.59E-08 | -0.493646 | 0.865 | 0.088 | 7.18E-05 |
| Macrophages | Down | Akap12 | 1.08E-05 | -0.5246881 | 0.923 | 0.211 | 0.02152184 |
| Macrophages | Down | LOC24906 | 2.14E-09 | -0.5252457 | 0.827 | 0.088 | 4.27E-06 |
| Macrophages | Down | Kcnab1 | 1.78E-11 | -0.5261355 | 0.923 | 0.088 | 3.57E-08 |
| Macrophages | Down | Gria4 | 2.41E-05 | -0.5284422 | 0.808 | 0.105 | 0.04819873 |
| Macrophages | Down | Galnt16 | 3.48E-09 | -0.5409648 | 0.923 | 0.123 | 6.97E-06 |
| Macrophages | Down | Btbd11 | 1.67E-07 | -0.5431332 | 0.904 | 0.14 | 0.00033324 |
| Macrophages | Down | Adra1a | 4.25E-06 | -0.5553387 | 0.846 | 0.123 | 0.00850781 |
| Macrophages | Down | AABR070322 | 2.90E-07 | -0.6065309 | 0.846 | 0.158 | 0.00058018 |
| Macrophages | Down | Cyp4f18 | 6.10E-10 | -0.6461471 | 0.846 | 0.105 | 1.22E-06 |
| Macrophages | Down | RT1-Db1 | 3.21E-08 | -0.6540698 | 0.885 | 0.105 | 6.43E-05 |
| Macrophages | Down | Mob3b | 5.10E-09 | -0.6634277 | 0.885 | 0.088 | 1.02E-05 |
| Macrophages | Down | Fcgr3a | 1.73E-07 | -0.7203719 | 0.808 | 0.123 | 0.00034601 |
| Macrophages | Down | Dgkb | 1.09E-06 | -0.8049233 | 0.981 | 0.228 | 0.00218965 |
| Macrophages | Down | Mx1 | 3.01E-08 | -0.8329831 | 0.962 | 0.193 | 6.01E-05 |

| | | | | | | | |
|---|---|---|---|---|---|---|---|
| Endothelial Cells | Up | Srpx | 1.46E-119 | 0.91971594 | 0.954 | 0.305 | 2.93E-116 |
| Endothelial Cells | Up | Plcb1 | 2.40E-50 | 0.76730204 | 0.97 | 0.493 | 4.80E-47 |
| Endothelial Cells | Up | Lims2 | 9.47E-39 | 0.74127148 | 0.938 | 0.522 | 1.89E-35 |
| Endothelial Cells | Up | Slc26a10 | 3.82E-30 | 0.70950556 | 0.963 | 0.607 | 7.64E-27 |
| Endothelial Cells | Up | Dnm3 | 1.08E-29 | 0.66893805 | 0.991 | 0.705 | 2.15E-26 |
| Endothelial Cells | Up | Unc13b | 1.97E-71 | 0.6561294 | 0.95 | 0.407 | 3.95E-68 |
| Endothelial Cells | Up | Rimbp2 | 1.11E-71 | 0.64551355 | 0.875 | 0.301 | 2.22E-68 |
| Endothelial Cells | Up | Lmo7 | 4.05E-14 | 0.55676211 | 0.875 | 0.574 | 8.11E-11 |
| Endothelial Cells | Up | Hmcn1 | 1.60E-21 | 0.55238364 | 0.991 | 0.654 | 3.20E-18 |
| Endothelial Cells | Up | Slc6a6 | 1.58E-30 | 0.53764212 | 0.941 | 0.536 | 3.17E-27 |
| Endothelial Cells | Up | Rasa4 | 1.74E-140 | 0.52172848 | 0.954 | 0.275 | 3.48E-137 |
| Endothelial Cells | Up | St6galnac3 | 7.87E-39 | 0.50270931 | 0.86 | 0.371 | 1.57E-35 |
| Endothelial Cells | Up | Epha4 | 2.62E-69 | 0.49630781 | 0.925 | 0.371 | 5.23E-66 |
| Endothelial Cells | Up | Meox2 | 5.04E-18 | 0.48253561 | 0.97 | 0.651 | 1.01E-14 |
| Endothelial Cells | Up | Ltbp1 | 9.48E-19 | 0.46780826 | 0.797 | 0.393 | 1.90E-15 |
| Endothelial Cells | Up | Vegfc | 1.17E-34 | 0.46684987 | 0.764 | 0.28 | 2.35E-31 |
| Endothelial Cells | Up | Adgrd1 | 9.01E-84 | 0.4306769 | 0.751 | 0.122 | 1.80E-80 |
| Endothelial Cells | Up | Fbln5 | 1.59E-35 | 0.42918078 | 0.781 | 0.284 | 3.19E-32 |
| Endothelial Cells | Up | Mcf2l | 5.19E-15 | 0.42462689 | 0.888 | 0.577 | 1.04E-11 |
| Endothelial Cells | Up | Specc1 | 9.56E-33 | 0.41241099 | 0.714 | 0.209 | 1.91E-29 |
| Endothelial Cells | Up | Myo10 | 5.17E-13 | 0.40883909 | 0.957 | 0.662 | 1.03E-09 |
| Endothelial Cells | Up | AABR070606 | 5.87E-149 | 0.40353442 | 0.959 | 0.286 | 1.17E-145 |
| Endothelial Cells | Up | Cyp7b1 | 3.98E-20 | 0.38966186 | 0.762 | 0.311 | 7.96E-17 |
| Endothelial Cells | Up | Dipk2b | 5.76E-25 | 0.33746593 | 0.867 | 0.421 | 1.15E-21 |
| Endothelial Cells | Up | Bcar3 | 5.44E-19 | 0.3297154 | 0.706 | 0.25 | 1.09E-15 |
| Endothelial Cells | Up | Timp3 | 1.44E-29 | 0.32859066 | 0.904 | 0.453 | 2.89E-26 |
| Endothelial Cells | Up | Thsd7a | 6.48E-113 | 0.3275399 | 0.971 | 0.345 | 1.30E-109 |
| Endothelial Cells | Up | Sybu | 6.60E-11 | 0.31704138 | 0.648 | 0.229 | 1.32E-07 |
| Endothelial Cells | Up | Entpd1 | 3.72E-47 | 0.31064244 | 0.802 | 0.266 | 7.44E-44 |
| Endothelial Cells | Up | Pkhd1l1 | 7.31E-42 | 0.31035417 | 0.748 | 0.224 | 1.46E-38 |
| Endothelial Cells | Up | Prom1 | 3.29E-47 | 0.30627024 | 0.891 | 0.36 | 6.58E-44 |
| Endothelial Cells | Up | Akr1c15 | 1.65E-09 | 0.30066457 | 0.941 | 0.671 | 3.30E-06 |
| Endothelial Cells | Up | Dapk1 | 3.93E-94 | 0.29585794 | 0.874 | 0.243 | 7.87E-91 |
| Endothelial Cells | Up | Trak2 | 9.62E-37 | 0.28783697 | 0.829 | 0.328 | 1.92E-33 |
| Endothelial Cells | Up | Chn1 | 3.60E-36 | 0.28710021 | 0.852 | 0.348 | 7.20E-33 |
| Endothelial Cells | Up | Enpp1 | 5.77E-08 | 0.28594508 | 0.626 | 0.225 | 0.00011532 |
| Endothelial Cells | Up | Fap | 2.72E-38 | 0.28390956 | 0.781 | 0.272 | 5.45E-35 |
| Endothelial Cells | Up | Col14a1 | 4.55E-26 | 0.28374387 | 0.635 | 0.12 | 9.11E-23 |
| Endothelial Cells | Up | Adamts9 | 2.13E-14 | 0.28057773 | 0.872 | 0.503 | 4.27E-11 |
| Endothelial Cells | Up | Flrt2 | 3.83E-09 | 0.27570291 | 0.647 | 0.234 | 7.66E-06 |
| Endothelial Cells | Up | Sh3kbp1 | 6.21E-68 | 0.27566343 | 0.876 | 0.31 | 1.24E-64 |

| | | | | | | | |
|---|---|---|---|---|---|---|---|
| Endothelial Cells | Up | Ccn3 | 2.89E-09 | 0.271092 | 0.531 | 0.089 | 5.79E-06 |
| Endothelial Cells | Up | Slc35f1 | 4.83E-32 | 0.26822716 | 0.685 | 0.168 | 9.66E-29 |
| Endothelial Cells | Up | Aatk | 9.73E-06 | 0.25621081 | 0.584 | 0.185 | 0.01946897 |
| Endothelial Cells | Up | Sfrp2 | 1.03E-39 | 0.2535801 | 0.665 | 0.117 | 2.07E-36 |
| Endothelial Cells | Up | Dclk1 | 5.81E-19 | 0.25291122 | 0.638 | 0.164 | 1.16E-15 |
| Endothelial Cells | Up | Lpl | 1.53E-15 | 0.25211992 | 0.809 | 0.401 | 3.06E-12 |
| Endothelial Cells | Down | Speg | 9.20E-06 | -0.284573 | 0.585 | 0.179 | 0.0184022 |
| Endothelial Cells | Down | Usp18 | 2.04E-23 | -0.3393574 | 0.821 | 0.353 | 4.07E-20 |
| Endothelial Cells | Down | Ldhb | 1.16E-05 | -0.3723207 | 0.758 | 0.37 | 0.02314645 |
| Endothelial Cells | Down | Cox5b | 1.60E-06 | -0.4893364 | 0.613 | 0.201 | 0.00320145 |
| Endothelial Cells | Down | Kit | 4.97E-09 | -0.4939798 | 0.731 | 0.316 | 9.94E-06 |
| Endothelial Cells | Down | Hspb7 | 2.99E-17 | -0.5467197 | 0.74 | 0.271 | 5.99E-14 |
| Cardiomyocytes | Up | Glb1l2 | 3.48E-48 | 0.73574806 | 0.988 | 0.345 | 6.96E-45 |
| Cardiomyocytes | Up | Dgkb | 3.31E-17 | 0.63704891 | 0.995 | 0.633 | 6.63E-14 |
| Cardiomyocytes | Up | Ivns1abp | 4.28E-20 | 0.61734467 | 0.995 | 0.634 | 8.57E-17 |
| Cardiomyocytes | Up | Gria3 | 1.09E-21 | 0.61518074 | 0.993 | 0.555 | 2.19E-18 |
| Cardiomyocytes | Up | Enah | 1.40E-20 | 0.59305689 | 0.995 | 0.555 | 2.79E-17 |
| Cardiomyocytes | Up | Miga2 | 2.10E-24 | 0.58044433 | 0.988 | 0.551 | 4.20E-21 |
| Cardiomyocytes | Up | Cluh | 8.16E-26 | 0.55140558 | 0.99 | 0.536 | 1.63E-22 |
| Cardiomyocytes | Up | Pfkfb2 | 2.89E-28 | 0.55128981 | 0.97 | 0.444 | 5.78E-25 |
| Cardiomyocytes | Up | Unc45b | 2.67E-28 | 0.53650271 | 0.985 | 0.44 | 5.33E-25 |
| Cardiomyocytes | Up | Slc38a3 | 1.24E-14 | 0.53532026 | 0.998 | 0.601 | 2.49E-11 |
| Cardiomyocytes | Up | Cabcoco1 | 6.01E-16 | 0.51471163 | 0.993 | 0.58 | 1.20E-12 |
| Cardiomyocytes | Up | Fbn1 | 2.30E-31 | 0.51362179 | 0.84 | 0.249 | 4.61E-28 |
| Cardiomyocytes | Up | Nxn | 5.97E-17 | 0.51075038 | 0.693 | 0.125 | 1.19E-13 |
| Cardiomyocytes | Up | Tgfb2 | 1.62E-75 | 0.50574266 | 0.873 | 0.118 | 3.24E-72 |
| Cardiomyocytes | Up | Trim55 | 2.91E-13 | 0.49062551 | 0.995 | 0.623 | 5.81E-10 |
| Cardiomyocytes | Up | Chst15 | 1.23E-08 | 0.48545515 | 0.673 | 0.174 | 2.45E-05 |
| Cardiomyocytes | Up | Ndrg4 | 1.81E-25 | 0.47384449 | 0.99 | 0.461 | 3.61E-22 |
| Cardiomyocytes | Up | Trim50 | 1.14E-42 | 0.47250491 | 0.988 | 0.353 | 2.28E-39 |
| Cardiomyocytes | Up | Adhfe1 | 2.49E-31 | 0.46663247 | 0.98 | 0.416 | 4.98E-28 |
| Cardiomyocytes | Up | Anks1b | 2.13E-110 | 0.46028693 | 0.983 | 0.161 | 4.26E-107 |
| Cardiomyocytes | Up | Abcc4 | 2.26E-66 | 0.4564661 | 0.97 | 0.253 | 4.52E-63 |
| Cardiomyocytes | Up | Asb15 | 2.63E-39 | 0.44687443 | 0.985 | 0.37 | 5.27E-36 |
| Cardiomyocytes | Up | Jph1 | 5.77E-87 | 0.44245983 | 0.943 | 0.168 | 1.15E-83 |
| Cardiomyocytes | Up | AABR070347 | 3.34E-11 | 0.43479103 | 0.978 | 0.681 | 6.68E-08 |
| Cardiomyocytes | Up | Pank1 | 8.27E-49 | 0.43298009 | 0.97 | 0.322 | 1.65E-45 |
| Cardiomyocytes | Up | Ccdc80 | 1.84E-64 | 0.43285191 | 0.918 | 0.2 | 3.68E-61 |
| Cardiomyocytes | Up | Des | 1.44E-21 | 0.43119903 | 0.985 | 0.484 | 2.89E-18 |
| Cardiomyocytes | Up | Prkaa2 | 2.66E-14 | 0.42581839 | 0.985 | 0.555 | 5.33E-11 |
| Cardiomyocytes | Up | Rcan1 | 3.15E-125 | 0.42269275 | 0.958 | 0.117 | 6.29E-122 |

| | | | | | | | |
|---|---|---|---|---|---|---|---|
| Cardiomyocytes | Up | Fgf12 | 1.26E-37 | 0.42203969 | 0.953 | 0.33 | 2.53E-34 |
| Cardiomyocytes | Up | Srpk3 | 4.14E-15 | 0.41906618 | 0.88 | 0.414 | 8.27E-12 |
| Cardiomyocytes | Up | Coq8a | 3.93E-33 | 0.41073144 | 0.97 | 0.377 | 7.85E-30 |
| Cardiomyocytes | Up | Arhgap44 | 1.09E-32 | 0.40925005 | 0.983 | 0.397 | 2.18E-29 |
| Cardiomyocytes | Up | Actn1 | 1.30E-89 | 0.40855837 | 0.94 | 0.165 | 2.61E-86 |
| Cardiomyocytes | Up | Myl6 | 2.85E-30 | 0.40554405 | 0.973 | 0.394 | 5.70E-27 |
| Cardiomyocytes | Up | Adamts19 | 1.67E-50 | 0.40462387 | 0.985 | 0.318 | 3.33E-47 |
| Cardiomyocytes | Up | Hspb6 | 7.47E-61 | 0.4041466 | 0.975 | 0.261 | 1.49E-57 |
| Cardiomyocytes | Up | Cobll1 | 1.99E-31 | 0.40049866 | 0.88 | 0.287 | 3.98E-28 |
| Cardiomyocytes | Up | Ppm1e | 6.84E-48 | 0.39199779 | 0.87 | 0.198 | 1.37E-44 |
| Cardiomyocytes | Up | Lrrc4c | 1.45E-71 | 0.38781938 | 0.81 | 0.068 | 2.90E-68 |
| Cardiomyocytes | Up | Lgr6 | 9.16E-26 | 0.37912833 | 0.978 | 0.438 | 1.83E-22 |
| Cardiomyocytes | Up | Rgs6 | 4.33E-36 | 0.36804828 | 0.993 | 0.374 | 8.66E-33 |
| Cardiomyocytes | Up | Ppara | 2.05E-07 | 0.36207588 | 0.998 | 0.678 | 0.00040986 |
| Cardiomyocytes | Up | Arl15 | 2.90E-27 | 0.35591898 | 0.945 | 0.408 | 5.80E-24 |
| Cardiomyocytes | Up | Myoz2 | 5.94E-07 | 0.3534311 | 0.983 | 0.71 | 0.00118894 |
| Cardiomyocytes | Up | Slc25a13 | 3.11E-27 | 0.34897281 | 0.99 | 0.424 | 6.22E-24 |
| Cardiomyocytes | Up | Dclk1 | 1.37E-37 | 0.34531261 | 0.726 | 0.063 | 2.74E-34 |
| Cardiomyocytes | Up | Sptb | 2.93E-22 | 0.33873335 | 0.995 | 0.485 | 5.86E-19 |
| Cardiomyocytes | Up | Slc25a4 | 3.21E-16 | 0.33870826 | 0.99 | 0.579 | 6.42E-13 |
| Cardiomyocytes | Up | Palmd | 2.99E-109 | 0.33853731 | 0.985 | 0.16 | 5.99E-106 |
| Cardiomyocytes | Up | Creb5 | 3.66E-48 | 0.33724839 | 0.82 | 0.139 | 7.31E-45 |
| Cardiomyocytes | Up | Ckmt2 | 3.63E-08 | 0.33650708 | 0.915 | 0.582 | 7.27E-05 |
| Cardiomyocytes | Up | Popdc2 | 5.60E-18 | 0.33606587 | 0.995 | 0.52 | 1.12E-14 |
| Cardiomyocytes | Up | Tmem116 | 1.51E-56 | 0.33605721 | 0.965 | 0.276 | 3.02E-53 |
| Cardiomyocytes | Up | Fgf13 | 6.63E-26 | 0.33602413 | 0.9 | 0.338 | 1.33E-22 |
| Cardiomyocytes | Up | Flna | 3.03E-63 | 0.3346278 | 0.833 | 0.11 | 6.05E-60 |
| Cardiomyocytes | Up | Mlf1 | 4.84E-47 | 0.33368633 | 0.98 | 0.316 | 9.69E-44 |
| Cardiomyocytes | Up | Ankrd9 | 8.43E-21 | 0.33362608 | 0.878 | 0.361 | 1.69E-17 |
| Cardiomyocytes | Up | Mylk | 2.05E-104 | 0.33281707 | 0.928 | 0.119 | 4.10E-101 |
| Cardiomyocytes | Up | Ppp1r9a | 8.41E-75 | 0.33246026 | 0.935 | 0.181 | 1.68E-71 |
| Cardiomyocytes | Up | Myo18b | 9.38E-08 | 0.3271106 | 0.978 | 0.615 | 0.00018768 |
| Cardiomyocytes | Up | Rnf43 | 1.90E-108 | 0.32549938 | 0.868 | 0.056 | 3.79E-105 |
| Cardiomyocytes | Up | Fgf9 | 1.60E-141 | 0.325157 | 0.938 | 0.071 | 3.21E-138 |
| Cardiomyocytes | Up | Phyh | 7.07E-09 | 0.32256642 | 0.97 | 0.678 | 1.41E-05 |
| Cardiomyocytes | Up | Inpp4b | 1.37E-07 | 0.32200074 | 0.935 | 0.654 | 0.00027323 |
| Cardiomyocytes | Up | Zeb2 | 2.07E-17 | 0.32104407 | 0.843 | 0.324 | 4.15E-14 |
| Cardiomyocytes | Up | Fbxo40 | 1.42E-50 | 0.31785964 | 0.988 | 0.3 | 2.84E-47 |
| Cardiomyocytes | Up | Adamts2 | 1.46E-104 | 0.31720953 | 0.895 | 0.084 | 2.93E-101 |
| Cardiomyocytes | Up | Astn2 | 5.02E-153 | 0.31583183 | 0.953 | 0.072 | 1.00E-149 |
| Cardiomyocytes | Up | Robo1 | 3.75E-75 | 0.31307389 | 0.835 | 0.084 | 7.50E-72 |

| | | | | | | | |
|---|---|---|---|---|---|---|---|
| Cardiomyocytes | Up | Pcdh19 | 1.08E-22 | 0.31220707 | 0.676 | 0.067 | 2.15E-19 |
| Cardiomyocytes | Up | Emcn | 1.44E-10 | 0.3105289 | 0.628 | 0.082 | 2.89E-07 |
| Cardiomyocytes | Up | Rbm24 | 2.67E-20 | 0.3097808 | 0.983 | 0.487 | 5.34E-17 |
| Cardiomyocytes | Up | Ano5 | 5.15E-77 | 0.3097417 | 0.98 | 0.227 | 1.03E-73 |
| Cardiomyocytes | Up | Slc38a1 | 5.76E-07 | 0.30829508 | 0.99 | 0.677 | 0.00115274 |
| Cardiomyocytes | Up | Ccn2 | 3.26E-88 | 0.3071826 | 0.943 | 0.162 | 6.53E-85 |
| Cardiomyocytes | Up | Cpq | 9.00E-89 | 0.30602797 | 0.928 | 0.149 | 1.80E-85 |
| Cardiomyocytes | Up | AABR070265 | 6.49E-32 | 0.30588806 | 0.978 | 0.377 | 1.30E-28 |
| Cardiomyocytes | Up | Frmd5 | 3.47E-45 | 0.3040076 | 0.918 | 0.253 | 6.93E-42 |
| Cardiomyocytes | Up | Ckm | 2.03E-13 | 0.3009723 | 0.935 | 0.508 | 4.07E-10 |
| Cardiomyocytes | Up | Rap1gap2 | 1.85E-07 | 0.2999436 | 0.95 | 0.572 | 0.00036989 |
| Cardiomyocytes | Up | Pcsk6 | 1.39E-29 | 0.29944432 | 0.968 | 0.393 | 2.79E-26 |
| Cardiomyocytes | Up | Kcng2 | 1.16E-15 | 0.29570641 | 0.983 | 0.508 | 2.31E-12 |
| Cardiomyocytes | Up | Frmd4b | 2.69E-142 | 0.29378795 | 0.973 | 0.102 | 5.37E-139 |
| Cardiomyocytes | Up | Tango2 | 1.15E-09 | 0.29357638 | 0.918 | 0.568 | 2.31E-06 |
| Cardiomyocytes | Up | Lgals1 | 7.20E-82 | 0.29160418 | 0.895 | 0.131 | 1.44E-78 |
| Cardiomyocytes | Up | Irs1 | 3.72E-46 | 0.29010817 | 0.975 | 0.308 | 7.43E-43 |
| Cardiomyocytes | Up | Acta1 | 3.66E-79 | 0.28530662 | 0.978 | 0.218 | 7.32E-76 |
| Cardiomyocytes | Up | Sctr | 1.20E-26 | 0.28260098 | 0.683 | 0.056 | 2.41E-23 |
| Cardiomyocytes | Up | Pcbp3 | 8.71E-17 | 0.27975705 | 0.935 | 0.441 | 1.74E-13 |
| Cardiomyocytes | Up | Lims2 | 8.23E-06 | 0.27864473 | 0.995 | 0.694 | 0.01645474 |
| Cardiomyocytes | Up | Samd12 | 9.39E-32 | 0.2785511 | 0.96 | 0.362 | 1.88E-28 |
| Cardiomyocytes | Up | Pygm | 1.83E-16 | 0.27665993 | 0.943 | 0.457 | 3.66E-13 |
| Cardiomyocytes | Up | Lmln | 2.04E-20 | 0.27314845 | 0.678 | 0.078 | 4.08E-17 |
| Cardiomyocytes | Up | Ntn1 | 1.56E-20 | 0.26844409 | 0.96 | 0.424 | 3.13E-17 |
| Cardiomyocytes | Up | Ralgapa2 | 6.86E-06 | 0.26594001 | 0.993 | 0.688 | 0.01371317 |
| Cardiomyocytes | Up | Pcsk5 | 5.70E-69 | 0.26480263 | 0.835 | 0.095 | 1.14E-65 |
| Cardiomyocytes | Up | Ky | 1.01E-28 | 0.26230139 | 0.993 | 0.428 | 2.01E-25 |
| Cardiomyocytes | Up | AABR070070 | 1.26E-36 | 0.26139577 | 0.915 | 0.295 | 2.52E-33 |
| Cardiomyocytes | Up | Asb18 | 3.68E-08 | 0.26004558 | 0.985 | 0.607 | 7.37E-05 |
| Cardiomyocytes | Up | Pcp4l1 | 1.76E-112 | 0.25962943 | 0.978 | 0.153 | 3.52E-109 |
| Cardiomyocytes | Up | Gramd1b | 4.26E-13 | 0.25569833 | 0.766 | 0.259 | 8.53E-10 |
| Cardiomyocytes | Up | Sema5a | 1.82E-38 | 0.25512747 | 0.91 | 0.271 | 3.63E-35 |
| Cardiomyocytes | Up | Fbxl2 | 3.40E-138 | 0.25375439 | 0.938 | 0.075 | 6.80E-135 |
| Cardiomyocytes | Up | Alpk2 | 2.01E-28 | 0.25268352 | 0.968 | 0.395 | 4.02E-25 |
| Cardiomyocytes | Up | Lbh | 5.01E-09 | 0.25259367 | 0.87 | 0.45 | 1.00E-05 |
| Cardiomyocytes | Up | Pip5k1b | 1.53E-55 | 0.25250443 | 0.983 | 0.291 | 3.07E-52 |
| Cardiomyocytes | Up | Ripor2 | 1.44E-30 | 0.25190908 | 0.958 | 0.361 | 2.88E-27 |
| Cardiomyocytes | Up | Lamc2 | 5.79E-87 | 0.25009023 | 0.948 | 0.176 | 1.16E-83 |
| Cardiomyocytes | Down | Gabrb2 | 2.46E-08 | -0.2722655 | 0.631 | 0.099 | 4.93E-05 |
| Cardiomyocytes | Down | Dcdc5 | 1.61E-32 | -0.7745068 | 0.973 | 0.307 | 3.21E-29 |

| | | | | | | | |
|---|---|---|---|---|---|---|---|
| Fibroblasts | Up | Heyl | 2.14E-45 | 0.80491083 | 0.851 | 0.489 | 4.28E-42 |
| Fibroblasts | Up | Cdh13 | 5.72E-38 | 0.73172201 | 0.737 | 0.351 | 1.14E-34 |
| Fibroblasts | Up | Ptprt | 7.00E-77 | 0.72895578 | 0.649 | 0.16 | 1.40E-73 |
| Fibroblasts | Up | Slco3a1 | 2.72E-41 | 0.66553359 | 0.852 | 0.495 | 5.44E-38 |
| Fibroblasts | Up | Crispld2 | 1.87E-41 | 0.65726233 | 0.865 | 0.503 | 3.74E-38 |
| Fibroblasts | Up | Kcnc2 | 1.24E-38 | 0.58532097 | 0.846 | 0.491 | 2.49E-35 |
| Fibroblasts | Up | Zbtb16 | 7.58E-48 | 0.57582237 | 0.904 | 0.506 | 1.52E-44 |
| Fibroblasts | Up | Col4a5 | 4.53E-26 | 0.55686927 | 0.888 | 0.597 | 9.07E-23 |
| Fibroblasts | Up | Rgs17 | 5.32E-114 | 0.54201759 | 0.739 | 0.234 | 1.06E-110 |
| Fibroblasts | Up | Vcan | 1.98E-37 | 0.50788012 | 0.813 | 0.429 | 3.96E-34 |
| Fibroblasts | Up | Sgip1 | 1.45E-50 | 0.50009979 | 0.747 | 0.327 | 2.90E-47 |
| Fibroblasts | Up | Enpp2 | 2.94E-70 | 0.48195141 | 0.734 | 0.266 | 5.87E-67 |
| Fibroblasts | Up | Uap1 | 2.81E-24 | 0.4676117 | 0.733 | 0.388 | 5.62E-21 |
| Fibroblasts | Up | Gucy1a2 | 4.89E-38 | 0.46069843 | 0.852 | 0.472 | 9.79E-35 |
| Fibroblasts | Up | Fgf10 | 2.21E-53 | 0.4558538 | 0.677 | 0.252 | 4.43E-50 |
| Fibroblasts | Up | Blnk | 8.41E-06 | 0.44814985 | 0.549 | 0.272 | 0.01682402 |
| Fibroblasts | Up | Col24a1 | 2.58E-09 | 0.4439557 | 0.498 | 0.169 | 5.15E-06 |
| Fibroblasts | Up | Adamts3 | 7.58E-104 | 0.43160197 | 0.756 | 0.248 | 1.52E-100 |
| Fibroblasts | Up | Tll2 | 1.13E-118 | 0.42199428 | 0.74 | 0.239 | 2.27E-115 |
| Fibroblasts | Up | Nova1 | 1.05E-59 | 0.4219357 | 0.641 | 0.179 | 2.10E-56 |
| Fibroblasts | Up | Zfp385b | 5.34E-19 | 0.41386084 | 0.713 | 0.391 | 1.07E-15 |
| Fibroblasts | Up | Colec12 | 1.42E-18 | 0.41373543 | 0.91 | 0.644 | 2.85E-15 |
| Fibroblasts | Up | Aldh1a1 | 1.07E-68 | 0.40715785 | 0.736 | 0.273 | 2.14E-65 |
| Fibroblasts | Up | Foxp2 | 1.71E-17 | 0.39056232 | 0.858 | 0.58 | 3.42E-14 |
| Fibroblasts | Up | Pcsk6 | 1.01E-17 | 0.39027675 | 0.883 | 0.628 | 2.03E-14 |
| Fibroblasts | Up | Gria1 | 8.35E-06 | 0.37687345 | 0.474 | 0.176 | 0.01669145 |
| Fibroblasts | Up | Gda | 1.55E-20 | 0.36402045 | 0.887 | 0.58 | 3.09E-17 |
| Fibroblasts | Up | Gpm6a | 4.25E-10 | 0.3570878 | 0.705 | 0.436 | 8.49E-07 |
| Fibroblasts | Up | Slc16a2 | 2.94E-44 | 0.35382911 | 0.71 | 0.305 | 5.88E-41 |
| Fibroblasts | Up | Lepr | 5.47E-116 | 0.34152416 | 0.732 | 0.223 | 1.09E-112 |
| Fibroblasts | Up | Crlf1 | 9.55E-25 | 0.33992861 | 0.535 | 0.155 | 1.91E-21 |
| Fibroblasts | Up | Egr1 | 2.99E-22 | 0.31900348 | 0.523 | 0.152 | 5.98E-19 |
| Fibroblasts | Up | Scara5 | 2.95E-11 | 0.30922701 | 0.564 | 0.242 | 5.90E-08 |
| Fibroblasts | Up | Negr1 | 9.56E-41 | 0.30831456 | 0.77 | 0.381 | 1.91E-37 |
| Fibroblasts | Up | Cald1 | 5.44E-17 | 0.30593833 | 0.812 | 0.522 | 1.09E-13 |
| Fibroblasts | Up | AC134204.1 | 9.50E-41 | 0.30332689 | 0.777 | 0.379 | 1.90E-37 |
| Fibroblasts | Up | Ppargc1b | 3.13E-14 | 0.29752733 | 0.626 | 0.312 | 6.27E-11 |
| Fibroblasts | Up | Astn2 | 1.70E-06 | 0.28991944 | 0.574 | 0.29 | 0.00339092 |
| Fibroblasts | Up | Cgnl1 | 3.90E-91 | 0.28945861 | 0.735 | 0.265 | 7.80E-88 |
| Fibroblasts | Up | Dgkb | 1.43E-25 | 0.28445808 | 0.717 | 0.354 | 2.87E-22 |
| Fibroblasts | Up | Nr4a1 | 3.37E-42 | 0.28299626 | 0.597 | 0.196 | 6.73E-39 |

| | | | | | | | |
|---|---|---|---|---|---|---|---|
| Fibroblasts | Up | Mid1 | 4.43E-08 | 0.27598417 | 0.668 | 0.4 | 8.86E-05 |
| Fibroblasts | Up | Syt1 | 4.22E-22 | 0.26944213 | 0.516 | 0.15 | 8.44E-19 |
| Fibroblasts | Up | Egfr | 8.29E-15 | 0.26806645 | 0.885 | 0.601 | 1.66E-11 |
| Fibroblasts | Up | Plekha7 | 2.87E-17 | 0.26471773 | 0.83 | 0.519 | 5.74E-14 |
| Fibroblasts | Up | Arhgap22 | 1.68E-27 | 0.26384901 | 0.67 | 0.308 | 3.37E-24 |
| Fibroblasts | Up | LOC1025530 | 5.17E-34 | 0.26304023 | 0.577 | 0.183 | 1.03E-30 |
| Fibroblasts | Up | Errfi1 | 6.28E-55 | 0.26269449 | 0.764 | 0.321 | 1.26E-51 |
| Fibroblasts | Up | Robo2 | 1.54E-17 | 0.26135361 | 0.49 | 0.123 | 3.08E-14 |
| Fibroblasts | Up | Slc22a23 | 1.35E-120 | 0.25796603 | 0.729 | 0.208 | 2.70E-117 |
| Fibroblasts | Up | Calcrl | 2.42E-11 | 0.25574411 | 0.811 | 0.549 | 4.85E-08 |
| Fibroblasts | Up | Trhde | 4.84E-13 | 0.25440275 | 0.507 | 0.184 | 9.67E-10 |
| Fibroblasts | Up | Ppl | 6.06E-37 | 0.25342804 | 0.622 | 0.225 | 1.21E-33 |
| Fibroblasts | Up | Asic2 | 1.49E-57 | 0.25260603 | 0.649 | 0.195 | 2.99E-54 |
| Fibroblasts | Down | Uqcrq | 6.56E-09 | -0.2501594 | 0.579 | 0.277 | 1.31E-05 |
| Fibroblasts | Down | Cdh2 | 1.79E-14 | -0.2513645 | 0.757 | 0.426 | 3.59E-11 |
| Fibroblasts | Down | Dsp | 7.97E-25 | -0.2646767 | 0.638 | 0.277 | 1.59E-21 |
| Fibroblasts | Down | Arhgap15 | 4.30E-20 | -0.281608 | 0.571 | 0.22 | 8.61E-17 |
| Fibroblasts | Down | Eef1a2 | 1.80E-17 | -0.2925551 | 0.555 | 0.206 | 3.60E-14 |
| Fibroblasts | Down | Lgals1 | 2.07E-11 | -0.298642 | 0.652 | 0.349 | 4.15E-08 |
| Fibroblasts | Down | Atp5mg | 1.32E-18 | -0.2992217 | 0.632 | 0.279 | 2.64E-15 |
| Fibroblasts | Down | Runx1 | 1.71E-34 | -0.3080664 | 0.643 | 0.229 | 3.42E-31 |
| Fibroblasts | Down | AABR070490 | 1.45E-11 | -0.3187705 | 0.615 | 0.291 | 2.89E-08 |
| Fibroblasts | Down | Uqcr10 | 6.39E-14 | -0.3572421 | 0.573 | 0.246 | 1.28E-10 |
| Fibroblasts | Down | Ckm | 7.97E-10 | -0.3892115 | 0.641 | 0.329 | 1.59E-06 |
| Fibroblasts | Down | Runx2 | 7.92E-06 | -0.4348416 | 0.599 | 0.314 | 0.01584573 |
| Fibroblasts | Down | Cox4i1 | 1.25E-13 | -0.4478284 | 0.718 | 0.401 | 2.51E-10 |
| Fibroblasts | Down | Tgfbi | 2.20E-48 | -0.4675254 | 0.688 | 0.254 | 4.39E-45 |
| T-cells | Up | Uap1 | 1.34E-15 | 0.96060482 | 0.914 | 0.184 | 2.69E-12 |
| T-cells | Up | Gxylt2 | 1.44E-15 | 0.74652904 | 0.981 | 0.245 | 2.88E-12 |
| T-cells | Up | Il1rl1 | 6.72E-08 | 0.69319541 | 0.314 | 0.037 | 0.0001345 |
| T-cells | Up | Thsd7b | 4.91E-08 | 0.65852194 | 0.695 | 0.037 | 9.83E-05 |
| T-cells | Up | C1qtnf7 | 3.38E-08 | 0.59701058 | 0.933 | 0.35 | 6.76E-05 |
| T-cells | Up | Egfr | 4.89E-14 | 0.54533971 | 0.933 | 0.233 | 9.78E-11 |
| T-cells | Up | Pid1 | 1.24E-06 | 0.52904221 | 0.971 | 0.429 | 0.00248321 |
| T-cells | Up | Cdon | 3.21E-26 | 0.51980402 | 0.952 | 0.117 | 6.42E-23 |
| T-cells | Up | Nrxn1 | 4.12E-07 | 0.51282677 | 0.781 | 0.172 | 0.00082376 |
| T-cells | Up | Errfi1 | 3.48E-16 | 0.5119014 | 0.838 | 0.098 | 6.96E-13 |
| T-cells | Up | Kcnn3 | 7.42E-20 | 0.50620443 | 0.905 | 0.123 | 1.48E-16 |
| T-cells | Up | Actn1 | 1.90E-11 | 0.4953168 | 0.857 | 0.184 | 3.80E-08 |
| T-cells | Up | Ighm | 4.22E-09 | 0.48628811 | 0.705 | 0.031 | 8.44E-06 |
| T-cells | Up | Ugdh | 5.71E-25 | 0.47009206 | 0.905 | 0.08 | 1.14E-21 |

| | | | | | | | |
|---|---|---|---|---|---|---|---|
| T-cells | Up | Tox | 1.69E-08 | 0.46976132 | 0.952 | 0.35 | 3.39E-05 |
| T-cells | Up | Myh10 | 2.33E-07 | 0.46484093 | 0.762 | 0.141 | 0.00046516 |
| T-cells | Up | Fap | 2.19E-16 | 0.45901586 | 0.914 | 0.172 | 4.37E-13 |
| T-cells | Up | Nox4 | 9.43E-19 | 0.44993373 | 0.924 | 0.153 | 1.89E-15 |
| T-cells | Up | Cd96 | 6.05E-17 | 0.43897641 | 0.981 | 0.233 | 1.21E-13 |
| T-cells | Up | Blk | 5.14E-10 | 0.43670287 | 0.305 | 0 | 1.03E-06 |
| T-cells | Up | Ftl1 | 1.60E-10 | 0.43065511 | 0.781 | 0.098 | 3.19E-07 |
| T-cells | Up | Eva1c | 1.15E-28 | 0.42010935 | 0.952 | 0.098 | 2.30E-25 |
| T-cells | Up | Slit3 | 6.31E-22 | 0.40791708 | 0.914 | 0.117 | 1.26E-18 |
| T-cells | Up | Rgs5 | 8.35E-17 | 0.40320416 | 0.857 | 0.104 | 1.67E-13 |
| T-cells | Up | Pparg | 4.21E-21 | 0.40071937 | 0.905 | 0.11 | 8.41E-18 |
| T-cells | Up | Col14a1 | 1.49E-11 | 0.39602577 | 0.943 | 0.258 | 2.99E-08 |
| T-cells | Up | Grik2 | 5.09E-06 | 0.39476592 | 0.371 | 0.074 | 0.01018347 |
| T-cells | Up | Serpine2 | 3.52E-34 | 0.38572811 | 0.933 | 0.043 | 7.05E-31 |
| T-cells | Up | Dmpk | 6.82E-20 | 0.37571665 | 0.943 | 0.166 | 1.36E-16 |
| T-cells | Up | LOC1009114 | 1.71E-24 | 0.36484316 | 0.838 | 0.037 | 3.43E-21 |
| T-cells | Up | LOC687780 | 1.56E-12 | 0.362896 | 0.81 | 0.104 | 3.12E-09 |
| T-cells | Up | Ankh | 9.04E-07 | 0.36170705 | 0.8 | 0.196 | 0.00180837 |
| T-cells | Up | Gata3 | 2.74E-08 | 0.36040875 | 0.333 | 0.074 | 5.48E-05 |
| T-cells | Up | Asb2 | 7.52E-07 | 0.32801725 | 0.8 | 0.196 | 0.00150348 |
| T-cells | Up | Lum | 4.51E-07 | 0.32751265 | 0.362 | 0.098 | 0.00090239 |
| T-cells | Up | Ebf2 | 1.02E-07 | 0.32017817 | 0.924 | 0.337 | 0.00020486 |
| T-cells | Up | Adamts3 | 1.75E-20 | 0.31621466 | 0.79 | 0.043 | 3.49E-17 |
| T-cells | Up | Bcl11a | 1.90E-08 | 0.31614898 | 0.562 | 0 | 3.80E-05 |
| T-cells | Up | Adamts15 | 9.12E-06 | 0.3155677 | 0.381 | 0.086 | 0.01824868 |
| T-cells | Up | Gcnt2 | 8.54E-24 | 0.31441552 | 0.905 | 0.086 | 1.71E-20 |
| T-cells | Up | LOC1003610 | 6.98E-22 | 0.30829231 | 0.924 | 0.123 | 1.40E-18 |
| T-cells | Up | Pde10a | 2.83E-11 | 0.30786587 | 0.829 | 0.147 | 5.65E-08 |
| T-cells | Up | Il33 | 1.87E-39 | 0.30652938 | 0.952 | 0.031 | 3.73E-36 |
| T-cells | Up | Mark1 | 5.10E-25 | 0.30250854 | 0.895 | 0.067 | 1.02E-21 |
| T-cells | Up | Has1 | 7.57E-14 | 0.29870358 | 0.552 | 0.018 | 1.51E-10 |
| T-cells | Up | Fbxo32 | 3.89E-11 | 0.29648507 | 0.8 | 0.11 | 7.79E-08 |
| T-cells | Up | Polr2m | 1.49E-07 | 0.29289005 | 0.8 | 0.178 | 0.00029815 |
| T-cells | Up | Crispld2 | 8.57E-21 | 0.29273857 | 0.895 | 0.104 | 1.71E-17 |
| T-cells | Up | Fermt3 | 9.09E-23 | 0.29185134 | 0.962 | 0.153 | 1.82E-19 |
| T-cells | Up | Sema3c | 2.15E-09 | 0.2890501 | 0.79 | 0.129 | 4.30E-06 |
| T-cells | Up | Sgip1 | 3.20E-07 | 0.28571462 | 0.352 | 0.092 | 0.0006406 |
| T-cells | Up | Sacs | 4.61E-13 | 0.28564323 | 0.924 | 0.221 | 9.22E-10 |
| T-cells | Up | Colgalt2 | 2.36E-13 | 0.28563611 | 0.933 | 0.227 | 4.71E-10 |
| T-cells | Up | Prkar2b | 1.00E-17 | 0.28208151 | 0.829 | 0.067 | 2.01E-14 |
| T-cells | Up | Gli2 | 2.84E-14 | 0.27879156 | 0.895 | 0.184 | 5.69E-11 |

| | | | | | | | |
|---|---|---|---|---|---|---|---|
| T-cells | Up | Ltbp2 | 1.00E-10 | 0.27607231 | 0.838 | 0.153 | 2.00E-07 |
| T-cells | Up | Uqcrq | 3.81E-08 | 0.27019004 | 0.762 | 0.117 | 7.62E-05 |
| T-cells | Up | Emid1 | 2.76E-09 | 0.26874019 | 0.324 | 0.074 | 5.52E-06 |
| T-cells | Up | Ank3 | 2.38E-10 | 0.26823534 | 0.867 | 0.202 | 4.76E-07 |
| T-cells | Up | Usp13 | 3.43E-29 | 0.26296971 | 0.933 | 0.074 | 6.86E-26 |
| T-cells | Up | Synpo2 | 4.01E-15 | 0.26147684 | 0.99 | 0.252 | 8.02E-12 |
| T-cells | Up | Rgcc | 9.32E-06 | 0.25711059 | 0.667 | 0.037 | 0.0186387 |
| T-cells | Up | Kank1 | 7.57E-26 | 0.25379442 | 0.933 | 0.098 | 1.51E-22 |
| T-cells | Up | Lrrc4c | 1.25E-23 | 0.25322083 | 0.905 | 0.086 | 2.49E-20 |
| T-cells | Up | Ezr | 9.67E-08 | 0.25190761 | 0.924 | 0.307 | 0.00019333 |
| T-cells | Up | Shb | 6.42E-16 | 0.25098938 | 0.905 | 0.166 | 1.28E-12 |
| T-cells | Up | Ifitm10 | 7.15E-20 | 0.25070782 | 0.819 | 0.043 | 1.43E-16 |
| T-cells | Up | Mki67 | 4.79E-10 | 0.2501086 | 0.305 | 0.043 | 9.57E-07 |
| T-cells | Down | Ndst3 | 3.77E-31 | -0.2557604 | 0.924 | 0.055 | 7.54E-28 |
| T-cells | Down | Kcng2 | 1.62E-19 | -0.2559087 | 0.829 | 0.043 | 3.24E-16 |
| T-cells | Down | Kcnt2 | 4.54E-18 | -0.2600838 | 0.99 | 0.215 | 9.08E-15 |
| T-cells | Down | Dipk2b | 1.04E-13 | -0.2700995 | 0.857 | 0.135 | 2.08E-10 |
| T-cells | Down | Anpep | 1.37E-26 | -0.270794 | 0.905 | 0.067 | 2.74E-23 |
| T-cells | Down | Kif20b | 1.77E-16 | -0.2804227 | 0.781 | 0.061 | 3.55E-13 |
| T-cells | Down | Stard10 | 7.89E-10 | -0.2826196 | 0.79 | 0.135 | 1.58E-06 |
| T-cells | Down | Rapgef5 | 1.28E-06 | -0.2839382 | 0.838 | 0.221 | 0.00255086 |
| T-cells | Down | Rbm24 | 6.35E-26 | -0.2934935 | 0.886 | 0.049 | 1.27E-22 |
| T-cells | Down | Dlgap1 | 5.95E-06 | -0.2946603 | 0.39 | 0.117 | 0.01190595 |
| T-cells | Down | Mcoln2 | 3.22E-22 | -0.2952186 | 0.733 | 0.061 | 6.44E-19 |
| T-cells | Down | Cryab | 1.34E-11 | -0.2974065 | 0.905 | 0.209 | 2.69E-08 |
| T-cells | Down | AABR070042 | 1.22E-07 | -0.2998705 | 0.695 | 0.067 | 0.00024419 |
| T-cells | Down | AABR070033 | 8.41E-30 | -0.3065471 | 0.695 | 0.012 | 1.68E-26 |
| T-cells | Down | Adap2 | 1.49E-08 | -0.3088405 | 0.724 | 0.086 | 2.97E-05 |
| T-cells | Down | Depdc1 | 1.06E-17 | -0.3173086 | 0.59 | 0.031 | 2.11E-14 |
| T-cells | Down | Pappa1 | 7.23E-26 | -0.3213281 | 0.905 | 0.067 | 1.45E-22 |
| T-cells | Down | Ly49i4 | 3.87E-13 | -0.3249492 | 0.752 | 0.061 | 7.73E-10 |
| T-cells | Down | Atp1b1 | 7.16E-17 | -0.3312018 | 0.905 | 0.141 | 1.43E-13 |
| T-cells | Down | Mctp2 | 1.73E-07 | -0.3458564 | 0.905 | 0.27 | 0.00034667 |
| T-cells | Down | Angpt1 | 6.65E-17 | -0.3548037 | 0.924 | 0.16 | 1.33E-13 |
| T-cells | Down | Nmnat2 | 5.41E-29 | -0.3563007 | 0.895 | 0.043 | 1.08E-25 |
| T-cells | Down | Mx1 | 2.64E-29 | -0.3659336 | 0.943 | 0.098 | 5.28E-26 |
| T-cells | Down | Slc39a8 | 1.60E-31 | -0.3807932 | 0.933 | 0.061 | 3.21E-28 |
| T-cells | Down | Slc16a7 | 2.55E-18 | -0.3836062 | 0.876 | 0.117 | 5.10E-15 |
| T-cells | Down | Mertk | 1.96E-05 | -0.3895283 | 0.733 | 0.129 | 0.03916637 |
| T-cells | Down | Tspan18 | 1.87E-06 | -0.4033106 | 0.886 | 0.276 | 0.00373598 |
| T-cells | Down | Ppp1r14c | 3.15E-06 | -0.4050297 | 0.371 | 0.08 | 0.00630139 |

| | | | | | | | |
|---|---|---|---|---|---|---|---|
| T-cells | Down | Tgfb2 | 3.57E-16 | -0.4131574 | 0.943 | 0.184 | 7.14E-13 |
| T-cells | Down | Itgb8 | 5.91E-18 | -0.4285384 | 0.924 | 0.147 | 1.18E-14 |
| T-cells | Down | Thsd7a | 1.70E-12 | -0.4388266 | 0.857 | 0.147 | 3.39E-09 |
| T-cells | Down | Zc3h12d | 6.96E-25 | -0.4487083 | 0.962 | 0.129 | 1.39E-21 |
| T-cells | Down | Ly49s6 | 2.61E-21 | -0.4620604 | 0.943 | 0.147 | 5.22E-18 |
| T-cells | Down | Slc26a10 | 1.68E-07 | -0.466758 | 0.81 | 0.178 | 0.00033671 |
| T-cells | Down | Ninj2 | 1.01E-13 | -0.5070844 | 0.867 | 0.141 | 2.02E-10 |
| Neural Cells | Up | Reln | 7.62E-31 | 0.82643728 | 0.939 | 0.226 | 1.52E-27 |
| Neural Cells | Up | Myh11 | 8.17E-13 | 0.65856618 | 0.718 | 0.079 | 1.63E-09 |
| Neural Cells | Up | Crip1 | 1.20E-06 | 0.63569708 | 0.74 | 0.222 | 0.00239748 |
| Neural Cells | Up | Hmga2 | 3.59E-07 | 0.60602012 | 0.621 | 0.033 | 0.00071706 |
| Neural Cells | Up | Aatk | 3.52E-11 | 0.5429216 | 0.977 | 0.498 | 7.03E-08 |
| Neural Cells | Up | Myl3 | 1.38E-11 | 0.49845801 | 0.975 | 0.46 | 2.77E-08 |
| Neural Cells | Up | Postn | 1.47E-10 | 0.4790811 | 0.827 | 0.285 | 2.94E-07 |
| Neural Cells | Up | Egfl8 | 5.72E-15 | 0.46923601 | 0.336 | 0.063 | 1.14E-11 |
| Neural Cells | Up | Lum | 5.92E-20 | 0.44462085 | 0.802 | 0.126 | 1.18E-16 |
| Neural Cells | Up | Shank3 | 1.52E-38 | 0.43164882 | 0.992 | 0.222 | 3.04E-35 |
| Neural Cells | Up | AABR070581 | 9.65E-15 | 0.43017857 | 0.967 | 0.381 | 1.93E-11 |
| Neural Cells | Up | AC134204.1 | 1.70E-51 | 0.42700169 | 0.906 | 0.067 | 3.39E-48 |
| Neural Cells | Up | S100a6 | 1.79E-05 | 0.42300139 | 0.667 | 0.117 | 0.0357915 |
| Neural Cells | Up | Megf10 | 3.29E-15 | 0.41400558 | 0.83 | 0.218 | 6.58E-12 |
| Neural Cells | Up | Shtn1 | 1.21E-44 | 0.40480314 | 0.753 | 0.017 | 2.42E-41 |
| Neural Cells | Up | Igfbp4 | 2.19E-43 | 0.40462019 | 0.878 | 0.071 | 4.39E-40 |
| Neural Cells | Up | LOC1009118 | 2.30E-11 | 0.39935692 | 0.763 | 0.167 | 4.59E-08 |
| Neural Cells | Up | Pdgfrb | 2.72E-33 | 0.3961926 | 0.959 | 0.222 | 5.43E-30 |
| Neural Cells | Up | P2rx7 | 1.89E-07 | 0.39539848 | 0.962 | 0.51 | 0.00037779 |
| Neural Cells | Up | Ebf2 | 1.80E-10 | 0.39229633 | 0.761 | 0.18 | 3.59E-07 |
| Neural Cells | Up | Tmem196 | 1.12E-19 | 0.38523552 | 0.88 | 0.23 | 2.24E-16 |
| Neural Cells | Up | Cd63 | 1.66E-20 | 0.38279639 | 0.802 | 0.121 | 3.32E-17 |
| Neural Cells | Up | Fabp4 | 8.86E-39 | 0.37283875 | 0.954 | 0.18 | 1.77E-35 |
| Neural Cells | Up | Acta2 | 1.18E-44 | 0.36814249 | 0.845 | 0.029 | 2.35E-41 |
| Neural Cells | Up | Emid1 | 1.78E-10 | 0.3564946 | 0.364 | 0.042 | 3.57E-07 |
| Neural Cells | Up | Art3 | 5.91E-09 | 0.34847181 | 0.969 | 0.464 | 1.18E-05 |
| Neural Cells | Up | Serpinf1 | 7.40E-22 | 0.34788916 | 0.761 | 0.059 | 1.48E-18 |
| Neural Cells | Up | Fap | 2.99E-38 | 0.34738884 | 0.888 | 0.109 | 5.98E-35 |
| Neural Cells | Up | Cst3 | 1.40E-52 | 0.34478829 | 0.936 | 0.096 | 2.79E-49 |
| Neural Cells | Up | Tuba1b | 3.27E-18 | 0.34372058 | 0.751 | 0.071 | 6.53E-15 |
| Neural Cells | Up | Gpx1 | 2.70E-62 | 0.33801524 | 0.934 | 0.054 | 5.40E-59 |
| Neural Cells | Up | AABR070251 | 7.64E-28 | 0.32670984 | 0.807 | 0.071 | 1.53E-24 |
| Neural Cells | Up | Mcam | 8.49E-08 | 0.31602459 | 0.967 | 0.515 | 0.00016976 |
| Neural Cells | Up | Dok6 | 4.77E-60 | 0.31475763 | 0.799 | 0.008 | 9.55E-57 |

| | | | | | | | |
|---|---|---|---|---|---|---|---|
| Neural Cells | Up | Spon1 | 2.48E-06 | 0.31434588 | 0.781 | 0.276 | 0.00496467 |
| Neural Cells | Up | Rab7b | 2.09E-15 | 0.31402131 | 0.758 | 0.113 | 4.18E-12 |
| Neural Cells | Up | Adcy5 | 2.14E-29 | 0.30875049 | 0.878 | 0.146 | 4.28E-26 |
| Neural Cells | Up | Neb | 7.74E-47 | 0.30638201 | 0.878 | 0.054 | 1.55E-43 |
| Neural Cells | Up | Trpm3 | 5.94E-10 | 0.30602059 | 0.791 | 0.23 | 1.19E-06 |
| Neural Cells | Up | Acacb | 1.19E-38 | 0.30568238 | 0.934 | 0.155 | 2.38E-35 |
| Neural Cells | Up | Pcdh19 | 1.06E-57 | 0.30418733 | 0.98 | 0.117 | 2.12E-54 |
| Neural Cells | Up | Crlf1 | 1.21E-14 | 0.29892401 | 0.715 | 0.063 | 2.43E-11 |
| Neural Cells | Up | Sntg2 | 7.51E-56 | 0.29695447 | 0.924 | 0.067 | 1.50E-52 |
| Neural Cells | Up | Kcnk5 | 2.09E-11 | 0.29388868 | 0.341 | 0.017 | 4.19E-08 |
| Neural Cells | Up | Ablim3 | 1.71E-09 | 0.29252076 | 0.74 | 0.159 | 3.43E-06 |
| Neural Cells | Up | Cdon | 4.69E-64 | 0.29020666 | 0.952 | 0.067 | 9.37E-61 |
| Neural Cells | Up | Atp5f1e | 6.64E-28 | 0.28930334 | 0.863 | 0.138 | 1.33E-24 |
| Neural Cells | Up | Magi2 | 2.46E-11 | 0.28590869 | 0.702 | 0.075 | 4.91E-08 |
| Neural Cells | Up | Egr1 | 5.96E-27 | 0.28391767 | 0.746 | 0.029 | 1.19E-23 |
| Neural Cells | Up | Ldhb | 2.04E-27 | 0.28083911 | 0.906 | 0.197 | 4.07E-24 |
| Neural Cells | Up | Sema5a | 2.40E-29 | 0.27988438 | 0.791 | 0.042 | 4.79E-26 |
| Neural Cells | Up | Sema3c | 1.26E-06 | 0.27971926 | 0.941 | 0.481 | 0.00251334 |
| Neural Cells | Up | Fhl1 | 3.29E-34 | 0.27894972 | 0.875 | 0.117 | 6.57E-31 |
| Neural Cells | Up | Snca | 1.04E-34 | 0.27793214 | 0.677 | 0.008 | 2.09E-31 |
| Neural Cells | Up | Slc12a2 | 1.29E-05 | 0.27777539 | 0.43 | 0.088 | 0.02572547 |
| Neural Cells | Up | Lilrb4 | 1.21E-08 | 0.27407018 | 0.382 | 0.038 | 2.42E-05 |
| Neural Cells | Up | Cpe | 1.57E-62 | 0.27376556 | 0.939 | 0.059 | 3.14E-59 |
| Neural Cells | Up | Igfbp5 | 7.05E-06 | 0.26972564 | 0.463 | 0.192 | 0.01409218 |
| Neural Cells | Up | Vegfc | 3.08E-20 | 0.26749805 | 0.776 | 0.088 | 6.16E-17 |
| Neural Cells | Up | Chn1 | 2.09E-26 | 0.26542974 | 0.868 | 0.155 | 4.17E-23 |
| Neural Cells | Up | Coro6 | 4.05E-08 | 0.25645459 | 0.382 | 0.025 | 8.10E-05 |
| Neural Cells | Up | Igfbp3 | 8.80E-52 | 0.25635461 | 0.908 | 0.067 | 1.76E-48 |
| Neural Cells | Up | Lmod1 | 2.80E-49 | 0.25614324 | 0.873 | 0.038 | 5.60E-46 |
| Neural Cells | Up | Palm2 | 8.39E-11 | 0.25544077 | 0.372 | 0.071 | 1.68E-07 |
| Neural Cells | Up | Slc22a23 | 3.85E-37 | 0.25047 | 0.87 | 0.092 | 7.70E-34 |
| Neural Cells | Down | Plcxd3 | 9.13E-11 | -0.256876 | 0.623 | 0.042 | 1.83E-07 |
| Neural Cells | Down | Dpyd | 2.60E-16 | -0.258777 | 0.86 | 0.213 | 5.20E-13 |
| Neural Cells | Down | Ripor2 | 2.87E-09 | -0.2621209 | 0.644 | 0.063 | 5.75E-06 |
| Neural Cells | Down | Kcnd3 | 2.29E-38 | -0.2623232 | 0.919 | 0.134 | 4.58E-35 |
| Neural Cells | Down | F13a1 | 1.60E-05 | -0.2675073 | 0.626 | 0.05 | 0.03194362 |
| Neural Cells | Down | Maob | 4.22E-06 | -0.2677059 | 0.417 | 0.059 | 0.00844025 |
| Neural Cells | Down | Hs6st2 | 7.55E-12 | -0.2727286 | 0.333 | 0.071 | 1.51E-08 |
| Neural Cells | Down | Sv2c | 6.08E-13 | -0.2833904 | 0.985 | 0.368 | 1.22E-09 |
| Neural Cells | Down | Trhde | 3.10E-06 | -0.2867885 | 0.939 | 0.427 | 0.00619712 |
| Neural Cells | Down | Col28a1 | 6.53E-13 | -0.2905966 | 0.679 | 0.046 | 1.31E-09 |

| | | | | | | | |
|---|---|---|---|---|---|---|---|
| Neural Cells | Down | Xkr4 | 7.18E-45 | -0.2913271 | 0.919 | 0.105 | 1.44E-41 |
| Neural Cells | Down | Mrc1 | 8.63E-19 | -0.2930133 | 0.761 | 0.075 | 1.73E-15 |
| Neural Cells | Down | Zfhx4 | 8.57E-06 | -0.3028071 | 0.476 | 0.192 | 0.01714541 |
| Neural Cells | Down | Kcnq1 | 9.50E-07 | -0.3129361 | 0.796 | 0.255 | 0.00190087 |
| Neural Cells | Down | Tdrd12 | 3.54E-06 | -0.3417645 | 0.761 | 0.234 | 0.0070868 |
| Lymphatic Endothelial Cells | Up | Klhl4 | 2.89E-23 | 0.58752119 | 0.968 | 0.35 | 5.79E-20 |
| Lymphatic Endothelial Cells | Up | Sema3a | 3.37E-07 | 0.51016108 | 0.989 | 0.687 | 0.00067346 |
| Lymphatic Endothelial Cells | Up | Mylk | 1.01E-21 | 0.41604833 | 0.846 | 0.218 | 2.03E-18 |
| Lymphatic Endothelial Cells | Up | Olfml2a | 2.14E-29 | 0.37360526 | 0.868 | 0.186 | 4.28E-26 |
| Lymphatic Endothelial Cells | Up | Pde1c | 1.33E-30 | 0.37215898 | 0.803 | 0.098 | 2.67E-27 |
| Lymphatic Endothelial Cells | Up | Adgrl3 | 9.55E-12 | 0.36630038 | 0.666 | 0.048 | 1.91E-08 |
| Lymphatic Endothelial Cells | Up | Lmo7 | 1.65E-28 | 0.36504201 | 0.968 | 0.302 | 3.30E-25 |
| Lymphatic Endothelial Cells | Up | Cryab | 4.99E-43 | 0.3633468 | 0.887 | 0.138 | 9.98E-40 |
| Lymphatic Endothelial Cells | Up | Ankh | 2.96E-09 | 0.360658 | 0.96 | 0.472 | 5.93E-06 |
| Lymphatic Endothelial Cells | Up | Bcar3 | 6.30E-16 | 0.35767535 | 0.642 | 0.048 | 1.26E-12 |
| Lymphatic Endothelial Cells | Up | Tbx1 | 7.08E-09 | 0.34391654 | 0.841 | 0.363 | 1.42E-05 |
| Lymphatic Endothelial Cells | Up | Ackr3 | 1.06E-09 | 0.33457852 | 0.792 | 0.265 | 2.13E-06 |
| Lymphatic Endothelial Cells | Up | AABR070546 | 4.96E-07 | 0.33194031 | 0.639 | 0.066 | 0.00099246 |
| Lymphatic Endothelial Cells | Up | Sox13 | 9.97E-11 | 0.31370162 | 0.836 | 0.313 | 1.99E-07 |
| Lymphatic Endothelial Cells | Up | Grin2b | 4.25E-40 | 0.31332148 | 0.962 | 0.233 | 8.50E-37 |
| Lymphatic Endothelial Cells | Up | Lyve1 | 2.16E-14 | 0.30313243 | 0.946 | 0.403 | 4.31E-11 |
| Lymphatic Endothelial Cells | Up | Asap3 | 1.40E-79 | 0.30218249 | 0.938 | 0.069 | 2.80E-76 |
| Lymphatic Endothelial Cells | Up | Shroom3 | 1.17E-19 | 0.28152885 | 0.763 | 0.111 | 2.35E-16 |
| Lymphatic Endothelial Cells | Up | Apba2 | 6.52E-17 | 0.27510054 | 0.925 | 0.353 | 1.30E-13 |
| Lymphatic Endothelial Cells | Up | Met | 1.85E-53 | 0.27292914 | 0.747 | 0.029 | 3.69E-50 |
| Lymphatic Endothelial Cells | Up | Ebf2 | 2.56E-49 | 0.27002803 | 0.914 | 0.141 | 5.12E-46 |
| Lymphatic Endothelial Cells | Up | Asb2 | 6.51E-16 | 0.25634858 | 0.358 | 0.066 | 1.30E-12 |
| Lymphatic Endothelial Cells | Up | Rgs5 | 2.90E-29 | 0.25485128 | 0.752 | 0.04 | 5.81E-26 |
| Lymphatic Endothelial Cells | Up | Tmem51 | 3.03E-30 | 0.25283774 | 0.914 | 0.228 | 6.05E-27 |
| Lymphatic Endothelial Cells | Up | Brca1 | 8.46E-41 | 0.25126694 | 0.79 | 0.032 | 1.69E-37 |
| Lymphatic Endothelial Cells | Down | Sv2c | 2.08E-47 | -0.2643139 | 0.825 | 0.08 | 4.17E-44 |
| Lymphatic Endothelial Cells | Down | Ptafr | 9.56E-09 | -0.2645017 | 0.412 | 0.069 | 1.91E-05 |
| Lymphatic Endothelial Cells | Down | Cobll1 | 6.55E-10 | -0.2651319 | 0.439 | 0.178 | 1.31E-06 |
| Lymphatic Endothelial Cells | Down | Slc16a2 | 1.49E-10 | -0.2746381 | 0.42 | 0.133 | 2.98E-07 |
| Lymphatic Endothelial Cells | Down | Prss12 | 1.77E-15 | -0.2767938 | 0.79 | 0.172 | 3.55E-12 |
| Lymphatic Endothelial Cells | Down | Col27a1 | 1.06E-74 | -0.2928163 | 0.951 | 0.09 | 2.11E-71 |
| Lymphatic Endothelial Cells | Down | Antxr1 | 1.57E-20 | -0.3113183 | 0.809 | 0.159 | 3.14E-17 |
| Lymphatic Endothelial Cells | Down | Adamts17 | 5.10E-18 | -0.3226598 | 0.356 | 0.093 | 1.02E-14 |
| Lymphatic Endothelial Cells | Down | Col13a1 | 1.55E-05 | -0.3450188 | 0.496 | 0.191 | 0.03095325 |
| Smooth Muscle Cells | Down | Cox4i1 | 1.45E-10 | -0.4347985 | 0.7 | 0.274 | 2.90E-07 |
| Smooth Muscle Cells | Down | Cox6a2 | 1.91E-05 | -0.3651652 | 0.611 | 0.23 | 0.03815222 |

| | | | | | | | |
|---|---|---|---|---|---|---|---|
| Smooth Muscle Cells | Down | Gpr176 | 1.09E-43 | -0.3539712 | 0.696 | 0.124 | 2.17E-40 |
| Smooth Muscle Cells | Down | Sorcs2 | 1.54E-21 | -0.31027 | 0.662 | 0.168 | 3.07E-18 |
| Smooth Muscle Cells | Down | Trpc6 | 4.39E-42 | -0.2898188 | 0.703 | 0.128 | 8.78E-39 |
| Smooth Muscle Cells | Down | Dmpk | 8.45E-18 | -0.2832556 | 0.698 | 0.22 | 1.69E-14 |
| Smooth Muscle Cells | Down | Sorcs1 | 3.12E-10 | -0.2777792 | 0.505 | 0.068 | 6.23E-07 |
| Smooth Muscle Cells | Down | LOC1025530 | 1.42E-59 | -0.2572762 | 0.776 | 0.156 | 2.84E-56 |
| Smooth Muscle Cells | Down | Atp5mc3 | 6.11E-17 | -0.2566503 | 0.655 | 0.178 | 1.22E-13 |
| Smooth Muscle Cells | Up | Ripor2 | 7.60E-53 | 0.25130952 | 0.693 | 0.088 | 1.52E-49 |
| Smooth Muscle Cells | Up | Abca8a | 1.09E-23 | 0.2547676 | 0.85 | 0.36 | 2.18E-20 |
| Smooth Muscle Cells | Up | Gpx3 | 4.92E-34 | 0.25681392 | 0.635 | 0.082 | 9.83E-31 |
| Smooth Muscle Cells | Up | Agap2 | 1.84E-06 | 0.26322884 | 0.703 | 0.344 | 0.00368189 |
| Smooth Muscle Cells | Up | Slc16a10 | 3.55E-40 | 0.26669383 | 0.625 | 0.066 | 7.11E-37 |
| Smooth Muscle Cells | Up | Pde8b | 5.26E-12 | 0.27348888 | 0.935 | 0.536 | 1.05E-08 |
| Smooth Muscle Cells | Up | Trpc4 | 9.59E-08 | 0.28669738 | 0.539 | 0.112 | 0.00019179 |
| Smooth Muscle Cells | Up | Ednra | 1.04E-06 | 0.29947814 | 0.92 | 0.644 | 0.00208885 |
| Smooth Muscle Cells | Up | Slfn4 | 1.79E-50 | 0.3037203 | 0.691 | 0.088 | 3.58E-47 |
| Smooth Muscle Cells | Up | Pde1a | 1.30E-21 | 0.30419342 | 0.737 | 0.256 | 2.60E-18 |
| Smooth Muscle Cells | Up | Pde4b | 2.17E-39 | 0.3067515 | 0.724 | 0.156 | 4.35E-36 |
| Smooth Muscle Cells | Up | Tgfb2 | 1.69E-41 | 0.3150155 | 0.746 | 0.174 | 3.38E-38 |
| Smooth Muscle Cells | Up | Mcam | 1.85E-21 | 0.32428351 | 0.892 | 0.426 | 3.69E-18 |
| Smooth Muscle Cells | Up | Itpkb | 1.23E-21 | 0.32726699 | 0.85 | 0.366 | 2.46E-18 |
| Smooth Muscle Cells | Up | Tspan5 | 2.71E-18 | 0.33018089 | 0.572 | 0.078 | 5.42E-15 |
| Smooth Muscle Cells | Up | Arhgap15 | 1.43E-18 | 0.33245521 | 0.904 | 0.454 | 2.87E-15 |
| Smooth Muscle Cells | Up | Angpt1 | 8.71E-48 | 0.33921016 | 0.807 | 0.21 | 1.74E-44 |
| Smooth Muscle Cells | Up | P2ry14 | 1.48E-36 | 0.34066723 | 0.882 | 0.31 | 2.96E-33 |
| Smooth Muscle Cells | Up | Pdgfra | 2.32E-76 | 0.34500146 | 0.761 | 0.106 | 4.65E-73 |
| Smooth Muscle Cells | Up | Akap6 | 1.58E-11 | 0.3529056 | 0.904 | 0.556 | 3.15E-08 |
| Smooth Muscle Cells | Up | Ldhb | 3.96E-21 | 0.35400953 | 0.744 | 0.26 | 7.93E-18 |
| Smooth Muscle Cells | Up | Klhl29 | 6.26E-43 | 0.35420599 | 0.659 | 0.084 | 1.25E-39 |
| Smooth Muscle Cells | Up | Trpc3 | 1.47E-17 | 0.3551426 | 0.865 | 0.412 | 2.94E-14 |
| Smooth Muscle Cells | Up | Ano4 | 5.54E-28 | 0.36430943 | 0.749 | 0.238 | 1.11E-24 |
| Smooth Muscle Cells | Up | Kcnj8 | 2.82E-16 | 0.36492481 | 0.846 | 0.406 | 5.64E-13 |
| Smooth Muscle Cells | Up | Art3 | 2.66E-17 | 0.37068803 | 0.935 | 0.514 | 5.31E-14 |

| | | | | | | | |
|---|---|---|---|---|---|---|---|
| Smooth Muscle Cells | Up | Ntng1 | 2.71E-16 | 0.37107884 | 0.884 | 0.442 | 5.43E-13 |
| Smooth Muscle Cells | Up | Chn1 | 1.30E-11 | 0.37319659 | 0.93 | 0.58 | 2.60E-08 |
| Smooth Muscle Cells | Up | Col25a1 | 2.93E-57 | 0.37408623 | 0.712 | 0.088 | 5.85E-54 |
| Smooth Muscle Cells | Up | AABR070275 | 7.78E-10 | 0.37418011 | 0.72 | 0.336 | 1.56E-06 |
| Smooth Muscle Cells | Up | Col4a5 | 1.65E-39 | 0.37515448 | 0.811 | 0.252 | 3.29E-36 |
| Smooth Muscle Cells | Up | Egr1 | 2.53E-13 | 0.38079908 | 0.531 | 0.056 | 5.07E-10 |
| Smooth Muscle Cells | Up | Kank4 | 1.38E-13 | 0.38269365 | 0.65 | 0.196 | 2.77E-10 |
| Smooth Muscle Cells | Up | Gli2 | 2.36E-21 | 0.38516031 | 0.896 | 0.434 | 4.72E-18 |
| Smooth Muscle Cells | Up | Mamdc2 | 2.66E-30 | 0.39273803 | 0.867 | 0.346 | 5.32E-27 |
| Smooth Muscle Cells | Up | AABR070592 | 3.09E-43 | 0.3939244 | 0.724 | 0.148 | 6.18E-40 |
| Smooth Muscle Cells | Up | C7 | 2.86E-20 | 0.39809924 | 0.858 | 0.422 | 5.71E-17 |
| Smooth Muscle Cells | Up | Med12l | 2.60E-32 | 0.41392567 | 0.846 | 0.316 | 5.19E-29 |
| Smooth Muscle Cells | Up | Dpysl3 | 1.07E-71 | 0.42277668 | 0.795 | 0.136 | 2.15E-68 |
| Smooth Muscle Cells | Up | Il1r1 | 3.22E-16 | 0.42400355 | 0.915 | 0.512 | 6.44E-13 |
| Smooth Muscle Cells | Up | Agtr1a | 4.52E-51 | 0.43221724 | 0.811 | 0.202 | 9.03E-48 |
| Smooth Muscle Cells | Up | Pld5 | 6.27E-17 | 0.4622036 | 0.739 | 0.266 | 1.25E-13 |
| Smooth Muscle Cells | Up | Calcrl | 1.16E-16 | 0.50669121 | 0.879 | 0.474 | 2.32E-13 |
| Smooth Muscle Cells | Up | AABR070067 | 2.20E-38 | 0.52820795 | 0.85 | 0.29 | 4.41E-35 |
