## Supplemental Table 8 for "Uncovering the Regional and Cell Specific Bioactivity of Injectable Extracellular Matrix Biomaterials in Myocardial Infarction through Spatial and Single Nucleus Transcriptomics"

| Supplementary Table 8. Differentially Expressed Genes Spatially Comparing ECM Hydrogel in the Infarct and Infarct Only Zones in the Chronic MI Model | | | | | | | |
|---|---|---|---|---|---|---|---|
| **Spatial Area** | **Direction** | **Gene** | **p_val** | **avg_log2FC** | **pct.1** | **pct.2** | **p_val_adj** |
| ECM Hydrogel in Infarct | Up | Postn | 1.29E-38 | 2.31577254 | 0.934 | 0.584 | 1.60E-34 |
| ECM Hydrogel in Infarct | Up | Tagln | 1.61E-43 | 2.04025646 | 0.885 | 0.29 | 1.99E-39 |
| ECM Hydrogel in Infarct | Up | Fn1 | 1.29E-44 | 1.85421355 | 0.984 | 0.725 | 1.60E-40 |
| ECM Hydrogel in Infarct | Up | Ccn3 | 2.48E-45 | 1.77081855 | 0.82 | 0.183 | 3.07E-41 |
| ECM Hydrogel in Infarct | Up | Ccn2 | 2.11E-37 | 1.69814858 | 0.907 | 0.473 | 2.61E-33 |
| ECM Hydrogel in Infarct | Up | Actg2 | 4.68E-32 | 1.48273289 | 0.628 | 0.111 | 5.79E-28 |
| ECM Hydrogel in Infarct | Up | Col8a1 | 1.57E-36 | 1.45359283 | 0.689 | 0.126 | 1.94E-32 |
| ECM Hydrogel in Infarct | Up | Tpm2 | 2.42E-34 | 1.41266256 | 0.923 | 0.443 | 2.99E-30 |
| ECM Hydrogel in Infarct | Up | Myh11 | 8.60E-15 | 1.34756753 | 0.536 | 0.183 | 1.06E-10 |
| ECM Hydrogel in Infarct | Up | Acta2 | 1.13E-23 | 1.25312824 | 0.667 | 0.214 | 1.40E-19 |
| ECM Hydrogel in Infarct | Up | Tnc | 1.96E-29 | 1.17470366 | 0.437 | 0.011 | 2.42E-25 |
| ECM Hydrogel in Infarct | Up | Ltbp2 | 2.48E-28 | 1.13959383 | 0.88 | 0.489 | 3.06E-24 |
| ECM Hydrogel in Infarct | Up | Myl9 | 1.05E-17 | 1.11372394 | 0.803 | 0.443 | 1.29E-13 |
| ECM Hydrogel in Infarct | Up | Aspn | 2.53E-18 | 0.98856245 | 0.585 | 0.206 | 3.12E-14 |
| ECM Hydrogel in Infarct | Up | Flna | 2.36E-21 | 0.97517776 | 0.934 | 0.656 | 2.91E-17 |
| ECM Hydrogel in Infarct | Up | Lox | 5.03E-24 | 0.96920266 | 0.76 | 0.317 | 6.22E-20 |
| ECM Hydrogel in Infarct | Up | Cthrc1 | 4.13E-27 | 0.93048332 | 0.607 | 0.122 | 5.10E-23 |
| ECM Hydrogel in Infarct | Up | Fndc1 | 1.48E-16 | 0.91206372 | 0.749 | 0.424 | 1.83E-12 |
| ECM Hydrogel in Infarct | Up | Serpine1 | 1.98E-17 | 0.88902035 | 0.404 | 0.076 | 2.45E-13 |
| ECM Hydrogel in Infarct | Up | Col12a1 | 3.35E-21 | 0.88779677 | 0.404 | 0.05 | 4.14E-17 |
| ECM Hydrogel in Infarct | Up | Actn1 | 1.41E-17 | 0.8517494 | 0.634 | 0.252 | 1.74E-13 |
| ECM Hydrogel in Infarct | Up | Csrp1 | 1.17E-10 | 0.82158319 | 0.82 | 0.553 | 1.45E-06 |
| ECM Hydrogel in Infarct | Up | Actb | 1.22E-17 | 0.81134589 | 0.563 | 0.179 | 1.50E-13 |
| ECM Hydrogel in Infarct | Up | Cald1 | 4.86E-15 | 0.8095804 | 0.678 | 0.34 | 6.01E-11 |
| ECM Hydrogel in Infarct | Up | Dkk3 | 4.00E-17 | 0.79625866 | 0.809 | 0.401 | 4.95E-13 |
| ECM Hydrogel in Infarct | Up | Col18a1 | 9.58E-11 | 0.78181876 | 0.607 | 0.305 | 1.18E-06 |
| ECM Hydrogel in Infarct | Up | NEWGENE-6: | 4.16E-19 | 0.77664146 | 0.617 | 0.198 | 5.13E-15 |
| ECM Hydrogel in Infarct | Up | Actn4 | 1.78E-14 | 0.75466834 | 0.71 | 0.347 | 2.20E-10 |
| ECM Hydrogel in Infarct | Up | Csrp2 | 9.39E-16 | 0.75463851 | 0.563 | 0.214 | 1.16E-11 |
| ECM Hydrogel in Infarct | Up | Fstl1 | 7.97E-15 | 0.74545483 | 0.825 | 0.531 | 9.85E-11 |
| Infarct Only | Up | Coq8a | 1.92E-05 | -0.2540664 | 0.563 | 0.821 | 0.23746784 |
| Infarct Only | Up | Acadvl | 9.75E-07 | -0.2582975 | 0.65 | 0.905 | 0.01204341 |
| Infarct Only | Up | Rrad | 2.52E-05 | -0.2643945 | 0.568 | 0.821 | 0.3107833 |
| Infarct Only | Up | Mrps36 | 5.36E-07 | -0.2662573 | 0.514 | 0.794 | 0.00662205 |
| Infarct Only | Up | Etfa | 1.68E-06 | -0.2721913 | 0.53 | 0.802 | 0.0207745 |

| | | | | | | | |
|---|---|---|---|---|---|---|---|
| Infarct Only | Up | Scn1b | 9.33E-07 | -0.2763349 | 0.464 | 0.729 | 0.01153394 |
| Infarct Only | Up | LOC1003624 | 4.63E-06 | -0.2785977 | 0.377 | 0.641 | 0.05727463 |
| Infarct Only | Up | Apobec2 | 2.75E-07 | -0.2951069 | 0.273 | 0.553 | 0.00340296 |
| Infarct Only | Up | Aldh6a1 | 4.50E-07 | -0.2982655 | 0.186 | 0.439 | 0.00555539 |
| Infarct Only | Up | Nrap | 1.78E-07 | -0.3008728 | 0.579 | 0.847 | 0.00220193 |
| Infarct Only | Up | Hadh | 1.16E-06 | -0.3017046 | 0.525 | 0.786 | 0.01431099 |
| Infarct Only | Up | Dsp | 5.13E-07 | -0.3052084 | 0.53 | 0.782 | 0.00634364 |
| Infarct Only | Up | Txlnb | 3.21E-08 | -0.3059541 | 0.137 | 0.401 | 0.00039714 |
| Infarct Only | Up | Pdhb | 4.11E-08 | -0.3195473 | 0.503 | 0.771 | 0.00050821 |
| Infarct Only | Up | Cpt1b | 3.97E-08 | -0.3338657 | 0.552 | 0.813 | 0.00049092 |
| Infarct Only | Up | Macrod1 | 2.19E-07 | -0.3438628 | 0.486 | 0.748 | 0.00271181 |
| Infarct Only | Up | Ndufc2 | 2.51E-08 | -0.3571421 | 0.557 | 0.813 | 0.00031035 |
| Infarct Only | Up | Acads | 5.64E-07 | -0.3708882 | 0.47 | 0.725 | 0.00696603 |
| Infarct Only | Up | Vegfa | 1.95E-08 | -0.3841774 | 0.224 | 0.496 | 0.00024078 |
| Infarct Only | Up | Akr1c15 | 2.26E-11 | -0.4384817 | 0.41 | 0.779 | 2.80E-07 |
| Infarct Only | Up | Igfbp3 | 4.10E-11 | -0.7204366 | 0.164 | 0.447 | 5.06E-07 |
| Infarct Only | Up | Pla2g2a | 1.09E-13 | -0.9655626 | 0.29 | 0.595 | 1.35E-09 |
| Infarct Only | Up | Ces1d | 1.64E-20 | -1.0415437 | 0.355 | 0.737 | 2.02E-16 |
