## Supplemental Table 10 for "Uncovering the Regional and Cell Specific Bioactivity of Injectable Extracellular Matrix Biomaterials in Myocardial Infarction through Spatial and Single Nucleus Transcriptomics"

| Supplementary Table 10. ECM Hydrogel Treated Chronic Infarcts without Visible ECM Compared to Saline Chronic Infarcts | | | | | | | |
|---|---|---|---|---|---|---|---|
| **Spatial Area** | **Direction** | **Gene** | **p_val** | **avg_log2FC** | **pct.1** | **pct.2** | **p_val_adj** |
| ECM Hydrogel Treatment with No Visible ECM | Up | Gnas.1 | 1.47E-48 | 0.78926188 | 0.258 | 0.05 | 3.71E-44 |
| ECM Hydrogel Treatment with No Visible ECM | Up | Eef1a1 | 3.53E-48 | 1.12573749 | 0.524 | 0.265 | 8.93E-44 |
| ECM Hydrogel Treatment with No Visible ECM | Up | Postn | 3.95E-35 | 0.64685311 | 0.741 | 0.536 | 9.99E-31 |
| ECM Hydrogel Treatment with No Visible ECM | Up | LOC1003620 | 2.64E-34 | 0.71136583 | 0.633 | 0.406 | 6.66E-30 |
| Saline Infarct | Up | AABR070718 | 3.11E-33 | -0.7612152 | 0.331 | 0.54 | 7.86E-29 |
| Saline Infarct | Up | RT1-A2 | 4.02E-26 | 0.75387772 | 0.451 | 0.251 | 1.02E-21 |
