## Supplemental Table 9 for "Uncovering the Regional and Cell Specific Bioactivity of Injectable Extracellular Matrix Biomaterials in Myocardial Infarction through Spatial and Single Nucleus Transcriptomics"

| Supplementary Table 9. Global Spatial Comparisons of Infarcts Treated with ECM Hydrogel or Saline in the Chronic MI Model | | | | | | | |
|---|---|---|---|---|---|---|---|
| **Sample** | **Direction** | **Gene** | **p_val** | **avg_log2FC** | **pct.1** | **pct.2** | **p_val_adj** |
| ECM Hydrogel | Up | Postn | 1.49E-114 | 1.15154864 | 0.85 | 0.593 | 3.78E-110 |
| ECM Hydrogel | Up | Tnn | 4.30E-114 | 1.42340489 | 0.287 | 0.012 | 1.09E-109 |
| ECM Hydrogel | Up | Gnas.1 | 1.15E-107 | 0.92481331 | 0.375 | 0.06 | 2.91E-103 |
| ECM Hydrogel | Up | Eef1a1 | 1.63E-103 | 1.17092996 | 0.62 | 0.28 | 4.12E-99 |
| ECM Hydrogel | Up | Col5a2 | 2.74E-99 | 0.96553229 | 0.345 | 0.057 | 6.91E-95 |
| ECM Hydrogel | Up | Ccn3 | 7.83E-97 | 1.58024285 | 0.55 | 0.224 | 1.98E-92 |
| ECM Hydrogel | Up | Ccn1 | 1.57E-93 | 1.04843659 | 0.355 | 0.07 | 3.96E-89 |
| ECM Hydrogel | Up | Cthrc1 | 7.76E-93 | 1.21973829 | 0.507 | 0.186 | 1.96E-88 |
| ECM Hydrogel | Up | Plod2 | 2.02E-92 | 1.22635564 | 0.501 | 0.192 | 5.11E-88 |
| ECM Hydrogel | Up | NEWGENE-621351.1 | 8.84E-91 | 1.46579452 | 0.614 | 0.355 | 2.23E-86 |
| ECM Hydrogel | Up | Mmp12 | 8.59E-87 | 1.13680762 | 0.254 | 0.021 | 2.17E-82 |
| ECM Hydrogel | Up | Col12a1 | 4.23E-83 | 1.15003477 | 0.346 | 0.08 | 1.07E-78 |
| ECM Hydrogel | Up | Ppial4d | 2.39E-82 | 0.73671591 | 0.347 | 0.073 | 6.03E-78 |
| ECM Hydrogel | Up | Fndc1 | 4.38E-75 | 0.95101727 | 0.706 | 0.443 | 1.11E-70 |
| ECM Hydrogel | Up | Col11a1 | 3.88E-71 | 0.90793681 | 0.259 | 0.042 | 9.80E-67 |
| ECM Hydrogel | Up | Tnc | 3.93E-70 | 1.06987232 | 0.321 | 0.08 | 9.94E-66 |
| ECM Hydrogel | Up | C3 | 6.13E-67 | 1.01049965 | 0.287 | 0.062 | 1.55E-62 |
| ECM Hydrogel | Up | Lox | 3.77E-66 | 0.87047974 | 0.639 | 0.367 | 9.53E-62 |
| ECM Hydrogel | Up | LOC498555 | 5.74E-66 | 0.75511404 | 0.52 | 0.23 | 1.45E-61 |
| ECM Hydrogel | Up | LOC100362027 | 6.58E-66 | 0.72005925 | 0.708 | 0.421 | 1.66E-61 |
| ECM Hydrogel | Up | Tnfrsf11b | 3.58E-63 | 0.55488992 | 0.237 | 0.036 | 9.04E-59 |
| ECM Hydrogel | Up | Nid1 | 3.78E-63 | 0.59143023 | 0.309 | 0.077 | 9.56E-59 |
| ECM Hydrogel | Up | F2r | 1.06E-59 | 0.6571994 | 0.328 | 0.097 | 2.67E-55 |
| ECM Hydrogel | Up | Pabpc1 | 1.18E-59 | 0.75771483 | 0.722 | 0.475 | 2.98E-55 |
| ECM Hydrogel | Up | Myadm | 1.28E-59 | 0.63405635 | 0.375 | 0.125 | 3.23E-55 |
| ECM Hydrogel | Up | Cebpb | 1.67E-58 | 0.52154479 | 0.295 | 0.073 | 4.23E-54 |
| ECM Hydrogel | Up | Hmox1 | 1.84E-58 | 1.03893202 | 0.344 | 0.109 | 4.65E-54 |
| ECM Hydrogel | Up | Serpine1 | 6.24E-56 | 0.92232654 | 0.421 | 0.177 | 1.58E-51 |
| ECM Hydrogel | Up | Ppia | 5.66E-55 | 0.62412625 | 0.457 | 0.196 | 1.43E-50 |
| ECM Hydrogel | Up | Cd200 | 2.14E-53 | 0.6045675 | 0.41 | 0.16 | 5.42E-49 |

| | | | | | | | |
|---|---|---|---|---|---|---|---|
| ECM Hydrogel | Up | Arf4 | 5.56E-53 | 0.72878358 | 0.594 | 0.362 | 1.41E-48 |
| ECM Hydrogel | Up | Fat1 | 2.69E-51 | 0.66940429 | 0.446 | 0.201 | 6.81E-47 |
| ECM Hydrogel | Up | Pdia3 | 9.80E-50 | 0.59608461 | 0.731 | 0.51 | 2.48E-45 |
| ECM Hydrogel | Up | Cemip | 1.08E-49 | 0.63533836 | 0.306 | 0.097 | 2.72E-45 |
| ECM Hydrogel | Up | Cdh11 | 2.73E-49 | 0.54406254 | 0.356 | 0.13 | 6.90E-45 |
| ECM Hydrogel | Up | LOC100359951 | 3.95E-49 | 0.61510799 | 0.547 | 0.292 | 9.99E-45 |
| ECM Hydrogel | Up | RGD1559482 | 4.01E-49 | 0.70793939 | 0.542 | 0.274 | 1.01E-44 |
| ECM Hydrogel | Up | Aspn | 7.40E-49 | 0.82547276 | 0.581 | 0.353 | 1.87E-44 |
| ECM Hydrogel | Up | Lgmn | 2.73E-48 | 0.63389809 | 0.652 | 0.409 | 6.90E-44 |
| ECM Hydrogel | Up | AABR07053749.2 | 3.19E-48 | 0.56812332 | 0.464 | 0.215 | 8.05E-44 |
| ECM Hydrogel | Up | Lgals3 | 6.49E-48 | 0.6851214 | 0.77 | 0.565 | 1.64E-43 |
| ECM Hydrogel | Up | Txndc5 | 9.43E-48 | 0.66314082 | 0.563 | 0.327 | 2.38E-43 |
| ECM Hydrogel | Up | Ppic | 3.60E-47 | 0.63022576 | 0.489 | 0.253 | 9.11E-43 |
| ECM Hydrogel | Up | Col8a1 | 2.25E-46 | 0.9639717 | 0.545 | 0.337 | 5.69E-42 |
| ECM Hydrogel | Up | Ost4 | 2.26E-46 | 0.5917898 | 0.585 | 0.337 | 5.71E-42 |
| ECM Hydrogel | Up | RT1-A2 | 3.55E-46 | 0.67531427 | 0.534 | 0.283 | 8.97E-42 |
| ECM Hydrogel | Up | Vmp1 | 9.95E-44 | 0.5743812 | 0.499 | 0.258 | 2.51E-39 |
| ECM Hydrogel | Up | Acta2 | 1.54E-43 | 0.88867289 | 0.486 | 0.262 | 3.90E-39 |
| ECM Hydrogel | Up | Txn1 | 1.85E-43 | 0.55337377 | 0.513 | 0.269 | 4.67E-39 |
| ECM Hydrogel | Up | Spp1 | 3.26E-43 | 1.08345507 | 0.523 | 0.303 | 8.25E-39 |
| ECM Hydrogel | Up | RT1-Da | 1.13E-41 | 0.61008225 | 0.76 | 0.527 | 2.85E-37 |
| ECM Hydrogel | Up | Mfap4 | 1.33E-41 | 0.57753008 | 0.423 | 0.202 | 3.35E-37 |
| ECM Hydrogel | Up | Rab31 | 6.60E-40 | 0.5442153 | 0.563 | 0.333 | 1.67E-35 |
| ECM Hydrogel | Up | Dusp1 | 1.00E-39 | 0.48007777 | 0.425 | 0.197 | 2.53E-35 |
| ECM Hydrogel | Up | Cmpk1 | 1.59E-39 | 0.37207448 | 0.342 | 0.134 | 4.02E-35 |
| ECM Hydrogel | Up | Igf2r | 2.79E-39 | 0.49106851 | 0.5 | 0.266 | 7.06E-35 |
| ECM Hydrogel | Up | Anpep | 1.35E-38 | 0.42374366 | 0.368 | 0.157 | 3.42E-34 |
| ECM Hydrogel | Up | LOC103693375 | 3.02E-38 | 0.47680558 | 0.725 | 0.498 | 7.64E-34 |
| ECM Hydrogel | Up | Gpnmb | 4.64E-38 | 0.61622799 | 0.675 | 0.443 | 1.17E-33 |
| ECM Hydrogel | Up | Sdcbp | 5.21E-38 | 0.45016579 | 0.546 | 0.302 | 1.32E-33 |
| ECM Hydrogel | Up | Ext1 | 7.45E-38 | 0.5039637 | 0.493 | 0.268 | 1.88E-33 |
| ECM Hydrogel | Up | Arpc2 | 1.68E-37 | 0.52318202 | 0.652 | 0.443 | 4.25E-33 |
| ECM Hydrogel | Up | Ran | 1.84E-37 | 0.50637252 | 0.568 | 0.337 | 4.66E-33 |

| | | | | | | | |
|---|---|---|---|---|---|---|---|
| ECM Hydrogel | Up | Aprt | 6.02E-37 | 0.40426183 | 0.412 | 0.19 | 1.52E-32 |
| ECM Hydrogel | Up | AC134224.1 | 1.22E-36 | 0.78910363 | 0.455 | 0.234 | 3.10E-32 |
| ECM Hydrogel | Up | Ptgfrn | 2.02E-36 | 0.43816 | 0.466 | 0.238 | 5.09E-32 |
| ECM Hydrogel | Up | Fzd1 | 6.07E-36 | 0.41257844 | 0.408 | 0.19 | 1.53E-31 |
| ECM Hydrogel | Up | Scd2 | 6.59E-36 | 0.44824178 | 0.496 | 0.268 | 1.66E-31 |
| ECM Hydrogel | Up | Spag9 | 7.99E-36 | 0.368965 | 0.346 | 0.145 | 2.02E-31 |
| ECM Hydrogel | Up | Pdgfrl | 1.27E-35 | 0.47602666 | 0.412 | 0.203 | 3.20E-31 |
| ECM Hydrogel | Up | Jun | 5.01E-35 | 0.45330345 | 0.508 | 0.28 | 1.27E-30 |
| ECM Hydrogel | Up | Tmed2 | 1.31E-34 | 0.45274467 | 0.659 | 0.449 | 3.30E-30 |
| ECM Hydrogel | Up | Rrbp1 | 1.76E-34 | 0.476073 | 0.386 | 0.179 | 4.45E-30 |
| ECM Hydrogel | Up | Nucb1 | 2.48E-34 | 0.46640609 | 0.67 | 0.459 | 6.27E-30 |
| ECM Hydrogel | Up | Cr1l | 3.27E-34 | 0.34696674 | 0.365 | 0.161 | 8.25E-30 |
| ECM Hydrogel | Up | Actr3 | 4.20E-34 | 0.50722503 | 0.571 | 0.356 | 1.06E-29 |
| ECM Hydrogel | Up | AABR07065438.1 | 4.89E-34 | 0.44077808 | 0.482 | 0.255 | 1.24E-29 |
| ECM Hydrogel | Up | LOC100911372 | 7.58E-34 | 0.5014712 | 0.614 | 0.414 | 1.92E-29 |
| ECM Hydrogel | Up | Etfa | 2.12E-33 | 0.50338994 | 0.449 | 0.237 | 5.35E-29 |
| ECM Hydrogel | Up | Sod2 | 2.68E-33 | 0.45385538 | 0.6 | 0.363 | 6.78E-29 |
| ECM Hydrogel | Up | Vldlr | 4.43E-33 | 0.46566811 | 0.446 | 0.237 | 1.12E-28 |
| ECM Hydrogel | Up | Smpdl3a | 7.83E-33 | 0.43661548 | 0.493 | 0.269 | 1.98E-28 |
| ECM Hydrogel | Up | Rab14 | 8.21E-33 | 0.35383473 | 0.442 | 0.218 | 2.08E-28 |
| ECM Hydrogel | Up | Npm1 | 8.82E-33 | 0.47251957 | 0.726 | 0.502 | 2.23E-28 |
| ECM Hydrogel | Up | Myh10 | 1.85E-32 | 0.4686622 | 0.476 | 0.263 | 4.67E-28 |
| ECM Hydrogel | Up | Olfml2b | 2.97E-32 | 0.43381035 | 0.422 | 0.218 | 7.51E-28 |
| ECM Hydrogel | Up | Ckap4 | 4.17E-32 | 0.50671493 | 0.466 | 0.263 | 1.05E-27 |
| ECM Hydrogel | Up | Pdcd6ip | 6.82E-32 | 0.35324349 | 0.386 | 0.182 | 1.72E-27 |
| ECM Hydrogel | Up | Wdr26 | 2.32E-31 | 0.36814505 | 0.418 | 0.208 | 5.87E-27 |
| ECM Hydrogel | Up | Ybx1 | 7.29E-31 | 0.48515177 | 0.56 | 0.33 | 1.84E-26 |
| ECM Hydrogel | Up | Mrc1 | 5.78E-30 | 0.4568043 | 0.52 | 0.302 | 1.46E-25 |
| ECM Hydrogel | Up | Ssr3 | 8.83E-30 | 0.39133787 | 0.453 | 0.247 | 2.23E-25 |
| ECM Hydrogel | Up | Hnrnpf | 9.11E-30 | 0.43546534 | 0.504 | 0.297 | 2.30E-25 |
| ECM Hydrogel | Up | Vcan | 1.02E-29 | 0.39439917 | 0.5 | 0.293 | 2.57E-25 |
| ECM Hydrogel | Up | Ube2d3 | 2.13E-29 | 0.37800481 | 0.55 | 0.329 | 5.39E-25 |
| ECM Hydrogel | Up | Slfn2 | 2.30E-29 | 0.43524452 | 0.482 | 0.27 | 5.81E-25 |

| | | | | | | | |
|---|---|---|---|---|---|---|---|
| ECM Hydrogel | Up | Rexo2 | 4.71E-29 | 0.31997147 | 0.443 | 0.233 | 1.19E-24 |
| ECM Hydrogel | Up | Tp53inp2 | 8.41E-29 | 0.33458742 | 0.429 | 0.226 | 2.13E-24 |
| ECM Hydrogel | Up | Ybx1-ps3 | 9.81E-29 | 0.34727714 | 0.564 | 0.33 | 2.48E-24 |
| ECM Hydrogel | Up | Sec61b | 1.28E-28 | 0.42343282 | 0.598 | 0.394 | 3.23E-24 |
| ECM Hydrogel | Up | RGD1310352 | 1.32E-28 | 0.33038084 | 0.419 | 0.218 | 3.34E-24 |
| ECM Hydrogel | Up | Mettl9 | 1.59E-27 | 0.28820025 | 0.432 | 0.226 | 4.01E-23 |
| ECM Hydrogel | Up | Hist1h2bq | 2.31E-27 | 0.32144286 | 0.464 | 0.258 | 5.85E-23 |
| ECM Hydrogel | Up | Cd68 | 3.50E-27 | 0.39975506 | 0.5 | 0.289 | 8.84E-23 |
| ECM Hydrogel | Up | Spcs2 | 4.94E-25 | 0.27961107 | 0.437 | 0.235 | 1.25E-20 |
| ECM Hydrogel | Up | Pnrc2 | 3.17E-24 | 0.32147309 | 0.482 | 0.281 | 8.02E-20 |
| ECM Hydrogel | Up | Ndufb4 | 5.25E-24 | 0.32970009 | 0.55 | 0.34 | 1.33E-19 |
| ECM Hydrogel | Up | Capza2 | 1.05E-23 | 0.33483297 | 0.522 | 0.318 | 2.66E-19 |
| ECM Hydrogel | Up | Lcp1 | 2.36E-23 | 0.34667304 | 0.521 | 0.319 | 5.96E-19 |
