## Supplemental Table 11 for "Uncovering the Regional and Cell Specific Bioactivity of Injectable Extracellular Matrix Biomaterials in Myocardial Infarction through Spatial and Single Nucleus Transcriptomics"

**Supplementary Table 11. Differentially Expressed Genes Spatially Comparing the Remote Zone of ECM Treated Hearts vs. Saline Treated Hearts in a Chronic MI Model**

| Spatial Area | Direction | Gene | p_val | avg_log2FC | pct.1 | pct.2 | p_val_adj |
|---|---|---|---|---|---|---|---|
| Saline Remote Zone | Up | Wdr78 | 7.41E-230 | -2.3961177 | 0.332 | 0.627 | 1.48E-226 |
| Saline Remote Zone | Up | Fam151a | 0 | -2.2294741 | 0.423 | 0.652 | 0 |
| Saline Remote Zone | Up | Lekr1 | 0 | -1.5426464 | 0.318 | 0.587 | 0 |
| Saline Remote Zone | Up | Fcer1a | 9.53E-255 | -1.2504006 | 0.501 | 0.924 | 1.91E-251 |
| Saline Remote Zone | Up | Sbspon | 0 | -1.1461542 | 0.39 | 0.814 | 0 |
| Saline Remote Zone | Up | Tlr12 | 0 | -1.1452929 | 0.449 | 0.815 | 0 |
| Saline Remote Zone | Up | Syt8 | 0 | -1.1452409 | 0.249 | 0.475 | 0 |
| Saline Remote Zone | Up | RGD1560242 | 0 | -1.1268344 | 0.49 | 0.856 | 0 |

| | | | | | | | |
|---|---|---|---|---|---|---|---|
| Saline Remote Zone | Up | Camk4 | 6.32E-260 | -1.1056972 | 0.449 | 0.929 | 1.26E-256 |
| Saline Remote Zone | Up | Clic3 | 5.18E-111 | -1.0711663 | 0.414 | 0.749 | 1.04E-107 |
| Saline Remote Zone | Up | Slc38a4 | 0 | -1.0679449 | 0.325 | 0.661 | 0 |
| Saline Remote Zone | Up | Rassf7 | 0 | -0.9976199 | 0.191 | 0.619 | 0 |
| Saline Remote Zone | Up | Dnaaf4 | 0 | -0.9874277 | 0.485 | 0.916 | 0 |
| Saline Remote Zone | Up | Nufip2 | 0 | -0.966748 | 0.212 | 0.555 | 0 |
| Saline Remote Zone | Up | Cenpt | 0 | -0.9635975 | 0.177 | 0.42 | 0 |
| Saline Remote Zone | Up | Slc2a6 | 0 | -0.9541994 | 0.35 | 0.573 | 0 |
| Saline Remote Zone | Up | AABR07034739.1 | 0 | -0.9430694 | 0.308 | 0.673 | 0 |

| | | | | | | | |
|---|---|---|---|---|---|---|---|
| Saline Remote Zone | Up | Prss46 | 0 | -0.9416955 | 0.376 | 0.684 | 0 |
| Saline Remote Zone | Up | Il18rap | 2.14E-242 | -0.8982683 | 0.501 | 0.898 | 4.28E-239 |
| Saline Remote Zone | Up | Tmem63c | 2.02E-230 | -0.8774108 | 0.401 | 0.633 | 4.04E-227 |
| Saline Remote Zone | Up | Fjx1 | 0 | -0.8703498 | 0.316 | 0.579 | 0 |
| Saline Remote Zone | Up | Zmynd15 | 0 | -0.8628397 | 0.254 | 0.519 | 0 |
| Saline Remote Zone | Up | Dpf1 | 0 | -0.8579761 | 0.449 | 0.83 | 0 |
| Saline Remote Zone | Up | Egr2 | 0 | -0.8448438 | 0.325 | 0.626 | 0 |
| Saline Remote Zone | Up | Cdhr1 | 0 | -0.8391083 | 0.26 | 0.483 | 0 |
| Saline Remote Zone | Up | Mcpt8l3 | 5.43E-226 | -0.8203359 | 0.465 | 0.703 | 1.09E-222 |

| | | | | | | | |
|---|---|---|---|---|---|---|---|
| Saline Remote Zone | Up | Habp2 | 0 | -0.8189342 | 0.277 | 0.563 | 0 |
| Saline Remote Zone | Up | Pmfbp1 | 4.49E-264 | -0.8155266 | 0.458 | 0.794 | 8.99E-261 |
| Saline Remote Zone | Up | LOC24906 | 0 | -0.8063722 | 0.456 | 0.797 | 0 |
| Saline Remote Zone | Up | Sall1 | 0 | -0.7956342 | 0.243 | 0.453 | 0 |
| Saline Remote Zone | Up | Clec2dl1 | 0 | -0.794534 | 0.347 | 0.64 | 0 |
| Saline Remote Zone | Up | Esyt3 | 3.09E-304 | -0.7899411 | 0.416 | 0.923 | 6.17E-301 |
| Saline Remote Zone | Up | Arnt | 0 | -0.7889115 | 0.323 | 0.606 | 0 |
| Saline Remote Zone | Up | Gpr75 | 0 | -0.7835961 | 0.502 | 0.948 | 0 |
| Saline Remote Zone | Up | Thy1 | 0 | -0.7800479 | 0.307 | 0.579 | 0 |

| | | | | | | | |
|---|---|---|---|---|---|---|---|
| Saline Remote Zone | Up | Nrg1 | 2.73E-251 | -0.7767564 | 0.483 | 0.924 | 5.47E-248 |
| Saline Remote Zone | Up | Scg2 | 0 | -0.7720385 | 0.115 | 0.43 | 0 |
| Saline Remote Zone | Up | RGD1564149 | 0 | -0.7704335 | 0.408 | 0.709 | 0 |
| Saline Remote Zone | Up | Cma1 | 8.95E-238 | -0.7612882 | 0.583 | 0.884 | 1.79E-234 |
| Saline Remote Zone | Up | Spi1 | 5.47E-235 | -0.7357008 | 0.491 | 0.727 | 1.09E-231 |
| Saline Remote Zone | Up | Ppp2r2b | 0 | -0.731304 | 0.525 | 0.89 | 0 |
| Saline Remote Zone | Up | Pcyox1 | 0 | -0.7271938 | 0.36 | 0.75 | 0 |
| Saline Remote Zone | Up | Ttll9 | 0 | -0.7258665 | 0.379 | 0.688 | 0 |
| Saline Remote Zone | Up | LOC102549542 | 1.04E-295 | -0.7226399 | 0.442 | 0.832 | 2.08E-292 |

| | | | | | | | |
|---|---|---|---|---|---|---|---|
| Saline Remote Zone | Up | Efemp2 | 4.35E-210 | -0.7195132 | 0.397 | 0.605 | 8.71E-207 |
| Saline Remote Zone | Up | Ccnt1 | 4.50E-140 | -0.6982932 | 0.441 | 0.643 | 9.01E-137 |
| Saline Remote Zone | Up | Fkbp1b | 0 | -0.6943776 | 0.235 | 0.51 | 0 |
| Saline Remote Zone | Up | Ccl1 | 1.46E-211 | -0.6873357 | 0.478 | 0.757 | 2.92E-208 |
| Saline Remote Zone | Up | Mal | 0 | -0.6862467 | 0.484 | 0.855 | 0 |
| Saline Remote Zone | Up | Mcpt1 | 1.82E-96 | -0.6849766 | 0.359 | 0.152 | 3.64E-93 |
| Saline Remote Zone | Up | LOC502684 | 0 | -0.6745427 | 0.418 | 0.79 | 0 |
| Saline Remote Zone | Up | Aacs | 2.22E-279 | -0.6714875 | 0.433 | 0.725 | 4.44E-276 |
| Saline Remote Zone | Up | Odf4 | 0 | -0.6699124 | 0.474 | 0.858 | 0 |

| | | | | | | | |
|---|---|---|---|---|---|---|---|
| Saline Remote Zone | Up | Sfrp4 | 0 | -0.6688127 | 0.406 | 0.666 | 0 |
| Saline Remote Zone | Up | Baz2a | 0 | -0.6624049 | 0.208 | 0.447 | 0 |
| Saline Remote Zone | Up | Raph1 | 0 | -0.6605293 | 0.263 | 0.534 | 0 |
| Saline Remote Zone | Up | Nub1 | 0 | -0.6581328 | 0.424 | 0.718 | 0 |
| Saline Remote Zone | Up | Rnf212 | 0 | -0.6542228 | 0.423 | 0.769 | 0 |
| Saline Remote Zone | Up | Tpsab1 | 6.47E-140 | -0.6401746 | 0.509 | 0.759 | 1.29E-136 |
| Saline Remote Zone | Up | Fgr | 0 | -0.6328572 | 0.395 | 0.664 | 0 |
| Saline Remote Zone | Up | Osr1 | 8.02E-263 | -0.6268131 | 0.488 | 0.729 | 1.60E-259 |
| Saline Remote Zone | Up | Lmod1 | 0 | -0.623588 | 0.374 | 0.595 | 0 |

| | | | | | | | |
|---|---|---|---|---|---|---|---|
| Saline Remote Zone | Up | Cd163 | 0 | -0.623114 | 0.341 | 0.698 | 0 |
| Saline Remote Zone | Up | Gdf6 | 0 | -0.62093 | 0.406 | 0.698 | 0 |
| Saline Remote Zone | Up | Cd7 | 0 | -0.6155049 | 0.485 | 0.768 | 0 |
| Saline Remote Zone | Up | Cpa3 | 5.02E-49 | -0.614918 | 0.522 | 0.737 | 1.00E-45 |
| Saline Remote Zone | Up | Kcnn4 | 0 | -0.6035362 | 0.459 | 0.78 | 0 |
| Saline Remote Zone | Up | Csf3r | 0 | -0.6027653 | 0.424 | 0.773 | 0 |
| Saline Remote Zone | Up | AABR07006097.1 | 1.59E-211 | -0.6018119 | 0.439 | 0.687 | 3.18E-208 |
| Saline Remote Zone | Up | Krt18 | 4.81E-280 | -0.6013275 | 0.452 | 0.765 | 9.61E-277 |
| Saline Remote Zone | Up | Paip2b | 0 | -0.5988715 | 0.266 | 0.497 | 0 |

| | | | | | | | |
|---|---|---|---|---|---|---|---|
| Saline Remote Zone | Up | Hoga1 | 0 | -0.5950096 | 0.336 | 0.791 | 0 |
| Saline Remote Zone | Up | Cfap300 | 0 | -0.5896538 | 0.397 | 0.879 | 0 |
| Saline Remote Zone | Up | Lag3 | 0 | -0.5861449 | 0.298 | 0.566 | 0 |
| Saline Remote Zone | Up | Pstpip1 | 5.41E-199 | -0.5856156 | 0.471 | 0.676 | 1.08E-195 |
| Saline Remote Zone | Up | Lrrc25 | 6.36E-207 | -0.5842263 | 0.466 | 0.675 | 1.27E-203 |
| Saline Remote Zone | Up | Rhox5 | 5.90E-260 | -0.5841704 | 0.478 | 0.739 | 1.18E-256 |
| Saline Remote Zone | Up | Plxdc2 | 2.59E-256 | -0.5825767 | 0.353 | 0.566 | 5.19E-253 |
| Saline Remote Zone | Up | Ccl24 | 1.05E-163 | -0.5682706 | 0.554 | 0.761 | 2.11E-160 |
| Saline Remote Zone | Up | Irf6 | 0 | -0.5672855 | 0.448 | 0.791 | 0 |

| | | | | | | | |
|---|---|---|---|---|---|---|---|
| Saline Remote Zone | Up | Clip3 | 9.81E-166 | -0.5596247 | 0.469 | 0.686 | 1.96E-162 |
| Saline Remote Zone | Up | P2rx1 | 2.40E-140 | -0.5522107 | 0.538 | 0.807 | 4.80E-137 |
| Saline Remote Zone | Up | Synpo2 | 3.00E-283 | -0.5496881 | 0.472 | 0.708 | 6.01E-280 |
| Saline Remote Zone | Up | Arnt2 | 1.69E-229 | -0.5467131 | 0.478 | 0.792 | 3.38E-226 |
| Saline Remote Zone | Up | Col20a1 | 0 | -0.5445756 | 0.445 | 0.862 | 0 |
| Saline Remote Zone | Up | Arhgef25 | 4.43E-263 | -0.5437569 | 0.323 | 0.532 | 8.85E-260 |
| Saline Remote Zone | Up | Bcorl1 | 1.89E-273 | -0.5409501 | 0.478 | 0.719 | 3.77E-270 |
| Saline Remote Zone | Up | Chd3 | 1.38E-167 | -0.5385419 | 0.521 | 0.749 | 2.76E-164 |
| Saline Remote Zone | Up | Ptafr | 3.31E-304 | -0.5366633 | 0.482 | 0.732 | 6.62E-301 |

| | | | | | | | |
|---|---|---|---|---|---|---|---|
| Saline Remote Zone | Up | Rcn1 | 2.04E-246 | -0.5358063 | 0.447 | 0.691 | 4.09E-243 |
| Saline Remote Zone | Up | Spn | 3.65E-280 | -0.5302028 | 0.321 | 0.521 | 7.31E-277 |
| Saline Remote Zone | Up | Pak1 | 0 | -0.5291653 | 0.532 | 0.871 | 0 |
| Saline Remote Zone | Up | Gpr68 | 0 | -0.5279099 | 0.396 | 0.765 | 0 |
| Saline Remote Zone | Up | Ect2 | 0 | -0.5271801 | 0.357 | 0.67 | 0 |
| Saline Remote Zone | Up | Clec9a | 2.26E-272 | -0.5213439 | 0.516 | 0.869 | 4.52E-269 |
| Saline Remote Zone | Up | Mafb | 0 | -0.5203347 | 0.434 | 0.772 | 0 |
| Saline Remote Zone | Up | Ephb3 | 2.16E-197 | -0.5145659 | 0.536 | 0.828 | 4.32E-194 |
| Saline Remote Zone | Up | Stard4 | 0 | -0.5137794 | 0.417 | 0.699 | 0 |

| | | | | | | | |
|---|---|---|---|---|---|---|---|
| Saline Remote Zone | Up | Tnfrsf18 | 0 | -0.5114811 | 0.415 | 0.831 | 0 |
| Saline Remote Zone | Up | Capn5 | 1.86E-235 | -0.510033 | 0.402 | 0.617 | 3.71E-232 |
| Saline Remote Zone | Up | Fmnl2 | 0 | -0.5095925 | 0.453 | 0.886 | 0 |
| Saline Remote Zone | Up | Lst1 | 7.29E-243 | -0.5095905 | 0.509 | 0.733 | 1.46E-239 |
| Saline Remote Zone | Up | Fosb | 0 | -0.5062228 | 0.263 | 0.476 | 0 |
| Saline Remote Zone | Up | Nkd2 | 5.73E-289 | -0.5060625 | 0.448 | 0.705 | 1.15E-285 |
| Saline Remote Zone | Up | AABR07005821.1 | 1.12E-299 | -0.4889181 | 0.551 | 0.889 | 2.24E-296 |
| Saline Remote Zone | Up | Zkscan1 | 0 | -0.4876742 | 0.374 | 0.736 | 0 |
| Saline Remote Zone | Up | Msln | 0 | -0.4874002 | 0.154 | 0.368 | 0 |

| | | | | | | | |
|---|---|---|---|---|---|---|---|
| Saline Remote Zone | Up | Dzip3 | 4.00E-250 | -0.4848832 | 0.395 | 0.631 | 8.01E-247 |
| Saline Remote Zone | Up | Rgs7 | 2.18E-306 | -0.4825995 | 0.472 | 0.854 | 4.36E-303 |
| Saline Remote Zone | Up | Cd3e | 0 | -0.4772402 | 0.378 | 0.687 | 0 |
| Saline Remote Zone | Up | Ldb2 | 0 | -0.4746422 | 0.465 | 0.78 | 0 |
| Saline Remote Zone | Up | Fgd2 | 8.05E-308 | -0.4737944 | 0.462 | 0.745 | 1.61E-304 |
| Saline Remote Zone | Up | Olfml2a | 0 | -0.4707314 | 0.359 | 0.652 | 0 |
| Saline Remote Zone | Up | Kif23 | 0 | -0.4696844 | 0.441 | 0.773 | 0 |
| Saline Remote Zone | Up | Il27ra | 0 | -0.4673075 | 0.436 | 0.757 | 0 |
| Saline Remote Zone | Up | Ch25h | 0 | -0.4627072 | 0.451 | 0.776 | 0 |

| | | | | | | | |
|---|---|---|---|---|---|---|---|
| Saline Remote Zone | Up | Akap7 | 0 | -0.4598169 | 0.294 | 0.541 | 0 |
| Saline Remote Zone | Up | Trh | 3.86E-295 | -0.4590746 | 0.448 | 0.69 | 7.73E-292 |
| Saline Remote Zone | Up | Dot1l | 1.90E-299 | -0.4582642 | 0.49 | 0.805 | 3.79E-296 |
| Saline Remote Zone | Up | Ccl20 | 1.09E-280 | -0.4577936 | 0.519 | 0.956 | 2.18E-277 |
| Saline Remote Zone | Up | Zdhhc17 | 1.10E-306 | -0.4556615 | 0.447 | 0.754 | 2.20E-303 |
| Saline Remote Zone | Up | Lox | 1.39E-291 | -0.4541626 | 0.47 | 0.781 | 2.79E-288 |
| Saline Remote Zone | Up | Srek1 | 3.87E-232 | -0.4515539 | 0.338 | 0.546 | 7.74E-229 |
| Saline Remote Zone | Up | Cd247 | 0 | -0.451058 | 0.205 | 0.421 | 0 |
| Saline Remote Zone | Up | Ro60 | 0 | -0.446156 | 0.516 | 0.824 | 0 |

| | | | | | | | |
|---|---|---|---|---|---|---|---|
| Saline Remote Zone | Up | Fibin | 4.06E-177 | -0.4430309 | 0.486 | 0.707 | 8.12E-174 |
| Saline Remote Zone | Up | Aldh3a1 | 9.37E-175 | -0.4426054 | 0.463 | 0.67 | 1.87E-171 |
| Saline Remote Zone | Up | Col18a1 | 8.35E-156 | -0.4412604 | 0.498 | 0.719 | 1.67E-152 |
| Saline Remote Zone | Up | Myo1g | 3.59E-227 | -0.4407561 | 0.558 | 0.794 | 7.19E-224 |
| Saline Remote Zone | Up | Abca4 | 0 | -0.4384343 | 0.413 | 0.732 | 0 |
| Saline Remote Zone | Up | Hapln1 | 0 | -0.4370877 | 0.104 | 0.321 | 0 |
| Saline Remote Zone | Up | Dusp2 | 0 | -0.4353162 | 0.39 | 0.672 | 0 |
| Saline Remote Zone | Up | Qprt | 9.73E-225 | -0.4338869 | 0.428 | 0.708 | 1.95E-221 |
| Saline Remote Zone | Up | Elfn1 | 1.95E-274 | -0.4298187 | 0.449 | 0.715 | 3.90E-271 |

| | | | | | | | |
|---|---|---|---|---|---|---|---|
| Saline Remote Zone | Up | Cdc6 | 0 | -0.42958 | 0.244 | 0.559 | 0 |
| Saline Remote Zone | Up | Ptpn6 | 1.52E-185 | -0.4276657 | 0.576 | 0.789 | 3.04E-182 |
| Saline Remote Zone | Up | Fcgr2b | 0 | -0.4276039 | 0.269 | 0.49 | 0 |
| Saline Remote Zone | Up | Matk | 1.97E-232 | -0.4272678 | 0.483 | 0.735 | 3.95E-229 |
| Saline Remote Zone | Up | Supt16h | 1.70E-278 | -0.426442 | 0.357 | 0.57 | 3.41E-275 |
| Saline Remote Zone | Up | AABR07008439.1 | 0 | -0.4257128 | 0.333 | 0.56 | 0 |
| Saline Remote Zone | Up | Tubb3 | 7.92E-283 | -0.4237944 | 0.341 | 0.569 | 1.58E-279 |
| Saline Remote Zone | Up | Ccr5 | 1.85E-217 | -0.4232596 | 0.503 | 0.789 | 3.70E-214 |
| Saline Remote Zone | Up | Ciita | 0 | -0.4228036 | 0.538 | 0.871 | 0 |

| | | | | | | | |
|---|---|---|---|---|---|---|---|
| Saline Remote Zone | Up | P2ry10 | 0 | -0.4224556 | 0.363 | 0.679 | 0 |
| Saline Remote Zone | Up | Tnfsf13 | 2.62E-253 | -0.4223089 | 0.421 | 0.652 | 5.24E-250 |
| Saline Remote Zone | Up | Map9 | 0 | -0.4188762 | 0.448 | 0.756 | 0 |
| Saline Remote Zone | Up | Gprc5a | 0 | -0.4179014 | 0.314 | 0.599 | 0 |
| Saline Remote Zone | Up | Ipo9 | 1.13E-168 | -0.4158478 | 0.462 | 0.666 | 2.26E-165 |
| Saline Remote Zone | Up | Kif20b | 0 | -0.415194 | 0.421 | 0.687 | 0 |
| Saline Remote Zone | Up | Fst | 0 | -0.4106381 | 0.384 | 0.702 | 0 |
| Saline Remote Zone | Up | RGD1311744 | 7.21E-216 | -0.4104954 | 0.335 | 0.542 | 1.44E-212 |
| Saline Remote Zone | Up | Xcl1 | 3.57E-232 | -0.4081132 | 0.505 | 0.853 | 7.13E-229 |

| | | | | | | | |
|---|---|---|---|---|---|---|---|
| Saline Remote Zone | Up | AABR07072108.1 | 0 | -0.407495 | 0.466 | 0.945 | 0 |
| Saline Remote Zone | Up | Gja4 | 0 | -0.4064006 | 0.268 | 0.536 | 0 |
| Saline Remote Zone | Up | Matn4 | 4.06E-149 | -0.4060134 | 0.505 | 0.72 | 8.12E-146 |
| Saline Remote Zone | Up | Slc39a6 | 1.15E-214 | -0.404455 | 0.507 | 0.779 | 2.29E-211 |
| Saline Remote Zone | Up | Apobec1 | 0 | -0.4042409 | 0.467 | 0.772 | 0 |
| Saline Remote Zone | Up | Adamts4 | 2.90E-228 | -0.4036346 | 0.548 | 0.954 | 5.80E-225 |
| Saline Remote Zone | Up | Rfx8 | 0 | -0.4026561 | 0.296 | 0.675 | 0 |
| Saline Remote Zone | Up | Cpz | 0 | -0.4016976 | 0.319 | 0.571 | 0 |
| Saline Remote Zone | Up | Golim4 | 1.32E-224 | -0.4000318 | 0.516 | 0.776 | 2.65E-221 |

| | | | | | | | |
|---|---|---|---|---|---|---|---|
| Saline Remote Zone | Up | Coro1a | 1.87E-135 | -0.3993074 | 0.589 | 0.802 | 3.73E-132 |
| Saline Remote Zone | Up | Fstl3 | 1.45E-219 | -0.3987284 | 0.489 | 0.726 | 2.90E-216 |
| Saline Remote Zone | Up | AC120486.10 | 2.18E-216 | -0.3977464 | 0.472 | 0.731 | 4.37E-213 |
| Saline Remote Zone | Up | Ino80 | 2.52E-267 | -0.3976875 | 0.45 | 0.704 | 5.05E-264 |
| Saline Remote Zone | Up | Inmt | 0 | -0.3973583 | 0.428 | 0.769 | 0 |
| Saline Remote Zone | Up | Nfkbiz | 0 | -0.3973526 | 0.459 | 0.764 | 0 |
| Saline Remote Zone | Up | Ice1 | 0 | -0.3973429 | 0.248 | 0.496 | 0 |
| Saline Remote Zone | Up | Arhgap9 | 2.80E-245 | -0.397167 | 0.496 | 0.764 | 5.59E-242 |
| Saline Remote Zone | Up | C1qtnf3 | 0 | -0.3951217 | 0.47 | 0.789 | 0 |

| | | | | | | | |
|---|---|---|---|---|---|---|---|
| Saline Remote Zone | Up | Trem1 | 0 | -0.3926886 | 0.363 | 0.707 | 0 |
| Saline Remote Zone | Up | Gng2 | 4.55E-210 | -0.3904793 | 0.539 | 0.784 | 9.11E-207 |
| Saline Remote Zone | Up | Gzma | 3.01E-124 | -0.3891413 | 0.551 | 0.896 | 6.03E-121 |
| Saline Remote Zone | Up | Lyve1 | 3.25E-198 | -0.3888605 | 0.427 | 0.632 | 6.51E-195 |
| Saline Remote Zone | Up | Lrrc17 | 8.49E-204 | -0.3879863 | 0.482 | 0.696 | 1.70E-200 |
| Saline Remote Zone | Up | Lxn | 1.45E-213 | -0.387102 | 0.51 | 0.765 | 2.91E-210 |
| Saline Remote Zone | Up | Sertad4 | 3.47E-233 | -0.3860097 | 0.509 | 0.763 | 6.94E-230 |
| Saline Remote Zone | Up | AABR07017902.1 | 0 | -0.3818136 | 0.429 | 0.74 | 0 |
| Saline Remote Zone | Up | Kif21a | 0 | -0.3811655 | 0.356 | 0.636 | 0 |

| | | | | | | | |
|---|---|---|---|---|---|---|---|
| Saline Remote Zone | Up | Clec10a | 2.90E-248 | -0.3810988 | 0.425 | 0.697 | 5.80E-245 |
| Saline Remote Zone | Up | Snx20 | 1.05E-127 | -0.3793006 | 0.523 | 0.763 | 2.11E-124 |
| Saline Remote Zone | Up | Gpr171 | 0 | -0.3790556 | 0.557 | 0.895 | 0 |
| Saline Remote Zone | Up | Enpp1 | 1.20E-286 | -0.3775384 | 0.389 | 0.687 | 2.39E-283 |
| Saline Remote Zone | Up | Ints6 | 3.10E-181 | -0.3752681 | 0.55 | 0.805 | 6.20E-178 |
| Saline Remote Zone | Up | S100a9 | 0 | -0.3748004 | 0.395 | 0.751 | 0 |
| Saline Remote Zone | Up | Cradd | 1.28E-211 | -0.374469 | 0.426 | 0.629 | 2.57E-208 |
| Saline Remote Zone | Up | Smarca1 | 0 | -0.3744155 | 0.306 | 0.633 | 0 |
| Saline Remote Zone | Up | Pole | 4.92E-214 | -0.3743688 | 0.45 | 0.65 | 9.85E-211 |

| | | | | | | | |
|---|---|---|---|---|---|---|---|
| Saline Remote Zone | Up | Foxc2 | 0 | -0.3743601 | 0.323 | 0.565 | 0 |
| Saline Remote Zone | Up | AC119762.7 | 1.58E-297 | -0.3743583 | 0.489 | 0.769 | 3.16E-294 |
| Saline Remote Zone | Up | Orai2 | 6.09E-284 | -0.3740168 | 0.479 | 0.724 | 1.22E-280 |
| Saline Remote Zone | Up | Nrgn | 1.50E-285 | -0.3718467 | 0.49 | 0.776 | 3.00E-282 |
| Saline Remote Zone | Up | AABR07001054.1 | 0 | -0.3716868 | 0.272 | 0.531 | 0 |
| Saline Remote Zone | Up | Postn | 1.30E-242 | -0.3645327 | 0.442 | 0.764 | 2.61E-239 |
| Saline Remote Zone | Up | Cnn1 | 2.52E-295 | -0.3639674 | 0.448 | 0.707 | 5.04E-292 |
| Saline Remote Zone | Up | Plac8 | 1.04E-221 | -0.3632244 | 0.507 | 0.728 | 2.08E-218 |
| Saline Remote Zone | Up | Cyb5d2 | 8.08E-239 | -0.3631547 | 0.423 | 0.679 | 1.62E-235 |

| | | | | | | | |
|---|---|---|---|---|---|---|---|
| Saline Remote Zone | Up | Acta2 | 9.05E-193 | -0.3613747 | 0.429 | 0.634 | 1.81E-189 |
| Saline Remote Zone | Up | Tmem100 | 1.05E-255 | -0.3601894 | 0.361 | 0.588 | 2.09E-252 |
| Saline Remote Zone | Up | Col16a1 | 0 | -0.3597514 | 0.389 | 0.704 | 0 |
| Saline Remote Zone | Up | Tspan6 | 3.30E-250 | -0.3578995 | 0.521 | 0.816 | 6.60E-247 |
| Saline Remote Zone | Up | Kcne1 | 6.50E-148 | -0.3545406 | 0.57 | 0.773 | 1.30E-144 |
| Saline Remote Zone | Up | Rgs4 | 2.30E-174 | -0.3478966 | 0.491 | 0.733 | 4.60E-171 |
| Saline Remote Zone | Up | Satb1 | 3.17E-285 | -0.3475058 | 0.386 | 0.619 | 6.33E-282 |
| Saline Remote Zone | Up | Asap1 | 0 | -0.3436881 | 0.27 | 0.529 | 0 |
| Saline Remote Zone | Up | AABR07051518.1 | 0 | -0.3405127 | 0.164 | 0.374 | 0 |

| | | | | | | | |
|---|---|---|---|---|---|---|---|
| Saline Remote Zone | Up | AC128848.1 | 6.81E-255 | -0.3403832 | 0.532 | 0.823 | 1.36E-251 |
| Saline Remote Zone | Up | Ncf4 | 0 | -0.3401421 | 0.555 | 0.844 | 0 |
| Saline Remote Zone | Up | Myl1 | 7.49E-151 | -0.3368188 | 0.666 | 0.898 | 1.50E-147 |
| Saline Remote Zone | Up | Mfap2 | 1.46E-228 | -0.3362019 | 0.337 | 0.537 | 2.91E-225 |
| Saline Remote Zone | Up | S100b | 0 | -0.3358627 | 0.342 | 0.636 | 0 |
| Saline Remote Zone | Up | C1qtnf7 | 0 | -0.3345598 | 0.361 | 0.653 | 0 |
| Saline Remote Zone | Up | AABR07030501.1 | 5.69E-73 | -0.33428 | 0.525 | 0.806 | 1.14E-69 |
| Saline Remote Zone | Up | Hcls1 | 3.06E-217 | -0.3320217 | 0.527 | 0.769 | 6.12E-214 |
| Saline Remote Zone | Up | Ly86 | 1.99E-217 | -0.3306578 | 0.505 | 0.752 | 3.99E-214 |

| | | | | | | | |
|---|---|---|---|---|---|---|---|
| Saline Remote Zone | Up | Bcl2a1 | 2.39E-206 | -0.3301717 | 0.537 | 0.799 | 4.79E-203 |
| Saline Remote Zone | Up | Igf1 | 1.99E-138 | -0.3299926 | 0.558 | 0.766 | 3.98E-135 |
| Saline Remote Zone | Up | Cxcl14 | 0 | -0.3286063 | 0.355 | 0.595 | 0 |
| Saline Remote Zone | Up | LOC100363469 | 5.26E-304 | -0.3276331 | 0.497 | 0.839 | 1.05E-300 |
| Saline Remote Zone | Up | Sla | 0 | -0.3253203 | 0.379 | 0.702 | 0 |
| Saline Remote Zone | Up | Myh11 | 4.55E-205 | -0.3252008 | 0.457 | 0.693 | 9.10E-202 |
| Saline Remote Zone | Up | Hcst | 1.18E-293 | -0.3251889 | 0.5 | 0.793 | 2.35E-290 |
| Saline Remote Zone | Up | Wdfy4 | 8.88E-233 | -0.3248325 | 0.367 | 0.579 | 1.78E-229 |
| Saline Remote Zone | Up | Cyp1b1 | 5.18E-231 | -0.3241357 | 0.497 | 0.739 | 1.04E-227 |

| | | | | | | | |
|---|---|---|---|---|---|---|---|
| Saline Remote Zone | Up | Arl11 | 3.57E-225 | -0.3236085 | 0.39 | 0.619 | 7.15E-222 |
| Saline Remote Zone | Up | Col6a1 | 1.06E-204 | -0.3209263 | 0.455 | 0.694 | 2.11E-201 |
| Saline Remote Zone | Up | Ccl9 | 1.44E-248 | -0.3196389 | 0.473 | 0.899 | 2.88E-245 |
| Saline Remote Zone | Up | S100a8 | 0 | -0.3194133 | 0.253 | 0.467 | 0 |
| Saline Remote Zone | Up | Gpnmb | 4.43E-208 | -0.3185833 | 0.464 | 0.719 | 8.87E-205 |
| Saline Remote Zone | Up | Tlr2 | 7.63E-165 | -0.3184805 | 0.518 | 0.736 | 1.53E-161 |
| Saline Remote Zone | Up | Ccr1 | 3.36E-229 | -0.3171188 | 0.484 | 0.792 | 6.72E-226 |
| Saline Remote Zone | Up | Msr1 | 0 | -0.3161711 | 0.372 | 0.667 | 0 |
| Saline Remote Zone | Up | Cacna1a | 7.77E-171 | -0.3159363 | 0.499 | 0.703 | 1.55E-167 |

| | | | | | | | |
|---|---|---|---|---|---|---|---|
| Saline Remote Zone | Up | Sycp2 | 0 | -0.3137132 | 0.392 | 0.773 | 0 |
| Saline Remote Zone | Up | Ctsk | 1.76E-181 | -0.3095736 | 0.441 | 0.668 | 3.52E-178 |
| Saline Remote Zone | Up | Myof | 0 | -0.3094196 | 0.446 | 0.761 | 0 |
| Saline Remote Zone | Up | Fam169b | 0 | -0.3061734 | 0.427 | 0.743 | 0 |
| Saline Remote Zone | Up | Rarres1 | 0 | -0.3060829 | 0.432 | 0.764 | 0 |
| Saline Remote Zone | Up | Cthrc1 | 0 | -0.3048038 | 0.309 | 0.621 | 0 |
| Saline Remote Zone | Up | Wapl | 0 | -0.302436 | 0.27 | 0.639 | 0 |
| Saline Remote Zone | Up | Cd68 | 1.03E-305 | -0.3008929 | 0.52 | 0.818 | 2.07E-302 |
| Saline Remote Zone | Up | Cemip | 0 | -0.3004285 | 0.326 | 0.607 | 0 |

| | | | | | | | |
|---|---|---|---|---|---|---|---|
| Saline Remote Zone | Up | Klre1 | 4.03E-162 | -0.2979657 | 0.495 | 0.899 | 8.06E-159 |
| Saline Remote Zone | Up | Cdo1 | 4.07E-267 | -0.2948249 | 0.418 | 0.659 | 8.14E-264 |
| Saline Remote Zone | Up | Olfm2 | 0 | -0.2920028 | 0.277 | 0.641 | 0 |
| Saline Remote Zone | Up | Zfp950 | 0 | -0.2917812 | 0.397 | 0.61 | 0 |
| Saline Remote Zone | Up | Cxcl1 | 3.96E-269 | -0.2909226 | 0.512 | 0.833 | 7.92E-266 |
| Saline Remote Zone | Up | Cercam | 8.26E-224 | -0.2904573 | 0.419 | 0.681 | 1.65E-220 |
| Saline Remote Zone | Up | Phip | 7.54E-135 | -0.2900659 | 0.548 | 0.77 | 1.51E-131 |
| Saline Remote Zone | Up | Bcl11b | 2.07E-250 | -0.2884641 | 0.477 | 0.749 | 4.13E-247 |
| Saline Remote Zone | Up | Adgre1 | 2.41E-236 | -0.2866308 | 0.46 | 0.694 | 4.81E-233 |

| | | | | | | | |
|---|---|---|---|---|---|---|---|
| Saline Remote Zone | Up | AABR07041096.1 | 4.42E-170 | -0.2864808 | 0.541 | 0.815 | 8.85E-167 |
| Saline Remote Zone | Up | Asic1 | 0 | -0.2861725 | 0.492 | 0.862 | 0 |
| Saline Remote Zone | Up | Tnfrsf14 | 1.58E-279 | -0.2858938 | 0.455 | 0.784 | 3.15E-276 |
| Saline Remote Zone | Up | Cep164 | 3.33E-150 | -0.2845324 | 0.565 | 0.832 | 6.66E-147 |
| Saline Remote Zone | Up | Ptprc | 0 | -0.2839079 | 0.503 | 0.801 | 0 |
| Saline Remote Zone | Up | Mcpt1l4 | 0 | -0.2836513 | 0.467 | 0.891 | 0 |
| Saline Remote Zone | Up | Cep350 | 0 | -0.2836194 | 0.456 | 0.747 | 0 |
| Saline Remote Zone | Up | Anxa4 | 2.01E-142 | -0.2834051 | 0.562 | 0.799 | 4.02E-139 |
| Saline Remote Zone | Up | Bin2 | 2.69E-223 | -0.2824858 | 0.462 | 0.692 | 5.38E-220 |

| | | | | | | | |
|---|---|---|---|---|---|---|---|
| Saline Remote Zone | Up | Sgo2 | 0 | -0.2813923 | 0.288 | 0.52 | 0 |
| Saline Remote Zone | Up | Snai1 | 1.46E-227 | -0.2769049 | 0.424 | 0.66 | 2.93E-224 |
| Saline Remote Zone | Up | Cacna1g | 1.46E-212 | -0.2764082 | 0.566 | 0.828 | 2.92E-209 |
| Saline Remote Zone | Up | Tagln | 5.39E-133 | -0.2756732 | 0.51 | 0.727 | 1.08E-129 |
| Saline Remote Zone | Up | Adcy7 | 0 | -0.2750435 | 0.347 | 0.712 | 0 |
| Saline Remote Zone | Up | H1f2 | 0 | -0.2738324 | 0.265 | 0.552 | 0 |
| Saline Remote Zone | Up | Ggn | 0 | -0.2738005 | 0.459 | 0.855 | 0 |
| Saline Remote Zone | Up | Lck | 5.11E-292 | -0.2691931 | 0.49 | 0.807 | 1.02E-288 |
| Saline Remote Zone | Up | Actg2 | 9.74E-227 | -0.2685638 | 0.51 | 0.723 | 1.95E-223 |

| | | | | | | | |
|---|---|---|---|---|---|---|---|
| Saline Remote Zone | Up | Fcgr2a | 3.94E-300 | -0.2683452 | 0.527 | 0.795 | 7.87E-297 |
| Saline Remote Zone | Up | Vcam1 | 1.99E-275 | -0.2663142 | 0.488 | 0.77 | 3.97E-272 |
| Saline Remote Zone | Up | Golga4 | 5.69E-195 | -0.2655211 | 0.549 | 0.814 | 1.14E-191 |
| Saline Remote Zone | Up | Slc4a4 | 0 | -0.2650079 | 0.534 | 0.866 | 0 |
| Saline Remote Zone | Up | Pmepa1 | 4.47E-114 | -0.2637802 | 0.545 | 0.757 | 8.94E-111 |
| Saline Remote Zone | Up | Hamp | 2.45E-173 | -0.2627225 | 0.616 | 0.859 | 4.90E-170 |
| Saline Remote Zone | Up | Hspa1b | 0 | -0.2625759 | 0.309 | 0.586 | 0 |
| Saline Remote Zone | Up | Cntfr | 1.61E-268 | -0.2618122 | 0.425 | 0.628 | 3.21E-265 |
| Saline Remote Zone | Up | Ptprf | 0 | -0.2616466 | 0.436 | 0.723 | 0 |

| | | | | | | | |
|---|---|---|---|---|---|---|---|
| Saline Remote Zone | Up | Rab7b | 0 | -0.260473 | 0.425 | 0.69 | 0 |
| Saline Remote Zone | Up | Aga | 2.84E-153 | -0.2600974 | 0.499 | 0.713 | 5.68E-150 |
| Saline Remote Zone | Up | Trib3 | 1.06E-169 | -0.2595926 | 0.497 | 0.715 | 2.11E-166 |
| Saline Remote Zone | Up | AC109737.1 | 0 | -0.2591749 | 0.515 | 0.909 | 0 |
| Saline Remote Zone | Up | Rxfp1 | 5.54E-102 | -0.2588997 | 0.615 | 0.832 | 1.11E-98 |
| Saline Remote Zone | Up | Grb10 | 7.27E-219 | -0.2585099 | 0.478 | 0.765 | 1.45E-215 |
| Saline Remote Zone | Up | AABR07030351.1 | 2.08E-216 | -0.2581404 | 0.434 | 0.698 | 4.16E-213 |
| Saline Remote Zone | Up | Efhd2 | 5.89E-179 | -0.2540402 | 0.516 | 0.774 | 1.18E-175 |
| Saline Remote Zone | Up | Itgax | 2.34E-136 | -0.2532927 | 0.47 | 0.689 | 4.69E-133 |

| | | | | | | | |
|---|---|---|---|---|---|---|---|
| Saline Remote Zone | Up | Kif20a | 0 | -0.2530335 | 0.504 | 0.941 | 0 |
| Saline Remote Zone | Up | AABR07030791.1 | 0 | -0.2528948 | 0.488 | 0.929 | 0 |
| Saline Remote Zone | Up | Tusc3 | 4.83E-266 | -0.2510395 | 0.376 | 0.621 | 9.66E-263 |
| Saline Remote Zone | Up | Dhh | 0 | -0.2500682 | 0.268 | 0.496 | 0 |
| ECM Remote Zone | Up | Stx17 | 1.18E-139 | 0.25338234 | 0.48 | 0.78 | 2.36E-136 |
| ECM Remote Zone | Up | Cdc20 | 0 | 0.2573736 | 0.34 | 0.568 | 0 |
| ECM Remote Zone | Up | Pttg1 | 0 | 0.2577235 | 0.39 | 0.601 | 0 |
| ECM Remote Zone | Up | Zdhhc23 | 0 | 0.25915141 | 0.432 | 0.744 | 0 |
| ECM Remote Zone | Up | Scn2a | 0 | 0.26430573 | 0.279 | 0.701 | 0 |

| | | | | | | | |
|---|---|---|---|---|---|---|---|
| ECM Remote Zone | Up | Kif4a | 1.61E-217 | 0.26445045 | 0.451 | 0.75 | 3.23E-214 |
| ECM Remote Zone | Up | Sema4a | 9.92E-118 | 0.26842104 | 0.525 | 0.775 | 1.98E-114 |
| ECM Remote Zone | Up | Ncaph | 1.31E-295 | 0.27297805 | 0.383 | 0.614 | 2.63E-292 |
| ECM Remote Zone | Up | Rbp4 | 3.23E-165 | 0.27514533 | 0.543 | 0.927 | 6.47E-162 |
| ECM Remote Zone | Up | Col11a1 | 0 | 0.27695105 | 0.219 | 0.47 | 0 |
| ECM Remote Zone | Up | Dnajc5 | 1.96E-244 | 0.27885722 | 0.464 | 0.745 | 3.92E-241 |
| ECM Remote Zone | Up | Akr1b8 | 6.48E-281 | 0.2894427 | 0.433 | 0.724 | 1.30E-277 |
| ECM Remote Zone | Up | Cd3g | 0 | 0.29207718 | 0.408 | 0.715 | 0 |
| ECM Remote Zone | Up | Zbtb47 | 0 | 0.29864115 | 0.24 | 0.461 | 0 |

| | | | | | | | |
|---|---|---|---|---|---|---|---|
| ECM Remote Zone | Up | Ckap2 | 0 | 0.30182764 | 0.42 | 0.731 | 0 |
| ECM Remote Zone | Up | Agr2 | 0 | 0.30922031 | 0.308 | 0.535 | 0 |
| ECM Remote Zone | Up | Dnajb14 | 1.25E-277 | 0.31105255 | 0.458 | 0.702 | 2.50E-274 |
| ECM Remote Zone | Up | Cyp26b1 | 0 | 0.33178372 | 0.425 | 0.771 | 0 |
| ECM Remote Zone | Up | Tbc1d9 | 0 | 0.33199805 | 0.35 | 0.659 | 0 |
| ECM Remote Zone | Up | Cdk12 | 1.29E-217 | 0.33470588 | 0.408 | 0.615 | 2.58E-214 |
| ECM Remote Zone | Up | Jaml | 0 | 0.33614061 | 0.517 | 0.86 | 0 |
| ECM Remote Zone | Up | Eef1a1 | 1.33E-197 | 0.33637889 | 0.361 | 0.561 | 2.65E-194 |
| ECM Remote Zone | Up | Lpal2 | 0 | 0.3364542 | 0.285 | 0.56 | 0 |

| | | | | | | | |
|---|---|---|---|---|---|---|---|
| ECM Remote Zone | Up | Fign | 8.24E-183 | 0.33673594 | 0.48 | 0.791 | 1.65E-179 |
| ECM Remote Zone | Up | Mllt3 | 9.33E-125 | 0.34316962 | 0.498 | 0.73 | 1.87E-121 |
| ECM Remote Zone | Up | Vcpip1 | 0 | 0.34642012 | 0.386 | 0.638 | 0 |
| ECM Remote Zone | Up | Pmaip1 | 0 | 0.34689896 | 0.416 | 0.811 | 0 |
| ECM Remote Zone | Up | Ifit1bl | 0 | 0.35031837 | 0.343 | 0.703 | 0 |
| ECM Remote Zone | Up | Tnn | 0 | 0.36005657 | 0.214 | 0.624 | 0 |
| ECM Remote Zone | Up | Casd1 | 0 | 0.36605613 | 0.178 | 0.392 | 0 |
| ECM Remote Zone | Up | Bpgm | 8.28E-215 | 0.3699372 | 0.429 | 0.725 | 1.66E-211 |
| ECM Remote Zone | Up | Tlr8 | 8.18E-298 | 0.370656 | 0.468 | 0.856 | 1.64E-294 |

| | | | | | | | |
|---|---|---|---|---|---|---|---|
| ECM Remote Zone | Up | Plk4 | 4.85E-253 | 0.37163647 | 0.476 | 0.856 | 9.69E-250 |
| ECM Remote Zone | Up | Zfhx2 | 5.94E-244 | 0.37208553 | 0.437 | 0.815 | 1.19E-240 |
| ECM Remote Zone | Up | Rgs1 | 3.70E-190 | 0.37216856 | 0.525 | 0.888 | 7.40E-187 |
| ECM Remote Zone | Up | Mkrn1 | 1.06E-219 | 0.37543147 | 0.432 | 0.727 | 2.13E-216 |
| ECM Remote Zone | Up | Phactr3 | 0 | 0.39727547 | 0.2 | 0.489 | 0 |
| ECM Remote Zone | Up | Ccdc66 | 2.79E-179 | 0.39792014 | 0.498 | 0.738 | 5.58E-176 |
| ECM Remote Zone | Up | Ccne1 | 0 | 0.40818319 | 0.319 | 0.573 | 0 |
| ECM Remote Zone | Up | Hsf2bp | 0 | 0.40882112 | 0.319 | 0.627 | 0 |
| ECM Remote Zone | Up | Ubxn11 | 0 | 0.42088877 | 0.208 | 0.541 | 0 |

| | | | | | | | |
|---|---|---|---|---|---|---|---|
| ECM Remote Zone | Up | Acss3 | 4.21E-282 | 0.42781458 | 0.321 | 0.559 | 8.42E-279 |
| ECM Remote Zone | Up | Dmxl1 | 0 | 0.43191542 | 0.381 | 0.725 | 0 |
| ECM Remote Zone | Up | Herc6 | 0 | 0.43253929 | 0.388 | 0.695 | 0 |
| ECM Remote Zone | Up | Col5a2 | 0 | 0.43728249 | 0.292 | 0.575 | 0 |
| ECM Remote Zone | Up | Osr2 | 0 | 0.44088528 | 0.216 | 0.496 | 0 |
| ECM Remote Zone | Up | Alas2 | 0 | 0.44329221 | 0.462 | 0.962 | 0 |
| ECM Remote Zone | Up | Ccnb1 | 7.14E-287 | 0.44646739 | 0.475 | 0.89 | 1.43E-283 |
| ECM Remote Zone | Up | Tpm4 | 6.99E-263 | 0.45660645 | 0.269 | 0.48 | 1.40E-259 |
| ECM Remote Zone | Up | Cd79b | 3.71E-89 | 0.45671691 | 0.5 | 0.859 | 7.43E-86 |

| | | | | | | | |
|---|---|---|---|---|---|---|---|
| ECM Remote Zone | Up | Tsc1 | 0 | 0.46052755 | 0.424 | 0.76 | 0 |
| ECM Remote Zone | Up | Spc25 | 0 | 0.46121028 | 0.371 | 0.666 | 0 |
| ECM Remote Zone | Up | Smc2 | 6.22E-302 | 0.46331723 | 0.353 | 0.64 | 1.24E-298 |
| ECM Remote Zone | Up | Ube2o | 0 | 0.47722846 | 0.237 | 0.539 | 0 |
| ECM Remote Zone | Up | Mospd1 | 0 | 0.49480212 | 0.331 | 0.597 | 0 |
| ECM Remote Zone | Up | Car2 | 0 | 0.50920654 | 0.359 | 0.811 | 0 |
| ECM Remote Zone | Up | Zbtb41 | 4.55E-291 | 0.51388476 | 0.484 | 0.896 | 9.09E-288 |
| ECM Remote Zone | Up | Abi2 | 0 | 0.51807586 | 0.467 | 0.814 | 0 |
| ECM Remote Zone | Up | Siglec10 | 0 | 0.52501132 | 0.462 | 0.871 | 0 |

| | | | | | | | |
|---|---|---|---|---|---|---|---|
| ECM Remote Zone | Up | Ubd | 2.23E-272 | 0.54404738 | 0.583 | 0.939 | 4.45E-269 |
| ECM Remote Zone | Up | Prss35 | 0 | 0.55556887 | 0.218 | 0.434 | 0 |
| ECM Remote Zone | Up | LOC688553 | 1.09E-70 | 0.55958639 | 0.592 | 0.922 | 2.19E-67 |
| ECM Remote Zone | Up | AABR07030603.1 | 0 | 0.57013505 | 0.119 | 0.32 | 0 |
| ECM Remote Zone | Up | Mpz | 0 | 0.59523337 | 0.424 | 0.733 | 0 |
| ECM Remote Zone | Up | Emb | 0 | 0.60444405 | 0.408 | 0.939 | 0 |
| ECM Remote Zone | Up | Clec1b | 2.90E-263 | 0.62253159 | 0.438 | 0.818 | 5.80E-260 |
| ECM Remote Zone | Up | Epb42 | 0 | 0.62631802 | 0.362 | 0.669 | 0 |
| ECM Remote Zone | Up | Vom2r44 | 5.81E-301 | 0.65391246 | 0.431 | 0.701 | 1.16E-297 |

| | | | | | | | |
|---|---|---|---|---|---|---|---|
| ECM Remote Zone | Up | Tnfrsf11b | 0 | 0.6563166 | 0.284 | 0.504 | 0 |
| ECM Remote Zone | Up | Tent5c | 0 | 0.66474649 | 0.343 | 0.614 | 0 |
| ECM Remote Zone | Up | LOC103694857 | 4.57E-201 | 0.67437832 | 0.484 | 0.963 | 9.14E-198 |
| ECM Remote Zone | Up | Lgals5 | 0 | 0.69062446 | 0.331 | 0.931 | 0 |
| ECM Remote Zone | Up | Prrx1 | 9.75E-253 | 0.7042126 | 0.465 | 0.764 | 1.95E-249 |
| ECM Remote Zone | Up | Ccdc18 | 0 | 0.70654378 | 0.249 | 0.462 | 0 |
| ECM Remote Zone | Up | Cdc25c | 0 | 0.71689746 | 0.259 | 0.551 | 0 |
| ECM Remote Zone | Up | Myef2 | 5.48E-250 | 0.72364507 | 0.439 | 0.695 | 1.10E-246 |
| ECM Remote Zone | Up | Brwd3 | 0 | 0.74458504 | 0.234 | 0.448 | 0 |

| | | | | | | | |
|---|---|---|---|---|---|---|---|
| ECM Remote Zone | Up | AABR07010085.2 | 2.04E-264 | 0.78044977 | 0.447 | 0.781 | 4.08E-261 |
| ECM Remote Zone | Up | Setbp1 | 0 | 0.80096577 | 0.332 | 0.538 | 0 |
| ECM Remote Zone | Up | Mmp12 | 0 | 0.82690294 | 0.233 | 0.454 | 0 |
| ECM Remote Zone | Up | Ahsp | 0 | 0.90576119 | 0.379 | 0.959 | 0 |
| ECM Remote Zone | Up | Cacna1d | 3.61E-168 | 0.92122122 | 0.468 | 0.705 | 7.21E-165 |
| ECM Remote Zone | Up | Fignl1 | 6.17E-279 | 0.98995482 | 0.433 | 0.811 | 1.23E-275 |
| ECM Remote Zone | Up | Troap | 0 | 0.99371747 | 0.416 | 0.881 | 0 |
| ECM Remote Zone | Up | Add2 | 0 | 1.01289185 | 0.355 | 0.941 | 0 |
| ECM Remote Zone | Up | Slc24a1 | 0 | 1.03873577 | 0.302 | 0.559 | 0 |

| | | | | | | | |
|---|---|---|---|---|---|---|---|
| ECM Remote Zone | Up | Hemgn | 0 | 1.05401733 | 0.27 | 0.726 | 0 |
| ECM Remote Zone | Up | Arg1 | 1.07E-87 | 1.09620618 | 0.511 | 0.799 | 2.14E-84 |
| ECM Remote Zone | Up | Prkd3 | 4.24E-172 | 1.1177838 | 0.49 | 0.741 | 8.48E-169 |
| ECM Remote Zone | Up | Cd52 | 0 | 1.129072 | 0.299 | 0.902 | 0 |
| ECM Remote Zone | Up | Tmem86b | 0 | 1.17533747 | 0.111 | 0.361 | 0 |
| ECM Remote Zone | Up | Tlr11 | 1.26E-140 | 1.21110246 | 0.465 | 0.902 | 2.51E-137 |
| ECM Remote Zone | Up | Zfp870 | 0 | 1.21670697 | 0.329 | 0.665 | 0 |
| ECM Remote Zone | Up | Ttll5 | 0 | 1.25108225 | 0.225 | 0.449 | 0 |
| ECM Remote Zone | Up | Setdb2 | 2.85E-209 | 1.27013869 | 0.415 | 0.673 | 5.70E-206 |

| | | | | | | | |
|---|---|---|---|---|---|---|---|
| ECM Remote Zone | Up | Chac1 | 2.23E-190 | 1.36282265 | 0.474 | 0.716 | 4.47E-187 |
| ECM Remote Zone | Up | Caprin2 | 7.66E-295 | 1.37510391 | 0.345 | 0.636 | 1.53E-291 |
| ECM Remote Zone | Up | Alox15 | 0 | 1.67931479 | 0.284 | 0.939 | 0 |
| ECM Remote Zone | Up | Spag5.1 | 0 | 1.72389058 | 0.128 | 0.355 | 0 |
| ECM Remote Zone | Up | Lpin1 | 0 | 1.99245925 | 0.173 | 0.376 | 0 |
| ECM Remote Zone | Up | AABR07068341.1 | 0 | 2.05459946 | 0.217 | 0.472 | 0 |
