## Supplemental Table 12 for "Uncovering the Regional and Cell Specific Bioactivity of Injectable Extracellular Matrix Biomaterials in Myocardial Infarction through Spatial and Single Nucleus Transcriptomics"

**Supplementary Table 12. Coarse Cluster Markers for Chronic Model**

| Cell Type | Gene | p_val | avg_log2FC | pct.1 | pct.2 | p_val_adj |
|---|---|---|---|---|---|---|
| CM | AABR07052585.1 | 0 | 5.26290828 | 0.967 | 0.256 | 0 |
| CM | Tnni3k | 0 | 4.67958481 | 0.929 | 0.237 | 0 |
| CM | Acacb | 0 | 4.39905795 | 0.953 | 0.265 | 0 |
| CM | Obscn | 0 | 4.86967414 | 0.958 | 0.283 | 0 |
| CM | Trim63 | 0 | 4.94616559 | 0.919 | 0.258 | 0 |
| CM | Myh7b | 0 | 4.67789568 | 0.883 | 0.227 | 0 |
| CM | Kcnq1 | 0 | 4.94777898 | 0.949 | 0.295 | 0 |
| CM | Mlip | 0 | 4.125542 | 0.918 | 0.267 | 0 |
| CM | Myocd | 0 | 4.22144727 | 0.904 | 0.28 | 0 |
| CM | Ldb3 | 0 | 4.35872583 | 0.987 | 0.365 | 0 |
| CM | Nexn | 0 | 3.54916915 | 0.982 | 0.361 | 0 |
| CM | Shroom3 | 0 | 4.27656552 | 0.862 | 0.246 | 0 |
| CM | Sgcd | 0 | 2.88842759 | 0.901 | 0.289 | 0 |
| CM | Fhl2 | 0 | 4.48677822 | 0.943 | 0.333 | 0 |
| CM | Actn2 | 0 | 3.989783 | 0.914 | 0.31 | 0 |
| CM | Cdh2 | 0 | 3.61936181 | 0.949 | 0.346 | 0 |
| CM | Mypn | 0 | 4.58855227 | 0.789 | 0.196 | 0 |
| CM | Kcnd3 | 0 | 4.72316014 | 0.892 | 0.305 | 0 |
| CM | Art1 | 0 | 6.0773845 | 0.59 | 0.006 | 0 |
| CM | Ppip5k2 | 0 | 3.4862169 | 0.928 | 0.346 | 0 |
| CM | Trdn | 0 | 3.82030514 | 0.878 | 0.297 | 0 |
| CM | Akap6 | 0 | 3.287206 | 0.96 | 0.38 | 0 |
| CM | Asb2 | 0 | 4.29825476 | 0.887 | 0.311 | 0 |
| CM | Rbm20 | 0 | 5.19388319 | 0.984 | 0.413 | 0 |
| CM | Fgf1 | 0 | 3.75866374 | 0.891 | 0.321 | 0 |

| | | | | | | |
|---|---|---|---|---|---|---|
| CM | Asb18 | 0 | 4.60192203 | 0.792 | 0.223 | 0 |
| CM | Mybpc3 | 0 | 4.81903586 | 0.992 | 0.424 | 0 |
| CM | Kcng2 | 0 | 4.86957562 | 0.788 | 0.22 | 0 |
| CM | Ptpn3 | 0 | 3.36476213 | 0.88 | 0.312 | 0 |
| CM | Myom2 | 0 | 4.18081048 | 0.918 | 0.353 | 0 |
| CM | Rilpl1 | 0 | 3.31175845 | 0.861 | 0.297 | 0 |
| CM | Ryr3 | 0 | 3.3372045 | 0.794 | 0.231 | 0 |
| CM | Coro6 | 0 | 4.39414948 | 0.765 | 0.206 | 0 |
| CM | Mylk3 | 0 | 4.01294403 | 0.841 | 0.283 | 0 |
| CM | Rnf207 | 0 | 4.4258 | 0.766 | 0.218 | 0 |
| CM | Cacnb2 | 0 | 2.72431918 | 0.919 | 0.373 | 0 |
| CM | Fhod3 | 0 | 4.04167212 | 0.977 | 0.433 | 0 |
| CM | Fbxl2 | 0 | 5.32626697 | 0.596 | 0.052 | 0 |
| CM | Macrod1 | 0 | 4.12522757 | 0.877 | 0.343 | 0 |
| CM | Rbfox1 | 0 | 4.09317509 | 0.835 | 0.302 | 0 |
| CM | Myom1 | 0 | 3.18412108 | 0.844 | 0.313 | 0 |
| CM | Sacs | 0 | 2.93685692 | 0.843 | 0.312 | 0 |
| CM | Phf24 | 0 | 4.02632798 | 0.586 | 0.058 | 0 |
| CM | Synpo2 | 0 | 2.54205787 | 0.906 | 0.379 | 0 |
| CM | Casq2 | 0 | 3.77308393 | 0.843 | 0.318 | 0 |
| CM | Dmpk | 0 | 3.08383269 | 0.885 | 0.361 | 0 |
| CM | Srpk3 | 0 | 4.75007771 | 0.7 | 0.179 | 0 |
| CM | Nrap | 0 | 4.20096081 | 0.863 | 0.343 | 0 |
| CM | Slco5a1 | 0 | 4.72778867 | 0.727 | 0.208 | 0 |
| CM | Scn5a | 0 | 4.73466684 | 0.805 | 0.291 | 0 |
| CM | Ivns1abp | 0 | 3.17785056 | 0.918 | 0.404 | 0 |
| CM | Ppargc1b | 0 | 3.25187856 | 0.89 | 0.383 | 0 |
| CM | Speg | 0 | 3.62706607 | 0.781 | 0.274 | 0 |
| CM | Pde4dip | 0 | 4.37264383 | 0.998 | 0.508 | 0 |

| | | | | | | |
|---|---|---|---|---|---|---|
| CM | Fsd2 | 0 | 4.69103899 | 0.734 | 0.246 | 0 |
| CM | Trim55 | 0 | 4.31986603 | 0.787 | 0.3 | 0 |
| CM | AABR07065190.1 | 0 | 4.25176675 | 0.54 | 0.056 | 0 |
| CM | Ccdc141 | 0 | 3.13519618 | 0.953 | 0.469 | 0 |
| CM | Naca | 0 | 3.15522714 | 0.797 | 0.315 | 0 |
| CM | Mtss1 | 0 | 2.41751171 | 0.928 | 0.447 | 0 |
| CM | Cux2 | 0 | 5.28739992 | 0.738 | 0.259 | 0 |
| CM | Alpk3 | 0 | 4.58371535 | 0.826 | 0.349 | 0 |
| CM | Cacna1c | 0 | 2.59572967 | 0.919 | 0.443 | 0 |
| CM | Pln | 0 | 3.11585196 | 0.934 | 0.472 | 0 |
| CM | Tbc1d4 | 0 | 2.37669003 | 0.88 | 0.418 | 0 |
| CM | Gba3 | 0 | 4.0398218 | 0.532 | 0.071 | 0 |
| CM | Rcan2 | 0 | 2.25831742 | 0.944 | 0.484 | 0 |
| CM | Vegfa | 0 | 3.40985267 | 0.992 | 0.533 | 0 |
| CM | Lmo7 | 0 | 2.25377126 | 0.926 | 0.479 | 0 |
| CM | Sypl2 | 0 | 3.66197538 | 0.464 | 0.02 | 0 |
| CM | Myo18b | 0 | 4.72306109 | 0.745 | 0.309 | 0 |
| CM | Apeg3 | 0 | 1.86185512 | 0.426 | 0.001 | 0 |
| CM | Cmya5 | 0 | 2.79560483 | 0.844 | 0.423 | 0 |
| CM | Sorbs2 | 0 | 1.92252148 | 0.933 | 0.521 | 0 |
| CM | Mybpc2 | 0 | 3.24472414 | 0.426 | 0.016 | 0 |
| CM | Hcn4 | 0 | 3.70405277 | 0.48 | 0.072 | 0 |
| CM | Popdc3 | 0 | 7.25281476 | 0.408 | 0 | 0 |
| CM | Atp2a2 | 0 | 2.85570963 | 0.975 | 0.569 | 0 |
| CM | Tnni3 | 0 | 3.65721025 | 0.994 | 0.589 | 0 |
| CM | AABR07031164.1 | 0 | 3.00317405 | 0.4 | 0.001 | 0 |
| CM | Myo3b | 0 | 4.76370157 | 0.436 | 0.041 | 0 |

| | | | | | | |
|---|---|---|---|---|---|---|
| CM | Ryr2 | 0 | 4.57644457 | 0.992 | 0.6 | 0 |
| CM | RGD1565355 | 0 | 2.02380991 | 0.957 | 0.566 | 0 |
| CM | Tnnt2 | 0 | 3.54061006 | 0.994 | 0.604 | 0 |
| CM | Dusp5 | 0 | 3.91160812 | 0.409 | 0.022 | 0 |
| CM | Gja3 | 0 | 4.68486164 | 0.479 | 0.096 | 0 |
| CM | Sorbs1 | 0 | 3.04683374 | 0.988 | 0.621 | 0 |
| CM | Myl3 | 0 | 2.57497478 | 0.913 | 0.552 | 0 |
| CM | AABR07068046.1 | 0 | 4.60747113 | 0.369 | 0.012 | 0 |
| CM | Aspdh | 0 | 4.30130174 | 0.355 | 0.001 | 0 |
| CM | Bcl11b | 0 | 2.07897409 | 0.363 | 0.012 | 0 |
| CM | Tpm1 | 0 | 2.01076141 | 0.962 | 0.622 | 0 |
| CM | AABR07030527.1 | 0 | 4.49087726 | 0.339 | 0 | 0 |
| CM | Ankrd1 | 0 | 2.98368717 | 0.921 | 0.586 | 0 |
| CM | AABR07049033.1 | 0 | 2.06846511 | 0.382 | 0.053 | 0 |
| CM | Slc8a1 | 0 | 2.2503782 | 0.982 | 0.656 | 0 |
| CM | Cacng6 | 0 | 4.49520169 | 0.328 | 0.006 | 0 |
| CM | AC130940.1 | 0 | 5.54202534 | 0.375 | 0.055 | 0 |
| CM | Myh6 | 0 | 3.56286796 | 0.984 | 0.677 | 0 |
| CM | Dmd | 0 | 3.26061014 | 0.99 | 0.706 | 0 |
| CM | Rd3l | 0 | 4.48848797 | 0.296 | 0.012 | 0 |
| CM | AABR07026021.1 | 0 | 4.26171036 | 0.283 | 0.001 | 0 |
| CM | Tnfrsf19 | 2.44E-307 | 0.90000636 | 0.491 | 0.063 | 4.87E-304 |
| CM | Kcnj3 | 4.15E-307 | 3.96231734 | 0.781 | 0.229 | 8.31E-304 |

| | | | | | | |
|---|---|---|---|---|---|---|
| CM | Oxct1 | 6.02E-307 | 2.72341951 | 0.865 | 0.342 | 1.20E-303 |
| CM | AC110709.2 | 3.80E-306 | 4.70225045 | 0.634 | 0.177 | 7.61E-303 |
| CM | Vash2 | 7.22E-306 | 4.69157076 | 0.657 | 0.121 | 1.44E-302 |
| CM | Svil | 1.95E-300 | 2.58691552 | 0.851 | 0.395 | 3.90E-297 |
| CM | Nav2 | 1.68E-299 | 2.19862249 | 0.863 | 0.381 | 3.35E-296 |
| CM | Kcnj12 | 2.30E-299 | 1.70474723 | 0.369 | 0.069 | 4.60E-296 |
| CM | Tesc | 2.79E-299 | 3.87475594 | 0.767 | 0.298 | 5.59E-296 |
| CM | Lrrc2 | 1.32E-297 | 4.07464481 | 0.758 | 0.314 | 2.64E-294 |
| CM | LOC102546683 | 2.33E-291 | 3.55854227 | 0.346 | 0.013 | 4.65E-288 |
| CM | Hs3st5 | 4.60E-290 | 4.07368333 | 0.756 | 0.251 | 9.21E-287 |
| CM | Jph2 | 1.84E-289 | 3.58223497 | 0.751 | 0.245 | 3.68E-286 |
| CM | Parm1 | 5.76E-289 | 3.5925663 | 0.792 | 0.331 | 1.15E-285 |
| CM | Polr2m | 6.77E-289 | 2.44899582 | 0.88 | 0.444 | 1.35E-285 |
| CM | Smyd1 | 1.92E-288 | 4.11273531 | 0.741 | 0.186 | 3.84E-285 |
| CM | Fam228b | 2.13E-288 | 3.48419383 | 0.274 | 0.001 | 4.27E-285 |
| CM | Cacna2d2 | 2.17E-288 | 4.10966837 | 0.613 | 0.139 | 4.34E-285 |
| CM | Myh7 | 1.09E-287 | 3.20588619 | 0.89 | 0.501 | 2.19E-284 |
| CM | Wwc1 | 2.47E-283 | 3.82442348 | 0.46 | 0.105 | 4.93E-280 |
| CM | Ppara | 6.78E-282 | 4.34608197 | 0.716 | 0.166 | 1.36E-278 |
| CM | Klhl40 | 7.22E-282 | 2.91856989 | 0.433 | 0.094 | 1.44E-278 |
| CM | Pla2g7 | 1.79E-280 | 3.63778711 | 0.517 | 0.1 | 3.58E-277 |
| CM | Wnk2 | 2.09E-279 | 4.47277646 | 0.712 | 0.205 | 4.18E-276 |
| CM | Gabrb2 | 3.18E-279 | 3.27544766 | 0.458 | 0.098 | 6.36E-276 |
| CM | Tmem163 | 5.27E-276 | 2.34331848 | 0.455 | 0.066 | 1.05E-272 |
| CM | Efcab6 | 1.68E-274 | 5.25426918 | 0.426 | 0.097 | 3.36E-271 |
| CM | Mapt | 3.24E-274 | 3.4544389 | 0.727 | 0.283 | 6.49E-271 |
| CM | Fbxo40 | 4.79E-274 | 4.45822915 | 0.62 | 0.159 | 9.58E-271 |

| | | | | | | |
|---|---|---|---|---|---|---|
| CM | Kcnk3 | 9.15E-273 | 3.87288334 | 0.542 | 0.091 | 1.83E-269 |
| CM | Ank3 | 1.09E-272 | 1.7834468 | 0.863 | 0.434 | 2.18E-269 |
| CM | Dgkz | 1.57E-272 | 2.41353756 | 0.816 | 0.399 | 3.13E-269 |
| CM | Sptb | 2.52E-272 | 4.41539264 | 0.724 | 0.314 | 5.05E-269 |
| CM | Rasgrp1 | 3.03E-272 | 1.11005557 | 0.335 | 0.013 | 6.07E-269 |
| CM | AABR07025 295.1 | 3.13E-272 | 2.13646904 | 0.88 | 0.411 | 6.25E-269 |
| CM | Slc30a3 | 3.28E-271 | 6.07373852 | 0.499 | 0.155 | 6.56E-268 |
| CM | Atp2b2 | 1.67E-269 | 3.93452851 | 0.688 | 0.251 | 3.35E-266 |
| CM | Fabp3 | 5.43E-269 | 3.27741033 | 0.778 | 0.351 | 1.09E-265 |
| CM | Pde4b | 1.20E-268 | 2.48947022 | 0.849 | 0.331 | 2.40E-265 |
| CM | Xirp2 | 1.79E-267 | 3.12035514 | 0.778 | 0.4 | 3.57E-264 |
| CM | AC111831.1 | 1.74E-266 | 4.19371715 | 0.61 | 0.134 | 3.47E-263 |
| CM | Nebl | 2.94E-266 | 1.45064394 | 0.904 | 0.461 | 5.88E-263 |
| CM | Rgs7 | 6.08E-266 | 2.74850039 | 0.572 | 0.116 | 1.22E-262 |
| CM | Corin | 3.27E-262 | 3.40180519 | 0.761 | 0.289 | 6.55E-259 |
| CM | Nnt | 1.28E-259 | 2.39325246 | 0.847 | 0.41 | 2.56E-256 |
| CM | Ppp1r14c | 2.62E-259 | 3.39805925 | 0.751 | 0.215 | 5.24E-256 |
| CM | Lpar3 | 3.29E-259 | 3.79717632 | 0.478 | 0.119 | 6.59E-256 |
| CM | Oxr1 | 1.46E-258 | 2.33993381 | 0.809 | 0.415 | 2.92E-255 |
| CM | Slc16a10 | 3.92E-257 | 3.40548479 | 0.7 | 0.168 | 7.83E-254 |
| CM | Asb14 | 4.38E-256 | 4.79206213 | 0.592 | 0.218 | 8.76E-253 |
| CM | Pfkfb2 | 1.28E-255 | 3.20719412 | 0.764 | 0.319 | 2.56E-252 |
| CM | Hcn2 | 1.97E-254 | 3.94936688 | 0.428 | 0.103 | 3.94E-251 |
| CM | Arhgap21 | 2.25E-254 | 1.78076374 | 0.894 | 0.525 | 4.49E-251 |
| CM | Pde3a | 2.94E-254 | 1.34811862 | 0.946 | 0.562 | 5.88E-251 |
| CM | Adk | 7.48E-254 | 2.18578334 | 0.858 | 0.405 | 1.50E-250 |
| CM | Trak2 | 1.71E-253 | 2.28851128 | 0.832 | 0.435 | 3.42E-250 |

| | | | | | | |
|---|---|---|---|---|---|---|
| CM | Abcc9 | 5.21E-252 | 1.64748735 | 0.873 | 0.427 | 1.04E-248 |
| CM | Tmem182 | 5.45E-252 | 3.9656024 | 0.73 | 0.238 | 1.09E-248 |
| CM | Neb | 7.16E-250 | 2.23173082 | 0.525 | 0.119 | 1.43E-246 |
| CM | Drc3 | 4.92E-249 | 3.85206009 | 0.518 | 0.078 | 9.85E-246 |
| CM | Dtna | 5.35E-249 | 2.9480021 | 0.763 | 0.278 | 1.07E-245 |
| CM | Alpk2 | 1.76E-248 | 4.45044643 | 0.68 | 0.287 | 3.53E-245 |
| CM | Eya1 | 2.22E-248 | 3.28133217 | 0.608 | 0.15 | 4.43E-245 |
| CM | Tpd52l1 | 2.77E-248 | 3.93389161 | 0.712 | 0.309 | 5.54E-245 |
| CM | Epha3 | 3.56E-248 | 3.26817736 | 0.53 | 0.12 | 7.13E-245 |
| CM | Cacna1h | 6.28E-248 | 1.06364752 | 0.525 | 0.11 | 1.26E-244 |
| CM | AC096301.1 | 1.97E-246 | 4.44384404 | 0.678 | 0.178 | 3.93E-243 |
| CM | Tmem196 | 2.55E-246 | 2.95519351 | 0.722 | 0.254 | 5.11E-243 |
| CM | Ppp1r12b | 1.15E-245 | 1.74057192 | 0.921 | 0.56 | 2.29E-242 |
| CM | Acsl1 | 9.78E-244 | 3.29299288 | 0.737 | 0.363 | 1.96E-240 |
| CM | Inpp4b | 1.52E-243 | 1.79984179 | 0.841 | 0.4 | 3.03E-240 |
| CM | Dhrs7c | 1.54E-241 | 4.66997972 | 0.334 | 0.078 | 3.08E-238 |
| CM | Cap2 | 8.37E-241 | 2.98704023 | 0.734 | 0.264 | 1.67E-237 |
| CM | Tnik | 3.71E-239 | 2.18645068 | 0.834 | 0.446 | 7.42E-236 |
| CM | Ppp2r3a | 5.16E-237 | 1.9766128 | 0.839 | 0.42 | 1.03E-233 |
| CM | Srl | 4.28E-235 | 4.19231835 | 0.642 | 0.177 | 8.57E-232 |
| CM | Rnf144b | 1.79E-233 | 2.67015138 | 0.727 | 0.304 | 3.58E-230 |
| CM | Rrad | 5.17E-233 | 4.27826488 | 0.68 | 0.314 | 1.03E-229 |
| CM | Rap1gap2 | 3.01E-232 | 3.10619876 | 0.721 | 0.309 | 6.02E-229 |
| CM | Abcc8 | 3.04E-232 | 4.61692864 | 0.459 | 0.137 | 6.09E-229 |
| CM | AABR07050449.1 | 4.74E-231 | 3.64782447 | 0.623 | 0.203 | 9.48E-228 |
| CM | Sctr | 2.07E-230 | 3.35249104 | 0.509 | 0.12 | 4.14E-227 |
| CM | Tafa2 | 6.05E-230 | 0.86928325 | 0.438 | 0.05 | 1.21E-226 |

| | | | | | | |
|---|---|---|---|---|---|---|
| CM | Chrna1 | 9.01E-230 | 1.51358769 | 0.32 | 0.001 | 1.80E-226 |
| CM | Pdlim3 | 2.23E-229 | 2.44312114 | 0.741 | 0.321 | 4.45E-226 |
| CM | Esrrg | 8.09E-229 | 2.53170849 | 0.737 | 0.269 | 1.62E-225 |
| CM | Vcl | 1.72E-226 | 2.00344252 | 0.805 | 0.402 | 3.43E-223 |
| CM | Dsp | 2.06E-225 | 2.70325015 | 0.726 | 0.398 | 4.12E-222 |
| CM | AABR07044900.1 | 3.14E-225 | 1.70593355 | 0.838 | 0.381 | 6.28E-222 |
| CM | Cxadr | 1.26E-224 | 4.41201436 | 0.502 | 0.123 | 2.52E-221 |
| CM | Tango2 | 2.35E-224 | 2.53021931 | 0.771 | 0.303 | 4.70E-221 |
| CM | AABR07031740.1 | 9.53E-224 | 3.23097905 | 0.682 | 0.24 | 1.91E-220 |
| CM | Pcdh7 | 2.42E-223 | 3.71120457 | 0.666 | 0.286 | 4.84E-220 |
| CM | Veph1 | 5.83E-223 | 3.93089033 | 0.654 | 0.264 | 1.17E-219 |
| CM | Pik3ap1 | 2.88E-221 | 3.72087596 | 0.615 | 0.169 | 5.77E-218 |
| CM | Ldhb | 9.89E-221 | 2.20910518 | 0.794 | 0.392 | 1.98E-217 |
| CM | Unc45b | 3.99E-220 | 4.83397991 | 0.513 | 0.14 | 7.97E-217 |
| CM | Pank1 | 1.31E-218 | 3.61191407 | 0.691 | 0.295 | 2.62E-215 |
| CM | Ankrd9 | 1.57E-218 | 5.67652395 | 0.459 | 0.157 | 3.14E-215 |
| CM | Fam189a2 | 2.51E-218 | 3.6021555 | 0.542 | 0.101 | 5.01E-215 |
| CM | Adamts20 | 2.94E-218 | 1.17920099 | 0.419 | 0.071 | 5.87E-215 |
| CM | Ube2ql1 | 5.22E-218 | 4.35841809 | 0.489 | 0.099 | 1.04E-214 |
| CM | AABR07007026.1 | 1.21E-217 | 4.2928549 | 0.603 | 0.167 | 2.41E-214 |
| CM | Arhgap26 | 1.22E-216 | 1.83597773 | 0.848 | 0.393 | 2.44E-213 |
| CM | Fbxo32 | 1.35E-216 | 2.56475344 | 0.78 | 0.37 | 2.70E-213 |
| CM | AABR07058170.1 | 1.64E-216 | 3.30747506 | 0.566 | 0.139 | 3.28E-213 |
| CM | AABR07035916.1 | 2.03E-216 | 1.43952747 | 0.936 | 0.581 | 4.07E-213 |

| | | | | | | |
|---|---|---|---|---|---|---|
| CM | Gmpr | 1.15E-215 | 3.60984962 | 0.65 | 0.232 | 2.30E-212 |
| CM | Dpf3 | 7.96E-214 | 3.34825297 | 0.698 | 0.222 | 1.59E-210 |
| CM | Bzw2 | 2.68E-213 | 2.67964872 | 0.723 | 0.332 | 5.37E-210 |
| CM | Limch1 | 4.77E-213 | 1.6344027 | 0.885 | 0.529 | 9.53E-210 |
| CM | Bcl11a | 5.60E-213 | 4.15967858 | 0.563 | 0.223 | 1.12E-209 |
| CM | Adra1a | 1.21E-212 | 2.80694082 | 0.689 | 0.243 | 2.42E-209 |
| CM | Grb14 | 2.67E-210 | 2.58718181 | 0.726 | 0.314 | 5.35E-207 |
| CM | Neurod4 | 4.48E-210 | 3.90326289 | 0.268 | 0 | 8.95E-207 |
| CM | Itga7 | 2.86E-208 | 2.50989404 | 0.697 | 0.322 | 5.72E-205 |
| CM | Casz1 | 2.16E-207 | 2.98455877 | 0.691 | 0.356 | 4.31E-204 |
| CM | Smpx | 1.59E-206 | 3.09256431 | 0.701 | 0.313 | 3.18E-203 |
| CM | Trim7 | 2.79E-206 | 3.32171773 | 0.669 | 0.304 | 5.58E-203 |
| CM | Masp1 | 2.50E-202 | 4.15165818 | 0.482 | 0.128 | 5.00E-199 |
| CM | Trabd2b | 5.96E-202 | 2.24046136 | 0.752 | 0.325 | 1.19E-198 |
| CM | Epb41l4b | 8.22E-201 | 2.80124174 | 0.683 | 0.229 | 1.64E-197 |
| CM | Myoz2 | 3.63E-200 | 3.76006121 | 0.63 | 0.157 | 7.25E-197 |
| CM | Serf1 | 1.36E-199 | 1.50720278 | 0.383 | 0.011 | 2.71E-196 |
| CM | Spsb4 | 4.09E-199 | 3.09026873 | 0.677 | 0.236 | 8.19E-196 |
| CM | Trim54 | 2.45E-198 | 3.6723545 | 0.671 | 0.313 | 4.91E-195 |
| CM | Mitf | 1.33E-196 | 1.92871796 | 0.793 | 0.363 | 2.66E-193 |
| CM | Nfatc2 | 3.71E-196 | 2.73594387 | 0.684 | 0.283 | 7.43E-193 |
| CM | Usp13 | 9.58E-195 | 2.65453499 | 0.72 | 0.372 | 1.92E-191 |
| CM | Me3 | 1.53E-194 | 3.42667591 | 0.66 | 0.319 | 3.06E-191 |
| CM | Ankh | 2.13E-191 | 2.08281024 | 0.774 | 0.398 | 4.27E-188 |
| CM | Ppargc1a | 5.83E-190 | 3.55521373 | 0.615 | 0.251 | 1.17E-186 |
| CM | Plbd1 | 3.04E-189 | 3.97896465 | 0.444 | 0.17 | 6.07E-186 |
| CM | B3galt2 | 6.72E-189 | 3.44632672 | 0.38 | 0.118 | 1.34E-185 |
| CM | Ehd4 | 1.98E-188 | 1.73134014 | 0.793 | 0.368 | 3.96E-185 |
| CM | Phyh | 1.51E-187 | 2.22040063 | 0.76 | 0.397 | 3.02E-184 |

| | | | | | | |
|---|---|---|---|---|---|---|
| CM | AABR07052585.2 | 7.45E-187 | 4.40975211 | 0.632 | 0.34 | 1.49E-183 |
| CM | Txlnb | 1.89E-185 | 3.93130227 | 0.606 | 0.297 | 3.78E-182 |
| CM | Fgf12 | 2.45E-185 | 2.66609448 | 0.708 | 0.227 | 4.91E-182 |
| CM | Fgf13 | 1.66E-183 | 3.90436278 | 0.583 | 0.157 | 3.32E-180 |
| CM | Ky | 8.32E-183 | 4.35655891 | 0.462 | 0.191 | 1.66E-179 |
| CM | Nppb | 8.37E-182 | 3.49006352 | 0.644 | 0.268 | 1.67E-178 |
| CM | Dip2c | 4.12E-181 | 1.79172432 | 0.825 | 0.484 | 8.23E-178 |
| CM | Rxfp1 | 3.54E-180 | 3.93386881 | 0.507 | 0.165 | 7.07E-177 |
| CM | Adcy5 | 6.40E-180 | 1.78069319 | 0.786 | 0.438 | 1.28E-176 |
| CM | Ckmt2 | 2.12E-179 | 2.56649817 | 0.7 | 0.366 | 4.25E-176 |
| CM | Slc38a1 | 1.00E-178 | 3.24146269 | 0.63 | 0.211 | 2.00E-175 |
| CM | Arhgap44 | 5.01E-178 | 2.65811718 | 0.669 | 0.267 | 1.00E-174 |
| CM | Enah | 4.57E-177 | 2.12122235 | 0.728 | 0.287 | 9.14E-174 |
| CM | Samd4a | 5.37E-177 | 1.32324133 | 0.925 | 0.653 | 1.07E-173 |
| CM | Rgs6 | 4.49E-176 | 2.80931755 | 0.623 | 0.241 | 8.99E-173 |
| CM | AABR07049292.1 | 3.53E-175 | 2.81245451 | 0.586 | 0.153 | 7.05E-172 |
| CM | Sgca | 9.12E-174 | 3.80548654 | 0.564 | 0.266 | 1.82E-170 |
| CM | Mpp7 | 1.15E-171 | 1.61589833 | 0.752 | 0.296 | 2.31E-168 |
| CM | Cacna1a | 2.08E-171 | 2.23972792 | 0.74 | 0.381 | 4.17E-168 |
| CM | Fam151a | 8.08E-171 | 1.25940948 | 0.391 | 0.108 | 1.62E-167 |
| CM | Slc12a7 | 6.44E-170 | 2.36881108 | 0.698 | 0.34 | 1.29E-166 |
| CM | Prox1 | 7.18E-170 | 2.19342449 | 0.619 | 0.329 | 1.44E-166 |
| CM | Trim50 | 3.52E-169 | 4.81566788 | 0.414 | 0.146 | 7.04E-166 |
| CM | Mtus2 | 5.41E-169 | 3.02441047 | 0.614 | 0.268 | 1.08E-165 |
| CM | Bdnf | 1.08E-168 | 4.0455149 | 0.55 | 0.165 | 2.16E-165 |
| CM | Adam19 | 3.24E-168 | 1.45706031 | 0.849 | 0.493 | 6.49E-165 |
| CM | Ptgfr | 9.84E-168 | 2.50789223 | 0.567 | 0.131 | 1.97E-164 |

| | | | | | | |
|---|---|---|---|---|---|---|
| CM | Cpe | 2.74E-167 | 3.0141193 | 0.625 | 0.257 | 5.48E-164 |
| CM | Bche | 2.90E-167 | 2.89449212 | 0.577 | 0.218 | 5.79E-164 |
| CM | Kif21a | 5.64E-166 | 3.79519611 | 0.516 | 0.207 | 1.13E-162 |
| CM | AABR07017 268.1 | 1.79E-165 | 4.98806966 | 0.419 | 0.155 | 3.58E-162 |
| CM | Erbb4 | 1.97E-165 | 4.036216 | 0.583 | 0.224 | 3.94E-162 |
| CM | Myl4 | 4.47E-165 | 2.25871506 | 0.35 | 0.083 | 8.94E-162 |
| CM | Angpt1 | 5.72E-165 | 2.29544834 | 0.732 | 0.306 | 1.14E-161 |
| CM | Gpcpd1 | 7.67E-165 | 2.14924803 | 0.739 | 0.385 | 1.53E-161 |
| CM | Optn | 2.59E-164 | 2.39138729 | 0.684 | 0.27 | 5.18E-161 |
| CM | Slc38a3 | 7.59E-164 | 4.39588202 | 0.469 | 0.136 | 1.52E-160 |
| CM | Nt5e | 1.78E-163 | 1.91267963 | 0.697 | 0.258 | 3.55E-160 |
| CM | AABR07001 519.1 | 2.11E-163 | 1.56809304 | 0.811 | 0.41 | 4.21E-160 |
| CM | Lrrc4b | 4.04E-163 | 4.32424781 | 0.511 | 0.183 | 8.08E-160 |
| CM | Atp1b1 | 1.51E-162 | 2.76326468 | 0.632 | 0.239 | 3.01E-159 |
| CM | Pam | 1.94E-162 | 1.42358147 | 0.895 | 0.585 | 3.88E-159 |
| CM | Lck | 2.60E-162 | 0.54354683 | 0.414 | 0.064 | 5.21E-159 |
| CM | Slc4a3 | 2.91E-162 | 2.81399706 | 0.621 | 0.269 | 5.82E-159 |
| CM | Tmod1 | 5.88E-162 | 1.79119764 | 0.725 | 0.288 | 1.18E-158 |
| CM | Pstpip2 | 1.35E-161 | 2.635097 | 0.501 | 0.184 | 2.69E-158 |
| CM | Gadd45b | 3.47E-161 | 1.55959513 | 0.375 | 0.107 | 6.94E-158 |
| CM | AABR07052 523.2 | 1.11E-160 | 5.18530559 | 0.453 | 0.165 | 2.23E-157 |
| CM | AABR07001 573.2 | 1.34E-160 | 0.63801147 | 0.291 | 0.04 | 2.69E-157 |
| CM | Pygm | 1.62E-160 | 3.98821705 | 0.589 | 0.281 | 3.24E-157 |
| CM | Slc20a2 | 2.23E-160 | 2.35166702 | 0.7 | 0.346 | 4.45E-157 |
| CM | Myl2 | 6.50E-160 | 1.71393885 | 0.733 | 0.472 | 1.30E-156 |

| | | | | | | |
|---|---|---|---|---|---|---|
| CM | Man1c1 | 8.77E-160 | 1.67163415 | 0.827 | 0.462 | 1.75E-156 |
| CM | Lsamp | 9.67E-160 | 3.7317692 | 0.536 | 0.164 | 1.93E-156 |
| CM | Adra1b | 9.32E-159 | 4.08636036 | 0.52 | 0.169 | 1.86E-155 |
| CM | AABR07044049.1 | 1.64E-158 | 2.02764037 | 0.714 | 0.328 | 3.28E-155 |
| CM | Scn1a | 2.85E-157 | 3.88304065 | 0.533 | 0.182 | 5.71E-154 |
| CM | LOC100912195 | 1.33E-156 | 2.10756658 | 0.455 | 0.148 | 2.66E-153 |
| CM | Ptprd | 5.63E-154 | 1.49795645 | 0.813 | 0.441 | 1.13E-150 |
| CM | Rimbp2 | 6.08E-153 | 2.29380024 | 0.662 | 0.31 | 1.22E-149 |
| CM | Flnc | 7.38E-153 | 3.57457625 | 0.613 | 0.294 | 1.48E-149 |
| CM | Slc25a20 | 1.27E-152 | 2.64501418 | 0.649 | 0.34 | 2.54E-149 |
| CM | Rbm24 | 1.07E-149 | 3.58257638 | 0.467 | 0.151 | 2.14E-146 |
| CM | Wipf3 | 1.70E-149 | 1.50771474 | 0.746 | 0.368 | 3.39E-146 |
| CM | Gal3st3 | 7.57E-149 | 4.34530115 | 0.499 | 0.202 | 1.51E-145 |
| CM | Lbh | 5.43E-148 | 1.71360489 | 0.757 | 0.423 | 1.09E-144 |
| CM | Cabcoco1 | 2.80E-146 | 3.95981364 | 0.546 | 0.178 | 5.60E-143 |
| CM | Ndrg4 | 4.11E-146 | 3.78929184 | 0.577 | 0.326 | 8.23E-143 |
| CM | Snta1 | 1.66E-145 | 2.41011853 | 0.614 | 0.22 | 3.33E-142 |
| CM | Pdgfc | 2.58E-145 | 0.66563777 | 0.362 | 0.087 | 5.16E-142 |
| CM | Lgr6 | 3.54E-145 | 2.35279549 | 0.657 | 0.355 | 7.08E-142 |
| CM | Rtn4rl1 | 9.91E-145 | 1.8947128 | 0.599 | 0.276 | 1.98E-141 |
| CM | Sphkap | 2.06E-144 | 4.36494994 | 0.467 | 0.158 | 4.12E-141 |
| CM | Rmdn1 | 5.34E-143 | 1.75132128 | 0.758 | 0.353 | 1.07E-139 |
| CM | Ddc | 7.77E-143 | 3.21554652 | 0.549 | 0.243 | 1.55E-139 |
| CM | Samd14 | 3.10E-142 | 1.56201348 | 0.444 | 0.143 | 6.20E-139 |
| CM | Ppp1r3a | 3.71E-142 | 3.644643 | 0.559 | 0.269 | 7.41E-139 |
| CM | Slc16a1 | 3.19E-139 | 2.72142976 | 0.635 | 0.265 | 6.39E-136 |
| CM | Ros1 | 1.43E-138 | 5.02598561 | 0.436 | 0.183 | 2.85E-135 |

| | | | | | | |
|---|---|---|---|---|---|---|
| CM | Samd12 | 5.01E-138 | 1.50018584 | 0.683 | 0.261 | 1.00E-134 |
| CM | Snx31 | 5.41E-138 | 1.15824401 | 0.369 | 0.055 | 1.08E-134 |
| CM | Ckm | 1.55E-137 | 2.7918975 | 0.632 | 0.382 | 3.10E-134 |
| CM | Gja1 | 3.77E-137 | 2.2566954 | 0.623 | 0.286 | 7.54E-134 |
| CM | Coq8a | 7.44E-137 | 3.53537642 | 0.576 | 0.26 | 1.49E-133 |
| CM | AABR07034767.1 | 1.05E-136 | 3.63300113 | 0.525 | 0.21 | 2.09E-133 |
| CM | Lhfpl2 | 7.12E-136 | 2.11706805 | 0.671 | 0.359 | 1.42E-132 |
| CM | Adhfe1 | 1.51E-135 | 3.67915467 | 0.521 | 0.257 | 3.03E-132 |
| CM | Pdlim5 | 2.92E-134 | 1.14511307 | 0.865 | 0.614 | 5.85E-131 |
| CM | AABR07031399.1 | 3.04E-134 | 1.87231929 | 0.649 | 0.305 | 6.09E-131 |
| CM | Lrrtm3 | 4.10E-134 | 1.41792978 | 0.547 | 0.224 | 8.20E-131 |
| CM | Pde7b | 5.53E-134 | 1.03111027 | 0.858 | 0.528 | 1.11E-130 |
| CM | Ablim2 | 3.87E-133 | 2.91273903 | 0.509 | 0.235 | 7.75E-130 |
| CM | Anks1b | 9.49E-133 | 2.6655668 | 0.501 | 0.155 | 1.90E-129 |
| CM | Popdc2 | 1.02E-132 | 3.63294348 | 0.546 | 0.258 | 2.04E-129 |
| CM | Gata4 | 3.04E-131 | 1.27227489 | 0.803 | 0.443 | 6.08E-128 |
| CM | Gnao1 | 7.17E-131 | 1.59314523 | 0.735 | 0.418 | 1.43E-127 |
| CM | Pcbp3 | 3.65E-130 | 2.25206096 | 0.63 | 0.332 | 7.31E-127 |
| CM | Art3 | 6.64E-130 | 2.27897186 | 0.606 | 0.272 | 1.33E-126 |
| CM | Usp2 | 7.20E-130 | 3.49401592 | 0.5 | 0.247 | 1.44E-126 |
| CM | Rxrg | 7.30E-129 | 1.90937102 | 0.57 | 0.309 | 1.46E-125 |
| CM | Gria3 | 1.46E-128 | 1.1658 | 0.837 | 0.444 | 2.92E-125 |
| CM | Lrrc7 | 5.69E-128 | 3.52749381 | 0.545 | 0.209 | 1.14E-124 |
| CM | Ppm1l | 4.07E-125 | 2.31026472 | 0.607 | 0.262 | 8.14E-122 |
| CM | Zfp622 | 2.48E-124 | 1.35013393 | 0.739 | 0.362 | 4.97E-121 |
| CM | Slc22a23 | 9.70E-123 | 2.34571272 | 0.647 | 0.375 | 1.94E-119 |
| CM | Prkaa2 | 3.74E-122 | 2.70789179 | 0.552 | 0.259 | 7.47E-119 |

| | | | | | | |
|---|---|---|---|---|---|---|
| CM | Frmd5 | 7.07E-121 | 2.6876559 | 0.539 | 0.212 | 1.41E-117 |
| CM | Hspb8 | 1.56E-120 | 2.18507315 | 0.605 | 0.301 | 3.13E-117 |
| CM | Ednra | 9.69E-120 | 1.34744891 | 0.726 | 0.399 | 1.94E-116 |
| CM | Glb1l2 | 2.21E-119 | 4.21065221 | 0.483 | 0.23 | 4.41E-116 |
| CM | Fry | 3.62E-118 | 0.89715249 | 0.902 | 0.606 | 7.25E-115 |
| CM | Ppm1e | 5.81E-117 | 2.08792176 | 0.579 | 0.293 | 1.16E-113 |
| CM | Actc1 | 1.55E-115 | 1.67292016 | 0.657 | 0.374 | 3.09E-112 |
| CM | Homer1 | 9.38E-115 | 2.02701536 | 0.601 | 0.261 | 1.88E-111 |
| CM | Sfxn5 | 8.11E-113 | 2.89659042 | 0.562 | 0.25 | 1.62E-109 |
| CM | Sema5a | 1.68E-111 | 1.46081868 | 0.666 | 0.311 | 3.36E-108 |
| CM | Msrb3 | 1.41E-110 | 1.24293169 | 0.759 | 0.42 | 2.82E-107 |
| CM | Nuak1 | 8.08E-108 | 0.8224197 | 0.754 | 0.459 | 1.62E-104 |
| CM | Fign | 1.21E-105 | 1.31040036 | 0.577 | 0.229 | 2.42E-102 |
| CM | Sh3kbp1 | 3.50E-103 | 1.5970939 | 0.63 | 0.363 | 7.00E-100 |
| CM | Shb | 1.65E-102 | 1.69500509 | 0.653 | 0.403 | 3.29E-99 |
| CM | Mb | 1.96E-100 | 0.9431554 | 0.838 | 0.554 | 3.92E-97 |
| CM | Efhd1 | 2.08E-100 | 1.26077882 | 0.525 | 0.197 | 4.17E-97 |
| CM | Iqgap2 | 2.30E-98 | 1.03628092 | 0.579 | 0.248 | 4.59E-95 |
| CM | Ptpn13 | 2.57E-98 | 1.62610948 | 0.596 | 0.267 | 5.13E-95 |
| CM | Chst15 | 3.72E-98 | 1.2514686 | 0.571 | 0.263 | 7.45E-95 |
| CM | Tox3 | 3.78E-97 | 1.74216547 | 0.577 | 0.318 | 7.56E-94 |
| CM | Crim1 | 1.99E-95 | 0.71637449 | 0.851 | 0.587 | 3.98E-92 |
| CM | Dlgap1 | 8.43E-95 | 1.17632271 | 0.666 | 0.386 | 1.69E-91 |
| CM | Clic5 | 1.20E-92 | 0.72748266 | 0.788 | 0.501 | 2.40E-89 |
| CM | Palm2 | 1.74E-92 | 0.77550232 | 0.7 | 0.364 | 3.47E-89 |
| CM | Ikzf2 | 2.02E-91 | 0.68820808 | 0.603 | 0.28 | 4.03E-88 |
| CM | Gbe1 | 2.80E-91 | 1.15337715 | 0.698 | 0.386 | 5.60E-88 |
| CM | Clybl | 1.04E-90 | 1.14818215 | 0.672 | 0.389 | 2.08E-87 |
| CM | Tspan18 | 2.57E-90 | 1.14821043 | 0.692 | 0.378 | 5.13E-87 |

| | | | | | | |
|---|---|---|---|---|---|---|
| CM | Tec | 2.34E-88 | 0.48056357 | 0.523 | 0.187 | 4.67E-85 |
| CM | Abcb4 | 6.04E-88 | 1.19361751 | 0.625 | 0.324 | 1.21E-84 |
| CM | Egln3 | 8.32E-86 | 1.55417903 | 0.572 | 0.285 | 1.66E-82 |
| CM | Itga9 | 1.11E-85 | 1.00142892 | 0.826 | 0.546 | 2.21E-82 |
| CM | Lrrc10 | 2.04E-85 | 2.86950507 | 0.398 | 0.141 | 4.08E-82 |
| CM | Ralgapa2 | 7.77E-85 | 0.67856851 | 0.817 | 0.53 | 1.55E-81 |
| CM | Tmem51 | 9.33E-82 | 0.66862243 | 0.564 | 0.306 | 1.87E-78 |
| CM | Ccdc3 | 3.12E-81 | 0.30769142 | 0.53 | 0.227 | 6.23E-78 |
| CM | Me1 | 4.29E-81 | 0.61473283 | 0.618 | 0.272 | 8.58E-78 |
| CM | Gramd1b | 2.41E-78 | 1.16511474 | 0.545 | 0.234 | 4.83E-75 |
| CM | Eva1c | 7.49E-78 | 1.2298206 | 0.583 | 0.296 | 1.50E-74 |
| CM | Col24a1 | 8.09E-78 | 0.35205374 | 0.591 | 0.324 | 1.62E-74 |
| CM | Csrp2 | 2.92E-77 | 0.4394524 | 0.521 | 0.252 | 5.84E-74 |
| CM | Sntb1 | 1.68E-75 | 1.36954782 | 0.569 | 0.312 | 3.35E-72 |
| CM | Lekr1 | 1.29E-74 | 1.05702644 | 0.579 | 0.275 | 2.58E-71 |
| CM | Ntn4 | 1.69E-74 | 0.51822196 | 0.698 | 0.442 | 3.37E-71 |
| CM | Esr1 | 1.10E-73 | 1.69182462 | 0.486 | 0.194 | 2.20E-70 |
| CM | Nrxn3 | 1.10E-70 | 0.3530738 | 0.513 | 0.199 | 2.19E-67 |
| CM | Mt1 | 4.70E-69 | 0.93556465 | 0.348 | 0.059 | 9.41E-66 |
| CM | Man1a1 | 3.02E-68 | 0.59686103 | 0.756 | 0.421 | 6.04E-65 |
| CM | Sox6 | 1.97E-63 | 0.44907959 | 0.81 | 0.542 | 3.93E-60 |
| CM | Foxp2 | 1.97E-63 | 0.58820487 | 0.644 | 0.34 | 3.94E-60 |
| CM | AC134204.1 | 7.50E-56 | 0.43713783 | 0.644 | 0.354 | 1.50E-52 |
| CM | Fhad1 | 5.84E-51 | 0.64801382 | 0.348 | 0.095 | 1.17E-47 |
| CM | Stxbp6 | 2.23E-43 | 0.38390274 | 0.779 | 0.5 | 4.45E-40 |
| EC | Cyyr1 | 0 | 3.40827821 | 0.949 | 0.499 | 0 |
| EC | Ccdc85a | 0 | 2.98711333 | 0.868 | 0.421 | 0 |
| EC | Ptprb | 0 | 3.08936996 | 0.868 | 0.425 | 0 |

| | | | | | | |
|---|---|---|---|---|---|---|
| EC | Adgrf5 | 0 | 3.19640165 | 0.923 | 0.487 | 0 |
| EC | Prkch | 0 | 3.11060229 | 0.854 | 0.426 | 0 |
| EC | Shank3 | 0 | 3.10584289 | 0.901 | 0.481 | 0 |
| EC | Flt1 | 0 | 3.31103311 | 0.863 | 0.456 | 0 |
| EC | Etl4 | 0 | 2.78385755 | 0.957 | 0.584 | 0 |
| EC | Arhgap31 | 0 | 2.8419961 | 0.938 | 0.572 | 0 |
| EC | Dach1 | 0 | 3.19107204 | 0.96 | 0.603 | 0 |
| EC | Itpkb | 0 | 2.70653495 | 0.902 | 0.57 | 0 |
| EC | Cdh13 | 0 | 2.19247113 | 0.97 | 0.718 | 0 |
| EC | Fli1 | 5.16E-300 | 2.51142441 | 0.896 | 0.579 | 1.03E-296 |
| EC | Kitlg | 5.25E-267 | 3.04870627 | 0.843 | 0.454 | 1.05E-263 |
| EC | AABR07001734.1 | 7.55E-255 | 2.53540035 | 0.337 | 0.026 | 1.51E-251 |
| EC | Adgrl4 | 1.39E-243 | 2.91854621 | 0.8 | 0.397 | 2.77E-240 |
| EC | Dnm3 | 8.34E-242 | 2.46283789 | 0.863 | 0.509 | 1.67E-238 |
| EC | Egfl7 | 2.15E-224 | 2.75114105 | 0.795 | 0.456 | 4.30E-221 |
| EC | Ablim3 | 1.05E-220 | 3.17919366 | 0.76 | 0.396 | 2.09E-217 |
| EC | Epas1 | 1.05E-211 | 2.3597338 | 0.852 | 0.577 | 2.10E-208 |
| EC | Myrip | 1.79E-198 | 3.37745361 | 0.721 | 0.369 | 3.57E-195 |
| EC | Emcn | 5.13E-197 | 2.59111352 | 0.761 | 0.443 | 1.03E-193 |
| EC | Eng | 8.62E-194 | 2.15835435 | 0.839 | 0.554 | 1.72E-190 |
| EC | AABR07007642.1 | 3.14E-187 | 3.58522292 | 0.719 | 0.341 | 6.29E-184 |
| EC | Cd109 | 6.93E-186 | 3.22638311 | 0.385 | 0.064 | 1.39E-182 |
| EC | Mcf2l | 1.51E-180 | 3.26756702 | 0.698 | 0.32 | 3.03E-177 |
| EC | Dll1 | 8.57E-176 | 3.7445655 | 0.373 | 0.089 | 1.71E-172 |
| EC | Lrg1 | 5.79E-175 | 2.65922381 | 0.383 | 0.055 | 1.16E-171 |
| EC | Myo10 | 5.41E-170 | 3.00825214 | 0.741 | 0.443 | 1.08E-166 |
| EC | Podxl | 1.92E-165 | 3.18715867 | 0.704 | 0.343 | 3.83E-162 |

| | | | | | | |
|---|---|---|---|---|---|---|
| EC | Eepd1 | 7.97E-151 | 2.73484405 | 0.723 | 0.428 | 1.59E-147 |
| EC | Akr1c15 | 2.26E-140 | 2.492418 | 0.698 | 0.381 | 4.52E-137 |
| EC | Meox2 | 1.15E-138 | 2.68624664 | 0.671 | 0.414 | 2.30E-135 |
| EC | Rasa4 | 1.23E-137 | 3.23336327 | 0.683 | 0.312 | 2.46E-134 |
| EC | Prss12 | 1.60E-133 | 0.89953087 | 0.275 | 0.02 | 3.19E-130 |
| EC | Kdr | 1.83E-133 | 2.72677463 | 0.709 | 0.408 | 3.67E-130 |
| EC | Fabp4 | 1.43E-130 | 3.09219394 | 0.657 | 0.397 | 2.86E-127 |
| EC | Sox17 | 7.94E-129 | 3.06325997 | 0.436 | 0.113 | 1.59E-125 |
| EC | Palmd | 2.18E-128 | 2.68864319 | 0.66 | 0.362 | 4.36E-125 |
| EC | Cadps2 | 3.47E-128 | 3.14565366 | 0.653 | 0.367 | 6.93E-125 |
| EC | Hmcn1 | 7.98E-122 | 1.77431763 | 0.728 | 0.448 | 1.60E-118 |
| EC | Mgmt | 3.41E-118 | 2.08185433 | 0.734 | 0.426 | 6.82E-115 |
| EC | Aqp1 | 1.82E-115 | 3.21012426 | 0.616 | 0.284 | 3.63E-112 |
| EC | Spns2 | 7.71E-107 | 2.77184706 | 0.581 | 0.234 | 1.54E-103 |
| EC | Cxcl12 | 2.06E-105 | 2.77269029 | 0.636 | 0.374 | 4.13E-102 |
| EC | Lcp1 | 5.21E-105 | 0.49408983 | 0.416 | 0.162 | 1.04E-101 |
| EC | Rsad2 | 1.17E-103 | 2.8827225 | 0.43 | 0.175 | 2.34E-100 |
| EC | Slc27a2 | 9.27E-99 | 2.06249447 | 0.277 | 0.023 | 1.85E-95 |
| EC | Cdk19 | 1.09E-95 | 2.06976715 | 0.719 | 0.444 | 2.19E-92 |
| EC | E2f7 | 5.54E-94 | 0.2678633 | 0.339 | 0.08 | 1.11E-90 |
| EC | Thsd7a | 6.26E-93 | 3.12978238 | 0.608 | 0.291 | 1.25E-89 |
| EC | Notch4 | 7.02E-90 | 3.20662668 | 0.54 | 0.276 | 1.40E-86 |
| EC | Nrp2 | 1.04E-80 | 2.0488151 | 0.628 | 0.363 | 2.07E-77 |
| EC | Wt1 | 1.55E-78 | 3.5756453 | 0.465 | 0.182 | 3.11E-75 |
| EC | Sox13 | 3.33E-63 | 1.8987993 | 0.606 | 0.34 | 6.67E-60 |
| EC | Tmem26 | 6.90E-48 | 2.2120137 | 0.435 | 0.173 | 1.38E-44 |
| Endocardiac Cells | Il1b | 5.88E-202 | 4.03272644 | 0.272 | 0.004 | 1.18E-198 |

| | | | | | | |
|---|---|---|---|---|---|---|
| Endocardiac Cells | Cgnl1 | 5.70E-99 | 5.92146474 | 0.986 | 0.39 | 1.14E-95 |
| Endocardiac Cells | LOC108348771 | 2.16E-81 | 3.61832078 | 0.259 | 0.004 | 4.33E-78 |
| Endocardiac Cells | Pkhd1l1 | 1.03E-77 | 4.14578912 | 0.946 | 0.43 | 2.06E-74 |
| Endocardiac Cells | Hmcn11 | 1.30E-77 | 3.91492797 | 0.959 | 0.495 | 2.59E-74 |
| Endocardiac Cells | LOC691995 | 1.23E-72 | 5.62257915 | 0.34 | 0.037 | 2.45E-69 |
| Endocardiac Cells | Tmem45b | 1.77E-66 | 2.84244662 | 0.272 | 0.008 | 3.54E-63 |
| Endocardiac Cells | Mest | 5.61E-60 | 6.146713 | 0.313 | 0.035 | 1.12E-56 |
| Endocardiac Cells | Ltbp1 | 1.75E-59 | 3.32087561 | 0.918 | 0.613 | 3.50E-56 |
| Endocardiac Cells | Nrg1 | 3.25E-59 | 7.23542098 | 0.844 | 0.4 | 6.50E-56 |
| Endocardiac Cells | Cdh11 | 1.83E-55 | 3.71418922 | 0.884 | 0.448 | 3.66E-52 |
| Endocardiac Cells | Gmds | 5.41E-54 | 2.5597729 | 0.939 | 0.633 | 1.08E-50 |
| Endocardiac Cells | Vwf | 9.96E-51 | 3.60587028 | 0.844 | 0.438 | 1.99E-47 |
| Endocardiac Cells | Pgm5 | 5.07E-48 | 3.91140474 | 0.837 | 0.479 | 1.01E-44 |
| Endocardiac Cells | Wnt9b | 1.19E-46 | 3.51783497 | 0.313 | 0.034 | 2.39E-43 |

| | | | | | | |
|---|---|---|---|---|---|---|
| Endocardiac Cells | Slc9a9 | 6.57E-41 | 2.73203949 | 0.837 | 0.447 | 1.31E-37 |
| Endocardiac Cells | Bmp6 | 2.30E-40 | 3.53535335 | 0.83 | 0.551 | 4.60E-37 |
| Endocardiac Cells | Chn2 | 3.38E-38 | 3.07153299 | 0.789 | 0.424 | 6.77E-35 |
| Endocardiac Cells | Gpm6a | 1.01E-37 | 2.89406977 | 0.837 | 0.539 | 2.02E-34 |
| Endocardiac Cells | Smoc1 | 9.60E-35 | 6.40192799 | 0.605 | 0.216 | 1.92E-31 |
| Endocardiac Cells | Eng1 | 1.92E-32 | 1.72897178 | 0.878 | 0.607 | 3.84E-29 |
| Endocardiac Cells | Zmat4 | 1.22E-31 | 6.67190817 | 0.612 | 0.218 | 2.45E-28 |
| Endocardiac Cells | Klhl29 | 3.38E-25 | 2.96182261 | 0.714 | 0.437 | 6.75E-22 |
| Endocardiac Cells | Gria2 | 5.87E-25 | 4.13145201 | 0.354 | 0.061 | 1.17E-21 |
| Endocardiac Cells | Sncaip | 3.93E-24 | 3.61376259 | 0.646 | 0.387 | 7.86E-21 |
| Endocardiac Cells | RGD1305807 | 2.55E-22 | 2.90377791 | 0.299 | 0.046 | 5.11E-19 |
| Endocardiac Cells | Npr3 | 2.60E-21 | 5.29603131 | 0.619 | 0.261 | 5.19E-18 |
| Endocardiac Cells | Emcn1 | 4.15E-21 | 2.13195642 | 0.769 | 0.503 | 8.29E-18 |
| Endocardiac Cells | Grm3 | 6.97E-19 | 3.20342417 | 0.299 | 0.028 | 1.39E-15 |

| | | | | | | |
|---|---|---|---|---|---|---|
| Endocardia c Cells | AABR07016 779.1 | 1.28E-14 | 1.95579288 | 0.299 | 0.045 | 2.57E-11 |
| Endocardia c Cells | LOC100911 486 | 1.77E-14 | 4.05213601 | 0.408 | 0.12 | 3.55E-11 |
| Endocardia c Cells | Tp53inp2 | 2.02E-09 | 1.71357567 | 0.082 | 0.403 | 4.04E-06 |
| Endocardia c Cells | Pfkfb3 | 4.36E-08 | 0.34272862 | 0.129 | 0.389 | 8.72E-05 |
| Endocardia c Cells | Ppp1r3c | 7.16E-08 | 3.65941198 | 0.034 | 0.303 | 0.00014318 |
| Endocardia c Cells | Gng10 | 6.40E-07 | 1.21376854 | 0.054 | 0.313 | 0.00128051 |
| Endocardia c Cells | LOC691083 | 2.34E-06 | 0.43754074 | 0.122 | 0.392 | 0.00468956 |
| Endocardia c Cells | LOC100911 847 | 2.87E-06 | 0.30629515 | 0.184 | 0.434 | 0.00574977 |
| Endocardia c Cells | Mamdc2 | 1.91E-05 | 0.56927861 | 0.156 | 0.441 | 0.0381729 |
| Fibs | C1qtnf7 | 0 | 3.45209933 | 0.893 | 0.444 | 0 |
| Fibs | C7 | 0 | 3.46621185 | 0.765 | 0.335 | 0 |
| Fibs | Tmeff2 | 0 | 3.47175249 | 0.81 | 0.413 | 0 |
| Fibs | Itgbl1 | 0 | 3.34154457 | 0.848 | 0.452 | 0 |
| Fibs | Ebf2 | 0 | 3.10335276 | 0.914 | 0.532 | 0 |
| Fibs | Zfp385d | 0 | 2.50581063 | 0.889 | 0.511 | 0 |
| Fibs | Gpc6 | 0 | 2.64914093 | 0.972 | 0.615 | 0 |
| Fibs | Col3a1 | 0 | 2.94588617 | 0.875 | 0.52 | 0 |
| Fibs | Bicc1 | 3.01E-307 | 2.29738479 | 0.86 | 0.581 | 6.02E-304 |
| Fibs | Dcn | 1.62E-303 | 3.13438247 | 0.792 | 0.488 | 3.25E-300 |
| Fibs | Trps1 | 1.90E-300 | 2.19849092 | 0.873 | 0.588 | 3.80E-297 |

| | | | | | | |
|---|---|---|---|---|---|---|
| Fibs | Cpq | 1.97E-297 | 3.21561991 | 0.766 | 0.314 | 3.94E-294 |
| Fibs | Fbn1 | 3.11E-297 | 2.88619029 | 0.813 | 0.488 | 6.23E-294 |
| Fibs | AC111804.2 | 3.13E-276 | 4.10358509 | 0.31 | 0.011 | 6.27E-273 |
| Fibs | Pdgfra | 2.18E-251 | 3.53593721 | 0.697 | 0.321 | 4.36E-248 |
| Fibs | Plod2 | 1.63E-244 | 2.51919483 | 0.784 | 0.503 | 3.25E-241 |
| Fibs | Gsn | 2.43E-239 | 2.81333668 | 0.765 | 0.453 | 4.85E-236 |
| Fibs | Adamtsl3 | 1.77E-222 | 2.92162534 | 0.743 | 0.462 | 3.55E-219 |
| Fibs | Adamts2 | 3.96E-222 | 2.74465908 | 0.711 | 0.422 | 7.92E-219 |
| Fibs | Clmp | 2.40E-220 | 3.6152878 | 0.674 | 0.327 | 4.80E-217 |
| Fibs | B4galnt3 | 8.56E-218 | 3.47190347 | 0.305 | 0.017 | 1.71E-214 |
| Fibs | Zeb2 | 2.68E-217 | 1.29415941 | 0.861 | 0.556 | 5.36E-214 |
| Fibs | Egr3 | 2.44E-216 | 2.95863735 | 0.309 | 0.02 | 4.88E-213 |
| Fibs | Dapk1 | 1.19E-215 | 2.50631725 | 0.752 | 0.394 | 2.38E-212 |
| Fibs | Abca8a | 4.29E-213 | 2.18181646 | 0.758 | 0.495 | 8.59E-210 |
| Fibs | Dcst2 | 2.23E-209 | 2.98169013 | 0.254 | 0.004 | 4.46E-206 |
| Fibs | Nrxn1 | 2.48E-209 | 2.82955889 | 0.781 | 0.514 | 4.96E-206 |
| Fibs | Ccdc80 | 7.13E-209 | 2.94450203 | 0.695 | 0.407 | 1.43E-205 |
| Fibs | Axl | 4.86E-208 | 3.04513649 | 0.682 | 0.384 | 9.72E-205 |
| Fibs | Col14a1 | 2.85E-206 | 3.49070689 | 0.68 | 0.403 | 5.70E-203 |
| Fibs | Fbln1 | 1.19E-202 | 3.64927229 | 0.676 | 0.259 | 2.38E-199 |
| Fibs | AABR07028 499.1 | 2.54E-199 | 5.41851474 | 0.278 | 0.017 | 5.07E-196 |
| Fibs | Pid1 | 5.12E-187 | 1.95651979 | 0.741 | 0.415 | 1.02E-183 |
| Fibs | Opcml | 5.69E-176 | 3.37260837 | 0.662 | 0.397 | 1.14E-172 |
| Fibs | Dclk1 | 7.01E-175 | 2.82633247 | 0.67 | 0.356 | 1.40E-171 |
| Fibs | Pde1a | 1.18E-174 | 2.83758056 | 0.679 | 0.401 | 2.36E-171 |
| Fibs | Mrc2 | 3.79E-174 | 3.89037855 | 0.627 | 0.354 | 7.58E-171 |
| Fibs | Aldh1a3 | 3.66E-166 | 3.87108638 | 0.421 | 0.12 | 7.32E-163 |

| | | | | | | |
|---|---|---|---|---|---|---|
| Fibs | Stk32b | 7.01E-165 | 1.40125734 | 0.387 | 0.088 | 1.40E-161 |
| Fibs | Pdpn | 1.99E-164 | 1.57256081 | 0.37 | 0.1 | 3.99E-161 |
| Fibs | Sntg2 | 1.91E-162 | 3.10530842 | 0.357 | 0.105 | 3.81E-159 |
| Fibs | Cxcl1 | 2.92E-160 | 1.44705142 | 0.349 | 0.063 | 5.84E-157 |
| Fibs | Crlf1 | 2.78E-159 | 3.25436204 | 0.453 | 0.182 | 5.55E-156 |
| Fibs | Robo1 | 1.03E-153 | 3.23177382 | 0.631 | 0.344 | 2.05E-150 |
| Fibs | Adgrd1 | 5.43E-153 | 3.90113451 | 0.563 | 0.256 | 1.09E-149 |
| Fibs | Has2 | 2.26E-148 | 3.83874763 | 0.413 | 0.139 | 4.51E-145 |
| Fibs | Ar | 1.87E-146 | 2.96675648 | 0.638 | 0.35 | 3.73E-143 |
| Fibs | Necab1 | 5.55E-145 | 3.19522272 | 0.395 | 0.088 | 1.11E-141 |
| Fibs | Vcan | 2.05E-144 | 3.12116958 | 0.601 | 0.334 | 4.10E-141 |
| Fibs | Gda | 1.24E-143 | 2.20442615 | 0.657 | 0.394 | 2.48E-140 |
| Fibs | AABR07003304.2 | 4.57E-141 | 2.18521595 | 0.335 | 0.053 | 9.15E-138 |
| Fibs | Npffr2 | 9.85E-140 | 3.91856328 | 0.402 | 0.135 | 1.97E-136 |
| Fibs | Htra3 | 5.84E-138 | 3.37242503 | 0.594 | 0.245 | 1.17E-134 |
| Fibs | Egfr | 3.06E-137 | 3.43484944 | 0.579 | 0.249 | 6.13E-134 |
| Fibs | AABR07001068.1 | 2.55E-134 | 3.66867124 | 0.43 | 0.138 | 5.10E-131 |
| Fibs | Ndst3 | 1.46E-133 | 1.89342584 | 0.417 | 0.148 | 2.93E-130 |
| Fibs | Uap1 | 4.79E-133 | 2.99936535 | 0.627 | 0.342 | 9.59E-130 |
| Fibs | Cacnb4 | 4.39E-129 | 1.76087941 | 0.368 | 0.091 | 8.77E-126 |
| Fibs | Tll2 | 6.09E-129 | 2.62318812 | 0.395 | 0.118 | 1.22E-125 |
| Fibs | Igfbp6 | 1.23E-127 | 2.60911953 | 0.438 | 0.172 | 2.46E-124 |
| Fibs | Adcy3 | 1.63E-127 | 1.24053034 | 0.414 | 0.162 | 3.27E-124 |
| Fibs | Sdc2 | 1.60E-121 | 2.43170555 | 0.666 | 0.413 | 3.21E-118 |
| Fibs | Tnnt3 | 4.56E-119 | 3.47698945 | 0.555 | 0.232 | 9.12E-116 |
| Fibs | Gfpt2 | 3.83E-118 | 3.73774211 | 0.524 | 0.215 | 7.66E-115 |
| Fibs | Efemp1 | 9.83E-116 | 2.93396167 | 0.44 | 0.16 | 1.97E-112 |

| | | | | | | |
|---|---|---|---|---|---|---|
| Fibs | Enpp2 | 1.64E-114 | 3.83804032 | 0.541 | 0.186 | 3.29E-111 |
| Fibs | RGD1563354 | 8.02E-114 | 3.67559055 | 0.533 | 0.264 | 1.60E-110 |
| Fibs | Ltbp2 | 2.04E-112 | 3.40105993 | 0.518 | 0.246 | 4.08E-109 |
| Fibs | Pi16 | 1.06E-107 | 3.2191537 | 0.582 | 0.3 | 2.11E-104 |
| Fibs | Col6a6 | 4.09E-102 | 3.49055366 | 0.513 | 0.258 | 8.17E-99 |
| Fibs | Scara5 | 8.45E-90 | 3.63754263 | 0.5 | 0.247 | 1.69E-86 |
| Fibs | Pappa1 | 3.93E-76 | 1.77865174 | 0.396 | 0.121 | 7.86E-73 |
| Fibs | Rab27b | 1.82E-63 | 1.14960568 | 0.391 | 0.128 | 3.65E-60 |
| Immune Cells | LOC102549869 | 0 | 8.25639011 | 0.457 | 0.002 | 0 |
| Immune Cells | AABR07003235.1 | 0 | 10.7827787 | 0.414 | 0 | 0 |
| Immune Cells | Clec4d | 0 | 6.63440644 | 0.405 | 0.003 | 0 |
| Immune Cells | Clec7a | 0 | 6.87841913 | 0.31 | 0.001 | 0 |
| Immune Cells | AABR07035722.1 | 0 | 11.2784487 | 0.293 | 0 | 0 |
| Immune Cells | RGD1560281 | 0 | 8.61126823 | 0.284 | 0 | 0 |
| Immune Cells | AABR07043667.1 | 0 | 6.65166458 | 0.284 | 0.001 | 0 |
| Immune Cells | Mmp8 | 0 | 7.00325211 | 0.276 | 0 | 0 |
| Immune Cells | Emb | 0 | 6.2843276 | 0.267 | 0.001 | 0 |
| Immune Cells | AABR07052608.1 | 3.09E-260 | 7.06375635 | 0.302 | 0.003 | 6.18E-257 |

| | | | | | | |
|---|---|---|---|---|---|---|
| Immune Cells | Ccr1 | 8.65E-248 | 8.52709705 | 0.44 | 0.001 | 1.73E-244 |
| Immune Cells | Tyrobp | 4.17E-241 | 7.32218225 | 0.483 | 0.015 | 8.33E-238 |
| Immune Cells | Spn | 6.11E-229 | 5.07716385 | 0.302 | 0.001 | 1.22E-225 |
| Immune Cells | Mmp12 | 3.40E-214 | 9.90338944 | 0.414 | 0.002 | 6.81E-211 |
| Immune Cells | Pipox | 1.59E-192 | 4.86531206 | 0.371 | 0.001 | 3.18E-189 |
| Immune Cells | AABR07009154.1 | 5.56E-184 | 3.84560109 | 0.267 | 0.001 | 1.11E-180 |
| Immune Cells | Asgr2 | 2.94E-169 | 8.14364409 | 0.509 | 0.019 | 5.87E-166 |
| Immune Cells | Arhgap30 | 1.03E-144 | 6.24324238 | 0.414 | 0.017 | 2.06E-141 |
| Immune Cells | Hmgb3 | 1.04E-136 | 1.60974698 | 0.328 | 0.012 | 2.09E-133 |
| Immune Cells | Fam205a1 | 1.50E-123 | 0.76217868 | 0.431 | 0.015 | 3.01E-120 |
| Immune Cells | Cx3cr11 | 1.50E-120 | 7.45385662 | 0.491 | 0.015 | 3.00E-117 |
| Immune Cells | Cass4 | 1.13E-98 | 5.71890982 | 0.293 | 0.027 | 2.25E-95 |
| Immune Cells | Nckap1l | 8.62E-94 | 6.75356407 | 0.526 | 0.056 | 1.72E-90 |
| Immune Cells | Lilrb2 | 4.59E-85 | 5.58746366 | 0.388 | 0.038 | 9.18E-82 |

| | | | | | | |
|---|---|---|---|---|---|---|
| Immune Cells | Ubash3a | 7.39E-80 | 6.06209429 | 0.293 | 0.036 | 1.48E-76 |
| Immune Cells | Spdef | 1.22E-78 | 0.929011 | 0.302 | 0.015 | 2.44E-75 |
| Immune Cells | Tlr7 | 2.43E-76 | 5.62777042 | 0.491 | 0.019 | 4.86E-73 |
| Immune Cells | AABR07029272.1 | 4.76E-76 | 5.87559255 | 0.534 | 0.054 | 9.53E-73 |
| Immune Cells | Glipr11 | 4.34E-75 | 5.08656528 | 0.474 | 0.053 | 8.68E-72 |
| Immune Cells | Hck | 5.85E-74 | 6.66842923 | 0.638 | 0.104 | 1.17E-70 |
| Immune Cells | Zc3h12d | 3.93E-71 | 5.30701307 | 0.336 | 0.054 | 7.86E-68 |
| Immune Cells | C5ar1 | 5.31E-71 | 6.89816943 | 0.328 | 0.014 | 1.06E-67 |
| Immune Cells | Tbx11 | 8.11E-70 | 1.18909056 | 0.457 | 0.055 | 1.62E-66 |
| Immune Cells | Nlrp3 | 1.11E-66 | 6.59781816 | 0.586 | 0.087 | 2.21E-63 |
| Immune Cells | Esco2 | 1.14E-65 | 2.7221348 | 0.448 | 0.052 | 2.29E-62 |
| Immune Cells | LOC6900451 | 2.81E-65 | 6.48086002 | 0.586 | 0.089 | 5.61E-62 |
| Immune Cells | Matk1 | 5.08E-65 | 6.63548231 | 0.345 | 0.02 | 1.02E-61 |
| Immune Cells | Hoxb51 | 3.45E-64 | 2.22958054 | 0.379 | 0.023 | 6.91E-61 |

| | | | | | | |
|---|---|---|---|---|---|---|
| Immune Cells | Cyth4 | 2.40E-62 | 6.14537403 | 0.578 | 0.075 | 4.80E-59 |
| Immune Cells | Kif11 | 9.75E-62 | 1.86085251 | 0.491 | 0.055 | 1.95E-58 |
| Immune Cells | Csf1r1 | 4.13E-61 | 6.39605197 | 0.517 | 0.068 | 8.25E-58 |
| Immune Cells | Cd41 | 5.39E-61 | 5.96022753 | 0.44 | 0.061 | 1.08E-57 |
| Immune Cells | Uroc1 | 1.45E-60 | 0.96869767 | 0.371 | 0.039 | 2.89E-57 |
| Immune Cells | Coro1a | 5.42E-59 | 5.628231 | 0.517 | 0.076 | 1.08E-55 |
| Immune Cells | Epsti1 | 1.64E-58 | 5.24624693 | 0.405 | 0.059 | 3.28E-55 |
| Immune Cells | RT1-Db1 | 1.65E-58 | 5.96473644 | 0.422 | 0.062 | 3.30E-55 |
| Immune Cells | Syk1 | 1.51E-56 | 6.31066228 | 0.466 | 0.047 | 3.01E-53 |
| Immune Cells | Ptprc | 4.03E-52 | 6.49700108 | 0.793 | 0.18 | 8.06E-49 |
| Immune Cells | Clec4e | 8.66E-52 | 6.13162817 | 0.345 | 0.047 | 1.73E-48 |
| Immune Cells | Lcp11 | 3.05E-51 | 5.70656104 | 0.733 | 0.204 | 6.11E-48 |
| Immune Cells | Kif20b | 1.71E-48 | 1.80177986 | 0.448 | 0.055 | 3.41E-45 |
| Immune Cells | Aoah | 1.40E-47 | 5.96110999 | 0.647 | 0.174 | 2.79E-44 |

| | | | | | | |
|---|---|---|---|---|---|---|
| Immune Cells | Plk41 | 1.59E-47 | 3.56362584 | 0.44 | 0.026 | 3.19E-44 |
| Immune Cells | RGD1559482 | 5.59E-46 | 5.63403717 | 0.534 | 0.075 | 1.12E-42 |
| Immune Cells | Acap11 | 2.16E-45 | 4.00643463 | 0.44 | 0.086 | 4.32E-42 |
| Immune Cells | AABR07006311.1 | 2.18E-45 | 8.2137702 | 0.31 | 0.002 | 4.36E-42 |
| Immune Cells | Melk1 | 1.95E-44 | 0.91555676 | 0.345 | 0.049 | 3.90E-41 |
| Immune Cells | Kcnk13 | 2.58E-44 | 6.68729953 | 0.534 | 0.102 | 5.16E-41 |
| Immune Cells | Itgam | 4.43E-44 | 6.67164358 | 0.647 | 0.19 | 8.86E-41 |
| Immune Cells | Lilrb4 | 3.03E-41 | 5.52320808 | 0.388 | 0.085 | 6.06E-38 |
| Immune Cells | Rspo31 | 7.40E-41 | 2.86826484 | 0.276 | 0.025 | 1.48E-37 |
| Immune Cells | Notum | 2.76E-39 | 0.35209414 | 0.397 | 0.047 | 5.51E-36 |
| Immune Cells | LOC100910636 | 4.67E-39 | 5.85221573 | 0.44 | 0.039 | 9.34E-36 |
| Immune Cells | Tbxas1 | 6.46E-39 | 6.17494298 | 0.621 | 0.179 | 1.29E-35 |
| Immune Cells | Csf3r | 1.01E-38 | 6.21114659 | 0.336 | 0.039 | 2.02E-35 |
| Immune Cells | F13a1 | 4.63E-37 | 5.26083031 | 0.474 | 0.09 | 9.25E-34 |

| | | | | | | |
|---|---|---|---|---|---|---|
| Immune Cells | AABR07072096.11 | 6.69E-37 | 3.80971762 | 0.302 | 0.019 | 1.34E-33 |
| Immune Cells | Selp | 8.20E-37 | 1.83168989 | 0.405 | 0.07 | 1.64E-33 |
| Immune Cells | Ikzf11 | 1.16E-36 | 5.91120267 | 0.491 | 0.091 | 2.33E-33 |
| Immune Cells | Ciita1 | 4.99E-36 | 6.05827946 | 0.388 | 0.137 | 9.97E-33 |
| Immune Cells | Fcgr2a | 8.54E-36 | 6.63845917 | 0.328 | 0.056 | 1.71E-32 |
| Immune Cells | Sgo2 | 3.62E-35 | 1.6615217 | 0.457 | 0.049 | 7.24E-32 |
| Immune Cells | Alox5 | 6.46E-35 | 6.73013846 | 0.379 | 0.128 | 1.29E-31 |
| Immune Cells | Arhgap15 | 6.71E-35 | 4.89511995 | 0.75 | 0.263 | 1.34E-31 |
| Immune Cells | Pstpip11 | 5.33E-34 | 4.00716269 | 0.647 | 0.154 | 1.07E-30 |
| Immune Cells | Kif4a | 6.00E-34 | 3.4800711 | 0.466 | 0.097 | 1.20E-30 |
| Immune Cells | RGD13058071 | 2.46E-33 | 1.87004607 | 0.397 | 0.046 | 4.92E-30 |
| Immune Cells | Themis | 6.42E-32 | 3.91997031 | 0.319 | 0.069 | 1.28E-28 |
| Immune Cells | Itgal | 2.83E-31 | 5.18049388 | 0.44 | 0.111 | 5.66E-28 |
| Immune Cells | Nrg4 | 7.73E-31 | 4.33862677 | 0.353 | 0.064 | 1.55E-27 |

| | | | | | | |
|---|---|---|---|---|---|---|
| Immune Cells | Fyb1 | 8.79E-31 | 5.54965737 | 0.586 | 0.169 | 1.76E-27 |
| Immune Cells | Pmel | 1.11E-30 | 0.7107995 | 0.319 | 0.038 | 2.22E-27 |
| Immune Cells | Dpep2 | 7.99E-30 | 0.86997518 | 0.371 | 0.078 | 1.60E-26 |
| Immune Cells | Plac8 | 8.17E-30 | 4.18861993 | 0.466 | 0.077 | 1.63E-26 |
| Immune Cells | Il71 | 9.42E-30 | 2.15757449 | 0.388 | 0.063 | 1.88E-26 |
| Immune Cells | Cfh | 6.11E-29 | 3.97101208 | 0.759 | 0.36 | 1.22E-25 |
| Immune Cells | Prkcb2 | 2.00E-28 | 5.35589232 | 0.483 | 0.143 | 3.99E-25 |
| Immune Cells | Dock8 | 2.74E-26 | 3.44652451 | 0.698 | 0.372 | 5.48E-23 |
| Immune Cells | Lrrc31 | 4.23E-26 | 3.80120444 | 0.345 | 0.071 | 8.46E-23 |
| Immune Cells | Nusap1 | 1.02E-25 | 2.61180402 | 0.405 | 0.048 | 2.03E-22 |
| Immune Cells | Inpp5d | 1.39E-24 | 4.05496106 | 0.629 | 0.248 | 2.78E-21 |
| Immune Cells | Lilrb3a | 1.58E-24 | 4.39477428 | 0.517 | 0.184 | 3.17E-21 |
| Immune Cells | Cenpf | 1.98E-24 | 2.51356744 | 0.5 | 0.141 | 3.97E-21 |
| Immune Cells | P2ry6 | 2.11E-24 | 4.9957506 | 0.388 | 0.083 | 4.22E-21 |

| | | | | | | |
|---|---|---|---|---|---|---|
| Immune Cells | Wdfy41 | 3.53E-24 | 5.09807091 | 0.405 | 0.104 | 7.06E-21 |
| Immune Cells | Lcp2 | 6.52E-24 | 3.81337058 | 0.595 | 0.24 | 1.30E-20 |
| Immune Cells | Cers6 | 8.87E-24 | 2.96604841 | 0.716 | 0.341 | 1.77E-20 |
| Immune Cells | Catsper3 | 1.04E-23 | 3.21255629 | 0.466 | 0.06 | 2.08E-20 |
| Immune Cells | Zeb22 | 1.08E-23 | 1.41213228 | 0.94 | 0.66 | 2.17E-20 |
| Immune Cells | S100a9 | 1.13E-23 | 9.13844162 | 0.414 | 0.091 | 2.26E-20 |
| Immune Cells | Trpc61 | 1.41E-23 | 1.64883501 | 0.5 | 0.146 | 2.83E-20 |
| Immune Cells | Stab1 | 1.82E-23 | 5.45394127 | 0.517 | 0.192 | 3.64E-20 |
| Immune Cells | Mbp | 2.99E-22 | 3.44058155 | 0.603 | 0.247 | 5.99E-19 |
| Immune Cells | Prkcq2 | 4.75E-22 | 4.63652962 | 0.448 | 0.195 | 9.50E-19 |
| Immune Cells | Ccl2 | 7.17E-22 | 2.04754381 | 0.474 | 0.154 | 1.43E-18 |
| Immune Cells | Runx1 | 7.56E-22 | 3.13263825 | 0.698 | 0.375 | 1.51E-18 |
| Immune Cells | Ccl211 | 1.24E-21 | 0.30476713 | 0.483 | 0.123 | 2.47E-18 |
| Immune Cells | Fcer1g | 1.54E-21 | 4.38479265 | 0.509 | 0.189 | 3.08E-18 |

| | | | | | | |
|---|---|---|---|---|---|---|
| Immune Cells | Slfn4 | 1.70E-21 | 4.78019313 | 0.681 | 0.399 | 3.41E-18 |
| Immune Cells | Pkmyt1 | 2.75E-21 | 2.07602488 | 0.431 | 0.08 | 5.51E-18 |
| Immune Cells | Frmd4b1 | 2.22E-19 | 2.48830386 | 0.741 | 0.395 | 4.45E-16 |
| Immune Cells | Kcnt2 | 2.29E-19 | 3.64197292 | 0.629 | 0.313 | 4.59E-16 |
| Immune Cells | Itga41 | 2.40E-19 | 4.30653362 | 0.526 | 0.193 | 4.80E-16 |
| Immune Cells | Pstpip21 | 6.20E-19 | 3.2435431 | 0.603 | 0.256 | 1.24E-15 |
| Immune Cells | Ptpro | 7.47E-19 | 5.28933205 | 0.491 | 0.178 | 1.49E-15 |
| Immune Cells | Anln | 1.85E-18 | 1.67421742 | 0.552 | 0.188 | 3.70E-15 |
| Immune Cells | Mme | 2.47E-18 | 0.82304715 | 0.466 | 0.117 | 4.93E-15 |
| Immune Cells | LOC1009114861 | 3.41E-18 | 0.37965413 | 0.474 | 0.12 | 6.83E-15 |
| Immune Cells | Slfn13 | 7.78E-18 | 2.8621946 | 0.56 | 0.298 | 1.56E-14 |
| Immune Cells | Ubash3b | 1.64E-17 | 2.08349231 | 0.707 | 0.443 | 3.27E-14 |
| Immune Cells | Mctp21 | 2.27E-16 | 4.71876422 | 0.5 | 0.234 | 4.54E-13 |
| Immune Cells | Aspm | 2.59E-16 | 2.00759714 | 0.431 | 0.111 | 5.17E-13 |

| | | | | | | |
|---|---|---|---|---|---|---|
| Immune Cells | Irf4 | 4.52E-16 | 3.67808878 | 0.362 | 0.092 | 9.03E-13 |
| Immune Cells | Pak1 | 7.90E-16 | 2.37390859 | 0.353 | 0.071 | 1.58E-12 |
| Immune Cells | Colec121 | 1.70E-15 | 2.2383005 | 0.707 | 0.308 | 3.41E-12 |
| Immune Cells | Tnfrsf1b | 2.00E-15 | 2.95409083 | 0.569 | 0.268 | 4.01E-12 |
| Immune Cells | Brca1 | 4.70E-15 | 2.15553904 | 0.586 | 0.216 | 9.41E-12 |
| Immune Cells | Lyn | 5.53E-15 | 2.01639775 | 0.716 | 0.457 | 1.11E-11 |
| Immune Cells | Susd41 | 2.68E-14 | 0.62886023 | 0.388 | 0.099 | 5.37E-11 |
| Immune Cells | Abca1 | 4.53E-14 | 1.72647628 | 0.776 | 0.491 | 9.07E-11 |
| Immune Cells | LOC103693323 | 5.21E-14 | 0.99363561 | 0.448 | 0.14 | 1.04E-10 |
| Immune Cells | Dipk1a | 6.33E-14 | 2.36341752 | 0.603 | 0.34 | 1.27E-10 |
| Immune Cells | Adgrg3 | 8.74E-14 | 1.06473804 | 0.405 | 0.124 | 1.75E-10 |
| Immune Cells | Dnmt3l | 9.25E-14 | 0.99045668 | 0.345 | 0.055 | 1.85E-10 |
| Immune Cells | Kcnip1 | 1.53E-13 | 1.51263075 | 0.491 | 0.226 | 3.06E-10 |
| Immune Cells | Nnat1 | 1.77E-13 | 1.61708263 | 0.483 | 0.208 | 3.54E-10 |

| | | | | | | |
|---|---|---|---|---|---|---|
| Immune Cells | Rhpn21 | 2.34E-13 | 1.78410927 | 0.5 | 0.209 | 4.69E-10 |
| Immune Cells | Sv2c | 3.94E-13 | 0.25052778 | 0.474 | 0.125 | 7.89E-10 |
| Immune Cells | Ankrd55 | 9.63E-13 | 1.36601025 | 0.569 | 0.309 | 1.93E-09 |
| Immune Cells | Tec1 | 1.07E-12 | 2.32179637 | 0.543 | 0.266 | 2.15E-09 |
| Immune Cells | Igf1 | 1.36E-12 | 2.42286618 | 0.629 | 0.355 | 2.73E-09 |
| Immune Cells | C1qtnf9 | 2.38E-12 | 0.42127656 | 0.509 | 0.179 | 4.77E-09 |
| Immune Cells | Tmem63c | 4.15E-12 | 0.47393279 | 0.431 | 0.163 | 8.30E-09 |
| Immune Cells | Satb2 | 5.93E-12 | 0.95543195 | 0.569 | 0.24 | 1.19E-08 |
| Immune Cells | S100a4 | 7.06E-12 | 1.27117955 | 0.509 | 0.198 | 1.41E-08 |
| Immune Cells | Mertk | 1.10E-11 | 2.72731463 | 0.517 | 0.238 | 2.19E-08 |
| Immune Cells | Rhoh | 1.29E-11 | 2.93727617 | 0.5 | 0.248 | 2.58E-08 |
| Immune Cells | Rab11fip4 | 1.56E-11 | 1.68964465 | 0.466 | 0.155 | 3.13E-08 |
| Immune Cells | Slc7a7 | 1.60E-11 | 2.85083307 | 0.526 | 0.276 | 3.19E-08 |
| Immune Cells | St8sia4 | 2.14E-11 | 1.83090806 | 0.612 | 0.359 | 4.28E-08 |

| | | | | | | |
|---|---|---|---|---|---|---|
| Immune Cells | Pdpn1 | 2.69E-11 | 0.52575862 | 0.448 | 0.193 | 5.37E-08 |
| Immune Cells | Spp1 | 3.64E-11 | 5.37126536 | 0.457 | 0.159 | 7.28E-08 |
| Immune Cells | Kntc11 | 5.95E-11 | 2.49013611 | 0.509 | 0.257 | 1.19E-07 |
| Immune Cells | Prph | 1.64E-10 | 1.13996459 | 0.534 | 0.2 | 3.28E-07 |
| Immune Cells | Vim | 2.54E-10 | 1.51893998 | 0.655 | 0.36 | 5.09E-07 |
| Immune Cells | AABR07054490.1 | 2.98E-10 | 0.64790115 | 0.586 | 0.234 | 5.96E-07 |
| Immune Cells | Tmem47 | 1.10E-09 | 0.66672936 | 0.552 | 0.288 | 2.21E-06 |
| Immune Cells | Sfrp4 | 1.39E-09 | 1.46534969 | 0.345 | 0.094 | 2.77E-06 |
| Immune Cells | Angptl1 | 1.40E-09 | 0.81252082 | 0.552 | 0.255 | 2.79E-06 |
| Immune Cells | Brip1 | 1.56E-09 | 1.34836781 | 0.552 | 0.278 | 3.13E-06 |
| Immune Cells | Arhgap261 | 2.20E-09 | 1.01436966 | 0.759 | 0.501 | 4.40E-06 |
| Immune Cells | Smox | 2.35E-09 | 1.59498402 | 0.509 | 0.247 | 4.69E-06 |
| Immune Cells | Ptgs1 | 5.78E-09 | 1.82990449 | 0.483 | 0.18 | 1.16E-05 |
| Immune Cells | Mark11 | 7.48E-09 | 0.25375303 | 0.543 | 0.234 | 1.50E-05 |

| | | | | | | |
|---|---|---|---|---|---|---|
| Immune Cells | Thsd7b1 | 1.64E-07 | 1.05793117 | 0.431 | 0.171 | 0.00032823 |
| Immune Cells | Aatk1 | 4.25E-07 | 0.29012743 | 0.5 | 0.225 | 0.0008499 |
| Immune Cells | Kcnk2 | 6.52E-07 | 0.44367458 | 0.509 | 0.217 | 0.00130433 |
| Immune Cells | Tnnt31 | 1.36E-06 | 0.8234536 | 0.638 | 0.343 | 0.00272963 |
| Immune Cells | Slit2 | 1.03E-05 | 0.34773706 | 0.578 | 0.288 | 0.02067989 |
| Immune Cells | Luzp2 | 1.21E-05 | 0.30992426 | 0.448 | 0.193 | 0.0241808 |
| Lymphatic ECs | Fam205a | 0 | 8.28010618 | 0.603 | 0.01 | 0 |
| Lymphatic ECs | Tbx1 | 1.09E-193 | 7.8676472 | 0.649 | 0.05 | 2.18E-190 |
| Lymphatic ECs | Kcnn2 | 4.05E-181 | 4.88416102 | 0.519 | 0.021 | 8.09E-178 |
| Lymphatic ECs | Apba2 | 7.99E-161 | 7.13364916 | 0.45 | 0.026 | 1.60E-157 |
| Lymphatic ECs | Vil11 | 5.59E-155 | 4.47449205 | 0.504 | 0.033 | 1.12E-151 |
| Lymphatic ECs | Rspo3 | 8.22E-155 | 5.88907419 | 0.55 | 0.017 | 1.64E-151 |
| Lymphatic ECs | AABR07013701.1 | 4.02E-144 | 4.08312097 | 0.45 | 0.011 | 8.05E-141 |
| Lymphatic ECs | AABR07060560.2 | 4.23E-144 | 6.47730202 | 0.511 | 0.022 | 8.46E-141 |

| | | | | | | |
|---|---|---|---|---|---|---|
| Lymphatic ECs | Nsg2 | 5.77E-144 | 6.1897049 | 0.55 | 0.04 | 1.15E-140 |
| Lymphatic ECs | Hoxb5 | 1.50E-128 | 5.87397591 | 0.542 | 0.018 | 3.00E-125 |
| Lymphatic ECs | Cx3cr1 | 1.78E-119 | 3.17023857 | 0.473 | 0.014 | 3.56E-116 |
| Lymphatic ECs | AABR07068316.2 | 6.84E-109 | 2.39590721 | 0.45 | 0.017 | 1.37E-105 |
| Lymphatic ECs | Rufy4 | 9.67E-108 | 3.58159283 | 0.473 | 0.029 | 1.93E-104 |
| Lymphatic ECs | Mmrn1 | 2.35E-107 | 8.16063141 | 0.763 | 0.064 | 4.70E-104 |
| Lymphatic ECs | Sh3gl3 | 2.56E-97 | 6.51674397 | 0.466 | 0.07 | 5.11E-94 |
| Lymphatic ECs | Il7 | 4.99E-97 | 6.34592212 | 0.588 | 0.057 | 9.99E-94 |
| Lymphatic ECs | Jaml | 5.39E-92 | 3.22388576 | 0.458 | 0.045 | 1.08E-88 |
| Lymphatic ECs | Reln1 | 6.38E-88 | 6.48054105 | 0.954 | 0.303 | 1.28E-84 |
| Lymphatic ECs | Brinp1 | 7.92E-88 | 2.32011251 | 0.519 | 0.051 | 1.58E-84 |
| Lymphatic ECs | Melk | 2.65E-85 | 1.67458033 | 0.519 | 0.044 | 5.31E-82 |
| Lymphatic ECs | Matk | 6.73E-83 | 2.29343986 | 0.473 | 0.016 | 1.35E-79 |
| Lymphatic ECs | Chdh1 | 9.33E-80 | 3.82122535 | 0.45 | 0.021 | 1.87E-76 |

| | | | | | | |
|---|---|---|---|---|---|---|
| Lymphatic ECs | Adh7 | 1.04E-76 | 3.16874017 | 0.473 | 0.021 | 2.08E-73 |
| Lymphatic ECs | Dtx1 | 5.78E-71 | 6.61976603 | 0.405 | 0.104 | 1.16E-67 |
| Lymphatic ECs | Pax3 | 9.67E-68 | 1.06983898 | 0.466 | 0.064 | 1.93E-64 |
| Lymphatic ECs | Gldn1 | 6.44E-66 | 0.75047988 | 0.511 | 0.012 | 1.29E-62 |
| Lymphatic ECs | AABR07063811.1 | 3.32E-65 | 3.75401073 | 0.466 | 0.065 | 6.64E-62 |
| Lymphatic ECs | Pkhd1l11 | 1.71E-64 | 3.88830866 | 0.931 | 0.431 | 3.42E-61 |
| Lymphatic ECs | Cmahp | 7.63E-62 | 5.20156851 | 0.573 | 0.094 | 1.53E-58 |
| Lymphatic ECs | Ccl21 | 1.91E-59 | 6.79718359 | 0.55 | 0.121 | 3.81E-56 |
| Lymphatic ECs | Mpp71 | 6.61E-57 | 3.23197715 | 0.939 | 0.397 | 1.32E-53 |
| Lymphatic ECs | Ldb22 | 8.58E-57 | 3.68936333 | 0.908 | 0.456 | 1.72E-53 |
| Lymphatic ECs | Sema3d | 5.39E-54 | 4.53988317 | 0.84 | 0.317 | 1.08E-50 |
| Lymphatic ECs | Slc38a4 | 3.44E-52 | 4.27923935 | 0.534 | 0.053 | 6.89E-49 |
| Lymphatic ECs | Flt4 | 3.51E-51 | 4.86688545 | 0.733 | 0.237 | 7.02E-48 |
| Lymphatic ECs | Chst9 | 3.70E-50 | 2.03065957 | 0.511 | 0.078 | 7.40E-47 |

| | | | | | | |
|---|---|---|---|---|---|---|
| Lymphatic ECs | Nrp21 | 9.25E-50 | 3.18295845 | 0.893 | 0.407 | 1.85E-46 |
| Lymphatic ECs | Npnt | 5.11E-47 | 2.48498239 | 0.534 | 0.105 | 1.02E-43 |
| Lymphatic ECs | Ralgapa21 | 9.40E-47 | 2.58253729 | 0.931 | 0.594 | 1.88E-43 |
| Lymphatic ECs | Cd4 | 2.13E-46 | 1.39141142 | 0.527 | 0.057 | 4.25E-43 |
| Lymphatic ECs | AABR07001573.21 | 1.45E-44 | 2.0669831 | 0.473 | 0.094 | 2.90E-41 |
| Lymphatic ECs | Cpne7 | 7.73E-43 | 4.97663814 | 0.595 | 0.135 | 1.55E-39 |
| Lymphatic ECs | Susd4 | 1.30E-42 | 3.62622777 | 0.466 | 0.097 | 2.61E-39 |
| Lymphatic ECs | Cdh12 | 2.37E-42 | 1.58413795 | 0.504 | 0.105 | 4.74E-39 |
| Lymphatic ECs | Tc2n | 2.80E-42 | 5.58951802 | 0.718 | 0.263 | 5.61E-39 |
| Lymphatic ECs | Nr5a2 | 1.33E-40 | 2.89144594 | 0.511 | 0.107 | 2.66E-37 |
| Lymphatic ECs | Rab11fip1 | 2.20E-39 | 1.71414041 | 0.473 | 0.068 | 4.41E-36 |
| Lymphatic ECs | Adgrl41 | 3.20E-39 | 2.16226343 | 0.901 | 0.472 | 6.40E-36 |
| Lymphatic ECs | AABR07038983.1 | 9.90E-39 | 0.73322602 | 0.481 | 0.075 | 1.98E-35 |
| Lymphatic ECs | Hapln3 | 1.63E-38 | 2.8699677 | 0.496 | 0.053 | 3.26E-35 |

| | | | | | | |
|---|---|---|---|---|---|---|
| Lymphatic ECs | Pde7b1 | 2.03E-35 | 2.02059153 | 0.931 | 0.603 | 4.06E-32 |
| Lymphatic ECs | Prox11 | 4.77E-35 | 3.90627628 | 0.687 | 0.394 | 9.53E-32 |
| Lymphatic ECs | Csf1r | 6.11E-35 | 0.59273171 | 0.481 | 0.068 | 1.22E-31 |
| Lymphatic ECs | Sema3a | 9.22E-35 | 5.5949225 | 0.634 | 0.242 | 1.84E-31 |
| Lymphatic ECs | Nxn | 1.11E-34 | 2.52195072 | 0.878 | 0.561 | 2.22E-31 |
| Lymphatic ECs | Mctp11 | 1.32E-34 | 3.33314259 | 0.794 | 0.35 | 2.64E-31 |
| Lymphatic ECs | Clnk | 4.00E-34 | 0.95058321 | 0.496 | 0.09 | 8.00E-31 |
| Lymphatic ECs | Ntn1 | 4.64E-34 | 2.52378584 | 0.878 | 0.563 | 9.28E-31 |
| Lymphatic ECs | Klhl4 | 3.20E-33 | 3.74482066 | 0.748 | 0.32 | 6.40E-30 |
| Lymphatic ECs | Efna5 | 4.70E-33 | 4.29550813 | 0.649 | 0.278 | 9.40E-30 |
| Lymphatic ECs | Plxdc21 | 7.46E-33 | 2.11192645 | 0.901 | 0.583 | 1.49E-29 |
| Lymphatic ECs | Olfml2a | 1.22E-32 | 3.39271834 | 0.565 | 0.195 | 2.43E-29 |
| Lymphatic ECs | Kalrn1 | 5.87E-32 | 1.6410978 | 0.893 | 0.616 | 1.17E-28 |
| Lymphatic ECs | Cysltr1 | 7.02E-30 | 0.29534858 | 0.45 | 0.098 | 1.40E-26 |

| | | | | | | |
|---|---|---|---|---|---|---|
| Lymphatic ECs | Abi3bp | 3.09E-29 | 1.94326517 | 0.832 | 0.512 | 6.17E-26 |
| Lymphatic ECs | AABR07049 085.11 | 2.65E-28 | 4.02388662 | 0.733 | 0.311 | 5.29E-25 |
| Lymphatic ECs | Ston2 | 4.03E-28 | 2.22918066 | 0.809 | 0.42 | 8.06E-25 |
| Lymphatic ECs | Frmd4b | 2.37E-27 | 2.56200728 | 0.725 | 0.395 | 4.74E-24 |
| Lymphatic ECs | Cenpe | 3.34E-27 | 0.7523626 | 0.473 | 0.088 | 6.68E-24 |
| Lymphatic ECs | Ifit3 | 1.09E-24 | 1.67355014 | 0.504 | 0.151 | 2.18E-21 |
| Lymphatic ECs | Ciita | 1.82E-24 | 0.56918544 | 0.443 | 0.135 | 3.65E-21 |
| Lymphatic ECs | Dapk11 | 1.99E-24 | 1.68124533 | 0.847 | 0.516 | 3.98E-21 |
| Lymphatic ECs | Tspan5 | 2.86E-24 | 2.27574551 | 0.794 | 0.435 | 5.71E-21 |
| Lymphatic ECs | Cp | 4.22E-24 | 3.03378062 | 0.687 | 0.285 | 8.43E-21 |
| Lymphatic ECs | Procr | 4.88E-24 | 1.22015009 | 0.504 | 0.131 | 9.77E-21 |
| Lymphatic ECs | Egfl71 | 8.69E-24 | 1.8847717 | 0.802 | 0.52 | 1.74E-20 |
| Lymphatic ECs | Casc4 | 3.59E-23 | 2.93623406 | 0.679 | 0.331 | 7.18E-20 |
| Lymphatic ECs | Galnt14 | 9.43E-23 | 0.27903165 | 0.511 | 0.186 | 1.89E-19 |

| | | | | | | |
|---|---|---|---|---|---|---|
| Lymphatic ECs | Msr1 | 2.67E-22 | 0.4259236 | 0.489 | 0.104 | 5.33E-19 |
| Lymphatic ECs | Abo | 8.08E-22 | 3.77661621 | 0.511 | 0.075 | 1.62E-18 |
| Lymphatic ECs | Sat1 | 1.50E-21 | 2.65638192 | 0.74 | 0.409 | 3.00E-18 |
| Lymphatic ECs | Icam11 | 2.80E-21 | 1.80670258 | 0.542 | 0.189 | 5.59E-18 |
| Lymphatic ECs | Nnat | 3.27E-21 | 2.07653405 | 0.481 | 0.207 | 6.53E-18 |
| Lymphatic ECs | Adgre41 | 3.30E-21 | 1.70557575 | 0.504 | 0.139 | 6.59E-18 |
| Lymphatic ECs | Flrt2 | 5.92E-21 | 2.99392168 | 0.626 | 0.356 | 1.18E-17 |
| Lymphatic ECs | Kntc1 | 1.23E-20 | 2.14348005 | 0.534 | 0.256 | 2.47E-17 |
| Lymphatic ECs | Lyve1 | 1.53E-20 | 4.9461624 | 0.42 | 0.139 | 3.07E-17 |
| Lymphatic ECs | Wnt5b | 2.13E-20 | 1.51787262 | 0.466 | 0.164 | 4.27E-17 |
| Lymphatic ECs | Trpc3 | 2.76E-20 | 1.84925631 | 0.588 | 0.247 | 5.53E-17 |
| Lymphatic ECs | AABR07027581.1 | 5.42E-20 | 2.36558185 | 0.687 | 0.393 | 1.08E-16 |
| Lymphatic ECs | Itgb41 | 6.79E-20 | 2.07255354 | 0.634 | 0.313 | 1.36E-16 |
| Lymphatic ECs | Fam189a1 | 1.08E-19 | 4.16567092 | 0.573 | 0.294 | 2.16E-16 |

| | | | | | | |
|---|---|---|---|---|---|---|
| Lymphatic ECs | Lmcd1 | 1.15E-19 | 2.13555128 | 0.748 | 0.44 | 2.30E-16 |
| Lymphatic ECs | Prkch1 | 2.99E-19 | 1.15067598 | 0.863 | 0.507 | 5.98E-16 |
| Lymphatic ECs | Shc3 | 3.10E-19 | 2.85023388 | 0.534 | 0.235 | 6.19E-16 |
| Lymphatic ECs | Ptpn18 | 5.32E-19 | 1.16255145 | 0.45 | 0.155 | 1.06E-15 |
| Lymphatic ECs | Mapk101 | 8.55E-19 | 2.93769463 | 0.542 | 0.187 | 1.71E-15 |
| Lymphatic ECs | Nr2f2 | 2.27E-18 | 3.21723834 | 0.626 | 0.287 | 4.55E-15 |
| Lymphatic ECs | AABR07054000.1 | 2.48E-18 | 2.22830019 | 0.481 | 0.165 | 4.96E-15 |
| Lymphatic ECs | Pappa2 | 2.52E-18 | 0.58797814 | 0.473 | 0.153 | 5.04E-15 |
| Lymphatic ECs | Wdfy4 | 3.25E-18 | 0.85868406 | 0.481 | 0.101 | 6.51E-15 |
| Lymphatic ECs | Kcnh81 | 8.27E-18 | 1.57822871 | 0.542 | 0.138 | 1.65E-14 |
| Lymphatic ECs | Slc39a8 | 8.98E-18 | 2.23685284 | 0.534 | 0.268 | 1.80E-14 |
| Lymphatic ECs | Emcn2 | 1.86E-17 | 1.35040045 | 0.84 | 0.502 | 3.72E-14 |
| Lymphatic ECs | RGD1564053 | 7.45E-17 | 1.10740059 | 0.511 | 0.155 | 1.49E-13 |
| Lymphatic ECs | Mcm6 | 8.95E-17 | 0.62697294 | 0.481 | 0.082 | 1.79E-13 |

| | | | | | | |
|---|---|---|---|---|---|---|
| Lymphatic ECs | Actn1 | 9.93E-17 | 1.93465112 | 0.725 | 0.462 | 1.99E-13 |
| Lymphatic ECs | Glis3 | 1.96E-16 | 2.00675619 | 0.725 | 0.436 | 3.91E-13 |
| Lymphatic ECs | Mafb | 4.01E-16 | 2.03020857 | 0.542 | 0.226 | 8.02E-13 |
| Lymphatic ECs | Ablim21 | 5.48E-16 | 2.48723351 | 0.611 | 0.296 | 1.10E-12 |
| Lymphatic ECs | Lck1 | 6.27E-16 | 0.61942592 | 0.473 | 0.144 | 1.25E-12 |
| Lymphatic ECs | Lrrtm41 | 1.78E-15 | 0.65112493 | 0.489 | 0.167 | 3.55E-12 |
| Lymphatic ECs | Tspan181 | 2.09E-15 | 1.49640046 | 0.779 | 0.449 | 4.18E-12 |
| Lymphatic ECs | Fam83b | 3.18E-15 | 0.3465434 | 0.504 | 0.107 | 6.36E-12 |
| Lymphatic ECs | Slc9a91 | 6.77E-15 | 1.57531766 | 0.763 | 0.45 | 1.35E-11 |
| Lymphatic ECs | Prkcq1 | 7.03E-15 | 0.93917108 | 0.481 | 0.193 | 1.41E-11 |
| Lymphatic ECs | St6galnac3 | 1.22E-14 | 1.64778976 | 0.702 | 0.361 | 2.43E-11 |
| Lymphatic ECs | Rhpn2 | 1.35E-14 | 1.07755721 | 0.519 | 0.207 | 2.70E-11 |
| Lymphatic ECs | Nebl1 | 2.21E-14 | 1.05298863 | 0.87 | 0.564 | 4.41E-11 |
| Lymphatic ECs | LOC682419 | 2.64E-14 | 0.58616646 | 0.45 | 0.129 | 5.27E-11 |

| | | | | | | |
|---|---|---|---|---|---|---|
| Lymphatic ECs | Timp3 | 5.12E-14 | 1.73948338 | 0.779 | 0.527 | 1.02E-10 |
| Lymphatic ECs | Pgm51 | 1.69E-13 | 1.00199231 | 0.733 | 0.483 | 3.37E-10 |
| Lymphatic ECs | Prkcb1 | 2.23E-13 | 0.70826245 | 0.481 | 0.142 | 4.47E-10 |
| Lymphatic ECs | Gbp2 | 2.29E-13 | 1.14835906 | 0.534 | 0.228 | 4.58E-10 |
| Lymphatic ECs | Mctp2 | 4.56E-13 | 1.71991152 | 0.519 | 0.233 | 9.12E-10 |
| Lymphatic ECs | Angpt2 | 4.89E-13 | 1.88413637 | 0.573 | 0.292 | 9.78E-10 |
| Lymphatic ECs | Adamts15 | 1.57E-12 | 1.51325123 | 0.618 | 0.247 | 3.14E-09 |
| Lymphatic ECs | Ednrb | 2.60E-12 | 1.6595863 | 0.573 | 0.299 | 5.20E-09 |
| Lymphatic ECs | Piezo2 | 2.62E-12 | 2.69024646 | 0.565 | 0.228 | 5.25E-09 |
| Lymphatic ECs | Ac1576 | 4.26E-12 | 1.29070174 | 0.519 | 0.199 | 8.52E-09 |
| Lymphatic ECs | Igfbp5 | 6.08E-12 | 2.27596709 | 0.588 | 0.264 | 1.22E-08 |
| Lymphatic ECs | Flvcr2 | 6.99E-12 | 0.89955682 | 0.557 | 0.234 | 1.40E-08 |
| Lymphatic ECs | Eln | 2.41E-11 | 1.28075344 | 0.634 | 0.35 | 4.81E-08 |
| Lymphatic ECs | Rsad21 | 2.68E-11 | 0.58733059 | 0.504 | 0.222 | 5.36E-08 |

| | | | | | | |
|---|---|---|---|---|---|---|
| Lymphatic ECs | Slco2b1 | 3.15E-11 | 1.13774927 | 0.71 | 0.459 | 6.30E-08 |
| Lymphatic ECs | Necab11 | 3.24E-11 | 0.337038 | 0.489 | 0.192 | 6.49E-08 |
| Lymphatic ECs | Ankrd6 | 4.49E-11 | 2.03454003 | 0.603 | 0.352 | 8.99E-08 |
| Lymphatic ECs | Camk4 | 6.72E-11 | 2.50671493 | 0.504 | 0.236 | 1.34E-07 |
| Lymphatic ECs | Thsd7b | 2.06E-10 | 3.12601416 | 0.466 | 0.169 | 4.12E-07 |
| Lymphatic ECs | Ube2ql11 | 1.12E-09 | 1.05798841 | 0.496 | 0.189 | 2.25E-06 |
| Lymphatic ECs | Mx11 | 2.24E-08 | 0.98113807 | 0.489 | 0.164 | 4.48E-05 |
| Lymphatic ECs | Tm4sf1 | 2.59E-07 | 0.32661929 | 0.565 | 0.291 | 0.00051754 |
| Lymphatic ECs | Colec12 | 8.87E-07 | 0.97624788 | 0.626 | 0.309 | 0.00177419 |
| Lymphatic ECs | Clec1a | 8.20E-06 | 0.8436186 | 0.55 | 0.287 | 0.01640761 |
| Lymphatic ECs | Usp18 | 1.34E-05 | 1.21522702 | 0.534 | 0.279 | 0.02681069 |
| Neuronal Cells | Rhov | 0 | 6.14769206 | 0.43 | 0.005 | 0 |
| Neuronal Cells | Tfap2b | 4.28E-248 | 6.44139409 | 0.407 | 0.008 | 8.57E-245 |
| Neuronal Cells | AABR07022098.1 | 8.07E-198 | 9.85723046 | 0.4 | 0.001 | 1.61E-194 |

| | | | | | | |
|---|---|---|---|---|---|---|
| Neuronal Cells | Slc22a2 | 1.38E-195 | 2.80849482 | 0.422 | 0.015 | 2.76E-192 |
| Neuronal Cells | Dusp2 | 2.40E-195 | 3.56416656 | 0.415 | 0.007 | 4.79E-192 |
| Neuronal Cells | Necab2 | 5.89E-182 | 3.68549814 | 0.385 | 0.009 | 1.18E-178 |
| Neuronal Cells | Slitrk6 | 2.74E-153 | 8.22957061 | 0.481 | 0.011 | 5.48E-150 |
| Neuronal Cells | Neto1 | 5.71E-149 | 7.06803164 | 0.407 | 0.032 | 1.14E-145 |
| Neuronal Cells | Tmsb15b2 | 6.33E-122 | 3.79488135 | 0.326 | 0.001 | 1.27E-118 |
| Neuronal Cells | Ptprz1 | 5.18E-118 | 7.1988767 | 0.304 | 0.03 | 1.04E-114 |
| Neuronal Cells | AABR07032787.1 | 2.66E-116 | 5.06060757 | 0.393 | 0.017 | 5.32E-113 |
| Neuronal Cells | Mest1 | 5.55E-108 | 2.10935147 | 0.43 | 0.033 | 1.11E-104 |
| Neuronal Cells | Erbb3 | 3.26E-105 | 7.20517793 | 0.667 | 0.105 | 6.53E-102 |
| Neuronal Cells | Pex5l | 3.22E-97 | 3.95274836 | 0.452 | 0.016 | 6.43E-94 |
| Neuronal Cells | AABR07072096.1 | 3.01E-94 | 0.68091358 | 0.437 | 0.015 | 6.03E-91 |
| Neuronal Cells | Scn7a | 2.81E-91 | 6.30559357 | 0.985 | 0.423 | 5.62E-88 |
| Neuronal Cells | Lgi4 | 1.43E-90 | 5.88139487 | 0.941 | 0.315 | 2.86E-87 |

| | | | | | | |
|---|---|---|---|---|---|---|
| Neuronal Cells | Tdrd12 | 2.10E-87 | 8.16595212 | 0.622 | 0.084 | 4.21E-84 |
| Neuronal Cells | Vil1 | 2.20E-87 | 4.41593597 | 0.378 | 0.035 | 4.41E-84 |
| Neuronal Cells | Igsf11 | 3.02E-86 | 8.21779204 | 0.756 | 0.08 | 6.04E-83 |
| Neuronal Cells | Cnksr2 | 1.27E-82 | 6.93006394 | 0.83 | 0.193 | 2.55E-79 |
| Neuronal Cells | Grik2 | 6.84E-80 | 6.14864982 | 0.941 | 0.422 | 1.37E-76 |
| Neuronal Cells | Rasgef1c | 8.14E-73 | 7.57434104 | 0.704 | 0.106 | 1.63E-69 |
| Neuronal Cells | Cadm2 | 1.23E-71 | 7.02028795 | 0.859 | 0.301 | 2.46E-68 |
| Neuronal Cells | Dab1 | 4.33E-68 | 7.69243432 | 0.378 | 0.026 | 8.66E-65 |
| Neuronal Cells | AABR07067469.1 | 3.09E-67 | 2.97391415 | 0.407 | 0.047 | 6.19E-64 |
| Neuronal Cells | Chdh | 1.84E-66 | 3.52949204 | 0.407 | 0.021 | 3.68E-63 |
| Neuronal Cells | Gldn | 1.46E-63 | 7.5625199 | 0.274 | 0.018 | 2.92E-60 |
| Neuronal Cells | L1cam | 5.68E-62 | 7.97510399 | 0.556 | 0.022 | 1.14E-58 |
| Neuronal Cells | Baiap2l2 | 6.64E-61 | 2.68460872 | 0.422 | 0.059 | 1.33E-57 |
| Neuronal Cells | Zeb21 | 9.84E-61 | 2.66029502 | 0.985 | 0.658 | 1.97E-57 |

| | | | | | | |
|---|---|---|---|---|---|---|
| Neuronal Cells | Abca8a1 | 1.69E-60 | 2.6676969 | 0.978 | 0.581 | 3.37E-57 |
| Neuronal Cells | Cdh19 | 1.80E-58 | 7.10755438 | 0.748 | 0.205 | 3.60E-55 |
| Neuronal Cells | Adam23 | 8.25E-58 | 5.5378522 | 0.807 | 0.234 | 1.65E-54 |
| Neuronal Cells | Ank31 | 3.92E-54 | 2.89503427 | 0.948 | 0.531 | 7.84E-51 |
| Neuronal Cells | Tp63 | 1.92E-51 | 3.33919168 | 0.415 | 0.033 | 3.84E-48 |
| Neuronal Cells | Xrra1 | 4.36E-51 | 2.64737143 | 0.4 | 0.065 | 8.72E-48 |
| Neuronal Cells | Gfra3 | 3.49E-50 | 6.39761699 | 0.422 | 0.137 | 6.97E-47 |
| Neuronal Cells | Afap1l2 | 4.50E-50 | 3.61292519 | 0.859 | 0.395 | 9.00E-47 |
| Neuronal Cells | Fbln7 | 1.42E-47 | 3.8095803 | 0.474 | 0.099 | 2.84E-44 |
| Neuronal Cells | Ntng1 | 2.25E-45 | 5.71245478 | 0.704 | 0.153 | 4.50E-42 |
| Neuronal Cells | Il1rapl1 | 1.40E-43 | 5.14145727 | 0.748 | 0.325 | 2.80E-40 |
| Neuronal Cells | Rassf4 | 5.48E-43 | 5.81577422 | 0.578 | 0.187 | 1.10E-39 |
| Neuronal Cells | Iqgap21 | 1.12E-42 | 4.29955688 | 0.807 | 0.319 | 2.24E-39 |
| Neuronal Cells | Ncam2 | 1.64E-41 | 6.57187864 | 0.563 | 0.154 | 3.28E-38 |

| | | | | | | |
|---|---|---|---|---|---|---|
| Neuronal Cells | Itgb4 | 7.55E-41 | 5.20081442 | 0.741 | 0.31 | 1.51E-37 |
| Neuronal Cells | Cadm1 | 7.91E-41 | 4.75470906 | 0.467 | 0.074 | 1.58E-37 |
| Neuronal Cells | Arhgef26 | 1.23E-40 | 5.26343463 | 0.741 | 0.296 | 2.46E-37 |
| Neuronal Cells | AABR07026483.1 | 1.70E-40 | 7.23955963 | 0.437 | 0.092 | 3.40E-37 |
| Neuronal Cells | Edaradd | 2.98E-39 | 2.79411304 | 0.385 | 0.066 | 5.95E-36 |
| Neuronal Cells | Matn2 | 3.15E-39 | 4.14785978 | 0.748 | 0.312 | 6.31E-36 |
| Neuronal Cells | Chl1 | 4.42E-39 | 6.58639879 | 0.4 | 0.15 | 8.85E-36 |
| Neuronal Cells | Ttyh1 | 5.34E-39 | 6.37572312 | 0.415 | 0.119 | 1.07E-35 |
| Neuronal Cells | Sox61 | 8.07E-39 | 2.32615734 | 0.904 | 0.601 | 1.61E-35 |
| Neuronal Cells | Alcam | 1.64E-38 | 3.83994637 | 0.793 | 0.379 | 3.29E-35 |
| Neuronal Cells | Alk | 1.67E-38 | 6.26790712 | 0.681 | 0.268 | 3.33E-35 |
| Neuronal Cells | Plk4 | 7.07E-38 | 0.67936831 | 0.378 | 0.026 | 1.41E-34 |
| Neuronal Cells | Adamts16 | 2.81E-37 | 2.10666411 | 0.422 | 0.038 | 5.62E-34 |
| Neuronal Cells | Gas7 | 6.25E-37 | 3.47049228 | 0.756 | 0.281 | 1.25E-33 |

| | | | | | | |
|---|---|---|---|---|---|---|
| Neuronal Cells | Fign1 | 3.33E-36 | 3.7496492 | 0.778 | 0.305 | 6.65E-33 |
| Neuronal Cells | Nrxn11 | 3.49E-36 | 1.8036914 | 0.889 | 0.604 | 6.98E-33 |
| Neuronal Cells | Dpp6 | 1.44E-35 | 2.25177814 | 0.341 | 0.041 | 2.88E-32 |
| Neuronal Cells | Adgrl3 | 3.39E-35 | 3.95592646 | 0.756 | 0.374 | 6.78E-32 |
| Neuronal Cells | LOC689599 | 2.62E-34 | 0.26649989 | 0.415 | 0.047 | 5.23E-31 |
| Neuronal Cells | LOC1025466831 | 1.49E-33 | 2.53291495 | 0.385 | 0.089 | 2.98E-30 |
| Neuronal Cells | Sorcs1 | 1.64E-33 | 4.56500406 | 0.637 | 0.253 | 3.27E-30 |
| Neuronal Cells | Gfra2 | 3.08E-33 | 5.10926329 | 0.444 | 0.183 | 6.17E-30 |
| Neuronal Cells | Egflam | 1.15E-32 | 3.09349789 | 0.756 | 0.362 | 2.29E-29 |
| Neuronal Cells | AC131411.1 | 1.64E-32 | 2.28763321 | 0.407 | 0.071 | 3.27E-29 |
| Neuronal Cells | Cobl | 2.19E-32 | 3.83745184 | 0.748 | 0.373 | 4.39E-29 |
| Neuronal Cells | Adgrg6 | 2.51E-31 | 3.90769103 | 0.689 | 0.373 | 5.02E-28 |
| Neuronal Cells | Col16a1 | 2.81E-31 | 4.28394428 | 0.644 | 0.284 | 5.61E-28 |
| Neuronal Cells | AC121209.1 | 7.23E-31 | 1.9205372 | 0.43 | 0.08 | 1.45E-27 |

| | | | | | | |
|---|---|---|---|---|---|---|
| Neuronal Cells | Stac3 | 2.60E-29 | 1.9106411 | 0.43 | 0.11 | 5.20E-26 |
| Neuronal Cells | Cdca2 | 6.37E-29 | 2.12318069 | 0.393 | 0.073 | 1.27E-25 |
| Neuronal Cells | Syk | 1.41E-28 | 0.86349549 | 0.437 | 0.046 | 2.82E-25 |
| Neuronal Cells | Zfp536 | 1.77E-28 | 5.78592614 | 0.637 | 0.189 | 3.53E-25 |
| Neuronal Cells | Gap43 | 2.17E-28 | 4.5131278 | 0.459 | 0.103 | 4.34E-25 |
| Neuronal Cells | Slc35f1 | 3.79E-27 | 4.53063443 | 0.667 | 0.318 | 7.57E-24 |
| Neuronal Cells | Lrrtm4 | 9.17E-27 | 6.78113345 | 0.563 | 0.165 | 1.83E-23 |
| Neuronal Cells | Col28a1 | 3.36E-26 | 4.76307792 | 0.481 | 0.177 | 6.72E-23 |
| Neuronal Cells | Egr31 | 4.15E-26 | 2.3382339 | 0.4 | 0.118 | 8.31E-23 |
| Neuronal Cells | Tmtc2 | 5.02E-26 | 2.64728522 | 0.763 | 0.424 | 1.00E-22 |
| Neuronal Cells | Ldb21 | 9.63E-26 | 2.0818975 | 0.778 | 0.459 | 1.93E-22 |
| Neuronal Cells | Insc | 2.12E-25 | 4.93717746 | 0.496 | 0.195 | 4.25E-22 |
| Neuronal Cells | Plxdc2 | 2.17E-25 | 1.89457402 | 0.852 | 0.584 | 4.33E-22 |
| Neuronal Cells | Arnt2 | 3.90E-25 | 3.02828783 | 0.43 | 0.092 | 7.80E-22 |

| | | | | | | |
|---|---|---|---|---|---|---|
| Neuronal Cells | Sgcd1 | 8.20E-25 | 1.95746236 | 0.793 | 0.433 | 1.64E-21 |
| Neuronal Cells | Grin3a | 1.00E-24 | 0.82309249 | 0.43 | 0.088 | 2.01E-21 |
| Neuronal Cells | Arhgap39 | 1.78E-24 | 3.89052291 | 0.637 | 0.346 | 3.56E-21 |
| Neuronal Cells | Myof | 1.93E-24 | 1.93874457 | 0.815 | 0.489 | 3.86E-21 |
| Neuronal Cells | Aatk | 4.98E-24 | 5.03870313 | 0.526 | 0.224 | 9.95E-21 |
| Neuronal Cells | Exoc3l2 | 5.64E-24 | 0.35013555 | 0.378 | 0.064 | 1.13E-20 |
| Neuronal Cells | Cdh1 | 5.69E-24 | 3.24170907 | 0.407 | 0.107 | 1.14E-20 |
| Neuronal Cells | LOC690045 | 8.17E-24 | 0.3701525 | 0.43 | 0.091 | 1.63E-20 |
| Neuronal Cells | Pde1c | 8.86E-24 | 2.78507373 | 0.689 | 0.334 | 1.77E-20 |
| Neuronal Cells | Kcnh8 | 1.10E-23 | 5.95334895 | 0.519 | 0.138 | 2.19E-20 |
| Neuronal Cells | Col11a1 | 2.32E-23 | 3.87940593 | 0.444 | 0.119 | 4.64E-20 |
| Neuronal Cells | Vangl2 | 7.86E-23 | 1.5873137 | 0.43 | 0.111 | 1.57E-19 |
| Neuronal Cells | Fam78b | 8.46E-23 | 3.21049936 | 0.659 | 0.38 | 1.69E-19 |
| Neuronal Cells | Fndc1 | 2.51E-22 | 2.38366151 | 0.748 | 0.412 | 5.02E-19 |

| | | | | | | |
|---|---|---|---|---|---|---|
| Neuronal Cells | Kif22 | 3.09E-22 | 1.0902201 | 0.415 | 0.122 | 6.18E-19 |
| Neuronal Cells | Sorcs2 | 5.08E-22 | 4.86216678 | 0.489 | 0.218 | 1.02E-18 |
| Neuronal Cells | Kank4 | 1.39E-21 | 3.94640397 | 0.326 | 0.056 | 2.77E-18 |
| Neuronal Cells | Fgf7 | 2.53E-21 | 2.54871749 | 0.4 | 0.096 | 5.07E-18 |
| Neuronal Cells | Gpm6b | 5.43E-21 | 2.61227555 | 0.696 | 0.351 | 1.09E-17 |
| Neuronal Cells | Hcn1 | 1.52E-20 | 4.89405848 | 0.481 | 0.097 | 3.03E-17 |
| Neuronal Cells | Map2 | 2.19E-20 | 4.77390177 | 0.504 | 0.206 | 4.37E-17 |
| Neuronal Cells | Dclk11 | 3.27E-20 | 1.88439117 | 0.741 | 0.463 | 6.54E-17 |
| Neuronal Cells | Kif23 | 3.23E-19 | 1.47450904 | 0.444 | 0.157 | 6.45E-16 |
| Neuronal Cells | Itga4 | 7.51E-19 | 4.0850576 | 0.533 | 0.192 | 1.50E-15 |
| Neuronal Cells | Grid2 | 7.98E-19 | 5.85222299 | 0.496 | 0.162 | 1.60E-15 |
| Neuronal Cells | Nlgn1 | 2.02E-18 | 5.86835281 | 0.519 | 0.214 | 4.05E-15 |
| Neuronal Cells | Sorbs21 | 4.10E-18 | 0.94012024 | 0.919 | 0.617 | 8.20E-15 |
| Neuronal Cells | Tmem200a | 5.90E-18 | 1.94415044 | 0.415 | 0.135 | 1.18E-14 |

| | | | | | | |
|---|---|---|---|---|---|---|
| Neuronal Cells | AABR07003030.2 | 1.00E-17 | 4.78664108 | 0.504 | 0.218 | 2.01E-14 |
| Neuronal Cells | AABR07044900.12 | 1.05E-17 | 1.83557612 | 0.748 | 0.489 | 2.10E-14 |
| Neuronal Cells | Trpm3 | 1.11E-17 | 3.98993725 | 0.563 | 0.25 | 2.22E-14 |
| Neuronal Cells | Vgll3 | 2.27E-17 | 3.72078532 | 0.57 | 0.264 | 4.54E-14 |
| Neuronal Cells | Gabra4 | 4.99E-17 | 1.99301611 | 0.43 | 0.152 | 9.98E-14 |
| Neuronal Cells | Cntn3 | 7.50E-17 | 0.72569418 | 0.407 | 0.097 | 1.50E-13 |
| Neuronal Cells | Dnajc6 | 8.34E-17 | 1.94028202 | 0.407 | 0.101 | 1.67E-13 |
| Neuronal Cells | Col5a3 | 8.96E-17 | 2.13913672 | 0.689 | 0.358 | 1.79E-13 |
| Neuronal Cells | Nav21 | 1.02E-16 | 1.69500486 | 0.77 | 0.495 | 2.03E-13 |
| Neuronal Cells | AABR07049033.11 | 5.17E-16 | 1.86377949 | 0.415 | 0.129 | 1.03E-12 |
| Neuronal Cells | Ryr31 | 8.16E-16 | 2.27737432 | 0.704 | 0.363 | 1.63E-12 |
| Neuronal Cells | Chrdl1 | 9.80E-16 | 2.0890132 | 0.467 | 0.126 | 1.96E-12 |
| Neuronal Cells | Mctp1 | 1.47E-15 | 2.26334297 | 0.659 | 0.353 | 2.93E-12 |
| Neuronal Cells | Abca8 | 1.85E-15 | 3.20770944 | 0.563 | 0.251 | 3.70E-12 |

| | | | | | | |
|---|---|---|---|---|---|---|
| Neuronal Cells | AABR07049085.1 | 2.40E-15 | 2.69663689 | 0.637 | 0.313 | 4.80E-12 |
| Neuronal Cells | Prkcq | 1.01E-14 | 3.25394873 | 0.459 | 0.194 | 2.02E-11 |
| Neuronal Cells | Chn21 | 4.96E-13 | 1.59352928 | 0.696 | 0.427 | 9.93E-10 |
| Neuronal Cells | Pde8b | 7.57E-13 | 3.6931484 | 0.459 | 0.204 | 1.51E-09 |
| Neuronal Cells | Sntb12 | 8.37E-13 | 1.68192939 | 0.63 | 0.37 | 1.67E-09 |
| Neuronal Cells | Prkcb | 1.34E-12 | 2.55454781 | 0.467 | 0.142 | 2.67E-09 |
| Neuronal Cells | Reln | 3.41E-12 | 1.37633089 | 0.563 | 0.312 | 6.82E-09 |
| Neuronal Cells | Kif6 | 6.33E-12 | 1.12437025 | 0.356 | 0.106 | 1.27E-08 |
| Neuronal Cells | Susd5 | 6.70E-12 | 0.76821734 | 0.422 | 0.168 | 1.34E-08 |
| Neuronal Cells | Itgb8 | 1.25E-11 | 2.41489056 | 0.615 | 0.309 | 2.51E-08 |
| Neuronal Cells | Mx1 | 1.59E-11 | 0.89269107 | 0.452 | 0.164 | 3.18E-08 |
| Neuronal Cells | Zswim5 | 1.69E-11 | 2.13865962 | 0.63 | 0.336 | 3.39E-08 |
| Neuronal Cells | Icam1 | 1.80E-11 | 0.94415695 | 0.467 | 0.19 | 3.60E-08 |
| Neuronal Cells | Ccn5 | 2.19E-11 | 1.29934409 | 0.422 | 0.095 | 4.39E-08 |

| | | | | | | |
|---|---|---|---|---|---|---|
| Neuronal Cells | Diaph3 | 7.48E-11 | 0.46341653 | 0.444 | 0.128 | 1.50E-07 |
| Neuronal Cells | Ikzf1 | 7.96E-11 | 0.71054971 | 0.415 | 0.092 | 1.59E-07 |
| Neuronal Cells | Mapk10 | 3.37E-10 | 2.640659 | 0.467 | 0.188 | 6.73E-07 |
| Neuronal Cells | Col12a1 | 6.64E-10 | 2.27967114 | 0.474 | 0.183 | 1.33E-06 |
| Neuronal Cells | Atf3 | 1.25E-09 | 1.81457772 | 0.4 | 0.09 | 2.49E-06 |
| Neuronal Cells | Pdgfc1 | 5.08E-09 | 0.87830511 | 0.437 | 0.149 | 1.02E-05 |
| Neuronal Cells | Mx2 | 6.05E-09 | 0.74970082 | 0.474 | 0.167 | 1.21E-05 |
| Neuronal Cells | RGD1306750 | 8.61E-09 | 3.00799862 | 0.319 | 0.064 | 1.72E-05 |
| Neuronal Cells | Caskin1 | 1.13E-08 | 3.24665076 | 0.407 | 0.141 | 2.26E-05 |
| Neuronal Cells | Pstpip1 | 1.91E-08 | 0.30847463 | 0.467 | 0.156 | 3.82E-05 |
| Neuronal Cells | Adgre4 | 2.70E-08 | 1.10688606 | 0.422 | 0.141 | 5.40E-05 |
| Neuronal Cells | Gabre | 8.72E-08 | 0.5591731 | 0.452 | 0.183 | 0.00017446 |
| Neuronal Cells | Lrg11 | 1.97E-07 | 0.64337088 | 0.393 | 0.117 | 0.00039358 |
| Neuronal Cells | Sh3gl2 | 2.31E-05 | 1.13291012 | 0.519 | 0.23 | 0.04613591 |

| | | | | | | |
|---|---|---|---|---|---|---|
| Proliferating Cells | Aunip | 0 | 9.42070984 | 0.5 | 0.001 | 0 |
| Proliferating Cells | LOC500827 | 0 | 7.00035955 | 0.389 | 0 | 0 |
| Proliferating Cells | Depdc1 | 1.52E-210 | 9.26756506 | 0.25 | 0 | 3.04E-207 |
| Proliferating Cells | Troap | 5.62E-140 | 9.9266192 | 0.722 | 0.017 | 1.12E-136 |
| Proliferating Cells | Klra22 | 2.58E-125 | 3.97110145 | 0.278 | 0.003 | 5.17E-122 |
| Proliferating Cells | Ly49i4 | 1.81E-116 | 2.8061123 | 0.278 | 0.003 | 3.62E-113 |
| Proliferating Cells | Depdc1b | 1.98E-89 | 7.3851645 | 0.417 | 0.021 | 3.96E-86 |
| Proliferating Cells | Top2a | 6.65E-87 | 8.74160953 | 0.889 | 0.018 | 1.33E-83 |
| Proliferating Cells | Fam111a | 2.52E-83 | 8.4062394 | 0.528 | 0.026 | 5.04E-80 |
| Proliferating Cells | Birc5 | 3.32E-82 | 8.5366965 | 0.556 | 0.013 | 6.64E-79 |
| Proliferating Cells | AABR07052608.11 | 6.02E-68 | 3.11924143 | 0.278 | 0.007 | 1.20E-64 |
| Proliferating Cells | Fcgr3a | 4.25E-62 | 6.25671224 | 0.389 | 0.012 | 8.50E-59 |
| Proliferating Cells | Neil3 | 2.18E-56 | 7.34319221 | 0.556 | 0.04 | 4.36E-53 |
| Proliferating Cells | Kif111 | 8.66E-54 | 8.8141087 | 0.75 | 0.059 | 1.73E-50 |

| | | | | | | |
|---|---|---|---|---|---|---|
| Proliferating Cells | Prc1 | 1.47E-53 | 7.7448539 | 0.639 | 0.045 | 2.95E-50 |
| Proliferating Cells | LOC1025498691 | 2.27E-49 | 1.69931699 | 0.278 | 0.01 | 4.54E-46 |
| Proliferating Cells | Gucy2e | 1.20E-45 | 5.07195703 | 0.417 | 0.023 | 2.40E-42 |
| Proliferating Cells | Nusap11 | 2.85E-43 | 7.60623291 | 0.806 | 0.051 | 5.70E-40 |
| Proliferating Cells | Ect2 | 6.06E-40 | 8.90207917 | 0.806 | 0.098 | 1.21E-36 |
| Proliferating Cells | Esco21 | 9.80E-40 | 8.08467849 | 0.583 | 0.057 | 1.96E-36 |
| Proliferating Cells | Iqgap3 | 5.80E-39 | 8.47440351 | 0.75 | 0.11 | 1.16E-35 |
| Proliferating Cells | Cdkn3 | 1.22E-38 | 8.49344291 | 0.611 | 0.026 | 2.43E-35 |
| Proliferating Cells | Diaph31 | 2.10E-38 | 7.05591044 | 0.972 | 0.13 | 4.19E-35 |
| Proliferating Cells | Kif20b1 | 6.54E-38 | 8.2530523 | 0.611 | 0.06 | 1.31E-34 |
| Proliferating Cells | Clspn | 3.05E-34 | 6.26461078 | 0.472 | 0.03 | 6.09E-31 |
| Proliferating Cells | Cenpf1 | 7.15E-34 | 8.5151528 | 0.861 | 0.143 | 1.43E-30 |
| Proliferating Cells | Sstr3 | 1.37E-33 | 2.0859862 | 0.278 | 0.003 | 2.74E-30 |
| Proliferating Cells | Ckap2 | 3.39E-33 | 8.51734455 | 0.528 | 0.07 | 6.78E-30 |

| | | | | | | |
|---|---|---|---|---|---|---|
| Proliferating Cells | Melk2 | 1.15E-32 | 7.47638623 | 0.583 | 0.051 | 2.29E-29 |
| Proliferating Cells | Arhgef39 | 2.31E-32 | 10.5001951 | 0.389 | 0.02 | 4.62E-29 |
| Proliferating Cells | Hmmr | 1.16E-31 | 8.07618549 | 0.667 | 0.044 | 2.32E-28 |
| Proliferating Cells | Sgo21 | 3.02E-29 | 7.83184055 | 0.611 | 0.054 | 6.03E-26 |
| Proliferating Cells | Cenpe1 | 4.86E-27 | 7.54451396 | 0.667 | 0.093 | 9.71E-24 |
| Proliferating Cells | Tpx2 | 9.20E-27 | 6.84170345 | 0.722 | 0.14 | 1.84E-23 |
| Proliferating Cells | Mki67 | 9.84E-27 | 6.96543076 | 0.778 | 0.16 | 1.97E-23 |
| Proliferating Cells | Aspm1 | 3.88E-26 | 8.11111744 | 0.806 | 0.113 | 7.77E-23 |
| Proliferating Cells | Kif4a1 | 5.47E-26 | 7.3809182 | 0.667 | 0.101 | 1.09E-22 |
| Proliferating Cells | Aurkb | 2.86E-24 | 7.62763578 | 0.472 | 0.099 | 5.73E-21 |
| Proliferating Cells | Kif231 | 2.87E-24 | 7.49566435 | 0.75 | 0.16 | 5.75E-21 |
| Proliferating Cells | Nav22 | 2.69E-23 | 4.3298212 | 1 | 0.498 | 5.38E-20 |
| Proliferating Cells | Arhgap11a | 1.06E-22 | 6.97239615 | 0.667 | 0.094 | 2.13E-19 |
| Proliferating Cells | Kif221 | 1.90E-21 | 7.12389847 | 0.639 | 0.125 | 3.80E-18 |

| | | | | | | |
|---|---|---|---|---|---|---|
| Proliferating Cells | Cdca21 | 2.05E-19 | 7.4677639 | 0.417 | 0.079 | 4.10E-16 |
| Proliferating Cells | Tbx12 | 3.79E-19 | 2.44877692 | 0.417 | 0.061 | 7.58E-16 |
| Proliferating Cells | E2f71 | 2.16E-17 | 6.85229525 | 0.639 | 0.131 | 4.31E-14 |
| Proliferating Cells | Ptprz11 | 1.05E-16 | 1.45445555 | 0.417 | 0.034 | 2.10E-13 |
| Proliferating Cells | Cit | 2.20E-16 | 5.99599599 | 0.778 | 0.184 | 4.41E-13 |
| Proliferating Cells | Itk | 3.56E-16 | 1.27546125 | 0.278 | 0.028 | 7.12E-13 |
| Proliferating Cells | Plk42 | 3.02E-15 | 6.10091307 | 0.444 | 0.032 | 6.04E-12 |
| Proliferating Cells | Bard1 | 5.95E-12 | 6.21404432 | 0.639 | 0.23 | 1.19E-08 |
| Proliferating Cells | Anln1 | 2.35E-11 | 6.21101844 | 0.444 | 0.194 | 4.70E-08 |
| Proliferating Cells | Epsti11 | 1.72E-10 | 2.03351981 | 0.417 | 0.064 | 3.43E-07 |
| Proliferating Cells | Cx3cr12 | 2.48E-10 | 2.19556304 | 0.278 | 0.023 | 4.96E-07 |
| Proliferating Cells | Klf5 | 5.17E-10 | 3.05271989 | 0.389 | 0.054 | 1.03E-06 |
| Proliferating Cells | Ndc80 | 6.36E-10 | 6.77453061 | 0.417 | 0.111 | 1.27E-06 |
| Proliferating Cells | Brip11 | 1.17E-09 | 5.69031891 | 0.639 | 0.281 | 2.34E-06 |

| | | | | | | |
|---|---|---|---|---|---|---|
| Proliferating Cells | Lef1 | 1.45E-09 | 3.01107976 | 0.417 | 0.089 | 2.90E-06 |
| Proliferating Cells | Dtx11 | 2.37E-09 | 3.11975046 | 0.417 | 0.109 | 4.74E-06 |
| Proliferating Cells | Slfn131 | 3.18E-09 | 3.39560202 | 0.722 | 0.301 | 6.35E-06 |
| Proliferating Cells | Ablim31 | 6.17E-09 | 1.86236387 | 0.861 | 0.47 | 1.23E-05 |
| Proliferating Cells | Cdh14 | 1.87E-08 | 3.77347851 | 0.417 | 0.113 | 3.73E-05 |
| Proliferating Cells | Mybpc21 | 4.65E-08 | 3.85647504 | 0.417 | 0.116 | 9.29E-05 |
| Proliferating Cells | Chst11 | 7.86E-08 | 2.14379072 | 0.806 | 0.472 | 0.00015711 |
| Proliferating Cells | Cd247 | 1.06E-07 | 0.81451851 | 0.278 | 0.006 | 0.00021233 |
| Proliferating Cells | Shank31 | 2.53E-07 | 1.3636207 | 0.861 | 0.567 | 0.00050511 |
| Proliferating Cells | Pcdh19 | 6.88E-07 | 2.19318384 | 0.75 | 0.415 | 0.00137546 |
| Proliferating Cells | Brca11 | 1.16E-06 | 4.90859258 | 0.556 | 0.222 | 0.00231773 |
| Proliferating Cells | Klrd1 | 1.44E-06 | 0.89039133 | 0.278 | 0.015 | 0.0028887 |
| Proliferating Cells | Cmtm8 | 2.28E-06 | 1.93176405 | 0.694 | 0.38 | 0.00455645 |
| Proliferating Cells | Adam12 | 3.03E-06 | 3.73182449 | 0.611 | 0.259 | 0.00606042 |

| | | | | | | |
|---|---|---|---|---|---|---|
| Proliferating Cells | AABR07049156.1 | 4.10E-06 | 4.6029034 | 0.417 | 0.154 | 0.00819898 |
| Proliferating Cells | LOC6900452 | 5.51E-06 | 0.41359949 | 0.417 | 0.097 | 0.01102484 |
| Proliferating Cells | Kank41 | 9.50E-06 | 2.88812219 | 0.444 | 0.06 | 0.0190032 |
| Proliferating Cells | Ptprb1 | 1.15E-05 | 1.09684878 | 0.833 | 0.515 | 0.02306906 |
| Proliferating Cells | Lrg12 | 1.59E-05 | 2.90776471 | 0.444 | 0.122 | 0.03175822 |
| Proliferating Cells | Cntn31 | 1.94E-05 | 2.55942013 | 0.417 | 0.102 | 0.03873363 |
| SMC | Dmp1 | 0 | 6.34665969 | 0.302 | 0.02 | 0 |
| SMC | Olr59 | 5.95E-295 | 7.10828742 | 0.45 | 0.039 | 1.19E-291 |
| SMC | Mgat3 | 9.91E-262 | 3.74749994 | 0.28 | 0.009 | 1.98E-258 |
| SMC | Mrvi1 | 1.05E-170 | 4.75687606 | 0.857 | 0.325 | 2.10E-167 |
| SMC | Stum | 2.62E-163 | 5.80077442 | 0.391 | 0.022 | 5.23E-160 |
| SMC | Notch3 | 1.32E-153 | 5.89197179 | 0.775 | 0.243 | 2.64E-150 |
| SMC | Gucy1a1 | 3.35E-130 | 3.61539337 | 0.817 | 0.419 | 6.70E-127 |
| SMC | Glipr1 | 4.02E-126 | 2.29272857 | 0.349 | 0.037 | 8.05E-123 |
| SMC | Pde3a1 | 6.64E-124 | 2.10425834 | 0.954 | 0.632 | 1.33E-120 |
| SMC | Mylk | 1.04E-123 | 2.92006475 | 0.868 | 0.507 | 2.09E-120 |
| SMC | Rasl12 | 1.04E-122 | 6.03808183 | 0.563 | 0.145 | 2.09E-119 |
| SMC | Pak3 | 9.89E-122 | 2.71042227 | 0.283 | 0.012 | 1.98E-118 |
| SMC | Gucy1a2 | 1.35E-117 | 3.0983728 | 0.826 | 0.463 | 2.70E-114 |
| SMC | Cyp4f18 | 1.53E-108 | 5.49780483 | 0.331 | 0.08 | 3.06E-105 |
| SMC | Pdgfrb | 2.19E-106 | 2.45798358 | 0.848 | 0.522 | 4.38E-103 |
| SMC | Ano1 | 2.20E-97 | 5.70804762 | 0.6 | 0.204 | 4.40E-94 |
| SMC | Col7a1 | 6.34E-93 | 3.03566375 | 0.331 | 0.025 | 1.27E-89 |

| | | | | | | |
|---|---|---|---|---|---|---|
| SMC | Acta2 | 1.32E-92 | 6.26642142 | 0.66 | 0.211 | 2.65E-89 |
| SMC | Cald1 | 5.66E-91 | 2.49343056 | 0.823 | 0.512 | 1.13E-87 |
| SMC | Myh11 | 1.10E-87 | 6.69645972 | 0.698 | 0.359 | 2.19E-84 |
| SMC | Acap1 | 1.84E-87 | 2.91678856 | 0.389 | 0.067 | 3.67E-84 |
| SMC | Trpc6 | 7.85E-80 | 4.84599958 | 0.411 | 0.131 | 1.57E-76 |
| SMC | Slco3a1 | 7.00E-76 | 1.83214889 | 0.854 | 0.588 | 1.40E-72 |
| SMC | Agap2 | 9.10E-76 | 5.15308003 | 0.313 | 0.045 | 1.82E-72 |
| SMC | Tbc1d1 | 1.86E-75 | 2.65417589 | 0.762 | 0.467 | 3.73E-72 |
| SMC | Mark1 | 1.67E-71 | 4.20966495 | 0.614 | 0.208 | 3.33E-68 |
| SMC | Kalrn | 7.14E-71 | 1.75525559 | 0.859 | 0.602 | 1.43E-67 |
| SMC | Ldb2 | 3.74E-69 | 2.03374696 | 0.74 | 0.443 | 7.47E-66 |
| SMC | Myo1b | 4.44E-63 | 2.77701098 | 0.735 | 0.479 | 8.89E-60 |
| SMC | Mcam | 2.90E-62 | 3.3293169 | 0.651 | 0.317 | 5.81E-59 |
| SMC | Rgs5 | 1.18E-58 | 5.14767963 | 0.576 | 0.316 | 2.35E-55 |
| SMC | Il34 | 4.65E-57 | 4.7170129 | 0.49 | 0.214 | 9.31E-54 |
| SMC | Dgkb | 3.40E-52 | 2.79135038 | 0.656 | 0.405 | 6.80E-49 |
| SMC | AABR07004228.1 | 9.08E-52 | 3.87186753 | 0.483 | 0.197 | 1.82E-48 |
| SMC | Pde5a | 2.76E-45 | 2.94778309 | 0.6 | 0.329 | 5.53E-42 |
| SMC | Lrrc4c | 1.28E-44 | 4.51834137 | 0.589 | 0.321 | 2.56E-41 |
| SMC | Flna | 6.56E-44 | 2.65875557 | 0.678 | 0.411 | 1.31E-40 |
| SMC | AABR07044900.11 | 8.26E-42 | 1.69815476 | 0.753 | 0.473 | 1.65E-38 |
| SMC | Actg2 | 2.34E-39 | 6.80318346 | 0.353 | 0.09 | 4.68E-36 |
| SMC | Tmcc3 | 3.68E-38 | 1.96153851 | 0.662 | 0.401 | 7.36E-35 |
| SMC | Sntb11 | 3.67E-26 | 1.64163458 | 0.629 | 0.354 | 7.34E-23 |
| SMC | Dlgap11 | 6.85E-10 | 0.25556304 | 0.221 | 0.476 | 1.37E-06 |
