## Supplemental Table 14 for "Uncovering the Regional and Cell Specific Bioactivity of Injectable Extracellular Matrix Biomaterials in Myocardial Infarction through Spatial and Single Nucleus Transcriptomics"

**Supplementary Table 14. Cell-Specific Differentially Expressed Genes with ECM (Up) or Saline (Down) Treatment in the Chronic MI Model**

| Subsetted Cell Type | Direction | Gene | p_val | avg_log2FC | pct.1 | pct.2 | p_val_adj |
|---|---|---|---|---|---|---|---|
| Endothelial Cells | Up | Tnnt2 | 1.33E-10 | 1.46750792 | 0.837 | 0.574 | 2.65E-07 |
| Endothelial Cells | Up | Actc1 | 7.99E-10 | 1.30270125 | 0.695 | 0.444 | 1.60E-06 |
| Endothelial Cells | Up | AABR07052585.1 | 7.99E-16 | 1.29671132 | 0.411 | 0.139 | 1.60E-12 |
| Endothelial Cells | Up | Tnnc1 | 5.08E-18 | 1.24475011 | 0.426 | 0.111 | 1.02E-14 |
| Endothelial Cells | Up | Epha4 | 1.86E-17 | 1.06277699 | 0.426 | 0.139 | 3.72E-14 |
| Endothelial Cells | Up | Rgs5 | 6.69E-06 | 0.58798057 | 0.39 | 0.759 | 0.01337602 |
| Endothelial Cells | Up | Clmn | 1.02E-05 | 0.56406286 | 0.277 | 0.704 | 0.02047709 |
| Endothelial Cells | Up | Macrod1 | 4.14E-07 | 0.56254873 | 0.34 | 0.75 | 0.00082893 |
| Endothelial Cells | Up | Specc1 | 1.06E-06 | 0.50136513 | 0.34 | 0.731 | 0.00211543 |
| Endothelial Cells | Up | Rsad2 | 7.37E-08 | 0.48115708 | 0.255 | 0.713 | 0.0001474 |
| Endothelial Cells | Up | Pdgfrb | 6.04E-10 | 0.46602874 | 0.362 | 0.759 | 1.21E-06 |
| Endothelial Cells | Up | Ptpn3 | 4.66E-06 | 0.45813766 | 0.199 | 0.62 | 0.00931937 |
| Endothelial Cells | Up | Cd83 | 1.74E-06 | 0.43156924 | 0.199 | 0.454 | 0.00348014 |
| Endothelial Cells | Up | Nceh1 | 1.35E-08 | 0.41432209 | 0.348 | 0.759 | 2.71E-05 |
| Endothelial Cells | Up | Cacna1a | 1.10E-06 | 0.36474127 | 0.27 | 0.676 | 0.00219502 |
| Endothelial Cells | Up | Grb14 | 6.33E-12 | 0.35454094 | 0.255 | 0.722 | 1.27E-08 |
| Endothelial Cells | Up | AC110709.2 | 2.99E-06 | 0.35048043 | 0.199 | 0.519 | 0.005978 |
| Endothelial Cells | Up | Pde8b | 2.42E-05 | 0.3473769 | 0.163 | 0.444 | 0.04837384 |
| Endothelial Cells | Up | Cux2 | 5.24E-09 | 0.34046558 | 0.227 | 0.611 | 1.05E-05 |
| Endothelial Cells | Up | Kcnn3 | 1.24E-11 | 0.33160891 | 0.255 | 0.741 | 2.49E-08 |
| Endothelial Cells | Up | Tesc | 3.79E-09 | 0.33017606 | 0.319 | 0.75 | 7.59E-06 |
| Endothelial Cells | Up | Parm1 | 1.66E-07 | 0.32013687 | 0.355 | 0.769 | 0.00033135 |
| Endothelial Cells | Up | Ralgps2 | 8.99E-23 | 0.31199019 | 0.213 | 0.778 | 1.80E-19 |
| Endothelial Cells | Up | Usp13 | 1.20E-06 | 0.30380984 | 0.262 | 0.685 | 0.00239974 |
| Endothelial Cells | Up | Iqub | 3.03E-07 | 0.28983902 | 0.348 | 0.759 | 0.00060582 |
| Endothelial Cells | Up | Ncam1 | 2.92E-13 | 0.27579499 | 0.248 | 0.731 | 5.83E-10 |
| Endothelial Cells | Up | Col6a2 | 1.06E-07 | 0.26432941 | 0.255 | 0.685 | 0.00021104 |
| Endothelial Cells | Up | Tmem26 | 2.63E-09 | 0.26224515 | 0.362 | 0.769 | 5.25E-06 |
| Endothelial Cells | Up | Trim50 | 2.57E-07 | 0.25724909 | 0.34 | 0.694 | 0.00051476 |
| Endothelial Cells | Up | Dlgap1 | 2.66E-09 | 0.25038932 | 0.277 | 0.704 | 5.31E-06 |
| Endothelial Cells | Down | Camk1d | 1.10E-08 | -0.25022249 | 0.262 | 0.704 | 2.20E-05 |
| Endothelial Cells | Down | Slit3 | 7.58E-17 | -0.25156874 | 0.248 | 0.75 | 1.52E-13 |
| Endothelial Cells | Down | Cacna1g | 5.38E-12 | -0.25186233 | 0.227 | 0.676 | 1.08E-08 |
| Endothelial Cells | Down | Kcnma1 | 1.56E-09 | -0.25930206 | 0.34 | 0.713 | 3.12E-06 |

| | | | | | | | |
|---|---|---|---|---|---|---|---|
| Endothelial Cells | Down | Sh3gl3 | 2.17E-08 | -0.27017611 | 0.035 | 0.454 | 4.35E-05 |
| Endothelial Cells | Down | Dip2c | 1.21E-07 | -0.27045886 | 0.39 | 0.806 | 0.00024287 |
| Endothelial Cells | Down | Col24a1 | 1.09E-08 | -0.275274 | 0.362 | 0.75 | 2.18E-05 |
| Endothelial Cells | Down | Itgb8 | 1.17E-15 | -0.27833427 | 0.284 | 0.769 | 2.34E-12 |
| Endothelial Cells | Down | Efhd1 | 7.78E-20 | -0.28162043 | 0.078 | 0.676 | 1.56E-16 |
| Endothelial Cells | Down | Cd274 | 7.67E-08 | -0.29703443 | 0.369 | 0.787 | 0.00015337 |
| Endothelial Cells | Down | Pcbp3 | 1.03E-15 | -0.29753358 | 0.291 | 0.759 | 2.05E-12 |
| Endothelial Cells | Down | Bdh1 | 4.72E-20 | -0.30417659 | 0.227 | 0.759 | 9.45E-17 |
| Endothelial Cells | Down | Abcc8 | 1.30E-19 | -0.30489874 | 0.028 | 0.546 | 2.60E-16 |
| Endothelial Cells | Down | Sulf1 | 1.04E-08 | -0.32078366 | 0.355 | 0.759 | 2.09E-05 |
| Endothelial Cells | Down | Glis3 | 2.20E-13 | -0.3340629 | 0.34 | 0.787 | 4.40E-10 |
| Endothelial Cells | Down | Rimbp2 | 4.73E-09 | -0.34690324 | 0.241 | 0.676 | 9.46E-06 |
| Endothelial Cells | Down | Slc25a13 | 2.51E-13 | -0.36325147 | 0.255 | 0.731 | 5.01E-10 |
| Endothelial Cells | Down | Hs3st5 | 8.51E-08 | -0.37303029 | 0.156 | 0.611 | 0.00017011 |
| Endothelial Cells | Down | Adam19 | 1.91E-06 | -0.37646374 | 0.397 | 0.769 | 0.00382796 |
| Endothelial Cells | Down | Ppip5k1 | 2.86E-09 | -0.38648079 | 0.277 | 0.741 | 5.72E-06 |
| Endothelial Cells | Down | Dipk1a | 1.44E-13 | -0.38917477 | 0.312 | 0.796 | 2.89E-10 |
| Endothelial Cells | Down | Gda | 2.79E-06 | -0.43347131 | 0.433 | 0.815 | 0.00557307 |
| Endothelial Cells | Down | Fam78b | 5.26E-07 | -0.44516277 | 0.461 | 0.806 | 0.00105273 |
| Endothelial Cells | Down | Matn2 | 2.85E-14 | -0.44913911 | 0.326 | 0.796 | 5.70E-11 |
| Endothelial Cells | Down | Antxr1 | 1.48E-14 | -0.45547021 | 0.355 | 0.769 | 2.96E-11 |
| Endothelial Cells | Down | Sgip1 | 4.36E-11 | -0.46038757 | 0.348 | 0.778 | 8.71E-08 |
| Endothelial Cells | Down | Fgf10 | 5.73E-08 | -0.4678067 | 0.142 | 0.593 | 0.00011452 |
| Endothelial Cells | Down | Ppm1l | 6.50E-19 | -0.49326884 | 0.333 | 0.796 | 1.30E-15 |
| Endothelial Cells | Down | Tox3 | 9.91E-06 | -0.50216586 | 0.482 | 0.824 | 0.01981258 |
| Endothelial Cells | Down | Rerg | 5.06E-07 | -0.63602291 | 0.191 | 0.63 | 0.0010112 |
| Endothelial Cells | Down | Tspan18 | 4.08E-06 | -0.66962877 | 0.355 | 0.741 | 0.00816258 |
| Endothelial Cells | Down | Papss2 | 3.88E-06 | -0.67215087 | 0.504 | 0.87 | 0.00776685 |
| Endothelial Cells | Down | Casz1 | 6.95E-12 | -0.76413441 | 0.39 | 0.778 | 1.39E-08 |
| Endothelial Cells | Down | Nxn | 1.06E-09 | -0.93530514 | 0.504 | 0.861 | 2.12E-06 |
| Endothelial Cells | Down | Bmp6 | 1.71E-07 | -0.95841258 | 0.61 | 0.926 | 0.00034271 |
| Cardiomyocytes | Up | Slc6a6 | 3.34E-11 | 1.20418476 | 0.289 | 0.868 | 6.67E-08 |
| Cardiomyocytes | Up | Atp2b2 | 2.24E-09 | 0.86380483 | 0.305 | 0.853 | 4.47E-06 |
| Cardiomyocytes | Up | Nrxn1 | 9.25E-16 | 0.71828171 | 0.242 | 0.865 | 1.85E-12 |
| Cardiomyocytes | Up | Nxn | 3.13E-17 | 0.70319742 | 0.18 | 0.812 | 6.26E-14 |
| Cardiomyocytes | Up | Cacna1g | 1.19E-07 | 0.66387688 | 0.133 | 0.653 | 0.00023767 |
| Cardiomyocytes | Up | Sema5a | 2.82E-12 | 0.64588078 | 0.227 | 0.809 | 5.63E-09 |
| Cardiomyocytes | Up | Ebf2 | 4.86E-19 | 0.63605731 | 0.203 | 0.853 | 9.72E-16 |

| | | | | | | | |
|---|---|---|---|---|---|---|---|
| Cardiomyocytes | Up | Smpx | 3.35E-06 | 0.63602569 | 0.328 | 0.826 | 0.00670159 |
| Cardiomyocytes | Up | Sox5 | 4.28E-29 | 0.62621705 | 0.109 | 0.829 | 8.55E-26 |
| Cardiomyocytes | Up | Timp3 | 4.17E-10 | 0.62002252 | 0.219 | 0.771 | 8.35E-07 |
| Cardiomyocytes | Up | Ndufa4 | 1.42E-12 | 0.58383268 | 0.164 | 0.753 | 2.84E-09 |
| Cardiomyocytes | Up | Ntn4 | 6.61E-11 | 0.58324434 | 0.281 | 0.874 | 1.32E-07 |
| Cardiomyocytes | Up | Epb41l4b | 1.16E-11 | 0.57662457 | 0.289 | 0.876 | 2.31E-08 |
| Cardiomyocytes | Up | Ncam1 | 1.48E-25 | 0.57613859 | 0.133 | 0.829 | 2.96E-22 |
| Cardiomyocytes | Up | Bche | 2.61E-19 | 0.55885662 | 0.141 | 0.8 | 5.23E-16 |
| Cardiomyocytes | Up | Tmem182 | 1.61E-07 | 0.55230179 | 0.336 | 0.853 | 0.00032252 |
| Cardiomyocytes | Up | Plxdc2 | 1.83E-15 | 0.55063593 | 0.195 | 0.818 | 3.66E-12 |
| Cardiomyocytes | Up | Adgrl3 | 3.65E-13 | 0.5456364 | 0.102 | 0.568 | 7.30E-10 |
| Cardiomyocytes | Up | Slc16a10 | 5.29E-13 | 0.52754168 | 0.242 | 0.844 | 1.06E-09 |
| Cardiomyocytes | Up | Khdrbs3 | 6.94E-09 | 0.52585518 | 0.102 | 0.638 | 1.39E-05 |
| Cardiomyocytes | Up | Slit3 | 2.28E-25 | 0.52448611 | 0.109 | 0.771 | 4.56E-22 |
| Cardiomyocytes | Up | Gnao1 | 9.33E-08 | 0.51624736 | 0.344 | 0.871 | 0.00018663 |
| Cardiomyocytes | Up | Npr3 | 3.31E-29 | 0.50530388 | 0.102 | 0.826 | 6.63E-26 |
| Cardiomyocytes | Up | Rerg | 3.33E-06 | 0.49642008 | 0.078 | 0.524 | 0.00666766 |
| Cardiomyocytes | Up | Chst15 | 9.05E-16 | 0.49558354 | 0.211 | 0.821 | 1.81E-12 |
| Cardiomyocytes | Up | Kit | 2.94E-17 | 0.49368042 | 0.109 | 0.647 | 5.87E-14 |
| Cardiomyocytes | Up | Ikzf2 | 1.63E-22 | 0.49322373 | 0.172 | 0.85 | 3.27E-19 |
| Cardiomyocytes | Up | AABR07058170.1 | 2.29E-16 | 0.47455839 | 0.109 | 0.694 | 4.59E-13 |
| Cardiomyocytes | Up | Fbxl2 | 2.01E-23 | 0.46651535 | 0.117 | 0.806 | 4.03E-20 |
| Cardiomyocytes | Up | Tmeff2 | 2.11E-27 | 0.4646207 | 0.133 | 0.847 | 4.22E-24 |
| Cardiomyocytes | Up | Trim7 | 4.03E-20 | 0.45693463 | 0.18 | 0.838 | 8.06E-17 |
| Cardiomyocytes | Up | Creb5 | 1.48E-14 | 0.44792196 | 0.227 | 0.832 | 2.96E-11 |
| Cardiomyocytes | Up | Srgap1 | 7.95E-26 | 0.44108553 | 0.148 | 0.844 | 1.59E-22 |
| Cardiomyocytes | Up | Dcn | 2.75E-25 | 0.43988485 | 0.125 | 0.829 | 5.50E-22 |
| Cardiomyocytes | Up | AABR07058158.1 | 5.63E-22 | 0.43902966 | 0.125 | 0.779 | 1.13E-18 |
| Cardiomyocytes | Up | Zfp385d | 7.53E-24 | 0.43866737 | 0.125 | 0.821 | 1.51E-20 |
| Cardiomyocytes | Up | Col18a1 | 6.38E-16 | 0.430773 | 0.086 | 0.638 | 1.28E-12 |
| Cardiomyocytes | Up | Itgbl1 | 1.81E-19 | 0.42243282 | 0.18 | 0.841 | 3.62E-16 |
| Cardiomyocytes | Up | Eva1c | 1.74E-06 | 0.41282449 | 0.266 | 0.768 | 0.00347337 |
| Cardiomyocytes | Up | Ppm1e | 4.89E-10 | 0.4114849 | 0.148 | 0.706 | 9.78E-07 |
| Cardiomyocytes | Up | AABR07031740.1 | 4.47E-06 | 0.41085712 | 0.219 | 0.721 | 0.00894313 |
| Cardiomyocytes | Up | Ednra | 1.12E-11 | 0.40660247 | 0.258 | 0.835 | 2.23E-08 |
| Cardiomyocytes | Up | Mgmt | 6.73E-09 | 0.40572265 | 0.109 | 0.659 | 1.35E-05 |
| Cardiomyocytes | Up | Col3a1 | 1.81E-16 | 0.40489427 | 0.188 | 0.809 | 3.62E-13 |
| Cardiomyocytes | Up | Lgr6 | 1.26E-06 | 0.39794364 | 0.273 | 0.779 | 0.00251441 |

| | | | | | | | |
|---|---|---|---|---|---|---|---|
| Cardiomyocytes | Up | Fbn1 | 2.67E-20 | 0.39775753 | 0.188 | 0.841 | 5.33E-17 |
| Cardiomyocytes | Up | Irs1 | 8.01E-13 | 0.39719725 | 0.242 | 0.832 | 1.60E-09 |
| Cardiomyocytes | Up | Dusp5 | 9.88E-34 | 0.39531084 | 0.07 | 0.809 | 1.98E-30 |
| Cardiomyocytes | Up | Samd12 | 1.88E-07 | 0.38840959 | 0.289 | 0.797 | 0.000376 |
| Cardiomyocytes | Up | Fgf13 | 1.28E-18 | 0.38831653 | 0.164 | 0.818 | 2.57E-15 |
| Cardiomyocytes | Up | Trps1 | 1.13E-18 | 0.38708775 | 0.211 | 0.856 | 2.26E-15 |
| Cardiomyocytes | Up | Fam189a2 | 2.54E-07 | 0.3868859 | 0.156 | 0.679 | 0.00050792 |
| Cardiomyocytes | Up | Myo1b | 1.40E-10 | 0.38138862 | 0.133 | 0.706 | 2.80E-07 |
| Cardiomyocytes | Up | Scn7a | 1.95E-20 | 0.37685212 | 0.109 | 0.765 | 3.90E-17 |
| Cardiomyocytes | Up | LOC103694210 | 7.68E-09 | 0.37180001 | 0.078 | 0.506 | 1.54E-05 |
| Cardiomyocytes | Up | Bicc1 | 3.94E-16 | 0.36876869 | 0.219 | 0.847 | 7.88E-13 |
| Cardiomyocytes | Up | Rtn4rl1 | 1.00E-20 | 0.3653382 | 0.172 | 0.85 | 2.01E-17 |
| Cardiomyocytes | Up | Gucy1a1 | 4.56E-33 | 0.36477668 | 0.094 | 0.829 | 9.13E-30 |
| Cardiomyocytes | Up | Ar | 1.20E-30 | 0.35999793 | 0.07 | 0.806 | 2.39E-27 |
| Cardiomyocytes | Up | Efna5 | 1.64E-27 | 0.35916526 | 0.062 | 0.691 | 3.28E-24 |
| Cardiomyocytes | Up | Cacna1d | 3.78E-25 | 0.35848404 | 0.086 | 0.779 | 7.56E-22 |
| Cardiomyocytes | Up | Tesc | 3.51E-06 | 0.3577978 | 0.398 | 0.885 | 0.0070124 |
| Cardiomyocytes | Up | Slco5a1 | 1.47E-08 | 0.35234174 | 0.32 | 0.856 | 2.95E-05 |
| Cardiomyocytes | Up | Nrg1 | 2.98E-34 | 0.346062 | 0.094 | 0.838 | 5.95E-31 |
| Cardiomyocytes | Up | Gpr176 | 1.12E-08 | 0.34522379 | 0.07 | 0.344 | 2.23E-05 |
| Cardiomyocytes | Up | Astn2 | 3.72E-30 | 0.34493562 | 0.102 | 0.838 | 7.44E-27 |
| Cardiomyocytes | Up | Adamts3 | 1.16E-09 | 0.33982685 | 0.086 | 0.612 | 2.31E-06 |
| Cardiomyocytes | Up | Bcl11a | 3.92E-17 | 0.33759792 | 0.203 | 0.85 | 7.84E-14 |
| Cardiomyocytes | Up | Fgf12 | 7.41E-12 | 0.3300791 | 0.289 | 0.859 | 1.48E-08 |
| Cardiomyocytes | Up | Tmtc2 | 1.68E-22 | 0.32687817 | 0.125 | 0.815 | 3.36E-19 |
| Cardiomyocytes | Up | Eepd1 | 4.45E-13 | 0.32630879 | 0.195 | 0.791 | 8.89E-10 |
| Cardiomyocytes | Up | Fli1 | 2.68E-12 | 0.31926569 | 0.148 | 0.726 | 5.37E-09 |
| Cardiomyocytes | Up | Prrx1 | 1.20E-15 | 0.31918498 | 0.086 | 0.553 | 2.41E-12 |
| Cardiomyocytes | Up | Slfn13 | 3.40E-31 | 0.31838826 | 0.055 | 0.768 | 6.80E-28 |
| Cardiomyocytes | Up | Pak1 | 6.34E-11 | 0.31211981 | 0.008 | 0.297 | 1.27E-07 |
| Cardiomyocytes | Up | Enox2 | 4.46E-17 | 0.30784125 | 0.125 | 0.753 | 8.92E-14 |
| Cardiomyocytes | Up | AC111831.1 | 7.41E-28 | 0.30512939 | 0.094 | 0.776 | 1.48E-24 |
| Cardiomyocytes | Up | Slc16a1 | 8.05E-14 | 0.30211147 | 0.234 | 0.835 | 1.61E-10 |
| Cardiomyocytes | Up | Hcn2 | 2.12E-05 | 0.29803992 | 0.086 | 0.591 | 0.04234022 |
| Cardiomyocytes | Up | Clybl | 2.17E-15 | 0.2980225 | 0.25 | 0.874 | 4.34E-12 |
| Cardiomyocytes | Up | Fstl1 | 5.94E-10 | 0.29365723 | 0.102 | 0.656 | 1.19E-06 |
| Cardiomyocytes | Up | Dapk1 | 6.04E-26 | 0.28806025 | 0.109 | 0.812 | 1.21E-22 |
| Cardiomyocytes | Up | Ttll7 | 1.10E-12 | 0.28785291 | 0.086 | 0.682 | 2.20E-09 |

| | | | | | | | |
|---|---|---|---|---|---|---|---|
| Cardiomyocytes | Up | Apba1 | 1.11E-25 | 0.28511962 | 0.07 | 0.756 | 2.22E-22 |
| Cardiomyocytes | Up | Ackr3 | 3.45E-32 | 0.28369227 | 0.062 | 0.794 | 6.89E-29 |
| Cardiomyocytes | Up | Cspg4 | 3.14E-09 | 0.28368752 | 0.086 | 0.647 | 6.28E-06 |
| Cardiomyocytes | Up | Lekr1 | 7.32E-11 | 0.2832595 | 0.141 | 0.712 | 1.46E-07 |
| Cardiomyocytes | Up | Lrrc4c | 1.19E-31 | 0.27669304 | 0.07 | 0.815 | 2.38E-28 |
| Cardiomyocytes | Up | Dlgap1 | 7.48E-18 | 0.2745227 | 0.203 | 0.85 | 1.50E-14 |
| Cardiomyocytes | Up | Abca1 | 1.87E-19 | 0.27362138 | 0.141 | 0.8 | 3.74E-16 |
| Cardiomyocytes | Up | Gabrb2 | 4.79E-17 | 0.27302247 | 0.086 | 0.721 | 9.57E-14 |
| Cardiomyocytes | Up | Ltbp1 | 5.63E-24 | 0.27193796 | 0.164 | 0.847 | 1.13E-20 |
| Cardiomyocytes | Up | St3gal5 | 6.83E-06 | 0.2680049 | 0.242 | 0.735 | 0.01366767 |
| Cardiomyocytes | Up | AABR07049292.1 | 8.57E-22 | 0.26790921 | 0.164 | 0.841 | 1.71E-18 |
| Cardiomyocytes | Up | Stk17b | 1.46E-30 | 0.26660747 | 0.078 | 0.788 | 2.93E-27 |
| Cardiomyocytes | Up | Gclc | 5.82E-31 | 0.26410694 | 0.094 | 0.821 | 1.16E-27 |
| Cardiomyocytes | Up | Il1rapl1 | 4.69E-23 | 0.26169031 | 0.047 | 0.606 | 9.38E-20 |
| Cardiomyocytes | Up | Sgip1 | 5.58E-29 | 0.26060142 | 0.055 | 0.744 | 1.12E-25 |
| Cardiomyocytes | Up | Zfp385b | 2.08E-05 | 0.25787236 | 0.055 | 0.562 | 0.04164003 |
| Cardiomyocytes | Up | Rnf213 | 6.53E-25 | 0.25740881 | 0.141 | 0.85 | 1.31E-21 |
| Cardiomyocytes | Up | Zfhx3 | 2.47E-19 | 0.25603389 | 0.117 | 0.782 | 4.94E-16 |
| Cardiomyocytes | Up | Col27a1 | 3.53E-28 | 0.25568513 | 0.078 | 0.797 | 7.06E-25 |
| Cardiomyocytes | Up | Pank1 | 2.82E-09 | 0.25032062 | 0.266 | 0.8 | 5.64E-06 |
| Cardiomyocytes | Up | Meox1 | 1.20E-14 | 0.25031956 | 0.023 | 0.403 | 2.40E-11 |
| Cardiomyocytes | Up | Cyyr1 | 2.56E-21 | 0.25004229 | 0.148 | 0.809 | 5.11E-18 |
| Cardiomyocytes | Down | Bard1 | 2.28E-40 | -0.25147639 | 0.039 | 0.803 | 4.57E-37 |
| Cardiomyocytes | Down | Slc9a3r2 | 1.48E-14 | -0.255437 | 0.031 | 0.647 | 2.97E-11 |
| Cardiomyocytes | Down | Samd14 | 6.24E-20 | -0.25606737 | 0.047 | 0.709 | 1.25E-16 |
| Cardiomyocytes | Down | Mgll | 1.50E-26 | -0.26005275 | 0.156 | 0.862 | 3.01E-23 |
| Cardiomyocytes | Down | Man1c1 | 9.55E-07 | -0.2617507 | 0.445 | 0.921 | 0.00191011 |
| Cardiomyocytes | Down | AABR07006724.1 | 3.61E-27 | -0.26483118 | 0.078 | 0.785 | 7.22E-24 |
| Cardiomyocytes | Down | Slc26a10 | 4.80E-24 | -0.26796015 | 0.031 | 0.721 | 9.60E-21 |
| Cardiomyocytes | Down | Hecw2 | 1.06E-16 | -0.26847748 | 0.172 | 0.791 | 2.12E-13 |
| Cardiomyocytes | Down | Atp5mc3 | 3.97E-26 | -0.28190544 | 0.164 | 0.859 | 7.94E-23 |
| Cardiomyocytes | Down | Dpt | 9.38E-32 | -0.29107059 | 0.039 | 0.771 | 1.88E-28 |
| Cardiomyocytes | Down | Enox1 | 1.59E-28 | -0.29130122 | 0.047 | 0.776 | 3.18E-25 |
| Cardiomyocytes | Down | Arhgap44 | 8.78E-09 | -0.29343146 | 0.258 | 0.779 | 1.76E-05 |
| Cardiomyocytes | Down | Pmepa1 | 1.04E-33 | -0.29481137 | 0.086 | 0.835 | 2.09E-30 |
| Cardiomyocytes | Down | Asb14 | 8.99E-16 | -0.31630293 | 0.156 | 0.768 | 1.80E-12 |
| Cardiomyocytes | Down | Flnc | 2.14E-20 | -0.32224802 | 0.211 | 0.879 | 4.28E-17 |
| Cardiomyocytes | Down | Dip2c | 1.24E-06 | -0.32436088 | 0.5 | 0.938 | 0.00247882 |

| | | | | | | | |
|---|---|---|---|---|---|---|---|
| Cardiomyocytes | Down | Rimbp2 | 1.13E-17 | -0.33402986 | 0.164 | 0.8 | 2.27E-14 |
| Cardiomyocytes | Down | Pdzd2 | 5.95E-07 | -0.33633182 | 0.477 | 0.938 | 0.00118959 |
| Cardiomyocytes | Down | Ablim2 | 5.00E-22 | -0.34178636 | 0.102 | 0.782 | 9.99E-19 |
| Cardiomyocytes | Down | Kcng2 | 1.52E-11 | -0.34648255 | 0.359 | 0.903 | 3.03E-08 |
| Cardiomyocytes | Down | Lbh | 7.10E-15 | -0.347166 | 0.32 | 0.912 | 1.42E-11 |
| Cardiomyocytes | Down | Cacna1a | 8.98E-10 | -0.35307139 | 0.32 | 0.85 | 1.80E-06 |
| Cardiomyocytes | Down | Fabp3 | 5.04E-06 | -0.36344226 | 0.398 | 0.85 | 0.01007819 |
| Cardiomyocytes | Down | Clic5 | 2.02E-07 | -0.36344721 | 0.422 | 0.906 | 0.00040318 |
| Cardiomyocytes | Down | Hspb8 | 5.03E-12 | -0.36484222 | 0.141 | 0.726 | 1.01E-08 |
| Cardiomyocytes | Down | Ncald | 1.41E-37 | -0.36675951 | 0.062 | 0.824 | 2.82E-34 |
| Cardiomyocytes | Down | LOC100361087 | 2.67E-25 | -0.36821643 | 0.172 | 0.862 | 5.34E-22 |
| Cardiomyocytes | Down | Bdnf | 9.19E-31 | -0.3720223 | 0.117 | 0.871 | 1.84E-27 |
| Cardiomyocytes | Down | Nfatc2 | 2.34E-16 | -0.37274359 | 0.289 | 0.885 | 4.67E-13 |
| Cardiomyocytes | Down | Casz1 | 1.57E-18 | -0.37562074 | 0.211 | 0.85 | 3.15E-15 |
| Cardiomyocytes | Down | Sned1 | 4.06E-22 | -0.37784563 | 0.047 | 0.735 | 8.12E-19 |
| Cardiomyocytes | Down | Itga7 | 2.03E-09 | -0.38363459 | 0.398 | 0.882 | 4.07E-06 |
| Cardiomyocytes | Down | Galnt17 | 3.33E-42 | -0.38914809 | 0.031 | 0.824 | 6.66E-39 |
| Cardiomyocytes | Down | AABR07040864.1 | 1.45E-21 | -0.39614674 | 0.109 | 0.771 | 2.90E-18 |
| Cardiomyocytes | Down | Akr1c15 | 1.64E-08 | -0.39984371 | 0.219 | 0.744 | 3.28E-05 |
| Cardiomyocytes | Down | Pstpip2 | 3.07E-09 | -0.42022559 | 0.109 | 0.653 | 6.15E-06 |
| Cardiomyocytes | Down | Atp5mc1 | 1.45E-30 | -0.43008494 | 0.086 | 0.818 | 2.90E-27 |
| Cardiomyocytes | Down | Pdgfd | 7.84E-39 | -0.43732166 | 0.078 | 0.859 | 1.57E-35 |
| Cardiomyocytes | Down | Rmdn1 | 4.08E-07 | -0.44195653 | 0.359 | 0.821 | 0.0008168 |
| Cardiomyocytes | Down | Crim1 | 3.22E-10 | -0.44787705 | 0.422 | 0.947 | 6.45E-07 |
| Cardiomyocytes | Down | Meox2 | 3.75E-37 | -0.45180841 | 0.047 | 0.818 | 7.51E-34 |
| Cardiomyocytes | Down | Man1a1 | 4.83E-16 | -0.45508821 | 0.281 | 0.903 | 9.66E-13 |
| Cardiomyocytes | Down | Mtss1 | 7.42E-06 | -0.47628975 | 0.711 | 0.979 | 0.01483957 |
| Cardiomyocytes | Down | Angpt1 | 1.27E-15 | -0.47850161 | 0.289 | 0.894 | 2.55E-12 |
| Cardiomyocytes | Down | Cux2 | 1.02E-14 | -0.48252004 | 0.328 | 0.926 | 2.03E-11 |
| Cardiomyocytes | Down | Cxcl12 | 3.39E-30 | -0.50130256 | 0.016 | 0.726 | 6.78E-27 |
| Cardiomyocytes | Down | Ube2ql1 | 4.34E-28 | -0.52125471 | 0.086 | 0.803 | 8.68E-25 |
| Cardiomyocytes | Down | Gpcpd1 | 6.88E-11 | -0.53110378 | 0.352 | 0.882 | 1.38E-07 |
| Cardiomyocytes | Down | Ppargc1b | 1.16E-06 | -0.53557289 | 0.578 | 0.926 | 0.00231009 |
| Cardiomyocytes | Down | Rxrg | 3.66E-35 | -0.55670742 | 0.102 | 0.862 | 7.32E-32 |
| Cardiomyocytes | Down | Abcb4 | 1.28E-22 | -0.55768397 | 0.227 | 0.897 | 2.57E-19 |
| Cardiomyocytes | Down | Ptprd | 5.71E-07 | -0.55903211 | 0.492 | 0.894 | 0.00114149 |
| Cardiomyocytes | Down | Kif21a | 1.26E-23 | -0.5751444 | 0.086 | 0.782 | 2.51E-20 |
| Cardiomyocytes | Down | Rrad | 3.36E-19 | -0.58463719 | 0.227 | 0.871 | 6.73E-16 |

| | | | | | | | |
|---|---|---|---|---|---|---|---|
| Cardiomyocytes | Down | AABR07052585.2 | 5.32E-16 | -0.58578977 | 0.25 | 0.856 | 1.06E-12 |
| Cardiomyocytes | Down | Drc3 | 4.32E-29 | -0.58879852 | 0.07 | 0.812 | 8.65E-26 |
| Cardiomyocytes | Down | Atp5me | 7.41E-35 | -0.61808802 | 0.078 | 0.853 | 1.48E-31 |
| Cardiomyocytes | Down | Mtus2 | 3.35E-32 | -0.63529855 | 0.125 | 0.865 | 6.70E-29 |
| Cardiomyocytes | Down | Nnt | 4.88E-08 | -0.64327319 | 0.5 | 0.941 | 9.75E-05 |
| Cardiomyocytes | Down | Actn2 | 1.04E-07 | -0.6599641 | 0.617 | 0.974 | 0.00020857 |
| Cardiomyocytes | Down | Slc25a20 | 5.39E-18 | -0.67121854 | 0.125 | 0.774 | 1.08E-14 |
| Cardiomyocytes | Down | Coro6 | 4.08E-06 | -0.67619009 | 0.43 | 0.824 | 0.00816213 |
| Fibroblasts | Up | Pdgfd | 4.57E-06 | 0.57954525 | 0.309 | 0.596 | 0.00913468 |
| Fibroblasts | Up | Prodh1 | 4.66E-10 | 0.49895353 | 0.257 | 0.577 | 9.32E-07 |
| Fibroblasts | Up | Macrod1 | 1.62E-06 | 0.46022432 | 0.275 | 0.543 | 0.00323376 |
| Fibroblasts | Up | Efemp1 | 8.38E-06 | 0.44984516 | 0.363 | 0.644 | 0.01676107 |
| Fibroblasts | Up | Pde7a | 5.46E-07 | 0.44963737 | 0.262 | 0.558 | 0.00109152 |
| Fibroblasts | Up | Cp | 6.19E-12 | 0.44747671 | 0.191 | 0.524 | 1.24E-08 |
| Fibroblasts | Up | Lrrc2 | 2.28E-06 | 0.42556768 | 0.245 | 0.524 | 0.00455183 |
| Fibroblasts | Up | Cntfr | 2.47E-12 | 0.41573958 | 0.164 | 0.49 | 4.95E-09 |
| Fibroblasts | Up | Tnnc1 | 2.09E-10 | 0.40363026 | 0.319 | 0.606 | 4.18E-07 |
| Fibroblasts | Up | Nox4 | 2.42E-13 | 0.4016377 | 0.223 | 0.534 | 4.84E-10 |
| Fibroblasts | Up | Igfbp3 | 2.93E-11 | 0.36074173 | 0.167 | 0.476 | 5.86E-08 |
| Fibroblasts | Up | Actn2 | 1.96E-08 | 0.35408117 | 0.208 | 0.5 | 3.93E-05 |
| Fibroblasts | Up | Tpt1 | 7.99E-09 | 0.35325228 | 0.255 | 0.548 | 1.60E-05 |
| Fibroblasts | Up | Parm1 | 2.56E-10 | 0.35073846 | 0.164 | 0.462 | 5.12E-07 |
| Fibroblasts | Up | Ppargc1a | 9.11E-08 | 0.34447963 | 0.225 | 0.514 | 0.0001821 |
| Fibroblasts | Up | Rgs5 | 3.99E-08 | 0.34351131 | 0.189 | 0.462 | 7.98E-05 |
| Fibroblasts | Up | RGD1565355 | 6.54E-06 | 0.3402625 | 0.355 | 0.639 | 0.01308295 |
| Fibroblasts | Up | Grik2 | 2.44E-10 | 0.33795131 | 0.314 | 0.596 | 4.87E-07 |
| Fibroblasts | Up | Ldhb | 3.72E-12 | 0.33687049 | 0.248 | 0.587 | 7.43E-09 |
| Fibroblasts | Up | Acyp2 | 9.16E-07 | 0.33560396 | 0.304 | 0.591 | 0.0018328 |
| Fibroblasts | Up | Fabp4 | 1.89E-07 | 0.33304245 | 0.257 | 0.534 | 0.00037761 |
| Fibroblasts | Up | Maob | 1.30E-06 | 0.31707996 | 0.208 | 0.49 | 0.00259782 |
| Fibroblasts | Up | Nppa | 2.60E-10 | 0.31425502 | 0.275 | 0.567 | 5.21E-07 |
| Fibroblasts | Up | Myh7 | 3.06E-06 | 0.31163979 | 0.382 | 0.668 | 0.00611377 |
| Fibroblasts | Up | Ptn | 4.19E-16 | 0.29465221 | 0.135 | 0.49 | 8.38E-13 |
| Fibroblasts | Up | AABR07033925.1 | 1.13E-26 | 0.29211895 | 0.091 | 0.529 | 2.26E-23 |
| Fibroblasts | Up | Actc1 | 1.54E-05 | 0.2914712 | 0.181 | 0.457 | 0.03076941 |
| Fibroblasts | Up | Cst3 | 9.37E-09 | 0.27745294 | 0.186 | 0.481 | 1.87E-05 |
| Fibroblasts | Up | Dach1 | 1.23E-05 | 0.27389876 | 0.363 | 0.63 | 0.02462428 |
| Fibroblasts | Up | Fbln5 | 4.28E-09 | 0.27189833 | 0.267 | 0.572 | 8.55E-06 |

| | | | | | | | |
|---|---|---|---|---|---|---|---|
| Fibroblasts | Up | Ldb2 | 2.81E-08 | 0.26858493 | 0.216 | 0.524 | 5.62E-05 |
| Fibroblasts | Up | Myl2 | 4.81E-08 | 0.264273 | 0.292 | 0.587 | 9.62E-05 |
| Fibroblasts | Up | Ckmt2 | 1.84E-06 | 0.2541063 | 0.27 | 0.543 | 0.00367613 |
| Fibroblasts | Up | Ndufa4 | 4.94E-14 | 0.25061785 | 0.238 | 0.577 | 9.87E-11 |
| Fibroblasts | Down | Lyn | 5.12E-08 | -0.259412 | 0.255 | 0.553 | 0.00010249 |
| Fibroblasts | Down | Itga9 | 6.92E-07 | -0.2610415 | 0.397 | 0.678 | 0.00138429 |
| Fibroblasts | Down | Alpk3 | 9.68E-09 | -0.26352151 | 0.142 | 0.442 | 1.94E-05 |
| Fibroblasts | Down | Cdkn1a | 2.51E-19 | -0.26733973 | 0.159 | 0.514 | 5.01E-16 |
| Fibroblasts | Down | Nav3 | 2.20E-07 | -0.27239305 | 0.368 | 0.644 | 0.00044038 |
| Fibroblasts | Down | Fhod3 | 6.63E-10 | -0.27514507 | 0.233 | 0.567 | 1.33E-06 |
| Fibroblasts | Down | Eif2ak2 | 5.29E-16 | -0.27631369 | 0.297 | 0.635 | 1.06E-12 |
| Fibroblasts | Down | Pkhd1l1 | 1.32E-10 | -0.27694837 | 0.279 | 0.596 | 2.64E-07 |
| Fibroblasts | Down | Pamr1 | 1.23E-16 | -0.28416335 | 0.228 | 0.572 | 2.45E-13 |
| Fibroblasts | Down | Lyve1 | 4.09E-32 | -0.29066156 | 0.11 | 0.543 | 8.19E-29 |
| Fibroblasts | Down | Ncam1 | 1.25E-23 | -0.29681253 | 0.26 | 0.63 | 2.51E-20 |
| Fibroblasts | Down | Bgn | 2.71E-16 | -0.29747754 | 0.169 | 0.553 | 5.42E-13 |
| Fibroblasts | Down | Cit | 6.33E-07 | -0.29751747 | 0.135 | 0.418 | 0.0012666 |
| Fibroblasts | Down | Ptprj | 1.11E-06 | -0.30283662 | 0.468 | 0.736 | 0.00222156 |
| Fibroblasts | Down | Ntn1 | 1.31E-07 | -0.303169 | 0.417 | 0.702 | 0.00026207 |
| Fibroblasts | Down | Atp1a1 | 2.05E-18 | -0.31905185 | 0.235 | 0.601 | 4.10E-15 |
| Fibroblasts | Down | Itpkb | 1.01E-06 | -0.33613357 | 0.395 | 0.673 | 0.00201832 |
| Fibroblasts | Down | Veph1 | 5.91E-18 | -0.33626145 | 0.083 | 0.447 | 1.18E-14 |
| Fibroblasts | Down | Ppp1r12b | 6.23E-10 | -0.3466974 | 0.444 | 0.76 | 1.25E-06 |
| Fibroblasts | Down | Syt1 | 1.50E-07 | -0.34843833 | 0.064 | 0.365 | 0.00029915 |
| Fibroblasts | Down | Maoa | 2.44E-19 | -0.34898006 | 0.279 | 0.635 | 4.89E-16 |
| Fibroblasts | Down | Cacna1a | 8.88E-17 | -0.35312823 | 0.321 | 0.692 | 1.78E-13 |
| Fibroblasts | Down | Fry | 1.82E-11 | -0.35665809 | 0.287 | 0.615 | 3.64E-08 |
| Fibroblasts | Down | Ephb1 | 4.41E-25 | -0.36426345 | 0.277 | 0.635 | 8.82E-22 |
| Fibroblasts | Down | Enpp2 | 2.41E-07 | -0.36526853 | 0.409 | 0.673 | 0.00048152 |
| Fibroblasts | Down | Scn5a | 1.77E-16 | -0.3738918 | 0.238 | 0.562 | 3.54E-13 |
| Fibroblasts | Down | Zfhx3 | 1.91E-08 | -0.38597411 | 0.453 | 0.74 | 3.82E-05 |
| Fibroblasts | Down | Sncaip | 2.42E-23 | -0.38982188 | 0.152 | 0.553 | 4.85E-20 |
| Fibroblasts | Down | Atp10a | 3.02E-18 | -0.39130652 | 0.282 | 0.678 | 6.04E-15 |
| Fibroblasts | Down | Prkaa2 | 6.22E-20 | -0.39502492 | 0.11 | 0.519 | 1.24E-16 |
| Fibroblasts | Down | Fam78b | 1.14E-05 | -0.40081002 | 0.206 | 0.481 | 0.02282346 |
| Fibroblasts | Down | Arhgap31 | 6.36E-14 | -0.40336543 | 0.304 | 0.673 | 1.27E-10 |
| Fibroblasts | Down | Has2 | 7.88E-08 | -0.40843345 | 0.248 | 0.534 | 0.00015755 |
| Fibroblasts | Down | Rhobtb1 | 1.87E-19 | -0.41619085 | 0.169 | 0.587 | 3.73E-16 |

| | | | | | | | |
|---|---|---|---|---|---|---|---|
| Fibroblasts | Down | Pi16 | 1.72E-13 | -0.41794954 | 0.311 | 0.668 | 3.45E-10 |
| Fibroblasts | Down | Pde7b | 1.56E-06 | -0.43796904 | 0.387 | 0.707 | 0.00311775 |
| Fibroblasts | Down | Prkd1 | 2.09E-08 | -0.46266908 | 0.532 | 0.808 | 4.19E-05 |
| Fibroblasts | Down | Fat3 | 6.93E-17 | -0.46390159 | 0.142 | 0.524 | 1.39E-13 |
| Fibroblasts | Down | Actn1 | 1.60E-21 | -0.46587105 | 0.287 | 0.668 | 3.20E-18 |
| Fibroblasts | Down | Prkch | 3.75E-13 | -0.46676995 | 0.159 | 0.505 | 7.49E-10 |
| Fibroblasts | Down | AABR07001519.1 | 3.79E-11 | -0.46788887 | 0.368 | 0.702 | 7.58E-08 |
| Fibroblasts | Down | Rbm20 | 7.52E-14 | -0.48366453 | 0.311 | 0.683 | 1.50E-10 |
| Fibroblasts | Down | Itga8 | 6.36E-10 | -0.49994283 | 0.196 | 0.514 | 1.27E-06 |
| Fibroblasts | Down | Vcan | 9.10E-08 | -0.51055497 | 0.397 | 0.697 | 0.00018197 |
| Fibroblasts | Down | Ednra | 8.80E-19 | -0.51645373 | 0.255 | 0.63 | 1.76E-15 |
| Fibroblasts | Down | Gask1b | 4.44E-13 | -0.56621763 | 0.424 | 0.779 | 8.88E-10 |
| Fibroblasts | Down | Sh3kbp1 | 1.12E-18 | -0.5791472 | 0.252 | 0.644 | 2.24E-15 |
| Fibroblasts | Down | Cilp | 2.68E-20 | -0.58103982 | 0.206 | 0.562 | 5.37E-17 |
| Fibroblasts | Down | Slc22a23 | 1.17E-17 | -0.58314462 | 0.294 | 0.625 | 2.35E-14 |
| Fibroblasts | Down | Loxl1 | 5.10E-17 | -0.59429669 | 0.326 | 0.668 | 1.02E-13 |
| Fibroblasts | Down | Myo1b | 2.83E-23 | -0.60486478 | 0.282 | 0.659 | 5.67E-20 |
| Fibroblasts | Down | Rgs3 | 1.93E-15 | -0.62352891 | 0.353 | 0.683 | 3.87E-12 |
| Fibroblasts | Down | Rem1 | 6.77E-25 | -0.72169695 | 0.267 | 0.697 | 1.35E-21 |
| Fibroblasts | Down | Cacna1g | 2.13E-16 | -0.79277925 | 0.225 | 0.577 | 4.27E-13 |
| Fibroblasts | Down | Nrk | 3.04E-29 | -0.80477493 | 0.083 | 0.548 | 6.08E-26 |
| Fibroblasts | Down | Slc44a5 | 1.49E-25 | -0.83096264 | 0.289 | 0.644 | 2.97E-22 |
| Fibroblasts | Down | Adamts14 | 3.25E-26 | -0.83399824 | 0.248 | 0.635 | 6.50E-23 |
| Fibroblasts | Down | Col5a3 | 4.97E-06 | -0.91998517 | 0.331 | 0.606 | 0.00993703 |
| Fibroblasts | Down | Opcml | 5.56E-15 | -1.00872318 | 0.534 | 0.889 | 1.11E-11 |
| Fibroblasts | Down | Rnf150 | 5.91E-23 | -1.01324709 | 0.248 | 0.692 | 1.18E-19 |
| Fibroblasts | Down | Unc5b | 1.79E-24 | -1.04680879 | 0.328 | 0.76 | 3.57E-21 |
| Fibroblasts | Down | Zfp385b | 7.97E-15 | -1.09205332 | 0.27 | 0.639 | 1.59E-11 |
| Fibroblasts | Down | Adamts17 | 8.74E-25 | -1.12841408 | 0.324 | 0.726 | 1.75E-21 |
| Fibroblasts | Down | Serpine2 | 9.36E-31 | -1.22517479 | 0.353 | 0.774 | 1.87E-27 |
| Macrophages | Down | Eln | 7.73E-06 | -1.75891402 | 0.1 | 0.803 | 0.01545799 |
| Macrophages | Down | Fbln1 | 2.45E-12 | -0.82441471 | 0.1 | 0.955 | 4.90E-09 |
| Macrophages | Down | Kcnma1 | 3.92E-17 | -0.7564369 | 0.04 | 0.97 | 7.84E-14 |
| Macrophages | Down | Abca9 | 7.92E-08 | -0.70296731 | 0.08 | 0.833 | 0.0001583 |
| Macrophages | Down | Crispld2 | 1.56E-09 | -0.60302926 | 0.06 | 0.864 | 3.11E-06 |
| Macrophages | Down | Ar | 3.90E-11 | -0.52615533 | 0.12 | 0.955 | 7.80E-08 |
| Macrophages | Down | Slc1a1 | 9.98E-11 | -0.51704951 | 0.02 | 0.803 | 2.00E-07 |

| | | | | | | | |
|---|---|---|---|---|---|---|---|
| Macrophages | Down | Cdh11 | 2.89E-08 | -0.51348124 | 0.14 | 0.909 | 5.78E-05 |
| Macrophages | Down | Grik2 | 3.34E-08 | -0.49127707 | 0.18 | 0.955 | 6.68E-05 |
| Macrophages | Down | Slc12a7 | 8.56E-07 | -0.49068175 | 0.12 | 0.848 | 0.0017113 |
| Macrophages | Down | Ltbp1 | 8.04E-08 | -0.48715209 | 0.24 | 0.985 | 0.0001608 |
| Macrophages | Down | Osbp2 | 1.19E-05 | -0.47071172 | 0 | 0.712 | 0.02370219 |
| Macrophages | Down | Polr2m | 1.48E-10 | -0.465326 | 0.12 | 0.939 | 2.96E-07 |
| Macrophages | Down | Dnm3 | 5.96E-08 | -0.46031963 | 0.18 | 0.939 | 0.00011924 |
| Macrophages | Down | Pln | 2.57E-06 | -0.45838966 | 0.24 | 0.939 | 0.00514793 |
| Macrophages | Down | Ankrd6 | 6.71E-14 | -0.4552634 | 0.02 | 0.909 | 1.34E-10 |
| Macrophages | Down | Itga7 | 2.83E-13 | -0.42792583 | 0 | 0.864 | 5.65E-10 |
| Macrophages | Down | Rhoh | 3.77E-09 | -0.4260999 | 0.06 | 0.833 | 7.55E-06 |
| Macrophages | Down | Cadps | 7.18E-16 | -0.40993179 | 0.02 | 0.939 | 1.44E-12 |
| Macrophages | Down | Nrk | 6.95E-06 | -0.40648241 | 0 | 0.591 | 0.01390011 |
| Macrophages | Down | Rnf207 | 1.32E-05 | -0.3974448 | 0 | 0.273 | 0.02643903 |
| Macrophages | Down | Afap1l2 | 4.41E-08 | -0.38559528 | 0.14 | 0.909 | 8.82E-05 |
| Macrophages | Down | Ptprd | 1.11E-05 | -0.38373767 | 0.16 | 0.848 | 0.02210848 |
| Macrophages | Down | Fam78b | 3.66E-06 | -0.38361357 | 0.04 | 0.773 | 0.00732365 |
| Macrophages | Down | Prima1 | 3.63E-16 | -0.38315123 | 0.02 | 0.909 | 7.26E-13 |
| Macrophages | Down | Gfra2 | 2.83E-10 | -0.37253979 | 0 | 0.652 | 5.65E-07 |
| Macrophages | Down | Vegfd | 7.44E-08 | -0.35952023 | 0.08 | 0.848 | 0.00014883 |
| Macrophages | Down | AABR07006311.1 | 3.08E-06 | -0.35283325 | 0.06 | 0.5 | 0.0061642 |
| Macrophages | Down | Pgm5 | 8.40E-06 | -0.35155385 | 0.2 | 0.894 | 0.01680801 |
| Macrophages | Down | Sparcl1 | 3.97E-08 | -0.35061145 | 0.16 | 0.924 | 7.94E-05 |
| Macrophages | Down | Cobl | 3.60E-06 | -0.34838514 | 0 | 0.667 | 0.00719794 |
| Macrophages | Down | AABR07027581.1 | 1.60E-05 | -0.3469166 | 0.08 | 0.788 | 0.03190978 |
| Macrophages | Down | Spsb4 | 2.38E-05 | -0.34452838 | 0.02 | 0.727 | 0.04764847 |
| Macrophages | Down | Tox | 4.77E-10 | -0.34272734 | 0.08 | 0.894 | 9.54E-07 |
| Macrophages | Down | Qrfpr | 2.33E-05 | -0.33366367 | 0 | 0.303 | 0.04655972 |
| Macrophages | Down | Fsd2 | 8.43E-07 | -0.32955165 | 0.02 | 0.773 | 0.00168638 |
| Macrophages | Down | Fgf10 | 1.93E-14 | -0.32924667 | 0.02 | 0.909 | 3.87E-11 |
| Macrophages | Down | Tmem168 | 4.16E-08 | -0.32693115 | 0.04 | 0.803 | 8.31E-05 |
| Macrophages | Down | Pstpip2 | 1.35E-05 | -0.32587166 | 0.24 | 0.879 | 0.02698267 |
| Macrophages | Down | Cnnm2 | 1.66E-11 | -0.32525804 | 0.08 | 0.924 | 3.32E-08 |

| | | | | | | | |
|---|---|---|---|---|---|---|---|
| Macrophages | Down | Eva1c | 8.70E-07 | -0.32355063 | 0.06 | 0.803 | 0.00173987 |
| Macrophages | Down | Clic5 | 5.37E-07 | -0.32262693 | 0.2 | 0.924 | 0.00107443 |
| Macrophages | Down | Lrrc2 | 1.32E-05 | -0.31937614 | 0 | 0.727 | 0.02643903 |
| Macrophages | Down | Slc12a2 | 1.11E-07 | -0.29954997 | 0.14 | 0.894 | 0.0002221 |
| Macrophages | Down | Art3 | 3.82E-08 | -0.29510435 | 0.02 | 0.803 | 7.64E-05 |
| Macrophages | Down | Dcdc5 | 4.92E-06 | -0.29459207 | 0 | 0.697 | 0.00984609 |
| Macrophages | Down | Btnl9 | 5.44E-06 | -0.29415197 | 0.02 | 0.727 | 0.01088981 |
| Macrophages | Down | Setbp1 | 8.03E-10 | -0.28282905 | 0.1 | 0.909 | 1.61E-06 |
| Macrophages | Down | Synpo2 | 3.26E-06 | -0.27773748 | 0.2 | 0.909 | 0.00651515 |
| Macrophages | Down | Mx1 | 3.15E-07 | -0.27454127 | 0.04 | 0.697 | 0.00063081 |
| Macrophages | Down | Mboat2 | 1.62E-06 | -0.27449492 | 0.02 | 0.758 | 0.00324895 |
| Macrophages | Down | Angpt2 | 2.66E-11 | -0.27402114 | 0.02 | 0.848 | 5.32E-08 |
| Macrophages | Down | Unc5c | 7.39E-07 | -0.27293802 | 0.06 | 0.773 | 0.00147748 |
| Macrophages | Down | AABR07035722.1 | 1.01E-06 | -0.27266874 | 0.04 | 0.485 | 0.00201121 |
| Macrophages | Down | Ntrk3 | 5.18E-10 | -0.27072729 | 0.02 | 0.833 | 1.04E-06 |
| Macrophages | Down | Egln3 | 7.12E-08 | -0.26369659 | 0 | 0.773 | 0.0001425 |
| Macrophages | Down | Samd5 | 1.73E-06 | -0.25570224 | 0.06 | 0.788 | 0.00345446 |
| Macrophages | Down | Mylk | 2.60E-09 | -0.25118761 | 0.16 | 0.955 | 5.21E-06 |
| Macrophages | Up | Gpm6b | 3.68E-06 | 0.25513422 | 0.1 | 0.818 | 0.00735418 |
| Macrophages | Up | AABR07001573.2 | 1.16E-06 | 0.25603694 | 0.04 | 0.485 | 0.00231461 |
| Macrophages | Up | Pkmyt1 | 7.08E-11 | 0.2564274 | 0.04 | 0.727 | 1.42E-07 |
| Macrophages | Up | Atp10a | 1.47E-11 | 0.25670136 | 0.12 | 0.97 | 2.93E-08 |
| Macrophages | Up | Brip1 | 7.11E-13 | 0.26008802 | 0.06 | 0.924 | 1.42E-09 |
| Macrophages | Up | Kif11 | 8.15E-12 | 0.26062635 | 0.06 | 0.818 | 1.63E-08 |
| Macrophages | Up | Hlf | 4.45E-11 | 0.26098406 | 0.04 | 0.864 | 8.89E-08 |
| Macrophages | Up | AABR07003235.1 | 2.28E-09 | 0.26999051 | 0.06 | 0.682 | 4.56E-06 |
| Macrophages | Up | Hs3st5 | 1.46E-05 | 0.27338676 | 0.04 | 0.758 | 0.02918543 |
| Macrophages | Up | LOC308990 | 9.81E-06 | 0.28356692 | 0.02 | 0.379 | 0.01961029 |
| Macrophages | Up | Mmp8 | 3.73E-06 | 0.28751468 | 0.04 | 0.455 | 0.00746436 |
| Macrophages | Up | Tmem63c | 2.06E-09 | 0.28877078 | 0.04 | 0.727 | 4.13E-06 |
| Macrophages | Up | Angptl1 | 2.45E-06 | 0.29170939 | 0.14 | 0.864 | 0.00489393 |
| Macrophages | Up | Satb2 | 4.42E-12 | 0.29209941 | 0.08 | 0.939 | 8.84E-09 |
| Macrophages | Up | Rgcc | 9.80E-07 | 0.29944171 | 0.14 | 0.879 | 0.00195992 |

| | | | | | | | |
|---|---|---|---|---|---|---|---|
| Macrophages | Up | Hivep3 | 9.14E-09 | 0.30055986 | 0.04 | 0.833 | 1.83E-05 |
| Macrophages | Up | Agmo | 5.83E-10 | 0.3046206 | 0.12 | 0.939 | 1.17E-06 |
| Macrophages | Up | Flnc | 1.97E-09 | 0.30636984 | 0.04 | 0.848 | 3.94E-06 |
| Macrophages | Up | Galnt14 | 5.67E-09 | 0.33073299 | 0.04 | 0.636 | 1.13E-05 |
| Macrophages | Up | AABR07072096.1 | 7.03E-06 | 0.3362187 | 0.06 | 0.485 | 0.01405321 |
| Macrophages | Up | Lbh | 5.15E-07 | 0.33816342 | 0.18 | 0.924 | 0.00102925 |
| Macrophages | Up | Notch4 | 7.84E-07 | 0.33914109 | 0.12 | 0.864 | 0.00156704 |
| Macrophages | Up | S100a4 | 1.61E-11 | 0.34166292 | 0.06 | 0.848 | 3.22E-08 |
| Macrophages | Up | Ccl21 | 4.91E-10 | 0.34243294 | 0.04 | 0.818 | 9.83E-07 |
| Macrophages | Up | Galnt17 | 5.91E-06 | 0.34570684 | 0.14 | 0.848 | 0.01182575 |
| Macrophages | Up | Sgo2 | 6.04E-11 | 0.347906 | 0.06 | 0.758 | 1.21E-07 |
| Macrophages | Up | Atp5me | 2.35E-09 | 0.35528347 | 0.12 | 0.924 | 4.70E-06 |
| Macrophages | Up | Thsd7b | 2.11E-07 | 0.35586937 | 0.06 | 0.712 | 0.00042268 |
| Macrophages | Up | LOC100911486 | 3.83E-10 | 0.36182986 | 0.04 | 0.803 | 7.66E-07 |
| Macrophages | Up | Crlf1 | 1.16E-12 | 0.37624237 | 0.04 | 0.848 | 2.32E-09 |
| Macrophages | Up | Irs1 | 3.52E-06 | 0.38876449 | 0.14 | 0.864 | 0.00703231 |
| Macrophages | Up | Odc1 | 1.20E-11 | 0.39271253 | 0.06 | 0.894 | 2.40E-08 |
| Macrophages | Up | Cx3cr1 | 2.34E-13 | 0.40226247 | 0.04 | 0.833 | 4.69E-10 |
| Macrophages | Up | Esco2 | 1.28E-10 | 0.40379435 | 0.06 | 0.742 | 2.56E-07 |
| Macrophages | Up | Gfpt2 | 2.04E-05 | 0.41514782 | 0.1 | 0.803 | 0.04088758 |
| Macrophages | Up | Fam189a1 | 3.33E-07 | 0.41572667 | 0.08 | 0.742 | 0.00066535 |
| Macrophages | Up | Plac8 | 8.82E-11 | 0.42942241 | 0.06 | 0.773 | 1.76E-07 |
| Macrophages | Up | Tlr7 | 2.94E-08 | 0.44948461 | 0.1 | 0.788 | 5.88E-05 |
| Macrophages | Up | LOC102549869 | 1.90E-12 | 0.45151213 | 0.04 | 0.773 | 3.80E-09 |
| Macrophages | Up | Cenpf | 1.08E-08 | 0.48941779 | 0.1 | 0.803 | 2.15E-05 |
| Macrophages | Up | Ccr1 | 1.17E-06 | 0.50507933 | 0.12 | 0.682 | 0.00233863 |
| Macrophages | Up | Ccl2 | 1.11E-06 | 0.51876953 | 0.08 | 0.773 | 0.0022181 |
| Macrophages | Up | Csrnp1 | 5.20E-06 | 0.52282759 | 0.02 | 0.712 | 0.01040479 |
| Macrophages | Up | Prom1 | 9.91E-08 | 0.54257008 | 0.08 | 0.848 | 0.00019821 |
| Macrophages | Up | Bgn | 6.76E-07 | 0.55056457 | 0.12 | 0.864 | 0.00135227 |
| Macrophages | Up | AABR07054490.1 | 5.71E-07 | 0.57620108 | 0.16 | 0.909 | 0.00114107 |
| Macrophages | Up | Unc5b | 1.60E-07 | 0.57984828 | 0.16 | 0.924 | 0.00031969 |
| Macrophages | Up | Kif4a | 5.01E-09 | 0.65115282 | 0.08 | 0.758 | 1.00E-05 |

| | | | | | | | |
|---|---|---|---|---|---|---|---|
| Macrophages | Up | Epha4 | 2.17E-05 | 0.66477773 | 0.16 | 0.864 | 0.04334241 |
| Macrophages | Up | Srpk3 | 1.33E-09 | 0.68022449 | 0.04 | 0.818 | 2.66E-06 |
| Macrophages | Up | Frem1 | 5.41E-09 | 0.70377122 | 0.14 | 0.939 | 1.08E-05 |
| Macrophages | Up | Brca1 | 3.68E-12 | 0.72594286 | 0.1 | 0.955 | 7.36E-09 |
| Macrophages | Up | Slit2 | 1.33E-06 | 0.7333508 | 0.16 | 0.894 | 0.00265913 |
| Macrophages | Up | LOC100910636 | 1.55E-08 | 0.73689578 | 0.08 | 0.712 | 3.09E-05 |
| Macrophages | Up | Aspm | 1.04E-05 | 0.74225158 | 0.06 | 0.712 | 0.02078964 |
| Macrophages | Up | Nusap1 | 3.53E-08 | 0.75360826 | 0.06 | 0.667 | 7.06E-05 |
| Macrophages | Up | Clec4d | 5.85E-06 | 0.88883866 | 0.08 | 0.652 | 0.01169805 |
| Macrophages | Up | Mmp12 | 2.28E-09 | 1.03392195 | 0.06 | 0.682 | 4.56E-06 |
| Macrophages | Up | Spp1 | 4.75E-09 | 2.41950572 | 0.04 | 0.773 | 9.51E-06 |
| Neuronal Cells | Down | Glis3 | 1.74E-16 | -1.05069252 | 0.091 | 0.966 | 3.48E-13 |
| Neuronal Cells | Down | Ackr3 | 5.06E-16 | -1.03927555 | 0.065 | 0.931 | 1.01E-12 |
| Neuronal Cells | Down | Lrrtm4 | 4.49E-08 | -0.79364771 | 0.26 | 0.966 | 8.98E-05 |
| Neuronal Cells | Down | Zfp536 | 1.21E-06 | -0.73230984 | 0.364 | 1 | 0.00242655 |
| Neuronal Cells | Down | C1qtnf7 | 2.55E-07 | -0.71533092 | 0.221 | 0.914 | 0.00050967 |
| Neuronal Cells | Down | Pparg | 5.16E-17 | -0.64805099 | 0.052 | 0.931 | 1.03E-13 |
| Neuronal Cells | Down | Dkk2 | 1.88E-10 | -0.62789725 | 0.026 | 0.793 | 3.76E-07 |
| Neuronal Cells | Down | Ccn5 | 2.73E-22 | -0.59359336 | 0.026 | 0.948 | 5.46E-19 |
| Neuronal Cells | Down | Tnni3 | 2.22E-06 | -0.5915623 | 0.338 | 1 | 0.00444684 |
| Neuronal Cells | Down | L1cam | 3.98E-08 | -0.55443423 | 0.247 | 0.966 | 7.97E-05 |
| Neuronal Cells | Down | Pln | 5.61E-14 | -0.54033155 | 0.169 | 1 | 1.12E-10 |
| Neuronal Cells | Down | Pip5k1b | 1.81E-10 | -0.50455106 | 0.234 | 1 | 3.61E-07 |
| Neuronal Cells | Down | Myh7 | 1.50E-07 | -0.50225705 | 0.312 | 1 | 0.0002993 |
| Neuronal Cells | Down | Sh3bp2 | 1.08E-20 | -0.48972171 | 0.065 | 0.983 | 2.16E-17 |
| Neuronal Cells | Down | Magi2 | 1.06E-06 | -0.48714826 | 0.026 | 0.741 | 0.00211393 |
| Neuronal Cells | Down | Atp5me | 1.04E-15 | -0.4749738 | 0.117 | 0.983 | 2.08E-12 |
| Neuronal Cells | Down | Rmdn1 | 6.20E-09 | -0.47361957 | 0.195 | 0.931 | 1.24E-05 |
| Neuronal Cells | Down | Fabp3 | 3.34E-21 | -0.46872018 | 0.065 | 1 | 6.67E-18 |
| Neuronal Cells | Down | Plekha7 | 1.38E-05 | -0.44330331 | 0.052 | 0.31 | 0.02750906 |
| Neuronal Cells | Down | Myl2 | 3.20E-10 | -0.43182217 | 0.234 | 1 | 6.40E-07 |
| Neuronal Cells | Down | Klhl29 | 2.57E-07 | -0.39782236 | 0.299 | 1 | 0.00051404 |
| Neuronal Cells | Down | Col27a1 | 9.21E-08 | -0.39329369 | 0.234 | 0.948 | 0.00018426 |

| | | | | | | | |
|---|---|---|---|---|---|---|---|
| Neuronal Cells | Down | Sdc2 | 3.25E-13 | -0.38716577 | 0.156 | 0.983 | 6.50E-10 |
| Neuronal Cells | Down | Herc6 | 4.37E-11 | -0.38609728 | 0.143 | 0.931 | 8.74E-08 |
| Neuronal Cells | Down | Spon1 | 4.70E-13 | -0.3813982 | 0.182 | 1 | 9.41E-10 |
| Neuronal Cells | Down | Htra3 | 5.90E-16 | -0.36941275 | 0.13 | 1 | 1.18E-12 |
| Neuronal Cells | Down | LOC108352650 | 1.81E-16 | -0.36937014 | 0.091 | 0.966 | 3.62E-13 |
| Neuronal Cells | Down | Adamts9 | 2.38E-16 | -0.36088699 | 0.104 | 0.983 | 4.76E-13 |
| Neuronal Cells | Down | Svil | 2.60E-10 | -0.35882767 | 0.195 | 0.966 | 5.19E-07 |
| Neuronal Cells | Down | LOC100911847 | 1.51E-17 | -0.35787887 | 0.091 | 0.983 | 3.02E-14 |
| Neuronal Cells | Down | Sox5 | 2.31E-05 | -0.35350538 | 0.377 | 1 | 0.04629352 |
| Neuronal Cells | Down | Col4a3 | 2.09E-10 | -0.34943674 | 0.156 | 0.931 | 4.18E-07 |
| Neuronal Cells | Down | Atp8b1 | 9.13E-07 | -0.34236717 | 0.325 | 1 | 0.00182658 |
| Neuronal Cells | Down | Tdrd12 | 8.94E-06 | -0.34110364 | 0.351 | 0.983 | 0.01787832 |
| Neuronal Cells | Down | Plcxd3 | 1.67E-07 | -0.3189403 | 0.026 | 0.759 | 0.0003348 |
| Neuronal Cells | Down | Tmsb4x | 4.57E-10 | -0.31805893 | 0.234 | 1 | 9.15E-07 |
| Neuronal Cells | Down | Prkaa2 | 4.01E-22 | -0.31720056 | 0.039 | 0.983 | 8.03E-19 |
| Neuronal Cells | Down | Cyp26b1 | 3.82E-08 | -0.31176533 | 0.013 | 0.741 | 7.63E-05 |
| Neuronal Cells | Down | Man1c1 | 1.31E-15 | -0.31102045 | 0.117 | 0.983 | 2.63E-12 |
| Neuronal Cells | Down | Tspan18 | 1.75E-09 | -0.3101601 | 0.156 | 0.914 | 3.49E-06 |
| Neuronal Cells | Down | Errfi1 | 1.85E-16 | -0.30092246 | 0.039 | 0.914 | 3.71E-13 |
| Neuronal Cells | Down | Hs3st5 | 6.98E-19 | -0.29678865 | 0.026 | 0.931 | 1.40E-15 |
| Neuronal Cells | Down | Ppp2r3a | 1.03E-05 | -0.28917747 | 0.364 | 1 | 0.02052069 |
| Neuronal Cells | Down | Abca8 | 1.01E-09 | -0.28604708 | 0.234 | 1 | 2.02E-06 |
| Neuronal Cells | Down | Skap1 | 3.20E-24 | -0.28448332 | 0.013 | 0.966 | 6.41E-21 |
| Neuronal Cells | Down | Nes | 1.02E-08 | -0.26949259 | 0.039 | 0.793 | 2.05E-05 |
| Neuronal Cells | Down | Rasl12 | 1.68E-17 | -0.25879364 | 0.026 | 0.914 | 3.37E-14 |
| Neuronal Cells | Down | Jph2 | 7.37E-16 | -0.25130408 | 0.013 | 0.879 | 1.47E-12 |
| Neuronal Cells | Up | Slc7a2 | 6.96E-21 | 0.2550198 | 0.065 | 1 | 1.39E-17 |
| Neuronal Cells | Up | Arhgap22 | 1.03E-19 | 0.25533912 | 0.078 | 1 | 2.07E-16 |
| Neuronal Cells | Up | Crip1 | 2.84E-10 | 0.25699458 | 0.195 | 0.983 | 5.69E-07 |
| Neuronal Cells | Up | Fgf12 | 4.89E-16 | 0.25799051 | 0.091 | 0.966 | 9.78E-13 |
| Neuronal Cells | Up | Ikzf2 | 1.27E-14 | 0.25828686 | 0.104 | 0.948 | 2.53E-11 |
| Neuronal Cells | Up | Nlgn1 | 1.30E-13 | 0.2590997 | 0.156 | 1 | 2.60E-10 |
| Neuronal Cells | Up | Bcl11a | 8.93E-06 | 0.26187518 | 0.013 | 0.345 | 0.01785232 |

| | | | | | | | |
|---|---|---|---|---|---|---|---|
| Neuronal Cells | Up | Notch3 | 4.85E-22 | 0.26447497 | 0.039 | 0.983 | 9.71E-19 |
| Neuronal Cells | Up | Arhgap15 | 1.30E-15 | 0.2660122 | 0.065 | 0.931 | 2.61E-12 |
| Neuronal Cells | Up | Susd5 | 3.59E-23 | 0.26910603 | 0.026 | 0.948 | 7.18E-20 |
| Neuronal Cells | Up | Rasgef1b | 2.45E-18 | 0.27223757 | 0.078 | 0.983 | 4.90E-15 |
| Neuronal Cells | Up | Atp10a | 2.17E-09 | 0.27668149 | 0.13 | 0.897 | 4.33E-06 |
| Neuronal Cells | Up | C1qtnf9 | 2.57E-20 | 0.27678611 | 0.065 | 0.983 | 5.14E-17 |
| Neuronal Cells | Up | Nxn | 2.96E-07 | 0.27916077 | 0.286 | 1 | 0.00059293 |
| Neuronal Cells | Up | Dnajc6 | 5.35E-21 | 0.28133187 | 0.026 | 0.914 | 1.07E-17 |
| Neuronal Cells | Up | Cadm1 | 4.53E-19 | 0.28327266 | 0.078 | 0.983 | 9.06E-16 |
| Neuronal Cells | Up | Usp2 | 4.85E-22 | 0.2888354 | 0.039 | 0.983 | 9.71E-19 |
| Neuronal Cells | Up | Cers6 | 1.56E-18 | 0.2895691 | 0.091 | 1 | 3.13E-15 |
| Neuronal Cells | Up | Prrx1 | 7.78E-13 | 0.2899104 | 0.104 | 0.931 | 1.56E-09 |
| Neuronal Cells | Up | Col11a1 | 4.43E-16 | 0.29128524 | 0.078 | 0.931 | 8.85E-13 |
| Neuronal Cells | Up | Me1 | 2.67E-08 | 0.30024797 | 0.065 | 0.81 | 5.34E-05 |
| Neuronal Cells | Up | Lypd1 | 1.22E-17 | 0.30498167 | 0.026 | 0.81 | 2.44E-14 |
| Neuronal Cells | Up | Rhov | 8.75E-24 | 0.30712925 | 0.026 | 0.966 | 1.75E-20 |
| Neuronal Cells | Up | Pdlim3 | 2.36E-10 | 0.30996585 | 0.13 | 0.914 | 4.72E-07 |
| Neuronal Cells | Up | Hspb8 | 1.89E-10 | 0.31529289 | 0.039 | 0.776 | 3.78E-07 |
| Neuronal Cells | Up | Rrad | 1.36E-23 | 0.31763109 | 0.039 | 1 | 2.72E-20 |
| Neuronal Cells | Up | Gnao1 | 3.77E-10 | 0.32398071 | 0.208 | 1 | 7.55E-07 |
| Neuronal Cells | Up | Chrdl1 | 2.16E-18 | 0.34340038 | 0.078 | 0.983 | 4.32E-15 |
| Neuronal Cells | Up | Cox8b | 2.78E-09 | 0.34718183 | 0.182 | 0.948 | 5.56E-06 |
| Neuronal Cells | Up | Nrxn3 | 1.21E-20 | 0.3489364 | 0.026 | 0.931 | 2.42E-17 |
| Neuronal Cells | Up | Tmem51 | 1.52E-20 | 0.35286812 | 0.039 | 0.966 | 3.04E-17 |
| Neuronal Cells | Up | Cntfr | 1.19E-06 | 0.35492358 | 0.039 | 0.534 | 0.00238096 |
| Neuronal Cells | Up | Inpp5d | 2.62E-19 | 0.35639372 | 0.052 | 0.966 | 5.24E-16 |
| Neuronal Cells | Up | Icam1 | 6.96E-21 | 0.35709814 | 0.065 | 1 | 1.39E-17 |
| Neuronal Cells | Up | Neto1 | 4.10E-07 | 0.35914902 | 0.104 | 0.81 | 0.00081911 |
| Neuronal Cells | Up | Spns2 | 2.48E-17 | 0.362157 | 0.104 | 1 | 4.97E-14 |
| Neuronal Cells | Up | Usp18 | 2.06E-14 | 0.36356919 | 0.065 | 0.914 | 4.12E-11 |
| Neuronal Cells | Up | Ar | 5.77E-06 | 0.3648242 | 0.195 | 0.879 | 0.01154832 |
| Neuronal Cells | Up | Ntrk3 | 1.49E-13 | 0.3670442 | 0.078 | 0.914 | 2.98E-10 |
| Neuronal Cells | Up | Ppip5k1 | 1.08E-08 | 0.36722381 | 0.208 | 0.966 | 2.17E-05 |

| | | | | | | | |
|---|---|---|---|---|---|---|---|
| Neuronal Cells | Up | Abca1 | 1.30E-05 | 0.36907023 | 0.325 | 1 | 0.02599504 |
| Neuronal Cells | Up | Hs6st2 | 1.36E-14 | 0.38350223 | 0.078 | 0.931 | 2.71E-11 |
| Neuronal Cells | Up | Apoe | 5.48E-13 | 0.38660883 | 0.117 | 0.948 | 1.10E-09 |
| Neuronal Cells | Up | Musk | 1.77E-19 | 0.38938061 | 0.065 | 0.983 | 3.53E-16 |
| Neuronal Cells | Up | Lmcd1 | 3.03E-13 | 0.39646796 | 0.156 | 1 | 6.06E-10 |
| Neuronal Cells | Up | Srpx | 4.74E-10 | 0.39741743 | 0.117 | 0.897 | 9.47E-07 |
| Neuronal Cells | Up | Grid2 | 2.82E-09 | 0.40028944 | 0.169 | 0.931 | 5.65E-06 |
| Neuronal Cells | Up | Mx2 | 1.08E-19 | 0.40128348 | 0.078 | 1 | 2.16E-16 |
| Neuronal Cells | Up | Tox3 | 2.45E-18 | 0.40601059 | 0.078 | 0.983 | 4.90E-15 |
| Neuronal Cells | Up | Negr1 | 2.29E-17 | 0.4085568 | 0.104 | 1 | 4.58E-14 |
| Neuronal Cells | Up | Sema5a | 3.08E-13 | 0.40940302 | 0.143 | 0.983 | 6.16E-10 |
| Neuronal Cells | Up | Gria4 | 1.27E-16 | 0.42403208 | 0.052 | 0.931 | 2.54E-13 |
| Neuronal Cells | Up | Chsy3 | 8.98E-13 | 0.42618686 | 0.169 | 1 | 1.80E-09 |
| Neuronal Cells | Up | Galnt15 | 8.85E-16 | 0.42784312 | 0.078 | 0.948 | 1.77E-12 |
| Neuronal Cells | Up | Ptk2b | 1.45E-07 | 0.4281214 | 0.039 | 0.603 | 0.00028934 |
| Neuronal Cells | Up | Rfx2 | 5.15E-17 | 0.43451455 | 0.078 | 0.966 | 1.03E-13 |
| Neuronal Cells | Up | Slco5a1 | 1.36E-23 | 0.44004148 | 0.039 | 1 | 2.72E-20 |
| Neuronal Cells | Up | Msrb3 | 1.50E-08 | 0.4410905 | 0.208 | 0.966 | 3.00E-05 |
| Neuronal Cells | Up | Map2 | 1.71E-08 | 0.44248306 | 0.182 | 0.931 | 3.42E-05 |
| Neuronal Cells | Up | Ndrg4 | 2.05E-09 | 0.44289152 | 0.078 | 0.845 | 4.11E-06 |
| Neuronal Cells | Up | Xirp2 | 3.52E-08 | 0.44385208 | 0.234 | 0.966 | 7.04E-05 |
| Neuronal Cells | Up | Pstpip1 | 2.79E-18 | 0.44398746 | 0.078 | 0.983 | 5.57E-15 |
| Neuronal Cells | Up | Sh3gl2 | 7.73E-10 | 0.44535035 | 0.182 | 0.966 | 1.55E-06 |
| Neuronal Cells | Up | Bche | 5.71E-11 | 0.44791619 | 0.117 | 0.914 | 1.14E-07 |
| Neuronal Cells | Up | Slc6a6 | 2.99E-06 | 0.45034809 | 0.169 | 0.862 | 0.00597002 |
| Neuronal Cells | Up | Lrrc2 | 4.26E-11 | 0.46291141 | 0.065 | 0.862 | 8.52E-08 |
| Neuronal Cells | Up | Myrip | 6.08E-07 | 0.46782708 | 0.234 | 0.948 | 0.00121545 |
| Neuronal Cells | Up | Maob | 2.22E-14 | 0.46868847 | 0.065 | 0.914 | 4.44E-11 |
| Neuronal Cells | Up | Clic5 | 5.89E-09 | 0.46898905 | 0.234 | 1 | 1.18E-05 |
| Neuronal Cells | Up | RGD1306750 | 3.70E-17 | 0.47844102 | 0.013 | 0.724 | 7.40E-14 |
| Neuronal Cells | Up | Kcnab1 | 2.22E-14 | 0.48535626 | 0.065 | 0.914 | 4.44E-11 |
| Neuronal Cells | Up | Kcnma1 | 7.91E-13 | 0.48684381 | 0.026 | 0.759 | 1.58E-09 |
| Neuronal Cells | Up | Caskin1 | 6.56E-13 | 0.49357966 | 0.078 | 0.845 | 1.31E-09 |

| | | | | | | | |
|---|---|---|---|---|---|---|---|
| Neuronal Cells | Up | Zmat4 | 4.77E-14 | 0.49765704 | 0.026 | 0.828 | 9.54E-11 |
| Neuronal Cells | Up | Mtus2 | 1.39E-12 | 0.50092319 | 0.026 | 0.776 | 2.79E-09 |
| Neuronal Cells | Up | Fbln7 | 7.05E-18 | 0.50320325 | 0.091 | 0.983 | 1.41E-14 |
| Neuronal Cells | Up | Slc9a9 | 1.62E-06 | 0.5109685 | 0.26 | 0.966 | 0.00324465 |
| Neuronal Cells | Up | Cacna1g | 3.90E-17 | 0.55179429 | 0.091 | 0.983 | 7.79E-14 |
| Neuronal Cells | Up | Pcsk5 | 5.36E-17 | 0.5616736 | 0.078 | 0.966 | 1.07E-13 |
| Neuronal Cells | Up | Fat3 | 3.92E-16 | 0.56683666 | 0.117 | 1 | 7.85E-13 |
| Neuronal Cells | Up | Atp5mc1 | 3.81E-18 | 0.59626686 | 0.065 | 0.966 | 7.62E-15 |
| Neuronal Cells | Up | Lepr | 9.88E-09 | 0.60732701 | 0.091 | 0.845 | 1.98E-05 |
| Neuronal Cells | Up | Zbtb7c | 6.36E-08 | 0.63722457 | 0.143 | 0.879 | 0.00012712 |
| Neuronal Cells | Up | Nrg1 | 5.96E-16 | 0.64505939 | 0.091 | 0.966 | 1.19E-12 |
| Neuronal Cells | Up | Atp5f1e | 3.88E-15 | 0.67182271 | 0.13 | 1 | 7.75E-12 |
| Neuronal Cells | Up | Adgrb3 | 8.60E-23 | 0.71201505 | 0.026 | 0.966 | 1.72E-19 |
| Neuronal Cells | Up | Bcat1 | 7.58E-12 | 0.79653761 | 0.117 | 0.931 | 1.52E-08 |
| Neuronal Cells | Up | Chst11 | 1.83E-06 | 0.94001018 | 0.273 | 1 | 0.0036545 |
| Neuronal Cells | Up | Myh11 | 2.32E-14 | 0.97955504 | 0.143 | 1 | 4.64E-11 |
| Lymphatic Endothelial Cells | Down | Rassf9 | 3.31E-09 | -0.59720192 | 0.172 | 0.925 | 6.61E-06 |
| Lymphatic Endothelial Cells | Down | Nrap | 8.81E-23 | -0.57139353 | 0.031 | 1 | 1.76E-19 |
| Lymphatic Endothelial Cells | Down | Ednrb | 1.55E-15 | -0.56857972 | 0.125 | 1 | 3.11E-12 |
| Lymphatic Endothelial Cells | Down | Prag1 | 2.39E-21 | -0.54039303 | 0.047 | 1 | 4.79E-18 |
| Lymphatic Endothelial Cells | Down | Filip1l | 1.52E-07 | -0.53304267 | 0.312 | 1 | 0.00030346 |
| Lymphatic Endothelial Cells | Down | Ccser1 | 1.41E-12 | -0.52562034 | 0.031 | 1 | 2.82E-09 |
| Lymphatic Endothelial Cells | Down | Mlip | 4.29E-10 | -0.52099358 | 0.078 | 1 | 8.58E-07 |
| Lymphatic Endothelial Cells | Down | Cap2 | 2.63E-21 | -0.50993858 | 0.047 | 1 | 5.26E-18 |
| Lymphatic Endothelial Cells | Down | Enox2 | 2.76E-08 | -0.50759524 | 0.266 | 1 | 5.52E-05 |
| Lymphatic Endothelial Cells | Down | Ablim3 | 6.43E-08 | -0.50673045 | 0.281 | 1 | 0.00012855 |
| Lymphatic Endothelial Cells | Down | Ube2ql1 | 8.15E-18 | -0.49023544 | 0.031 | 1 | 1.63E-14 |
| Lymphatic Endothelial Cells | Down | B2m | 1.40E-14 | -0.48841449 | 0.141 | 1 | 2.80E-11 |
| Lymphatic Endothelial Cells | Down | Cpq | 3.91E-06 | -0.4616731 | 0.203 | 1 | 0.00782325 |
| Lymphatic Endothelial Cells | Down | Adcy5 | 1.19E-12 | -0.44906654 | 0.047 | 1 | 2.37E-09 |
| Lymphatic Endothelial Cells | Down | Rasa4 | 3.53E-20 | -0.44695804 | 0.031 | 1 | 7.07E-17 |
| Lymphatic Endothelial Cells | Down | Tbx1 | 5.11E-06 | -0.44494297 | 0.312 | 1 | 0.01021113 |
| Lymphatic Endothelial Cells | Down | Slc26a10 | 1.48E-15 | -0.44297114 | 0.062 | 1 | 2.96E-12 |

| | | | | | | | |
|---|---|---|---|---|---|---|---|
| Lymphatic Endothelial Cells | Down | Dipk2b | 1.85E-10 | -0.43931235 | 0.172 | 1 | 3.71E-07 |
| Lymphatic Endothelial Cells | Down | Podxl | 6.77E-09 | -0.43838051 | 0.266 | 1 | 1.35E-05 |
| Lymphatic Endothelial Cells | Down | Grb14 | 1.27E-11 | -0.43520923 | 0.016 | 1 | 2.55E-08 |
| Lymphatic Endothelial Cells | Down | Papss2 | 1.09E-17 | -0.42359836 | 0.047 | 1 | 2.19E-14 |
| Lymphatic Endothelial Cells | Down | Slc7a2 | 9.71E-23 | -0.3889069 | 0.031 | 1 | 1.94E-19 |
| Lymphatic Endothelial Cells | Down | Camk1d | 1.55E-14 | -0.37958688 | 0.109 | 1 | 3.10E-11 |
| Lymphatic Endothelial Cells | Down | AABR07060560.3 | 1.32E-06 | -0.37597019 | 0 | 1 | 0.00264059 |
| Lymphatic Endothelial Cells | Down | Thsd7b | 1.08E-11 | -0.35527938 | 0.047 | 1 | 2.15E-08 |
| Lymphatic Endothelial Cells | Down | Gbp1 | 1.81E-05 | -0.3424905 | 0 | 1 | 0.03622601 |
| Lymphatic Endothelial Cells | Down | Lama5 | 1.07E-22 | -0.33666728 | 0.031 | 1 | 2.14E-19 |
| Lymphatic Endothelial Cells | Down | Kcng2 | 1.07E-22 | -0.33023154 | 0.031 | 1 | 2.14E-19 |
| Lymphatic Endothelial Cells | Down | Igfbp5 | 1.73E-09 | -0.32942463 | 0.203 | 1 | 3.47E-06 |
| Lymphatic Endothelial Cells | Down | Pdcd1lg2 | 8.19E-06 | -0.30642555 | 0.047 | 1 | 0.01637042 |
| Lymphatic Endothelial Cells | Down | Tpt1 | 1.39E-13 | -0.294236 | 0.062 | 1 | 2.77E-10 |
| Lymphatic Endothelial Cells | Down | Tmem50b | 1.56E-21 | -0.29358983 | 0.016 | 1 | 3.11E-18 |
| Lymphatic Endothelial Cells | Down | Ptprf | 7.81E-18 | -0.29281771 | 0.031 | 1 | 1.56E-14 |
| Lymphatic Endothelial Cells | Down | Enah | 7.66E-23 | -0.28905368 | 0.016 | 1 | 1.53E-19 |
| Lymphatic Endothelial Cells | Down | Cd44 | 3.61E-24 | -0.28758538 | 0.016 | 1 | 7.22E-21 |
| Lymphatic Endothelial Cells | Down | Nr2f2 | 6.18E-07 | -0.28564789 | 0.266 | 1 | 0.00123522 |
| Lymphatic Endothelial Cells | Down | Rhobtb1 | 3.14E-15 | -0.28092178 | 0.109 | 1 | 6.28E-12 |
| Lymphatic Endothelial Cells | Down | Gxylt2 | 8.75E-12 | -0.27979112 | 0.172 | 1 | 1.75E-08 |
| Lymphatic Endothelial Cells | Down | Ror2 | 1.37E-17 | -0.2772758 | 0.078 | 1 | 2.74E-14 |
| Lymphatic Endothelial Cells | Down | Vcam1 | 1.56E-21 | -0.27493698 | 0.016 | 1 | 3.11E-18 |
| Lymphatic Endothelial Cells | Down | Pln | 2.02E-09 | -0.27132361 | 0.219 | 1 | 4.05E-06 |
| Lymphatic Endothelial Cells | Down | Blnk | 1.94E-21 | -0.2614425 | 0.031 | 1 | 3.88E-18 |
| Lymphatic Endothelial Cells | Down | Pawr | 2.14E-15 | -0.25837307 | 0.094 | 1 | 4.27E-12 |
| Lymphatic Endothelial Cells | Down | Speg | 2.88E-20 | -0.25764314 | 0.016 | 1 | 5.75E-17 |
| Lymphatic Endothelial Cells | Down | Sncaip | 1.13E-11 | -0.25240127 | 0.188 | 1 | 2.27E-08 |
| Lymphatic Endothelial Cells | Up | Prkcq | 3.67E-16 | 0.25653022 | 0.031 | 1 | 7.34E-13 |
| Lymphatic Endothelial Cells | Up | Smox | 1.06E-19 | 0.26132389 | 0.062 | 1 | 2.11E-16 |
| Lymphatic Endothelial Cells | Up | Gask1b | 1.96E-08 | 0.26203612 | 0.062 | 1 | 3.92E-05 |
| Lymphatic Endothelial Cells | Up | Mafb | 1.01E-19 | 0.26502723 | 0.062 | 1 | 2.02E-16 |
| Lymphatic Endothelial Cells | Up | Aspa | 3.99E-21 | 0.26944873 | 0.047 | 1 | 7.98E-18 |

| | | | | | | | |
|---|---|---|---|---|---|---|---|
| Lymphatic Endothelial Cells | Up | AABR07007642.1 | 4.43E-13 | 0.27163426 | 0.141 | 1 | 8.85E-10 |
| Lymphatic Endothelial Cells | Up | Procr | 6.69E-19 | 0.27588775 | 0.031 | 1 | 1.34E-15 |
| Lymphatic Endothelial Cells | Up | Hspb7 | 9.33E-11 | 0.27705042 | 0.062 | 1 | 1.87E-07 |
| Lymphatic Endothelial Cells | Up | Asic2 | 1.50E-13 | 0.27764677 | 0.031 | 1 | 3.00E-10 |
| Lymphatic Endothelial Cells | Up | Mitf | 3.92E-12 | 0.28079861 | 0.172 | 1 | 7.84E-09 |
| Lymphatic Endothelial Cells | Up | Adgre4 | 1.34E-19 | 0.28397321 | 0.031 | 1 | 2.67E-16 |
| Lymphatic Endothelial Cells | Up | Alpl | 2.00E-08 | 0.28492561 | 0.031 | 1 | 4.00E-05 |
| Lymphatic Endothelial Cells | Up | Wnt5b | 1.25E-18 | 0.30082704 | 0.031 | 1 | 2.49E-15 |
| Lymphatic Endothelial Cells | Up | Nppa | 6.44E-20 | 0.31084883 | 0.047 | 1 | 1.29E-16 |
| Lymphatic Endothelial Cells | Up | LOC103694210 | 4.62E-16 | 0.31204499 | 0.094 | 1 | 9.24E-13 |
| Lymphatic Endothelial Cells | Up | Ano1 | 2.46E-19 | 0.32151087 | 0.031 | 1 | 4.92E-16 |
| Lymphatic Endothelial Cells | Up | AABR07060560.2 | 1.42E-21 | 0.33121384 | 0.031 | 1 | 2.84E-18 |
| Lymphatic Endothelial Cells | Up | Prune2 | 5.11E-06 | 0.33138399 | 0.219 | 1 | 0.01021517 |
| Lymphatic Endothelial Cells | Up | Ccn1 | 1.02E-10 | 0.33247986 | 0.031 | 1 | 2.03E-07 |
| Lymphatic Endothelial Cells | Up | Mctp2 | 4.13E-22 | 0.33825323 | 0.031 | 1 | 8.26E-19 |
| Lymphatic Endothelial Cells | Up | Atp8b1 | 5.17E-15 | 0.34866344 | 0.109 | 1 | 1.03E-11 |
| Lymphatic Endothelial Cells | Up | Myrip | 6.93E-11 | 0.34879859 | 0.188 | 1 | 1.39E-07 |
| Lymphatic Endothelial Cells | Up | AABR07034767.1 | 2.67E-17 | 0.35152504 | 0.078 | 1 | 5.33E-14 |
| Lymphatic Endothelial Cells | Up | Nkd1 | 9.38E-15 | 0.35178024 | 0.047 | 1 | 1.88E-11 |
| Lymphatic Endothelial Cells | Up | Trpc3 | 1.31E-12 | 0.35755389 | 0.156 | 1 | 2.63E-09 |
| Lymphatic Endothelial Cells | Up | Apoo | 2.78E-17 | 0.36095033 | 0.078 | 1 | 5.56E-14 |
| Lymphatic Endothelial Cells | Up | Mbp | 1.99E-19 | 0.36141783 | 0.047 | 1 | 3.98E-16 |
| Lymphatic Endothelial Cells | Up | Icam1 | 1.06E-19 | 0.36764988 | 0.062 | 1 | 2.11E-16 |
| Lymphatic Endothelial Cells | Up | Carmil1 | 2.12E-08 | 0.375508 | 0.109 | 1 | 4.24E-05 |
| Lymphatic Endothelial Cells | Up | Ablim2 | 7.56E-09 | 0.37692182 | 0.219 | 1 | 1.51E-05 |
| Lymphatic Endothelial Cells | Up | Lepr | 1.06E-19 | 0.37997451 | 0.062 | 1 | 2.11E-16 |
| Lymphatic Endothelial Cells | Up | Cobl | 1.40E-18 | 0.38052267 | 0.062 | 1 | 2.81E-15 |
| Lymphatic Endothelial Cells | Up | Loxl2 | 3.20E-16 | 0.39395009 | 0.078 | 1 | 6.40E-13 |
| Lymphatic Endothelial Cells | Up | Abca8a | 1.24E-06 | 0.3992754 | 0.266 | 1 | 0.00248288 |
| Lymphatic Endothelial Cells | Up | Rnf213 | 9.97E-06 | 0.40501953 | 0.281 | 1 | 0.01993097 |
| Lymphatic Endothelial Cells | Up | Spon1 | 6.35E-16 | 0.40645206 | 0.109 | 1 | 1.27E-12 |
| Lymphatic Endothelial Cells | Up | Ankh | 2.14E-07 | 0.41121605 | 0.25 | 1 | 0.00042715 |
| Lymphatic Endothelial Cells | Up | Mx1 | 1.53E-17 | 0.41247127 | 0.047 | 1 | 3.06E-14 |

| | | | | | | | |
|---|---|---|---|---|---|---|---|
| Lymphatic Endothelial Cells | Up | Eif2ak2 | 5.09E-11 | 0.41802948 | 0.172 | 1 | 1.02E-07 |
| Lymphatic Endothelial Cells | Up | Adgrd1 | 1.05E-15 | 0.42608723 | 0.094 | 1 | 2.10E-12 |
| Lymphatic Endothelial Cells | Up | Slit2 | 1.06E-19 | 0.42629436 | 0.062 | 1 | 2.11E-16 |
| Lymphatic Endothelial Cells | Up | Fndc1 | 1.34E-05 | 0.4354181 | 0.312 | 1 | 0.02675244 |
| Lymphatic Endothelial Cells | Up | Pfkfb3 | 5.71E-10 | 0.43772054 | 0.125 | 1 | 1.14E-06 |
| Lymphatic Endothelial Cells | Up | Rsad2 | 3.77E-17 | 0.44699372 | 0.047 | 1 | 7.54E-14 |
| Lymphatic Endothelial Cells | Up | Hk2 | 2.67E-17 | 0.44827514 | 0.078 | 1 | 5.33E-14 |
| Lymphatic Endothelial Cells | Up | Tm4sf1 | 6.61E-16 | 0.48239202 | 0.109 | 1 | 1.32E-12 |
| Lymphatic Endothelial Cells | Up | Gramd1b | 4.63E-17 | 0.48355775 | 0.094 | 1 | 9.26E-14 |
| Lymphatic Endothelial Cells | Up | Cpeb1 | 2.84E-13 | 0.49178213 | 0.094 | 1 | 5.67E-10 |
| Lymphatic Endothelial Cells | Up | Shc3 | 1.74E-17 | 0.50187906 | 0.078 | 1 | 3.47E-14 |
| Lymphatic Endothelial Cells | Up | Zfp385b | 2.43E-11 | 0.50355492 | 0.125 | 1 | 4.85E-08 |
| Lymphatic Endothelial Cells | Up | AABR07051518.1 | 1.06E-19 | 0.50693851 | 0.062 | 1 | 2.11E-16 |
| Lymphatic Endothelial Cells | Up | Clic5 | 2.11E-08 | 0.50956415 | 0.234 | 1 | 4.23E-05 |
| Lymphatic Endothelial Cells | Up | Hs3st1 | 3.63E-09 | 0.5147943 | 0.078 | 1 | 7.26E-06 |
| Lymphatic Endothelial Cells | Up | Aox3 | 4.45E-13 | 0.52932054 | 0.109 | 1 | 8.90E-10 |
| Lymphatic Endothelial Cells | Up | Ralgps2 | 4.62E-16 | 0.53010181 | 0.094 | 1 | 9.24E-13 |
| Lymphatic Endothelial Cells | Up | Opcml | 4.81E-16 | 0.53453724 | 0.094 | 1 | 9.61E-13 |
| Lymphatic Endothelial Cells | Up | Ar | 8.08E-14 | 0.55038742 | 0.125 | 1 | 1.62E-10 |
| Lymphatic Endothelial Cells | Up | Sdc2 | 2.72E-12 | 0.55944995 | 0.109 | 1 | 5.45E-09 |
| Lymphatic Endothelial Cells | Up | Dgki | 3.07E-19 | 0.58396235 | 0.016 | 1 | 6.14E-16 |
| Lymphatic Endothelial Cells | Up | Slc7a1 | 1.51E-10 | 0.59311019 | 0.125 | 1 | 3.02E-07 |
| Lymphatic Endothelial Cells | Up | Smad6 | 6.16E-20 | 0.5966768 | 0.047 | 1 | 1.23E-16 |
| Lymphatic Endothelial Cells | Up | Kcnab1 | 2.86E-18 | 0.62944054 | 0.047 | 1 | 5.72E-15 |
| Lymphatic Endothelial Cells | Up | Ankrd55 | 2.92E-12 | 0.66734442 | 0.109 | 1 | 5.84E-09 |
| Lymphatic Endothelial Cells | Up | Eda | 7.56E-15 | 0.72489326 | 0.109 | 1 | 1.51E-11 |
| Lymphatic Endothelial Cells | Up | Cfh | 8.92E-13 | 0.87327644 | 0.141 | 1 | 1.78E-09 |
| Lymphatic Endothelial Cells | Up | Cmahp | 1.90E-09 | 0.89903413 | 0.172 | 1 | 3.80E-06 |
| Lymphatic Endothelial Cells | Up | Pde1c | 3.63E-12 | 0.95483145 | 0.125 | 1 | 7.25E-09 |
| Smooth Muscle Cells | Down | Nrp2 | 2.19E-19 | -0.53393256 | 0.101 | 0.801 | 4.38E-16 |
| Smooth Muscle Cells | Down | Cav1 | 2.76E-16 | -0.43464391 | 0.273 | 0.914 | 5.52E-13 |
| Smooth Muscle Cells | Down | Col8a1 | 9.66E-07 | -0.41080102 | 0.348 | 0.871 | 0.00193111 |
| Smooth Muscle Cells | Down | Ccl19 | 3.08E-14 | -0.39940327 | 0.011 | 0.446 | 6.16E-11 |

| | | | | | | | |
|---|---|---|---|---|---|---|---|
| Smooth Muscle Cells | Down | Gpr176 | 3.10E-21 | -0.34959034 | 0.094 | 0.801 | 6.19E-18 |
| Smooth Muscle Cells | Down | Col19a1 | 1.26E-06 | -0.31695378 | 0.18 | 0.43 | 0.00251486 |
| Smooth Muscle Cells | Down | Col9a1 | 6.63E-23 | -0.3163121 | 0.03 | 0.452 | 1.33E-19 |
| Smooth Muscle Cells | Down | Chn1 | 1.01E-12 | -0.31062742 | 0.33 | 0.941 | 2.02E-09 |
| Smooth Muscle Cells | Down | Clic5 | 6.50E-26 | -0.30741634 | 0.232 | 0.968 | 1.30E-22 |
| Smooth Muscle Cells | Down | Ptn | 2.66E-14 | -0.29888051 | 0.105 | 0.64 | 5.32E-11 |
| Smooth Muscle Cells | Down | Cacna1g | 5.06E-23 | -0.28910953 | 0.124 | 0.849 | 1.01E-19 |
| Smooth Muscle Cells | Down | Bace2 | 3.06E-19 | -0.28616183 | 0.064 | 0.769 | 6.12E-16 |
| Smooth Muscle Cells | Down | Slc12a2 | 1.05E-16 | -0.2830189 | 0.307 | 0.968 | 2.10E-13 |
| Smooth Muscle Cells | Down | Nostrin | 2.69E-20 | -0.27970111 | 0.157 | 0.747 | 5.37E-17 |
| Smooth Muscle Cells | Down | Sgip1 | 5.43E-14 | -0.27873097 | 0.24 | 0.876 | 1.09E-10 |
| Smooth Muscle Cells | Down | Mcf2l | 1.25E-22 | -0.27235647 | 0.184 | 0.898 | 2.51E-19 |
| Smooth Muscle Cells | Down | Sybu | 1.16E-06 | -0.27052316 | 0.037 | 0.613 | 0.00231919 |
| Smooth Muscle Cells | Down | Flnc | 1.72E-13 | -0.26998158 | 0.015 | 0.317 | 3.44E-10 |
| Smooth Muscle Cells | Down | Pcsk5 | 1.04E-10 | -0.26125711 | 0.109 | 0.737 | 2.08E-07 |
| Smooth Muscle Cells | Down | Flnb | 1.77E-21 | -0.25454316 | 0.243 | 0.952 | 3.54E-18 |
| Smooth Muscle Cells | Down | Medag | 6.75E-06 | -0.25136811 | 0.026 | 0.398 | 0.01349327 |
| Smooth Muscle Cells | Up | Uqcrq | 1.37E-21 | 0.25471985 | 0.105 | 0.828 | 2.75E-18 |
| Smooth Muscle Cells | Up | Nfatc2 | 2.30E-06 | 0.25857333 | 0.037 | 0.344 | 0.00460448 |
| Smooth Muscle Cells | Up | Itgb8 | 2.71E-15 | 0.26062864 | 0.131 | 0.801 | 5.42E-12 |
| Smooth Muscle Cells | Up | Cox8b | 5.34E-14 | 0.26419879 | 0.18 | 0.833 | 1.07E-10 |
| Smooth Muscle Cells | Up | Phyh | 1.70E-08 | 0.2655836 | 0.15 | 0.409 | 3.40E-05 |
| Smooth Muscle Cells | Up | Kcnab1 | 5.58E-09 | 0.276669 | 0.221 | 0.801 | 1.12E-05 |
| Smooth Muscle Cells | Up | Ndufa4 | 2.08E-05 | 0.28550106 | 0.135 | 0.699 | 0.04158997 |
| Smooth Muscle Cells | Up | Ankh | 5.71E-18 | 0.28715416 | 0.157 | 0.849 | 1.14E-14 |
| Smooth Muscle Cells | Up | Agmo | 4.93E-08 | 0.29298784 | 0.27 | 0.844 | 9.85E-05 |
| Smooth Muscle Cells | Up | Thbs1 | 1.60E-26 | 0.29445845 | 0.049 | 0.597 | 3.20E-23 |
| Smooth Muscle Cells | Up | Pfkfb3 | 1.92E-05 | 0.30382116 | 0.12 | 0.688 | 0.03848939 |
| Smooth Muscle Cells | Up | Galnt16 | 1.07E-06 | 0.31068233 | 0.232 | 0.79 | 0.00213108 |
| Smooth Muscle Cells | Up | Ccn1 | 9.67E-20 | 0.31300629 | 0.101 | 0.812 | 1.93E-16 |
| Smooth Muscle Cells | Up | Aox3 | 9.66E-14 | 0.32023096 | 0.064 | 0.333 | 1.93E-10 |
| Smooth Muscle Cells | Up | Nppa | 9.25E-16 | 0.32435667 | 0.086 | 0.769 | 1.85E-12 |
| Smooth Muscle Cells | Up | Tmem50b | 2.55E-40 | 0.32581079 | 0.075 | 0.903 | 5.11E-37 |

| | | | | | | | |
|---|---|---|---|---|---|---|---|
| Smooth Muscle Cells | Up | Trpc6 | 1.05E-08 | 0.32795895 | 0.165 | 0.763 | 2.09E-05 |
| Smooth Muscle Cells | Up | Pde10a | 2.51E-23 | 0.35786431 | 0.154 | 0.887 | 5.02E-20 |
| Smooth Muscle Cells | Up | Lrrtm3 | 9.46E-31 | 0.36818011 | 0.191 | 0.978 | 1.89E-27 |
| Smooth Muscle Cells | Up | Igfbp5 | 2.20E-10 | 0.39087042 | 0.228 | 0.839 | 4.40E-07 |
| Smooth Muscle Cells | Up | Rtn4rl1 | 1.01E-17 | 0.4126974 | 0.127 | 0.806 | 2.03E-14 |
| Smooth Muscle Cells | Up | Cobl | 1.17E-20 | 0.41530982 | 0.213 | 0.93 | 2.34E-17 |
| Smooth Muscle Cells | Up | Postn | 8.06E-47 | 0.45865176 | 0.06 | 0.903 | 1.61E-43 |
| Smooth Muscle Cells | Up | Stk38l | 3.13E-07 | 0.56297117 | 0.292 | 0.866 | 0.00062585 |
| Smooth Muscle Cells | Up | Srgap1 | 5.66E-20 | 0.58456646 | 0.236 | 0.957 | 1.13E-16 |
| Smooth Muscle Cells | Up | Pi16 | 5.58E-10 | 0.60064433 | 0.079 | 0.71 | 1.12E-06 |
| Smooth Muscle Cells | Up | Trdn | 4.03E-07 | 0.72422872 | 0.187 | 0.446 | 0.00080538 |
