## Supplemental Table 15 for "Uncovering the Regional and Cell Specific Bioactivity of Injectable Extracellular Matrix Biomaterials in Myocardial Infarction through Spatial and Single Nucleus Transcriptomics"

| Supplementary Table 15. Differentially Expressed Genes Spatially Comparing ECM Hydrogel in the Infarct and Infarct Only Zones with Integrated Subacute and Chronic MI | | | | | | | |
|---|---|---|---|---|---|---|---|
| **Spatial Area** | **Direction** | **Gene** | **p_val** | **avg_log2FC** | **pct.1** | **pct.2** | **p_val_adj** |
| ECM Hydrogel in Infarct | Up | Col8a1 | 4.67E-75 | 1.2365319 | 0.805 | 0.407 | 1.18E-70 |
| ECM Hydrogel in Infarct | Up | Tnc | 2.18E-48 | 1.15267302 | 0.499 | 0.171 | 5.52E-44 |
| ECM Hydrogel in Infarct | Up | Ltbp2 | 2.44E-87 | 1.14769842 | 0.962 | 0.76 | 6.16E-83 |
| ECM Hydrogel in Infarct | Up | Ccn2 | 7.77E-69 | 1.084305 | 0.962 | 0.754 | 1.96E-64 |
| ECM Hydrogel in Infarct | Up | Serpine1 | 2.16E-54 | 1.08203497 | 0.627 | 0.269 | 5.47E-50 |
| ECM Hydrogel in Infarct | Up | Thbs4 | 3.89E-37 | 1.03721615 | 0.519 | 0.234 | 9.82E-33 |
| ECM Hydrogel in Infarct | Up | Xirp2 | 2.11E-38 | 1.00830847 | 0.711 | 0.438 | 5.32E-34 |
| ECM Hydrogel in Infarct | Up | Cthrc1 | 7.89E-35 | 0.89965275 | 0.809 | 0.591 | 1.99E-30 |
| ECM Hydrogel in Infarct | Up | Cilp | 1.30E-33 | 0.85862532 | 0.727 | 0.473 | 3.29E-29 |
| ECM Hydrogel in Infarct | Up | Lox | 1.76E-46 | 0.85635513 | 0.878 | 0.658 | 4.46E-42 |
| ECM Hydrogel in Infarct | Up | Clec11a | 4.23E-45 | 0.80269884 | 0.763 | 0.458 | 1.07E-40 |
| ECM Hydrogel in Infarct | Up | Csrp2 | 1.80E-38 | 0.79150061 | 0.745 | 0.47 | 4.54E-34 |
| ECM Hydrogel in Infarct | Up | Nppb | 1.99E-25 | 0.79141403 | 0.805 | 0.604 | 5.02E-21 |
| ECM Hydrogel in Infarct | Up | Fibin | 9.00E-51 | 0.75745383 | 0.901 | 0.667 | 2.27E-46 |
| ECM Hydrogel in Infarct | Up | Col12a1 | 7.24E-31 | 0.73010185 | 0.501 | 0.226 | 1.83E-26 |
| ECM Hydrogel in Infarct | Up | Pdlim3 | 3.42E-27 | 0.7292766 | 0.667 | 0.444 | 8.64E-23 |
| ECM Hydrogel in Infarct | Up | Col8a2 | 2.56E-37 | 0.72441304 | 0.666 | 0.385 | 6.46E-33 |
| ECM Hydrogel in Infarct | Up | Aspn | 2.00E-33 | 0.67952366 | 0.835 | 0.618 | 5.06E-29 |
| ECM Hydrogel in Infarct | Up | Plod2 | 1.48E-28 | 0.63545068 | 0.667 | 0.42 | 3.73E-24 |
| ECM Hydrogel in Infarct | Up | Pmepa1 | 6.59E-38 | 0.62535466 | 0.811 | 0.507 | 1.66E-33 |
| ECM Hydrogel in Infarct | Up | Serpine2 | 3.74E-24 | 0.61336617 | 0.606 | 0.351 | 9.44E-20 |
| ECM Hydrogel in Infarct | Up | Ckap4 | 3.38E-25 | 0.56463716 | 0.685 | 0.442 | 8.55E-21 |
| ECM Hydrogel in Infarct | Up | C1qtnf3 | 2.10E-24 | 0.53931314 | 0.321 | 0.117 | 5.30E-20 |
| ECM Hydrogel in Infarct | Up | Enpp1 | 1.68E-28 | 0.53150359 | 0.672 | 0.411 | 4.24E-24 |
| ECM Hydrogel in Infarct | Up | Col16a1 | 4.37E-22 | 0.51942017 | 0.746 | 0.541 | 1.10E-17 |
| ECM Hydrogel in Infarct | Up | Fat1 | 3.03E-24 | 0.50150858 | 0.615 | 0.364 | 7.67E-20 |
| ECM Hydrogel in Infarct | Up | Ext1 | 2.93E-22 | 0.49500125 | 0.727 | 0.489 | 7.41E-18 |

| | | | | | | | |
|---|---|---|---|---|---|---|---|
| ECM Hydrogel in Infarct | Up | Nexn | 9.31E-22 | 0.49362768 | 0.437 | 0.202 | 2.35E-17 |
| ECM Hydrogel in Infarct | Up | Sdc1 | 5.15E-26 | 0.48707955 | 0.402 | 0.17 | 1.30E-21 |
| ECM Hydrogel in Infarct | Up | Tmem119 | 2.71E-23 | 0.4778117 | 0.606 | 0.36 | 6.85E-19 |
| ECM Hydrogel in Infarct | Up | Pls3 | 3.84E-20 | 0.45732279 | 0.837 | 0.631 | 9.70E-16 |
| ECM Hydrogel in Infarct | Up | Ddah1 | 3.70E-24 | 0.45217202 | 0.427 | 0.189 | 9.35E-20 |
| ECM Hydrogel in Infarct | Up | Spp1 | 9.79E-56 | 0.44679206 | 0.754 | 0.398 | 2.47E-51 |
| ECM Hydrogel in Infarct | Up | Cgref1 | 1.06E-26 | 0.42975746 | 0.35 | 0.122 | 2.68E-22 |
| ECM Hydrogel in Infarct | Up | Pabpc4 | 7.12E-18 | 0.40572971 | 0.779 | 0.558 | 1.80E-13 |
| ECM Hydrogel in Infarct | Up | Cdh2 | 7.85E-18 | 0.40027922 | 0.579 | 0.357 | 1.98E-13 |
| ECM Hydrogel in Infarct | Up | Ctsk | 9.58E-19 | 0.3967721 | 0.735 | 0.523 | 2.42E-14 |
| ECM Hydrogel in Infarct | Up | Rbp1 | 1.55E-16 | 0.37031069 | 0.746 | 0.518 | 3.92E-12 |
| ECM Hydrogel in Infarct | Up | Map1a | 3.37E-15 | 0.33800172 | 0.48 | 0.276 | 8.51E-11 |
| ECM Hydrogel in Infarct | Up | P3h1 | 2.72E-15 | 0.33242281 | 0.571 | 0.359 | 6.88E-11 |
| ECM Hydrogel in Infarct | Up | Spink8 | 6.36E-15 | 0.32184842 | 0.536 | 0.326 | 1.61E-10 |
| ECM Hydrogel in Infarct | Up | Fam162a | 7.31E-13 | 0.28068606 | 0.679 | 0.472 | 1.85E-08 |
| Infarct Only | Up | Lyve1 | 2.29E-24 | -0.8641376 | 0.195 | 0.403 | 5.79E-20 |
| Infarct Only | Up | Pla2g2a | 5.04E-26 | -1.1175613 | 0.246 | 0.473 | 1.27E-21 |
| Infarct Only | Up | Igfbp3 | 1.69E-38 | -1.272262 | 0.329 | 0.576 | 4.28E-34 |
| Infarct Only | Up | Cfd | 1.91E-60 | -1.3005305 | 0.507 | 0.775 | 4.84E-56 |
