## Supplemental Table 16 for "Uncovering the Regional and Cell Specific Bioactivity of Injectable Extracellular Matrix Biomaterials in Myocardial Infarction through Spatial and Single Nucleus Transcriptomics"

**Supplementary Table 16. Differentially Expressed Genes Spatially Comparing ECM Hydrogel in the Infarct between Subacute and Chronic MI**

| Spatial Sample | Direction | Gene | p_val | avg_log2FC | pct.1 | pct.2 | p_val_adj |
|---|---|---|---|---|---|---|---|
| Subacute | Up | Dbp | 4.31E-92 | 1.97244472 | 0.961 | 0.167 | 1.09E-87 |
| Subacute | Up | Cthrc1 | 5.20E-45 | 1.37996056 | 0.963 | 0.533 | 1.31E-40 |
| Subacute | Up | Mfap4 | 3.91E-37 | 1.21167492 | 0.765 | 0.228 | 9.87E-33 |
| Subacute | Up | NEWGENE-6: | 1.04E-23 | 1.01388527 | 0.947 | 0.598 | 2.63E-19 |
| Subacute | Up | P3h3 | 2.60E-42 | 1.01203085 | 0.97 | 0.547 | 6.56E-38 |
| Subacute | Up | Mdk | 5.61E-41 | 0.93792517 | 0.702 | 0.12 | 1.42E-36 |
| Subacute | Up | Myadm | 2.20E-27 | 0.92017866 | 0.645 | 0.185 | 5.56E-23 |
| Subacute | Up | Nr1d1 | 4.31E-37 | 0.91670116 | 0.803 | 0.243 | 1.09E-32 |
| Subacute | Up | RT1-T24-3 | 2.88E-35 | 0.89412958 | 0.844 | 0.297 | 7.27E-31 |
| Subacute | Up | Ifi27l2b | 5.62E-28 | 0.78927454 | 0.98 | 0.75 | 1.42E-23 |
| Subacute | Up | Nr1d2 | 5.38E-33 | 0.78223296 | 0.667 | 0.152 | 1.36E-28 |
| Subacute | Up | Pdlim3 | 5.86E-19 | 0.76271548 | 0.805 | 0.42 | 1.48E-14 |
| Subacute | Up | LOC1009113 | 4.98E-26 | 0.7436241 | 0.89 | 0.457 | 1.26E-21 |
| Subacute | Up | Lgals3bp | 6.60E-27 | 0.72705773 | 0.959 | 0.656 | 1.67E-22 |
| Subacute | Up | Scg2 | 1.19E-09 | 0.71495363 | 0.43 | 0.192 | 3.00E-05 |
| Subacute | Up | C1qtnf3 | 2.54E-24 | 0.7056787 | 0.458 | 0.076 | 6.42E-20 |
| Subacute | Up | Tef | 8.49E-35 | 0.70267678 | 0.692 | 0.145 | 2.15E-30 |
| Subacute | Up | Fbln1 | 1.39E-14 | 0.696795 | 0.97 | 0.746 | 3.52E-10 |
| Subacute | Up | Itm2a | 1.11E-23 | 0.69552488 | 0.915 | 0.478 | 2.81E-19 |
| Subacute | Up | Cemip | 5.94E-11 | 0.66981662 | 0.542 | 0.261 | 1.50E-06 |
| Subacute | Up | RT1-T24-4 | 1.54E-22 | 0.65471976 | 0.787 | 0.326 | 3.90E-18 |
| Subacute | Up | Isg15 | 2.74E-23 | 0.64962948 | 0.527 | 0.127 | 6.93E-19 |
| Subacute | Up | Lrrc17 | 1.57E-24 | 0.62725293 | 0.763 | 0.29 | 3.96E-20 |
| Subacute | Up | Reg3b | 3.30E-15 | 0.62428583 | 0.323 | 0.065 | 8.34E-11 |
| Subacute | Up | Fndc1 | 2.55E-24 | 0.58903896 | 0.984 | 0.757 | 6.45E-20 |
| Subacute | Up | Col16a1 | 4.47E-17 | 0.56873202 | 0.88 | 0.507 | 1.13E-12 |
| Subacute | Up | Herpud1 | 3.93E-20 | 0.56665308 | 0.895 | 0.493 | 9.94E-16 |
| Subacute | Up | LOC1083496 | 1.88E-19 | 0.55401547 | 0.669 | 0.261 | 4.76E-15 |
| Subacute | Up | Gpx7 | 4.10E-18 | 0.55298415 | 0.923 | 0.529 | 1.04E-13 |
| Subacute | Up | Aspn | 4.69E-20 | 0.55242465 | 0.957 | 0.616 | 1.19E-15 |
| Subacute | Up | Myh10 | 3.90E-19 | 0.5497183 | 0.838 | 0.388 | 9.85E-15 |
| Subacute | Up | Cpz | 9.89E-16 | 0.54836238 | 0.615 | 0.239 | 2.50E-11 |
| Subacute | Up | LOC1003601 | 1.46E-17 | 0.54637873 | 0.935 | 0.543 | 3.70E-13 |
| Subacute | Up | Mfap2 | 1.78E-18 | 0.53677659 | 0.72 | 0.293 | 4.49E-14 |

| | | | | | | | |
|---|---|---|---|---|---|---|---|
| Subacute | Up | C1qtnf2 | 3.10E-16 | 0.53633847 | 0.819 | 0.399 | 7.84E-12 |
| Subacute | Up | RT1-S3 | 2.35E-21 | 0.53284593 | 0.671 | 0.221 | 5.94E-17 |
| Subacute | Up | AABR070546 | 1.97E-19 | 0.5258589 | 0.978 | 0.736 | 4.98E-15 |
| Subacute | Up | C1qtnf6 | 9.96E-18 | 0.52475341 | 0.88 | 0.464 | 2.52E-13 |
| Subacute | Up | Txndc5 | 1.36E-16 | 0.51686111 | 0.903 | 0.471 | 3.44E-12 |
| Subacute | Up | Id3 | 5.51E-18 | 0.51615451 | 0.966 | 0.717 | 1.39E-13 |
| Subacute | Up | Pmp22 | 1.14E-16 | 0.51529834 | 0.963 | 0.721 | 2.89E-12 |
| Subacute | Up | Ly6e | 5.89E-18 | 0.51251995 | 0.943 | 0.591 | 1.49E-13 |
| Subacute | Up | Thbd | 1.54E-19 | 0.50785977 | 0.623 | 0.203 | 3.89E-15 |
| Subacute | Up | LOC1025554 | 1.61E-18 | 0.50699138 | 0.86 | 0.395 | 4.07E-14 |
| Subacute | Up | Mx2 | 2.43E-20 | 0.49641057 | 0.391 | 0.062 | 6.14E-16 |
| Subacute | Up | LOC1003604 | 2.44E-14 | 0.48884604 | 0.921 | 0.551 | 6.18E-10 |
| Subacute | Up | Axl | 7.74E-15 | 0.48852685 | 0.945 | 0.598 | 1.96E-10 |
| Subacute | Up | Adamtsl2 | 1.14E-08 | 0.48012815 | 0.677 | 0.399 | 0.00028688 |
| Subacute | Up | Fam20c | 8.97E-15 | 0.47904206 | 0.858 | 0.464 | 2.27E-10 |
| Subacute | Up | Maged2 | 7.40E-16 | 0.47115802 | 0.74 | 0.326 | 1.87E-11 |
| Subacute | Up | Csrp2 | 1.03E-11 | 0.46874105 | 0.868 | 0.525 | 2.60E-07 |
| Subacute | Up | Tbc1d2b | 4.33E-18 | 0.46701152 | 0.639 | 0.228 | 1.10E-13 |
| Subacute | Up | Lamb1 | 2.34E-11 | 0.45172342 | 0.935 | 0.598 | 5.91E-07 |
| Subacute | Up | Ptgfrn | 2.16E-16 | 0.45148711 | 0.805 | 0.402 | 5.47E-12 |
| Subacute | Up | Scd2 | 2.23E-13 | 0.44889429 | 0.793 | 0.406 | 5.64E-09 |
| Subacute | Up | Adamts2 | 9.92E-12 | 0.4404407 | 0.961 | 0.746 | 2.51E-07 |
| Subacute | Up | Tspan4 | 1.44E-16 | 0.43709557 | 0.398 | 0.094 | 3.64E-12 |
| Subacute | Up | Fkbp10 | 2.60E-14 | 0.43628024 | 0.937 | 0.634 | 6.57E-10 |
| Subacute | Up | Adamts8 | 2.25E-16 | 0.42923219 | 0.412 | 0.101 | 5.70E-12 |
| Subacute | Up | Aldh1a1 | 6.85E-16 | 0.42402294 | 0.515 | 0.174 | 1.73E-11 |
| Subacute | Up | Dchs1 | 8.13E-16 | 0.42114655 | 0.647 | 0.254 | 2.05E-11 |
| Subacute | Up | Olfml3 | 1.79E-11 | 0.42035691 | 0.884 | 0.54 | 4.53E-07 |
| Subacute | Up | C1qtnf7 | 1.41E-18 | 0.41881448 | 0.767 | 0.304 | 3.55E-14 |
| Subacute | Up | Rcn3 | 4.70E-13 | 0.41763907 | 0.939 | 0.616 | 1.19E-08 |
| Subacute | Up | Cavin2 | 1.76E-16 | 0.41544577 | 0.751 | 0.322 | 4.46E-12 |
| Subacute | Up | Arf4 | 1.16E-15 | 0.41050068 | 0.919 | 0.525 | 2.93E-11 |
| Subacute | Up | Grcc10 | 5.81E-16 | 0.40986669 | 0.832 | 0.395 | 1.47E-11 |
| Subacute | Up | C1qtnf1 | 5.10E-14 | 0.40368393 | 0.842 | 0.435 | 1.29E-09 |
| Subacute | Up | AC128960.1 | 1.31E-12 | 0.40275674 | 0.797 | 0.42 | 3.30E-08 |
| Subacute | Up | C1qtnf5 | 1.75E-10 | 0.39955135 | 0.927 | 0.612 | 4.43E-06 |
| Subacute | Up | Tax1bp3 | 7.77E-16 | 0.39812206 | 0.801 | 0.362 | 1.96E-11 |
| Subacute | Up | RGD1308134 | 4.07E-15 | 0.39748529 | 0.438 | 0.127 | 1.03E-10 |

| | | | | | | | |
|---|---|---|---|---|---|---|---|
| Subacute | Up | Manf | 6.11E-15 | 0.39616144 | 0.803 | 0.384 | 1.54E-10 |
| Subacute | Up | Ext1 | 1.08E-12 | 0.39302616 | 0.862 | 0.486 | 2.72E-08 |
| Subacute | Up | AABR070654 | 3.28E-15 | 0.39254677 | 0.736 | 0.326 | 8.29E-11 |
| Subacute | Up | Rbm3 | 8.83E-14 | 0.38959248 | 0.815 | 0.391 | 2.23E-09 |
| Subacute | Up | Vkorc1 | 4.16E-11 | 0.38936173 | 0.937 | 0.645 | 1.05E-06 |
| Subacute | Up | Smtnl2 | 3.96E-16 | 0.38777714 | 0.316 | 0.051 | 1.00E-11 |
| Subacute | Up | Mx1 | 1.11E-18 | 0.38742894 | 0.353 | 0.051 | 2.80E-14 |
| Subacute | Up | Tmem119 | 7.27E-13 | 0.38086749 | 0.74 | 0.366 | 1.84E-08 |
| Subacute | Up | Wfdc1 | 5.49E-11 | 0.38006808 | 0.884 | 0.572 | 1.39E-06 |
| Subacute | Up | Cyp46a1 | 1.20E-14 | 0.38006521 | 0.481 | 0.156 | 3.03E-10 |
| Subacute | Up | Smim7 | 9.02E-17 | 0.37793012 | 0.643 | 0.239 | 2.28E-12 |
| Subacute | Up | Cys1 | 6.27E-16 | 0.37761734 | 0.369 | 0.083 | 1.58E-11 |
| Subacute | Up | Etfa | 2.35E-12 | 0.37749831 | 0.627 | 0.283 | 5.94E-08 |
| Subacute | Up | Gask1b | 2.02E-12 | 0.37627529 | 0.945 | 0.63 | 5.09E-08 |
| Subacute | Up | Rtp4 | 3.38E-14 | 0.3726664 | 0.402 | 0.112 | 8.55E-10 |
| Subacute | Up | Dab2ip | 1.01E-16 | 0.3717144 | 0.485 | 0.134 | 2.56E-12 |
| Subacute | Up | Galnt16 | 6.15E-12 | 0.37068881 | 0.819 | 0.417 | 1.55E-07 |
| Subacute | Up | Oas1a | 1.50E-14 | 0.36703209 | 0.42 | 0.116 | 3.79E-10 |
| Subacute | Up | Ncor2 | 1.86E-12 | 0.36621932 | 0.765 | 0.37 | 4.71E-08 |
| Subacute | Up | Niban2 | 3.91E-14 | 0.36612993 | 0.704 | 0.304 | 9.89E-10 |
| Subacute | Up | AC094643.2 | 1.04E-14 | 0.36395973 | 0.688 | 0.29 | 2.64E-10 |
| Subacute | Up | Uap1 | 3.39E-07 | 0.36205967 | 0.769 | 0.489 | 0.00856623 |
| Subacute | Up | Ctsk | 2.50E-09 | 0.36200111 | 0.856 | 0.518 | 6.32E-05 |
| Subacute | Up | Lsp1 | 1.91E-14 | 0.3617641 | 0.696 | 0.293 | 4.82E-10 |
| Subacute | Up | Tcf21 | 3.61E-15 | 0.361006 | 0.458 | 0.138 | 9.13E-11 |
| Subacute | Up | Add1 | 5.25E-12 | 0.36030723 | 0.858 | 0.453 | 1.33E-07 |
| Subacute | Up | Cnih1 | 7.13E-12 | 0.35979403 | 0.773 | 0.377 | 1.80E-07 |
| Subacute | Up | Pamr1 | 1.13E-12 | 0.35428282 | 0.327 | 0.083 | 2.85E-08 |
| Subacute | Up | Sdf4 | 8.39E-10 | 0.35328161 | 0.935 | 0.623 | 2.12E-05 |
| Subacute | Up | Thbs2 | 1.61E-11 | 0.35294453 | 0.878 | 0.493 | 4.08E-07 |
| Subacute | Up | LOC1009097 | 1.50E-13 | 0.35130839 | 0.542 | 0.196 | 3.79E-09 |
| Subacute | Up | Ppial4d | 6.34E-13 | 0.35036772 | 0.375 | 0.109 | 1.60E-08 |
| Subacute | Up | Fbln5 | 7.05E-09 | 0.34982399 | 0.748 | 0.438 | 0.00017806 |
| Subacute | Up | Mical2 | 7.57E-15 | 0.34890876 | 0.57 | 0.207 | 1.91E-10 |
| Subacute | Up | Marcksl1 | 4.29E-13 | 0.34885191 | 0.55 | 0.21 | 1.08E-08 |
| Subacute | Up | Sin3b | 6.14E-12 | 0.34527537 | 0.708 | 0.341 | 1.55E-07 |
| Subacute | Up | Emp1 | 5.59E-07 | 0.34421724 | 0.925 | 0.707 | 0.01413525 |
| Subacute | Up | Mtch1 | 4.59E-10 | 0.33620803 | 0.97 | 0.707 | 1.16E-05 |

| | | | | | | | |
|---|---|---|---|---|---|---|---|
| Subacute | Up | Crtapl1 | 6.05E-12 | 0.33618462 | 0.73 | 0.341 | 1.53E-07 |
| Subacute | Up | Nedd4 | 2.46E-12 | 0.33434156 | 0.984 | 0.746 | 6.22E-08 |
| Subacute | Up | Mmp23 | 1.81E-09 | 0.33361135 | 0.88 | 0.547 | 4.57E-05 |
| Subacute | Up | LOC498555 | 3.80E-09 | 0.33291497 | 0.71 | 0.366 | 9.59E-05 |
| Subacute | Up | Osr1 | 1.29E-10 | 0.33172672 | 0.767 | 0.399 | 3.27E-06 |
| Subacute | Up | Tmem229b | 2.32E-16 | 0.33124242 | 0.4 | 0.091 | 5.87E-12 |
| Subacute | Up | Siglec1 | 1.30E-11 | 0.33021953 | 0.359 | 0.105 | 3.30E-07 |
| Subacute | Up | Sfxn3 | 1.97E-15 | 0.33008722 | 0.688 | 0.279 | 4.97E-11 |
| Subacute | Up | Hspa5 | 2.79E-09 | 0.32981873 | 0.982 | 0.775 | 7.06E-05 |
| Subacute | Up | Stt3a | 1.24E-11 | 0.3287839 | 0.734 | 0.351 | 3.14E-07 |
| Subacute | Up | Tuba1c | 9.97E-13 | 0.32723089 | 0.653 | 0.275 | 2.52E-08 |
| Subacute | Up | Atp8b2 | 4.11E-11 | 0.32665289 | 0.432 | 0.159 | 1.04E-06 |
| Subacute | Up | Rhoq | 1.13E-08 | 0.32643485 | 0.844 | 0.507 | 0.00028487 |
| Subacute | Up | Pkd1 | 3.51E-09 | 0.32616795 | 0.919 | 0.587 | 8.88E-05 |
| Subacute | Up | Cpt1a | 2.39E-12 | 0.32585371 | 0.367 | 0.109 | 6.05E-08 |
| Subacute | Up | Snd1 | 9.84E-14 | 0.3247082 | 0.631 | 0.257 | 2.49E-09 |
| Subacute | Up | Arl2bp | 3.95E-13 | 0.32397234 | 0.673 | 0.283 | 9.98E-09 |
| Subacute | Up | Nipsnap3b | 1.90E-14 | 0.32355604 | 0.278 | 0.04 | 4.80E-10 |
| Subacute | Up | Col8a2 | 6.12E-11 | 0.32206477 | 0.795 | 0.435 | 1.55E-06 |
| Subacute | Up | Ddx47 | 1.17E-14 | 0.32066591 | 0.367 | 0.087 | 2.97E-10 |
| Subacute | Up | AC141489.1 | 9.05E-09 | 0.31993499 | 0.923 | 0.591 | 0.00022867 |
| Subacute | Up | Snai1 | 2.31E-14 | 0.31966066 | 0.544 | 0.192 | 5.84E-10 |
| Subacute | Up | Anxa5 | 1.50E-09 | 0.31832631 | 0.984 | 0.757 | 3.80E-05 |
| Subacute | Up | Slc12a4 | 1.70E-13 | 0.31566289 | 0.645 | 0.257 | 4.31E-09 |
| Subacute | Up | Tsc22d1 | 6.31E-09 | 0.31523609 | 0.982 | 0.779 | 0.00015952 |
| Subacute | Up | Cercam | 8.96E-13 | 0.31504974 | 0.777 | 0.359 | 2.26E-08 |
| Subacute | Up | Clec11a | 8.52E-09 | 0.31459866 | 0.864 | 0.583 | 0.00021523 |
| Subacute | Up | Hmgn2 | 1.57E-08 | 0.31437732 | 0.933 | 0.594 | 0.00039658 |
| Subacute | Up | Fabp4 | 2.26E-09 | 0.31344939 | 0.884 | 0.551 | 5.72E-05 |
| Subacute | Up | Calu | 2.32E-08 | 0.3118526 | 0.976 | 0.725 | 0.00058703 |
| Subacute | Up | Usp18 | 1.35E-11 | 0.30883438 | 0.29 | 0.069 | 3.41E-07 |
| Subacute | Up | Sema3c | 1.08E-14 | 0.30765143 | 0.645 | 0.25 | 2.73E-10 |
| Subacute | Up | Pik3ip1 | 3.96E-14 | 0.30739773 | 0.434 | 0.123 | 1.00E-09 |
| Subacute | Up | Trim72 | 2.15E-09 | 0.30587319 | 0.629 | 0.315 | 5.44E-05 |
| Subacute | Up | Adarb1 | 3.82E-12 | 0.30550592 | 0.6 | 0.254 | 9.64E-08 |
| Subacute | Up | Tnfsf12 | 1.20E-10 | 0.3038205 | 0.625 | 0.279 | 3.04E-06 |
| Subacute | Up | Ppa1 | 1.81E-08 | 0.30347547 | 0.947 | 0.678 | 0.00045711 |
| Subacute | Up | Ssr2 | 2.73E-08 | 0.30307715 | 0.947 | 0.627 | 0.00068986 |

| | | | | | | | |
|---|---|---|---|---|---|---|---|
| Subacute | Up | Anxa1 | 8.34E-10 | 0.30244482 | 0.955 | 0.681 | 2.11E-05 |
| Subacute | Up | Fkbp11 | 2.11E-11 | 0.30123734 | 0.791 | 0.388 | 5.33E-07 |
| Subacute | Up | Per2 | 2.09E-13 | 0.30091332 | 0.28 | 0.051 | 5.29E-09 |
| Subacute | Up | Steap3 | 9.24E-14 | 0.30069932 | 0.588 | 0.225 | 2.33E-09 |
| Subacute | Up | Smo | 3.06E-13 | 0.29995206 | 0.389 | 0.109 | 7.72E-09 |
| Subacute | Up | Marveld1 | 1.84E-09 | 0.29886239 | 0.824 | 0.475 | 4.66E-05 |
| Subacute | Up | Senp3 | 2.47E-14 | 0.2976939 | 0.29 | 0.047 | 6.25E-10 |
| Subacute | Up | Gpx8 | 1.68E-10 | 0.29742428 | 0.815 | 0.457 | 4.24E-06 |
| Subacute | Up | Hmgb1 | 2.21E-13 | 0.29684594 | 0.329 | 0.072 | 5.59E-09 |
| Subacute | Up | Pdgfrl | 7.07E-11 | 0.29671327 | 0.726 | 0.351 | 1.79E-06 |
| Subacute | Up | Pigt | 1.40E-09 | 0.29579304 | 0.836 | 0.493 | 3.54E-05 |
| Subacute | Up | Ric8a | 2.95E-14 | 0.29573889 | 0.475 | 0.152 | 7.46E-10 |
| Subacute | Up | Ube2l6 | 7.61E-15 | 0.29511021 | 0.379 | 0.091 | 1.92E-10 |
| Subacute | Up | Fkbp9 | 7.47E-09 | 0.29464166 | 0.901 | 0.576 | 0.00018872 |
| Subacute | Up | Nr1h2 | 3.83E-12 | 0.29433332 | 0.515 | 0.196 | 9.68E-08 |
| Subacute | Up | Gbp2 | 2.49E-12 | 0.29377198 | 0.355 | 0.098 | 6.30E-08 |
| Subacute | Up | Anpep | 3.30E-12 | 0.29369784 | 0.515 | 0.196 | 8.35E-08 |
| Subacute | Up | Hnrnpab | 2.37E-09 | 0.29042111 | 0.917 | 0.587 | 6.00E-05 |
| Subacute | Up | Nucb1 | 2.12E-07 | 0.28966571 | 0.899 | 0.616 | 0.0053677 |
| Subacute | Up | Copz2 | 3.86E-10 | 0.28726706 | 0.795 | 0.417 | 9.76E-06 |
| Subacute | Up | Ephx1 | 3.23E-14 | 0.28685019 | 0.564 | 0.199 | 8.17E-10 |
| Subacute | Up | Rcn2 | 7.31E-15 | 0.28676206 | 0.54 | 0.181 | 1.85E-10 |
| Subacute | Up | Cd48 | 1.41E-09 | 0.28661429 | 0.746 | 0.38 | 3.56E-05 |
| Subacute | Up | Jund | 1.52E-09 | 0.28636228 | 0.862 | 0.525 | 3.85E-05 |
| Subacute | Up | Rbbp4 | 4.05E-13 | 0.28592132 | 0.525 | 0.188 | 1.02E-08 |
| Subacute | Up | Add3 | 2.88E-11 | 0.28491584 | 0.712 | 0.322 | 7.27E-07 |
| Subacute | Up | Tgfb3 | 1.29E-07 | 0.284517 | 0.959 | 0.699 | 0.00326339 |
| Subacute | Up | Ost4 | 2.16E-08 | 0.28444108 | 0.884 | 0.518 | 0.00054579 |
| Subacute | Up | Slc39a7 | 2.10E-09 | 0.28377573 | 0.868 | 0.504 | 5.32E-05 |
| Subacute | Up | Dynlt1 | 2.22E-14 | 0.28349459 | 0.487 | 0.152 | 5.61E-10 |
| Subacute | Up | Ndel1 | 4.95E-14 | 0.28315457 | 0.529 | 0.188 | 1.25E-09 |
| Subacute | Up | Sh3bgrl3 | 8.14E-09 | 0.28298631 | 0.941 | 0.601 | 0.00020585 |
| Subacute | Up | Ncstn | 4.55E-12 | 0.28284909 | 0.531 | 0.207 | 1.15E-07 |
| Subacute | Up | RGD1359290 | 4.69E-11 | 0.28186907 | 0.57 | 0.243 | 1.19E-06 |
| Subacute | Up | Trappc3 | 2.26E-12 | 0.28023351 | 0.564 | 0.221 | 5.71E-08 |
| Subacute | Up | Rtcb | 1.62E-13 | 0.28013925 | 0.584 | 0.221 | 4.08E-09 |
| Subacute | Up | Tln1 | 1.14E-07 | 0.27871052 | 0.929 | 0.58 | 0.00288189 |
| Subacute | Up | Spry1 | 3.95E-14 | 0.277864 | 0.4 | 0.105 | 9.98E-10 |

| | | | | | | | |
|---|---|---|---|---|---|---|---|
| Subacute | Up | Slfn4 | 2.44E-13 | 0.27776954 | 0.296 | 0.058 | 6.17E-09 |
| Subacute | Up | Rab31 | 3.10E-09 | 0.27765497 | 0.852 | 0.511 | 7.84E-05 |
| Subacute | Up | Eva1b | 8.78E-09 | 0.27714326 | 0.87 | 0.525 | 0.0002218 |
| Subacute | Up | Hnrnpf | 3.56E-11 | 0.27669624 | 0.767 | 0.377 | 9.00E-07 |
| Subacute | Up | Tmem120a | 7.93E-13 | 0.27604309 | 0.611 | 0.25 | 2.00E-08 |
| Subacute | Up | Lrrc59 | 4.02E-09 | 0.27591918 | 0.708 | 0.366 | 0.00010158 |
| Subacute | Up | Cdh2 | 1.39E-08 | 0.27552349 | 0.698 | 0.366 | 0.0003508 |
| Subacute | Up | Sppl3 | 1.33E-11 | 0.2751224 | 0.623 | 0.268 | 3.35E-07 |
| Subacute | Up | Cgref1 | 4.70E-11 | 0.27459471 | 0.452 | 0.167 | 1.19E-06 |
| Subacute | Up | Furin | 1.41E-10 | 0.27456189 | 0.692 | 0.322 | 3.56E-06 |
| Subacute | Up | Tmed3 | 4.97E-07 | 0.2738544 | 0.911 | 0.674 | 0.01254889 |
| Subacute | Up | Igsf10 | 3.97E-08 | 0.27379765 | 0.89 | 0.587 | 0.00100459 |
| Subacute | Up | MGC108823 | 1.40E-12 | 0.27293797 | 0.3 | 0.065 | 3.55E-08 |
| Subacute | Up | Man2b2 | 2.63E-12 | 0.27289397 | 0.675 | 0.297 | 6.64E-08 |
| Subacute | Up | Tmem86a | 9.99E-13 | 0.27254886 | 0.452 | 0.149 | 2.53E-08 |
| Subacute | Up | Hnrnpl | 5.04E-09 | 0.27235592 | 0.803 | 0.442 | 0.00012738 |
| Subacute | Up | Fam3a | 7.54E-12 | 0.27172908 | 0.347 | 0.098 | 1.91E-07 |
| Subacute | Up | Usp9x | 4.94E-11 | 0.27087592 | 0.702 | 0.322 | 1.25E-06 |
| Subacute | Up | Myrf | 8.71E-12 | 0.27023188 | 0.32 | 0.083 | 2.20E-07 |
| Subacute | Up | Sulf2 | 7.66E-09 | 0.26994129 | 0.913 | 0.638 | 0.00019359 |
| Subacute | Up | Rgcc | 1.31E-10 | 0.2699337 | 0.357 | 0.12 | 3.32E-06 |
| Subacute | Up | LOC1083496 | 4.07E-13 | 0.26968591 | 0.507 | 0.174 | 1.03E-08 |
| Subacute | Up | Id1 | 1.41E-10 | 0.26951198 | 0.669 | 0.308 | 3.57E-06 |
| Subacute | Up | Sel1l | 9.10E-15 | 0.26893232 | 0.59 | 0.21 | 2.30E-10 |
| Subacute | Up | Fcgr1a | 3.85E-11 | 0.26828651 | 0.371 | 0.12 | 9.74E-07 |
| Subacute | Up | Plxnb2 | 2.48E-08 | 0.26818051 | 0.872 | 0.565 | 0.00062597 |
| Subacute | Up | Enpp1 | 5.89E-10 | 0.26765782 | 0.799 | 0.446 | 1.49E-05 |
| Subacute | Up | Ppic | 1.40E-09 | 0.26765661 | 0.813 | 0.438 | 3.55E-05 |
| Subacute | Up | Trak2 | 1.54E-10 | 0.26761709 | 0.659 | 0.301 | 3.89E-06 |
| Subacute | Up | Ptn | 8.68E-10 | 0.26641563 | 0.619 | 0.297 | 2.19E-05 |
| Subacute | Up | Acsf2 | 2.71E-08 | 0.26633867 | 0.357 | 0.141 | 0.00068454 |
| Subacute | Up | Col4a5 | 6.76E-10 | 0.26606523 | 0.639 | 0.312 | 1.71E-05 |
| Subacute | Up | Tmed2 | 2.43E-08 | 0.2650601 | 0.963 | 0.667 | 0.00061479 |
| Subacute | Up | Igf2r | 2.79E-08 | 0.26202506 | 0.815 | 0.453 | 0.00070617 |
| Subacute | Up | Grb10 | 1.91E-10 | 0.26113622 | 0.789 | 0.409 | 4.83E-06 |
| Subacute | Up | Poglut3 | 3.55E-13 | 0.25984622 | 0.467 | 0.159 | 8.97E-09 |
| Subacute | Up | Glud1 | 2.87E-07 | 0.25879625 | 0.858 | 0.514 | 0.0072648 |
| Subacute | Up | Tmem214 | 1.78E-12 | 0.25828312 | 0.564 | 0.217 | 4.50E-08 |

| | | | | | | | |
|---|---|---|---|---|---|---|---|
| Subacute | Up | Ddit4 | 2.61E-09 | 0.25712675 | 0.552 | 0.25 | 6.60E-05 |
| Subacute | Up | Irf2bp1 | 6.09E-10 | 0.25631941 | 0.325 | 0.101 | 1.54E-05 |
| Subacute | Up | Sh2b1 | 2.33E-10 | 0.25612015 | 0.448 | 0.178 | 5.89E-06 |
| Subacute | Up | Cstb | 3.35E-07 | 0.25547749 | 0.907 | 0.591 | 0.00846756 |
| Subacute | Up | Eif4a1 | 5.20E-07 | 0.25466752 | 0.972 | 0.707 | 0.01314594 |
| Subacute | Up | Aldh1a2 | 2.67E-11 | 0.25430703 | 0.256 | 0.051 | 6.74E-07 |
| Subacute | Up | Rasl10a | 2.25E-11 | 0.2542989 | 0.422 | 0.145 | 5.69E-07 |
| Subacute | Up | Ppt1 | 3.16E-12 | 0.25347687 | 0.564 | 0.225 | 7.98E-08 |
| Subacute | Up | Fzd4 | 7.15E-13 | 0.2530024 | 0.335 | 0.08 | 1.81E-08 |
| Subacute | Up | LOC1009109 | 9.87E-11 | 0.25268921 | 0.546 | 0.228 | 2.49E-06 |
| Subacute | Up | B4galt2 | 6.12E-10 | 0.25261888 | 0.28 | 0.076 | 1.55E-05 |
| Subacute | Up | Ehmt2 | 1.22E-10 | 0.25227236 | 0.665 | 0.304 | 3.09E-06 |
| Subacute | Up | Stat3 | 4.40E-09 | 0.25199245 | 0.649 | 0.315 | 0.00011116 |
| Subacute | Up | Phgdh | 1.56E-11 | 0.25079722 | 0.252 | 0.047 | 3.93E-07 |
| Subacute | Up | Sgce | 2.46E-09 | 0.25058801 | 0.3 | 0.094 | 6.22E-05 |
| Subacute | Up | Twsg1 | 1.96E-15 | 0.25026098 | 0.639 | 0.225 | 4.95E-11 |
| Subacute | Up | Morf4l2 | 1.24E-08 | 0.2502363 | 0.941 | 0.62 | 0.00031223 |
| Chronic | Up | Comp | 1.64E-16 | -1.2517378 | 0.118 | 0.33 | 4.14E-12 |
