## Supplemental Table 17 for "Uncovering the Regional and Cell Specific Bioactivity of Injectable Extracellular Matrix Biomaterials in Myocardial Infarction through Spatial and Single Nucleus Transcriptomics"

**Supplementary Table 17. ECM Hydrogel Treated Subacute Remote Zones Vs. ECM Hydrogel Treated Chronic Remote Zones**

| Spatial Area | Directio | Gene | p_val | avg_log2FC | pct.1 | pct.2 | p_val_adj |
|---|---|---|---|---|---|---|---|
| Subacute Remote | Up | Klrb1c | 1.64E-128 | 1.91716601 | 0.287 | 0.489 | 3.29E-125 |
| Subacute Remote | Up | Reg3g | 1.39E-243 | 1.87074151 | 0.538 | 0.896 | 2.79E-240 |
| Subacute Remote | Up | Atp7a | 6.47E-181 | 1.77264148 | 0.36 | 0.591 | 1.29E-177 |
| Subacute Remote | Up | LOC1025525 | 8.16E-172 | 1.62730503 | 0.324 | 0.593 | 1.63E-168 |
| Subacute Remote | Up | Rdh12 | 3.83E-100 | 1.51444035 | 0.532 | 0.763 | 7.65E-97 |
| Subacute Remote | Up | Scg2 | 0 | 1.48307805 | 0.311 | 0.908 | 0 |
| Subacute Remote | Up | Ptgs2 | 6.57E-294 | 1.47887014 | 0.305 | 0.566 | 1.31E-290 |
| Subacute Remote | Up | Cdk1 | 0 | 1.46459178 | 0.306 | 0.716 | 0 |
| Subacute Remote | Up | Kif20b | 5.71E-185 | 1.44133281 | 0.33 | 0.562 | 1.14E-181 |
| Subacute Remote | Up | Ccnb1 | 3.77E-119 | 1.41376421 | 0.357 | 0.595 | 7.53E-116 |
| Subacute Remote | Up | Ccl2 | 0 | 1.3687936 | 0.22 | 0.548 | 0 |
| Subacute Remote | Up | LOC1003595 | 0 | 1.36338488 | 0.216 | 0.449 | 0 |

| | | | | | | | |
|---|---|---|---|---|---|---|---|
| Subacute Remote | Up | Apba2 | 5.16E-229 | 1.34927519 | 0.508 | 0.767 | 1.03E-225 |
| Subacute Remote | Up | Knstrn | 1.64E-190 | 1.31983536 | 0.433 | 0.791 | 3.29E-187 |
| Subacute Remote | Up | Col7a1 | 4.36E-125 | 1.29607717 | 0.491 | 0.733 | 8.72E-122 |
| Subacute Remote | Up | Thbs4 | 0 | 1.2880526 | 0.249 | 0.475 | 0 |
| Subacute Remote | Up | Pkib | 5.21E-174 | 1.28630476 | 0.356 | 0.58 | 1.04E-170 |
| Subacute Remote | Up | Ccl12 | 2.67E-271 | 1.26191728 | 0.311 | 0.568 | 5.34E-268 |
| Subacute Remote | Up | Esyt3 | 1.52E-113 | 1.26063897 | 0.421 | 0.65 | 3.04E-110 |
| Subacute Remote | Up | Fcer1a | 1.44E-27 | 1.26055704 | 0.534 | 0.823 | 2.88E-24 |
| Subacute Remote | Up | Mcpt1 | 2.50E-214 | 1.23981223 | 0.507 | 0.866 | 5.00E-211 |
| Subacute Remote | Up | Cemip | 0 | 1.2360757 | 0.316 | 0.903 | 0 |
| Subacute Remote | Up | Evc2 | 1.73E-134 | 1.23439243 | 0.444 | 0.691 | 3.45E-131 |
| Subacute Remote | Up | Gdf6 | 4.22E-233 | 1.21198069 | 0.394 | 0.628 | 8.45E-230 |
| Subacute Remote | Up | Agr2 | 1.60E-171 | 1.19059829 | 0.563 | 0.914 | 3.19E-168 |
| Subacute Remote | Up | Reg3b | 4.99E-68 | 1.16569724 | 0.567 | 0.907 | 9.98E-65 |

| | | | | | | | |
|---|---|---|---|---|---|---|---|
| Subacute Remote | Up | NEWGENE-6 | 0 | 1.15293733 | 0.287 | 0.831 | 0 |
| Subacute Remote | Up | Has2 | 1.43E-169 | 1.14462304 | 0.44 | 0.796 | 2.85E-166 |
| Subacute Remote | Up | Cdh11 | 0 | 1.1352649 | 0.332 | 0.645 | 0 |
| Subacute Remote | Up | Myrf | 0 | 1.13012415 | 0.383 | 0.734 | 0 |
| Subacute Remote | Up | Kif20a | 0 | 1.12565358 | 0.44 | 0.823 | 0 |
| Subacute Remote | Up | Clmp | 3.45E-193 | 1.11647153 | 0.414 | 0.637 | 6.90E-190 |
| Subacute Remote | Up | Ngef | 6.96E-266 | 1.11539998 | 0.427 | 0.694 | 1.39E-262 |
| Subacute Remote | Up | Aspm | 0 | 1.07803922 | 0.358 | 0.731 | 0 |
| Subacute Remote | Up | Cxcl13 | 0 | 1.07528885 | 0.335 | 0.692 | 0 |
| Subacute Remote | Up | Trpv2 | 1.30E-249 | 1.0728709 | 0.346 | 0.564 | 2.59E-246 |
| Subacute Remote | Up | Cyp46a1 | 0 | 1.06664798 | 0.323 | 0.768 | 0 |
| Subacute Remote | Up | Crabp2.1 | 0 | 1.06608536 | 0.388 | 0.788 | 0 |
| Subacute Remote | Up | Nub1 | 1.33E-153 | 1.05510235 | 0.501 | 0.753 | 2.66E-150 |
| Subacute Remote | Up | Mmp16 | 1.40E-163 | 1.05270202 | 0.477 | 0.747 | 2.81E-160 |

| | | | | | | | |
|---|---|---|---|---|---|---|---|
| Subacute Remote | Up | N4bp2 | 2.96E-120 | 1.04246145 | 0.537 | 0.757 | 5.91E-117 |
| Subacute Remote | Up | Ncaph | 5.58E-200 | 1.0397637 | 0.453 | 0.719 | 1.12E-196 |
| Subacute Remote | Up | Fermt3 | 0 | 1.03702263 | 0.236 | 0.573 | 0 |
| Subacute Remote | Up | Pik3cd | 0 | 1.03188257 | 0.398 | 0.739 | 0 |
| Subacute Remote | Up | Kif23 | 2.10E-292 | 1.0315743 | 0.449 | 0.835 | 4.20E-289 |
| Subacute Remote | Up | Matn4 | 0 | 1.02404172 | 0.251 | 0.649 | 0 |
| Subacute Remote | Up | Cilp2 | 1.88E-91 | 1.02347936 | 0.491 | 0.775 | 3.75E-88 |
| Subacute Remote | Up | Lrrc17 | 0 | 1.0184584 | 0.352 | 0.677 | 0 |
| Subacute Remote | Up | Tc2n | 3.37E-248 | 1.00897381 | 0.387 | 0.647 | 6.74E-245 |
| Subacute Remote | Up | Mfap4 | 0 | 1.00416144 | 0.199 | 0.729 | 0 |
| Subacute Remote | Up | Mki67 | 0 | 0.9994006 | 0.359 | 0.872 | 0 |
| Subacute Remote | Up | AABR070400 | 2.83E-306 | 0.99476534 | 0.513 | 0.821 | 5.65E-303 |
| Subacute Remote | Up | Cdc25c | 5.59E-253 | 0.99402675 | 0.375 | 0.649 | 1.12E-249 |
| Subacute Remote | Up | Fam111a | 4.08E-275 | 0.99325481 | 0.451 | 0.772 | 8.16E-272 |

| | | | | | | | |
|---|---|---|---|---|---|---|---|
| Subacute Remote | Up | Adamts8 | 4.19E-192 | 0.98750692 | 0.434 | 0.746 | 8.39E-189 |
| Subacute Remote | Up | Adgre1 | 1.79E-176 | 0.98742137 | 0.446 | 0.689 | 3.58E-173 |
| Subacute Remote | Up | Trh | 7.03E-130 | 0.98620203 | 0.358 | 0.574 | 1.41E-126 |
| Subacute Remote | Up | Lilrb4 | 0 | 0.98325401 | 0.25 | 0.549 | 0 |
| Subacute Remote | Up | Gem | 0 | 0.98134822 | 0.326 | 0.562 | 0 |
| Subacute Remote | Up | C1qtnf3 | 2.07E-95 | 0.96689073 | 0.546 | 0.806 | 4.13E-92 |
| Subacute Remote | Up | Ptprv | 8.56E-178 | 0.96398942 | 0.426 | 0.75 | 1.71E-174 |
| Subacute Remote | Up | Slc2a10 | 0 | 0.95932235 | 0.229 | 0.623 | 0 |
| Subacute Remote | Up | Tac3 | 1.96E-187 | 0.95048715 | 0.457 | 0.707 | 3.92E-184 |
| Subacute Remote | Up | Osr2 | 1.76E-149 | 0.94878972 | 0.467 | 0.749 | 3.51E-146 |
| Subacute Remote | Up | Hmcn1 | 1.23E-188 | 0.94666015 | 0.462 | 0.691 | 2.46E-185 |
| Subacute Remote | Up | Arhgap30 | 2.17E-219 | 0.94523159 | 0.541 | 0.809 | 4.34E-216 |
| Subacute Remote | Up | Kctd15 | 0 | 0.91341867 | 0.368 | 0.709 | 0 |
| Subacute Remote | Up | Scara3 | 4.27E-249 | 0.90917341 | 0.462 | 0.696 | 8.54E-246 |

| | | | | | | | |
|---|---|---|---|---|---|---|---|
| Subacute Remote | Up | Sfrp2 | 0 | 0.90805771 | 0.272 | 0.728 | 0 |
| Subacute Remote | Up | Mfap2 | 0 | 0.9017947 | 0.369 | 0.832 | 0 |
| Subacute Remote | Up | Clec11a | 0 | 0.89761099 | 0.285 | 0.505 | 0 |
| Subacute Remote | Up | Stmn4 | 1.78E-151 | 0.89611607 | 0.429 | 0.632 | 3.56E-148 |
| Subacute Remote | Up | Top2a | 0 | 0.88636113 | 0.319 | 0.575 | 0 |
| Subacute Remote | Up | Lox | 1.13E-246 | 0.87651189 | 0.447 | 0.812 | 2.25E-243 |
| Subacute Remote | Up | Cthrc1 | 0 | 0.86667877 | 0.344 | 0.934 | 0 |
| Subacute Remote | Up | LOC690045 | 7.92E-187 | 0.86221496 | 0.468 | 0.767 | 1.58E-183 |
| Subacute Remote | Up | Cpz | 0 | 0.86091848 | 0.331 | 0.577 | 0 |
| Subacute Remote | Up | Rasa3 | 3.14E-304 | 0.85614398 | 0.329 | 0.585 | 6.28E-301 |
| Subacute Remote | Up | Cd86 | 1.09E-147 | 0.85108738 | 0.501 | 0.735 | 2.17E-144 |
| Subacute Remote | Up | Ckap2 | 0 | 0.84684509 | 0.424 | 0.865 | 0 |
| Subacute Remote | Up | Ltbp2 | 1.30E-251 | 0.84637262 | 0.532 | 0.983 | 2.61E-248 |
| Subacute Remote | Up | Hck | 2.40E-307 | 0.84258774 | 0.483 | 0.793 | 4.81E-304 |

| | | | | | | | |
|---|---|---|---|---|---|---|---|
| Subacute Remote | Up | Haus7 | 3.61E-265 | 0.84008778 | 0.373 | 0.615 | 7.22E-262 |
| Subacute Remote | Up | Tbx18 | 0 | 0.83960014 | 0.218 | 0.514 | 0 |
| Subacute Remote | Up | Man2a1 | 0 | 0.83585583 | 0.38 | 0.755 | 0 |
| Subacute Remote | Up | Parvg | 3.22E-114 | 0.83190173 | 0.468 | 0.728 | 6.43E-111 |
| Subacute Remote | Up | Slc20a1 | 0 | 0.82713536 | 0.338 | 0.638 | 0 |
| Subacute Remote | Up | Adarb1 | 3.82E-224 | 0.82650536 | 0.389 | 0.628 | 7.63E-221 |
| Subacute Remote | Up | Nipsnap3b | 2.38E-264 | 0.82441663 | 0.48 | 0.887 | 4.75E-261 |
| Subacute Remote | Up | Cd83 | 0 | 0.82118917 | 0.375 | 0.765 | 0 |
| Subacute Remote | Up | Sall1 | 1.03E-221 | 0.8177929 | 0.41 | 0.651 | 2.05E-218 |
| Subacute Remote | Up | Scarb1 | 1.38E-304 | 0.81374335 | 0.376 | 0.672 | 2.77E-301 |
| Subacute Remote | Up | Cenpw | 3.01E-203 | 0.81175546 | 0.296 | 0.504 | 6.03E-200 |
| Subacute Remote | Up | Ly86 | 2.65E-224 | 0.80080096 | 0.453 | 0.766 | 5.29E-221 |
| Subacute Remote | Up | Ube2c | 0 | 0.79963605 | 0.293 | 0.696 | 0 |
| Subacute Remote | Up | Dse | 0 | 0.79888842 | 0.209 | 0.591 | 0 |

| | | | | | | | |
|---|---|---|---|---|---|---|---|
| Subacute Remote | Up | Gata3 | 1.53E-292 | 0.79884991 | 0.523 | 0.816 | 3.05E-289 |
| Subacute Remote | Up | Pttg1 | 5.39E-193 | 0.79210179 | 0.502 | 0.784 | 1.08E-189 |
| Subacute Remote | Up | Endou | 1.92E-294 | 0.78977618 | 0.27 | 0.487 | 3.84E-291 |
| Subacute Remote | Up | Prc1 | 0 | 0.78760755 | 0.429 | 0.803 | 0 |
| Subacute Remote | Up | Trps1 | 7.83E-265 | 0.7875326 | 0.51 | 0.85 | 1.57E-261 |
| Subacute Remote | Up | Gabre | 1.07E-154 | 0.78675887 | 0.427 | 0.707 | 2.14E-151 |
| Subacute Remote | Up | Ccn3 | 0 | 0.77925352 | 0.273 | 0.754 | 0 |
| Subacute Remote | Up | Ccn5 | 0 | 0.77526378 | 0.362 | 0.951 | 0 |
| Subacute Remote | Up | Slc1a7 | 0 | 0.77380398 | 0.398 | 0.781 | 0 |
| Subacute Remote | Up | AABR070357 | 0 | 0.77363313 | 0.424 | 0.811 | 0 |
| Subacute Remote | Up | Bcat1 | 0 | 0.77268632 | 0.439 | 0.829 | 0 |
| Subacute Remote | Up | Cdkn2b | 1.91E-129 | 0.76791325 | 0.54 | 0.765 | 3.82E-126 |
| Subacute Remote | Up | Plod2 | 0 | 0.76684861 | 0.304 | 0.663 | 0 |
| Subacute Remote | Up | Tcf21 | 0 | 0.76498232 | 0.292 | 0.658 | 0 |

| | | | | | | | |
|---|---|---|---|---|---|---|---|
| Subacute Remote | Up | Hmgb2 | 0 | 0.76126764 | 0.343 | 0.668 | 0 |
| Subacute Remote | Up | RGD1559482 | 0 | 0.75848808 | 0.402 | 0.853 | 0 |
| Subacute Remote | Up | Nnat | 5.48E-74 | 0.75817397 | 0.501 | 0.726 | 1.10E-70 |
| Subacute Remote | Up | Lilrb3a | 0 | 0.75329871 | 0.449 | 0.892 | 0 |
| Subacute Remote | Up | Ptn | 4.08E-120 | 0.74869228 | 0.514 | 0.789 | 8.17E-117 |
| Subacute Remote | Up | Irf8 | 0 | 0.74795388 | 0.302 | 0.617 | 0 |
| Subacute Remote | Up | Iqgap2 | 1.31E-269 | 0.74785405 | 0.511 | 0.756 | 2.62E-266 |
| Subacute Remote | Up | Sfrp4 | 1.80E-278 | 0.74220351 | 0.513 | 0.937 | 3.60E-275 |
| Subacute Remote | Up | P2ry6 | 0 | 0.73643122 | 0.329 | 0.619 | 0 |
| Subacute Remote | Up | Cip2a | 7.56E-102 | 0.73546907 | 0.547 | 0.762 | 1.51E-98 |
| Subacute Remote | Up | Anpep | 6.39E-246 | 0.73459901 | 0.409 | 0.694 | 1.28E-242 |
| Subacute Remote | Up | Vcan | 0 | 0.73270283 | 0.374 | 0.873 | 0 |
| Subacute Remote | Up | C5ar1 | 0 | 0.7289089 | 0.396 | 0.783 | 0 |
| Subacute Remote | Up | A2m | 9.39E-138 | 0.7285124 | 0.557 | 0.877 | 1.88E-134 |

| | | | | | | | |
|---|---|---|---|---|---|---|---|
| Subacute Remote | Up | Angptl4 | 0 | 0.72643917 | 0.312 | 0.56 | 0 |
| Subacute Remote | Up | Rassf2 | 2.53E-255 | 0.72451671 | 0.341 | 0.564 | 5.05E-252 |
| Subacute Remote | Up | Gpc3 | 0 | 0.72061314 | 0.273 | 0.724 | 0 |
| Subacute Remote | Up | Chst12 | 2.36E-149 | 0.71992614 | 0.469 | 0.68 | 4.71E-146 |
| Subacute Remote | Up | Tsku | 2.75E-243 | 0.71963079 | 0.464 | 0.831 | 5.50E-240 |
| Subacute Remote | Up | Ccdc15 | 5.36E-198 | 0.71865783 | 0.534 | 0.846 | 1.07E-194 |
| Subacute Remote | Up | Cercam | 3.74E-269 | 0.71642126 | 0.438 | 0.83 | 7.48E-266 |
| Subacute Remote | Up | Capn6 | 2.44E-79 | 0.71609978 | 0.565 | 0.839 | 4.88E-76 |
| Subacute Remote | Up | Il1rn | 3.73E-91 | 0.71499694 | 0.405 | 0.611 | 7.46E-88 |
| Subacute Remote | Up | Dclk1 | 0 | 0.71294175 | 0.382 | 0.675 | 0 |
| Subacute Remote | Up | Mcpt1l1 | 0 | 0.71199955 | 0.306 | 0.703 | 0 |
| Subacute Remote | Up | Slc1a5 | 0 | 0.71064162 | 0.339 | 0.632 | 0 |
| Subacute Remote | Up | Clec4a1 | 1.67E-265 | 0.71059624 | 0.419 | 0.742 | 3.35E-262 |
| Subacute Remote | Up | Adamtsl2 | 0 | 0.70865767 | 0.332 | 0.766 | 0 |

| | | | | | | | |
|---|---|---|---|---|---|---|---|
| Subacute Remote | Up | Rassf5 | 2.96E-233 | 0.70743489 | 0.446 | 0.717 | 5.92E-230 |
| Subacute Remote | Up | Trem2 | 1.21E-251 | 0.70498689 | 0.424 | 0.757 | 2.43E-248 |
| Subacute Remote | Up | Cp | 1.49E-241 | 0.70478054 | 0.436 | 0.772 | 2.99E-238 |
| Subacute Remote | Up | Fcgr2b | 6.89E-303 | 0.70314331 | 0.394 | 0.796 | 1.38E-299 |
| Subacute Remote | Up | Nfkbiz | 0 | 0.70149995 | 0.36 | 0.619 | 0 |
| Subacute Remote | Up | Mrc1 | 1.06E-294 | 0.69981246 | 0.428 | 0.817 | 2.12E-291 |
| Subacute Remote | Up | Pcdh18 | 0 | 0.69863534 | 0.373 | 0.689 | 0 |
| Subacute Remote | Up | Fam13a | 1.11E-209 | 0.69720474 | 0.54 | 0.819 | 2.22E-206 |
| Subacute Remote | Up | Lima1 | 6.54E-178 | 0.69177116 | 0.487 | 0.787 | 1.31E-174 |
| Subacute Remote | Up | Tbxas1 | 0 | 0.69057349 | 0.411 | 0.816 | 0 |
| Subacute Remote | Up | LOC1003604 | 9.57E-105 | 0.69051358 | 0.495 | 0.8 | 1.91E-101 |
| Subacute Remote | Up | Arrb2 | 2.43E-170 | 0.68987871 | 0.501 | 0.788 | 4.85E-167 |
| Subacute Remote | Up | Ctsk | 3.04E-95 | 0.68972249 | 0.543 | 0.798 | 6.08E-92 |
| Subacute Remote | Up | Ptgs1 | 0 | 0.68791252 | 0.355 | 0.718 | 0 |

| | | | | | | | |
|---|---|---|---|---|---|---|---|
| Subacute Remote | Up | Plpp5 | 0 | 0.68398355 | 0.286 | 0.685 | 0 |
| Subacute Remote | Up | Sertad4 | 0 | 0.68104215 | 0.376 | 0.734 | 0 |
| Subacute Remote | Up | Tmem63c | 1.02E-90 | 0.68013 | 0.481 | 0.72 | 2.04E-87 |
| Subacute Remote | Up | Cfd | 0 | 0.67992531 | 0.312 | 0.607 | 0 |
| Subacute Remote | Up | Grb10 | 1.30E-192 | 0.67801401 | 0.447 | 0.749 | 2.60E-189 |
| Subacute Remote | Up | Pgf | 7.17E-174 | 0.67634177 | 0.47 | 0.798 | 1.43E-170 |
| Subacute Remote | Up | Ms4a7 | 5.58E-136 | 0.67515031 | 0.479 | 0.712 | 1.12E-132 |
| Subacute Remote | Up | Cebpa | 0 | 0.67429098 | 0.318 | 0.588 | 0 |
| Subacute Remote | Up | Pdgfrl | 2.59E-255 | 0.6719497 | 0.402 | 0.726 | 5.19E-252 |
| Subacute Remote | Up | Adh6 | 1.39E-117 | 0.67084749 | 0.53 | 0.766 | 2.79E-114 |
| Subacute Remote | Up | Rab7b | 3.42E-170 | 0.66980955 | 0.509 | 0.84 | 6.85E-167 |
| Subacute Remote | Up | Dnm1 | 0 | 0.66810542 | 0.333 | 0.668 | 0 |
| Subacute Remote | Up | Bcl11b | 4.98E-284 | 0.66657366 | 0.442 | 0.794 | 9.95E-281 |
| Subacute Remote | Up | Pcdha13 | 4.84E-84 | 0.66606934 | 0.529 | 0.752 | 9.68E-81 |

| | | | | | | | |
|---|---|---|---|---|---|---|---|
| Subacute Remote | Up | Ptprc | 0 | 0.66556524 | 0.389 | 0.766 | 0 |
| Subacute Remote | Up | Chd3 | 0 | 0.66269904 | 0.36 | 0.708 | 0 |
| Subacute Remote | Up | Napsa | 3.24E-265 | 0.6587238 | 0.38 | 0.629 | 6.48E-262 |
| Subacute Remote | Up | Prps2 | 0 | 0.65811606 | 0.363 | 0.711 | 0 |
| Subacute Remote | Up | Meox1 | 3.37E-244 | 0.65767957 | 0.412 | 0.7 | 6.74E-241 |
| Subacute Remote | Up | Lbp | 0 | 0.65713134 | 0.315 | 0.574 | 0 |
| Subacute Remote | Up | Plvap | 1.93E-218 | 0.65499334 | 0.466 | 0.82 | 3.86E-215 |
| Subacute Remote | Up | Fibin | 4.91E-180 | 0.65417676 | 0.448 | 0.749 | 9.82E-177 |
| Subacute Remote | Up | Tmem229b | 7.73E-169 | 0.6505416 | 0.534 | 0.9 | 1.55E-165 |
| Subacute Remote | Up | Il17ra | 0 | 0.64985362 | 0.348 | 0.777 | 0 |
| Subacute Remote | Up | Atp8b2 | 0 | 0.64927583 | 0.268 | 0.791 | 0 |
| Subacute Remote | Up | Mmp23 | 4.47E-251 | 0.64763675 | 0.447 | 0.828 | 8.94E-248 |
| Subacute Remote | Up | Jag1 | 1.81E-265 | 0.64737778 | 0.452 | 0.766 | 3.63E-262 |
| Subacute Remote | Up | Vegfd | 1.47E-219 | 0.64350803 | 0.434 | 0.73 | 2.93E-216 |

| | | | | | | | |
|---|---|---|---|---|---|---|---|
| Subacute Remote | Up | Acta2 | 0 | 0.6413821 | 0.229 | 0.664 | 0 |
| Subacute Remote | Up | Mga | 2.07E-204 | 0.63923776 | 0.454 | 0.7 | 4.15E-201 |
| Subacute Remote | Up | Spi1 | 0 | 0.63894906 | 0.409 | 0.791 | 0 |
| Subacute Remote | Up | Fxyd6 | 0 | 0.63885474 | 0.277 | 0.696 | 0 |
| Subacute Remote | Up | Pla2g2d | 0 | 0.63874863 | 0.35 | 0.646 | 0 |
| Subacute Remote | Up | Pcsk5 | 1.10E-251 | 0.63860121 | 0.43 | 0.814 | 2.21E-248 |
| Subacute Remote | Up | Snx20 | 4.73E-205 | 0.63858739 | 0.439 | 0.655 | 9.46E-202 |
| Subacute Remote | Up | Slco2a1 | 1.49E-226 | 0.6375825 | 0.458 | 0.699 | 2.98E-223 |
| Subacute Remote | Up | Pamr1 | 1.17E-293 | 0.6372944 | 0.494 | 0.911 | 2.34E-290 |
| Subacute Remote | Up | Tasor | 3.98E-264 | 0.63725579 | 0.337 | 0.576 | 7.95E-261 |
| Subacute Remote | Up | Il4i1 | 1.94E-55 | 0.63705011 | 0.53 | 0.773 | 3.88E-52 |
| Subacute Remote | Up | Fads1 | 0 | 0.63531695 | 0.266 | 0.494 | 0 |
| Subacute Remote | Up | Mis18a | 3.57E-255 | 0.63365036 | 0.523 | 0.861 | 7.14E-252 |
| Subacute Remote | Up | Rnd3 | 1.94E-206 | 0.63334441 | 0.468 | 0.803 | 3.88E-203 |

| | | | | | | | |
|---|---|---|---|---|---|---|---|
| Subacute Remote | Up | Ssc5d | 1.85E-278 | 0.63275312 | 0.376 | 0.665 | 3.71E-275 |
| Subacute Remote | Up | Dpf1 | 2.50E-79 | 0.63209382 | 0.47 | 0.726 | 5.00E-76 |
| Subacute Remote | Up | Tgfbr1 | 1.13E-211 | 0.63142217 | 0.386 | 0.599 | 2.26E-208 |
| Subacute Remote | Up | Unc93b1 | 6.93E-289 | 0.63071987 | 0.469 | 0.86 | 1.39E-285 |
| Subacute Remote | Up | Actn1 | 2.96E-196 | 0.63024847 | 0.458 | 0.764 | 5.92E-193 |
| Subacute Remote | Up | Fcgr3a | 4.96E-133 | 0.62905655 | 0.55 | 0.922 | 9.93E-130 |
| Subacute Remote | Up | Cgref1 | 1.23E-235 | 0.62903793 | 0.442 | 0.822 | 2.45E-232 |
| Subacute Remote | Up | Tes | 0 | 0.62835791 | 0.375 | 0.791 | 0 |
| Subacute Remote | Up | Sncaip | 2.82E-184 | 0.62800152 | 0.427 | 0.652 | 5.63E-181 |
| Subacute Remote | Up | Ppic | 0 | 0.62772104 | 0.41 | 0.822 | 0 |
| Subacute Remote | Up | Ugt1a6 | 0 | 0.62696867 | 0.418 | 0.822 | 0 |
| Subacute Remote | Up | Myh10 | 1.81E-261 | 0.62673352 | 0.438 | 0.788 | 3.62E-258 |
| Subacute Remote | Up | Irf5 | 9.41E-255 | 0.62456066 | 0.425 | 0.753 | 1.88E-251 |
| Subacute Remote | Up | Aldh1a3 | 0 | 0.62327307 | 0.391 | 0.783 | 0 |

| | | | | | | | |
|---|---|---|---|---|---|---|---|
| Subacute Remote | Up | Ect2 | 3.40E-189 | 0.62224938 | 0.507 | 0.842 | 6.80E-186 |
| Subacute Remote | Up | Ptgfrn | 0 | 0.62204778 | 0.331 | 0.738 | 0 |
| Subacute Remote | Up | Col8a2 | 0 | 0.62181759 | 0.402 | 0.763 | 0 |
| Subacute Remote | Up | Nfatc4 | 0 | 0.62125358 | 0.382 | 0.725 | 0 |
| Subacute Remote | Up | Apcdd1 | 0 | 0.62071637 | 0.333 | 0.648 | 0 |
| Subacute Remote | Up | Tnfaip8l2 | 3.76E-163 | 0.620707 | 0.45 | 0.684 | 7.53E-160 |
| Subacute Remote | Up | Fkbp1b | 3.80E-232 | 0.62036313 | 0.5 | 0.813 | 7.60E-229 |
| Subacute Remote | Up | Fcer1g | 4.39E-78 | 0.61906411 | 0.63 | 0.969 | 8.78E-75 |
| Subacute Remote | Up | Kcnn4 | 3.08E-287 | 0.6180756 | 0.429 | 0.741 | 6.16E-284 |
| Subacute Remote | Up | Kctd17 | 0 | 0.6177876 | 0.369 | 0.696 | 0 |
| Subacute Remote | Up | Sema4a | 0 | 0.61747218 | 0.329 | 0.664 | 0 |
| Subacute Remote | Up | C1qb | 4.49E-54 | 0.615559 | 0.669 | 0.947 | 8.98E-51 |
| Subacute Remote | Up | Flrt3 | 6.85E-162 | 0.61528826 | 0.503 | 0.834 | 1.37E-158 |
| Subacute Remote | Up | Igf2 | 0 | 0.61480328 | 0.396 | 0.84 | 0 |

| | | | | | | | |
|---|---|---|---|---|---|---|---|
| Subacute Remote | Up | Cgnl1 | 8.61E-205 | 0.61456675 | 0.507 | 0.796 | 1.72E-201 |
| Subacute Remote | Up | Igsf10 | 2.05E-194 | 0.61299707 | 0.489 | 0.846 | 4.10E-191 |
| Subacute Remote | Up | Nckap1l | 2.78E-172 | 0.61292143 | 0.476 | 0.795 | 5.56E-169 |
| Subacute Remote | Up | Slc11a1 | 1.10E-205 | 0.61257188 | 0.492 | 0.76 | 2.20E-202 |
| Subacute Remote | Up | Lyve1 | 0 | 0.61252896 | 0.502 | 0.916 | 0 |
| Subacute Remote | Up | C1qtnf6 | 0 | 0.61247971 | 0.371 | 0.885 | 0 |
| Subacute Remote | Up | Evc | 0 | 0.61227212 | 0.361 | 0.721 | 0 |
| Subacute Remote | Up | Rcn3 | 5.64E-253 | 0.6120326 | 0.403 | 0.824 | 1.13E-249 |
| Subacute Remote | Up | Tmem237 | 1.44E-197 | 0.61135244 | 0.401 | 0.631 | 2.89E-194 |
| Subacute Remote | Up | Ptprf | 0 | 0.60971258 | 0.379 | 0.712 | 0 |
| Subacute Remote | Up | Itm2a | 2.81E-183 | 0.60775639 | 0.478 | 0.789 | 5.62E-180 |
| Subacute Remote | Up | Procr | 0 | 0.60749558 | 0.25 | 0.532 | 0 |
| Subacute Remote | Up | Col8a1 | 2.31E-226 | 0.60578368 | 0.501 | 0.868 | 4.63E-223 |
| Subacute Remote | Up | Camk2a | 8.66E-121 | 0.60386328 | 0.353 | 0.567 | 1.73E-117 |

| | | | | | | | |
|---|---|---|---|---|---|---|---|
| Subacute Remote | Up | Coro1a | 9.49E-131 | 0.60181654 | 0.52 | 0.77 | 1.90E-127 |
| Subacute Remote | Up | Cd276 | 3.50E-289 | 0.60176749 | 0.458 | 0.842 | 7.01E-286 |
| Subacute Remote | Up | Cnn3 | 2.54E-158 | 0.59971643 | 0.447 | 0.674 | 5.07E-155 |
| Subacute Remote | Up | C1qtnf7 | 8.66E-161 | 0.59917619 | 0.571 | 0.923 | 1.73E-157 |
| Subacute Remote | Up | Hic2 | 0 | 0.59869044 | 0.411 | 0.772 | 0 |
| Subacute Remote | Up | Nfam1 | 5.64E-227 | 0.59775766 | 0.477 | 0.813 | 1.13E-223 |
| Subacute Remote | Up | Pstpip1 | 0 | 0.59765909 | 0.387 | 0.742 | 0 |
| Subacute Remote | Up | Fcrla | 1.87E-55 | 0.59706743 | 0.568 | 0.853 | 3.74E-52 |
| Subacute Remote | Up | Cfh | 1.82E-113 | 0.59626046 | 0.532 | 0.811 | 3.65E-110 |
| Subacute Remote | Up | Kctd14 | 1.38E-114 | 0.59617697 | 0.507 | 0.772 | 2.75E-111 |
| Subacute Remote | Up | LOC691995 | 8.80E-171 | 0.5961397 | 0.521 | 0.848 | 1.76E-167 |
| Subacute Remote | Up | Smarca1 | 1.04E-89 | 0.59527477 | 0.497 | 0.751 | 2.08E-86 |
| Subacute Remote | Up | Slc16a3 | 9.79E-285 | 0.59459405 | 0.392 | 0.727 | 1.96E-281 |
| Subacute Remote | Up | Osr1 | 0 | 0.59457687 | 0.363 | 0.86 | 0 |

| | | | | | | | |
|---|---|---|---|---|---|---|---|
| Subacute Remote | Up | Rarres1 | 0 | 0.59357108 | 0.409 | 0.828 | 0 |
| Subacute Remote | Up | Nbeal1 | 2.34E-154 | 0.59241251 | 0.517 | 0.777 | 4.67E-151 |
| Subacute Remote | Up | Mycn | 0 | 0.59212979 | 0.347 | 0.722 | 0 |
| Subacute Remote | Up | Plac9 | 3.33E-242 | 0.59149136 | 0.362 | 0.594 | 6.66E-239 |
| Subacute Remote | Up | Colec11 | 1.35E-170 | 0.59030273 | 0.528 | 0.903 | 2.69E-167 |
| Subacute Remote | Up | Il2ra | 1.81E-160 | 0.5892567 | 0.386 | 0.604 | 3.63E-157 |
| Subacute Remote | Up | Clec10a | 9.34E-176 | 0.58736674 | 0.52 | 0.858 | 1.87E-172 |
| Subacute Remote | Up | Slc44a1 | 1.74E-124 | 0.58684774 | 0.483 | 0.736 | 3.48E-121 |
| Subacute Remote | Up | Pkp1 | 0 | 0.58632152 | 0.421 | 0.779 | 0 |
| Subacute Remote | Up | Smpd3 | 0 | 0.58605346 | 0.329 | 0.736 | 0 |
| Subacute Remote | Up | Col16a1 | 0 | 0.58424603 | 0.427 | 0.847 | 0 |
| Subacute Remote | Up | Fnbp1l | 6.72E-194 | 0.58122957 | 0.461 | 0.706 | 1.34E-190 |
| Subacute Remote | Up | Notch2 | 2.49E-211 | 0.58102915 | 0.472 | 0.815 | 4.99E-208 |
| Subacute Remote | Up | B4galt5 | 1.24E-203 | 0.58088172 | 0.43 | 0.633 | 2.48E-200 |

| | | | | | | | |
|---|---|---|---|---|---|---|---|
| Subacute Remote | Up | Pcolce | 2.77E-72 | 0.58087767 | 0.587 | 0.95 | 5.55E-69 |
| Subacute Remote | Up | Crlf1 | 2.23E-252 | 0.58050292 | 0.394 | 0.662 | 4.46E-249 |
| Subacute Remote | Up | Sdc1 | 0 | 0.58039975 | 0.388 | 0.813 | 0 |
| Subacute Remote | Up | Rgs1 | 3.16E-105 | 0.5803447 | 0.504 | 0.752 | 6.31E-102 |
| Subacute Remote | Up | Alox5ap | 1.18E-245 | 0.57973577 | 0.463 | 0.812 | 2.36E-242 |
| Subacute Remote | Up | Ptafr | 6.10E-212 | 0.57844512 | 0.469 | 0.751 | 1.22E-208 |
| Subacute Remote | Up | Ccl22 | 1.23E-196 | 0.57664196 | 0.277 | 0.492 | 2.47E-193 |
| Subacute Remote | Up | NEWGENE-6 | 2.15E-307 | 0.57621844 | 0.4 | 0.744 | 4.29E-304 |
| Subacute Remote | Up | Cd247 | 1.42E-189 | 0.5729888 | 0.468 | 0.688 | 2.84E-186 |
| Subacute Remote | Up | Tmed3 | 1.15E-95 | 0.57205733 | 0.573 | 0.857 | 2.29E-92 |
| Subacute Remote | Up | Tmem132e | 3.62E-204 | 0.57201686 | 0.448 | 0.741 | 7.23E-201 |
| Subacute Remote | Up | Pawr | 1.45E-160 | 0.57173351 | 0.45 | 0.653 | 2.90E-157 |
| Subacute Remote | Up | Ccl6 | 1.47E-200 | 0.57128246 | 0.486 | 0.806 | 2.94E-197 |
| Subacute Remote | Up | Cdkn3 | 0 | 0.5710524 | 0.467 | 0.855 | 0 |

| | | | | | | | |
|---|---|---|---|---|---|---|---|
| Subacute Remote | Up | Dnase2 | 1.05E-267 | 0.56944687 | 0.388 | 0.672 | 2.10E-264 |
| Subacute Remote | Up | Slit3 | 9.41E-281 | 0.5690322 | 0.313 | 0.553 | 1.88E-277 |
| Subacute Remote | Up | Lrrk1 | 3.62E-149 | 0.56860809 | 0.5 | 0.742 | 7.24E-146 |
| Subacute Remote | Up | Pak1 | 1.41E-250 | 0.56766833 | 0.438 | 0.793 | 2.82E-247 |
| Subacute Remote | Up | Cspg4 | 7.11E-177 | 0.56719143 | 0.465 | 0.74 | 1.42E-173 |
| Subacute Remote | Up | Mnda | 0 | 0.56631141 | 0.419 | 0.801 | 0 |
| Subacute Remote | Up | Pkd2 | 0 | 0.56550613 | 0.38 | 0.741 | 0 |
| Subacute Remote | Up | Twistnb | 4.31E-224 | 0.56517752 | 0.434 | 0.662 | 8.62E-221 |
| Subacute Remote | Up | Fgf18 | 1.63E-165 | 0.56400882 | 0.437 | 0.644 | 3.26E-162 |
| Subacute Remote | Up | Gpx7 | 7.92E-154 | 0.56373589 | 0.503 | 0.824 | 1.58E-150 |
| Subacute Remote | Up | Myo1g | 6.36E-172 | 0.56344112 | 0.449 | 0.71 | 1.27E-168 |
| Subacute Remote | Up | Cpa3 | 0 | 0.56301959 | 0.465 | 0.88 | 0 |
| Subacute Remote | Up | Colec12 | 1.10E-277 | 0.56223043 | 0.366 | 0.617 | 2.20E-274 |
| Subacute Remote | Up | Olfml2b | 0 | 0.56215401 | 0.39 | 0.731 | 0 |

| | | | | | | | |
|---|---|---|---|---|---|---|---|
| Subacute Remote | Up | Syt5 | 0 | 0.56010572 | 0.433 | 0.777 | 0 |
| Subacute Remote | Up | Ppt1 | 3.18E-234 | 0.55958081 | 0.468 | 0.801 | 6.36E-231 |
| Subacute Remote | Up | Serpine1 | 0 | 0.55947144 | 0.448 | 0.818 | 0 |
| Subacute Remote | Up | Atp2b4 | 4.72E-196 | 0.55921157 | 0.395 | 0.603 | 9.45E-193 |
| Subacute Remote | Up | Cdc20 | 0 | 0.55911167 | 0.445 | 0.813 | 0 |
| Subacute Remote | Up | Armcx2 | 0 | 0.55888181 | 0.4 | 0.764 | 0 |
| Subacute Remote | Up | Gpm6b | 2.42E-223 | 0.55876411 | 0.423 | 0.673 | 4.84E-220 |
| Subacute Remote | Up | Sfxn1 | 0 | 0.55609346 | 0.435 | 0.787 | 0 |
| Subacute Remote | Up | Basp1 | 1.03E-238 | 0.55565185 | 0.499 | 0.782 | 2.06E-235 |
| Subacute Remote | Up | Adamts2 | 3.29E-113 | 0.55535406 | 0.573 | 0.922 | 6.58E-110 |
| Subacute Remote | Up | Entpd1 | 1.75E-186 | 0.55416894 | 0.446 | 0.704 | 3.50E-183 |
| Subacute Remote | Up | Esco2 | 0 | 0.55352386 | 0.493 | 0.883 | 0 |
| Subacute Remote | Up | Numbl | 2.81E-229 | 0.55083601 | 0.451 | 0.755 | 5.63E-226 |
| Subacute Remote | Up | Tmem178a | 0 | 0.54925027 | 0.265 | 0.509 | 0 |

| | | | | | | | |
|---|---|---|---|---|---|---|---|
| Subacute Remote | Up | Loxl1 | 4.96E-17 | 0.54633644 | 0.657 | 0.98 | 9.92E-14 |
| Subacute Remote | Up | Plk2 | 0 | 0.54506247 | 0.3 | 0.575 | 0 |
| Subacute Remote | Up | Calhm2 | 0 | 0.54414987 | 0.368 | 0.83 | 0 |
| Subacute Remote | Up | Serpinf1 | 7.92E-16 | 0.54260408 | 0.663 | 0.914 | 1.58E-12 |
| Subacute Remote | Up | Prrx2 | 0 | 0.54243516 | 0.438 | 0.881 | 0 |
| Subacute Remote | Up | Fndc1 | 7.50E-100 | 0.54067497 | 0.606 | 0.93 | 1.50E-96 |
| Subacute Remote | Up | Itgbl1 | 2.05E-307 | 0.53877465 | 0.434 | 0.799 | 4.10E-304 |
| Subacute Remote | Up | Rai14 | 0 | 0.53834807 | 0.325 | 0.72 | 0 |
| Subacute Remote | Up | Ugdh | 0 | 0.53767024 | 0.356 | 0.666 | 0 |
| Subacute Remote | Up | Myo1f | 2.39E-262 | 0.53650999 | 0.469 | 0.819 | 4.77E-259 |
| Subacute Remote | Up | Cys1 | 0 | 0.53541412 | 0.473 | 0.817 | 0 |
| Subacute Remote | Up | Lcp1 | 8.54E-198 | 0.53515611 | 0.534 | 0.858 | 1.71E-194 |
| Subacute Remote | Up | Gask1b | 1.54E-136 | 0.53507588 | 0.569 | 0.925 | 3.09E-133 |
| Subacute Remote | Up | Arl4d | 0 | 0.53506307 | 0.36 | 0.631 | 0 |

| | | | | | | | |
|---|---|---|---|---|---|---|---|
| Subacute Remote | Up | Slfn13 | 0 | 0.53440472 | 0.308 | 0.785 | 0 |
| Subacute Remote | Up | Tgfb3 | 1.73E-137 | 0.53427775 | 0.563 | 0.897 | 3.46E-134 |
| Subacute Remote | Up | Plxna1 | 0 | 0.53407598 | 0.252 | 0.556 | 0 |
| Subacute Remote | Up | Twsg1 | 4.62E-148 | 0.53256631 | 0.475 | 0.699 | 9.25E-145 |
| Subacute Remote | Up | Tnfrsf11b | 0 | 0.5320448 | 0.378 | 0.837 | 0 |
| Subacute Remote | Up | Pon3 | 6.03E-171 | 0.53153717 | 0.438 | 0.71 | 1.21E-167 |
| Subacute Remote | Up | Fn1 | 5.20E-28 | 0.53056275 | 0.598 | 0.937 | 1.04E-24 |
| Subacute Remote | Up | Ltbp1 | 4.73E-106 | 0.53034141 | 0.544 | 0.811 | 9.45E-103 |
| Subacute Remote | Up | Klrb1b | 0 | 0.52994933 | 0.479 | 0.769 | 0 |
| Subacute Remote | Up | Zbp1 | 1.39E-196 | 0.52931293 | 0.528 | 0.879 | 2.79E-193 |
| Subacute Remote | Up | Scara5 | 0 | 0.52921835 | 0.433 | 0.812 | 0 |
| Subacute Remote | Up | LOC1025495 | 2.53E-288 | 0.52884613 | 0.507 | 0.849 | 5.07E-285 |
| Subacute Remote | Up | Sfxn3 | 2.05E-232 | 0.5284699 | 0.482 | 0.81 | 4.10E-229 |
| Subacute Remote | Up | Aspn | 2.71E-209 | 0.52839346 | 0.535 | 0.879 | 5.42E-206 |

| | | | | | | | |
|---|---|---|---|---|---|---|---|
| Subacute Remote | Up | Lsp1 | 0 | 0.52791827 | 0.33 | 0.845 | 0 |
| Subacute Remote | Up | Runx1t1 | 3.76E-178 | 0.52776768 | 0.442 | 0.651 | 7.52E-175 |
| Subacute Remote | Up | Nid1 | 7.83E-198 | 0.52668426 | 0.532 | 0.87 | 1.57E-194 |
| Subacute Remote | Up | Cpq | 6.81E-243 | 0.52657159 | 0.447 | 0.762 | 1.36E-239 |
| Subacute Remote | Up | Ibtk | 4.08E-168 | 0.52635836 | 0.529 | 0.767 | 8.16E-165 |
| Subacute Remote | Up | Sh3gl1 | 2.30E-255 | 0.52568589 | 0.437 | 0.775 | 4.61E-252 |
| Subacute Remote | Up | Pros1 | 0 | 0.52517513 | 0.284 | 0.627 | 0 |
| Subacute Remote | Up | Acsl4 | 0 | 0.52429904 | 0.32 | 0.555 | 0 |
| Subacute Remote | Up | Map4k1 | 1.58E-218 | 0.52421697 | 0.459 | 0.745 | 3.16E-215 |
| Subacute Remote | Up | Nptxr | 0 | 0.52419964 | 0.295 | 0.593 | 0 |
| Subacute Remote | Up | Crtapl1 | 8.45E-170 | 0.52414308 | 0.492 | 0.793 | 1.69E-166 |
| Subacute Remote | Up | Sulf2 | 2.01E-73 | 0.52403806 | 0.588 | 0.842 | 4.03E-70 |
| Subacute Remote | Up | Trem1 | 3.64E-178 | 0.52363494 | 0.518 | 0.757 | 7.28E-175 |
| Subacute Remote | Up | H2aj | 0 | 0.52331595 | 0.314 | 0.613 | 0 |

| | | | | | | | |
|---|---|---|---|---|---|---|---|
| Subacute Remote | Up | Tmem97 | 2.03E-171 | 0.52315195 | 0.429 | 0.63 | 4.06E-168 |
| Subacute Remote | Up | Akap12 | 2.57E-161 | 0.52268743 | 0.526 | 0.805 | 5.14E-158 |
| Subacute Remote | Up | Acap3 | 0 | 0.52028728 | 0.43 | 0.811 | 0 |
| Subacute Remote | Up | P3h4 | 0 | 0.51965873 | 0.412 | 0.785 | 0 |
| Subacute Remote | Up | Fbln1 | 2.07E-97 | 0.51965678 | 0.495 | 0.944 | 4.15E-94 |
| Subacute Remote | Up | Slamf8 | 8.27E-291 | 0.51955935 | 0.466 | 0.849 | 1.65E-287 |
| Subacute Remote | Up | Evi2a | 3.01E-294 | 0.51882608 | 0.51 | 0.848 | 6.02E-291 |
| Subacute Remote | Up | Cd44 | 0 | 0.51863964 | 0.374 | 0.759 | 0 |
| Subacute Remote | Up | Rhox5 | 0 | 0.5174824 | 0.438 | 0.714 | 0 |
| Subacute Remote | Up | Marcks | 1.01E-48 | 0.51716317 | 0.597 | 0.879 | 2.01E-45 |
| Subacute Remote | Up | Bmp1 | 2.18E-299 | 0.51535344 | 0.403 | 0.789 | 4.36E-296 |
| Subacute Remote | Up | Cmklr1 | 0 | 0.51527683 | 0.364 | 0.647 | 0 |
| Subacute Remote | Up | Piezo1 | 0 | 0.5150499 | 0.305 | 0.638 | 0 |
| Subacute Remote | Up | Arl11 | 1.01E-268 | 0.51366831 | 0.475 | 0.839 | 2.02E-265 |

| | | | | | | | |
|---|---|---|---|---|---|---|---|
| Subacute Remote | Up | Shc1 | 3.44E-188 | 0.51317101 | 0.435 | 0.692 | 6.88E-185 |
| Subacute Remote | Up | RT1-Da | 3.02E-38 | 0.51313268 | 0.653 | 0.924 | 6.04E-35 |
| Subacute Remote | Up | P3h3 | 2.47E-216 | 0.51196781 | 0.518 | 0.93 | 4.94E-213 |
| Subacute Remote | Up | Ecrg4 | 7.82E-219 | 0.51182779 | 0.457 | 0.822 | 1.56E-215 |
| Subacute Remote | Up | Niban2 | 1.63E-176 | 0.51153779 | 0.463 | 0.742 | 3.26E-173 |
| Subacute Remote | Up | Nnmt | 0 | 0.51075139 | 0.423 | 0.756 | 0 |
| Subacute Remote | Up | Itgax | 2.12E-209 | 0.51023914 | 0.512 | 0.807 | 4.23E-206 |
| Subacute Remote | Up | Maged2 | 3.90E-58 | 0.50911197 | 0.568 | 0.793 | 7.80E-55 |
| Subacute Remote | Up | Ankrd13a | 2.79E-88 | 0.50898588 | 0.548 | 0.754 | 5.57E-85 |
| Subacute Remote | Up | Slc39a6 | 7.84E-254 | 0.50758024 | 0.41 | 0.662 | 1.57E-250 |
| Subacute Remote | Up | Parpbp | 1.21E-170 | 0.50725662 | 0.553 | 0.778 | 2.42E-167 |
| Subacute Remote | Up | Vsir | 4.92E-98 | 0.50703362 | 0.524 | 0.741 | 9.84E-95 |
| Subacute Remote | Up | Iffo1 | 5.52E-206 | 0.50582148 | 0.438 | 0.692 | 1.10E-202 |
| Subacute Remote | Up | Tmem176a | 5.60E-110 | 0.50553244 | 0.602 | 0.924 | 1.12E-106 |

| | | | | | | | |
|---|---|---|---|---|---|---|---|
| Subacute Remote | Up | Pdlim2 | 2.11E-234 | 0.50411411 | 0.46 | 0.736 | 4.22E-231 |
| Subacute Remote | Up | Lasp1 | 1.40E-219 | 0.50398165 | 0.426 | 0.765 | 2.79E-216 |
| Subacute Remote | Up | Igfbp4 | 4.92E-230 | 0.50247396 | 0.386 | 0.744 | 9.83E-227 |
| Subacute Remote | Up | Nrg1 | 5.99E-246 | 0.50241346 | 0.435 | 0.721 | 1.20E-242 |
| Subacute Remote | Up | Tpst1 | 4.56E-285 | 0.50155595 | 0.478 | 0.852 | 9.11E-282 |
| Subacute Remote | Up | Tmeff2 | 0 | 0.49944344 | 0.385 | 0.723 | 0 |
| Subacute Remote | Up | Dact2 | 1.13E-223 | 0.49851113 | 0.503 | 0.758 | 2.26E-220 |
| Subacute Remote | Up | Racgap1 | 0 | 0.49839173 | 0.408 | 0.856 | 0 |
| Subacute Remote | Up | Ptpro | 0 | 0.49753471 | 0.43 | 0.861 | 0 |
| Subacute Remote | Up | Nckap5 | 1.76E-241 | 0.49680113 | 0.426 | 0.704 | 3.52E-238 |
| Subacute Remote | Up | Melk | 7.64E-233 | 0.49674845 | 0.478 | 0.781 | 1.53E-229 |
| Subacute Remote | Up | Lama5 | 1.58E-105 | 0.49668768 | 0.489 | 0.697 | 3.15E-102 |
| Subacute Remote | Up | Arhgap45 | 0 | 0.49647885 | 0.378 | 0.676 | 0 |
| Subacute Remote | Up | Ccdc34 | 1.19E-298 | 0.4962108 | 0.394 | 0.676 | 2.38E-295 |

| | | | | | | | |
|---|---|---|---|---|---|---|---|
| Subacute Remote | Up | Fam167a | 0 | 0.49540488 | 0.421 | 0.921 | 0 |
| Subacute Remote | Up | Emp1 | 1.02E-88 | 0.49481041 | 0.488 | 0.913 | 2.03E-85 |
| Subacute Remote | Up | Lgals3 | 9.32E-30 | 0.49421499 | 0.659 | 0.904 | 1.86E-26 |
| Subacute Remote | Up | Smc4 | 8.53E-118 | 0.49346318 | 0.612 | 0.927 | 1.71E-114 |
| Subacute Remote | Up | Spn | 1.82E-281 | 0.4931537 | 0.456 | 0.757 | 3.63E-278 |
| Subacute Remote | Up | Slc15a3 | 1.21E-184 | 0.4924862 | 0.475 | 0.767 | 2.42E-181 |
| Subacute Remote | Up | Apobec1 | 1.97E-93 | 0.49171618 | 0.481 | 0.712 | 3.95E-90 |
| Subacute Remote | Up | Gpr176 | 4.02E-278 | 0.49168276 | 0.412 | 0.651 | 8.04E-275 |
| Subacute Remote | Up | Gpnmb | 6.96E-238 | 0.49121587 | 0.496 | 0.879 | 1.39E-234 |
| Subacute Remote | Up | Cmtm7 | 4.32E-156 | 0.49117971 | 0.556 | 0.885 | 8.63E-153 |
| Subacute Remote | Up | Anxa1 | 1.64E-109 | 0.49047775 | 0.533 | 0.849 | 3.28E-106 |
| Subacute Remote | Up | Cd68 | 5.97E-276 | 0.48959263 | 0.462 | 0.864 | 1.19E-272 |
| Subacute Remote | Up | Igfbp3 | 2.31E-302 | 0.48942082 | 0.358 | 0.788 | 4.61E-299 |
| Subacute Remote | Up | Gm2a | 1.81E-113 | 0.48933825 | 0.589 | 0.917 | 3.63E-110 |

| | | | | | | | |
|---|---|---|---|---|---|---|---|
| Subacute Remote | Up | Prph | 7.20E-200 | 0.48915518 | 0.45 | 0.706 | 1.44E-196 |
| Subacute Remote | Up | Fam91a1 | 1.22E-261 | 0.48907206 | 0.417 | 0.719 | 2.43E-258 |
| Subacute Remote | Up | AABR070721 | 5.05E-107 | 0.48891335 | 0.537 | 0.764 | 1.01E-103 |
| Subacute Remote | Up | Ltbr | 9.29E-104 | 0.48869833 | 0.537 | 0.765 | 1.86E-100 |
| Subacute Remote | Up | Tmem119 | 8.85E-273 | 0.48857783 | 0.469 | 0.878 | 1.77E-269 |
| Subacute Remote | Up | Lrrcc1 | 2.13E-153 | 0.48854263 | 0.524 | 0.812 | 4.27E-150 |
| Subacute Remote | Up | Serpini1 | 1.83E-153 | 0.48821438 | 0.531 | 0.868 | 3.66E-150 |
| Subacute Remote | Up | Pld4 | 7.71E-289 | 0.48783448 | 0.448 | 0.815 | 1.54E-285 |
| Subacute Remote | Up | Spp1 | 3.55E-271 | 0.48725549 | 0.334 | 0.578 | 7.10E-268 |
| Subacute Remote | Up | Dpysl3 | 9.08E-267 | 0.48713183 | 0.492 | 0.826 | 1.82E-263 |
| Subacute Remote | Up | Acvrl1 | 1.66E-305 | 0.48682172 | 0.326 | 0.621 | 3.32E-302 |
| Subacute Remote | Up | Aard | 6.23E-129 | 0.48509494 | 0.562 | 0.791 | 1.25E-125 |
| Subacute Remote | Up | Plat | 5.91E-154 | 0.48490886 | 0.436 | 0.738 | 1.18E-150 |
| Subacute Remote | Up | Cyp1b1 | 1.10E-266 | 0.48423725 | 0.448 | 0.823 | 2.20E-263 |

| | | | | | | | |
|---|---|---|---|---|---|---|---|
| Subacute Remote | Up | Islr | 2.89E-59 | 0.48313253 | 0.597 | 0.955 | 5.78E-56 |
| Subacute Remote | Up | Tmem176b | 2.88E-157 | 0.48289128 | 0.508 | 0.852 | 5.76E-154 |
| Subacute Remote | Up | C6 | 4.90E-266 | 0.48281794 | 0.5 | 0.847 | 9.80E-263 |
| Subacute Remote | Up | Efhd2 | 1.02E-168 | 0.48262788 | 0.452 | 0.675 | 2.04E-165 |
| Subacute Remote | Up | Fcgr1a | 3.10E-206 | 0.48092031 | 0.532 | 0.885 | 6.19E-203 |
| Subacute Remote | Up | Inpp5d | 0 | 0.48050264 | 0.421 | 0.739 | 0 |
| Subacute Remote | Up | Dbn1 | 3.28E-273 | 0.48040538 | 0.474 | 0.87 | 6.57E-270 |
| Subacute Remote | Up | Ergic3 | 1.00E-71 | 0.4794819 | 0.609 | 0.866 | 2.00E-68 |
| Subacute Remote | Up | Ccl4 | 2.11E-175 | 0.479455 | 0.484 | 0.819 | 4.22E-172 |
| Subacute Remote | Up | Ints6 | 4.54E-153 | 0.47902038 | 0.473 | 0.721 | 9.09E-150 |
| Subacute Remote | Up | Olfml3 | 9.46E-192 | 0.47864019 | 0.509 | 0.864 | 1.89E-188 |
| Subacute Remote | Up | Hoxb4 | 3.39E-307 | 0.47815152 | 0.452 | 0.773 | 6.77E-304 |
| Subacute Remote | Up | Spag5.1 | 3.82E-135 | 0.47773716 | 0.459 | 0.722 | 7.64E-132 |
| Subacute Remote | Up | C1qtnf5 | 6.86E-283 | 0.47760304 | 0.524 | 0.934 | 1.37E-279 |

| | | | | | | | |
|---|---|---|---|---|---|---|---|
| Subacute Remote | Up | Tgfb1 | 0 | 0.47743041 | 0.242 | 0.665 | 0 |
| Subacute Remote | Up | Sat1 | 1.02E-215 | 0.47676122 | 0.434 | 0.736 | 2.04E-212 |
| Subacute Remote | Up | Heph | 0 | 0.47665547 | 0.336 | 0.587 | 0 |
| Subacute Remote | Up | Laptm5 | 0 | 0.47621717 | 0.447 | 0.874 | 0 |
| Subacute Remote | Up | Myo9b | 3.47E-123 | 0.47608303 | 0.537 | 0.811 | 6.94E-120 |
| Subacute Remote | Up | Pid1 | 4.22E-158 | 0.47582907 | 0.533 | 0.863 | 8.43E-155 |
| Subacute Remote | Up | Cpxm1 | 3.66E-236 | 0.47543677 | 0.496 | 0.9 | 7.32E-233 |
| Subacute Remote | Up | Ptpn6 | 6.19E-228 | 0.47528467 | 0.479 | 0.831 | 1.24E-224 |
| Subacute Remote | Up | Rasl10a | 3.46E-214 | 0.47477812 | 0.466 | 0.737 | 6.93E-211 |
| Subacute Remote | Up | Slc29a3 | 2.23E-140 | 0.47366599 | 0.488 | 0.755 | 4.46E-137 |
| Subacute Remote | Up | Tnfrsf1a | 6.71E-106 | 0.47361992 | 0.564 | 0.862 | 1.34E-102 |
| Subacute Remote | Up | Limd2 | 3.85E-193 | 0.4735293 | 0.457 | 0.72 | 7.70E-190 |
| Subacute Remote | Up | Aif1 | 6.10E-79 | 0.47201197 | 0.617 | 0.94 | 1.22E-75 |
| Subacute Remote | Up | LOC1009113 | 2.39E-53 | 0.47143828 | 0.52 | 0.857 | 4.78E-50 |

| | | | | | | | |
|---|---|---|---|---|---|---|---|
| Subacute Remote | Up | Tnnt3 | 0 | 0.4705051 | 0.337 | 0.812 | 0 |
| Subacute Remote | Up | Spc25 | 0 | 0.47014973 | 0.409 | 0.836 | 0 |
| Subacute Remote | Up | Dpep1 | 1.41E-275 | 0.4694413 | 0.393 | 0.636 | 2.82E-272 |
| Subacute Remote | Up | Cd53 | 0 | 0.4691832 | 0.449 | 0.85 | 0 |
| Subacute Remote | Up | LOC312273 | 9.30E-30 | 0.46898133 | 0.42 | 0.154 | 1.86E-26 |
| Subacute Remote | Up | Tspan17 | 3.65E-233 | 0.4679507 | 0.505 | 0.849 | 7.31E-230 |
| Subacute Remote | Up | Adamts15 | 0 | 0.46780303 | 0.384 | 0.777 | 0 |
| Subacute Remote | Up | Fkbp10 | 2.91E-143 | 0.46761293 | 0.508 | 0.805 | 5.81E-140 |
| Subacute Remote | Up | Hmgcs2 | 2.26E-246 | 0.46704764 | 0.484 | 0.831 | 4.52E-243 |
| Subacute Remote | Up | Sulf1 | 3.00E-229 | 0.46663761 | 0.467 | 0.827 | 6.01E-226 |
| Subacute Remote | Up | Pycard | 2.78E-127 | 0.46485843 | 0.533 | 0.807 | 5.57E-124 |
| Subacute Remote | Up | Atp10a | 4.36E-267 | 0.46431456 | 0.416 | 0.681 | 8.71E-264 |
| Subacute Remote | Up | Matk | 2.50E-184 | 0.46390245 | 0.486 | 0.81 | 4.99E-181 |
| Subacute Remote | Up | LOC1083480 | 0 | 0.46299408 | 0.42 | 0.797 | 0 |

| | | | | | | | |
|---|---|---|---|---|---|---|---|
| Subacute Remote | Up | Lst1 | 0 | 0.46228323 | 0.358 | 0.657 | 0 |
| Subacute Remote | Up | Fxyd5 | 9.97E-103 | 0.46149313 | 0.599 | 0.948 | 1.99E-99 |
| Subacute Remote | Up | Vcam1 | 3.15E-225 | 0.46089678 | 0.402 | 0.637 | 6.31E-222 |
| Subacute Remote | Up | Sirpa | 4.59E-92 | 0.46081212 | 0.552 | 0.783 | 9.18E-89 |
| Subacute Remote | Up | LOC685048 | 5.75E-214 | 0.46078091 | 0.423 | 0.666 | 1.15E-210 |
| Subacute Remote | Up | Arpc5 | 1.15E-74 | 0.46074114 | 0.59 | 0.835 | 2.31E-71 |
| Subacute Remote | Up | Slc25a24 | 0 | 0.4606041 | 0.37 | 0.87 | 0 |
| Subacute Remote | Up | Igfbp6 | 1.43E-37 | 0.46035311 | 0.623 | 0.87 | 2.87E-34 |
| Subacute Remote | Up | Rhoj | 1.39E-164 | 0.45993372 | 0.449 | 0.662 | 2.77E-161 |
| Subacute Remote | Up | Mxra8 | 5.00E-112 | 0.459455 | 0.54 | 0.839 | 1.00E-108 |
| Subacute Remote | Up | Ier3 | 1.93E-103 | 0.45917776 | 0.539 | 0.776 | 3.87E-100 |
| Subacute Remote | Up | Chrne | 7.42E-166 | 0.45913312 | 0.502 | 0.824 | 1.48E-162 |
| Subacute Remote | Up | Plek | 3.28E-239 | 0.45846072 | 0.483 | 0.733 | 6.57E-236 |
| Subacute Remote | Up | AC128848.1 | 1.72E-158 | 0.45843879 | 0.541 | 0.765 | 3.45E-155 |

| | | | | | | | |
|---|---|---|---|---|---|---|---|
| Subacute Remote | Up | Orai2 | 6.36E-260 | 0.45732924 | 0.455 | 0.757 | 1.27E-256 |
| Subacute Remote | Up | Loxl2 | 1.67E-154 | 0.45706363 | 0.544 | 0.918 | 3.34E-151 |
| Subacute Remote | Up | Arap1 | 0 | 0.45645843 | 0.363 | 0.788 | 0 |
| Subacute Remote | Up | RGD1309362 | 1.31E-109 | 0.45636314 | 0.625 | 0.91 | 2.63E-106 |
| Subacute Remote | Up | Arhgef40 | 1.55E-200 | 0.45599953 | 0.458 | 0.778 | 3.10E-197 |
| Subacute Remote | Up | Ephb3 | 0 | 0.45517854 | 0.448 | 0.806 | 0 |
| Subacute Remote | Up | Slc66a3 | 0 | 0.45516591 | 0.401 | 0.757 | 0 |
| Subacute Remote | Up | Myo1d | 5.67E-199 | 0.45488455 | 0.444 | 0.693 | 1.13E-195 |
| Subacute Remote | Up | Csf1 | 0 | 0.45442535 | 0.448 | 0.808 | 0 |
| Subacute Remote | Up | Uap1 | 8.22E-308 | 0.45319824 | 0.45 | 0.93 | 1.64E-304 |
| Subacute Remote | Up | Zfp385a | 1.39E-244 | 0.45272922 | 0.465 | 0.778 | 2.78E-241 |
| Subacute Remote | Up | Cilp | 1.74E-276 | 0.45226038 | 0.483 | 0.91 | 3.47E-273 |
| Subacute Remote | Up | LOC1003602 | 5.90E-116 | 0.45212938 | 0.487 | 0.706 | 1.18E-112 |
| Subacute Remote | Up | Ext1 | 1.57E-99 | 0.45199377 | 0.579 | 0.852 | 3.13E-96 |

| | | | | | | | |
|---|---|---|---|---|---|---|---|
| Subacute Remote | Up | AABR070715 | 4.20E-275 | 0.45183616 | 0.146 | 0.467 | 8.40E-272 |
| Subacute Remote | Up | Dlgap4 | 1.78E-300 | 0.45162056 | 0.423 | 0.81 | 3.55E-297 |
| Subacute Remote | Up | Timp1 | 1.08E-111 | 0.45100562 | 0.545 | 0.844 | 2.15E-108 |
| Subacute Remote | Up | Thbs2 | 8.21E-118 | 0.45055326 | 0.555 | 0.857 | 1.64E-114 |
| Subacute Remote | Up | AABR070568 | 0 | 0.44998349 | 0.472 | 0.839 | 0 |
| Subacute Remote | Up | Gfpt2 | 3.96E-255 | 0.44942024 | 0.469 | 0.777 | 7.93E-252 |
| Subacute Remote | Up | Parva | 4.53E-68 | 0.44907628 | 0.568 | 0.79 | 9.07E-65 |
| Subacute Remote | Up | Htra1 | 5.07E-48 | 0.44793223 | 0.556 | 0.791 | 1.01E-44 |
| Subacute Remote | Up | Bicc1 | 1.61E-147 | 0.44789394 | 0.604 | 0.947 | 3.22E-144 |
| Subacute Remote | Up | Prdx4 | 3.97E-89 | 0.44760424 | 0.605 | 0.874 | 7.93E-86 |
| Subacute Remote | Up | Dpp7 | 7.38E-42 | 0.44733956 | 0.643 | 0.923 | 1.48E-38 |
| Subacute Remote | Up | P4ha2 | 8.00E-174 | 0.44698751 | 0.468 | 0.743 | 1.60E-170 |
| Subacute Remote | Up | Susd6 | 2.01E-124 | 0.44693024 | 0.609 | 0.928 | 4.02E-121 |
| Subacute Remote | Up | Glb1 | 1.13E-155 | 0.44627202 | 0.481 | 0.721 | 2.26E-152 |

| | | | | | | | |
|---|---|---|---|---|---|---|---|
| Subacute Remote | Up | Mrgprf | 4.40E-224 | 0.44587042 | 0.448 | 0.766 | 8.81E-221 |
| Subacute Remote | Up | Colgalt1 | 1.88E-116 | 0.444777 | 0.531 | 0.801 | 3.77E-113 |
| Subacute Remote | Up | Pkd1 | 8.30E-257 | 0.44344215 | 0.435 | 0.853 | 1.66E-253 |
| Subacute Remote | Up | St3gal2 | 6.01E-133 | 0.44216974 | 0.507 | 0.74 | 1.20E-129 |
| Subacute Remote | Up | Hsd17b11 | 1.51E-215 | 0.44212694 | 0.433 | 0.699 | 3.01E-212 |
| Subacute Remote | Up | Olfml1 | 0 | 0.44157463 | 0.428 | 0.823 | 0 |
| Subacute Remote | Up | Pls3 | 2.88E-73 | 0.44094258 | 0.581 | 0.807 | 5.77E-70 |
| Subacute Remote | Up | Mtap | 0 | 0.44011533 | 0.366 | 0.701 | 0 |
| Subacute Remote | Up | Cybb | 0 | 0.43959283 | 0.551 | 0.951 | 0 |
| Subacute Remote | Up | Fcna | 0 | 0.43912262 | 0.403 | 0.77 | 0 |
| Subacute Remote | Up | Fam180a | 4.37E-206 | 0.43851473 | 0.497 | 0.837 | 8.74E-203 |
| Subacute Remote | Up | Ccl19 | 0 | 0.4383864 | 0.374 | 0.694 | 0 |
| Subacute Remote | Up | Ch25h | 3.90E-285 | 0.43829347 | 0.475 | 0.87 | 7.81E-282 |
| Subacute Remote | Up | Kdelr3 | 1.27E-154 | 0.4374643 | 0.517 | 0.821 | 2.55E-151 |

| | | | | | | | |
|---|---|---|---|---|---|---|---|
| Subacute Remote | Up | Mgat2 | 2.90E-100 | 0.43711041 | 0.547 | 0.782 | 5.79E-97 |
| Subacute Remote | Up | P3h1 | 1.05E-158 | 0.43509604 | 0.469 | 0.705 | 2.10E-155 |
| Subacute Remote | Up | Tf | 1.53E-131 | 0.43506582 | 0.561 | 0.906 | 3.05E-128 |
| Subacute Remote | Up | Marcksl1 | 1.04E-137 | 0.43477125 | 0.56 | 0.857 | 2.08E-134 |
| Subacute Remote | Up | Cotl1 | 1.72E-183 | 0.43378945 | 0.516 | 0.876 | 3.44E-180 |
| Subacute Remote | Up | C3 | 3.72E-307 | 0.43370206 | 0.394 | 0.682 | 7.44E-304 |
| Subacute Remote | Up | Prcp | 0 | 0.43297036 | 0.364 | 0.721 | 0 |
| Subacute Remote | Up | Smc2 | 0 | 0.43279706 | 0.329 | 0.667 | 0 |
| Subacute Remote | Up | Ptprj | 2.58E-273 | 0.43229052 | 0.459 | 0.793 | 5.15E-270 |
| Subacute Remote | Up | Serp1 | 5.39E-189 | 0.43183793 | 0.462 | 0.748 | 1.08E-185 |
| Subacute Remote | Up | Snai1 | 0 | 0.43175583 | 0.34 | 0.645 | 0 |
| Subacute Remote | Up | Ccdc186 | 1.22E-214 | 0.43166208 | 0.533 | 0.846 | 2.44E-211 |
| Subacute Remote | Up | Ccl21 | 0 | 0.43135409 | 0.283 | 0.735 | 0 |
| Subacute Remote | Up | Flt3lg | 0 | 0.42984918 | 0.417 | 0.774 | 0 |

| | | | | | | | |
|---|---|---|---|---|---|---|---|
| Subacute Remote | Up | Galnt16 | 0 | 0.42960817 | 0.473 | 0.865 | 0 |
| Subacute Remote | Up | Wfikkn2 | 8.39E-115 | 0.42944843 | 0.511 | 0.757 | 1.68E-111 |
| Subacute Remote | Up | Csf1r | 9.86E-122 | 0.42869005 | 0.635 | 0.948 | 1.97E-118 |
| Subacute Remote | Up | Fgfrl1 | 8.25E-209 | 0.42836344 | 0.47 | 0.727 | 1.65E-205 |
| Subacute Remote | Up | Vat1 | 7.67E-86 | 0.42594249 | 0.577 | 0.854 | 1.53E-82 |
| Subacute Remote | Up | Folr2 | 1.82E-303 | 0.42588551 | 0.49 | 0.899 | 3.64E-300 |
| Subacute Remote | Up | Myd88 | 0 | 0.42588008 | 0.338 | 0.604 | 0 |
| Subacute Remote | Up | Fcmr | 0 | 0.4258039 | 0.376 | 0.815 | 0 |
| Subacute Remote | Up | LOC1009119 | 1.66E-215 | 0.4257285 | 0.444 | 0.884 | 3.31E-212 |
| Subacute Remote | Up | Slamf9 | 2.47E-161 | 0.42363826 | 0.524 | 0.868 | 4.93E-158 |
| Subacute Remote | Up | Kirrel1 | 0 | 0.42279357 | 0.195 | 0.43 | 0 |
| Subacute Remote | Up | Nes | 3.92E-228 | 0.42223249 | 0.439 | 0.731 | 7.84E-225 |
| Subacute Remote | Up | Glrx | 1.62E-286 | 0.42199921 | 0.426 | 0.726 | 3.24E-283 |
| Subacute Remote | Up | Cdkn2c | 0 | 0.42125639 | 0.278 | 0.572 | 0 |

| | | | | | | | |
|---|---|---|---|---|---|---|---|
| Subacute Remote | Up | Gsdmd | 0 | 0.42072794 | 0.358 | 0.648 | 0 |
| Subacute Remote | Up | Smc6 | 4.98E-80 | 0.42026514 | 0.607 | 0.863 | 9.95E-77 |
| Subacute Remote | Up | Stom | 7.24E-152 | 0.42024713 | 0.443 | 0.662 | 1.45E-148 |
| Subacute Remote | Up | Mast3 | 4.90E-295 | 0.41937546 | 0.369 | 0.653 | 9.81E-292 |
| Subacute Remote | Up | Cyba | 1.32E-115 | 0.41859677 | 0.531 | 0.864 | 2.64E-112 |
| Subacute Remote | Up | AC128960.1 | 2.96E-132 | 0.41857304 | 0.437 | 0.711 | 5.92E-129 |
| Subacute Remote | Up | Ino80d | 0 | 0.41781452 | 0.493 | 0.764 | 0 |
| Subacute Remote | Up | Ctsz | 1.13E-264 | 0.41780602 | 0.456 | 0.858 | 2.26E-261 |
| Subacute Remote | Up | Sec22b | 1.52E-215 | 0.41753111 | 0.455 | 0.746 | 3.04E-212 |
| Subacute Remote | Up | Ms4a6a | 0 | 0.41677663 | 0.361 | 0.811 | 0 |
| Subacute Remote | Up | Pdpn | 2.09E-239 | 0.41673661 | 0.452 | 0.787 | 4.18E-236 |
| Subacute Remote | Up | Zfp36l2 | 9.37E-89 | 0.41613102 | 0.547 | 0.814 | 1.87E-85 |
| Subacute Remote | Up | Dact3 | 4.88E-288 | 0.41390528 | 0.404 | 0.736 | 9.76E-285 |
| Subacute Remote | Up | Rbp1 | 1.37E-103 | 0.41367708 | 0.607 | 0.894 | 2.74E-100 |

| | | | | | | | |
|---|---|---|---|---|---|---|---|
| Subacute Remote | Up | Spty2d1 | 1.14E-183 | 0.41345606 | 0.497 | 0.756 | 2.28E-180 |
| Subacute Remote | Up | Ppp1r18 | 6.35E-144 | 0.41296703 | 0.484 | 0.695 | 1.27E-140 |
| Subacute Remote | Up | Srpx | 0 | 0.41290712 | 0.399 | 0.782 | 0 |
| Subacute Remote | Up | Twist1 | 0 | 0.41221313 | 0.376 | 0.649 | 0 |
| Subacute Remote | Up | Igf1 | 2.31E-90 | 0.41213124 | 0.652 | 0.975 | 4.62E-87 |
| Subacute Remote | Up | Enpp1 | 1.46E-197 | 0.41109393 | 0.505 | 0.819 | 2.91E-194 |
| Subacute Remote | Up | Man2b1 | 5.14E-189 | 0.41096328 | 0.473 | 0.797 | 1.03E-185 |
| Subacute Remote | Up | Susd2 | 0 | 0.41090033 | 0.322 | 0.591 | 0 |
| Subacute Remote | Up | Rif1 | 2.85E-222 | 0.41078244 | 0.501 | 0.822 | 5.69E-219 |
| Subacute Remote | Up | Chpf | 1.69E-66 | 0.41067967 | 0.606 | 0.876 | 3.37E-63 |
| Subacute Remote | Up | Klrd1 | 8.35E-195 | 0.41018833 | 0.495 | 0.768 | 1.67E-191 |
| Subacute Remote | Up | Cebpb | 0 | 0.41013059 | 0.381 | 0.744 | 0 |
| Subacute Remote | Up | Slfn4 | 7.82E-273 | 0.41004146 | 0.523 | 0.933 | 1.56E-269 |
| Subacute Remote | Up | Taf7 | 6.39E-260 | 0.40982025 | 0.391 | 0.625 | 1.28E-256 |

| | | | | | | | |
|---|---|---|---|---|---|---|---|
| Subacute Remote | Up | Ttc3 | 2.39E-112 | 0.40911354 | 0.592 | 0.889 | 4.79E-109 |
| Subacute Remote | Up | LOC1083482 | 4.53E-235 | 0.40902963 | 0.442 | 0.732 | 9.06E-232 |
| Subacute Remote | Up | Avp | 3.18E-220 | 0.4085947 | 0.39 | 0.609 | 6.37E-217 |
| Subacute Remote | Up | Ar | 1.68E-174 | 0.40657168 | 0.522 | 0.875 | 3.36E-171 |
| Subacute Remote | Up | Aldh1a2 | 0 | 0.40611274 | 0.439 | 0.8 | 0 |
| Subacute Remote | Up | Pafah1b3 | 0 | 0.40553985 | 0.435 | 0.881 | 0 |
| Subacute Remote | Up | Pmp22 | 1.56E-50 | 0.40544616 | 0.645 | 0.973 | 3.12E-47 |
| Subacute Remote | Up | Adamts12 | 0 | 0.40535511 | 0.364 | 0.721 | 0 |
| Subacute Remote | Up | Palm | 4.51E-229 | 0.40522734 | 0.43 | 0.739 | 9.03E-226 |
| Subacute Remote | Up | Tnc | 0 | 0.40472512 | 0.491 | 0.866 | 0 |
| Subacute Remote | Up | Tap1 | 1.21E-80 | 0.40364054 | 0.618 | 0.883 | 2.42E-77 |
| Subacute Remote | Up | Cnpy3 | 1.66E-274 | 0.40245878 | 0.427 | 0.747 | 3.32E-271 |
| Subacute Remote | Up | Klri1 | 2.95E-178 | 0.40209119 | 0.484 | 0.703 | 5.90E-175 |
| Subacute Remote | Up | Map1b | 2.66E-83 | 0.40195216 | 0.605 | 0.889 | 5.32E-80 |

| | | | | | | | |
|---|---|---|---|---|---|---|---|
| Subacute Remote | Up | Resf1 | 9.11E-251 | 0.40162504 | 0.512 | 0.849 | 1.82E-247 |
| Subacute Remote | Up | Steap3 | 1.80E-176 | 0.40157921 | 0.475 | 0.748 | 3.59E-173 |
| Subacute Remote | Up | PCOLCE2 | 3.58E-170 | 0.40133674 | 0.524 | 0.825 | 7.16E-167 |
| Subacute Remote | Up | AABR070070 | 0 | 0.40018071 | 0.246 | 0.503 | 0 |
| Subacute Remote | Up | Cst6 | 1.47E-214 | 0.40016993 | 0.55 | 0.842 | 2.95E-211 |
| Subacute Remote | Up | F13a1 | 0 | 0.39987784 | 0.481 | 0.819 | 0 |
| Subacute Remote | Up | Wapl | 6.85E-198 | 0.39984622 | 0.505 | 0.785 | 1.37E-194 |
| Subacute Remote | Up | Col4a5 | 0 | 0.39903591 | 0.425 | 0.848 | 0 |
| Subacute Remote | Up | Junb | 5.17E-230 | 0.39883706 | 0.359 | 0.596 | 1.03E-226 |
| Subacute Remote | Up | Fcgrt | 3.20E-18 | 0.39821529 | 0.709 | 0.915 | 6.40E-15 |
| Subacute Remote | Up | Ca3 | 4.03E-225 | 0.39787146 | 0.494 | 0.856 | 8.06E-222 |
| Subacute Remote | Up | Rcn2 | 1.37E-297 | 0.39773887 | 0.352 | 0.634 | 2.74E-294 |
| Subacute Remote | Up | Egr1 | 8.68E-243 | 0.39755685 | 0.412 | 0.664 | 1.74E-239 |
| Subacute Remote | Up | Plxdc2 | 8.45E-274 | 0.3966201 | 0.524 | 0.894 | 1.69E-270 |

| | | | | | | | |
|---|---|---|---|---|---|---|---|
| Subacute Remote | Up | Olfml2a | 0 | 0.39633886 | 0.484 | 0.845 | 0 |
| Subacute Remote | Up | Aoc3 | 1.00E-47 | 0.39620352 | 0.594 | 0.89 | 2.01E-44 |
| Subacute Remote | Up | Tyrobp | 1.98E-55 | 0.39613786 | 0.649 | 0.923 | 3.96E-52 |
| Subacute Remote | Up | Cap1 | 1.41E-121 | 0.39520902 | 0.532 | 0.788 | 2.81E-118 |
| Subacute Remote | Up | Pheta2 | 0 | 0.39520516 | 0.499 | 0.787 | 0 |
| Subacute Remote | Up | Clic1 | 4.05E-126 | 0.39504311 | 0.559 | 0.88 | 8.10E-123 |
| Subacute Remote | Up | Glmp | 2.13E-114 | 0.39323106 | 0.507 | 0.732 | 4.25E-111 |
| Subacute Remote | Up | Cd14 | 0 | 0.39309147 | 0.44 | 0.838 | 0 |
| Subacute Remote | Up | C1qtnf2 | 1.03E-246 | 0.39235343 | 0.515 | 0.908 | 2.05E-243 |
| Subacute Remote | Up | Pdlim7 | 2.07E-214 | 0.39171718 | 0.478 | 0.772 | 4.15E-211 |
| Subacute Remote | Up | Nbl1 | 4.61E-106 | 0.39053086 | 0.575 | 0.867 | 9.21E-103 |
| Subacute Remote | Up | Pld3 | 3.40E-136 | 0.39008382 | 0.559 | 0.865 | 6.81E-133 |
| Subacute Remote | Up | Tcirg1 | 0 | 0.3894294 | 0.409 | 0.738 | 0 |
| Subacute Remote | Up | Tmbim1 | 3.49E-170 | 0.38889138 | 0.46 | 0.686 | 6.99E-167 |

| | | | | | | | |
|---|---|---|---|---|---|---|---|
| Subacute Remote | Up | Gng2 | 3.66E-250 | 0.38883215 | 0.479 | 0.826 | 7.32E-247 |
| Subacute Remote | Up | Oaf | 1.16E-28 | 0.38882668 | 0.689 | 0.927 | 2.33E-25 |
| Subacute Remote | Up | C1qc | 9.20E-91 | 0.38837723 | 0.682 | 0.996 | 1.84E-87 |
| Subacute Remote | Up | Septin8 | 5.12E-247 | 0.38800776 | 0.419 | 0.79 | 1.02E-243 |
| Subacute Remote | Up | Sdc2 | 5.41E-76 | 0.3876976 | 0.588 | 0.804 | 1.08E-72 |
| Subacute Remote | Up | AABR070546 | 5.85E-24 | 0.38756978 | 0.718 | 0.984 | 1.17E-20 |
| Subacute Remote | Up | Chrm2 | 2.96E-288 | 0.38742262 | 0.504 | 0.818 | 5.92E-285 |
| Subacute Remote | Up | Psd4 | 1.62E-243 | 0.38718055 | 0.465 | 0.734 | 3.24E-240 |
| Subacute Remote | Up | Agtr1a | 0 | 0.38678846 | 0.389 | 0.856 | 0 |
| Subacute Remote | Up | Glt8d1 | 5.38E-287 | 0.38508107 | 0.412 | 0.726 | 1.08E-283 |
| Subacute Remote | Up | Chpf2 | 9.23E-110 | 0.38497314 | 0.535 | 0.758 | 1.85E-106 |
| Subacute Remote | Up | Efemp1 | 4.79E-299 | 0.38367873 | 0.379 | 0.718 | 9.58E-296 |
| Subacute Remote | Up | Sult1a1 | 2.22E-223 | 0.38293346 | 0.412 | 0.676 | 4.43E-220 |
| Subacute Remote | Up | Dram1 | 0 | 0.3824821 | 0.389 | 0.747 | 0 |

| | | | | | | | |
|---|---|---|---|---|---|---|---|
| Subacute Remote | Up | Slc7a6 | 0 | 0.38241129 | 0.31 | 0.639 | 0 |
| Subacute Remote | Up | Ganab | 6.44E-177 | 0.38024124 | 0.405 | 0.629 | 1.29E-173 |
| Subacute Remote | Up | Smoc2 | 0 | 0.37951407 | 0.408 | 0.777 | 0 |
| Subacute Remote | Up | Tlr12 | 0 | 0.37949961 | 0.492 | 0.829 | 0 |
| Subacute Remote | Up | Cwf19l2 | 9.09E-173 | 0.37890606 | 0.567 | 0.88 | 1.82E-169 |
| Subacute Remote | Up | Dok3 | 0 | 0.37774219 | 0.424 | 0.856 | 0 |
| Subacute Remote | Up | Gna15 | 0 | 0.37739596 | 0.375 | 0.831 | 0 |
| Subacute Remote | Up | F2r | 0 | 0.37618799 | 0.32 | 0.702 | 0 |
| Subacute Remote | Up | Rbm3 | 5.25E-259 | 0.37593846 | 0.406 | 0.796 | 1.05E-255 |
| Subacute Remote | Up | Ccn2 | 9.01E-110 | 0.37584152 | 0.468 | 0.852 | 1.80E-106 |
| Subacute Remote | Up | Kpna3 | 0 | 0.37445937 | 0.421 | 0.719 | 0 |
| Subacute Remote | Up | Tnfsf13 | 3.11E-236 | 0.37409002 | 0.435 | 0.714 | 6.21E-233 |
| Subacute Remote | Up | Marveld1 | 7.56E-134 | 0.37402284 | 0.535 | 0.823 | 1.51E-130 |
| Subacute Remote | Up | Ppp1r9b | 1.08E-202 | 0.37327726 | 0.489 | 0.778 | 2.15E-199 |

| | | | | | | | |
|---|---|---|---|---|---|---|---|
| Subacute Remote | Up | Cyth4 | 0 | 0.37259822 | 0.354 | 0.792 | 0 |
| Subacute Remote | Up | Mrvi1 | 0 | 0.37202974 | 0.284 | 0.523 | 0 |
| Subacute Remote | Up | AC111804.2 | 2.80E-271 | 0.37163123 | 0.421 | 0.692 | 5.61E-268 |
| Subacute Remote | Up | Ankrd12 | 1.43E-146 | 0.37146633 | 0.595 | 0.933 | 2.87E-143 |
| Subacute Remote | Up | Necap2 | 2.23E-213 | 0.37041877 | 0.432 | 0.705 | 4.46E-210 |
| Subacute Remote | Up | Ggcx | 1.35E-198 | 0.3703789 | 0.464 | 0.768 | 2.69E-195 |
| Subacute Remote | Up | Rbm22 | 3.39E-217 | 0.37015139 | 0.446 | 0.711 | 6.79E-214 |
| Subacute Remote | Up | Fam172a | 2.99E-127 | 0.36979266 | 0.538 | 0.785 | 5.98E-124 |
| Subacute Remote | Up | Birc5 | 1.64E-275 | 0.36959251 | 0.482 | 0.882 | 3.27E-272 |
| Subacute Remote | Up | Mvb12a | 1.08E-102 | 0.36934178 | 0.566 | 0.825 | 2.16E-99 |
| Subacute Remote | Up | Nrgn | 0 | 0.36918282 | 0.494 | 0.852 | 0 |
| Subacute Remote | Up | Coro1b | 2.63E-228 | 0.36897964 | 0.43 | 0.77 | 5.27E-225 |
| Subacute Remote | Up | Alox5 | 0 | 0.368028 | 0.464 | 0.798 | 0 |
| Subacute Remote | Up | Adamts10 | 0 | 0.36791073 | 0.375 | 0.823 | 0 |

| | | | | | | | |
|---|---|---|---|---|---|---|---|
| Subacute Remote | Up | Cep290 | 7.89E-230 | 0.36765202 | 0.472 | 0.779 | 1.58E-226 |
| Subacute Remote | Up | Cxcl16 | 1.86E-232 | 0.36724091 | 0.464 | 0.784 | 3.71E-229 |
| Subacute Remote | Up | Bin1 | 5.10E-239 | 0.36681051 | 0.462 | 0.781 | 1.02E-235 |
| Subacute Remote | Up | Stt3a | 1.74E-232 | 0.36677794 | 0.447 | 0.823 | 3.49E-229 |
| Subacute Remote | Up | C1qtnf1 | 3.86E-192 | 0.36675828 | 0.558 | 0.958 | 7.72E-189 |
| Subacute Remote | Up | Aph1b | 9.65E-88 | 0.36562335 | 0.56 | 0.845 | 1.93E-84 |
| Subacute Remote | Up | Sh3rf3 | 3.68E-171 | 0.36555182 | 0.395 | 0.632 | 7.35E-168 |
| Subacute Remote | Up | Gbp2 | 9.45E-236 | 0.36528085 | 0.496 | 0.838 | 1.89E-232 |
| Subacute Remote | Up | Apbb1ip | 0 | 0.36522552 | 0.349 | 0.615 | 0 |
| Subacute Remote | Up | Msn | 2.62E-84 | 0.3639581 | 0.57 | 0.906 | 5.23E-81 |
| Subacute Remote | Up | Snx18 | 0 | 0.36383892 | 0.417 | 0.793 | 0 |
| Subacute Remote | Up | Pard3b | 3.86E-157 | 0.36366578 | 0.502 | 0.759 | 7.71E-154 |
| Subacute Remote | Up | Rab32 | 0 | 0.36363481 | 0.337 | 0.668 | 0 |
| Subacute Remote | Up | Fstl1 | 1.53E-26 | 0.36360302 | 0.737 | 0.996 | 3.06E-23 |

| | | | | | | | |
|---|---|---|---|---|---|---|---|
| Subacute Remote | Up | Cfp | 0 | 0.36262693 | 0.312 | 0.645 | 0 |
| Subacute Remote | Up | Emp3 | 2.20E-56 | 0.36244256 | 0.671 | 0.947 | 4.40E-53 |
| Subacute Remote | Up | Mcam | 0 | 0.36219097 | 0.365 | 0.806 | 0 |
| Subacute Remote | Up | Siglec1 | 1.46E-256 | 0.36141119 | 0.56 | 0.972 | 2.93E-253 |
| Subacute Remote | Up | Ramp1 | 1.12E-299 | 0.36094422 | 0.413 | 0.704 | 2.23E-296 |
| Subacute Remote | Up | Clec3b | 4.60E-156 | 0.36049532 | 0.487 | 0.74 | 9.19E-153 |
| Subacute Remote | Up | Tgfb1i1 | 1.21E-258 | 0.36034578 | 0.462 | 0.819 | 2.43E-255 |
| Subacute Remote | Up | LOC1009109 | 4.36E-280 | 0.35943274 | 0.467 | 0.821 | 8.72E-277 |
| Subacute Remote | Up | Adam15 | 4.24E-135 | 0.35891056 | 0.572 | 0.892 | 8.47E-132 |
| Subacute Remote | Up | Ptprcap | 5.98E-225 | 0.35852693 | 0.504 | 0.787 | 1.20E-221 |
| Subacute Remote | Up | Pdgfa | 1.13E-70 | 0.35824477 | 0.62 | 0.869 | 2.25E-67 |
| Subacute Remote | Up | Rhbdf1 | 7.38E-278 | 0.35773955 | 0.43 | 0.777 | 1.48E-274 |
| Subacute Remote | Up | Arhgef25 | 6.10E-252 | 0.35730099 | 0.426 | 0.746 | 1.22E-248 |
| Subacute Remote | Up | Pi16 | 1.76E-90 | 0.35711723 | 0.477 | 0.837 | 3.53E-87 |

| | | | | | | | |
|---|---|---|---|---|---|---|---|
| Subacute Remote | Up | Ywhaz | 4.17E-75 | 0.35651543 | 0.578 | 0.816 | 8.35E-72 |
| Subacute Remote | Up | Bace2 | 1.24E-190 | 0.35633066 | 0.518 | 0.845 | 2.47E-187 |
| Subacute Remote | Up | Oasl2 | 3.18E-142 | 0.35609242 | 0.663 | 0.992 | 6.35E-139 |
| Subacute Remote | Up | Sox9 | 4.14E-182 | 0.3554045 | 0.465 | 0.714 | 8.27E-179 |
| Subacute Remote | Up | Fgl2 | 1.80E-209 | 0.35350305 | 0.516 | 0.873 | 3.60E-206 |
| Subacute Remote | Up | Myof | 0 | 0.35280241 | 0.382 | 0.856 | 0 |
| Subacute Remote | Up | Nucb2 | 1.75E-72 | 0.35253287 | 0.611 | 0.853 | 3.49E-69 |
| Subacute Remote | Up | Osbpl5 | 6.47E-272 | 0.35248206 | 0.441 | 0.801 | 1.29E-268 |
| Subacute Remote | Up | Ngfr | 7.87E-100 | 0.35185568 | 0.524 | 0.736 | 1.57E-96 |
| Subacute Remote | Up | Oas1a | 3.02E-74 | 0.34998121 | 0.645 | 0.964 | 6.04E-71 |
| Subacute Remote | Up | AABR070605 | 1.13E-288 | 0.34943363 | 0.485 | 0.772 | 2.26E-285 |
| Subacute Remote | Up | Erp29 | 5.21E-138 | 0.3491824 | 0.505 | 0.879 | 1.04E-134 |
| Subacute Remote | Up | Rgs10 | 9.51E-184 | 0.34916219 | 0.444 | 0.688 | 1.90E-180 |
| Subacute Remote | Up | Gprc5a | 3.68E-286 | 0.3490439 | 0.441 | 0.785 | 7.36E-283 |

| | | | | | | | |
|---|---|---|---|---|---|---|---|
| Subacute Remote | Up | Cald1 | 6.70E-98 | 0.34897641 | 0.523 | 0.774 | 1.34E-94 |
| Subacute Remote | Up | Zfp951 | 1.47E-187 | 0.34853667 | 0.483 | 0.744 | 2.94E-184 |
| Subacute Remote | Up | Anxa2 | 2.24E-24 | 0.34853469 | 0.742 | 0.978 | 4.47E-21 |
| Subacute Remote | Up | Col12a1 | 0 | 0.34835321 | 0.256 | 0.657 | 0 |
| Subacute Remote | Up | Pigt | 8.60E-134 | 0.34769368 | 0.502 | 0.771 | 1.72E-130 |
| Subacute Remote | Up | Os9 | 4.30E-150 | 0.34702338 | 0.521 | 0.816 | 8.61E-147 |
| Subacute Remote | Up | Zmat1 | 1.27E-231 | 0.34678714 | 0.512 | 0.819 | 2.55E-228 |
| Subacute Remote | Up | Axl | 1.17E-80 | 0.34677456 | 0.6 | 0.939 | 2.34E-77 |
| Subacute Remote | Up | Slc35e4 | 1.40E-164 | 0.34618156 | 0.439 | 0.647 | 2.79E-161 |
| Subacute Remote | Up | Serpinb8 | 9.15E-100 | 0.34572708 | 0.486 | 0.691 | 1.83E-96 |
| Subacute Remote | Up | Xpnpep1 | 5.84E-62 | 0.34536242 | 0.626 | 0.883 | 1.17E-58 |
| Subacute Remote | Up | Fkbp11 | 1.93E-207 | 0.34526622 | 0.511 | 0.813 | 3.85E-204 |
| Subacute Remote | Up | Ano6 | 0 | 0.34460658 | 0.345 | 0.718 | 0 |
| Subacute Remote | Up | Serpinb1a | 7.81E-170 | 0.34382014 | 0.527 | 0.815 | 1.56E-166 |

| | | | | | | | |
|---|---|---|---|---|---|---|---|
| Subacute Remote | Up | Rarres2 | 2.82E-45 | 0.34217396 | 0.591 | 0.808 | 5.63E-42 |
| Subacute Remote | Up | Prkcd | 6.96E-201 | 0.34214823 | 0.493 | 0.782 | 1.39E-197 |
| Subacute Remote | Up | Rac2 | 5.05E-211 | 0.34171424 | 0.461 | 0.747 | 1.01E-207 |
| Subacute Remote | Up | Pdcd4 | 5.00E-59 | 0.34073866 | 0.618 | 0.837 | 1.00E-55 |
| Subacute Remote | Up | Slc39a13 | 1.46E-175 | 0.34017297 | 0.492 | 0.751 | 2.92E-172 |
| Subacute Remote | Up | Aebp1 | 2.89E-15 | 0.34008059 | 0.694 | 0.911 | 5.78E-12 |
| Subacute Remote | Up | Sod3 | 3.18E-83 | 0.33946987 | 0.546 | 0.834 | 6.36E-80 |
| Subacute Remote | Up | Mybpc2 | 1.74E-189 | 0.33933631 | 0.427 | 0.653 | 3.47E-186 |
| Subacute Remote | Up | RGD1311744 | 4.56E-274 | 0.33912166 | 0.392 | 0.64 | 9.12E-271 |
| Subacute Remote | Up | Mt2A | 2.06E-116 | 0.33854541 | 0.619 | 0.968 | 4.12E-113 |
| Subacute Remote | Up | Asgr2 | 7.43E-164 | 0.33712991 | 0.501 | 0.803 | 1.49E-160 |
| Subacute Remote | Up | Il17b | 9.24E-45 | 0.33702939 | 0.519 | 0.729 | 1.85E-41 |
| Subacute Remote | Up | LOC1025554 | 6.47E-159 | 0.33683433 | 0.459 | 0.79 | 1.29E-155 |
| Subacute Remote | Up | Adamtsl3 | 0 | 0.33613854 | 0.255 | 0.643 | 0 |

| | | | | | | | |
|---|---|---|---|---|---|---|---|
| Subacute Remote | Up | Naglu | 1.19E-216 | 0.33612383 | 0.443 | 0.696 | 2.38E-213 |
| Subacute Remote | Up | Tinagl1 | 0 | 0.33607203 | 0.294 | 0.614 | 0 |
| Subacute Remote | Up | Yipf5 | 4.88E-132 | 0.33549803 | 0.566 | 0.792 | 9.76E-129 |
| Subacute Remote | Up | Nrxn1 | 1.10E-238 | 0.3347134 | 0.455 | 0.771 | 2.20E-235 |
| Subacute Remote | Up | Gucy1b1 | 6.80E-251 | 0.33463159 | 0.408 | 0.669 | 1.36E-247 |
| Subacute Remote | Up | Rab31 | 2.58E-229 | 0.3345389 | 0.511 | 0.867 | 5.17E-226 |
| Subacute Remote | Up | Cd37 | 4.24E-125 | 0.33342062 | 0.504 | 0.736 | 8.49E-122 |
| Subacute Remote | Up | Id3 | 3.70E-48 | 0.33328493 | 0.58 | 0.816 | 7.40E-45 |
| Subacute Remote | Up | Txndc5 | 2.63E-94 | 0.33322948 | 0.577 | 0.885 | 5.26E-91 |
| Subacute Remote | Up | Shtn1 | 9.93E-82 | 0.33274561 | 0.54 | 0.789 | 1.99E-78 |
| Subacute Remote | Up | Hcls1 | 0 | 0.33271617 | 0.462 | 0.848 | 0 |
| Subacute Remote | Up | Tkt | 3.63E-110 | 0.33162285 | 0.515 | 0.732 | 7.26E-107 |
| Subacute Remote | Up | Kazald1 | 5.68E-216 | 0.33136276 | 0.463 | 0.759 | 1.14E-212 |
| Subacute Remote | Up | Lyl1 | 0 | 0.33112777 | 0.353 | 0.64 | 0 |

| | | | | | | | |
|---|---|---|---|---|---|---|---|
| Subacute Remote | Up | Mylk | 4.49E-164 | 0.33019485 | 0.436 | 0.678 | 8.97E-161 |
| Subacute Remote | Up | Heyl | 2.28E-84 | 0.32938329 | 0.57 | 0.818 | 4.57E-81 |
| Subacute Remote | Up | Ccdc18 | 1.11E-216 | 0.32934363 | 0.492 | 0.784 | 2.21E-213 |
| Subacute Remote | Up | Gnb1 | 6.95E-207 | 0.32913863 | 0.439 | 0.692 | 1.39E-203 |
| Subacute Remote | Up | Stat6 | 6.18E-165 | 0.32884348 | 0.515 | 0.823 | 1.24E-161 |
| Subacute Remote | Up | Tcn2 | 5.97E-135 | 0.32852124 | 0.51 | 0.765 | 1.19E-131 |
| Subacute Remote | Up | Naaa | 3.38E-265 | 0.32816363 | 0.448 | 0.766 | 6.76E-262 |
| Subacute Remote | Up | Ptbp1 | 2.91E-128 | 0.32779164 | 0.51 | 0.787 | 5.81E-125 |
| Subacute Remote | Up | Klra2 | 1.86E-212 | 0.32757276 | 0.495 | 0.782 | 3.72E-209 |
| Subacute Remote | Up | Gpx8 | 6.49E-237 | 0.3269443 | 0.519 | 0.885 | 1.30E-233 |
| Subacute Remote | Up | Npdc1 | 4.70E-155 | 0.32648397 | 0.504 | 0.759 | 9.41E-152 |
| Subacute Remote | Up | Rab8b | 4.57E-235 | 0.32585626 | 0.422 | 0.702 | 9.13E-232 |
| Subacute Remote | Up | Prpf40a | 1.15E-89 | 0.32558974 | 0.618 | 0.89 | 2.31E-86 |
| Subacute Remote | Up | Pltp | 2.99E-41 | 0.32537059 | 0.702 | 0.959 | 5.98E-38 |

| | | | | | | | |
|---|---|---|---|---|---|---|---|
| Subacute Remote | Up | C1qa | 1.01E-86 | 0.32506825 | 0.673 | 0.987 | 2.01E-83 |
| Subacute Remote | Up | Prf1 | 7.39E-287 | 0.32496287 | 0.49 | 0.766 | 1.48E-283 |
| Subacute Remote | Up | Prelp | 5.63E-66 | 0.32487746 | 0.608 | 0.928 | 1.13E-62 |
| Subacute Remote | Up | Pdgfrb | 1.46E-103 | 0.32417661 | 0.562 | 0.886 | 2.92E-100 |
| Subacute Remote | Up | Ift20 | 1.26E-119 | 0.32394457 | 0.559 | 0.828 | 2.52E-116 |
| Subacute Remote | Up | Rnf19a | 2.01E-120 | 0.32319526 | 0.493 | 0.695 | 4.01E-117 |
| Subacute Remote | Up | Camk4 | 1.25E-304 | 0.32297726 | 0.398 | 0.695 | 2.49E-301 |
| Subacute Remote | Up | Tlr7 | 3.47E-251 | 0.32292853 | 0.546 | 0.95 | 6.93E-248 |
| Subacute Remote | Up | Pou2f2 | 7.42E-225 | 0.32281211 | 0.555 | 0.91 | 1.48E-221 |
| Subacute Remote | Up | Il33 | 2.11E-163 | 0.32177924 | 0.473 | 0.702 | 4.23E-160 |
| Subacute Remote | Up | Cradd | 2.67E-133 | 0.32134471 | 0.505 | 0.843 | 5.34E-130 |
| Subacute Remote | Up | Efemp2 | 1.69E-219 | 0.32098183 | 0.446 | 0.787 | 3.38E-216 |
| Subacute Remote | Up | Gucy1a1 | 3.75E-265 | 0.32012637 | 0.355 | 0.597 | 7.51E-262 |
| Subacute Remote | Up | Aida | 2.45E-80 | 0.31877565 | 0.597 | 0.871 | 4.91E-77 |

| | | | | | | | |
|---|---|---|---|---|---|---|---|
| Subacute Remote | Up | Tgfbi | 7.35E-14 | 0.31869898 | 0.658 | 0.95 | 1.47E-10 |
| Subacute Remote | Up | Scarf2 | 5.22E-196 | 0.31861594 | 0.533 | 0.916 | 1.04E-192 |
| Subacute Remote | Up | Ltbp3 | 1.15E-128 | 0.31851868 | 0.542 | 0.962 | 2.31E-125 |
| Subacute Remote | Up | Tbx1 | 1.22E-156 | 0.31810785 | 0.515 | 0.735 | 2.45E-153 |
| Subacute Remote | Up | Selenof | 3.10E-30 | 0.31800042 | 0.684 | 0.892 | 6.20E-27 |
| Subacute Remote | Up | S100a10 | 1.05E-29 | 0.31775247 | 0.713 | 0.982 | 2.11E-26 |
| Subacute Remote | Up | Rims2 | 2.19E-151 | 0.31749941 | 0.526 | 0.814 | 4.38E-148 |
| Subacute Remote | Up | St3gal4 | 3.97E-148 | 0.31709705 | 0.583 | 0.883 | 7.95E-145 |
| Subacute Remote | Up | Dusp6 | 5.31E-171 | 0.31633652 | 0.547 | 0.817 | 1.06E-167 |
| Subacute Remote | Up | Ythdc2 | 0 | 0.3160735 | 0.487 | 0.805 | 0 |
| Subacute Remote | Up | Olr1 | 7.09E-46 | 0.3160327 | 0.53 | 0.795 | 1.42E-42 |
| Subacute Remote | Up | Ifngr1 | 1.08E-58 | 0.31579094 | 0.608 | 0.869 | 2.16E-55 |
| Subacute Remote | Up | Ackr3 | 8.82E-144 | 0.31537711 | 0.447 | 0.84 | 1.76E-140 |
| Subacute Remote | Up | Map7d3 | 1.22E-214 | 0.31438065 | 0.461 | 0.712 | 2.45E-211 |

| | | | | | | | |
|---|---|---|---|---|---|---|---|
| Subacute Remote | Up | Mfap5 | 9.51E-48 | 0.3138269 | 0.625 | 0.989 | 1.90E-44 |
| Subacute Remote | Up | Mx1 | 1.25E-65 | 0.31364019 | 0.663 | 0.987 | 2.49E-62 |
| Subacute Remote | Up | Samd9 | 9.15E-98 | 0.31361912 | 0.615 | 0.977 | 1.83E-94 |
| Subacute Remote | Up | Nlgn2 | 1.27E-50 | 0.31340845 | 0.512 | 0.74 | 2.54E-47 |
| Subacute Remote | Up | Hps1 | 1.06E-262 | 0.31218987 | 0.453 | 0.787 | 2.13E-259 |
| Subacute Remote | Up | Shisa5 | 7.27E-120 | 0.31189103 | 0.633 | 0.961 | 1.45E-116 |
| Subacute Remote | Up | Tubb6 | 0 | 0.31178051 | 0.391 | 0.779 | 0 |
| Subacute Remote | Up | Fam20c | 3.00E-106 | 0.31093134 | 0.545 | 0.767 | 6.00E-103 |
| Subacute Remote | Up | Pmepa1 | 1.27E-30 | 0.31051412 | 0.692 | 0.943 | 2.53E-27 |
| Subacute Remote | Up | Lpin1 | 0 | 0.31038981 | 0.266 | 0.525 | 0 |
| Subacute Remote | Up | Siva1 | 4.03E-122 | 0.31008099 | 0.524 | 0.751 | 8.05E-119 |
| Subacute Remote | Up | Eif4e2 | 4.18E-201 | 0.30976385 | 0.451 | 0.717 | 8.35E-198 |
| Subacute Remote | Up | Fbln2 | 3.90E-121 | 0.30927336 | 0.622 | 0.939 | 7.80E-118 |
| Subacute Remote | Up | Sec61a1 | 1.78E-99 | 0.30902956 | 0.592 | 0.877 | 3.55E-96 |

| | | | | | | | |
|---|---|---|---|---|---|---|---|
| Subacute Remote | Up | Ext2 | 4.06E-240 | 0.3089593 | 0.397 | 0.72 | 8.13E-237 |
| Subacute Remote | Up | Scap | 1.86E-97 | 0.30751748 | 0.544 | 0.751 | 3.71E-94 |
| Subacute Remote | Up | Il1b | 1.17E-83 | 0.30656846 | 0.505 | 0.753 | 2.34E-80 |
| Subacute Remote | Up | Bod1l1 | 1.23E-200 | 0.30524729 | 0.611 | 0.966 | 2.45E-197 |
| Subacute Remote | Up | Slc39a1 | 6.15E-129 | 0.30455472 | 0.538 | 0.79 | 1.23E-125 |
| Subacute Remote | Up | Tmem100 | 0 | 0.30418404 | 0.37 | 0.882 | 0 |
| Subacute Remote | Up | Chi3l1 | 0 | 0.30283 | 0.293 | 0.714 | 0 |
| Subacute Remote | Up | Cma1 | 0 | 0.30268979 | 0.441 | 0.78 | 0 |
| Subacute Remote | Up | Lpal2 | 3.64E-218 | 0.30218343 | 0.352 | 0.56 | 7.28E-215 |
| Subacute Remote | Up | Slc2a6 | 6.07E-209 | 0.30205119 | 0.441 | 0.727 | 1.21E-205 |
| Subacute Remote | Up | Sfrp1 | 0 | 0.301877 | 0.299 | 0.558 | 0 |
| Subacute Remote | Up | Baz1a | 2.58E-266 | 0.30110652 | 0.436 | 0.765 | 5.17E-263 |
| Subacute Remote | Up | Csk | 9.17E-136 | 0.30105852 | 0.497 | 0.725 | 1.83E-132 |
| Subacute Remote | Up | Ccl3 | 1.15E-264 | 0.30060267 | 0.481 | 0.759 | 2.29E-261 |

| | | | | | | | |
|---|---|---|---|---|---|---|---|
| Subacute Remote | Up | Aspa | 4.47E-291 | 0.30007041 | 0.371 | 0.588 | 8.95E-288 |
| Subacute Remote | Up | Il17re | 7.63E-84 | 0.29937678 | 0.474 | 0.753 | 1.53E-80 |
| Subacute Remote | Up | Scpep1 | 1.17E-13 | 0.29936399 | 0.775 | 0.985 | 2.35E-10 |
| Subacute Remote | Up | Zdhhc23 | 1.08E-187 | 0.29924246 | 0.472 | 0.777 | 2.16E-184 |
| Subacute Remote | Up | Ifit2 | 1.20E-136 | 0.29922986 | 0.634 | 0.898 | 2.40E-133 |
| Subacute Remote | Up | Anxa4 | 5.82E-91 | 0.29895659 | 0.559 | 0.814 | 1.16E-87 |
| Subacute Remote | Up | Top1 | 7.30E-156 | 0.29678442 | 0.55 | 0.896 | 1.46E-152 |
| Subacute Remote | Up | Mdk | 1.19E-208 | 0.29558825 | 0.468 | 0.762 | 2.39E-205 |
| Subacute Remote | Up | Arpc3 | 3.47E-99 | 0.29551273 | 0.577 | 0.826 | 6.94E-96 |
| Subacute Remote | Up | Elk3 | 1.48E-113 | 0.29455597 | 0.593 | 0.879 | 2.96E-110 |
| Subacute Remote | Up | LOC685067 | 0 | 0.29416268 | 0.553 | 0.879 | 0 |
| Subacute Remote | Up | Plod3 | 9.27E-167 | 0.29398055 | 0.488 | 0.737 | 1.85E-163 |
| Subacute Remote | Up | Hs2st1 | 2.82E-157 | 0.29386953 | 0.504 | 0.724 | 5.65E-154 |
| Subacute Remote | Up | Ap3s1 | 7.52E-304 | 0.29237948 | 0.453 | 0.806 | 1.50E-300 |

| | | | | | | | |
|---|---|---|---|---|---|---|---|
| Subacute Remote | Up | Rab3d | 4.23E-305 | 0.29223342 | 0.42 | 0.741 | 8.47E-302 |
| Subacute Remote | Up | Spon1 | 4.96E-225 | 0.29214353 | 0.502 | 0.829 | 9.93E-222 |
| Subacute Remote | Up | Pea15 | 2.53E-62 | 0.29142199 | 0.589 | 0.81 | 5.07E-59 |
| Subacute Remote | Up | Dock8 | 0 | 0.28987889 | 0.454 | 0.8 | 0 |
| Subacute Remote | Up | Ece1 | 2.81E-187 | 0.28938455 | 0.378 | 0.598 | 5.62E-184 |
| Subacute Remote | Up | Elovl1 | 2.67E-71 | 0.28928142 | 0.599 | 0.826 | 5.34E-68 |
| Subacute Remote | Up | Aldh3a1 | 5.64E-257 | 0.28920768 | 0.389 | 0.618 | 1.13E-253 |
| Subacute Remote | Up | Gbp1 | 4.33E-45 | 0.28877079 | 0.64 | 0.954 | 8.66E-42 |
| Subacute Remote | Up | Npw | 2.63E-271 | 0.28802614 | 0.35 | 0.605 | 5.25E-268 |
| Subacute Remote | Up | Sema3c | 3.89E-154 | 0.28689083 | 0.502 | 0.772 | 7.79E-151 |
| Subacute Remote | Up | Zc3h13 | 3.05E-153 | 0.28677522 | 0.526 | 0.77 | 6.10E-150 |
| Subacute Remote | Up | Gsta1 | 8.13E-124 | 0.28674172 | 0.567 | 0.852 | 1.63E-120 |
| Subacute Remote | Up | Lgmn | 1.31E-27 | 0.28668583 | 0.661 | 0.878 | 2.61E-24 |
| Subacute Remote | Up | Irf7 | 1.91E-44 | 0.28629362 | 0.625 | 0.885 | 3.82E-41 |

| | | | | | | | |
|---|---|---|---|---|---|---|---|
| Subacute Remote | Up | Pcf11 | 6.11E-104 | 0.28593968 | 0.547 | 0.828 | 1.22E-100 |
| Subacute Remote | Up | Stk38l | 3.85E-288 | 0.28570412 | 0.345 | 0.602 | 7.70E-285 |
| Subacute Remote | Up | Ddias | 0 | 0.2854652 | 0.552 | 0.815 | 0 |
| Subacute Remote | Up | Galk1 | 4.56E-258 | 0.2848689 | 0.451 | 0.754 | 9.13E-255 |
| Subacute Remote | Up | Bnc2 | 3.59E-167 | 0.28310857 | 0.484 | 0.843 | 7.19E-164 |
| Subacute Remote | Up | Dusp2 | 3.24E-186 | 0.28248467 | 0.537 | 0.797 | 6.47E-183 |
| Subacute Remote | Up | Capg | 3.40E-105 | 0.28235644 | 0.515 | 0.746 | 6.80E-102 |
| Subacute Remote | Up | Sh3bgrl3 | 5.26E-38 | 0.28226612 | 0.66 | 0.886 | 1.05E-34 |
| Subacute Remote | Up | Twf1 | 0 | 0.28193945 | 0.354 | 0.749 | 0 |
| Subacute Remote | Up | Nucb1 | 2.84E-68 | 0.28174789 | 0.505 | 0.786 | 5.67E-65 |
| Subacute Remote | Up | Xdh | 5.32E-21 | 0.28172553 | 0.729 | 0.944 | 1.06E-17 |
| Subacute Remote | Up | MGC105649 | 2.76E-222 | 0.2796187 | 0.517 | 0.839 | 5.51E-219 |
| Subacute Remote | Up | Slfn2 | 6.12E-154 | 0.27945293 | 0.612 | 0.911 | 1.22E-150 |
| Subacute Remote | Up | Dchs1 | 1.25E-232 | 0.27939022 | 0.446 | 0.777 | 2.49E-229 |

| | | | | | | | |
|---|---|---|---|---|---|---|---|
| Subacute Remote | Up | Ramp3 | 1.42E-285 | 0.27925636 | 0.456 | 0.825 | 2.85E-282 |
| Subacute Remote | Up | Cd9 | 3.29E-100 | 0.27897973 | 0.499 | 0.815 | 6.57E-97 |
| Subacute Remote | Up | Tnfaip6 | 9.87E-285 | 0.27849211 | 0.409 | 0.664 | 1.97E-281 |
| Subacute Remote | Up | Col18a1 | 1.64E-132 | 0.27841397 | 0.561 | 0.878 | 3.27E-129 |
| Subacute Remote | Up | Pias1 | 6.59E-147 | 0.27820516 | 0.505 | 0.804 | 1.32E-143 |
| Subacute Remote | Up | S100a4 | 6.87E-24 | 0.27804387 | 0.704 | 0.934 | 1.37E-20 |
| Subacute Remote | Up | Cacnb3 | 6.75E-279 | 0.27716478 | 0.484 | 0.845 | 1.35E-275 |
| Subacute Remote | Up | Smim14 | 8.08E-222 | 0.27683555 | 0.375 | 0.582 | 1.62E-218 |
| Subacute Remote | Up | Grk6 | 3.76E-306 | 0.27679891 | 0.395 | 0.723 | 7.52E-303 |
| Subacute Remote | Up | Adss | 8.87E-211 | 0.27622228 | 0.494 | 0.778 | 1.77E-207 |
| Subacute Remote | Up | Flna | 2.81E-68 | 0.27570088 | 0.561 | 0.917 | 5.62E-65 |
| Subacute Remote | Up | Thbd | 5.22E-143 | 0.27548276 | 0.612 | 0.965 | 1.04E-139 |
| Subacute Remote | Up | Fst | 0 | 0.2750644 | 0.319 | 0.632 | 0 |
| Subacute Remote | Up | Cnpy4 | 7.21E-139 | 0.27473607 | 0.531 | 0.788 | 1.44E-135 |

| | | | | | | | |
|---|---|---|---|---|---|---|---|
| Subacute Remote | Up | Mmp14 | 3.90E-28 | 0.27392351 | 0.603 | 0.828 | 7.79E-25 |
| Subacute Remote | Up | Fat1 | 4.87E-110 | 0.2731381 | 0.504 | 0.708 | 9.74E-107 |
| Subacute Remote | Up | Mcub | 1.70E-225 | 0.27206129 | 0.447 | 0.751 | 3.40E-222 |
| Subacute Remote | Up | Sbk2 | 0 | 0.27201976 | 0.475 | 0.82 | 0 |
| Subacute Remote | Up | RGD1310951 | 4.12E-179 | 0.27157915 | 0.491 | 0.76 | 8.25E-176 |
| Subacute Remote | Up | AABR070179 | 3.70E-297 | 0.2703715 | 0.488 | 0.714 | 7.39E-294 |
| Subacute Remote | Up | Fkbp7 | 3.31E-130 | 0.27009419 | 0.507 | 0.769 | 6.62E-127 |
| Subacute Remote | Up | Pald1 | 8.99E-157 | 0.26980875 | 0.475 | 0.688 | 1.80E-153 |
| Subacute Remote | Up | Plxnb2 | 1.38E-61 | 0.26948964 | 0.674 | 0.946 | 2.77E-58 |
| Subacute Remote | Up | Tmem43 | 0 | 0.26886675 | 0.453 | 0.834 | 0 |
| Subacute Remote | Up | LOC1003601 | 1.23E-74 | 0.26790351 | 0.489 | 0.79 | 2.47E-71 |
| Subacute Remote | Up | Kidins220 | 4.14E-189 | 0.26772125 | 0.471 | 0.756 | 8.28E-186 |
| Subacute Remote | Up | Mical1 | 1.38E-185 | 0.26770484 | 0.48 | 0.753 | 2.76E-182 |
| Subacute Remote | Up | Emilin1 | 4.07E-86 | 0.26671166 | 0.552 | 0.779 | 8.14E-83 |

| | | | | | | | |
|---|---|---|---|---|---|---|---|
| Subacute Remote | Up | Ccdc85a | 2.97E-283 | 0.26652716 | 0.282 | 0.55 | 5.94E-280 |
| Subacute Remote | Up | Kcnj8 | 8.81E-112 | 0.26633951 | 0.496 | 0.72 | 1.76E-108 |
| Subacute Remote | Up | Myoz1 | 4.42E-168 | 0.26632442 | 0.451 | 0.68 | 8.85E-165 |
| Subacute Remote | Up | Golgb1 | 4.82E-178 | 0.26620478 | 0.496 | 0.784 | 9.64E-175 |
| Subacute Remote | Up | Lxn | 1.77E-295 | 0.26593567 | 0.466 | 0.806 | 3.55E-292 |
| Subacute Remote | Up | Ror1 | 4.45E-240 | 0.2646486 | 0.485 | 0.802 | 8.89E-237 |
| Subacute Remote | Up | Rab8a | 1.12E-219 | 0.26376964 | 0.446 | 0.753 | 2.23E-216 |
| Subacute Remote | Up | Cntfr | 8.91E-130 | 0.26335778 | 0.418 | 0.652 | 1.78E-126 |
| Subacute Remote | Up | Frzb | 8.43E-306 | 0.26192117 | 0.339 | 0.563 | 1.69E-302 |
| Subacute Remote | Up | Jaml | 1.90E-280 | 0.26163736 | 0.519 | 0.859 | 3.81E-277 |
| Subacute Remote | Up | Rad18 | 1.56E-251 | 0.26140739 | 0.467 | 0.729 | 3.12E-248 |
| Subacute Remote | Up | Thoc2 | 2.39E-211 | 0.25920173 | 0.562 | 0.945 | 4.79E-208 |
| Subacute Remote | Up | Pdgfra | 8.43E-215 | 0.25887924 | 0.514 | 0.914 | 1.69E-211 |
| Subacute Remote | Up | Cxcr4 | 1.40E-171 | 0.25848932 | 0.487 | 0.735 | 2.81E-168 |

| | | | | | | | |
|---|---|---|---|---|---|---|---|
| Subacute Remote | Up | Thbs3 | 0 | 0.25776782 | 0.36 | 0.888 | 0 |
| Subacute Remote | Up | Vkorc1 | 4.39E-38 | 0.25742085 | 0.664 | 0.891 | 8.77E-35 |
| Subacute Remote | Up | Atad2 | 1.99E-190 | 0.25719809 | 0.431 | 0.686 | 3.97E-187 |
| Subacute Remote | Up | Slc39a7 | 4.91E-106 | 0.2567515 | 0.547 | 0.82 | 9.83E-103 |
| Subacute Remote | Up | Pcdhb9 | 3.54E-189 | 0.25537884 | 0.489 | 0.71 | 7.08E-186 |
| Subacute Remote | Up | Csf2ra | 5.64E-146 | 0.25474411 | 0.535 | 0.786 | 1.13E-142 |
| Subacute Remote | Up | Fzd1 | 3.31E-194 | 0.25458202 | 0.391 | 0.598 | 6.62E-191 |
| Subacute Remote | Up | Vwf | 3.50E-52 | 0.2537364 | 0.622 | 0.829 | 6.99E-49 |
| Subacute Remote | Up | Gzmbl2 | 0 | 0.25373244 | 0.549 | 0.899 | 0 |
| Subacute Remote | Up | Coro1c | 1.17E-262 | 0.25358176 | 0.423 | 0.751 | 2.34E-259 |
| Subacute Remote | Up | Actn4 | 9.86E-17 | 0.25294155 | 0.757 | 0.963 | 1.97E-13 |
| Subacute Remote | Up | Mgst1 | 4.98E-64 | 0.25291813 | 0.587 | 0.789 | 9.95E-61 |
| Subacute Remote | Up | Rsf1 | 2.63E-154 | 0.251734 | 0.52 | 0.814 | 5.26E-151 |
| Subacute Remote | Up | Fabp5 | 5.20E-46 | 0.25022445 | 0.64 | 0.908 | 1.04E-42 |

| | | | | | | | |
|---|---|---|---|---|---|---|---|
| Chronic Remote | Up | AABR070252 | 7.16E-179 | -0.251919 | 0.396 | 0.633 | 1.43E-175 |
| Chronic Remote | Up | AC111831.1 | 2.66E-257 | -0.253902 | 0.414 | 0.668 | 5.32E-254 |
| Chronic Remote | Up | Gapt | 2.16E-172 | -0.2622199 | 0.546 | 0.76 | 4.31E-169 |
| Chronic Remote | Up | Ldb2 | 6.66E-292 | -0.273384 | 0.431 | 0.724 | 1.33E-288 |
| Chronic Remote | Up | Pwwp3b | 2.99E-296 | -0.2751143 | 0.535 | 0.879 | 5.97E-293 |
| Chronic Remote | Up | LOC1003616 | 3.48E-265 | -0.2786879 | 0.538 | 0.786 | 6.97E-262 |
| Chronic Remote | Up | Lsamp | 0 | -0.2798042 | 0.486 | 0.882 | 0 |
| Chronic Remote | Up | Rcn1 | 0 | -0.2805946 | 0.532 | 0.867 | 0 |
| Chronic Remote | Up | AABR070392 | 3.22E-280 | -0.2828325 | 0.487 | 0.717 | 6.44E-277 |
| Chronic Remote | Up | Zfhx2 | 0 | -0.2907598 | 0.469 | 0.763 | 0 |
| Chronic Remote | Up | Kdm6b | 0 | -0.2924901 | 0.464 | 0.774 | 0 |
| Chronic Remote | Up | Mill1 | 2.10E-203 | -0.3026659 | 0.461 | 0.664 | 4.19E-200 |
| Chronic Remote | Up | L1cam | 0 | -0.3184086 | 0.359 | 0.642 | 0 |
| Chronic Remote | Up | Mefv | 2.22E-243 | -0.3189967 | 0.509 | 0.766 | 4.44E-240 |

| | | | | | | | |
|---|---|---|---|---|---|---|---|
| Chronic Remote | Up | RGD1565166 | 1.26E-183 | -0.3353515 | 0.544 | 0.834 | 2.53E-180 |
| Chronic Remote | Up | Elf4 | 0 | -0.3367187 | 0.486 | 0.805 | 0 |
| Chronic Remote | Up | Camkk2 | 2.31E-154 | -0.3513206 | 0.548 | 0.778 | 4.61E-151 |
| Chronic Remote | Up | Nol11 | 2.04E-251 | -0.3628301 | 0.339 | 0.557 | 4.09E-248 |
| Chronic Remote | Up | Spdya | 1.70E-239 | -0.3767073 | 0.53 | 0.803 | 3.39E-236 |
| Chronic Remote | Up | Tsc1 | 5.74E-199 | -0.3867372 | 0.537 | 0.745 | 1.15E-195 |
| Chronic Remote | Up | LOC688553 | 8.52E-257 | -0.3922616 | 0.33 | 0.541 | 1.70E-253 |
| Chronic Remote | Up | LOC1036948 | 1.96E-297 | -0.3959281 | 0.317 | 0.578 | 3.92E-294 |
| Chronic Remote | Up | Stx17 | 6.27E-197 | -0.4037181 | 0.495 | 0.731 | 1.25E-193 |
| Chronic Remote | Up | Acpp | 4.09E-259 | -0.4090896 | 0.488 | 0.784 | 8.18E-256 |
| Chronic Remote | Up | Opn4 | 3.45E-265 | -0.4126144 | 0.341 | 0.582 | 6.90E-262 |
| Chronic Remote | Up | Slc24a1 | 7.26E-141 | -0.4138939 | 0.513 | 0.753 | 1.45E-137 |
| Chronic Remote | Up | Atp6v1g2 | 3.18E-265 | -0.4228535 | 0.387 | 0.598 | 6.37E-262 |
| Chronic Remote | Up | C1qtnf4 | 8.97E-290 | -0.4307379 | 0.286 | 0.527 | 1.79E-286 |

| | | | | | | | |
|---|---|---|---|---|---|---|---|
| Chronic Remote | Up | Tbx3 | 0 | -0.4371933 | 0.563 | 0.901 | 0 |
| Chronic Remote | Up | Gpt2 | 3.47E-221 | -0.4458265 | 0.353 | 0.594 | 6.94E-218 |
| Chronic Remote | Up | Tshz2 | 4.12E-195 | -0.4498695 | 0.558 | 0.821 | 8.25E-192 |
| Chronic Remote | Up | Igsf9 | 1.99E-219 | -0.4634344 | 0.49 | 0.791 | 3.98E-216 |
| Chronic Remote | Up | Esm1 | 0 | -0.4641842 | 0.284 | 0.555 | 0 |
| Chronic Remote | Up | Zfp26 | 0 | -0.4680973 | 0.491 | 0.858 | 0 |
| Chronic Remote | Up | Fjx1 | 4.35E-271 | -0.4694302 | 0.384 | 0.695 | 8.69E-268 |
| Chronic Remote | Up | Ttll5 | 0 | -0.4902187 | 0.534 | 0.828 | 0 |
| Chronic Remote | Up | Sncg | 9.62E-285 | -0.4985733 | 0.405 | 0.684 | 1.92E-281 |
| Chronic Remote | Up | Siglec10 | 0 | -0.5020217 | 0.435 | 0.719 | 0 |
| Chronic Remote | Up | Rergl | 0 | -0.5032418 | 0.473 | 0.774 | 0 |
| Chronic Remote | Up | Mcpt1l4 | 4.15E-262 | -0.5362028 | 0.432 | 0.722 | 8.30E-259 |
| Chronic Remote | Up | AABR070608 | 0 | -0.5364133 | 0.312 | 0.759 | 0 |
| Chronic Remote | Up | AC109737.1 | 2.21E-173 | -0.5521458 | 0.479 | 0.717 | 4.41E-170 |

| | | | | | | | |
|---|---|---|---|---|---|---|---|
| Chronic Remote | Up | Atp10d | 5.72E-279 | -0.5788009 | 0.416 | 0.707 | 1.14E-275 |
| Chronic Remote | Up | Mlxip | 1.86E-278 | -0.6178376 | 0.51 | 0.75 | 3.72E-275 |
| Chronic Remote | Up | Folr1 | 5.65E-192 | -0.7624192 | 0.554 | 0.873 | 1.13E-188 |
| Chronic Remote | Up | Hemgn | 8.89E-152 | -0.7990182 | 0.554 | 0.807 | 1.78E-148 |
| Chronic Remote | Up | Snorc | 1.28E-208 | -0.8094345 | 0.257 | 0.462 | 2.55E-205 |
| Chronic Remote | Up | AABR070683 | 5.71E-175 | -0.8252112 | 0.459 | 0.717 | 1.14E-171 |
| Chronic Remote | Up | Foxred1 | 5.92E-245 | -0.8278574 | 0.463 | 0.748 | 1.18E-241 |
| Chronic Remote | Up | LOC299282 | 2.60E-76 | -0.8590547 | 0.498 | 0.725 | 5.20E-73 |
| Chronic Remote | Up | Mybph | 0 | -0.9473239 | 0.481 | 0.803 | 0 |
| Chronic Remote | Up | Mttp | 2.20E-292 | -0.9726463 | 0.436 | 0.678 | 4.40E-289 |
| Chronic Remote | Up | Map10 | 7.51E-284 | -1.0431979 | 0.48 | 0.73 | 1.50E-280 |
| Chronic Remote | Up | Fmnl2 | 1.15E-302 | -1.0728517 | 0.505 | 0.827 | 2.30E-299 |
| Chronic Remote | Up | Plekhh2 | 0 | -1.2106767 | 0.32 | 0.604 | 0 |
| Chronic Remote | Up | Chad | 4.42E-131 | -1.2677922 | 0.422 | 0.63 | 8.83E-128 |

| | | | | | | | |
|---|---|---|---|---|---|---|---|
| Chronic Remote | Up | LOC1036917 | 3.54E-230 | -1.3117466 | 0.237 | 0.44 | 7.08E-227 |
| Chronic Remote | Up | Ccl11 | 0 | -1.6723543 | 0.251 | 0.522 | 0 |
| Chronic Remote | Up | AABR070504 | 1.67E-265 | -2.0873584 | 0.176 | 0.415 | 3.35E-262 |
| Chronic Remote | Up | AABR070010 | 1.27E-217 | -2.5680923 | 0.112 | 0.329 | 2.54E-214 |
