## Supplemental Table 18 for "Uncovering the Regional and Cell Specific Bioactivity of Injectable Extracellular Matrix Biomaterials in Myocardial Infarction through Spatial and Single Nucleus Transcriptomics"

**Supplementary Table 18. Cell-Specific Differentially Expressed Genes in the Subacute (Up) or Chronic (Down) with ECM Hydrogel Administration**

| Subsetted Cell Type | Direction | Gene | p_val | avg_log2FC | pct.1 | pct.2 | p_val_adj |
|---|---|---|---|---|---|---|---|
| Macrophages | Up | Slc26a10 | 1.14E-13 | 0.95741347 | 0.172 | 0.956 | 2.28E-10 |
| Macrophages | Up | Slc7a8 | 3.42E-12 | 0.86145843 | 0.201 | 0.978 | 6.84E-09 |
| Macrophages | Up | Scarb1 | 7.75E-11 | 0.78182089 | 0.235 | 1 | 1.55E-07 |
| Macrophages | Up | Mylk3 | 3.11E-16 | 0.73329437 | 0.162 | 0.978 | 6.22E-13 |
| Macrophages | Up | Flrt2 | 3.52E-08 | 0.73139023 | 0.225 | 0.933 | 7.03E-05 |
| Macrophages | Up | Acsl1 | 1.96E-05 | 0.6780788 | 0.206 | 0.844 | 0.03913298 |
| Macrophages | Up | Slc7a11 | 7.42E-20 | 0.67727185 | 0.015 | 0.467 | 1.48E-16 |
| Macrophages | Up | Itga4 | 6.36E-11 | 0.6661043 | 0.24 | 1 | 1.27E-07 |
| Macrophages | Up | Lrrc2 | 2.60E-14 | 0.65846219 | 0.132 | 0.911 | 5.20E-11 |
| Macrophages | Up | Mpc1 | 1.23E-14 | 0.65408478 | 0.162 | 0.956 | 2.46E-11 |
| Macrophages | Up | Cemip | 6.51E-07 | 0.64185893 | 0.054 | 0.711 | 0.00130255 |
| Macrophages | Up | Lhfpl2 | 2.31E-07 | 0.6417728 | 0.275 | 0.978 | 0.00046246 |
| Macrophages | Up | Tnni3k | 2.78E-10 | 0.63505254 | 0.216 | 0.956 | 5.56E-07 |
| Macrophages | Up | Srgap3 | 2.71E-07 | 0.61450414 | 0.172 | 0.844 | 0.0005423 |
| Macrophages | Up | Pik3ap1 | 4.95E-06 | 0.60939053 | 0.304 | 0.978 | 0.00990124 |
| Macrophages | Up | Tgfb3 | 6.15E-16 | 0.59798053 | 0.118 | 0.911 | 1.23E-12 |
| Macrophages | Up | Kcnd3 | 9.55E-13 | 0.59530988 | 0.098 | 0.844 | 1.91E-09 |
| Macrophages | Up | LOC690097 | 9.44E-08 | 0.57904354 | 0.255 | 0.956 | 0.0001888 |
| Macrophages | Up | Gm2a | 1.09E-14 | 0.57788139 | 0.113 | 0.889 | 2.18E-11 |
| Macrophages | Up | Podxl | 6.99E-07 | 0.57331191 | 0.201 | 0.867 | 0.00139809 |
| Macrophages | Up | Atp1b1 | 4.22E-10 | 0.56023217 | 0.245 | 1 | 8.44E-07 |
| Macrophages | Up | Serpine1 | 6.94E-19 | 0.55850125 | 0.078 | 0.889 | 1.39E-15 |
| Macrophages | Up | Ntrk3 | 1.62E-16 | 0.55670514 | 0.113 | 0.911 | 3.24E-13 |
| Macrophages | Up | S100a6 | 1.19E-20 | 0.54317419 | 0.142 | 1 | 2.37E-17 |
| Macrophages | Up | Kcnn4 | 1.75E-12 | 0.54106872 | 0.186 | 0.956 | 3.50E-09 |
| Macrophages | Up | Slc1a7 | 3.28E-08 | 0.52914354 | 0.078 | 0.733 | 6.55E-05 |

| | | | | | | | |
|---|---|---|---|---|---|---|---|
| Macrophages | Up | Ccn2 | 2.14E-14 | 0.52677557 | 0.196 | 1 | 4.28E-11 |
| Macrophages | Up | Pax5 | 1.89E-10 | 0.52677243 | 0.039 | 0.511 | 3.78E-07 |
| Macrophages | Up | Atp6v0c | 1.07E-05 | 0.52250467 | 0.245 | 0.889 | 0.02133289 |
| Macrophages | Up | Macrod1 | 1.60E-14 | 0.51522348 | 0.132 | 0.911 | 3.19E-11 |
| Macrophages | Up | AABR070311 | 1.26E-30 | 0.50859193 | 0.074 | 0.956 | 2.52E-27 |
| Macrophages | Up | Ckm | 8.12E-15 | 0.50720703 | 0.162 | 0.956 | 1.62E-11 |
| Macrophages | Up | Atp5mc1 | 6.63E-20 | 0.48984044 | 0.132 | 0.978 | 1.33E-16 |
| Macrophages | Up | Ndufa5 | 3.27E-09 | 0.48581439 | 0.123 | 0.822 | 6.55E-06 |
| Macrophages | Up | Fabp3 | 3.64E-09 | 0.47395836 | 0.162 | 0.867 | 7.28E-06 |
| Macrophages | Up | Casq2 | 5.84E-15 | 0.4710038 | 0.162 | 0.956 | 1.17E-11 |
| Macrophages | Up | Pygm | 3.22E-08 | 0.46728599 | 0.078 | 0.733 | 6.44E-05 |
| Macrophages | Up | Cux2 | 5.21E-27 | 0.46598387 | 0.083 | 0.956 | 1.04E-23 |
| Macrophages | Up | Parm1 | 9.15E-13 | 0.46397486 | 0.098 | 0.844 | 1.83E-09 |
| Macrophages | Up | Trim63 | 6.13E-11 | 0.46178328 | 0.137 | 0.867 | 1.23E-07 |
| Macrophages | Up | P2rx7 | 2.73E-10 | 0.45911335 | 0.088 | 0.8 | 5.46E-07 |
| Macrophages | Up | Cd68 | 1.47E-37 | 0.45530113 | 0.054 | 1 | 2.95E-34 |
| Macrophages | Up | Atp5f1e | 2.97E-19 | 0.44117671 | 0.108 | 0.933 | 5.94E-16 |
| Macrophages | Up | Atp6v0d2 | 6.07E-34 | 0.44060714 | 0.069 | 1 | 1.21E-30 |
| Macrophages | Up | Slc25a20 | 6.63E-20 | 0.43895769 | 0.132 | 0.978 | 1.33E-16 |
| Macrophages | Up | Piezo2 | 5.50E-06 | 0.43663072 | 0.093 | 0.733 | 0.01099768 |
| Macrophages | Up | Tm4sf1 | 8.04E-23 | 0.42404946 | 0.098 | 0.956 | 1.61E-19 |
| Macrophages | Up | Cst3 | 2.60E-09 | 0.42124395 | 0.162 | 0.867 | 5.19E-06 |
| Macrophages | Up | Ddit3 | 4.86E-19 | 0.4185936 | 0.093 | 0.911 | 9.72E-16 |
| Macrophages | Up | Sh3tc2 | 1.68E-05 | 0.41231855 | 0.098 | 0.378 | 0.03364022 |
| Macrophages | Up | LOC1036942 | 7.91E-13 | 0.41143866 | 0.059 | 0.733 | 1.58E-09 |
| Macrophages | Up | Ccdc88b | 4.23E-08 | 0.41048574 | 0.083 | 0.711 | 8.46E-05 |
| Macrophages | Up | Actn2 | 1.51E-05 | 0.40646136 | 0.167 | 0.8 | 0.03021253 |
| Macrophages | Up | Col8a2 | 1.10E-07 | 0.39991794 | 0.044 | 0.689 | 0.00022065 |
| Macrophages | Up | Dok3 | 2.59E-08 | 0.39510737 | 0.25 | 0.956 | 5.19E-05 |
| Macrophages | Up | Ednra | 3.08E-08 | 0.39309727 | 0.137 | 0.822 | 6.16E-05 |
| Macrophages | Up | Lims2 | 4.38E-11 | 0.38746739 | 0.206 | 0.956 | 8.76E-08 |

| | | | | | | | |
|---|---|---|---|---|---|---|---|
| Macrophages | Up | Hvcn1 | 6.82E-13 | 0.38742089 | 0.201 | 0.978 | 1.36E-09 |
| Macrophages | Up | Atp1a3 | 1.47E-14 | 0.38734124 | 0.029 | 0.467 | 2.95E-11 |
| Macrophages | Up | Hspb7 | 2.69E-25 | 0.38698377 | 0.083 | 0.956 | 5.38E-22 |
| Macrophages | Up | Slc4a3 | 7.43E-18 | 0.38339213 | 0.118 | 0.933 | 1.49E-14 |
| Macrophages | Up | Prodh1 | 4.30E-08 | 0.37818058 | 0.069 | 0.733 | 8.60E-05 |
| Macrophages | Up | Speg | 8.67E-10 | 0.37656952 | 0.108 | 0.8 | 1.73E-06 |
| Macrophages | Up | Vwf | 3.58E-09 | 0.37443333 | 0.162 | 0.867 | 7.16E-06 |
| Macrophages | Up | Ccn5 | 9.63E-32 | 0.37286299 | 0.078 | 1 | 1.93E-28 |
| Macrophages | Up | Crispld2 | 1.79E-13 | 0.3703199 | 0.127 | 0.889 | 3.58E-10 |
| Macrophages | Up | Ca4 | 1.92E-18 | 0.36766326 | 0.108 | 0.911 | 3.85E-15 |
| Macrophages | Up | Eef1a2 | 4.70E-30 | 0.36262414 | 0.059 | 0.956 | 9.40E-27 |
| Macrophages | Up | AABR070490 | 3.06E-20 | 0.35744965 | 0.147 | 1 | 6.12E-17 |
| Macrophages | Up | Enpp3 | 2.51E-10 | 0.35656355 | 0.069 | 0.778 | 5.01E-07 |
| Macrophages | Up | Igfbp3 | 1.08E-29 | 0.35638587 | 0.088 | 1 | 2.16E-26 |
| Macrophages | Up | Pcsk5 | 4.38E-15 | 0.35627474 | 0.162 | 0.956 | 8.76E-12 |
| Macrophages | Up | Galnt16 | 1.17E-15 | 0.35231346 | 0.137 | 0.933 | 2.34E-12 |
| Macrophages | Up | Treml4 | 2.89E-13 | 0.34633309 | 0.059 | 0.711 | 5.78E-10 |
| Macrophages | Up | Cp | 2.57E-10 | 0.34450877 | 0.23 | 0.978 | 5.14E-07 |
| Macrophages | Up | LOC1003644 | 4.17E-09 | 0.34404056 | 0.221 | 0.933 | 8.34E-06 |
| Macrophages | Up | Grk3 | 7.06E-07 | 0.34184816 | 0.181 | 0.844 | 0.00141255 |
| Macrophages | Up | Islr | 8.00E-33 | 0.33953102 | 0.074 | 1 | 1.60E-29 |
| Macrophages | Up | Cytip | 4.08E-11 | 0.33599212 | 0.147 | 0.867 | 8.17E-08 |
| Macrophages | Up | Erbb4 | 1.74E-09 | 0.33533925 | 0.059 | 0.756 | 3.48E-06 |
| Macrophages | Up | Homer1 | 2.47E-16 | 0.33470543 | 0.132 | 0.933 | 4.94E-13 |
| Macrophages | Up | LOC1009119 | 4.97E-28 | 0.33152785 | 0.049 | 0.911 | 9.94E-25 |
| Macrophages | Up | Sgca | 5.11E-22 | 0.33073253 | 0.088 | 0.933 | 1.02E-18 |
| Macrophages | Up | Spink8 | 7.99E-22 | 0.32729824 | 0.059 | 0.889 | 1.60E-18 |
| Macrophages | Up | Cobl | 7.15E-09 | 0.32473004 | 0.049 | 0.711 | 1.43E-05 |
| Macrophages | Up | Ddah1 | 3.27E-11 | 0.32037024 | 0.078 | 0.8 | 6.55E-08 |
| Macrophages | Up | Jag2 | 2.41E-10 | 0.31389342 | 0.069 | 0.778 | 4.82E-07 |
| Macrophages | Up | Hk3 | 6.35E-21 | 0.31372883 | 0.118 | 0.911 | 1.27E-17 |

| | | | | | | | |
|---|---|---|---|---|---|---|---|
| Macrophages | Up | Nox4 | 2.35E-08 | 0.31367572 | 0.157 | 0.844 | 4.70E-05 |
| Macrophages | Up | Gask1b | 9.16E-08 | 0.31277613 | 0.26 | 0.956 | 0.00018324 |
| Macrophages | Up | Des | 4.80E-11 | 0.31064712 | 0.24 | 1 | 9.60E-08 |
| Macrophages | Up | Tmem163 | 4.67E-09 | 0.30962479 | 0.029 | 0.444 | 9.35E-06 |
| Macrophages | Up | Slc12a7 | 3.19E-07 | 0.30955531 | 0.26 | 0.933 | 0.00063846 |
| Macrophages | Up | Kcnq1 | 1.72E-10 | 0.30820861 | 0.127 | 0.844 | 3.43E-07 |
| Macrophages | Up | Gxylt2 | 2.06E-06 | 0.30454868 | 0.216 | 0.867 | 0.0041234 |
| Macrophages | Up | Dsp | 1.72E-05 | 0.29855821 | 0.108 | 0.378 | 0.03443756 |
| Macrophages | Up | Cd63 | 1.42E-17 | 0.29632345 | 0.137 | 0.956 | 2.84E-14 |
| Macrophages | Up | Arhgap44 | 1.32E-09 | 0.29467921 | 0.069 | 0.756 | 2.65E-06 |
| Macrophages | Up | Acyp2 | 2.61E-07 | 0.29389195 | 0.113 | 0.778 | 0.00052123 |
| Macrophages | Up | Ctsd | 1.73E-10 | 0.2934702 | 0.25 | 1 | 3.46E-07 |
| Macrophages | Up | Cadps | 1.25E-11 | 0.29233029 | 0.054 | 0.778 | 2.50E-08 |
| Macrophages | Up | Tmem82 | 3.79E-24 | 0.29103303 | 0.064 | 0.889 | 7.58E-21 |
| Macrophages | Up | Npr3 | 6.82E-06 | 0.28432339 | 0.049 | 0.689 | 0.01363352 |
| Macrophages | Up | Wt1 | 1.63E-18 | 0.28431804 | 0.098 | 0.911 | 3.27E-15 |
| Macrophages | Up | Ckmt2 | 2.31E-17 | 0.28371492 | 0.172 | 1 | 4.62E-14 |
| Macrophages | Up | Plaur | 5.80E-12 | 0.28343223 | 0.235 | 1 | 1.16E-08 |
| Macrophages | Up | Lilrb2 | 8.12E-20 | 0.28065288 | 0.152 | 1 | 1.62E-16 |
| Macrophages | Up | Spp1 | 2.14E-15 | 0.27576803 | 0.093 | 0.867 | 4.28E-12 |
| Macrophages | Up | Aldh1a2 | 2.85E-34 | 0.26969414 | 0.049 | 0.956 | 5.70E-31 |
| Macrophages | Up | Sntg2 | 1.08E-05 | 0.26912687 | 0.044 | 0.644 | 0.02158963 |
| Macrophages | Up | Ndst3 | 1.57E-05 | 0.26865987 | 0.039 | 0.6 | 0.03135663 |
| Macrophages | Up | AABR070010 | 7.04E-11 | 0.26792576 | 0.054 | 0.756 | 1.41E-07 |
| Macrophages | Up | Dcdc5 | 5.78E-26 | 0.26372411 | 0.039 | 0.867 | 1.16E-22 |
| Macrophages | Up | Wnk2 | 1.65E-24 | 0.26123352 | 0.088 | 0.956 | 3.29E-21 |
| Macrophages | Up | Fsd2 | 4.78E-27 | 0.26075464 | 0.103 | 1 | 9.55E-24 |
| Macrophages | Up | Vash2 | 1.17E-20 | 0.26021812 | 0.044 | 0.844 | 2.34E-17 |
| Macrophages | Up | Atf3 | 1.18E-17 | 0.26017663 | 0.025 | 0.556 | 2.36E-14 |
| Macrophages | Up | Cox6a2 | 4.69E-06 | 0.25686289 | 0.225 | 0.867 | 0.00938678 |
| Macrophages | Up | Cthrc1 | 3.33E-40 | 0.25240716 | 0.044 | 1 | 6.66E-37 |

| | | | | | | | |
|---|---|---|---|---|---|---|---|
| Macrophages | Down | Corin | 3.27E-14 | -0.2517668 | 0.069 | 0.822 | 6.54E-11 |
| Macrophages | Down | Emb | 3.79E-12 | -0.2544409 | 0.069 | 0.733 | 7.58E-09 |
| Macrophages | Down | Fcgr2a | 1.48E-20 | -0.2549396 | 0.127 | 0.956 | 2.95E-17 |
| Macrophages | Down | Nav3 | 1.15E-06 | -0.2635464 | 0.343 | 1 | 0.00229672 |
| Macrophages | Down | Fgr | 2.33E-15 | -0.2645818 | 0.152 | 0.933 | 4.66E-12 |
| Macrophages | Down | C1qc | 1.27E-11 | -0.2680433 | 0.23 | 0.978 | 2.55E-08 |
| Macrophages | Down | Wipf3 | 4.29E-15 | -0.2762498 | 0.157 | 0.933 | 8.58E-12 |
| Macrophages | Down | Hopx | 5.28E-07 | -0.2766475 | 0.015 | 0.378 | 0.00105512 |
| Macrophages | Down | Diaph3 | 1.28E-10 | -0.2801946 | 0.059 | 0.756 | 2.55E-07 |
| Macrophages | Down | Asgr2 | 1.03E-09 | -0.2828849 | 0.211 | 0.911 | 2.06E-06 |
| Macrophages | Down | Msr1 | 6.06E-16 | -0.2838516 | 0.172 | 0.978 | 1.21E-12 |
| Macrophages | Down | Itgb4 | 2.11E-13 | -0.2854274 | 0.029 | 0.756 | 4.21E-10 |
| Macrophages | Down | LOC1036933 | 3.77E-10 | -0.2870167 | 0.015 | 0.289 | 7.54E-07 |
| Macrophages | Down | Ccl6 | 1.62E-22 | -0.2872663 | 0.088 | 0.933 | 3.25E-19 |
| Macrophages | Down | Eml6 | 9.56E-16 | -0.2913989 | 0.127 | 0.911 | 1.91E-12 |
| Macrophages | Down | AABR070440 | 2.25E-11 | -0.2916703 | 0.181 | 0.911 | 4.49E-08 |
| Macrophages | Down | Chi3l1 | 1.15E-25 | -0.2934397 | 0.01 | 0.778 | 2.30E-22 |
| Macrophages | Down | AABR070579 | 3.34E-37 | -0.2960359 | 0.025 | 0.889 | 6.67E-34 |
| Macrophages | Down | Clu | 3.50E-09 | -0.3117504 | 0.088 | 0.778 | 7.00E-06 |
| Macrophages | Down | Lrg1 | 2.36E-12 | -0.3143238 | 0.02 | 0.667 | 4.71E-09 |
| Macrophages | Down | Kdr | 4.81E-10 | -0.3242748 | 0.162 | 0.867 | 9.63E-07 |
| Macrophages | Down | Mcm6 | 6.52E-08 | -0.3364425 | 0.02 | 0.289 | 0.00013035 |
| Macrophages | Down | LOC1025498 | 8.34E-09 | -0.3392903 | 0.015 | 0.622 | 1.67E-05 |
| Macrophages | Down | AABR070179 | 5.26E-07 | -0.3407622 | 0.01 | 0.311 | 0.00105178 |
| Macrophages | Down | Mcf2l | 8.94E-08 | -0.3457793 | 0.211 | 0.867 | 0.00017875 |
| Macrophages | Down | Hivep3 | 5.79E-11 | -0.3561706 | 0.083 | 0.8 | 1.16E-07 |
| Macrophages | Down | Prkcz | 1.96E-10 | -0.3635518 | 0.054 | 0.733 | 3.92E-07 |
| Macrophages | Down | Pcdha13 | 4.31E-13 | -0.4083865 | 0.176 | 0.933 | 8.62E-10 |
| Macrophages | Down | Mki67 | 1.62E-13 | -0.4152457 | 0.054 | 0.667 | 3.25E-10 |
| Macrophages | Down | Mctp2 | 1.56E-07 | -0.435167 | 0.23 | 0.889 | 0.00031287 |
| Macrophages | Down | Smad6 | 3.78E-08 | -0.4389282 | 0.172 | 0.844 | 7.55E-05 |

| | | | | | | | |
|---|---|---|---|---|---|---|---|
| Macrophages | Down | Nudt4 | 1.43E-17 | -0.4455702 | 0.113 | 0.911 | 2.86E-14 |
| Macrophages | Down | Slc12a2 | 1.89E-11 | -0.4496028 | 0.167 | 0.889 | 3.78E-08 |
| Macrophages | Down | AABR070408 | 1.70E-05 | -0.4545804 | 0.088 | 0.711 | 0.03396115 |
| Macrophages | Down | Hs6st2 | 1.67E-26 | -0.4729081 | 0.049 | 0.911 | 3.35E-23 |
| Macrophages | Down | Adgrg6 | 1.99E-07 | -0.4747908 | 0.309 | 0.978 | 0.00039742 |
| Macrophages | Down | B2m | 1.29E-05 | -0.4891587 | 0.377 | 1 | 0.02578973 |
| Macrophages | Down | Optn | 8.25E-16 | -0.4922568 | 0.191 | 1 | 1.65E-12 |
| Macrophages | Down | Adamts15 | 6.62E-23 | -0.503597 | 0.132 | 1 | 1.32E-19 |
| Macrophages | Down | Lpcat2 | 1.31E-05 | -0.5128134 | 0.358 | 0.978 | 0.02629082 |
| Macrophages | Down | Dagla | 3.51E-09 | -0.5155447 | 0.132 | 0.822 | 7.02E-06 |
| Macrophages | Down | Ptafr | 2.50E-09 | -0.5372642 | 0.284 | 1 | 5.00E-06 |
| Macrophages | Down | AABR070032 | 2.00E-25 | -0.537834 | 0.015 | 0.711 | 3.99E-22 |
| Macrophages | Down | Entpd1 | 1.23E-05 | -0.56273 | 0.275 | 0.867 | 0.02459913 |
| Macrophages | Down | Lyz2 | 3.11E-08 | -0.5638899 | 0.289 | 1 | 6.22E-05 |
| Macrophages | Down | Ifit1bl | 2.60E-23 | -0.5670381 | 0.02 | 0.844 | 5.19E-20 |
| Macrophages | Down | Aldoa | 2.65E-14 | -0.5757455 | 0.206 | 0.978 | 5.31E-11 |
| Macrophages | Down | Slc43a2 | 1.25E-06 | -0.5840893 | 0.358 | 1 | 0.00250066 |
| Macrophages | Down | Apobec1 | 1.10E-07 | -0.584197 | 0.328 | 1 | 0.00022074 |
| Macrophages | Down | Alox5 | 1.54E-10 | -0.6418799 | 0.211 | 0.911 | 3.08E-07 |
| Macrophages | Down | Cd44 | 3.16E-10 | -0.6588746 | 0.289 | 1 | 6.32E-07 |
| Macrophages | Down | Kif4a | 1.18E-27 | -0.6834285 | 0.025 | 0.667 | 2.37E-24 |
| Macrophages | Down | Fth1 | 2.98E-09 | -0.8489619 | 0.255 | 0.956 | 5.95E-06 |
| Macrophages | Down | Nlrp3 | 8.32E-11 | -0.946304 | 0.275 | 1 | 1.66E-07 |
| Macrophages | Down | Nrg1 | 1.49E-11 | -1.1887044 | 0.098 | 0.822 | 2.98E-08 |
| Macrophages | Down | S100a9 | 6.79E-34 | -2.7992579 | 0.01 | 0.844 | 1.36E-30 |
| Endothelial Cells | Up | Col1a1 | 4.75E-10 | 0.65522268 | 0.444 | 1 | 9.49E-07 |
| Endothelial Cells | Up | Slco2a1 | 1.29E-52 | 0.56287232 | 0.081 | 0.577 | 2.57E-49 |
| Endothelial Cells | Up | Myl4 | 1.59E-165 | 0.54678435 | 0.115 | 1 | 3.18E-162 |
| Endothelial Cells | Up | LOC1001348 | 3.12E-142 | 0.54407274 | 0.128 | 0.98 | 6.25E-139 |
| Endothelial Cells | Up | Postn | 1.61E-75 | 0.44012686 | 0.24 | 0.998 | 3.23E-72 |
| Endothelial Cells | Up | Nppa | 1.14E-54 | 0.4211824 | 0.29 | 0.991 | 2.28E-51 |

| | | | | | | | |
|---|---|---|---|---|---|---|---|
| Endothelial Cells | Up | Scarb1 | 6.77E-36 | 0.41207532 | 0.222 | 0.858 | 1.35E-32 |
| Endothelial Cells | Up | AABR070490 | 3.95E-30 | 0.39784785 | 0.262 | 0.871 | 7.91E-27 |
| Endothelial Cells | Up | Myl7 | 1.08E-198 | 0.37222157 | 0.082 | 1 | 2.16E-195 |
| Endothelial Cells | Up | Bgn | 4.81E-81 | 0.37219349 | 0.235 | 1 | 9.63E-78 |
| Endothelial Cells | Up | Fam160a1 | 6.01E-09 | 0.35955827 | 0.282 | 0.777 | 1.20E-05 |
| Endothelial Cells | Up | Mfap5 | 7.87E-109 | 0.34270265 | 0.147 | 0.948 | 1.57E-105 |
| Endothelial Cells | Up | Nav2 | 2.92E-29 | 0.33370628 | 0.303 | 0.91 | 5.84E-26 |
| Endothelial Cells | Up | Ftl1 | 1.25E-81 | 0.33182846 | 0.172 | 0.925 | 2.50E-78 |
| Endothelial Cells | Up | Col18a1 | 3.02E-118 | 0.3288371 | 0.141 | 0.956 | 6.04E-115 |
| Endothelial Cells | Up | Apoe | 1.18E-32 | 0.32695006 | 0.18 | 0.805 | 2.35E-29 |
| Endothelial Cells | Up | Lum | 1.46E-85 | 0.32150534 | 0.153 | 0.914 | 2.91E-82 |
| Endothelial Cells | Up | Fn1 | 4.12E-77 | 0.30527044 | 0.237 | 0.986 | 8.23E-74 |
| Endothelial Cells | Up | Lyz2 | 2.08E-130 | 0.29082732 | 0.053 | 0.863 | 4.16E-127 |
| Endothelial Cells | Up | Ptn | 3.61E-67 | 0.28697721 | 0.136 | 0.855 | 7.21E-64 |
| Endothelial Cells | Up | Piezo2 | 1.36E-149 | 0.28482222 | 0.123 | 0.984 | 2.72E-146 |
| Endothelial Cells | Up | Clu | 2.18E-30 | 0.27373396 | 0.171 | 0.789 | 4.36E-27 |
| Endothelial Cells | Up | Plaur | 4.77E-64 | 0.26940844 | 0.118 | 0.832 | 9.54E-61 |
| Endothelial Cells | Up | Acta1 | 2.56E-135 | 0.26907979 | 0.107 | 0.945 | 5.12E-132 |
| Endothelial Cells | Up | Ifitm10 | 1.80E-13 | 0.2648323 | 0.094 | 0.601 | 3.59E-10 |
| Endothelial Cells | Down | Igfbp3 | 1.13E-91 | 0.2574822 | 0.066 | 0.81 | 2.27E-88 |
| Endothelial Cells | Down | Enox1 | 3.97E-55 | -0.2515376 | 0.033 | 0.642 | 7.94E-52 |
| Endothelial Cells | Down | Cacna1g | 9.81E-12 | -0.2571468 | 0.067 | 0.629 | 1.96E-08 |
| Endothelial Cells | Down | Tpm1 | 2.26E-07 | -0.2580883 | 0.591 | 0.926 | 0.00045102 |
| Endothelial Cells | Down | Cacna1c | 1.70E-10 | -0.2623867 | 0.149 | 0.676 | 3.39E-07 |
| Endothelial Cells | Down | Ngf | 5.22E-12 | -0.2652962 | 0.075 | 0.362 | 1.04E-08 |
| Endothelial Cells | Down | Pdzrn3 | 4.91E-16 | -0.2705596 | 0.288 | 0.8 | 9.82E-13 |
| Endothelial Cells | Down | Aox1 | 1.87E-20 | -0.2756494 | 0.062 | 0.349 | 3.74E-17 |
| Endothelial Cells | Down | Tnnt2 | 5.35E-07 | -0.2765181 | 0.559 | 0.899 | 0.00106967 |
| Endothelial Cells | Down | AABR070592 | 6.43E-111 | -0.2836063 | 0.092 | 0.888 | 1.29E-107 |
| Endothelial Cells | Down | Abca8a | 2.48E-14 | -0.296629 | 0.147 | 0.44 | 4.97E-11 |
| Endothelial Cells | Down | Ebf2 | 8.38E-07 | -0.3118354 | 0.203 | 0.679 | 0.00167699 |

| | | | | | | | |
|---|---|---|---|---|---|---|---|
| Endothelial Cells | Down | AABR070359 | 3.25E-10 | -0.3124765 | 0.459 | 0.895 | 6.50E-07 |
| Endothelial Cells | Down | Cacnb2 | 1.08E-09 | -0.3126883 | 0.336 | 0.794 | 2.17E-06 |
| Endothelial Cells | Down | Ldb2 | 1.10E-42 | -0.3191273 | 0.228 | 0.868 | 2.21E-39 |
| Endothelial Cells | Down | Adamts17 | 2.29E-96 | -0.3249273 | 0.079 | 0.851 | 4.59E-93 |
| Endothelial Cells | Down | Zfp385b | 1.79E-57 | -0.3291837 | 0.05 | 0.72 | 3.58E-54 |
| Endothelial Cells | Down | Irs1 | 3.28E-09 | -0.3344903 | 0.103 | 0.64 | 6.57E-06 |
| Endothelial Cells | Down | Carmil1 | 6.37E-43 | -0.3628897 | 0.092 | 0.748 | 1.27E-39 |
| Endothelial Cells | Down | Bnc2 | 1.46E-40 | -0.3698964 | 0.182 | 0.818 | 2.93E-37 |
| Endothelial Cells | Down | Pde3a | 3.32E-35 | -0.3861379 | 0.264 | 0.86 | 6.64E-32 |
| Endothelial Cells | Down | Mctp1 | 7.28E-17 | -0.3977549 | 0.106 | 0.675 | 1.46E-13 |
| Endothelial Cells | Down | Pcsk5 | 1.08E-34 | -0.4043869 | 0.168 | 0.789 | 2.16E-31 |
| Endothelial Cells | Down | Bmper | 2.63E-13 | -0.4353584 | 0.086 | 0.417 | 5.26E-10 |
| Endothelial Cells | Down | Arhgap24 | 1.38E-26 | -0.6125856 | 0.154 | 0.747 | 2.76E-23 |
| Cardiomyocytes | Up | Mfap5 | 1.26E-11 | 0.42016435 | 0.266 | 0.889 | 2.52E-08 |
| Cardiomyocytes | Up | Uqcr10 | 7.47E-08 | 0.40885201 | 0.316 | 0.836 | 0.00014949 |
| Cardiomyocytes | Up | Gpx3 | 4.86E-13 | 0.40074147 | 0.291 | 0.898 | 9.72E-10 |
| Cardiomyocytes | Up | Pik3c2g | 9.61E-11 | 0.37676152 | 0.228 | 0.797 | 1.92E-07 |
| Cardiomyocytes | Up | Postn | 3.38E-12 | 0.36765236 | 0.278 | 0.898 | 6.75E-09 |
| Cardiomyocytes | Up | Acta1 | 5.15E-10 | 0.32566786 | 0.291 | 0.879 | 1.03E-06 |
| Cardiomyocytes | Up | AABR070428 | 6.04E-15 | 0.3171442 | 0.228 | 0.872 | 1.21E-11 |
| Cardiomyocytes | Up | Fth1 | 3.34E-13 | 0.31710737 | 0.291 | 0.902 | 6.68E-10 |
| Cardiomyocytes | Up | Epha3 | 2.94E-10 | 0.31159294 | 0.139 | 0.718 | 5.88E-07 |
| Cardiomyocytes | Up | Spp1 | 9.03E-16 | 0.28879032 | 0.241 | 0.879 | 1.81E-12 |
| Cardiomyocytes | Up | Mid1 | 4.55E-13 | 0.26442853 | 0.215 | 0.839 | 9.10E-10 |
| Cardiomyocytes | Down | Mcam | 7.52E-26 | -0.2531079 | 0.165 | 0.882 | 1.50E-22 |
| Cardiomyocytes | Down | Gfpt2 | 4.66E-26 | -0.2543612 | 0.177 | 0.885 | 9.33E-23 |
| Cardiomyocytes | Down | Des | 5.57E-06 | -0.2546297 | 0.443 | 0.918 | 0.01113565 |
| Cardiomyocytes | Down | Scg2 | 2.30E-21 | -0.255873 | 0.152 | 0.807 | 4.61E-18 |
| Cardiomyocytes | Down | Cmtm8 | 3.63E-10 | -0.2562957 | 0.101 | 0.708 | 7.27E-07 |
| Cardiomyocytes | Down | AABR070294 | 3.80E-20 | -0.2567102 | 0.114 | 0.787 | 7.60E-17 |
| Cardiomyocytes | Down | Cgnl1 | 9.29E-14 | -0.2587961 | 0.215 | 0.849 | 1.86E-10 |

| | | | | | | | |
|---|---|---|---|---|---|---|---|
| Cardiomyocytes | Down | Mbp | 7.95E-15 | -0.2606549 | 0.152 | 0.764 | 1.59E-11 |
| Cardiomyocytes | Down | Zfp385b | 2.95E-09 | -0.2634394 | 0.165 | 0.721 | 5.89E-06 |
| Cardiomyocytes | Down | Xkr4 | 1.11E-09 | -0.2640602 | 0.215 | 0.78 | 2.22E-06 |
| Cardiomyocytes | Down | Myl7 | 2.02E-23 | -0.2669351 | 0.19 | 0.895 | 4.04E-20 |
| Cardiomyocytes | Down | Dok5 | 2.44E-26 | -0.2686772 | 0.177 | 0.889 | 4.88E-23 |
| Cardiomyocytes | Down | Ptprb | 7.67E-07 | -0.2703411 | 0.203 | 0.705 | 0.00153454 |
| Cardiomyocytes | Down | Prdm6 | 1.26E-17 | -0.2704952 | 0.152 | 0.823 | 2.52E-14 |
| Cardiomyocytes | Down | Uap1 | 1.47E-10 | -0.2724248 | 0.241 | 0.839 | 2.93E-07 |
| Cardiomyocytes | Down | Tnnc1 | 4.62E-06 | -0.2736495 | 0.456 | 0.931 | 0.00924801 |
| Cardiomyocytes | Down | Cd63 | 4.99E-15 | -0.2825823 | 0.203 | 0.833 | 9.98E-12 |
| Cardiomyocytes | Down | Vcan | 5.89E-19 | -0.2826604 | 0.215 | 0.902 | 1.18E-15 |
| Cardiomyocytes | Down | Ccdc148 | 7.60E-11 | -0.28296 | 0.165 | 0.731 | 1.52E-07 |
| Cardiomyocytes | Down | Plxnd1 | 3.84E-21 | -0.2830691 | 0.19 | 0.882 | 7.68E-18 |
| Cardiomyocytes | Down | Samd5 | 4.50E-15 | -0.2867757 | 0.101 | 0.738 | 9.00E-12 |
| Cardiomyocytes | Down | AABR070030 | 1.52E-10 | -0.289932 | 0.19 | 0.774 | 3.03E-07 |
| Cardiomyocytes | Down | Fat3 | 2.10E-22 | -0.2900148 | 0.19 | 0.879 | 4.19E-19 |
| Cardiomyocytes | Down | AABR070546 | 9.05E-09 | -0.2902228 | 0.165 | 0.721 | 1.81E-05 |
| Cardiomyocytes | Down | Adamts2 | 1.43E-15 | -0.2932868 | 0.228 | 0.885 | 2.85E-12 |
| Cardiomyocytes | Down | Camk1d | 1.40E-17 | -0.2932873 | 0.177 | 0.83 | 2.79E-14 |
| Cardiomyocytes | Down | Frmd5 | 2.10E-06 | -0.29334 | 0.418 | 0.925 | 0.00420163 |
| Cardiomyocytes | Down | Dpt | 7.99E-18 | -0.2991089 | 0.241 | 0.902 | 1.60E-14 |
| Cardiomyocytes | Down | Ppp1r16b | 1.09E-17 | -0.3013688 | 0.165 | 0.803 | 2.19E-14 |
| Cardiomyocytes | Down | AABR070490 | 4.54E-22 | -0.3021732 | 0.203 | 0.889 | 9.08E-19 |
| Cardiomyocytes | Down | Inpp5d | 5.91E-21 | -0.3034149 | 0.177 | 0.892 | 1.18E-17 |
| Cardiomyocytes | Down | Tmem38a | 1.18E-05 | -0.3040408 | 0.354 | 0.813 | 0.02367126 |
| Cardiomyocytes | Down | Kcnq3 | 4.60E-17 | -0.3077729 | 0.165 | 0.856 | 9.19E-14 |
| Cardiomyocytes | Down | Serf2 | 1.80E-07 | -0.3102391 | 0.38 | 0.921 | 0.00035961 |
| Cardiomyocytes | Down | Tbc1d10c | 1.10E-11 | -0.310958 | 0.139 | 0.757 | 2.20E-08 |
| Cardiomyocytes | Down | Abca1 | 7.25E-08 | -0.3114812 | 0.241 | 0.8 | 0.00014494 |
| Cardiomyocytes | Down | Pcdh17 | 2.32E-22 | -0.3116178 | 0.165 | 0.862 | 4.65E-19 |
| Cardiomyocytes | Down | Gda | 6.34E-15 | -0.3118829 | 0.241 | 0.895 | 1.27E-11 |

| | | | | | | | |
|---|---|---|---|---|---|---|---|
| Cardiomyocytes | Down | Tox | 2.13E-06 | -0.3124177 | 0.367 | 0.856 | 0.00426375 |
| Cardiomyocytes | Down | Med12l | 3.00E-08 | -0.3144326 | 0.101 | 0.662 | 6.01E-05 |
| Cardiomyocytes | Down | Adamtsl2 | 6.42E-15 | -0.3267027 | 0.278 | 0.898 | 1.28E-11 |
| Cardiomyocytes | Down | Ankrd6 | 2.74E-11 | -0.3290761 | 0.304 | 0.892 | 5.48E-08 |
| Cardiomyocytes | Down | Tpt1 | 2.20E-12 | -0.3333899 | 0.278 | 0.872 | 4.41E-09 |
| Cardiomyocytes | Down | Cald1 | 1.20E-06 | -0.3334562 | 0.266 | 0.761 | 0.00240024 |
| Cardiomyocytes | Down | Kcnj12 | 6.06E-18 | -0.3339612 | 0.203 | 0.856 | 1.21E-14 |
| Cardiomyocytes | Down | AABR070651 | 8.48E-15 | -0.3346045 | 0.165 | 0.793 | 1.70E-11 |
| Cardiomyocytes | Down | Ghitm | 1.03E-19 | -0.3373701 | 0.203 | 0.882 | 2.06E-16 |
| Cardiomyocytes | Down | Slc9a3r2 | 3.93E-21 | -0.3383515 | 0.177 | 0.875 | 7.86E-18 |
| Cardiomyocytes | Down | Atp5me | 1.35E-13 | -0.3398421 | 0.291 | 0.885 | 2.71E-10 |
| Cardiomyocytes | Down | Enpp1 | 4.05E-17 | -0.3465583 | 0.152 | 0.816 | 8.10E-14 |
| Cardiomyocytes | Down | Tenm3 | 3.81E-13 | -0.3471568 | 0.139 | 0.757 | 7.62E-10 |
| Cardiomyocytes | Down | Csrp2 | 1.20E-06 | -0.3507266 | 0.139 | 0.656 | 0.00239775 |
| Cardiomyocytes | Down | Slc25a13 | 9.09E-09 | -0.3530832 | 0.392 | 0.931 | 1.82E-05 |
| Cardiomyocytes | Down | Mcf2l | 1.24E-05 | -0.3554331 | 0.203 | 0.685 | 0.02474271 |
| Cardiomyocytes | Down | Fgf13 | 3.44E-08 | -0.358595 | 0.38 | 0.925 | 6.88E-05 |
| Cardiomyocytes | Down | Tmtc2 | 4.19E-11 | -0.360193 | 0.278 | 0.875 | 8.38E-08 |
| Cardiomyocytes | Down | Pla2g5 | 3.21E-11 | -0.3609892 | 0.316 | 0.885 | 6.43E-08 |
| Cardiomyocytes | Down | Sema3a | 6.52E-15 | -0.3626955 | 0.165 | 0.79 | 1.30E-11 |
| Cardiomyocytes | Down | Ikzf2 | 1.40E-09 | -0.3636869 | 0.228 | 0.77 | 2.80E-06 |
| Cardiomyocytes | Down | Pola2 | 2.65E-10 | -0.3647638 | 0.329 | 0.925 | 5.30E-07 |
| Cardiomyocytes | Down | Ppm1l | 1.08E-05 | -0.3666313 | 0.291 | 0.751 | 0.02151721 |
| Cardiomyocytes | Down | Ccdc80 | 3.82E-11 | -0.3709744 | 0.304 | 0.911 | 7.65E-08 |
| Cardiomyocytes | Down | Ubc | 5.01E-11 | -0.3757287 | 0.316 | 0.872 | 1.00E-07 |
| Cardiomyocytes | Down | Prkch | 3.51E-15 | -0.3760235 | 0.215 | 0.839 | 7.03E-12 |
| Cardiomyocytes | Down | Hmcn1 | 9.37E-06 | -0.380373 | 0.291 | 0.77 | 0.01874577 |
| Cardiomyocytes | Down | Crispld2 | 1.67E-20 | -0.3818709 | 0.203 | 0.885 | 3.34E-17 |
| Cardiomyocytes | Down | Mmp2 | 5.00E-27 | -0.3866162 | 0.165 | 0.892 | 1.00E-23 |
| Cardiomyocytes | Down | Ar | 1.38E-10 | -0.3913177 | 0.152 | 0.728 | 2.77E-07 |
| Cardiomyocytes | Down | Csrp3 | 2.62E-07 | -0.3931235 | 0.418 | 0.892 | 0.00052416 |

| | | | | | | | |
|---|---|---|---|---|---|---|---|
| Cardiomyocytes | Down | Uqcrq | 4.40E-16 | -0.4065785 | 0.253 | 0.898 | 8.79E-13 |
| Cardiomyocytes | Down | Abi3bp | 9.97E-14 | -0.4094273 | 0.241 | 0.889 | 1.99E-10 |
| Cardiomyocytes | Down | Ndufa4 | 9.70E-08 | -0.4112907 | 0.241 | 0.774 | 0.00019394 |
| Cardiomyocytes | Down | Epha4 | 4.37E-22 | -0.4134728 | 0.19 | 0.879 | 8.75E-19 |
| Cardiomyocytes | Down | Eda | 1.11E-15 | -0.4138832 | 0.291 | 0.895 | 2.21E-12 |
| Cardiomyocytes | Down | Dapk2 | 5.09E-18 | -0.4157659 | 0.203 | 0.836 | 1.02E-14 |
| Cardiomyocytes | Down | Fli1 | 1.65E-11 | -0.4176975 | 0.291 | 0.885 | 3.30E-08 |
| Cardiomyocytes | Down | Ubash3b | 3.52E-09 | -0.4197765 | 0.038 | 0.63 | 7.04E-06 |
| Cardiomyocytes | Down | Masp1 | 1.23E-07 | -0.4206679 | 0.215 | 0.767 | 0.0002454 |
| Cardiomyocytes | Down | Fhl1 | 4.06E-08 | -0.4210974 | 0.304 | 0.826 | 8.11E-05 |
| Cardiomyocytes | Down | Csdc2 | 1.23E-09 | -0.4270772 | 0.19 | 0.751 | 2.47E-06 |
| Cardiomyocytes | Down | LOC687780 | 6.95E-16 | -0.4306192 | 0.203 | 0.866 | 1.39E-12 |
| Cardiomyocytes | Down | Slc12a2 | 1.02E-07 | -0.4366447 | 0.19 | 0.725 | 0.00020361 |
| Cardiomyocytes | Down | Fbn1 | 3.84E-11 | -0.4410277 | 0.342 | 0.931 | 7.67E-08 |
| Cardiomyocytes | Down | Sned1 | 8.69E-20 | -0.4419781 | 0.228 | 0.892 | 1.74E-16 |
| Cardiomyocytes | Down | Pkhd1l1 | 1.25E-16 | -0.443319 | 0.139 | 0.793 | 2.50E-13 |
| Cardiomyocytes | Down | Frmd4b | 3.23E-15 | -0.4503533 | 0.203 | 0.839 | 6.46E-12 |
| Cardiomyocytes | Down | Meox2 | 8.90E-18 | -0.4548855 | 0.177 | 0.833 | 1.78E-14 |
| Cardiomyocytes | Down | Atp5po | 5.13E-08 | -0.4554011 | 0.291 | 0.8 | 0.00010255 |
| Cardiomyocytes | Down | Cox6a2 | 2.33E-10 | -0.4577223 | 0.342 | 0.905 | 4.67E-07 |
| Cardiomyocytes | Down | Col24a1 | 1.81E-14 | -0.4632455 | 0.165 | 0.803 | 3.61E-11 |
| Cardiomyocytes | Down | AC111831.1 | 1.68E-05 | -0.4660361 | 0.127 | 0.58 | 0.03364772 |
| Cardiomyocytes | Down | Slit2 | 9.09E-19 | -0.4715594 | 0.165 | 0.833 | 1.82E-15 |
| Cardiomyocytes | Down | Slc9a9 | 2.06E-07 | -0.472187 | 0.354 | 0.862 | 0.0004129 |
| Cardiomyocytes | Down | Myo1b | 2.52E-11 | -0.4838693 | 0.278 | 0.862 | 5.03E-08 |
| Cardiomyocytes | Down | Gsn | 7.84E-11 | -0.4853471 | 0.266 | 0.833 | 1.57E-07 |
| Cardiomyocytes | Down | Pgm5 | 1.48E-06 | -0.4961277 | 0.266 | 0.751 | 0.00296666 |
| Cardiomyocytes | Down | Baiap2l1 | 1.74E-16 | -0.5060308 | 0.266 | 0.908 | 3.48E-13 |
| Cardiomyocytes | Down | Tbx5 | 1.11E-09 | -0.5064683 | 0.291 | 0.823 | 2.23E-06 |
| Cardiomyocytes | Down | Ndufa11 | 3.85E-21 | -0.5267912 | 0.241 | 0.908 | 7.69E-18 |
| Cardiomyocytes | Down | Slc7a1 | 1.84E-08 | -0.5348073 | 0.329 | 0.902 | 3.68E-05 |

| | | | | | | | |
|---|---|---|---|---|---|---|---|
| Cardiomyocytes | Down | Nppb | 2.91E-10 | -0.5350223 | 0.354 | 0.918 | 5.83E-07 |
| Cardiomyocytes | Down | Chst15 | 1.90E-07 | -0.5433455 | 0.392 | 0.915 | 0.0003809 |
| Cardiomyocytes | Down | Vwf | 4.63E-09 | -0.5475132 | 0.139 | 0.705 | 9.25E-06 |
| Cardiomyocytes | Down | Lyn | 2.44E-22 | -0.5532793 | 0.241 | 0.905 | 4.89E-19 |
| Cardiomyocytes | Down | Cpq | 4.93E-15 | -0.5541179 | 0.291 | 0.905 | 9.87E-12 |
| Cardiomyocytes | Down | Twf2 | 9.58E-09 | -0.5555585 | 0.38 | 0.905 | 1.92E-05 |
| Cardiomyocytes | Down | Arhgap24 | 4.56E-08 | -0.5679515 | 0.278 | 0.777 | 9.11E-05 |
| Cardiomyocytes | Down | Il1r1 | 1.48E-13 | -0.5735953 | 0.19 | 0.813 | 2.96E-10 |
| Cardiomyocytes | Down | Col8a1 | 1.02E-06 | -0.5831109 | 0.342 | 0.849 | 0.00203654 |
| Cardiomyocytes | Down | Flt1 | 5.68E-13 | -0.5969546 | 0.266 | 0.882 | 1.14E-09 |
| Cardiomyocytes | Down | Neb | 8.49E-18 | -0.6373248 | 0.177 | 0.83 | 1.70E-14 |
| Cardiomyocytes | Down | LOC1083526 | 4.85E-23 | -0.6512131 | 0.203 | 0.908 | 9.69E-20 |
| Cardiomyocytes | Down | Nppa | 1.78E-12 | -0.6579202 | 0.354 | 0.931 | 3.57E-09 |
| Cardiomyocytes | Down | Timp3 | 4.44E-07 | -0.6665574 | 0.392 | 0.862 | 0.00088808 |
| Cardiomyocytes | Down | Tmsb4x | 1.06E-14 | -0.6769858 | 0.291 | 0.911 | 2.13E-11 |
| Cardiomyocytes | Down | AABR070492 | 3.47E-11 | -0.6996113 | 0.215 | 0.797 | 6.93E-08 |
| Cardiomyocytes | Down | Tec | 1.76E-10 | -0.7032537 | 0.152 | 0.744 | 3.51E-07 |
| Cardiomyocytes | Down | Erbb4 | 1.11E-05 | -0.7078453 | 0.494 | 0.931 | 0.02210356 |
| Cardiomyocytes | Down | Ncam1 | 6.54E-18 | -0.7147299 | 0.228 | 0.892 | 1.31E-14 |
| Cardiomyocytes | Down | Fgf12 | 6.06E-07 | -0.7419282 | 0.354 | 0.823 | 0.00121203 |
| Cardiomyocytes | Down | Itpkb | 8.11E-09 | -0.7612317 | 0.228 | 0.751 | 1.62E-05 |
| Cardiomyocytes | Down | Zeb2 | 2.15E-10 | -0.837718 | 0.304 | 0.859 | 4.30E-07 |
| Cardiomyocytes | Down | Col3a1 | 2.43E-07 | -0.8574336 | 0.468 | 0.961 | 0.00048651 |
| Cardiomyocytes | Down | Dsg2 | 6.63E-19 | -0.9014969 | 0.228 | 0.921 | 1.33E-15 |
| Cardiomyocytes | Down | Coq8a | 8.17E-10 | -0.9192441 | 0.405 | 0.918 | 1.63E-06 |
| Cardiomyocytes | Down | Col4a3 | 5.04E-10 | -0.9222834 | 0.253 | 0.833 | 1.01E-06 |
| Cardiomyocytes | Down | Slco5a1 | 2.12E-11 | -1.1761582 | 0.405 | 0.905 | 4.23E-08 |
| Fibroblasts | Up | Myl4 | 2.13E-75 | 0.76451092 | 0.131 | 1 | 4.27E-72 |
| Fibroblasts | Up | Nrk | 4.25E-07 | 0.69251813 | 0.149 | 0.702 | 0.00085024 |
| Fibroblasts | Up | Ltbp2 | 9.25E-61 | 0.54836586 | 0.096 | 0.915 | 1.85E-57 |
| Fibroblasts | Up | Loxl1 | 1.37E-12 | 0.54585205 | 0.385 | 0.99 | 2.74E-09 |

| | | | | | | | |
|---|---|---|---|---|---|---|---|
| Fibroblasts | Up | Ccn2 | 1.30E-64 | 0.54491954 | 0.161 | 1 | 2.61E-61 |
| Fibroblasts | Up | Adamts14 | 1.32E-35 | 0.50639245 | 0.188 | 0.915 | 2.64E-32 |
| Fibroblasts | Up | Egr1 | 2.20E-62 | 0.5032723 | 0.081 | 0.901 | 4.39E-59 |
| Fibroblasts | Up | Apoe | 1.00E-77 | 0.49382372 | 0.125 | 0.998 | 2.00E-74 |
| Fibroblasts | Up | Adamts17 | 8.61E-20 | 0.47136106 | 0.346 | 0.992 | 1.72E-16 |
| Fibroblasts | Up | Npas2 | 2.41E-12 | 0.46944686 | 0.388 | 1 | 4.83E-09 |
| Fibroblasts | Up | LOC1001348 | 3.24E-66 | 0.4658193 | 0.081 | 0.917 | 6.48E-63 |
| Fibroblasts | Up | Lgals1 | 1.99E-66 | 0.4614307 | 0.155 | 0.998 | 3.99E-63 |
| Fibroblasts | Up | Nppa | 7.05E-54 | 0.45912568 | 0.194 | 0.996 | 1.41E-50 |
| Fibroblasts | Up | Crip1 | 5.50E-54 | 0.41967452 | 0.149 | 0.948 | 1.10E-50 |
| Fibroblasts | Up | Fth1 | 1.69E-68 | 0.41509578 | 0.143 | 0.99 | 3.39E-65 |
| Fibroblasts | Up | Myl7 | 2.15E-96 | 0.40362223 | 0.084 | 1 | 4.29E-93 |
| Fibroblasts | Up | Sv2c | 6.28E-84 | 0.39362397 | 0.075 | 0.959 | 1.26E-80 |
| Fibroblasts | Up | Ehd4 | 1.81E-25 | 0.38936107 | 0.251 | 0.925 | 3.62E-22 |
| Fibroblasts | Up | AABR070490 | 6.65E-70 | 0.38726038 | 0.14 | 0.99 | 1.33E-66 |
| Fibroblasts | Up | Myl3 | 2.49E-24 | 0.37370307 | 0.304 | 0.967 | 4.98E-21 |
| Fibroblasts | Up | Ntrk3 | 3.48E-38 | 0.36897701 | 0.23 | 0.967 | 6.97E-35 |
| Fibroblasts | Up | Tex22 | 2.29E-84 | 0.35445447 | 0.06 | 0.944 | 4.58E-81 |
| Fibroblasts | Up | LOC1003644 | 8.94E-57 | 0.35388833 | 0.182 | 0.994 | 1.79E-53 |
| Fibroblasts | Up | Galnt16 | 1.40E-24 | 0.34764112 | 0.218 | 0.886 | 2.80E-21 |
| Fibroblasts | Up | Nbl1 | 2.88E-80 | 0.34580245 | 0.11 | 0.988 | 5.76E-77 |
| Fibroblasts | Up | LOC1083526 | 5.58E-23 | 0.3426262 | 0.09 | 0.764 | 1.12E-19 |
| Fibroblasts | Up | Ckm | 1.52E-48 | 0.3290126 | 0.122 | 0.901 | 3.04E-45 |
| Fibroblasts | Up | Myl2 | 1.68E-20 | 0.32729841 | 0.325 | 0.973 | 3.37E-17 |
| Fibroblasts | Up | Mb | 4.75E-07 | 0.32688228 | 0.463 | 0.981 | 0.00095055 |
| Fibroblasts | Up | Epha4 | 7.49E-23 | 0.32522136 | 0.072 | 0.749 | 1.50E-19 |
| Fibroblasts | Up | Lsamp | 6.26E-70 | 0.3230944 | 0.063 | 0.907 | 1.25E-66 |
| Fibroblasts | Up | Actc1 | 1.64E-34 | 0.31782004 | 0.257 | 0.977 | 3.27E-31 |
| Fibroblasts | Up | Sgca | 2.78E-24 | 0.31781071 | 0.084 | 0.766 | 5.55E-21 |
| Fibroblasts | Up | Plaur | 1.01E-41 | 0.31714491 | 0.081 | 0.836 | 2.02E-38 |
| Fibroblasts | Up | Agtr1a | 5.95E-48 | 0.31441763 | 0.072 | 0.851 | 1.19E-44 |

| | | | | | | | |
|---|---|---|---|---|---|---|---|
| Fibroblasts | Up | Cox6a2 | 9.10E-33 | 0.31410028 | 0.179 | 0.892 | 1.82E-29 |
| Fibroblasts | Up | Gfod1 | 1.38E-43 | 0.31389722 | 0.125 | 0.884 | 2.75E-40 |
| Fibroblasts | Up | Tpt1 | 3.76E-59 | 0.31113973 | 0.179 | 0.996 | 7.51E-56 |
| Fibroblasts | Up | Cacna1e | 1.86E-49 | 0.31099141 | 0.078 | 0.861 | 3.73E-46 |
| Fibroblasts | Up | Trdn | 1.41E-45 | 0.31098431 | 0.149 | 0.915 | 2.81E-42 |
| Fibroblasts | Up | Mfap5 | 4.61E-26 | 0.30668388 | 0.313 | 0.992 | 9.22E-23 |
| Fibroblasts | Up | Cox8b | 2.78E-39 | 0.29522403 | 0.23 | 0.973 | 5.55E-36 |
| Fibroblasts | Up | Optn | 1.95E-07 | 0.28768277 | 0.122 | 0.683 | 0.00038967 |
| Fibroblasts | Up | Col27a1 | 5.98E-45 | 0.28279511 | 0.113 | 0.878 | 1.20E-41 |
| Fibroblasts | Up | Fam20a | 1.99E-09 | 0.28101135 | 0.385 | 0.923 | 3.98E-06 |
| Fibroblasts | Up | Uqcrq | 1.20E-15 | 0.27883846 | 0.084 | 0.718 | 2.41E-12 |
| Fibroblasts | Up | Dmpk | 1.31E-34 | 0.27728752 | 0.272 | 0.998 | 2.62E-31 |
| Fibroblasts | Up | Atp5f1e | 2.34E-89 | 0.27701813 | 0.09 | 0.988 | 4.68E-86 |
| Fibroblasts | Up | Zfp385b | 6.61E-08 | 0.2765814 | 0.296 | 0.816 | 0.00013218 |
| Fibroblasts | Up | LOC685963 | 2.07E-55 | 0.27611315 | 0.054 | 0.857 | 4.13E-52 |
| Fibroblasts | Up | Adgrd1 | 2.15E-07 | 0.27166503 | 0.316 | 0.83 | 0.00042947 |
| Fibroblasts | Up | Cd63 | 3.91E-34 | 0.26802191 | 0.104 | 0.828 | 7.82E-31 |
| Fibroblasts | Up | Trim54 | 7.49E-06 | 0.2654525 | 0.057 | 0.435 | 0.01497458 |
| Fibroblasts | Up | Pgam2 | 1.43E-75 | 0.26499948 | 0.104 | 0.967 | 2.87E-72 |
| Fibroblasts | Up | Cacna1g | 3.58E-15 | 0.26414686 | 0.128 | 0.749 | 7.17E-12 |
| Fibroblasts | Up | Cox6a1 | 1.36E-92 | 0.26413841 | 0.063 | 0.963 | 2.72E-89 |
| Fibroblasts | Up | Ddah1 | 9.00E-73 | 0.26354634 | 0.051 | 0.89 | 1.80E-69 |
| Fibroblasts | Up | Bgn | 2.53E-34 | 0.26237422 | 0.272 | 1 | 5.05E-31 |
| Fibroblasts | Up | Gask1b | 1.68E-05 | 0.26131456 | 0.367 | 0.845 | 0.03367346 |
| Fibroblasts | Up | Ntrk2 | 3.67E-36 | 0.26129599 | 0.099 | 0.83 | 7.34E-33 |
| Fibroblasts | Up | Lmod1 | 3.78E-44 | 0.26112964 | 0.069 | 0.834 | 7.56E-41 |
| Fibroblasts | Up | Specc1 | 3.16E-51 | 0.26092334 | 0.149 | 0.936 | 6.32E-48 |
| Fibroblasts | Up | Col16a1 | 1.33E-28 | 0.26052178 | 0.14 | 0.834 | 2.66E-25 |
| Fibroblasts | Up | Slc1a7 | 6.47E-16 | 0.25866617 | 0.101 | 0.373 | 1.29E-12 |
| Fibroblasts | Up | Cilp | 2.74E-82 | 0.25443699 | 0.039 | 0.886 | 5.49E-79 |
| Fibroblasts | Up | Smyd1 | 3.18E-93 | 0.25364854 | 0.063 | 0.969 | 6.37E-90 |

| | | | | | | | |
|---|---|---|---|---|---|---|---|
| Fibroblasts | Up | Prag1 | 1.15E-07 | 0.25208153 | 0.045 | 0.398 | 0.0002308 |
| Fibroblasts | Up | Adam12 | 1.87E-60 | 0.25182748 | 0.173 | 0.998 | 3.74E-57 |
| Fibroblasts | Up | Col24a1 | 4.00E-84 | 0.25074194 | 0.087 | 0.971 | 7.99E-81 |
| Fibroblasts | Down | Obscn | 1.00E-39 | -0.2502966 | 0.087 | 0.83 | 2.01E-36 |
| Fibroblasts | Down | AABR070070 | 2.41E-12 | -0.2517278 | 0.072 | 0.366 | 4.81E-09 |
| Fibroblasts | Down | Palm2 | 1.95E-44 | -0.2524242 | 0.215 | 0.963 | 3.90E-41 |
| Fibroblasts | Down | Prkch | 1.61E-14 | -0.2539502 | 0.066 | 0.696 | 3.22E-11 |
| Fibroblasts | Down | Syn3 | 6.29E-31 | -0.2547627 | 0.104 | 0.81 | 1.26E-27 |
| Fibroblasts | Down | Kcnn3 | 7.85E-18 | -0.2574014 | 0.388 | 0.986 | 1.57E-14 |
| Fibroblasts | Down | Fgf1 | 9.46E-62 | -0.2608587 | 0.045 | 0.867 | 1.89E-58 |
| Fibroblasts | Down | Pcsk5 | 2.48E-34 | -0.2632388 | 0.301 | 1 | 4.96E-31 |
| Fibroblasts | Down | Boc | 1.02E-12 | -0.2637706 | 0.46 | 0.994 | 2.04E-09 |
| Fibroblasts | Down | Mitf | 5.16E-20 | -0.2664282 | 0.275 | 0.882 | 1.03E-16 |
| Fibroblasts | Down | Ppp1r16b | 3.58E-73 | -0.2676823 | 0.024 | 0.88 | 7.16E-70 |
| Fibroblasts | Down | Palmd | 2.19E-32 | -0.2727481 | 0.039 | 0.764 | 4.39E-29 |
| Fibroblasts | Down | Myrip | 2.08E-15 | -0.2800219 | 0.027 | 0.675 | 4.17E-12 |
| Fibroblasts | Down | Dpt | 2.36E-06 | -0.283093 | 0.501 | 0.934 | 0.004726 |
| Fibroblasts | Down | Aldh1a1 | 9.77E-06 | -0.2839193 | 0.096 | 0.644 | 0.01954679 |
| Fibroblasts | Down | Plxnd1 | 5.18E-19 | -0.2854037 | 0.122 | 0.762 | 1.04E-15 |
| Fibroblasts | Down | Chsy3 | 9.37E-13 | -0.2875412 | 0.454 | 0.979 | 1.87E-09 |
| Fibroblasts | Down | Fgf10 | 2.24E-06 | -0.2901566 | 0.093 | 0.648 | 0.00447464 |
| Fibroblasts | Down | Tnik | 7.39E-11 | -0.2914678 | 0.29 | 0.822 | 1.48E-07 |
| Fibroblasts | Down | Oxr1 | 6.88E-16 | -0.2920066 | 0.203 | 0.797 | 1.38E-12 |
| Fibroblasts | Down | Ldlrad3 | 5.66E-06 | -0.2954868 | 0.248 | 0.733 | 0.0113133 |
| Fibroblasts | Down | Tnnt3 | 9.04E-18 | -0.3021202 | 0.376 | 0.946 | 1.81E-14 |
| Fibroblasts | Down | Nxn | 2.42E-11 | -0.3024734 | 0.269 | 0.805 | 4.84E-08 |
| Fibroblasts | Down | Adamts9 | 2.80E-95 | -0.3032323 | 0.048 | 0.957 | 5.59E-92 |
| Fibroblasts | Down | Adgrf5 | 7.55E-11 | -0.3048474 | 0.113 | 0.406 | 1.51E-07 |
| Fibroblasts | Down | Nav3 | 8.52E-36 | -0.3071288 | 0.2 | 0.907 | 1.70E-32 |
| Fibroblasts | Down | Adgrl4 | 9.69E-09 | -0.3085331 | 0.081 | 0.663 | 1.94E-05 |
| Fibroblasts | Down | Pdlim5 | 1.32E-14 | -0.3098282 | 0.403 | 0.938 | 2.64E-11 |

| | | | | | | | |
|---|---|---|---|---|---|---|---|
| Fibroblasts | Down | Dip2c | 4.69E-17 | -0.3098488 | 0.215 | 0.814 | 9.37E-14 |
| Fibroblasts | Down | Pdgfd | 1.62E-61 | -0.3316946 | 0.155 | 0.969 | 3.24E-58 |
| Fibroblasts | Down | Dsp | 6.53E-15 | -0.3472864 | 0.06 | 0.362 | 1.31E-11 |
| Fibroblasts | Down | Egfr | 4.77E-07 | -0.3559446 | 0.493 | 0.909 | 0.00095374 |
| Fibroblasts | Down | Col6a2 | 6.58E-06 | -0.3619314 | 0.573 | 0.992 | 0.01315546 |
| Fibroblasts | Down | Tbx18 | 3.38E-56 | -0.3620811 | 0.185 | 0.979 | 6.77E-53 |
| Fibroblasts | Down | Vwf | 1.16E-09 | -0.3627663 | 0.042 | 0.393 | 2.32E-06 |
| Fibroblasts | Down | Nt5e | 4.06E-65 | -0.3750998 | 0.143 | 0.969 | 8.11E-62 |
| Fibroblasts | Down | Mrvi1 | 3.82E-46 | -0.3776245 | 0.09 | 0.855 | 7.64E-43 |
| Fibroblasts | Down | AABR070592 | 1.12E-17 | -0.3901281 | 0.355 | 0.925 | 2.23E-14 |
| Fibroblasts | Down | Ntn1 | 8.67E-20 | -0.3999239 | 0.307 | 0.901 | 1.73E-16 |
| Fibroblasts | Down | Slfn13 | 2.88E-61 | -0.4077733 | 0.158 | 0.967 | 5.77E-58 |
| Fibroblasts | Down | Colec12 | 3.86E-19 | -0.4152715 | 0.43 | 0.99 | 7.72E-16 |
| Fibroblasts | Down | AABR070076 | 4.96E-35 | -0.4165601 | 0.012 | 0.754 | 9.92E-32 |
| Fibroblasts | Down | Gli2 | 1.03E-09 | -0.4218478 | 0.481 | 0.903 | 2.05E-06 |
| Fibroblasts | Down | Chn1 | 4.95E-70 | -0.4342591 | 0.104 | 0.944 | 9.90E-67 |
| Fibroblasts | Down | Ltbp1 | 5.85E-07 | -0.4345156 | 0.472 | 0.865 | 0.00117071 |
| Fibroblasts | Down | Prodh1 | 4.57E-20 | -0.4384631 | 0.039 | 0.708 | 9.14E-17 |
| Fibroblasts | Down | Shank3 | 1.82E-13 | -0.4433125 | 0.14 | 0.735 | 3.65E-10 |
| Fibroblasts | Down | Bmp6 | 3.76E-07 | -0.4519777 | 0.46 | 0.882 | 0.00075224 |
| Fibroblasts | Down | Efcc1 | 1.00E-05 | -0.4530281 | 0.397 | 0.814 | 0.02008148 |
| Fibroblasts | Down | Ccdc85a | 1.02E-11 | -0.4782374 | 0.09 | 0.689 | 2.05E-08 |
| Fibroblasts | Down | Adamts5 | 5.27E-11 | -0.4927998 | 0.313 | 0.822 | 1.05E-07 |
| Fibroblasts | Down | Cyyr1 | 3.43E-13 | -0.4955998 | 0.116 | 0.385 | 6.87E-10 |
| Fibroblasts | Down | Tbx20 | 6.64E-17 | -0.6131448 | 0.454 | 0.992 | 1.33E-13 |
