## Supplemental Table 4 for "Uncovering the Regional and Cell Specific Bioactivity of Injectable Extracellular Matrix Biomaterials in Myocardial Infarction through Spatial and Single Nucleus Transcriptomics"

| Supplementary Table 4. Differentially Expressed Genes Spatially Comparing the Remote Zone of ECM Treated Hearts vs. Saline Treated Hearts in a Subacute MI Model | | | | | | | |
|---|---|---|---|---|---|---|---|
| **Spatial Area** | **Direction** | **Gene** | **p_val** | **avg_log2FC** | **pct.1** | **pct.2** | **p_val_adj** |
| ECM Remote Zone | Up | AABR07012711.1 | 0 | 1.80313401 | 0.022 | 0.366 | 0 |
| ECM Remote Zone | Up | Reg3g | 0 | 0.81801563 | 0.513 | 0.932 | 0 |
| ECM Remote Zone | Up | Ino80d | 2.80E-285 | 0.79216175 | 0.425 | 0.721 | 5.59E-282 |
| ECM Remote Zone | Up | Gprin3 | 3.01E-84 | 0.74442644 | 0.387 | 0.608 | 6.02E-81 |
| ECM Remote Zone | Up | Tdp1 | 0 | 0.71131435 | 0.293 | 0.54 | 0 |
| ECM Remote Zone | Up | Slc9b1 | 1.46E-107 | 0.703751 | 0.142 | 0.405 | 2.92E-104 |
| ECM Remote Zone | Up | Fam13a | 1.52E-232 | 0.6584811 | 0.521 | 0.744 | 3.04E-229 |
| ECM Remote Zone | Up | Col9a2 | 0 | 0.65584602 | 0.127 | 0.532 | 0 |

| | | | | | | | |
|---|---|---|---|---|---|---|---|
| ECM Remote Zone | Up | Agr2 | 0 | 0.61357445 | 0.545 | 0.858 | 0 |
| ECM Remote Zone | Up | Bcl11b | 0 | 0.61009885 | 0.254 | 0.516 | 0 |
| ECM Remote Zone | Up | Fscn1 | 0 | 0.56438453 | 0.03 | 0.26 | 0 |
| ECM Remote Zone | Up | S100a8 | 0 | 0.52734312 | 0.34 | 0.677 | 0 |
| ECM Remote Zone | Up | Slpi | 4.86E-286 | 0.50358674 | 0.502 | 0.838 | 9.72E-283 |
| ECM Remote Zone | Up | Slc8a2 | 1.30E-98 | 0.44899931 | 0.077 | 0.371 | 2.61E-95 |
| ECM Remote Zone | Up | Ctrb1 | 6.97E-30 | 0.44237566 | 0.418 | 0.618 | 1.39E-26 |
| ECM Remote Zone | Up | Ccl4 | 3.57E-220 | 0.43858843 | 0.504 | 0.796 | 7.14E-217 |
| ECM Remote Zone | Up | LOC685067 | 1.04E-243 | 0.43175652 | 0.534 | 0.863 | 2.07E-240 |

| | | | | | | | |
|---|---|---|---|---|---|---|---|
| ECM Remote Zone | Up | Xcl1 | 5.38E-201 | 0.42937548 | 0.463 | 0.733 | 1.08E-197 |
| ECM Remote Zone | Up | Il1r2 | 1.26E-117 | 0.42177959 | 0.452 | 0.665 | 2.52E-114 |
| ECM Remote Zone | Up | Rassf5 | 1.13E-235 | 0.41062988 | 0.39 | 0.598 | 2.27E-232 |
| ECM Remote Zone | Up | Cass4 | 6.66E-263 | 0.38663366 | 0.322 | 0.53 | 1.33E-259 |
| ECM Remote Zone | Up | Zbp1 | 1.14E-298 | 0.38061444 | 0.529 | 0.793 | 2.28E-295 |
| ECM Remote Zone | Up | Tbc1d7 | 0 | 0.37668489 | 0.43 | 0.812 | 0 |
| ECM Remote Zone | Up | Angptl4 | 0 | 0.37236096 | 0.282 | 0.531 | 0 |
| ECM Remote Zone | Up | Rec114 | 6.17E-268 | 0.33434274 | 0.551 | 0.924 | 1.23E-264 |
| ECM Remote Zone | Up | AC111831.1 | 1.71E-185 | 0.33353313 | 0.536 | 0.769 | 3.42E-182 |

| | | | | | | | |
|---|---|---|---|---|---|---|---|
| ECM Remote Zone | Up | Oas1a | 2.16E-80 | 0.3310651 | 0.625 | 0.978 | 4.32E-77 |
| ECM Remote Zone | Up | AABR07052502.1 | 2.04E-178 | 0.32483959 | 0.143 | 0.53 | 4.07E-175 |
| ECM Remote Zone | Up | Mmp24 | 2.41E-105 | 0.29564971 | 0.251 | 0.5 | 4.83E-102 |
| ECM Remote Zone | Up | Gata3 | 6.85E-246 | 0.28336522 | 0.498 | 0.753 | 1.37E-242 |
| ECM Remote Zone | Up | Gzma | 4.65E-302 | 0.27547574 | 0.528 | 0.75 | 9.29E-299 |
| ECM Remote Zone | Up | Itih3 | 1.39E-182 | 0.27108219 | 0.379 | 0.619 | 2.78E-179 |
| ECM Remote Zone | Up | Mx1 | 9.26E-129 | 0.27019928 | 0.659 | 0.997 | 1.85E-125 |
| ECM Remote Zone | Up | Zfhx2 | 4.65E-134 | 0.26635954 | 0.491 | 0.708 | 9.30E-131 |
| ECM Remote Zone | Up | MGC108823 | 1.96E-281 | 0.26511084 | 0.61 | 0.976 | 3.92E-278 |

| | | | | | | | |
|---|---|---|---|---|---|---|---|
| ECM Remote Zone | Up | Nrde2 | 0 | 0.25493829 | 0.465 | 0.855 | 0 |
| Saline Remote Zone | Up | Slc37a2 | 1.64E-283 | -0.2500265 | 0.468 | 0.701 | 3.29E-280 |
| Saline Remote Zone | Up | Cmtm5 | 9.18E-193 | -0.2508667 | 0.353 | 0.553 | 1.84E-189 |
| Saline Remote Zone | Up | Vps33a | 0 | -0.2510835 | 0.438 | 0.787 | 0 |
| Saline Remote Zone | Up | Rab13 | 1.47E-228 | -0.2522572 | 0.485 | 0.755 | 2.94E-225 |
| Saline Remote Zone | Up | RT1-Ba | 4.67E-81 | -0.2528653 | 0.602 | 0.815 | 9.35E-78 |
| Saline Remote Zone | Up | Cic | 2.16E-150 | -0.2530492 | 0.565 | 0.798 | 4.31E-147 |
| Saline Remote Zone | Up | Ttc26 | 1.32E-249 | -0.2531855 | 0.499 | 0.719 | 2.63E-246 |
| Saline Remote Zone | Up | Aldh3a1 | 0 | -0.2538453 | 0.361 | 0.631 | 0 |

| | | | | | | | |
|---|---|---|---|---|---|---|---|
| Saline Remote Zone | Up | Ddost | 6.54E-106 | -0.2539802 | 0.68 | 0.918 | 1.31E-102 |
| Saline Remote Zone | Up | Ptprj | 7.55E-263 | -0.2541812 | 0.47 | 0.706 | 1.51E-259 |
| Saline Remote Zone | Up | Csrp1 | 2.65E-71 | -0.2548808 | 0.605 | 0.808 | 5.30E-68 |
| Saline Remote Zone | Up | Plpp1 | 4.41E-226 | -0.254951 | 0.562 | 0.796 | 8.83E-223 |
| Saline Remote Zone | Up | Prps2 | 0 | -0.2550316 | 0.369 | 0.676 | 0 |
| Saline Remote Zone | Up | RGD1304694 | 9.80E-278 | -0.2551388 | 0.491 | 0.758 | 1.96E-274 |
| Saline Remote Zone | Up | Cope | 3.32E-34 | -0.2574119 | 0.67 | 0.883 | 6.64E-31 |
| Saline Remote Zone | Up | Arhgef25 | 7.50E-154 | -0.2587648 | 0.489 | 0.705 | 1.50E-150 |
| Saline Remote Zone | Up | Tnfsf13 | 0 | -0.2593277 | 0.371 | 0.659 | 0 |

| | | | | | | | |
|---|---|---|---|---|---|---|---|
| Saline Remote Zone | Up | Arf6 | 3.02E-245 | -0.2617186 | 0.531 | 0.808 | 6.04E-242 |
| Saline Remote Zone | Up | Eif3b | 1.08E-64 | -0.2628086 | 0.684 | 0.91 | 2.16E-61 |
| Saline Remote Zone | Up | Cenpf | 6.05E-265 | -0.2649293 | 0.442 | 0.7 | 1.21E-261 |
| Saline Remote Zone | Up | Triobp | 3.58E-228 | -0.265849 | 0.449 | 0.704 | 7.17E-225 |
| Saline Remote Zone | Up | Gba | 2.49E-218 | -0.2671295 | 0.524 | 0.822 | 4.98E-215 |
| Saline Remote Zone | Up | Ccl21 | 9.33E-275 | -0.2673197 | 0.341 | 0.567 | 1.87E-271 |
| Saline Remote Zone | Up | Ggcx | 2.64E-191 | -0.2674546 | 0.511 | 0.792 | 5.28E-188 |
| Saline Remote Zone | Up | Slc48a1 | 6.03E-122 | -0.2676157 | 0.59 | 0.814 | 1.21E-118 |
| Saline Remote Zone | Up | Cxcr4 | 2.03E-204 | -0.2676285 | 0.533 | 0.818 | 4.07E-201 |

| | | | | | | | |
|---|---|---|---|---|---|---|---|
| Saline Remote Zone | Up | Vwf | 3.53E-98 | -0.2687165 | 0.539 | 0.751 | 7.05E-95 |
| Saline Remote Zone | Up | Rab34 | 1.09E-197 | -0.2695006 | 0.515 | 0.8 | 2.18E-194 |
| Saline Remote Zone | Up | Zfhx3 | 9.59E-299 | -0.2702648 | 0.477 | 0.792 | 1.92E-295 |
| Saline Remote Zone | Up | Rsf1 | 0 | -0.2707871 | 0.485 | 0.828 | 0 |
| Saline Remote Zone | Up | AC128836.1 | 0 | -0.2721106 | 0.443 | 0.707 | 0 |
| Saline Remote Zone | Up | Cyth4 | 0 | -0.2730434 | 0.395 | 0.696 | 0 |
| Saline Remote Zone | Up | Ptbp1 | 9.30E-128 | -0.2734605 | 0.551 | 0.816 | 1.86E-124 |
| Saline Remote Zone | Up | Prf1 | 0 | -0.2735956 | 0.508 | 0.888 | 0 |
| Saline Remote Zone | Up | Rin2 | 9.04E-143 | -0.2736316 | 0.494 | 0.706 | 1.81E-139 |

| | | | | | | | |
|---|---|---|---|---|---|---|---|
| Saline Remote Zone | Up | Stmn4 | 0 | -0.2748894 | 0.404 | 0.762 | 0 |
| Saline Remote Zone | Up | Pkn1 | 2.69E-137 | -0.2761659 | 0.521 | 0.762 | 5.38E-134 |
| Saline Remote Zone | Up | Abca1 | 4.51E-223 | -0.2776806 | 0.449 | 0.705 | 9.03E-220 |
| Saline Remote Zone | Up | Stk38l | 0 | -0.2780808 | 0.411 | 0.68 | 0 |
| Saline Remote Zone | Up | Prdx4 | 9.04E-131 | -0.2781608 | 0.646 | 0.913 | 1.81E-127 |
| Saline Remote Zone | Up | Gucy1a1 | 0 | -0.2801179 | 0.367 | 0.635 | 0 |
| Saline Remote Zone | Up | Sirpa | 3.33E-83 | -0.2806469 | 0.594 | 0.804 | 6.67E-80 |
| Saline Remote Zone | Up | C5ar1 | 1.83E-268 | -0.2806825 | 0.477 | 0.765 | 3.65E-265 |
| Saline Remote Zone | Up | Tpsb2 | 7.69E-254 | -0.2810833 | 0.342 | 0.552 | 1.54E-250 |

| | | | | | | | |
|---|---|---|---|---|---|---|---|
| Saline Remote Zone | Up | Stard10 | 0 | -0.2818034 | 0.294 | 0.69 | 0 |
| Saline Remote Zone | Up | Ptprcap | 0 | -0.2823182 | 0.488 | 0.876 | 0 |
| Saline Remote Zone | Up | Lxn | 0 | -0.2826482 | 0.342 | 0.679 | 0 |
| Saline Remote Zone | Up | Pmaip1 | 1.07E-225 | -0.2831861 | 0.458 | 0.69 | 2.15E-222 |
| Saline Remote Zone | Up | Farp1 | 2.65E-257 | -0.2852223 | 0.446 | 0.726 | 5.30E-254 |
| Saline Remote Zone | Up | Mettl18 | 0 | -0.2852894 | 0.398 | 0.689 | 0 |
| Saline Remote Zone | Up | Susd2 | 0 | -0.2854647 | 0.317 | 0.661 | 0 |
| Saline Remote Zone | Up | Cybc1 | 2.80E-291 | -0.2856118 | 0.464 | 0.768 | 5.60E-288 |
| Saline Remote Zone | Up | Serp1 | 1.09E-159 | -0.2860209 | 0.512 | 0.726 | 2.18E-156 |

| | | | | | | | |
|---|---|---|---|---|---|---|---|
| Saline Remote Zone | Up | Sbno2 | 0 | -0.2875218 | 0.382 | 0.686 | 0 |
| Saline Remote Zone | Up | Sult1a1 | 0 | -0.2879655 | 0.374 | 0.7 | 0 |
| Saline Remote Zone | Up | Reck | 0 | -0.2881065 | 0.352 | 0.62 | 0 |
| Saline Remote Zone | Up | Folr2 | 1.08E-184 | -0.2884917 | 0.543 | 0.835 | 2.17E-181 |
| Saline Remote Zone | Up | Usp30 | 4.22E-302 | -0.2891254 | 0.406 | 0.674 | 8.44E-299 |
| Saline Remote Zone | Up | Man1b1 | 5.18E-118 | -0.2910203 | 0.56 | 0.792 | 1.04E-114 |
| Saline Remote Zone | Up | Plpp5 | 0 | -0.292809 | 0.244 | 0.467 | 0 |
| Saline Remote Zone | Up | Rassf4 | 0 | -0.2932042 | 0.372 | 0.622 | 0 |
| Saline Remote Zone | Up | Ugt1a6 | 1.84E-234 | -0.2947583 | 0.484 | 0.72 | 3.68E-231 |

| | | | | | | | |
|---|---|---|---|---|---|---|---|
| Saline Remote Zone | Up | Crlf2 | 0 | -0.2966481 | 0.424 | 0.746 | 0 |
| Saline Remote Zone | Up | Ptcd3 | 3.12E-195 | -0.2970096 | 0.571 | 0.833 | 6.23E-192 |
| Saline Remote Zone | Up | Abhd8 | 0 | -0.2973853 | 0.414 | 0.774 | 0 |
| Saline Remote Zone | Up | Arl4d | 0 | -0.2984532 | 0.268 | 0.557 | 0 |
| Saline Remote Zone | Up | Mki67 | 0 | -0.2986047 | 0.324 | 0.719 | 0 |
| Saline Remote Zone | Up | Gpx8 | 1.23E-130 | -0.2987832 | 0.589 | 0.817 | 2.47E-127 |
| Saline Remote Zone | Up | Eri3 | 2.47E-134 | -0.2988425 | 0.626 | 0.913 | 4.94E-131 |
| Saline Remote Zone | Up | Alcam | 2.47E-262 | -0.2990699 | 0.518 | 0.829 | 4.93E-259 |
| Saline Remote Zone | Up | Marveld1 | 2.90E-107 | -0.3004135 | 0.575 | 0.793 | 5.79E-104 |

| | | | | | | | |
|---|---|---|---|---|---|---|---|
| Saline Remote Zone | Up | Prg4 | 5.97E-236 | -0.3006054 | 0.477 | 0.692 | 1.19E-232 |
| Saline Remote Zone | Up | LOC100910636 | 0 | -0.3006616 | 0.46 | 0.772 | 0 |
| Saline Remote Zone | Up | Dynlt1 | 4.79E-117 | -0.3013606 | 0.527 | 0.732 | 9.59E-114 |
| Saline Remote Zone | Up | Coro1a | 5.73E-168 | -0.3014238 | 0.518 | 0.75 | 1.15E-164 |
| Saline Remote Zone | Up | Rab3d | 7.49E-187 | -0.3021561 | 0.523 | 0.746 | 1.50E-183 |
| Saline Remote Zone | Up | Cpeb1 | 0 | -0.3032693 | 0.492 | 0.842 | 0 |
| Saline Remote Zone | Up | Mfsd4b | 1.03E-261 | -0.303361 | 0.417 | 0.7 | 2.05E-258 |
| Saline Remote Zone | Up | A3galt2 | 6.59E-268 | -0.3035629 | 0.383 | 0.591 | 1.32E-264 |
| Saline Remote Zone | Up | Ypel3 | 2.16E-56 | -0.3040749 | 0.641 | 0.848 | 4.32E-53 |

| | | | | | | | |
|---|---|---|---|---|---|---|---|
| Saline Remote Zone | Up | Tmem43 | 2.90E-177 | -0.3041037 | 0.459 | 0.661 | 5.79E-174 |
| Saline Remote Zone | Up | Fasn | 0 | -0.30628 | 0.443 | 0.741 | 0 |
| Saline Remote Zone | Up | Cd53 | 1.06E-304 | -0.3090595 | 0.413 | 0.67 | 2.11E-301 |
| Saline Remote Zone | Up | Pik3r2 | 8.21E-212 | -0.309697 | 0.557 | 0.879 | 1.64E-208 |
| Saline Remote Zone | Up | Ticam2 | 0 | -0.3099797 | 0.422 | 0.739 | 0 |
| Saline Remote Zone | Up | Mcrip1 | 2.31E-89 | -0.3100112 | 0.577 | 0.808 | 4.62E-86 |
| Saline Remote Zone | Up | Alox5 | 1.81E-256 | -0.3100461 | 0.47 | 0.696 | 3.62E-253 |
| Saline Remote Zone | Up | Coro1b | 1.68E-127 | -0.3107198 | 0.532 | 0.736 | 3.35E-124 |
| Saline Remote Zone | Up | Scyl1 | 6.56E-121 | -0.3118299 | 0.62 | 0.9 | 1.31E-117 |

| | | | | | | | |
|---|---|---|---|---|---|---|---|
| Saline Remote Zone | Up | Gramd1a | 2.12E-222 | -0.3126398 | 0.468 | 0.716 | 4.24E-219 |
| Saline Remote Zone | Up | Ubd | 3.43E-199 | -0.3128885 | 0.554 | 0.926 | 6.86E-196 |
| Saline Remote Zone | Up | Cd82 | 4.57E-274 | -0.3135108 | 0.515 | 0.852 | 9.14E-271 |
| Saline Remote Zone | Up | Napsa | 0 | -0.3136278 | 0.317 | 0.609 | 0 |
| Saline Remote Zone | Up | Gimap4 | 0 | -0.3153414 | 0.347 | 0.703 | 0 |
| Saline Remote Zone | Up | Plod3 | 1.69E-282 | -0.3153677 | 0.403 | 0.68 | 3.38E-279 |
| Saline Remote Zone | Up | Tmed3 | 2.38E-71 | -0.3176665 | 0.643 | 0.872 | 4.76E-68 |
| Saline Remote Zone | Up | Mcub | 0 | -0.318712 | 0.469 | 0.802 | 0 |
| Saline Remote Zone | Up | Golim4 | 6.55E-296 | -0.3190764 | 0.486 | 0.785 | 1.31E-292 |

| | | | | | | | |
|---|---|---|---|---|---|---|---|
| Saline Remote Zone | Up | Plek | 2.91E-304 | -0.3198253 | 0.445 | 0.709 | 5.82E-301 |
| Saline Remote Zone | Up | Klhl6 | 4.24E-273 | -0.3204073 | 0.364 | 0.58 | 8.47E-270 |
| Saline Remote Zone | Up | P2rx1 | 0 | -0.320501 | 0.485 | 0.749 | 0 |
| Saline Remote Zone | Up | Lurap1l | 1.45E-170 | -0.3217529 | 0.469 | 0.693 | 2.90E-167 |
| Saline Remote Zone | Up | Rara | 2.53E-250 | -0.3217664 | 0.409 | 0.651 | 5.06E-247 |
| Saline Remote Zone | Up | Ppp1r12c | 1.92E-103 | -0.3218054 | 0.581 | 0.794 | 3.83E-100 |
| Saline Remote Zone | Up | Hoxb4 | 3.20E-292 | -0.3225933 | 0.533 | 0.761 | 6.41E-289 |
| Saline Remote Zone | Up | LOC362863 | 2.04E-241 | -0.3227672 | 0.389 | 0.654 | 4.08E-238 |
| Saline Remote Zone | Up | Fam167b | 0 | -0.3235878 | 0.493 | 0.829 | 0 |

| | | | | | | | |
|---|---|---|---|---|---|---|---|
| Saline Remote Zone | Up | Arhgap45 | 0 | -0.3243613 | 0.373 | 0.728 | 0 |
| Saline Remote Zone | Up | Slc4a2 | 4.06E-307 | -0.3251438 | 0.492 | 0.844 | 8.12E-304 |
| Saline Remote Zone | Up | Mras | 0 | -0.3251582 | 0.423 | 0.717 | 0 |
| Saline Remote Zone | Up | Itgb4 | 0 | -0.325521 | 0.386 | 0.702 | 0 |
| Saline Remote Zone | Up | Pld4 | 8.61E-160 | -0.3261336 | 0.554 | 0.772 | 1.72E-156 |
| Saline Remote Zone | Up | Ext1 | 4.80E-98 | -0.3264177 | 0.65 | 0.858 | 9.61E-95 |
| Saline Remote Zone | Up | Cep85l | 5.87E-253 | -0.3265263 | 0.459 | 0.7 | 1.17E-249 |
| Saline Remote Zone | Up | Xylt2 | 6.44E-144 | -0.3273984 | 0.518 | 0.73 | 1.29E-140 |
| Saline Remote Zone | Up | Cst6 | 2.18E-140 | -0.3281542 | 0.446 | 0.682 | 4.36E-137 |

| | | | | | | | |
|---|---|---|---|---|---|---|---|
| Saline Remote Zone | Up | Tnfaip8l3 | 1.56E-253 | -0.3287035 | 0.379 | 0.596 | 3.12E-250 |
| Saline Remote Zone | Up | Hm13 | 1.39E-134 | -0.3297906 | 0.47 | 0.691 | 2.77E-131 |
| Saline Remote Zone | Up | Enc1 | 3.92E-194 | -0.3299737 | 0.519 | 0.812 | 7.84E-191 |
| Saline Remote Zone | Up | Adamts12 | 2.47E-195 | -0.330297 | 0.456 | 0.71 | 4.95E-192 |
| Saline Remote Zone | Up | Klri1 | 1.10E-304 | -0.3318486 | 0.485 | 0.694 | 2.20E-301 |
| Saline Remote Zone | Up | Uvssa | 1.03E-177 | -0.3319349 | 0.274 | 0.555 | 2.06E-174 |
| Saline Remote Zone | Up | Arfip2 | 1.18E-195 | -0.3333604 | 0.443 | 0.653 | 2.36E-192 |
| Saline Remote Zone | Up | Prima1 | 0 | -0.3336596 | 0.416 | 0.716 | 0 |
| Saline Remote Zone | Up | Calhm2 | 0 | -0.3343752 | 0.389 | 0.759 | 0 |

| | | | | | | | |
|---|---|---|---|---|---|---|---|
| Saline Remote Zone | Up | Dlgap4 | 0 | -0.335255 | 0.497 | 0.836 | 0 |
| Saline Remote Zone | Up | Nckap5 | 0 | -0.335883 | 0.375 | 0.7 | 0 |
| Saline Remote Zone | Up | Fstl3 | 6.12E-210 | -0.3367712 | 0.486 | 0.716 | 1.22E-206 |
| Saline Remote Zone | Up | Ndrg1 | 1.61E-258 | -0.3368022 | 0.387 | 0.597 | 3.21E-255 |
| Saline Remote Zone | Up | Flrt3 | 0 | -0.3371711 | 0.463 | 0.842 | 0 |
| Saline Remote Zone | Up | Aida | 6.94E-200 | -0.3375731 | 0.618 | 0.891 | 1.39E-196 |
| Saline Remote Zone | Up | AABR07027309.1 | 7.47E-204 | -0.3381317 | 0.45 | 0.694 | 1.49E-200 |
| Saline Remote Zone | Up | Pltp | 2.84E-37 | -0.3401327 | 0.715 | 0.922 | 5.67E-34 |
| Saline Remote Zone | Up | Snx20 | 9.36E-208 | -0.3409553 | 0.408 | 0.613 | 1.87E-204 |

| | | | | | | | |
|---|---|---|---|---|---|---|---|
| Saline Remote Zone | Up | P2ry6 | 0 | -0.3417056 | 0.411 | 0.702 | 0 |
| Saline Remote Zone | Up | Skap2 | 4.31E-212 | -0.342034 | 0.483 | 0.712 | 8.62E-209 |
| Saline Remote Zone | Up | Zcchc2 | 0 | -0.3429581 | 0.273 | 0.49 | 0 |
| Saline Remote Zone | Up | Dbn1 | 2.80E-189 | -0.3433828 | 0.52 | 0.824 | 5.60E-186 |
| Saline Remote Zone | Up | Cdr2l | 2.93E-192 | -0.34362 | 0.477 | 0.697 | 5.86E-189 |
| Saline Remote Zone | Up | Nckap1l | 0 | -0.3443759 | 0.352 | 0.648 | 0 |
| Saline Remote Zone | Up | Ppp1r18 | 1.41E-160 | -0.3446476 | 0.573 | 0.839 | 2.81E-157 |
| Saline Remote Zone | Up | Hs2st1 | 5.32E-245 | -0.3447642 | 0.503 | 0.796 | 1.06E-241 |
| Saline Remote Zone | Up | Cpa3 | 2.22E-238 | -0.3461583 | 0.276 | 0.51 | 4.44E-235 |

| | | | | | | | |
|---|---|---|---|---|---|---|---|
| Saline Remote Zone | Up | Myl4 | 1.65E-295 | -0.3462447 | 0.508 | 0.837 | 3.30E-292 |
| Saline Remote Zone | Up | Cnn1 | 6.63E-247 | -0.3465524 | 0.332 | 0.537 | 1.33E-243 |
| Saline Remote Zone | Up | Dchs1 | 1.64E-210 | -0.3468042 | 0.562 | 0.867 | 3.29E-207 |
| Saline Remote Zone | Up | Gzmk | 6.31E-305 | -0.3468336 | 0.451 | 0.7 | 1.26E-301 |
| Saline Remote Zone | Up | Specc1 | 0 | -0.3469863 | 0.386 | 0.77 | 0 |
| Saline Remote Zone | Up | Cers5 | 1.46E-274 | -0.3481085 | 0.448 | 0.755 | 2.92E-271 |
| Saline Remote Zone | Up | Ppp1r12b | 0 | -0.3484487 | 0.507 | 0.872 | 0 |
| Saline Remote Zone | Up | Mal | 6.02E-234 | -0.3498298 | 0.333 | 0.577 | 1.20E-230 |
| Saline Remote Zone | Up | St3gal4 | 8.41E-224 | -0.350155 | 0.523 | 0.799 | 1.68E-220 |

| | | | | | | | |
|---|---|---|---|---|---|---|---|
| Saline Remote Zone | Up | Gusb | 2.98E-182 | -0.3519921 | 0.482 | 0.713 | 5.97E-179 |
| Saline Remote Zone | Up | Sfr1 | 1.25E-121 | -0.3521783 | 0.646 | 0.858 | 2.49E-118 |
| Saline Remote Zone | Up | Trip6 | 5.16E-221 | -0.3523788 | 0.493 | 0.762 | 1.03E-217 |
| Saline Remote Zone | Up | Gng2 | 5.34E-250 | -0.3526359 | 0.457 | 0.691 | 1.07E-246 |
| Saline Remote Zone | Up | Slc35c2 | 0 | -0.3533476 | 0.435 | 0.728 | 0 |
| Saline Remote Zone | Up | Nrgn | 0 | -0.3543124 | 0.339 | 0.68 | 0 |
| Saline Remote Zone | Up | Atp1b2 | 0 | -0.3545099 | 0.21 | 0.425 | 0 |
| Saline Remote Zone | Up | Ikbip | 1.20E-231 | -0.3557786 | 0.506 | 0.81 | 2.40E-228 |
| Saline Remote Zone | Up | Yipf1 | 2.57E-261 | -0.3575448 | 0.479 | 0.764 | 5.13E-258 |

| | | | | | | | |
|---|---|---|---|---|---|---|---|
| Saline Remote Zone | Up | Sh3bp1 | 0 | -0.3581218 | 0.341 | 0.642 | 0 |
| Saline Remote Zone | Up | Cdkn1c | 0 | -0.358562 | 0.41 | 0.713 | 0 |
| Saline Remote Zone | Up | Alox15 | 0 | -0.3590709 | 0.371 | 0.693 | 0 |
| Saline Remote Zone | Up | Nelfe | 0 | -0.3599346 | 0.433 | 0.768 | 0 |
| Saline Remote Zone | Up | Rhox5 | 0 | -0.3609496 | 0.35 | 0.611 | 0 |
| Saline Remote Zone | Up | Olfml1 | 3.71E-306 | -0.3611651 | 0.384 | 0.637 | 7.43E-303 |
| Saline Remote Zone | Up | Sfrp1 | 0 | -0.3619249 | 0.232 | 0.626 | 0 |
| Saline Remote Zone | Up | Atxn7l3b | 1.03E-176 | -0.3623086 | 0.533 | 0.773 | 2.07E-173 |
| Saline Remote Zone | Up | Jag1 | 3.18E-298 | -0.3626019 | 0.401 | 0.679 | 6.35E-295 |

| | | | | | | | |
|---|---|---|---|---|---|---|---|
| Saline Remote Zone | Up | Smim29 | 2.70E-276 | -0.3639674 | 0.418 | 0.659 | 5.41E-273 |
| Saline Remote Zone | Up | Scamp2 | 1.19E-159 | -0.3647397 | 0.569 | 0.831 | 2.38E-156 |
| Saline Remote Zone | Up | Nagk | 0 | -0.3650093 | 0.42 | 0.706 | 0 |
| Saline Remote Zone | Up | Mrvi1 | 0 | -0.3650143 | 0.245 | 0.447 | 0 |
| Saline Remote Zone | Up | LOC100359539 | 0 | -0.3654267 | 0.281 | 0.521 | 0 |
| Saline Remote Zone | Up | Ncln | 1.73E-168 | -0.3658149 | 0.541 | 0.843 | 3.45E-165 |
| Saline Remote Zone | Up | Lama5 | 6.05E-124 | -0.3658514 | 0.53 | 0.733 | 1.21E-120 |
| Saline Remote Zone | Up | Laptm5 | 1.98E-214 | -0.3666377 | 0.457 | 0.72 | 3.97E-211 |
| Saline Remote Zone | Up | Hsd17b7 | 0 | -0.3679148 | 0.385 | 0.67 | 0 |

| | | | | | | | |
|---|---|---|---|---|---|---|---|
| Saline Remote Zone | Up | Bin2 | 2.88E-155 | -0.3683254 | 0.48 | 0.702 | 5.76E-152 |
| Saline Remote Zone | Up | Gsdmd | 5.91E-268 | -0.3688854 | 0.439 | 0.709 | 1.18E-264 |
| Saline Remote Zone | Up | Timp1 | 1.28E-62 | -0.3690031 | 0.666 | 0.894 | 2.56E-59 |
| Saline Remote Zone | Up | Mylk | 1.62E-226 | -0.3707235 | 0.348 | 0.574 | 3.24E-223 |
| Saline Remote Zone | Up | Plac9 | 4.57E-257 | -0.3707562 | 0.466 | 0.737 | 9.13E-254 |
| Saline Remote Zone | Up | Prph | 6.72E-293 | -0.3709046 | 0.453 | 0.761 | 1.34E-289 |
| Saline Remote Zone | Up | Hmcn1 | 0 | -0.3712251 | 0.369 | 0.638 | 0 |
| Saline Remote Zone | Up | Fam3c | 3.10E-271 | -0.37142 | 0.384 | 0.595 | 6.19E-268 |
| Saline Remote Zone | Up | Sbsn | 1.33E-213 | -0.3763988 | 0.449 | 0.673 | 2.66E-210 |

| | | | | | | | |
|---|---|---|---|---|---|---|---|
| Saline Remote Zone | Up | Gfra3 | 0 | -0.3775509 | 0.459 | 0.838 | 0 |
| Saline Remote Zone | Up | Tmem176a | 1.36E-87 | -0.377761 | 0.649 | 0.904 | 2.72E-84 |
| Saline Remote Zone | Up | Leprot | 3.27E-230 | -0.3788942 | 0.493 | 0.792 | 6.54E-227 |
| Saline Remote Zone | Up | Ufm1 | 1.72E-245 | -0.3792721 | 0.457 | 0.674 | 3.45E-242 |
| Saline Remote Zone | Up | Rab11fip1 | 5.82E-270 | -0.3798222 | 0.464 | 0.753 | 1.16E-266 |
| Saline Remote Zone | Up | Fcer1g | 7.31E-76 | -0.3800782 | 0.625 | 0.842 | 1.46E-72 |
| Saline Remote Zone | Up | Man2b2 | 1.07E-306 | -0.3807441 | 0.452 | 0.755 | 2.14E-303 |
| Saline Remote Zone | Up | Ms4a7 | 0 | -0.3808482 | 0.376 | 0.674 | 0 |
| Saline Remote Zone | Up | Serpinb1a | 4.20E-214 | -0.3821208 | 0.511 | 0.757 | 8.40E-211 |

| | | | | | | | |
|---|---|---|---|---|---|---|---|
| Saline Remote Zone | Up | AABR07028769.1 | 0 | -0.3829977 | 0.453 | 0.757 | 0 |
| Saline Remote Zone | Up | Kpna3 | 1.99E-304 | -0.383281 | 0.468 | 0.733 | 3.98E-301 |
| Saline Remote Zone | Up | Tes | 0 | -0.3833929 | 0.348 | 0.581 | 0 |
| Saline Remote Zone | Up | Ddah2 | 1.15E-99 | -0.3834792 | 0.642 | 0.92 | 2.30E-96 |
| Saline Remote Zone | Up | Cfd | 0 | -0.3835427 | 0.307 | 0.696 | 0 |
| Saline Remote Zone | Up | Lrrk1 | 8.42E-233 | -0.3851901 | 0.537 | 0.809 | 1.68E-229 |
| Saline Remote Zone | Up | Cbfa2t3 | 0 | -0.3873355 | 0.454 | 0.733 | 0 |
| Saline Remote Zone | Up | Adm | 0 | -0.3874627 | 0.344 | 0.568 | 0 |
| Saline Remote Zone | Up | Zfp260.1 | 8.59E-171 | -0.3880059 | 0.529 | 0.734 | 1.72E-167 |

| | | | | | | | |
|---|---|---|---|---|---|---|---|
| Saline Remote Zone | Up | Os9 | 2.80E-150 | -0.3886042 | 0.498 | 0.74 | 5.60E-147 |
| Saline Remote Zone | Up | Cep135 | 0 | -0.3887143 | 0.34 | 0.607 | 0 |
| Saline Remote Zone | Up | Entpd2 | 0 | -0.388797 | 0.27 | 0.521 | 0 |
| Saline Remote Zone | Up | Myoz1 | 0 | -0.3891022 | 0.382 | 0.782 | 0 |
| Saline Remote Zone | Up | Mfsd11 | 0 | -0.3899221 | 0.413 | 0.741 | 0 |
| Saline Remote Zone | Up | Inpp5d | 0 | -0.3904155 | 0.386 | 0.653 | 0 |
| Saline Remote Zone | Up | Acap3 | 3.17E-224 | -0.3905596 | 0.461 | 0.692 | 6.33E-221 |
| Saline Remote Zone | Up | Ncbp2 | 5.37E-234 | -0.3906112 | 0.524 | 0.814 | 1.07E-230 |
| Saline Remote Zone | Up | St3gal2 | 6.73E-157 | -0.3906475 | 0.565 | 0.774 | 1.35E-153 |

| | | | | | | | |
|---|---|---|---|---|---|---|---|
| Saline Remote Zone | Up | Senp2 | 1.41E-92 | -0.3916594 | 0.267 | 0.505 | 2.82E-89 |
| Saline Remote Zone | Up | Hps1 | 6.27E-209 | -0.3927434 | 0.525 | 0.752 | 1.25E-205 |
| Saline Remote Zone | Up | Lck | 0 | -0.3929477 | 0.481 | 0.788 | 0 |
| Saline Remote Zone | Up | LOC108348250 | 0 | -0.3946707 | 0.479 | 0.857 | 0 |
| Saline Remote Zone | Up | Bax | 0 | -0.3956017 | 0.457 | 0.842 | 0 |
| Saline Remote Zone | Up | Plod1 | 3.05E-149 | -0.3963462 | 0.578 | 0.824 | 6.10E-146 |
| Saline Remote Zone | Up | Ighm | 3.91E-247 | -0.3965632 | 0.458 | 0.772 | 7.83E-244 |
| Saline Remote Zone | Up | Glb1 | 2.19E-207 | -0.3967418 | 0.552 | 0.815 | 4.39E-204 |
| Saline Remote Zone | Up | Col20a1 | 2.27E-108 | -0.3968717 | 0.508 | 0.729 | 4.54E-105 |

| | | | | | | | |
|---|---|---|---|---|---|---|---|
| Saline Remote Zone | Up | Aph1b | 1.52E-121 | -0.3999881 | 0.616 | 0.861 | 3.04E-118 |
| Saline Remote Zone | Up | Spon1 | 5.46E-148 | -0.3999932 | 0.546 | 0.771 | 1.09E-144 |
| Saline Remote Zone | Up | Armcx2 | 0 | -0.4014517 | 0.473 | 0.749 | 0 |
| Saline Remote Zone | Up | Cys1 | 0 | -0.4016915 | 0.492 | 0.9 | 0 |
| Saline Remote Zone | Up | Rbbp9 | 0 | -0.4017806 | 0.419 | 0.757 | 0 |
| Saline Remote Zone | Up | Msrb1 | 8.66E-159 | -0.4043612 | 0.573 | 0.811 | 1.73E-155 |
| Saline Remote Zone | Up | Galk1 | 9.34E-294 | -0.4074503 | 0.486 | 0.813 | 1.87E-290 |
| Saline Remote Zone | Up | LOC103694857 | 0 | -0.410789 | 0.231 | 0.524 | 0 |
| Saline Remote Zone | Up | Blnk | 9.10E-194 | -0.411087 | 0.506 | 0.714 | 1.82E-190 |

| | | | | | | | |
|---|---|---|---|---|---|---|---|
| Saline Remote Zone | Up | Fzd2 | 4.78E-195 | -0.411971 | 0.549 | 0.833 | 9.55E-192 |
| Saline Remote Zone | Up | P3h1 | 2.91E-216 | -0.4125279 | 0.525 | 0.808 | 5.82E-213 |
| Saline Remote Zone | Up | Jund | 2.59E-79 | -0.4127397 | 0.51 | 0.72 | 5.19E-76 |
| Saline Remote Zone | Up | Ahr | 6.17E-186 | -0.4129441 | 0.51 | 0.731 | 1.23E-182 |
| Saline Remote Zone | Up | Cdc20 | 0 | -0.4134515 | 0.308 | 0.592 | 0 |
| Saline Remote Zone | Up | Dpysl3 | 1.53E-268 | -0.4150631 | 0.443 | 0.675 | 3.07E-265 |
| Saline Remote Zone | Up | Bbs9 | 6.04E-272 | -0.4158094 | 0.46 | 0.776 | 1.21E-268 |
| Saline Remote Zone | Up | Alg8 | 3.78E-287 | -0.4159684 | 0.392 | 0.676 | 7.55E-284 |
| Saline Remote Zone | Up | Tubb6 | 0 | -0.4169566 | 0.446 | 0.69 | 0 |

| | | | | | | | |
|---|---|---|---|---|---|---|---|
| Saline Remote Zone | Up | Lrg1 | 3.77E-182 | -0.4172791 | 0.506 | 0.761 | 7.53E-179 |
| Saline Remote Zone | Up | Cdkn3 | 0 | -0.4187554 | 0.284 | 0.518 | 0 |
| Saline Remote Zone | Up | Fcna | 0 | -0.4188995 | 0.347 | 0.646 | 0 |
| Saline Remote Zone | Up | Slc25a1 | 2.41E-267 | -0.4189748 | 0.463 | 0.747 | 4.83E-264 |
| Saline Remote Zone | Up | Vcam1 | 5.95E-231 | -0.4198694 | 0.476 | 0.789 | 1.19E-227 |
| Saline Remote Zone | Up | Ifngr2 | 1.48E-301 | -0.4200847 | 0.416 | 0.712 | 2.96E-298 |
| Saline Remote Zone | Up | Pxn | 2.45E-272 | -0.4213466 | 0.475 | 0.801 | 4.90E-269 |
| Saline Remote Zone | Up | Crlf1 | 1.65E-300 | -0.4239213 | 0.457 | 0.758 | 3.30E-297 |
| Saline Remote Zone | Up | Atp10a | 4.96E-229 | -0.4261931 | 0.387 | 0.589 | 9.93E-226 |

| | | | | | | | |
|---|---|---|---|---|---|---|---|
| Saline Remote Zone | Up | Agtr1a | 0 | -0.4269926 | 0.32 | 0.58 | 0 |
| Saline Remote Zone | Up | Nfatc4 | 0 | -0.4272415 | 0.446 | 0.826 | 0 |
| Saline Remote Zone | Up | Gna15 | 4.75E-257 | -0.4279123 | 0.44 | 0.676 | 9.50E-254 |
| Saline Remote Zone | Up | Casp12 | 9.60E-279 | -0.4279741 | 0.413 | 0.67 | 1.92E-275 |
| Saline Remote Zone | Up | Irf8 | 0 | -0.4285658 | 0.424 | 0.753 | 0 |
| Saline Remote Zone | Up | Snx18 | 6.78E-217 | -0.4299759 | 0.462 | 0.688 | 1.36E-213 |
| Saline Remote Zone | Up | Limd2 | 0 | -0.4305609 | 0.472 | 0.804 | 0 |
| Saline Remote Zone | Up | Thbs3 | 4.42E-234 | -0.4315505 | 0.399 | 0.616 | 8.85E-231 |
| Saline Remote Zone | Up | Egfl8 | 7.07E-270 | -0.4325386 | 0.417 | 0.69 | 1.41E-266 |

| | | | | | | | |
|---|---|---|---|---|---|---|---|
| Saline Remote Zone | Up | B3galnt1 | 2.47E-223 | -0.4325453 | 0.562 | 0.845 | 4.93E-220 |
| Saline Remote Zone | Up | LOC100910669 | 9.66E-170 | -0.4340133 | 0.515 | 0.743 | 1.93E-166 |
| Saline Remote Zone | Up | RT1-A2 | 0 | -0.4347982 | 0.317 | 0.586 | 0 |
| Saline Remote Zone | Up | Mdk | 0 | -0.4348182 | 0.512 | 0.858 | 0 |
| Saline Remote Zone | Up | Cd8a | 2.76E-273 | -0.4369268 | 0.451 | 0.71 | 5.52E-270 |
| Saline Remote Zone | Up | Sigmar1 | 1.19E-281 | -0.4372563 | 0.517 | 0.829 | 2.38E-278 |
| Saline Remote Zone | Up | Lima1 | 1.89E-208 | -0.4372961 | 0.52 | 0.773 | 3.79E-205 |
| Saline Remote Zone | Up | Ahsp | 0 | -0.4392654 | 0.483 | 0.784 | 0 |
| Saline Remote Zone | Up | Socs1 | 0 | -0.4394362 | 0.354 | 0.675 | 0 |

| | | | | | | | |
|---|---|---|---|---|---|---|---|
| Saline Remote Zone | Up | Alox5ap | 0 | -0.4394463 | 0.407 | 0.748 | 0 |
| Saline Remote Zone | Up | Tubb2b | 0 | -0.4395575 | 0.431 | 0.797 | 0 |
| Saline Remote Zone | Up | Ccl12 | 0 | -0.4399812 | 0.355 | 0.822 | 0 |
| Saline Remote Zone | Up | C6 | 4.07E-230 | -0.4414916 | 0.467 | 0.697 | 8.14E-227 |
| Saline Remote Zone | Up | Slitrk6 | 2.29E-159 | -0.4429905 | 0.449 | 0.699 | 4.58E-156 |
| Saline Remote Zone | Up | Sdf2l1 | 0 | -0.4437175 | 0.413 | 0.833 | 0 |
| Saline Remote Zone | Up | Mt3 | 0 | -0.4442952 | 0.23 | 0.447 | 0 |
| Saline Remote Zone | Up | Gap43 | 0 | -0.4449802 | 0.481 | 0.772 | 0 |
| Saline Remote Zone | Up | Irf5 | 6.02E-255 | -0.4454901 | 0.427 | 0.646 | 1.20E-251 |

| | | | | | | | |
|---|---|---|---|---|---|---|---|
| Saline Remote Zone | Up | Arl11 | 6.92E-263 | -0.448122 | 0.426 | 0.646 | 1.38E-259 |
| Saline Remote Zone | Up | Cnpy3 | 0 | -0.4509049 | 0.364 | 0.647 | 0 |
| Saline Remote Zone | Up | Csf1 | 6.07E-288 | -0.4512413 | 0.446 | 0.709 | 1.21E-284 |
| Saline Remote Zone | Up | Itgb2 | 3.20E-161 | -0.4514317 | 0.481 | 0.696 | 6.39E-158 |
| Saline Remote Zone | Up | Cd68 | 0 | -0.4517706 | 0.469 | 0.79 | 0 |
| Saline Remote Zone | Up | Dpep1 | 0 | -0.4535523 | 0.428 | 0.806 | 0 |
| Saline Remote Zone | Up | Cxcl16 | 1.45E-147 | -0.4548217 | 0.521 | 0.723 | 2.89E-144 |
| Saline Remote Zone | Up | Rnd1 | 0 | -0.4570568 | 0.305 | 0.623 | 0 |
| Saline Remote Zone | Up | Epha3 | 0 | -0.4603897 | 0.403 | 0.726 | 0 |

| | | | | | | | |
|---|---|---|---|---|---|---|---|
| Saline Remote Zone | Up | Chd3 | 4.97E-303 | -0.4614852 | 0.427 | 0.715 | 9.94E-300 |
| Saline Remote Zone | Up | Icam1 | 0 | -0.4638344 | 0.335 | 0.655 | 0 |
| Saline Remote Zone | Up | Slc25a24 | 8.14E-273 | -0.4642241 | 0.379 | 0.595 | 1.63E-269 |
| Saline Remote Zone | Up | Sec22b | 1.96E-209 | -0.4645557 | 0.445 | 0.682 | 3.92E-206 |
| Saline Remote Zone | Up | Atf6b | 1.04E-293 | -0.4652757 | 0.362 | 0.608 | 2.09E-290 |
| Saline Remote Zone | Up | Castor1 | 5.08E-237 | -0.4653151 | 0.518 | 0.788 | 1.02E-233 |
| Saline Remote Zone | Up | Rad51 | 0 | -0.465778 | 0.247 | 0.551 | 0 |
| Saline Remote Zone | Up | Cnpy4 | 1.30E-269 | -0.4668567 | 0.473 | 0.701 | 2.59E-266 |
| Saline Remote Zone | Up | Vcan | 4.84E-161 | -0.4690246 | 0.473 | 0.682 | 9.69E-158 |

| | | | | | | | |
|---|---|---|---|---|---|---|---|
| Saline Remote Zone | Up | B4galt5 | 1.29E-194 | -0.4691022 | 0.505 | 0.785 | 2.59E-191 |
| Saline Remote Zone | Up | Glipr1 | 5.72E-285 | -0.4732573 | 0.37 | 0.58 | 1.14E-281 |
| Saline Remote Zone | Up | AC115420.3 | 0 | -0.4737461 | 0.057 | 0.491 | 0 |
| Saline Remote Zone | Up | Fam149a | 0 | -0.474956 | 0.287 | 0.519 | 0 |
| Saline Remote Zone | Up | Arhgap9 | 0 | -0.4767876 | 0.404 | 0.731 | 0 |
| Saline Remote Zone | Up | LOC679894 | 4.30E-274 | -0.4794658 | 0.452 | 0.723 | 8.60E-271 |
| Saline Remote Zone | Up | Ctsz | 3.38E-126 | -0.4798079 | 0.535 | 0.738 | 6.77E-123 |
| Saline Remote Zone | Up | Prc1 | 0 | -0.4801993 | 0.281 | 0.553 | 0 |
| Saline Remote Zone | Up | Hsbp1l1 | 5.02E-301 | -0.481027 | 0.349 | 0.644 | 1.00E-297 |

| | | | | | | | |
|---|---|---|---|---|---|---|---|
| Saline Remote Zone | Up | Egr1 | 7.86E-278 | -0.4815592 | 0.44 | 0.721 | 1.57E-274 |
| Saline Remote Zone | Up | Plxdc2 | 1.48E-98 | -0.4818489 | 0.573 | 0.828 | 2.95E-95 |
| Saline Remote Zone | Up | Pstpip1 | 0 | -0.4862376 | 0.371 | 0.675 | 0 |
| Saline Remote Zone | Up | Spi1 | 1.46E-272 | -0.4872075 | 0.428 | 0.66 | 2.93E-269 |
| Saline Remote Zone | Up | RGD1564664 | 0 | -0.4874184 | 0.321 | 0.544 | 0 |
| Saline Remote Zone | Up | Tmeff2 | 2.92E-260 | -0.4875613 | 0.436 | 0.666 | 5.83E-257 |
| Saline Remote Zone | Up | Cercam | 1.53E-203 | -0.487656 | 0.482 | 0.724 | 3.05E-200 |
| Saline Remote Zone | Up | Irf3 | 1.59E-239 | -0.4878301 | 0.412 | 0.681 | 3.18E-236 |
| Saline Remote Zone | Up | Twist1 | 0 | -0.4881863 | 0.312 | 0.633 | 0 |

| | | | | | | | |
|---|---|---|---|---|---|---|---|
| Saline Remote Zone | Up | Asgr2 | 8.19E-289 | -0.4892326 | 0.456 | 0.755 | 1.64E-285 |
| Saline Remote Zone | Up | Amdhd2 | 0 | -0.4894709 | 0.434 | 0.735 | 0 |
| Saline Remote Zone | Up | Dagla | 0 | -0.490028 | 0.407 | 0.656 | 0 |
| Saline Remote Zone | Up | Ccl22 | 1.67E-150 | -0.4902726 | 0.448 | 0.65 | 3.34E-147 |
| Saline Remote Zone | Up | Man2b1 | 5.44E-161 | -0.4924552 | 0.506 | 0.747 | 1.09E-157 |
| Saline Remote Zone | Up | Rasa3 | 0 | -0.492651 | 0.375 | 0.653 | 0 |
| Saline Remote Zone | Up | Ano6 | 0 | -0.4938349 | 0.408 | 0.709 | 0 |
| Saline Remote Zone | Up | Kdelr3 | 1.79E-182 | -0.4946358 | 0.55 | 0.764 | 3.59E-179 |
| Saline Remote Zone | Up | Ngfr | 0 | -0.4973912 | 0.495 | 0.779 | 0 |

| | | | | | | | |
|---|---|---|---|---|---|---|---|
| Saline Remote Zone | Up | Spats2l | 0 | -0.4974448 | 0.243 | 0.452 | 0 |
| Saline Remote Zone | Up | Rasa4 | 1.27E-302 | -0.499392 | 0.512 | 0.859 | 2.55E-299 |
| Saline Remote Zone | Up | Dse | 0 | -0.4998657 | 0.222 | 0.572 | 0 |
| Saline Remote Zone | Up | Sncg | 1.99E-299 | -0.5006247 | 0.367 | 0.609 | 3.98E-296 |
| Saline Remote Zone | Up | Chpf2 | 2.15E-167 | -0.5008225 | 0.517 | 0.757 | 4.29E-164 |
| Saline Remote Zone | Up | Sat1 | 3.32E-206 | -0.5015321 | 0.482 | 0.758 | 6.64E-203 |
| Saline Remote Zone | Up | Sh3bp2 | 0 | -0.5031927 | 0.291 | 0.545 | 0 |
| Saline Remote Zone | Up | Fos | 2.01E-259 | -0.5040651 | 0.415 | 0.616 | 4.03E-256 |
| Saline Remote Zone | Up | Spsb1 | 4.69E-185 | -0.5058379 | 0.47 | 0.674 | 9.39E-182 |

| | | | | | | | |
|---|---|---|---|---|---|---|---|
| Saline Remote Zone | Up | Sod3 | 4.78E-104 | -0.5071349 | 0.568 | 0.799 | 9.57E-101 |
| Saline Remote Zone | Up | Slc8b1 | 0 | -0.5074397 | 0.338 | 0.652 | 0 |
| Saline Remote Zone | Up | Lyl1 | 0 | -0.5082543 | 0.265 | 0.498 | 0 |
| Saline Remote Zone | Up | Itpr2 | 6.80E-305 | -0.5096088 | 0.446 | 0.712 | 1.36E-301 |
| Saline Remote Zone | Up | AABR07017902.1 | 2.01E-250 | -0.511175 | 0.471 | 0.785 | 4.02E-247 |
| Saline Remote Zone | Up | Cd14 | 3.41E-255 | -0.5126518 | 0.414 | 0.632 | 6.81E-252 |
| Saline Remote Zone | Up | Chpf | 2.18E-52 | -0.5134022 | 0.663 | 0.884 | 4.35E-49 |
| Saline Remote Zone | Up | Tbx18 | 0 | -0.515812 | 0.334 | 0.553 | 0 |
| Saline Remote Zone | Up | Serpine2 | 8.91E-241 | -0.5159326 | 0.416 | 0.687 | 1.78E-237 |

| | | | | | | | |
|---|---|---|---|---|---|---|---|
| Saline Remote Zone | Up | Mast3 | 0 | -0.5253884 | 0.362 | 0.733 | 0 |
| Saline Remote Zone | Up | Fyb1 | 7.94E-307 | -0.5273478 | 0.428 | 0.724 | 1.59E-303 |
| Saline Remote Zone | Up | Map3k7cl | 0 | -0.5284806 | 0.377 | 0.683 | 0 |
| Saline Remote Zone | Up | Fbln5 | 1.60E-181 | -0.5285504 | 0.464 | 0.677 | 3.19E-178 |
| Saline Remote Zone | Up | Stx2 | 0 | -0.530714 | 0.4 | 0.678 | 0 |
| Saline Remote Zone | Up | Fkbp1b | 5.55E-206 | -0.5317451 | 0.497 | 0.745 | 1.11E-202 |
| Saline Remote Zone | Up | Ube2c | 0 | -0.5327365 | 0.278 | 0.551 | 0 |
| Saline Remote Zone | Up | Clec4a1 | 3.19E-201 | -0.5328455 | 0.506 | 0.764 | 6.39E-198 |
| Saline Remote Zone | Up | Col6a1 | 5.20E-225 | -0.5336665 | 0.533 | 0.776 | 1.04E-221 |

| | | | | | | | |
|---|---|---|---|---|---|---|---|
| Saline Remote Zone | Up | Itgbl1 | 6.92E-232 | -0.5351955 | 0.474 | 0.692 | 1.38E-228 |
| Saline Remote Zone | Up | AABR07012100.2 | 5.56E-231 | -0.5398976 | 0.219 | 0.448 | 1.11E-227 |
| Saline Remote Zone | Up | Apobec1 | 6.25E-166 | -0.5414705 | 0.49 | 0.723 | 1.25E-162 |
| Saline Remote Zone | Up | Il18 | 1.62E-177 | -0.5451597 | 0.448 | 0.702 | 3.24E-174 |
| Saline Remote Zone | Up | Cebpa | 1.42E-279 | -0.5456806 | 0.361 | 0.564 | 2.85E-276 |
| Saline Remote Zone | Up | Cpxm1 | 1.20E-176 | -0.5460526 | 0.522 | 0.739 | 2.41E-173 |
| Saline Remote Zone | Up | Vegfd | 0 | -0.5463433 | 0.36 | 0.624 | 0 |
| Saline Remote Zone | Up | Zfp536 | 0 | -0.54677 | 0.476 | 0.808 | 0 |
| Saline Remote Zone | Up | Ccl3 | 0 | -0.5472801 | 0.321 | 0.703 | 0 |

| | | | | | | | |
|---|---|---|---|---|---|---|---|
| Saline Remote Zone | Up | Scara3 | 3.50E-260 | -0.5521524 | 0.376 | 0.646 | 7.00E-257 |
| Saline Remote Zone | Up | F13a1 | 0 | -0.5530398 | 0.297 | 0.539 | 0 |
| Saline Remote Zone | Up | Nnat | 0 | -0.5545308 | 0.472 | 0.818 | 0 |
| Saline Remote Zone | Up | Il17ra | 0 | -0.5552419 | 0.367 | 0.615 | 0 |
| Saline Remote Zone | Up | Myrf | 0 | -0.5563182 | 0.407 | 0.64 | 0 |
| Saline Remote Zone | Up | Pld1 | 0 | -0.5585551 | 0.357 | 0.704 | 0 |
| Saline Remote Zone | Up | Sostdc1 | 1.11E-256 | -0.5612169 | 0.437 | 0.733 | 2.22E-253 |
| Saline Remote Zone | Up | Ltc4s | 0 | -0.5642413 | 0.394 | 0.725 | 0 |
| Saline Remote Zone | Up | Pdpn | 1.42E-282 | -0.564374 | 0.491 | 0.816 | 2.84E-279 |

| | | | | | | | |
|---|---|---|---|---|---|---|---|
| Saline Remote Zone | Up | Chka | 0 | -0.5651018 | 0.338 | 0.695 | 0 |
| Saline Remote Zone | Up | Ptprc | 0 | -0.5655609 | 0.306 | 0.714 | 0 |
| Saline Remote Zone | Up | Edil3 | 3.92E-144 | -0.5688029 | 0.089 | 0.347 | 7.83E-141 |
| Saline Remote Zone | Up | Wasf3 | 0 | -0.5719549 | 0.146 | 0.439 | 0 |
| Saline Remote Zone | Up | Naaa | 0 | -0.5723172 | 0.323 | 0.555 | 0 |
| Saline Remote Zone | Up | Pik3cd | 1.98E-282 | -0.5748268 | 0.438 | 0.709 | 3.96E-279 |
| Saline Remote Zone | Up | Dnah7 | 5.03E-218 | -0.5767554 | 0.318 | 0.526 | 1.01E-214 |
| Saline Remote Zone | Up | Ttc7a | 1.75E-251 | -0.5780111 | 0.389 | 0.596 | 3.49E-248 |
| Saline Remote Zone | Up | Trem2 | 2.22E-187 | -0.5800276 | 0.497 | 0.704 | 4.45E-184 |

| | | | | | | | |
|---|---|---|---|---|---|---|---|
| Saline Remote Zone | Up | Tbxas1 | 0 | -0.5828753 | 0.262 | 0.518 | 0 |
| Saline Remote Zone | Up | Mtmr11 | 0 | -0.5845182 | 0.342 | 0.567 | 0 |
| Saline Remote Zone | Up | Slfn13 | 0 | -0.5874025 | 0.338 | 0.61 | 0 |
| Saline Remote Zone | Up | Large1 | 4.22E-302 | -0.5926097 | 0.492 | 0.786 | 8.45E-299 |
| Saline Remote Zone | Up | NEWGENE-621351 | 0 | -0.5938607 | 0.437 | 0.706 | 0 |
| Saline Remote Zone | Up | Tspan33 | 2.82E-300 | -0.5946322 | 0.488 | 0.806 | 5.65E-297 |
| Saline Remote Zone | Up | Kctd14 | 0 | -0.5966756 | 0.361 | 0.619 | 0 |
| Saline Remote Zone | Up | Hbb | 0 | -0.5986107 | 0.522 | 0.825 | 0 |
| Saline Remote Zone | Up | Igsf11 | 6.98E-146 | -0.5993634 | 0.513 | 0.718 | 1.40E-142 |

| | | | | | | | |
|---|---|---|---|---|---|---|---|
| Saline Remote Zone | Up | AC109737.1 | 1.25E-110 | -0.5997132 | 0.275 | 0.526 | 2.50E-107 |
| Saline Remote Zone | Up | Itk | 1.15E-265 | -0.6032015 | 0.532 | 0.746 | 2.31E-262 |
| Saline Remote Zone | Up | Ptgs2 | 4.84E-272 | -0.6074681 | 0.345 | 0.553 | 9.67E-269 |
| Saline Remote Zone | Up | AABR07007068.1 | 0 | -0.6145351 | 0.28 | 0.588 | 0 |
| Saline Remote Zone | Up | LOC100911486 | 7.16E-305 | -0.6249551 | 0.446 | 0.709 | 1.43E-301 |
| Saline Remote Zone | Up | Rab38 | 4.18E-254 | -0.6250117 | 0.376 | 0.585 | 8.36E-251 |
| Saline Remote Zone | Up | Ccl2 | 0 | -0.6321086 | 0.208 | 0.484 | 0 |
| Saline Remote Zone | Up | LOC690045 | 0 | -0.6331186 | 0.428 | 0.771 | 0 |
| Saline Remote Zone | Up | Mcpt1l1 | 0 | -0.6336768 | 0.235 | 0.64 | 0 |

| | | | | | | | |
|---|---|---|---|---|---|---|---|
| Saline Remote Zone | Up | Arhgef19 | 4.47E-187 | -0.6340673 | 0.49 | 0.707 | 8.94E-184 |
| Saline Remote Zone | Up | Ccl7 | 0 | -0.6348508 | 0.244 | 0.521 | 0 |
| Saline Remote Zone | Up | Nkd2 | 0 | -0.6374344 | 0.297 | 0.581 | 0 |
| Saline Remote Zone | Up | Pamr1 | 0 | -0.6375049 | 0.505 | 0.803 | 0 |
| Saline Remote Zone | Up | Gas2l3 | 0 | -0.6408884 | 0.3 | 0.578 | 0 |
| Saline Remote Zone | Up | Ldlrad4 | 1.16E-214 | -0.6420746 | 0.434 | 0.643 | 2.32E-211 |
| Saline Remote Zone | Up | Cacnb3 | 9.36E-296 | -0.6445418 | 0.381 | 0.635 | 1.87E-292 |
| Saline Remote Zone | Up | Numbl | 3.02E-275 | -0.6452586 | 0.414 | 0.683 | 6.04E-272 |
| Saline Remote Zone | Up | Dact2 | 0 | -0.6498058 | 0.265 | 0.546 | 0 |

| | | | | | | | |
|---|---|---|---|---|---|---|---|
| Saline Remote Zone | Up | Sdc1 | 0 | -0.6503899 | 0.445 | 0.758 | 0 |
| Saline Remote Zone | Up | Cst7 | 4.89E-196 | -0.6616573 | 0.415 | 0.645 | 9.78E-193 |
| Saline Remote Zone | Up | Unc5cl | 2.11E-242 | -0.6657184 | 0.519 | 0.773 | 4.21E-239 |
| Saline Remote Zone | Up | Fdxacb1 | 1.46E-218 | -0.6679407 | 0.481 | 0.727 | 2.93E-215 |
| Saline Remote Zone | Up | Lsp1 | 1.83E-265 | -0.6681454 | 0.36 | 0.592 | 3.66E-262 |
| Saline Remote Zone | Up | Aard | 0 | -0.6716858 | 0.554 | 0.863 | 0 |
| Saline Remote Zone | Up | Spn | 0 | -0.6718375 | 0.304 | 0.606 | 0 |
| Saline Remote Zone | Up | Cpz | 4.83E-295 | -0.6804243 | 0.481 | 0.799 | 9.67E-292 |
| Saline Remote Zone | Up | Tmem178a | 0 | -0.683373 | 0.298 | 0.634 | 0 |

| | | | | | | | |
|---|---|---|---|---|---|---|---|
| Saline Remote Zone | Up | Sulf1 | 0 | -0.683994 | 0.437 | 0.714 | 0 |
| Saline Remote Zone | Up | Slc16a3 | 0 | -0.6857664 | 0.455 | 0.839 | 0 |
| Saline Remote Zone | Up | Ldb2 | 6.84E-282 | -0.6916443 | 0.512 | 0.797 | 1.37E-278 |
| Saline Remote Zone | Up | Ncf4 | 0 | -0.6960539 | 0.38 | 0.646 | 0 |
| Saline Remote Zone | Up | Tns3 | 6.91E-222 | -0.6988423 | 0.494 | 0.725 | 1.38E-218 |
| Saline Remote Zone | Up | Slc1a7 | 0 | -0.7073854 | 0.238 | 0.459 | 0 |
| Saline Remote Zone | Up | Angpt2 | 0 | -0.7102633 | 0.412 | 0.711 | 0 |
| Saline Remote Zone | Up | Rubcnl | 1.25E-169 | -0.710727 | 0.344 | 0.551 | 2.50E-166 |
| Saline Remote Zone | Up | Apbb1ip | 3.02E-295 | -0.7114945 | 0.42 | 0.656 | 6.04E-292 |

| | | | | | | | |
|---|---|---|---|---|---|---|---|
| Saline Remote Zone | Up | Adh6 | 3.96E-213 | -0.7179669 | 0.334 | 0.613 | 7.92E-210 |
| Saline Remote Zone | Up | Rgn | 0 | -0.7245222 | 0.243 | 0.45 | 0 |
| Saline Remote Zone | Up | AABR07066861.1 | 3.43E-246 | -0.7279299 | 0.529 | 0.782 | 6.86E-243 |
| Saline Remote Zone | Up | Myh4 | 7.16E-171 | -0.7294909 | 0.486 | 0.711 | 1.43E-167 |
| Saline Remote Zone | Up | Nr4a1 | 0 | -0.7329264 | 0.334 | 0.585 | 0 |
| Saline Remote Zone | Up | Slc7a6 | 0 | -0.7349274 | 0.332 | 0.596 | 0 |
| Saline Remote Zone | Up | Camk4 | 1.59E-155 | -0.7352298 | 0.435 | 0.674 | 3.18E-152 |
| Saline Remote Zone | Up | Plvap | 0 | -0.735771 | 0.376 | 0.639 | 0 |
| Saline Remote Zone | Up | Ca3 | 1.84E-236 | -0.7434715 | 0.513 | 0.73 | 3.67E-233 |

| | | | | | | | |
|---|---|---|---|---|---|---|---|
| Saline Remote Zone | Up | Inhba | 2.26E-297 | -0.7457033 | 0.445 | 0.747 | 4.52E-294 |
| Saline Remote Zone | Up | Gng8 | 9.08E-275 | -0.7551035 | 0.371 | 0.622 | 1.82E-271 |
| Saline Remote Zone | Up | Il21r | 0 | -0.7599079 | 0.259 | 0.526 | 0 |
| Saline Remote Zone | Up | Esm1 | 0 | -0.770015 | 0.262 | 0.639 | 0 |
| Saline Remote Zone | Up | Ttc37 | 0 | -0.7706908 | 0.228 | 0.437 | 0 |
| Saline Remote Zone | Up | Acap1 | 3.46E-254 | -0.7737195 | 0.412 | 0.647 | 6.91E-251 |
| Saline Remote Zone | Up | Dok3 | 4.93E-215 | -0.7795759 | 0.443 | 0.647 | 9.86E-212 |
| Saline Remote Zone | Up | Prex1 | 6.48E-198 | -0.780327 | 0.417 | 0.653 | 1.30E-194 |
| Saline Remote Zone | Up | Slc11a1 | 0 | -0.7835552 | 0.237 | 0.484 | 0 |

| | | | | | | | |
|---|---|---|---|---|---|---|---|
| Saline Remote Zone | Up | Gldn | 0 | -0.7947576 | 0.282 | 0.612 | 0 |
| Saline Remote Zone | Up | Armcx5 | 2.45E-251 | -0.8112562 | 0.28 | 0.553 | 4.91E-248 |
| Saline Remote Zone | Up | Uchl1 | 2.74E-287 | -0.8235211 | 0.344 | 0.554 | 5.48E-284 |
| Saline Remote Zone | Up | Ccl19 | 0 | -0.8240008 | 0.245 | 0.567 | 0 |
| Saline Remote Zone | Up | Fgfrl1 | 0 | -0.8353701 | 0.39 | 0.782 | 0 |
| Saline Remote Zone | Up | Lpar4 | 0 | -0.8418671 | 0.36 | 0.572 | 0 |
| Saline Remote Zone | Up | Pxdn | 2.26E-237 | -0.8514294 | 0.425 | 0.642 | 4.51E-234 |
| Saline Remote Zone | Up | Mrc2 | 0 | -0.8567905 | 0.348 | 0.572 | 0 |
| Saline Remote Zone | Up | Phldb2 | 0 | -0.8578433 | 0.547 | 0.867 | 0 |

| | | | | | | | |
|---|---|---|---|---|---|---|---|
| Saline Remote Zone | Up | Zpbp2 | 0 | -0.8592705 | 0.349 | 0.728 | 0 |
| Saline Remote Zone | Up | Adamts8 | 0 | -0.8653444 | 0.391 | 0.629 | 0 |
| Saline Remote Zone | Up | Smpd3 | 6.78E-270 | -0.8719905 | 0.3 | 0.531 | 1.36E-266 |
| Saline Remote Zone | Up | Apcdd1 | 0 | -0.8815239 | 0.268 | 0.5 | 0 |
| Saline Remote Zone | Up | Fcgr2a | 2.11E-287 | -0.8823296 | 0.402 | 0.648 | 4.23E-284 |
| Saline Remote Zone | Up | Alox12 | 1.14E-203 | -0.8884708 | 0.43 | 0.708 | 2.28E-200 |
| Saline Remote Zone | Up | Ceacam1 | 0 | -0.8885963 | 0.326 | 0.631 | 0 |
| Saline Remote Zone | Up | Braf | 1.78E-284 | -0.8929977 | 0.185 | 0.463 | 3.55E-281 |
| Saline Remote Zone | Up | Mmp17 | 4.26E-150 | -0.8975969 | 0.41 | 0.619 | 8.51E-147 |

| | | | | | | | |
|---|---|---|---|---|---|---|---|
| Saline Remote Zone | Up | Aldh1a3 | 0 | -0.8984719 | 0.291 | 0.563 | 0 |
| Saline Remote Zone | Up | Cemip | 0 | -0.9117976 | 0.319 | 0.663 | 0 |
| Saline Remote Zone | Up | Dctn4 | 0 | -0.9177202 | 0.239 | 0.593 | 0 |
| Saline Remote Zone | Up | Slc27a3 | 0 | -0.920614 | 0.404 | 0.681 | 0 |
| Saline Remote Zone | Up | Adgrg3 | 1.90E-256 | -0.9217356 | 0.418 | 0.69 | 3.80E-253 |
| Saline Remote Zone | Up | Frzb | 5.81E-307 | -0.9583058 | 0.389 | 0.636 | 1.16E-303 |
| Saline Remote Zone | Up | Tnc | 2.32E-247 | -0.9636815 | 0.506 | 0.739 | 4.64E-244 |
| Saline Remote Zone | Up | Clec4a2 | 7.23E-271 | -0.9716714 | 0.425 | 0.625 | 1.45E-267 |
| Saline Remote Zone | Up | C1qtnf3 | 9.13E-297 | -0.9723635 | 0.477 | 0.708 | 1.83E-293 |

| | | | | | | | |
|---|---|---|---|---|---|---|---|
| Saline Remote Zone | Up | Arnt2 | 1.40E-125 | -0.9929596 | 0.373 | 0.583 | 2.79E-122 |
| Saline Remote Zone | Up | Fam180a | 4.78E-240 | -1.0016007 | 0.392 | 0.634 | 9.57E-237 |
| Saline Remote Zone | Up | Tram2 | 2.15E-266 | -1.0813337 | 0.354 | 0.627 | 4.30E-263 |
| Saline Remote Zone | Up | Tshz2 | 0 | -1.1089078 | 0.491 | 0.692 | 0 |
| Saline Remote Zone | Up | AABR07013288.4 | 5.74E-158 | -1.1225704 | 0.31 | 0.559 | 1.15E-154 |
| Saline Remote Zone | Up | AABR07060872.1 | 0 | -1.1255415 | 0.228 | 0.541 | 0 |
| Saline Remote Zone | Up | Mpz | 1.12E-189 | -1.1267321 | 0.496 | 0.71 | 2.25E-186 |
| Saline Remote Zone | Up | Tmem104 | 1.67E-243 | -1.2180066 | 0.256 | 0.603 | 3.34E-240 |
| Saline Remote Zone | Up | Lilrb4 | 0 | -1.3026533 | 0.292 | 0.527 | 0 |

| | | | | | | | |
|---|---|---|---|---|---|---|---|
| Saline Remote Zone | Up | Trh | 1.61E-179 | -1.3408591 | 0.292 | 0.502 | 3.21E-176 |
| Saline Remote Zone | Up | Cd3g | 1.22E-132 | -1.3580301 | 0.228 | 0.448 | 2.44E-129 |
| Saline Remote Zone | Up | Olfm2 | 0 | -1.4891839 | 0.301 | 0.557 | 0 |
| Saline Remote Zone | Up | Psmb8 | 1.51E-89 | -1.6223973 | 0.208 | 0.45 | 3.02E-86 |
| Saline Remote Zone | Up | AC125248.1 | 0 | -1.6801104 | 0.044 | 0.317 | 0 |
| Saline Remote Zone | Up | Ccl17 | 2.80E-87 | -1.6958191 | 0.139 | 0.364 | 5.61E-84 |
| Saline Remote Zone | Up | AABR07062799.2 | 0 | -2.2253732 | 0.029 | 0.469 | 0 |
| Saline Remote Zone | Up | Adipoq | 3.87E-121 | -2.2611341 | 0.271 | 0.49 | 7.74E-118 |
| Saline Remote Zone | Up | Dntt | 0 | -4.5757165 | 0.029 | 0.384 | 0 |

| | | | | | | | |
|---|---|---|---|---|---|---|---|
| Saline Remote Zone | Up | Krt5 | 0 | -4.9020682 | 0.01 | 0.257 | 0 |
