## Supplemental Table 3 for "Uncovering the Regional and Cell Specific Bioactivity of Injectable Extracellular Matrix Biomaterials in Myocardial Infarction through Spatial and Single Nucleus Transcriptomics"

| Supplementary Table 3. ECM Hydrogel Treated Subacute Infarcts without Visible ECM Compared to Saline Subacute Infarcts | | | | | | | |
|---|---|---|---|---|---|---|---|
| **Spatial Area** | **Direction** | **Gene** | **p_val** | **avg_log2FC** | **pct.1** | **pct.2** | **p_val_adj** |
| Saline Infarct | Up | Hbb | 4.86E-80 | -1.9667269 | 0.126 | 0.39 | 1.23E-75 |
| Saline Infarct | Up | RT1-Bb | 3.52E-92 | -1.0186772 | 0.365 | 0.694 | 8.91E-88 |
| Saline Infarct | Up | Gnas.1 | 1.54E-85 | -0.9986309 | 0.221 | 0.492 | 3.89E-81 |
| Saline Infarct | Up | RT1-A2 | 1.28E-73 | -0.9966319 | 0.211 | 0.453 | 3.22E-69 |
| ECM Hydrogel Treatment with No Visible ECM | Up | Itga7 | 7.11E-20 | 0.26293936 | 0.658 | 0.453 | 1.80E-15 |
| ECM Hydrogel Treatment with No Visible ECM | Up | Pfkm | 8.39E-23 | 0.28284756 | 0.747 | 0.539 | 2.12E-18 |
| ECM Hydrogel Treatment with No Visible ECM | Up | Tmem256 | 2.82E-24 | 0.28620611 | 0.736 | 0.507 | 7.12E-20 |
| ECM Hydrogel Treatment with No Visible ECM | Up | Polr2f | 4.63E-24 | 0.2873761 | 0.541 | 0.338 | 1.17E-19 |
| ECM Hydrogel Treatment with No Visible ECM | Up | Mrps36 | 2.48E-21 | 0.28846551 | 0.656 | 0.45 | 6.28E-17 |
| ECM Hydrogel Treatment with No Visible ECM | Up | Sh3bgr | 2.72E-23 | 0.30008982 | 0.575 | 0.369 | 6.88E-19 |
| ECM Hydrogel Treatment with No Visible ECM | Up | Depp1 | 8.35E-27 | 0.3006658 | 0.499 | 0.295 | 2.11E-22 |
| ECM Hydrogel Treatment with No Visible ECM | Up | Itgb1bp2 | 1.11E-23 | 0.3009704 | 0.545 | 0.345 | 2.82E-19 |

| | | | | | | | |
|---|---|---|---|---|---|---|---|
| ECM Hydrogel Treatment with No Visible ECM | Up | Tomm5 | 3.36E-22 | 0.30462801 | 0.625 | 0.424 | 8.49E-18 |
| ECM Hydrogel Treatment with No Visible ECM | Up | Ndufs3 | 4.56E-24 | 0.30748199 | 0.788 | 0.585 | 1.15E-19 |
| ECM Hydrogel Treatment with No Visible ECM | Up | LOC1083496 | 7.24E-25 | 0.31225333 | 0.63 | 0.415 | 1.83E-20 |
| ECM Hydrogel Treatment with No Visible ECM | Up | Cpt2 | 9.62E-26 | 0.31579564 | 0.648 | 0.429 | 2.43E-21 |
| ECM Hydrogel Treatment with No Visible ECM | Up | Mrps16 | 5.22E-26 | 0.31972209 | 0.654 | 0.434 | 1.32E-21 |
| ECM Hydrogel Treatment with No Visible ECM | Up | Tmem38a | 1.56E-24 | 0.33457857 | 0.575 | 0.372 | 3.95E-20 |
| ECM Hydrogel Treatment with No Visible ECM | Up | Ndufb6 | 5.51E-28 | 0.36024196 | 0.621 | 0.406 | 1.39E-23 |
| ECM Hydrogel Treatment with No Visible ECM | Up | Pet100 | 3.00E-34 | 0.36218546 | 0.559 | 0.326 | 7.58E-30 |
| ECM Hydrogel Treatment with No Visible ECM | Up | Timm8b | 9.49E-34 | 0.38285135 | 0.817 | 0.604 | 2.40E-29 |
| ECM Hydrogel Treatment with No Visible ECM | Up | RT1-S3 | 2.33E-35 | 0.39000979 | 0.667 | 0.431 | 5.89E-31 |

| | | | | | | | |
|---|---|---|---|---|---|---|---|
| ECM Hydrogel Treatment with No Visible ECM | Up | Cpt1b | 2.10E-30 | 0.39619272 | 0.665 | 0.451 | 5.31E-26 |
| ECM Hydrogel Treatment with No Visible ECM | Up | Usp18 | 1.52E-50 | 0.39777074 | 0.283 | 0.079 | 3.84E-46 |
| ECM Hydrogel Treatment with No Visible ECM | Up | Naa38 | 3.37E-41 | 0.39996274 | 0.798 | 0.552 | 8.51E-37 |
| ECM Hydrogel Treatment with No Visible ECM | Up | Nrap | 2.15E-28 | 0.40086511 | 0.694 | 0.486 | 5.43E-24 |
| ECM Hydrogel Treatment with No Visible ECM | Up | Oasl2 | 1.05E-44 | 0.40441749 | 0.342 | 0.127 | 2.66E-40 |
| ECM Hydrogel Treatment with No Visible ECM | Up | Cltb | 1.57E-37 | 0.40592065 | 0.782 | 0.538 | 3.96E-33 |
| ECM Hydrogel Treatment with No Visible ECM | Up | Rtp4 | 4.33E-46 | 0.40934554 | 0.376 | 0.147 | 1.09E-41 |
| ECM Hydrogel Treatment with No Visible ECM | Up | Mx1 | 1.55E-50 | 0.41942077 | 0.293 | 0.084 | 3.92E-46 |
| ECM Hydrogel Treatment with No Visible ECM | Up | RT1-CE10 | 2.18E-50 | 0.47089559 | 0.825 | 0.596 | 5.52E-46 |
| ECM Hydrogel Treatment with No Visible ECM | Up | Mx2 | 2.11E-57 | 0.49405647 | 0.362 | 0.114 | 5.32E-53 |

| | | | | | | | |
|---|---|---|---|---|---|---|---|
| ECM Hydrogel Treatment with No Visible ECM | Up | Rrad | 2.37E-49 | 0.49503837 | 0.691 | 0.415 | 5.98E-45 |
| ECM Hydrogel Treatment with No Visible ECM | Up | LOC1025554 | 9.08E-67 | 0.504826 | 0.367 | 0.105 | 2.29E-62 |
| ECM Hydrogel Treatment with No Visible ECM | Up | LOC691427 | 9.62E-60 | 0.52169932 | 0.498 | 0.213 | 2.43E-55 |
| ECM Hydrogel Treatment with No Visible ECM | Up | Fam162a | 4.52E-44 | 0.52542358 | 0.748 | 0.524 | 1.14E-39 |
| ECM Hydrogel Treatment with No Visible ECM | Up | RT1-T24-4 | 3.31E-57 | 0.5284831 | 0.801 | 0.534 | 8.37E-53 |
| ECM Hydrogel Treatment with No Visible ECM | Up | Oas1a | 1.14E-78 | 0.5602471 | 0.381 | 0.094 | 2.88E-74 |
| ECM Hydrogel Treatment with No Visible ECM | Up | Bst2 | 9.50E-63 | 0.56166271 | 0.65 | 0.344 | 2.40E-58 |
| ECM Hydrogel Treatment with No Visible ECM | Up | AC094217.1 | 4.32E-73 | 0.60092901 | 0.503 | 0.195 | 1.09E-68 |
| ECM Hydrogel Treatment with No Visible ECM | Up | Psmb6 | 6.24E-76 | 0.60592992 | 0.548 | 0.225 | 1.58E-71 |
| ECM Hydrogel Treatment with No Visible ECM | Up | Ly6e | 2.27E-77 | 0.61230162 | 0.91 | 0.705 | 5.74E-73 |

| | | | | | | | |
|---|---|---|---|---|---|---|---|
| ECM Hydrogel Treatment with No Visible ECM | Up | LOC1083511 | 9.47E-64 | 0.66326926 | 0.824 | 0.604 | 2.39E-59 |
| ECM Hydrogel Treatment with No Visible ECM | Up | Hspa8 | 3.33E-79 | 0.67624014 | 0.841 | 0.604 | 8.41E-75 |
| ECM Hydrogel Treatment with No Visible ECM | Up | Ndufa3 | 4.20E-83 | 0.70837778 | 0.798 | 0.495 | 1.06E-78 |
| ECM Hydrogel Treatment with No Visible ECM | Up | Acadl | 2.29E-75 | 0.78446354 | 0.855 | 0.643 | 5.80E-71 |
| ECM Hydrogel Treatment with No Visible ECM | Up | Isg15 | 1.25E-104 | 0.79473918 | 0.489 | 0.134 | 3.15E-100 |
| ECM Hydrogel Treatment with No Visible ECM | Up | LOC1083511 | 5.75E-163 | 0.94301856 | 0.935 | 0.657 | 1.45E-158 |
| ECM Hydrogel Treatment with No Visible ECM | Up | RT1-T24-3 | 8.16E-143 | 0.95350224 | 0.819 | 0.424 | 2.06E-138 |
| ECM Hydrogel Treatment with No Visible ECM | Up | LOC1036934 | 3.64E-147 | 1.12363639 | 0.697 | 0.281 | 9.19E-143 |
| ECM Hydrogel Treatment with No Visible ECM | Up | Ifi27l2b | 2.19E-206 | 1.26976138 | 0.961 | 0.739 | 5.52E-202 |
