## Supplemental Table 2 for "Uncovering the Regional and Cell Specific Bioactivity of Injectable Extracellular Matrix Biomaterials in Myocardial Infarction through Spatial and Single Nucleus Transcriptomics"

**Supplementary Table 2. Global Spatial Comparisons of Infarcts Treated with ECM Hydrogel or Saline in the Subacute MI Model**

| Sample | Direction | Gene | p_val | avg_log2FC | pct.1 | pct.2 | p_val_adj |
|---|---|---|---|---|---|---|---|
| ECM Hydrogel | Up | RT1-T24-3 | 9.02E-81 | 1.017492106 | 0.858 | 0.459 | 2.28E-76 |
| ECM Hydrogel | Up | Cfd | 9.40E-47 | 0.799050003 | 0.838 | 0.537 | 2.38E-42 |
| ECM Hydrogel | Up | Isg15 | 1.14E-56 | 0.767292707 | 0.488 | 0.133 | 2.89E-52 |
| ECM Hydrogel | Up | Oas1a | 3.87E-63 | 0.682862968 | 0.463 | 0.1 | 9.79E-59 |
| ECM Hydrogel | Up | Bst2 | 4.22E-43 | 0.658746357 | 0.71 | 0.367 | 1.07E-38 |
| ECM Hydrogel | Up | Psmb6 | 1.08E-40 | 0.640210576 | 0.576 | 0.25 | 2.72E-36 |
| ECM Hydrogel | Up | LOC10835113 | 1.21E-49 | 0.634984865 | 0.918 | 0.636 | 3.06E-45 |
| ECM Hydrogel | Up | AC094217.1 | 3.64E-41 | 0.630353974 | 0.54 | 0.223 | 9.19E-37 |
| ECM Hydrogel | Up | LOC10255545 | 2.27E-43 | 0.595559095 | 0.423 | 0.129 | 5.75E-39 |
| ECM Hydrogel | Up | Siglec1 | 1.47E-31 | 0.546152677 | 0.452 | 0.184 | 3.71E-27 |
| ECM Hydrogel | Up | Trem2 | 4.86E-28 | 0.519437567 | 0.837 | 0.548 | 1.23E-23 |
| ECM Hydrogel | Up | LOC10834968 | 1.33E-23 | 0.458092053 | 0.708 | 0.445 | 3.37E-19 |
| ECM Hydrogel | Up | RT1-Bb | 2.37E-34 | -0.8963394 | 0.475 | 0.755 | 6.00E-30 |
| Saline | Up | Col5a2 | 1.94E-40 | -0.90711907 | 0.277 | 0.545 | 4.91E-36 |
| Saline | Up | Gnas.1 | 1.69E-56 | -1.16107538 | 0.199 | 0.545 | 4.28E-52 |
| Saline | Up | Hbb | 8.14E-46 | -2.07712688 | 0.106 | 0.42 | 2.06E-41 |
