## Supplemental Table 1 for "Uncovering the Regional and Cell Specific Bioactivity of Injectable Extracellular Matrix Biomaterials in Myocardial Infarction through Spatial and Single Nucleus Transcriptomics"

| Supplementary Table 1. Differentially Expressed Genes Spatially Comparing ECM Hydrogel in the Infarct and Infarct Only Zones in the Subacute MI Model | | | | | | | |
|---|---|---|---|---|---|---|---|
| **Spatial Area** | **Direction** | **Gene** | **p_val** | **avg_log2FC** | **pct.1** | **pct.2** | **p_val_adj** |
| ECM Hydrogel in Infarct | Up | Postn | 7.60E-37 | 1.383992068 | 1 | 0.984 | 9.41E-33 |
| ECM Hydrogel in Infarct | Up | Ltbp2 | 7.36E-36 | 1.362402936 | 0.971 | 0.833 | 9.12E-32 |
| ECM Hydrogel in Infarct | Up | Thbs4 | 8.16E-17 | 1.209466579 | 0.625 | 0.296 | 1.01E-12 |
| ECM Hydrogel in Infarct | Up | Cthrc1 | 4.43E-08 | 1.156578234 | 0.779 | 0.642 | 0.000548953 |
| ECM Hydrogel in Infarct | Up | Ccn2 | 1.82E-17 | 0.969417244 | 0.981 | 0.918 | 2.26E-13 |
| ECM Hydrogel in Infarct | Up | Fn1 | 5.85E-24 | 0.959514876 | 1 | 0.969 | 7.24E-20 |
| ECM Hydrogel in Infarct | Up | Scg2 | 1.39E-07 | 0.90466593 | 0.423 | 0.206 | 0.001720054 |
| ECM Hydrogel in Infarct | Up | Col1a1 | 1.14E-22 | 0.889485622 | 1 | 1 | 1.41E-18 |
| ECM Hydrogel in Infarct | Up | NEWGENE-621351.1 | 5.24E-17 | 0.841428916 | 0.99 | 0.936 | 6.48E-13 |
| ECM Hydrogel in Infarct | Up | Serpine1 | 7.30E-15 | 0.808631017 | 0.644 | 0.286 | 9.04E-11 |
| ECM Hydrogel in Infarct | Up | Lox | 2.56E-07 | 0.793349582 | 0.731 | 0.58 | 0.003174825 |
| ECM Hydrogel in Infarct | Up | Cilp | 7.74E-11 | 0.792733319 | 0.731 | 0.49 | 9.58E-07 |
| ECM Hydrogel in Infarct | Up | Spp1 | 6.70E-20 | 0.787762964 | 0.74 | 0.383 | 8.29E-16 |
| ECM Hydrogel in Infarct | Up | Bgn | 3.31E-26 | 0.786933996 | 1 | 1 | 4.10E-22 |
| ECM Hydrogel in Infarct | Up | Fibin | 1.01E-19 | 0.755761069 | 0.952 | 0.718 | 1.25E-15 |
| ECM Hydrogel in Infarct | Up | Timp1 | 3.16E-09 | 0.72789077 | 0.779 | 0.562 | 3.91E-05 |
| ECM Hydrogel in Infarct | Up | Clec11a | 2.50E-15 | 0.647156317 | 0.76 | 0.395 | 3.10E-11 |
| ECM Hydrogel in Infarct | Up | Fndc1 | 1.31E-14 | 0.644535317 | 0.923 | 0.837 | 1.62E-10 |
| ECM Hydrogel in Infarct | Up | Dkk3 | 3.41E-14 | 0.620780466 | 0.981 | 0.79 | 4.23E-10 |
| ECM Hydrogel in Infarct | Up | Col8a1 | 2.82E-13 | 0.610227089 | 0.75 | 0.444 | 3.49E-09 |
| ECM Hydrogel in Infarct | Up | Aspn | 1.63E-10 | 0.600239687 | 0.865 | 0.665 | 2.01E-06 |
| ECM Hydrogel in Infarct | Up | Sparc | 5.85E-20 | 0.578919031 | 1 | 1 | 7.24E-16 |
| ECM Hydrogel in Infarct | Up | Cemip | 1.40E-07 | 0.55862763 | 0.404 | 0.195 | 0.001734956 |
| ECM Hydrogel in Infarct | Up | Prnp | 2.28E-09 | 0.542954852 | 0.875 | 0.675 | 2.82E-05 |
| ECM Hydrogel in Infarct | Up | Pdlim3 | 1.36E-07 | 0.539874611 | 0.596 | 0.381 | 0.001679295 |
| ECM Hydrogel in Infarct | Up | Tnc | 1.77E-18 | 0.534800878 | 0.404 | 0.082 | 2.19E-14 |
| ECM Hydrogel in Infarct | Up | Csrp2 | 5.82E-08 | 0.534440725 | 0.587 | 0.356 | 0.000720595 |
| ECM Hydrogel in Infarct | Up | Loxl1 | 7.10E-13 | 0.531772867 | 1 | 0.971 | 8.78E-09 |
| ECM Hydrogel in Infarct | Up | Chpf | 8.38E-11 | 0.513487845 | 0.894 | 0.677 | 1.04E-06 |
| ECM Hydrogel in Infarct | Up | P3h3 | 1.49E-10 | 0.505607781 | 0.923 | 0.747 | 1.85E-06 |
| ECM Hydrogel in Infarct | Up | Fhl1 | 1.46E-10 | 0.503378148 | 0.894 | 0.71 | 1.80E-06 |
| ECM Hydrogel in Infarct | Up | Plod2 | 2.47E-10 | 0.503243739 | 0.712 | 0.447 | 3.05E-06 |
| ECM Hydrogel in Infarct | Up | Col3a1 | 3.82E-14 | 0.503040296 | 1 | 1 | 4.73E-10 |
| ECM Hydrogel in Infarct | Up | Col6a2 | 7.42E-12 | 0.499286061 | 1 | 0.981 | 9.18E-08 |

| | | | | | | | |
|---|---|---|---|---|---|---|---|
| ECM Hydrogel in Infarct | Up | Gask1b | 1.77E-12 | 0.49193685 | 0.942 | 0.704 | 2.20E-08 |
| ECM Hydrogel in Infarct | Up | Fkbp10 | 1.81E-07 | 0.479324845 | 0.779 | 0.58 | 0.002246065 |
| ECM Hydrogel in Infarct | Up | Col8a2 | 4.70E-10 | 0.464912554 | 0.673 | 0.368 | 5.81E-06 |
| ECM Hydrogel in Infarct | Up | C1qtnf5 | 2.12E-06 | 0.462660529 | 0.788 | 0.619 | 0.02625023 |
| ECM Hydrogel in Infarct | Up | P4hb | 6.12E-10 | 0.45163606 | 1 | 0.984 | 7.58E-06 |
| ECM Hydrogel in Infarct | Up | Dpt | 4.86E-08 | 0.434295855 | 1 | 0.994 | 0.000601387 |
| ECM Hydrogel in Infarct | Up | Lum | 3.09E-10 | 0.42342176 | 1 | 1 | 3.82E-06 |
| ECM Hydrogel in Infarct | Up | Islr | 3.87E-09 | 0.423308695 | 0.99 | 0.969 | 4.79E-05 |
| ECM Hydrogel in Infarct | Up | Ltbp3 | 1.39E-08 | 0.423133977 | 0.942 | 0.877 | 0.000171748 |
| ECM Hydrogel in Infarct | Up | Mmp2 | 1.83E-09 | 0.415610127 | 1 | 0.99 | 2.27E-05 |
| ECM Hydrogel in Infarct | Up | Enpp1 | 1.08E-09 | 0.409713063 | 0.654 | 0.348 | 1.33E-05 |
| ECM Hydrogel in Infarct | Up | Vim | 7.23E-11 | 0.399618879 | 1 | 0.998 | 8.95E-07 |
| ECM Hydrogel in Infarct | Up | Fat1 | 6.12E-08 | 0.375949614 | 0.596 | 0.329 | 0.000757519 |
| ECM Hydrogel in Infarct | Up | Lgals1 | 9.08E-09 | 0.366141822 | 0.981 | 0.996 | 0.000112382 |
| ECM Hydrogel in Infarct | Up | Serpinh1 | 9.00E-07 | 0.360889851 | 1 | 0.975 | 0.011135709 |
| ECM Hydrogel in Infarct | Up | Sfrp4 | 2.21E-06 | 0.360735388 | 0.356 | 0.167 | 0.027408011 |
| ECM Hydrogel in Infarct | Up | Lmna | 3.70E-08 | 0.360588763 | 1 | 0.953 | 0.000457919 |
| ECM Hydrogel in Infarct | Up | Tmem119 | 1.32E-09 | 0.35229026 | 0.51 | 0.224 | 1.64E-05 |
| ECM Hydrogel in Infarct | Up | Selenom | 2.66E-06 | 0.340849445 | 0.913 | 0.844 | 0.032945332 |
| ECM Hydrogel in Infarct | Up | Sdc2 | 2.09E-06 | 0.323382791 | 0.663 | 0.42 | 0.025906601 |
| ECM Hydrogel in Infarct | Up | Ccn1 | 2.69E-06 | 0.321776841 | 0.394 | 0.187 | 0.03332668 |
| ECM Hydrogel in Infarct | Up | Fxyd5 | 1.09E-06 | 0.316748181 | 0.837 | 0.654 | 0.013532845 |
| ECM Hydrogel in Infarct | Up | Col12a1 | 1.95E-06 | 0.311250236 | 0.375 | 0.173 | 0.024124385 |
| ECM Hydrogel in Infarct | Up | Pkd2 | 6.04E-07 | 0.302052149 | 0.644 | 0.391 | 0.00747681 |
| ECM Hydrogel in Infarct | Up | Sdc1 | 3.39E-07 | 0.278010749 | 0.26 | 0.086 | 0.004195055 |
| ECM Hydrogel in Infarct | Up | Ddah1 | 9.56E-07 | 0.261653802 | 0.385 | 0.171 | 0.011833773 |
| Infarct Only | Up | Gsta1 | 3.49E-06 | -0.35523648 | 0.26 | 0.498 | 0.043196559 |
| Infarct Only | Up | Htra3 | 1.16E-06 | -0.39730794 | 0.952 | 0.961 | 0.014391214 |
| Infarct Only | Up | Cadm3 | 5.27E-07 | -0.40133606 | 0.212 | 0.469 | 0.006520248 |
| Infarct Only | Up | Scara5 | 1.45E-06 | -0.43112871 | 0.337 | 0.588 | 0.017901975 |
| Infarct Only | Up | Serping1 | 1.90E-06 | -0.44065152 | 0.962 | 0.992 | 0.023560396 |
| Infarct Only | Up | Ly6e | 7.33E-08 | -0.48705772 | 0.5 | 0.724 | 0.000907736 |
| Infarct Only | Up | Cfd | 3.34E-08 | -0.70983356 | 0.279 | 0.551 | 0.000413805 |
| Infarct Only | Up | Igfbp3 | 9.23E-11 | -0.78784316 | 0.327 | 0.636 | 1.14E-06 |
| Infarct Only | Up | Gsn | 3.54E-26 | -1.0069881 | 0.99 | 1 | 4.38E-22 |
